# Group A *Streptococcus* remains viable inside fibrin clots and gains access to human plasminogen for subsequent fibrinolysis and dissemination

**DOI:** 10.1101/2023.10.04.560727

**Authors:** Henry M. Vu, Zhong Liang, Yun-Juan Bao, Paulina G. Carles, Jessica C. Keane, Madelyn G. Cerney, Caitlyn N. Dahnke, Ana L. Flores-Mireles, Victoria A. Ploplis, Francis J. Castellino, Shaun W. Lee

## Abstract

Group A *Streptococcus* (GAS) is a Gram-positive bacterial pathogen that causes a wide spectrum of illnesses ranging from pharyngitis and rheumatic fever to more invasive and severe diseases such as necrotizing fasciitis and toxic shock syndrome. Invasive outcomes of GAS infections often result from entry of the bacteria via an open wound into tissue and blood systems. The coagulation cascade serves as an innate defense mechanism that initiates fibrin clots to sequester bacteria and restrict its growth and prevent dissemination into deeper tissues. GAS, especially skin-tropic bacterial strains, utilize the specific virulence factors Plasminogen binding M-protein (PAM) and streptokinase (SK) to manipulate hemostasis and ultimately activate human plasminogen to cause fibrinolysis and escape from the fibrin clot. A major unresolved question regarding this process is to understand the temporal dynamics of how GAS that is enmeshed in a fibrin clot accesses host plasminogen for dissolution of the clot and eventual dissemination. Using fluorescently labeled plasminogen and fibrinogen, we established conditions to observe the process of fibrin clot dissolution by GAS (an AP53 CovR+S-strain) that is sequestered in a fibrin clot using real-time imaging microscopy. We hypothesized that initiation of fibrinolysis by GAS inside a fibrin clot would be determined by the rate of hPg access into the fibrin clot where bacteria are trapped. Our live imaging studies show that GAS trapped inside a fibrin clot, has limited access to hPg; however, at 4.25 h post incubation, when sufficient hPg is accessible to the bacterium, fibrinolysis quickly occurs. If hPg is bound to the bacterial surface prior to being trapped inside a clot, dissolution and bacterial dissemination occurs at a much faster rate of 2.5 h post incubation. During the time which bacteria are trapped in the clot without access to hPg, we did not observe any growth of GAS; however, we demonstrate that the bacteria continue to remain viable inside the fibrin clot. We performed RNA-seq analysis of GAS and the isogenic GAS SK-deficient mutant to understand SK-dependent transcriptional changes during the lag-phase of the GAS bacteria inside the fibrin clot. We observed a dramatic change in the transcription profile of wt GAS inside the fibrin clot over time prior to escape from the fibrin clot (22 gene expression changes at 4h, to 802 gene expression changes at 8h). Furthermore, we also identified gene expression changes that were distinct between wt GAS and the GAS SK-deficient mutant. Our findings reveal for the first time that GAS can engage a latent, growth suspended phase whereby physical structures such as fibrin clots and Neutrophil extracellular traps that immobilize an invading pathogen allow bacteria to remain viable and transcriptionally active for an extended time during host infection. GAS that is trapped in a fibrin clot will therefore enter a state in which the bacteria suspend growth, but remain viable, until sufficient access to hPg allow it to initiate fibrinolysis and escape into surrounding tissues. The viability of GAS while trapped and its readiness to avoid immune defenses allow GAS to act quickly to disseminate when host conditions are more favorable for the bacteria.

## Introduction

Group A *Streptococcus* (*Streptococcus pyogenes* or GAS) is a Gram-positive bacterial pathogen responsible for a broad spectrum of infections from noninvasive clinical conditions, including pharyngitis, impetigo, scarlet fever, and cellulitis, in addition to invasive diseases such as toxic shock syndrome and necrotizing fasciitis (Cunningham, 2000; Carapetis et al., 2005; Plummer and Pavia, 2020; Strus et al., 2017; Walker et al., 2014). Worldwide, GAS is responsible for over 1.7 million invasive infections and 500,000 deaths annually (Carapetis et al., 2005; Nelson et al., 2016).

An important determinant in the ability of GAS to invade deeper host tissues is the ability of the bacteria to subvert several human defenses. To bypass human physical barriers against pathogens, invasive GAS infections can penetrate via an open wound at the skin and tissue levels. The wound creates an entry point for GAS and allows for bacterial colonization and dissemination (Dupuy et al., 1999 ; Nelson et al., 2016). As a result of the initial breach of the epithelial barrier, the host initiates coagulation cascades to produce a blood clot containing fibrin (Fn) and other host hemostatic components (Koivisto et al., 2011). Fibrin clots are normally able to sequester bacteria such as GAS to limit growth and spread of bacteria into deeper tissues (Shannon et al., 2010). To overcome this host response, skin-trophic strains of GAS have evolved two highly specialized virulence factors, human plasminogen (hPg)-binding M-protein (PAM) and a streptokinase variant (SK2b), in concert to initiate fibrinolysis and allow for dissemination into deeper tissues and blood systems (Ringdahl et al., 1998; McKay et al., 2004; Sun et al., 2004; Chandrahas et al., 2015; Sumitomo et al., 2016; Vu et al., 2021). PAM, an M-protein variant, has an increased binding affinity for hPg, which allows for localized recruitment of hPg to the surface of GAS, whilst secretion of GAS enzyme SK2b, rapidly activates PAM-bound hPg to human plasmin (hPm) (Robbins et al., 1967; Berge and Sjobring, 1993; Ringdahl et al., 1998; Li et al., 1999; Sanderson-Smith et al., 2007; Bhattacharya et al., 2014; Zhang et al., 2013; Chandrahas et al., 2015; Yuan et al., 2017; Ayinuola et al., 2020). hPm is a protease that exhibits activity against a wide spectrum of substrates including fibrin, host integrins, and other cell surface proteins (Grainger et al., 1995; Plow et al., 1995; Rømer et al., 1996; Castellino and Ploplis, 2005; Deryugina and Quigley, 2012; Sulniute et al., 2016; Vu et al., 2021). Previously, we demonstrated that the activation of hPg to hPm by a skin-trophic GAS AP53 strain initiates rapid and efficient fibrinolysis of fibrin clots, and also triggers a compromise in cell-cell junctions through the degradation and redistribution of host integrins across cell layers. Therefore, activation of host hPm during GAS infection plays a critical role in the ability of GAS to not only maintain infection, but allows it to subvert innate defenses to gain entry and access to deeper tissues and blood systems.

An important, unresolved question of hPg activation and consequent fibrinolysis during GAS infection, is the overall temporal dynamics of bacterial and cellular events that ultimately result in the ability of the bacterial pathogen to escape the enmeshed fibrin clot and disseminate into deeper tissues. Specifically, we sought to determine the duration needed for GAS during infection conditions to obtain access to hPg while enmeshed in a Fn clot. In this study, we utilized real time live imaging (RTLI) microscopy with fluorescently labeled plasminogen and fibrinogen to examine the events of fibrin clot dissolution by GAS (AP53/covR^+^covS^−^ strain) that is sequestered by Fn. We hypothesized clot dissolution by GAS trapped inside the clot would be determined by the rate of hPg access into the clot where GAS are sequestered. We obtain data to show that GAS initially trapped inside a fibrin clot has limited access to hPg and remains trapped in the clot. However, strikingly, after 4.5 hours of incubation, the bacteria are able to gain access to hPg that is diffused into the clot, and is able to subsequently initiate rapid fibrinolysis and resume growth and dissemination.

Our findings demonstrate for the first time that when GAS that is trapped in a Fn clot, it will enter a state in which the bacteria will suspend its own growth, but still remain viable. With sufficient access to hPg over time, it proceeds to initiate fibrinolysis to escape from the fibrin clot and disseminate to other locations. Despite being trapped in a physical structure such as a fibrin clot that will immobilize bacteria, we propose that the bacteria can enter into a latent suspended growth phase in which it remains viable and transcriptionally active. Our study reveals a key step in the progress of invasive GAS infection, namely that GAS can remain hidden from immune surveillance by remaining viable while trapped in a clot and can be ready to disseminate when the conditions are more favorable.

## Materials and Methods

### Bacterial Cultures

The *Streptococcus pyogenes* isolate used in this study is a clinical isolate from a patient with necrotizing fasciitis AP53/covR^+^covS^−^ (AP53R^+^S^−^) provided by Dr. Gunnar Lindahl (Lund, Sweden). This strain has enhanced invasive capabilities due to mutations in the sensor (S) component in the two-component control of virulence (cov) responder (R)/extracellular sensor (S) gene regulatory system, *covRS* or *csrRS* (Agrahari et al., 2013; Liang et al., 2013). The mutations in this two-component system help the bacteria avoid immune defenses of the host (Mayfield et al. 2014). SK is one virulence factor under control of the CovRS system. An isogenic mutant (AP53R^+^S^−^/ΔSK) was also used in this study and has been discussed in Zhang et al., 2012. The GAS strains were grown overnight for 16-18hrs in Todd-Hewitt (TH) broth at 37°C prior to experimentation.

### Real-time Live Imaging (RTLI)

An inverted Nikon Eclipse Ti-E microscope fitted with an environmental chamber (set for 37° C with 5% CO_2_) at 60x was used to record in real-time the interactions of GAS and the Fn clot. An iXon Ultra 897 electron multiplying charge-coupled device (Andor) and a Neo sCMOS (Andor) were used to capture the images. Images were obtained in the FITC channel with a 480/30 nm excitation filter and 535/45 nm filter. The images were analyzed and reconstructed by ImageJ/FIJI(NIH).

### RTLI of an Fn Clot Infection Model with GAS

Overnight GAS cultures in 4ml TH broth were centrifuged and supernatant was removed. PBS was then added and vortexed with the centrifuged GAS pellet. After mixing, the GAS and PBS combination was used to dilute stock Fibrinogen (Fg) to a concentration of 1g/mL. This Fg-GAS mixture solution was then placed into an optical imaging dish (Mattek). To convert Fg to Fn, thrombin (5 NIH units/mg) was added to the imaging dish. This dish was set at 37° C with 5% CO_2_ for 1 hr in an incubator. The samples were observed visually with microscopy to ensure the mixture and presence of bacteria and the formation of the Fn clot prior to live imaging and addition of any other components to the experiment. To label the Fn clot for microscopy, DyLight 488 NHS-Ester, a fluorescent labeling reagent (50 µg/mL in PBS), was added to the Fn clot. This labeled clot was allowed to incubate for 2 h in the dark at room temperature. After 2h, to remove excess reagent and bacteria not entrapped in the clot, two PBS washes were performed. The samples were then observed by microscopy to ensure that the fibrin clots contained enmeshed GAS bacteria. Then, TH media and hPg (7μg/mL) was added to the Fn clot immediately preceding real-time imaging. Images were obtained every 10 min for 10 h. With certain experiments, hPg (at a concentration of 7μg/mL) was added to the bacteria before the bacteria PBS solution was converted from Fg to Fn.

### Determination of GAS viability after entrapment in Fn clot

Fn clot with GAS enmeshed was formed following the aforementioned procedure for the GAS strains (AP53R^+^S^−^ and AP53R^+^S^−^/ΔSK). After clot formation occurred on the imaging device, the clot was placed into an incubator set at 37° C with 5% CO_2_. Triplicates were performed for each timepoint (4 and 8 h) and treatment. At each timepoint, the dish was removed from the incubator and the clot was extracted from the dish into TH media. The clot was then mechanically broken down in the TH solution using vortexing and pipette mixing of solution. This solution was then placed onto TH agar plates to quantify growth of GAS at timepoints and treatment conditions. These plates were placed at 37° C for 24 hrs. The plates were scored for colony growth. Statistical analysis were performed via Graphpad Prism.

### RNA Isolation and RNA Seq Transcriptome Analysis

AP53R^+^S^−^ and AP53R^+^S^−^/ΔSK were grown overnight and entrapped into a clot in the aforementioned method. After clot formation, hPg (7μg/mL) was also added to clot. The clot was then incubated at 37°C with 2 timepoints of 4 and 8 h. Three replicates of each GAS strain at timepoint were processed for RNA sequencing. The clot was mechanically minced to be homogenous in TH media for RNA Isolation. RNA Isolation and analysis of the RNA-Seq were performed as previously described (Bao et al., 2015; Bao et al., 2016). Briefly, a DNeasy blood and tissue kit (Qiagen, Valencia, CA) were used to extract and purify the RNA from GAS cells (Qiagen, Valencia, CA). The sequencing library was constructed using an NEBNext rRNA Depeletion Kit (Bacteria) (New England Biolabs, Ipswich, MA). From the kit, NEBNext RNase H-based RNA depletion workflow was used to target removal of rRNA from gram-positive and gram-negative organisms. Quality control of the RNA preparation was checked using an Agilent 2100 Bioanalyzer System and Qubit RNA IQ Assay (Agilent Technologies, Santa Clara, Ca; Invitrogen, Waltham, MA). This analysis provided an RNA integrity number (RIN) of >7.0. The Genomics and Bioinformatics Core Facility at the University of Notre Dame performed the RNA sequencing using an Illumina Miseq platform.

The sequencing reads were aligned to the GAS genome of AP53 (Bao et al., 2016) using bwa (version 0.7.17) (Li and Durbin, 2010). The reads which were aligned to multiple loci were removed. Expression levels of genes were normalized using the method GeTMM (Smid et al., 2018). The differential expression of genes was performed between different treatments or time points. The differential expression was calculated using edgeR (version 3.40.2) (Robinson et al., 2010). The Benjamini-Hochberg multiple testing correction was applied to evaluate the false-discovery rate (FDR) (Benjamini and Hochberg, 1995). Volcano plots were graphed by the platform https://www.bioinformatics.com.cn/en.

## Results and Discussion

Severe disease due to invasive outcomes of Group A *Streptococcus* (GAS) are often a result of bacterial entry via an open wound into tissue and blood systems. In these types of skin-based GAS infections, the coagulation cascade provides a critical first line of defense against bacterial spread at the site of infection. The rapid initiation of fibrin clots at the site of localized infection to sequester bacteria and restrict its growth and prevent dissemination into deeper tissues is an important arm of the innate response to curtail the spread of bacteria into deeper tissues. It has been widely established that GAS, especially skin-tropic bacterial strains, have evolved specific virulence factors to manipulate hemostasis and ultimately activate human plasminogen to cause fibrinolysis and escape from the fibrin clot at these infection sites. However, the timing of events involved in the dissemination of GAS from these fibrin clots, and whether the bacteria remain viable while enmeshed in the fibrin clots during infection are major unresolved questions that we explored using real-time live imaging of GAS and *in vitro* fibrin clots.

### GAS trapped inside fibrin clots gain access to hPg in solution

We established conditions to observe the process of fibrin clot dissolution by skin-trophic GAS (an AP53 CovR+S-strain) that is sequestered in a fibrin clot using real-time imaging microscopy. We hypothesized that initiation of fibrinolysis by GAS inside a fibrin clot would be determined by the rate of hPg access into the fibrin clot where the bacteria are trapped. Overnight GAS cultures were rinsed in PBS prior to the addition of Fibrinogen (Fg) to a concentration of 1g/mL. To convert Fg to Fn, thrombin (5 NIH units/mg) was added to the imaging dish. This dish was set at 37° C with 5% CO_2_ for 1 h in an incubator. The samples were observed visually with microscopy to ensure the mixture and presence of bacteria and the formation of the Fn clot with enmeshed GAS bacteria prior to time-lapse live imaging and addition of any other components to the experiment. To visualize the clot structure distinct from GAS, we labeled the Fn clot using DyLight 488 NHS-Ester, a fluorescent labeling reagent (50 µg/mL in PBS). This labeled clot was allowed to incubate for 2 h in the dark at room temperature. After 2h, we removed any excess reagent and bacteria not entrapped in the clot with two PBS washes and the samples were then observed by microscopy to ensure that the fibrin clots contained enmeshed GAS bacteria. TH media and hPg (7μg/mL) was added to the Fn clot immediately preceding the initiation of real-time imaging. Images were obtained every 10 min for 10 h.

Previously, we demonstrated that GAS strains containing PAM and SK have the ability to recruit and activate hPg on the surface to initiate rapid fibrinolysis observed in live imaging studies (Vu et al. 2020). In our first experiment, we initiated a fibrin clot that contained enmeshed GAS bacteria, but did not have any hPg previously bound to GAS surfaces. hPg was then added in solution to the dishes containing enmeshed bacteria at a physiologically relevant concentration of 7μg/mL. Live imaging shows that GAS trapped inside a fibrin clot that has limited access to hPg is significantly delayed in its ability to initiate fibrinolysis. However, in our live imaging conditions, by 4.5 h post incubation, we observed a rapid, dramatic dissolution of fibrin clots, indicating that GAS can eventually gain access to hPg in solution while enmeshed in the fibrin clot (**Figure 1, Movie S1**). Next, we performed an identical experiment in which GAS is trapped inside a fibrin clot, but we wanted to observe if the rapid fibrinolytic event would occur more quickly when hPg is pre-bound to the bacterial surface prior to being trapped inside a clot. Under these conditions, dissolution of the fibrin clot was initated at 2.5 h post incubation, with a gradual loss of fibrin clot structure occurring through 4h. These data show that fibrinolysis is accelerated when hPg is pre-bound to GAS prior to their immobilization in the fibrin clot, and that access to hPg by GAS inside the fibrin clot determines the overall timing of the fibrinolytic event (**Figure 2, Movie S2)**.

**Figure 1.**
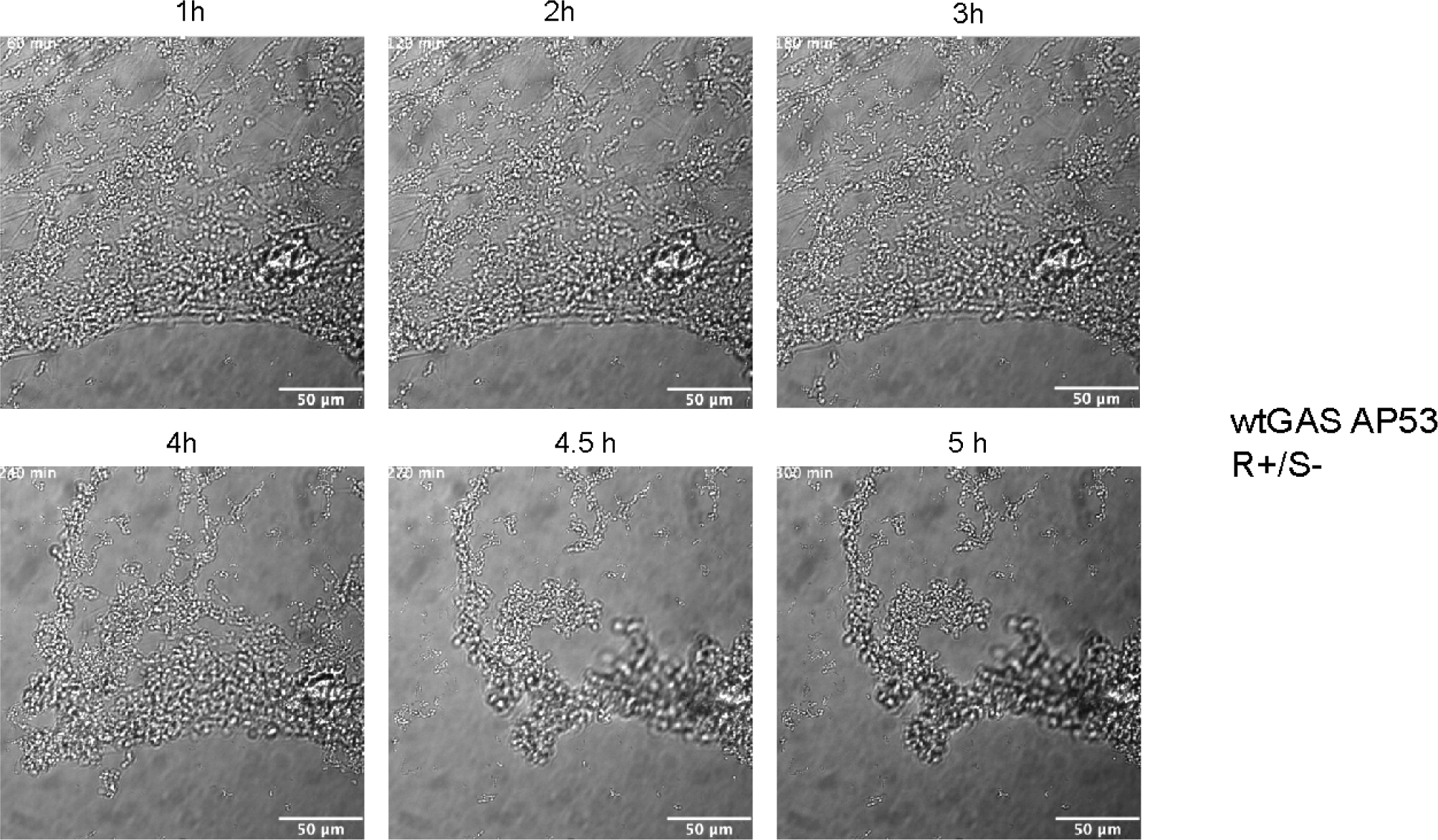
Wt GAS AP53/covR^+^covS^−^ (AP53R^+^S^−^) triggers rapid fibrin clot dissolution at 4.5 h post incubation with hPg in solution. GAS bacteria are enmeshed in fibrin clots prior to initiation of live imaging. Image is acquired in DIC every 10 minutes for 10h for live imaging movie reconstruction. Bacteria remain trapped in the clot until 4.5h, at which point rapid fibrinolysis occurs. Time lapse images (Supplemental movie S1) are shown in panels.

**Figure 2.**
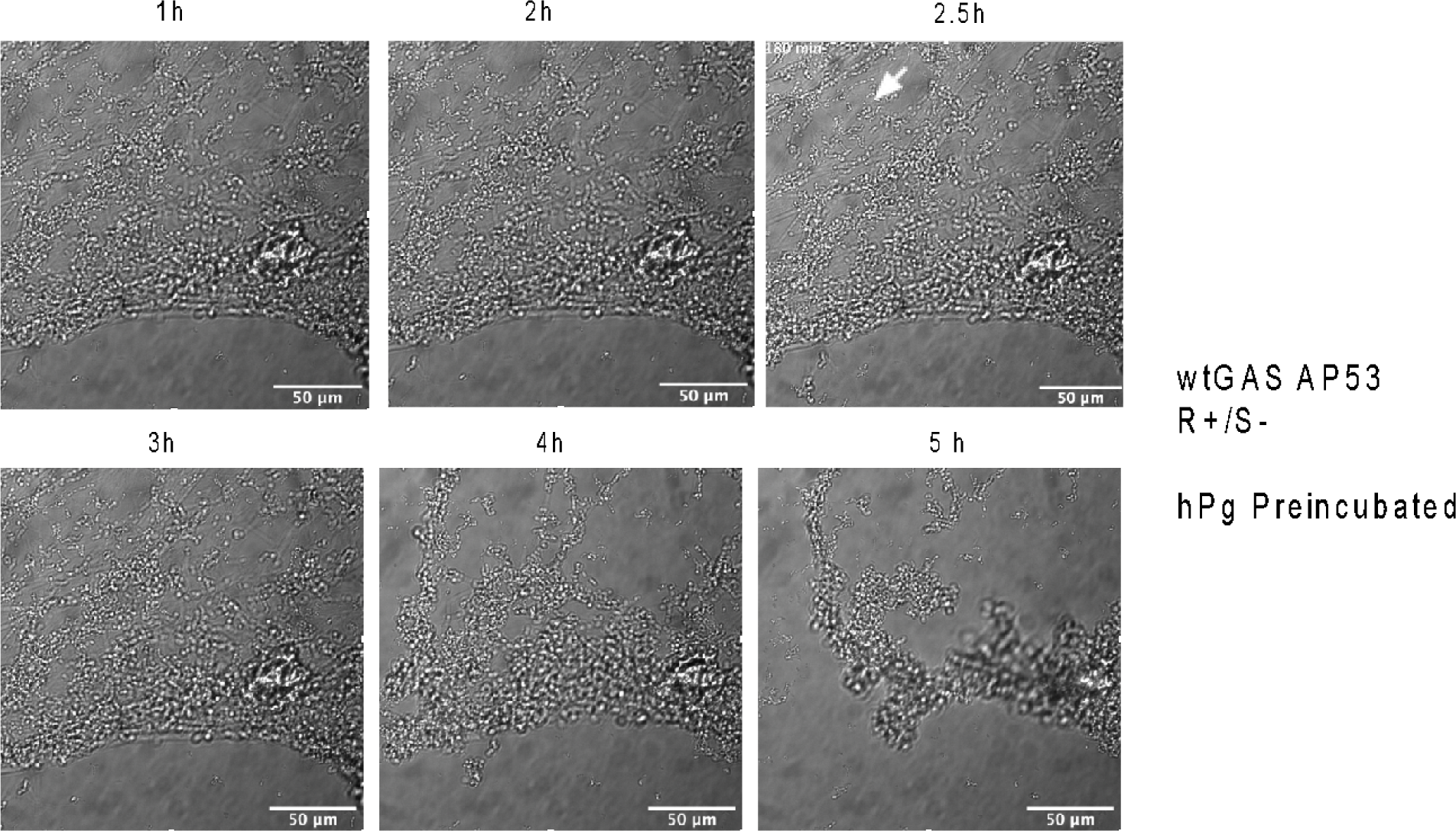
Wt GAS AP53/covR^+^covS^−^ (AP53R^+^S^−^) initiates fibrin clot dissolution at 2.5 h when bacteria are pre-incubated with hPg prior to entrapment in the fibrin clot. GAS bacteria with hPg bound are enmeshed in fibrin clots prior to initiation of live imaging. Image is acquired in DIC every 10 minutes for 10h for live imaging movie reconstruction. Bacteria remain trapped in the clot until 2.5h, at which point structural changes in the fibrin clot are observed (arrow), following by a pronounced fibrinolytic event. Time lapse images (Supplemental movie S2) are shown in panels.

### Escape from fibrin clots by GAS requires SK for hPg activation

We next sought to confirm the critical function of GAS protein SK in the ability of GAS to initiate fibrinolysis via hPg activation and conversion to hPm. In our live imaging study using conditions identical to observe GAS escape from fibrin clots, we observed no dissemination of GAS from enmeshed fibrin clots in our AP53R^+^S^−^/ΔSK isogenic mutant, over the course of 10h (**Figure 3, Movie S3, S4)**. Fluorescently labeled fibrin was imaged along with DIC capture of the SK-mutant inside the fibrin clot, confirming that there was no fibrinolytic activity observed over the course of 10h, and that the fibrin structures retained their original structure throughout the 10h with bacteria intertwined inside the fibrin clot in DIC images. This finding importantly confirms that the fibrinolysis event that occurs 4.5h after GAS are initially trapped in the clot requires SK activation of hPg by GAS.

**Figure 3.**
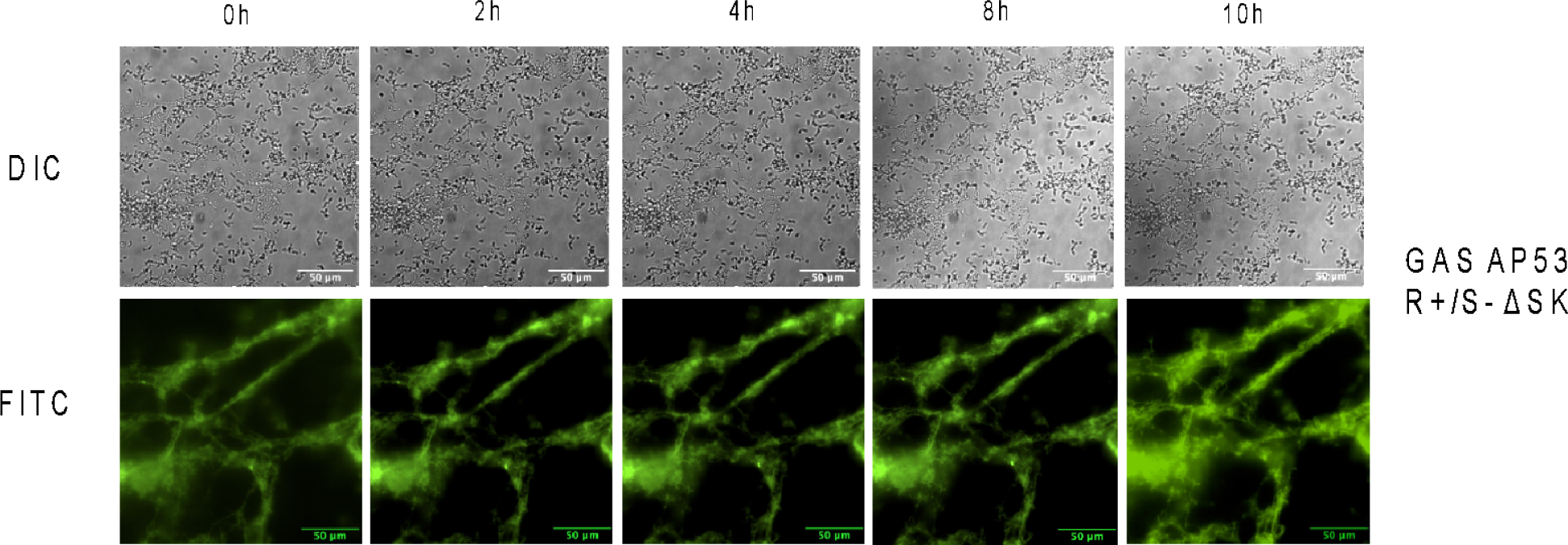
SK is required for hPg activation and fibrinolysis by GAS. GAS AP53R^+^S^−^/ΔSK isogenic mutant remains enmeshed inside a fibrin clot over the course of 10h in the presence of hPg and TH media (Supplemental movie S3). DIC images (top panels) show immobilized GAS bacteria without growth or movement over the course of 10h. Fluorescently labeled fibrin was imaged along with DIC capture of the SK-mutant inside the fibrin clot (bottom panels), confirming that there was no fibrinolytic activity observed over the course of 10h. Fibrin structures retained their original structure throughout the 10h.

### GAS remain viable while encased in fibrin clots in solution

Live imaging time-lapse images showed that during the period when GAS are sequestered inside the fibrin clot, there was no significant growth of GAS, indicating that rapid growth of the bacterium is only observed after the process of fibrinolysis and subsequent escape from the fibrin clot **(figure S1, Movie S5)**. Therefore, we wanted to determine if GAS that are trapped inside the clot without access to hPg would continue to remain viable and metabolically active in some form. Fn clots with GAS enmeshed were formed following the aforementioned procedure for the GAS strains (AP53R^+^S^−^ and AP53R^+^S^−^/ΔSK), and allowed to incubate for 4 and 8h. Following incubation, the dish was removed from the incubator and the clot was extracted from the dish into TH media. The clot was mechanically broken down in TH solution prior to plating on TH plates to quantify growth of GAS at time points and treatment conditions. These plates were placed at 37° C for 24 h prior to counting, and triplicates were performed for each timepoint (4 and 8 h) and treatment. Quantification of bacteria revealed that GAS continued to remain viable while enmeshed in the clot without appreciable growth or reproduction observed (**Figure 4).** An average of 1 x 10^7^ CFUs were recovered from the collected fibrin clots at 4h and 8h, indicating that there was not significant growth over the course of 8h, even though most of the bacteria remained viable inside the fibrin clot. A similar trend occurred with AP53R^+^S^−^/ΔSK strain, with an average of 5 x 10^6^ CFUs recovered from the fibrin clot. The bacteria displayed no visible sign of growth or division during the time course we observed, even though we performed our experiment in TH broth which is permissible for GAS growth. Significantly, our finding demonstrates that when GAS are enmeshed in a fibrin clot, they enter a state in which they suspend growth and division, but remain viable. This is the first observation that GAS can remain viable for up to 8h while not engaging in growth in the presence of nutrient-rich conditions.

**Figure 4.**
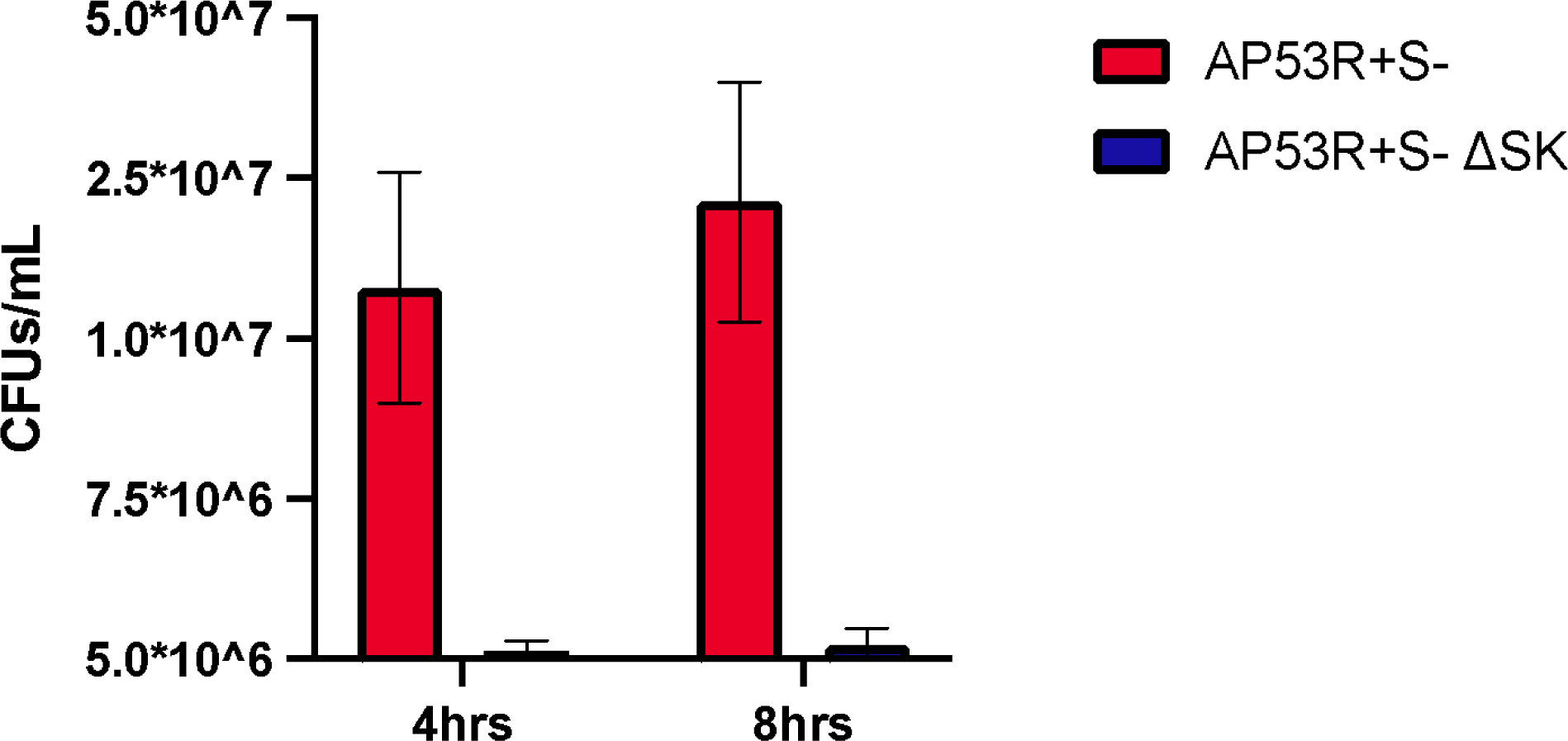
GAS AP53/covR^+^covS^−^ (AP53R^+^S^−^) and ΔSK mutant remains **viable while encased in fibrin clots in solution.** GAS bacteria isolated from mechanically broken fibrin clots after incubation for 10h was plated on TH plates to quantify growth of GAS at time points and treatment conditions. Bacteria CFU counts were quantified and performed in triplicate for significance. CFU of wt GAS are in red; ΔSK mutant shown in blue. Error bars indicated in graph of triplicate CFU sampling (p<0.0001). Timelapse movie shows that the bacteria displayed no visible sign of growth or division during the 10h time course (figure S1).

### GAS remain transcriptionally active while encased in fibrin clots in solution

We performed RNA-seq analysis of wt GAS and the isogenic GAS SK-deficient mutant to understand SK-dependent transcriptional changes during the lag-phase of the GAS bacteria trapped inside the fibrin clot. We observed a dramatic change in the transcription profile of wt GAS inside the fibrin clot over time prior to escape from the fibrin clot (22 gene expression changes at 4h, to 802 gene expression changes at 8h). Furthermore, we also identified gene expression changes that were distinct between wt GAS and the GAS SK-deficient mutant (**Figure 5, Supp data file 1)**. The detection of active transcription by GAS while trapped in a fibrin clot shows for the first time that GAS can engage a latent, growth suspended phase whereby physical structures such as fibrin clots and Neutrophil extracellular traps that immobilize an invading pathogen allow bacteria to remain viable and transcriptionally active for an extended time during host infection. GAS that is trapped in a fibrin clot will therefore enter a state in which the bacteria suspend growth, but remain viable, until sufficient access to hPg allow it to initiate fibrinolysis and escape into surrounding tissues.

**Figure 5.**
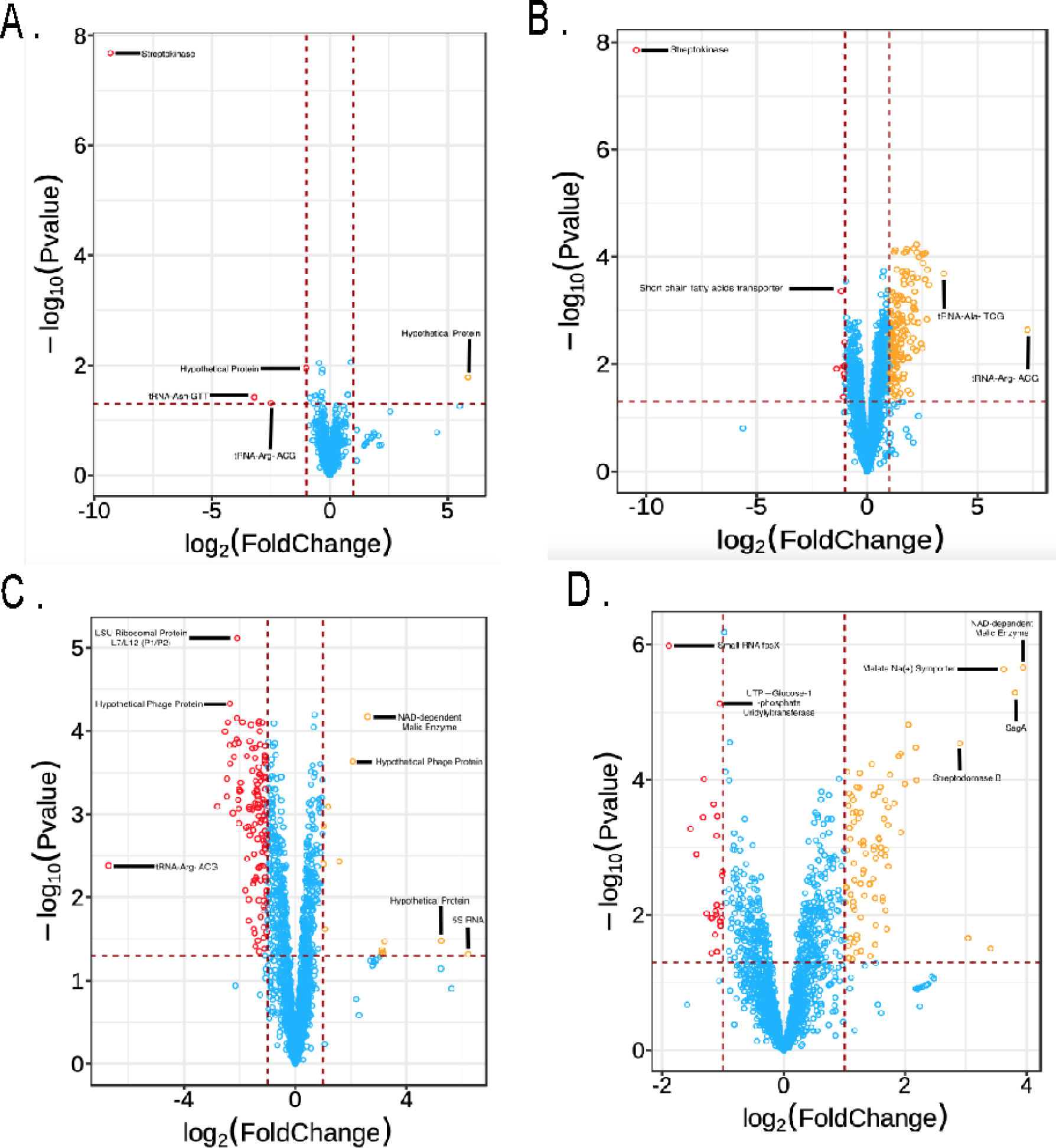
Transcriptional analysis of gene expression changes in GAS trapped inside a fibrin clot without access to plasminogen. RNA-seq analysis of wt GAS and the isogenic GAS ΔSK mutant mutant was compared along with 4 and 8h timepoints of each GAS strain in a trapped clot. Graphs show the following: **A.** Transcriptional differences at 4h between wt GAS and GAS ΔSK mutant ; **B.** Transcriptional differences at 8h between wt GAS and GAS ΔSK mutant. **C.** Transcriptional changes between 4h and 8h in wt GAS; **D**. Transcriptional changes between 4h and 8h in GAS ΔSK mutant. X-axis in graphs show fold changes in transcript. Upregulated genes are identified in orange, downregulated genes are identified in red, and genes expression changes that fall below significance threshold are in blue. Gene transcript names of distinct upregulated and downregulated genes are noted in each graph. A complete profile of all gene expression changes are listed in excel **(Supp data file 1)**.

## Conclusion

In this study, we provide the first real-time live image capture of the human pathogen GAS trapped inside a fibrin clot over the course of 10h. Our data reveal the striking finding that GAS can enter a latent state whereby growth and bacterial division are halted; however, the bacteria inside the fibrin clot can remain transcriptionally active during the course of latency. It has not previously been reported to our knowledge that GAS can enter a latent phase inside fibrin clots. In our live imaging conditions, we incubated the fibrin-enmeshed GAS in the presence of TH broth, thereby providing nutrient rich conditions for access by GAS trapped in the clot. We speculated therefore, in our conditions, that GAS would have the ability to grow inside the trapped clot over time and that we would observe a phenotype where the bacterial growth would allow eventual escape from the fibrin structure. However, our live imaging reveals that GAS trapped inside the clot in the absence of hPg do not have the ability to escape the fibrin clot over the course of infection. Thus, hPg activation by GAS remains a critical event by which GAS can initiate fibrinolysis to escape from the enmeshed clots to disseminate into deeper tissues. Our imaging demonstrates that when hPg is absent, or when SK is inactivated in the GAS mutant, fibrinolysis does not occur over the timecourse observed.

We also reveal with our live-imaging studies that hPg added exogenously to the enmeshed fibrin clot will eventually diffuse into areas where GAS and SK can activate the hPg to initiate fibrinolysis. We observed the fibrinolysis event by GAS at 4.5h post incubation, at which time, fibrinolysis happened rapidly, presumably by rapid amplified hPm generation at that time. These data therefore confirm that GAS trapped inside fibrin clots can gain access to sufficient hPg in solution for SK-dependent activation and initation of fibrinolysis. Given that we used a concentration of hPg that mimics physiological levels of circulating hPg, our results likely reflect conditions in which skin-trophic GAS can initiate an infection at a site of skin breach, and persist in a viable state, especially as innate defenses initiate hemostasis to contain the spread of bacteria and bleeding.

To better understand the state of viable GAS trapped inside a fibrin clot without access to plasminogen, we performed RNA-seq analysis of wt GAS and the isogenic GAS SK-deficient mutant to understand both global and SK-dependent transcriptional changes during the lag-phase of the GAS bacteria trapped inside the fibrin clot. We hypothesized that genes involved in transcription and RNA maintenance would continue to be active. Indeed, a survey of transcriptional changes indicated that many genes involved in transcription (rRNA subunit genes, tRNA, ribosomal subunit genes) were maintained over the course of incubation **(Supp data file 1)**. The RNA-seq survey also revealed distinctions between transcripts activated in wt GAS vs. the isogenic SK-mutant GAS. Interestingly, we found that when GAS enter a latent state inside the fibrin clot without the addition of hPg, SK is highly downregulated, at both the 4h and 8h timepoints (−9.3 fold at 4h, −10.4 fold at 8h). These results suggest that GAS may be able to sense environmental conditions in which lack of access to hPg causes GAS to downregulate SK until at which point, access to hPg on the surface, especially as it interacts with PAM protein would initiate events to trigger rapid changes in expression of SK, thereby leading to rapid hPg activation and subsequent fibrinolysis. This is consistent with the often dramatic and rapid fibrinolysis that we observe in live imaging during GAS infection. We are currently pursuing studies to identify and analyze the profile of genes that are regulated in a possible SK-dependent manner, and better understand possible mechanisms of GAS latency and reactivation, especially in the context of a skin-based infection that involves the host activation of hemostasis and innate defenses.

In summary, we provide the first live-image progress of fibrin clot dissolution by wt GAS that is trapped inside a fibrin clot. Our findings demonstrate that GAS bacteria trapped inside a fibrin clot eventually gain access to hPg in solution, and utilize SK to initiate a delayed, but rapid fibrinolytic event. Importantly, we reveal for the first time that GAS can engage a latent, growth suspended phase whereby physical structures such as fibrin clots and Neutrophil extracellular traps that immobilize an invading pathogen would allow bacteria to remain viable and transcriptionally active for an extended time during host infection. GAS that is trapped in a fibrin clot will therefore enter a state in which the bacteria suspend growth, but remain viable, until sufficient access to hPg allow it to initiate fibrinolysis and escape into surrounding tissues. The potential of GAS to sense and respond to specific host environments such as fibrin clots, and regulate and maintain transcription are novel findings that have significance for understanding how GAS persists during infection and can avoid immune defenses to efficiently disseminate into deeper tissues when conditions present a favorable opportunity for the bacteria.

## Author Contributions

HV, JL, YB, AF, VP, FC, and SL: designed the overall project and experimental aims; HV, ZL, YN, PC, CP, JK, MC: performed experimental work and analyzed the results; HV and SL: wrote the paper, with all authors contributing to the proofreading and editing of the paper.

## Supplemental data

**Supplemental figure 1.**
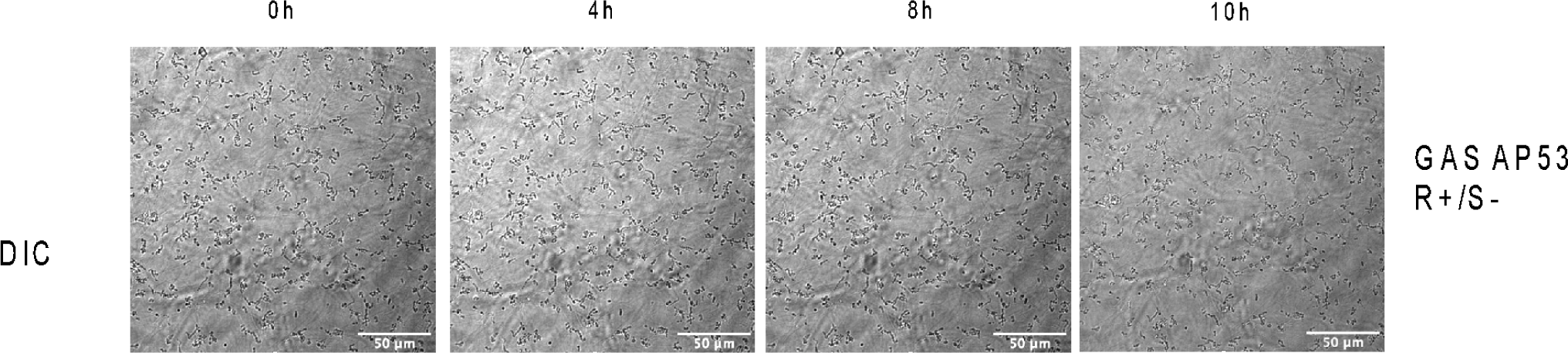

Supplemental movies S1-S4 (separate file).

Supplemental file 1 (separate file).

## Supporting information

Supplemental RNA Seq

