## Supplemental RNA Seq for "Group A *Streptococcus* remains viable inside fibrin clots and gains access to human plasminogen for subsequent fibrinolysis and dissemination"

| GeneID | wildSK_4hr_Rp1 | wildSK_4hr_Rp2 | wildSK_4hr_Rp3 | wildSK_8hr_Rp1 | wildSK_8hr_Rp2 | wildSK_8hr_Rp3 |
|---|---|---|---|---|---|---|
| Total reads (paired-ends) | 30198572 | 24317014 | 22951630 | 23214358 | 22207224 | 21786316 |
| Mapped to gene regions | 25725930 | 20690510 | 19602576 | 19071597 | 18217436 | 17765409 |
| Mapping to intergenic region | 1837965 | 1534744 | 1365115 | 1603197 | 1307155 | 1431180 |
| Unmapped | 1062256 | 808147 | 792129 | 1203150 | 1455485 | 1411531 |

| deltaSK_4hr_Rp1 | deltaSK_4hr_Rp2 | deltaSK_4hr_Rp3 | deltaSK_8hr_Rp1 | deltaSK_8hr_Rp2 | deltaSK_8hr_Rp3 |
|---|---|---|---|---|---|
| 22141574 | 22755506 | 23177688 | 20755188 | 24172614 | 25221062 |
| 18724070 | 19680913 | 18591369 | 17131675 | 16457297 | 15853810 |
| 1359034 | 1489619 | 1382064 | 1508180 | 5870427 | 5461816 |
| 848887 | 500896 | 1756652 | 1010094 | 788658 | 2202286 |

| GeneID | wildSK_4hr_Rp1 | wildSK_4hr_Rp2 | wildSK_4hr_Rp3 | wildSK_8hr_Rp1 | wildSK_8hr_Rp2 | wildSK_8hr_Rp3 |
|---|---|---|---|---|---|---|
| AP53_1 | 272.9033626 | 251.3140447 | 240.6567726 | 265.0367002 | 299.0519331 | 231.4085007 |
| AP53_10 | 128.0037628 | 107.287773 | 111.1355239 | 150.7362014 | 143.8467242 | 103.8028558 |
| AP53_100 | 594.8515006 | 729.2956898 | 701.1685766 | 512.968433 | 447.2763518 | 508.1882142 |
| AP53_1000 | 163.0364496 | 126.289466 | 159.0191582 | 214.3399505 | 182.7788205 | 175.012605 |
| AP53_1001 | 120.4718588 | 116.9325458 | 147.2283544 | 176.6668698 | 161.8190282 | 173.1194272 |
| AP53_1002 | 360.[illegible] | [illegible] | [illegible] | [illegible] | [illegible] | [illegible]2415 |
| AP53_1003 | 86.9[illegible] | [illegible] | [illegible] | [illegible] | [illegible] | [illegible]1083 |
| AP53_1004 | 541.[illegible] | [illegible] | [illegible] | [illegible] | [illegible] | [illegible]9856 |
| AP53_1005 | 268.[illegible] | [illegible] | [illegible] | [illegible] | [illegible] | [illegible]2266 |
| AP53_1006 | 64.3[illegible] | [illegible] | [illegible] | [illegible] | [illegible] | [illegible]0838 |
| AP53_1007 | 49.0[illegible] | [illegible] | [illegible] | [illegible] | [illegible] | [illegible]0105 |
| AP53_1008 | 56.6[illegible] | [illegible] | [illegible] | [illegible] | [illegible] | [illegible]5031 |
| AP53_1009 | 51.9[illegible] | [illegible] | [illegible] | [illegible] | [illegible] | [illegible]0206 |
| AP53_101 | 330.[illegible] | [illegible] | [illegible] | [illegible] | [illegible] | [illegible]4814 |
| AP53_1010 | 60.2[illegible] | [illegible] | [illegible] | [illegible] | [illegible] | [illegible]0252 |
| AP53_1011 | 50.0[illegible] | [illegible] | [illegible] | [illegible] | [illegible] | [illegible]4346 |
| AP53_1012 | 58.7[illegible] | [illegible] | [illegible] | [illegible] | [illegible] | [illegible]9685 |
| AP53_1013 | 64.8[illegible] | [illegible] | [illegible] | [illegible] | [illegible] | [illegible]6233 |
| AP53_1014 | 87.2[illegible] | [illegible] | [illegible] | [illegible] | [illegible] | [illegible]7427 |
| AP53_1015 | 79.3[illegible] | [illegible] | [illegible] | [illegible] | [illegible] | [illegible]2653 |
| AP53_1016 | 61.2[illegible] | [illegible] | [illegible] | [illegible] | [illegible] | [illegible]5973 |
| AP53_1017 | 121.[illegible] | [illegible] | [illegible] | [illegible] | [illegible] | [illegible]7558 |
| AP53_1018 | 69.5[illegible] | [illegible] | [illegible] | [illegible] | [illegible] | [illegible]5421 |
| AP53_1019 | 183.[illegible] | [illegible] | [illegible] | [illegible] | [illegible] | [illegible]6242 |
| AP53_102 | 78.1[illegible] | [illegible] | [illegible] | [illegible] | [illegible] | [illegible]7976 |
| AP53_1020 | 69.5[illegible] | [illegible] | [illegible] | [illegible] | [illegible] | [illegible]7748 |
| AP53_1021 | 101.[illegible] | [illegible] | [illegible] | [illegible] | [illegible] | [illegible]0275 |
| AP53_1022 | 110.[illegible] | [illegible] | [illegible] | [illegible] | [illegible] | [illegible]4936 |
| AP53_1023 | 232.[illegible] | [illegible] | [illegible] | [illegible] | [illegible] | [illegible]7352 |
| AP53_1024 | 202.[illegible] | [illegible] | [illegible] | [illegible] | [illegible] | [illegible]6275 |
| AP53_1025 | 81.6[illegible] | [illegible] | [illegible] | [illegible] | [illegible] | [illegible]0491 |
| AP53_1026 | 61.5[illegible] | [illegible] | [illegible] | [illegible] | [illegible] | [illegible]9546 |
| AP53_1027 | 25.6[illegible] | [illegible] | [illegible] | [illegible] | [illegible] | [illegible]8904 |
| AP53_1028 | 40.4[illegible] | [illegible] | [illegible] | [illegible] | [illegible] | [illegible]0392 |

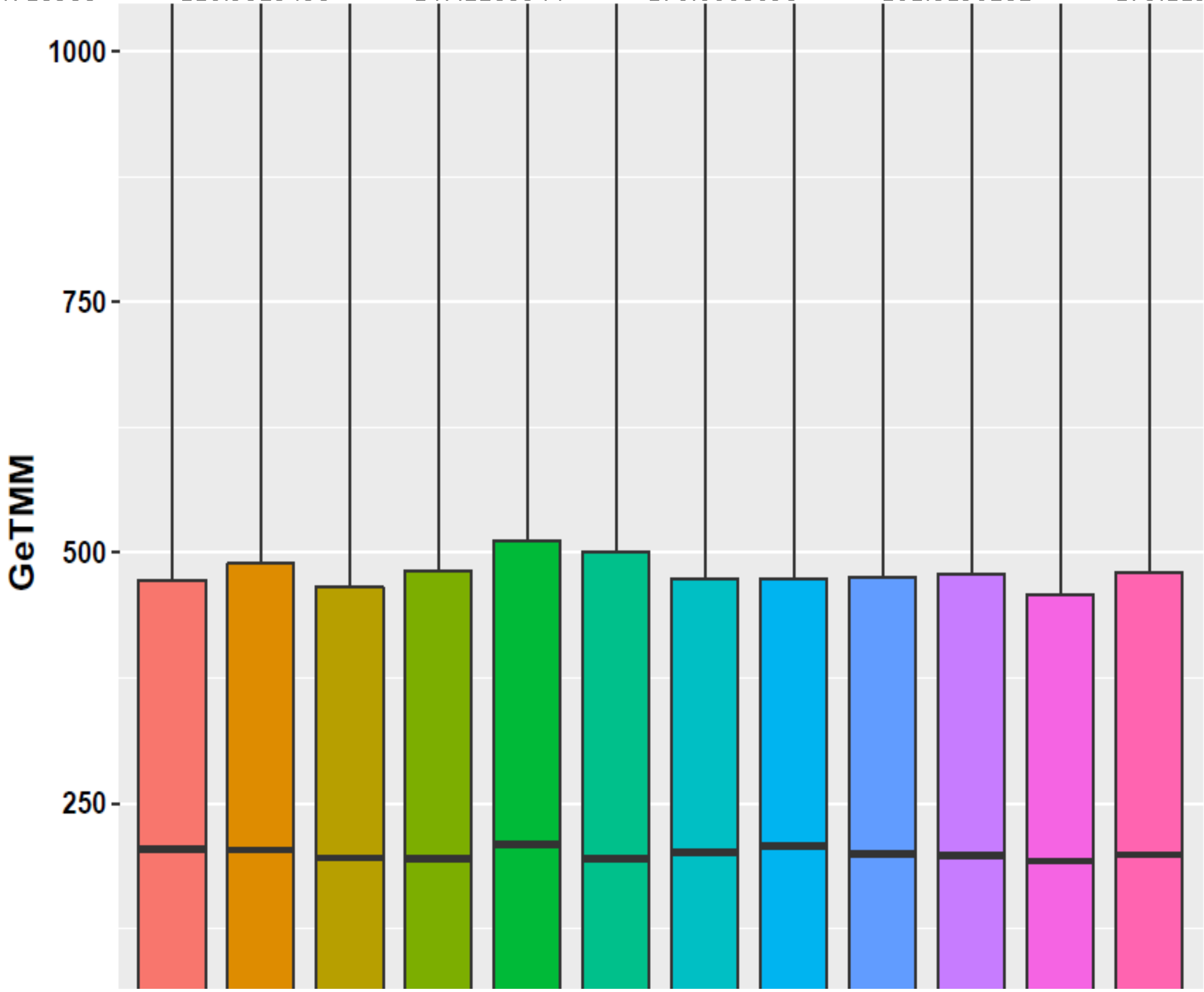

| | | | | | | |
|---|---|---|---|---|---|---|
| AP53_1029 | 38.7[illegible] | [illegible] | [illegible] | [illegible] | [illegible] | [illegible]1664 |
| AP53_103 | 129.[illegible] | [illegible] | [illegible] | [illegible] | [illegible] | [illegible]9517 |
| AP53_1030 | 42.4[illegible] | [illegible] | [illegible] | [illegible] | [illegible] | [illegible]6674 |
| AP53_1031 | 76.2[illegible] | [illegible] | [illegible] | [illegible] | [illegible] | [illegible]4078 |
| AP53_1032 | 73.9[illegible] | [illegible] | [illegible] | [illegible] | [illegible] | [illegible]5326 |
| AP53_1033 | 65.4[illegible] | [illegible] | [illegible] | [illegible] | [illegible] | [illegible]8304 |
| AP53_1034 | 83.[illegible] | [illegible] | [illegible] | [illegible] | [illegible] | [illegible]2895 |
| AP53_1035 | 33.2[illegible] | [illegible] | [illegible] | [illegible] | [illegible] | [illegible]2922 |
| AP53_1036 | 58.0[illegible] | [illegible] | [illegible] | [illegible] | [illegible] | [illegible]0773 |
| AP53_1037 | 59.4[illegible] | [illegible] | [illegible] | [illegible] | [illegible] | [illegible]5949 |
| AP53_1038 | 58.8[illegible] | [illegible] | [illegible] | [illegible] | [illegible] | [illegible]2849 |
| AP53_1039 | 54.49651735 | 70.84747098 | 70.87664036 | 30.50828875 | 38.9099643 | 34.61642988 |
| AP53_104 | 175.5381253 | 165.6977858 | 180.2473954 | 215.8586527 | 169.3240122 | 212.6558284 |
| AP53_1040 | 58.13940003 | 50.104502 | 34.9374354 | 29.22137876 | 28.53729031 | 22.61626467 |
| AP53_1041 | 66.02668701 | 71.59945714 | 71.94357362 | 41.75912495 | 41.53524636 | 31.07670757 |
| AP53_1042 | 84.10480528 | 100.4063944 | 106.9899474 | 51.58886038 | 57.29199808 | 50.54707838 |
| AP53_1043 | 68.77751183 | 78.01782155 | 95.47639854 | 44.59770172 | 50.47689717 | 42.97720047 |
| AP53_1044 | 72.46573892 | 67.56460667 | 94.45099813 | 48.11000631 | 61.63036765 | 42.03533603 |
| AP53_1045 | 77.92816791 | 80.69271829 | 107.5997709 | 46.61599187 | 57.70031522 | 43.96356245 |
| AP53_1046 | 82.36395515 | 60.59046401 | 63.30703188 | 40.62622755 | 35.63842999 | 35.97018746 |
| AP53_1047 | 82.70546289 | 74.35157054 | 80.29924925 | 45.43722818 | 46.44607417 | 34.70807562 |
| AP53_1048 | 84.08849739 | 91.02886819 | 79.79802707 | 47.47086371 | 51.64989853 | 53.36157036 |
| AP53_1049 | 48.02529081 | 38.63739734 | 27.61767358 | 22.2281548 | 24.34178479 | 19.05827833 |
| AP53_105 | 142.340766 | 123.8536979 | 154.5812121 | 166.413556 | 150.8559948 | 135.5970421 |
| AP53_1050 | 76.09703608 | 65.32339635 | 70.40838686 | 46.47915864 | 58.10300934 | 38.49916486 |
| AP53_1051 | 248.5994718 | 299.7661588 | 279.8655531 | 267.8677503 | 258.2295506 | 221.3030677 |
| AP53_1052 | 171.5709175 | 221.5295512 | 220.0941469 | 145.666408 | 156.1600378 | 135.7792773 |
| AP53_1053 | 111.3305825 | 142.2398088 | 124.8991703 | 107.3457232 | 94.78746187 | 84.66560045 |
| AP53_1054 | 49.83326802 | 35.32940099 | 30.21159873 | 20.17398286 | 20.45780717 | 11.04107733 |
| AP53_1055 | 206.3597233 | 213.2682656 | 236.5250164 | 192.2172937 | 200.4475605 | 190.5730448 |
| AP53_1056 | 150.9366283 | 142.6363919 | 160.7928421 | 151.9721845 | 145.9478561 | 139.6250178 |
| AP53_1057 | 56.25732019 | 43.46707201 | 29.35655348 | 23.31536644 | 22.73430844 | 22.46565691 |
| AP53_1058 | 20.96849648 | 21.21607086 | 25.62590963 | 17.22987329 | 18.18287244 | 17.66393134 |
| AP53_1059 | 122.8250825 | 114.314543 | 119.6976649 | 125.7018733 | 118.4076028 | 111.8591275 |
| AP53_106 | 154.3194354 | 146.327168 | 168.6606193 | 176.4107195 | 167.9536399 | 161.1543121 |

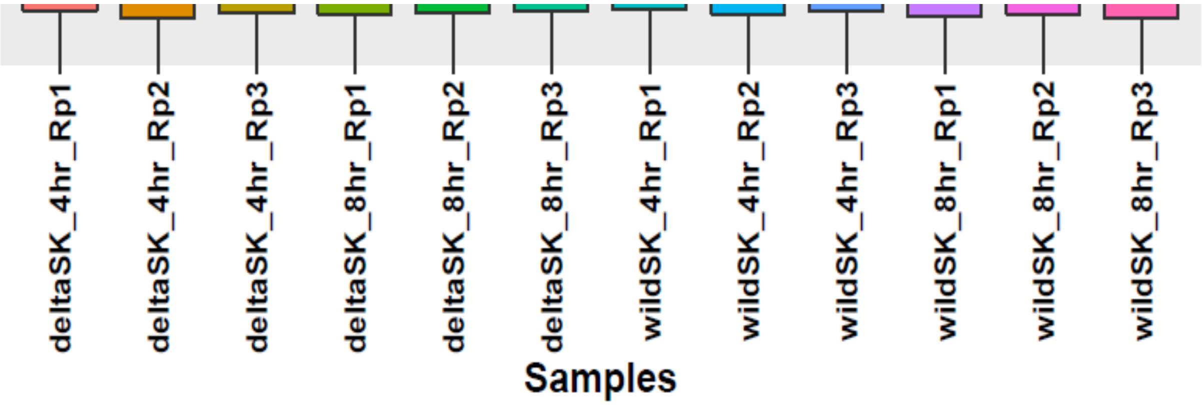

| | | | | | | |
|---|---|---|---|---|---|---|
| AP53_1060 | 50.98222673 | 37.94744382 | 31.89002088 | 55.88793208 | 46.06327686 | 43.32388098 |
| AP53_1061 | 36.24889328 | 32.01156267 | 30.16998495 | 39.77751619 | 44.78763732 | 37.44371366 |
| AP53_1062 | 236.8047764 | 256.9526914 | 211.9920152 | 267.8975875 | 250.215064 | 215.0391642 |
| AP53_1063 | 95.86314105 | 85.494837 | 91.4751188 | 117.6635994 | 108.1628036 | 108.3561092 |
| AP53_1064 | 77.60275593 | 70.8449266 | 69.31074608 | 99.39013501 | 112.3540865 | 102.4418687 |
| AP53_1065 | 43.17043393 | 47.08932801 | 44.78425223 | 50.7329974 | 49.26488851 | 38.6836248 |
| AP53_1066 | 314.6460473 | 238.4767965 | 265.1140412 | 308.3589267 | 238.5278161 | 307.7537439 |
| AP53_1067 | 62.40378144 | 53.29663014 | 58.95919928 | 62.11756158 | 49.82801256 | 60.80496385 |
| AP53_1068 | 82.93674762 | 68.60215308 | 81.97197217 | 116.6729922 | 94.15990719 | 106.0351753 |
| AP53_1069 | 40.03658021 | 40.2472889 | 41.45702714 | 55.86106288 | 60.1516358 | 57.80394322 |
| AP53_107 | 253.6776927 | 250.0835112 | 266.663601 | 295.1965259 | 270.3187036 | 302.7167916 |
| AP53_1070 | 56.46110684 | 59.58407624 | 53.71893819 | 83.59188674 | 77.70283692 | 92.51695375 |
| AP53_1071 | 129.441106 | 169.9809889 | 242.0612239 | 128.8148678 | 127.4870666 | 130.0141938 |
| AP53_1072 | 643.9063259 | 711.2432179 | 811.8446073 | 524.342571 | 448.015629 | 519.3865991 |
| AP53_1073 | 612.985653 | 706.8350487 | 902.5443433 | 479.7369676 | 457.5165353 | 503.9029025 |
| AP53_1074 | 682.0928561 | 775.5250098 | 914.467329 | 503.8075406 | 469.4514617 | 572.4422194 |
| AP53_1075 | 654.3497562 | 746.7553063 | 933.9638146 | 492.758649 | 448.2041894 | 546.3967815 |
| AP53_1076 | 521.1869151 | 587.2714934 | 652.2289589 | 364.5867351 | 310.5006069 | 383.2171477 |
| AP53_1077 | 145.1006251 | 133.4197627 | 139.41394 | 97.6863284 | 88.38896326 | 94.26215213 |
| AP53_1078 | 191.7356742 | 190.0593359 | 183.180567 | 151.2011519 | 122.0917814 | 156.1456849 |
| AP53_1079 | 103.5931626 | 143.0916026 | 148.4118743 | 74.80829487 | 76.47339952 | 78.55328486 |
| AP53_108 | 254.1542959 | 219.7862397 | 251.6130163 | 293.9590532 | 254.5179986 | 234.0348847 |
| AP53_1080 | 181.0711318 | 242.441847 | 257.5936313 | 144.3300189 | 161.393237 | 165.234172 |
| AP53_1081 | 21.44413711 | 17.89820612 | 13.62483864 | 21.24247192 | 11.64654575 | 16.88143329 |
| AP53_1082 | 32.45916105 | 27.56064348 | 25.87689108 | 26.42176756 | 23.02109763 | 25.60558936 |
| AP53_1083 | 30.77572268 | 22.11990998 | 20.24177115 | 23.81148337 | 18.21519755 | 20.88269216 |
| AP53_1084 | 39.16175078 | 27.68438515 | 29.53723268 | 27.20506309 | 15.75848945 | 23.71546337 |
| AP53_1085 | 42.61696683 | 30.23190585 | 24.83742011 | 23.92650421 | 22.84514743 | 22.89768878 |
| AP53_1086 | 22.26252753 | 20.19748158 | 18.64941545 | 18.83672205 | 16.28164578 | 17.51931436 |
| AP53_1087 | 20.05946738 | 17.58530858 | 17.71310172 | 18.31890398 | 16.08727564 | 16.61180271 |
| AP53_1088 | 18.71026944 | 16.00928562 | 18.04932772 | 15.44144623 | 17.55211755 | 16.35689288 |
| AP53_1089 | 106.9832822 | 92.80469685 | 83.86323095 | 105.65154 | 109.0658849 | 98.53901928 |
| AP53_109 | 78.82819289 | 92.89556806 | 88.92664721 | 36.20752569 | 32.87780423 | 34.34029401 |
| AP53_1090 | 115.3383669 | 108.8043689 | 96.90512799 | 101.2461803 | 112.3907361 | 118.5495434 |
| AP53_1091 | 246.3540196 | 321.1733654 | 314.5662009 | 227.0539259 | 232.7850482 | 251.3936205 |

| | | | | | | |
|---|---|---|---|---|---|---|
| AP53_1092 | 109.2312375 | 99.73839779 | 68.15174597 | 64.56105929 | 70.25072578 | 60.91850223 |
| AP53_1093 | 4.952886993 | 3.622256001 | 6.235533458 | 4.822396197 | 6.018671731 | 4.899040389 |
| AP53_1094 | 11.83793031 | 10.79842355 | 16.67338232 | 15.80973478 | 9.606045819 | 11.57845395 |
| AP53_1095 | 12.80818172 | 11.80231757 | 16.60637546 | 12.21925718 | 12.25035909 | 8.006609055 |
| AP53_1096 | 41.8154856 | 38.38844504 | 45.47312798 | 48.22328557 | 49.86231 | 54.95445306 |
| AP53_1097 | 66.01684663 | 67.30599251 | 68.6155875 | 80.14460794 | 79.41657331 | 94.5250127 |
| AP53_1098 | 21.84952574 | 18.03735641 | 19.88416447 | 19.86401432 | 20.43408522 | 16.44895194 |
| AP53_1099 | 58.83297031 | 51.09287412 | 56.5274913 | 63.70121206 | 56.53916336 | 62.37812479 |
| AP53_11 | 172.1450403 | 167.8565252 | 186.4457054 | 193.8189877 | 195.0243754 | 165.3330554 |
| AP53_110 | 212.2314236 | 206.2738471 | 237.5508771 | 303.4225866 | 270.1622918 | 259.6056241 |
| AP53_1100 | 82.40251248 | 70.91995959 | 105.7405955 | 79.73788083 | 65.64973945 | 88.21635886 |
| AP53_1101 | 110.4030769 | 120.9002167 | 151.3675387 | 106.0749691 | 110.2398742 | 113.362328 |
| AP53_1102 | 162.113236 | 149.6407397 | 100.1687843 | 125.7082041 | 89.72204271 | 107.956972 |
| AP53_1103 | 142.8690331 | 152.6153462 | 175.2569646 | 177.5525854 | 193.3073142 | 201.6796922 |
| AP53_1104 | 283.4771282 | 317.4120733 | 300.0408472 | 271.4570659 | 230.6298399 | 265.631693 |
| AP53_1105 | 141.1522845 | 188.6816932 | 148.8033967 | 121.5249717 | 101.9507326 | 126.9967336 |
| AP53_1106 | 135.3178514 | 147.7710986 | 154.679852 | 128.4913337 | 120.9781291 | 124.7990991 |
| AP53_1107 | 154.9180075 | 187.557697 | 170.0319447 | 100.145594 | 97.36600393 | 103.8837306 |
| AP53_1108 | 328.23801 | 330.8826081 | 314.0383331 | 377.0936514 | 411.3987594 | 384.6054766 |
| AP53_1109 | 117.3674236 | 106.9890736 | 112.1881928 | 113.8483968 | 128.8323035 | 112.0311309 |
| AP53_111 | 127.5169321 | 75.59625103 | 96.04582879 | 220.0587326 | 150.1670062 | 193.5318514 |
| AP53_1110 | 171.9555041 | 160.5302613 | 165.3873267 | 191.9779094 | 180.0954039 | 184.2942951 |
| AP53_1111 | 321.3739384 | 346.9716276 | 362.9613965 | 328.0904324 | 348.3205912 | 337.1222452 |
| AP53_1112 | 631.3212168 | 876.4092568 | 826.6609227 | 610.3960931 | 596.7530039 | 688.8773657 |
| AP53_1113 | 225.8580272 | 222.650259 | 231.2457884 | 254.7840322 | 286.0035553 | 266.7687741 |
| AP53_1114 | 190.5541363 | 165.8633024 | 170.3366658 | 190.3185384 | 213.6805629 | 176.7043739 |
| AP53_1115 | 35.06442584 | 33.49756008 | 33.53764629 | 30.59390989 | 41.88177224 | 36.04810453 |
| AP53_1116 | 224.129929 | 225.5803047 | 198.2096674 | 166.4949861 | 159.4839945 | 196.0478306 |
| AP53_1117 | 414.2341988 | 468.2605627 | 431.2156761 | 316.5332955 | 257.1972624 | 321.508801 |
| AP53_1118 | 17.42928461 | 16.3877129 | 16.85468962 | 19.14584197 | 19.47525981 | 18.55120638 |
| AP53_1119 | 193.5428626 | 252.0813316 | 221.2470423 | 162.8068036 | 138.8461126 | 159.3795815 |
| AP53_112 | 68.95908788 | 47.08932801 | 51.13114324 | 94.7722578 | 72.46034167 | 75.14057343 |
| AP53_1120 | 453.5435 | 503.4935841 | 470.5012214 | 462.5216347 | 460.7040739 | 449.3334691 |
| AP53_1121 | 411.8113207 | 484.953096 | 476.0089595 | 408.8758718 | 392.3365504 | 439.6685117 |
| AP53_1122 | 352.6452314 | 407.0892045 | 365.9971388 | 333.861117 | 298.0092467 | 353.4520344 |

| | | | | | | |
|---|---|---|---|---|---|---|
| AP53_1123 | 58.41332788 | 58.54638958 | 57.01040575 | 52.66362831 | 61.78584952 | 51.90142789 |
| AP53_1124 | 27.12443702 | 28.38260866 | 26.07302356 | 25.04356493 | 27.89700978 | 20.32561963 |
| AP53_1125 | 121.5657387 | 143.5037306 | 141.5174888 | 105.7772193 | 109.7004304 | 104.8254001 |
| AP53_1126 | 335.6640369 | 330.6859567 | 341.3409911 | 486.6449702 | 385.5438677 | 438.9777903 |
| AP53_1127 | 578.8020908 | 481.3765151 | 489.2501841 | 818.0210406 | 705.5410862 | 636.6096249 |
| AP53_1128 | 1327.255038 | 1773.961827 | 1548.24923 | 863.2717733 | 613.0123702 | 815.5425555 |
| AP53_1129 | 1327.338975 | 1793.505537 | 1743.256289 | 1034.457619 | 877.6895598 | 1140.796132 |
| AP53_113 | 39.4034948 | 30.44529094 | 33.76815402 | 49.19716163 | 45.24082794 | 47.44169239 |
| AP53_1130 | 4707.67041 | 5980.015361 | 5311.061731 | 2520.095678 | 1828.590872 | 2160.917338 |
| AP53_1131 | 408.9727888 | 432.9516833 | 322.3423966 | 350.0282317 | 344.8787535 | 326.9324241 |
| AP53_1132 | 610.4139467 | 659.3310867 | 645.1225342 | 569.4960127 | 542.7318048 | 552.7247652 |
| AP53_1133 | 874.9207943 | 887.2802318 | 723.6397219 | 976.5610055 | 952.3784522 | 922.4497478 |
| AP53_1134 | 461.3480372 | 457.1739855 | 434.2917317 | 387.8535162 | 371.2866791 | 346.0985658 |
| AP53_1135 | 145.6043388 | 129.1258985 | 125.9062837 | 191.7495031 | 179.0842517 | 192.2734042 |
| AP53_1136 | 44.4038749 | 39.32735087 | 24.35424795 | 50.26459276 | 43.11938322 | 39.62702057 |
| AP53_1137 | 36.94682624 | 30.59956333 | 36.15575804 | 40.02734514 | 42.57986736 | 37.23147662 |
| AP53_1138 | 78.55034127 | 84.74829988 | 94.93213566 | 100.5440761 | 114.4789469 | 97.74944381 |
| AP53_1139 | 85.6610185 | 77.80719976 | 94.14362359 | 106.2972981 | 104.8307024 | 104.6442407 |
| AP53_114 | 38.04230966 | 29.17562561 | 45.18007264 | 58.24631488 | 50.44920609 | 56.03688926 |
| AP53_1140 | 108.5656468 | 69.62780979 | 44.05858148 | 76.87949311 | 82.96777352 | 69.45632262 |
| AP53_1141 | 50.65330914 | 44.11102863 | 23.83359455 | 39.12155246 | 41.861013 | 34.60095193 |
| AP53_1142 | 275.9479192 | 380.2995372 | 387.6116971 | 165.5724308 | 146.1655785 | 175.8164804 |
| AP53_1143 | 892.2216727 | 961.3796722 | 881.200239 | 581.020456 | 481.2055867 | 473.6699164 |
| AP53_1144 | 114.2474698 | 141.5913997 | 122.1276904 | 75.34291084 | 78.56516135 | 73.68149138 |
| AP53_1145 | 127.9893366 | 127.6050886 | 128.380365 | 142.1239163 | 197.2256831 | 165.1847303 |
| AP53_1146 | 723.4848402 | 728.7353302 | 745.5790972 | 842.302496 | 966.313956 | 886.6958816 |
| AP53_1147 | 647.4758165 | 548.5968777 | 559.9028063 | 712.4882664 | 818.7692705 | 681.5950025 |
| AP53_1148 | 2231.231063 | 2250.163615 | 2306.636602 | 2372.058903 | 2853.062596 | 2654.139128 |
| AP53_1149 | 2175.15303 | 2165.646673 | 2059.852513 | 2215.930209 | 2570.155468 | 2225.148677 |
| AP53_115 | 33.4704105 | 27.23399882 | 35.24686518 | 48.90194057 | 38.65543993 | 44.77770249 |
| AP53_1150 | 206.4461334 | 261.4360628 | 248.959428 | 147.709304 | 133.1431558 | 161.979452 |
| AP53_1151 | 222.0678636 | 240.1190833 | 253.8293903 | 219.9755503 | 209.9861161 | 213.5617361 |
| AP53_1152 | 99.86906475 | 102.3057482 | 106.7374249 | 136.9674423 | 162.2121433 | 137.6185965 |
| AP53_1153 | 113.8620195 | 114.553846 | 129.1073789 | 138.8591762 | 164.1383048 | 141.1070071 |
| AP53_1154 | 96.52466253 | 86.00536043 | 90.44113209 | 118.9272046 | 125.7570973 | 114.6058679 |

| | | | | | | |
|---|---|---|---|---|---|---|
| AP53_1155 | 2655.08142 | 3070.668899 | 2613.228509 | 2203.771725 | 1741.007036 | 2008.46737 |
| AP53_1156 | 147.0034142 | 151.1535069 | 140.0877878 | 140.2502452 | 151.7090844 | 133.6644334 |
| AP53_1157 | 12.71257698 | 12.72632188 | 13.61137536 | 12.83147896 | 15.20101101 | 13.83657129 |
| AP53_1158 | 15.67939742 | 13.04667772 | 14.353927 | 12.10208136 | 16.43314807 | 14.75398589 |
| AP53_1159 | 229.6830212 | 297.4062822 | 272.3324881 | 240.2759005 | 264.7975885 | 285.0066979 |
| AP53_116 | 241.3994308 | 209.5483075 | 236.6353416 | 257.86618 | 275.3436913 | 236.2799392 |
| AP53_1160 | 154.4579232 | 154.008815 | 139.2766366 | 177.9684968 | 193.2298769 | 162.4586598 |
| AP53_1161 | 158.4463238 | 156.7307328 | 122.4706744 | 208.2276179 | 196.5314342 | 187.5148363 |
| AP53_1162 | 42.11328687 | 41.22963652 | 45.5891666 | 46.46840495 | 67.42474439 | 55.75089441 |
| AP53_1163 | 1458.43583 | 1828.085525 | 1760.804705 | 1048.131593 | 998.1588843 | 1052.920536 |
| AP53_1164 | 749.1146055 | 649.9804302 | 654.2717682 | 733.9016252 | 775.0163984 | 789.5300936 |
| AP53_1165 | 734.7905582 | 726.7619497 | 701.3778911 | 799.0992172 | 774.3604677 | 719.6980455 |
| AP53_1166 | 311.1048288 | 304.8732134 | 287.9708637 | 304.7338286 | 284.9288012 | 292.9699022 |
| AP53_1167 | 233.9037608 | 268.9395344 | 274.0685432 | 213.4138028 | 247.2175987 | 258.5011283 |
| AP53_1168 | 152.8970946 | 160.1801649 | 168.6687748 | 133.9746826 | 140.1147176 | 148.3862284 |
| AP53_1169 | 124.6850687 | 115.1877408 | 84.53437721 | 121.3578336 | 112.528669 | 118.5043462 |
| AP53_117 | 117.7632821 | 129.6886411 | 155.0201705 | 131.5357367 | 130.2012899 | 130.1789913 |
| AP53_1170 | 250.5316232 | 232.1445835 | 271.6540162 | 282.9059354 | 305.753539 | 272.5457497 |
| AP53_1171 | 84.6140505 | 69.54731521 | 94.41124602 | 107.0670372 | 126.8558401 | 89.30098622 |
| AP53_1172 | 132.7201386 | 132.9825939 | 147.8458409 | 174.0429145 | 202.087649 | 162.3367177 |
| AP53_1173 | 880.7760945 | 1089.861887 | 1065.2193 | 758.6772882 | 747.0757091 | 807.6567534 |
| AP53_1174 | 10.62656835 | 11.14540308 | 8.521220154 | 9.548899638 | 20.88699193 | 14.56058158 |
| AP53_1175 | 291.0867977 | 334.7336058 | 312.0896881 | 342.3497541 | 329.4596618 | 335.1282244 |
| AP53_1176 | 983.4919545 | 802.6419676 | 769.0935435 | 1694.802467 | 1819.666635 | 1172.361665 |
| AP53_1177 | 1019.220808 | 974.4976901 | 1075.163175 | 1418.646931 | 1710.759417 | 1632.321293 |
| AP53_1178 | 113.9888179 | 113.0618922 | 120.8463949 | 151.7933502 | 154.1271679 | 155.5541349 |
| AP53_1179 | 178.1420277 | 155.885863 | 190.2853066 | 195.369869 | 187.4824239 | 178.1856508 |
| AP53_118 | 182.0945833 | 235.4892548 | 268.6166557 | 183.1665333 | 182.8341303 | 212.1672903 |
| AP53_1180 | 82.29211529 | 58.93840272 | 57.99090773 | 102.3545669 | 101.2488084 | 71.87545697 |
| AP53_1181 | 11.90908522 | 11.27717139 | 14.36167132 | 17.67807931 | 18.18287244 | 16.37805128 |
| AP53_1184 | 14.51290417 | 17.44176349 | 25.48656196 | 10.50954939 | 12.30575443 | 9.723378741 |
| AP53_1185 | 2.576137782 | 2.229080616 | 3.59158866 | 3.551243667 | 4.628367531 | 2.177682289 |
| AP53_1186 | 3.662945909 | 3.536882964 | 4.374069849 | 3.619199564 | 5.25691127 | 4.224893417 |
| AP53_1187 | 3.496959881 | 4.907572646 | 3.451593941 | 4.368411565 | 4.790111558 | 4.678022438 |
| AP53_1188 | 3.590935177 | 3.522560881 | 3.764901065 | 3.545994632 | 5.086014473 | 4.117382569 |

| | | | | | | |
|---|---|---|---|---|---|---|
| AP53_1189 | 3.671967943 | 2.810371035 | 3.971783453 | 4.377826245 | 3.364876395 | 4.026834923 |
| AP53_119 | 55.19360151 | 55.78951298 | 54.35264257 | 43.82190433 | 43.29902709 | 38.51947644 |
| AP53_1190 | 4.101008142 | 3.723642301 | 4.266523484 | 4.113448612 | 5.030748932 | 4.148385346 |
| AP53_1191 | 6.684454286 | 6.660277163 | 7.065615831 | 10.92102891 | 8.485340473 | 6.352262047 |
| AP53_1192 | 3.401306916 | 3.622256001 | 5.836331572 | 2.693026448 | 4.242670237 | 5.204020176 |
| AP53_1193 | 4.235027024 | 3.678560498 | 6.887618363 | 6.490373141 | 5.099997383 | 7.118452469 |
| AP53_1194 | 3.597536161 | 2.328593143 | 2.023098129 | 2.871645549 | 0.909143622 | 2.943988557 |
| AP53_1195 | 8.497037789 | 8.624419049 | 9.35120913 | 6.045569576 | 9.293468137 | 8.286782604 |
| AP53_1196 | 15.71846569 | 17.55400985 | 13.07232637 | 15.29994147 | 13.38072921 | 7.397714835 |
| AP53_1197 | 11.94653517 | 8.748089964 | 6.982869517 | 13.41432042 | 15.36967331 | 11.4056412 |
| AP53_1198 | 15.21484299 | 13.89590606 | 17.31424956 | 13.75985777 | 15.38278097 | 15.74718246 |
| AP53_1199 | 6.845041776 | 8.484563605 | 10.47879776 | 9.15005125 | 9.784897122 | 4.125709689 |
| AP53_12 | 129.8802799 | 115.5406786 | 110.9871319 | 164.0674574 | 154.4371525 | 129.8496439 |
| AP53_120 | 59.35557113 | 56.67445821 | 66.87769943 | 54.05663711 | 52.92206352 | 58.52470738 |
| AP53_1200 | 10.66601367 | 10.45533135 | 14.93758334 | 11.49870026 | 13.0398328 | 10.6889879 |
| AP53_1201 | 10.98883773 | 12.95230934 | 18.42449771 | 13.59337159 | 10.92809 | 14.84880424 |
| AP53_1202 | 10.25703543 | 11.05741305 | 16.01441887 | 10.37292464 | 9.888930627 | 10.67411056 |
| AP53_1203 | 8.762953752 | 8.866417673 | 10.82206522 | 8.337472072 | 10.13174982 | 9.842588608 |
| AP53_1204 | 8.941449098 | 6.752166526 | 10.21719116 | 5.806877185 | 6.370870517 | 6.995280972 |
| AP53_1205 | 9.628314961 | 5.663891201 | 9.887432311 | 7.848248504 | 9.102456144 | 5.661973952 |
| AP53_1206 | 11.1784312 | 7.769476639 | 13.79225159 | 9.567770981 | 8.608316422 | 6.637011239 |
| AP53_1207 | 9.062729392 | 10.48142162 | 8.356399648 | 10.08452457 | 5.41617477 | 9.353949457 |
| AP53_1208 | 7.084378901 | 7.058755284 | 4.647938266 | 8.24677696 | 8.920486139 | 5.63635416 |
| AP53_1209 | 7.228537774 | 4.043448559 | 8.431143831 | 7.348413252 | 8.551118306 | 4.828039804 |
| AP53_121 | 76.69947094 | 75.88626322 | 89.59627248 | 60.62194892 | 56.74571442 | 63.08313258 |
| AP53_1210 | 3.156547857 | 4.20649084 | 5.847406205 | 4.91446301 | 3.558368586 | 4.727264779 |
| AP53_1211 | 2.988722349 | 3.622256001 | 3.776449841 | 3.41807203 | 3.426772114 | 4.051129553 |
| AP53_1212 | 13.45782541 | 14.28495324 | 18.08440769 | 13.58975922 | 18.88287035 | 12.3841021 |
| AP53_1213 | 135.6328236 | 110.910029 | 131.7561389 | 206.8928255 | 209.5070969 | 183.908421 |
| AP53_1214 | 163.4986607 | 142.6289473 | 141.3658334 | 257.8278169 | 261.721253 | 239.3007012 |
| AP53_1215 | 68.74032349 | 59.62936913 | 70.61267367 | 97.43363629 | 114.7414019 | 90.77340978 |
| AP53_1216 | 2397.523229 | 3072.145557 | 2037.148399 | 1552.186846 | 1323.713114 | 1323.088191 |
| AP53_1217 | 330.6209151 | 295.1769023 | 320.3816732 | 384.7573209 | 351.5355339 | 337.7642744 |
| AP53_1218 | 442.786287 | 449.2628688 | 491.2341444 | 476.6834796 | 474.0919766 | 461.9955028 |
| AP53_1219 | 419.5841893 | 408.498645 | 434.2917317 | 414.0704266 | 380.1073997 | 409.3203746 |

| | | | | | | |
|---|---|---|---|---|---|---|
| AP53_122 | 88.3185903 | 65.38636473 | 96.52218465 | 57.72743872 | 54.74132469 | 57.30293396 |
| AP53_1220 | 411.1682103 | 429.3424984 | 433.5851701 | 431.1076163 | 413.4276855 | 453.8765096 |
| AP53_1221 | 352.1793089 | 348.5211803 | 349.6692618 | 346.3434787 | 333.7465143 | 377.9358595 |
| AP53_1222 | 281.4235676 | 206.7223807 | 191.6608429 | 254.8842939 | 211.9713716 | 208.6169046 |
| AP53_1223 | 447.0121163 | 377.607064 | 309.8330804 | 442.6320716 | 387.312751 | 453.7060883 |
| AP53_1224 | 318.6205497 | 297.2490491 | 268.7378808 | 343.8454053 | 299.2321785 | 349.8703896 |
| AP53_1225 | 227.0445044 | 225.9885272 | 147.071741 | 249.917129 | 215.6690704 | 233.5564255 |
| AP53_1226 | 269.4144127 | 274.7728333 | 283.2590834 | 317.0799807 | 317.3175645 | 346.6515425 |
| AP53_1227 | 180.921987 | 170.055387 | 169.0259445 | 178.1852086 | 181.318328 | 188.0020763 |
| AP53_1228 | 1207.34587 | 1010.026017 | 968.1079557 | 1437.766987 | 1061.793932 | 1355.089274 |
| AP53_1229 | 108.6057895 | 99.6351608 | 101.0160157 | 118.4571475 | 115.4006304 | 110.2207044 |
| AP53_123 | 74.63861209 | 73.70692691 | 89.09246231 | 59.06866418 | 49.43411567 | 60.40038985 |
| AP53_1230 | 191.758956 | 183.6262022 | 175.5972106 | 248.7320054 | 271.1152866 | 279.1835751 |
| AP53_1231 | 129.699819 | 122.7454435 | 117.1524003 | 144.4389531 | 157.1084874 | 142.1456522 |
| AP53_1232 | 247.8842578 | 286.5816046 | 154.3930402 | 197.8550043 | 184.4735059 | 204.1165399 |
| AP53_1233 | 4525.925513 | 3930.980737 | 4132.478247 | 7123.492248 | 6810.14917 | 7163.45618 |
| AP53_1234 | 400.7276595 | 419.1331483 | 420.974777 | 619.3049655 | 679.2738347 | 629.5659272 |
| AP53_1235 | 961.5466155 | 793.1664724 | 715.5063778 | 1337.78279 | 1197.693206 | 1154.950353 |
| AP53_1236 | 263.5584265 | 346.8058037 | 343.6003292 | 165.1074826 | 170.0520375 | 186.986351 |
| AP53_1237 | 789.4672239 | 968.9320044 | 916.4981037 | 510.5710515 | 509.0533506 | 526.9545578 |
| AP53_1238 | 545.7789404 | 561.9436241 | 589.6983646 | 427.22941 | 419.702939 | 405.2890913 |
| AP53_1239 | 426.2191547 | 351.6571355 | 368.5815045 | 294.3410016 | 249.3691824 | 274.34125 |
| AP53_124 | 77.89506334 | 65.53913661 | 82.72895726 | 73.56384662 | 46.94693047 | 61.97355766 |
| AP53_1240 | 1011.489181 | 1084.692002 | 1106.723976 | 793.4326975 | 678.8272379 | 743.265251 |
| AP53_1241 | 747.1568672 | 829.0438422 | 890.3654866 | 624.9607549 | 560.4112811 | 623.3895769 |
| AP53_1242 | 963.642577 | 1047.029562 | 1070.692193 | 588.4647506 | 470.7821173 | 557.7044342 |
| AP53_1243 | 637.2145808 | 662.6419791 | 584.544636 | 392.6457926 | 319.272591 | 402.4759467 |
| AP53_1244 | 568.8127357 | 536.8883255 | 525.9026859 | 533.8600986 | 512.4879787 | 507.5993127 |
| AP53_1245 | 529.4297648 | 508.8416795 | 510.2496328 | 508.0154078 | 463.8496091 | 503.2670965 |
| AP53_1246 | 352.2504104 | 350.7751131 | 368.5748352 | 328.6868079 | 323.304584 | 348.8131804 |
| AP53_1247 | 282.046835 | 323.5882027 | 338.6216691 | 275.0734157 | 291.127991 | 310.100128 |
| AP53_1248 | 804.2731526 | 1134.155491 | 1093.07926 | 621.3309516 | 552.6527126 | 630.4248359 |
| AP53_1249 | 278.1504266 | 204.2952384 | 127.3535085 | 278.2636163 | 240.2004072 | 222.3541716 |
| AP53_125 | 72.10219842 | 73.20770023 | 91.42983825 | 66.81945321 | 54.48481778 | 70.38026445 |
| AP53_1250 | 370.1997951 | 384.0038553 | 321.6509562 | 347.433659 | 386.1877488 | 405.1477476 |

| | | | | | | |
|---|---|---|---|---|---|---|
| AP53_1251 | 87.01805113 | 72.6581939 | 69.53110589 | 88.12306711 | 109.3111508 | 87.28060192 |
| AP53_1252 | 6786.588443 | 7279.993646 | 5918.81267 | 8148.809492 | 8179.386095 | 7341.519423 |
| AP53_1253 | 27.48586479 | 25.37312337 | 28.11557872 | 34.01717618 | 32.8857693 | 35.75983548 |
| AP53_1254 | 374.7833227 | 380.5381165 | 355.6996145 | 391.2625457 | 380.3318163 | 382.1369838 |
| AP53_1255 | 461.8749807 | 353.8261659 | 345.1711642 | 437.3370872 | 392.5392289 | 398.0436874 |
| AP53_1256 | 192.8279382 | 189.0817632 | 224.3211206 | 221.751492 | 214.254847 | 233.5564255 |
| AP53_1257 | 166.5372322 | 142.9791111 | 145.9609448 | 167.4585683 | 166.7916005 | 167.7290822 |
| AP53_1258 | 355.5447067 | 321.0198208 | 285.7440949 | 327.8376094 | 317.2928758 | 335.2856948 |
| AP53_1259 | 360.8767493 | 380.1601847 | 383.9083643 | 370.239528 | 398.7075225 | 352.177623 |
| AP53_126 | 87.17705194 | 88.01103094 | 94.84502742 | 69.42601386 | 62.95205297 | 81.68905185 |
| AP53_1260 | 390.0714824 | 422.3649737 | 444.5864374 | 427.2478554 | 429.7301881 | 485.9328692 |
| AP53_1261 | 49.3177646 | 61.76438933 | 51.88910537 | 46.12897438 | 47.2205352 | 52.07667707 |
| AP53_1262 | 2760.365664 | 2581.697261 | 2025.034857 | 1505.220955 | 700.4554874 | 1187.559374 |
| AP53_1263 | 1405.852311 | 1314.030472 | 1127.355333 | 1176.810966 | 550.6297967 | 918.3417197 |
| AP53_1264 | 1758.140483 | 1649.362993 | 1318.29949 | 1007.939697 | 527.5564493 | 888.0992736 |
| AP53_1265 | 6886.390946 | 7650.504742 | 5836.514402 | 3307.291439 | 1751.544985 | 2883.699697 |
| AP53_1266 | 2724.937986 | 3093.660818 | 2041.535533 | 1685.929048 | 914.9281149 | 1501.445642 |
| AP53_1267 | 4658.299386 | 4679.075565 | 3206.755883 | 1840.382899 | 984.0111581 | 1467.897466 |
| AP53_1268 | 49.80131115 | 44.6798097 | 40.39304059 | 42.43084339 | 34.16564402 | 38.73441625 |
| AP53_1269 | 179.8950234 | 155.4360486 | 150.6755683 | 166.4903946 | 119.761451 | 151.6464654 |
| AP53_127 | 110.3894825 | 127.2394163 | 136.6562781 | 89.2643718 | 73.06021967 | 96.44196035 |
| AP53_1270 | 349.2650671 | 330.8832343 | 273.6109014 | 351.5840195 | 244.0511733 | 325.7411792 |
| AP53_1271 | 196.6972882 | 188.5183012 | 213.8729427 | 230.7325215 | 231.1783871 | 280.5730131 |
| AP53_1272 | 163.8385188 | 160.0059519 | 196.0095073 | 191.9144097 | 187.0004618 | 211.6411981 |
| AP53_1273 | 121.1335577 | 124.4765544 | 143.3522012 | 137.8194055 | 130.8185284 | 148.399272 |
| AP53_1274 | 89.19549817 | 71.65980706 | 93.00914947 | 116.9793137 | 96.94903912 | 125.3477814 |
| AP53_1275 | 70.67774888 | 57.7471197 | 75.16587664 | 78.37150868 | 79.3052975 | 92.73563954 |
| AP53_1276 | 45.27127361 | 41.75528858 | 48.82296334 | 51.9970001 | 50.5540116 | 58.20070767 |
| AP53_1277 | 1036.978809 | 947.0510609 | 1160.183677 | 1299.451442 | 1214.958879 | 1535.897447 |
| AP53_1278 | 648.9235727 | 634.9814769 | 632.8385821 | 600.9825146 | 556.9565353 | 780.6966988 |
| AP53_1279 | 83.6399484 | 85.93105774 | 108.3550608 | 87.89328076 | 90.53205567 | 81.51577204 |
| AP53_128 | 93.30872737 | 102.9483284 | 102.6327038 | 69.41123806 | 64.715946 | 85.54313587 |
| AP53_1280 | 747.7920963 | 922.4241046 | 888.419448 | 832.6536275 | 901.5409987 | 967.8954179 |
| AP53_1281 | 468.2001955 | 456.4813254 | 476.5558299 | 582.7414555 | 624.3946814 | 591.6373032 |
| AP53_1282 | 421.5205446 | 286.4368591 | 175.2660054 | 317.2838925 | 283.2798281 | 289.3328469 |

| | | | | | | |
|---|---|---|---|---|---|---|
| AP53_1283 | 281.872409 | 278.2551201 | 253.5943287 | 248.0149119 | 262.1455943 | 283.7925669 |
| AP53_1284 | 176.6093987 | 142.3333534 | 104.4965885 | 121.6795345 | 125.7826941 | 113.141521 |
| AP53_1285 | 365.9109927 | 301.0368931 | 348.6370347 | 332.5496814 | 288.6368006 | 351.5479421 |
| AP53_1286 | 707.9629597 | 618.6975138 | 777.4001326 | 790.2066048 | 646.8768497 | 883.3135225 |
| AP53_1287 | 412.4046159 | 408.9598049 | 456.4320945 | 538.6958114 | 452.7178711 | 511.6511006 |
| AP53_1288 | 603.8257508 | 593.6653198 | 628.1606302 | 681.8974477 | 587.5159634 | 657.1839313 |
| AP53_1289 | 252.7640327 | 280.0636346 | 275.1413456 | 229.8659899 | 215.2313345 | 254.6368374 |
| AP53_129 | 334.5623177 | 311.2903138 | 343.2353366 | 434.327504 | 373.7177745 | 424.9724015 |
| AP53_1290 | 102.548198 | 113.0935082 | 116.7219446 | 115.8933624 | 102.3453694 | 131.2279829 |
| AP53_1291 | 69.83073527 | 61.97587791 | 72.37256113 | 110.809611 | 83.16974631 | 106.7245392 |
| AP53_1292 | 56.58379299 | 42.44259556 | 55.6625819 | 79.67938568 | 59.99735688 | 69.05387839 |
| AP53_1293 | 57.36036786 | 48.93510368 | 54.93256777 | 80.13376188 | 62.24550629 | 77.66896033 |
| AP53_1294 | 44.85165875 | 30.84297189 | 44.71914861 | 59.66558172 | 46.20729961 | 54.41034957 |
| AP53_1295 | 118.0588962 | 92.34498286 | 113.4501944 | 229.3653978 | 146.1168588 | 204.8440539 |
| AP53_1296 | 63.26075237 | 54.58001275 | 62.4764129 | 98.82647587 | 78.64729811 | 98.40818995 |
| AP53_1297 | 73.72111956 | 52.01422497 | 69.71540842 | 92.59447762 | 74.46756213 | 96.61813409 |
| AP53_1298 | 153.27223 | 111.6099904 | 129.9717413 | 202.3223107 | 148.6237792 | 227.0700495 |
| AP53_1299 | 127.7564061 | 110.2579388 | 118.267356 | 170.9274208 | 137.4211238 | 180.5008052 |
| AP53_13 | 710.6346347 | 586.1250483 | 556.995055 | 860.688702 | 892.7890827 | 677.7715878 |
| AP53_130 | 219.3588465 | 192.8573161 | 225.1949885 | 245.8256516 | 228.0326651 | 230.9462453 |
| AP53_1300 | 465.8218465 | 358.4602612 | 451.5095165 | 824.8067352 | 533.3945001 | 925.1999187 |
| AP53_1301 | 326.88296 | 218.4637088 | 251.3979317 | 477.9274203 | 309.2268325 | 449.4917624 |
| AP53_1302 | 253.4656025 | 196.6665794 | 231.0273119 | 336.2591096 | 237.9807881 | 356.5968592 |
| AP53_1303 | 390.8215694 | 283.4309777 | 348.0319835 | 575.4593071 | 389.0597835 | 556.4191747 |
| AP53_1304 | 616.5108088 | 400.8255166 | 530.2604493 | 1159.22121 | 710.6613755 | 1098.825052 |
| AP53_1305 | 156.8653112 | 151.6539216 | 163.3565206 | 207.063433 | 217.5401005 | 177.3051122 |
| AP53_1306 | 69.07269428 | 71.22701623 | 64.33332694 | 60.93185124 | 53.05215084 | 62.12771131 |
| AP53_1307 | 20.00466009 | 13.53892407 | 16.83925175 | 20.53511502 | 20.86559133 | 21.02082904 |
| AP53_1308 | 148.2346585 | 148.3632645 | 123.109436 | 175.4890728 | 144.6003714 | 146.8882447 |
| AP53_1309 | 59.78360781 | 62.25284106 | 66.4655172 | 93.97733672 | 92.34391219 | 77.06256636 |
| AP53_131 | 1127.706556 | 1234.96238 | 1223.604797 | 1159.559882 | 1146.721905 | 1266.970028 |
| AP53_1310 | 283.5007873 | 281.4079299 | 313.8242885 | 367.4283815 | 365.0751676 | 329.8534876 |
| AP53_1311 | 242.9385333 | 235.9641052 | 263.8767352 | 260.7816892 | 237.7834839 | 240.8575171 |
| AP53_1312 | 55.21233606 | 69.88611329 | 84.32192481 | 58.32019606 | 81.98274228 | 75.82438631 |
| AP53_1313 | 12.04227894 | 15.9188619 | 31.50354209 | 6.013750789 | 7.703795956 | 8.435990602 |

| | | | | | | |
|---|---|---|---|---|---|---|
| AP53_1314 | 17.45913284 | 20.8751815 | 26.20247662 | 8.102782351 | 12.1448949 | 7.726613464 |
| AP53_1315 | 11.59206096 | 12.47665956 | 19.6165589 | 4.70210967 | 5.892597551 | 5.342794048 |
| AP53_1316 | 12.92138902 | 14.37384893 | 22.02228619 | 7.131657592 | 9.60502769 | 8.940655585 |
| AP53_1317 | 15.74033105 | 15.15981215 | 18.69575785 | 8.777271384 | 11.62805917 | 9.441763026 |
| AP53_1318 | 15.69833961 | 22.6555648 | 31.12005939 | 8.665034548 | 13.1018404 | 11.83634374 |
| AP53_1319 | 13.05348686 | 15.33775344 | 22.05825932 | 8.215903759 | 10.15400575 | 8.967463052 |
| AP53_132 | 1342.319308 | 1299.224121 | 1315.116101 | 1409.124983 | 1445.141262 | 1446.50227 |
| AP53_1320 | 10.65426716 | 13.80922619 | 24.16649963 | 5.559476861 | 7.369053732 | 7.650819285 |
| AP53_1321 | 6.349465884 | 7.617940455 | 15.49512409 | 4.362782168 | 4.723797789 | 5.948677909 |
| AP53_1322 | 7.304238936 | 9.193035919 | 10.20943681 | 3.502260996 | 4.681567158 | 4.800618888 |
| AP53_1323 | 7.624907811 | 5.83324343 | 5.002569919 | 4.067019533 | 4.518168304 | 2.260031619 |
| AP53_1324 | 10.47342858 | 14.489024 | 18.74027741 | 5.981932002 | 8.421540921 | 9.22794074 |
| AP53_1325 | 7.255535114 | 12.90618945 | 8.50494586 | 4.267460877 | 6.132262863 | 4.156219139 |
| AP53_1326 | 14.31236008 | 19.97135741 | 38.92124881 | 6.557536729 | 11.16089828 | 7.921362603 |
| AP53_1327 | 13.58264012 | 14.69749917 | 25.42990684 | 8.159344262 | 7.691747479 | 7.511759771 |
| AP53_1328 | 14.71695379 | 18.20415836 | 29.82427054 | 8.308782802 | 7.273148977 | 8.991326875 |
| AP53_1329 | 16.21903987 | 19.27111567 | 23.96092313 | 10.25662149 | 10.78432435 | 6.406840997 |
| AP53_133 | 1247.081913 | 1314.55695 | 1365.648184 | 1647.619228 | 1895.436499 | 1784.493212 |
| AP53_1330 | 14.8752057 | 12.53544773 | 10.18368496 | 17.3079902 | 16.97068095 | 17.90650718 |
| AP53_1331 | 115.8785332 | 124.8725095 | 109.5170454 | 157.4711781 | 185.1144014 | 158.1145667 |
| AP53_1332 | 28.41042211 | 27.14509919 | 27.3906362 | 28.41660495 | 31.93077921 | 28.43359586 |
| AP53_1333 | 6.847525914 | 9.543668505 | 11.75900549 | 4.696478401 | 6.469860401 | 5.923234462 |
| AP53_1334 | 6.525431927 | 8.714932754 | 13.4232227 | 4.525198613 | 5.670895861 | 5.849112578 |
| AP53_1335 | 5.357840018 | 5.923185913 | 13.87073401 | 6.600697348 | 6.404030546 | 5.299590861 |
| AP53_1338 | 8.336359655 | 7.794095671 | 10.41779266 | 5.603617593 | 4.915645516 | 5.432279268 |
| AP53_1339 | 26.58750534 | 24.83832686 | 19.06208015 | 14.91240495 | 10.1015958 | 7.414489699 |
| AP53_134 | 418.9187594 | 414.604063 | 471.1204726 | 466.0331634 | 456.7816469 | 438.0550761 |
| AP53_1340 | 17.48402574 | 15.4851444 | 20.29841789 | 17.24498722 | 22.16795199 | 16.94428969 |
| AP53_1341 | 22.53319332 | 20.8751815 | 22.69449478 | 22.78406126 | 24.53107909 | 16.49501751 |
| AP53_1342 | 27.87143804 | 29.23224141 | 29.9465847 | 25.24290455 | 31.55951194 | 32.13710705 |
| AP53_1343 | 26.66730602 | 16.68988841 | 28.49068488 | 22.92724867 | 25.1337933 | 23.88315893 |
| AP53_1344 | 26.94881633 | 19.099168 | 23.68863991 | 24.23723803 | 29.31299436 | 21.23240232 |
| AP53_1345 | 20.31549832 | 17.89820612 | 25.2609586 | 17.78108699 | 20.96378235 | 21.08896378 |
| AP53_1346 | 19.67883028 | 15.4797265 | 22.98147254 | 18.08503719 | 18.56621505 | 19.10097799 |
| AP53_1347 | 16.94388862 | 19.79486378 | 27.55212701 | 17.04680323 | 21.87066629 | 21.28517548 |

| | | | | | | |
|---|---|---|---|---|---|---|
| AP53_1348 | 21.54044762 | 23.82639503 | 30.54728316 | 22.94629519 | 23.53267758 | 27.27369152 |
| AP53_1349 | 9.581512458 | 8.999393791 | 12.6663535 | 10.4088937 | 11.59487518 | 7.736858816 |
| AP53_135 | 424.206953 | 357.9641224 | 448.2233408 | 514.3712844 | 525.9484986 | 458.5695117 |
| AP53_1350 | 13.54366555 | 16.19361506 | 14.92808408 | 16.33015029 | 15.9724056 | 13.79249018 |
| AP53_1351 | 13.17365819 | 15.38525229 | 16.81882816 | 13.61188037 | 17.49554737 | 12.84159041 |
| AP53_1352 | 15.51632988 | 19.52868453 | 22.65869905 | 17.6635772 | 25.08709357 | 22.69857844 |
| AP53_1353 | 20.42187577 | 19.81832019 | 25.58146865 | 21.20813603 | 23.01770519 | 26.88065839 |
| AP53_1354 | 12.01067191 | 10.6100727 | 12.66746167 | 9.396814443 | 11.95965311 | 11.32263665 |
| AP53_1355 | 15.01580311 | 14.17404522 | 20.0316035 | 14.71964766 | 18.44639233 | 23.49501979 |
| AP53_1356 | 15.62358561 | 12.59165181 | 20.32089676 | 16.12151887 | 17.37474478 | 18.75429747 |
| AP53_1357 | 17.08249429 | 16.12488155 | 20.46592172 | 20.47692921 | 30.93043463 | 20.38632936 |
| AP53_1358 | 8551.487355 | 8031.447118 | 6396.550741 | 10943.90164 | 10759.62385 | 7586.233268 |
| AP53_1359 | 1500.028678 | 1268.936648 | 1515.552262 | 2240.367173 | 2386.996986 | 2338.693606 |
| AP53_136 | 652.7269505 | 606.7751269 | 614.3725633 | 698.5061602 | 653.0318028 | 635.8150575 |
| AP53_1360 | 586.307189 | 527.5229162 | 579.1265615 | 846.1413238 | 1009.636004 | 620.4951156 |
| AP53_1361 | 481.590174 | 437.7210409 | 431.0453099 | 782.3815532 | 934.1317802 | 821.5851418 |
| AP53_1362 | 592.02121 | 487.9312991 | 493.1763755 | 889.7958866 | 994.1990588 | 827.0305961 |
| AP53_1363 | 659.3236399 | 517.1114326 | 571.9807742 | 1145.862462 | 1229.765716 | 1047.728676 |
| AP53_1364 | 359.4150244 | 306.5778045 | 368.7962438 | 505.6289166 | 571.886991 | 498.0022822 |
| AP53_1365 | 684.9486608 | 672.0893219 | 757.5645866 | 1019.969284 | 1068.038994 | 945.5419343 |
| AP53_1366 | 706.119547 | 651.19209 | 562.054546 | 852.6562646 | 823.1733668 | 762.9818633 |
| AP53_1367 | 1328.611387 | 1110.033424 | 1172.721812 | 1205.466752 | 856.0456602 | 976.1672609 |
| AP53_1368 | 18.0978756 | 16.64279784 | 19.73280097 | 11.43756406 | 16.66490291 | 14.77004069 |
| AP53_1369 | 26.69664843 | 27.33901814 | 30.14324669 | 23.51480365 | 25.57120704 | 25.1978186 |
| AP53_137 | 71.56180038 | 47.38302444 | 54.50335716 | 85.78173047 | 97.46674868 | 62.87581566 |
| AP53_1370 | 182.1751392 | 172.6322576 | 142.8965665 | 141.1167738 | 140.7520932 | 126.9248189 |
| AP53_1371 | 1926.006289 | 2222.127162 | 2064.346995 | 1898.15482 | 1836.616815 | 1925.506027 |
| AP53_1372 | 501.6906935 | 500.3312971 | 572.9413901 | 597.7053121 | 494.0353787 | 495.4623705 |
| AP53_1373 | 485.8507251 | 577.2960806 | 544.5203993 | 460.3798324 | 401.0095594 | 468.2307682 |
| AP53_1374 | 243.9062273 | 238.6626617 | 248.8221624 | 267.9938842 | 273.5927536 | 293.775207 |
| AP53_1375 | 134.8807587 | 120.7562836 | 123.1911757 | 154.6222093 | 161.4494334 | 144.1392421 |
| AP53_1376 | 95.08708564 | 89.25489678 | 92.98894673 | 111.0186413 | 122.8354049 | 96.90381189 |
| AP53_1377 | 1424.428814 | 1411.295028 | 1367.635642 | 1754.479685 | 1540.900296 | 1680.9291 |
| AP53_1378 | 1132.148999 | 1132.431613 | 1094.895247 | 1235.851487 | 1199.353063 | 1277.079193 |
| AP53_1379 | 530.9930742 | 529.3586275 | 586.0776125 | 648.1618277 | 642.3075639 | 671.0009333 |

| | | | | | | |
|---|---|---|---|---|---|---|
| AP53_138 | 1380.631592 | 1648.385213 | 813.2854479 | 1146.54227 | 1071.577283 | 849.8313634 |
| AP53_1380 | 299.6470888 | 291.2200946 | 288.9468289 | 370.3996468 | 327.936652 | 341.4691229 |
| AP53_1381 | 185.2364581 | 163.8079846 | 164.8337226 | 226.1697008 | 249.5864171 | 227.0068225 |
| AP53_1382 | 698.7306042 | 601.2255008 | 564.2375724 | 793.5011588 | 773.216549 | 640.61191 |
| AP53_1383 | 586.8463893 | 668.4087488 | 703.7022383 | 677.5828754 | 787.8558579 | 758.2159208 |
| AP53_1384 | 933.7700425 | 1061.861643 | 1042.412886 | 933.9863601 | 962.5795562 | 1079.98487 |
| AP53_1385 | 87.63068684 | 75.49799129 | 89.37176948 | 106.1513053 | 112.8351629 | 106.0492288 |
| AP53_1386 | 151.8669002 | 153.5251129 | 179.9726298 | 154.3146901 | 173.1780851 | 164.9096171 |
| AP53_1387 | 179.8967022 | 171.3310396 | 207.0260245 | 192.0540941 | 217.5688221 | 218.7204115 |
| AP53_1388 | 177.4958932 | 161.025744 | 187.3119121 | 181.1252645 | 194.5457152 | 202.0825115 |
| AP53_139 | 105.5235833 | 107.2438127 | 95.88756875 | 129.6441711 | 157.8309982 | 155.6053736 |
| AP53_1391 | 167.3146007 | 150.601004 | 158.1686064 | 173.355346 | 201.9663922 | 198.8179499 |
| AP53_1392 | 259.4873784 | 292.4802982 | 312.8120191 | 264.0073949 | 264.57397 | 293.5455257 |
| AP53_1393 | 1351.827808 | 1376.102447 | 1355.976704 | 1581.417363 | 1707.181235 | 1721.532356 |
| AP53_1394 | 1059.892087 | 1062.779319 | 1016.600679 | 1138.14992 | 1293.694844 | 1176.310773 |
| AP53_1395 | 307.1305734 | 215.2085308 | 110.0367403 | 200.0251571 | 157.0955695 | 125.0339849 |
| AP53_1396 | 30.11890739 | 29.98891015 | 22.04392814 | 20.66741227 | 16.77334745 | 18.4246519 |
| AP53_1397 | 422.8859452 | 433.6001026 | 430.4036627 | 476.7244313 | 417.2446729 | 404.172406 |
| AP53_1398 | 936.514583 | 957.4946607 | 983.418792 | 1143.55826 | 976.0599367 | 1102.222097 |
| AP53_1399 | 1409.537253 | 1523.654069 | 1353.086737 | 1441.654173 | 1143.770625 | 1266.129241 |
| AP53_14 | 1435.491059 | 1658.29075 | 1618.540627 | 1609.557788 | 1667.600821 | 1780.024693 |
| AP53_140 | 770.1798662 | 874.2232336 | 927.9380252 | 975.1634206 | 1021.069304 | 1124.589509 |
| AP53_1400 | 236.5635124 | 239.5167386 | 241.4730691 | 233.9085829 | 213.1517527 | 240.8004216 |
| AP53_1401 | 226.4600997 | 244.647319 | 238.4669107 | 209.0053955 | 210.0252445 | 203.117075 |
| AP53_1402 | 253.7420461 | 304.8679637 | 292.3300564 | 215.3116462 | 214.3839717 | 231.4060339 |
| AP53_1403 | 269.8951573 | 311.8928141 | 319.5419584 | 244.5650218 | 253.0073414 | 272.0185563 |
| AP53_1404 | 69.37968404 | 61.81983575 | 70.29231971 | 83.73516901 | 82.13809578 | 67.16655374 |
| AP53_1405 | 519.0664432 | 472.5875611 | 486.6138351 | 683.042102 | 604.5805087 | 637.8114277 |
| AP53_1406 | 460.0980277 | 456.7827008 | 507.0024824 | 614.2569386 | 560.4124119 | 588.914885 |
| AP53_1407 | 96.93201431 | 102.2925095 | 100.0003918 | 93.44032337 | 91.98109073 | 102.5816457 |
| AP53_1408 | 362.6059483 | 426.9734261 | 406.4404141 | 221.9479731 | 193.3824246 | 235.7644169 |
| AP53_1409 | 535.0571763 | 511.6732186 | 530.9441943 | 612.8593006 | 603.3404404 | 665.9762205 |
| AP53_141 | 34.27142398 | 33.19138344 | 37.89090677 | 36.36953948 | 41.39710038 | 40.31526124 |
| AP53_1410 | 554.4951373 | 542.0860244 | 543.7485146 | 583.529309 | 599.5931145 | 607.6656853 |
| AP53_1411 | 382.1382855 | 410.1198739 | 423.8015933 | 499.520784 | 547.7758683 | 480.8514643 |

| | | | | | | |
|---|---|---|---|---|---|---|
| AP53_1412 | 435.0081121 | 476.20878 | 462.4251657 | 446.7783234 | 487.0284177 | 517.2980813 |
| AP53_1413 | 199.543339 | 114.3870316 | 82.94938949 | 186.7974936 | 144.1019224 | 140.1177868 |
| AP53_1414 | 370.2925853 | 334.4289113 | 334.682207 | 421.606714 | 406.8079778 | 460.8530512 |
| AP53_1415 | 306.7789078 | 264.1991263 | 277.0503202 | 299.054175 | 316.3508987 | 327.5795358 |
| AP53_1416 | 830.8652207 | 712.2302554 | 832.2099226 | 1055.776894 | 1027.14812 | 1115.821663 |
| AP53_1417 | 481.441378 | 446.9716058 | 461.9908352 | 496.4967197 | 508.4277475 | 559.6382057 |
| AP53_1418 | 355.3233918 | 259.8256439 | 218.1005863 | 266.2767723 | 273.8190768 | 266.7452104 |
| AP53_1419 | 811.5345982 | 833.4034078 | 780.0935995 | 899.3360078 | 743.4798925 | 829.3529039 |
| AP53_142 | 32.18440952 | 27.26429248 | 25.98847202 | 25.84643509 | 24.81733988 | 25.80298692 |
| AP53_1420 | 142.2147876 | 115.8448012 | 103.7733287 | 154.8678093 | 106.4811563 | 131.4646838 |
| AP53_1421 | 101.1914971 | 89.42625615 | 83.38401249 | 134.9467858 | 97.78506361 | 122.7316118 |
| AP53_1422 | 102.6603816 | 86.4501009 | 86.5976115 | 128.1445318 | 103.6761199 | 122.4745866 |
| AP53_1423 | 203.9218788 | 227.4900606 | 226.264217 | 322.9987643 | 298.5099264 | 376.2266402 |
| AP53_1424 | 182.0529359 | 162.9483385 | 174.7639952 | 240.3276634 | 231.4989137 | 247.7283392 |
| AP53_1425 | 148.3247643 | 130.0252533 | 137.3090026 | 190.49457 | 184.2385902 | 209.7629389 |
| AP53_1426 | 65.47515812 | 72.74697468 | 82.13778404 | 84.63797406 | 102.8847532 | 98.07843359 |
| AP53_1427 | 2093.303058 | 2147.399349 | 2049.922651 | 2817.929349 | 2518.744195 | 2331.661692 |
| AP53_1428 | 250.7231713 | 292.1391584 | 229.5730206 | 144.4750534 | 134.5814466 | 151.6572503 |
| AP53_1429 | 235.151893 | 320.1008935 | 325.1079025 | 176.4950694 | 191.7936516 | 225.1785653 |
| AP53_143 | 28.85135173 | 30.67737798 | 32.9157233 | 29.46655393 | 27.65592451 | 35.05325195 |
| AP53_1430 | 238.2556497 | 332.3479068 | 347.1371932 | 167.0631906 | 176.4174382 | 194.8548417 |
| AP53_1431 | 951.6251158 | 886.1096365 | 918.7381119 | 1232.005566 | 1242.291982 | 1200.76509 |
| AP53_1432 | 1437.246532 | 1260.234609 | 1223.947393 | 1917.65467 | 1779.254678 | 2043.716856 |
| AP53_1433 | 1317.791585 | 1195.363395 | 1176.516958 | 1692.494297 | 1601.314461 | 1800.105091 |
| AP53_1434 | 172.2796581 | 160.5778046 | 164.4977122 | 164.7951142 | 172.7016355 | 169.4092239 |
| AP53_1435 | 444.7211506 | 413.2678465 | 407.1044268 | 532.7626832 | 517.7818133 | 502.266979 |
| AP53_1436 | 323.1586077 | 370.3724534 | 283.1934201 | 129.6979062 | 114.3256193 | 126.9884633 |
| AP53_1437 | 293.0638059 | 273.7334965 | 211.2375492 | 252.7308107 | 275.180504 | 233.0147598 |
| AP53_1438 | 336.6561257 | 339.3592858 | 368.4167139 | 403.8000799 | 394.5837599 | 419.3857412 |
| AP53_1439 | 336.0308339 | 317.9145073 | 337.6072829 | 343.6466093 | 374.0140364 | 414.0257823 |
| AP53_144 | 35.52607655 | 31.60049579 | 32.60391989 | 32.32328059 | 34.86284683 | 35.80742643 |
| AP53_1440 | 114.2577485 | 111.2032592 | 113.6711402 | 122.7250624 | 148.0691913 | 108.5350448 |
| AP53_1441 | 506.7096576 | 444.3152545 | 502.6153549 | 535.6077929 | 520.1565767 | 550.275009 |
| AP53_1442 | 493.3482751 | 443.4632047 | 490.357487 | 552.8234169 | 488.9586792 | 532.7920335 |
| AP53_1443 | 456.4687742 | 389.0976853 | 441.6689826 | 513.4375706 | 444.6910409 | 462.1073098 |

| | | | | | | |
|---|---|---|---|---|---|---|
| AP53_1444 | 572.9304349 | 500.8916819 | 549.6596149 | 599.7375317 | 525.3740383 | 580.891789 |
| AP53_1445 | 520.8815255 | 433.410805 | 446.4811423 | 512.9096275 | 425.7427351 | 490.3044646 |
| AP53_1446 | 430.9142921 | 377.4248704 | 403.7098928 | 389.3346807 | 362.8730897 | 392.5831189 |
| AP53_1447 | 280.5167437 | 258.509865 | 286.8189753 | 327.731092 | 294.1942729 | 316.0460739 |
| AP53_1448 | 249.5934193 | 215.0238815 | 238.0485892 | 318.9237902 | 269.0876454 | 269.141394 |
| AP53_1449 | 62.97804479 | 48.58084519 | 21.77012261 | 34.60199528 | 36.43705027 | 26.93845738 |
| AP53_145 | 32.53771594 | 28.77681156 | 30.68365496 | 26.15548504 | 33.35210214 | 34.47374255 |
| AP53_1450 | 136.7063741 | 136.6525288 | 146.7739565 | 142.9744702 | 134.0775035 | 167.005401 |
| AP53_1451 | 86.64381827 | 79.68963202 | 71.35502594 | 83.301585 | 77.70785486 | 79.65985411 |
| AP53_1452 | 468.357656 | 502.0072481 | 436.1397816 | 567.4603137 | 491.943417 | 470.7597777 |
| AP53_1453 | 1353.11637 | 1081.104099 | 1033.00428 | 1746.291191 | 1596.694495 | 1780.618218 |
| AP53_1454 | 2062.937348 | 2004.81618 | 1985.335893 | 2399.282511 | 2514.880584 | 2717.979659 |
| AP53_1455 | 928.7199885 | 834.1947978 | 897.9724691 | 1081.857668 | 1232.306674 | 1154.950353 |
| AP53_1456 | 866.4320625 | 882.9523415 | 1004.306782 | 1038.909758 | 1153.834883 | 1058.937154 |
| AP53_1457 | 1162.09738 | 1180.142266 | 1232.414077 | 1273.560313 | 1424.174269 | 1418.944859 |
| AP53_1458 | 582.1712892 | 546.5078741 | 538.1441023 | 552.5272744 | 613.9939333 | 545.9667667 |
| AP53_1459 | 50.92131607 | 45.76691709 | 49.47349104 | 56.96270001 | 60.74426275 | 51.85296717 |
| AP53_146 | 54.52333953 | 57.46285264 | 50.84547786 | 48.04555634 | 58.78355017 | 58.07243621 |
| AP53_1460 | 527.7653426 | 611.4778196 | 577.369303 | 478.2844006 | 537.6984147 | 524.6594953 |
| AP53_1461 | 502.6957195 | 478.5768534 | 549.301795 | 481.0899389 | 494.592497 | 483.5575548 |
| AP53_1462 | 195.5391249 | 211.8337306 | 200.7976692 | 139.6403908 | 147.7187098 | 143.0408662 |
| AP53_1463 | 250.3171607 | 302.1739841 | 292.3783645 | 186.4133892 | 201.4742412 | 192.9461444 |
| AP53_1464 | 454.0707355 | 494.3861976 | 511.5471056 | 338.9146304 | 331.6555934 | 360.4750433 |
| AP53_1465 | 517.3680239 | 513.6906913 | 550.2826911 | 463.1617539 | 432.6810588 | 465.4965436 |
| AP53_1466 | 871.3972261 | 839.8894521 | 887.0774795 | 934.4586043 | 956.8212662 | 986.9117859 |
| AP53_1467 | 611.2604527 | 543.4625917 | 560.8135912 | 578.440097 | 647.6981602 | 660.1861628 |
| AP53_1468 | 470.2774023 | 333.1534675 | 270.4330445 | 343.3883519 | 353.7395184 | 331.3920048 |
| AP53_1469 | 591.207421 | 553.5283195 | 589.2192294 | 529.1003044 | 598.3297574 | 648.7333689 |
| AP53_147 | 57.62348631 | 55.36971286 | 53.13169139 | 50.53611639 | 51.37572265 | 52.5187062 |
| AP53_1470 | 424.1216358 | 372.9186067 | 285.0035135 | 459.3562076 | 467.2886392 | 389.5479865 |
| AP53_1471 | 342.6158668 | 472.4034654 | 477.7266095 | 215.3537817 | 183.3961503 | 212.9554233 |
| AP53_1472 | 593.3003339 | 885.8159971 | 943.6089336 | 387.9553952 | 316.5974812 | 427.6270588 |
| AP53_1473 | 758.1334099 | 1132.669919 | 1205.830372 | 548.1979307 | 441.2377046 | 578.6486982 |
| AP53_1474 | 74.34908065 | 71.23770135 | 71.37490199 | 69.54420202 | 70.14548125 | 61.97641095 |
| AP53_1475 | 49.97720076 | 47.18515489 | 47.87499375 | 53.24579427 | 54.32413742 | 54.42138517 |

| | | | | | | |
|---|---|---|---|---|---|---|
| AP53_1476 | 229.107317 | 284.7467346 | 224.1402765 | 154.2159518 | 147.7963059 | 152.6082581 |
| AP53_1477 | 1240.352078 | 1487.295495 | 1497.355015 | 841.1763166 | 669.2523401 | 889.392562 |
| AP53_1478 | 1208.254304 | 1477.379098 | 1524.692621 | 665.389194 | 587.925459 | 747.1777309 |
| AP53_1479 | 1244.624167 | 1446.004595 | 1519.444219 | 661.4411169 | 576.3317846 | 768.1948124 |
| AP53_148 | 100.7310125 | 98.0795471 | 102.8356341 | 57.6189285 | 57.81220982 | 63.71093185 |
| AP53_1480 | 1022.292205 | 1175.768884 | 1155.754351 | 516.7118303 | 470.3646889 | 588.9090679 |
| AP53_1481 | 1443.219489 | 1602.208421 | 1667.517433 | 616.0833609 | 497.4340313 | 624.3527881 |
| AP53_1482 | 1364.339767 | 1742.243443 | 1702.508122 | 512.2347851 | 447.8855869 | 583.818638 |
| AP53_1483 | 907.8125535 | 1048.413949 | 946.3629161 | 403.3604021 | 354.5140616 | 421.9903851 |
| AP53_1484 | 769.4359768 | 953.9610436 | 1022.850836 | 409.5071819 | 355.4083502 | 449.58478 |
| AP53_1485 | 819.3499641 | 973.1794455 | 902.291775 | 395.9698799 | 351.3035713 | 454.7876753 |
| AP53_1486 | 928.6440009 | 1071.028654 | 873.5180245 | 526.335348 | 514.0984948 | 456.4883234 |
| AP53_1487 | 1159.748248 | 1523.175366 | 1309.045333 | 767.9398705 | 748.5897909 | 836.7731386 |
| AP53_1488 | 1186.462464 | 1334.788846 | 1266.074343 | 681.7734738 | 615.7431204 | 679.919233 |
| AP53_1489 | 1082.051664 | 1552.685523 | 1226.316257 | 595.3512039 | 549.2972299 | 636.555748 |
| AP53_149 | 3618.959083 | 3399.401013 | 3596.353882 | 4405.671707 | 4876.625123 | 4906.968966 |
| AP53_1490 | 9344.132213 | 8303.625249 | 8595.558551 | 16279.52252 | 17122.5663 | 17879.72372 |
| AP53_1491 | 22227.28679 | 18323.92844 | 20308.09776 | 39063.2185 | 37279.70941 | 38035.30947 |
| AP53_1492 | 13753.29181 | 11741.23758 | 11859.74978 | 23049.10761 | 23024.35166 | 23260.24409 |
| AP53_1493 | 8048.837517 | 7516.821658 | 7564.940536 | 12339.18624 | 13131.51939 | 11712.89124 |
| AP53_1494 | 439.9445905 | 379.917461 | 446.9726527 | 687.6167198 | 633.5423155 | 676.2290066 |
| AP53_1495 | 470.4184703 | 394.2416692 | 486.0047303 | 686.52318 | 619.1561339 | 682.8681701 |
| AP53_1496 | 356.9243673 | 343.4389842 | 367.9158252 | 417.3799755 | 425.05803 | 480.3081293 |
| AP53_1497 | 184.4816543 | 170.4271448 | 140.6727566 | 164.8324545 | 132.3713114 | 146.5452082 |
| AP53_1498 | 111.1053028 | 94.68408709 | 106.0918932 | 107.4705438 | 110.1120927 | 115.4469518 |
| AP53_1499 | 800.2839107 | 956.8491081 | 992.136314 | 821.9052598 | 764.246332 | 921.6319732 |
| AP53_15 | 199.5764198 | 176.8972435 | 186.8691559 | 238.0807839 | 270.3239286 | 239.6756354 |
| AP53_150 | 4354.317865 | 4119.486101 | 4758.090771 | 6213.779153 | 6998.697037 | 7552.293805 |
| AP53_1500 | 478.8945968 | 564.3904483 | 582.9170429 | 435.3749692 | 404.034085 | 472.9754086 |
| AP53_1501 | 307.6356467 | 309.5773643 | 263.0802087 | 238.8299268 | 216.9222683 | 251.5571828 |
| AP53_1502 | 50.03811142 | 48.56506194 | 47.11570754 | 50.42452846 | 54.27924144 | 50.4960671 |
| AP53_1503 | 197.5479057 | 191.4483038 | 219.5376174 | 269.0359069 | 278.771719 | 327.7727823 |
| AP53_1504 | 89.48336431 | 70.72210094 | 81.97958141 | 97.90554838 | 112.7403615 | 100.0072029 |
| AP53_1505 | 1037.810615 | 1188.988184 | 1107.179854 | 978.6225216 | 939.1093675 | 950.9997944 |
| AP53_1506 | 756.246515 | 781.9779917 | 727.502757 | 787.6556154 | 842.9190566 | 864.9785682 |

| | | | | | | |
|---|---|---|---|---|---|---|
| AP53_1507 | 482.212794 | 424.6915248 | 349.9343455 | 426.5529686 | 317.8350047 | 310.9233183 |
| AP53_1508 | 360.6798068 | 324.9859919 | 348.9015516 | 442.2379846 | 441.5675883 | 462.9388447 |
| AP53_1509 | 172.6817357 | 158.6143156 | 164.8033328 | 204.4979622 | 219.9337006 | 208.6107447 |
| AP53_151 | 870.4257116 | 950.8154753 | 1025.976977 | 929.6343017 | 1135.109326 | 1087.63242 |
| AP53_1510 | 117.0316785 | 104.5059391 | 124.4460748 | 157.7507772 | 172.4510047 | 121.4454438 |
| AP53_1511 | 148.8160872 | 151.1960943 | 162.4500749 | 169.6696131 | 202.6333134 | 178.9214751 |
| AP53_1512 | 292.6510011 | 300.0004166 | 320.8408844 | 304.3188585 | 289.0319099 | 319.7498683 |
| AP53_1513 | 250.3699489 | 218.2078174 | 241.1405348 | 238.3696577 | 196.6409225 | 232.7783965 |
| AP53_1514 | 230.5973134 | 193.0167779 | 231.6444387 | 254.3613476 | 215.9836883 | 225.0630978 |
| AP53_1515 | 213.9625547 | 196.9921849 | 222.6961027 | 226.8126719 | 208.1051179 | 217.2273666 |
| AP53_1516 | 137.9593759 | 129.3867242 | 134.8131573 | 135.7854464 | 128.0265733 | 135.8568795 |
| AP53_1517 | 140.9581219 | 114.8501693 | 128.4915639 | 155.5388947 | 137.1870753 | 151.4684111 |
| AP53_1518 | 135.9997613 | 115.3409402 | 114.8907034 | 157.1414716 | 142.8213579 | 147.1361163 |
| AP53_1519 | 158.8739488 | 157.637182 | 187.9586707 | 193.649699 | 193.8937624 | 222.1814446 |
| AP53_152 | 849.5486971 | 854.280481 | 851.4900589 | 845.4888146 | 927.693342 | 962.3054993 |
| AP53_1520 | 190.3206384 | 176.8152082 | 189.30895 | 225.9690453 | 212.3923866 | 235.0932873 |
| AP53_1521 | 155.2884304 | 116.6471425 | 133.4892922 | 172.2812095 | 161.098493 | 164.2660282 |
| AP53_1522 | 293.37851 | 230.155194 | 266.1982794 | 361.0910683 | 309.9942253 | 324.5298762 |
| AP53_1523 | 29.51824542 | 26.00594052 | 27.11297322 | 24.74033088 | 35.68194455 | 30.06055552 |
| AP53_1524 | 1657.628143 | 1466.705715 | 1602.957709 | 1980.185929 | 2063.930244 | 2144.929408 |
| AP53_1525 | 745.4643868 | 667.4426152 | 749.5275379 | 846.6103618 | 887.7469559 | 942.2617301 |
| AP53_1526 | 575.2188238 | 415.0679229 | 434.1330573 | 609.535662 | 579.2849244 | 596.0326114 |
| AP53_1527 | 1424.651729 | 1370.834159 | 1465.550268 | 1650.984596 | 1546.756349 | 1733.202389 |
| AP53_1528 | 52.71818069 | 48.33507743 | 48.39450537 | 52.52928232 | 56.88320836 | 53.74293514 |
| AP53_1529 | 284.1465291 | 313.9744164 | 327.054807 | 270.3624341 | 304.138084 | 311.3509631 |
| AP53_153 | 1444.032566 | 1282.650138 | 1366.590682 | 1651.525596 | 1713.168484 | 1741.516011 |
| AP53_1530 | 127.8430206 | 126.3246432 | 130.677966 | 148.7906893 | 198.5857309 | 202.3069018 |
| AP53_1531 | 714.929487 | 619.6615956 | 587.1780373 | 1114.518387 | 1253.033789 | 1145.250052 |
| AP53_1532 | 1571.257144 | 1542.342762 | 1552.433584 | 2451.751861 | 3072.666093 | 3055.864289 |
| AP53_1533 | 251.275909 | 223.5233807 | 218.9082091 | 335.3779722 | 376.5723391 | 362.0887852 |
| AP53_1534 | 339.8934289 | 312.9539191 | 304.1801338 | 473.3943829 | 510.7015477 | 478.3768072 |
| AP53_1535 | 1175.836826 | 1134.557747 | 1068.110065 | 1747.241674 | 1598.615667 | 1573.056055 |
| AP53_1536 | 1679.883347 | 1660.107789 | 1727.290058 | 2242.645888 | 2147.966033 | 2127.441878 |
| AP53_1537 | 1438.457426 | 1188.80105 | 974.5677009 | 1471.153603 | 1333.384577 | 1276.952399 |
| AP53_1538 | 310.0048303 | 291.0976641 | 281.51717 | 368.3400664 | 373.905977 | 412.8720922 |

| | | | | | | |
|---|---|---|---|---|---|---|
| AP53_1539 | 233.6187728 | 224.4179835 | 262.4949102 | 276.2318684 | 290.0185085 | 313.097112 |
| AP53_154 | 840.6758931 | 813.2193978 | 972.6987514 | 975.1580252 | 1173.339992 | 1079.611534 |
| AP53_1540 | 593.9671875 | 616.4144692 | 615.2799726 | 579.6058199 | 574.3676215 | 649.9195407 |
| AP53_1541 | 251.6950505 | 253.6729123 | 271.0412 | 238.2760489 | 283.006308 | 251.976344 |
| AP53_1542 | 545.0645107 | 567.4536935 | 622.7176103 | 470.0306121 | 503.4635347 | 519.0979234 |
| AP53_1543 | 1787.543767 | 1730.170579 | 1579.877791 | 1799.403329 | 1598.213878 | 1354.398291 |
| AP53_1544 | 2226.962628 | 2652.904956 | 2564.94631 | 2087.977533 | 2245.717792 | 2215.491009 |
| AP53_1545 | 2833.404645 | 3351.445686 | 3026.454689 | 2500.397716 | 2423.833132 | 2219.14013 |
| AP53_1546 | 647.9366259 | 619.7474984 | 648.4093123 | 773.8785327 | 875.7511771 | 866.8287313 |
| AP53_1547 | 333.1337173 | 318.344556 | 258.1718437 | 270.3578714 | 251.5187155 | 248.2704208 |
| AP53_1548 | 132.7417424 | 126.3354185 | 135.8751239 | 160.9848813 | 182.0018946 | 160.9380411 |
| AP53_1549 | 366.5238365 | 387.0639269 | 230.0397446 | 259.2340234 | 232.4175162 | 225.4004868 |
| AP53_155 | 662.6839464 | 591.7382522 | 635.2922137 | 863.8779285 | 946.5063605 | 800.2356311 |
| AP53_1550 | 220.7293085 | 199.9832355 | 243.140052 | 234.654982 | 245.6175799 | 239.5619271 |
| AP53_1551 | 252.5020118 | 230.232997 | 251.9948292 | 255.5137204 | 240.992339 | 255.6820849 |
| AP53_1552 | 291.1553222 | 256.1850508 | 266.0114668 | 280.9794721 | 264.5374827 | 272.356685 |
| AP53_1553 | 339.8868338 | 275.7134665 | 220.2806859 | 322.7747205 | 321.4543546 | 272.2824195 |
| AP53_1554 | 458.1133622 | 454.9360992 | 465.7203259 | 453.6145507 | 493.5028914 | 457.527772 |
| AP53_1555 | 487.2299479 | 542.094393 | 499.7883416 | 514.2398121 | 514.6916115 | 524.8427815 |
| AP53_1556 | 290.673885 | 284.3958061 | 290.2788022 | 355.6864299 | 391.5912015 | 377.1792392 |
| AP53_1557 | 475.786447 | 477.3278467 | 448.4827389 | 626.5312888 | 639.0884384 | 574.2342124 |
| AP53_1558 | 206.0826413 | 238.0963451 | 243.5137629 | 206.5166567 | 189.7926565 | 231.1098231 |
| AP53_1559 | 2876.842112 | 2699.962406 | 2130.579562 | 2940.44974 | 1517.301345 | 2342.976497 |
| AP53_156 | 813.0675624 | 842.7373001 | 646.1356327 | 1096.085673 | 1078.932614 | 1040.512375 |
| AP53_1560 | 166.9256779 | 168.5302266 | 151.0579936 | 148.3391861 | 138.4450288 | 165.3454158 |
| AP53_1561 | 275.3828816 | 276.7540705 | 295.015785 | 240.9378795 | 213.7663376 | 244.3190616 |
| AP53_1562 | 257.715451 | 233.1520635 | 224.4204216 | 189.7778956 | 166.2585835 | 187.122467 |
| AP53_1563 | 310.1739546 | 283.2039018 | 300.8303873 | 350.5572826 | 363.7864054 | 377.4708354 |
| AP53_1564 | 341.580338 | 436.6674407 | 434.7587838 | 245.1809581 | 281.3055276 | 280.792011 |
| AP53_1565 | 200.8994028 | 364.7312208 | 323.6957006 | 85.91072555 | 84.00274403 | 95.12583688 |
| AP53_1566 | 166.8733501 | 145.7464096 | 134.5102771 | 163.1204591 | 168.8582754 | 131.3078371 |
| AP53_1567 | 345.18194 | 348.3458386 | 357.675951 | 417.361973 | 427.5451973 | 432.7272022 |
| AP53_1568 | 20.3795562 | 16.44700022 | 13.47273997 | 21.04511788 | 23.85068674 | 21.04111942 |
| AP53_1569 | 152.6046059 | 149.4108155 | 130.2724137 | 162.7757517 | 171.2002294 | 141.4161259 |
| AP53_157 | 42.08527548 | 33.01597273 | 32.44032322 | 39.12771916 | 63.70960552 | 45.64522877 |

| | | | | | | |
|---|---|---|---|---|---|---|
| AP53_1570 | 238.070553 | 218.2047015 | 176.8889105 | 244.2651932 | 258.9725912 | 237.5864187 |
| AP53_1571 | 81.00598496 | 75.39004359 | 75.28337406 | 100.1893417 | 101.2032071 | 96.2819889 |
| AP53_1572 | 80.46733943 | 80.05924998 | 97.49406222 | 87.40166301 | 108.9240644 | 107.2920274 |
| AP53_1573 | 457.1675783 | 443.3477627 | 386.179334 | 432.6578471 | 499.340714 | 443.7753196 |
| AP53_1574 | 123.0320469 | 115.1691652 | 115.5787315 | 132.2956641 | 131.2399327 | 142.9285476 |
| AP53_1575 | 344.1300305 | 343.9849538 | 320.2564338 | 376.7901238 | 283.804334 | 326.9462847 |
| AP53_1576 | 263.6303207 | 260.2243164 | 280.1165189 | 292.8769397 | 244.8687281 | 285.5073413 |
| AP53_1577 | 164.1362037 | 167.9054974 | 140.3677356 | 164.7836249 | 127.73701 | 165.6383579 |
| AP53_1578 | 294.1732336 | 253.5865545 | 273.6955901 | 340.5591209 | 349.0728207 | 305.4714591 |
| AP53_1579 | 313.641197 | 467.3515187 | 371.6026643 | 371.8428327 | 407.2963427 | 391.3978268 |
| AP53_158 | 13.43080167 | 14.24754027 | 14.3505094 | 16.92759481 | 20.08197245 | 15.87573088 |
| AP53_1580 | 289.1789767 | 296.7081767 | 254.4204167 | 278.0962005 | 227.9909557 | 263.0764196 |
| AP53_1581 | 129.9571936 | 83.56697647 | 64.89110994 | 108.9564906 | 80.43146674 | 81.39967108 |
| AP53_1582 | 323.4756291 | 354.017226 | 310.0254616 | 311.0899613 | 274.4443168 | 307.7134897 |
| AP53_1583 | 392.0099663 | 440.5039635 | 350.768432 | 291.506373 | 309.2190308 | 261.2120756 |
| AP53_1584 | 667.9079185 | 666.3917956 | 687.0580687 | 719.6189103 | 708.6242445 | 732.9219569 |
| AP53_1585 | 232.3703891 | 301.0748195 | 289.6007147 | 224.9221173 | 189.285457 | 210.1079683 |
| AP53_1586 | 183.7296532 | 213.5962571 | 248.5147637 | 203.6771892 | 165.8747202 | 170.9201672 |
| AP53_1587 | 54.74511549 | 45.12071062 | 48.76546099 | 53.91070957 | 59.95077508 | 42.11183631 |
| AP53_1588 | 1004.958653 | 1459.614837 | 1366.828758 | 736.0321865 | 692.352743 | 758.4357265 |
| AP53_1589 | 668.9549065 | 602.4383664 | 642.8180164 | 905.5372299 | 832.5161064 | 756.5075 |
| AP53_159 | 56.83349758 | 45.6022966 | 52.15475991 | 55.32650726 | 57.70031522 | 58.61808327 |
| AP53_1590 | 871.1221837 | 732.6957963 | 789.1635789 | 1183.506755 | 1050.63943 | 1110.561035 |
| AP53_1591 | 1099.085541 | 1289.340392 | 1155.743348 | 1290.218227 | 1231.490458 | 1321.613339 |
| AP53_1592 | 908.3248022 | 913.5210871 | 880.346176 | 979.5804867 | 927.6841904 | 951.7931092 |
| AP53_1593 | 857.5612547 | 710.1921607 | 347.4333854 | 461.3441316 | 425.0751513 | 392.2410433 |
| AP53_1594 | 2090.741178 | 2791.354828 | 2754.855934 | 1360.368308 | 1231.644284 | 1394.671743 |
| AP53_1595 | 595.6932529 | 739.5316129 | 690.8591097 | 493.8942459 | 510.1017263 | 551.1021965 |
| AP53_1596 | 147.642884 | 177.8527696 | 220.9223157 | 159.965771 | 179.889218 | 195.0883084 |
| AP53_1597 | 447.6168805 | 468.9839788 | 477.3150248 | 365.4778461 | 357.0187179 | 399.1987341 |
| AP53_1598 | 272.5133642 | 303.1375491 | 216.9098377 | 219.0007579 | 212.8406235 | 213.8071689 |
| AP53_1599 | 819.0816536 | 1009.187604 | 949.759488 | 776.4149854 | 796.1152618 | 765.7243923 |
| AP53_16 | 615.71065 | 707.4928688 | 689.6043494 | 483.9849665 | 402.8729071 | 496.2887532 |
| AP53_160 | 35.59358226 | 28.97804801 | 30.21159873 | 37.48253137 | 38.96329809 | 26.54262699 |
| AP53_1600 | 91.03552497 | 111.6990716 | 109.5438287 | 79.29557712 | 75.64590762 | 79.37865442 |

| | | | | | | |
|---|---|---|---|---|---|---|
| AP53_1601 | 240.2769392 | 251.8920697 | 230.4644149 | 194.2599918 | 184.5901874 | 225.5973358 |
| AP53_1602 | 40.89830583 | 41.56062148 | 45.31739809 | 40.53713495 | 26.34921515 | 29.88750956 |
| AP53_1603 | 135.5957254 | 126.3170616 | 102.496397 | 95.6579519 | 82.11987223 | 94.54132976 |
| AP53_1604 | 157.4225392 | 189.3800667 | 176.8267102 | 142.4573273 | 139.958204 | 158.1107859 |
| AP53_1605 | 247.1421001 | 222.2036721 | 187.4327585 | 206.8552086 | 183.6227678 | 175.634432 |
| AP53_1606 | 17.45385286 | 24.14837334 | 19.65378197 | 26.21046939 | 16.42323963 | 20.09087531 |
| AP53_1607 | 41.28926418 | 40.22930128 | 39.45229722 | 35.79382479 | 31.90298401 | 41.753104 |
| AP53_1608 | 201.6765005 | 189.2473729 | 202.1712724 | 206.8132732 | 193.5136157 | 189.6934007 |
| AP53_1609 | 170.9462409 | 155.0471187 | 186.0897846 | 226.3959477 | 210.0441566 | 204.9975996 |
| AP53_161 | 68.545824 | 62.29948765 | 65.46998626 | 82.70117832 | 76.77582662 | 88.99184123 |
| AP53_1610 | 187.0991861 | 178.1572591 | 193.2223786 | 228.9240384 | 208.0921068 | 210.398042 |
| AP53_1611 | 24.39200755 | 18.93615018 | 19.06920217 | 25.97799204 | 30.9168841 | 22.58029507 |
| AP53_1612 | 20.94827614 | 18.13107375 | 21.25542807 | 22.10762383 | 25.82696527 | 19.61944044 |
| AP53_1613 | 13.98668264 | 10.42668082 | 12.91757609 | 14.5545675 | 18.47742365 | 12.06943595 |
| AP53_1616 | 46.75328627 | 40.21443397 | 23.12112147 | 16.58213369 | 24.59017035 | 14.20591304 |
| AP53_1617 | 106.7263947 | 98.88990583 | 96.59337376 | 56.88249344 | 62.31026139 | 62.49262608 |
| AP53_1618 | 186.5939038 | 178.6808967 | 181.6165454 | 123.607331 | 114.4171705 | 131.5605723 |
| AP53_1619 | 92.31413544 | 88.57441089 | 84.36446437 | 81.44408825 | 78.60947495 | 87.27043647 |
| AP53_162 | 40.62866955 | 37.79095951 | 43.0333597 | 47.23438828 | 44.32205333 | 52.37354519 |
| AP53_1620 | 151.1667146 | 154.873531 | 133.5573724 | 96.09506568 | 97.32271604 | 99.74456623 |
| AP53_1621 | 94.39934886 | 94.87973783 | 109.2003109 | 89.77085765 | 110.5831468 | 105.8907311 |
| AP53_1622 | 205.4778793 | 189.8736052 | 210.9542446 | 137.7827485 | 156.6827985 | 170.8274084 |
| AP53_1623 | 135.4887465 | 127.9694917 | 118.9449656 | 82.74397744 | 94.34749198 | 89.1568749 |
| AP53_1624 | 83.84211406 | 72.50127902 | 75.11914955 | 80.37326994 | 91.03652105 | 76.53863101 |
| AP53_1625 | 74.5889164 | 74.70903002 | 78.99074251 | 92.0437968 | 99.70275056 | 97.50599137 |
| AP53_1626 | 97.23626308 | 117.8526863 | 125.8367036 | 161.4167077 | 197.284166 | 182.3637356 |
| AP53_1627 | 4.427736813 | 2.972107488 | 5.035266455 | 6.944654282 | 5.656893649 | 4.22726562 |
| AP53_1628 | 4.682894528 | 7.858453697 | 2.560304977 | 5.738167733 | 6.903327843 | 4.346679903 |
| AP53_1629 | 2.302423143 | 4.636487681 | 2.719043885 | 3.046967066 | 6.109445141 | 2.930904163 |
| AP53_163 | 9.834853822 | 14.96879301 | 14.30549211 | 10.31350148 | 8.653923397 | 12.73778713 |
| AP53_1630 | 2895.341041 | 3709.743161 | 3957.834745 | 1819.522615 | 1629.833107 | 1817.691947 |
| AP53_1631 | 2592.775699 | 2955.469171 | 2817.485034 | 1365.227563 | 1281.229463 | 1346.076748 |
| AP53_1632 | 113.9532613 | 133.341018 | 127.1952091 | 151.121716 | 175.3637031 | 146.0142473 |
| AP53_1633 | 97.88704308 | 100.7372971 | 87.86390991 | 106.4735697 | 131.9495548 | 104.1641605 |
| AP53_1634 | 267.8672778 | 324.0593905 | 274.3255271 | 275.6189915 | 250.7166802 | 282.8782067 |

| | | | | | | |
|---|---|---|---|---|---|---|
| AP53_1635 | 177.043469 | 157.5793853 | 120.3772707 | 154.4511601 | 146.3062432 | 147.2278722 |
| AP53_1636 | 327.933481 | 404.7616505 | 347.0693902 | 301.909432 | 282.367595 | 296.4009255 |
| AP53_1637 | 69.24580881 | 73.42558029 | 80.81034772 | 60.01023274 | 62.14076407 | 58.3288493 |
| AP53_1638 | 188.5317501 | 212.6106783 | 209.7297941 | 146.2151668 | 157.2862386 | 177.6064208 |
| AP53_1639 | 156.7497899 | 197.6070015 | 207.7384596 | 124.0475307 | 131.6364203 | 136.9772454 |
| AP53_164 | 13.03258383 | 13.12213495 | 14.96329182 | 10.38012889 | 11.20705346 | 16.93564906 |
| AP53_1640 | 93.44249768 | 77.90202516 | 92.20422988 | 70.78812376 | 77.58025576 | 72.32101182 |
| AP53_1641 | 365.7173667 | 382.8388507 | 344.1630577 | 250.6854634 | 224.2929173 | 270.6176331 |
| AP53_1642 | 293.3066932 | 362.9183024 | 373.6127588 | 203.8729845 | 189.5172059 | 222.8100202 |
| AP53_1643 | 351.678134 | 491.2207083 | 510.537776 | 191.8460745 | 165.6729908 | 204.9777912 |
| AP53_1644 | 91.8600509 | 77.39404591 | 55.32576722 | 41.3785651 | 30.51929043 | 45.15461918 |
| AP53_1645 | 108.6393355 | 91.43834061 | 70.01209618 | 42.6501791 | 31.46954532 | 48.50327814 |
| AP53_1646 | 61.07480337 | 44.22965222 | 20.51208545 | 16.39303919 | 14.46973849 | 19.66790952 |
| AP53_1647 | 49.34960812 | 46.79933663 | 30.95919287 | 20.03221919 | 19.11980648 | 22.61105104 |
| AP53_1648 | 2788.234426 | 2957.04881 | 2531.396289 | 2139.271816 | 1996.317769 | 1903.866496 |
| AP53_1649 | 117.5266739 | 147.269035 | 162.7818976 | 135.6734092 | 216.3128586 | 184.5485364 |
| AP53_165 | 6.358436005 | 4.548879629 | 7.025953192 | 6.200223681 | 5.920004981 | 4.686038633 |
| AP53_1650 | 90.6244458 | 119.1132555 | 122.778532 | 107.2737113 | 176.8108154 | 170.8274084 |
| AP53_1651 | 403.26099 | 591.434892 | 603.6424799 | 266.2173935 | 273.352139 | 299.1666811 |
| AP53_1652 | 268.8492438 | 340.532089 | 330.9922667 | 277.2010554 | 324.2243907 | 302.907326 |
| AP53_1653 | 135.6497551 | 127.6655262 | 125.8112381 | 107.7210579 | 101.5867335 | 114.9047655 |
| AP53_1654 | 240.7021629 | 284.2457915 | 295.6244091 | 209.1354688 | 194.9891882 | 239.8978045 |
| AP53_1655 | 764.7838811 | 885.2114859 | 876.4048022 | 778.2318145 | 793.3998965 | 906.319375 |
| AP53_1656 | 578.8675518 | 626.8227765 | 609.9865648 | 577.0697679 | 561.4466946 | 613.7889031 |
| AP53_1657 | 271.2080694 | 304.3241796 | 309.2983673 | 266.8172209 | 285.0433996 | 305.2564336 |
| AP53_1658 | 344.2122598 | 352.7787564 | 290.6355798 | 279.9147078 | 285.1074399 | 305.3269412 |
| AP53_1659 | 667.4196266 | 940.7176978 | 701.6767622 | 464.3988511 | 411.330357 | 428.2243173 |
| AP53_166 | 199.7432022 | 201.4376809 | 202.8792776 | 200.0747665 | 170.1782173 | 174.5312722 |
| AP53_1660 | 348.2415003 | 566.0711792 | 524.014971 | 255.2272701 | 193.5535422 | 273.9826898 |
| AP53_1661 | 51.72553536 | 42.77120122 | 49.79990802 | 51.01506704 | 38.81715788 | 53.80760484 |
| AP53_1662 | 38.49497429 | 38.6733057 | 33.1597566 | 28.28607386 | 25.89763765 | 32.20998302 |
| AP53_1663 | 144.4800977 | 150.3332067 | 125.0824127 | 94.80348735 | 74.25795314 | 85.29086189 |
| AP53_1664 | 148.9659638 | 163.0143195 | 173.7433814 | 159.7055765 | 178.8218042 | 196.3213853 |
| AP53_1665 | 152.7117391 | 135.9865541 | 145.4404175 | 179.8317045 | 187.2932067 | 178.6123569 |
| AP53_1666 | 926.5697458 | 735.905361 | 676.9031174 | 1160.45525 | 1121.440944 | 1172.980522 |

| | | | | | | |
|---|---|---|---|---|---|---|
| AP53_1667 | 1192.297379 | 1215.354997 | 1357.99095 | 1603.752358 | 1705.301168 | 1889.962854 |
| AP53_1668 | 10640.44339 | 10773.52181 | 11333.68683 | 9751.300212 | 8968.206754 | 10716.57177 |
| AP53_1669 | 166.2463915 | 195.3768392 | 173.8574299 | 137.3132846 | 139.0330942 | 162.4740351 |
| AP53_167 | 132.2036514 | 138.8141977 | 119.3845434 | 85.45705123 | 81.29503615 | 77.99986886 |
| AP53_1670 | 74.82875214 | 103.3296185 | 85.46702272 | 69.93769436 | 63.86335198 | 87.25224565 |
| AP53_1671 | 75.73538396 | 50.84426738 | 60.64452785 | 137.4049454 | 86.09667806 | 138.0235775 |
| AP53_1672 | 203.8070856 | 137.2780367 | 182.8029657 | 345.8895094 | 225.810158 | 386.1134548 |
| AP53_1673 | 136.5797408 | 157.2928446 | 132.8706112 | 100.0082302 | 85.46434925 | 111.6674486 |
| AP53_1674 | 158.6717081 | 195.9207648 | 178.6569541 | 113.9152607 | 92.59160831 | 122.5242365 |
| AP53_1675 | 75.89197302 | 84.53468781 | 62.05885726 | 45.49964975 | 40.53506596 | 39.78655731 |
| AP53_1676 | 185.855935 | 207.9108885 | 245.9169039 | 150.2645612 | 110.2062609 | 157.7087014 |
| AP53_1677 | 243.4046309 | 230.5386686 | 240.5843438 | 280.518555 | 238.5162599 | 292.7465602 |
| AP53_1678 | 299.4977406 | 267.7786288 | 283.0139447 | 393.9024444 | 320.3777228 | 385.7473072 |
| AP53_1679 | 249.8646203 | 233.3315166 | 244.4425453 | 328.2672819 | 267.2380751 | 330.2896875 |
| AP53_168 | 497.0152005 | 644.8946694 | 597.4918541 | 361.7884634 | 315.7615675 | 415.5479771 |
| AP53_1681 | 420.4148335 | 469.2789413 | 399.2604649 | 287.5197182 | 260.842179 | 268.6324799 |
| AP53_1682 | 59.57343316 | 67.95618927 | 64.91786782 | 42.83823841 | 47.58037541 | 48.60490992 |
| AP53_1683 | 64.36701195 | 73.92575021 | 78.9470974 | 54.48955703 | 56.73410127 | 62.77643726 |
| AP53_1684 | 94.93199899 | 106.0726309 | 112.4480214 | 74.84776065 | 81.68723291 | 78.2622497 |
| AP53_1685 | 77.88579722 | 100.105984 | 103.7891286 | 65.28058095 | 63.57915398 | 70.64320441 |
| AP53_1686 | 450.8625617 | 319.4072903 | 102.5841599 | 173.5710095 | 131.9660414 | 126.3132205 |
| AP53_1687 | 106.3473602 | 103.1463772 | 93.20131549 | 84.76123325 | 76.36806426 | 77.67429571 |
| AP53_1688 | 67.79773684 | 62.20369263 | 58.6026288 | 53.81541543 | 46.71083192 | 51.07601718 |
| AP53_1689 | 9.997363646 | 6.481931791 | 6.360336574 | 6.681945321 | 11.16492168 | 10.12318872 |
| AP53_169 | 438.1245577 | 468.1069293 | 434.5822278 | 630.0646108 | 683.0699081 | 669.4931926 |
| AP53_1690 | 2128.122516 | 2292.51073 | 2481.009531 | 2713.925455 | 3049.861177 | 3251.710324 |
| AP53_1691 | 1188.672885 | 1244.053718 | 1398.459896 | 1212.102969 | 1387.196571 | 1398.500419 |
| AP53_1692 | 3005.307138 | 2946.832086 | 2340.97577 | 2190.203171 | 1955.859095 | 1916.632759 |
| AP53_1693 | 11611.63634 | 9499.75567 | 7480.088317 | 10050.28481 | 8971.344297 | 8548.199413 |
| AP53_1696 | 364.7874719 | 364.2063093 | 326.4436791 | 333.4165585 | 339.2864977 | 302.3524028 |
| AP53_1697 | 243.9472139 | 266.8395254 | 320.0990818 | 381.8784782 | 369.3143425 | 425.4608648 |
| AP53_1698 | 669.2291798 | 575.2115846 | 632.2736794 | 863.7126002 | 788.433459 | 874.7170125 |
| AP53_1699 | 371.9432807 | 296.3208818 | 354.7742166 | 450.2082128 | 418.0358881 | 422.5835228 |
| AP53_17 | 405.9724295 | 474.0754489 | 462.1151083 | 421.8715218 | 402.4324715 | 417.3228611 |
| AP53_170 | 548.7441824 | 511.5430419 | 531.3045321 | 863.2132933 | 944.984084 | 894.7399098 |

| | | | | | | |
|---|---|---|---|---|---|---|
| AP53_1700 | 346.3140131 | 273.2975537 | 296.0875261 | 404.3210376 | 369.3069285 | 379.3873938 |
| AP53_1701 | 301.6684706 | 269.0818743 | 330.0579955 | 389.1679064 | 378.9979412 | 390.6368264 |
| AP53_1702 | 225.3304782 | 193.8309433 | 227.9968651 | 304.8471741 | 313.6936091 | 299.4407087 |
| AP53_1703 | 273.0249287 | 221.6854925 | 261.9052779 | 351.157552 | 342.5830842 | 344.1040851 |
| AP53_1704 | 342.3788488 | 241.4837334 | 274.4779692 | 462.562202 | 401.0109053 | 373.8259708 |
| AP53_1705 | 192.7988672 | 176.3563023 | 184.321269 | 234.7635119 | 225.5900619 | 234.4353267 |
| AP53_1706 | 4039.295386 | 4599.990013 | 4513.957033 | 6464.841305 | 6999.654025 | 7197.178348 |
| AP53_1707 | 3150.699543 | 3148.720068 | 3055.12417 | 3024.290477 | 2849.686859 | 2818.921504 |
| AP53_1708 | 53.7757734 | 53.78802061 | 65.80334517 | 139.1309163 | 98.56943454 | 112.4182419 |
| AP53_1709 | 49.53960572 | 41.50615147 | 44.63593346 | 138.0086865 | 82.47155478 | 87.78021241 |
| AP53_171 | 174.8956041 | 214.8276444 | 202.9406045 | 147.5304994 | 126.8232042 | 140.908854 |
| AP53_1710 | 19.79276035 | 15.75999102 | 19.08100972 | 41.27957332 | 37.51413683 | 31.17299384 |
| AP53_1711 | 9.942281753 | 6.256624001 | 8.926154169 | 18.081749 | 16.19928636 | 9.991718738 |
| AP53_1712 | 28.22721611 | 26.65774167 | 30.96420083 | 39.51780141 | 46.38250166 | 33.70279031 |
| AP53_1713 | 703.5934251 | 629.7931279 | 438.7901499 | 1696.680211 | 788.0759965 | 1215.867274 |
| AP53_1714 | 8.56715588 | 7.749943071 | 8.78244149 | 9.0542949 | 13.81334496 | 7.668063218 |
| AP53_1715 | 53.83607055 | 58.12655512 | 58.4683293 | 64.82273072 | 58.4989355 | 64.43679006 |
| AP53_1716 | 111.0300747 | 117.4289494 | 123.2183799 | 167.7837084 | 138.4770164 | 167.8911527 |
| AP53_1717 | 97.31093328 | 98.04456153 | 95.71522198 | 125.9321964 | 97.4101866 | 117.4661702 |
| AP53_1718 | 105.0230606 | 96.06618396 | 98.84365489 | 128.3010877 | 112.4991914 | 128.3634206 |
| AP53_1719 | 78.04420515 | 62.65253828 | 79.2446762 | 158.107627 | 104.7500652 | 146.4188761 |
| AP53_172 | 259.9596362 | 288.6011409 | 263.2975959 | 220.3539813 | 192.6961621 | 222.7998368 |
| AP53_1720 | 68.29031749 | 49.23454758 | 65.26291958 | 95.32043681 | 78.26284903 | 89.63444771 |
| AP53_1721 | 82.62901746 | 52.53963844 | 58.58791342 | 109.4756599 | 73.11667212 | 87.99560397 |
| AP53_1722 | 367.8244189 | 379.5060874 | 378.2917156 | 545.5137435 | 584.5283777 | 553.8019602 |
| AP53_1723 | 426.364382 | 402.0267744 | 335.2395473 | 508.0317913 | 552.6035386 | 487.6747916 |
| AP53_1724 | 333.2712879 | 325.160655 | 307.1512537 | 408.2962113 | 419.399464 | 418.2644483 |
| AP53_1725 | 71.65003362 | 63.79495643 | 52.08118885 | 97.77580884 | 90.93245462 | 72.7257936 |
| AP53_1726 | 1422.429213 | 896.3548748 | 410.6729183 | 1380.890065 | 1140.199649 | 1027.368843 |
| AP53_1727 | 693.798868 | 716.5463561 | 749.2636123 | 775.7400154 | 746.8220533 | 787.3901137 |
| AP53_1728 | 147.0287535 | 133.9379069 | 125.604127 | 123.5581133 | 118.928394 | 110.7743306 |
| AP53_1729 | 334.7507397 | 395.0522951 | 355.4583413 | 266.874112 | 239.7108684 | 252.7332399 |
| AP53_173 | 88.16526949 | 86.2933518 | 104.9611808 | 88.38117685 | 93.10941168 | 87.49505057 |
| AP53_1730 | 301.8900871 | 336.9651306 | 361.1975512 | 318.0049144 | 271.5867198 | 297.5494399 |
| AP53_1731 | 277.0905219 | 248.6708544 | 260.6647901 | 288.8522303 | 288.129276 | 301.8643753 |

| | | | | | | |
|---|---|---|---|---|---|---|
| AP53_1732 | 295.7692769 | 278.9137121 | 330.8170061 | 310.9260615 | 285.5147362 | 332.8041677 |
| AP53_1733 | 156.9833961 | 115.2536 | 153.8045026 | 217.3657061 | 199.7911984 | 195.254837 |
| AP53_1734 | 204.5017409 | 119.3309505 | 136.6311066 | 247.6373983 | 194.3047627 | 215.7013738 |
| AP53_1735 | 141.4711109 | 142.3144135 | 156.764629 | 249.0237281 | 244.3778056 | 248.3127138 |
| AP53_1736 | 10154.78648 | 8868.401328 | 9108.547104 | 14805.95107 | 14146.02965 | 15667.23845 |
| AP53_1737 | 11056.25967 | 8441.723624 | 8147.164943 | 14533.64898 | 14149.69889 | 15487.71342 |
| AP53_1738 | 1246.472563 | 1060.614335 | 1088.006783 | 1634.250236 | 1676.640809 | 1688.663809 |
| AP53_1739 | 841.1319871 | 580.9722287 | 362.4410951 | 773.2833085 | 722.7967294 | 665.1629903 |
| AP53_174 | 89.35361303 | 78.4873513 | 85.4923964 | 99.98085787 | 95.32467596 | 81.84705775 |
| AP53_1740 | 78.76710752 | 73.58899033 | 53.26781881 | 63.70121206 | 72.79528932 | 62.66735875 |
| AP53_1741 | 2032.904198 | 2258.37008 | 962.106133 | 1174.227423 | 1125.555457 | 899.475092 |
| AP53_1742 | 1443.016002 | 1644.620446 | 1420.187479 | 1097.75905 | 718.4859949 | 928.4461785 |
| AP53_1743 | 1068.959645 | 1224.450122 | 1092.642967 | 1224.617074 | 782.1458062 | 1095.683534 |
| AP53_1744 | 32.16299627 | 23.41591083 | 28.9945438 | 31.40830981 | 27.42655072 | 27.82449006 |
| AP53_1745 | 38.36238265 | 29.70570948 | 33.13458206 | 34.75532761 | 24.54105856 | 33.65994072 |
| AP53_1746 | 43.24718137 | 27.96381633 | 37.16026643 | 41.58545792 | 29.47241591 | 36.08675751 |
| AP53_1747 | 35.80323069 | 28.90872253 | 31.36802356 | 34.50313584 | 18.75706841 | 28.92339635 |
| AP53_1748 | 47.1831696 | 35.70324015 | 45.23618412 | 59.61061614 | 32.20779771 | 48.45446399 |
| AP53_1749 | 36.96603991 | 28.54119301 | 38.86517223 | 50.35746799 | 25.75450073 | 35.80784295 |
| AP53_175 | 84.03844471 | 49.98713281 | 50.98207285 | 63.05529068 | 63.21578653 | 49.45900776 |
| AP53_1750 | 33.53417635 | 26.87584586 | 29.57095099 | 47.23101231 | 31.97155071 | 44.81404803 |
| AP53_1751 | 36.48489591 | 26.55381442 | 31.79858932 | 57.53406253 | 36.6157299 | 51.10550304 |
| AP53_1752 | 74.25139679 | 77.90602876 | 82.27839653 | 107.3281543 | 87.84519651 | 103.1161571 |
| AP53_1753 | 33.85123587 | 29.93927106 | 26.92516628 | 42.62450948 | 32.89828589 | 39.38849254 |
| AP53_1754 | 3164.752561 | 2948.063603 | 3105.067868 | 3239.584581 | 2770.198498 | 3156.589508 |
| AP53_1755 | 2075.365737 | 2186.798747 | 2173.580871 | 2324.007541 | 2005.181322 | 2357.076205 |
| AP53_1756 | 516.0233007 | 707.6237577 | 716.7845208 | 292.6973989 | 285.3598519 | 321.4661598 |
| AP53_1757 | 238.807498 | 205.6213977 | 187.1950717 | 250.1730173 | 207.5236123 | 251.0565007 |
| AP53_1758 | 448.4949732 | 401.7860019 | 377.6002262 | 511.3889088 | 359.5018768 | 496.0826676 |
| AP53_1759 | 98.91712872 | 87.90819605 | 95.96625478 | 113.7278324 | 123.9287541 | 112.987587 |
| AP53_176 | 70.79951164 | 52.52271201 | 66.84316218 | 74.05822731 | 71.27685998 | 74.64646541 |
| AP53_1760 | 24.3324264 | 15.476912 | 14.76248574 | 22.69836577 | 14.65649718 | 18.31815102 |
| AP53_1761 | 23.55967867 | 20.21724279 | 24.94213383 | 25.19455972 | 35.12536289 | 21.30017561 |
| AP53_1762 | 51.37281637 | 50.89269681 | 48.71620295 | 55.86106288 | 82.94420313 | 73.04362719 |
| AP53_1763 | 147.4453722 | 139.2650895 | 148.1553106 | 166.88617 | 187.0768006 | 177.5064752 |

| | | | | | | |
|---|---|---|---|---|---|---|
| AP53_1764 | 575.0396161 | 458.7201247 | 473.5420464 | 794.48695 | 721.0452843 | 748.4165882 |
| AP53_1765 | 898.731219 | 819.1599424 | 865.157026 | 1160.324694 | 1223.213569 | 1399.774808 |
| AP53_1766 | 559.0702013 | 461.2778369 | 368.6730548 | 580.3600234 | 590.6825496 | 583.0722979 |
| AP53_1767 | 584.4981571 | 495.9239978 | 475.5421838 | 607.8742701 | 606.6474506 | 611.8379865 |
| AP53_1768 | 352.8815878 | 170.3199556 | 96.80042858 | 289.8418785 | 201.9164692 | 163.3679999 |
| AP53_1769 | 988.1887049 | 832.8915626 | 740.5963325 | 1149.991049 | 1103.997204 | 1092.216631 |
| AP53_177 | 468.2617023 | 580.3659059 | 642.1643152 | 371.2785796 | 348.3137983 | 397.4631701 |
| AP53_1770 | 199.0267697 | 182.3945214 | 159.8600267 | 239.232489 | 236.8062711 | 206.9598793 |
| AP53_1771 | 7.084378901 | 5.944214976 | 3.485953699 | 3.038286248 | 8.267767641 | 6.57574652 |
| AP53_1772 | 1528.064065 | 1922.920764 | 1677.336114 | 2015.04761 | 2487.702172 | 2022.359791 |
| AP53_1773 | 2235.78732 | 2235.751587 | 2209.08533 | 3077.60972 | 3291.692736 | 2916.463577 |
| AP53_1774 | 1696.246294 | 1349.89407 | 875.1293098 | 1829.49683 | 1964.450601 | 1542.388316 |
| AP53_1775 | 2529.549396 | 2236.974578 | 1752.954191 | 3086.931463 | 3255.180764 | 2695.79827 |
| AP53_1776 | 1032.901042 | 1132.131812 | 1045.767714 | 1100.740301 | 1278.890814 | 1264.580747 |
| AP53_1777 | 203.9870881 | 199.2582523 | 203.7857839 | 219.2203375 | 239.2305661 | 243.2339393 |
| AP53_1778 | 577.8220219 | 537.7984792 | 584.440954 | 771.6086534 | 778.2917636 | 838.6186678 |
| AP53_1779 | 411.0808757 | 360.0413251 | 390.85307 | 497.2799964 | 555.8537609 | 536.3812362 |
| AP53_178 | 2469.406038 | 1984.247353 | 2252.505927 | 3345.841757 | 3669.682017 | 3170.169078 |
| AP53_1780 | 410.3913683 | 394.9416027 | 478.1894816 | 496.9309652 | 593.18159 | 601.8885346 |
| AP53_1781 | 516.4951643 | 445.3018497 | 542.5518649 | 616.691998 | 673.8464329 | 660.451097 |
| AP53_1782 | 370.2764093 | 303.2507446 | 344.4830339 | 383.5599023 | 434.0340041 | 403.5717647 |
| AP53_1783 | 366.7088526 | 312.1177254 | 325.4040946 | 463.7455662 | 506.2919816 | 452.2293533 |
| AP53_1784 | 241.9343068 | 192.205959 | 222.3974914 | 319.971216 | 312.7643465 | 275.4162628 |
| AP53_1785 | 50.38636153 | 39.47734076 | 26.27095542 | 25.02340103 | 42.05777452 | 30.26477125 |
| AP53_1786 | 70.08148793 | 60.64478088 | 45.47312798 | 66.05252203 | 91.85162368 | 65.90757429 |
| AP53_1787 | 71.50794953 | 55.30906278 | 37.3287542 | 55.74712715 | 80.77391412 | 57.94876621 |
| AP53_1788 | 150.9366283 | 127.0025561 | 134.0872808 | 160.4986767 | 186.6774904 | 146.0929081 |
| AP53_1789 | 53.662296 | 46.84017813 | 46.95585781 | 54.27578019 | 56.70362443 | 53.0644851 |
| AP53_179 | 587.1179014 | 530.8251521 | 439.441436 | 605.5462326 | 627.1438004 | 505.4144395 |
| AP53_1790 | 27.26553722 | 23.00450302 | 27.56145849 | 38.01284449 | 37.66300245 | 31.01230831 |
| AP53_1791 | 37.18102237 | 35.43936952 | 34.70251205 | 45.06400241 | 45.17870468 | 41.83469625 |
| AP53_1792 | 31.65831821 | 26.26135601 | 29.50351438 | 25.39139222 | 39.77503347 | 35.4914176 |
| AP53_1793 | 40.1877494 | 36.88115201 | 45.18007264 | 43.55008484 | 52.3005531 | 49.95859369 |
| AP53_1794 | 46.04846286 | 35.98372994 | 40.33746423 | 50.78278444 | 55.76080882 | 59.38301704 |
| AP53_1795 | 44.23711598 | 38.19833601 | 43.09906392 | 51.1970962 | 55.71842451 | 63.7292317 |

| | | | | | | |
|---|---|---|---|---|---|---|
| AP53_1796 | 29.79245576 | 23.44493937 | 24.9527414 | 30.30731007 | 32.8562029 | 33.44770737 |
| AP53_1797 | 16.2757498 | 14.489024 | 10.67823748 | 13.42533382 | 21.06705221 | 14.84401893 |
| AP53_1798 | 21.03938389 | 18.23618538 | 18.7520268 | 25.39139222 | 28.08940295 | 29.68803786 |
| AP53_1799 | 33.4528539 | 27.61437516 | 27.36815414 | 26.08841083 | 37.93446329 | 31.14085673 |
| AP53_18 | 35.49569012 | 29.20803251 | 28.71300752 | 37.0123204 | 38.92481582 | 30.89370708 |
| AP53_180 | 8.346283893 | 8.693414402 | 6.797609714 | 7.617417666 | 9.333874521 | 6.869306633 |
| AP53_1800 | 31.07564978 | 24.9780107 | 31.13833488 | 31.0512322 | 43.83224956 | 36.07439557 |
| AP53_1801 | 13.41494476 | 11.136192 | 13.35802092 | 13.01046543 | 14.02535615 | 10.14310842 |
| AP53_1802 | 12.59137656 | 6.452143501 | 12.74551821 | 16.26636064 | 17.23584784 | 10.87640217 |
| AP53_1803 | 113.5484637 | 109.8157174 | 75.61154217 | 79.92044764 | 89.86114665 | 71.27062036 |
| AP53_1804 | 84.98968056 | 84.26421823 | 99.12737587 | 90.87654774 | 91.44647444 | 99.22331803 |
| AP53_1805 | 170.1791019 | 127.8813858 | 153.2472399 | 191.1100922 | 179.0529816 | 211.5879183 |
| AP53_1806 | 92.57002636 | 94.87333525 | 101.2916272 | 100.6380294 | 102.2890358 | 107.2741036 |
| AP53_1807 | 591.1593888 | 609.3610804 | 583.7695087 | 664.4681233 | 671.3047729 | 659.5508737 |
| AP53_1808 | 208.1122278 | 156.9878717 | 117.4732552 | 261.3094733 | 244.6743806 | 206.6571989 |
| AP53_1809 | 233.0115517 | 224.9533005 | 228.5595691 | 260.0799877 | 261.2668407 | 262.1195218 |
| AP53_181 | 111.523621 | 91.26791459 | 76.20336286 | 106.8554423 | 109.4002825 | 108.9275766 |
| AP53_1810 | 191.5018839 | 171.6430113 | 165.6997923 | 216.5923309 | 212.9733578 | 206.0010499 |
| AP53_1811 | 234.4951978 | 239.2073262 | 274.0211248 | 236.0698652 | 234.1845877 | 257.0374949 |
| AP53_1812 | 239.0886742 | 304.2943566 | 307.8681098 | 203.2472791 | 193.7510332 | 217.8383208 |
| AP53_1813 | 108.7045681 | 146.1457292 | 140.6572793 | 83.01260782 | 78.13667152 | 90.26086592 |
| AP53_1814 | 963.2432558 | 1036.559029 | 596.5552568 | 459.2650724 | 348.5944791 | 421.617771 |
| AP53_1815 | 694.8713045 | 770.816077 | 536.8601094 | 527.4638544 | 362.8331586 | 455.7555974 |
| AP53_1816 | 43.68988305 | 37.17671525 | 43.84366157 | 49.21388541 | 54.55453046 | 47.71654949 |
| AP53_1817 | 40.75085209 | 35.90200638 | 40.50493104 | 48.23615513 | 58.87174275 | 50.73965725 |
| AP53_1818 | 43.58158092 | 38.76284344 | 44.77790526 | 51.49726084 | 58.35049066 | 43.29744787 |
| AP53_1819 | 90.67511062 | 84.91376218 | 90.87552605 | 114.9862516 | 121.9049313 | 116.4771674 |
| AP53_182 | 508.0680402 | 522.7156893 | 481.1197097 | 430.807288 | 400.9606218 | 413.62385 |
| AP53_1820 | 103.208122 | 91.04297172 | 111.1516282 | 139.8000368 | 145.4750411 | 138.343051 |
| AP53_1821 | 71.37511743 | 37.67146241 | 22.79602449 | 82.79132736 | 62.94579842 | 61.94865618 |
| AP53_1822 | 148.6332947 | 155.2027126 | 148.3163717 | 183.3119965 | 170.5523238 | 163.494387 |
| AP53_1823 | 54.92462684 | 58.43003605 | 56.1879266 | 59.87938913 | 63.20389119 | 64.02792973 |
| AP53_1824 | 17.2153657 | 13.29268257 | 13.44277558 | 17.39349192 | 14.79096046 | 14.28479667 |
| AP53_1825 | 93.15416083 | 96.32244018 | 115.1431849 | 119.0977206 | 111.781373 | 136.5449931 |
| AP53_1826 | 120.7493026 | 99.45105087 | 114.1327063 | 166.1725557 | 125.6773206 | 143.695718 |

| | | | | | | |
|---|---|---|---|---|---|---|
| AP53_1827 | 193.2759691 | 165.3239691 | 156.9767567 | 287.1214862 | 317.4278555 | 288.4746289 |
| AP53_1828 | 50.25232534 | 46.56023444 | 44.46875768 | 72.46532161 | 89.62044994 | 60.20291206 |
| AP53_1829 | 954.6591252 | 1282.917846 | 1368.037467 | 649.5549617 | 579.4156509 | 622.2783655 |
| AP53_183 | 781.9748925 | 930.7167749 | 965.0333239 | 722.0443097 | 717.2740903 | 800.1087379 |
| AP53_1830 | 48.05851481 | 46.53353212 | 48.03484348 | 52.61884631 | 57.82602397 | 52.19219219 |
| AP53_1831 | 80.33179647 | 56.04545548 | 39.34147746 | 62.68790607 | 66.01781379 | 49.92199399 |
| AP53_1832 | 435.3304022 | 425.1283893 | 466.5163279 | 424.1220066 | 417.5011795 | 436.0849697 |
| AP53_1833 | 432.8860465 | 422.8684025 | 459.3763621 | 429.6918604 | 385.2681741 | 434.5405096 |
| AP53_1834 | 222.3288265 | 191.2815005 | 168.0892667 | 205.7768772 | 199.8883543 | 185.4330584 |
| AP53_1835 | 188.1403429 | 202.6174415 | 231.8131675 | 154.4877018 | 163.299238 | 180.6059724 |
| AP53_1836 | 121.9632636 | 128.8976381 | 142.0800185 | 104.1206775 | 111.750333 | 116.4757905 |
| AP53_1837 | 363.4935146 | 402.3997121 | 419.7311952 | 263.4639824 | 253.7888196 | 314.6901612 |
| AP53_1838 | 128.0722873 | 120.440012 | 151.1928668 | 117.2840498 | 115.385478 | 104.4298163 |
| AP53_1839 | 135.0992274 | 138.3676897 | 136.8346637 | 123.7285023 | 118.7364481 | 122.5616496 |
| AP53_184 | 260.8645421 | 228.7333923 | 254.9858933 | 299.3927269 | 315.0889762 | 328.0170243 |
| AP53_1840 | 528.041093 | 572.1397527 | 623.7589835 | 363.6542808 | 323.7674789 | 389.4170836 |
| AP53_1841 | 190.1008172 | 183.7447014 | 161.5245587 | 224.1162834 | 242.4201391 | 218.9945246 |
| AP53_1842 | 122.0038455 | 113.2309491 | 106.2781685 | 138.974951 | 151.2262815 | 133.9637023 |
| AP53_1843 | 320.2012756 | 326.3135192 | 342.2003049 | 352.8798962 | 374.6883915 | 342.8765339 |
| AP53_1844 | 311.9719756 | 318.6784782 | 282.0861981 | 318.0704418 | 339.4136189 | 326.386945 |
| AP53_1845 | 377.206926 | 350.9139936 | 327.8665166 | 380.1779408 | 356.9797624 | 340.3319637 |
| AP53_1846 | 72.20506065 | 74.06249944 | 81.50105703 | 61.49047232 | 66.34352249 | 56.4880657 |
| AP53_1847 | 50.54234899 | 56.45482365 | 56.5648487 | 48.45778949 | 48.74470286 | 47.05337105 |
| AP53_1848 | 354.6828032 | 373.1958611 | 414.0787335 | 283.0132638 | 278.7232314 | 247.8620292 |
| AP53_1849 | 314.6835402 | 311.316275 | 424.7990675 | 279.9563758 | 277.9317475 | 235.4541496 |
| AP53_185 | 294.3985866 | 254.0290043 | 290.3817811 | 360.1501983 | 386.7764332 | 400.3916419 |
| AP53_1850 | 284.5137169 | 271.4967117 | 386.6365313 | 252.7048083 | 240.0139162 | 214.5840548 |
| AP53_1851 | 528.8401747 | 635.8543812 | 693.1333183 | 449.2750164 | 385.1509951 | 467.8035354 |
| AP53_1852 | 192.4525431 | 177.2280617 | 196.7037787 | 186.0808792 | 161.4674209 | 171.1021497 |
| AP53_1853 | 210.7122752 | 203.3567895 | 212.9958647 | 221.9854507 | 202.3144322 | 195.5425389 |
| AP53_1854 | 468.0794271 | 460.6302214 | 407.5943294 | 541.0952703 | 570.344517 | 535.0426611 |
| AP53_1855 | 1938.882647 | 2116.072095 | 1928.986116 | 2684.702831 | 2860.89953 | 2967.762953 |
| AP53_1856 | 150.5352609 | 151.7310989 | 136.7938664 | 130.0098851 | 152.1886872 | 127.9047436 |
| AP53_1857 | 326.1766119 | 360.845693 | 169.40075 | 176.9336696 | 197.5872139 | 197.5743431 |
| AP53_1858 | 207.2180829 | 194.1330055 | 210.0796221 | 206.5631931 | 249.0199575 | 212.7675273 |

| | | | | | | |
|---|---|---|---|---|---|---|
| AP53_1859 | 345.7258899 | 399.4409265 | 372.2180858 | 317.1102762 | 326.1827732 | 288.1038085 |
| AP53_186 | 305.0226962 | 275.9245899 | 305.2894745 | 397.0161103 | 417.7354507 | 444.8693819 |
| AP53_1860 | 126.338434 | 112.2857192 | 120.9343216 | 146.416798 | 171.8997613 | 125.1349362 |
| AP53_1861 | 151.9669257 | 152.3329302 | 160.6999932 | 220.7276041 | 233.6176718 | 235.7974759 |
| AP53_1862 | 263.1957455 | 250.3884461 | 213.8414722 | 332.4156431 | 360.2027031 | 255.1947414 |
| AP53_1863 | 1161.538616 | 1188.27753 | 1133.416265 | 1089.672427 | 1121.60395 | 1009.653383 |
| AP53_1864 | 676.9035143 | 554.2191537 | 522.7539757 | 992.061628 | 960.5798559 | 835.9866605 |
| AP53_1865 | 860.424056 | 865.8533418 | 701.3007223 | 3325.331959 | 1618.814399 | 2321.316804 |
| AP53_1866 | 16.30883059 | 23.42392214 | 17.62343259 | 18.62035429 | 23.75895333 | 26.8666215 |
| AP53_1867 | 229.456121 | 264.194984 | 164.026875 | 104.7859406 | 108.2812228 | 77.47237529 |
| AP53_187 | 131.4065891 | 129.2232466 | 134.1714293 | 136.9346084 | 148.7349111 | 158.1615966 |
| AP53_188 | 260.6064853 | 225.9898777 | 143.9613094 | 196.6554807 | 179.2741732 | 195.7214525 |
| AP53_189 | 378.4291077 | 380.1294381 | 377.445348 | 364.0924303 | 365.8415296 | 417.2422901 |
| AP53_19 | 249.1208136 | 199.8277894 | 182.7082399 | 274.2673398 | 257.540185 | 245.386898 |
| AP53_190 | 458.5428259 | 503.5372257 | 499.0373717 | 449.0911299 | 458.6173176 | 517.1015403 |
| AP53_191 | 373.6823046 | 320.1035008 | 299.2477975 | 412.7619005 | 410.8766552 | 408.5643304 |
| AP53_192 | 523.2758157 | 610.4628619 | 662.4300439 | 480.8560751 | 457.468999 | 512.7182063 |
| AP53_193 | 89.09710937 | 87.96676934 | 88.04426934 | 70.83953375 | 76.84052197 | 69.72320734 |
| AP53_194 | 78.43301267 | 77.28607897 | 93.95242501 | 72.89148888 | 82.43468619 | 75.32697616 |
| AP53_195 | 118.4303078 | 111.3953708 | 130.570371 | 100.2634462 | 105.3281534 | 108.7223052 |
| AP53_196 | 433.0474194 | 393.6184854 | 390.1072686 | 422.9077437 | 445.4803748 | 431.3924565 |
| AP53_197 | 855.7642697 | 864.129973 | 922.767309 | 1095.743954 | 1116.979364 | 1170.117149 |
| AP53_198 | 1130.904954 | 1124.348263 | 1189.296918 | 1546.18316 | 1546.279421 | 1552.358195 |
| AP53_199 | 50.82177913 | 57.24931435 | 62.63380224 | 76.38061074 | 69.95231902 | 71.93225157 |
| AP53_2 | 324.0402386 | 315.6619293 | 309.6489585 | 348.4672684 | 370.8474396 | 302.4669897 |
| AP53_20 | 111.1416604 | 116.2699457 | 106.6730523 | 175.336445 | 160.9071971 | 135.0115575 |
| AP53_200 | 1669.768428 | 1739.004859 | 1024.844566 | 1475.33393 | 1104.602767 | 1124.327403 |
| AP53_201 | 112.0839181 | 97.10728853 | 90.3133962 | 115.9000002 | 95.18024463 | 109.9868557 |
| AP53_202 | 160.0707362 | 140.2471664 | 138.5957092 | 153.7333383 | 125.8530271 | 146.0456222 |
| AP53_203 | 82.88006048 | 62.15385063 | 75.19958686 | 92.2843954 | 77.29325715 | 81.60422417 |
| AP53_204 | 101.7388421 | 77.98965628 | 95.70641835 | 106.5766258 | 90.36734439 | 101.444201 |
| AP53_205 | 112.9333342 | 91.39230525 | 109.4999574 | 128.4505724 | 107.1226241 | 136.0184879 |
| AP53_206 | 126.5577836 | 86.12655907 | 112.6744051 | 131.1472346 | 108.6123581 | 129.7285449 |
| AP53_207 | 140.0211646 | 116.3288093 | 127.3616758 | 164.0300162 | 137.2835916 | 165.6827014 |
| AP53_208 | 145.2937059 | 149.8593999 | 148.8766508 | 157.5427416 | 165.4181897 | 162.8133134 |

| | | | | | | |
|---|---|---|---|---|---|---|
| AP53_209 | 193.0023642 | 178.149136 | 149.9136149 | 179.7915752 | 166.4926653 | 167.2919019 |
| AP53_21 | 78.47033977 | 73.39134199 | 55.98394213 | 68.60857816 | 91.50314094 | 68.71175832 |
| AP53_210 | 300.5268102 | 311.6665321 | 311.8950048 | 304.0730584 | 277.0686963 | 320.0855863 |
| AP53_211 | 382.8272982 | 460.2132263 | 476.3777346 | 423.2480408 | 427.3579037 | 489.9948026 |
| AP53_212 | 631.1883375 | 595.0955838 | 624.1654005 | 757.6116661 | 710.0276307 | 830.2969483 |
| AP53_213 | 643.1157493 | 629.4530292 | 701.2918619 | 807.3160603 | 764.9860796 | 837.4959181 |
| AP53_214 | 485.3638929 | 497.9829482 | 555.3350031 | 581.7958179 | 566.7081419 | 667.8745773 |
| AP53_215 | 659.9646148 | 694.7555354 | 744.6731564 | 813.8021053 | 785.1341449 | 867.21756 |
| AP53_216 | 481.3020646 | 438.4665666 | 402.7891421 | 548.9840733 | 496.5415211 | 510.8013486 |
| AP53_217 | 296.9714707 | 273.7390606 | 239.9664127 | 292.4242004 | 285.3498782 | 271.239479 |
| AP53_218 | 1171.193127 | 1100.917157 | 1086.165738 | 1225.371932 | 1138.4725 | 1161.75642 |
| AP53_219 | 3785.483963 | 4121.497371 | 4034.452516 | 2954.110622 | 2832.628007 | 3237.003122 |
| AP53_22 | 338.3375586 | 373.9498103 | 357.9660925 | 348.8829082 | 456.4184023 | 412.4238784 |
| AP53_220 | 2811.156002 | 3047.401666 | 3212.243424 | 2489.542447 | 2303.040307 | 2688.801212 |
| AP53_221 | 3584.769839 | 4411.834632 | 4520.535796 | 3072.590511 | 2822.177713 | 3347.25221 |
| AP53_222 | 6536.422164 | 8449.895613 | 8312.492786 | 5995.181458 | 6007.016758 | 6252.686143 |
| AP53_223 | 16.23503498 | 20.43323898 | 15.10579936 | 16.05951303 | 23.49786593 | 11.7424045 |
| AP53_224 | 317.9115032 | 338.0576734 | 308.7821093 | 321.4872359 | 336.18438 | 333.8056589 |
| AP53_225 | 247.2297045 | 289.9866247 | 286.4882396 | 221.847828 | 235.4164583 | 235.2318662 |
| AP53_226 | 184.7318008 | 180.2326257 | 181.3401802 | 203.5266423 | 186.8360949 | 206.7782561 |
| AP53_227 | 175.5186499 | 206.5720851 | 221.1381128 | 199.9874416 | 193.4657628 | 210.5278928 |
| AP53_228 | 280.0072869 | 275.1773693 | 238.600264 | 262.3777196 | 265.0499658 | 272.0317545 |
| AP53_229 | 289.0682997 | 295.8402327 | 307.2187506 | 299.0693025 | 323.9898633 | 352.9559018 |
| AP53_23 | 365.7767075 | 362.2554744 | 342.6991967 | 371.1506046 | 470.4552687 | 422.1106305 |
| AP53_230 | 50.15993275 | 34.15269944 | 34.5275414 | 36.67645543 | 46.46734069 | 36.20015559 |
| AP53_231 | 354.5014932 | 399.4911443 | 372.3550154 | 286.7152538 | 259.8429164 | 313.6894266 |
| AP53_232 | 426.4131412 | 470.0500946 | 481.4121379 | 378.4382954 | 329.6988111 | 388.4231215 |
| AP53_233 | 433.9420244 | 496.0985138 | 491.8064164 | 352.1684701 | 320.9491861 | 376.0819416 |
| AP53_234 | 449.979823 | 467.2710241 | 471.3009401 | 351.3533898 | 330.7161449 | 364.7601822 |
| AP53_235 | 457.4179762 | 478.9342289 | 503.4734185 | 356.803637 | 338.4448908 | 379.9564052 |
| AP53_236 | 531.4273112 | 451.5868395 | 506.2743162 | 602.6481534 | 575.8401867 | 621.2363805 |
| AP53_237 | 84.59024199 | 80.97637016 | 87.36371067 | 70.88172913 | 69.53437157 | 67.16655374 |
| AP53_238 | 660.2207123 | 631.0554366 | 641.6860798 | 769.316672 | 671.4396964 | 718.5622345 |
| AP53_239 | 402.671289 | 362.2256001 | 396.9116623 | 478.1623189 | 469.7931618 | 447.0579492 |
| AP53_24 | 391.479454 | 393.2431588 | 353.1035974 | 416.5379709 | 503.74555 | 489.1107479 |

| | | | | | | |
|---|---|---|---|---|---|---|
| AP53_240 | 482.502947 | 446.9090219 | 503.282215 | 600.9570068 | 584.5273699 | 592.6425752 |
| AP53_241 | 563.9001891 | 537.1896966 | 561.3264268 | 710.2951549 | 667.0856183 | 719.3337791 |
| AP53_242 | 535.5750197 | 492.2034355 | 541.4546265 | 676.609155 | 640.4779069 | 704.3566965 |
| AP53_243 | 16.88443638 | 15.2134752 | 18.8822492 | 19.46673403 | 22.91041928 | 13.73861327 |
| AP53_244 | 8.169888571 | 10.28253316 | 10.07053291 | 9.464891723 | 9.671463335 | 10.43937639 |
| AP53_245 | 46.71848017 | 42.24048797 | 42.32021568 | 45.72241879 | 52.43850621 | 48.36379555 |
| AP53_246 | 412.9012169 | 467.5125078 | 412.736918 | 511.1265552 | 558.1178303 | 524.3688154 |
| AP53_247 | 419.6212473 | 474.5252472 | 437.0962266 | 408.7583016 | 414.6214892 | 422.4961213 |
| AP53_248 | 435.1132667 | 480.2478441 | 456.8404371 | 423.8472526 | 456.8902744 | 402.9993224 |
| AP53_249 | 400.4688594 | 430.275945 | 417.5214308 | 460.3343082 | 433.3555874 | 492.3328646 |
| AP53_25 | 463.4715559 | 558.1798598 | 515.9651145 | 504.8083275 | 649.8394801 | 655.3937384 |
| AP53_250 | 387.1094264 | 398.4481601 | 382.1462071 | 408.1431194 | 413.724631 | 444.2614202 |
| AP53_251 | 256.0988916 | 233.2042911 | 187.0241826 | 245.7188168 | 261.1599235 | 237.845199 |
| AP53_252 | 153.0925711 | 156.5165133 | 141.9214214 | 155.9659441 | 185.5141776 | 175.4997049 |
| AP53_253 | 154.8051983 | 138.7255333 | 145.4905445 | 171.6614086 | 186.1255983 | 170.8204815 |
| AP53_254 | 302.8312615 | 394.7198129 | 284.4820207 | 199.5189732 | 183.9792231 | 199.3536017 |
| AP53_255 | 831.0026873 | 822.3681275 | 814.9605635 | 843.3074725 | 703.4679419 | 729.2710159 |
| AP53_256 | 257.5529718 | 223.089866 | 243.0855231 | 385.6130109 | 350.6070468 | 362.2057095 |
| AP53_257 | 283.1656181 | 226.3961018 | 263.777043 | 461.3365629 | 408.7304848 | 407.1273565 |
| AP53_258 | 208.9623428 | 156.2431117 | 153.1505719 | 285.2043247 | 256.5438003 | 289.9977415 |
| AP53_259 | 128.2958541 | 153.1334484 | 149.9672029 | 144.4216459 | 145.2719997 | 141.6550234 |
| AP53_26 | 766.6265895 | 791.1961809 | 825.4909505 | 962.248471 | 1149.171634 | 1177.75771 |
| AP53_260 | 286.5237689 | 312.8913677 | 278.729972 | 244.2589237 | 286.574042 | 260.6152745 |
| AP53_261 | 245.9499566 | 274.2611255 | 263.7136884 | 266.6284267 | 245.8863106 | 256.6169423 |
| AP53_262 | 228.1334138 | 260.8024321 | 255.1382535 | 198.4979383 | 209.8658777 | 202.2497579 |
| AP53_263 | 117.975001 | 125.4700776 | 121.0368041 | 107.113268 | 118.6521559 | 111.2173455 |
| AP53_264 | 783.8141581 | 896.4499367 | 789.9683355 | 787.7702188 | 670.2506572 | 665.1655483 |
| AP53_265 | 764.5353904 | 902.6322746 | 849.4107181 | 903.8429806 | 753.8614861 | 858.8088328 |
| AP53_266 | 661.5586047 | 704.1014473 | 630.8792835 | 537.0231905 | 508.548383 | 553.660969 |
| AP53_267 | 1111.798913 | 1463.707779 | 1173.239072 | 947.2800329 | 932.2758354 | 994.7795899 |
| AP53_269 | 300.1522533 | 282.9311233 | 261.650755 | 270.3285596 | 288.9129865 | 260.3397827 |
| AP53_27 | 233.1894159 | 227.0719842 | 238.0271158 | 252.3791536 | 322.6692137 | 287.8636372 |
| AP53_270 | 308.4745121 | 261.7894228 | 244.203563 | 301.2835131 | 297.564937 | 277.2679262 |
| AP53_271 | 604.5964293 | 636.4580577 | 652.108827 | 647.5789178 | 671.8923749 | 685.439651 |
| AP53_272 | 742.9924751 | 783.2361877 | 723.1771819 | 887.2461309 | 887.8734061 | 930.5369211 |

| | | | | | | |
|---|---|---|---|---|---|---|
| AP53_273 | 226.0186048 | 209.3544746 | 218.3833569 | 258.8463579 | 254.0345009 | 241.5938911 |
| AP53_274 | 1231.968204 | 1133.531853 | 1309.92081 | 1600.710421 | 1480.417511 | 1497.440495 |
| AP53_275 | 635.0893813 | 534.9818637 | 532.4879523 | 697.3557683 | 669.6331895 | 663.0509066 |
| AP53_276 | 397.0607364 | 296.6200193 | 280.4424491 | 460.5146912 | 422.5804964 | 430.4196603 |
| AP53_277 | 729.8637339 | 671.2047295 | 695.993207 | 843.2725722 | 810.0823133 | 844.3161539 |
| AP53_278 | 700.5944706 | 739.210206 | 730.051086 | 757.6412933 | 747.8694492 | 738.9837943 |
| AP53_279 | 1091.146734 | 1144.726981 | 1096.837977 | 1496.168551 | 1359.031966 | 1327.768576 |
| AP53_28 | 560.3290445 | 589.6895432 | 595.3904285 | 674.8574766 | 846.0005097 | 870.5896608 |
| AP53_280 | 466.3025795 | 450.2503158 | 462.9196579 | 502.1853128 | 502.5511325 | 516.4536772 |
| AP53_281 | 268.9436139 | 288.5436122 | 277.5537119 | 262.7217764 | 275.8080586 | 269.6342473 |
| AP53_282 | 573.537741 | 513.6228868 | 446.3899393 | 455.8287058 | 408.6174137 | 403.8768387 |
| AP53_283 | 851.5962468 | 916.9662321 | 1040.789576 | 773.8854759 | 757.777797 | 855.7762292 |
| AP53_284 | 516.4818334 | 521.4259873 | 473.5015381 | 463.1055569 | 449.4087732 | 488.2578772 |
| AP53_285 | 160.9524103 | 170.5330787 | 191.0171082 | 151.4540653 | 146.5242189 | 167.8357384 |
| AP53_286 | 317.1955287 | 331.1776915 | 371.3547272 | 258.9561845 | 243.3667863 | 293.2017729 |
| AP53_287 | 228.1913052 | 231.2414923 | 240.2169359 | 199.6288768 | 173.2179848 | 201.183831 |
| AP53_288 | 195.6312131 | 208.0209875 | 240.6138041 | 186.8630596 | 166.5110319 | 203.6407438 |
| AP53_289 | 122.3798004 | 126.5607518 | 148.5100275 | 115.4339598 | 115.5233101 | 104.3913908 |
| AP53_29 | 579.4665562 | 566.7505426 | 603.7714319 | 809.7376909 | 915.588933 | 951.8736768 |
| AP53_290 | 175.5230238 | 140.6976714 | 173.0417527 | 158.1109388 | 144.3591115 | 152.313477 |
| AP53_291 | 246.3843026 | 227.7296599 | 287.7490585 | 226.682574 | 212.6869036 | 232.9590945 |
| AP53_292 | 240.0347779 | 232.5458292 | 245.2655307 | 184.7285243 | 157.8062075 | 188.2741082 |
| AP53_293 | 155.8060206 | 154.76912 | 188.822492 | 164.4669723 | 142.5151502 | 152.16555 |
| AP53_294 | 191.9944107 | 182.9734124 | 174.0056014 | 252.4339139 | 253.3546467 | 226.3242593 |
| AP53_295 | 108.0791715 | 127.5496528 | 135.3093943 | 111.6500934 | 188.6634212 | 120.1397883 |
| AP53_296 | 173.4348648 | 221.3977032 | 203.7165279 | 168.6431408 | 236.1620928 | 165.0345569 |
| AP53_297 | 141.9327621 | 138.3772586 | 122.2733271 | 139.5561644 | 167.5509249 | 132.5231707 |
| AP53_298 | 140.1989011 | 142.3840305 | 132.9310344 | 167.2629369 | 174.4769468 | 173.576858 |
| AP53_299 | 293.6494954 | 332.2382043 | 320.9558045 | 379.4444006 | 412.5210243 | 368.2565821 |
| AP53_3 | 50.58353874 | 54.88266668 | 48.52165856 | 57.19475217 | 68.65411837 | 49.40349821 |
| AP53_30 | 848.7130448 | 971.2907459 | 983.5225625 | 1054.806662 | 1360.072087 | 1450.818028 |
| AP53_300 | 260.7414264 | 220.6557614 | 162.1775404 | 285.1241751 | 160.9268391 | 226.241886 |
| AP53_301 | 713.6778487 | 812.2620686 | 789.4945011 | 748.8034776 | 580.6242847 | 689.6725521 |
| AP53_302 | 529.1876678 | 542.8066928 | 535.8401215 | 754.908609 | 729.0386563 | 709.6182815 |
| AP53_303 | 102.4688638 | 93.2759852 | 109.3119347 | 117.5196918 | 137.2293721 | 121.3796972 |

| | | | | | | |
|---|---|---|---|---|---|---|
| AP53_304 | 164.2635011 | 174.094679 | 195.4784849 | 185.4629607 | 227.5662248 | 228.6906666 |
| AP53_305 | 93.05935339 | 88.51815307 | 107.7320353 | 91.82267669 | 115.8480889 | 108.0240803 |
| AP53_306 | 72.07201945 | 74.33257908 | 85.59952973 | 72.83405738 | 92.005799 | 90.88891024 |
| AP53_307 | 174.5208081 | 213.3974718 | 195.6232357 | 162.4627668 | 146.2929032 | 167.8657117 |
| AP53_308 | 237.8044013 | 270.7187733 | 262.3142847 | 238.1666 | 224.4489326 | 239.6484711 |
| AP53_309 | 331.9874894 | 424.9423754 | 423.4659088 | 265.4408082 | 277.9959165 | 296.5795879 |
| AP53_31 | 247.9611859 | 227.1891929 | 212.7991467 | 248.3092599 | 316.7101129 | 249.7743411 |
| AP53_310 | 101.5427642 | 112.5685711 | 132.8535688 | 98.96132352 | 102.2592313 | 90.65136274 |
| AP53_311 | 4251.885987 | 3466.259961 | 3281.562384 | 5912.886202 | 5725.321961 | 5461.691556 |
| AP53_312 | 679.0896954 | 590.5852175 | 685.1793559 | 926.785816 | 1058.085064 | 853.7851258 |
| AP53_313 | 748.613336 | 606.6937079 | 590.9787732 | 987.9487293 | 965.8879071 | 1016.22535 |
| AP53_314 | 770.8386522 | 665.9492847 | 694.8667707 | 1147.598174 | 1187.657072 | 1152.161512 |
| AP53_315 | 534.4179939 | 420.8256527 | 423.2980666 | 681.6178178 | 686.3697627 | 596.7646532 |
| AP53_316 | 608.5722989 | 565.0719361 | 613.844756 | 998.3433759 | 1069.538597 | 1017.906346 |
| AP53_317 | 718.2281082 | 405.9609873 | 361.6999737 | 876.1597686 | 786.4653531 | 688.2875633 |
| AP53_318 | 1018.822241 | 1022.381756 | 1070.990685 | 1594.896824 | 1650.516574 | 1780.231698 |
| AP53_319 | 627.9815183 | 551.5783412 | 539.5422536 | 863.565772 | 903.7211472 | 773.8370057 |
| AP53_32 | 69.3756447 | 46.51739285 | 44.91987706 | 62.81028602 | 74.58167679 | 37.60041525 |
| AP53_320 | 272.5630444 | 305.9253925 | 272.5517799 | 246.6592387 | 259.5706057 | 264.6536676 |
| AP53_321 | 619.2357369 | 709.8404196 | 631.523965 | 505.1251192 | 489.2975322 | 506.7508669 |
| AP53_322 | 16.65145309 | 15.00648915 | 11.32934952 | 14.81164546 | 15.45544158 | 13.08439359 |
| AP53_323 | 223.860598 | 265.6321067 | 252.2559031 | 230.0558266 | 182.6807721 | 202.0306221 |
| AP53_324 | 989.2530501 | 677.0860658 | 639.1976827 | 1525.717632 | 842.8053657 | 1262.418159 |
| AP53_325 | 91.23351703 | 36.22256001 | 46.07268806 | 115.1076447 | 55.5789801 | 65.94534367 |
| AP53_326 | 822.8928406 | 726.6741003 | 862.8082641 | 1155.394392 | 873.2842641 | 1076.459429 |
| AP53_327 | 478.1739771 | 458.7012965 | 498.8598131 | 428.1442883 | 478.4904189 | 499.5047034 |
| AP53_328 | 439.3441979 | 457.5897217 | 478.7165144 | 442.5796607 | 498.3209042 | 477.937213 |
| AP53_329 | 187.873954 | 172.7994256 | 200.1518416 | 215.4799569 | 239.9543003 | 236.934773 |
| AP53_33 | 646.9488508 | 708.5838699 | 704.9606691 | 836.6002253 | 911.0223783 | 999.2955146 |
| AP53_330 | 1046.029553 | 1058.982314 | 1117.715746 | 1297.350183 | 1281.464782 | 1462.784123 |
| AP53_331 | 50.49632575 | 48.07721601 | 61.1098247 | 43.08842316 | 42.42670237 | 46.62802078 |
| AP53_332 | 21.72097305 | 22.41698053 | 32.49171939 | 19.48270346 | 18.25148706 | 21.77440593 |
| AP53_333 | 167.198154 | 166.623776 | 186.1856809 | 156.9057827 | 143.2968149 | 180.0381588 |
| AP53_334 | 158.1326815 | 140.801313 | 167.5538972 | 150.6867489 | 136.8065936 | 181.4957908 |
| AP53_335 | 292.220885 | 271.6243578 | 342.1972623 | 351.4280447 | 290.9759807 | 376.5173737 |

| | | | | | | |
|---|---|---|---|---|---|---|
| AP53_336 | 986.5103462 | 1026.642597 | 1019.190089 | 1115.400361 | 896.5387624 | 1150.175538 |
| AP53_337 | 49.97720076 | 45.65192483 | 46.55623349 | 54.90272815 | 57.19748023 | 50.7868314 |
| AP53_338 | 9055.616275 | 7261.123155 | 4919.98937 | 9019.330291 | 8530.767044 | 8200.240575 |
| AP53_339 | 126.9009965 | 128.9186182 | 116.4903039 | 106.2108158 | 113.743029 | 118.6845785 |
| AP53_34 | 66.45974697 | 58.14674106 | 38.65892074 | 51.89644199 | 70.89725264 | 49.41080209 |
| AP53_340 | 93.05626869 | 107.4602614 | 101.0015212 | 73.60184607 | 83.2728021 | 82.43167959 |
| AP53_341 | 15.51949777 | 13.29688912 | 14.72337406 | 15.32054467 | 20.30037151 | 13.21689377 |
| AP53_342 | 571.2202178 | 700.1303384 | 734.6094095 | 598.1083501 | 528.3134604 | 744.0658486 |
| AP53_343 | 137.0424314 | 147.8401012 | 129.710876 | 130.1498907 | 117.4736957 | 133.3824649 |
| AP53_344 | 232.865328 | 274.0076185 | 183.5641442 | 121.2787173 | 102.6833607 | 130.0820598 |
| AP53_345 | 290.8662026 | 335.2996456 | 259.4566912 | 215.1911409 | 163.7342951 | 223.9059833 |
| AP53_346 | 535.4529215 | 620.2180396 | 640.0281415 | 480.2307353 | 367.9037924 | 504.9703631 |
| AP53_347 | 159.4144095 | 170.4896815 | 157.0528961 | 158.2312618 | 130.7427348 | 170.9831002 |
| AP53_348 | 358.4987705 | 449.7883352 | 424.5024528 | 243.3020461 | 237.1845929 | 273.6596661 |
| AP53_349 | 399.3148461 | 401.1501673 | 360.1712486 | 309.2717323 | 279.6034353 | 321.3102706 |
| AP53_35 | 571.5955592 | 473.7867338 | 381.455456 | 554.4422721 | 527.6199995 | 541.1280828 |
| AP53_350 | 471.8596953 | 465.891117 | 426.0195082 | 370.794934 | 354.7680445 | 386.4257572 |
| AP53_351 | 503.3573354 | 577.6249267 | 585.6752814 | 502.209634 | 454.1851638 | 552.0711721 |
| AP53_352 | 528.8504385 | 449.5410342 | 382.6802505 | 568.4108153 | 445.8525389 | 440.5997377 |
| AP53_353 | 845.7674272 | 825.0694224 | 758.9964838 | 1081.485224 | 1029.868916 | 990.1130239 |
| AP53_354 | 969.2387069 | 994.1986245 | 804.1531005 | 896.5637999 | 727.0654435 | 708.9301147 |
| AP53_355 | 256.3586429 | 250.5044481 | 239.2583841 | 194.5274429 | 205.681848 | 194.9899051 |
| AP53_356 | 298.3455672 | 303.7102786 | 277.4166802 | 233.6008084 | 251.1065318 | 267.766376 |
| AP53_357 | 840.7268149 | 891.6405329 | 849.350243 | 1062.914513 | 1143.281339 | 1108.41838 |
| AP53_358 | 135.7769358 | 150.2558761 | 159.5526514 | 128.434432 | 147.2767324 | 147.8928028 |
| AP53_359 | 230.734285 | 214.6109282 | 172.4901534 | 267.3691899 | 287.0510905 | 201.029965 |
| AP53_36 | 463.2443823 | 455.4118572 | 468.486709 | 633.3934963 | 667.668434 | 610.0195223 |
| AP53_360 | 3074.383466 | 2943.210545 | 2998.607553 | 3288.00648 | 2996.042258 | 3282.045058 |
| AP53_361 | 3812.962883 | 4282.829504 | 4620.732671 | 4445.1128 | 4124.391969 | 5107.737658 |
| AP53_362 | 578.7325332 | 695.7116546 | 696.2965377 | 576.6527978 | 615.7633494 | 668.8009706 |
| AP53_363 | 773.1066526 | 875.2616865 | 930.4685124 | 768.8404354 | 804.6475014 | 781.2789895 |
| AP53_364 | 492.3240815 | 547.0873084 | 570.956962 | 483.7385748 | 501.1097923 | 507.079726 |
| AP53_365 | 311.1657159 | 234.4665002 | 187.4896274 | 335.4650996 | 325.238109 | 312.5938595 |
| AP53_366 | 397.6476636 | 346.7303938 | 389.1841364 | 494.1917263 | 514.5416181 | 475.763089 |
| AP53_367 | 1195.360827 | 1244.69542 | 1375.164496 | 1435.494411 | 1467.576247 | 1544.16434 |

| | | | | | | |
|---|---|---|---|---|---|---|
| AP53_368 | 665.4626212 | 753.0615064 | 737.065882 | 834.5218316 | 667.1975834 | 801.4113583 |
| AP53_369 | 90.06344658 | 122.8058302 | 118.6084987 | 67.99973985 | 62.80602436 | 80.004916 |
| AP53_37 | 508.2804661 | 373.9979321 | 257.7427016 | 548.0308821 | 503.2109949 | 487.7207709 |
| AP53_370 | 141.2032943 | 198.4694434 | 193.0709981 | 122.8132269 | 118.7063777 | 140.916193 |
| AP53_371 | 258.3291026 | 310.1174836 | 280.8222701 | 199.0521992 | 198.2298214 | 225.0042767 |
| AP53_372 | 246.7486567 | 290.9523864 | 295.8959522 | 192.9247938 | 200.5285609 | 216.1811205 |
| AP53_373 | 524.6290593 | 616.4861651 | 623.834149 | 432.8708694 | 435.1078265 | 455.013069 |
| AP53_374 | 390.1810791 | 484.5166077 | 501.0565147 | 343.5556507 | 369.9394979 | 423.7368519 |
| AP53_375 | 9.401561166 | 10.62528427 | 6.042319745 | 8.463797406 | 7.919651108 | 5.800747823 |
| AP53_376 | 14.6798791 | 14.39178223 | 17.43756705 | 11.70162594 | 15.03442876 | 12.90876414 |
| AP53_377 | 25.59867081 | 22.25318454 | 24.79390697 | 24.07003988 | 24.34612456 | 22.95565761 |
| AP53_378 | 25.64724514 | 22.37545479 | 22.94551802 | 28.49835582 | 24.70415581 | 23.6512836 |
| AP53_379 | 39.14119343 | 40.18289323 | 22.35658306 | 18.05610113 | 21.49619587 | 22.71450727 |
| AP53_38 | 441.6802098 | 391.8644651 | 403.9270337 | 551.9768417 | 570.7824803 | 534.7538614 |
| AP53_380 | 31.80465867 | 36.8365017 | 45.82945909 | 28.40393028 | 39.98177376 | 36.63630204 |
| AP53_381 | 43.90667389 | 42.11925582 | 51.2894583 | 33.85518963 | 41.04536787 | 28.1162318 |
| AP53_382 | 54.42091065 | 49.30031544 | 28.64216503 | 34.7345452 | 28.651799 | 41.87005948 |
| AP53_383 | 100.0250809 | 110.3079469 | 99.04274111 | 69.14762787 | 73.80645204 | 91.93638059 |
| AP53_384 | 26.94324954 | 24.50802996 | 34.87189853 | 26.2917962 | 29.42788292 | 34.68756257 |
| AP53_385 | 19.20705622 | 18.60695714 | 21.14811911 | 19.06581732 | 21.43664962 | 21.49972462 |
| AP53_386 | 4.604846286 | 6.037093335 | 10.32229623 | 7.617417666 | 9.333874521 | 8.853772993 |
| AP53_387 | 3.558290312 | 3.951552001 | 3.570461668 | 2.154421158 | 2.159904848 | 1.665286456 |
| AP53_39 | 1004.049883 | 999.4301801 | 939.2515134 | 1324.886916 | 1090.631117 | 1239.417326 |
| AP53_390 | 71.88165052 | 59.92660328 | 61.02742943 | 70.1479529 | 73.72063803 | 63.67389295 |
| AP53_391 | 41.21078726 | 34.41550196 | 33.47037792 | 37.16462951 | 42.82713416 | 31.20260331 |
| AP53_392 | 1.644587959 | 1.379907048 | 0.719323779 | 0.403037972 | 2.02031916 | 2.616878717 |
| AP53_393 | 9.943431815 | 9.616890657 | 11.42064831 | 8.742823695 | 10.89107437 | 9.219465019 |
| AP53_394 | 16.49856138 | 15.43746986 | 18.28264575 | 19.21447044 | 21.17789711 | 17.60379416 |
| AP53_395 | 19.84325209 | 16.77676464 | 17.6896861 | 17.81852086 | 22.10654492 | 18.31815102 |
| AP53_396 | 12.37122883 | 2.595049075 | 3.607355072 | 5.053013377 | 6.332343637 | 5.741510021 |
| AP53_397 | 12.31528658 | 9.771667351 | 4.566869575 | 6.692304926 | 12.23467696 | 12.78010536 |
| AP53_398 | 8.327913495 | 7.090373448 | 3.696099844 | 5.762585468 | 8.304801314 | 4.092352888 |
| AP53_4 | 33.38513557 | 38.03368801 | 26.43514889 | 27.93053144 | 27.57735654 | 23.35564255 |
| AP53_40 | 1195.120615 | 894.1309208 | 666.883166 | 1665.944973 | 993.2604149 | 1477.17786 |
| AP53_401 | 21.01540587 | 18.47593665 | 18.95828511 | 23.63046793 | 21.29877407 | 24.71106279 |

| | | | | | | |
|---|---|---|---|---|---|---|
| AP53_402 | 19.26065514 | 24.79852185 | 20.62522605 | 27.67010691 | 22.19242893 | 28.1817708 |
| AP53_403 | 14.23866944 | 17.92063495 | 21.46613594 | 16.03666877 | 19.65026215 | 19.76432084 |
| AP53_404 | 13.81453886 | 6.664951041 | 8.761363631 | 10.83366068 | 11.20064942 | 14.2881578 |
| AP53_405 | 67.95733359 | 64.42351503 | 59.74369783 | 51.01707641 | 59.42674435 | 52.0387612 |
| AP53_406 | 126.1744566 | 144.6802542 | 142.4104708 | 122.4184031 | 120.331966 | 130.3509007 |
| AP53_407 | 236.8705021 | 228.7509547 | 286.1709768 | 238.183228 | 228.0756667 | 275.6049085 |
| AP53_408 | 656.3284653 | 739.4897081 | 768.1947229 | 661.2574014 | 638.0434058 | 767.314805 |
| AP53_409 | 671.5786499 | 637.7354836 | 776.3545879 | 696.4556911 | 675.1175664 | 829.0574229 |
| AP53_41 | 128.346704 | 111.2285308 | 116.1390063 | 169.7089796 | 143.856121 | 192.3405857 |
| AP53_410 | 1175.255864 | 1062.100481 | 901.9500709 | 1081.009061 | 1008.359194 | 1080.539035 |
| AP53_411 | 1275.533474 | 1195.607235 | 1253.331304 | 1365.236834 | 1441.584605 | 1588.838591 |
| AP53_412 | 1126.932815 | 925.7186041 | 864.9038456 | 1157.04451 | 1130.182079 | 1208.645695 |
| AP53_413 | 906.2998101 | 785.0361064 | 897.1697546 | 1099.150872 | 1178.280823 | 1215.181309 |
| AP53_414 | 797.0541227 | 943.9330823 | 1003.766381 | 765.8169283 | 737.9103493 | 780.6415747 |
| AP53_415 | 566.06824 | 585.3284357 | 646.6162077 | 596.3005487 | 674.2138489 | 659.1570265 |
| AP53_416 | 365.8631161 | 409.9504116 | 396.6597403 | 360.6765597 | 411.911177 | 383.1546588 |
| AP53_417 | 248.4977729 | 236.2320483 | 260.3672732 | 260.9807822 | 300.9137893 | 274.2387269 |
| AP53_418 | 470.1694243 | 478.1377921 | 512.9977419 | 492.5124015 | 470.0609246 | 433.8203385 |
| AP53_419 | 742.1166125 | 803.1618441 | 895.9372082 | 805.8907639 | 744.6459599 | 812.9298372 |
| AP53_42 | 1784.601472 | 2211.756548 | 1825.601849 | 1272.527831 | 1126.820262 | 1330.289026 |
| AP53_420 | 724.9452757 | 811.3853442 | 866.8725856 | 680.1900391 | 675.8737768 | 710.4102823 |
| AP53_421 | 451.4886339 | 434.106776 | 430.1082394 | 411.8372319 | 386.3624489 | 421.1529392 |
| AP53_422 | 1053.514157 | 1001.961696 | 981.1057917 | 1347.345047 | 1233.661483 | 1354.773043 |
| AP53_423 | 954.7553765 | 889.2679181 | 931.7732286 | 1334.769381 | 1244.580163 | 1312.155914 |
| AP53_424 | 91.80174326 | 126.6860817 | 85.21220154 | 73.35291086 | 55.69864516 | 61.0605034 |
| AP53_425 | 177.6205755 | 262.4123236 | 241.1566272 | 93.96382491 | 86.05811357 | 101.3717431 |
| AP53_426 | 19.8592691 | 17.0716597 | 15.10579936 | 15.54679403 | 13.63806988 | 17.87541625 |
| AP53_427 | 20.95301396 | 19.64202417 | 22.75350532 | 25.07267181 | 20.16599744 | 22.38033514 |
| AP53_428 | 22.27726209 | 18.02832758 | 19.42997857 | 22.4839809 | 21.37528516 | 23.42206333 |
| AP53_429 | 4.604846286 | 6.664951041 | 6.64655172 | 4.401174651 | 2.375895333 | 2.198178122 |
| AP53_43 | 1651.300081 | 2192.556757 | 2157.310523 | 1151.481414 | 1012.962894 | 1206.193274 |
| AP53_430 | 118.4575667 | 125.5967616 | 134.966176 | 141.5641937 | 140.4179841 | 167.9721838 |
| AP53_431 | 92.06925236 | 78.88855617 | 93.50344558 | 114.017117 | 112.2675816 | 114.5324779 |
| AP53_432 | 156.800564 | 172.3844723 | 170.8957302 | 180.9519096 | 175.2785089 | 185.995443 |
| AP53_433 | 312.3329284 | 365.2212211 | 369.1041524 | 228.4153933 | 199.4950089 | 234.8124169 |

| | | | | | | |
|---|---|---|---|---|---|---|
| AP53_434 | 379.1340459 | 421.1776157 | 417.3018942 | 332.0961395 | 313.4119769 | 338.3780846 |
| AP53_435 | 234.8471606 | 250.3918 | 196.9348658 | 231.3437958 | 213.4534537 | 231.4871529 |
| AP53_436 | 42.90879493 | 55.00811277 | 67.74722139 | 29.23857286 | 31.3700466 | 38.46018721 |
| AP53_437 | 79.15067355 | 84.14684872 | 98.52053551 | 57.03738727 | 52.8536038 | 59.05275126 |
| AP53_438 | 34.61741838 | 31.17180868 | 26.11601228 | 53.16695272 | 49.65716854 | 37.99081321 |
| AP53_439 | 4.786824747 | 4.132608221 | 4.464256857 | 6.166065464 | 7.523085368 | 4.469377053 |
| AP53_44 | 2506.20973 | 3373.225901 | 3245.71339 | 1618.701254 | 1445.118754 | 1709.752807 |
| AP53_440 | 4.463881603 | 4.731109879 | 4.130973704 | 6.909222373 | 6.372663866 | 4.859917618 |
| AP53_441 | 16.68082073 | 15.25782936 | 18.25027188 | 17.13487148 | 17.80189797 | 15.1031286 |
| AP53_442 | 13.89151957 | 13.8284302 | 14.55482796 | 16.8246915 | 14.54629795 | 11.80379336 |
| AP53_443 | 20.57790684 | 15.64814592 | 17.44719827 | 19.42441505 | 17.60708148 | 17.31065271 |
| AP53_444 | 8.238357808 | 7.380346602 | 9.629947094 | 9.574670816 | 7.636806426 | 6.926550855 |
| AP53_445 | 7.918940203 | 7.156699735 | 11.3522371 | 10.56692282 | 10.02812965 | 10.38028558 |
| AP53_446 | 11.00510544 | 11.14829049 | 12.52294248 | 14.12094697 | 14.28878577 | 11.3420676 |
| AP53_447 | 17.05392824 | 15.06427061 | 18.74168656 | 21.21199846 | 19.41311146 | 18.18178761 |
| AP53_448 | 1.644587959 | 1.655888457 | 1.72637707 | 2.176405047 | 3.636574489 | 2.616878717 |
| AP53_449 | 4.126984879 | 2.733778114 | 1.710090494 | 3.513274395 | 5.443425964 | 4.147505891 |
| AP53_45 | 2399.403996 | 2902.195414 | 3227.910965 | 1457.14104 | 1311.3708 | 1492.281697 |
| AP53_450 | 2.325977344 | 2.927449862 | 2.008955414 | 2.770757207 | 3.732681743 | 2.529872519 |
| AP53_451 | 1.074464133 | 1.352308907 | 0.201410658 | 0.677103793 | 0.678827238 | 0 |
| AP53_452 | 1.131699511 | 1.326114061 | 1.689801285 | 1.319778579 | 1.725831961 | 1.179813117 |
| AP53_453 | 1.365844237 | 1.473460068 | 2.048243982 | 0 | 2.013470621 | 1.862862816 |
| AP53_454 | 800.1490328 | 796.3224974 | 753.6447821 | 1099.120463 | 1154.678411 | 1134.818524 |
| AP53_455 | 1328.520726 | 1246.056064 | 1241.045085 | 2035.128384 | 2019.843791 | 1881.381864 |
| AP53_456 | 1680.507255 | 1575.68136 | 1339.609753 | 2220.592665 | 2311.098187 | 2308.087029 |
| AP53_457 | 3.180389944 | 4.341069448 | 4.17319749 | 4.347164427 | 6.867512912 | 4.134057429 |
| AP53_458 | 14.13241071 | 14.86746866 | 16.79674705 | 16.23280547 | 15.07097786 | 13.87531588 |
| AP53_459 | 19.2585861 | 17.46807567 | 22.02340842 | 24.67948403 | 15.2260315 | 22.08449983 |
| AP53_46 | 2682.074498 | 3252.004107 | 3497.481589 | 1565.132723 | 1462.103523 | 1779.89171 |
| AP53_460 | 195.3597381 | 240.7029965 | 242.1470996 | 132.1116046 | 119.9431586 | 134.2183321 |
| AP53_461 | 245.438307 | 289.622418 | 298.5455256 | 214.2725729 | 188.158568 | 214.8219529 |
| AP53_462 | 451.399632 | 514.850793 | 559.9079946 | 402.6279176 | 341.2518358 | 419.5813238 |
| AP53_463 | 48.42157443 | 47.97900492 | 46.44370769 | 47.70166521 | 51.9541022 | 51.29885713 |
| AP53_464 | 47.68719819 | 38.77489698 | 44.29600952 | 41.38525849 | 45.71816896 | 41.9818123 |
| AP53_465 | 87.85171718 | 77.2047994 | 80.66156311 | 74.25244488 | 70.99811449 | 83.36086117 |

| | | | | | | |
|---|---|---|---|---|---|---|
| AP53_466 | 119.4836431 | 102.6432964 | 111.2740884 | 101.4230207 | 101.1452584 | 112.6855522 |
| AP53_467 | 447.7027942 | 417.3051987 | 391.806671 | 504.8406218 | 442.4231566 | 489.4717706 |
| AP53_468 | 324.1368129 | 365.9320946 | 371.1208847 | 312.2888173 | 314.944265 | 323.9452421 |
| AP53_469 | 1214.749595 | 1219.028462 | 869.745448 | 692.403734 | 573.0868412 | 564.3399603 |
| AP53_47 | 2427.199623 | 2361.866708 | 2296.081503 | 1306.155058 | 1100.539535 | 1208.604029 |
| AP53_470 | 240.0381371 | 298.8896133 | 305.982732 | 209.4879139 | 218.8048273 | 241.9490242 |
| AP53_471 | 252.2414943 | 277.0777741 | 263.9376313 | 210.7833381 | 214.8069753 | 240.2688987 |
| AP53_472 | 2439.809491 | 2908.950204 | 2454.111404 | 1883.985497 | 1858.382809 | 2141.814581 |
| AP53_473 | 565.7594102 | 678.9808226 | 648.3350801 | 628.8247681 | 564.6161921 | 716.1160132 |
| AP53_474 | 462.1228912 | 471.0604611 | 523.7064441 | 569.2880348 | 493.9773592 | 588.5762832 |
| AP53_475 | 536.4138035 | 567.7353596 | 537.6998141 | 567.1988938 | 541.0652396 | 492.9737701 |
| AP53_476 | 781.1332781 | 787.9799977 | 694.8667707 | 852.890354 | 694.2551296 | 802.9242699 |
| AP53_477 | 589.9050452 | 632.7890588 | 542.2186929 | 474.3290252 | 510.9068159 | 472.9939416 |
| AP53_478 | 123.83907 | 122.2423481 | 110.1403429 | 126.5460971 | 116.3233082 | 113.7622389 |
| AP53_479 | 240.1980134 | 278.2635635 | 285.1087586 | 238.5646579 | 228.7283852 | 271.9540882 |
| AP53_48 | 3412.891835 | 3784.785053 | 3847.118711 | 1774.306329 | 1530.165137 | 1842.487416 |
| AP53_480 | 48.19025183 | 45.48879629 | 49.18167235 | 63.7737293 | 48.14937385 | 36.63630204 |
| AP53_481 | 329.1492239 | 349.1650714 | 220.6297738 | 282.5239416 | 304.038115 | 315.7946035 |
| AP53_482 | 171.9469198 | 162.153758 | 127.9171946 | 181.1612807 | 161.4020082 | 153.9504182 |
| AP53_483 | 869.9275875 | 1059.677173 | 1027.770682 | 844.6122006 | 843.7631732 | 885.5612165 |
| AP53_484 | 111.6407501 | 115.575238 | 113.067661 | 86.88748833 | 88.51816972 | 84.40955304 |
| AP53_485 | 127.9615939 | 123.2263628 | 143.650342 | 124.2713331 | 135.7654476 | 130.8690982 |
| AP53_486 | 282.6224408 | 254.6445969 | 278.0411195 | 256.2414665 | 319.2609353 | 252.6759956 |
| AP53_487 | 626.5530212 | 604.7626136 | 674.7792716 | 815.9460861 | 958.1213168 | 840.4913335 |
| AP53_488 | 298.8268623 | 251.6891943 | 160.452413 | 262.8861456 | 279.5664826 | 213.6686096 |
| AP53_489 | 197.6246531 | 293.7477128 | 261.4741937 | 114.8658219 | 120.0069581 | 135.9686567 |
| AP53_49 | 2661.463853 | 3310.941375 | 3343.647901 | 1601.754247 | 1537.09218 | 1883.324398 |
| AP53_490 | 291.7601826 | 364.1499236 | 297.8674812 | 174.169081 | 173.153979 | 169.2997864 |
| AP53_491 | 221.6801782 | 239.2500088 | 253.3053731 | 174.671619 | 160.6911352 | 175.5107845 |
| AP53_492 | 214.2020997 | 275.7985719 | 289.6285265 | 134.5179534 | 126.1204439 | 140.4086276 |
| AP53_493 | 310.7160749 | 305.6205706 | 321.6418115 | 328.7182881 | 300.9067255 | 320.5774597 |
| AP53_494 | 145.8923653 | 160.2315596 | 160.6190761 | 110.606896 | 106.6158074 | 127.0633158 |
| AP53_495 | 101.894471 | 102.6562764 | 99.31259582 | 78.8753886 | 94.60251932 | 75.22134355 |
| AP53_496 | 8761.052139 | 10398.31278 | 9170.091702 | 5966.733024 | 4714.871275 | 5321.202701 |
| AP53_497 | 1335.769442 | 1929.560023 | 1796.03775 | 1244.01095 | 1233.828875 | 1314.779859 |

| | | | | | | |
|---|---|---|---|---|---|---|
| AP53_498 | 0 | 0 | 0 | 0 | 1.339790601 | 0 |
| AP53_499 | 199.6737352 | 204.35854 | 225.5237182 | 195.6945002 | 174.1142436 | 174.2447424 |
| AP53_5 | 245.1894968 | 259.8287073 | 272.1074235 | 225.7922727 | 222.0786958 | 231.4882472 |
| AP53_50 | 2574.2926 | 3168.161589 | 3044.037496 | 1442.770798 | 1333.797142 | 1620.094444 |
| AP53_500 | 270.4483784 | 298.6187847 | 299.6990594 | 228.9034009 | 222.7401874 | 232.0909734 |
| AP53_501 | 255.6542438 | 289.9951323 | 317.8372081 | 217.1120772 | 219.8645998 | 218.2573772 |
| AP53_502 | 145.3404609 | 135.8346 | 131.905998 | 119.0977206 | 113.9460006 | 110.5631258 |
| AP53_503 | 211.2234503 | 218.2701358 | 225.3687808 | 204.3402517 | 198.5647877 | 201.0353763 |
| AP53_504 | 179.9102735 | 242.9438676 | 191.8085222 | 161.2714266 | 177.2054824 | 181.0514926 |
| AP53_505 | 813.4327042 | 890.1349724 | 819.616342 | 865.5416265 | 841.6812332 | 944.1634014 |
| AP53_506 | 167.757879 | 157.7208818 | 150.1935051 | 250.0899173 | 252.6689034 | 203.5411419 |
| AP53_507 | 19.34976682 | 19.15922828 | 20.66359347 | 22.889515 | 22.57420767 | 24.04977375 |
| AP53_508 | 48.40130402 | 40.34717268 | 48.73549868 | 75.8034994 | 69.24533321 | 77.41758716 |
| AP53_509 | 50.50334288 | 37.4161003 | 42.48144342 | 69.98036323 | 54.35819644 | 71.30743846 |
| AP53_51 | 2167.280915 | 2728.976173 | 2560.104605 | 1224.674686 | 1260.257524 | 1314.924667 |
| AP53_510 | 52.68735167 | 41.51149822 | 47.63423848 | 82.45711829 | 67.57037997 | 68.44020841 |
| AP53_511 | 59.77666085 | 51.25492241 | 62.31142238 | 84.59565508 | 70.97987306 | 84.21769932 |
| AP53_512 | 49.5967177 | 47.23818785 | 50.28368829 | 56.23207866 | 53.58550628 | 54.82898627 |
| AP53_513 | 58.42941749 | 49.8641128 | 50.84669069 | 55.19034686 | 57.95647644 | 52.74244992 |
| AP53_514 | 90.32169292 | 86.32479254 | 89.57597847 | 95.6330006 | 95.71781263 | 112.2200747 |
| AP53_515 | 112.8807753 | 121.9420553 | 115.3739347 | 132.6839617 | 142.319992 | 154.8816122 |
| AP53_516 | 102.9031098 | 115.5021253 | 105.0993116 | 139.8123515 | 142.4096293 | 149.5694312 |
| AP53_517 | 58.83380796 | 52.41421126 | 48.79799702 | 46.57038757 | 48.48765985 | 46.6817397 |
| AP53_518 | 145.3404609 | 138.7915437 | 157.9943301 | 165.21678 | 163.2129265 | 178.1346727 |
| AP53_519 | 484.0878717 | 504.2008885 | 465.0172851 | 425.2432176 | 451.0786912 | 451.8838556 |
| AP53_52 | 2572.891701 | 3096.154543 | 3113.010078 | 1510.252769 | 1542.186248 | 1652.844661 |
| AP53_520 | 520.9855311 | 468.854781 | 430.4171923 | 493.0253589 | 470.1099021 | 501.8856182 |
| AP53_521 | 514.8886593 | 498.8898136 | 501.3176253 | 498.4357595 | 514.9233064 | 513.9127723 |
| AP53_522 | 272.381002 | 282.7410014 | 232.5723072 | 293.5979534 | 229.4243943 | 252.4794211 |
| AP53_523 | 631.8119977 | 580.4700754 | 512.6493635 | 785.5731648 | 544.5259275 | 674.3234652 |
| AP53_524 | 48.21079147 | 52.85810609 | 47.23559484 | 57.18660999 | 58.31987976 | 50.54452782 |
| AP53_525 | 116.4127604 | 118.68631 | 123.0569997 | 111.4537615 | 98.90595639 | 111.2492586 |
| AP53_526 | 502.1943543 | 540.644461 | 542.9737832 | 473.1645535 | 470.2754778 | 525.0129364 |
| AP53_527 | 443.8959006 | 491.368304 | 498.9287407 | 430.692301 | 398.9054938 | 411.6076295 |
| AP53_528 | 472.0355056 | 503.9448416 | 487.4036205 | 456.418112 | 408.4391495 | 481.7612041 |

| | | | | | | |
|---|---|---|---|---|---|---|
| AP53_529 | 516.1382923 | 537.4272339 | 593.6671824 | 530.8306803 | 487.8289913 | 588.4846492 |
| AP53_53 | 2906.575232 | 3622.373798 | 4018.879499 | 1620.232307 | 1416.982873 | 1748.862174 |
| AP53_530 | 520.2794664 | 513.8360782 | 576.9024033 | 597.5329603 | 552.4961491 | 603.4746134 |
| AP53_531 | 137.8242351 | 145.2011633 | 157.638417 | 127.3565395 | 122.0359482 | 138.8405781 |
| AP53_532 | 451.939783 | 381.6846963 | 358.4114372 | 488.2430109 | 446.4012645 | 435.8015929 |
| AP53_533 | 151.3771564 | 114.4091812 | 152.2489073 | 156.2114974 | 164.0733884 | 153.5380293 |
| AP53_534 | 242.4451569 | 199.8398636 | 247.9994611 | 262.2507626 | 260.3726724 | 282.6032749 |
| AP53_535 | 2337.224136 | 2522.08942 | 1771.198383 | 1587.691651 | 1308.10789 | 1302.904811 |
| AP53_536 | 392.3651668 | 452.0667776 | 424.8866878 | 389.8737761 | 393.1901398 | 436.8353976 |
| AP53_537 | 455.4192976 | 410.6189402 | 268.983934 | 378.2753188 | 413.6320636 | 353.1739517 |
| AP53_538 | 306.5858185 | 283.9848705 | 307.8402902 | 382.0290871 | 378.3568657 | 324.7133297 |
| AP53_539 | 374.9230845 | 394.0945697 | 428.631542 | 471.8014193 | 571.1581433 | 591.1410921 |
| AP53_54 | 2898.231563 | 3237.728668 | 3530.610361 | 1601.151321 | 1648.152603 | 1895.749041 |
| AP53_540 | 7318.331353 | 6368.550576 | 5999.997463 | 9270.047073 | 8447.449039 | 9192.23768 |
| AP53_541 | 391.1695739 | 391.051132 | 353.8732261 | 376.9508087 | 437.6649297 | 377.7395563 |
| AP53_542 | 811.04712 | 785.6790113 | 781.951356 | 881.6910675 | 929.8681864 | 923.5330793 |
| AP53_543 | 886.6931839 | 858.7581531 | 778.0143446 | 916.6218993 | 983.532125 | 863.9795751 |
| AP53_544 | 278.6232973 | 273.776264 | 280.39523 | 278.5861682 | 345.7360295 | 282.3150334 |
| AP53_545 | 3920.403479 | 5697.077592 | 5235.441394 | 2754.888406 | 2619.518021 | 2800.954528 |
| AP53_546 | 132.2622253 | 101.880235 | 104.1204152 | 230.1512102 | 156.0553153 | 181.0434295 |
| AP53_547 | 118.6432775 | 93.29618005 | 98.03792165 | 190.9509421 | 134.1308777 | 163.254683 |
| AP53_548 | 149.12383 | 127.3306944 | 121.7467406 | 224.1691861 | 160.8655717 | 200.9739748 |
| AP53_549 | 79.33101522 | 75.02732231 | 83.68014598 | 111.2025659 | 81.24083404 | 101.8380376 |
| AP53_55 | 2314.062464 | 2471.619365 | 2415.759494 | 1436.507494 | 1390.56447 | 1660.170991 |
| AP53_550 | 107.1939547 | 93.73381756 | 103.1567088 | 133.2676872 | 109.1185012 | 128.2270571 |
| AP53_551 | 106.4496934 | 88.6276663 | 92.10614028 | 134.7612418 | 100.6669938 | 121.9822329 |
| AP53_552 | 64.02083499 | 50.55744546 | 56.24499763 | 102.8711546 | 67.29597152 | 73.71107046 |
| AP53_553 | 103.9485408 | 72.36743088 | 79.57880898 | 164.5561898 | 102.0970725 | 126.4099751 |
| AP53_554 | 6.781109256 | 3.969595617 | 8.277150336 | 7.652200395 | 7.439202607 | 6.022405815 |
| AP53_555 | 441.4915332 | 384.4283196 | 344.4487049 | 420.1239503 | 364.3573531 | 393.1465299 |
| AP53_556 | 558.4410144 | 545.0302323 | 562.6231859 | 583.5458883 | 560.0324712 | 608.6671857 |
| AP53_557 | 141.5478975 | 155.4962944 | 165.7936662 | 113.2826789 | 107.0998104 | 117.9879681 |
| AP53_558 | 189.5616708 | 219.7147892 | 231.7256202 | 152.6958406 | 152.9351981 | 173.6195428 |
| AP53_559 | 733.9845898 | 1167.903147 | 1073.885009 | 414.2131155 | 360.7555359 | 466.5577555 |
| AP53_56 | 1761.910742 | 2096.000908 | 2132.35413 | 1028.487898 | 1037.401303 | 1194.225265 |

| | | | | | | |
|---|---|---|---|---|---|---|
| AP53_560 | 723.7219188 | 1028.235716 | 1011.772536 | 448.8291562 | 379.3905787 | 464.775179 |
| AP53_561 | 928.1556081 | 1105.64425 | 1015.109717 | 569.895692 | 425.1926972 | 505.8030064 |
| AP53_562 | 491.3551065 | 682.5220692 | 719.9305831 | 322.1917069 | 276.5557336 | 300.151603 |
| AP53_563 | 1433.178302 | 2147.630542 | 2181.181328 | 811.7505258 | 695.3930517 | 926.3230405 |
| AP53_564 | 992.8411302 | 1382.411674 | 1363.350125 | 594.5527822 | 535.8550628 | 646.6558244 |
| AP53_565 | 1839.680274 | 2851.680893 | 2675.465748 | 975.051117 | 866.8073818 | 1122.328507 |
| AP53_566 | 1726.982999 | 2291.663257 | 2148.501104 | 1052.55541 | 944.497754 | 1200.827281 |
| AP53_567 | 186.6497694 | 222.1650347 | 208.4600312 | 131.1324345 | 154.998886 | 149.9645964 |
| AP53_568 | 895.854382 | 1038.835683 | 983.943318 | 870.17419 | 859.3408489 | 942.563917 |
| AP53_569 | 310.6443923 | 296.9099998 | 238.0961709 | 279.1709684 | 297.1216045 | 282.0413729 |
| AP53_57 | 1598.799168 | 1947.629858 | 2060.976205 | 945.781632 | 980.8543191 | 1064.794177 |
| AP53_570 | 562.3811238 | 614.5262081 | 578.1401393 | 720.3321132 | 766.2052067 | 788.4374423 |
| AP53_571 | 153.111139 | 116.2744176 | 105.3629506 | 186.6267328 | 193.4657628 | 144.2554393 |
| AP53_572 | 412.0186214 | 496.6837428 | 441.3914574 | 500.0411508 | 581.9246497 | 599.0035384 |
| AP53_573 | 109.2658569 | 156.3815349 | 153.1415522 | 72.72055818 | 76.12423264 | 78.43116385 |
| AP53_574 | 135.2458283 | 200.2990139 | 191.5724256 | 94.64256252 | 96.17424551 | 109.4520944 |
| AP53_575 | 50.52539674 | 46.80184737 | 44.47818702 | 52.26059034 | 51.58548256 | 51.75604574 |
| AP53_576 | 60.59477961 | 72.22048398 | 81.03266015 | 62.19906931 | 60.25249514 | 70.00657716 |
| AP53_577 | 228.350351 | 293.0899507 | 273.208789 | 224.3967234 | 179.4448604 | 229.7677829 |
| AP53_578 | 238.9667428 | 307.2582713 | 260.9984078 | 252.6234777 | 195.4291869 | 258.672007 |
| AP53_579 | 188.8508562 | 198.4386718 | 177.2609336 | 211.2388531 | 158.0085731 | 190.9474577 |
| AP53_58 | 1803.800611 | 1895.957593 | 2032.700499 | 1100.482798 | 1021.464338 | 1123.445039 |
| AP53_580 | 198.8739467 | 189.8710907 | 217.4699642 | 195.251595 | 188.0072826 | 185.0440834 |
| AP53_581 | 215.5485999 | 203.8414939 | 230.7864768 | 194.2313475 | 186.094687 | 194.6546939 |
| AP53_582 | 284.9992042 | 291.5982912 | 317.8349372 | 274.8391778 | 272.8459664 | 275.3130225 |
| AP53_583 | 2782.029591 | 3895.582843 | 1873.887213 | 1711.121618 | 1698.793927 | 1747.054844 |
| AP53_584 | 1219.142568 | 1206.540544 | 664.0309654 | 1265.372687 | 1086.263834 | 1116.498874 |
| AP53_585 | 530.5037887 | 475.0260554 | 473.8557465 | 629.7860527 | 621.2806838 | 586.9492115 |
| AP53_586 | 919.195769 | 945.3500769 | 998.6974704 | 1202.865737 | 1209.183951 | 1195.828702 |
| AP53_587 | 549.9327489 | 504.6796325 | 597.4143129 | 817.0186027 | 783.9553499 | 695.2792011 |
| AP53_588 | 398.561564 | 416.0637629 | 449.8348042 | 444.9284658 | 416.5855576 | 450.1765956 |
| AP53_589 | 118.1412187 | 115.3853184 | 112.2537454 | 116.7784203 | 111.0367411 | 114.714447 |
| AP53_59 | 2340.732368 | 2814.401595 | 2825.926936 | 1465.730563 | 1358.652751 | 1728.679294 |
| AP53_590 | 192.4855361 | 190.6867371 | 214.5412083 | 189.6870351 | 186.8411947 | 198.5546209 |
| AP53_591 | 100.9887464 | 103.693836 | 113.5189549 | 97.39683284 | 104.5469934 | 97.33226512 |

| | | | | | | |
|---|---|---|---|---|---|---|
| AP53_592 | 148.8270283 | 141.6103863 | 169.5081118 | 140.9369648 | 134.6678413 | 138.6164562 |
| AP53_593 | 117.8278748 | 108.4951916 | 105.0212718 | 98.39164485 | 100.6624022 | 85.21211323 |
| AP53_594 | 104.3211311 | 111.7297933 | 117.7058421 | 87.08113208 | 92.25985312 | 81.99103678 |
| AP53_596 | 237.8074189 | 217.128374 | 143.2892968 | 285.5927068 | 390.5681001 | 279.7443349 |
| AP53_597 | 231.6519611 | 253.7057672 | 235.5271574 | 282.9326562 | 321.4039167 | 268.7908282 |
| AP53_598 | 424.7148405 | 500.2163049 | 460.1873878 | 594.8840463 | 913.7903562 | 568.7349745 |
| AP53_599 | 922.8399759 | 619.4057761 | 325.9076213 | 918.1104237 | 1034.787271 | 742.3430701 |
| AP53_6 | 66.04319015 | 55.97338746 | 58.27658386 | 74.39232457 | 85.74659847 | 60.73913233 |
| AP53_60 | 1642.325405 | 1781.447454 | 1467.917497 | 952.5619263 | 871.5730324 | 1117.851289 |
| AP53_600 | 1422.181623 | 1294.028127 | 1146.771357 | 2147.955308 | 2591.237587 | 2185.401597 |
| AP53_601 | 280.1471931 | 295.9643811 | 354.2553593 | 329.4370375 | 461.32459 | 389.9084354 |
| AP53_604 | 64.10775338 | 78.55660407 | 97.43598547 | 60.32962701 | 99.33076289 | 72.14399762 |
| AP53_605 | 78.25018513 | 88.96058094 | 111.5505184 | 72.32699602 | 112.5049338 | 79.0370572 |
| AP53_606 | 52.64753391 | 59.98684111 | 83.68850734 | 40.46628177 | 59.73145184 | 50.48309336 |
| AP53_607 | 42.72502469 | 49.11606649 | 61.32235218 | 25.13949349 | 29.54716772 | 26.87752516 |
| AP53_608 | 32.28104962 | 37.32406476 | 50.61767857 | 24.92118125 | 39.3005243 | 30.90518462 |
| AP53_609 | 22.44190651 | 25.20075381 | 30.38793102 | 13.63428843 | 22.31941696 | 20.10007233 |
| AP53_61 | 1446.752116 | 1751.03418 | 1720.080039 | 845.2697344 | 899.4460902 | 1033.624194 |
| AP53_610 | 17.68503511 | 25.43793534 | 35.01849172 | 10.26552751 | 17.54174995 | 12.3416365 |
| AP53_611 | 28.04233315 | 44.58161232 | 54.61327462 | 20.39992195 | 20.45184627 | 8.92422742 |
| AP53_612 | 29.97709339 | 54.6566153 | 64.31181907 | 9.720797021 | 13.44212352 | 11.60754124 |
| AP53_613 | 37.66528116 | 55.1325939 | 89.93760544 | 9.548899638 | 15.40415655 | 9.112105892 |
| AP53_614 | 40.60966803 | 65.71399863 | 121.4411114 | 13.46210296 | 16.43617131 | 14.85645319 |
| AP53_615 | 49.59731056 | 66.01765824 | 99.49383641 | 13.49116579 | 23.09543798 | 17.21630735 |
| AP53_616 | 41.00596589 | 47.89760001 | 69.28693097 | 11.47159318 | 16.83042739 | 21.95150329 |
| AP53_617 | 19.22411556 | 38.26227698 | 56.61008305 | 6.409477648 | 10.05059745 | 8.358767941 |
| AP53_618 | 29.77271305 | 41.96820746 | 55.99563557 | 6.420811826 | 12.87430968 | 5.68494342 |
| AP53_619 | 31.30808189 | 49.63832297 | 79.52524004 | 8.284669419 | 11.5382672 | 11.33980777 |
| AP53_62 | 1687.973756 | 1846.31563 | 2025.615762 | 1059.644405 | 1125.375496 | 1289.871981 |
| AP53_620 | 27.13264447 | 39.31965622 | 53.96532784 | 7.299631964 | 12.11290239 | 8.512151961 |
| AP53_621 | 11.42490272 | 14.19631645 | 16.13574023 | 6.112742571 | 7.84251165 | 5.643470769 |
| AP53_622 | 13.40752471 | 14.96613225 | 22.90740805 | 7.921058231 | 9.735347302 | 8.000301659 |
| AP53_623 | 15.53049573 | 15.85591306 | 22.94371413 | 6.387771628 | 7.844937419 | 9.331888256 |
| AP53_624 | 16.34720431 | 13.61968256 | 34.44122255 | 10.49510878 | 13.57654476 | 13.92179478 |
| AP53_625 | 9.638050365 | 10.78252949 | 17.0964861 | 7.348413252 | 9.077340971 | 8.804072583 |

| | | | | | | |
|---|---|---|---|---|---|---|
| AP53_626 | 9.328783423 | 10.49205186 | 11.19912711 | 8.171942323 | 7.022350737 | 8.21158494 |
| AP53_627 | 9.867527755 | 10.20145568 | 16.80134827 | 4.145533424 | 6.92680855 | 7.850636152 |
| AP53_628 | 6.775359769 | 8.376467002 | 9.677152717 | 5.02537971 | 7.601450841 | 4.674944792 |
| AP53_629 | 16.15132652 | 14.70527809 | 14.09123075 | 13.51681078 | 16.59074033 | 11.75642528 |
| AP53_63 | 1673.993948 | 1678.495142 | 1888.190194 | 1075.388696 | 1103.289327 | 1147.642689 |
| AP53_630 | 24.5675223 | 27.14067177 | 38.14761651 | 19.50353315 | 26.80875686 | 20.97295552 |
| AP53_631 | 23.83924847 | 19.61788206 | 32.48415261 | 23.21926722 | 21.47617146 | 22.3708393 |
| AP53_632 | 23.39163938 | 20.03801192 | 24.42639897 | 13.1458981 | 16.24852431 | 17.53865523 |
| AP53_633 | 20.03635407 | 16.5904855 | 25.36851802 | 17.83212278 | 20.20936052 | 21.25464851 |
| AP53_634 | 21.46854012 | 15.27221449 | 28.57853934 | 17.61384866 | 14.21867863 | 12.12958649 |
| AP53_635 | 18.19198533 | 12.78969403 | 22.4722077 | 13.79285503 | 14.98029244 | 13.79515077 |
| AP53_636 | 19.69608423 | 15.86239121 | 21.40584839 | 15.80443686 | 17.05111071 | 16.66864927 |
| AP53_637 | 30.18732565 | 26.88518898 | 31.21865202 | 23.13437958 | 26.77596327 | 31.75146177 |
| AP53_638 | 17.1671901 | 11.94708997 | 10.07053291 | 14.25481668 | 11.3137873 | 12.85484282 |
| AP53_639 | 28.78028929 | 23.88730984 | 30.61986357 | 28.7082858 | 30.73069252 | 29.95265234 |
| AP53_64 | 229.6477919 | 307.8747542 | 311.831924 | 93.45939671 | 103.8956242 | 118.7669792 |
| AP53_640 | 19.11011209 | 18.25617024 | 23.26293102 | 20.99021757 | 22.74071247 | 16.85269894 |
| AP53_641 | 27.87143804 | 23.25869643 | 29.81407769 | 20.49129898 | 22.32984335 | 23.86180199 |
| AP53_642 | 37.63739676 | 31.17011437 | 41.13682631 | 33.10562823 | 36.25839213 | 45.62639092 |
| AP53_643 | 20.98468337 | 15.97215244 | 24.79967848 | 15.39478111 | 20.84587581 | 19.61628771 |
| AP53_644 | 27.55696603 | 22.85181638 | 32.37439774 | 23.81981462 | 26.46211324 | 24.26028055 |
| AP53_645 | 24.94291738 | 15.56228504 | 24.05738417 | 17.55454277 | 22.94184646 | 14.92590083 |
| AP53_646 | 13.13100699 | 15.168197 | 16.99402428 | 11.1087341 | 9.811174922 | 11.16262328 |
| AP53_647 | 13.52004287 | 7.412989025 | 11.7684716 | 6.29863993 | 13.22134446 | 10.86308956 |
| AP53_648 | 182.0832969 | 157.2059104 | 177.1154975 | 186.2740746 | 195.8699426 | 212.6432031 |
| AP53_649 | 79.04986124 | 71.15720677 | 70.15804593 | 119.9978388 | 116.9091354 | 73.6796741 |
| AP53_65 | 443.6107055 | 390.2112492 | 212.9296924 | 190.6093553 | 172.9614606 | 181.9268423 |
| AP53_650 | 20.75423678 | 14.08088248 | 21.06301601 | 15.02026019 | 15.53653889 | 20.64017016 |
| AP53_651 | 14.76553972 | 13.38659826 | 19.70321656 | 14.35165647 | 10.88337148 | 9.955516859 |
| AP53_652 | 212.8265495 | 229.22379 | 276.8105456 | 303.8286248 | 350.147212 | 246.9036253 |
| AP53_653 | 215.7423212 | 190.4587735 | 201.6797182 | 255.206105 | 309.4882197 | 239.3944622 |
| AP53_654 | 334.2242178 | 304.5628041 | 322.5424528 | 434.9089744 | 469.4076608 | 432.6643443 |
| AP53_655 | 712.9433065 | 670.7800787 | 598.9695532 | 667.3460311 | 656.0720642 | 594.3069297 |
| AP53_657 | 203.0995242 | 207.7176458 | 242.7996803 | 215.8997976 | 279.1384418 | 238.4517935 |
| AP53_658 | 112.091653 | 100.2792977 | 116.4736635 | 132.7479804 | 169.4090782 | 119.2286672 |

| | | | | | | |
|---|---|---|---|---|---|---|
| AP53_659 | 153.62905 | 148.1663131 | 168.3469155 | 180.4820948 | 224.179135 | 182.5837151 |
| AP53_66 | 1978.903173 | 1754.657958 | 1319.627139 | 666.2527702 | 468.2167359 | 538.2718223 |
| AP53_660 | 136.5304625 | 145.5290347 | 144.9732929 | 170.9042538 | 214.4631544 | 155.6492191 |
| AP53_661 | 281.059276 | 244.4312554 | 219.9226672 | 306.3147855 | 266.0403807 | 299.645441 |
| AP53_662 | 245.8507656 | 232.8021709 | 236.5030673 | 337.8249443 | 303.3899649 | 313.9933371 |
| AP53_663 | 214.3695487 | 191.8698027 | 132.9768095 | 247.2454758 | 219.3331947 | 194.2834199 |
| AP53_664 | 290.4818129 | 202.6640842 | 162.2685869 | 351.3274395 | 246.0215068 | 259.2191182 |
| AP53_665 | 286.1293254 | 249.504823 | 251.0756872 | 421.922165 | 376.7939734 | 409.9392387 |
| AP53_666 | 1291.893528 | 1438.915615 | 1372.835529 | 951.8663908 | 1207.986493 | 1010.499585 |
| AP53_667 | 2049.331023 | 1830.007638 | 1825.055214 | 1392.166434 | 1584.187353 | 1416.881227 |
| AP53_668 | 1987.854581 | 2177.700308 | 2189.333854 | 1444.205964 | 1686.319997 | 1459.651334 |
| AP53_669 | 310.4159773 | 303.7049966 | 279.751557 | 290.2148195 | 327.6498515 | 269.3303471 |
| AP53_67 | 1594.144942 | 1728.231781 | 1402.239162 | 715.6765086 | 513.2935467 | 616.6610348 |
| AP53_670 | 282.195187 | 269.1672294 | 255.2802228 | 264.8645054 | 303.8976578 | 264.3384731 |
| AP53_671 | 250.8032546 | 238.3412813 | 241.0331479 | 253.9139222 | 259.4513275 | 246.4551236 |
| AP53_672 | 355.8397927 | 325.5374545 | 303.1644618 | 356.7414454 | 352.9814384 | 384.9166232 |
| AP53_673 | 927.8361332 | 913.316899 | 834.3966543 | 986.9975677 | 975.5164232 | 1043.491866 |
| AP53_674 | 265.0594447 | 310.2450217 | 289.7566969 | 288.9326494 | 280.8252592 | 315.3077747 |
| AP53_675 | 324.5551871 | 310.9358156 | 334.7743 | 333.9425042 | 304.5161623 | 354.5379229 |
| AP53_676 | 319.3633189 | 304.3680689 | 324.1067428 | 282.0114265 | 259.8707674 | 275.6445582 |
| AP53_677 | 304.6027258 | 291.8340425 | 265.3228592 | 272.0855397 | 274.5152406 | 305.8794666 |
| AP53_678 | 356.7618872 | 372.7793336 | 343.8527514 | 310.6318139 | 316.7860443 | 339.8130089 |
| AP53_679 | 162.5051323 | 138.5365845 | 137.2139316 | 159.7320838 | 138.4823315 | 151.9854015 |
| AP53_68 | 1891.854328 | 2665.980417 | 2639.738439 | 960.2442651 | 807.300596 | 1049.143478 |
| AP53_680 | 1814.199797 | 1938.907393 | 1445.553067 | 1228.137307 | 1125.721836 | 1252.089237 |
| AP53_681 | 786.309188 | 854.187782 | 680.7310686 | 591.8446223 | 553.8046984 | 581.9794219 |
| AP53_682 | 1799.296698 | 1980.560873 | 1299.561168 | 1358.69858 | 1208.381751 | 1468.25588 |
| AP53_683 | 184.9487443 | 154.8187843 | 126.492497 | 190.0330644 | 186.0097915 | 167.1456206 |
| AP53_684 | 168.5433935 | 143.4697475 | 150.5643401 | 186.5354566 | 193.1108204 | 174.0822979 |
| AP53_685 | 454.3990994 | 430.0849684 | 462.6341785 | 531.3668904 | 527.6338985 | 543.1917712 |
| AP53_686 | 293.4266275 | 323.8380135 | 356.2885091 | 293.6062135 | 318.9317626 | 372.4690707 |
| AP53_687 | 380.8043419 | 364.8474235 | 490.6147836 | 392.1559465 | 471.017209 | 519.0578936 |
| AP53_688 | 491.0319751 | 474.8788627 | 627.6482593 | 472.89217 | 577.7768914 | 629.9996316 |
| AP53_689 | 648.1225478 | 589.0731449 | 774.7894211 | 663.4585568 | 773.3647431 | 871.0509153 |
| AP53_69 | 1479.987298 | 1976.247325 | 1935.521033 | 740.1631393 | 642.5544406 | 740.1586452 |

| | | | | | | |
|---|---|---|---|---|---|---|
| AP53_690 | 501.706482 | 447.9147005 | 542.7246971 | 503.9595933 | 580.528695 | 618.6138196 |
| AP53_691 | 442.9992453 | 296.717442 | 330.6887493 | 523.4778849 | 452.5648336 | 477.6976317 |
| AP53_692 | 413.9504224 | 264.9596204 | 266.8691221 | 489.8288828 | 433.2178048 | 382.1305428 |
| AP53_693 | 515.7823834 | 434.7419192 | 519.5355761 | 744.1071224 | 703.5952587 | 761.8460499 |
| AP53_694 | 268.2442464 | 247.7923915 | 277.8630776 | 391.4433084 | 406.4155115 | 380.1491721 |
| AP53_695 | 528.418762 | 554.5634132 | 384.1188981 | 392.4969786 | 360.6735929 | 348.4474661 |
| AP53_696 | 492.2868482 | 621.7602425 | 554.9493041 | 436.0971614 | 414.5088821 | 400.7095536 |
| AP53_697 | 168.3788838 | 173.5110636 | 181.5228454 | 145.8043522 | 146.2424163 | 154.0964775 |
| AP53_698 | 273.4557039 | 333.4638062 | 332.6657755 | 223.5958419 | 238.4760614 | 234.0348847 |
| AP53_699 | 29.9630409 | 26.89401031 | 40.24764351 | 39.884196 | 33.76700558 | 41.59224016 |
| AP53_7 | 74.84846467 | 70.77044686 | 84.93132228 | 92.53655212 | 98.49678608 | 86.58776694 |
| AP53_70 | 1571.894622 | 2067.662348 | 1898.295453 | 785.492888 | 667.5134506 | 775.0423897 |
| AP53_700 | 24.81059421 | 27.89553472 | 26.56537129 | 27.4343778 | 26.13874996 | 21.26589946 |
| AP53_701 | 143.4689103 | 156.7829649 | 116.7425072 | 94.81410043 | 89.95441866 | 106.8382334 |
| AP53_702 | 127.3881657 | 141.505509 | 102.3975088 | 86.44173417 | 79.49790296 | 94.59373068 |
| AP53_703 | 97.62691483 | 114.9491753 | 97.48032459 | 61.42006456 | 57.26963933 | 60.26727028 |
| AP53_704 | 288.6181985 | 345.848488 | 311.1452328 | 195.8422017 | 195.7637757 | 227.394158 |
| AP53_705 | 424.7522174 | 457.4862287 | 434.0955525 | 322.4486973 | 278.1428488 | 332.5814952 |
| AP53_706 | 270.8733109 | 239.8603133 | 238.5193026 | 288.4803553 | 196.0184955 | 273.540793 |
| AP53_707 | 341.9174105 | 300.3612805 | 242.4381418 | 323.2391049 | 239.4317453 | 308.9380272 |
| AP53_708 | 737.697794 | 838.7113309 | 658.1147566 | 752.2737898 | 542.6695558 | 781.0193481 |
| AP53_709 | 40.86464467 | 34.31610948 | 28.70985259 | 38.40881162 | 34.04060564 | 38.1360337 |
| AP53_71 | 50.59976442 | 54.2496015 | 35.48106362 | 31.09953466 | 43.01870286 | 37.91431258 |
| AP53_710 | 160.2915773 | 166.8693527 | 165.0756634 | 161.6728759 | 141.0867628 | 146.9333046 |
| AP53_711 | 4.604846286 | 9.780091202 | 9.818769586 | 6.347848055 | 6.364005355 | 4.12158398 |
| AP53_712 | 1.439014464 | 2.1733536 | 3.398804857 | 3.385518963 | 0.424267024 | 0.915907551 |
| AP53_713 | 3.500760334 | 4.617846831 | 4.240224383 | 3.415216497 | 3.126178069 | 3.16014886 |
| AP53_714 | 0 | 0.7244512 | 0.755289968 | 0.846379741 | 1.272801071 | 0 |
| AP53_715 | 9.951150871 | 7.612877019 | 9.217097917 | 7.172709666 | 6.328050522 | 12.72956257 |
| AP53_716 | 487.8598959 | 489.5464724 | 409.6407323 | 482.1032318 | 450.8588781 | 422.1828979 |
| AP53_717 | 340.1972569 | 340.3128712 | 254.9837582 | 333.5274512 | 287.0023993 | 291.3427482 |
| AP53_718 | 239.811939 | 226.8269291 | 194.7636067 | 244.295016 | 240.5372942 | 241.3632306 |
| AP53_719 | 278.258503 | 289.0166817 | 248.4011161 | 311.6462144 | 306.2584387 | 320.3042829 |
| AP53_720 | 84.21191495 | 81.37671015 | 76.35671185 | 113.8554665 | 133.2082217 | 96.35849304 |
| AP53_721 | 156.2118475 | 166.0420999 | 145.3244056 | 174.4654298 | 219.6278636 | 161.5206309 |

| | | | | | | |
|---|---|---|---|---|---|---|
| AP53_722 | 196.5466432 | 178.76994 | 149.2119794 | 190.3660663 | 212.0449911 | 162.6520772 |
| AP53_723 | 146.3514608 | 135.9739176 | 122.0858451 | 131.4275823 | 121.8407863 | 113.0089009 |
| AP53_724 | 173.9723316 | 160.5487817 | 159.593109 | 149.8433729 | 137.9723971 | 132.9092751 |
| AP53_725 | 225.4526019 | 239.7210784 | 210.1948092 | 211.3684102 | 200.0971286 | 207.9176995 |
| AP53_726 | 148.8079016 | 173.3871027 | 161.3143297 | 118.9304078 | 114.0354613 | 127.0103534 |
| AP53_727 | 189.5661721 | 218.5312795 | 209.3711747 | 159.1731296 | 151.9280009 | 167.1313207 |
| AP53_728 | 240.7326446 | 244.1700613 | 232.1734547 | 194.3668505 | 168.2507451 | 188.5468858 |
| AP53_729 | 258.2900144 | 278.5844161 | 274.0558206 | 217.8017199 | 193.7228943 | 226.6454867 |
| AP53_730 | 176.8433926 | 217.0972775 | 208.7153405 | 151.9509919 | 130.3077403 | 148.824227 |
| AP53_731 | 124.8712143 | 159.7119202 | 151.0965288 | 95.3040861 | 90.87106966 | 90.32904572 |
| AP53_732 | 84.32624761 | 92.83324665 | 87.28994061 | 68.73812608 | 63.0339578 | 62.87051118 |
| AP53_733 | 185.7195535 | 233.0463499 | 192.2804461 | 133.4832627 | 129.1203014 | 133.6342222 |
| AP53_734 | 233.4737854 | 359.683666 | 334.1826842 | 209.0706447 | 207.8164088 | 223.9956362 |
| AP53_735 | 363.7547782 | 424.846455 | 438.8050498 | 264.8136418 | 247.275239 | 274.5041948 |
| AP53_736 | 409.8313194 | 472.4870727 | 499.473256 | 317.8155926 | 290.9623248 | 349.0523677 |
| AP53_737 | 214.6916741 | 211.6098586 | 192.7207628 | 185.5209786 | 195.4365515 | 166.2667659 |
| AP53_738 | 12419.73092 | 9243.852424 | 10732.17411 | 19668.39005 | 18802.42352 | 19146.15762 |
| AP53_739 | 1036.696315 | 1026.56006 | 919.6649165 | 977.6428443 | 738.0013227 | 959.4533311 |
| AP53_74 | 130.6780703 | 134.2192696 | 122.0711895 | 111.1731227 | 111.8000941 | 88.86778671 |
| AP53_740 | 1372.591836 | 1333.133664 | 1316.971448 | 1208.06043 | 894.3212806 | 1147.559614 |
| AP53_741 | 851.6616217 | 854.5567218 | 897.9215975 | 909.7142791 | 709.7669827 | 875.0343977 |
| AP53_742 | 1036.815251 | 1130.090209 | 1138.58564 | 957.1614445 | 745.8300004 | 1017.132297 |
| AP53_743 | 843.9638444 | 887.9701853 | 747.2927803 | 767.0760927 | 558.3211421 | 738.3446334 |
| AP53_744 | 327.8545502 | 304.2144127 | 316.0730566 | 388.562319 | 335.2193442 | 368.243591 |
| AP53_745 | 33.78555699 | 30.23796314 | 40.06320701 | 36.06313678 | 50.54311499 | 42.60961216 |
| AP53_746 | 278.3873545 | 261.4261318 | 257.4488388 | 299.3157493 | 303.1120961 | 295.5773176 |
| AP53_747 | 505.568389 | 580.2085143 | 480.7694915 | 634.453819 | 721.3013444 | 654.6436429 |
| AP53_748 | 354.1346072 | 345.8548846 | 392.293032 | 397.3221605 | 515.9527805 | 497.7620518 |
| AP53_749 | 411.1469898 | 354.567116 | 488.205049 | 462.5263764 | 579.1850969 | 546.1425883 |
| AP53_75 | 194.4322863 | 239.0997238 | 216.4307509 | 173.0216287 | 174.1119649 | 164.5515609 |
| AP53_750 | 498.2491154 | 456.7478103 | 528.5161019 | 523.5687624 | 631.58926 | 634.2990273 |
| AP53_751 | 850.8572746 | 780.4452408 | 668.5889739 | 698.9098261 | 729.9160586 | 721.4680105 |
| AP53_752 | 665.2093039 | 668.3646556 | 637.1584404 | 759.4795534 | 830.9377192 | 824.0635496 |
| AP53_753 | 489.0536864 | 396.5207211 | 409.7274846 | 566.4532301 | 528.1151355 | 533.4951414 |
| AP53_754 | 604.3724673 | 539.7932134 | 585.162242 | 751.8973497 | 733.7111422 | 798.0737711 |

| | | | | | | |
|---|---|---|---|---|---|---|
| AP53_755 | 273.3858507 | 227.6943351 | 240.7751478 | 320.0422832 | 354.163837 | 294.9735907 |
| AP53_756 | 462.3796999 | 465.7441039 | 488.777188 | 443.7113237 | 464.1611783 | 527.0836593 |
| AP53_757 | 501.1982085 | 529.1320888 | 565.3621737 | 499.5498377 | 521.041297 | 559.0833727 |
| AP53_758 | 212.6711903 | 223.1394428 | 188.645816 | 133.0449557 | 148.1212942 | 137.1718853 |
| AP53_759 | 415.1837057 | 454.9929875 | 341.7441882 | 295.4634731 | 338.0912282 | 320.92449 |
| AP53_76 | 152.8741248 | 182.6043172 | 176.6045661 | 153.7797308 | 156.1053078 | 143.5819779 |
| AP53_760 | 1826.732212 | 2747.472166 | 2129.880134 | 1004.286409 | 899.1927964 | 988.1776692 |
| AP53_761 | 349.4667998 | 392.9251163 | 277.737321 | 390.6754803 | 316.2259559 | 320.4769589 |
| AP53_762 | 471.1219634 | 657.2908305 | 513.1380355 | 341.278834 | 359.4792569 | 350.5503916 |
| AP53_763 | 181.6668016 | 182.7030588 | 199.6912989 | 168.4502118 | 197.8533047 | 179.6072368 |
| AP53_764 | 125.4731372 | 132.9845304 | 157.1471531 | 144.8687184 | 182.3361534 | 156.2012878 |
| AP53_765 | 300.4928158 | 325.6444009 | 322.4066155 | 353.8034916 | 401.9755022 | 385.2857309 |
| AP53_766 | 1162.528567 | 1361.231526 | 782.6852314 | 512.4183786 | 428.2939649 | 391.5116684 |
| AP53_767 | 90.44685579 | 95.57926167 | 91.94396546 | 82.04240953 | 95.94091628 | 90.43060554 |
| AP53_768 | 404.5090514 | 360.4757179 | 339.8804857 | 344.8077465 | 340.5204025 | 353.8411925 |
| AP53_769 | 414.8705852 | 375.7578017 | 369.1657854 | 338.392202 | 358.1454083 | 365.8445822 |
| AP53_77 | 50.27556784 | 61.57835201 | 47.55966518 | 83.05101205 | 90.15674253 | 72.12771964 |
| AP53_770 | 365.5282418 | 309.3172931 | 281.1140533 | 286.2947729 | 264.4689021 | 288.540424 |
| AP53_771 | 540.6631286 | 561.094557 | 567.6226255 | 539.6251695 | 503.5966381 | 605.7201937 |
| AP53_772 | 307.0782671 | 316.930198 | 331.1797912 | 266.3856779 | 285.4390244 | 309.2461805 |
| AP53_773 | 631.9685719 | 809.5428093 | 718.0057119 | 499.1194286 | 524.0310845 | 625.2524958 |
| AP53_774 | 479.7724716 | 481.7600481 | 488.0233251 | 433.2127883 | 452.3281935 | 446.4647597 |
| AP53_775 | 551.7930532 | 620.1169953 | 613.6127449 | 523.8278998 | 576.6156937 | 604.080761 |
| AP53_776 | 229.2871606 | 257.1327229 | 247.0358892 | 220.5022503 | 238.9234732 | 246.3751316 |
| AP53_777 | 284.3448809 | 268.2948551 | 247.1492573 | 269.489884 | 249.1399207 | 270.3146164 |
| AP53_778 | 250.8018922 | 220.6336745 | 227.3647958 | 261.9877886 | 263.8044063 | 271.0490639 |
| AP53_779 | 719.6660199 | 757.0265231 | 691.0121874 | 629.1811879 | 582.7965915 | 656.6741311 |
| AP53_78 | 160.784967 | 182.0983303 | 168.3629667 | 171.4979474 | 170.8003856 | 143.2663028 |
| AP53_780 | 677.5868512 | 800.9210491 | 636.503455 | 555.8748559 | 521.7198733 | 599.873188 |
| AP53_781 | 715.8412702 | 697.268622 | 620.5131531 | 702.2268799 | 619.7477492 | 655.0190716 |
| AP53_782 | 119.537755 | 121.2886223 | 92.4398917 | 95.71011155 | 75.78102813 | 79.60907142 |
| AP53_783 | 128.9707583 | 154.4147507 | 149.5244099 | 103.4989236 | 99.75659155 | 110.5598049 |
| AP53_784 | 155.634949 | 169.0270036 | 164.202944 | 143.1114206 | 119.6514597 | 140.3364118 |
| AP53_785 | 106.5967593 | 106.1564245 | 94.96640471 | 116.158323 | 112.8504083 | 102.8609061 |
| AP53_786 | 18.17702481 | 16.77676464 | 9.938025897 | 11.13657553 | 12.95130914 | 9.641132116 |

| | | | | | | |
|---|---|---|---|---|---|---|
| AP53_787 | 54.65662145 | 47.60057736 | 54.88894063 | 52.20612574 | 53.61308215 | 47.80322293 |
| AP53_788 | 61.84784925 | 62.10295461 | 56.51652521 | 53.17599612 | 60.86037305 | 57.16526439 |
| AP53_789 | 80.3117954 | 58.07063375 | 47.34742646 | 63.09376248 | 62.98604509 | 60.38473498 |
| AP53_79 | 331.5849548 | 376.0912371 | 367.3249856 | 295.4935684 | 277.3942986 | 302.7152075 |
| AP53_790 | 561.7455309 | 600.5883524 | 475.5872789 | 488.9752704 | 463.5078942 | 501.4345852 |
| AP53_791 | 275.5877472 | 319.643354 | 306.1518879 | 295.3283813 | 312.2087105 | 328.3283862 |
| AP53_792 | 195.3891199 | 177.9890196 | 155.0769678 | 219.5928355 | 206.1392805 | 179.988438 |
| AP53_793 | 114.7229393 | 138.2836096 | 121.6776889 | 133.2648856 | 142.0894277 | 148.820576 |
| AP53_794 | 1153.456969 | 1285.171943 | 904.9449464 | 790.513437 | 586.3028752 | 725.5802605 |
| AP53_795 | 1497.574598 | 1719.508783 | 1446.945626 | 1348.125814 | 1070.524781 | 1296.804434 |
| AP53_796 | 1645.595707 | 1927.194331 | 1701.748648 | 1654.258207 | 1326.240662 | 1570.33324 |
| AP53_797 | 2060.688291 | 2429.498846 | 2293.897842 | 1910.803024 | 1568.748966 | 1835.802727 |
| AP53_798 | 85.76954698 | 84.47136947 | 89.26665306 | 92.9757596 | 81.8635329 | 75.93169052 |
| AP53_799 | 138.0756182 | 148.490543 | 145.5649757 | 109.6702961 | 107.8409635 | 108.8542247 |
| AP53_8 | 144.9767539 | 128.4905755 | 128.4822935 | 160.9051595 | 181.8287244 | 130.8439359 |
| AP53_80 | 408.743042 | 588.3226155 | 581.3763538 | 203.9711109 | 160.5292251 | 211.1286304 |
| AP53_800 | 288.8058423 | 345.4315041 | 336.9623199 | 360.5449456 | 347.0632819 | 359.5631007 |
| AP53_801 | 184.2452448 | 192.7234242 | 189.3282666 | 197.4256315 | 186.1471566 | 190.0917056 |
| AP53_802 | 87.11104461 | 98.87228351 | 83.93292745 | 71.4654443 | 75.83026099 | 75.20762003 |
| AP53_803 | 88.05505436 | 93.52006402 | 78.03873778 | 79.59684081 | 75.0646734 | 68.56386307 |
| AP53_804 | 388.7121903 | 531.8305249 | 496.7535437 | 295.0344954 | 275.0226503 | 328.5919656 |
| AP53_805 | 30.39413728 | 34.39450558 | 37.69391056 | 27.13161225 | 23.79067422 | 22.2556975 |
| AP53_806 | 91.57781174 | 108.033545 | 69.99955302 | 67.53077612 | 55.54859493 | 65.40114396 |
| AP53_807 | 122.5971526 | 126.9377043 | 112.6615793 | 115.1885735 | 88.36924302 | 97.50219029 |
| AP53_808 | 199.4079024 | 228.5288413 | 207.3937395 | 209.105583 | 159.1916424 | 200.1347794 |
| AP53_809 | 196.6880434 | 215.7559878 | 202.102709 | 206.3401605 | 160.2965267 | 179.1011638 |
| AP53_81 | 627.1942742 | 899.5520824 | 914.7890674 | 286.9361443 | 246.9453465 | 305.1446458 |
| AP53_810 | 191.495822 | 183.9692076 | 133.1036721 | 182.7212949 | 151.378474 | 158.0594745 |
| AP53_811 | 3437.155943 | 2935.22788 | 2509.850144 | 3612.929108 | 3164.644095 | 3476.575713 |
| AP53_814 | 49.21581744 | 46.38020911 | 47.67518159 | 57.36573798 | 58.85863154 | 52.33757434 |
| AP53_815 | 384.7420669 | 440.2434216 | 371.7114459 | 383.5443905 | 356.1065473 | 411.963524 |
| AP53_816 | 11235.84834 | 13147.84691 | 14165.67827 | 8500.095399 | 7682.85492 | 9071.654741 |
| AP53_817 | 473.8102968 | 497.5888106 | 503.3542048 | 599.1904794 | 631.7510339 | 601.362314 |
| AP53_818 | 595.3795374 | 612.2464937 | 586.0161577 | 702.4951847 | 718.4587693 | 709.6128444 |
| AP53_819 | 465.6731878 | 394.3667449 | 191.1628272 | 185.6075009 | 95.48995828 | 142.4171741 |

| | | | | | | |
|---|---|---|---|---|---|---|
| AP53_82 | 294.7856516 | 313.4142159 | 212.7193714 | 358.1157558 | 370.1555902 | 333.3302891 |
| AP53_820 | 1309.783946 | 1380.432927 | 918.5062881 | 542.8184215 | 311.577563 | 456.725119 |
| AP53_821 | 1156.242792 | 1261.618349 | 883.857105 | 582.9989043 | 299.5639459 | 513.3152986 |
| AP53_822 | 1030.58462 | 1011.029396 | 785.6110292 | 653.9816793 | 323.9186494 | 547.5534272 |
| AP53_823 | 57.0582317 | 54.13626241 | 49.32730119 | 40.34923054 | 33.20082309 | 38.03514266 |
| AP53_824 | 53.7086159 | 46.88273546 | 44.1686681 | 40.54899138 | 34.58663493 | 37.12385739 |
| AP53_825 | 65.77409379 | 65.30439758 | 57.69636261 | 52.28638169 | 41.5271104 | 46.45147179 |
| AP53_826 | 309.8142557 | 266.0881275 | 215.4683441 | 340.6950896 | 185.3300329 | 286.2112824 |
| AP53_827 | 322.9350256 | 289.2241402 | 268.4494728 | 378.6242326 | 265.2979238 | 328.839863 |
| AP53_828 | 709.2265119 | 797.0404896 | 793.7684223 | 775.0480351 | 609.2558891 | 732.1792303 |
| AP53_829 | 1005.669036 | 1109.365995 | 958.0933596 | 984.717808 | 723.4926258 | 974.4087099 |
| AP53_83 | 9.93057905 | 9.952524475 | 8.928347867 | 10.32961856 | 13.55485699 | 11.70488883 |
| AP53_830 | 185.0967625 | 227.0262759 | 194.4995081 | 177.610668 | 161.7575798 | 192.5999931 |
| AP53_831 | 986.1126664 | 1592.664752 | 1350.838369 | 319.1915932 | 281.388117 | 354.8456201 |
| AP53_832 | 1067.512828 | 1694.384471 | 1167.356364 | 306.361716 | 281.6854829 | 335.432372 |
| AP53_833 | 77.81854918 | 77.86092025 | 68.52606556 | 76.25634945 | 81.51693591 | 77.76321877 |
| AP53_834 | 131.7617232 | 129.9498446 | 112.611146 | 150.0807939 | 119.7463936 | 106.0854914 |
| AP53_835 | 960.6941635 | 1146.333486 | 1059.143477 | 894.4207316 | 725.4766918 | 904.7274588 |
| AP53_836 | 680.5136299 | 692.5010447 | 633.9787794 | 603.3168408 | 536.8827218 | 643.3898274 |
| AP53_837 | 700.7128311 | 738.4462802 | 664.9984856 | 650.8147248 | 533.7407723 | 693.2448744 |
| AP53_838 | 458.8537455 | 372.0056913 | 279.5831699 | 337.2823266 | 248.4790535 | 310.1873573 |
| AP53_839 | 620.6202262 | 623.1147928 | 607.126499 | 558.3065403 | 464.2040154 | 559.416588 |
| AP53_84 | 8.117966096 | 7.927125594 | 7.89725681 | 8.695329858 | 9.491201502 | 10.18912352 |
| AP53_840 | 273.2182868 | 306.9617214 | 305.841404 | 193.2948327 | 169.6494761 | 235.4129949 |
| AP53_841 | 277.3971363 | 311.4199315 | 309.3353823 | 221.4657015 | 190.865061 | 248.8889558 |
| AP53_842 | 229.7210109 | 241.0098785 | 259.3637249 | 180.3906708 | 170.2191319 | 213.4583032 |
| AP53_843 | 204.5511835 | 212.3565062 | 197.3085685 | 164.3087775 | 138.9105931 | 176.8161295 |
| AP53_844 | 270.695503 | 246.3538802 | 245.1527774 | 218.1673812 | 176.2011762 | 234.963546 |
| AP53_845 | 37.64772977 | 32.89400044 | 32.32096711 | 58.86532971 | 48.69515209 | 44.72269303 |
| AP53_846 | 46.09325708 | 40.50725193 | 42.37852662 | 58.06099155 | 52.59590418 | 54.09913083 |
| AP53_847 | 27.92465906 | 24.08310746 | 25.68666333 | 157.8383841 | 62.53160817 | 106.0719961 |
| AP53_848 | 25.53973229 | 19.59184223 | 21.00832251 | 202.3043401 | 81.27603754 | 116.7840991 |
| AP53_849 | 67.09918873 | 56.75557688 | 59.30105236 | 73.12720959 | 53.53037647 | 59.82184748 |
| AP53_85 | 8.766106461 | 7.576829066 | 8.869460177 | 9.317942099 | 13.07832293 | 8.066708706 |
| AP53_850 | 50.15993275 | 46.45687061 | 46.1965716 | 54.05187021 | 55.67101686 | 52.09527076 |

| | | | | | | |
|---|---|---|---|---|---|---|
| AP53_851 | 75.46831413 | 59.71234135 | 40.96875888 | 85.4929031 | 54.59759476 | 52.73407112 |
| AP53_852 | 68.05890719 | 85.43136853 | 77.83477441 | 47.80140766 | 55.16301576 | 52.33757434 |
| AP53_853 | 91.98461239 | 98.31332872 | 98.0034788 | 74.06854901 | 80.63143045 | 73.54067458 |
| AP53_854 | 63.92253725 | 79.30834191 | 77.11908096 | 59.69204487 | 58.05759271 | 65.55969839 |
| AP53_855 | 115.3048611 | 110.6714812 | 84.92994642 | 99.89982152 | 89.32174892 | 79.75216282 |
| AP53_856 | 105.0797522 | 112.6569487 | 95.15988145 | 153.0194914 | 181.2947194 | 124.1922221 |
| AP53_857 | 297.2309187 | 285.6586025 | 243.4507923 | 311.6282649 | 341.6081035 | 288.0371333 |
| AP53_858 | 604.4663921 | 650.995218 | 540.0850219 | 728.2802419 | 703.0597916 | 669.5923203 |
| AP53_859 | 289.8869828 | 230.5753303 | 241.5625674 | 294.3358512 | 271.5308951 | 211.6062273 |
| AP53_86 | 8.038632524 | 9.617714209 | 8.204011723 | 10.2149279 | 14.48359839 | 9.15907551 |
| AP53_860 | 392.6060101 | 344.3069933 | 255.0710643 | 379.7453783 | 308.9927706 | 339.2268233 |
| AP53_861 | 391.4823448 | 432.4995819 | 396.5849772 | 379.1367107 | 340.5034792 | 365.0745878 |
| AP53_862 | 50.16419334 | 45.49403392 | 31.07255102 | 47.8007211 | 42.03101291 | 36.16173854 |
| AP53_863 | 205.2651347 | 210.4789469 | 199.6123487 | 193.8360745 | 203.4966474 | 194.2214673 |
| AP53_864 | 197.2326635 | 180.9310381 | 178.6707452 | 201.432311 | 192.3952109 | 202.7471339 |
| AP53_865 | 141.3093144 | 143.0191409 | 148.0568414 | 154.4783156 | 166.5037339 | 155.2796908 |
| AP53_866 | 1190.602425 | 1380.865086 | 1138.667876 | 1419.215668 | 1091.899746 | 1261.083312 |
| AP53_867 | 106.1672894 | 105.1029201 | 87.93733201 | 99.751898 | 101.9587736 | 102.4944164 |
| AP53_868 | 393.0136199 | 411.8032605 | 336.9906806 | 461.681749 | 409.3623387 | 408.4947678 |
| AP53_869 | 423.7216582 | 483.086718 | 439.0006383 | 361.2613444 | 385.2973119 | 389.053405 |
| AP53_87 | 10.20060886 | 7.519620052 | 10.89912106 | 7.713840674 | 9.022387339 | 9.738763834 |
| AP53_870 | 491.9224298 | 515.6840056 | 460.7713967 | 434.6633874 | 436.3365453 | 470.9816157 |
| AP53_871 | 198.6541919 | 183.8074051 | 190.894934 | 205.9696064 | 224.1371642 | 227.078055 |
| AP53_872 | 214.8137056 | 218.9037596 | 242.5493042 | 230.7039211 | 275.7298265 | 268.1626232 |
| AP53_873 | 271.3954382 | 258.0942033 | 302.9983354 | 315.0589259 | 359.2391808 | 379.4596424 |
| AP53_874 | 208.0640211 | 191.2736332 | 192.3214392 | 284.4701237 | 290.1823784 | 264.7060609 |
| AP53_875 | 187.9074371 | 136.2435644 | 88.19837693 | 150.1641475 | 163.9586756 | 129.9997814 |
| AP53_876 | 168.3798398 | 166.547518 | 152.4890694 | 151.0119643 | 154.8797935 | 135.7471402 |
| AP53_877 | 259.6941436 | 288.4523195 | 267.0369643 | 224.5257368 | 240.4179801 | 246.5317825 |
| AP53_878 | 218.5246251 | 219.042995 | 215.6352859 | 188.0776695 | 210.3758342 | 192.0134759 |
| AP53_879 | 343.6103574 | 365.2410822 | 336.9283371 | 293.0965782 | 301.1649776 | 324.0546005 |
| AP53_88 | 8.495147458 | 9.592733133 | 11.45957193 | 10.27329892 | 11.46983954 | 7.453592484 |
| AP53_880 | 502.1874583 | 486.2938652 | 440.2690265 | 457.2692665 | 430.897952 | 470.2063136 |
| AP53_881 | 593.1349738 | 604.389407 | 617.1746523 | 564.7514269 | 567.890018 | 617.3960112 |
| AP53_882 | 549.9330614 | 479.0266893 | 481.4857705 | 555.3393452 | 548.7360364 | 506.3339228 |

| | | | | | | |
|---|---|---|---|---|---|---|
| AP53_883 | 420.2497841 | 367.006978 | 383.2341299 | 400.6761692 | 374.7126353 | 398.9693292 |
| AP53_884 | 385.0252127 | 374.1191989 | 346.9605082 | 345.6467664 | 351.0127104 | 372.0336828 |
| AP53_885 | 262.8399892 | 255.7966755 | 275.1038808 | 250.5342809 | 269.7336529 | 267.3623188 |
| AP53_886 | 2161.859454 | 2643.787163 | 2422.613232 | 2289.00345 | 2135.037922 | 2187.916751 |
| AP53_887 | 1035.212966 | 1551.474089 | 1388.766402 | 681.9033849 | 671.8940938 | 749.9272314 |
| AP53_888 | 550.595749 | 539.3622204 | 504.1401098 | 646.9444951 | 607.5462836 | 609.1834807 |
| AP53_889 | 488.1597547 | 449.5074807 | 440.6059558 | 553.1260881 | 535.6625734 | 475.1728375 |
| AP53_89 | 9.731891635 | 9.559768414 | 7.786494517 | 7.678496616 | 9.272640105 | 9.631192805 |
| AP53_890 | 93.09797925 | 102.0531256 | 98.95393206 | 107.8214191 | 115.8433439 | 96.36940319 |
| AP53_891 | 3839.866196 | 3575.891124 | 3418.820041 | 4623.349333 | 4839.189672 | 3387.942031 |
| AP53_892 | 5867.263329 | 6105.300169 | 5091.373709 | 8255.369198 | 8282.615876 | 7344.207813 |
| AP53_893 | 1125.915212 | 1038.729569 | 1166.277029 | 1472.812114 | 1581.176208 | 1446.350589 |
| AP53_894 | 806.0603049 | 644.8617227 | 755.2203562 | 1045.688518 | 1135.315094 | 995.7645596 |
| AP53_895 | 545.7682613 | 488.9799189 | 573.5065731 | 668.4672645 | 723.8514934 | 645.2475237 |
| AP53_896 | 544.7305354 | 548.8933793 | 611.8115629 | 528.2845137 | 643.8664358 | 583.0091387 |
| AP53_897 | 461.83643 | 403.7168961 | 410.9464054 | 432.2948645 | 476.4647242 | 433.8071219 |
| AP53_898 | 95.89303555 | 84.07788481 | 88.68566078 | 93.79950387 | 126.4589451 | 93.16651003 |
| AP53_899 | 315.3508993 | 320.0645809 | 299.7331006 | 267.0030061 | 287.8801145 | 281.6867223 |
| AP53_9 | 122.0841304 | 101.7737089 | 116.3390193 | 147.1608646 | 161.4951897 | 110.2043602 |
| AP53_90 | 182.9991979 | 200.9782543 | 161.4610441 | 178.538217 | 167.999068 | 167.4555504 |
| AP53_900 | 51.65224405 | 45.72858634 | 47.55529429 | 53.60405024 | 57.5117521 | 48.07303125 |
| AP53_901 | 199.3467699 | 215.6573439 | 192.7419255 | 203.5081443 | 200.2086792 | 213.8613533 |
| AP53_902 | 24.14797374 | 27.09227482 | 22.57678039 | 27.2089973 | 22.0508414 | 20.98044195 |
| AP53_903 | 16.33624039 | 14.90299612 | 18.5585535 | 15.31544293 | 14.86954902 | 19.53936109 |
| AP53_904 | 36.20904772 | 47.30604418 | 37.9581625 | 38.19559855 | 33.36116767 | 39.76760991 |
| AP53_905 | 37.00977253 | 37.6253435 | 30.97289896 | 39.42288659 | 36.9573715 | 36.53912352 |
| AP53_906 | 35.47292605 | 34.18427358 | 30.16039263 | 30.18276228 | 30.94991983 | 28.3155148 |
| AP53_907 | 37.11664894 | 32.47539863 | 24.93759119 | 28.6747619 | 25.30972245 | 24.63475482 |
| AP53_908 | 48.0901415 | 46.00574715 | 41.702335 | 40.76367383 | 38.58291566 | 39.33705508 |
| AP53_909 | 50.31220942 | 42.93044149 | 41.68081676 | 39.81119521 | 39.12684774 | 47.66111516 |
| AP53_91 | 394.6145529 | 442.2602089 | 424.0981566 | 419.7110163 | 339.0308216 | 361.6342747 |
| AP53_910 | 151.9250422 | 112.619232 | 89.49041744 | 117.2107701 | 98.35281003 | 115.0435394 |
| AP53_911 | 133.3435435 | 136.0186184 | 109.9007488 | 139.403722 | 111.2623254 | 124.3577151 |
| AP53_912 | 75.21248933 | 76.95281637 | 72.84352138 | 88.3996618 | 62.60295638 | 83.85642467 |
| AP53_913 | 187.3233586 | 177.6663817 | 134.9255865 | 176.6714993 | 134.4473364 | 146.1895159 |

| | | | | | | |
|---|---|---|---|---|---|---|
| AP53_914 | 233.9370811 | 259.2360511 | 221.4326468 | 274.7302888 | 187.7668247 | 245.5622407 |
| AP53_915 | 244.3677478 | 262.021665 | 206.4853396 | 313.3095729 | 202.718271 | 265.6311136 |
| AP53_916 | 214.1969302 | 228.9716228 | 172.5035326 | 324.6063648 | 196.9654162 | 267.8436382 |
| AP53_917 | 216.0316596 | 239.1779532 | 186.8895994 | 342.5562735 | 205.7649445 | 275.3631734 |
| AP53_918 | 837.1046421 | 840.4046714 | 788.2128642 | 1202.727314 | 1365.680496 | 1255.863208 |
| AP53_919 | 788.6871821 | 797.8862535 | 701.0592102 | 1049.616019 | 1190.899089 | 1069.051845 |
| AP53_92 | 69.75658235 | 68.57181657 | 75.82812156 | 98.04596996 | 101.9921122 | 99.02683621 |
| AP53_920 | 691.8878285 | 665.3587101 | 643.0754586 | 932.8195315 | 1118.591298 | 1066.608978 |
| AP53_921 | 235.973668 | 241.9604824 | 197.2830363 | 263.5764549 | 347.4983639 | 238.5290566 |
| AP53_922 | 643.7946323 | 698.3375849 | 618.9898476 | 897.6173933 | 947.9802833 | 933.0302757 |
| AP53_923 | 371.21524 | 332.6120686 | 284.0950336 | 503.5217018 | 517.7546345 | 416.6575929 |
| AP53_924 | 445.662162 | 405.3817488 | 392.6859517 | 589.1965405 | 589.5308652 | 526.7451152 |
| AP53_925 | 43.39923121 | 40.31464838 | 38.42814231 | 41.00869204 | 44.91194853 | 42.15474084 |
| AP53_926 | 430.7210588 | 533.7022258 | 437.8362307 | 431.1825542 | 408.7621405 | 423.638007 |
| AP53_929 | 118.2059571 | 106.7686332 | 101.1941899 | 101.7299145 | 98.79243005 | 91.4484782 |
| AP53_93 | 410.8078335 | 350.6682074 | 401.2147333 | 622.8959694 | 472.7358167 | 564.7265001 |
| AP53_930 | 72.53870762 | 71.0429564 | 82.26975675 | 88.18730846 | 83.75852209 | 72.38624194 |
| AP53_931 | 83.21387449 | 72.59552511 | 68.99534657 | 85.165131 | 77.57186689 | 73.4627579 |
| AP53_932 | 70.37903366 | 63.710602 | 67.35115272 | 79.95587337 | 69.60787575 | 65.43434088 |
| AP53_933 | 605.7614162 | 568.9185797 | 382.9921697 | 641.6007839 | 658.1019815 | 544.8434122 |
| AP53_934 | 377.5580003 | 363.5202467 | 326.6413726 | 303.9887084 | 278.1772075 | 288.4238151 |
| AP53_935 | 632.6909986 | 727.0264486 | 678.1608957 | 432.5198014 | 400.7152528 | 487.6179985 |
| AP53_936 | 413.5896866 | 401.2181206 | 379.5109946 | 277.4134068 | 212.2832531 | 272.6171887 |
| AP53_937 | 426.4573205 | 419.8367193 | 376.6173787 | 279.8810829 | 216.0587033 | 260.0678992 |
| AP53_938 | 393.1762432 | 433.1718555 | 445.3027598 | 284.4614209 | 222.7645706 | 302.0740306 |
| AP53_939 | 721.1147798 | 829.2681936 | 810.9500757 | 420.7498567 | 305.6251461 | 362.0722823 |
| AP53_94 | 451.0097693 | 415.7047505 | 431.3639223 | 597.6962551 | 504.0673605 | 607.3290351 |
| AP53_940 | 328.5965762 | 304.4000358 | 281.2491173 | 251.1180732 | 183.466821 | 230.8747142 |
| AP53_941 | 405.4854957 | 434.2722719 | 388.5211596 | 305.9239573 | 224.3524021 | 318.277874 |
| AP53_942 | 515.8583178 | 565.1758001 | 552.9047423 | 390.9121627 | 299.937017 | 438.847747 |
| AP53_943 | 153.4450402 | 159.6928793 | 151.5157451 | 146.6325421 | 154.4993162 | 133.3022159 |
| AP53_944 | 205.0674678 | 197.1940245 | 189.9844766 | 191.4120337 | 187.3302089 | 170.0971166 |
| AP53_945 | 226.0615986 | 227.5539349 | 226.9050073 | 246.1628656 | 228.6575959 | 208.5376877 |
| AP53_946 | 394.3653043 | 474.5724302 | 456.6802136 | 321.8310958 | 283.5851135 | 294.4970457 |
| AP53_947 | 315.5546702 | 366.2225722 | 373.9987567 | 291.2183083 | 254.3474157 | 288.3816753 |

| | | | | | | |
|---|---|---|---|---|---|---|
| AP53_948 | 266.4325147 | 316.3504931 | 341.3910656 | 234.983626 | 210.6754674 | 236.8975531 |
| AP53_949 | 201.6309895 | 236.2476477 | 225.4932242 | 161.1931044 | 128.1269806 | 150.6676883 |
| AP53_95 | 112.2128332 | 103.4058766 | 79.66321559 | 90.16171553 | 86.99706969 | 86.77018904 |
| AP53_950 | 1155.639308 | 1042.631818 | 849.5398275 | 1060.795942 | 851.3383194 | 955.9361033 |
| AP53_951 | 908.571594 | 862.7656524 | 666.6305458 | 888.5685154 | 858.1942811 | 873.6348948 |
| AP53_952 | 143.2618844 | 157.7693725 | 122.2196494 | 130.7186488 | 110.3951367 | 138.12626 |
| AP53_953 | 42.71107426 | 41.24888267 | 35.4272819 | 44.88293869 | 39.57982918 | 34.66724998 |
| AP53_954 | 1263.380904 | 1624.628256 | 934.2355914 | 621.7635787 | 539.1454793 | 564.8096565 |
| AP53_955 | 104.5837515 | 111.6682674 | 112.4120355 | 99.62877313 | 115.5326678 | 96.28556726 |
| AP53_956 | 135.8429654 | 146.3069446 | 133.5352664 | 102.3931402 | 118.8701919 | 105.8382059 |
| AP53_957 | 231.3445381 | 290.6025523 | 288.9207323 | 182.8020168 | 175.7247814 | 185.2168603 |
| AP53_958 | 165.235826 | 178.7201172 | 160.2046244 | 126.1028164 | 130.160452 | 135.7055683 |
| AP53_959 | 153.934733 | 193.7405858 | 180.1351647 | 121.9054456 | 140.1255039 | 140.6805037 |
| AP53_96 | 100.8939198 | 69.16458628 | 57.00301647 | 73.61906801 | 76.68826579 | 73.27260408 |
| AP53_960 | 194.5032088 | 215.0106286 | 219.3722803 | 144.2003692 | 156.1555941 | 158.9851465 |
| AP53_961 | 151.7576876 | 174.0151363 | 156.8247346 | 115.4050214 | 128.2547746 | 138.8095025 |
| AP53_962 | 147.3329425 | 186.2211098 | 187.3215953 | 136.1803301 | 142.9453511 | 164.2175269 |
| AP53_963 | 186.4245728 | 217.23509 | 230.4026422 | 146.842492 | 159.9002222 | 162.0110519 |
| AP53_964 | 111.9072484 | 128.7765886 | 111.8589628 | 92.38515705 | 109.1638081 | 107.3519102 |
| AP53_965 | 113.2389481 | 119.4931295 | 114.413507 | 99.31284713 | 108.9866164 | 100.2971006 |
| AP53_966 | 54.40657427 | 49.22298565 | 38.48874906 | 37.56534739 | 42.54293991 | 36.63630204 |
| AP53_967 | 429.9562032 | 439.8223731 | 409.4231102 | 588.2025724 | 604.030533 | 474.3722664 |
| AP53_968 | 164.1679524 | 181.7883333 | 182.8238868 | 208.0297986 | 221.8193506 | 186.4210546 |
| AP53_969 | 271.2176414 | 351.7640336 | 296.4321102 | 292.7039361 | 325.0892137 | 332.7693943 |
| AP53_97 | 155.0555298 | 195.8444632 | 170.7894923 | 142.7911467 | 131.6242766 | 146.4575615 |
| AP53_970 | 93.51340527 | 96.01004272 | 88.30302849 | 97.85437359 | 108.3541911 | 88.06645759 |
| AP53_971 | 233.1654534 | 233.3686687 | 209.2082181 | 229.5042243 | 280.4684324 | 230.6134624 |
| AP53_972 | 243.5477747 | 266.6999745 | 245.3363997 | 238.9427831 | 267.693304 | 251.6674618 |
| AP53_973 | 59.70845909 | 58.03851367 | 67.57224471 | 50.62872214 | 56.06695958 | 52.07856587 |
| AP53_974 | 47.96714881 | 49.34936808 | 45.31739809 | 38.412619 | 41.04874109 | 36.32317125 |
| AP53_975 | 41.34185201 | 39.86482847 | 38.47388678 | 31.61064667 | 38.25435263 | 30.66519204 |
| AP53_976 | 652.9628997 | 626.684141 | 572.9615581 | 752.9615655 | 686.2023469 | 673.3204483 |
| AP53_977 | 7635.004645 | 7054.821344 | 6662.028214 | 13013.89335 | 10039.87567 | 11529.98368 |
| AP53_978 | 9978.942453 | 9078.995444 | 8584.603232 | 15818.20573 | 13065.65127 | 16395.56538 |
| AP53_979 | 8973.18255 | 7428.039639 | 7330.508747 | 13305.74782 | 9907.012129 | 11405.08436 |

| | | | | | | |
|---|---|---|---|---|---|---|
| AP53_98 | 17.70423856 | 20.30658667 | 20.02662794 | 23.08308384 | 24.17036377 | 16.79163844 |
| AP53_980 | 3362.63418 | 2184.622841 | 1565.248544 | 4282.681488 | 3156.27728 | 3091.696822 |
| AP53_981 | 5256.447084 | 4312.61538 | 4577.353399 | 7260.831796 | 5668.872953 | 6620.036107 |
| AP53_982 | 3367.917997 | 3855.82605 | 2935.075016 | 2989.65798 | 2688.728367 | 3038.606063 |
| AP53_983 | 304.7210862 | 292.1047219 | 329.513569 | 324.8433913 | 288.6333019 | 341.4095754 |
| AP53_984 | 334.1907151 | 318.4628337 | 346.6455546 | 381.1971521 | 437.0431372 | 428.1712034 |
| AP53_985 | 309.5138489 | 292.2422074 | 321.034901 | 302.3958685 | 304.030573 | 303.5833378 |
| AP53_986 | 119.4939763 | 101.030055 | 115.6940292 | 174.3935931 | 170.5619213 | 136.3211239 |
| AP53_987 | 477.6083849 | 422.1551115 | 396.540769 | 672.7960533 | 738.437515 | 579.6808436 |
| AP53_988 | 653.1206982 | 659.0436061 | 700.7652257 | 864.597057 | 960.1364777 | 880.3180005 |
| AP53_989 | 465.4337624 | 373.1939267 | 368.9626788 | 599.2684967 | 659.7153962 | 559.4739957 |
| AP53_99 | 496.8500034 | 512.6000034 | 500.2560752 | 456.0167481 | 396.1940281 | 460.6929383 |
| AP53_990 | 426.9761489 | 378.7844846 | 247.627211 | 421.5273387 | 462.2995319 | 425.8970112 |
| AP53_991 | 2556.070254 | 2259.37769 | 2158.256689 | 3388.065097 | 3259.941582 | 3409.386379 |
| AP53_992 | 703.848279 | 619.834215 | 546.0015544 | 964.1448357 | 995.3395615 | 764.4381087 |
| AP53_993 | 2366.53253 | 3140.387502 | 3148.850612 | 2351.753112 | 2299.6797 | 2479.824265 |
| AP53_994 | 1431.657609 | 1745.884525 | 1458.111864 | 1254.995853 | 1137.838969 | 1316.251284 |
| AP53_995 | 254.5432027 | 235.1497338 | 243.775846 | 268.6378282 | 252.1545343 | 260.8525903 |
| AP53_996 | 434.9537268 | 388.2824739 | 408.5875086 | 439.7079266 | 436.0370121 | 396.9425145 |
| AP53_997 | 408.8530366 | 347.5500221 | 367.012576 | 448.1090335 | 439.271145 | 405.9082134 |
| AP53_998 | 302.8740978 | 251.2905537 | 292.5451373 | 415.4367735 | 366.8776217 | 324.6927227 |
| AP53_999 | 229.961531 | 224.6152111 | 186.0592361 | 229.0592586 | 234.6093161 | 203.5548684 |
| AP53_pseudo11 | 15.28771906 | 13.29375241 | 15.95683031 | 16.34858252 | 18.43896924 | 20.23473825 |
| AP53_pseudo13 | 4.261554403 | 4.134400165 | 7.882974989 | 2.654455742 | 3.882752196 | 3.013783201 |
| AP53_pseudo13 | 134.3375349 | 123.6025201 | 133.1053321 | 160.3564078 | 163.7017993 | 155.2111027 |
| AP53_pseudo16 | 7.662561676 | 7.420833458 | 7.928546544 | 7.039729906 | 10.55953481 | 6.571273223 |
| AP53_pseudo16 | 31.55768783 | 29.58597908 | 25.88056534 | 35.51243667 | 39.87516642 | 25.49169268 |
| AP53_pseudo16 | 3614.094075 | 3256.044988 | 3324.518496 | 5057.695533 | 5550.677836 | 4933.280558 |
| AP53_pseudo26 | 240.1218863 | 236.3775261 | 245.4440969 | 220.0700026 | 235.036022 | 216.3054105 |
| AP53_pseudo38 | 72.26066479 | 76.2054821 | 74.66003445 | 62.63776221 | 61.36689043 | 60.54179545 |
| AP53_pseudo39 | 20.93936008 | 16.94189027 | 22.89658565 | 17.39410333 | 22.98391435 | 23.58281647 |
| AP53_pseudo59 | 86.39319565 | 80.87509762 | 101.2775185 | 71.40367267 | 86.85903066 | 62.94782805 |
| AP53_pseudo60 | 135.2963137 | 152.4408582 | 163.1243967 | 145.7067417 | 241.2431809 | 196.1553315 |
| AP53_pseudo65 | 311.7579719 | 282.2490567 | 327.5415902 | 391.3179468 | 474.086894 | 375.431412 |
| AP53_pseudo73 | 104.3851391 | 120.5248027 | 117.8478654 | 85.14516792 | 84.0906775 | 85.41609745 |

| | | | | | | |
|---|---|---|---|---|---|---|
| AP53_pseudo81 | 352.6893632 | 362.4451307 | 332.327586 | 390.6170742 | 399.3252653 | 406.0523476 |
| AP53_pseudo92 | 43.00994904 | 45.83702389 | 42.34116253 | 53.17404319 | 59.99671963 | 51.2091062 |
| AP53_rna1 | 5.683196365 | 11.06223951 | 5.072167141 | 12.51807478 | 167.0498408 | 13.50978893 |
| AP53_rna10 | 5.276386369 | 12.677896 | 3.147041534 | 2.821265802 | 2.121335118 | 3.816281463 |
| AP53_rna11 | 13.00191892 | 17.89820612 | 9.596625478 | 6.571889751 | 10.18240857 | 9.051321681 |
| AP53_rna12 | 8.867440482 | 11.7478573 | 3.061986357 | 3.431269219 | 6.19200521 | 3.71313872 |
| AP53_rna13 | 14.00122181 | 14.09742876 | 7.348767258 | 8.235046125 | 3.440002895 | 7.426277441 |
| AP53_rna14 | 6.235409777 | 10.83056957 | 4.770252431 | 11.77375682 | 158.5362347 | 12.77755104 |
| AP53_rna15 | 10.40821421 | 10.12246882 | 11.79493923 | 6.260891232 | 4.184551466 | 6.022405815 |
| AP53_rna16 | 9.847140301 | 10.8518 | 6.398117499 | 11.15682385 | 178.0869598 | 15.1578915 |
| AP53_rna17 | 0.295182454 | 0 | 0 | 1.736163571 | 0 | 0 |
| AP53_rna18 | 24.60123358 | 21.43581633 | 12.4157255 | 16.69570995 | 9.763953421 | 5.269605088 |
| AP53_rna19 | 20.81642841 | 30.36740647 | 22.96909218 | 16.00005537 | 6.276827199 | 4.516804361 |
| AP53_rna2 | 8.042710978 | 14.29054422 | 6.207862752 | 7.652200395 | 3.487126222 | 3.764003634 |
| AP53_rna20 | 23.18193164 | 42.27619332 | 18.62358826 | 14.60874621 | 3.487126222 | 12.79761236 |
| AP53_rna21 | 32.00929735 | 45.58741699 | 16.57953589 | 19.81767198 | 12.41757142 | 12.06317262 |
| AP53_rna22 | 14.66612002 | 25.60389173 | 9.311794128 | 12.52178246 | 6.276827199 | 12.79761236 |
| AP53_rna23 | 6.715400833 | 4.829674668 | 3.147041534 | 2.821265802 | 1.414223412 | 1.526512585 |
| AP53_rna24 | 11.78298902 | 11.76167831 | 10.6629172 | 7.766778797 | 4.791721679 | 5.818706795 |
| AP53_rna25 | 11.20097745 | 9.985678705 | 7.348767258 | 4.117523063 | 4.816004052 | 6.683649697 |
| AP53_rna26 | 14.9346366 | 20.55875027 | 14.08513724 | 11.66631534 | 8.256006947 | 8.911532929 |
| AP53_rna27 | 28.46915102 | 35.2435719 | 25.7206854 | 24.70513838 | 16.51201389 | 14.85255488 |
| AP53_rna28 | 14.46792921 | 22.90832173 | 14.08513724 | 13.03882303 | 11.00800926 | 12.62467165 |
| AP53_rna29 | 19.57059671 | 22.21650347 | 14.09874607 | 19.18460745 | 15.83930222 | 19.53936109 |
| AP53_rna3 | 10.02553052 | 12.393504 | 6.491748487 | 11.13933667 | 172.9151097 | 16.06622956 |
| AP53_rna30 | 11.66768485 | 15.85960735 | 11.02315089 | 7.548792281 | 6.19200521 | 8.168905185 |
| AP53_rna31 | 10.88131485 | 16.67230159 | 9.311794128 | 6.956545814 | 5.579401955 | 7.528007269 |
| AP53_rna32 | 20.42995183 | 31.22282637 | 15.3185571 | 10.72875728 | 6.453639233 | 10.06208295 |
| AP53_rna33 | 35.93646932 | 41.70489341 | 20.82150723 | 20.58761531 | 6.880005789 | 7.426277441 |
| AP53_rna34 | 18.44554904 | 23.215368 | 7.724556493 | 6.347848055 | 3.471275648 | 5.62034179 |
| AP53_rna35 | 6.212400885 | 10.3672297 | 5.796762447 | 12.21358107 | 156.7385409 | 13.29011756 |
| AP53_rna36 | 11.3544155 | 16.07686225 | 10.55336668 | 9.739164139 | 4.184551466 | 3.764003634 |
| AP53_rna37 | 9.60928667 | 11.74988 | 7.240796389 | 11.47159318 | 173.4761239 | 16.48255118 |
| AP53_rna38 | 0 | 0 | 0 | 0 | 0.870291331 | 0.46969618 |
| AP53_rna39 | 14.19301937 | 13.09966554 | 8.691007853 | 6.956545814 | 2.789700978 | 7.528007269 |

| | | | | | | |
|---|---|---|---|---|---|---|
| AP53_rna4 | 0 | 0 | 0.387328189 | 0.434040893 | 0.870291331 | 0.46969618 |
| AP53_rna40 | 15.07925016 | 17.14194389 | 11.48891783 | 15.73551067 | 4.302426155 | 3.096025525 |
| AP53_rna41 | 15.86805139 | 28.78225038 | 11.02315089 | 19.21510763 | 2.752002316 | 6.683649697 |
| AP53_rna42 | 10.07310125 | 11.47047734 | 7.552899682 | 14.81164546 | 3.535558531 | 6.869306633 |
| AP53_rna43 | 14.19827605 | 15.9379264 | 14.60227272 | 15.79908849 | 8.485340473 | 14.04391578 |
| AP53_rna44 | 12.13439224 | 8.810892975 | 6.736369986 | 5.49003075 | 2.752002316 | 6.683649697 |
| AP53_rna45 | 2.365503229 | 9.527029481 | 4.345503927 | 4.173927488 | 2.092275733 | 6.775206542 |
| AP53_rna46 | 7.248369153 | 11.26924089 | 5.594740505 | 8.150323428 | 1.885631216 | 4.749150265 |
| AP53_rna47 | 16.53853243 | 25.71291584 | 11.48891783 | 9.298256306 | 5.019497181 | 10.06208295 |
| AP53_rna48 | 20.34332777 | 34.53548187 | 18.62358826 | 17.39136453 | 10.46137867 | 16.56161599 |
| AP53_rna49 | 19.66653101 | 41.05223468 | 23.91751566 | 16.22227836 | 7.778228767 | 19.08140731 |
| AP53_rna5 | 30.27844133 | 18.45861962 | 14.27808433 | 14.60874621 | 4.881976711 | 15.05601454 |
| AP53_rna50 | 7.811792806 | 12.93662857 | 7.552899682 | 3.627341746 | 7.273148977 | 5.233757434 |
| AP53_rna51 | 5.338062983 | 10.42514718 | 5.464656265 | 14.20970651 | 157.5865097 | 12.59449157 |
| AP53_rna52 | 8.515811624 | 11.90878685 | 9.932580403 | 2.086963744 | 1.394850489 | 6.022405815 |
| AP53_rna53 | 9.37143304 | 12.438408 | 6.600984639 | 12.04867027 | 172.3891588 | 15.91483988 |
| AP53_rna54 | 0 | 0 | 0 | 0.868081785 | 0.870291331 | 0.93939236 |
| AP53_rna55 | 3.266951757 | 1.762178595 | 1.224794543 | 0 | 1.376001158 | 0 |
| AP53_rna56 | 0.466707394 | 0 | 0 | 0 | 2.064001737 | 1.485255488 |
| AP53_rna57 | 40.292405 | 51.91900268 | 59.16438084 | 56.42531604 | 86.97473985 | 59.53399082 |
| AP53_rna58 | 9.80661709 | 15.02565452 | 11.18948101 | 6.896427516 | 13.82796225 | 4.749150265 |
| AP53_rna59 | 11.51211571 | 19.31869867 | 3.776449841 | 11.28506321 | 9.899563886 | 8.395819218 |
| AP53_rna6 | 20.34332777 | 30.96284581 | 19.24437453 | 7.652200395 | 6.974252444 | 10.53921018 |
| AP53_rna60 | 20.1462025 | 19.31869867 | 14.47639106 | 9.169113857 | 19.09201606 | 9.15907551 |
| AP53_rna61 | 0.972854849 | 4.897698255 | 1.914819638 | 1.43050097 | 0.717071026 | 3.096025525 |
| AP53_rna62 | 0 | 1.811128 | 0 | 0 | 0.707111706 | 3.05302517 |
| AP53_rna63 | 7.467318301 | 10.57307157 | 5.511575443 | 4.117523063 | 2.752002316 | 1.485255488 |
| AP53_rna64 | 10.88131485 | 19.05405896 | 10.55336668 | 16.69570995 | 7.671677688 | 9.786409449 |
| AP53_rna65 | 25.69167287 | 34.9856921 | 15.47423349 | 15.48255623 | 9.313178568 | 16.08423016 |
| AP53_rna66 | 25.07433423 | 36.91723924 | 17.38201571 | 13.21743705 | 9.066528177 | 11.2920109 |
| AP53_rna67 | 17.97782454 | 25.00845239 | 19.86516081 | 14.60874621 | 4.881976711 | 11.2920109 |
| AP53_rna68 | 25.07433423 | 17.26774094 | 16.76122943 | 12.52178246 | 6.276827199 | 10.53921018 |
| AP53_rna69 | 0 | 0 | 0 | 0 | 1.740582661 | 0 |
| AP53_rna7 | 23.18193164 | 32.74916384 | 16.14044316 | 5.565236651 | 6.974252444 | 9.033608722 |
| AP53_rna70 | 9.086008683 | 12.139048 | 6.866272438 | 10.77210579 | 165.7972414 | 13.94677407 |

| | | | | | | |
|---|---|---|---|---|---|---|
| AP53_rna71 | 8.98891227 | 10.12246882 | 10.55336668 | 4.869582069 | 4.184551466 | 5.269605088 |
| AP53_rna72 | 5.31505409 | 8.861375106 | 5.706188034 | 12.58574005 | 157.6882659 | 14.09557923 |
| AP53_rna73 | 50.81209694 | 63.4519327 | 59.38141819 | 44.94568278 | 38.03773316 | 54.32279268 |
| AP53_rna74 | 7.630123206 | 13.64663889 | 13.70060873 | 8.266964909 | 7.104005978 | 12.78010536 |
| AP53_rna75 | 1.400122181 | 1.17478573 | 1.224794543 | 0 | 0 | 0 |
| AP53_rna76 | 1.400122181 | 1.17478573 | 1.837191814 | 2.058761531 | 0 | 0.742627744 |
| AP53_rna77 | 0 | 0.371513436 | 0 | 0 | 0.435145665 | 0 |
| AP53_rna78 | 9.133579409 | 10.567408 | 7.084744743 | 11.52405473 | 174.563089 | 14.49556165 |
| AP53_rna79 | 5.677207749 | 8.336150796 | 13.65729805 | 8.347854976 | 4.881976711 | 3.011202907 |
| AP53_rna8 | 30.32459749 | 40.28655454 | 18.79014067 | 15.48255623 | 12.41757142 | 12.73334888 |
| AP53_rna80 | 5.384080767 | 9.990766051 | 5.072167141 | 12.11208316 | 154.160716 | 12.48465589 |
| AP53_rna81 | 212.42219 | 306.6512614 | 232.1740669 | 197.5659011 | 172.9614606 | 170.1329643 |
| AP53_rna82 | 10.21497592 | 3.673273691 | 1.914819638 | 5.72200388 | 6.453639233 | 6.96605743 |
| AP53_rna83 | 7.000610907 | 5.286535785 | 10.41075362 | 8.235046125 | 2.752002316 | 4.455766464 |
| AP53_rna84 | 2.878028929 | 4.829674668 | 5.035266455 | 1.410632901 | 0.707111706 | 1.526512585 |
| AP53_rna85 | 23.33536969 | 23.4957146 | 20.20910996 | 22.64637684 | 24.08002026 | 20.79357683 |
| AP53_rna9 | 24.60123358 | 18.45861962 | 11.79493923 | 12.52178246 | 9.763953421 | 12.04481163 |

| deltaSK_4hr_Rp1 | deltaSK_4hr_Rp2 | deltaSK_4hr_Rp3 | deltaSK_8hr_Rp1 | deltaSK_8hr_Rp2 | deltaSK_8hr_Rp3 | function |
|---|---|---|---|---|---|---|
| 235.5312121 | 220.9672345 | 260.6943505 | 258.7567053 | 238.1247468 | 180.4773261 | Chromoson |
| 95.40247125 | 88.31072961 | 134.7870638 | 105.5444063 | 139.3063485 | 151.6882718 | FIG011146 |
| 631.3649905 | 736.3045971 | 511.5066942 | 497.4414951 | 519.8736031 | 520.4389347 | tRNA dihyd |
| 137.1875511 | 113.6247191 | 134.0398035 | 187.1368098 | 94.18908287 | 47.79945881 | Paratox |
| 138.969051 | 129.9363051 | 131.2985338 | 198.3979302 | 425.8594677 | 650.7627024 | Phage prote |
| 370.9454536 | 361.5571304 | 394.8089742 | 541.4193623 | 1260.959111 | 1911.879336 | Streptococ |
| 87.41117273 | 70.60252855 | 125.7392273 | 93.63124083 | 74.75939557 | 80.93108544 | Streptococ |
| 546.5801294 | 521.7817007 | 466.6724123 | 773.9409237 | 782.0637095 | 792.4023497 | Phage lysin |
| 223.9198499 | 199.8897716 | 226.3556906 | 249.3100027 | 224.9810277 | 316.5138702 | Phage prote |
| 108.5639495 | 87.45369547 | 39.10695343 | 37.52158514 | 71.04050743 | 111.7248103 | Phage lysin |
| 106.3825872 | 69.79285193 | 35.12164043 | 30.97124625 | 49.8594693 | 68.62238394 | Phage holir |
| 83.81364525 | 48.62721623 | 31.69782941 | 25.20940366 | 25.46559669 | 39.20007373 | Phage prote |
| 96.68666936 | 63.52469101 | 42.74581079 | 29.84619894 | 36.521578 | 64.36821591 | FIG011165 |
| 356.4268359 | 435.6822113 | 280.888529 | 260.1606219 | 370.8827913 | 311.7529683 | Chaperonir |
| 97.86419509 | 74.13356739 | 41.99461493 | 39.88979493 | 40.25542449 | 51.06534522 | Phage hyalu |
| 75.46410893 | 47.14894886 | 41.04093288 | 29.220972 | 25.82777098 | 35.35006648 | Hyaluronat |
| 110.220588 | 85.35168691 | 34.58012028 | 28.64210336 | 56.15535866 | 58.57788732 | Phage hyalu |
| 122.7411241 | 81.71152578 | 33.48686286 | 35.00529039 | 52.74498701 | 80.38999891 | Phage-relat |
| 188.9247074 | 131.9425605 | 65.20397752 | 64.99395573 | 108.0458357 | 132.600476 | Phage tail le |
| 138.8615021 | 97.38503128 | 56.44928311 | 49.06506493 | 49.08445164 | 76.80115967 | Phage prote |
| 139.7774032 | 92.71184745 | 48.38509981 | 37.81160967 | 59.26159482 | 100.3031179 | Phage prote |
| 150.0277652 | 90.19462302 | 60.85876265 | 33.74774121 | 37.52663495 | 89.89423523 | Phage majo |
| 132.6117188 | 79.57768599 | 40.5480766 | 29.41139667 | 49.914048 | 67.93521035 | Phage prote |
| 431.2593924 | 313.4531971 | 139.3951685 | 152.0653568 | 314.9196088 | 330.8112257 | Phage prote |
| 70.68131767 | 57.15582212 | 73.45966977 | 75.02355085 | 66.87756536 | 52.75593678 | Transcriptic |
| 149.0915545 | 106.2206204 | 34.4813355 | 34.58241128 | 76.67329859 | 74.15284382 | Phage prote |
| 227.0573589 | 175.5247997 | 46.46885823 | 34.09617185 | 78.53512262 | 83.77274639 | DNA packag |
| 278.446481 | 214.9114878 | 46.88478664 | 35.78326596 | 110.0404916 | 112.1720915 | conserved h |
| 446.0579846 | 298.3314925 | 126.1687413 | 77.90787019 | 137.7346992 | 181.6915735 | Phage capsi |
| 383.8825857 | 290.5394687 | 97.27875172 | 82.0992618 | 175.9080618 | 234.1992423 | Prophage C |
| 142.888296 | 99.78123166 | 36.4968689 | 29.61424263 | 69.70715736 | 112.5855022 | Phage porta |
| 106.2172652 | 67.68493547 | 38.92312474 | 38.10065842 | 57.59242326 | 77.17439895 | Phage prote |
| 23.06171658 | 21.50653808 | 11.318152 | 15.42536778 | 17.22261461 | 37.37429774 | hypothetica |
| 70.98477581 | 52.22652797 | 25.03343056 | 20.16504275 | 37.25384022 | 44.1114353 | Phage term |

| | | | | | | |
|---|---|---|---|---|---|---|
| 76.95431931 | 56.7197825 | 34.94479431 | 28.88548438 | 58.52376926 | 56.16993514 | Phage term |
| 117.4619359 | 107.3055307 | 117.9997731 | 108.3792768 | 103.7466819 | 85.10986627 | putative co |
| 48.0683972 | 37.22285436 | 27.58799551 | 22.65600893 | 15.84480544 | 30.67513116 | Phage endo |
| 107.0400904 | 115.325245 | 80.19691256 | 111.4216868 | 128.2288985 | 103.2595676 | FIG011147 |
| 48.78991613 | 43.60010325 | 52.28712399 | 66.17980371 | 26.91727993 | 50.24374932 | hypothetica |
| 42.92813508 | 34.94812437 | 47.32402306 | 62.90657798 | 17.05038846 | 60.29249918 | hypothetica |
| 94.53657266 | 87.79793975 | 71.98030281 | 92.04040822 | 57.81057637 | 76.91635698 | putative me |
| 19.73957302 | 16.51352074 | 22.32259436 | 19.77772472 | 7.682783735 | 12.81579693 | FIG011188 |
| 52.18060691 | 46.98351395 | 45.32919877 | 49.553994 | 20.25379478 | 46.54157832 | conserved h |
| 47.91195952 | 34.99851946 | 42.28613765 | 47.7181808 | 26.39836055 | 46.27491534 | FIG011146 |
| 69.44962593 | 56.53747041 | 53.07891329 | 49.86241697 | 53.40216489 | 74.01333051 | hypothetica |
| 69.43236407 | 66.86904154 | 46.9133097 | 59.27109593 | 64.50032885 | 75.11727084 | putative rep |
| 161.2335503 | 150.1912632 | 179.4735532 | 161.4154557 | 167.2561916 | 146.4246409 | Signal pepti |
| 38.84184788 | 27.74500529 | 28.2021159 | 31.72325986 | 28.62802654 | 46.25793233 | Phage prote |
| 76.47633236 | 64.14947776 | 51.58567062 | 54.09757509 | 48.74066949 | 71.70975584 | FIG011158 |
| 100.3442677 | 102.4120861 | 73.89136379 | 82.95726066 | 81.93043807 | 124.7656813 | FIG011175 |
| 88.52792406 | 84.02184476 | 59.75766601 | 67.04720155 | 59.90820251 | 59.9060088 | FIG011159 |
| 86.68982617 | 78.74701634 | 62.13665449 | 79.63346117 | 89.07536276 | 99.21818287 | hypothetica |
| 94.53807671 | 95.99042409 | 66.99475358 | 95.53345566 | 88.20098387 | 84.40949886 | hypothetica |
| 69.54130549 | 69.43754287 | 41.93375317 | 46.79320374 | 51.22631862 | 100.4874986 | hypothetica |
| 86.68982617 | 71.55059783 | 58.99586731 | 59.77330015 | 54.76791446 | 112.1228932 | hypothetica |
| 81.25359125 | 74.14001379 | 75.38258304 | 52.16540816 | 49.08445164 | 107.9148355 | hypothetica |
| 27.67574384 | 32.06763525 | 29.97595409 | 25.71868454 | 17.45224947 | 44.25510041 | hypothetica |
| 136.0737676 | 110.3349552 | 135.0927034 | 109.7989371 | 112.7654457 | 91.17116494 | Fibronectin |
| 79.99312187 | 70.32385593 | 65.06019071 | 50.29584961 | 61.56354951 | 55.52360094 | hypothetica |
| 304.5520829 | 301.1586013 | 240.8357446 | 244.3175017 | 752.9820203 | 870.9821841 | hypothetica |
| 212.8346373 | 202.6577626 | 181.2319089 | 174.6601539 | 271.4284063 | 284.3619918 | hypothetica |
| 137.1301308 | 140.4885346 | 123.7163731 | 112.809759 | 188.1969707 | 290.6784717 | conserved h |
| 28.2038818 | 24.32799644 | 18.11679536 | 22.83429871 | 12.37197137 | 23.67650653 | Phage trans |
| 234.7154829 | 223.3371262 | 201.7460594 | 217.4012772 | 205.8102446 | 171.3576293 | hypothetica |
| 156.6159146 | 132.7155598 | 133.3278306 | 151.4256937 | 145.7262831 | 111.6527763 | hypothetica |
| 43.03036364 | 21.94353595 | 32.25673321 | 31.79663076 | 10.74301206 | 19.33910351 | FIG011152 |
| 20.07976915 | 24.41641995 | 19.87245171 | 18.97182511 | 26.29524195 | 33.3762049 | hypothetica |
| 124.1713224 | 111.269129 | 122.158579 | 122.5121713 | 116.6082334 | 129.2200458 | Pleiotropic |
| 154.710753 | 142.356055 | 158.7511126 | 134.9600166 | 178.975703 | 138.1910725 | NPQTN spec |

| | | | | | | |
|---|---|---|---|---|---|---|
| 35.69736737 | 28.18958318 | 45.5690358 | 29.07559403 | 31.78802582 | 46.25615017 | hypothetica |
| 39.26668859 | 28.35864234 | 42.60910075 | 40.41988159 | 28.55822641 | 34.82459319 | Phage integ |
| 251.8708309 | 252.4480756 | 268.416536 | 206.6263411 | 235.3208203 | 177.8677271 | lipoprotein |
| 96.99721559 | 77.49913316 | 102.4311495 | 100.1039968 | 83.89215416 | 64.89762164 | Maltodextr |
| 80.52352019 | 59.29911306 | 92.55998344 | 95.70774754 | 68.95478387 | 57.70968797 | 4-alpha-glu |
| 49.18961177 | 44.79150142 | 46.8671359 | 42.61541404 | 70.33513188 | 66.85374174 | Maltose op |
| 215.5203174 | 175.5240516 | 204.3572605 | 236.2357069 | 226.9683922 | 193.9215479 | Maltose/ma |
| 52.08256309 | 45.52192933 | 43.9089452 | 51.40232001 | 46.87889478 | 39.44244991 | Maltose/ma |
| 71.35418798 | 59.82339748 | 66.18989162 | 87.51753804 | 72.76462078 | 48.76705131 | Maltose/ma |
| 45.92893675 | 26.54311195 | 50.17714054 | 58.41286841 | 38.83125507 | 33.27252733 | Mobile elen |
| 277.6422003 | 253.6260633 | 262.1682449 | 245.1384101 | 297.8123974 | 328.3275211 | FIG011151 |
| 64.95180904 | 52.44658356 | 68.68147127 | 89.73577718 | 65.40916814 | 51.59863694 | Mobile elen |
| 200.8666704 | 211.5952827 | 148.3022978 | 170.219874 | 191.5490796 | 143.6235956 | Transposase |
| 692.5554456 | 799.1363507 | 576.5891061 | 534.170963 | 626.7246447 | 706.6599523 | putative est |
| 709.6520162 | 867.4878944 | 579.7316236 | 503.4052129 | 614.391376 | 557.5248845 | Poly(glycer |
| 729.7157796 | 877.5598107 | 660.4566124 | 577.2341339 | 756.8822443 | 1070.694298 | D-alanine--|
| 755.1758928 | 933.0765727 | 552.0212541 | 543.2771275 | 663.6592731 | 588.1996161 | D-alanyl tra |
| 581.4825057 | 682.2658162 | 431.4416665 | 380.849474 | 567.0450179 | 538.4406165 | D-alanine--|
| 124.0609211 | 124.7482605 | 105.1703906 | 81.66607474 | 107.9857936 | 198.0441119 | FIG011141 |
| 171.7087742 | 206.9393875 | 175.881781 | 151.1479868 | 215.0845188 | 257.8775567 | Excinucleas |
| 137.6885081 | 172.3808926 | 100.9302059 | 83.80628911 | 93.15890408 | 100.8201469 | Glutamine / |
| 257.1831353 | 227.2610611 | 229.0067576 | 199.6256512 | 228.7677328 | 181.4774229 | Multiple su |
| 227.496674 | 300.7097602 | 187.5330724 | 172.2004999 | 168.1987039 | 153.9452206 | Glutamate |
| 17.39026679 | 10.16938113 | 13.59842674 | 17.95845514 | 26.5633503 | 47.68230328 | FIG011143 |
| 29.81593645 | 25.4194333 | 26.54168156 | 37.90951799 | 75.71810192 | 77.63751525 | PTS system, |
| 20.905585 | 18.33570233 | 20.64430925 | 28.20914133 | 54.18923461 | 99.14766532 | Putative un |
| 30.68528102 | 27.08258074 | 28.22464156 | 38.92495151 | 81.16464682 | 98.87252098 | PTS system, |
| 25.4314736 | 22.66776388 | 29.15512425 | 25.71512634 | 44.17600647 | 53.72216273 | PTS system, |
| 24.0064134 | 19.04864801 | 22.65247279 | 24.79076965 | 42.51525435 | 48.08289033 | Transcriptic |
| 17.70335965 | 17.69700272 | 21.74015444 | 21.27855528 | 33.02778445 | 33.8629105 | Outer surfa |
| 18.34715704 | 16.8268911 | 19.68558665 | 21.41632299 | 31.48972529 | 31.67400385 | Beta-glucos |
| 104.8555988 | 88.33653637 | 82.03220945 | 101.9312625 | 47.39188434 | 35.08399737 | Ribosyl nico |
| 68.269828 | 99.26545273 | 75.71982468 | 46.46786967 | 103.3314641 | 128.2516195 | Fibronectin |
| 99.50771593 | 117.724105 | 115.6373455 | 96.08112338 | 146.6976819 | 356.4462216 | FIG011140 |
| 303.4086823 | 355.1959245 | 269.5057424 | 287.1852184 | 433.063824 | 462.7013408 | GTP-bindin |

| | | | | | | |
|---|---|---|---|---|---|---|
| 88.76402371 | 82.60203444 | 102.396374 | 66.45463678 | 34.70156581 | 70.10755719 | FIG011149 |
| 3.959391099 | 3.83770514 | 6.376330983 | 5.857379844 | 12.89893729 | 16.82581373 | Mobile elen |
| 13.29757765 | 17.19898763 | 11.41158014 | 15.89831538 | 48.15832991 | 47.96856067 | Mobile elen |
| 15.2082572 | 13.3224403 | 11.42871011 | 11.28171559 | 21.06405607 | 21.69465202 | Mobile elen |
| 47.89366553 | 40.83001757 | 44.505424 | 53.94571812 | 47.43143574 | 37.29432939 | Quinolinate |
| 63.60394855 | 60.95916729 | 74.36764006 | 90.9593698 | 59.93469884 | 62.31604607 | Malonate p |
| 20.0393672 | 13.79186939 | 25.67372643 | 18.87836784 | 14.02412904 | 13.32997431 | Mobile elen |
| 62.26663476 | 54.59351973 | 54.49690189 | 59.19484886 | 54.25123602 | 35.54084162 | Mobile elen |
| 163.4702451 | 157.5661453 | 175.6074602 | 169.6408417 | 220.3651605 | 185.9838553 | FIG011142 |
| 242.6723577 | 224.6757258 | 263.3432762 | 221.1902368 | 318.151693 | 339.604525 | FIG011142 |
| 93.35827434 | 104.6375795 | 75.54866462 | 72.78845422 | 88.35201295 | 103.6607881 | Mobile elen |
| 140.52593 | 148.5320711 | 122.3508467 | 132.6526313 | 145.6440286 | 124.9504389 | Ribosomal s |
| 112.1611118 | 102.0272336 | 126.6473378 | 96.9933218 | 147.4142875 | 166.7104076 | Dihydrolipo |
| 167.4034448 | 145.8875748 | 190.4494592 | 197.8933508 | 135.0244522 | 118.0666384 | Transporter |
| 309.4564098 | 364.7137883 | 270.0768299 | 273.8762364 | 267.3763947 | 223.3055525 | ATP-depenc |
| 144.7837044 | 161.1508849 | 139.2576131 | 121.7985313 | 234.4332019 | 292.7635781 | Phenylaceti |
| 142.2602276 | 156.887438 | 130.8189736 | 133.6865208 | 155.9220709 | 160.0748224 | Uridine pho |
| 172.8497094 | 178.5692714 | 135.2219346 | 112.2106121 | 130.6751626 | 132.2974485 | 23S rRNA (U |
| 317.7656774 | 340.1413494 | 353.3578501 | 412.8656257 | 342.5770199 | 302.0438285 | Cell envelop |
| 99.37105954 | 108.2892525 | 111.6200981 | 116.9423938 | 96.73228518 | 108.8206083 | Shikimate k |
| 80.47103616 | 85.9203304 | 108.2904615 | 109.5151778 | 76.10706446 | 65.64897566 | Short chain |
| 164.9622946 | 166.7838985 | 161.0952505 | 194.6461567 | 156.2445255 | 134.2963767 | 5-Enolpyruv |
| 320.7352715 | 348.4922307 | 300.1779536 | 324.4267424 | 444.4429416 | 400.5144589 | Inner memb |
| 744.0344277 | 874.0900368 | 690.8785332 | 710.7493503 | 1129.797518 | 1053.403095 | Methionine |
| 220.5047809 | 239.3800605 | 244.1488019 | 255.8903629 | 249.0921237 | 200.2236889 | FIG009439 |
| 162.4006603 | 173.8346144 | 196.3027162 | 158.1792635 | 176.7040259 | 147.3446527 | Ribosomal- |
| 32.61957071 | 29.84658231 | 37.13963319 | 45.03782841 | 24.13694136 | 19.3599768 | Protein G-r |
| 192.5772414 | 204.4617909 | 149.3793954 | 180.9519595 | 167.6818362 | 184.6577415 | UDP-N-acet |
| 389.9057521 | 454.9066085 | 307.8133125 | 316.7709455 | 397.3011153 | 478.0082015 | S-adenosyl |
| 18.55775675 | 16.47613819 | 19.18577961 | 19.80913563 | 13.68869186 | 11.88974221 | internalin, |
| 234.8902465 | 262.3723885 | 175.3060994 | 161.183426 | 236.1681195 | 211.9838188 | Biotin oper |
| 48.50254097 | 42.59948888 | 53.63742972 | 53.56699984 | 47.2114923 | 34.90618374 | Transcriptio |
| 457.4261246 | 503.3843658 | 421.9465322 | 460.0956206 | 507.3022443 | 532.2881913 | FIG011143 |
| 432.1920625 | 524.6766631 | 396.66516 | 448.798237 | 646.6810411 | 574.7451942 | DNA polym |
| 365.8859004 | 407.2040744 | 343.6507993 | 323.2067554 | 643.4202817 | 647.4784852 | GAF domai |

| | | | | | | |
|---|---|---|---|---|---|---|
| 56.60462831 | 54.54143837 | 64.03558889 | 56.51326294 | 50.46608806 | 48.75504564 | FIG011156 |
| 25.16654069 | 17.22336031 | 27.39612957 | 20.32503512 | 22.05438375 | 25.87897225 | FIG011139 |
| 149.0549721 | 164.9837222 | 137.6698853 | 125.7693339 | 203.1955386 | 201.3596384 | Uridine kin |
| 371.3515099 | 399.0294558 | 334.4631163 | 444.7843013 | 578.1117901 | 517.9270026 | ATP-depend |
| 494.2065389 | 531.0635722 | 580.735045 | 562.8467176 | 555.2895141 | 446.5900881 | Peptidogly |
| 1544.229073 | 1904.81093 | 1034.24845 | 979.9159321 | 2126.140457 | 2497.07212 | Non-phosp |
| 1596.853941 | 2011.661294 | 1166.376685 | 1186.462312 | 2203.195525 | 1998.066997 | Phosphoen |
| 33.96054948 | 26.85699619 | 39.89218778 | 36.63524848 | 21.0752481 | 17.60439217 | 3-ketoacyl- |
| 5731.578561 | 6819.452512 | 3888.21929 | 2720.791664 | 4483.195324 | 4901.962888 | Phosphoca |
| 343.0577508 | 383.0519122 | 340.8825958 | 291.5296468 | 294.1739339 | 283.7494453 | Glutaredox |
| 623.9780406 | 725.1988555 | 446.0389386 | 505.2611353 | 382.7769154 | 226.4876567 | Ribonucleo |
| 756.0551575 | 730.8087073 | 461.5784918 | 710.0260063 | 689.5196778 | 569.9076708 | FIG011161 |
| 450.0837832 | 509.9944635 | 324.4825756 | 329.488266 | 256.4049042 | 172.8384977 | Ribonucleo |
| 131.5021239 | 129.8421338 | 135.4151856 | 175.56256 | 95.2406377 | 81.91328129 | Chloride ch |
| 32.64477559 | 28.38559166 | 39.99176617 | 34.20535955 | 19.43343595 | 41.63053515 | Mobile elen |
| 39.45097269 | 31.71632059 | 41.04873089 | 33.39494672 | 32.53395423 | 27.13525234 | Mobile elen |
| 94.00140592 | 82.84937422 | 115.2622062 | 95.77275088 | 134.4744718 | 128.2081879 | Membrane- |
| 95.087739 | 86.62536536 | 106.4853181 | 87.54958987 | 89.58878653 | 89.33101257 | FIG011141 |
| 38.73004384 | 32.8613886 | 49.33813966 | 37.13482005 | 33.1989382 | 27.2229769 | Acetyl-CoA: |
| 62.03046055 | 53.47913181 | 83.7779043 | 52.91758114 | 36.81333873 | 47.73156185 | Transcriptio |
| 33.43485817 | 22.35209427 | 40.14171244 | 24.01644067 | 15.70702452 | 25.01022188 | FIG011143 |
| 355.2839188 | 427.6710306 | 246.9690776 | 187.267808 | 251.1009863 | 236.5427373 | Alanyl-tRNA |
| 903.506053 | 1014.461427 | 772.7870203 | 529.4938257 | 793.4390052 | 879.7223744 | Foldase pro |
| 138.0130612 | 155.4091097 | 97.34621272 | 93.87782423 | 127.7948759 | 104.6146638 | O-methyltr |
| 153.6521598 | 119.6251944 | 174.5824946 | 198.096613 | 108.3548496 | 105.9274234 | Major facili |
| 805.2165008 | 725.1174167 | 880.0408409 | 983.9181021 | 658.1882515 | 495.8941212 | Oligoendop |
| 674.7493168 | 516.1904131 | 749.190263 | 734.4168971 | 344.1110591 | 240.6065154 | Competenc |
| 2517.576345 | 2119.664181 | 3186.533352 | 2583.244875 | 2111.493992 | 1862.928908 | Ribosomal |
| 2436.626131 | 2028.40797 | 3211.603895 | 2247.767363 | 1750.581302 | 1457.111491 | Glucosamir |
| 29.32882296 | 28.30392146 | 36.58749831 | 32.22545005 | 21.72441471 | 19.81836779 | Butyrate-ac |
| 236.1782562 | 275.6999274 | 202.6155443 | 172.2817953 | 213.8536575 | 201.0335906 | S-adenosyl |
| 237.9915082 | 259.2612962 | 194.2933205 | 226.6074645 | 246.0252613 | 190.852178 | FIG011150 |
| 112.4708253 | 87.9156373 | 117.2376699 | 145.0532681 | 175.4083957 | 180.1633732 | FIG011139 |
| 141.3007699 | 123.5204233 | 148.6833796 | 157.1308704 | 194.1903618 | 168.3165602 | FIG011147 |
| 102.3350315 | 72.5368444 | 115.3798534 | 127.9885283 | 136.429604 | 220.8148367 | FIG011167 |

| | | | | | | |
|---|---|---|---|---|---|---|
| 3010.078081 | 3130.581279 | 2085.9088 | 1909.657668 | 4577.198013 | 4985.871306 | Manganese |
| 135.326739 | 147.4706796 | 144.1327517 | 151.8726367 | 134.8337156 | 127.708896 | DNA polym |
| 14.12253403 | 10.86274802 | 16.51648238 | 12.39133405 | 11.70677296 | 9.135227149 | Late compe |
| 15.47924846 | 10.95164121 | 18.0258215 | 16.65992069 | 9.861310646 | 11.18548627 | Late compe |
| 268.4691584 | 298.9281535 | 275.684064 | 302.5745219 | 339.1000335 | 392.5114927 | 1-acyl-sn-gl |
| 237.98208 | 208.535439 | 227.8575846 | 225.13732 | 149.2859269 | 103.1435559 | YheO-like P |
| 162.7783646 | 166.0832343 | 154.1846939 | 182.9640692 | 138.8009744 | 107.1607997 | tRNA (adeni |
| 157.524162 | 135.3631112 | 154.1733109 | 192.4335364 | 112.31367 | 86.2247569 | COG2827: |
| 50.43326354 | 32.28088952 | 57.30768943 | 65.85556661 | 34.94047622 | 36.63951974 | Kup system |
| 1699.077372 | 2007.206424 | 1233.013627 | 1086.167274 | 1369.965225 | 1322.75044 | Cold-shock |
| 654.1067859 | 632.5893598 | 707.5811943 | 775.2089715 | 495.5337494 | 465.6766296 | Peptide cha |
| 713.3047861 | 718.6150137 | 759.354088 | 680.9892633 | 545.7683252 | 639.6549051 | membrane |
| 289.4870598 | 322.1155527 | 260.777296 | 293.1623283 | 255.6266573 | 228.3886921 | UDP-N-acet |
| 270.1915714 | 270.4647439 | 273.9973799 | 278.3858996 | 293.6628511 | 242.5520597 | D-alanine--I |
| 163.9466465 | 165.2187622 | 159.7437717 | 140.8339954 | 152.2357967 | 135.7338675 | Recombina |
| 103.7969605 | 80.93902888 | 119.4739772 | 112.4420319 | 86.53966397 | 131.4840271 | Formate/ni |
| 152.7935516 | 125.9881748 | 146.0806975 | 114.8605127 | 103.1578148 | 102.6290355 | Endoribonu |
| 277.9886303 | 250.9400777 | 269.4595614 | 265.8585574 | 234.0867329 | 175.7698457 | Substrate-s |
| 98.98477748 | 79.36739725 | 99.99587294 | 77.39196242 | 105.0407265 | 75.36562398 | hypothetic |
| 149.8197184 | 110.3755086 | 179.6366349 | 152.4363213 | 165.8715952 | 179.606966 | Transcripti |
| 1174.573849 | 1336.903026 | 999.8196918 | 913.6704643 | 959.008631 | 941.2040282 | Phosphogly |
| 9.746193475 | 7.25368444 | 16.95546233 | 15.02984553 | 13.59261738 | 3.607243541 | FIG011179 |
| 347.5634725 | 317.147203 | 331.5103046 | 391.5392181 | 488.0127211 | 424.1087763 | Dihydroorc |
| 899.4644332 | 625.1300289 | 1242.440379 | 1311.289239 | 895.0295871 | 835.7787818 | FIG011142 |
| 1270.371591 | 1033.759052 | 1543.128873 | 1803.710797 | 1240.311135 | 1170.89781 | Lead, cadm |
| 105.1510423 | 106.6648351 | 118.7153542 | 136.9317484 | 57.9357462 | 32.94672086 | Paratox |
| 175.4651897 | 178.4702636 | 183.0856478 | 181.4386022 | 213.3330554 | 200.41894 | Streptococ |
| 248.1373692 | 243.6491269 | 228.4535693 | 173.5864274 | 177.0505044 | 165.3906007 | Transcripti |
| 62.00809111 | 42.55392418 | 74.01280098 | 66.51781345 | 54.32567275 | 61.31440595 | Streptococ |
| 15.05738881 | 11.26465034 | 20.81592143 | 19.98696438 | 12.47663894 | 15.74139141 | Phage lysin, |
| 22.14256634 | 12.73691983 | 19.18587686 | 18.31762424 | 11.16613118 | 12.28709699 | Phage prote |
| 3.322565957 | 2.527799123 | 2.48128717 | 5.123810976 | 4.668171624 | 2.670297167 | FIG011165 |
| 4.559298841 | 4.445018748 | 5.837708451 | 5.704983981 | 6.676807563 | 5.210462892 | Phage hyalu |
| 5.832651404 | 3.717838261 | 5.15934308 | 4.530165135 | 6.702930493 | 5.618655838 | Hyaluronat |
| 4.431612056 | 3.412411182 | 5.894005533 | 5.853797349 | 5.643961412 | 4.087106214 | Phage hyalu |

| | | | | | | |
|---|---|---|---|---|---|---|
| 3.822860372 | 3.590365359 | 4.240648116 | 4.105674399 | 4.31604661 | 3.724967622 | FIG022125 |
| 41.74012299 | 41.86121628 | 46.12411384 | 42.96938023 | 37.43262854 | 31.36712574 | V-type ATP s |
| 4.815747953 | 2.859329808 | 5.704435007 | 4.754192551 | 5.195707858 | 4.807035558 | Phage minc |
| 8.940560547 | 5.576757392 | 8.45438572 | 7.977029911 | 7.125162335 | 5.510604764 | Phage prote |
| 7.918782198 | 3.571890065 | 4.215368544 | 3.954941597 | 3.346667157 | 2.283806787 | conserved h |
| 6.154493937 | 3.908721149 | 7.019599973 | 6.928427822 | 6.103766007 | 3.332227934 | Phage majo |
| 3.817984274 | 1.543566778 | 4.032091651 | 1.962602597 | 4.382540325 | 2.512187466 | FIG018217 |
| 9.238579231 | 4.303277079 | 9.984226945 | 4.797473015 | 14.02412904 | 15.55163669 | FIG033577 |
| 7.309645106 | 11.78723722 | 14.88772302 | 5.988454079 | 8.684172213 | 14.30014404 | FIG042840 |
| 10.9070019 | 7.858158143 | 12.02019775 | 8.121965465 | 6.853300795 | 4.739976351 | FIG048485 |
| 16.20803374 | 12.03998499 | 13.8647362 | 13.92567925 | 8.037220151 | 10.22504372 | FIG040428 |
| 5.136507372 | 6.229891141 | 8.863336602 | 6.600945672 | 4.068260857 | 7.966576469 | FIG011151 |
| 96.46516496 | 96.82569684 | 144.5537753 | 111.1866982 | 199.6329865 | 226.7410226 | tRNA(Ile)-ly |
| 55.44334763 | 47.38457579 | 62.94173654 | 44.79167187 | 41.1514623 | 37.24214342 | V-type ATP s |
| 11.49898885 | 12.00600554 | 14.11681086 | 10.23991912 | 13.9491671 | 14.05482164 | hypothetica |
| 12.11813639 | 10.12035518 | 15.27307444 | 14.43206758 | 24.54222582 | 18.42270808 | conserved h |
| 10.77668856 | 11.2850241 | 15.88583478 | 10.6738036 | 12.69552734 | 10.50207693 | Phage prote |
| 7.800591419 | 7.740872201 | 9.388153695 | 4.647755404 | 5.567788544 | 6.299216332 | Phage prote |
| 7.073087012 | 8.392207726 | 7.933694899 | 6.93579947 | 5.655037147 | 5.398357532 | Recombina |
| 9.502538638 | 5.572148502 | 9.677019962 | 5.994858842 | 5.890134197 | 5.298431746 | FIG011145 |
| 6.197307807 | 3.872135897 | 11.45347773 | 7.964184452 | 6.829141098 | 12.2332607 | Phage prote |
| 9.435144747 | 8.359742706 | 17.15783681 | 9.743417149 | 8.354800279 | 16.24904233 | hypothetica |
| 5.279188132 | 10.07456172 | 8.270957233 | 3.992302719 | 3.020581639 | 6.441506323 | FIG011141 |
| 4.54255723 | 6.944418824 | 10.8772705 | 6.95785727 | 7.001193102 | 9.659263435 | DNA replica |
| 72.35787234 | 61.58831445 | 75.60171845 | 51.51831818 | 44.2987176 | 50.24374932 | V-type ATP s |
| 3.320779632 | 3.548845613 | 4.942563959 | 4.431683284 | 0.950021645 | 0.972459664 | Phage prote |
| 3.045685461 | 2.871250091 | 3.566850307 | 4.051013053 | 3.775727049 | 4.444639363 | Phage excis |
| 25.42932875 | 14.38817688 | 28.62217172 | 21.67155544 | 17.14499156 | 21.79587999 | Cro-like pha |
| 153.0964558 | 112.8206991 | 195.4604429 | 181.7951517 | 138.2155829 | 127.5234209 | Phage CI-lik |
| 177.7835263 | 115.5558053 | 216.940515 | 224.3644264 | 166.1494502 | 198.4192459 | Phage prote |
| 75.11411776 | 54.40897948 | 87.4995637 | 93.75162013 | 63.07545281 | 66.99166576 | Phage integ |
| 2556.905913 | 2614.033389 | 2456.245049 | 1530.318043 | 1621.7076 | 2524.202276 | DNA-bindir |
| 315.862445 | 318.7365165 | 277.5560232 | 345.1376853 | 373.0418325 | 303.3081847 | YfaA |
| 461.149295 | 489.2192831 | 431.6203447 | 528.1473011 | 659.0242674 | 660.1420729 | FIG006988 |
| 416.1264279 | 451.6227367 | 375.4956337 | 460.3270571 | 487.802663 | 444.9756278 | Hypothetic |

| | | | | | | |
|---|---|---|---|---|---|---|
| 65.94924251 | 69.07119517 | 79.40118943 | 46.87433075 | 50.64508549 | 51.94430699 | V-type ATP s |
| 392.2607079 | 453.7959583 | 415.3314535 | 476.7900323 | 563.2995005 | 497.3807029 | FIG011153 |
| 323.3836254 | 378.6951374 | 315.5803141 | 377.3298336 | 392.7110531 | 383.6663486 | DNA repair |
| 180.9719142 | 181.3883128 | 199.6013778 | 188.3098441 | 160.8219231 | 204.3032456 | Arginine pa |
| 322.9485957 | 341.0326748 | 380.3606457 | 395.8863355 | 277.4338571 | 411.9259274 | RNA bindin |
| 260.5850802 | 281.6515101 | 265.8686412 | 308.5669898 | 318.3911028 | 366.3131909 | Octaprenyl |
| 159.2555087 | 150.6146977 | 207.4287149 | 200.9850438 | 161.1606162 | 212.6985388 | Exodeoxyril |
| 264.1365606 | 297.484904 | 288.6508564 | 355.7389173 | 385.6478616 | 387.3826793 | Exodeoxyril |
| 176.0470316 | 185.1768073 | 186.9758711 | 193.2027315 | 199.2312058 | 185.743208 | Methylenet |
| 871.9630366 | 955.6911127 | 958.9098955 | 1131.899496 | 1574.038458 | 1779.108932 | putative ph |
| 106.381257 | 100.851936 | 103.358809 | 109.0093511 | 93.65731113 | 86.07716373 | Deoxyriboc |
| 64.9673062 | 59.79751572 | 71.63319582 | 47.47180096 | 51.35442328 | 43.81617058 | V-type ATP s |
| 190.180059 | 171.2757734 | 218.9508054 | 295.6240255 | 133.83026 | 158.0729877 | Glutamate t |
| 128.0837086 | 108.5043496 | 134.2387194 | 153.3401121 | 74.24827095 | 67.13793577 | ABC transpo |
| 240.2373405 | 232.6831242 | 345.6078558 | 199.5907369 | 258.0439743 | 727.9471005 | FIG011141 |
| 4132.730224 | 4196.876014 | 4921.652665 | 6001.681725 | 4111.096949 | 2909.251735 | ATP-depend |
| 552.0224748 | 432.0953011 | 539.8758505 | 698.9619776 | 524.2994453 | 462.3747141 | Mutator mu |
| 974.872652 | 820.82741 | 1007.943069 | 1161.264832 | 946.3093488 | 1411.998595 | FIG011140 |
| 343.7972357 | 386.749265 | 242.0809116 | 204.5337293 | 201.4879953 | 184.6861244 | Isoleucyl-tR |
| 848.7181743 | 984.1644227 | 763.1968576 | 603.2507714 | 819.7297433 | 694.5950103 | Cell division |
| 506.202153 | 582.9324586 | 520.6285614 | 434.696651 | 599.1277915 | 787.9894685 | FIG001583 |
| 337.6196081 | 325.0504004 | 288.792635 | 239.6376254 | 403.5319389 | 482.3399935 | Cell division |
| 68.08672544 | 58.89946571 | 73.5574103 | 48.44754823 | 51.92859743 | 40.57065366 | V-type ATP s |
| 1080.931849 | 1211.089286 | 961.5894189 | 806.6446036 | 1363.427106 | 1522.087354 | FIG001960 |
| 763.7382617 | 939.8216815 | 770.0575037 | 669.4110359 | 1337.551307 | 1579.807032 | Hypothetic |
| 943.4149155 | 1130.074579 | 720.3148457 | 641.3666342 | 1195.206397 | 1397.415697 | Cell division |
| 539.2777697 | 634.6628471 | 500.7237509 | 428.1895636 | 709.8366852 | 982.8339922 | Cell division |
| 530.8685424 | 555.569729 | 532.4045404 | 560.0027735 | 399.8011889 | 378.1793225 | Cell division |
| 474.42499 | 521.0328901 | 439.0400516 | 512.5890307 | 438.0345413 | 469.813851 | UDP-N-acet |
| 360.8377531 | 385.6568915 | 302.4513781 | 356.607347 | 347.1235083 | 354.3016177 | UDP-N-acet |
| 314.3002363 | 343.1395723 | 282.6304243 | 324.9197633 | 303.3886582 | 404.58181 | FIG011144 |
| 1003.054343 | 1294.497172 | 775.6851235 | 802.9324736 | 1385.108865 | 1210.367011 | GTP-bindin |
| 155.9390956 | 128.0275304 | 207.9318648 | 190.6441969 | 142.2693952 | 337.174207 | Rhodanese- |
| 56.26503141 | 49.21688521 | 70.45549622 | 53.50674449 | 45.98438101 | 59.23473604 | hypothetica |
| 358.3004538 | 350.1246017 | 415.853008 | 433.9128582 | 487.0268368 | 691.1306851 | Glucokinas |

| | | | | | | |
|---|---|---|---|---|---|---|
| 73.83099873 | 48.99792725 | 92.50092611 | 89.70248341 | 59.19007403 | 58.51919038 | FIG011145 |
| 7295.358073 | 6308.493639 | 7636.506672 | 8925.678721 | 9350.811148 | 10296.77128 | Non-specifi |
| 23.26379076 | 26.99596912 | 26.85488793 | 31.81482105 | 24.51874044 | 23.6554303 | Late compe |
| 332.3688862 | 348.3783444 | 334.9324131 | 391.691865 | 404.5104198 | 387.7142656 | Ribosomal |
| 319.276117 | 339.0017015 | 399.7809133 | 339.1857667 | 285.6857359 | 391.6585213 | Hypothetic |
| 202.7208243 | 199.5972168 | 216.9265308 | 201.9518073 | 279.2905298 | 316.5356207 | FIG011149 |
| 135.4451826 | 126.4054641 | 153.4173768 | 158.9025808 | 126.7764058 | 127.5513096 | LacI family |
| 284.4811061 | 302.3801084 | 325.5506138 | 305.0996199 | 321.5740895 | 358.153709 | Lon-like pro |
| 381.1638212 | 437.0860889 | 430.0569949 | 336.0837277 | 459.717791 | 364.6960926 | Phosphopa |
| 78.67952178 | 69.74115352 | 86.71721436 | 67.05116846 | 67.42479066 | 79.06600822 | V-type ATP |
| 389.5389983 | 455.2349423 | 450.2686457 | 460.9517223 | 562.3867856 | 671.8896998 | 16S rRNA (g |
| 57.89562816 | 55.79054875 | 61.54116925 | 49.52952778 | 68.80720713 | 75.41116212 | Aspartate--a |
| 1661.695242 | 1732.116537 | 1795.082115 | 800.1970362 | 1976.198862 | 2551.938672 | Carbamate |
| 949.3264389 | 905.3518506 | 839.9101185 | 744.4216482 | 2099.620584 | 2571.348349 | Acetylornit |
| 1121.318641 | 1177.177337 | 1105.764701 | 573.2165786 | 1230.225887 | 1666.760715 | Arginine/or |
| 4875.486429 | 4862.61896 | 4766.838956 | 1763.911343 | 3568.12015 | 3240.0975 | Ornithine c |
| 1842.992362 | 2079.792522 | 1604.489523 | 827.5067091 | 1732.250577 | 1648.347565 | Acetyltrans |
| 2590.133711 | 2645.643486 | 2726.790059 | 855.1929667 | 1577.51709 | 1617.068243 | Arginine de |
| 40.88463761 | 39.25617328 | 33.81983482 | 43.09079887 | 66.85864514 | 53.29821514 | Transcriptio |
| 151.559477 | 147.415068 | 137.6015075 | 158.48223 | 171.2363908 | 153.9112321 | Arginine pa |
| 104.9574183 | 102.3891368 | 114.8804418 | 74.86164653 | 81.77968807 | 83.41313976 | V-type ATP |
| 293.8895318 | 294.5987321 | 262.5077242 | 278.1366459 | 381.8318514 | 559.1141242 | Putative un |
| 203.3367296 | 215.3135331 | 184.6249966 | 261.8402621 | 188.7024474 | 142.0781578 | Hypothetic |
| 175.5284386 | 193.4085774 | 150.1221667 | 213.5999782 | 162.7090127 | 113.7003877 | Two-compo |
| 126.0593182 | 123.5035219 | 108.7195563 | 149.9523733 | 115.9744372 | 85.02788346 | Two-compo |
| 92.6647738 | 90.80538299 | 80.02991667 | 122.0182124 | 70.28787058 | 63.72377962 | Peptide me |
| 67.5380751 | 74.04802866 | 63.97068485 | 80.13960605 | 64.18735984 | 55.26812425 | Thiol:disulf |
| 46.97809186 | 44.16483817 | 42.05624709 | 55.80305781 | 34.62835576 | 32.47823796 | Cytochrom |
| 1061.422362 | 1123.489208 | 907.2335864 | 1525.78153 | 1065.274646 | 769.1978754 | Magnesium |
| 627.3160273 | 629.7331482 | 543.2739488 | 758.6358564 | 535.2659451 | 588.9195467 | Oxidoreduc |
| 95.20812751 | 96.11131883 | 84.17766724 | 94.54712204 | 115.7637913 | 94.14905642 | FIG011148 |
| 81.91582351 | 81.13830274 | 99.08527617 | 61.17560632 | 81.07154405 | 74.62038176 | V-type ATP |
| 920.0853663 | 939.864192 | 787.7693355 | 1003.984213 | 789.7315827 | 701.2957663 | Valyl-tRNA |
| 552.2929372 | 467.3096172 | 521.5124499 | 607.5994934 | 466.6155956 | 360.472176 | hypothetica |
| 223.9594042 | 194.8420237 | 304.991548 | 239.3033218 | 112.0132358 | 334.4430083 | hypothetica |

| | | | | | | |
|---|---|---|---|---|---|---|
| 227.6049974 | 246.6985405 | 283.1017077 | 266.4381708 | 203.1798816 | 249.6962087 | FIG011141 |
| 101.919385 | 109.8293162 | 144.9655539 | 120.2498156 | 89.16046274 | 131.4612923 | Aldose 1-ep |
| 318.4549303 | 323.1550134 | 260.3670975 | 307.38579 | 216.2062539 | 150.8513525 | 2-keto-3-de |
| 671.5038826 | 727.7795849 | 516.1978339 | 762.6374003 | 525.9440125 | 376.631636 | 3-dehydroc |
| 441.8991007 | 478.7313206 | 386.9226772 | 474.2828002 | 639.2527996 | 664.0056283 | Uncharacte |
| 599.7251155 | 651.6012549 | 566.3483069 | 616.3151094 | 590.2296716 | 525.3806566 | hypothetica |
| 289.9154149 | 293.6207027 | 235.5253536 | 242.7085212 | 314.6079615 | 310.554222 | SAM-depen |
| 311.5598773 | 336.9115308 | 303.3358831 | 396.3307345 | 343.5212074 | 267.3779002 | putative to: |
| 110.7548445 | 115.3781445 | 105.4123619 | 134.662671 | 138.9776829 | 134.5777968 | Shikimate/( |
| 67.11845772 | 65.05920219 | 69.11550254 | 101.4304268 | 126.7145932 | 105.1280978 | Beta-galact |
| 54.03968943 | 50.51446305 | 49.88389752 | 65.49938365 | 79.96303394 | 57.28802448 | DNA-bindir |
| 55.6931395 | 50.92086477 | 52.03151313 | 68.90612388 | 81.02775912 | 65.36056433 | Two-compo |
| 38.10423909 | 36.10084533 | 42.87590528 | 51.77902694 | 57.24899409 | 39.69753659 | Integral me |
| 111.6186852 | 105.1558507 | 103.1278545 | 159.6521565 | 220.6967294 | 158.3199304 | Multiple su |
| 56.17466207 | 54.98167607 | 49.95096065 | 80.62125879 | 91.30661101 | 75.12172228 | Multiple su |
| 68.74602085 | 59.98972463 | 68.12581351 | 82.40041693 | 92.71851622 | 83.36975439 | Multiple su |
| 122.1059999 | 117.3091595 | 125.3939219 | 166.9842128 | 120.2150256 | 118.3214574 | N-acetylma |
| 92.32141197 | 81.26485495 | 106.2111947 | 151.4554785 | 90.98581279 | 96.56603528 | FIG011142 |
| 522.2021233 | 528.8844337 | 775.6764159 | 644.8613351 | 555.6034504 | 557.067161 | Hypoxanthi |
| 198.645219 | 217.56102 | 187.8707004 | 225.7506523 | 210.1417616 | 169.6969772 | FIG013576 |
| 407.9572201 | 451.100754 | 362.0667413 | 659.0155665 | 674.7326283 | 735.5016376 | Beta-glucos |
| 246.5334139 | 262.0313264 | 206.0719974 | 303.0675246 | 291.9699559 | 263.8997354 | Hyaluronog |
| 229.1654491 | 248.3359141 | 176.0641288 | 268.1657958 | 223.2706008 | 200.5695722 | Transcriptio |
| 371.3152044 | 377.7044777 | 322.1913795 | 358.026292 | 248.1911247 | 197.4614484 | hypothetica |
| 507.1181097 | 537.8569484 | 517.9494406 | 688.8401488 | 541.9315266 | 491.8985604 | Alpha-manr |
| 166.9601645 | 142.3161119 | 170.5967273 | 177.6647448 | 170.4055786 | 133.5238754 | Putative tw |
| 61.73847006 | 64.98627243 | 59.30386127 | 58.8838673 | 78.40480992 | 88.61574548 | RNA methyl |
| 14.27977118 | 13.65516005 | 19.30116003 | 12.61213472 | 15.77139758 | 5.930409756 | Paratox |
| 121.8958608 | 122.5519495 | 155.1826734 | 186.1262718 | 312.7433301 | 337.0659318 | Streptococ |
| 59.6366218 | 52.89353343 | 77.52738981 | 79.08160534 | 206.8994265 | 279.0087376 | FIG011175 |
| 1187.1559 | 1263.868962 | 1136.993714 | 1129.700162 | 1083.821135 | 1035.044551 | Adenylosuc |
| 329.6843855 | 300.9484233 | 390.0944033 | 372.5006147 | 483.0997033 | 578.7940838 | Phage prote |
| 288.6961739 | 287.1146467 | 292.8911375 | 280.8353476 | 459.9914325 | 613.949906 | Phage prote |
| 98.96507902 | 65.75831839 | 101.8654796 | 90.85906153 | 51.23342266 | 48.74393591 | Phage lysin |
| 23.13117958 | 16.85368128 | 9.125260052 | 8.435748006 | 13.30446979 | 16.13088794 | Phage holir |

| | | | | | | |
|---|---|---|---|---|---|---|
| 22.06750679 | 17.05704469 | 17.04561968 | 14.84508884 | 12.1896932 | 15.04931259 | Phage prote |
| 17.59729377 | 12.44208373 | 11.08825204 | 7.632343434 | 8.18074194 | 10.3278818 | FIG011165 |
| 19.43815535 | 13.39260021 | 10.71805603 | 11.21190037 | 12.14235401 | 9.968871089 | Phage hyalu |
| 17.10848006 | 13.67901603 | 12.69362186 | 10.17645791 | 11.08945019 | 10.79310171 | Hyaluronat |
| 28.3304124 | 23.91517779 | 14.32379412 | 13.60371814 | 17.38207434 | 19.36668156 | Phage endo |
| 22.19909654 | 14.59841931 | 11.94443887 | 9.350419068 | 13.47255242 | 13.28621321 | Phage-relat |
| 1324.757649 | 1304.499414 | 1276.643639 | 1352.240709 | 1001.604908 | 991.5921561 | Nucleoside- |
| 19.50919846 | 14.8343519 | 9.674052461 | 9.571674856 | 13.09520789 | 14.92059087 | Phage tail le |
| 12.73536106 | 7.453098445 | 7.232824199 | 5.476403124 | 8.602429669 | 8.598414832 | Phage prote |
| 9.939436966 | 6.828468456 | 6.451346641 | 5.684779937 | 5.687026121 | 4.712517177 | Phage prote |
| 8.124464853 | 4.082160074 | 5.027023357 | 3.211531523 | 3.952254548 | 5.089626554 | Phage majo |
| 17.7428353 | 16.77982641 | 13.94558015 | 10.60494035 | 20.07664789 | 23.87522524 | Phage minc |
| 11.71181233 | 7.792123201 | 6.234494654 | 5.541466157 | 7.25954915 | 11.99094522 | Phage minc |
| 25.25449458 | 20.67191151 | 10.4616432 | 10.23146579 | 19.45689975 | 25.71031497 | Phage minc |
| 21.99028726 | 14.359512 | 10.55885871 | 9.092921386 | 13.70127139 | 12.14525163 | Phage prote |
| 21.98694838 | 19.11288281 | 13.11537504 | 12.54723712 | 15.31866403 | 19.14047593 | Phage majo |
| 23.40526758 | 18.81312738 | 15.01606546 | 11.45979448 | 14.41100156 | 19.60248742 | Phage prote |
| 1424.500931 | 1254.423978 | 1347.883438 | 1936.681705 | 1292.44815 | 904.4991405 | Transcriptio |
| 15.30371391 | 9.005978996 | 19.93393625 | 11.31988015 | 6.838732588 | 15.12957845 | Phage prote |
| 130.8682953 | 123.6625939 | 166.3768344 | 148.7101863 | 249.2973465 | 321.0311141 | hypothetica |
| 29.25815109 | 23.7007099 | 30.37832906 | 31.19145654 | 33.90572563 | 44.02885984 | Phage minc |
| 12.48670247 | 9.22274848 | 7.307663112 | 7.348986971 | 9.013512077 | 8.424101682 | Phage minc |
| 10.74131843 | 8.402782965 | 8.543243696 | 7.798592499 | 5.637422168 | 9.551286999 | Phage term |
| 12.25170076 | 9.390251869 | 6.491921149 | 7.488336954 | 7.532456729 | 9.85915081 | Phage prote |
| 10.59478446 | 9.646566548 | 12.34654271 | 8.211348798 | 7.447296111 | 12.61291362 | FIG011147 |
| 18.85424333 | 16.4647123 | 28.03263719 | 11.77561558 | 7.947006456 | 22.96857112 | hypothetica |
| 467.6636549 | 461.5798644 | 524.8141947 | 441.1224333 | 520.6861255 | 504.3494235 | Nicotine ad |
| 18.51015339 | 14.43936559 | 20.86607429 | 14.8830697 | 13.00737968 | 11.43045297 | conserved h |
| 22.59292362 | 17.42946925 | 20.56175648 | 20.05389194 | 7.164934173 | 5.429182391 | Phage prote |
| 28.89660872 | 19.85218899 | 22.91925781 | 23.4594135 | 13.77809169 | 9.872456006 | Phage prote |
| 26.66324133 | 20.29195267 | 23.8863151 | 20.86817951 | 12.42644345 | 13.48313273 | Phage prote |
| 23.39640195 | 15.71631629 | 16.49492039 | 25.81119779 | 10.7093349 | 6.394659004 | FIG011151 |
| 23.55671343 | 20.86704179 | 27.37756348 | 23.70516078 | 15.83901633 | 11.31539901 | FIG011146 |
| 25.17766648 | 18.53719357 | 22.74513239 | 17.53481979 | 20.47283111 | 8.760448599 | Phage Holli |
| 26.3215859 | 17.59784711 | 20.78514851 | 21.41356073 | 8.849027901 | 11.88866181 | Single-strar |

| | | | | | | |
|---|---|---|---|---|---|---|
| 27.17022159 | 25.98430959 | 30.24964759 | 26.37737891 | 15.18345704 | 16.167322 | Single-strar |
| 12.00113575 | 7.809349708 | 14.82607613 | 10.80853604 | 5.731600564 | 5.492484398 | hypothetica |
| 423.3553979 | 381.0216175 | 469.2134889 | 425.3075275 | 450.257037 | 370.5370958 | Streptococ |
| 12.48372723 | 14.42203142 | 12.14371133 | 13.14558916 | 8.315483572 | 8.748323411 | FIG011168 |
| 22.04197107 | 14.90619689 | 16.95972571 | 20.77256357 | 11.94219648 | 9.116391629 | hypothetica |
| 21.00198757 | 20.49954298 | 24.19254991 | 31.85673781 | 7.682783735 | 8.73804336 | hypothetica |
| 24.21144902 | 17.88408405 | 24.59421421 | 29.75736658 | 17.45224947 | 11.70429103 | DNA replica |
| 13.77992808 | 8.662536536 | 11.49304864 | 12.47617714 | 4.947094338 | 4.114448763 | putative ph |
| 18.24762854 | 17.42461154 | 20.33576659 | 22.69792569 | 11.5241756 | 6.990434688 | hypothetica |
| 17.15736143 | 13.47112825 | 17.85640588 | 15.91888773 | 9.816890328 | 7.416934423 | phage prote |
| 30.65335045 | 13.68840451 | 27.0540343 | 16.84039648 | 9.183542565 | 9.0762902 | conserved h |
| 7707.944622 | 6362.947103 | 10068.33446 | 7381.773599 | 10726.17979 | 9162.701345 | hypothetica |
| 1792.7463 | 1281.665593 | 2536.051245 | 2014.938666 | 932.9318108 | 636.5883039 | Phage antir |
| 620.1439346 | 623.842092 | 702.3553039 | 613.7200539 | 711.8013766 | 734.3102118 | Thiol-activa |
| 658.2627118 | 433.6375156 | 954.981031 | 582.8100445 | 251.0911667 | 119.1696815 | FIG011170 |
| 441.0900611 | 377.1915909 | 840.7972169 | 650.1149796 | 751.5949016 | 1105.009897 | putative cI- |
| 590.5358502 | 433.6539124 | 878.2492963 | 656.04232 | 567.5616964 | 763.1467258 | Phage prote |
| 628.6911639 | 510.6476522 | 1094.176972 | 893.892334 | 706.4433103 | 818.5279161 | Chromosom |
| 397.9964406 | 314.5959684 | 579.0795156 | 474.7317616 | 399.1720846 | 490.3198831 | Phage integ |
| 840.002402 | 703.6540682 | 989.1857278 | 1000.891962 | 803.3930787 | 429.263971 | Regulatory |
| 633.7695011 | 580.7090574 | 681.5594022 | 738.8794497 | 675.6005984 | 753.4304252 | Cysteinyl-tF |
| 1203.222174 | 1159.486263 | 1158.8628 | 898.3643583 | 1762.856011 | 2579.947889 | Ribosomal s |
| 14.41075679 | 16.21187581 | 18.889078 | 13.20189134 | 15.38863889 | 10.59192553 | Competenc |
| 26.80203206 | 29.4058678 | 25.90755721 | 26.56469941 | 20.43334638 | 15.09585952 | ComF operc |
| 54.36136969 | 33.34407645 | 84.78290012 | 73.51803243 | 47.75784484 | 55.13233033 | FIG011147 |
| 146.5912951 | 159.8446197 | 120.0837153 | 118.5870839 | 97.14534125 | 82.390224 | FIG000605 |
| 1982.87315 | 2296.534204 | 1851.269392 | 1766.892366 | 1094.48948 | 853.3116763 | Cysteine syr |
| 543.8820726 | 629.6505128 | 484.3630097 | 457.940606 | 841.7593897 | 690.8116771 | General stre |
| 521.9613568 | 616.369021 | 448.2787977 | 456.7246609 | 731.9153906 | 835.7030989 | Streptococ |
| 246.6663651 | 279.2950974 | 276.1417534 | 291.969235 | 346.6188005 | 425.8979498 | Two compc |
| 131.0026595 | 133.7998031 | 138.4632309 | 141.2917643 | 153.7295721 | 134.5632594 | Sensor histi |
| 97.08221293 | 94.50200572 | 112.8287465 | 94.36351882 | 98.42388744 | 122.3250416 | Transporter |
| 1416.173491 | 1486.594688 | 1205.37246 | 1550.776435 | 1526.348098 | 1480.611062 | Serine/thre |
| 1186.406696 | 1289.310595 | 1087.522048 | 1205.4035 | 1316.337683 | 1457.109413 | Protein seri |
| 680.1892736 | 641.9455902 | 541.671741 | 629.4866108 | 685.6686074 | 711.0686808 | 16S rRNA (c |

| | | | | | | |
|---|---|---|---|---|---|---|
| 1555.475075 | 1106.316407 | 1560.131462 | 840.9752129 | 740.4740133 | 1269.372438 | FIG011149 |
| 365.6345396 | 333.5687386 | 300.2357475 | 356.547859 | 532.8180154 | 596.0325801 | Methionyl- |
| 201.7048296 | 157.6178487 | 226.8723569 | 232.5992663 | 144.6478658 | 135.4495642 | Helicase Pri |
| 605.5982957 | 549.9213908 | 724.8548762 | 680.5433539 | 516.6489138 | 454.0120891 | DNA-direct |
| 707.0127804 | 773.2872414 | 830.9912661 | 831.9830369 | 826.1931945 | 776.5029258 | Guanylate k |
| 986.0361222 | 1082.666535 | 944.6528753 | 1129.609611 | 1010.846662 | 1134.233893 | Hydrolase ( |
| 87.66277283 | 78.89695902 | 78.10660482 | 105.8011273 | 59.95197906 | 39.45562656 | Transcriptic |
| 175.2530485 | 172.6810408 | 142.3450537 | 172.5372182 | 114.6791279 | 67.34692459 | 3-ketoacyl- |
| 204.4286538 | 184.902099 | 189.9728343 | 229.7722267 | 117.3502927 | 87.89183061 | Butyrate-ac |
| 183.9317138 | 163.592565 | 159.0843433 | 191.6689591 | 106.11166 | 91.6263283 | Butyrate-ac |
| 109.3578691 | 84.18242849 | 109.7286281 | 161.4562085 | 86.59218164 | 81.04110581 | Cystathioni |
| 162.5490562 | 144.6679835 | 179.918468 | 180.2883287 | 113.2480006 | 104.832904 | D-beta-hydr |
| 315.3741084 | 336.7782061 | 277.0873418 | 298.770691 | 280.9703766 | 235.6775123 | S-ribosylho |
| 1369.367532 | 1440.320202 | 1380.324861 | 1741.781768 | 1750.692131 | 1698.402788 | Hypothetic |
| 1009.94297 | 988.0460244 | 1119.392426 | 1214.529855 | 738.6891241 | 639.2048939 | FIG001721 |
| 126.1193202 | 110.3025868 | 173.4169327 | 104.1919911 | 81.77739833 | 167.2333234 | Cell divisior |
| 20.25734981 | 23.2089787 | 21.754541 | 15.33568541 | 29.3365211 | 34.93693267 | FIG005686 |
| 441.1112764 | 432.6245094 | 380.4864417 | 430.509287 | 524.1155536 | 345.4195889 | RecU Hollic |
| 910.6053405 | 1014.255181 | 824.9270212 | 1051.3811 | 1197.281485 | 1035.58951 | Multimodu |
| 1293.140684 | 1682.56204 | 1260.254655 | 1239.848521 | 1590.468299 | 1514.251975 | Aminopept |
| 1472.36557 | 1784.278151 | 1546.68592 | 1726.436085 | 2190.118484 | 2254.14775 | Cell divisior |
| 908.5900553 | 942.9224437 | 828.058189 | 1155.43796 | 888.9818047 | 677.2182672 | Leucyl-tRN |
| 240.5006143 | 273.2353024 | 222.7474049 | 229.5364841 | 250.2414589 | 261.7790692 | NAD synthe |
| 223.7747035 | 257.4958679 | 237.222772 | 220.7518207 | 230.0014756 | 219.4217783 | Nicotinate |
| 295.6804414 | 331.0334533 | 241.1892041 | 219.8911327 | 203.5084047 | 216.2228829 | Histidine tr |
| 319.804936 | 363.0160899 | 309.9703399 | 277.0989628 | 336.8540799 | 330.2295053 | Thioredoxir |
| 66.51777047 | 52.59727184 | 74.40553127 | 77.66672678 | 57.06885577 | 76.90643229 | Hypothetic |
| 498.1169447 | 535.1785929 | 601.9522632 | 525.6714989 | 544.5207393 | 472.3722819 | ABC-type m |
| 520.2758095 | 536.2899718 | 567.9832985 | 474.3444486 | 512.3098064 | 352.9810866 | Cysteine AB |
| 107.3786866 | 107.4996034 | 79.35156369 | 87.19189139 | 100.9176326 | 137.4668981 | FIG011160 |
| 395.6209446 | 462.6841418 | 319.8312699 | 273.2106366 | 382.0260401 | 419.4904463 | ATP-depenc |
| 540.4980063 | 600.7177154 | 516.3948832 | 629.2131104 | 601.8307245 | 709.5550497 | Phospho-N- |
| 47.47956021 | 38.04137645 | 40.32091651 | 42.26849276 | 37.29596827 | 36.07459156 | Ascorbate-s |
| 538.4771895 | 581.3574071 | 562.455338 | 598.830418 | 423.0139816 | 385.4604236 | Cell divisior |
| 394.0327364 | 433.9449553 | 464.5865602 | 473.8837234 | 389.5851106 | 309.8364541 | Cell divisior |

| | | | | | | |
|---|---|---|---|---|---|---|
| 442.6552392 | 466.4677071 | 456.9862928 | 526.5504057 | 471.6173525 | 385.3606383 | rRNA small |
| 104.4723546 | 90.71347471 | 133.2712398 | 138.5069482 | 56.14572363 | 77.21670948 | FIG011152 |
| 333.7662252 | 322.8440415 | 352.8515312 | 432.6824911 | 258.2054032 | 234.8081983 | Gamma-glu |
| 282.4075585 | 267.5803593 | 291.1967729 | 324.819117 | 226.903216 | 244.9980919 | Glutamate ! |
| 743.0238545 | 771.6624466 | 742.7662314 | 1061.646321 | 700.7777217 | 505.8425944 | ABC transpo |
| 436.0824874 | 432.7439578 | 468.5784224 | 554.3885051 | 442.922064 | 382.1806173 | Methionine |
| 255.0025818 | 218.2625498 | 264.3964817 | 240.5989072 | 142.0691544 | 180.2000537 | FIG011139 |
| 788.1700288 | 826.1511941 | 724.0467177 | 897.8402891 | 1375.550977 | 1270.999885 | Transketola |
| 41.04143032 | 23.99695605 | 29.48196046 | 25.4083175 | 21.95605579 | 26.58056416 | Ascorbate-s |
| 108.579395 | 88.81546181 | 104.5718281 | 127.4565854 | 278.9823251 | 322.2143328 | Transaldola |
| 85.3644721 | 80.75043308 | 87.15769313 | 114.4851515 | 238.3148296 | 224.9714119 | Trans-acting |
| 87.01842175 | 78.19654919 | 87.25123047 | 118.5138049 | 219.8196333 | 194.9323192 | NADH pero: |
| 259.1539918 | 204.1777842 | 233.7234665 | 356.1760269 | 540.4323983 | 752.6255988 | Glycerol up |
| 184.495999 | 149.7023993 | 190.6199283 | 235.4397413 | 253.5736402 | 331.8218511 | Alpha-glyce |
| 152.369382 | 124.0992959 | 156.6157755 | 190.70804 | 206.9188859 | 256.2130354 | Glycerol kir |
| 77.20812643 | 61.55557212 | 83.66590176 | 74.79696565 | 124.3472775 | 121.0036963 | FIG011165 |
| 2506.466713 | 2198.234326 | 2569.87991 | 2334.143179 | 4002.559597 | 3932.643795 | FIG011163 |
| 243.4565132 | 232.4552828 | 232.3610026 | 156.5517886 | 101.8216997 | 106.3297951 | FIG011141 |
| 309.5312218 | 372.4535838 | 238.4626439 | 235.0043439 | 211.3230009 | 205.9698171 | Glycyl-tRN/ |
| 35.09682481 | 28.61921793 | 29.36955647 | 30.868589 | 29.08708245 | 29.89813231 | Ascorbate-s |
| 337.6092567 | 398.7116449 | 248.6720053 | 210.3533712 | 185.558476 | 154.0808312 | Glycyl-tRN/ |
| 894.377513 | 801.9736002 | 913.7897595 | 1100.703982 | 1094.748725 | 1000.584509 | INTEGRAL M |
| 1279.844891 | 1138.254207 | 1444.410432 | 1831.893264 | 2238.882081 | 2596.740518 | oxidoreduc |
| 1201.173814 | 1167.890543 | 1369.732064 | 1577.99398 | 1545.737306 | 1486.795189 | N-acetylglu |
| 164.7514282 | 164.3568444 | 146.5487429 | 169.6063847 | 160.7515791 | 160.115007 | Sodium-dep |
| 409.213749 | 435.6875897 | 462.7034303 | 515.0216674 | 393.3273569 | 328.6483254 | FIG011161 |
| 312.6932643 | 368.4105885 | 204.7326821 | 132.3953553 | 207.3774412 | 329.7134297 | DegV family |
| 230.4961657 | 214.1136853 | 261.6957549 | 224.9325471 | 198.5545237 | 167.8033176 | Transcriptic |
| 373.5937463 | 382.3351126 | 354.5894709 | 393.7956164 | 337.7724228 | 271.7912781 | Hydrolase (I |
| 339.7388207 | 340.2658769 | 321.4712295 | 375.7780585 | 268.1059465 | 231.8041716 | INTEGRAL M |
| 35.75993047 | 27.09464482 | 38.31399306 | 39.86777041 | 39.17872067 | 55.29085898 | 3-keto-L-gu |
| 113.2385854 | 101.763148 | 84.67392467 | 98.9151709 | 55.46543035 | 25.62431215 | hypothetica |
| 452.0021446 | 444.7331833 | 448.7248009 | 529.525432 | 439.2907856 | 439.6173943 | Tagatose 1,( |
| 458.7479123 | 460.071652 | 437.5336376 | 555.7911629 | 506.4508549 | 364.5033711 | Tagatose-6- |
| 395.0183213 | 384.7756272 | 394.7699035 | 500.1137409 | 514.9301427 | 432.2130901 | Galactose-6 |

| | | | | | | |
|---|---|---|---|---|---|---|
| 484.495632 | 490.9688665 | 469.4263322 | 598.1607296 | 631.5993664 | 564.9944994 | Galactose-6 |
| 389.5122722 | 404.8497044 | 392.1897842 | 493.4381139 | 470.9261882 | 570.7606703 | PTS system, |
| 363.3323597 | 375.496694 | 366.5250371 | 421.6645266 | 436.9478636 | 360.3757549 | PTS system, |
| 251.0955621 | 258.6229262 | 226.8179151 | 325.0798631 | 312.6493173 | 241.6787942 | PTS system, |
| 217.2272938 | 222.4745473 | 195.5486005 | 269.7038526 | 329.7252814 | 318.5883809 | Lactose pho |
| 23.98925195 | 22.64998524 | 40.79528023 | 26.66830588 | 19.34504859 | 26.00852906 | Copper cha |
| 41.95854734 | 33.77916591 | 36.06481977 | 33.58231111 | 43.01706803 | 44.03306364 | L-xylulose 5 |
| 141.1757084 | 146.6011224 | 153.9131759 | 150.5048884 | 146.3825017 | 143.5458515 | Copper-trar |
| 70.01870575 | 70.30983602 | 84.35560164 | 68.6910909 | 88.61035217 | 90.43874877 | Negative tra |
| 494.8155758 | 529.9599176 | 516.8920896 | 461.1141762 | 435.5238031 | 358.028711 | putative est |
| 1211.370631 | 997.9189205 | 1130.502004 | 1449.284167 | 1017.432582 | 1756.04051 | Ribosome-b |
| 1995.981347 | 1978.954396 | 2172.443645 | 2550.546767 | 1636.457095 | 1419.254252 | Translation |
| 886.9036062 | 869.6880369 | 1056.647543 | 1033.041462 | 739.8630611 | 722.7140496 | ribosomal p |
| 964.4916742 | 929.643921 | 994.4196944 | 1002.103565 | 944.2063606 | 807.3510749 | COG2740: I |
| 1280.668128 | 1239.59391 | 1452.305094 | 1325.892283 | 1255.849856 | 1238.235074 | Transcriptio |
| 576.7100598 | 513.2359537 | 678.0214117 | 513.8952238 | 449.1227325 | 528.3444264 | FIG000325 |
| 49.64950743 | 45.36111394 | 48.29976454 | 53.49909302 | 45.50050755 | 61.35585896 | Mobile elen |
| 62.94589424 | 53.43547537 | 66.0919874 | 66.6449733 | 96.49794322 | 112.8880836 | L-ribulose-5 |
| 567.985859 | 612.3432666 | 627.7129851 | 560.502021 | 794.89028 | 825.1350832 | tRNA (guani |
| 503.9824943 | 538.4624273 | 507.1271636 | 494.2289298 | 591.5420126 | 577.4224827 | aminoglyco |
| 193.2182856 | 198.0255852 | 169.5582193 | 143.3022256 | 156.6718787 | 180.2949613 | ABC transpo |
| 264.5920366 | 301.0129173 | 226.5968862 | 206.111119 | 246.6392281 | 215.0930756 | ABC transpo |
| 466.5293969 | 513.1377268 | 414.6142243 | 404.0344547 | 489.4421035 | 419.6070836 | Histidine tr |
| 500.679977 | 513.2235722 | 454.9825772 | 488.8804587 | 544.7961657 | 499.0597621 | Hypothetic |
| 863.8133254 | 847.2122649 | 681.6872707 | 980.2068878 | 747.0928779 | 586.3492502 | Cell envelop |
| 515.5805964 | 518.5486299 | 469.8423597 | 619.5543879 | 458.9816952 | 462.816708 | Acetyltrans |
| 275.8204397 | 253.7062486 | 293.7666774 | 279.0464056 | 116.7827473 | 123.7170762 | ATPase YjeE |
| 563.9733917 | 528.9331087 | 469.9746416 | 632.6914935 | 265.4390387 | 158.9642073 | Guanine-hy |
| 48.86855025 | 44.45799489 | 48.17945579 | 45.41035809 | 43.59486087 | 45.13700758 | Transcriptio |
| 314.2967577 | 308.7589222 | 354.7050367 | 384.8052945 | 143.0875158 | 109.6092421 | Hydrolase (I |
| 451.2988139 | 506.9580332 | 336.4055681 | 255.505955 | 384.7865109 | 303.223304 | PTS system, |
| 891.1856144 | 1038.091795 | 554.0989949 | 469.2703958 | 724.8500947 | 641.8545931 | PTS system, |
| 1102.794616 | 1271.574064 | 696.2785635 | 582.9704425 | 800.5286553 | 705.3296862 | PTS system, |
| 71.13706008 | 67.71112934 | 81.85146051 | 72.65990949 | 115.3484614 | 141.3524148 | Putative reg |
| 50.57127044 | 44.07220969 | 47.7877529 | 58.10272874 | 43.37091759 | 50.13741334 | Mobile elen |

| | | | | | | |
|---|---|---|---|---|---|---|
| 261.0223935 | 261.7910337 | 208.0786452 | 170.4055044 | 155.1345204 | 142.2393467 | Seryl-tRNA s |
| 1364.064155 | 1754.356738 | 940.5285225 | 774.2581795 | 1207.744896 | 1497.850233 | Acetyl-coen |
| 1419.133452 | 1802.590782 | 992.0340649 | 740.8813486 | 849.2119661 | 848.4065629 | Acetyl-coen |
| 1400.910889 | 1826.477742 | 1025.941351 | 719.8222449 | 937.1570292 | 1115.654172 | Biotin carbo |
| 258.7962446 | 195.6335964 | 107.5372136 | 71.15692494 | 87.16535588 | 133.9465229 | Probable L- |
| 1090.657803 | 1421.602113 | 805.9607879 | 546.8005704 | 596.8112333 | 794.3501133 | (3R)-hydrox |
| 1587.834375 | 1987.125859 | 891.7924384 | 594.6098264 | 699.9971858 | 923.522233 | Biotin carbo |
| 1546.301178 | 2055.089991 | 875.8370176 | 584.2029648 | 860.8290205 | 1024.483496 | 3-oxoacyl-[a |
| 911.1447762 | 1066.656764 | 570.8783477 | 421.529655 | 526.7462867 | 638.9364301 | 3-oxoacyl-[a |
| 915.6439787 | 1277.741535 | 609.3224763 | 489.7769881 | 743.8254665 | 794.9106921 | Malonyl Co |
| 818.7140928 | 1117.750235 | 635.4277768 | 447.4249328 | 658.0952405 | 897.8123798 | Enoyl-[acyl- |
| 822.9198461 | 1079.815704 | 817.8157093 | 519.2435885 | 815.7181403 | 1068.919019 | Acyl carrier |
| 1234.599058 | 1575.838766 | 1120.84704 | 902.0725415 | 1645.069172 | 1869.005636 | 3-oxoacyl-[a |
| 1268.315847 | 1492.18294 | 889.5072257 | 719.1878262 | 1422.298152 | 1545.705634 | Transcriptio |
| 1174.619359 | 1522.399077 | 974.2388721 | 752.6878324 | 1386.263907 | 1375.156193 | Trans-2,cis- |
| 3466.481639 | 3068.147931 | 3650.993304 | 4837.505518 | 2779.287131 | 1986.630293 | Glycine bet |
| 9522.304253 | 8237.588998 | 12777.15586 | 17632.23577 | 12481.59325 | 12081.93808 | Chaperone |
| 20996.34543 | 20398.82369 | 28648.90499 | 34149.34535 | 22687.62341 | 16238.87878 | Chaperone |
| 12889.99792 | 12130.87712 | 19666.27785 | 20362.40623 | 15519.9521 | 14447.16671 | Heat shock |
| 8390.787523 | 7880.434313 | 11917.64745 | 10632.6641 | 17317.22218 | 16281.88746 | Heat-induci |
| 439.4507342 | 427.2356506 | 470.0994434 | 642.4665681 | 564.419526 | 517.9866114 | putative N-a |
| 482.981853 | 431.1213698 | 494.1913622 | 647.9859575 | 471.0523988 | 378.8540775 | D-alanyl-D-a |
| 350.7080997 | 360.0101943 | 391.864636 | 495.0415568 | 397.8336402 | 456.7923243 | Phosphogly |
| 159.1675222 | 162.0745117 | 144.7520903 | 156.1577467 | 84.91610133 | 122.5947483 | Integral me |
| 107.5481117 | 103.4352909 | 78.76644155 | 112.8876843 | 67.12013387 | 58.89034804 | hypothetica |
| 918.0838111 | 1057.937282 | 750.8426669 | 950.3412426 | 851.1243914 | 765.1285625 | Aspartyl-tR |
| 192.8837854 | 161.8374121 | 215.855941 | 226.7595553 | 213.136537 | 220.9858698 | Cationic am |
| 4466.550605 | 4081.849306 | 4650.59771 | 7482.622297 | 3912.37166 | 2406.536026 | Glycine bet |
| 570.8972344 | 633.5700265 | 395.359367 | 489.9683539 | 512.1244831 | 439.432448 | Aspartyl-tR |
| 296.0526889 | 290.6740478 | 249.5904654 | 246.6531581 | 192.4499292 | 232.4146503 | Aspartyl-tR |
| 52.2891015 | 48.39627555 | 45.91037689 | 52.77220317 | 45.55244877 | 48.54237368 | Mobile elen |
| 220.4588964 | 165.5451982 | 231.6033444 | 350.2329755 | 229.5843418 | 238.7582968 | Pyruvate,pl |
| 78.67417125 | 69.15179166 | 87.4419011 | 92.87530561 | 77.79222281 | 69.96102608 | Nicotinami |
| 1093.338066 | 1285.486284 | 1085.667712 | 944.3752942 | 1215.488996 | 1352.07662 | GTP-sensing |
| 715.7405954 | 734.7862936 | 797.936004 | 817.9610726 | 722.717042 | 751.7457367 | Aspartate ar |

| | | | | | | |
|---|---|---|---|---|---|---|
| 403.0712581 | 332.2283549 | 349.5903569 | 321.46824 | 411.2691935 | 418.7200937 | Universal st |
| 344.4926804 | 336.3563242 | 432.8881464 | 426.5276608 | 435.9292984 | 416.7084177 | Hydrolase ( |
| 169.540635 | 153.4049133 | 199.3506183 | 199.6165946 | 165.789595 | 133.5048196 | L-asparagin |
| 1053.117137 | 887.8559155 | 1085.616649 | 1122.32354 | 877.9486821 | 790.7830057 | DNA polym |
| 127.9136052 | 96.30423597 | 146.0475069 | 115.9856377 | 96.41439522 | 75.00215856 | ABC transp |
| 161.58525 | 155.2834706 | 216.580923 | 177.8209237 | 207.7527953 | 189.4907117 | putative AB |
| 287.2208293 | 322.8861051 | 280.3503725 | 279.6436117 | 314.5495276 | 253.1028872 | ATP-depend |
| 223.0201541 | 237.6036634 | 199.57553 | 206.5361374 | 228.8074352 | 207.3716208 | Duplicated |
| 206.8651032 | 213.451115 | 218.6921251 | 188.2599883 | 170.7360485 | 147.0570355 | Transmemb |
| 210.5276237 | 224.7909481 | 200.3012802 | 190.9473557 | 200.3042269 | 211.9372699 | Substrate-s |
| 120.5872722 | 127.5081388 | 139.3931002 | 113.2601398 | 107.8271724 | 108.3172247 | Putative AB |
| 117.247728 | 118.8186425 | 140.0887968 | 118.6649584 | 115.5412013 | 116.2708754 | Putative AB |
| 107.0596647 | 99.14235364 | 132.1485522 | 118.769112 | 118.8582635 | 123.1061901 | Heme ABC t |
| 158.8865331 | 169.5149892 | 186.8241879 | 174.1517237 | 169.4493152 | 139.7984439 | Heme ABC t |
| 933.4785489 | 770.5130853 | 947.9659687 | 930.7040264 | 674.5237012 | 807.3377193 | Succinyl-Co |
| 161.9189635 | 180.4978969 | 216.4481471 | 197.6440609 | 182.2946617 | 158.3274216 | Heme trans |
| 119.5850877 | 138.5996878 | 162.2186438 | 137.7140228 | 122.0708971 | 97.64763455 | Cell surface |
| 228.8999707 | 255.2053867 | 314.0228369 | 259.1599298 | 225.480738 | 168.4346882 | Streptococ |
| 25.17766648 | 26.19386048 | 34.73802038 | 26.77191235 | 30.20581639 | 26.7966663 | hypothetica |
| 1700.806941 | 1486.050878 | 1627.169359 | 2032.848408 | 1379.809642 | 1192.465212 | Immunoger |
| 758.3474636 | 689.9762616 | 750.3865389 | 928.3092459 | 523.1038345 | 446.8544899 | Alanine rac |
| 444.6496021 | 380.6254079 | 472.7373337 | 497.0387283 | 300.7763205 | 364.6260665 | Holo-[acyl- |
| 1436.052297 | 1448.445645 | 1350.040287 | 1648.869669 | 1628.084514 | 1465.346986 | Protein exp |
| 51.57683009 | 56.33758881 | 51.54250491 | 57.71505416 | 51.16210039 | 52.6894768 | Mobile elem |
| 315.6058037 | 294.4585297 | 271.443453 | 345.9035647 | 292.4075006 | 264.8066788 | Mannose-6- |
| 1448.020391 | 1286.723023 | 1545.841907 | 1661.150193 | 1305.394698 | 1612.309033 | Peroxide st |
| 146.725639 | 108.2561176 | 152.7547671 | 233.0995302 | 115.2735868 | 112.8866341 | Fructokinas |
| 695.7667364 | 514.9868471 | 790.5210763 | 1251.691824 | 859.0432156 | 869.623133 | Secreted En |
| 1832.36585 | 1402.180709 | 2008.880516 | 3422.333073 | 1747.937966 | 1558.804322 | PTS system, |
| 242.2157505 | 208.961522 | 259.2298924 | 368.756673 | 267.7556837 | 281.4487358 | Sucrose-6-p |
| 337.4089806 | 287.6769199 | 334.0375058 | 505.7257127 | 347.1886554 | 277.5577059 | Sucrose ope |
| 1067.200117 | 1049.5585 | 1335.236496 | 1378.431551 | 2718.953558 | 2984.944546 | Transcriptio |
| 1749.198074 | 1835.061269 | 1948.678879 | 2104.272311 | 2931.927568 | 2459.00639 | Alkaline sho |
| 1116.633438 | 997.8170916 | 1396.404386 | 1119.344716 | 1196.60504 | 1218.816113 | Translation |
| 304.51317 | 287.5881772 | 360.8983852 | 369.4450707 | 504.1036929 | 656.0398125 | dCMP deam |

| | | | | | | |
|---|---|---|---|---|---|---|
| 262.5585047 | 226.7451777 | 259.0449943 | 290.5236604 | 319.6247868 | 271.8214907 | Aminopept |
| 945.2419763 | 875.9359571 | 1139.601802 | 1090.942051 | 1104.462294 | 1169.979763 | M protein t |
| 561.9260912 | 571.5923636 | 576.1363948 | 556.9480378 | 568.1023102 | 646.62071 | Excinucleas |
| 266.3224718 | 234.2978453 | 274.8478474 | 253.8299359 | 218.5270952 | 175.7733706 | Magnesium |
| 571.5986712 | 558.8262782 | 568.1014611 | 523.9342386 | 453.6389503 | 368.9130087 | Integral me |
| 1784.695538 | 1548.253608 | 1539.049383 | 1402.901047 | 1243.800005 | 1031.001736 | SSU ribosor |
| 2804.987167 | 2969.904622 | 2201.915416 | 2354.931643 | 3366.355355 | 2879.947202 | Single-strar |
| 3530.307396 | 3576.353087 | 2635.242086 | 2567.677699 | 2790.020954 | 2145.874275 | SSU ribosor |
| 619.4579433 | 645.5551048 | 765.6409505 | 680.5170213 | 826.8414797 | 1312.025454 | hypothetica |
| 274.6274803 | 282.8528716 | 283.2308743 | 238.5573193 | 229.3327574 | 245.6593084 | A/G-specific |
| 143.9925498 | 126.9333096 | 154.0423586 | 153.924118 | 162.0788628 | 143.4510312 | Transcriptic |
| 257.322713 | 276.0084498 | 286.3169081 | 199.749331 | 189.793213 | 299.7398531 | Thioredoxir |
| 665.8539153 | 591.7457964 | 802.1778383 | 751.7223683 | 1161.746244 | 1558.978861 | Co-activato |
| 197.0686157 | 205.6296472 | 193.2506681 | 200.1776302 | 160.8324068 | 132.7397736 | Phosphatid |
| 220.9977377 | 234.3196377 | 223.3665485 | 239.8134944 | 218.002668 | 172.2276158 | Recombina |
| 242.784641 | 253.5335419 | 239.7100641 | 240.9472112 | 207.341354 | 238.1884995 | Colicin V pr |
| 205.5808116 | 205.8379677 | 276.8441957 | 212.6978504 | 176.5134067 | 206.0481525 | FIG011142 |
| 432.2444901 | 454.050121 | 441.8422293 | 462.565654 | 348.0593142 | 248.8484435 | Ribonuclea |
| 467.0481743 | 499.5089818 | 469.5961286 | 490.1352183 | 461.9888085 | 360.0294522 | Putative sig |
| 306.3542561 | 288.6960495 | 306.8530189 | 378.0865131 | 208.7229298 | 157.8317094 | RecD-like D |
| 472.7660527 | 426.8785784 | 461.3113553 | 586.1639757 | 698.1577283 | 666.5878792 | Putative me |
| 219.0356851 | 248.6622645 | 203.5267303 | 209.8246218 | 255.1315662 | 215.676456 | DNA polym |
| 1776.80737 | 1904.327412 | 1859.958154 | 2712.637145 | 6127.839569 | 4881.024853 | Pyruvate fo |
| 750.5394924 | 693.2937716 | 995.7671765 | 1029.616065 | 2225.050882 | 3546.243568 | Co-activato |
| 125.4501811 | 139.7924975 | 136.6666854 | 148.4691649 | 152.4201393 | 244.8721677 | hypothetica |
| 263.0934199 | 286.6120645 | 197.3053807 | 234.488704 | 349.6918788 | 371.264853 | Beta-lactam |
| 188.4733261 | 209.696985 | 177.026494 | 169.602226 | 180.7715821 | 188.4040512 | C3-degradir |
| 315.7403796 | 323.1319214 | 274.8685457 | 355.634726 | 203.5090101 | 132.272906 | Integral me |
| 426.9986443 | 502.8665865 | 338.068801 | 286.6740557 | 289.3034111 | 247.7034595 | Aquaporin |
| 348.9821207 | 512.8482157 | 185.5368489 | 104.9935003 | 107.0262479 | 148.0364704 | transporter |
| 128.4282495 | 130.9454898 | 158.2046142 | 137.2989199 | 87.37032392 | 77.19266941 | Transcriptic |
| 352.6719662 | 361.516579 | 340.5819334 | 422.1478238 | 275.053529 | 186.0801355 | Xaa-Pro dip |
| 18.83386036 | 17.62776016 | 27.02591161 | 24.50601081 | 12.73542529 | 10.3203377 | Transcriptic |
| 157.2353393 | 137.2977391 | 177.5410596 | 171.8925859 | 117.4885434 | 107.5618185 | putative rep |
| 32.71365761 | 30.14440993 | 53.67308886 | 50.44853889 | 40.55502234 | 57.65676151 | FIG011154 |

| | | | | | | |
|---|---|---|---|---|---|---|
| 216.0243784 | 203.8406222 | 235.1515851 | 240.32723 | 172.580932 | 255.0372715 | Transcriptio |
| 79.25220233 | 64.2069019 | 84.11664371 | 104.4700277 | 58.82152985 | 63.2335967 | Transcriptio |
| 98.82316968 | 92.5338214 | 86.48425153 | 109.9057454 | 97.46769683 | 101.6154195 | DNA polym |
| 446.6730026 | 437.9258301 | 512.371782 | 492.6095559 | 343.6466241 | 345.7451224 | Transcriptio |
| 116.182748 | 115.6962668 | 108.0600562 | 132.5444503 | 108.8416251 | 84.92481936 | NAD(FAD)-u |
| 320.357162 | 344.3557158 | 257.2618477 | 283.7596255 | 319.0489357 | 337.7994932 | Deoxyribos |
| 254.8623518 | 296.4152122 | 231.468204 | 244.0218172 | 293.3316207 | 328.8772898 | Nucleoside |
| 153.0152376 | 168.3459264 | 151.7307104 | 131.6755373 | 217.1043053 | 294.428371 | Uridine pho |
| 273.4953314 | 259.9093571 | 306.8852049 | 281.7873263 | 192.4576127 | 150.5326561 | Predicted r |
| 402.4501086 | 414.91075 | 418.0831032 | 467.3029473 | 718.3782189 | 557.1473536 | SSU ribosor |
| 19.7969555 | 13.88274605 | 23.65493769 | 18.62291798 | 10.47134968 | 17.08287477 | Mobile elen |
| 248.1834071 | 268.4145213 | 241.5493272 | 259.7043927 | 324.5584149 | 400.7781287 | TsaD/Kae1/ |
| 64.01945045 | 65.07883082 | 70.07888869 | 68.36859752 | 64.4319844 | 94.82623111 | Ribosomal- |
| 284.5323801 | 333.6175981 | 287.3341192 | 304.0096693 | 453.8521142 | 429.5517109 | Inactive hor |
| 328.4751992 | 332.0837221 | 273.344395 | 282.1389916 | 289.6620107 | 280.8429832 | Protein of u |
| 635.6198973 | 656.583431 | 573.6063645 | 713.571052 | 695.0918317 | 680.1624418 | Ribonuclea |
| 296.4340561 | 348.0691518 | 274.7569625 | 236.0484941 | 483.62943 | 574.770588 | Glutamine s |
| 250.9743068 | 286.0623053 | 226.8376722 | 203.7097083 | 374.7835388 | 462.7287236 | Transcriptio |
| 47.25447203 | 43.39069931 | 58.37767477 | 45.49540368 | 45.98999708 | 55.48657533 | FIG011145 |
| 1409.741697 | 1717.844576 | 1144.831075 | 867.3555759 | 1302.423324 | 1201.115234 | Phosphogly |
| 598.9933566 | 626.9431574 | 776.1988643 | 715.3514309 | 1036.043604 | 828.0875063 | Acid phospl |
| 62.60005717 | 45.41188232 | 56.44928311 | 58.61639757 | 42.05762488 | 32.36755219 | Mobile elen |
| 803.1298228 | 779.25412 | 1075.495962 | 977.1866615 | 984.862169 | 891.5981697 | Putative sto |
| 1156.998286 | 1245.298604 | 1215.672 | 1334.084125 | 1621.997915 | 1599.79719 | Dihydroxya |
| 945.7103002 | 966.2958071 | 1104.528713 | 958.22195 | 982.010898 | 1163.348714 | FIG001802 |
| 481.4116797 | 374.6969375 | 654.3508736 | 398.6263942 | 258.0439743 | 480.1069379 | LSU ribosor |
| 2699.281214 | 3231.092875 | 2130.590143 | 1742.230363 | 2665.786586 | 2892.604425 | Fructose-bi |
| 677.3521589 | 762.0275125 | 663.0184992 | 582.7995141 | 566.4412501 | 503.0527228 | Hydrolases |
| 200.7411287 | 220.4213359 | 184.6697976 | 189.1294703 | 164.9237575 | 182.38481 | hypothetica |
| 476.7846957 | 502.0114561 | 472.7570936 | 400.6252733 | 325.2234844 | 423.5125457 | CTP synthas |
| 206.1357991 | 252.607042 | 338.9477044 | 219.0482565 | 384.0858341 | 686.7692485 | DNA-directe |
| 916.0994901 | 1096.396663 | 964.5366159 | 879.0747708 | 1642.81531 | 1624.563547 | Cell divisior |
| 651.205718 | 780.8921564 | 461.376382 | 620.4119649 | 1201.400388 | 1034.945681 | Secreted an |
| 32.80638339 | 24.69706845 | 37.35856346 | 41.12119728 | 25.64412167 | 16.40612223 | Mobile elen |
| 104.5728543 | 113.5810376 | 98.42879051 | 83.72843137 | 115.7836072 | 97.92186037 | Potassium e |

| | | | | | | |
|---|---|---|---|---|---|---|
| 212.3249943 | 238.6022564 | 219.759776 | 194.1472259 | 197.3877413 | 216.128727 | Hypothetic |
| 35.00935288 | 47.14894886 | 30.55901041 | 39.52750494 | 44.95102413 | 27.50184173 | hypothetica |
| 97.47011109 | 116.4484106 | 110.3007085 | 83.5972113 | 92.10746764 | 106.1525657 | Substrate-s |
| 158.7482925 | 194.3892611 | 158.5639807 | 150.0608511 | 207.8870893 | 193.2512523 | Novel pyrid |
| 179.4079295 | 191.4875976 | 210.1848736 | 147.3469694 | 134.6877353 | 152.5802179 | tRNA pseud |
| 21.6276417 | 17.74422807 | 21.5044888 | 20.87815236 | 21.95605579 | 18.80088684 | Competenc |
| 37.82435072 | 34.94465856 | 38.92432611 | 35.61191752 | 40.80316427 | 43.17032763 | Glycerate ki |
| 186.6216929 | 189.1813984 | 180.0270045 | 178.6936818 | 159.3045023 | 126.3126846 | Type I restri |
| 161.5988669 | 165.1397857 | 164.6065958 | 181.4273235 | 227.5150663 | 219.2546327 | Type I restri |
| 66.43079071 | 65.49064521 | 71.82105586 | 87.77369602 | 65.82034018 | 67.65085149 | Malonate p |
| 184.4610442 | 192.5621523 | 165.6294498 | 200.729184 | 208.6694601 | 168.4071324 | Type I restri |
| 18.42488927 | 16.57215529 | 27.62581606 | 23.21441488 | 24.20203655 | 29.94726445 | Transcriptio |
| 19.29932711 | 16.96159818 | 24.98574826 | 20.46969485 | 22.31599113 | 22.9528822 | FIG005250 |
| 12.50723543 | 7.564395222 | 18.38935258 | 14.46578737 | 14.95470209 | 15.21399512 | Salivaricin A |
| 36.20014719 | 18.60298663 | 33.24418422 | 16.82230798 | 22.83929586 | 34.04270362 | Lantibiotic |
| 79.82942907 | 70.0029525 | 91.6806511 | 61.51440976 | 62.50156614 | 63.59208869 | 6-phospho- |
| 142.4821164 | 131.0618602 | 174.6479839 | 113.3362546 | 163.3835811 | 144.97896 | PTS system, |
| 77.0960682 | 68.49941627 | 101.1826773 | 60.8283371 | 100.3915954 | 113.5698334 | PTS system, |
| 45.39013301 | 40.72200165 | 42.67643397 | 52.37203907 | 46.55432527 | 41.57627091 | Quinolinate |
| 112.4563643 | 105.4143166 | 154.6454664 | 81.7591521 | 122.1723973 | 119.2982682 | Tagatose 1,( |
| 102.0245681 | 81.31926234 | 136.3107113 | 91.35176476 | 126.289544 | 149.3049738 | Tagatose-6- |
| 191.1557191 | 150.1273701 | 253.1778478 | 145.6890114 | 184.5803681 | 210.2058256 | Galactose-6 |
| 119.1693657 | 101.9911994 | 157.5617353 | 74.29525916 | 95.01102247 | 107.9362223 | Galactose-6 |
| 75.38189566 | 68.8350287 | 103.4590958 | 68.22960037 | 105.2461963 | 125.8820138 | Lactose pho |
| 82.81726382 | 78.25415818 | 120.9627495 | 87.96275807 | 138.4590573 | 215.8387731 | DNA-damag |
| 136.7403997 | 137.0967948 | 178.9960686 | 158.6437099 | 431.4172696 | 533.6603945 | Toxin to DN |
| 6.091370922 | 4.835789627 | 9.925148679 | 4.696826728 | 5.034302732 | 9.791089611 | hypothetica |
| 4.294932718 | 4.528430117 | 8.747588666 | 4.346554905 | 3.99331132 | 5.790805006 | Integrase |
| 1.58375644 | 2.828936932 | 3.548240653 | 4.396229818 | 5.890134197 | 8.038999891 | Protein of u |
| 14.47406548 | 13.63468499 | 11.21508936 | 13.46527345 | 22.10425637 | 18.50034743 | Mobile elen |
| 3730.653146 | 4324.506388 | 2029.958081 | 1982.638123 | 2624.332056 | 2297.711858 | SSU ribosor |
| 2830.459747 | 3309.70854 | 1721.730196 | 1514.953581 | 1700.891548 | 1500.298587 | LSU ribosor |
| 142.5380796 | 137.346938 | 174.6070993 | 179.1941502 | 151.0947468 | 149.7118096 | Transcriptio |
| 105.8961406 | 103.5044972 | 127.9771575 | 127.4646515 | 160.7878842 | 212.8670089 | hypothetica |
| 289.1045292 | 311.4787702 | 278.509355 | 281.4659335 | 526.1442753 | 797.6676634 | DegV family |

| | | | | | | |
|---|---|---|---|---|---|---|
| 142.4397096 | 135.785068 | 165.991947 | 150.5230862 | 147.0704315 | 193.9845626 | Hypothetic |
| 347.0907185 | 416.2614695 | 376.9762797 | 311.6938711 | 634.6678735 | 636.5822504 | 23S rRNA (g |
| 77.40163051 | 80.05867883 | 76.21562715 | 68.58779603 | 85.91624317 | 98.82529941 | Hypothetic |
| 206.232632 | 224.1283366 | 173.2046327 | 184.7690793 | 133.3105252 | 146.9447625 | COG1939: I |
| 179.9401848 | 222.9050752 | 157.8995891 | 153.4918781 | 137.3049884 | 113.9007853 | Cysteinyl-tF |
| 13.59639962 | 14.38191207 | 11.41158014 | 20.21850978 | 50.75147075 | 52.70853702 | Mobile elen |
| 87.82649347 | 85.92946957 | 74.98643174 | 66.13376025 | 73.94540766 | 72.03779123 | FIG011146 |
| 312.9959892 | 368.3612895 | 270.5242522 | 260.6012521 | 281.1476426 | 355.5392735 | Serine acety |
| 328.4243932 | 380.8858644 | 251.5639652 | 234.2612502 | 271.3529287 | 253.0203153 | FIG011140 |
| 448.0850146 | 581.2384483 | 326.0379932 | 241.607145 | 369.6176569 | 330.237819 | Polyribonu |
| 59.89597399 | 40.2285956 | 51.57095 | 60.00341109 | 103.5823325 | 142.8328479 | Transaldola |
| 62.90267425 | 53.67463671 | 71.70110805 | 68.49198629 | 180.5880999 | 273.1512354 | Transport p |
| 22.33930136 | 20.67936354 | 26.31470341 | 27.38002781 | 26.04059329 | 51.83039403 | PTS system |
| 27.84829373 | 24.50098216 | 34.46352573 | 27.5964208 | 40.55361681 | 55.8458908 | Transcriptic |
| 2780.196443 | 2825.444417 | 2271.590834 | 2058.290377 | 1077.021946 | 1276.191233 | SSU ribosor |
| 154.593539 | 136.9899808 | 171.3939854 | 212.4297617 | 140.3668796 | 125.0844386 | FIG011149 |
| 5.248495178 | 5.482435914 | 7.032717996 | 7.774340521 | 15.5243847 | 16.47527594 | Mobile elen |
| 137.0133478 | 96.12537636 | 148.5310041 | 175.0824084 | 109.8121918 | 72.91185948 | FIG011181 |
| 588.7711135 | 690.0612824 | 449.6273909 | 351.3409684 | 473.2220011 | 523.0251758 | Peptide def |
| 330.8388452 | 339.4203335 | 311.6071713 | 330.4256527 | 470.885314 | 383.8511412 | FIG011146 |
| 112.3027294 | 125.7305303 | 109.402207 | 113.2362226 | 108.0544432 | 150.660977 | Transcriptic |
| 265.0901468 | 313.4686823 | 218.2720283 | 254.0851903 | 273.3595477 | 265.928581 | DNA polym |
| 790.9059623 | 916.8274331 | 697.584425 | 945.5326813 | 697.332258 | 618.8666273 | Prolyl-tRNA |
| 575.5446319 | 630.4862217 | 468.9130586 | 647.2663366 | 554.1400855 | 506.935505 | Intramemb |
| 286.4507402 | 328.7378912 | 225.918856 | 336.2148087 | 329.6252458 | 310.4115713 | Phosphatid |
| 295.4022511 | 309.2342393 | 273.2145303 | 309.4213087 | 272.2812701 | 306.0447259 | Undecaprer |
| 825.8900385 | 967.0687407 | 754.1994383 | 457.7904616 | 1112.849125 | 1329.56492 | Preprotein |
| 207.7580496 | 204.5303939 | 178.9800683 | 164.0953838 | 173.7044205 | 147.8003626 | membrane |
| 524.687586 | 680.5435923 | 322.7063697 | 282.0282491 | 699.8765777 | 607.6028581 | putative tra |
| 38.33070903 | 36.5815458 | 33.04516965 | 43.28049295 | 38.6950497 | 29.68087009 | Pullulanase |
| 29.43785204 | 27.86870955 | 27.0105173 | 28.49786522 | 32.91760251 | 39.93350782 | Glucan 1,6- |
| 112.2455953 | 110.5962998 | 99.45826072 | 86.16067698 | 158.5245994 | 172.2111154 | Multiple su |
| 182.6076771 | 166.5485249 | 181.8573068 | 201.3644135 | 150.5487775 | 148.7783107 | Leucine ricł |
| 0.502779822 | 0.071275811 | 0.07314452 | 0 | 0 | 0.364580494 | Streptokina |
| 744.1515055 | 589.6804347 | 902.6436526 | 1004.622675 | 567.5223895 | 658.7362917 | D-tyrosyl-tF |

| | | | | | | |
|---|---|---|---|---|---|---|
| 1304.287634 | 1206.184798 | 1528.402482 | 1763.145594 | 1562.822408 | 1481.267206 | GTP pyroph |
| 10196.02011 | 12643.07622 | 12113.12191 | 8873.21134 | 8245.799088 | 11234.95006 | Collagen-lik |
| 184.2469448 | 182.6411663 | 157.7774994 | 164.63107 | 167.3139569 | 160.0310227 | Ribonucleo |
| 124.6569585 | 120.4072619 | 96.77019962 | 91.29267565 | 61.7514069 | 45.381451 | FIG011169 |
| 83.35560209 | 88.09408866 | 64.93789711 | 83.87543731 | 122.4527899 | 133.2781561 | hypothetica |
| 56.21465165 | 53.13611697 | 60.90969219 | 91.78941378 | 82.70640203 | 76.04658322 | Exodeoxyril |
| 145.8620128 | 155.3091118 | 148.8500007 | 236.7722541 | 278.1856247 | 268.6834463 | PTS system, |
| 128.0308706 | 147.7962384 | 122.4465593 | 103.5312122 | 162.0964931 | 206.6022972 | Ribosomal |
| 159.4713245 | 182.1708863 | 130.7006313 | 113.7075165 | 141.6966245 | 191.2422458 | Ribosomal |
| 69.80633479 | 70.87357918 | 62.66435432 | 38.9687038 | 47.14608476 | 53.37998335 | FIG011153 |
| 183.0225527 | 187.5209714 | 127.1150596 | 145.0934005 | 320.6452993 | 296.3617627 | 6-aminohex |
| 215.0773677 | 238.9631026 | 204.2192495 | 259.5100963 | 229.809047 | 190.1374377 | Para-amino |
| 252.3116741 | 292.4565311 | 249.7763442 | 343.1889282 | 270.5098668 | 210.6515712 | Anthranilat |
| 216.5590366 | 238.4570164 | 190.5663742 | 265.887028 | 236.0231079 | 199.9297372 | Chromosom |
| 528.708613 | 657.1565217 | 415.189028 | 414.766783 | 583.7571319 | 595.7510812 | tRNA-guani |
| 415.889191 | 481.3013325 | 319.7159081 | 281.1401113 | 304.9202068 | 325.8163722 | Dipeptide-b |
| 66.11940228 | 68.07189754 | 50.90824919 | 52.19961018 | 46.43449351 | 43.83227701 | Dipeptide t |
| 72.54066904 | 79.95819308 | 59.51014101 | 68.32340137 | 54.17203714 | 50.75718909 | Dipeptide t |
| 105.6625565 | 116.5822268 | 76.12829759 | 83.72794662 | 74.94536422 | 77.01541873 | Dipeptide t |
| 98.8900552 | 112.7213307 | 80.5646638 | 78.70443334 | 69.61067687 | 68.56228376 | Dipeptide t |
| 151.5205788 | 132.0639712 | 263.831191 | 89.12754481 | 47.47272338 | 182.377311 | FIG011156 |
| 93.06491118 | 100.8018587 | 94.12780946 | 86.87533438 | 99.36027346 | 92.78020505 | Streptococ |
| 56.43744446 | 63.42839049 | 59.2599268 | 59.12952972 | 73.86650377 | 69.78480361 | Laminin-bir |
| 3.751002094 | 7.030983602 | 5.942029801 | 6.266555661 | 6.716819698 | 7.933223577 | hypothetica |
| 493.7056132 | 459.8533698 | 560.9259808 | 647.2814335 | 582.5946837 | 480.7940319 | Zinc finger |
| 2574.924012 | 2476.547652 | 2660.970485 | 3391.431805 | 1533.377817 | 1088.876255 | Fibronectir |
| 1433.822965 | 1470.804377 | 1300.735461 | 1462.469685 | 838.3203014 | 628.4738305 | C5a peptida |
| 2725.303238 | 2879.255558 | 2632.809021 | 1882.918366 | 1372.769745 | 1758.953443 | Antiphagoc |
| 8158.787906 | 8459.417402 | 10357.16223 | 8413.481685 | 5634.849175 | 7938.465436 | Antiphagoc |
| 344.6004883 | 368.3621997 | 370.438982 | 331.9812124 | 471.9460833 | 406.1652005 | M protein t |
| 294.1261959 | 325.7393649 | 316.2311881 | 431.7725714 | 479.8589486 | 356.2521083 | Hypothetic |
| 615.6525033 | 619.6512254 | 690.8762564 | 829.6905492 | 480.7202833 | 442.2560299 | Immunoger |
| 340.2852929 | 329.9068639 | 375.1325701 | 379.6060574 | 225.4280302 | 156.0052139 | Two-compo |
| 454.5035614 | 517.2675438 | 402.7569679 | 523.0509152 | 895.0190415 | 835.517551 | Ribose-pho |
| 596.2842995 | 553.4762719 | 693.519764 | 866.5253912 | 499.352488 | 351.1479813 | Substrate-s |

| | | | | | | |
|---|---|---|---|---|---|---|
| 302.8752567 | 286.4262597 | 344.0224987 | 320.7264758 | 235.9656207 | 189.5433919 | Two-compo |
| 322.6416136 | 317.0360354 | 360.7828214 | 338.3797581 | 385.4701123 | 348.2906603 | ABC transpo |
| 227.5682031 | 203.1246567 | 257.2653677 | 252.4575679 | 192.7600954 | 150.5079424 | ABC transpo |
| 279.2353553 | 251.6839871 | 362.8310558 | 296.8927466 | 202.3254134 | 141.5852227 | periplasmic |
| 298.8020483 | 272.1833147 | 408.2268791 | 337.9940888 | 232.6966596 | 236.5550153 | FIG011155 |
| 207.4880911 | 184.468278 | 235.2460745 | 221.2917029 | 168.7711852 | 159.5619675 | FIG011164 |
| 4933.702023 | 3768.732857 | 5688.821967 | 6990.144531 | 5285.463458 | 5864.908338 | FIG011148 |
| 3192.215496 | 3922.258783 | 3154.262188 | 2904.84111 | 1663.746815 | 1686.230787 | Peptidylpro |
| 72.67923388 | 41.33606475 | 75.33935633 | 89.58071033 | 133.1331702 | 97.45910827 | Streptopair |
| 56.76119571 | 35.21400191 | 65.16021796 | 64.18054809 | 96.29901941 | 78.52632099 | Strepotoco |
| 205.0964589 | 214.3463752 | 178.7147084 | 162.1814269 | 259.1659047 | 323.2605533 | Metal-depe |
| 18.61608447 | 18.74928961 | 29.71014901 | 28.92256459 | 53.39010529 | 28.20701716 | FIG011143 |
| 10.79833936 | 6.429402117 | 17.22802796 | 8.326192837 | 10.26311262 | 11.41903394 | hypothetica |
| 32.46418894 | 24.66510848 | 44.82652782 | 35.72262663 | 33.88748618 | 35.40306891 | RopB; Rgg-l |
| 505.9286656 | 446.1815969 | 554.3533065 | 1486.286643 | 4300.952892 | 5488.242959 | Streptodor |
| 6.261362668 | 6.578923097 | 9.376957328 | 10.22379027 | 5.479194602 | 6.07598829 | FIG011153 |
| 60.55539328 | 53.71282213 | 60.33906565 | 69.39147065 | 71.60555298 | 96.70443987 | Low tempe |
| 114.8332493 | 116.9848116 | 122.5400416 | 158.7527434 | 187.9542363 | 173.6246808 | Glycerol de |
| 100.0678396 | 93.2407464 | 98.2166899 | 111.4664399 | 106.5330699 | 75.34309315 | Transaldola |
| 97.59948183 | 97.02777896 | 112.4383275 | 115.7237502 | 98.97276775 | 80.81389158 | Pyruvate fo |
| 76.16594188 | 74.24604591 | 89.94808593 | 87.84037739 | 91.08268819 | 77.24832079 | PTS system, |
| 254.5059767 | 237.6635969 | 256.1784742 | 216.4035608 | 280.3521238 | 414.5693548 | tRNA-speci |
| 53.8169664 | 54.62564302 | 72.02959842 | 57.6204879 | 59.28258043 | 61.07298461 | PTS system, |
| 59.20584821 | 52.5835629 | 78.23011465 | 56.32241472 | 46.05681257 | 43.38806016 | PTS system, |
| 426.4518487 | 368.3481591 | 469.5475537 | 592.5499364 | 851.2774989 | 928.1111418 | Deoxyribor |
| 411.5858603 | 327.3285794 | 463.2533251 | 505.4634384 | 512.6072612 | 547.1524123 | Transcriptio |
| 380.8995623 | 273.4517202 | 504.230252 | 369.5474193 | 310.1832944 | 374.9196364 | Pyruvate fo |
| 72.56913836 | 51.37124279 | 119.1573354 | 69.44293369 | 49.81705539 | 65.69182747 | FIG011142 |
| 579.8159169 | 402.2311456 | 892.5274061 | 722.769987 | 362.3930023 | 1213.003331 | Preprotein |
| 740.0138577 | 807.6982415 | 725.0307999 | 798.4016008 | 777.8037623 | 645.2439014 | Multimodu |
| 141.4157325 | 127.3392871 | 137.9165837 | 114.5212098 | 126.7692924 | 162.9954702 | Hypothetic |
| 388.7627136 | 371.0524048 | 335.6716299 | 315.4065924 | 339.9098276 | 355.7885499 | FIG011154 |
| 106.2323116 | 78.3885063 | 113.8496247 | 83.71199284 | 63.82184719 | 54.46570565 | S-layer hom |
| 360.1482983 | 373.417607 | 344.1071901 | 329.4159596 | 402.2341642 | 380.0675195 | Similar to ri |
| 269.9624322 | 266.5165446 | 281.5830001 | 309.6247783 | 195.6558288 | 174.1905781 | Hypothetic |

| | | | | | | |
|---|---|---|---|---|---|---|
| 306.4568711 | 288.3629712 | 316.3740393 | 405.4789302 | 580.0211481 | 536.6434377 | Dipeptidase |
| 172.7734298 | 129.6596094 | 181.0775705 | 224.3908969 | 248.5458142 | 206.4561336 | transcriptio |
| 135.9539236 | 97.82965419 | 160.9212308 | 200.2589706 | 238.0320149 | 196.2328906 | Transcriptio |
| 118.9577059 | 91.15463446 | 167.3766045 | 246.6773398 | 245.2041051 | 171.4986643 | hypothetica |
| 8779.309273 | 9080.67954 | 10761.86015 | 13572.88815 | 14108.78425 | 13401.96597 | Heat shock |
| 8102.465291 | 8363.510623 | 11046.26841 | 12774.18239 | 8911.307496 | 9004.923022 | Heat shock |
| 1111.175178 | 1076.326618 | 1210.12223 | 1588.081833 | 1051.431113 | 827.203225 | ATP-depenc |
| 428.4369693 | 330.042642 | 612.249553 | 574.2694145 | 199.7800928 | 307.0741354 | Transcriptio |
| 72.71800844 | 67.88111077 | 100.201767 | 73.972023 | 68.25871828 | 90.73807324 | Streptococ |
| 63.7670356 | 58.72939244 | 89.13044702 | 69.89619776 | 50.63448695 | 52.35927561 | hypothetica |
| 1384.622358 | 1381.417977 | 1565.63747 | 966.5240555 | 1272.442226 | 2922.412902 | Cold shock |
| 1369.652895 | 1318.825857 | 1128.468175 | 1071.825906 | 2685.103243 | 3205.81989 | Alkyl hydro |
| 1228.170576 | 1054.377211 | 919.8850189 | 1205.737004 | 2817.88938 | 3868.552384 | Alkyl hydro |
| 42.93406083 | 31.84229959 | 42.69048696 | 46.44516099 | 126.82864 | 183.4492155 | Imidazolon |
| 48.7525616 | 36.67914288 | 53.00681786 | 48.32389497 | 126.8510437 | 191.8910461 | Urocanate h |
| 59.76040966 | 45.10582774 | 67.84666218 | 68.50791466 | 218.7857624 | 261.9374131 | Glutamate f |
| 57.21225416 | 40.1555641 | 57.49106756 | 58.19570572 | 193.3316775 | 258.8634893 | Formimino |
| 77.45647534 | 50.27531285 | 77.52021368 | 99.36818741 | 295.5974754 | 386.8588612 | Formate--te |
| 61.91771407 | 40.98878469 | 56.73294785 | 85.78907433 | 209.6572055 | 228.949909 | Conserved p |
| 44.34518031 | 34.96880374 | 77.4161597 | 43.04641697 | 19.14293614 | 39.19012447 | hypothetica |
| 49.31563021 | 34.27419869 | 51.6971751 | 81.611558 | 210.3619356 | 247.5850469 | putative ca |
| 53.55192008 | 39.04617879 | 58.55161884 | 95.22313613 | 273.4978006 | 318.9011824 | Histidine ar |
| 85.8309341 | 78.67712135 | 70.34713093 | 116.4755924 | 185.4765662 | 181.9159702 | Formimino |
| 26.56075434 | 26.98606504 | 30.49441412 | 40.97786574 | 68.17231742 | 75.42934776 | Transcriptio |
| 3003.940535 | 3243.638949 | 2743.208354 | 2880.303305 | 2734.847595 | 2456.605318 | SSU ribosor |
| 2087.317961 | 2238.782461 | 1790.155734 | 2165.966687 | 2065.168185 | 1740.373975 | Translation |
| 675.1513607 | 795.4644389 | 482.090148 | 353.5417413 | 550.0254533 | 584.8944818 | Neutral enc |
| 180.513234 | 151.5191819 | 203.0451457 | 282.6581464 | 572.4892249 | 759.9297687 | Trehalose-6 |
| 331.6737931 | 315.0248287 | 384.4285782 | 685.4319305 | 1877.542085 | 2092.61122 | PTS system, |
| 94.75920883 | 86.57180946 | 115.2026186 | 120.7577833 | 84.39225887 | 71.18568181 | Trehalose o |
| 72.45685712 | 49.5063963 | 76.60974137 | 56.78463515 | 28.46898195 | 32.15599956 | hypothetica |
| 21.95662337 | 14.64474927 | 22.72633476 | 14.15452782 | 20.08000294 | 11.87579529 | FIG011156 |
| 25.04545067 | 11.33036756 | 28.88102857 | 30.24537956 | 21.00357931 | 15.42366258 | FIG011147 |
| 54.24365806 | 42.63050793 | 65.31988475 | 84.49004181 | 40.74009486 | 49.74131183 | Transcriptio |
| 158.4998602 | 140.2141943 | 178.4872571 | 196.8965021 | 118.18766 | 91.50273405 | Glyoxalase f |

| | | | | | | |
|---|---|---|---|---|---|---|
| 478.8266908 | 433.6157428 | 549.6203619 | 634.8107811 | 329.8314217 | 262.8324657 | UPF0246 pr |
| 864.344734 | 833.1947581 | 1072.811742 | 1333.254982 | 920.8722 | 1024.678376 | Ribonucleo |
| 411.9686448 | 348.2354688 | 599.584247 | 571.9539399 | 246.1957102 | 317.0532836 | Acetyltrans |
| 524.2132292 | 475.5193133 | 542.0578596 | 618.2785285 | 435.2784 | 390.5485284 | Putative ox |
| 113.12546 | 82.10973407 | 189.9197455 | 125.6065662 | 58.50065257 | 125.096682 | FIG011151 |
| 829.7544141 | 790.447089 | 973.9949196 | 1038.731745 | 850.8953753 | 809.0820184 | Ribonucleo |
| 628.0474148 | 682.4723035 | 481.3421411 | 419.4735951 | 434.0374444 | 352.3761619 | Glucose-6-p |
| 180.2436656 | 152.9922943 | 205.7917546 | 199.1689374 | 170.9649208 | 184.355911 | Competenc |
| 3.654822553 | 2.820877282 | 14.88772302 | 3.28777871 | 2.517151366 | 29.88858934 | hypothetica |
| 1791.352749 | 1869.625312 | 2723.005895 | 1988.000984 | 4375.445765 | 4677.988613 | Ortholog of |
| 2436.672864 | 2308.692447 | 3334.451501 | 2758.206354 | 3927.042756 | 3809.283032 | Putative Ho |
| 983.8646949 | 843.6169331 | 1762.651266 | 1233.590228 | 1055.64294 | 1903.009919 | Hypothetic |
| 1869.070758 | 1867.287443 | 2976.229335 | 2426.052263 | 2037.779761 | 1667.941368 | Transcriptio |
| 1093.878425 | 1132.901744 | 1219.840709 | 1247.821696 | 1099.336304 | 1183.737431 | RecA protei |
| 212.7612425 | 179.8851013 | 217.3525631 | 244.6094067 | 167.118666 | 144.711478 | Molybdopt |
| 577.435904 | 521.579994 | 530.338643 | 838.9863723 | 546.0710866 | 364.4418266 | DNA-3-meth |
| 400.7938407 | 354.1304635 | 431.0083095 | 547.3195665 | 296.2826297 | 177.5447463 | Holliday jun |
| 2380.20016 | 1793.534766 | 2451.70411 | 3194.112967 | 1281.85558 | 853.6043423 | Transcriptio |
| 439.1324674 | 393.6802077 | 447.9089133 | 591.3847236 | 388.6716726 | 268.9213354 | Multidrug r |
| 509.6533015 | 450.0667942 | 568.421223 | 638.3733616 | 438.7748984 | 347.4647911 | DNA misma |
| 320.1332678 | 282.1569908 | 405.7712233 | 354.999374 | 267.7659096 | 217.356553 | Phage integ |
| 365.9137274 | 283.2211164 | 414.6334248 | 465.9545666 | 285.9169308 | 260.8487985 | hypothetica |
| 220.6123228 | 194.3052385 | 244.5715592 | 256.0889733 | 163.2058017 | 105.1978501 | Phage trans |
| 28.69123985 | 30.29376907 | 41.84025539 | 40.08639508 | 19.9183282 | 21.55384029 | conserved h |
| 55.02329074 | 53.22484433 | 70.58238787 | 97.25336581 | 54.24590944 | 73.65630055 | kilA protein |
| 52.08122138 | 39.2152315 | 67.30491448 | 85.15933962 | 40.4946726 | 66.76621304 | Phage antir |
| 148.5993696 | 118.7455008 | 197.9209433 | 207.3735979 | 91.62430973 | 128.5247513 | FIG011181 |
| 50.82266035 | 50.14253291 | 47.70241763 | 56.30973378 | 45.96797852 | 46.68149408 | Mobile elem |
| 592.8288309 | 413.2676805 | 576.5891061 | 493.7432352 | 589.9058642 | 699.3016382 | FIG011180 |
| 28.34090471 | 26.60744775 | 27.58799551 | 45.31201786 | 18.08374534 | 14.10350858 | FIG011184 |
| 42.8042281 | 33.13169379 | 21.5771391 | 41.58595774 | 23.08295834 | 11.9498647 | FIG011157 |
| 29.94289519 | 27.25798606 | 19.40444107 | 39.49737727 | 15.95244678 | 11.61886703 | phage prote |
| 36.71435383 | 43.5770588 | 36.50875713 | 55.95201586 | 25.70240377 | 18.08774975 | FIG011144 |
| 40.72516559 | 39.89526442 | 43.77699507 | 58.17355391 | 28.04825808 | 24.07282934 | FIG011140 |
| 40.97831347 | 35.6639485 | 38.34716535 | 58.34739749 | 33.08909887 | 21.7840032 | Phage prote |

| | | | | | | |
|---|---|---|---|---|---|---|
| 28.15879567 | 24.56546773 | 25.15223069 | 39.69360848 | 21.90055817 | 14.13607441 | DNA primas |
| 14.47225712 | 10.83883882 | 13.06953845 | 23.37076196 | 12.18648454 | 14.89987049 | FIG011143 |
| 21.02573204 | 15.17437435 | 15.29414074 | 25.26568861 | 17.26418644 | 9.355732632 | Phage prote |
| 26.83069733 | 27.2724312 | 24.28742265 | 34.15698167 | 27.37180009 | 16.55088213 | FIG011154 |
| 30.66956915 | 20.58089038 | 37.05684232 | 28.93021606 | 20.41289894 | 24.72311474 | DNA recom |
| 4.355330209 | 3.536171165 | 9.273810797 | 6.411168484 | 2.945067098 | 3.517062452 | hypothetica |
| 32.93824743 | 25.74390459 | 29.58516328 | 44.16457869 | 23.12690728 | 18.6179292 | FIG011155 |
| 12.82711827 | 11.04038747 | 10.92996745 | 14.53299113 | 10.0602843 | 10.29789242 | FIG011163 |
| 16.20875731 | 14.24291163 | 12.09627495 | 20.60732727 | 15.33889114 | 11.61886703 | Phage enco |
| 98.83332536 | 93.78260866 | 100.2073925 | 71.56885974 | 51.49844106 | 116.4117471 | hypothetica |
| 85.8797506 | 90.82997816 | 87.91095599 | 100.8544311 | 129.6474765 | 122.7551509 | DNA misma |
| 148.0467976 | 140.7635285 | 160.5824327 | 150.5230862 | 200.0369246 | 219.0336202 | biofilm forr |
| 96.0016746 | 89.99205764 | 103.8401685 | 92.46636346 | 133.4021261 | 136.5528749 | Arginine pa |
| 632.3231828 | 591.5632597 | 634.5539981 | 693.8936913 | 535.0553345 | 416.2391743 | Arginyl-tRN |
| 139.1552988 | 126.1882871 | 157.0558418 | 154.8995215 | 110.1779147 | 146.9263815 | bacteriocin |
| 228.4201298 | 210.1449714 | 234.7203868 | 246.3751934 | 184.1257299 | 186.4025834 | Transporter |
| 87.88434172 | 78.44125718 | 89.0660245 | 113.2585249 | 56.53477019 | 69.89264414 | Integral me |
| 163.1323186 | 143.6460581 | 190.3477533 | 189.4905088 | 118.3387599 | 128.9532406 | membrane |
| 247.7015247 | 264.4245189 | 219.5820612 | 263.7971236 | 233.7920569 | 225.7448457 | Integral me |
| 272.4713051 | 326.5111884 | 229.9191335 | 247.1465391 | 316.5315684 | 343.0708701 | Aspartyl-tRI |
| 132.1157011 | 147.3726708 | 106.7040648 | 96.95042338 | 132.7924087 | 135.2227558 | Histidyl-tRN |
| 789.0222656 | 805.9120876 | 582.4719939 | 499.0801883 | 711.6440824 | 975.5524048 | LSU ribosor |
| 675.3137459 | 679.8878426 | 460.6261502 | 552.4595471 | 1015.851811 | 1219.516283 | LSU ribosor |
| 39.0144879 | 43.23903603 | 45.47412633 | 52.09353626 | 40.79995395 | 31.56778006 | Cadmium r |
| 40.64507677 | 37.6913426 | 45.95870838 | 43.76777473 | 31.62254937 | 36.46006588 | Cadmium e |
| 39.38822834 | 32.8613886 | 50.63678736 | 44.42618606 | 39.20381527 | 37.32392807 | FIG011151 |
| 99.63153061 | 78.64419624 | 107.2439994 | 128.4423054 | 71.10401042 | 61.73375414 | FtsK/SpoIIII |
| 468.3695711 | 501.3504895 | 389.2850086 | 415.5047765 | 461.4592913 | 482.5409685 | UTP--glucos |
| 113.3055602 | 100.6313386 | 115.7872289 | 151.2798818 | 87.8147702 | 52.09351919 | FIG011145 |
| 32.53899594 | 22.86009642 | 43.98645437 | 33.97086677 | 27.48729292 | 48.23399935 | Transcriptio |
| 146.3706574 | 154.1988755 | 143.4334098 | 151.4952268 | 170.7649819 | 174.7266613 | FIG028593 |
| 49.65890519 | 50.52722245 | 54.41439574 | 57.00718572 | 65.04836675 | 60.76206693 | Phosphohy |
| 14.82047311 | 10.66979271 | 15.83243327 | 14.28438588 | 11.70821782 | 10.14094023 | FIG014387 |
| 104.1562271 | 98.22697679 | 120.7158868 | 107.1020274 | 96.86666272 | 82.85091724 | FIG04612: I |
| 100.6565204 | 97.26653524 | 113.4809795 | 95.30252834 | 72.86314154 | 54.5423812 | FIG015389 |

| | | | | | | |
|---|---|---|---|---|---|---|
| 189.3194868 | 153.8457346 | 220.3500852 | 292.1636901 | 166.6799189 | 130.6602621 | Phage infec |
| 52.85119804 | 40.43861157 | 82.36340024 | 56.39358474 | 52.28321366 | 42.11442359 | Transcriptio |
| 1282.532176 | 1535.884713 | 810.2923006 | 782.3600981 | 1224.993922 | 875.0293754 | SSU ribosor |
| 899.0504108 | 1045.5291 | 729.6301777 | 821.6994449 | 884.7363814 | 744.2003365 | Glycerol-3-p |
| 52.58238973 | 45.31953638 | 52.05451655 | 57.86043213 | 44.30585952 | 54.81619635 | Mobile elen |
| 47.68673236 | 42.31315923 | 69.83051035 | 49.51797322 | 31.28459555 | 52.34183995 | FIG011148 |
| 437.2772461 | 478.4822108 | 443.5300816 | 432.2797932 | 500.0559597 | 522.2251196 | Replicative |
| 425.7263171 | 457.3343958 | 433.3296908 | 423.3676066 | 612.0278513 | 664.1491632 | LSU ribosor |
| 168.6616494 | 164.4847245 | 208.8855978 | 166.3596071 | 207.8082247 | 285.817553 | Phosphoest |
| 195.9304373 | 214.4172314 | 186.2023746 | 169.5176032 | 147.8892041 | 162.9389709 | tRNA uridir |
| 122.5170076 | 142.3364494 | 141.6557482 | 100.22851 | 130.3979394 | 173.2935354 | MutT/nudix |
| 364.3910204 | 400.5559541 | 344.8624271 | 298.2246364 | 350.3632409 | 437.8460636 | tRNA-specif |
| 144.4470717 | 128.887826 | 136.0110916 | 101.0740338 | 72.39956617 | 58.58780054 | L-serine deh |
| 130.0748416 | 116.6031851 | 146.485474 | 111.9200546 | 85.01224614 | 84.46474971 | L-serine deh |
| 262.6924015 | 233.2301337 | 276.3326811 | 309.5678497 | 235.0818004 | 307.7597125 | Transcriptio |
| 532.2194201 | 663.6118721 | 388.1822479 | 422.0201917 | 688.7147156 | 580.7687238 | Immunodo |
| 160.3924125 | 160.5183315 | 184.6003309 | 205.061344 | 212.9566621 | 318.0222335 | Transmemb |
| 99.53430151 | 106.0991174 | 132.4707122 | 127.1803733 | 108.0207292 | 89.97385999 | ATPase com |
| 327.3285184 | 338.9672645 | 362.4274381 | 325.5303508 | 353.8988963 | 302.9698084 | ATPase com |
| 259.5260553 | 270.303274 | 329.6067849 | 282.4557417 | 239.835851 | 248.9425104 | CDP-diacylg |
| 326.2445648 | 327.5151643 | 382.3649018 | 304.6510137 | 266.433848 | 265.0871967 | Transcriptio |
| 73.8104164 | 72.95294723 | 70.70225825 | 66.28424028 | 90.2240711 | 70.94884788 | FIG009210 |
| 45.87169254 | 52.41296805 | 53.35963216 | 42.06432747 | 72.76326421 | 67.94407739 | FIG001621 |
| 402.3872611 | 399.0821743 | 404.7836009 | 205.1573915 | 189.3724891 | 216.0959733 | Hyaluronar |
| 390.0049357 | 411.0421183 | 388.001276 | 224.5386197 | 218.7485736 | 199.3292963 | UDP-glucos |
| 295.6252574 | 254.0850501 | 288.0146451 | 396.3238524 | 221.7340501 | 172.4358412 | Lipid A expo |
| 351.4430957 | 342.7653743 | 339.0797074 | 183.6123763 | 143.0461162 | 88.04618928 | FIG011171 |
| 679.717108 | 788.9075487 | 659.9410335 | 379.6252552 | 334.0317633 | 311.4782991 | UTP--glucos |
| 184.8863496 | 211.1452927 | 186.0605771 | 173.0882754 | 203.0246739 | 190.0524431 | FIG002958 |
| 208.6352318 | 230.2504223 | 187.4212358 | 194.2958842 | 200.062372 | 159.3094491 | DNA recom |
| 465.1734626 | 463.2493367 | 535.5961743 | 548.0022585 | 557.9266003 | 617.6492017 | Putative su |
| 2090.814979 | 2159.052811 | 2433.718283 | 3020.442749 | 1927.925976 | 1822.241114 | Inosine-5'-n |
| 140.308745 | 134.9483111 | 168.1417691 | 142.5121909 | 147.74276 | 156.1829158 | Tryptophar |
| 268.484425 | 234.2479523 | 298.3747822 | 192.7711884 | 129.9569291 | 348.3566619 | FIG011160 |
| 203.8746262 | 176.1739647 | 204.3480676 | 240.395213 | 169.484732 | 127.76761 | hypothetica |

| | | | | | | |
|---|---|---|---|---|---|---|
| 355.1427172 | 367.4998625 | 306.9765777 | 304.9545221 | 321.7758496 | 231.8655987 | ABC transpo |
| 311.9334742 | 275.6826775 | 279.8203839 | 440.7620609 | 275.3019025 | 219.2147449 | Lipid A expo |
| 111.2685694 | 92.94398922 | 156.6272808 | 122.9775774 | 89.46055586 | 92.86027523 | ABC transpo |
| 157.990536 | 153.3893362 | 182.0201374 | 227.8289313 | 225.4602532 | 176.723607 | Mobile elen |
| 254.9847868 | 254.8007778 | 300.390828 | 252.7832145 | 577.969418 | 677.1601314 | LSU m3Psi1 |
| 1128.115923 | 1335.270558 | 1107.08587 | 979.5439355 | 969.802896 | 1072.753307 | Serine prot |
| 580.9145242 | 522.5826867 | 698.6260422 | 809.7238607 | 442.8592532 | 362.9965395 | Chromoson |
| 781.0265553 | 864.1063529 | 672.6126221 | 3753.756108 | 17255.54808 | 11395.28235 | Streptolysi |
| 21.90863075 | 16.76407071 | 23.38613158 | 22.59173656 | 15.37979485 | 12.39345817 | paratox? |
| 334.211237 | 251.3843956 | 252.5466185 | 84.97590562 | 72.40555208 | 68.52757224 | small RNA f |
| 139.7697248 | 124.5166685 | 132.6328685 | 160.0185955 | 86.6759585 | 78.837753 | FIG053235 |
| 176.9768102 | 148.7248722 | 203.2823656 | 166.3338631 | 97.2465109 | 176.0217258 | Deoxyuridi |
| 356.3800834 | 347.9052394 | 330.3473151 | 387.7364955 | 378.5773477 | 364.3667847 | DNA repair |
| 194.0572995 | 199.1668891 | 238.1334129 | 234.5855271 | 223.0420856 | 321.5599956 | Phosphate: |
| 487.3390298 | 474.6138165 | 412.3420956 | 479.6789914 | 462.4583275 | 394.685838 | Carbonic ar |
| 342.8799279 | 290.7186945 | 358.7710917 | 410.1756618 | 310.7439293 | 317.2352172 | DUF124 do |
| 611.7834264 | 686.9530115 | 498.7412951 | 533.3012883 | 718.7511611 | 658.839405 | Glutamyl-tl |
| 72.52123029 | 73.68117101 | 76.870166 | 63.76491467 | 52.64826706 | 46.99860738 | Histidine ki |
| 73.1562262 | 72.63288228 | 76.60974137 | 70.70260571 | 75.73685015 | 56.39508685 | Sensory tra |
| 102.65563 | 115.8044358 | 103.5610908 | 100.5615323 | 74.79914007 | 67.20864281 | Response re |
| 429.5939343 | 437.43747 | 472.4267384 | 424.3582949 | 374.514366 | 379.3403074 | Ribonuclea |
| 893.2887112 | 948.072803 | 960.307172 | 1077.481755 | 959.4943242 | 757.1117387 | Inner meml |
| 1185.532566 | 1242.516507 | 1262.771785 | 1545.466966 | 1259.072511 | 992.289339 | RNA-bindin |
| 62.96397553 | 59.79866685 | 60.57971846 | 77.73821019 | 111.0984661 | 78.91945015 | FIG011156 |
| 302.0002055 | 293.7582706 | 288.5658415 | 325.9957137 | 308.8046609 | 225.5268241 | DNA polym |
| 95.61196284 | 90.02926861 | 149.1376122 | 129.3540872 | 128.9527322 | 213.132744 | Acyl carrier |
| 1389.482316 | 949.2655037 | 1341.521693 | 962.6929184 | 681.9466481 | 1570.731256 | LSU ribosor |
| 95.90150696 | 76.90963289 | 110.9082572 | 102.7345904 | 98.67019129 | 146.4979448 | N-acetylma |
| 142.3941017 | 123.337364 | 147.8311421 | 141.3371234 | 212.4464311 | 240.8502638 | ABC transpo |
| 68.77746034 | 56.84312526 | 71.69837739 | 71.44444096 | 79.63608228 | 85.02306738 | ABC transpo |
| 91.76639298 | 84.87945547 | 89.60850615 | 91.45586399 | 92.00233679 | 95.6989247 | ABC transpo |
| 102.6216842 | 94.58235593 | 110.3443 | 104.7668644 | 122.3335564 | 115.9471138 | FIG011141 |
| 102.5547203 | 88.06290011 | 100.7890713 | 108.1040119 | 108.9835759 | 110.1738346 | N-acetylneu |
| 112.6339244 | 107.0015016 | 135.1072755 | 144.6682656 | 157.6347948 | 167.3293428 | N-acetylma |
| 154.487722 | 149.1631607 | 154.9953498 | 167.5698911 | 150.1239979 | 113.4021375 | Sialic acid u |

| | | | | | | |
|---|---|---|---|---|---|---|
| 162.6949797 | 164.4855375 | 148.8208373 | 161.3893712 | 140.3369095 | 168.7733216 | Putative de |
| 77.50710086 | 55.74481165 | 99.07425199 | 92.1114819 | 55.85610252 | 70.54632557 | Phosphorib |
| 316.1677987 | 329.13275 | 261.1946528 | 323.3542721 | 332.0175645 | 349.27339 | Ribonuclea |
| 465.8747465 | 517.3422464 | 410.3588534 | 522.2096588 | 545.9675363 | 475.5717633 | SSU rRNA (a |
| 637.9653947 | 625.196142 | 642.5851629 | 678.6333572 | 610.9411816 | 585.8662893 | Ribosome s |
| 675.9988459 | 676.654975 | 707.8966337 | 729.3895625 | 654.1336062 | 547.2703772 | Ribulose-ph |
| 503.6495597 | 538.302454 | 490.0424564 | 577.3090104 | 545.6516103 | 527.8689265 | Thiamin pyr |
| 669.398565 | 743.6708813 | 709.6481306 | 814.1838359 | 710.5668966 | 567.9913661 | DNA recom |
| 406.4637534 | 396.6737977 | 389.6056866 | 439.7400273 | 362.9426673 | 373.5694997 | 3'->5' exorib |
| 244.6903699 | 235.239577 | 277.2351017 | 272.2129802 | 255.5897517 | 195.4481848 | PurR: transc |
| 1125.463715 | 1140.242123 | 1084.752602 | 1167.04644 | 1039.691804 | 919.3571357 | Surface excl |
| 3856.561696 | 4207.303425 | 2960.607122 | 3364.204249 | 2287.050899 | 1807.901171 | SSU ribosor |
| 423.8524984 | 340.140565 | 394.937209 | 535.3996741 | 270.1620865 | 194.3567385 | Phosphorib |
| 2923.493336 | 3328.555623 | 2021.284976 | 2682.890251 | 3194.178506 | 1961.746391 | SSU ribosor |
| 4231.052179 | 4959.212166 | 3184.689116 | 3617.453248 | 3882.332223 | 3356.340456 | Translation |
| 8439.072036 | 9298.462927 | 6724.763135 | 6988.200825 | 7316.75849 | 7234.980629 | NAD-depen |
| 26.39594066 | 16.92526369 | 27.29415887 | 16.43889355 | 18.12348984 | 20.09749973 | FIG011145 |
| 316.4948092 | 343.6592075 | 346.7272335 | 315.4784353 | 202.7764148 | 237.6712417 | Amino acid |
| 292.3511379 | 305.1265308 | 258.8406152 | 266.3129394 | 245.9011309 | 205.3866436 | Amino acid |
| 181.6632776 | 190.1870113 | 195.4747984 | 196.7004697 | 213.7696679 | 210.0533071 | FIG011147 |
| 196.0464221 | 225.0239714 | 185.5338173 | 199.1387435 | 238.0595904 | 226.8146398 | Undecaprer |
| 231.8893779 | 240.5710147 | 284.7228183 | 229.5472361 | 260.8046297 | 349.3325248 | Negative reg |
| 284.1845 | 306.110021 | 289.6187654 | 307.4708991 | 233.7104185 | 220.2104059 | Undecaprer |
| 395.1227406 | 324.2745919 | 364.0687785 | 519.9183821 | 233.2978926 | 190.6568995 | Amidophos |
| 31.67512879 | 26.94225649 | 35.32880304 | 31.40164156 | 20.10125162 | 34.93136857 | hypothetica |
| 337.3955839 | 409.6026712 | 312.400813 | 280.8227218 | 454.4036083 | 465.1359081 | Iron-sulfur |
| 406.6981204 | 482.8007566 | 348.0432556 | 348.2524096 | 482.1702095 | 457.2777907 | Iron-sulfur |
| 431.4477812 | 466.8783445 | 346.8190325 | 330.9712546 | 392.387587 | 332.81263 | Cysteine de |
| 420.6853043 | 454.3979946 | 358.4529478 | 324.1074639 | 281.0084856 | 259.0065277 | Putative iro |
| 439.3668499 | 470.9578585 | 325.2951742 | 329.3299715 | 331.3252372 | 283.2770205 | Iron-sulfur |
| 488.2311096 | 493.0296175 | 469.8103135 | 598.251748 | 431.8933425 | 309.2668549 | D-alanyl-D- |
| 78.841014 | 82.1760674 | 76.16827148 | 60.34565261 | 73.51919326 | 60.24360016 | D-alanyl-D- |
| 662.3091047 | 656.7602186 | 651.9345995 | 737.0543599 | 852.5304331 | 788.9415761 | Oligopeptic |
| 392.7779194 | 366.5571972 | 395.3542321 | 428.9102794 | 329.7769745 | 247.3476713 | Oligopeptic |
| 415.5851209 | 347.2798921 | 396.0199812 | 590.5688003 | 215.0503541 | 175.5142352 | Phosphorib |

| | | | | | | |
|---|---|---|---|---|---|---|
| 497.6275493 | 466.9627826 | 483.6422167 | 545.2012626 | 466.7629537 | 399.1532551 | Oligopeptic |
| 561.1466444 | 539.8136423 | 575.561318 | 688.1680009 | 506.9035191 | 406.8477046 | Oligopeptic |
| 579.973665 | 553.8980951 | 535.220893 | 642.484723 | 504.231185 | 415.3265772 | Oligopeptic |
| 21.11675253 | 14.14468466 | 25.53658046 | 20.14938666 | 12.76195743 | 9.043874877 | Transposase |
| 13.45341492 | 6.928698578 | 12.31305407 | 12.9996043 | 12.66695526 | 15.55935463 | Competenc |
| 47.59648983 | 39.58183361 | 46.63572671 | 53.06295911 | 39.47532619 | 47.47901391 | Mobile elen |
| 422.0101775 | 441.7493824 | 528.8450055 | 458.8799714 | 702.2852311 | 660.9500808 | FIG001553 |
| 414.7913313 | 485.4361392 | 466.8578397 | 411.9746642 | 617.1744723 | 636.1693815 | GTP-bindin |
| 442.5292265 | 527.488402 | 553.2186331 | 421.8388806 | 631.5215079 | 584.1933416 | FIG004454 |
| 399.4669086 | 441.4720693 | 458.779414 | 436.5845133 | 730.4510813 | 838.761055 | Nicotinate- |
| 581.8806768 | 517.8738599 | 521.600094 | 844.9870555 | 467.7084938 | 334.8134819 | Phosphorib |
| 379.6216193 | 397.6704273 | 416.8123834 | 404.8379984 | 404.6740346 | 380.0254494 | Hydrolase (I |
| 177.984057 | 187.8473994 | 238.3414176 | 211.8157025 | 185.5859743 | 271.7947582 | Iojap protei |
| 151.0316121 | 138.4049789 | 191.7194546 | 159.6144596 | 164.3537445 | 193.5194732 | FIG145533 |
| 150.306641 | 143.5338371 | 179.9034063 | 166.8939097 | 138.5006261 | 148.7432839 | FIG007079 |
| 317.0826177 | 337.6706449 | 292.4025627 | 218.8151348 | 258.3608375 | 399.0068461 | FIG007491 |
| 776.3224626 | 836.4324202 | 773.9320117 | 590.2055867 | 920.8312998 | 834.1535207 | Cysteine AB |
| 243.8535624 | 236.3577921 | 319.8220782 | 279.4412038 | 293.4623598 | 355.7685058 | Methionine |
| 257.8177032 | 249.6459367 | 328.7915186 | 332.4003813 | 231.0149346 | 190.1619763 | Methionine |
| 151.1767511 | 147.0938347 | 207.7138123 | 229.4064369 | 139.4763379 | 219.6804624 | Methionine |
| 146.4888068 | 144.3012323 | 132.0620303 | 123.6139019 | 74.75443838 | 53.12424436 | Branched-c |
| 937.7004455 | 776.2824674 | 793.1030508 | 1441.128437 | 568.4283899 | 289.5052625 | IMP cyclohy |
| 283.39464 | 270.1654173 | 228.5449282 | 247.7175999 | 117.9965978 | 98.89954804 | Dicarboxyla |
| 250.7966309 | 295.187345 | 224.5785448 | 263.0410841 | 265.274192 | 246.9759411 | Trk system p |
| 239.0311713 | 253.323986 | 196.4949492 | 232.681201 | 221.0916033 | 169.7459083 | Potassium u |
| 122.042408 | 121.7684338 | 128.3492704 | 109.3669918 | 104.6859985 | 91.87428447 | rRNA small |
| 817.5929749 | 936.9797382 | 748.4082374 | 674.1082684 | 1235.344812 | 1103.201464 | LemA prote |
| 831.0218991 | 968.8399391 | 746.705615 | 802.1159003 | 1793.503181 | 1736.854157 | Heat shock |
| 602.7171979 | 666.7073274 | 581.7991121 | 532.960539 | 771.4531568 | 832.1268146 | COG1399 p |
| 1175.506909 | 1366.084173 | 1138.63451 | 1008.669178 | 2042.084662 | 2161.051674 | Response re |
| 258.3922628 | 247.1747925 | 326.3794915 | 246.1222603 | 233.4635009 | 260.1712692 | Ribonucleo |
| 282.8800169 | 222.4173081 | 235.2591075 | 369.5275397 | 133.5620652 | 71.54709903 | putative ch |
| 243.8639684 | 233.389438 | 260.6467093 | 262.0867708 | 229.9880737 | 252.6151669 | Helicase loa |
| 623.0318938 | 646.092019 | 586.676033 | 686.332778 | 708.3163593 | 594.6456232 | Helicase loa |
| 714.2125436 | 681.1962169 | 675.8043956 | 871.0721953 | 811.814148 | 788.1715111 | GTP-bindin |

| | | | | | | |
|---|---|---|---|---|---|---|
| 214.6273611 | 217.5028632 | 236.1486398 | 219.0135305 | 190.3507389 | 160.4687103 | SWF/SNF fa |
| 1241.712325 | 1348.319194 | 1327.821943 | 1329.440311 | 1691.14251 | 1611.799481 | FIG011142 |
| 528.2524865 | 513.7071328 | 589.8686133 | 595.5915566 | 453.7929574 | 404.9895713 | UDP-N-acet |
| 313.5050791 | 272.0582204 | 343.9048606 | 392.0654232 | 236.0931637 | 279.6173875 | putative ary |
| 675.352507 | 715.7084226 | 705.1457567 | 822.3317391 | 797.1765434 | 773.8498079 | FIG004453 |
| 756.6617723 | 781.7159057 | 747.9154352 | 794.5980602 | 736.5716469 | 634.5067771 | Transcriptio |
| 1256.155392 | 1286.135558 | 1106.154598 | 1325.886736 | 1260.067994 | 950.6508881 | Inner memb |
| 678.3506551 | 580.1257318 | 639.0578625 | 1077.397515 | 418.032036 | 287.6990423 | Phosphorib |
| 508.1644318 | 550.578048 | 467.2023616 | 506.9845677 | 586.9022605 | 588.4461479 | Predicted n |
| 280.4407744 | 326.8482688 | 263.8889251 | 267.5415606 | 314.9386117 | 248.767832 | FIG011178 |
| 459.6687104 | 506.4041793 | 409.6798271 | 380.6117133 | 382.4472364 | 422.7695601 | FIG005986 |
| 950.0472869 | 1160.137469 | 734.4016671 | 812.1875305 | 773.314865 | 829.3276953 | FIG005935 |
| 510.5170449 | 556.8620709 | 536.019295 | 481.911818 | 722.0868219 | 1084.272516 | FIG027054 |
| 163.2763243 | 202.9480541 | 151.6675078 | 160.1582467 | 253.0164564 | 259.9782229 | Glutamate r |
| 324.068339 | 361.5230446 | 311.0470702 | 306.1660052 | 389.751433 | 405.9206252 | unknown d |
| 223.0001883 | 249.6545875 | 180.5635526 | 190.0190331 | 275.5499561 | 285.4075967 | FIG009886 |
| 186.5541676 | 225.5393441 | 189.7701317 | 191.264544 | 295.5266465 | 356.4043621 | FIG042801 |
| 129.9443537 | 126.4879431 | 132.2655607 | 120.0576818 | 153.9162243 | 144.7181406 | Tyrosine rec |
| 721.5748852 | 592.7159526 | 682.3085823 | 1117.82185 | 473.7248174 | 274.0122524 | Phosphorib |
| 152.0406182 | 148.2683966 | 147.9005533 | 138.5124182 | 172.2759818 | 161.2931255 | Segregation |
| 244.621511 | 248.5569587 | 235.8773616 | 228.4725415 | 280.955133 | 256.8984748 | Segregation |
| 201.9453751 | 231.8319685 | 217.5656424 | 190.3968827 | 217.1121388 | 212.6498934 | Ribosomal |
| 142.7180519 | 183.5951493 | 160.0007278 | 163.6096892 | 179.1582485 | 163.2921853 | Protein Yid |
| 197.2339304 | 181.6393932 | 226.7665971 | 233.9250702 | 205.3500337 | 173.0801069 | tRNA (cytid |
| 128.5538472 | 108.0078758 | 160.7689309 | 129.9771848 | 102.0329984 | 110.8569533 | Substrate-sp |
| 218.3955726 | 192.3412727 | 232.3539918 | 173.8462375 | 260.8356935 | 260.8918423 | Membrane- |
| 137.5856164 | 122.0958122 | 166.1169225 | 140.8130072 | 119.7852481 | 104.8735979 | COG1242: |
| 138.4288737 | 129.0436997 | 168.5196359 | 160.2049515 | 135.2077652 | 164.6908626 | SAM-depen |
| 323.6201642 | 356.495498 | 358.3034432 | 400.4356149 | 475.6669421 | 433.7446907 | Hemolysins |
| 56.39132778 | 49.2920829 | 67.93463509 | 48.84699798 | 59.19882349 | 116.3218924 | FIG011143 |
| 1094.605743 | 919.4045028 | 1068.121351 | 1803.671989 | 748.9903419 | 436.5311673 | Phosphorib |
| 166.6243754 | 165.8944497 | 193.9884094 | 252.7577734 | 548.4505742 | 753.0281929 | Pyruvate fo |
| 763.7841964 | 840.3724537 | 737.6929973 | 723.9545122 | 1287.450776 | 1306.273478 | Manganese- |
| 538.6224882 | 504.0756123 | 563.0131645 | 701.7834755 | 735.186016 | 531.4774584 | FIG011147 |
| 111.6219395 | 98.61611881 | 118.8758523 | 123.1904122 | 80.41695463 | 56.05468774 | ABC-type Fe |

| | | | | | | |
|---|---|---|---|---|---|---|
| 197.2766615 | 182.4566094 | 213.5496541 | 252.8976997 | 183.2077157 | 180.0610366 | ABC-type Fe |
| 100.4743233 | 99.19977129 | 103.3563815 | 104.1925315 | 75.37856625 | 52.60245909 | Ferrichrom |
| 85.92333788 | 77.91920795 | 84.78206506 | 91.65830367 | 68.45494405 | 57.05841877 | Ferrichrom |
| 186.6335389 | 222.5065194 | 153.8552482 | 187.1285929 | 298.8652462 | 258.1402921 | UDP-N-acet |
| 251.5993258 | 274.2136726 | 196.4109526 | 233.1211775 | 297.2806568 | 271.9615789 | Membrane |
| 381.685302 | 455.6983327 | 312.8903121 | 353.879055 | 563.115528 | 468.6545532 | Uracil phos |
| 211.0966637 | 179.8059094 | 255.564252 | 274.9282797 | 253.6579044 | 342.062143 | FIG011146 |
| 101.9289401 | 110.4171965 | 87.67214667 | 97.69399595 | 135.4227435 | 121.6156394 | hypothetica |
| 3612.250982 | 3354.436299 | 4320.682982 | 4780.899927 | 2977.158862 | 2561.666082 | ATP-depend |
| 644.6921594 | 548.3627748 | 771.3566637 | 776.9061933 | 511.9721717 | 363.2841527 | hypothetica |
| 652.7018874 | 534.4288873 | 857.1618232 | 907.5000349 | 549.2827975 | 633.2608405 | Thymidylat |
| 752.772453 | 650.7200819 | 924.8955438 | 1049.123109 | 793.3526642 | 684.4162236 | DNA polym |
| 480.40612 | 370.5558129 | 633.6656035 | 607.7380665 | 370.8603013 | 290.2972183 | hypothetica |
| 646.4605831 | 542.9272899 | 712.5805609 | 927.9541916 | 574.2880842 | 397.8391423 | hypothetica |
| 407.9639273 | 321.893441 | 560.4607395 | 569.5424278 | 302.5056575 | 382.0385828 | DNA replica |
| 1108.189575 | 1058.013681 | 1219.651723 | 1660.98874 | 1674.529702 | 1670.116184 | rRNA small |
| 625.1606526 | 511.6800684 | 614.4784559 | 756.0564448 | 459.2206407 | 358.6866745 | FIG011146 |
| 41.26102303 | 40.11796526 | 50.08282261 | 58.80921467 | 54.76791446 | 36.49282845 | hypothetica |
| 343.2980625 | 338.7988754 | 252.8313469 | 302.3280268 | 326.481724 | 358.5968166 | Cytoplasmi |
| 613.2730819 | 741.3082632 | 531.693968 | 533.0582584 | 620.0314933 | 526.5882702 | Arsenate re |
| 14.7063098 | 11.50658871 | 13.82431423 | 17.99052381 | 9.816890328 | 15.0731248 | FIG011151 |
| 252.1959983 | 287.392204 | 198.8580853 | 182.6452968 | 282.9113973 | 257.481011 | Exodeoxyril |
| 712.6903979 | 631.0459991 | 636.1289772 | 855.9072646 | 2478.092077 | 3013.451755 | Lactate 2-m |
| 48.70051052 | 32.21844839 | 39.91770734 | 54.49493212 | 128.1104188 | 159.7751228 | hypothetica |
| 862.3909988 | 873.1415939 | 777.0533744 | 890.4080895 | 1125.488633 | 1065.056039 | Serine endc |
| 495.2135989 | 525.9216084 | 590.0621928 | 495.9161685 | 367.2953601 | 385.7739582 | FIG011184 |
| 483.189692 | 488.9203121 | 545.7252773 | 470.5964191 | 310.92769 | 185.9932248 | FIG011152 |
| 200.630785 | 182.2190933 | 184.4186181 | 236.7027427 | 124.6423206 | 79.8957981 | Membrane |
| 745.3944171 | 675.1816992 | 701.2284496 | 1148.060287 | 937.3649757 | 774.0568341 | Adenylosuc |
| 1104.277745 | 1155.078453 | 1078.44527 | 1433.890424 | 954.5378136 | 660.169672 | Methionyl- |
| 44.99308067 | 64.65121018 | 44.71956195 | 55.78549201 | 41.05245046 | 26.49215873 | hypothetica |
| 23.90575758 | 26.68808426 | 17.95421943 | 21.77377976 | 22.96781888 | 17.06391486 | FIG011157 |
| 159.2190646 | 181.7140948 | 147.4934354 | 156.1333593 | 101.8865422 | 60.23303912 | Ribonucleo |
| 145.5501317 | 149.7388908 | 123.9806341 | 144.0686765 | 96.96437686 | 59.79930901 | Ribonucleo |
| 272.6805964 | 285.7472747 | 227.3944397 | 322.057597 | 210.0571471 | 147.6212497 | Ribonucleo |

| | | | | | | |
|---|---|---|---|---|---|---|
| 926.9076533 | 1011.166501 | 900.1684771 | 997.9821175 | 1110.364607 | 1245.884844 | C3 family AI |
| 49.39811752 | 49.10309401 | 50.98782564 | 57.23046092 | 47.3184502 | 49.18038954 | Mobile elem |
| 6019.874225 | 5317.829929 | 8616.213304 | 7081.889574 | 4249.027783 | 9268.702968 | FIG011143 |
| 121.9124143 | 123.6837542 | 133.377841 | 122.941078 | 130.6787912 | 141.9911725 | Mobile elem |
| 48.34624921 | 31.38093416 | 63.02795896 | 50.07218995 | 32.3569872 | 86.67046757 | Transcriptio |
| 104.496871 | 100.5381998 | 102.5416249 | 85.84141556 | 111.8355545 | 105.9059422 | hypothetica |
| 14.8351869 | 8.454980281 | 18.27201027 | 16.34674062 | 15.78158319 | 17.04471496 | hypothetica |
| 651.0370222 | 843.0681094 | 511.4996266 | 601.4286626 | 721.3077036 | 953.1957014 | FIG011142 |
| 125.3728152 | 149.4461692 | 129.316664 | 118.7903416 | 101.9310648 | 111.7998638 | FIG011154 |
| 204.6854842 | 234.7500407 | 210.8937051 | 130.5420563 | 150.6084946 | 221.3268957 | FIG011143 |
| 266.3837548 | 309.9419457 | 233.9651464 | 220.3618058 | 337.7347333 | 515.4620531 | 3-oxoacyl-[a |
| 579.4628864 | 744.2866392 | 434.9282861 | 486.3606776 | 760.1842892 | 977.591108 | NAD-depen |
| 136.9274626 | 143.631626 | 127.4357935 | 168.4728579 | 193.4323592 | 187.411438 | Glycerol-3-p |
| 427.2363359 | 515.433809 | 342.2292454 | 309.5320886 | 401.9175446 | 465.2505794 | N-acetylglu |
| 355.0182679 | 380.4198072 | 324.0493982 | 311.1025695 | 274.3422865 | 339.0502522 | ADP-ribose |
| 409.8267039 | 396.6835988 | 527.0537099 | 464.8165911 | 294.5656701 | 372.4978028 | Holliday jur |
| 421.5808808 | 462.3216374 | 369.9924101 | 384.2339751 | 308.9983098 | 327.0628825 | FIG011156 |
| 557.7962874 | 643.6915403 | 478.7761241 | 572.8994892 | 820.9797678 | 965.0264955 | 5'-methylth |
| 370.3767253 | 343.0017098 | 385.2415988 | 437.2127681 | 382.1698182 | 475.9934146 | FIG011162 |
| 850.5358658 | 826.4890588 | 868.3930722 | 948.615485 | 824.8914789 | 625.9068549 | Mn-depend |
| 752.615319 | 809.9209039 | 823.1690132 | 747.0763533 | 1035.539885 | 1082.421609 | Manganese |
| 222.8384577 | 244.3172805 | 237.4602075 | 183.7817837 | 207.6961921 | 207.9509889 | Manganese |
| 277.7408662 | 291.9374681 | 281.8219849 | 223.0893815 | 246.0767175 | 353.7159952 | Manganese |
| 919.6384977 | 947.0687249 | 1051.401624 | 1177.435096 | 956.9460899 | 936.887162 | Peptidyl-pr |
| 145.5002176 | 143.504469 | 147.5081908 | 165.9320949 | 151.2682427 | 144.7170336 | Cell divisior |
| 204.2639716 | 186.7152106 | 193.1268514 | 210.4178374 | 121.8301261 | 75.23679385 | Integral me |
| 477.9473201 | 472.6735946 | 595.8658182 | 577.0051636 | 502.2751421 | 471.4653189 | Low molecu |
| 2933.518442 | 2970.051743 | 2717.629773 | 3316.263972 | 2452.010325 | 1969.838037 | LSU ribosor |
| 4187.383168 | 4680.045663 | 3491.861493 | 4706.116235 | 6731.911201 | 6688.62267 | LSU ribosor |
| 658.464663 | 754.2538294 | 682.7011477 | 686.7224561 | 849.0196178 | 857.4933217 | Uridylate ki |
| 883.9234194 | 990.9728893 | 919.4036081 | 821.4371172 | 967.3331501 | 847.660674 | Ribosome r |
| 542.2427642 | 578.3714298 | 513.4008726 | 539.5372958 | 636.1070334 | 700.1800255 | S1 RNA binc |
| 226.9429816 | 181.8470243 | 262.3231392 | 295.2370025 | 120.2280333 | 140.8007187 | Peptide me |
| 386.4805645 | 376.10018 | 379.6886305 | 497.3743804 | 298.3243894 | 267.4083992 | FIG011143 |
| 1448.453796 | 1449.9299 | 1306.35676 | 1733.804977 | 1325.629018 | 1088.325518 | Group B str |

| | | | | | | |
|---|---|---|---|---|---|---|
| 730.0018473 | 759.6485466 | 663.2551979 | 848.5926127 | 1597.412367 | 1825.233842 | 67kDa Myo |
| 112.6678014 | 148.8348585 | 97.91894368 | 92.84060833 | 186.4370112 | 159.6348411 | Phosphate s |
| 311.3778286 | 246.690036 | 452.7462911 | 382.380406 | 228.9439066 | 424.0213559 | MORN moti |
| 182.2144779 | 210.1238745 | 142.9712498 | 161.5194679 | 218.0167727 | 185.2738256 | Hypothetic |
| 251.5882971 | 336.9540341 | 241.8283402 | 247.2879272 | 297.2270529 | 324.9499354 | Metal-depe |
| 288.5697395 | 320.2199443 | 238.0119983 | 244.8635358 | 302.7355738 | 302.2013746 | Diacylglyce |
| 560.5120617 | 677.0631107 | 453.2124957 | 487.6531403 | 582.3156082 | 496.052 | GTP-bindin |
| 436.4792905 | 565.5896968 | 340.5215515 | 473.608511 | 580.8635484 | 547.562068 | MutT/nudix |
| 7.654822792 | 2.881324653 | 9.408213852 | 6.105874747 | 8.507971617 | 4.689416603 | FIG011157 |
| 14.3494711 | 10.75882055 | 16.02012253 | 12.90839292 | 13.04526366 | 14.43243269 | Competenc |
| 24.65848634 | 22.54661408 | 24.90709779 | 19.70883621 | 19.63378066 | 18.23190904 | Peptide AB( |
| 19.24564787 | 24.46970764 | 26.54035011 | 24.11434077 | 21.12495387 | 16.02712004 | Peptide AB( |
| 35.26497672 | 30.80397992 | 34.40718209 | 24.179264 | 9.424214715 | 15.54206646 | Bacteriocin |
| 404.1521712 | 356.7497606 | 435.1117281 | 475.5156239 | 349.0302494 | 332.4574575 | Acyltransfe |
| 56.10255863 | 50.87824425 | 44.83139192 | 27.01073405 | 19.9665566 | 38.8324571 | hypothetica |
| 53.77405586 | 57.38282923 | 46.88478664 | 33.65330965 | 34.24496626 | 41.12976688 | hypothetica |
| 45.25018399 | 43.06678879 | 51.52698941 | 30.21218544 | 31.61803638 | 71.77678474 | FIG011191 |
| 115.4946913 | 161.4629098 | 122.3321391 | 96.42673893 | 120.5810491 | 234.1548317 | putative rol |
| 34.70785389 | 47.81772828 | 34.14409526 | 37.99932688 | 45.74253153 | 98.99087632 | Mobile elen |
| 22.17259016 | 19.35588427 | 21.56107956 | 21.78833199 | 24.90390073 | 14.91446032 | Mobile elen |
| 9.766498045 | 6.548465119 | 8.601795522 | 4.57940606 | 7.853512262 | 11.38858318 | Mobile elen |
| 3.455468596 | 2.571760847 | 1.759458175 | 2.331333994 | 9.281423583 | 5.846545375 | hypothetica |
| 875.8838253 | 981.0834625 | 1141.5808 | 1101.843563 | 1632.991234 | 1574.74518 | Alcohol deh |
| 62.52670424 | 57.58458287 | 70.51321879 | 62.13338142 | 41.85922036 | 39.47148946 | Response re |
| 32.35133941 | 25.6052344 | 37.29458255 | 34.98872046 | 29.64921483 | 38.02717926 | Histidine ki |
| 0.754169733 | 0.374198007 | 2.304052372 | 2.180669552 | 0.467470968 | 2.392559491 | FIG011162 |
| 10.05946398 | 11.43318833 | 12.72545849 | 9.903594644 | 8.716535588 | 7.288104297 | Competenc |
| 19.03595832 | 18.0365524 | 21.87887057 | 19.5677128 | 14.60228255 | 12.12007992 | Competenc |
| 17.29628743 | 17.16387173 | 19.0993815 | 20.00477384 | 9.04187267 | 14.54424322 | Bacteriocin |
| 4.018486489 | 3.987722043 | 6.980934799 | 3.280768521 | 2.344332019 | 3.899514872 | FIG011160 |
| 7.734624473 | 8.040906007 | 9.376957328 | 6.815860183 | 3.652796401 | 9.347674292 | hypothetica |
| 9.603629475 | 6.186209602 | 8.064183302 | 5.84605029 | 2.924180098 | 6.414095658 | unknown |
| 27.31979858 | 20.82411908 | 30.24068738 | 31.59790182 | 23.06969227 | 27.63406213 | hypothetica |
| 575.8052542 | 636.1167427 | 886.7271291 | 1190.969781 | 1773.063602 | 2924.571561 | Alcohol deh |
| 18.17599672 | 16.03275219 | 21.54057016 | 23.48098141 | 14.09942638 | 17.13008366 | FIG011140 |

| | | | | | | |
|---|---|---|---|---|---|---|
| 23.7563466 | 23.87671128 | 18.29949288 | 28.18096037 | 27.18523475 | 23.96240352 | hypothetica |
| 19.17178848 | 16.9570781 | 21.221535 | 15.42536778 | 32.03406317 | 17.45309187 | hypothetica |
| 9.502538638 | 10.37276875 | 9.999587294 | 12.82233697 | 13.74364646 | 7.235099902 | hypothetica |
| 57.21251637 | 55.57811503 | 54.35650198 | 48.93150835 | 49.4581049 | 44.99336444 | Positive tra |
| 132.4387631 | 139.6816082 | 128.2088273 | 131.5417799 | 109.3373944 | 88.83677416 | Formamido |
| 248.8417315 | 312.1831917 | 240.7851095 | 257.4643852 | 256.0324326 | 250.2037214 | Dephospho |
| 706.2415692 | 788.2312961 | 680.3852858 | 740.6755127 | 959.1160634 | 964.3590688 | ATPase |
| 654.9111428 | 743.0500592 | 611.7837955 | 721.7512144 | 668.7324889 | 672.2563163 | Multidrug r |
| 103.3493156 | 96.78326867 | 111.8858548 | 141.173504 | 124.0124378 | 146.4780562 | Multi antim |
| 1007.188905 | 934.822408 | 994.2423463 | 917.5042927 | 665.8088402 | 1248.58897 | Preprotein |
| 1194.977058 | 1165.124629 | 1283.667562 | 1485.720673 | 1056.841652 | 1012.528697 | 3'-to-5' exo |
| 868.0203564 | 794.7821743 | 982.4863323 | 1055.846649 | 478.3846171 | 410.8392733 | tmRNA-bin |
| 822.484146 | 782.6990765 | 808.5279478 | 1045.263925 | 830.9148522 | 773.8809389 | Glutaminyl |
| 1025.775583 | 1086.02656 | 762.886674 | 829.3813198 | 1005.413184 | 813.2974311 | Pyrrolidone |
| 640.4219001 | 648.6413644 | 629.5804465 | 669.2157283 | 543.646043 | 409.6247679 | FIG001614 |
| 404.6914481 | 406.7630807 | 416.8616859 | 412.94995 | 348.930172 | 304.2831976 | hypothetica |
| 283.9998198 | 228.1663264 | 275.9046792 | 289.5844479 | 182.9309207 | 146.5361388 | Bactopreno |
| 524.3993545 | 496.8102204 | 449.5462184 | 454.4515347 | 515.7763013 | 424.7590617 | Lactoylglut |
| 825.9789216 | 883.9365998 | 725.1226445 | 762.7392724 | 793.2224266 | 771.2189197 | Oxygen-inse |
| 2041.815705 | 2280.086546 | 1421.488466 | 1313.178033 | 2529.707797 | 2211.700577 | SSU riboson |
| 857.8534881 | 889.2309121 | 738.2069234 | 754.2104677 | 709.094056 | 603.6467252 | Proline dip |
| 419.8377101 | 401.0483943 | 383.9420445 | 436.2775474 | 477.2066807 | 554.4864072 | Catabolite |
| 947.0197666 | 1177.402229 | 1101.669149 | 1189.325387 | 1094.30321 | 886.8851606 | Glycosyltra |
| 850.2458729 | 1036.747111 | 881.5511481 | 1115.810877 | 1301.741718 | 891.6515441 | Glycosyltra |
| 98.68020893 | 143.0587765 | 109.5901833 | 71.86144894 | 101.6929152 | 112.3398703 | hypothetica |
| 267.7232722 | 320.4382265 | 163.797501 | 112.9021469 | 126.2864163 | 112.2420548 | Threonyl-tR |
| 15.8375644 | 16.14364815 | 17.34652118 | 15.99333355 | 12.21921092 | 13.11498472 | Daunorubic |
| 22.19909654 | 21.3715598 | 19.36753647 | 21.91983487 | 20.53742746 | 17.57480102 | Daunorubic |
| 15.5353208 | 18.17573983 | 16.86707042 | 21.18412269 | 20.53303778 | 19.9440837 | ABC-type m |
| 3.167512879 | 3.143263257 | 2.580538657 | 5.128934787 | 5.10478297 | 3.617549951 | Hypothetic |
| 2302.130174 | 2766.974039 | 1352.544887 | 1300.989542 | 2929.378862 | 2597.289319 | LSU riboson |
| 131.2905143 | 146.2889492 | 133.4906583 | 142.6192049 | 114.4705802 | 84.27306674 | 3-ketoacyl- |
| 87.06853311 | 91.76666409 | 89.21002778 | 96.91608402 | 66.16961654 | 45.17106309 | Long-chain- |
| 158.5187545 | 170.2285342 | 158.4174804 | 173.0243061 | 226.616408 | 186.144011 | FIG011142 |
| 312.2740018 | 359.0216725 | 281.7360242 | 265.5669067 | 328.9693797 | 334.6191305 | Two-compo |

| | | | | | | |
|---|---|---|---|---|---|---|
| 387.2302053 | 415.7331559 | 336.5142566 | 345.5573105 | 396.5936625 | 374.499751 | Two-compo |
| 206.4749136 | 223.1134827 | 203.9939702 | 210.8562079 | 285.9987382 | 357.4377544 | Zn-depende |
| 54.43734256 | 71.3697653 | 49.29275681 | 40.12431977 | 82.78485869 | 96.57240129 | Ribonuclea |
| 85.96576268 | 108.2827554 | 78.20890992 | 62.09364149 | 92.79457179 | 105.0860791 | Chromosom |
| 31.39629843 | 26.56278809 | 38.90116593 | 37.21573657 | 32.00859311 | 38.77968257 | Positive tra |
| 3.700874155 | 3.618533303 | 5.320698261 | 4.02861839 | 6.274713405 | 7.182611586 | Shikimate 5 |
| 3322.385985 | 3927.567887 | 2058.615408 | 1844.972249 | 2882.578816 | 2701.85762 | LSU riboson |
| 6.852170719 | 5.131858379 | 5.398065557 | 4.261651354 | 8.254201663 | 10.00773456 | FIG011141 |
| 11.89433408 | 16.87098442 | 15.00925545 | 11.8877643 | 15.14605936 | 20.50765278 | putative L-g |
| 14.39340959 | 15.76408625 | 15.54009791 | 12.7499573 | 15.6950891 | 16.73773534 | FIG011156 |
| 18.7279199 | 19.13461508 | 18.95083076 | 13.78401224 | 23.9532124 | 28.13649962 | Archaeal S-a |
| 7.522843088 | 6.384753491 | 8.215386739 | 7.784990302 | 10.5531571 | 15.5127576 | FIG011161 |
| 7.102907669 | 9.667915776 | 12.12071187 | 8.825764407 | 16.6589654 | 21.13282547 | Dolichyl-ph |
| 10.79087937 | 10.83041485 | 13.45423328 | 11.91120125 | 15.41200917 | 15.09915783 | UDP-glucos |
| 17.64532609 | 18.64118904 | 18.60965377 | 20.68118866 | 30.35197357 | 43.98509866 | ABC transpo |
| 1.58375644 | 0.673556412 | 2.304052372 | 1.831762424 | 1.963378066 | 2.87107139 | FIG011153 |
| 2.390575758 | 0.889602809 | 2.738779235 | 4.493002172 | 3.704486916 | 5.687971621 | hypothetica |
| 3204.787147 | 3773.820917 | 1860.62702 | 1556.813034 | 2604.153271 | 2623.028222 | LSU riboson |
| 3.483454062 | 3.577371226 | 3.506166653 | 3.045129349 | 3.113284912 | 4.934424486 | FIG011160 |
| 1.900507728 | 1.676407071 | 2.365493769 | 1.709644929 | 1.570702452 | 2.143733304 | FIG011146 |
| 2.201153018 | 1.70482075 | 2.460259312 | 1.614434679 | 4.592308018 | 4.76889824 | FIG011167 |
| 2.147466359 | 2.930160664 | 2.733621458 | 4.346554905 | 4.99163915 | 4.76889824 | FIG011151 |
| 744.3655266 | 780.5251732 | 999.7990426 | 968.6577334 | 1276.390136 | 1113.520876 | Plasmid sta |
| 1364.670132 | 1403.220514 | 1758.466324 | 1752.206468 | 2162.025729 | 1395.988094 | Plasmid sta |
| 1486.211441 | 1521.625168 | 2178.795707 | 2151.071919 | 1899.568278 | 2094.707585 | hypothetica |
| 4.067234437 | 2.323813303 | 7.970048866 | 5.488159792 | 12.91092969 | 22.67811253 | FIG011179 |
| 14.77384739 | 16.302746 | 15.34602046 | 15.44695179 | 15.45793925 | 13.64830205 | Mobile elen |
| 19.68594453 | 20.12276011 | 22.60985972 | 24.30937049 | 14.12898234 | 17.65574742 | Mobile elen |
| 3491.784162 | 3950.731407 | 1842.752908 | 1859.107079 | 2329.216137 | 2373.746056 | LSU riboson |
| 231.8476532 | 281.9483508 | 176.1387405 | 144.9571392 | 203.4236823 | 223.3391323 | Hydrolase ( |
| 271.6574228 | 309.4685552 | 218.1728137 | 229.3366555 | 259.8798603 | 314.1787321 | Hydrolase ( |
| 513.97032 | 624.5791768 | 449.4417363 | 422.0139697 | 511.4277062 | 580.0674873 | Signal recog |
| 45.42880314 | 42.53055767 | 43.04434683 | 48.08376363 | 33.19658966 | 27.85442945 | Multidrug r |
| 44.41281404 | 36.13074847 | 38.45553603 | 36.43968666 | 26.41127789 | 16.59295351 | Beta-glucos |
| 75.89789315 | 75.21560709 | 72.82437996 | 73.00502415 | 46.88711226 | 37.83249144 | PTS system, |

| | | | | | | |
|---|---|---|---|---|---|---|
| 106.0950103 | 91.75020013 | 111.1329345 | 94.5575045 | 68.03621676 | 52.25349929 | 6-phospho- |
| 401.7617439 | 427.0562267 | 387.6737531 | 402.3142912 | 282.452146 | 295.4775787 | ABC transpo |
| 352.7409692 | 383.7183003 | 311.7436442 | 288.5177958 | 287.201117 | 225.1454122 | possible pe |
| 1172.588902 | 1054.806612 | 1064.162035 | 569.6076615 | 751.3696828 | 1012.218304 | FIG011164 |
| 2685.233499 | 2815.41752 | 1692.611377 | 1265.294819 | 1279.995829 | 1552.261726 | SSU riboson |
| 273.2035546 | 339.834946 | 218.1072361 | 258.5077941 | 221.2735364 | 154.0242982 | Transcriptio |
| 262.4045772 | 317.0174758 | 199.9475586 | 249.7971908 | 349.9116251 | 311.2359376 | Putative me |
| 2760.609302 | 3473.133193 | 2662.598368 | 1859.943384 | 2066.725109 | 2793.33161 | Phage shock |
| 605.7995693 | 707.891185 | 609.6626296 | 654.0746534 | 1147.408242 | 1196.996744 | HPr kinase/ |
| 486.4568818 | 511.6869899 | 465.6755696 | 539.1017719 | 809.4403648 | 799.8031911 | Prolipoprot |
| 500.0478053 | 521.7585885 | 509.2294573 | 440.8351775 | 580.2070916 | 527.2638164 | General stre |
| 712.4688935 | 713.4987786 | 720.701137 | 705.5487715 | 931.8494357 | 1306.056398 | FIG011153 |
| 590.4910852 | 498.6208136 | 454.1408491 | 458.3262402 | 384.8221008 | 411.6814155 | FIG011148 |
| 122.7539376 | 117.5926319 | 127.3566813 | 106.1710842 | 111.0039314 | 102.5037527 | PROTEASE ( |
| 268.9801729 | 306.8894674 | 259.2193187 | 264.2605511 | 302.9277253 | 279.0226302 | peptidase, U |
| 4017.645793 | 4639.729895 | 2321.082324 | 1884.166758 | 2084.59532 | 2239.385752 | LSU riboson |
| 48.98595499 | 57.74832496 | 43.88416029 | 48.98899506 | 60.27114062 | 80.85738262 | FIG011140 |
| 295.1140521 | 223.12742 | 290.3105989 | 311.9156015 | 274.0433328 | 551.6905207 | bacterial se |
| 152.9841327 | 123.724192 | 164.7152334 | 157.4536211 | 140.7783847 | 121.0126047 | FIG011165 |
| 1010.45569 | 1101.2467 | 899.4641078 | 966.8983703 | 1034.834286 | 826.4592308 | Lysyl-tRNA s |
| 114.1251733 | 113.4081029 | 103.3072784 | 106.3152477 | 152.6347068 | 138.5458868 | 2-haloalkan |
| 143.0710745 | 106.462931 | 143.2943341 | 137.5583128 | 119.2185816 | 96.62259484 | Phosphogly |
| 304.7741299 | 218.0638885 | 357.0417157 | 259.9957791 | 181.9806044 | 120.9618265 | Transcriptio |
| 700.5594975 | 549.3186932 | 723.4601892 | 785.6117247 | 574.0792142 | 436.4792095 | Substrate-s |
| 197.1580896 | 145.1121359 | 262.0432143 | 194.697575 | 99.00143108 | 92.10762243 | Phage lysin, |
| 265.2792036 | 313.9521277 | 192.3883731 | 155.917873 | 369.3020647 | 467.6257526 | Substrate-s |
| 3329.012447 | 3844.081196 | 2001.249159 | 1891.756845 | 2228.524168 | 2752.343369 | SSU riboson |
| 311.8020491 | 361.9664095 | 274.9382494 | 180.8865394 | 353.8682185 | 441.6739589 | Additional |
| 251.4213348 | 279.455749 | 199.3869321 | 185.12249 | 181.7351822 | 185.6506537 | Glutathione |
| 270.5583918 | 314.3001319 | 198.7014766 | 160.4929177 | 198.1702927 | 180.8774975 | Oligoendop |
| 302.560287 | 299.3006514 | 282.4106379 | 286.8190488 | 265.9293528 | 245.5643054 | Phosphoen |
| 143.5442307 | 165.446586 | 120.2986403 | 130.0766823 | 108.9790319 | 133.3528217 | Cell division |
| 89.29690564 | 76.90963289 | 85.44602733 | 90.41891115 | 68.9271023 | 51.74037164 | FIG011145 |
| 9377.284823 | 10152.5892 | 6887.00639 | 5815.537467 | 7100.207519 | 7335.249221 | Translation |
| 1887.612319 | 2133.194867 | 1575.862539 | 1344.571541 | 1476.568951 | 1236.592009 | Triosephos |

| | | | | | | |
|---|---|---|---|---|---|---|
| 0.833556021 | 0.827174541 | 0.4244307 | 0.482042743 | 0.516678438 | 0.528881572 | hypothetica |
| 223.7632581 | 234.0281564 | 208.4552237 | 182.6871796 | 216.5434449 | 172.7799612 | tRNA-deper |
| 232.6248706 | 244.4478657 | 284.0240044 | 242.0683892 | 288.9649169 | 330.2040815 | GTP-bindin |
| 3082.242515 | 3861.43058 | 2136.891703 | 1707.653883 | 1919.130922 | 2521.216777 | LSU riboson |
| 292.2822509 | 328.2745564 | 268.6179458 | 226.2226594 | 225.6903086 | 234.9397718 | tRNA-deper |
| 321.0332961 | 356.9350054 | 267.1335387 | 226.4601101 | 248.185531 | 204.8456269 | Promiscuou |
| 129.245838 | 131.9047974 | 123.5548084 | 112.8496493 | 102.376142 | 68.90571335 | FIG011142 |
| 216.7629781 | 225.6776108 | 185.9964858 | 194.783032 | 171.1396134 | 142.3032642 | Deoxyguano |
| 207.7299144 | 243.1764908 | 192.1651122 | 184.0282249 | 262.6969412 | 266.7203065 | FIG011143 |
| 775.7749245 | 854.0956493 | 723.2508022 | 986.9798207 | 787.8109392 | 619.3357019 | Calcium-tra |
| 169.7767822 | 151.2458751 | 187.7108691 | 206.2387934 | 240.9869162 | 188.323258 | Transcriptio |
| 22.58596055 | 19.91721844 | 23.38105976 | 26.23798063 | 21.20818759 | 17.37991329 | Heparinase |
| 49.76694506 | 42.84703747 | 50.15097937 | 74.54069718 | 71.65613378 | 65.13350277 | PTS system, |
| 47.2699719 | 41.91017676 | 48.2615108 | 61.55002475 | 67.55224149 | 67.45367725 | PTS system, |
| 2693.533597 | 3451.895266 | 1997.580014 | 1382.847894 | 2415.524116 | 2659.277862 | LSU riboson |
| 51.593538 | 42.81008854 | 53.2335045 | 61.58317843 | 58.29907876 | 63.25164025 | PTS system, |
| 60.81624728 | 52.61036877 | 61.36843493 | 79.72745951 | 61.94457797 | 61.95054291 | Unsaturated |
| 50.33308137 | 43.5966308 | 51.47821121 | 60.97510535 | 40.61234081 | 54.92398898 | PTS system, |
| 60.54133107 | 52.60517942 | 59.03590795 | 64.97572372 | 55.04867557 | 69.39325378 | 2-deoxy-D-g |
| 92.06509397 | 84.08963621 | 98.12679121 | 108.7929926 | 86.3335869 | 102.8353374 | predicted 4 |
| 125.752158 | 115.1902463 | 119.900402 | 151.4220375 | 104.3999834 | 111.9202081 | 2-dehydro-3 |
| 112.4317661 | 98.96831247 | 109.0947062 | 162.4393093 | 127.0639012 | 147.508064 | 4-hydroxy-2 |
| 54.88409413 | 54.75361803 | 48.01347846 | 40.34942114 | 43.33908264 | 53.62420434 | Beta-phosp |
| 148.3963623 | 158.0852937 | 156.2229796 | 156.9147505 | 189.2255697 | 163.7023383 | Epoxyqueuo |
| 468.6138507 | 530.4024257 | 486.2368512 | 466.015535 | 447.801228 | 418.9555712 | Peptide cha |
| 3286.203592 | 3894.756082 | 2238.320671 | 1660.376836 | 2930.398182 | 3282.591622 | SSU riboson |
| 440.5722459 | 457.3380003 | 415.9861828 | 468.8043052 | 317.030311 | 322.0820086 | Cell divisior |
| 485.5490711 | 540.2356979 | 442.7496768 | 525.6567266 | 392.0422653 | 383.2139384 | Cell divisior |
| 239.6552197 | 226.8487162 | 205.1041338 | 248.9296011 | 281.3557813 | 253.6835343 | Hydroxyacy |
| 452.3953689 | 487.1441723 | 525.5950058 | 615.0411716 | 1315.155323 | 1698.041693 | 2,3-butaneo |
| 50.73886372 | 47.98049999 | 50.21980818 | 57.90889145 | 46.85097924 | 53.80600456 | Mobile elem |
| 114.7450855 | 121.9356149 | 100.3499103 | 113.9266874 | 142.7280079 | 115.830224 | DinG family |
| 502.2656729 | 556.6656048 | 459.0911187 | 538.2067786 | 522.1697694 | 471.0288378 | Aspartate ar |
| 452.0584083 | 499.1068016 | 383.3450521 | 439.4597958 | 448.1661869 | 357.5475007 | Asparaginyl |
| 499.0165913 | 572.7724158 | 470.2749319 | 464.9099378 | 410.3922906 | 449.4531757 | FIG000506 |

| | | | | | | |
|---|---|---|---|---|---|---|
| 547.3676014 | 624.6512581 | 494.5372778 | 575.6566268 | 626.6548703 | 646.0791323 | FIG002813 |
| 4129.097684 | 5035.609958 | 2227.943715 | 1845.314488 | 2252.936424 | 2408.75887 | LSU ribosor |
| 564.1090371 | 606.7842245 | 468.6776006 | 611.2301983 | 532.5663003 | 411.8004138 | FIG001886 |
| 158.375644 | 145.9661307 | 142.4210914 | 131.0142523 | 127.1982485 | 112.1751004 | FIG011171 |
| 380.7154046 | 365.0388579 | 424.0260104 | 418.9979049 | 287.3533169 | 255.0746622 | Candidate z |
| 139.9751542 | 112.81837 | 156.3983102 | 133.9238752 | 77.88337887 | 52.78720883 | Transcriptio |
| 223.2304702 | 183.6844466 | 252.9331093 | 240.4646122 | 151.3273644 | 114.053311 | Galactosam |
| 2275.512126 | 2082.863526 | 1548.13781 | 1306.236101 | 1627.347038 | 2111.854512 | LSU ribosor |
| 424.1340096 | 463.2309148 | 360.4227658 | 443.0648284 | 483.9039124 | 469.3470239 | FIG146085 |
| 302.3918962 | 300.7055183 | 418.2623073 | 385.5249315 | 273.4331186 | 379.0388449 | Flavodoxin |
| 330.0881843 | 247.9869275 | 281.312668 | 351.5055683 | 375.1085462 | 246.8819177 | Chorismate |
| 422.9494929 | 353.2998037 | 433.2056284 | 599.6955224 | 358.7245625 | 275.0636424 | Chloride ch |
| 3402.592021 | 4113.822829 | 2340.036249 | 1906.290013 | 1907.364044 | 2358.303669 | LSU ribosor |
| 6078.621052 | 5612.621711 | 6698.972684 | 8094.968719 | 4860.206996 | 4826.691766 | LSU ribosor |
| 377.8509443 | 374.4619149 | 433.7681755 | 416.0028874 | 512.5450108 | 518.6212693 | hydrolase, I |
| 787.158577 | 781.7117072 | 813.0827397 | 943.4279926 | 934.7911389 | 943.7798221 | DNA gyrase |
| 781.6595479 | 719.1513006 | 831.7188288 | 897.4361608 | 857.262082 | 765.9419287 | Septation ri |
| 268.3069733 | 245.7086834 | 280.7179102 | 326.4847144 | 228.0341714 | 186.0003896 | FIG011140 |
| 5506.169005 | 6467.732758 | 3709.746269 | 3256.167773 | 4093.192951 | 3931.504241 | Enolase (EC |
| 106.9160499 | 89.38964752 | 131.0111798 | 151.2793068 | 278.7749108 | 267.9877961 | Streptolysir |
| 89.73124304 | 87.79766492 | 111.9847777 | 145.8145159 | 260.1336887 | 233.5409175 | Streptolysir |
| 103.8356871 | 110.1876833 | 151.5283185 | 178.849563 | 342.6159737 | 356.2536927 | Streptolysir |
| 63.03664245 | 69.71197719 | 83.35650859 | 97.02899969 | 168.2478926 | 191.9211731 | Streptolysir |
| 2560.610412 | 2773.886409 | 1812.525442 | 1627.232487 | 1341.171624 | 1848.747903 | LSU ribosor |
| 87.8012342 | 99.26094497 | 99.52899917 | 137.4625223 | 239.5665692 | 283.3042286 | Streptolysir |
| 73.42870766 | 81.54114749 | 89.43912389 | 133.040667 | 257.7252441 | 277.1236407 | Export ABC |
| 39.50966863 | 42.21670066 | 53.96139678 | 76.24223919 | 140.3084272 | 162.062817 | Streptolysir |
| 72.47914859 | 73.77820327 | 80.03647878 | 126.6027756 | 241.3428534 | 302.5401098 | Streptolysir |
| 3.688199928 | 4.090548075 | 6.407159336 | 8.531496222 | 8.87554468 | 7.708630032 | Asparagine- |
| 340.5163269 | 351.8488152 | 346.2553173 | 429.6096153 | 640.6948752 | 665.5780096 | Endonuclea |
| 514.673425 | 551.2944959 | 424.5526922 | 589.3448805 | 689.0634037 | 850.5935813 | Substrate-s |
| 144.696645 | 165.8275945 | 136.1896071 | 122.6130713 | 163.2640566 | 148.3460164 | DNA ligase ( |
| 203.0552245 | 262.2458641 | 200.6822859 | 191.9321742 | 258.118002 | 305.3523346 | Transcriptio |
| 1080.553825 | 1276.355383 | 697.429671 | 596.1554071 | 902.2614656 | 841.0499128 | ATP synthas |
| 2187.116554 | 2612.20386 | 1329.28957 | 1215.167556 | 1229.328008 | 1418.494497 | SSU ribosor |

| | | | | | | |
|---|---|---|---|---|---|---|
| 1006.182334 | 1223.571118 | 612.2705866 | 576.1237716 | 1090.619548 | 1090.983102 | ATP synthas |
| 1053.629966 | 1230.44469 | 737.897209 | 578.9479419 | 826.7606545 | 1041.903107 | ATP synthas |
| 700.657388 | 787.659628 | 384.0173099 | 351.3095197 | 839.8651312 | 697.3495576 | ATP synthas |
| 1987.944937 | 2545.230864 | 1201.098379 | 1074.366899 | 1715.789456 | 1859.47821 | ATP synthas |
| 1301.826098 | 1704.466795 | 904.3484466 | 736.8452546 | 1311.025532 | 1532.572008 | ATP synthas |
| 2616.338623 | 3387.419079 | 1593.166243 | 1233.686134 | 2045.597138 | 2325.225011 | ATP synthas |
| 2133.855439 | 2721.749393 | 1698.351813 | 1321.504575 | 1988.238104 | 2865.70104 | ATP synthas |
| 225.3157495 | 208.5031294 | 181.0677957 | 157.7758035 | 154.4524078 | 224.8240303 | FIG011148 |
| 953.0553198 | 1111.113908 | 893.2224165 | 940.2039065 | 1017.900392 | 925.3381832 | UDP-N-acet |
| 234.2953971 | 276.4075944 | 261.6379471 | 258.7727869 | 218.1531184 | 277.2178931 | Competenc |
| 2060.074166 | 2625.924666 | 1349.447094 | 1142.992207 | 1450.094943 | 1319.786185 | SSU riboson |
| 612.429453 | 657.3576589 | 635.8042041 | 813.3933475 | 578.9531354 | 376.2052635 | DNA-entry n |
| 138.9746276 | 128.4808856 | 123.7852137 | 156.1577467 | 86.38863488 | 65.81931161 | hypothetica |
| 448.0446968 | 463.3170041 | 484.4961328 | 615.105822 | 438.6186598 | 322.1629206 | DNA-entry n |
| 152.4138022 | 196.9055719 | 119.6650649 | 91.53548435 | 106.462483 | 115.0408605 | Phenylalany |
| 193.2301342 | 233.0208441 | 141.8572295 | 123.6325437 | 138.5381106 | 140.005899 | Phenylalany |
| 48.64394779 | 43.65643413 | 47.87308817 | 51.02766753 | 68.56240864 | 66.67265783 | Salt-stress in |
| 74.39972112 | 83.4548578 | 70.13964081 | 80.65434549 | 132.1094698 | 132.1137967 | FIG011152 |
| 285.7820116 | 330.8306467 | 252.4830649 | 216.5969774 | 554.941427 | 639.1452768 | ABC transpo |
| 315.2598581 | 308.5472319 | 236.9351076 | 248.4789835 | 512.8554812 | 509.8276052 | ABC transpo |
| 185.0278559 | 201.1078142 | 166.6075929 | 185.5474559 | 222.5161808 | 251.4463882 | YpfJ protein |
| 2107.192367 | 2327.683023 | 1317.30011 | 1161.685309 | 1048.707133 | 1034.515992 | LSU riboson |
| 187.1403248 | 219.1487834 | 187.9586604 | 172.5820941 | 177.3816844 | 139.3139184 | ATP-depend |
| 195.360177 | 226.7107538 | 190.5204516 | 183.1612648 | 199.2595957 | 191.3124218 | ATP-depend |
| 278.7060687 | 307.0771024 | 293.6433979 | 269.0189833 | 257.1952816 | 224.5580412 | Amino acid |
| 2582.865163 | 2257.022846 | 2246.216752 | 1657.900228 | 1768.371363 | 4778.436037 | SSU riboson |
| 1040.698137 | 782.1789808 | 1115.523141 | 972.347938 | 1079.695673 | 2099.773484 | Large-condu |
| 493.9141098 | 445.225309 | 566.3329803 | 552.0796715 | 628.1492106 | 564.9218338 | DNA primas |
| 961.4685716 | 1025.21354 | 1124.844595 | 1243.964714 | 1599.675545 | 1473.092412 | RNA polyme |
| 590.3347012 | 539.9180515 | 606.597859 | 703.0401445 | 777.8799646 | 672.288044 | probably an |
| 453.0654826 | 482.2427576 | 423.2988849 | 501.3887253 | 707.9183509 | 700.3802361 | dTDP-4-deh |
| 117.9384341 | 116.6681349 | 117.2553198 | 115.9957608 | 176.3980449 | 188.7076922 | Alpha-D-Glo |
| 2602.817201 | 2966.949961 | 1608.906218 | 1566.233837 | 1540.344338 | 1828.703589 | LSU riboson |
| 197.4857708 | 219.5230802 | 210.9133986 | 217.6911228 | 297.4738728 | 271.2839352 | Alpha-L-Rha |
| 107.8490859 | 113.2982204 | 115.1252437 | 97.4661648 | 138.9749325 | 142.7822369 | rhamnose-c |

| | | | | | | |
|---|---|---|---|---|---|---|
| 159.2029794 | 167.0542773 | 162.2866739 | 140.5262516 | 227.4490461 | 266.9167936 | ABC-transp |
| 112.3241546 | 100.6592638 | 110.0185008 | 90.00713578 | 116.4587049 | 125.0710474 | putative gly |
| 114.3189141 | 118.2234239 | 117.5264652 | 88.18897444 | 122.2894414 | 131.9804879 | Alpha-L-Rha |
| 211.0921083 | 111.8103644 | 304.8261288 | 359.8104762 | 175.0211304 | 364.9131736 | putative rep |
| 512.6199415 | 224.5188041 | 482.2052464 | 572.7061293 | 306.9280813 | 711.2053985 | hypothetica |
| 781.3198436 | 394.4046992 | 921.6209488 | 959.2765361 | 600.7001939 | 835.0032625 | conserved h |
| 566.5888663 | 300.378095 | 1048.142225 | 788.8026939 | 192.4110504 | 604.1810856 | Phage prote |
| 51.7638289 | 45.32916487 | 76.48241216 | 59.00203176 | 52.80453639 | 56.90765712 | Peptidyl-tR |
| 1627.717679 | 1747.559121 | 1092.623527 | 1011.465906 | 797.4884724 | 1217.78668 | SSU ribosor |
| 1666.537659 | 1083.369265 | 2229.916099 | 2454.869507 | 1984.991723 | 3543.746528 | Phage prote |
| 457.7959992 | 269.1662904 | 507.1831017 | 437.44123 | 220.1127072 | 179.7803636 | Phage lysin |
| 113.7152136 | 67.70678438 | 121.6124705 | 102.9606746 | 38.89535694 | 41.52848285 | Phage prote |
| 126.3682586 | 79.3509116 | 129.3652902 | 109.1371738 | 51.76178536 | 36.25982468 | FIG011165 |
| 92.35669443 | 60.934757 | 69.97425196 | 61.27028959 | 37.44332027 | 39.21386167 | Phage hyalu |
| 69.85497154 | 37.13915218 | 39.88890669 | 28.34870418 | 21.44523066 | 28.89015586 | Hyaluronat |
| 54.82946271 | 35.24682629 | 47.93708963 | 38.80444083 | 20.06434602 | 22.09549678 | Phage endo |
| 33.03087361 | 18.10128257 | 27.61276773 | 19.03037227 | 15.35560277 | 12.35566131 | Phage tail fi |
| 1679.041458 | 2059.868012 | 1014.765099 | 1007.169044 | 948.5369113 | 905.7053566 | LSU ribosor |
| 35.20874777 | 23.79866475 | 23.78550627 | 17.50023208 | 14.49194198 | 13.90799908 | Phage tail le |
| 46.7005104 | 45.94000145 | 34.32447252 | 24.42349899 | 18.62692011 | 15.97493568 | hypothetica |
| 56.60753215 | 46.83773468 | 25.54988769 | 13.23947099 | 14.38514622 | 19.89851458 | Phage base |
| 78.05076608 | 64.31600203 | 33.99363423 | 17.6600685 | 29.19895585 | 31.12535855 | Phage majo |
| 95.898323 | 88.85287475 | 54.22687039 | 25.96198711 | 51.63529716 | 78.80751862 | hypothetica |
| 80.61677516 | 68.65548694 | 47.90003615 | 30.02437658 | 36.16749068 | 88.398777 | hypothetica |
| 61.38692316 | 44.68109754 | 36.78867093 | 15.28991775 | 18.01115415 | 36.04262348 | hypothetica |
| 51.97181326 | 40.58776827 | 24.42742903 | 11.55966578 | 19.82439989 | 19.51213566 | hypothetica |
| 45.87432446 | 39.01981975 | 26.97330277 | 14.21194984 | 14.55607876 | 22.52306004 | hypothetica |
| 70.72436164 | 56.33758881 | 25.60058191 | 14.19858175 | 22.69831256 | 26.26498642 | Phage majo |
| 1851.486695 | 2079.792522 | 1389.01443 | 1287.467304 | 1166.941099 | 1423.094385 | LSU ribosor |
| 50.10322417 | 41.89070178 | 20.44525283 | 13.89143251 | 15.91139101 | 22.11472089 | Prophage C |
| 19.4770047 | 12.93817957 | 13.23666451 | 7.724856687 | 6.495013298 | 6.952922886 | Portal prot |
| 16.8805929 | 15.00812872 | 15.26185124 | 11.0159716 | 12.11373642 | 13.93240882 | Phage term |
| 20.31989394 | 18.23685758 | 13.69389617 | 11.92373653 | 15.3736207 | 21.99349027 | Phage prote |
| 20.905585 | 26.71773769 | 17.41863593 | 14.65409939 | 13.35097085 | 21.30334971 | Phage-asso |
| 16.45142348 | 15.71631629 | 10.7522444 | 7.099854357 | 14.76338545 | 14.8004843 | hypothetica |

| | | | | | | |
|---|---|---|---|---|---|---|
| 14.19919567 | 5.96136135 | 11.40108674 | 6.316422152 | 8.12432303 | 12.82081879 | hypothetica |
| 15.5143488 | 10.10334618 | 8.55790881 | 5.420521459 | 7.412753921 | 8.408137641 | hypothetica |
| 9.56852849 | 7.530734887 | 10.50023867 | 5.819661868 | 6.033297181 | 9.31602852 | Phage integ |
| 14.89203816 | 10.55573482 | 18.05414172 | 9.15881212 | 14.79859587 | 24.44695862 | putative tra |
| 1730.044563 | 1881.477901 | 1193.091285 | 1162.68488 | 1101.071446 | 1000.809285 | Preprotein |
| 31.56036383 | 29.72433732 | 35.23697486 | 22.63155749 | 31.65591446 | 33.78710099 | B-cell recep |
| 29.71295267 | 26.84291189 | 28.5457816 | 22.04599024 | 26.41004124 | 36.10435792 | putaive imr |
| 18.19635058 | 18.39143395 | 24.02097154 | 17.73301921 | 15.87412053 | 23.73215393 | hypothetica |
| 22.84961581 | 19.55541645 | 26.96268921 | 16.91933231 | 11.69034268 | 13.5006105 | Single-strar |
| 20.11798721 | 14.44202037 | 25.06435351 | 13.11937952 | 21.75635154 | 22.54179023 | hypothetica |
| 16.32637811 | 14.26110182 | 20.4093528 | 13.34246704 | 13.33157946 | 15.13515412 | conserved h |
| 19.59054174 | 14.45007279 | 18.95656359 | 18.40443763 | 14.23681725 | 14.09682445 | Phage prote |
| 32.20304761 | 29.162498 | 36.91603912 | 25.44114478 | 20.07008689 | 19.42758307 | hypothetica |
| 13.89260035 | 11.02899389 | 12.1670134 | 12.85447315 | 15.50035315 | 12.34057001 | FIG011169 |
| 27.18068484 | 23.14970913 | 33.45546316 | 26.73382997 | 17.90919181 | 25.12187466 | conserved h |
| 288.868252 | 382.7992906 | 195.8877202 | 129.3413468 | 233.3009335 | 280.1383882 | Adenylate k |
| 19.00507728 | 16.97362159 | 18.38633793 | 16.8522143 | 13.35097085 | 14.8721498 | hypothetica |
| 31.25835078 | 23.9880617 | 22.70704246 | 20.72783796 | 22.9921905 | 15.0731248 | hypothetica |
| 39.21404321 | 27.25707991 | 42.07658741 | 39.74451311 | 23.47359754 | 24.84377273 | Helicase loa |
| 19.57872134 | 17.38694833 | 20.63668955 | 21.99557242 | 13.44991273 | 11.47298213 | replication |
| 28.76980378 | 26.3696582 | 32.58146542 | 25.98070194 | 24.86067081 | 26.07729271 | Replicative |
| 21.99661722 | 13.09693024 | 23.29652954 | 9.837242648 | 17.08866094 | 10.79310171 | hypothetica |
| 16.82741217 | 13.75177675 | 25.20057282 | 12.59336667 | 18.40666936 | 20.09749973 | hypothetica |
| 8.471255375 | 12.42685474 | 16.31590575 | 8.94581649 | 8.447091677 | 14.02151144 | hypothetica |
| 175.5330054 | 154.2818382 | 188.3658816 | 203.3256291 | 96.45094747 | 70.42498863 | Phage antir |
| 64.05414934 | 46.79969739 | 86.73477151 | 68.38579717 | 62.39179186 | 41.08822167 | hypothetica |
| 332.5888523 | 276.650225 | 343.5563023 | 164.6076918 | 165.4078781 | 351.0179747 | Translation |
| 20.52191443 | 19.92209107 | 22.26168911 | 18.83361366 | 13.55007397 | 13.02091532 | conserved c |
| 15.8375644 | 7.174840044 | 18.40737493 | 16.72478735 | 19.84719131 | 12.88861396 | hypothetica |
| 264.9069532 | 242.8641013 | 365.5763097 | 302.6322022 | 447.8851331 | 419.1273447 | Phage trans |
| 213.1421834 | 175.1589449 | 303.1147983 | 263.438199 | 303.6491573 | 337.2084305 | Phage prote |
| 353.2353938 | 280.3485407 | 430.8298091 | 444.3692488 | 523.0375575 | 486.0014813 | Pathogenic |
| 630.8431352 | 550.5831305 | 715.8326922 | 576.2935767 | 617.7505823 | 672.5358806 | Phage integ |
| 241.5911224 | 184.8022019 | 278.2143239 | 251.9462886 | 212.4171959 | 210.0708484 | Glycosyl tra |
| 103.9166506 | 84.37180322 | 156.0490207 | 111.9945973 | 91.96876202 | 117.0591212 | Membrane |

| | | | | | | |
|---|---|---|---|---|---|---|
| 168.0480026 | 133.8268144 | 204.8941678 | 176.9849708 | 164.6674657 | 177.8792691 | Heteropoly |
| 1682.84274 | 1719.929177 | 1115.338583 | 528.3930069 | 678.120578 | 1026.003127 | LSU ribosor |
| 156.8839295 | 114.5495838 | 173.5334675 | 169.8877054 | 152.0339248 | 188.00627 | membrane |
| 251.848328 | 242.5243317 | 280.3094304 | 282.1722264 | 282.379963 | 424.8059746 | Tripeptide a |
| 263.8946313 | 244.5188841 | 242.5191813 | 282.7432061 | 451.8178604 | 463.4754692 | Pore formir |
| 166.7743524 | 140.2562166 | 205.5144896 | 172.6297315 | 207.939586 | 191.8397701 | Ferredoxin |
| 203.696976 | 146.0925501 | 197.2935412 | 225.9173656 | 160.5277664 | 287.4321659 | FIG011145 |
| 284.029624 | 276.1086756 | 315.3202246 | 396.2900998 | 328.3252483 | 303.2332007 | Cytidylate k |
| 1409.095842 | 1406.121948 | 1226.484828 | 1208.342263 | 1026.835636 | 789.1391136 | Translation |
| 2097.037504 | 1831.427099 | 1747.484085 | 1649.696341 | 963.2451782 | 843.1814658 | LSU ribosor |
| 2220.558508 | 2171.732972 | 1716.864625 | 1780.015136 | 1497.730234 | 1139.695714 | LSU ribosor |
| 300.8623029 | 271.973915 | 318.1425042 | 273.0396522 | 222.2505723 | 179.4419619 | Lipoteichoi |
| 1610.109108 | 1818.970377 | 1083.244557 | 623.2496576 | 967.3660289 | 1472.059488 | SSU ribosor |
| 291.8520243 | 265.2735963 | 323.9390746 | 270.9403235 | 304.5260103 | 291.2065553 | LSU m5C19 |
| 260.801115 | 241.5096813 | 297.3535537 | 257.3266253 | 277.7879883 | 264.9535008 | 3-dehydroc |
| 328.8024937 | 325.6388413 | 347.3610679 | 392.9577745 | 366.7832494 | 375.7044165 | Chorismate |
| 966.6471322 | 997.2967721 | 1158.271381 | 1091.18409 | 956.027337 | 1197.211585 | FIG011169 |
| 299.2632457 | 307.8794998 | 278.2947869 | 321.0255033 | 372.6064936 | 386.4869515 | Glutathion |
| 315.9333855 | 357.7859985 | 239.8810582 | 327.1802884 | 259.7558398 | 157.1473512 | Dihydrofola |
| 327.4174027 | 354.4189693 | 260.6870684 | 278.2534349 | 185.2520665 | 117.1670562 | Histone ace |
| 281.3350022 | 307.9325488 | 265.1655864 | 297.74553 | 240.294801 | 210.5753777 | Cysteine de |
| 341.2310788 | 377.6572595 | 312.5717271 | 328.4960614 | 307.9352462 | 278.189095 | tRNA S(4)U |
| 174.5071771 | 155.23053 | 145.6043491 | 153.5535415 | 170.96422 | 213.798228 | Capsule bio |
| 2638.315513 | 3255.119227 | 1571.129712 | 1184.491999 | 2271.536388 | 2236.945927 | SSU ribosor |
| 1592.504809 | 1859.614415 | 1465.838119 | 1351.840669 | 1835.29102 | 2349.684825 | LSU ribosor |
| 767.7586264 | 853.7276031 | 647.3541641 | 634.0586813 | 996.8196527 | 1114.028379 | FIG139598 |
| 1682.337198 | 1672.665091 | 1508.002277 | 1350.83133 | 2016.669756 | 4019.705022 | LSU ribosor |
| 135.9434216 | 127.4309842 | 180.4261864 | 136.5113439 | 144.5818733 | 156.3651372 | Cysteine an |
| 155.5807797 | 136.3107955 | 153.6938465 | 152.0482535 | 119.7275644 | 105.4790345 | Lipoprotein |
| 457.6362884 | 477.8924323 | 482.4390872 | 500.5583782 | 392.8739341 | 341.7928321 | Ribosomal |
| 325.762316 | 290.300233 | 370.9524319 | 336.560027 | 296.6506284 | 348.9341763 | Uracil phos |
| 444.4699322 | 386.4342817 | 408.8540934 | 425.2741761 | 348.7800892 | 288.3991211 | Uracil perm |
| 541.5099146 | 499.3871443 | 566.6498163 | 545.185947 | 466.1381784 | 426.323558 | Aspartate ca |
| 694.0011944 | 622.5576812 | 699.2830861 | 744.8321502 | 439.0678981 | 321.1146217 | Carbamoyl- |
| 1837.511665 | 2493.622043 | 1244.9244 | 888.9607419 | 1839.428064 | 1543.526505 | DNA-direct |

| | | | | | | |
|---|---|---|---|---|---|---|
| 470.8646979 | 422.5744948 | 482.6935327 | 516.3356008 | 400.9814995 | 339.4181139 | Carbamoyl- |
| 338.4158807 | 292.6051905 | 435.4278597 | 383.158041 | 330.0697842 | 349.9998537 | periplasmic |
| 259.8830715 | 216.6464359 | 374.2869887 | 315.1868002 | 203.2137719 | 191.3926451 | ABC transpo |
| 477.6951856 | 466.5467651 | 619.8919872 | 644.492332 | 791.5259779 | 977.2717865 | ABC transpo |
| 256.689106 | 244.2282922 | 300.4954671 | 363.8805563 | 439.6200506 | 429.1755002 | Glycerophc |
| 471.6461485 | 548.5167091 | 372.5475451 | 381.2481353 | 529.0332985 | 589.4530415 | SSU ribosor |
| 607.7665337 | 784.4406367 | 460.0616574 | 489.0805672 | 761.5452672 | 571.5226485 | KH domain |
| 173.963306 | 203.3201549 | 173.228837 | 193.7079731 | 280.1395843 | 277.064052 | Cell surface |
| 335.1890495 | 421.1738193 | 325.6967166 | 300.3270183 | 554.1414809 | 538.4330151 | FIG011152 |
| 41.76344036 | 36.11523366 | 36.73069792 | 32.1813056 | 43.63809468 | 48.72955413 | Cobalt-zinc |
| 75.56741985 | 72.13627707 | 95.67932551 | 84.24852519 | 97.12669917 | 89.61282413 | Transcriptio |
| 1912.662137 | 2344.289597 | 1173.870031 | 870.7261383 | 1447.420574 | 1266.765661 | LSU ribosor |
| 31.31104685 | 30.43907235 | 37.07670484 | 22.94966715 | 37.12364273 | 29.91524385 | Transcriptio |
| 132.925685 | 138.7214738 | 141.0532871 | 103.5528122 | 151.168762 | 215.3801416 | 16S rRNA pı |
| 121.6376872 | 138.9348124 | 114.6832625 | 99.17041648 | 128.7461027 | 189.2789114 | tRNA (Guan |
| 104.1164354 | 133.3750225 | 84.78355858 | 71.66562958 | 121.420994 | 139.5288652 | Thioredoxir |
| 313.2517695 | 349.6769068 | 231.8738258 | 243.525809 | 286.8868006 | 293.207149 | Membrane |
| 456.2555484 | 511.0354143 | 349.5352178 | 349.0458983 | 468.7883862 | 496.3690923 | 2-dehydrop |
| 266.8429968 | 306.0719579 | 292.8857162 | 239.0526928 | 296.4865869 | 518.0594993 | Transcriptio |
| 295.2872204 | 316.3425637 | 279.3815083 | 250.1801837 | 213.9694583 | 247.2521348 | 1-phosphof |
| 946.8618431 | 1019.090506 | 706.8648069 | 699.7953396 | 715.4380258 | 789.0668576 | PTS system, |
| 34.26841419 | 30.05400834 | 41.12261894 | 35.13555995 | 29.62289713 | 27.61937097 | Peptidoglyc |
| 42.35627688 | 52.26588905 | 73.89042374 | 37.91322227 | 34.70156581 | 73.84662691 | hypothetica |
| 164.5496097 | 184.6001218 | 145.4970021 | 170.1365777 | 283.8578796 | 220.220908 | Peptidoglyc |
| 4.355330209 | 5.893618608 | 6.451346641 | 5.495287272 | 10.30773484 | 8.038999891 | Mobile elen |
| 2.37563466 | 1.571631629 | 2.822464156 | 3.205584242 | 5.890134197 | 3.014624959 | hypothetica |
| 4.399323444 | 3.86014786 | 4.385783901 | 4.284824384 | 4.248244937 | 4.642404908 | Immunoglc |
| 1.18781733 | 1.178723722 | 2.016045825 | 0.915881212 | 0.490844516 | 0 | Phage prote |
| 11.00576509 | 9.323238475 | 11.75457227 | 8.382641602 | 7.653846696 | 19.07559296 | Nucleoside |
| 473.3810587 | 507.5008905 | 584.8120332 | 432.2670854 | 974.5775847 | 1615.71238 | Hypothetic |
| 290.2807298 | 318.9912394 | 341.6592148 | 272.3694799 | 545.721974 | 700.5008386 | DegV family |
| 216.1061206 | 216.557579 | 235.9223849 | 270.1281411 | 274.2395814 | 272.1889665 | tRNA nucle |
| 277.3091747 | 276.24062 | 293.7629968 | 362.2851529 | 392.5815215 | 447.4101537 | COG0488: A |
| 93.07280995 | 69.75460927 | 80.42089649 | 129.7289278 | 59.17029787 | 34.68883515 | FIG011146 |
| 179.9933414 | 158.9986451 | 143.1539693 | 206.8420781 | 132.9937843 | 87.13806451 | Peptide def |

| | | | | | | |
|---|---|---|---|---|---|---|
| 194.204888 | 173.6055683 | 178.4696305 | 215.7985389 | 140.3039914 | 101.5058518 | 5'-nucleotic |
| 144.8934097 | 170.3003914 | 150.1178738 | 110.0936185 | 185.0609684 | 223.0307149 | GTP pyroph |
| 174.2842288 | 210.1616824 | 151.8091547 | 148.5123974 | 204.2617539 | 196.1984615 | Regulation |
| 220.223067 | 248.8990334 | 213.2790085 | 228.6360398 | 373.9494768 | 401.1187111 | Regulation |
| 159.4860267 | 182.448454 | 129.8601326 | 137.2807927 | 178.0081147 | 148.5435127 | Mevalonate |
| 218.4075547 | 241.7319124 | 168.451829 | 181.8387651 | 227.8765145 | 165.8203231 | Diphosphor |
| 256.6809993 | 273.2686119 | 182.5654752 | 192.1182779 | 234.3855176 | 211.5588877 | Phosphome |
| 278.0212441 | 314.1834501 | 215.5336264 | 234.2435585 | 267.435891 | 283.6792537 | Isopentenyl |
| 227.525573 | 257.0318675 | 170.2186297 | 175.1784065 | 231.5495634 | 220.9781425 | Hydroxyme |
| 156.5979582 | 174.2827217 | 120.7981744 | 105.7001685 | 154.9165397 | 138.1190415 | Hydroxyme |
| 93.61131813 | 100.0792569 | 83.6371011 | 70.84995376 | 124.3239039 | 107.0909628 | Thymidylat |
| 222.2983436 | 255.437478 | 193.0546051 | 165.8517424 | 305.1515307 | 341.5364261 | Dihydrofola |
| 360.096201 | 443.641279 | 284.9344767 | 249.698141 | 604.5137728 | 732.6772708 | FIG011141 |
| 438.5846492 | 519.558417 | 342.6687841 | 293.9308534 | 546.3937495 | 627.4341378 | ATP-depend |
| 484.7086584 | 613.9578957 | 408.0476751 | 368.5505997 | 647.4239171 | 715.8729403 | GTP-bindin |
| 213.6155359 | 207.2272349 | 207.1324501 | 186.4261435 | 242.5721933 | 222.125995 | Hypothetic |
| 10715.49245 | 10071.91355 | 14943.02382 | 15481.53265 | 9145.106498 | 6538.664115 | putative AT |
| 940.0428026 | 957.5923941 | 971.8755648 | 885.0304765 | 1386.162137 | 1929.18368 | Ribose 5-ph |
| 137.9366251 | 116.9166502 | 132.0782454 | 101.8608429 | 137.5691253 | 112.7089512 | Transcriptio |
| 1349.407529 | 1329.934913 | 1103.156323 | 1148.796152 | 1971.979994 | 1842.055241 | Phosphope |
| 853.0737068 | 868.9946856 | 723.9113119 | 861.052949 | 1493.102272 | 1385.907175 | Arsenate re |
| 1107.397697 | 1151.132855 | 986.5781586 | 1036.574003 | 1999.664201 | 2303.62008 | Purine nucl |
| 781.0314846 | 724.4693529 | 664.3125118 | 667.1309534 | 1130.592285 | 1404.207239 | Purine nucl |
| 333.25126 | 304.227248 | 323.3645517 | 360.7109351 | 456.2054509 | 351.209539 | Histidine pr |
| 38.56102636 | 30.06599637 | 29.45179989 | 35.04241159 | 38.41391867 | 15.29157588 | hypothetica |
| 273.3290915 | 277.2732887 | 248.1205008 | 311.1569946 | 243.4718826 | 168.8988553 | Methionine |
| 554.2262759 | 532.2475382 | 662.7947751 | 668.0304528 | 1321.825088 | 1898.70848 | FIG011145 |
| 407.5944603 | 285.9553132 | 465.767678 | 459.4472504 | 271.1331614 | 288.4121714 | Orotidine 5 |
| 463.588135 | 353.0932392 | 500.4401752 | 529.0304334 | 284.6898195 | 235.3321516 | Orotate pho |
| 224.2194755 | 234.6746972 | 190.1774632 | 194.400659 | 272.0322971 | 263.918656 | Zinc ABC tra |
| 516.9576948 | 406.6475321 | 518.4687912 | 578.8746943 | 330.2928173 | 274.4862643 | 6-aminohex |
| 751.6794019 | 707.8890794 | 685.1755743 | 630.4951705 | 562.2896627 | 558.2638813 | Cystine ABC |
| 692.6197517 | 610.4738709 | 620.9049521 | 749.5875726 | 607.7424178 | 618.6695769 | Cystine trar |
| 462.0500439 | 418.9335502 | 447.0072983 | 470.3764243 | 345.7526786 | 305.9798238 | Uracil-DNA |
| 603.3999712 | 607.511791 | 581.4981538 | 742.2751705 | 473.1137736 | 379.2868566 | Dihydroorc |

| | | | | | | |
|---|---|---|---|---|---|---|
| 263.6880465 | 238.0214069 | 302.2184583 | 280.4993207 | 257.4410678 | 294.4189797 | Acyl-phospl |
| 484.5563741 | 521.8784165 | 432.6124181 | 510.7517258 | 426.0530402 | 359.9616797 | Topoisomer |
| 545.1212409 | 592.4667915 | 450.2174531 | 558.0620595 | 452.2952641 | 321.6825414 | Topoisomer |
| 225.708447 | 236.5259647 | 177.3177147 | 140.8100413 | 102.6467831 | 99.42973549 | Branched-c |
| 391.6197742 | 403.7256314 | 482.1753237 | 279.9979705 | 461.5213375 | 913.7837214 | FIG011144 |
| 191.2152996 | 196.9161982 | 154.1682102 | 167.5523864 | 204.6388535 | 164.4005033 | Zinc ABC tra |
| 2344.235309 | 3325.681993 | 1849.466278 | 1408.907815 | 3143.602716 | 2854.894831 | SSU ribosor |
| 331.4911994 | 345.9145654 | 297.336818 | 267.8725842 | 433.4983254 | 346.4331389 | FIG015049 |
| 572.4148275 | 663.9068807 | 481.1547666 | 392.4091661 | 892.8894412 | 979.4018636 | Rhodanese |
| 220.1807733 | 180.9292997 | 242.3188739 | 204.1744946 | 214.0560964 | 184.7989609 | hypothetica |
| 169.9161948 | 143.6398209 | 159.6583268 | 166.9885745 | 142.0024601 | 116.5343395 | Glutathion |
| 377.6971628 | 332.7917006 | 393.0956679 | 437.0839051 | 284.2038348 | 226.5777527 | Exfoliative t |
| 985.6870587 | 899.2929455 | 886.7868011 | 445.8323459 | 304.4899881 | 492.5590611 | FIG011153 |
| 96.50355906 | 97.7554873 | 110.909401 | 96.53387975 | 93.39134998 | 108.3925152 | tRNA dimet |
| 357.5693585 | 368.1566071 | 342.2408227 | 361.5739741 | 307.5958969 | 322.4338 | GTP-bindin |
| 375.2456885 | 413.665306 | 358.7800797 | 359.5265777 | 287.4681847 | 218.4181102 | Cystathioni |
| 71.02157784 | 52.30586514 | 73.71167549 | 83.5741606 | 57.06067503 | 51.9708782 | HIT family p |
| 303.6214362 | 329.6370597 | 253.2673827 | 260.9966009 | 222.4950692 | 214.7190939 | Ribonuclea |
| 564.0657249 | 619.2228617 | 428.9513032 | 541.2678379 | 438.0257967 | 365.3016127 | Short chain |
| 314.0866231 | 347.4862604 | 267.135646 | 300.8116773 | 214.7195008 | 159.9346484 | Single-stran |
| 685.6841464 | 851.497298 | 639.2613283 | 642.4933173 | 744.8352742 | 659.6162049 | Adenine ph |
| 474.5017649 | 527.3237701 | 475.7160764 | 460.2704793 | 461.1355062 | 376.651826 | Chromosom |
| 612.9643645 | 683.4085845 | 516.7705409 | 620.1226764 | 630.0740203 | 476.0996739 | Endonuclea |
| 279.1975872 | 293.3254839 | 261.0823362 | 253.5671129 | 240.5781158 | 188.3372682 | Putative tR |
| 265.2641489 | 285.7620703 | 259.7709541 | 247.6013467 | 252.2530222 | 264.8590648 | FIG137478 |
| 249.6240502 | 250.1407197 | 228.3759228 | 260.5173225 | 193.6241946 | 190.9534798 | Glycine/D-a |
| 715.6394616 | 839.7390375 | 647.0255761 | 665.3719095 | 861.65216 | 1078.473416 | Glucose-1-p |
| 201.806101 | 194.1846525 | 176.1802719 | 166.5635999 | 178.2035032 | 139.8670864 | Tyrosyl-tRN |
| 694.2132394 | 790.1020824 | 732.1300961 | 660.1745787 | 671.0191601 | 917.5828158 | dTDP-4-deh |
| 675.9947155 | 686.762259 | 591.9157023 | 703.0905973 | 719.4337782 | 809.7496936 | dTDP-gluco |
| 103.6912235 | 106.0604237 | 119.7962324 | 90.66647973 | 119.6549274 | 170.0071518 | Mutator mu |
| 156.285407 | 175.3127159 | 127.5942099 | 129.2926726 | 182.5841937 | 153.1286654 | Hypothetic |
| 171.7385889 | 200.345253 | 153.0644023 | 147.6858454 | 241.4105482 | 204.2601655 | TPR-repeat- |
| 108.3335394 | 113.4655285 | 103.1805667 | 113.928309 | 106.0794283 | 94.35976732 | Fibronectir |
| 21.25567853 | 13.64837993 | 17.8260894 | 11.08698309 | 18.60042378 | 20.09749973 | hypothetica |

| | | | | | | |
|---|---|---|---|---|---|---|
| 60.97224491 | 50.87744432 | 56.37663281 | 58.4183692 | 60.25742892 | 65.48284446 | ABC transpo |
| 60.01890783 | 50.83415406 | 66.40437836 | 60.57448844 | 53.34972123 | 78.44955066 | ABC transpo |
| 57.34074699 | 51.31097728 | 67.25465125 | 55.17007775 | 58.82373809 | 90.08128336 | ABC transpo |
| 347.7037117 | 424.2585772 | 291.9717996 | 302.3721519 | 403.7340137 | 362.0956307 | Multimodu |
| 517.8940733 | 532.1987248 | 491.5949431 | 465.7107174 | 456.1487199 | 644.7524597 | Ribonuclea |
| 308.1071211 | 340.1202799 | 313.8875623 | 324.9630438 | 294.7315241 | 271.8532024 | Tributyrin e |
| 176.9738847 | 166.5352781 | 189.6932659 | 167.8835469 | 151.8465788 | 147.8733466 | FIG011152 |
| 131.0085319 | 141.2985794 | 140.8696171 | 150.3139637 | 158.6446522 | 136.5113189 | ABC transpo |
| 997.5704262 | 1019.419636 | 723.1799799 | 632.0998136 | 1268.354387 | 1422.442543 | Acetoin deh |
| 1569.673336 | 1659.953562 | 1171.527851 | 1219.712464 | 2042.207107 | 2349.361615 | Acetoin deh |
| 1844.503404 | 2030.949332 | 1364.49413 | 1449.157919 | 2377.358419 | 2360.301681 | Dihydrolipo |
| 2215.912284 | 2449.633795 | 1431.118244 | 1702.978311 | 2543.776663 | 2667.703833 | Dihydrolipo |
| 92.13689759 | 96.13081798 | 87.02514437 | 81.8156914 | 90.66616824 | 73.084084 | Hyaluronat |
| 160.2953487 | 179.4993821 | 133.6210736 | 117.7323667 | 188.8412721 | 190.1954292 | Lipoate-pro |
| 120.7831834 | 117.9587255 | 180.9567286 | 135.2686098 | 189.6493758 | 260.1632382 | Ribosome-a |
| 508.6935108 | 662.9403928 | 325.036098 | 303.7120642 | 674.038093 | 671.7703378 | DNA-directe |
| 351.2459831 | 358.2605735 | 349.5701277 | 372.7358993 | 399.8151697 | 384.8975705 | Putative am |
| 207.7619776 | 212.3105941 | 203.296621 | 192.1715043 | 211.4575707 | 185.67757 | proposed a |
| 94.13312923 | 97.8396028 | 82.0615836 | 72.81900622 | 92.56912781 | 92.34942656 | Hypothetic |
| 85.64200183 | 86.41510585 | 89.03465075 | 75.28026765 | 89.24445752 | 120.8370046 | uncharacte |
| 501.3009598 | 603.3744171 | 381.2645955 | 374.0928968 | 537.5399018 | 631.2037746 | Phosphoglu |
| 37.22567706 | 46.63486375 | 35.12064877 | 30.12992398 | 23.12015292 | 21.60011653 | FIG011154 |
| 79.56590708 | 92.88051111 | 83.12312019 | 67.78006848 | 99.78335208 | 94.83674275 | Hypothetic |
| 122.4729581 | 136.3124325 | 98.69789284 | 96.91555774 | 127.8151298 | 133.4762233 | Acyl-ACP th |
| 221.9122258 | 253.4333042 | 177.7282751 | 205.0137239 | 296.8935616 | 316.3582898 | Hypothetic |
| 187.4987482 | 231.2011647 | 177.4884704 | 188.9927392 | 215.785485 | 195.0695708 | COG4478, i |
| 799.2092025 | 1011.428188 | 464.6422346 | 381.2021352 | 694.8295086 | 739.4170187 | DNA-directe |
| 161.7241576 | 148.5416408 | 165.0623119 | 158.9969784 | 125.9927753 | 166.8666692 | Translation |
| 2671.706613 | 5778.171038 | 6675.024046 | 2885.235162 | 3540.251135 | 5674.155809 | Collagen-lik |
| 55.80856026 | 44.86218326 | 50.56114927 | 58.73269995 | 46.33156705 | 55.0288683 | Mobile elen |
| 400.2678714 | 424.2119283 | 414.1373873 | 436.2352706 | 591.8733321 | 725.3848797 | Translation |
| 12149.69739 | 14651.3442 | 13230.80064 | 9166.742311 | 13165.76683 | 15517.76155 | Collagen-lik |
| 536.5246131 | 500.9844931 | 575.4291893 | 588.4223129 | 738.6873777 | 711.2587746 | Peptide me |
| 651.762356 | 568.7457517 | 622.1754599 | 725.7011721 | 849.7384775 | 775.8225998 | Putative me |
| 208.11898 | 198.0034496 | 233.2934155 | 178.4033403 | 278.3295807 | 611.8414882 | PTS system, |

| | | | | | | |
|---|---|---|---|---|---|---|
| 264.9546634 | 266.275621 | 329.4417178 | 302.5410889 | 341.177172 | 483.1636615 | DNA bindin |
| 866.1409456 | 835.1688807 | 856.6719603 | 611.8533268 | 1593.688342 | 1835.367399 | PTS system, |
| 817.5702687 | 838.0289096 | 814.2435747 | 591.5914199 | 1643.347441 | 1981.539038 | PTS system, |
| 699.7793399 | 686.8485763 | 655.5362919 | 563.0678408 | 1464.993039 | 2027.590145 | PTS system, |
| 45.66977661 | 51.49236718 | 46.62564164 | 39.43284927 | 63.79193824 | 53.05739928 | Two-compo |
| 41.12949233 | 43.9220246 | 39.73833292 | 35.66016582 | 54.94472457 | 51.04612098 | Two-compo |
| 55.36340563 | 56.33556354 | 55.08599711 | 45.19047126 | 70.0402777 | 72.96141019 | Ferric iron / |
| 245.4822481 | 203.6713177 | 218.2174924 | 275.7077606 | 693.4179443 | 936.1725892 | Succinate-s |
| 272.2263156 | 279.7976576 | 251.8340782 | 318.1884478 | 545.6158811 | 608.5281345 | Excinucleas |
| 784.3928038 | 893.0933564 | 624.4325816 | 735.165546 | 1184.473752 | 1153.756464 | Oxygen-inse |
| 957.7008888 | 1107.833103 | 848.2491363 | 982.4872117 | 1532.771659 | 1896.22048 | Acetylornit |
| 10.62584193 | 8.084111572 | 10.76942051 | 11.47046119 | 14.49010649 | 8.732459945 | Late compe |
| 212.2716692 | 265.6708019 | 185.5991991 | 230.6663793 | 353.2797264 | 278.956799 | GTPase and |
| 1647.391204 | 2283.289016 | 879.0442684 | 444.0104367 | 1016.959297 | 864.6738655 | LSU ribosor |
| 1425.510612 | 1929.757524 | 1035.917055 | 412.7471233 | 609.2909308 | 725.1573262 | LSU ribosor |
| 77.84239782 | 79.19192381 | 79.15426523 | 87.23101641 | 54.80302853 | 45.70717778 | D-alanyl-D-a |
| 128.2793378 | 123.184585 | 125.5098435 | 120.8050172 | 199.0902057 | 179.4374898 | Polysacchar |
| 1095.357183 | 1408.233589 | 1070.946258 | 869.6571603 | 2168.426634 | 2089.809731 | Dihydrofola |
| 665.9898875 | 761.3950767 | 693.6851831 | 602.2271231 | 699.3653356 | 779.9891176 | GTP cyclohy |
| 727.7480822 | 851.47906 | 726.3263278 | 658.0467739 | 812.5707858 | 1225.110088 | Dihydropte |
| 311.8680389 | 326.7684095 | 356.7057081 | 275.527598 | 349.3176808 | 502.270014 | Dihydroneo |
| 639.2875545 | 674.9546132 | 574.6334209 | 537.9019717 | 631.8080074 | 767.0746303 | 2-amino-4-l |
| 7.990989939 | 7.929813081 | 9.118164706 | 9.966123827 | 12.59187756 | 8.796474045 | Late compe |
| 280.2606835 | 351.2809073 | 220.0214336 | 238.3766506 | 321.4368279 | 378.0502651 | UDP-N-acet |
| 287.2152587 | 343.0647327 | 247.5390085 | 251.0703977 | 317.7612656 | 357.1612809 | Putrescine t |
| 238.938047 | 273.226676 | 189.0974907 | 212.8991799 | 229.1595606 | 215.3086858 | Spermidine |
| 201.5467655 | 199.5183319 | 159.104157 | 158.7763182 | 140.3170965 | 174.281793 | Spermidine |
| 238.2712901 | 264.0604538 | 194.4414253 | 209.7316809 | 180.8720911 | 188.8491315 | ABC transpo |
| 38.66648605 | 37.45014106 | 40.61151771 | 53.55017177 | 58.01693743 | 41.00976296 | Two-compo |
| 46.31101029 | 44.61110012 | 49.85987263 | 59.72828449 | 50.53597628 | 47.03753341 | Two-compo |
| 27.10934446 | 25.45052119 | 27.82506491 | 143.3642888 | 409.7004004 | 436.4416495 | Malate Na(+ |
| 21.90388084 | 25.25115085 | 21.68415356 | 141.5966481 | 440.3720725 | 475.7269344 | NAD-depen |
| 52.94271527 | 49.57375194 | 57.18657987 | 80.80689094 | 196.6743857 | 286.0735533 | Alcohol deh |
| 8.136730332 | 8.939556053 | 10.50563329 | 7.394270336 | 8.465942117 | 7.559609989 | Late compe |
| 48.72774443 | 43.82274436 | 44.54501252 | 55.24362866 | 47.05874411 | 53.96550853 | Mobile elen |

| | | | | | | |
|---|---|---|---|---|---|---|
| 45.59298841 | 41.67205076 | 61.01084134 | 65.22184389 | 91.62430973 | 170.3212351 | Class B acid |
| 80.30357995 | 93.59050971 | 71.33093645 | 58.25076202 | 74.88500684 | 81.29458305 | Voltage-gat |
| 104.4506686 | 113.8474619 | 97.32092434 | 84.79719709 | 149.5040566 | 135.6826323 | Lipase/Acyl |
| 80.85493403 | 89.74843774 | 80.64183302 | 61.21942838 | 91.45208358 | 84.62105148 | hypothetica |
| 95.86781002 | 90.03442894 | 103.9764911 | 81.60111863 | 103.1295659 | 135.9788279 | Permease |
| 109.5373397 | 94.92101158 | 116.8773703 | 141.457248 | 112.6131384 | 97.30022996 | DNA repair |
| 281.9359524 | 265.9580075 | 278.9095121 | 289.4500451 | 287.2286705 | 212.1499001 | Glutamine a |
| 619.727578 | 648.023925 | 572.8945849 | 697.5180914 | 849.549123 | 766.6962454 | Redox-sensi |
| 229.0985608 | 207.8347343 | 220.0965891 | 204.6520777 | 285.1975897 | 344.0830557 | FIG011143 |
| 11.8781733 | 8.80655654 | 10.56686088 | 9.632543782 | 5.754728813 | 3.291831852 | Late compe |
| 295.0588793 | 276.49849 | 277.9569564 | 303.2151417 | 299.5718075 | 327.0654277 | Cysteine de |
| 391.144251 | 436.0196372 | 349.0040431 | 409.4297112 | 629.962161 | 547.6722326 | Ribose-pho |
| 38.568162 | 31.67692765 | 44.03963316 | 35.59123881 | 37.02951377 | 36.65450728 | Adenylate c |
| 182.2733974 | 205.57503 | 221.1170261 | 185.9565961 | 377.336722 | 411.4604185 | GTP pyroph |
| 178.2435563 | 191.9756484 | 204.0614126 | 191.9739544 | 238.7721067 | 265.4454713 | NAD kinase |
| 157.4841254 | 154.5611237 | 149.4811196 | 172.1371444 | 257.7746368 | 260.0696322 | Similar to ri |
| 1307.934761 | 1341.804172 | 1246.256376 | 1278.206026 | 3273.992062 | 2868.797017 | Phosphate a |
| 98.85908252 | 93.86133338 | 92.86611088 | 104.7448275 | 135.5665807 | 162.2155335 | Short chain |
| 398.3491741 | 395.2311887 | 374.7390919 | 394.7846233 | 457.2963607 | 285.1223548 | Na+ driven r |
| 436.0870098 | 481.384947 | 336.7045423 | 399.5201995 | 402.856092 | 356.958061 | L-proline gl |
| 9.22187294 | 6.962924937 | 12.4535489 | 7.883534483 | 11.43232798 | 5.596772076 | Late compe |
| 469.9618323 | 491.7447017 | 364.7102153 | 427.4952031 | 438.539199 | 441.5132455 | L-proline gl |
| 202.1220871 | 186.9666651 | 190.8359475 | 228.0209139 | 144.3202596 | 120.1560883 | GMP reduct |
| 263.9321943 | 214.6006281 | 257.0562347 | 295.3480857 | 167.7979811 | 163.991454 | Xanthine pl |
| 326.4092419 | 265.4515197 | 292.7600003 | 402.0889714 | 212.7145815 | 144.2042562 | Xanthine pe |
| 212.820434 | 192.4118978 | 232.8565134 | 250.1262812 | 193.2014199 | 150.8275619 | Hypothetic |
| 125.9341814 | 98.86069922 | 167.0066348 | 131.1778252 | 75.68505769 | 105.349797 | 4-oxalocrot |
| 172.2126739 | 185.7006845 | 185.9006466 | 147.4086709 | 230.7485905 | 219.2742997 | Thymidine l |
| 285.5160915 | 320.0453186 | 289.5041805 | 249.9846886 | 281.7992907 | 284.1563156 | Peptide cha |
| 215.7302522 | 212.338659 | 219.576191 | 188.9986301 | 190.6580343 | 160.3493371 | Protein-N(5 |
| 360.2442947 | 376.1544736 | 338.7775686 | 321.9996586 | 355.0026736 | 339.6171401 | TsaC protei |
| 9.393314056 | 9.53817816 | 11.01178134 | 8.842991013 | 11.64486301 | 6.791568873 | Late compe |
| 493.0621869 | 514.4751749 | 463.022975 | 446.4162729 | 375.9023833 | 377.460326 | Phosphinot |
| 630.4408987 | 699.0947564 | 606.4496798 | 611.4763968 | 453.9198883 | 387.368515 | Serine hydr |
| 500.1463971 | 497.0887645 | 508.2909156 | 465.4699362 | 307.7565005 | 255.4725119 | Pseudourid |

| | | | | | | |
|---|---|---|---|---|---|---|
| 369.4111896 | 402.2591154 | 360.7915609 | 348.3096249 | 327.4914613 | 298.5483585 | Pneumococ |
| 369.579894 | 386.2387225 | 354.9923613 | 359.5032862 | 231.4395918 | 200.3458582 | ABC transpo |
| 276.4699827 | 281.8568529 | 237.6694023 | 268.149666 | 204.4503756 | 179.5167293 | ABC transpo |
| 2447.232927 | 2743.270971 | 2174.541437 | 2542.101454 | 4166.700693 | 3825.165479 | NADH perox |
| 1495.104707 | 1773.068804 | 1132.91941 | 838.0871554 | 892.1996972 | 943.7246671 | L-lactate de |
| 525.180513 | 539.9474115 | 576.8663428 | 582.7832803 | 660.5144858 | 627.4589722 | DNA gyrase |
| 461.2532255 | 452.6927743 | 508.7531961 | 441.8943672 | 351.4446737 | 343.2652953 | Sortase A, L |
| 8.816788427 | 8.911313358 | 6.983416468 | 8.120183942 | 6.477123515 | 6.008530846 | Late compe |
| 103.8622883 | 114.0002363 | 135.454905 | 117.2062479 | 87.49837031 | 94.51650234 | FIG012802 |
| 3895.644902 | 3460.339938 | 4459.896575 | 3795.869683 | 3660.718403 | 3762.251949 | FIG011140 |
| 5784.158638 | 5414.538245 | 8227.77102 | 7395.460415 | 7930.043938 | 9864.454424 | FIG011154 |
| 1324.207934 | 1073.569158 | 1712.471767 | 1384.66778 | 1172.79547 | 838.7400626 | Transcriptio |
| 831.4356387 | 660.8095382 | 1148.978891 | 895.2000234 | 563.1366396 | 444.0899133 | Hemolysin I |
| 656.0199293 | 518.5315237 | 799.2867395 | 638.873874 | 371.7062011 | 290.3883634 | FIG011150 |
| 668.0877872 | 533.5217335 | 809.9517533 | 579.1087777 | 419.6633893 | 340.1661615 | 50S ribosor |
| 483.1656956 | 334.9242251 | 548.6088337 | 402.8489634 | 308.1164896 | 364.9523246 | Ribonuclea |
| 113.0200384 | 78.86323585 | 137.871521 | 116.6681659 | 43.06764789 | 47.61450652 | Rossmann f |
| 324.0454901 | 337.3031374 | 318.5238824 | 304.433755 | 245.394605 | 209.0989162 | DNA topois |
| 109.3302833 | 96.83278744 | 166.22623 | 119.8031714 | 167.9954942 | 162.8869938 | Cell divisior |
| 177.350839 | 204.0649998 | 182.838873 | 158.9831538 | 199.424879 | 167.0367666 | Adenine-spe |
| 54.84489893 | 49.51886957 | 52.6945311 | 60.18647965 | 51.73345379 | 56.88974791 | Mobile elen |
| 230.6852364 | 213.3078651 | 195.156828 | 236.0484941 | 188.5542588 | 188.7553594 | tRNA:m(5)U |
| 26.41884386 | 24.136989 | 27.21880524 | 32.30418331 | 46.25224684 | 38.18960903 | Oxaloacetat |
| 17.19506992 | 13.92016585 | 18.27881548 | 19.53879919 | 22.06462969 | 24.30840443 | FIG011140 |
| 44.53468963 | 47.01462137 | 47.4201548 | 47.43794996 | 112.7683812 | 123.8486949 | Biotin carbo |
| 39.95099136 | 40.81239694 | 40.34230692 | 49.41385637 | 77.18106878 | 80.28338087 | Methylmalo |
| 36.93642137 | 34.94882537 | 38.38004527 | 30.55007162 | 34.34247735 | 33.38228768 | 2-(5"-tripho |
| 27.57920697 | 24.04867363 | 33.99470375 | 24.08135945 | 25.38850947 | 22.17655142 | Transcriptio |
| 40.43993473 | 36.97364152 | 50.24606519 | 63.25059994 | 107.7340785 | 106.0701375 | Mg2+/citrat |
| 49.27242257 | 33.61545428 | 44.80101834 | 62.75482379 | 59.62851903 | 70.15516109 | FIG011148 |
| 406.7753382 | 461.9178975 | 362.0595982 | 366.0770318 | 672.0264221 | 676.4143455 | Acetate kina |
| 104.3839472 | 84.17754254 | 128.0494561 | 141.9615879 | 178.1914335 | 270.4027236 | Biotin carbo |
| 122.3811794 | 106.3162572 | 148.6483414 | 200.6612474 | 343.0661941 | 354.3930767 | Oxaloacetat |
| 77.07614673 | 64.96077399 | 91.03566927 | 109.4986871 | 189.3569068 | 219.7326637 | FIG011142 |
| 152.532659 | 119.0167641 | 184.3018203 | 214.2984195 | 296.7941406 | 297.5600688 | Citrate lyas |

| | | | | | | |
|---|---|---|---|---|---|---|
| 225.6852927 | 199.6396934 | 286.5509458 | 407.5052555 | 987.194451 | 1000.801169 | Citrate lyas |
| 230.961898 | 209.2638278 | 286.9649886 | 466.2928691 | 1106.676681 | 1128.252397 | Citrate lyas |
| 206.4627566 | 181.7555334 | 222.6215991 | 462.0217347 | 1030.010514 | 1043.195608 | Apo-citrate |
| 223.0882727 | 192.2460366 | 242.1682917 | 430.9959717 | 1055.706275 | 1011.919916 | Oxaloaceta |
| 900.4851073 | 778.9203365 | 950.2410865 | 1425.591286 | 798.1047931 | 615.979778 | [Citrate [pr |
| 825.6191179 | 708.0151679 | 912.5049155 | 1139.788569 | 954.4944173 | 1137.693245 | FIG011146 |
| 70.56340573 | 58.19705239 | 82.87744819 | 97.75445016 | 90.19875469 | 69.44581589 | Hypothetic |
| 692.948896 | 593.654132 | 848.2075154 | 1090.745255 | 746.3586478 | 784.8158087 | Tyrosine re |
| 216.424056 | 196.2178716 | 253.6237564 | 290.251797 | 235.6053679 | 274.5508225 | Pleiotropic |
| 687.4901274 | 767.3845874 | 836.1366257 | 980.4323792 | 983.0456856 | 899.1027325 | Signal reco |
| 342.7304506 | 316.1185372 | 456.2630026 | 412.7892691 | 348.0690413 | 371.0985695 | Signal reco |
| 416.3987961 | 425.0150598 | 509.6010083 | 517.2959979 | 557.4982524 | 363.6526131 | Transcripti |
| 39.59391099 | 39.15927343 | 44.43736154 | 39.58599548 | 50.23454758 | 45.74493662 | FIG011140 |
| 491.9068466 | 571.277539 | 454.87566 | 486.0323177 | 706.3262012 | 618.7021077 | GMP syntha |
| 97.97763139 | 107.694913 | 109.3284035 | 93.68664515 | 80.13632415 | 67.43394083 | ABC transp |
| 411.1604267 | 381.0442287 | 373.7764649 | 498.0683198 | 433.9294713 | 343.2215031 | Pyrroline-5 |
| 78.50663642 | 96.3258095 | 95.38281325 | 71.5963055 | 99.85783065 | 69.36878938 | Substrate-s |
| 68.88518494 | 79.23416204 | 84.77158779 | 71.05209956 | 50.74890709 | 51.66935051 | Pyridoxal k |
| 62.15214397 | 74.1602064 | 66.91556357 | 59.49980073 | 44.41968815 | 29.45732821 | Predicted t |
| 564.9665671 | 499.1668952 | 696.8025288 | 637.7126882 | 776.4899624 | 1140.933281 | Anaerobic r |
| 314.4027518 | 324.9333453 | 276.972501 | 277.7592603 | 454.3391219 | 539.0409242 | Cardiolipin |
| 610.1869514 | 669.1990545 | 464.3348058 | 473.4629016 | 754.8254559 | 902.7998935 | Formate--te |
| 315.0743694 | 345.8051828 | 258.860284 | 252.406087 | 463.5882091 | 629.1108518 | Lipoate-pro |
| 363.2943343 | 367.0347878 | 272.7010557 | 244.6711238 | 454.9160334 | 541.6754688 | NAD-depen |
| 363.111055 | 401.639194 | 256.8179755 | 278.273487 | 556.8922536 | 530.7741978 | COG2110, I |
| 647.6279711 | 678.633369 | 450.8677619 | 387.3104873 | 777.5684663 | 727.3122199 | Biotinyl-lip |
| 431.6181174 | 436.3926587 | 448.1511305 | 539.3922891 | 466.5228948 | 411.7164762 | Glutamyl a |
| 257.3485314 | 255.6615175 | 193.1044974 | 211.2302615 | 417.7927019 | 394.5265937 | Luciferase-l |
| 324.1949432 | 343.7944188 | 278.2949657 | 319.8715133 | 716.7311628 | 721.952434 | Probable N |
| 442.827383 | 525.9614522 | 411.4178678 | 472.9098473 | 1154.733716 | 1173.002457 | Protein:pro |
| 158.0328395 | 164.715159 | 153.3940322 | 140.5937133 | 130.891871 | 91.70028014 | Phosphopa |
| 200.2320642 | 207.9389539 | 175.4624499 | 198.474478 | 173.7913222 | 142.7805887 | Phosphopa |
| 221.3924791 | 247.159753 | 196.5114141 | 214.2197951 | 269.8094805 | 252.0649571 | Substrate-s |
| 453.0151492 | 495.489099 | 402.9839872 | 337.709219 | 545.4284825 | 500.8591555 | Phosphoglu |
| 360.6397109 | 371.4272993 | 295.2148357 | 318.2902544 | 572.1492946 | 826.6435546 | Nucleoside |

| | | | | | | |
|---|---|---|---|---|---|---|
| 317.7773837 | 340.7120286 | 231.8850229 | 280.1822525 | 537.0253807 | 637.6285618 | Putative de |
| 200.5574936 | 202.1284357 | 170.8786239 | 182.6743897 | 427.63988 | 457.9555515 | Putative de |
| 84.18915811 | 67.33200767 | 95.41202138 | 60.93020274 | 41.95428919 | 68.11994644 | FIG014356 |
| 854.4162946 | 959.4117068 | 1029.320628 | 932.5288534 | 1086.346593 | 1690.423033 | Nucleoside- |
| 687.2284674 | 729.7206547 | 841.1563506 | 803.0164658 | 512.290646 | 663.9904718 | Cytidine de |
| 136.5390021 | 150.9718868 | 135.2990754 | 131.7388733 | 429.2658407 | 452.9042615 | Ribosomal |
| 28.94095644 | 32.40530361 | 36.14435252 | 39.85724102 | 68.11067231 | 49.75276806 | Pantothena |
| 1211.979768 | 1213.178723 | 1013.192261 | 657.3209006 | 855.3280342 | 1576.623088 | SSU ribosor |
| 104.5931598 | 118.1061389 | 128.6947697 | 100.1810571 | 132.8088229 | 164.5511534 | Two compo |
| 126.0670126 | 158.141067 | 176.2651266 | 107.3819839 | 139.007167 | 279.2212629 | Two compo |
| 279.1604731 | 326.1228516 | 228.8474146 | 219.5516664 | 427.2319954 | 418.0327455 | Lysyl amino |
| 163.0978535 | 173.3120383 | 147.0788661 | 129.5677825 | 171.4803595 | 225.0366827 | Phosphate |
| 177.0301665 | 187.4776386 | 141.9041267 | 146.0341822 | 168.0123918 | 134.804178 | Phosphate |
| 53.3397216 | 52.04176431 | 78.05520819 | 61.86518376 | 42.23115084 | 59.15490486 | FIG009688 |
| 226.6308189 | 247.4146956 | 168.6858641 | 172.9921901 | 177.9494523 | 149.5313972 | Phosphate |
| 160.3018342 | 168.3663478 | 134.2032667 | 139.9813042 | 116.0780951 | 117.8012231 | Phosphate |
| 186.2436659 | 196.0509711 | 146.6544104 | 169.3206036 | 183.9408361 | 176.6261034 | Phosphate |
| 227.7540403 | 266.198506 | 196.1075441 | 193.1273393 | 259.247429 | 210.2240196 | Phosphate |
| 120.5394489 | 123.8603965 | 107.1410715 | 115.4387578 | 135.9538221 | 127.3914742 | tRNA and rF |
| 122.2443183 | 136.1881552 | 106.6434583 | 123.0693613 | 179.9887649 | 181.3359956 | Inositol-1-m |
| 34.71246991 | 39.39843672 | 43.74543271 | 29.86023678 | 38.99860541 | 71.85544423 | FIG006438 |
| 445.2115325 | 461.5940302 | 563.6565455 | 452.3774757 | 1223.838994 | 1228.47828 | Transcriptio |
| 177.1668377 | 185.2099652 | 186.5393398 | 185.5911061 | 195.2645774 | 202.2028188 | Riboflavin k |
| 302.41695 | 358.0656331 | 318.2482102 | 340.0247808 | 491.3104027 | 521.1043234 | tRNA pseud |
| 170.1969839 | 191.3029121 | 171.7015105 | 155.9189164 | 201.9742986 | 225.0150687 | Phenylalany |
| 102.8910224 | 111.3151171 | 106.3317592 | 102.693437 | 89.07674982 | 71.21798561 | FIG011158 |
| 248.6348668 | 226.2361266 | 297.4899972 | 253.1162622 | 151.5309341 | 148.4316908 | membrane |
| 287.1453887 | 269.7046237 | 322.2431438 | 268.0448331 | 214.491654 | 222.6883764 | Type I restri |
| 67.38622394 | 65.47571131 | 62.3483387 | 56.62000576 | 56.48900032 | 52.83881896 | ABC transpo |
| 55.09306589 | 52.58921219 | 48.66079839 | 43.13252546 | 42.62376313 | 49.04133481 | ABC transpo |
| 49.00019924 | 41.33130692 | 47.58313683 | 33.9027852 | 29.50490795 | 23.98375658 | Transcriptio |
| 641.1993361 | 610.1455874 | 643.4012416 | 647.2797874 | 664.5209032 | 699.6559485 | Transcriptio |
| 6955.799985 | 6940.633813 | 8359.243358 | 9371.656171 | 12789.78199 | 16687.82944 | General stre |
| 8175.965334 | 8540.527756 | 10641.35185 | 12230.43165 | 15662.18917 | 24858.80739 | FIG011146 |
| 7586.721265 | 7926.262181 | 9809.720579 | 9137.23803 | 13869.52081 | 20243.31825 | General stre |

| | | | | | | |
|---|---|---|---|---|---|---|
| 21.23673408 | 16.90694631 | 18.20550473 | 18.595164 | 10.41185338 | 7.612689291 | FIG107367 |
| 1815.537938 | 1412.971674 | 2501.176853 | 2030.348731 | 1460.067657 | 3921.883518 | FIG011142 |
| 4730.56662 | 4810.630878 | 5044.278423 | 5229.621859 | 7329.431477 | 10496.41182 | FIG011143 |
| 3260.439311 | 2888.583192 | 3965.052055 | 3005.85593 | 3083.686321 | 4083.666661 | FIG011141 |
| 300.5654201 | 332.7060798 | 262.3311168 | 329.409178 | 278.5304511 | 220.5525099 | ATP-depend |
| 353.8851691 | 264.2907065 | 397.9793319 | 410.4643146 | 250.9651282 | 211.3655414 | Sodium/gly |
| 297.6847056 | 295.5582975 | 284.3211812 | 283.9231757 | 264.0552879 | 194.0969695 | Cobalt-zinc |
| 121.421327 | 104.2271983 | 153.3445088 | 137.7371745 | 152.9608493 | 133.1264614 | CAMP facto |
| 469.2800891 | 419.4960838 | 634.3824197 | 564.0383866 | 407.8759594 | 378.8990988 | Amino acid |
| 798.5903305 | 678.7951844 | 929.9155373 | 965.1643439 | 629.2159229 | 447.3129225 | ABC-type pc |
| 443.0077592 | 354.5718459 | 575.3455638 | 504.505034 | 315.7001833 | 353.6784298 | ABC transpo |
| 464.3218646 | 514.0851122 | 357.3865161 | 405.3844318 | 406.9880887 | 349.1705794 | Deoxyadeno |
| 341.4974823 | 293.2776879 | 505.7394956 | 399.0625281 | 248.227084 | 359.4222496 | Alkylphospl |
| 2438.330472 | 2323.365117 | 2488.300395 | 3371.16951 | 2252.335393 | 1939.85718 | Glucosamin |
| 674.2885616 | 565.70289 | 796.099644 | 807.2557307 | 381.1697954 | 309.0260711 | Signal pepti |
| 3065.46965 | 3714.446265 | 2462.585892 | 2710.53308 | 3726.695352 | 3181.141369 | Pyruvate ki |
| 1510.659989 | 1773.66534 | 1431.225526 | 1393.873655 | 1677.177955 | 1940.954685 | 6-phosphof |
| 244.8942576 | 252.2946164 | 234.3045737 | 238.093787 | 255.6178135 | 245.3755487 | DNA polym |
| 432.2122413 | 426.1149625 | 491.6550465 | 387.9200101 | 560.6711073 | 664.0278741 | Transcriptio |
| 359.9817212 | 338.4066902 | 421.1372636 | 395.0474756 | 324.0837785 | 264.5452003 | ABC-type m |
| 267.3042357 | 252.4808216 | 295.9124361 | 317.6919258 | 302.0754573 | 312.1249099 | ABC transpo |
| 202.489055 | 203.775457 | 216.8281871 | 176.8320896 | 217.1208817 | 266.0712891 | Immunorea |
| 20.45818178 | 13.09165984 | 26.28566791 | 25.43090689 | 13.98462445 | 12.25260044 | Phage holin |
| 6.06631644 | 4.403800708 | 4.063187473 | 3.625853641 | 5.90527593 | 4.908129754 | DNA modifi |
| 133.5228506 | 117.5701353 | 131.8183809 | 146.4000891 | 93.71354536 | 64.85308566 | D-beta-hydr |
| 7.524938004 | 5.937275042 | 10.54743975 | 7.81164293 | 10.15970237 | 11.20781202 | Lanthionine |
| 36.88048213 | 27.1463645 | 41.16680986 | 30.61477058 | 30.06851723 | 36.82199251 | Streptococ |
| 3835.947667 | 3319.7385 | 4538.787713 | 5436.849813 | 3207.826641 | 2302.997809 | Antiphagoc |
| 229.6552281 | 224.883535 | 214.2860745 | 216.0430848 | 301.6046745 | 264.0104974 | Transmemb |
| 77.51401919 | 80.29513298 | 85.02184428 | 64.58340961 | 82.93466544 | 100.4202829 | Mobile elem |
| 19.89048442 | 17.69632464 | 20.31920202 | 23.94272145 | 17.16023349 | 16.06217498 | FIG011144 |
| 97.40102104 | 89.94019184 | 89.29250238 | 71.18894875 | 64.34525388 | 59.37897647 | Lipoteichoi |
| 234.0263037 | 122.2836923 | 271.2291511 | 218.1529535 | 115.1953811 | 112.6996614 | Phage holin |
| 331.5696031 | 259.0858081 | 460.2439665 | 370.0099642 | 387.7833674 | 428.945646 | Lipoteichoi |
| 116.972573 | 128.6147981 | 105.2270211 | 84.86479845 | 122.655978 | 137.8221798 | 4-diphosph |

| | | | | | | |
|---|---|---|---|---|---|---|
| 374.5823943 | 315.0407037 | 458.4366023 | 458.2181458 | 496.4966654 | 879.4178669 | Nucleoside |
| 44.8044108 | 43.292901 | 51.20306721 | 54.40811066 | 58.09847361 | 67.76368867 | UDP-N-acet |
| 9.432899781 | 52.42611302 | 15.79526909 | 3.22175403 | 255.5399993 | 138.1385108 | Small Subu |
| 3.959391099 | 9.822697679 | 8.064183302 | 1.526468687 | 14.72533549 | 1.674791644 | tRNA-Gly-G |
| 11.17945722 | 12.75795087 | 21.6309858 | 11.63707893 | 42.27037482 | 31.21023487 | tRNA-Leu-T |
| 11.55714159 | 6.371479576 | 14.3847594 | 8.16867027 | 50.14573708 | 28.51672259 | tRNA-Arg-A |
| 12.19920501 | 14.01725507 | 16.3463175 | 17.07994693 | 46.96188076 | 26.88719558 | tRNA-Pro-T |
| 9.654477963 | 57.07504336 | 15.15056423 | 3.770916649 | 257.0704139 | 136.9736324 | Small Subu |
| 11.71545859 | 18.08452833 | 11.26776297 | 12.04446525 | 36.30904642 | 50.38140343 | tRNA-Ala-T |
| 11.51822865 | 55.80266434 | 12.97933635 | 6.509568945 | 308.9683685 | 55.39269553 | Large Subu |
| 0 | 0 | 1.654191447 | 0.469682673 | 2.013721093 | 0.515320506 | 5S RNA |
| 24.73263481 | 18.73040434 | 27.83800263 | 12.04446525 | 12.90988317 | 38.81845838 | tRNA-Val-T |
| 28.63778768 | 25.18916446 | 22.53552594 | 9.786128019 | 29.04723713 | 60.29249918 | tRNA-Asp-G |
| 12.36631741 | 12.91752024 | 15.90743007 | 15.05558157 | 29.85410483 | 41.29623232 | tRNA-Ala-T |
| 15.62061146 | 24.54328845 | 29.1636218 | 8.280569862 | 34.69531102 | 43.77400626 | tRNA-Lys-T |
| 30.13000056 | 20.69953852 | 27.73292306 | 13.40313969 | 33.76052527 | 63.2335967 | tRNA-Leu-T |
| 25.38349362 | 12.91752024 | 16.57023966 | 9.033348941 | 34.69531102 | 37.99253373 | tRNA-Thr-T |
| 4.619289616 | 7.203311631 | 10.7522444 | 3.052937373 | 13.0891871 | 12.56093733 | tRNA-Gly-G |
| 6.148701472 | 12.20325735 | 18.21556699 | 8.404557005 | 32.56897732 | 39.01279359 | tRNA-Leu-T |
| 9.630951322 | 8.282923448 | 13.077054 | 3.713031941 | 64.47309053 | 20.36908756 | tRNA-Arg-A |
| 16.69364896 | 16.5658469 | 19.615581 | 5.940851105 | 85.96412071 | 41.55293863 | tRNA-Pro-T |
| 28.25079055 | 25.4859183 | 27.46181341 | 19.30776609 | 60.49327013 | 53.77439116 | tRNA-Met-C |
| 16.69364896 | 12.10581119 | 20.2694337 | 9.653883046 | 39.79820403 | 62.73678969 | tRNA-Met-C |
| 16.36548321 | 24.62222885 | 43.54658983 | 19.53879919 | 64.79147616 | 58.28274921 | tRNA-Pseuc |
| 11.12556177 | 54.16284208 | 13.77909006 | 5.960797144 | 314.8909552 | 54.85288665 | Large Subu |
| 12.19920501 | 12.74295915 | 16.3463175 | 10.39648943 | 28.6547069 | 29.33148609 | tRNA-Met-C |
| 9.112023352 | 8.396388153 | 19.88428759 | 4.51667447 | 23.39916325 | 19.82219151 | tRNA-Phe-G |
| 14.72224296 | 20.58616077 | 20.44440837 | 3.869920614 | 29.86546917 | 29.72165453 | tRNA-Gly-T |
| 16.05158554 | 24.84877035 | 28.11566611 | 5.198244717 | 17.51120977 | 27.70195908 | tRNA-Ile-GA |
| 5.399169681 | 8.572536156 | 12.64610563 | 3.746786777 | 11.37866833 | 16.44340887 | tRNA-Pseuc |
| 10.50913667 | 51.4209389 | 14.44138888 | 2.672591412 | 256.4425515 | 138.7812023 | Small Subu |
| 10.41374097 | 13.56339625 | 13.91900132 | 15.05558157 | 41.15025261 | 61.11842383 | tRNA-Ala-T |
| 11.73092322 | 54.50379522 | 12.49615181 | 6.566338442 | 301.5245694 | 53.2542219 | Large Subu |
| 0.81218279 | 0 | 0.827095723 | 0 | 3.524011913 | 1.545961517 | 5S RNA |
| 5.206870487 | 16.79277631 | 8.616524624 | 6.022232627 | 20.17169245 | 28.90736262 | tRNA-Val-T |

| | | | | | | |
|---|---|---|---|---|---|---|
| 0.81218279 | 0.805964938 | 0 | 0 | 1.006860546 | 1.030641012 | 5S RNA |
| 14.0530501 | 18.59395166 | 12.26664502 | 6.965857106 | 23.22869824 | 31.42003479 | tRNA-Gly-T( |
| 17.33571238 | 20.38873464 | 25.50025531 | 5.940851105 | 48.55380892 | 34.2200671 | tRNA-Ile-GA |
| 13.19797033 | 15.71631629 | 13.4403055 | 3.052937373 | 37.63141292 | 33.49583288 | tRNA-Glu-T |
| 10.55837626 | 23.05059722 | 31.18150877 | 17.09644929 | 58.90134197 | 60.29249918 | tRNA-Pseuc |
| 10.27301474 | 12.74295915 | 7.192379702 | 8.16867027 | 23.87892242 | 26.88719558 | tRNA-Met-C |
| 5.206870487 | 6.458760118 | 9.27933421 | 10.5389071 | 20.97856015 | 18.99626687 | tRNA-Phe-G |
| 5.865764591 | 9.895458403 | 13.73897896 | 7.462735802 | 22.5424889 | 33.49583288 | tRNA-Tyr-G |
| 20.07578586 | 21.25023047 | 27.94069144 | 14.70569833 | 58.0717456 | 33.11841504 | tRNA-Trp-C |
| 33.84465816 | 32.9396766 | 40.43138477 | 21.07781419 | 86.3348437 | 85.07023857 | tRNA-His-G |
| 33.65482434 | 41.25533025 | 51.07316091 | 23.66026464 | 83.44356779 | 86.25176966 | tRNA-Gln-T |
| 25.38349362 | 14.85514827 | 15.24462049 | 7.527790784 | 22.59229555 | 28.90736262 | tRNA-Val-TA |
| 13.5750552 | 11.78723722 | 19.58444516 | 5.233606926 | 29.45067098 | 34.45285668 | tRNA-Leu-C |
| 9.242975624 | 48.81376851 | 12.9585677 | 3.880749173 | 253.0285496 | 137.1343053 | Small Subu |
| 12.36631741 | 13.56339625 | 11.26776297 | 9.033348941 | 41.15025261 | 55.3369513 | tRNA-Ala-T( |
| 11.84545106 | 55.52665465 | 13.26258246 | 6.774493263 | 303.1066303 | 54.64526785 | Large Subu |
| 0 | 0 | 0.827095723 | 0 | 4.027442186 | 0 | 5S RNA |
| 0.642063421 | 0 | 0 | 0.742606388 | 3.183856322 | 3.25905401 | tRNA-Asn-G |
| 1.284126843 | 1.274295915 | 0.6538527 | 0.742606388 | 3.183856322 | 2.444290507 | tRNA-Arg-A |
| 66.64975017 | 38.6359442 | 98.78624545 | 72.50726262 | 65.44593552 | 51.91854096 | tRNA-Arg-C |
| 17.59729377 | 11.05962998 | 12.54428514 | 14.9254716 | 16.72507241 | 41.68370314 | tRNA-Tyr-G |
| 15.8375644 | 7.203311631 | 16.1283666 | 12.21174949 | 9.816890328 | 18.42270808 | tRNA-Gln-T |
| 39.70238746 | 29.06442053 | 27.17519304 | 16.56113972 | 22.59229555 | 52.85917737 | tRNA-Asp-G |
| 18.47715846 | 16.3711628 | 20.83247353 | 26.71320202 | 45.81215486 | 46.89416603 | tRNA-Arg-C |
| 2.007578586 | 1.992209107 | 6.133322511 | 1.547968246 | 3.318385463 | 5.944330905 | tRNA-Thr-G |
| 0 | 0.654846512 | 3.360076376 | 0 | 7.362667746 | 1.674791644 | tRNA-Glu-T |
| 6.420634215 | 8.282923448 | 7.192379702 | 5.198244717 | 14.32735345 | 7.332871522 | tRNA-Ile-GA |
| 12.36631741 | 16.14690029 | 19.88428759 | 9.033348941 | 43.5708557 | 38.81845838 | tRNA-Thr-T( |
| 19.70038498 | 22.99948725 | 37.17391815 | 10.05235477 | 32.32390718 | 58.82195042 | tRNA-Leu-T |
| 23.43091719 | 28.41854452 | 31.81486015 | 11.29168618 | 29.04723713 | 50.38140343 | tRNA-Lys-T |
| 22.78005838 | 25.18916446 | 23.86114511 | 13.55002341 | 26.62663404 | 37.99253373 | tRNA-Asp-G |
| 18.2240467 | 23.25153642 | 23.19833553 | 9.786128019 | 12.90988317 | 22.29996545 | tRNA-Val-TA |
| 0 | 0 | 0 | 0 | 0 | 0 | 5S RNA |
| 24.73263481 | 25.83504047 | 20.54709718 | 12.04446525 | 25.01289864 | 56.9888006 | tRNA-Lys-T |
| 11.69820097 | 53.35104888 | 12.76273638 | 6.433876283 | 305.3174589 | 52.54831799 | Large Subu |

| | | | | | | |
|---|---|---|---|---|---|---|
| 10.41374097 | 13.56339625 | 7.290905451 | 7.527790784 | 35.50217872 | 56.9888006 | tRNA-Ala-T( |
| 9.939364198 | 48.18553468 | 14.6025651 | 4.759409363 | 251.3411694 | 139.4238938 | Small Subu |
| 57.88902848 | 62.32332321 | 82.31028474 | 48.00480835 | 91.39863409 | 81.08301614 | tRNA-Ser-G |
| 11.60193671 | 18.64028211 | 11.25234879 | 15.9746723 | 34.92986558 | 28.74409845 | tRNA-Leu-A |
| 0.642063421 | 0 | 0 | 0.742606388 | 3.183856322 | 1.629527005 | tRNA-Arg-A |
| 0.642063421 | 0 | 2.615410801 | 0.742606388 | 4.775784484 | 4.073817512 | tRNA-Asn-G |
| 0 | 0 | 1.654191447 | 0 | 1.006860546 | 1.030641012 | 5S RNA |
| 12.05814562 | 52.65290673 | 14.57884378 | 5.922950813 | 306.1084893 | 55.68336185 | Large Subu |
| 9.762882162 | 14.20927226 | 8.616524624 | 16.56113972 | 30.66097253 | 53.68510201 | tRNA-Ala-T( |
| 21.43865424 | 17.24961544 | 32.45342061 | 17.42408159 | 37.35207052 | 65.43941984 | tRNA-Leu-T |
| 10.25590446 | 51.10682198 | 15.98868055 | 2.562758888 | 254.2057916 | 136.9334642 | Small Subu |
| 258.3909479 | 268.0385449 | 365.2080821 | 183.6780951 | 744.7388854 | 759.02475 | tRNA-Lys-C |
| 2.676771447 | 1.328139405 | 5.451842232 | 13.93171421 | 9.125560023 | 7.642711164 | tRNA-Cys-G |
| 14.76745869 | 7.645775491 | 17.0001702 | 5.940851105 | 54.12555748 | 23.62814157 | tRNA-Asn-G |
| 3.959391099 | 1.964539536 | 4.704106926 | 1.526468687 | 4.908445164 | 6.699166576 | tRNA-Glu-T |
| 26.9666637 | 23.57447443 | 22.23099181 | 16.33734054 | 44.57398851 | 39.10864812 | tRNA-Arg-T( |
| 16.27147027 | 14.85514827 | 24.5239547 | 12.79724433 | 44.3777234 | 42.12215696 | tRNA-Thr-T( |

nal replication initiator protein DnaA
59: hypothetical protein
rouridine synthase B (EC 1.-.-.-)

ein
cal phospholipase A2; _Toximoron (Other)
cal pyrogenic exotoxin K (SpeK)

ein

n
ein
22: hypothetical protein
n (heat shock protein 33)
uronidase
e lyase (phage associated)
uronidase; Phage endopeptidase
ed protein
ength tape-measure protein
ein
ein
r tail protein
ein
ein
onal regulator, RofA-like Protein (RALP)
ein
ging protein, phage associated
hypothetical protein - phage associated
d protein
lp protease-like protein
al protein
ein
al phage protein
inase, large subunit

inase, small subunit
llagen binding protein
nuclease
33: hypothetical protein
al protein - phage associated
al protein
ethyltransferase
89: hypothetical protein
hypothetical protein - phage associated
22: hypothetical protein
al protein
plication protein
idase I (EC 3.4.21.89)
ein
22: hypothetical protein
10: hypothetical protein
15: hypothetical protein
al phage protein
al phage protein
al phage protein
al phage protein
al protein, phage associated
al phage protein
n-binding protein/major pilin backbone protein T-antigen
al protein
al protein
al protein
hypothetical protein - phage associated
scriptional regulator, Cro/CI family
al phage protein
al protein
40: hypothetical protein
al protein
regulator of exopolysaccharide synthesis, competence and biofilm formation Ftr, XRE family
cific sortase B

al protein
grase: site-specific recombinase
, putative
in phosphorylase (EC 2.4.1.1)
canotransferase (amylomaltase) (EC 2.4.1.25)
eron transcriptional repressor MalR, LacI family
altodextrin ABC transporter, substrate binding periplasmic protein MalE
altodextrin ABC transporter, permease protein MalF
altodextrin ABC transporter, permease protein MalG
nent protein
82: hypothetical protein
nent protein
e (maltose area)
terase
ophosphate chain) D-alanine transfer protein DltD
poly(phosphoribitol) ligase subunit 2 (EC 6.1.1.13)
nsfer protein DltB
poly(phosphoribitol) ligase subunit 1 (EC 6.1.1.13)
14: hypothetical protein
se ABC subunit B
ABC transporter, glutamine-binding protein/permease protein
gar metabolism regulator
transport ATP-binding protein
24: hypothetical protein
cellobiose-specific IIC component (EC 2.7.1.69)
characterized protein SPs0857
cellobiose-specific IIA component (EC 2.7.1.69)
cellobiose-specific IIB component (EC 2.7.1.69)
onal antiterminator of lichenan operon, BglG family
ce protein of unknown function, cellobiose operon
sidase (EC 3.2.1.21)
otinamide transporter, PnuC-like
n-binding protein
98: hypothetical protein
g protein Obg

47: hypothetical protein
nent protein
nent protein
nent protein
e phosphoribosyltransferase [decarboxylating] (EC 2.4.2.19)
ermease
nent protein
nent protein
99: hypothetical protein
39: hypothetical protein
nent protein
small subunit pseudouridine synthase A (EC 4.2.1.70)
pamide acyltransferases
r, MFS superfamily
dent Zn protease
ic acid degradation protein PaaD, thioesterase
osphorylase (EC 2.4.2.3)
Uracil-5-) -methyltransferase RumA (EC 2.1.1.-) ## LSU rRNA m(5)U1939
pe-associated transcriptional attenuator LytR-CpsA-Psr, subfamily F1 (as in PMID19099556)
kinase I (EC 2.7.1.71)
fatty acids transporter
vylshikimate-3-phosphate synthase (EC 2.5.1.19)
brane protein YihY, formerly thought to be RNase BN
e aminopeptidase (EC 3.4.11.18)
: Cytosolic protein containing multiple CBS domains
protein-S5p-alanine acetyltransferase
elated alpha 2 macroglobulin-binding protein (GRAB)
ylglucosamine 1-carboxyvinyltransferase (EC 2.5.1.7)
methionine synthetase (EC 2.5.1.6)
putative
on repressor / Biotin-protein ligase (EC 6.3.4.15)
onal regulators, LysR family
35: hypothetical protein
erase III subunits gamma and tau (EC 2.7.7.7)
n-containing proteins

56: hypothetical protein
65: hypothetical protein
ase (EC 2.7.1.48) [C1]
dent RNA helicase YfmL
can N-acetylglucosamine deacetylase (EC 3.5.1.-)
horylating glyceraldehyde-3-phosphate dehydrogenase (NADP) (EC 1.2.1.9)
olpyruvate-protein phosphotransferase of PTS system (EC 2.7.3.9)
CoA thiolase (EC 2.3.1.16) @ Acetyl-CoA acetyltransferase (EC 2.3.1.9)
rrier protein of PTS system
in-like protein NrdH, required for reduction of Ribonucleotide reductase class Ib
otide reductase of class Ib (aerobic), alpha subunit (EC 1.17.4.1)
45: hypothetical protein
otide reductase of class Ib (aerobic), beta subunit (EC 1.17.4.1)
annel protein
nent protein
nent protein
-bound protease, CAAX family
75: hypothetical protein
acetoacetyl-CoA transferase, alpha subunit (EC 2.8.3.8)
onal regulator, Cro/CI family
22: hypothetical protein
A synthetase (EC 6.1.1.7)
otein prsA 1 precursor (EC 5.2.1.8)
ansferase family protein [C1]
tator:Oxalate:Formate Antiporter
peptidase F
e protein CoiA
small subunit pseudouridine synthase A (EC 4.2.1.70)
ne-6-phosphate deaminase (EC 3.5.99.6)
etoacetate CoA-transferase subunit B (EC 2.8.3.9)
methionine:tRNA ribosyltransferase-isomerase (EC 5.-.-.-)
59: hypothetical protein
92: hypothetical protein
64: hypothetical protein
92: hypothetical protein

superoxide dismutase (EC 1.15.1.1)
erase III delta subunit (EC 2.7.7.7)
:tence protein ComEC, DNA transport
:tence protein ComEA, DNA receptor
ycerol-3-phosphate acyltransferase (EC 2.3.1.51)
AS domain
ine37-N(6))-methyltransferase TrmN6 (EC 2.1.1.223)
utative endonuclease containing a URI domain
potassium uptake protein
DEAD-box protein A
in release factor 3
protein, putative
ylmuramoylalanyl-D-glutamyl-2,6-diaminopimelate--D-alanyl-D-alanine ligase (EC 6.3.2.10)
D-alanine ligase A (EC 6.3.2.4)
tion protein RecR
trite family of transporters
clease L-PSP
pecific component NiaX of predicted niacin ECF transporter
al protein
onal repressor for NAD biosynthesis in gram-positives
cerate mutase (EC 5.4.2.1)
45: hypothetical protein
otate dehydrogenase, catalytic subunit (EC 1.3.3.1)
17: hypothetical protein
ium, zinc and mercury transporting ATPase (EC 3.6.3.3) (EC 3.6.3.5); Copper-translocating P-type ATPase (EC 3.6.3.4)

cal extracellular nuclease 2; Mitogenic factor 2
onal regulator pfoR
cal pyrogenic exotoxin C (SpeC); _Toximoron (Superantigen)
, N-acetylmuramoyl-L-alanine amidase (EC 3.5.1.28)
ein
22: hypothetical protein
ronidase
e lyase (phage associated)
ronidase; Phage endopeptidase

: Phage-associated protein
synthase subunit G (EC 3.6.3.14)
›r tail protein
ein
ıypothetical protein - phage associated
r tail protein
: Phage-associated protein
: Phage protein
: Phage protein
: Phage protein
: Phage-associated protein
09: hypothetical protein
sidine synthetase
synthase subunit I (EC 3.6.3.14)
al protein
ıypothetical protein - phage associated
ein
ein
tional DNA repair protein RecT (prophage associated)
09: hypothetical protein
ein
al protein
74: hypothetical protein
ation protein dnaD
synthase subunit K (EC 3.6.3.14)
ein
ionase
age transcriptional repressor protein
te repressor
ein
grase
ng protein HBsu

: Lipase/Acylhydrolase with GDSL-like motif
al protein DUF194, DegV family

synthase subunit E (EC 3.6.3.14)
48: hypothetical protein
protein RecN
thway regulatory protein ArgR, repressor of arg regulon
g methyltransferase FtsJ like
diphosphate synthase (EC 2.5.1.90) / Dimethylallyltransferase (EC 2.5.1.1) / (2E,6E)-farnesyl diphosphate synthase (EC 2.5.1.10) / Geranyl
bonuclease VII small subunit (EC 3.1.11.6)
bonuclease VII large subunit (EC 3.1.11.6)
etrahydrofolate dehydrogenase (NADP+) (EC 1.5.1.5) / Methenyltetrahydrofolate cyclohydrolase (EC 3.5.4.9)
osphomannomutase
lipyrimidine photolyase (EC 4.1.99.3)
synthase subunit C (EC 3.6.3.14)
transport ATP-binding protein
orter membrane-spanning permease - glutamine transport
88: hypothetical protein
dent Clp protease, ATP-binding subunit ClpE
utT protein (7,8-dihydro-8-oxoguanine-triphosphatase) (EC 3.6.1.-)
27: hypothetical protein
RNA synthetase (EC 6.1.1.5)
n initiation protein DivIVA
: hypothetical protein, contains S4-like RNA binding domain
n protein YlmG/Ycf19 (putative), YggT family
synthase subunit F (EC 3.6.3.14)
: FtsZ-interacting protein related to cell division
al protein YggS, proline synthase co-transcribed bacterial homolog PROSC
n protein FtsZ (EC 3.4.24.-)
n protein FtsA
n protein FtsQ
ylglucosamine--N-acetylmuramyl-(pentapeptide) pyrophosphoryl-undecaprenol N-acetylglucosamine transferase (EC 2.4.1.227)
ylmuramoylalanine--D-glutamate ligase (EC 6.3.2.9)
39: hypothetical protein
g protein TypA/BipA
-like domain protein
al protein
e (EC 2.7.1.2)

76: hypothetical protein
c DNA-binding protein Dps / Iron-binding ferritin-like antioxidant protein / Ferroxidase (EC 1.16.3.1)
:tence protein ComC, processing protease
RNA large subunit methyltransferase N (EC 2.1.1.-)
al DUF1027 domain protein
63: hypothetical protein
regulatory protein
otease with PDZ domain
ntetheine adenylyltransferase (EC 2.7.7.3)
synthase subunit A (EC 3.6.3.14)
guanine(966)-N(2))-methyltransferase (EC 2.1.1.171) ## SSU rRNA m(2)G966
ammonia ligase (EC 6.3.1.1)
kinase (EC 2.7.2.2)
hine deacetylase/Succinyl-diaminopimelate desuccinylase and related deacylases
nithine antiporter ArcD
arbamoyltransferase (EC 2.1.3.3)
ferase (EC 2.3.1.-)
iminase (EC 3.5.3.6)
onal regulator ArcR essential for anaerobic expression of the ADI pathway, Crp/Fnr family
thway regulatory protein ArgR, repressor of arg regulon
synthase subunit B (EC 3.6.3.14)
characterized protein gbs2121
al protein SpyM3_1200
onent sensor kinase yesM (EC 2.7.3.-), associated with MetSO reductase
onent response regulator yesN, associated with MetSO reductase
thionine sulfoxide reductase MsrA (EC 1.8.4.11) / Peptide methionine sulfoxide reductase MsrB (EC 1.8.4.12)
ide oxidoreductase associated with MetSO reductase
e c-type biogenesis protein CcdA homolog, associated with MetSO reductase
and cobalt transport protein CorA
ctase, Gfo/Idh/MocA family
15: hypothetical protein
synthase subunit D (EC 3.6.3.14)
synthetase (EC 6.1.1.9)
al protein
al protein

59: hypothetical protein
imerase (EC 5.1.3.3)
eoxy-D-arabino-heptulosonate-7-phosphate synthase I beta (EC 2.5.1.54)
quinate synthase (EC 4.2.3.4)
erized conserved protein, contains double-stranded beta-helix domain
al protein
dent methyltransferase HI0095 (UbiE paralog)
xic anion resistance protein
quinate 5-dehydrogenase I beta (EC 1.1.1.282)
osidase (EC 3.2.1.23)
g response regulator, AraC family
onent sensor kinase YesM (EC 2.7.3.-)
mbrane protein
gar ABC transporter, substrate-binding protein
gar ABC transporter, membrane-spanning permease protein MsmG
gar ABC transporter, membrane-spanning permease protein MsmF
nnosamine kinase (EC 2.7.1.60)
49: hypothetical protein
ine-guanine phosphoribosyltransferase (EC 2.4.2.8)
: hypothetical protein
sidase (EC 3.2.1.21)
glucosaminidase (EC 3.2.1.35)
onal repressor of arabinoside utilization operon, GntR family
al protein
nosidase (EC 3.2.1.24)
o-component responsible histidine kinase
ltransferase, TrmA family

cal extracellular nuclease 3; Mitogenic factor 3
56: hypothetical protein
cinate synthetase (EC 6.3.4.4)
ein
ein

n

ein
22: hypothetical protein
uronidase
e lyase (phage associated)
peptidase
ed protein
-binding protein
ength tape-measure protein
ein
ein
r tail shaft protein
r capsid protein
r capsid protein #Fam0017
r capsid protein
ein
r capsid protein
ein
on antitermination protein NusG
ein
al protein, phage associated
r capsid protein
r capsid protein #Fam0015
inase large subunit
ein
33: hypothetical protein
al protein
enine dinucleotide glycohydrolase (NADGH) (EC 3.2.2.5)
hypothetical protein - phage associated
ein
ein
ein
46: hypothetical protein
22: hypothetical protein
day junction resolvase
nded DNA-binding protein

ıded DNA-binding protein
al protein (phage associated)
cal NAD glycohydrolase inhibitor
17: hypothetical protein
al phage protein
al phage protein
ation protein DnaC
age replication protein
al protein, phage associated
ein
ıypothetical protein - phage associated
al protein - phage associated
epressor protein
ated cytolysin
62: hypothetical protein
like repressor - phage associated
ein
ne segregation ATPase
;rase: site-specific recombinase
protein recX
RNA synthetase related protein
subunit interface protein
e protein F homolog, phosphoribosyltransferase domain; protein YhgH required for utilization of DNA as sole source of carbon and energ
on protein A, DNA transporter ATPase
43: hypothetical protein
: protein co-occurring with transport systems (COG1739)
ıthase (EC 2.5.1.47)
ess protein 13
cal lipoprotein rotamase A; Peptidyl-prolyl cis-trans isomerase (EC 5.2.1.8)
onent transcriptional regulator VraR
dine kinase VraS
associated with VraSR
onine protein kinase PrkC, regulator of stationary phase
ne/threonine phosphatase PrpC, regulation of stationary phase
ytosine(967)-C(5))-methyltransferase (EC 2.1.1.176)

03: hypothetical protein
tRNA formyltransferase (EC 2.1.2.9)
A essential for oriC/DnaA-independent DNA replication
ed RNA polymerase omega subunit (EC 2.7.7.6)
kinase (EC 2.7.4.8)
HAD superfamily)
onal regulators, LysR family
CoA thiolase (EC 2.3.1.16) @ Acetyl-CoA acetyltransferase (EC 2.3.1.9)
etoacetate CoA-transferase subunit A (EC 2.8.3.9)
etoacetate CoA-transferase subunit B (EC 2.8.3.9)
ne beta-lyase (EC 4.4.1.8)
roxybutyrate permease
mocysteine lyase (EC 4.4.1.21) / Autoinducer-2 production protein LuxS
al protein SPy1643
: Predicted N6-adenine-specific DNA methylase
n protein GpsB, coordinates the switch between cylindrical and septal cell wall synthesis by re-localization of PBP1
: hypothetical protein
lay junction resolvase
lar transpeptidase-transglycosylase (EC 2.4.1.129) (EC 3.4.-.-) / Penicillin-binding protein 1A/1B (PBP1)
idase C (EC 3.4.22.40)
n protein FtsH (EC 3.4.24.-)
A synthetase (EC 6.1.1.4)
etase (EC 6.3.1.5)
phosphoribosyltransferase (EC 2.4.2.11)
ansport protein (permease)
n reductase (EC 1.8.1.9)
al protein SPy1656
ultidrug transport system, ATPase component
C transporter, permease protein
65: hypothetical protein
dent RNA helicase YqfR
-acetylmuramoyl-pentapeptide-transferase (EC 2.7.8.13)
specific PTS system, EIIC component
n protein FtsI [Peptidoglycan synthetase] (EC 2.4.1.129)
n protein FtsL

subunit methyltransferase H
63: hypothetical protein
tamyl phosphate reductase (EC 1.2.1.41)
5-kinase (EC 2.7.2.11)
orter permease protein
e ABC transporter ATP-binding protein
61: hypothetical protein
ise (EC 2.2.1.1)
specific PTS system, EIIB component (EC 2.7.1.69)
ise (EC 2.2.1.2)
g positive regulator
xidase (EC 1.11.1.1)
take facilitator protein
erophosphate oxidase (EC 1.1.3.21)
nase (EC 2.7.1.30)
32: hypothetical protein
27: hypothetical protein
18: hypothetical protein
A synthetase beta chain (EC 6.1.1.14)
specific PTS system, EIIA component (EC 2.7.1.-)
A synthetase alpha chain (EC 6.1.1.14)
MEMBRANE PROTEIN (Rhomboid family)
ctase of aldo/keto reductase family, subgroup 1
cosamine-6-phosphate deacetylase (EC 3.5.1.25)
pendent phosphate transporter
30: hypothetical protein
y protein
onal regulator, TetR family
HAD superfamily)
MEMBRANE PROTEIN (Rhomboid family)
lonate 6-phosphate decarboxylase
al protein
6-diphosphate aldolase (EC 4.1.2.40)
phosphate kinase (EC 2.7.1.144)
6-phosphate isomerase, LacB subunit (EC 5.3.1.26)

-phosphate isomerase, LacA subunit (EC 5.3.1.26)
galactose-specific IIC component (EC 2.7.1.69)
galactose-specific IIB component (EC 2.7.1.69)
galactose-specific IIA component (EC 2.7.1.69)
osphotransferase system repressor
perone
-phosphate 3-epimerase (EC 5.1.3.-)
nslocating P-type ATPase (EC 3.6.3.4)
anscriptional regulator-copper transport operon
terase
binding factor A
initiation factor 2
protein L7Ae family protein
Predicted nucleic-acid-binding protein implicated in transcription termination
on termination protein NusA
: clustered with transcription termination protein NusA
nent protein
-phosphate 4-epimerase (EC 5.1.3.4)
ine46-N7-)-methyltransferase (EC 2.1.1.33)
oside phosphotransferase family protein
orter, permease protein EscB
orter, ATP-binding protein EcsA
iad (HIT) nucleotide-binding protein, similarity with At5g48545 and yeast YDL125C (HNT1)
al protein in cluster with Ecs transporter (in Streptococci)
oe-associated transcriptional attenuator LytR-CpsA-Psr, subfamily F2 (as in PMID19099556)
ferase (EC 2.3.1.-)
, predicted to have essential role in cell wall biosynthesis
ypoxanthine permease
onal antiterminator with PTS regulation domain, SPy0181 ortholog
HAD superfamily)
mannose-specific IIB component (EC 2.7.1.69) / PTS system, mannose-specific IIA component (EC 2.7.1.69)
mannose-specific IIC component (EC 2.7.1.69)
mannose-specific IID component (EC 2.7.1.69)
gulator of the mannose operon, ManO
nent protein

synthetase (EC 6.1.1.11)
ızyme A carboxyl transferase alpha chain (EC 6.4.1.2)
ızyme A carboxyl transferase beta chain (EC 6.4.1.2)
oxylase of acetyl-CoA carboxylase (EC 6.3.4.14)
ascorbate-6-phosphate lactonase UlaG (EC 3.1.1.-) (L-ascorbate utilization protein G)
ymyristoyl-[acyl carrier protein] dehydratase (EC 4.2.1.-)
oxyl carrier protein of acetyl-CoA carboxylase
acyl-carrier-protein] synthase, KASII (EC 2.3.1.41)
acyl-carrier protein] reductase (EC 1.1.1.100)
A-acyl carrier protein transacylase (EC 2.3.1.39)
-carrier-protein] reductase [FMN] (EC 1.3.1.9)
protein
acyl-carrier-protein] synthase, KASIII (EC 2.3.1.41)
onal regulator of fatty acid biosynthesis FabT
3-Decenoyl-ACP isomerase
aine ABC transport system, ATP-binding protein OpuAA (EC 3.6.3.32)
protein DnaJ
protein DnaK
protein GrpE
ible transcription repressor HrcA
acetyl-muramidase
alanine carboxypeptidase (EC 3.4.16.4)
cerate mutase family
mbrane protein
al protein
NA(Asn) amidotransferase subunit B (EC 6.3.5.6) @ Glutamyl-tRNA(Gln) amidotransferase subunit B (EC 6.3.5.7)
ino acid transporter - APC Superfamily
aine ABC transport system, permease protein OpuAB / Glycine betaine ABC transport system, glycine betaine-binding protein OpuAC
NA(Asn) amidotransferase subunit A (EC 6.3.5.6) @ Glutamyl-tRNA(Gln) amidotransferase subunit A (EC 6.3.5.7)
NA(Asn) amidotransferase subunit C (EC 6.3.5.6) @ Glutamyl-tRNA(Gln) amidotransferase subunit C (EC 6.3.5.7)
nent protein
osphate dikinase (EC 2.7.9.1)
dase (EC 3.5.1.19)
transcriptional pleiotropic repressor codY
minotransferase (EC 2.6.1.1)

:ress protein family
HAD superfamily)
ase (EC 3.5.1.1)
erase I (EC 2.7.7.7)
orter permease protein
C transporter (ATP-binding protein)
dent DNA helicase RecG (EC 3.6.1.-)
ATPase component BL0693 of energizing module of predicted ECF transporter
orane component BL0694 of energizing module of predicted ECF transporter
pecific component BL0695 of predicted ECF transporter
C transporter ATP-binding protein, spy1790 homolog
C transporter (ATP-binding protein), spy1791 homolog
:ransporter (Streptococcus), ATP-binding protein
:ransporter (Streptococcus), permease protein
oA synthetase, alpha subunit-related enzymes
porter IsdDEF, lipoprotein IsdE
protein Shp, transfers heme from hemoglobin to apo-SiaA/HtsA
cal cell surface hemoprotein receptor Shr
al protein
nic secreted protein
emase (EC 5.1.1.1)
:arrier protein] synthase (EC 2.7.8.7)
ort cytoplasm protein SecA ATPase RNA helicase (TC 3.A.5.1.1)
nent protein
-phosphate isomerase (EC 5.3.1.8)
ress regulator PerR, FUR family
se (EC 2.7.1.4)
do-beta-N-acetylglucosaminidase (EndoS)
sucrose-specific IIB component (EC 2.7.1.69) / PTS system, sucrose-specific IIC component (EC 2.7.1.69) / PTS system, sucrose-specific II/
phosphate hydrolase (EC 3.2.1.26)
eron repressor ScrR, LacI family
on termination protein NusB
ock protein
elongation factor P
ninase (EC 3.5.4.12); Late competence protein ComEB

idase YpdF (MP-, MA-, MS-, AP-, NP- specific)
rans-acting positive regulator (Mga)
se ABC subunit A
and cobalt transport protein CorA
mbrane protein
nal protein S18p @ SSU ribosomal protein S18p, zinc-independent
nded DNA-binding protein
nal protein S6p
al protein
c adenine glycosylase (EC 3.2.2.-)
onal regulator, XRE family
n
r of prophage gene expression IbrA
ylglycerophosphatase B (EC 3.1.3.27)
tion inhibitory protein MutS2
roduction protein
25: hypothetical protein
se HIII (EC 3.1.26.4)
gnal peptidase I (EC 3.4.21.89)
NA helicase YrrC
embrane protein precursor SPs0273
erase IV (EC 2.7.7.7)
rmate-lyase (EC 2.3.1.54)
r of prophage gene expression IbrB
al protein
nase, class C
ng proteinase
mbrane protein
(Major Intrinsic Protein Family)

onal regulator, CrP/Fnr family
eptidyl-peptidase (EC 3.4.14.11)
onal regulator, Cro/CI family
pressor - phage associated
69: hypothetical protein

ɔnal regulatory protein
ɔnal regulator, MerR family
erase III epsilon subunit (EC 2.7.7.7)
ɔnal regulator OrfX
ıtilizing dehydrogenases
e-phosphate aldolase (EC 4.1.2.4)
permease NupC
ɔsphorylase (EC 2.4.2.3)
egulator for deoxynucleoside utilization, GntR family
nal protein S14p (S29e) @ SSU ribosomal protein S14p (S29e), zinc-independent
nent protein
Qri7 protein, required for threonylcarbamoyladenosine t(6)A37 formation in tRNA
protein-S18p-alanine acetyltransferase (EC 2.3.1.-)
molog of metal-dependent proteases, putative molecular chaperone
ınknown function DUF1447
se J1 (endonuclease and 5' exonuclease)
synthetase type I (EC 6.3.1.2)
ɔnal regulator, MerR family
49: hypothetical protein
ɣcerate kinase (EC 2.7.2.3)
hatase (EC 3.1.3.2)
nent protein
ɔmatin/prohibitin-family membrane protease subunit YbbK
ıcetone kinase family protein
: Putative alkaline-shock protein
nal protein L28p
sphosphate aldolase class II (EC 4.1.2.13)
of the alpha/beta superfamily
al protein
se (EC 6.3.4.2)
ed RNA polymerase delta subunit (EC 2.7.7.6)
ı trigger factor (EC 5.2.1.8)
tigen GbpB/SagA/PcsB, putative peptidoglycan hydrolase
nent protein
efflux system KefA protein / Small-conductance mechanosensitive channel

al protein ywlG
al protein
pecific component PdxU2 of predicted pyridoxin-related ECF transporter
oxal kinase, thiD family (EC 2.7.1.35)
ouridine synthase A (EC 4.2.1.70)
:e-specific sigma factor ComX
inase (EC 2.7.1.31)
ction-modification system, restriction subunit R (EC 3.1.21.3)
ction-modification system, specificity subunit S (EC 3.1.21.3)
ermease
ction-modification system, DNA-methyltransferase subunit M (EC 2.1.1.72)
onal regulator DegU, LuxR family
14: hypothetical protein
A modification enzyme
salivaricin A
-beta-galactosidase (EC 3.2.1.85)
lactose-specific IIB component (EC 2.7.1.69) / PTS system, lactose-specific IIC component (EC 2.7.1.69)
lactose-specific IIA component (EC 2.7.1.69)
e phosphoribosyltransferase [decarboxylating] (EC 2.4.2.19)
6-diphosphate aldolase (EC 4.1.2.40)
phosphate kinase (EC 2.7.1.144)
i-phosphate isomerase, LacB subunit (EC 5.3.1.26)
i-phosphate isomerase, LacA subunit (EC 5.3.1.26)
osphotransferase system repressor
ge-inducible protein J
IA-damage-inducible protein J
al protein

ınknown function DUF208
nent protein
mal protein S9p (S16e)
mal protein L13p (L13Ae)
onal regulator, Cro/CI family
al protein
y protein in cluster with TrmH family tRNA/rRNA methyltransferase YacO

al protein DUF901, similar to C-terminal domain of ribosome protection-type Tc-resistance proteins
;uanosine-2'-O-) -methyltransferase rlmB (EC 2.1.1.-) ## LSU rRNA Gm2251
al protein, Spy1939 homolog
Ribonuclease III family protein
RNA synthetase (EC 6.1.1.16)
nent protein
32: hypothetical protein
yltransferase (EC 2.3.1.30)
29: hypothetical protein
cleotide nucleotidyltransferase (EC 2.7.7.8)
ise (EC 2.2.1.2)
rotein SgaT, putative
IIB component
on antiterminator, BglG family (EC 2.7.1.69)
mal protein S15p (S13e)
48: hypothetical protein
nent protein
58: hypothetical protein
formylase (EC 3.5.1.88)
77: hypothetical protein
onal regulator, MarR family
erase III alpha subunit (EC 2.7.7.7)
synthetase (EC 6.1.1.15), bacterial type
rane protease RasP/YluC, implicated in cell division based on FtsL cleavage
ate cytidylyltransferase (EC 2.7.7.41)
nyl diphosphate synthase (EC 2.5.1.31)
translocase subunit YajC (TC 3.A.5.1.1)
protein
ansport accessory protein
(EC 3.2.1.41)
-alpha-glucosidase (EC 3.2.1.70)
gar ABC transporter, ATP-binding protein
n protein
ise
RNA(Tyr) deacylase

nosphokinase (EC 2.7.6.5), (p)ppGpp synthetase I
ke surface protein
tide reduction protein NrdI
67: hypothetical protein
al protein
bonuclease III (EC 3.1.11.2)
maltose and glucose-specific IIC component (EC 2.7.1.69) / PTS system, maltose and glucose-specific IIB component (EC 2.7.1.69) / PTS s
RNA small subunit methyltransferase E (EC 2.1.1.-)
protein L11 methyltransferase (EC 2.1.1.-)
20: hypothetical protein
kanoate-cyclic-dimer hydrolase (EC 3.5.2.12)
benzoate synthase, aminase component (EC 2.6.1.85) / Aminodeoxychorismate lyase (EC 4.1.3.38)
e synthase, amidotransferase component (EC 4.1.3.27) @ Para-aminobenzoate synthase, amidotransferase component (EC 2.6.1.85) # Tr
ne segregation helicase
ne transglycosylase (EC 2.4.2.29)
binding ABC transporter, periplasmic substrate-binding component (TC 3.A.1.5.2)
ransport system permease protein DppB (TC 3.A.1.5.2)
ransport system permease protein DppC (TC 3.A.1.5.2)
ransport ATP-binding protein DppD (TC 3.A.1.5.2)
ransport ATP-binding protein DppF (TC 3.A.1.5.2)
96: hypothetical protein
cal histidine triad protein
nding surface protein
al protein
protein
n-binding protein
ase (EC 3.4.21.-)
ytic M protein
ytic M protein
rans-acting positive regulator (Mga)
al mga-associated protein
nic secreted protein
onent system histidine kinase
sphate pyrophosphokinase (EC 2.7.6.1)
pecific component BioY of biotin ECF transporter

onent system response regulator
orter permease protein
orter ATP-binding protein
: component of efflux system
31: hypothetical protein
35: hypothetical protein
45: hypothetical protein
oline cis-trans-isomerase (EC 5.2.1.8); Proteinase maturation protein (RopA)
n inhibitor
ccal cysteine protease (Streptopain) (EC 3.4.22.10) / Streptococcal pyrogenic exotoxin B (SpeB)
ndent hydrolase (EC 3.-.-.-)
05: hypothetical protein
al protein
ike transcription regulator
nase B; Mitogenic factor 1
19: hypothetical protein
rature requirement C protein
hydrogenase (EC 1.1.1.6)
ise (EC 2.2.1.2)
rmate-lyase (EC 2.3.1.54)
cellobiose-specific IIC component (EC 2.7.1.69)
fic adenosine-34 deaminase (EC 3.5.4.-)
cellobiose-specific IIB component (EC 2.7.1.69)
cellobiose-specific IIA component (EC 2.7.1.69)
nucleoside regulator DeoR (transcriptional repressor)
onal regulator, DeoR family
rmate-lyase activating enzyme (EC 1.97.1.4)
81: hypothetical protein
translocase subunit SecE (TC 3.A.5.1.1)
lar transpeptidase-transglycosylase (EC 2.4.1.129) (EC 3.4.-.-)
al protein SPy2060
77: hypothetical protein
nology domain / putative murein endopeptidase
ibosomal large subunit pseudouridine synthase D, Bacillus subtilis YhcT type
al protein SPy2065

e (EC 3.4.-.-)
onal regulator, MutR family
onal regulator, MutR family
al protein
protein 60 family chaperone GroEL
protein 60 family co-chaperone GroES
dent Clp protease ATP-binding subunit ClpA
onal regulator CtsR
cal pyrogenic exotoxin G (SpeG)
al protein
protein CspA
peroxide reductase protein C (EC 1.6.4.-)
peroxide reductase protein F (EC 1.6.4.-)
epropionase (EC 3.5.2.7)
hydratase (EC 4.2.1.49)
formiminotransferase (EC 2.1.2.5) @ Glutamate formyltransferase
tetrahydrofolate cyclodeaminase (EC 4.3.1.4)
etrahydrofolate ligase (EC 6.3.4.3)
protein
al protein
tionic amino acid transporter protein
nmonia-lyase (EC 4.3.1.3)
glutamase (EC 3.5.3.8)
onal regulator, LuxR family
mal protein S2p (SAe)
elongation factor Ts
lopeptidase O (EC 3.4.24.-)
-phosphate hydrolase (EC 3.2.1.93)
trehalose-specific IIB component (EC 2.7.1.69) / PTS system, trehalose-specific IIC component (EC 2.7.1.69) / PTS system, trehalose-speci
peron transcriptional repressor
al cytosolic protein
95: hypothetical protein
17: hypothetical protein
onal regulator, HxlR family
family protein

rotein YaaA
otide reductase of class III (anaerobic), activating protein (EC 1.97.1.4)
ferase (EC 2.3.1.-)
idoreductase
81: hypothetical protein
otide reductase of class III (anaerobic), large subunit (EC 1.17.4.2)
phosphate isomerase (EC 5.3.1.9)
e-induced protein Ccs4
al protein
f S. aureus MRSA252 (BX571856) SAR1694
olliday junction resolvase YggF
al protein possible functionally linked with Alanyl-tRNA synthetase
onal regulator SpxA2
n
erin binding motif, CinA N-terminal domain / C-terminal domain of CinA type S
hyladenine glycosylase (EC 3.2.2.20)
nction DNA helicase RuvA
onal regulator, RofA-like Protein (RALP)
esistance protein B
tch repair protein MutL
grase: site-specific recombinase
al protein
scriptional repressor
hypothetical protein - phage associated
n, putative phage-related DNA binding protein
epressor protein
51: hypothetical protein
nent protein
80: hypothetical protein
42: hypothetical protein
67: hypothetical protein
ein
19: hypothetical protein
73: hypothetical protein
ein

se, phage associated
28: hypothetical protein
ein
61: hypothetical protein
bination and repair protein RecO
al protein
81: hypothetical protein
40: hypothetical protein
ded transcriptional regulator, ArpU family
al protein
atch repair protein MutS
nation protein YmcA
thway regulatory protein ArgR, repressor of arg regulon
IA synthetase (EC 6.1.1.19)
immunity protein
[illegible]
mbrane protein (Rhomboid family)
protein, putative
mbrane protein
NA synthetase (EC 6.1.1.12)
NA synthetase (EC 6.1.1.21)
mal protein L32p
mal protein L33p @ LSU ribosomal protein L33p, zinc-independent
esistance protein
fflux system accessory protein
19: hypothetical protein
E family
se-1-phosphate uridylyltransferase (EC 2.7.7.9)
48: hypothetical protein
onal regulator
: membrane protein
drolase (MutT/nudix family protein)
: Transcriptional regulator, PadR family
Integral membrane protein (putative)
: hypothetical membrane associated protein

tion protein
onal regulator, TetR family
nal protein S4p (S9e)
phosphate dehydrogenase [NAD(P)+] (EC 1.1.1.94)
nent protein
55: hypothetical protein
DNA helicase (DnaB) (EC 3.6.4.12)
nal protein L9p
terase, DHH family protein
e 5-carboxymethylaminomethyl modification enzyme GidA
family protein
fic 2-thiouridylase MnmA
hydratase, beta subunit (EC 4.3.1.17)
hydratase, alpha subunit (EC 4.3.1.17)
onal regulator, MarR family
minant antigen A
brane component of general energizing module of ECF transporters
ponent of general energizing module of ECF transporters
ponent of general energizing module of ECF transporters
glycerol--glycerol-3-phosphate 3-phosphatidyltransferase (EC 2.7.8.5)
onal regulator in cluster with unspecified monosaccharide ABC transport system
: peptidase, M16 family
: Zinc protease
synthase (EC 2.4.1.212)
e dehydrogenase in hyaluronic acid synthesis (EC 1.1.1.22)
ort ATP-binding/permease protein MsbA
67: hypothetical protein
se-1-phosphate uridylyltransferase (EC 2.7.7.9)
: hypothetical protein
bination and repair protein RecF
gar uptake protein SpyM3_1856/SPs1852
monophosphate dehydrogenase (EC 1.1.1.205)
yl-tRNA synthetase (EC 6.1.1.2)
71: hypothetical protein
al protein BH3604

orter ATP-binding protein uup
ort ATP-binding/permease protein MsbA
orter, permease protein
nent protein
.915 methyltransferase RlmH
ease, DegP/HtrA, do-like (EC 3.4.21.-)
ne (plasmid) partitioning protein ParB / Stage 0 sporulation protein J
n S biosynthesis protein B (SagA)

asX
: Diacylglucosamine hydrolase like
ne 5'-triphosphate nucleotidohydrolase (EC 3.6.1.23)
protein RadA
acyl-ACP acyltransferase PlsX
nhydrase (EC 4.2.1.1)
main-containing protein
RNA synthetase (EC 6.1.1.17) @ Glutamyl-tRNA(Gln) synthetase (EC 6.1.1.24)
nase of the competence regulon ComD
nsduction protein kinase (EC 2.7.3.-)
egulator of the competence regulon ComE
se P protein component (EC 3.1.26.5)
orane protein translocase component YidC, short form OxaI-like
g protein Jag
31: hypothetical protein
erase III beta subunit (EC 2.7.7.7)
protein
mal protein L34p
nnosamine-6-phosphate 2-epimerase (EC 5.1.3.9)
orter, predicted N-acetylneuraminate-binding protein
orter, predicted N-acetylneuraminate transport system permease protein 1
orter, predicted N-acetylneuraminate transport system permease protein 2
61: hypothetical protein
uraminate lyase (EC 4.1.3.3)
nnosamine kinase (EC 2.7.1.60)
utilization regulator, RpiR family

oxyribonuclease YcfH
osylaminoimidazole-succinocarboxamide synthase (EC 6.3.2.6)
se M5 (EC 3.1.26.8)
denine(1518)-N(6)/adenine(1519)-N(6))-dimethyltransferase (EC 2.1.1.182)
mall subunit-stimulated GTPase EngC
osphate 3-epimerase (EC 5.1.3.1)
rophosphokinase (EC 2.7.6.2)
bination protein RmuC
onuclease Bsu YhaM
cription regulator associated with purine metabolism
lusion protein
nal protein S12p (S23e)
osylformylglycinamidine synthase, synthetase subunit (EC 6.3.5.3) / Phosphoribosylformylglycinamidine synthase, glutamine amidotra
nal protein S7p (S5e)
elongation factor G
dent glyceraldehyde-3-phosphate dehydrogenase (EC 1.2.1.12) / Plasmin(ogen) receptor
98: hypothetical protein
transport ATP-binding protein
ABC transporter, glutamine-binding protein/permease protein
29: hypothetical protein
yl-diphosphatase (EC 3.6.1.27)
gulator of genetic competence MecA
nyl-phosphate N-acetylglucosaminyl 1-phosphate transferase (EC 2.7.8.-)
phoribosyltransferase (EC 2.4.2.14)
al cytosolic protein
cluster assembly ATPase protein SufC
cluster assembly protein SufD
sulfurase (EC 2.8.1.7), SufS subfamily
n-sulfur cluster assembly scaffold protein for SUF system, SufE2
cluster assembly protein SufB
alanine carboxypeptidase (EC 3.4.16.4)
alanine carboxypeptidase (EC 3.4.16.4)
le ABC transporter, periplasmic oligopeptide-binding protein OppA (TC 3.A.1.5.1)
le transport system permease protein OppB (TC 3.A.1.5.1)
osylformylglycinamidine cyclo-ligase (EC 6.3.3.1)

de transport system permease protein OppC (TC 3.A.1.5.1)
de transport ATP-binding protein OppD (TC 3.A.1.5.1)
de transport ATP-binding protein OppF (TC 3.A.1.5.1)
e
e-specific sigma factor ComX
nent protein
: Hydrolase, HAD subfamily IIIA
g protein YqeH, required for biogenesis of 30S ribosome subunit
: RNA binding protein
nucleotide adenylyltransferase (EC 2.7.7.18)
osylglycinamide formyltransferase (EC 2.1.2.2)
HAD superfamily), YqeK
in
: Methyltransferase (EC 2.1.1.-)
: UPF0348 protein family
: hypothetical protein YeeN
C transporter, substrate-binding protein
e ABC transporter substrate-binding protein
e ABC transporter ATP-binding protein
e ABC transporter permease protein
hain amino acid transport system carrier protein
ydrolase (EC 3.5.4.10) / Phosphoribosylaminoimidazolecarboxamide formyltransferase (EC 2.1.2.3)
ate/amino acid:cation (Na+ or H+) symporter
potassium uptake protein TrkA
uptake protein TrkH
subunit 7-methylguanosine (m7G) methyltransferase GidB
ein
protein HtpX (EC 3.4.24.-)
rotein in cluster with ribosomal protein L32p, Firmicutes subfamily
egulator CsrR
tide reductase transcriptional regulator NrdR
oline binding protein
ader DnaB
ader DnaI
g protein EngA

mily helicase
42: hypothetical protein
:ylmuramate--alanine ligase (EC 6.3.2.8)
ylalkylamine n-acetyltransferase
: protein YceG like
on elongation factor GreA
orane protein translocase component YidC, OxaA protein
oosylamine--glycine ligase (EC 6.3.4.13)
ucleoside phosphatase
: rRNA methylase
: HD family hydrolase
: membrane protein
: hypothetical protein
racemase (EC 5.1.1.3)
omain / Nucleoside 5-triphosphatase RdgB (dHAPTP, dITP, XTP-specific) (EC 3.6.1.15)
: phosphoesterase
: CBS domain containing protein
combinase XerD
oosylaminoimidazole carboxylase catalytic subunit (EC 4.1.1.21)
ı and condensation protein A
ı and condensation protein B
large subunit pseudouridine synthase B (EC 4.2.1.70)
D
ine(34)-2'-O)-methyltransferase (EC 2.1.1.207)
pecific component RibU of riboflavin ECF transporter
-associated phospholipid phosphatase
Predicted Fe-S oxidoreductase
dent methyltransferase, MraW methylase family (EC 2.1.1.-)
; and related proteins containing CBS domains
17: hypothetical protein
oosylaminoimidazole carboxylase ATPase subunit (EC 4.1.1.21)
rmate-lyase activating enzyme (EC 1.97.1.4)
-dependent inorganic pyrophosphatase (EC 3.6.1.1)
25: hypothetical protein
e3+-siderophore transport system, permease 2 component

e3+-siderophore transport system, permease component
e-binding periplasmic protein precursor (TC 3.A.1.14.3)
e transport ATP-binding protein FhuC (TC 3.A.1.14.3)
ylmuramoylalanyl-D-glutamate--L-lysine ligase (EC 6.3.2.7)
protein involved in the export of O-antigen, teichoic acid lipoteichoic acids
phoribosyltransferase (EC 2.4.2.9)
57: hypothetical protein
al protein
dent Clp protease proteolytic subunit (EC 3.4.21.92)
al protein
e kinase (EC 2.7.4.9)
erase III delta prime subunit (EC 2.7.7.7)
al protein
al
ation intiation control protein YabA
subunit methyltransferase I
28: hypothetical protein
al protein
c copper homeostasis protein CutC
ductase (EC 1.20.4.1)
98: hypothetical protein
bonuclease III (EC 3.1.11.2)
onooxygenase (EC 1.13.12.4)
al protein
peptidase ScpC (EC 3.4.21.-)
51: hypothetical protein
81: hypothetical protein
protein, putative
cinate lyase (EC 4.3.2.2)
tRNA synthetase (EC 6.1.1.10)
al protein
97: hypothetical protein
tide reductase of class Ib (aerobic), beta subunit (EC 1.17.4.1)
tide reduction protein NrdI
tide reductase of class Ib (aerobic), alpha subunit (EC 1.17.4.1)

DP-ribosyltransferase (EC 2.4.2.-)
nent protein
23: hypothetical protein
nent protein
onal regulator
al protein
al protein
29: hypothetical protein
64: hypothetical protein
04: hypothetical protein
acyl-carrier protein] reductase (EC 1.1.1.100)
dent oxidoreductase
phosphate transporter
cosamine-1-phosphate uridyltransferase (EC 2.7.7.23) / Glucosamine-1-phosphate N-acetyltransferase (EC 2.3.1.157)
pyrophosphatase (EC 3.6.1.13)
nction DNA helicase RuvB
35: hypothetical protein
nioadenosine nucleosidase (EC 3.2.2.16) / S-adenosylhomocysteine nucleosidase (EC 3.2.2.9)
68: hypothetical protein
ent transcriptional regulator MntR
ABC transporter, periplasmic-binding protein SitA
ABC transporter, ATP-binding protein SitB
ABC transporter, inner membrane permease protein SitD
olyl cis-trans isomerase (EC 5.2.1.8)
n protein FtsK
mbrane protein
ular weight protein tyrosine phosphatase (EC 3.1.3.48)
mal protein L11p (L12e)
mal protein L1p (L10Ae)
nase (EC 2.7.4.-)
ecycling factor
ding domain
thionine sulfoxide reductase MsrA (EC 1.8.4.11)
18: hypothetical protein
eptococcal surface immunogenic protein

sin-crossreactive streptococcal antigen
starvation-inducible protein PhoH, predicted ATPase
if family protein
al protein VC0266 (sugar utilization related?)
ndent hydrolase YbeY, involved in rRNA and/or ribosome maturation and assembly
rol kinase (EC 2.7.1.107)
g protein Era
‹ family protein
81: hypothetical protein
:e-stimulating peptide ABC transporter ATP-binding protein ComA
: transporter ATP binding/permease protein
: transporter ATP binding/permease protein
BlpU
rase family
al protein
al protein
21: hypothetical protein
le in replication
nent protein
nent protein
nent protein
al protein
ıydrogenase (EC 1.1.1.1); Acetaldehyde dehydrogenase (EC 1.2.1.10)
:gulator of the competence regulon ComE
nase of the competence regulon ComD
20: hypothetical protein
:e-stimulating peptide ABC transporter permease protein ComB
:e-stimulating peptide ABC transporter ATP-binding protein ComA
-like peptide M BlpM
77: hypothetical protein
al protein

al protein
ıydrogenase (EC 1.1.1.1)
77: hypothetical protein

al protein
al protein
al protein
nscriptional regulator, MutR family
pyrimidine-DNA glycosylase (EC 3.2.2.23)
-CoA kinase (EC 2.7.1.24)

esistance efflux pump PmrA
icrobial extrusion (MATE) family transporter
translocase subunit SecG (TC 3.A.5.1.1)
ribonuclease RNase R
ding protein SmpB
-peptide cyclotransferase (EC 2.3.2.5)
e-carboxylate peptidase (EC 3.4.19.3)
: Membrane protein
al protein
l glucosyl transferase (EC 2.4.1.-)
athione lyase (EC 4.4.1.5)
ensitive NAD(P)H nitroreductase (EC 1.-.-.-) / Dihydropteridine reductase (EC 1.5.1.34)
nal protein S10p (S20e)
eptidase (EC 3.4.13.9)
control protein A
nsferase LafB, responsible for the formation of Gal-Glc-DAG
nsferase LafA, responsible for the formation of Glc-DAG
al protein
RNA synthetase (EC 6.1.1.3)
cin resistance ATP-binding protein drrA
cin resistance transmembrane protein
ultidrug transport system, permease component
al protein
nal protein L3p (L3e)
CoA thiolase (EC 2.3.1.16) @ Acetyl-CoA acetyltransferase (EC 2.3.1.9)
-fatty-acid--CoA ligase (EC 6.2.1.3)
64: hypothetical protein
onent response regulator SA14-24

onent sensor kinase SA14-24
ent hydrolase YycJ/WalJ, required for cell wall metabolism and coordination of cell division with DNA replication
se III (EC 3.1.26.3)
ne partition protein smc
nscriptional regulator, MutR family
-dehydrogenase I alpha (EC 1.1.1.25)
mal protein L4p (L1e)
71: hypothetical protein
glutamate ligase
25: hypothetical protein
adenosylmethionine synthetase (EC 2.5.1.6)
63: hypothetical protein
nosphate mannoosyltransferase, involved in cell wall biogenesis
se dehydrogenase (EC 1.1.1.22)
orter membrane-spanning permease - macrolide efflux
29: hypothetical protein
al protein
mal protein L23p (L23Ae)
26: hypothetical protein
00: hypothetical protein
48: hypothetical protein
04: hypothetical protein
bilization system antitoxin protein
bilization system toxin protein
al protein
88: hypothetical protein
nent protein
nent protein
mal protein L2p (L8e)
HAD superfamily)
HAD superfamily)
gnition particle receptor protein FtsY (=alpha subunit) (TC 3.A.5.1.1)
resistance protein B
side bgl operon antiterminator, BglG family
beta-glucoside-specific IIB component (EC 2.7.1.69) / PTS system, beta-glucoside-specific IIC component (EC 2.7.1.69) / PTS system, beta

-beta-glucosidase (EC 3.2.1.86)
orter, substrate-binding protein
rmease
44: hypothetical protein
nal protein S19p (S15e)
on accessory protein (S1 RNA-binding domain)
etallopeptidase (Zinc) SprT family
k protein C, putative; stress-responsive transcriptional regulator
phosphorylase (EC 2.7.1.-) (EC 2.7.4.-)
tein diacylglyceryl transferase (EC 2.4.99.-)
ess protein
01: hypothetical protein
01: hypothetical protein
EC 3.4.-.-)
U32 family large subunit [C1]
nal protein L22p (L17e)
33: hypothetical protein
ryl-tRNA synthetase related
74: hypothetical protein
synthetase (class II) (EC 6.1.1.6)
noic acid dehalogenase (EC 3.8.1.2)
ycerate mutase family 5
onal regulator
pecific component ThiT of thiamin ECF transporter
, glycosyl hydrolase, family 25
pecific component CbrT of predicted cobalamin ECF transporter
nal protein S3p (S3e)
lipoprotein component of predicted cobalamin ECF transporter
e peroxidase family protein
peptidase F
olpyruvate carboxylase (EC 4.1.1.31)
n protein FtsW
57: hypothetical protein
elongation factor Tu
phate isomerase (EC 5.3.1.1)

al protein
ndent lipid II-Ala--L-alanine ligase @ tRNA-dependent lipid II-Ser--L-alanine ligase
g and nucleic acid-binding protein YchF
nal protein L16p (L10e)
ndent lipid II--L-alanine ligase @ tRNA-dependent lipid II--L-serine ligase
us sugar phosphatase YidA, haloacid dehalogenase-like phosphatase family
93: hypothetical protein
osinetriphosphate triphosphohydrolase (EC 3.1.5.1)
78: hypothetical protein
ansporting ATPase (EC 3.6.3.8)
onal regulator RegR, rpressor of hyaluronate and KDG utilization
II/III-like
N-acetylgalactosamine-specific IID component (EC 2.7.1.69)
N-acetylgalactosamine-specific IIC component (EC 2.7.1.69)
nal protein L29p (L35e)
N-acetylgalactosamine-specific IIB component (EC 2.7.1.69)
d glucuronyl hydrolase (EC 3.2.1.-)
N-acetylgalactosamine- and galactosamine-specific IIA component (EC 2.7.1.69)
gluconate 3-dehydrogenase (EC 1.1.1.125)
-deoxy-L-threo-5-hexosulose-uronate ketol-isomerase (EC 5.3.1.17)
3-deoxygluconate kinase (EC 2.7.1.45)
2-oxoglutarate aldolase (EC 4.1.3.16) @ 2-dehydro-3-deoxyphosphogluconate aldolase (EC 4.1.2.14)
hoglucomutase (EC 5.4.2.6) (EC 2.7.1.41)
osine (oQ) reductase QueG
ain release factor 2; programmed frameshift-containing
nal protein S17p (S11e)
n transporter, ATP-binding protein FtsE (TC 3.A.5.1.1)
n protein FtsX
ylglutathione hydrolase (EC 3.1.2.6)
diol dehydrogenase, S-alcohol forming, (R)-acetoin-specific (EC 1.1.1.4) / Acetoin (diacetyl) reductase (EC 1.1.1.5)
nent protein
/ ATP-dependent helicase YoaA
minotransferase (EC 2.6.1.1)
-tRNA synthetase (EC 6.1.1.22)
: Predicted P-loop-containing kinase

: LPPG:FO 2-phospho-L-lactate transferase like, CofD-like
nal protein L14p (L23e)
: Cytoplasmic hypothetical protein
56: hypothetical protein
zinc-binding lipoprotein ZinT
onal regulator, GntR family
nine-6-phosphate isomerase (EC 5.3.1.-)
nal protein L31p @ LSU ribosomal protein L31p, zinc-independent
: 3'-to-5' oligoribonuclease A, Bacillus type

e mutase I (EC 5.4.99.5)
annel protein
nal protein L24p (L26e)
nal protein L19p
haloacid dehalogenase-like family
subunit B (EC 5.99.1.3)
ing formation regulator EzrA
91: hypothetical protein
4.2.1.11)
n S biosynthesis protein B (SagB)
n S biosynthesis protein C (SagC)
n S biosynthesis protein D (SagD)
n S self-immunity protein (SagE)
nal protein L5p (L11e)
n S biosynthesis protein (SagF)
transporter ATP-binding protein; Streptolysin S export protein (SagG)
n S export transmembrane permease (SagH)
n S export transmembrane permease (SagI)
-rich ORFan next to Sag operon
ase/Exonuclease/phosphatase family protein
pecific component QueT (COG4708) of predicted queuosine-regulated ECF transporter
(EC 6.5.1.2)
on regulator [contains diacylglycerol kinase catalytic domain]
se C chain (EC 3.6.3.14)
nal protein S14p (S29e) @ SSU ribosomal protein S14p (S29e), zinc-dependent

se A chain (EC 3.6.3.14)
se B chain (EC 3.6.3.14)
se delta chain (EC 3.6.3.14)
se alpha chain (EC 3.6.3.14)
se gamma chain (EC 3.6.3.14)
se beta chain (EC 3.6.3.14)
se epsilon chain (EC 3.6.3.14)
65: hypothetical protein
ylglucosamine 1-carboxyvinyltransferase (EC 2.5.1.7)
e-associated EpuA protein
mal protein S8p (S15Ae)
nuclease (Competence-specific nuclease) (EC 3.1.30.-)
al protein
nuclease (Competence-specific nuclease) (EC 3.1.30.-)
yl-tRNA synthetase alpha chain (EC 6.1.1.20)
yl-tRNA synthetase beta chain (EC 6.1.1.20)
nduced protein
89: hypothetical protein
orter permease protein
orter ATP-binding protein
n, zinc metalloprotease superfamily
mal protein L6p (L9e)
dent nuclease, subunit B
dent nuclease, subunit A
ABC transporter, amino acid-binding protein
mal protein S21p
uctance mechanosensitive channel
se (EC 2.7.7.-)
erase sigma factor RpoD
omatic ring hydroxylating enzyme, evidenced by COGnitor; PaaD-like protein (DUF59) involved in Fe-S cluster assembly
hydrorhamnose reductase (EC 1.1.1.133)
cNAc alpha-1,2-L-rhamnosyltransferase (EC 2.4.1.-)
mal protein L18p (L5e)
a alpha-1,3-L-rhamnosyltransferase (EC 2.4.1.-)
containing polysacharide translocation permease

orter (ATP-binding protein) -possibly involved in cell wall localization and side chain formation of rhamnose-glucose polysaccharide
ycosyltransferase - possibly involved in cell wall localization and side chain formation of rhamnose-glucose polysaccharide
a alpha-1,2-L-rhamnosyltransferase/alpha-L-Rha alpha-1,3-L- rhamnosyltransferase (EC 2.4.1.-)
pressor C1
al protein - phage associated
hypothetical protein
ein
NA hydrolase (EC 3.1.1.29)
mal protein S5p (S2e)
ein

ein
22: hypothetical protein
uronidase
e lyase (phage associated)
peptidase
bers
mal protein L30p (L7e)
ength tape-measure protein
al protein
plate
r tail protein
al phage protein
al phage protein
al phage protein
al phage protein
al phage protein
r capsid protein
mal protein L15p (L27Ae)
lp protease-like protein
ein, phage associated
inase, large subunit
ein
ciated homing endonuclease
al phage protein

al phage protein
al phage protein
;rase: site-specific recombinase
anscriptional activator - phage associated
translocase secY subunit (TC 3.A.5.1.1)
tor associated protein-related protein
nunity repressor protein - phage associated
al protein
ıded DNA-binding protein
al phage protein
ıypothetical protein - phage associated
ein
al phage protein
46: hypothetical protein
ıypothetical protein - phage associated
tinase (EC 2.7.4.3)
al phage protein
al phage protein
ıder DnaI
initiator protein A
DNA helicase (DnaB) (EC 3.6.4.12)
al protein
al phage protein
al phage protein
epressor protein
al phage protein
initiation factor 1
domain protein
al phage protein
criptional regulator, Cro/CI family
ein
ity island SaPIn2
;rase: site-specific recombinase
ansferase, group 2 family protein
protein

saccharide repeat unit export protein
nal protein L36p
protein, related to Actinobacillus protein (1944168)
aminopeptidase (EC 3.4.11.4)
ng protein ebsA

55: hypothetical protein
kinase (EC 2.7.4.14)
initiation factor 3
nal protein L35p
nal protein L20p
c acid synthase LtaS Type IIc
nal protein S13p (S18e)
62 methyltransferase RlmI
quinate dehydratase I (EC 4.2.1.10)
e synthase (EC 4.2.3.5)
36: hypothetical protein
e reductase (EC 1.8.1.7)
ate synthase (EC 6.3.2.12) / Folylpolyglutamate synthase (EC 6.3.2.17)
etyltransferase Gcn5
sulfurase (EC 2.8.1.7)
4-thiouridine synthase (former ThiI)
synthesis protein capA
nal protein S11p (S14e)
nal protein L21p
: Potential ribosomal protein
nal protein L27p
d methionine metabolism regulator CmbR, LysR family
n signal peptidase (EC 3.4.23.36)
large subunit pseudouridine synthase D (EC 4.2.1.70)
phoribosyltransferase (EC 2.4.2.9) / Pyrimidine operon regulatory protein PyrR
nease
arbamoyltransferase (EC 2.1.3.2)
-phosphate synthase small chain (EC 6.3.5.5)
ed RNA polymerase alpha subunit (EC 2.7.7.6)

-phosphate synthase large chain (EC 6.3.5.5)
: component of efflux system
orter ATP-binding protein
orter permease protein
osphoryl diester phosphodiesterase (EC 3.1.4.46)
mal protein S16p
RNA binding protein YlqC
protein
36: hypothetical protein
-cadmium resistance protein CzcD
on-repair coupling factor
mal protein L17p
onal regulator, TetR family
rocessing protein RimM
ine37-N1) -methyltransferase (EC 2.1.1.31)
n reductase (EC 1.8.1.9)
protein
pantoate 2-reductase (EC 1.1.1.169)
onal repressor of the fructose operon, DeoR family
fructokinase (EC 2.7.1.56)
fructose-specific IIA component (EC 2.7.1.69) / PTS system, fructose-specific IIB component (EC 2.7.1.69) / PTS system, fructose-specific
can hydrolase, Autolysin1 (EC 3.5.1.28)
al protein
can hydrolase, Autolysin2 (EC 3.5.1.28)
nent protein
al protein
obulin G-endopeptidase (IdeS) / Mac/ Secreted immunoglobulin binding protein (Sib38)
ein
diphosphate kinase (EC 2.7.4.6)
al protein SP1558
/ protein
otidyltransferase (EC 2.7.7.21) (EC 2.7.7.25)
ATPase components of ABC transporters with duplicated ATPase domains
10: hypothetical protein
formylase (EC 3.5.1.88)

dase (EC 3.1.3.5)
nosphokinase (EC 2.7.6.5)
of D-alanyl-lipoteichoic acid biosynthesis, DltR
of D-alanyl-lipoteichoic acid biosynthesis, sensor histidine kinase
kinase (EC 2.7.1.36)
mevalonate decarboxylase (EC 4.1.1.33)
evalonate kinase (EC 2.7.4.2)
l-diphosphate delta-isomerase, FMN-dependent (EC 5.3.3.2)
ethylglutaryl-CoA reductase (EC 1.1.1.34)
ethylglutaryl-CoA synthase (EC 2.3.3.10)
e synthase (EC 2.1.1.45)
ate reductase (EC 1.5.1.3)
89: hypothetical protein
dent Clp protease ATP-binding subunit ClpX
g protein EngB
al protein SPy0887
P-dependent Clp proteinase (ATP-binding subunit)
nosphate isomerase A (EC 5.3.1.6)
onal repressor AdcR for Zn(2+)-responsive expression
ntomutase (EC 5.4.2.7)
ductase (EC 1.20.4.1)
eoside phosphorylase (EC 2.4.2.1); N-Ribosylnicotinamide phosphorylase (EC 2.4.2.1) ## possible
eoside phosphorylase (EC 2.4.2.1)
rotein kinase
al protein
biosynthesis and transport regulator MtaR, LysR family
62: hypothetical protein
'-phosphate decarboxylase (EC 4.1.1.23)
osphoribosyltransferase (EC 2.4.2.10)
ansporter, ATP-binding protein ZnuC
kanoate-cyclic-dimer hydrolase (EC 3.5.2.12)
C transporter, periplasmic cystine-binding protein FliY
nsport system permease protein
glycosylase, family 1
otase (EC 3.5.2.3)

hate:glycerol-3-phosphate O-acyltransferase PlsY
rase IV subunit B (EC 5.99.1.-)
rase IV subunit A (EC 5.99.1.-)
hain amino acid aminotransferase (EC 2.6.1.42)
90: hypothetical protein
ansporter, inner membrane permease protein ZnuB
nal protein S1p
: hypothetical protein
domain protein UPF0176, Firmicutes subgroup
al protein
e S-transferase domain protein (class beta)
toxin A
06: hypothetical protein
hylallyltransferase (EC 2.5.1.75)
g protein HflX
ne beta-lyase (EC 4.4.1.8)
rotein
se Z (EC 3.1.26.11)
dehydrogenase
nded-DNA-specific exonuclease RecJ (EC 3.1.-.-)
osphoribosyltransferase (EC 2.4.2.7)
ne replication initiation protein dnaD
ase III (EC 4.2.99.18)
NA-m1A22 methylase
: Hypothetical protein
amino acid oxidases family
phosphate thymidylyltransferase (EC 2.7.7.24)
IA synthetase (EC 6.1.1.1)
hydrorhamnose 3,5-epimerase (EC 5.1.3.13)
se 4,6-dehydratase (EC 4.2.1.46)
utT protein (7,8-dihydro-8-oxoguanine-triphosphatase) (EC 3.6.1.-)
al protein SPy1011
-containing protein
n/fibrinogen-binding protein
al protein

ɔrter substrate-binding protein
ɔrter permease protein
ɔrter ATP-binding protein
lar transpeptidase-transglycosylase (EC 2.4.1.129) (EC 3.4.-.-)
se J2 (endoribonuclease in RNA processing)
ɛsterase
26: hypothetical protein
ɔrter ATP-binding protein
ıydrogenase E1 component alpha-subunit (EC 1.2.4.-)
ıydrogenase E1 component beta-subunit (EC 1.2.4.-)
ɔamide acetyltransferase component (E2) of acetoin dehydrogenase complex (EC 2.3.1.-)
ɔamide dehydrogenase of acetoin dehydrogenase (EC 1.8.1.4)
e lyase precursor (EC 4.2.2.1)
ɔtein ligase A
ıssociated heat shock protein implicated in the recycling of the 50S subunit (S4 paralog)
ed RNA polymerase beta subunit (EC 2.7.7.6)
nidotransferase similar to cobyric acid synthase
mino acid ligase found clustered with an amidotransferase
al protein YbbP, contains nucleotide-binding domain of DisA bacterial checkpoint controller
rized secreted protein, YBBR Bacillus subtilis homolog
ıcosamine mutase (EC 5.4.2.10)
18: hypothetical protein
al radical SAM family enzyme in heat shock gene cluster, similarity with CPO of BS HemN-type
ioesterase
al NagD-like phosphatase
ntegral membrane protein
ed RNA polymerase beta' subunit (EC 2.7.7.6)
elongation factor LepA
ĸe surface protein
nent protein
elongation factor LepA
ĸe surface protein
thionine sulfoxide reductase MsrB (EC 1.8.4.12)
ɛmbrane protein YeiH
IIA component, putative

g protein, FIG046916
mannose/fructose family IIB component
mannose-specific IIC component (EC 2.7.1.69)
mannose-specific IID component (EC 2.7.1.69)
onent sensor kinase, associated with ferric iron transporter, SPy1061 homolog
onent response regulator, associated with ferric iron transporter, SPy1062 homolog
ABC transporter, iron-binding protein
emialdehyde dehydrogenase [NAD] (EC 1.2.1.24); Succinate-semialdehyde dehydrogenase [NADP+] (EC 1.2.1.16)
se ABC subunit C
ensitive NAD(P)H nitroreductase (EC 1.-.-.-) / Dihydropteridine reductase (EC 1.5.1.34)
hine deacetylase/Succinyl-diaminopimelate desuccinylase and related deacylases
etence protein ComGA, access of DNA to ComEA
tRNA-U34 5-formylation enzyme TrmE
mal protein L10p (P0)
mal protein L7/L12 (P1/P2)
alanine carboxypeptidase (EC 3.4.16.4)
ride deacetylase
ate synthase (EC 6.3.2.12) / Folylpolyglutamate synthase (EC 6.3.2.17)
ydrolase I (EC 3.5.4.16) type 1
eroate synthase (EC 2.5.1.15)
opterin aldolase (EC 4.1.2.25)
hydroxy-6-hydroxymethyldihydropteridine pyrophosphokinase (EC 2.7.6.3)
etence protein ComGB, access of DNA to ComEA
ylenolpyruvoylglucosamine reductase (EC 1.1.1.158)
transport ATP-binding protein PotA (TC 3.A.1.11.1)
e Putrescine ABC transporter permease component PotB (TC 3.A.1.11.1)
e Putrescine ABC transporter permease component potC (TC_3.A.1.11.1)
orter, periplasmic spermidine putrescine-binding protein PotD (TC 3.A.1.11.1)
onent response regulator, malate (EC 2.7.3.-)
onent sensor histidine kinase, malate (EC 2.7.3.-)
-) symporter
dent malic enzyme (EC 1.1.1.38)
hydrogenase, zinc-containing
etence protein ComGC, access of DNA to ComEA, FIG007487
nent protein

phosphatase precursor (EC 3.1.3.2)
ed chloride channel family protein
hydrolase family protein
al protein

protein RadC
amidotransferase, class I
itive transcriptional regulator (AT-rich DNA-binding protein)
33: hypothetical protein
etence protein ComGD, access of DNA to ComEA, FIG038316
sulfurase (EC 2.8.1.7)
sphate pyrophosphokinase (EC 2.7.6.1)
yclase (EC 4.6.1.1)
iosphokinase (EC 2.7.6.5)
(EC 2.7.1.23)
ibosomal large subunit pseudouridine synthase D, Bacillus subtilis YjbO type
acetyltransferase (EC 2.3.1.8)
dehydrogenase
multidrug efflux pump
ycine betaine ABC transport system permease protein ProV (TC 3.A.1.12.1)
etence protein ComGE, FIG075573
ycine betaine binding ABC transporter protein ProX (TC 3.A.1.12.1) / Osmotic adaptation
tase (EC 1.7.1.7)
iosphoribosyltransferase (EC 2.4.2.22)
ermease
al similar to thiamin biosynthesis lipoprotein ApbE
conate tautomerase (EC 5.3.2.-); Xylose transport system permease protein xylH
kinase (EC 2.7.1.21)
ain release factor 1
)-glutamine methyltransferase PrmC, methylates polypeptide chain release factors RF1 and RF2
n (YrdC domain) required for threonylcarbamoyladenosine t(6)A37 modification in tRNA
etence protein ComGF, access of DNA to ComEA, FIG012620
chricin N-acetyltransferase (EC 2.3.1.-)
oxymethyltransferase (EC 2.1.2.1)
ylate synthases, 23S RNA-specific

ccal vaccine antigen A homolog
orter, ATP-binding/permease protein
orter, ATP-binding/permease protein
xidase (EC 1.11.1.1)
ehydrogenase (EC 1.1.1.27)
subunit A (EC 5.99.1.3)
PXTG specific
etence protein ComGG, FIG068335
59: hypothetical protein
48: hypothetical protein
12: hypothetical protein
on regulator [contains diacylglycerol kinase catalytic domain]
III
71: hypothetical protein
mal subunit maturation GTPase RbgA (B. subtilis YlqF)
se HII (EC 3.1.26.4)
fold nucleotide-binding protein Smf possibly involved in DNA uptake
omerase I (EC 5.99.1.2)
n protein DivIC (FtsB), stabilizes FtsL against RasP cleavage
ecific methyltransferase (EC 2.1.1.72)
ment protein
J-54 MTase gid
te decarboxylase alpha chain (EC 4.1.1.3)
41: hypothetical protein
oxyl carrier protein of oxaloacetate decarboxylase; Biotin carboxyl carrier protein
onyl-CoA decarboxylase, beta chain (EC 4.1.1.41); Oxaloacetate decarboxylase beta chain (EC 4.1.1.3)
osphoribosyl)-3'-dephosphocoenzyme-A synthase (EC 2.7.8.25)
onal regulator, GntR family
te complex transporter
46: hypothetical protein
ase (EC 2.7.2.1)
oxyl carrier protein of oxaloacetate decarboxylase; Biotin carboxyl carrier protein
te decarboxylase beta chain (EC 4.1.1.3)
13: hypothetical protein
e gamma chain, acyl carrier protein (EC 4.1.3.6)

e beta chain (EC 4.1.3.6)
e alpha chain (EC 4.1.3.6)
lyase phosphoribosyl-dephospho-CoA transferase (EC 2.7.7.61)
te decarboxylase alpha chain (EC 4.1.1.3)
o-3S]-lyase] ligase (EC 6.2.1.22)
60: hypothetical protein
al protein SPy0110
combinase XerC
regulator of exopolysaccharide synthesis, competence and biofilm formation Ftr, XRE family
gnition particle, subunit Ffh SRP54 (TC 3.A.5.1.1)
gnition particle associated protein
onal regulator in cluster with beta-lactamase, GntR family
68: hypothetical protein
ase [glutamine-hydrolyzing], amidotransferase subunit (EC 6.3.5.2) / GMP synthase [glutamine-hydrolyzing], ATP pyrophosphatase subun
orter ATP-binding protein
-carboxylate reductase (EC 1.5.1.2)
pecific component PdxU of predicted pyridoxine ECF transporter
inase (EC 2.7.1.35)
ranscriptional regulator of pyridoxine metabolism
ibonucleoside-triphosphate reductase (EC 1.17.4.2)
synthetase (EC 2.7.8.-)
etrahydrofolate ligase (EC 6.3.4.3)
otein ligase A
dent protein deacetylase of SIR2 family
Macro domain, possibly ADP-ribose binding module
oyl attachment domain protein, GcvH-like
minopeptidase (EC 3.4.11.7)
ike monooxygenase (EC 1.14.-.-)
ADH-dependent flavin oxidoreductase in cluster with COG2110
otein lipoyl transferase
ntothenoylcysteine synthetase (EC 6.3.2.5)
ntothenoylcysteine decarboxylase (EC 4.1.1.36)
pecific component PanT of predicted pantothenate ECF transporter
ucomutase (EC 5.4.2.2)
ABC transporter, permease protein 2

oxyribose-specific ABC transporter, permease protein
oxyribose-specific ABC transporter, ATP-binding protein
: hypothetical protein
-binding protein
aminase (EC 3.5.4.5)
RNA small subunit methyltransferase C (EC 2.1.1.52) ## SSU rRNA m2G1207
ate kinase (EC 2.7.1.33)
nal protein S20p
onent system sensor histidine kinase CiaH (EC 2.7.3.-)
onent system response regulator CiaR
peptidase (EC 3.4.11.15)
transport system regulatory protein PhoU
transport ATP-binding protein PstB (TC 3.A.1.7.1)
: Thioredoxin
transport ATP-binding protein PstB (TC 3.A.1.7.1)
transport system permease protein PstA (TC 3.A.1.7.1)
transport system permease protein PstC (TC 3.A.1.7.1)
ABC transporter, periplasmic phosphate-binding protein PstS (TC 3.A.1.7.1)
RNA cytosine-C5-methylases
nonophosphatase (EC 3.1.3.25)
: hypothetical protein
onal regulator SpxA1
kinase (EC 2.7.1.26) / FMN adenylyltransferase (EC 2.7.7.2)
ouridine synthase B (EC 4.2.1.70)
yl-tRNA synthetase domain protein (Bsu YtpR)
60: hypothetical protein
protein
ction-modification system specificity subunit
orter permease protein
orter ATP-binding protein YvcR
onal regulator, TetR family
onal regulator, GntR family
ess protein, Gls24 family
97: hypothetical protein
ess protein, Gls24 family

: tRNA-binding protein
32: hypothetical protein
74: hypothetical protein
77: hypothetical protein
dent DNA helicase UvrD/PcrA
'cine symporter GlyP
-cadmium resistance protein
r
ABC transporter, amino acid-binding protein
olar amino acid transport system, ATPase component
orter membrane-spanning permease - glutamine transport
osine kinase (EC 2.7.1.76) / Deoxyguanosine kinase (EC 2.7.1.113)
honate utilization operon protein PhnA
ne--fructose-6-phosphate aminotransferase [isomerizing] (EC 2.6.1.16)
idase I (EC 3.4.21.89)
nase (EC 2.7.1.40)
fructokinase (EC 2.7.1.11)
erase III alpha subunit (EC 2.7.7.7)
onal regulator, GntR family
ultidrug transport system, ATPase component
orter, ATP-binding protein
active protein Se23.5 (Fragment)
n
cation methylase
roxybutyrate dehydrogenase (EC 1.1.1.30)
e biosynthesis protein LanM/pseudogene causing stop codon at 177th aa
cal mitogenic exotoxin Z (SmeZ)
ytic M protein/pseudogene causing stop codon at 85th aa
brane histidine kinase CsrS / deletion of T at 1407 causing a early stop codon at 469th aa)
nent protein
99: hypothetical protein
c acid synthase LtaS Type Iic/interrupted by large phage
n
c acid synthase LtaS Type Iic (The first GCT-Ala is overlapped with the last codon of the former half interrupted protein)
ocytidyl-2-C-methyl-D-erythritol kinase (EC 2.7.1.148)

diphosphate kinase (EC 2.7.4.6)
ylmuramoylpentapeptide-lysine N(6)-alanyltransferase (EC 2.3.2.10) (EC 2.3.2.-)
nit Ribosomal RNA; ssuRNA; SSU rRNA
CC
AA
CG
GG
nit Ribosomal RNA; ssuRNA; SSU rRNA
GC
nit Ribosomal RNA; lsuRNA; LSU rRNA

AC
TC
GC
T
AG
GT
CC
AA
CG
GG
CAT
CAT
lo-TGA
nit Ribosomal RNA; lsuRNA; LSU rRNA
CAT
AA
CC
AT
lo-GCT
nit Ribosomal RNA; ssuRNA; SSU rRNA
GC
nit Ribosomal RNA; lsuRNA; LSU rRNA

AC

CC
AT
TC
lo-TGA
CAT
GAA
TA
CA
TG
TG
AC
AA
nit Ribosomal RNA; ssuRNA; SSU rRNA
GC
nit Ribosomal RNA; lsuRNA; LSU rRNA

TT
CG
CT
TA
TG
TC
CG
GT
TC
AT
GT
AG
TT
TC
AC

TT
nit Ribosomal RNA; lsuRNA; LSU rRNA

GC
nit Ribosomal RNA; ssuRNA; SSU rRNA
GA
AG
CG
TT

nit Ribosomal RNA; lsuRNA; LSU rRNA
GC
AG
nit Ribosomal RNA; ssuRNA; SSU rRNA
TT
CA
TT
TC
CT
GT

lgeranyl diphosphate synthase (EC 2.5.1.29)

[illegible]y

\ component (EC 2.7.1.69)

ystem, maltose and glucose-specific IIA component (EC 2.7.1.69)

pAb@PabAb

ific IIA component (EC 2.7.1.69)

nsferase subunit (EC 6.3.5.3)

a-glucoside-specific IIA component (EC 2.7.1.69)

IIC component (EC 2.7.1.69)

it (EC 6.3.5.2)

| genes | logFC.4hr | logCPM.4hr | statistic.4hr | Pvalue.4hr | FDR.4hr | function |
|---|---|---|---|---|---|---|
| AP53_1665 | -9.308305994 | 6.355545902 | 185.2022996 | 2.09E-08 | 4.05E-05 | Streptokinase |
| AP53_148 | 0.897575343 | 6.799100627 | 9.975776883 | 0.008724724 | 0.9997444 | Probable L-ascorbate-6-phosphate |
| AP53_1066 | -0.458726182 | 7.930079305 | 9.855853384 | 0.009031388 | 0.9997444 | Maltose/maltodextrin ABC transpo |
| AP53_429 | -1.006125157 | 2.085177102 | 9.119989882 | 0.011219752 | 0.9997444 | Hypothetical protein |
| AP53_1562 | -0.314499411 | 7.62799416 | 8.924747725 | 0.01190249 | 0.9997444 | C3-degrading proteinase |
| AP53_1583 | -0.341436697 | 8.319343603 | 8.47190186 | 0.013685424 | 0.9997444 | Protein of unknown function DUF1 |
| AP53_498 | 5.860543732 | -1.006521509 | 8.428437154 | 0.016722277 | 0.9997444 | hypothetical protein |
| AP53_1617 | -0.323455838 | 6.251830487 | 6.137210949 | 0.030009941 | 0.9997444 | 6-phospho-beta-galactosidase (EC 3 |
| AP53_597 | 0.758241098 | 8.519968898 | 5.814223733 | 0.033793034 | 0.9997444 | hypothetical protein - phage associ |
| AP53_1040 | -0.595107464 | 5.15098903 | 5.710767536 | 0.035124038 | 0.9997444 | Phage protein |
| AP53_811 | 0.767921183 | 11.91569231 | 5.707083835 | 0.035172579 | 0.9997444 | Collagen-like surface protein |
| AP53_1267 | -0.655708977 | 11.23135134 | 5.653121328 | 0.035892975 | 0.9997444 | Arginine deiminase (EC 3.5.3.6) |
| AP53_125 | -0.428106995 | 5.993230058 | 5.651393193 | 0.035916337 | 0.9997444 | hypothetical protein |
| AP53_119 | -0.349861549 | 5.4753407 | 5.533152734 | 0.037559059 | 0.9997444 | V-type ATP synthase subunit G (EC 3 |
| AP53_rna55 | -3.218626534 | 0.474471177 | 5.492788488 | 0.038140415 | 0.9997444 | tRNA-Asn-GTT |
| AP53_1646 | -0.859975446 | 4.87587575 | 5.447813874 | 0.038800964 | 0.9997444 | PTS system IIB component |
| AP53_1662 | -0.387829401 | 4.998466761 | 5.232177271 | 0.04216597 | 0.9997444 | Glucan 1,6-alpha-glucosidase (EC 3. |
| AP53_1661 | -0.418505499 | 5.406063771 | 5.077726736 | 0.044793509 | 0.9997444 | Pullulanase (EC 3.2.1.41) |
| AP53_347 | -0.255204054 | 7.310843593 | 4.893495713 | 0.048190715 | 0.9997444 | Glycerol-3-phosphate transporter |
| AP53_rna75 | -2.498416606 | -0.117416991 | 5.129465378 | 0.04854508 | 0.9997444 | tRNA-Arg-ACG |
| AP53_1054 | -0.707638585 | 4.635450138 | 4.826038461 | 0.04951189 | 0.9997444 | Phage transcriptional regulator, Cr |
| AP53_1632 | 0.279386428 | 7.214749221 | 4.624728339 | 0.05372329 | 0.9997444 | Transcriptional regulator, Cro/CI fa |
| AP53_rna38 | 5.51578949 | -0.468952949 | 4.788073631 | 0.055131311 | 0.9997444 | 5S RNA |
| AP53_1212 | 0.578612292 | 4.201745088 | 4.548751531 | 0.05542461 | 0.9997444 | Cro-like phage transcriptional repre |
| AP53_352 | -0.309110643 | 8.762418006 | 4.454854664 | 0.057618322 | 0.9997444 | FIG01116268: hypothetical protei |
| AP53_1383 | 0.23863172 | 9.539422849 | 4.339367661 | 0.060462704 | 0.9997444 | Guanylate kinase (EC 2.7.4.8) |
| AP53_1647 | -0.550148534 | 5.062154392 | 4.302207852 | 0.061413984 | 0.9997444 | Transcription antiterminator, BglG |
| AP53_1445 | -0.239481271 | 8.853964934 | 4.25605326 | 0.062620975 | 0.9997444 | PTS system, galactose-specific IIC co |
| AP53_598 | 0.59855622 | 9.372826546 | 4.199476519 | 0.064140101 | 0.9997444 | conserved hypothetical protein |
| AP53_922 | 0.224300675 | 9.687420644 | 4.160782727 | 0.065204905 | 0.9997444 | Signal recognition particle, subunit |
| AP53_rna56 | 2.555685965 | 0.251749348 | 4.018131712 | 0.069321116 | 0.9997444 | tRNA-Arg-ACG |
| AP53_1618 | -0.2871535 | 7.218910504 | 3.89068556 | 0.073269368 | 0.9997444 | PTS system, lactose-specific IIB com |
| AP53_1133 | -0.351413113 | 9.591269898 | 3.874455636 | 0.073791654 | 0.9997444 | FIG01116145: hypothetical protei |
| AP53_1581 | -0.483409794 | 6.347730772 | 3.856425991 | 0.074377198 | 0.9997444 | Ribosomal-protein-S18p-alanine ac |

| | | | | | | |
|---|---|---|---|---|---|---|
| AP53_1234 | 0.296545779 | 9.054293806 | 3.856235586 | 0.074383412 | 0.9997444 | Mutator mutT protein (7,8-dihydro |
| AP53_1067 | -0.303588619 | 5.709482512 | 3.817147787 | 0.075672609 | 0.9997444 | Maltose/maltodextrin ABC transpo |
| AP53_1627 | 0.743573287 | 2.557416665 | 3.737860083 | 0.078372665 | 0.9997444 | hypothetical protein |
| AP53_147 | -0.231396524 | 5.637764626 | 3.730087952 | 0.078643634 | 0.9997444 | Transcriptional antiterminator with |
| AP53_972 | 0.218423837 | 8.01282363 | 3.686699451 | 0.080177595 | 0.9997444 | Type I restriction-modification syst |
| AP53_123 | -0.273679825 | 5.959595513 | 3.585880017 | 0.083885999 | 0.9997444 | V-type ATP synthase subunit C (EC 3 |
| AP53_522 | -0.230034274 | 7.975341605 | 3.540911588 | 0.085607795 | 0.9997444 | Hydroxyacylglutathione hydrolase |
| AP53_1644 | -0.566147552 | 6.060227028 | 3.497453375 | 0.087313192 | 0.9997444 | Transaldolase (EC 2.2.1.2) |
| AP53_1116 | -0.245813252 | 7.551347705 | 3.459294396 | 0.088845186 | 0.9997444 | UDP-N-acetylglucosamine 1-carbox |
| AP53_193 | -0.249038723 | 6.180584908 | 3.447404062 | 0.089329299 | 0.9997444 | Histidine kinase of the competence |
| AP53_890 | 0.264641121 | 6.723217123 | 3.434000454 | 0.089878923 | 0.9997444 | FIG01280259: hypothetical protein |
| AP53_1751 | 0.671883088 | 6.490343622 | 3.422474465 | 0.090354886 | 0.9997444 | Histidine ammonia-lyase (EC 4.3.1.[illegible] |
| AP53_1299 | -0.348696209 | 6.93179985 | 3.29711514 | 0.095737602 | 0.9997444 | FIG01114249: hypothetical protein |
| AP53_1867 | 0.349734488 | 7.390902448 | 3.239550616 | 0.098341627 | 0.9997444 | small RNA fasX |
| AP53_673 | 0.222722845 | 9.971696134 | 3.217747348 | 0.099350739 | 0.9997444 | FIG01116936: hypothetical protein |
| AP53_1663 | -0.381624521 | 6.881238382 | 3.200543895 | 0.100156007 | 0.9997444 | Multiple sugar ABC transporter, ATP |
| AP53_1265 | -0.490142327 | 12.06436896 | 3.20046546 | 0.100159697 | 0.9997444 | Ornithine carbamoyltransferase (EC |
| AP53_1222 | -0.274822473 | 7.686486073 | 3.166485781 | 0.101774172 | 0.9997444 | Arginine pathway regulatory protei |
| AP53_158 | 0.447755825 | 4.055778573 | 3.129779617 | 0.103554546 | 0.9997444 | Mobile element protein |
| AP53_1749 | 0.612824768 | 6.229693034 | 3.120718701 | 0.103999954 | 0.9997444 | Conserved protein |
| AP53_1395 | -0.626080954 | 7.290890495 | 3.057873268 | 0.107155518 | 0.9997444 | Cell division protein GpsB, coordin |
| AP53_1750 | 0.587885159 | 6.210996007 | 3.023097584 | 0.108952747 | 0.9997444 | putative cationic amino acid transp |
| AP53_1746 | 0.67189003 | 6.287957149 | 2.992267617 | 0.110577469 | 0.9997444 | Glutamate formiminotransferase (E |
| AP53_1153 | 0.209903218 | 7.176369711 | 2.990808004 | 0.110655134 | 0.9997444 | FIG01114764: hypothetical protein |
| AP53_1449 | -0.609093016 | 5.031016365 | 2.977665707 | 0.111357488 | 0.9997444 | Copper chaperone |
| AP53_1266 | -0.507985516 | 10.8204009 | 2.9233742 | 0.11431841 | 0.9997444 | Acetyltransferase (EC 2.3.1.-) |
| AP53_141 | 0.256339575 | 5.279381164 | 2.89840294 | 0.115713174 | 0.9997444 | Ascorbate-specific PTS system, EIIC |
| AP53_1177 | 0.326113744 | 10.37121398 | 2.896856782 | 0.115800229 | 0.9997444 | Lead, cadmium, zinc and mercury t |
| AP53_1444 | -0.16814049 | 9.09676772 | 2.894958245 | 0.115907237 | 0.9997444 | Galactose-6-phosphate isomerase, [illegible] |
| AP53_1748 | 0.679485888 | 6.709134603 | 2.87425231 | 0.117082334 | 0.9997444 | Formate--tetrahydrofolate ligase (E |
| AP53_450 | 0.53886785 | 1.699702403 | 2.847944746 | 0.11859685 | 0.9997444 | FIG01116026: hypothetical protein |
| AP53_939 | -0.410022483 | 9.212780987 | 2.841736415 | 0.118957821 | 0.9997444 | Biotinyl-lipoyl attachment domain |
| AP53_819 | -0.717376299 | 8.051631786 | 2.832065753 | 0.119522839 | 0.9997444 | PTS system, IIA component, putativ |
| AP53_1364 | 0.319865013 | 8.777863786 | 2.82025526 | 0.120217432 | 0.9997444 | Phage integrase: site-specific recom |
| AP53_1027 | -0.602734888 | 4.361305901 | 2.815230277 | 0.120514487 | 0.9997444 | hypothetical phage protein |

| | | | | | | |
|---|---|---|---|---|---|---|
| AP53_rna34 | -0.891542308 | 3.37489484 | 2.770798505 | 0.123181441 | 0.9997444 | tRNA-Pseudo-GCT |
| AP53_380 | 0.407445066 | 5.255508454 | 2.716513705 | 0.126540845 | 0.9997444 | hypothetical protein |
| AP53_223 | 0.4475148 | 4.257604621 | 2.707845514 | 0.127087852 | 0.9997444 | FIG01114598: hypothetical protei |
| AP53_1747 | 0.688212253 | 6.135558656 | 2.677875757 | 0.129002093 | 0.9997444 | Formiminotetrahydrofolate cyclod |
| AP53_1553 | -0.280484674 | 7.983589658 | 2.676578821 | 0.129085745 | 0.9997444 | FIG01114225: hypothetical protei |
| AP53_1559 | -0.476074668 | 11.45855556 | 2.67393246 | 0.129256646 | 0.9997444 | Pyruvate formate-lyase (EC 2.3.1.54 |
| AP53_940 | -0.372696597 | 8.107856263 | 2.663858512 | 0.129909823 | 0.9997444 | Luciferase-like monooxygenase (EC |
| AP53_235 | -0.220624956 | 8.612244144 | 2.658566132 | 0.130254632 | 0.9997444 | Iron-sulfur cluster assembly protei |
| AP53_1379 | 0.178922824 | 9.287515947 | 2.617384733 | 0.132977345 | 0.9997444 | 16S rRNA (cytosine(967)-C(5))-meth |
| AP53_127 | -0.215954122 | 6.639887807 | 2.616777949 | 0.133017994 | 0.9997444 | V-type ATP synthase subunit B (EC 3 |
| AP53_700 | 0.318050597 | 4.841515719 | 2.595911505 | 0.134425441 | 0.9997444 | Transcriptional regulator, TetR fam |
| AP53_145 | 0.281086893 | 5.127435458 | 2.580089501 | 0.13550515 | 0.9997444 | L-xylulose 5-phosphate 3-epimeras |
| AP53_477 | -0.193620089 | 8.971203091 | 2.579218809 | 0.135564883 | 0.9997444 | FIG01114801: hypothetical protei |
| AP53_1658 | -0.170011117 | 8.238727787 | 2.566268037 | 0.136457289 | 0.9997444 | Undecaprenyl diphosphate synthas |
| AP53_1254 | -0.129443675 | 8.539635484 | 2.551949516 | 0.137452577 | 0.9997444 | Ribosomal RNA large subunit methy |
| AP53_405 | -0.200938392 | 5.795434844 | 2.544565716 | 0.137969405 | 0.9997444 | Positive transcriptional regulator, I |
| AP53_1560 | -0.275625495 | 7.297212437 | 2.532737808 | 0.138802414 | 0.9997444 | hypothetical protein |
| AP53_801 | 0.138540886 | 7.615810176 | 2.508820399 | 0.140506336 | 0.9997444 | proposed amino acid ligase found c |
| AP53_1173 | 0.209917126 | 9.926256259 | 2.489554544 | 0.141898115 | 0.9997444 | Phosphoglycerate mutase (EC 5.4.2 |
| AP53_1206 | -0.604654522 | 3.161078979 | 2.422547844 | 0.146876536 | 0.9997444 | Phage protein |
| AP53_4 | -0.320048815 | 4.818853348 | 2.414547555 | 0.147485618 | 0.9997444 | hypothetical protein |
| AP53_763 | 0.189941159 | 7.618129502 | 2.4004462 | 0.148567006 | 0.9997444 | hypothetical protein |
| AP53_860 | -0.223779684 | 8.307023261 | 2.392581215 | 0.149174516 | 0.9997444 | Cysteine desulfurase (EC 2.8.1.7) |
| AP53_1773 | 0.274336836 | 11.49108682 | 2.388138685 | 0.14951906 | 0.9997444 | Putative Holliday junction resolvas |
| AP53_451 | 1.155022649 | 0.454461434 | 2.373279227 | 0.15067886 | 0.9997444 | FIG01114600: hypothetical protei |
| AP53_776 | 0.184690611 | 7.944744549 | 2.3727934 | 0.150716972 | 0.9997444 | Putative tRNA-m1A22 methylase |
| AP53_930 | 0.258738719 | 6.355157634 | 2.369461281 | 0.150978698 | 0.9997444 | Substrate-specific component PdxU |
| AP53_428 | -0.240175786 | 4.324315563 | 2.350057079 | 0.152514354 | 0.9997444 | ABC-type multidrug transport syste |
| AP53_121 | -0.208739985 | 6.021714868 | 2.326282008 | 0.154423093 | 0.9997444 | V-type ATP synthase subunit K (EC 3 |
| AP53_1511 | 0.206022712 | 7.467094329 | 2.300809387 | 0.156501972 | 0.9997444 | putative ABC transporter (ATP-bind |
| AP53_893 | 0.303298002 | 10.31994222 | 2.289906738 | 0.15740265 | 0.9997444 | Transcription regulator [contains d |
| AP53_711 | -0.532200826 | 2.784300559 | 2.275177043 | 0.158630004 | 0.9997444 | Mobile element protein |
| AP53_pseudo603 | 0.477069098 | 7.457202431 | 2.273727163 | 0.158751474 | 0.9997444 | Phage holin |
| AP53_189 | -0.134842637 | 8.5345132 | 2.270704188 | 0.159005119 | 0.9997444 | DNA repair protein RadA |
| AP53_134 | 0.156440493 | 8.862852606 | 2.268841146 | 0.159161696 | 0.9997444 | Nicotine adenine dinucleotide glyc |

| | | | | | | |
|---|---|---|---|---|---|---|
| AP53_1552 | -0.14415326 | 7.991986028 | 2.26709555 | 0.159308581 | 0.9997444 | Colicin V production protein |
| AP53_41 | -0.189170328 | 7.04652198 | 2.235442135 | 0.162002364 | 0.9997444 | Multi antimicrobial extrusion (MAT |
| AP53_1363 | 0.353220943 | 9.67730007 | 2.232274564 | 0.162275124 | 0.9997444 | Chromosome segregation ATPase |
| AP53_1213 | 0.286289268 | 7.298142231 | 2.214568667 | 0.16381064 | 0.9997444 | Phage CI-like repressor |
| AP53_1032 | -0.392763681 | 5.685132842 | 2.20997777 | 0.164211808 | 0.9997444 | hypothetical protein - phage associ |
| AP53_1724 | 0.262839714 | 8.526789921 | 2.191022252 | 0.165881581 | 0.9997444 | Pyruvate formate-lyase activating e |
| AP53_708 | 0.258391107 | 9.580698848 | 2.190904553 | 0.165892017 | 0.9997444 | PTS system, fructose-specific IIA cor |
| AP53_234 | -0.170759269 | 8.561751269 | 2.186491641 | 0.166283893 | 0.9997444 | Putative iron-sulfur cluster assembl |
| AP53_rna72 | 1.870431469 | 5.80827402 | 2.208186098 | 0.16648356 | 0.9997444 | Small Subunit Ribosomal RNA; ssuR |
| AP53_1768 | -0.687416083 | 7.359813173 | 2.18362038 | 0.166539504 | 0.9997444 | FIG01115181: hypothetical protei |
| AP53_rna54 | 4.551327126 | -0.548503406 | 2.244861038 | 0.166817347 | 0.9997444 | 5S RNA |
| AP53_233 | -0.191409353 | 8.635923669 | 2.177867136 | 0.167053198 | 0.9997444 | Cysteine desulfurase (EC 2.8.1.7), Su |
| AP53_196 | 0.138554705 | 8.712936276 | 2.171497179 | 0.167624327 | 0.9997444 | Ribonuclease P protein component |
| AP53_495 | -0.271963387 | 6.404686299 | 2.161997745 | 0.168480703 | 0.9997444 | FIG01114557: hypothetical protei |
| AP53_455 | 0.246424096 | 10.67283373 | 2.153869573 | 0.169217919 | 0.9997444 | Plasmid stabilization system toxin p |
| AP53_77 | 0.305660074 | 6.050714383 | 2.150712471 | 0.16950538 | 0.9997444 | HIT family protein |
| AP53_1689 | -0.450504267 | 2.901079833 | 2.149909952 | 0.169578551 | 0.9997444 | hypothetical protein |
| AP53_1467 | -0.18989453 | 9.113828949 | 2.145570225 | 0.169974936 | 0.9997444 | Acetyltransferase (EC 2.3.1.-) |
| AP53_936 | -0.376885727 | 8.458939196 | 2.144986036 | 0.170028386 | 0.9997444 | Lipoate-protein ligase A |
| AP53_941 | -0.376272792 | 8.638612252 | 2.141465456 | 0.170350955 | 0.9997444 | Probable NADH-dependent flavin o |
| AP53_357 | 0.176701952 | 9.9521086 | 2.139256446 | 0.170553754 | 0.9997444 | Peptidyl-prolyl cis-trans isomerase |
| AP53_820 | -0.49649111 | 9.906872654 | 2.13778128 | 0.170689356 | 0.9997444 | PTS system, mannose/fructose fami |
| AP53_280 | 0.1455816 | 8.991995718 | 2.136195881 | 0.170835244 | 0.9997444 | Predicted nucleoside phosphatase |
| AP53_1701 | 0.151369813 | 8.428105587 | 2.114578736 | 0.172840532 | 0.9997444 | ABC transporter permease protein |
| AP53_1851 | 0.196303843 | 9.03965394 | 2.10738895 | 0.173514183 | 0.9997444 | UTP--glucose-1-phosphate uridylylt |
| AP53_1406 | 0.190216412 | 9.002072289 | 2.089083642 | 0.175244625 | 0.9997444 | Cysteine ABC transporter, permease |
| AP53_970 | 0.206270844 | 6.591033471 | 2.085366532 | 0.175598722 | 0.9997444 | FIG01115860: hypothetical protei |
| AP53_1605 | -0.176771222 | 7.544310561 | 2.074380798 | 0.176650638 | 0.9997444 | tRNA pseudouridine synthase A (EC |
| AP53_1264 | -0.473224685 | 10.19077399 | 2.047052871 | 0.17930283 | 0.9997444 | Arginine/ornithine antiporter ArcD |
| AP53_1370 | -0.222737344 | 7.087991316 | 2.040825616 | 0.179914364 | 0.9997444 | FIG000605: protein co-occurring w |
| AP53_rna14 | 1.90565629 | 5.834101851 | 2.056748001 | 0.180446835 | 0.9997444 | Small Subunit Ribosomal RNA; ssuR |
| AP53_953 | -0.29233718 | 5.365556629 | 2.03078237 | 0.180906325 | 0.9997444 | Pantothenate kinase (EC 2.7.1.33) |
| AP53_rna80 | 1.918878835 | 5.803466699 | 2.050978097 | 0.181008417 | 0.9997444 | Small Subunit Ribosomal RNA; ssuR |
| AP53_122 | -0.222778326 | 6.032187503 | 2.02427073 | 0.181553251 | 0.9997444 | V-type ATP synthase subunit E (EC 3 |
| AP53_230 | -0.33934631 | 5.128972314 | 2.015510083 | 0.182428339 | 0.9997444 | hypothetical cytosolic protein |

| | | | | | | |
|---|---|---|---|---|---|---|
| AP53_169 | 0.175714316 | 9.092234106 | 2.012078529 | 0.182772598 | 0.9997444 | Zinc finger protein |
| AP53_rna57 | 0.431049482 | 5.964443146 | 1.980065735 | 0.186025045 | 0.9997444 | tRNA-Arg-CCT |
| AP53_652 | 0.280906169 | 8.254094096 | 1.979895278 | 0.186042563 | 0.9997444 | Phage transcriptional regulator, Cr |
| AP53_1310 | 0.216108355 | 8.517374264 | 1.968237658 | 0.187245714 | 0.9997444 | Phage protein |
| AP53_pseudo1183 | 0.425515809 | 4.144345891 | 1.947151292 | 0.189447742 | 0.9997444 | Phage holin |
| AP53_216 | -0.149079829 | 8.772890398 | 1.937852023 | 0.19042954 | 0.9997444 | 3'->5' exoribonuclease Bsu YhaM |
| AP53_600 | 0.366362414 | 10.95078582 | 1.935535397 | 0.190675152 | 0.9997444 | Phage protein |
| AP53_310 | -0.209567584 | 6.753458318 | 1.935067656 | 0.190724793 | 0.9997444 | hypothetical protein |
| AP53_rna4 | 2.067283104 | -0.831064152 | 1.925679451 | 0.191724711 | 0.9997444 | 5S RNA |
| AP53_rna29 | 0.596412157 | 4.81700164 | 1.920547635 | 0.192274172 | 0.9997444 | tRNA-Pseudo-TGA |
| AP53_877 | 0.134888937 | 8.076302094 | 1.920167604 | 0.192314944 | 0.9997444 | Peptide chain release factor 1 |
| AP53_1493 | 0.285321484 | 13.43323349 | 1.918870779 | 0.192454157 | 0.9997444 | Heat-inducible transcription repres |
| AP53_1186 | 0.358373973 | 2.275852303 | 1.917392886 | 0.192612967 | 0.9997444 | Phage hyaluronidase |
| AP53_1262 | -0.505674014 | 10.79217666 | 1.916753865 | 0.192681688 | 0.9997444 | Carbamate kinase (EC 2.7.2.2) |
| AP53_182 | -0.153810471 | 8.834217995 | 1.901827419 | 0.194295998 | 0.9997444 | UTP--glucose-1-phosphate uridylylt |
| AP53_126 | -0.199584288 | 6.282394857 | 1.900657365 | 0.194423284 | 0.9997444 | V-type ATP synthase subunit A (EC 3 |
| AP53_452 | 0.618139924 | 1.143102125 | 1.893624573 | 0.195190648 | 0.9997444 | FIG01116748: hypothetical protei |
| AP53_1641 | -0.199058095 | 8.263963792 | 1.87966907 | 0.196725095 | 0.9997444 | Serine acetyltransferase (EC 2.3.1.3 |
| AP53_1620 | -0.238927901 | 6.896683767 | 1.879202566 | 0.19677666 | 0.9997444 | Tagatose 1,6-diphosphate aldolase |
| AP53_1725 | 0.374179867 | 6.193449723 | 1.876487522 | 0.197077115 | 0.9997444 | FIG01114281: hypothetical protei |
| AP53_rna1 | 1.831017799 | 5.84424604 | 1.892337995 | 0.197385999 | 0.9997444 | Small Subunit Ribosomal RNA; ssuR |
| AP53_554 | -0.423061955 | 2.74527286 | 1.873042743 | 0.197459188 | 0.9997444 | Asparagine-rich ORFan next to Sag c |
| AP53_1139 | 0.161849882 | 6.556102348 | 1.867410302 | 0.198085988 | 0.9997444 | FIG01114175: hypothetical protei |
| AP53_1263 | -0.513762763 | 10.24910459 | 1.863321979 | 0.198542581 | 0.9997444 | Acetylornithine deacetylase/Succin |
| AP53_1592 | 0.158756752 | 9.927094547 | 1.859881597 | 0.198927876 | 0.9997444 | FIG001802: Putative alkaline-shock |
| AP53_1447 | -0.165107249 | 8.138222723 | 1.856152107 | 0.199346653 | 0.9997444 | PTS system, galactose-specific IIA co |
| AP53_714 | 1.588194703 | -0.182630266 | 1.854915352 | 0.19948578 | 0.9997444 | Phage protein |
| AP53_1849 | 0.178380885 | 8.259439319 | 1.849435787 | 0.200103725 | 0.9997444 | UDP-glucose dehydrogenase in hyal |
| AP53_1645 | -0.520709388 | 6.523461692 | 1.843245184 | 0.200804871 | 0.9997444 | Transport protein SgaT, putative |
| AP53_1802 | 0.41961296 | 3.801703709 | 1.837012378 | 0.201514046 | 0.9997444 | Phage encoded transcriptional regu |
| AP53_671 | 0.131111664 | 8.008832027 | 1.825511109 | 0.202831301 | 0.9997444 | 3-dehydroquinate dehydratase I (EC |
| AP53_78 | 0.162468867 | 7.419845793 | 1.82514463 | 0.202873459 | 0.9997444 | Tyrosyl-tRNA synthetase (EC 6.1.1.1 |
| AP53_1405 | 0.14557884 | 9.099489907 | 1.823178115 | 0.203099876 | 0.9997444 | ABC-type multidrug transport syste |
| AP53_292 | -0.140246696 | 7.715056802 | 1.819348971 | 0.2035417 | 0.9997444 | Ribosomal large subunit pseudouri |
| AP53_800 | 0.125019285 | 8.470305784 | 1.818847958 | 0.203599602 | 0.9997444 | Putative amidotransferase similar t |

| | | | | | | |
|---|---|---|---|---|---|---|
| AP53_1705 | 0.180349806 | 7.659559924 | 1.817161268 | 0.203794692 | 0.9997444 | FIG01116435: hypothetical protei |
| AP53_384 | 0.434639269 | 5.312484241 | 1.814696302 | 0.204080241 | 0.9997444 | Mobile element protein |
| AP53_383 | 0.368076628 | 6.861804942 | 1.801891244 | 0.205572092 | 0.9997444 | putative role in replication |
| AP53_rna51 | 1.741939668 | 5.804361528 | 1.814351555 | 0.206153558 | 0.9997444 | Small Subunit Ribosomal RNA; ssuR |
| AP53_1149 | 0.262251468 | 11.11120345 | 1.795672975 | 0.206301712 | 0.9997444 | Glucosamine-6-phosphate deamina |
| AP53_1676 | -0.362043772 | 7.594097753 | 1.795447648 | 0.206328214 | 0.9997444 | 6-aminohexanoate-cyclic-dimer hy |
| AP53_354 | -0.21422285 | 9.752665909 | 1.787981406 | 0.207208912 | 0.9997444 | Manganese ABC transporter, peripla |
| AP53_793 | 0.141147729 | 7.110767067 | 1.779805898 | 0.208178945 | 0.9997444 | ABC transporter ATP-binding protei |
| AP53_rna35 | 1.770328344 | 5.821934084 | 1.783182601 | 0.209801666 | 0.9997444 | Small Subunit Ribosomal RNA; ssuR |
| AP53_36 | 0.156869086 | 9.058116692 | 1.765329549 | 0.209911272 | 0.9997444 | Low molecular weight protein tyro |
| AP53_1633 | 0.235791543 | 6.935103688 | 1.76482122 | 0.209972446 | 0.9997444 | hypothetical protein |
| AP53_1446 | -0.13296013 | 8.608058006 | 1.754786675 | 0.211184849 | 0.9997444 | PTS system, galactose-specific IIB co |
| AP53_1268 | -0.243123042 | 5.460772353 | 1.74731997 | 0.212092986 | 0.9997444 | Transcriptional regulator ArcR esse |
| AP53_1457 | 0.152268707 | 10.3336213 | 1.74248668 | 0.212683579 | 0.9997444 | Transcription termination protein |
| AP53_1239 | -0.269000298 | 8.396129883 | 1.731203921 | 0.214070713 | 0.9997444 | Cell division protein YlmG/Ycf19 (p |
| AP53_99 | -0.176554052 | 8.792586277 | 1.727234672 | 0.214561541 | 0.9997444 | Deoxyadenosine kinase (EC 2.7.1.76 |
| AP53_1181 | 0.328018841 | 3.955616634 | 1.717971774 | 0.215712761 | 0.9997444 | Phage lysin, N-acetylmuramoyl-L-al |
| AP53_299 | 0.133220818 | 8.536673103 | 1.713403216 | 0.21628356 | 0.9997444 | Hemolysins and related proteins co |
| AP53_270 | -0.142518472 | 8.047303966 | 1.708849259 | 0.216854522 | 0.9997444 | Helicase loader DnaB |
| AP53_96 | -0.308043792 | 6.056549802 | 1.699697139 | 0.218008029 | 0.9997444 | FIG009688: Thioredoxin |
| AP53_601 | 0.407676096 | 8.445616324 | 1.69070886 | 0.2191488 | 0.9997444 | Phage lysin |
| AP53_665 | 0.154135729 | 8.344185197 | 1.687913094 | 0.219505242 | 0.9997444 | Cytidylate kinase (EC 2.7.4.14) |
| AP53_1679 | -0.172618665 | 7.966748178 | 1.679103524 | 0.220633441 | 0.9997444 | Chromosome segregation helicase |
| AP53_938 | -0.315862483 | 8.540954766 | 1.677676816 | 0.220816874 | 0.9997444 | COG2110, Macro domain, possibly |
| AP53_282 | -0.156689577 | 8.804343559 | 1.675160417 | 0.221140904 | 0.9997444 | FIG005986: HD family hydrolase |
| AP53_327 | 0.166339772 | 8.892621413 | 1.666218043 | 0.222297504 | 0.9997444 | FIG01118451: hypothetical protei |
| AP53_1468 | -0.38348055 | 8.166070866 | 1.666184229 | 0.222301893 | 0.9997444 | ATPase YjeE, predicted to have esse |
| AP53_1492 | 0.258585108 | 14.03692929 | 1.661835356 | 0.222867289 | 0.9997444 | Heat shock protein GrpE |
| AP53_1035 | -0.40597242 | 4.328280742 | 1.655464309 | 0.223699038 | 0.9997444 | FIG01118889: hypothetical protei |
| AP53_1744 | 0.4735488 | 5.749783198 | 1.649816133 | 0.224439866 | 0.9997444 | Imidazolonepropionase (EC 3.5.2.7 |
| AP53_154 | 0.172788102 | 9.987487702 | 1.644738328 | 0.225108669 | 0.9997444 | M protein trans-acting positive regu |
| AP53_1711 | 0.453984963 | 3.491323419 | 1.642052163 | 0.225463537 | 0.9997444 | hypothetical protein |
| AP53_pseudo1680 | 0.273256558 | 5.028817522 | 1.641678486 | 0.225512962 | 0.9997444 | Streptococcal mitogenic exotoxin Z |
| AP53_1312 | 0.348344562 | 6.203251073 | 1.637940812 | 0.226008129 | 0.9997444 | Phage lysin |
| AP53_1828 | 0.313957643 | 5.847888352 | 1.637654875 | 0.226046069 | 0.9997444 | Transcriptional regulator, TetR fam |

| | | | | | | |
|---|---|---|---|---|---|---|
| AP53_1697 | 0.172085583 | 8.455477189 | 1.637156483 | 0.22611222 | 0.9997444 | Hypothetical mga-associated prote |
| AP53_1507 | -0.212312257 | 8.566607648 | 1.631139809 | 0.226912841 | 0.9997444 | Universal stress protein family |
| AP53_95 | -0.258061423 | 6.349589747 | 1.630611045 | 0.226983382 | 0.9997444 | FIG014356: hypothetical protein |
| AP53_1549 | -0.263183296 | 8.063844327 | 1.625690377 | 0.22764124 | 0.9997444 | Thioredoxin |
| AP53_1686 | -0.673453208 | 7.500481105 | 1.617455431 | 0.228747885 | 0.9997444 | FIG01115696: hypothetical protei |
| AP53_1854 | 0.131656494 | 9.006424022 | 1.606173109 | 0.230275721 | 0.9997444 | Putative sugar uptake protein SpyM |
| AP53_rna82 | -0.747119266 | 2.664243267 | 1.585042136 | 0.233174085 | 0.9997444 | tRNA-Cys-GCA |
| AP53_1085 | -0.338904568 | 4.943668624 | 1.583070563 | 0.233446991 | 0.9997444 | PTS system, cellobiose-specific IIB c |
| AP53_870 | -0.146683172 | 8.825899477 | 1.577153416 | 0.234268611 | 0.9997444 | L-proline glycine betaine binding A |
| AP53_822 | -0.469297434 | 9.76644687 | 1.575223608 | 0.23453741 | 0.9997444 | PTS system, mannose-specific IID co |
| AP53_552 | -0.332285745 | 6.275588373 | 1.574116663 | 0.234691779 | 0.9997444 | Streptolysin S export transmembra |
| AP53_rna2 | 0.529036886 | 3.836238549 | 1.560649184 | 0.23658084 | 0.9997444 | tRNA-Ala-TGC |
| AP53_1362 | 0.274159851 | 9.411929701 | 1.558180788 | 0.236929285 | 0.9997444 | Phage protein |
| AP53_764 | 0.187380618 | 7.227939036 | 1.556977368 | 0.237099412 | 0.9997444 | Glutathione S-transferase domain p |
| AP53_1626 | 0.409679425 | 7.678876515 | 1.552895481 | 0.237677688 | 0.9997444 | Toxin to DNA-damage-inducible pro |
| AP53_987 | 0.232604944 | 9.004300497 | 1.552061106 | 0.237796125 | 0.9997444 | Amino acid ABC transporter, aminc |
| AP53_1396 | -0.333007056 | 4.564997894 | 1.551391207 | 0.237891273 | 0.9997444 | FIG005686: hypothetical protein |
| AP53_1653 | -0.16351641 | 6.900789998 | 1.550049326 | 0.238082019 | 0.9997444 | Transcriptional regulator, MarR fam |
| AP53_1448 | -0.145539217 | 8.01963308 | 1.548307877 | 0.238329868 | 0.9997444 | Lactose phosphotransferase system |
| AP53_1443 | -0.132167798 | 8.795064872 | 1.536242932 | 0.24005654 | 0.9997444 | Galactose-6-phosphate isomerase, |
| AP53_1033 | -0.468703781 | 5.62939968 | 1.53462815 | 0.240288912 | 0.9997444 | hypothetical protein |
| AP53_1584 | -0.115515915 | 9.405547829 | 1.528252028 | 0.241209416 | 0.9997444 | Ribonuclease J1 (endonuclease and |
| AP53_272 | -0.119065335 | 9.628509917 | 1.525598477 | 0.2415939 | 0.9997444 | GTP-binding protein EngA |
| AP53_1357 | 0.411514713 | 4.280988049 | 1.523685036 | 0.241871657 | 0.9997444 | conserved hypothetical protein - pl |
| AP53_rna48 | 0.544621158 | 5.123122914 | 1.511121377 | 0.24370613 | 0.9997444 | tRNA-His-GTG |
| AP53_rna50 | 0.667119768 | 3.729627845 | 1.50788769 | 0.244181324 | 0.9997444 | tRNA-Leu-CAA |
| AP53_1068 | -0.242607903 | 6.318886915 | 1.499022243 | 0.245490532 | 0.9997444 | Maltose/maltodextrin ABC transpo |
| AP53_rna83 | 0.795813434 | 3.752216362 | 1.495787461 | 0.245970587 | 0.9997444 | tRNA-Asn-GTT |
| AP53_1259 | 0.150130179 | 8.599922287 | 1.495201673 | 0.246057656 | 0.9997444 | Phosphopantetheine adenylyltrans |
| AP53_180 | -0.473745695 | 2.720633585 | 1.488262345 | 0.247092256 | 0.9997444 | hypothetical protein |
| AP53_421 | -0.127013462 | 8.761816951 | 1.488254054 | 0.247093495 | 0.9997444 | Catabolite control protein A |
| AP53_1102 | -0.273495158 | 6.954638951 | 1.488216812 | 0.247099064 | 0.9997444 | Dihydrolipoamide acyltransferases |
| AP53_440 | 0.382295369 | 2.590914792 | 1.486359792 | 0.247376937 | 0.9997444 | FIG01114171: hypothetical protei |
| AP53_1403 | 0.13949383 | 8.232437424 | 1.481823237 | 0.248057538 | 0.9997444 | Thioredoxin reductase (EC 1.8.1.9) |
| AP53_896 | 0.238179197 | 9.148982694 | 1.48114155 | 0.248160027 | 0.9997444 | 50S ribosomal subunit maturation |

| | | | | | | |
|---|---|---|---|---|---|---|
| AP53_rna49 | 0.574076356 | 5.221521578 | 1.476627604 | 0.248840129 | 0.9997444 | tRNA-Gln-TTG |
| AP53_1338 | 0.295706957 | 3.122952067 | 1.476558311 | 0.248850589 | 0.9997444 | FIG01114733: hypothetical protei |
| AP53_824 | -0.21422061 | 5.45028148 | 1.465719409 | 0.250494071 | 0.9997444 | Two-component response regulato |
| AP53_975 | 0.204473526 | 5.218803085 | 1.458294181 | 0.251628433 | 0.9997444 | Transcriptional regulator, TetR fam |
| AP53_1284 | -0.247648407 | 6.949306293 | 1.457872327 | 0.251693089 | 0.9997444 | Aldose 1-epimerase (EC 5.1.3.3) |
| AP53_325 | -0.522580678 | 6.18675495 | 1.452724052 | 0.252483962 | 0.9997444 | hypothetical protein |
| AP53_rna78 | 1.564177801 | 5.81260375 | 1.463964954 | 0.252662893 | 0.9997444 | Large Subunit Ribosomal RNA; lsuR |
| AP53_128 | -0.189223074 | 6.379202322 | 1.451096499 | 0.252734686 | 0.9997444 | V-type ATP synthase subunit D (EC 3 |
| AP53_1196 | -0.447347844 | 3.612372154 | 1.446840163 | 0.253391971 | 0.9997444 | FIG042840: Phage protein |
| AP53_1269 | -0.15483094 | 7.264681156 | 1.444449064 | 0.253762234 | 0.9997444 | Arginine pathway regulatory protei |
| AP53_1745 | 0.451623348 | 5.866330514 | 1.433026689 | 0.255541166 | 0.9997444 | Urocanate hydratase (EC 4.2.1.49) |
| AP53_937 | -0.285958876 | 8.459560788 | 1.432038381 | 0.255695881 | 0.9997444 | NAD-dependent protein deacetylas |
| AP53_1224 | -0.130538844 | 8.260536247 | 1.429150052 | 0.256148767 | 0.9997444 | Octaprenyl diphosphate synthase ( |
| AP53_895 | 0.29549432 | 9.171990467 | 1.424840631 | 0.256826508 | 0.9997444 | FIG01115071: hypothetical protei |
| AP53_rna16 | 1.565358162 | 5.830256733 | 1.435834547 | 0.256990248 | 0.9997444 | Large Subunit Ribosomal RNA; lsuR |
| AP53_670 | 0.127271055 | 8.14152311 | 1.41612282 | 0.258205031 | 0.9997444 | LSU m5C1962 methyltransferase Rl |
| AP53_1603 | -0.168682047 | 6.695972091 | 1.411546132 | 0.258932765 | 0.9997444 | Substrate-specific component PdxU |
| AP53_349 | -0.131558865 | 8.400918934 | 1.408060388 | 0.259488904 | 0.9997444 | ADP-ribose pyrophosphatase (EC 3. |
| AP53_40 | -0.393230835 | 10.27616672 | 1.407941075 | 0.259507969 | 0.9997444 | Alcohol dehydrogenase (EC 1.1.1.1) |
| AP53_1733 | 0.182430549 | 7.51982531 | 1.405619438 | 0.259879319 | 0.9997444 | transcriptional regulator, MutR fam |
| AP53_817 | 0.129223896 | 9.179772291 | 1.401174726 | 0.260592283 | 0.9997444 | Peptide methionine sulfoxide redu |
| AP53_646 | 0.300560089 | 3.949294806 | 1.39463392 | 0.261646337 | 0.9997444 | hypothetical phage protein |
| AP53_766 | -0.254485723 | 9.494367576 | 1.384286028 | 0.263325827 | 0.9997444 | FIG01115306: hypothetical protei |
| AP53_1331 | 0.264989332 | 7.374790994 | 1.376329244 | 0.264627271 | 0.9997444 | hypothetical protein, phage associa |
| AP53_699 | 0.239198721 | 5.236852529 | 1.370544824 | 0.265578923 | 0.9997444 | Cobalt-zinc-cadmium resistance pr |
| AP53_rna53 | 1.503668009 | 5.811675989 | 1.379605922 | 0.265949875 | 0.9997444 | Large Subunit Ribosomal RNA; lsuR |
| AP53_1148 | 0.204870064 | 11.24128382 | 1.366044565 | 0.266322549 | 0.9997444 | Ribosomal small subunit pseudouri |
| AP53_859 | -0.213960873 | 7.980890273 | 1.363707636 | 0.266709831 | 0.9997444 | FIG01114333: hypothetical protei |
| AP53_rna23 | 0.614863238 | 2.598266444 | 1.347857981 | 0.269356966 | 0.9997444 | tRNA-Gly-GCC |
| AP53_1712 | 0.247963605 | 5.106717319 | 1.347845898 | 0.269358997 | 0.9997444 | RopB; Rgg-like transcription regula |
| AP53_1225 | -0.214363518 | 7.639382578 | 1.346660514 | 0.269558424 | 0.9997444 | Exodeoxyribonuclease VII small sub |
| AP53_1188 | 0.335577197 | 2.169798669 | 1.341377256 | 0.270449738 | 0.9997444 | Phage hyaluronidase; Phage endop |
| AP53_767 | 0.13466993 | 6.583723387 | 1.335516618 | 0.271443205 | 0.9997444 | tRNA dimethylallyltransferase (EC 2 |
| AP53_934 | -0.220647418 | 8.425851427 | 1.329681722 | 0.272437302 | 0.9997444 | Cardiolipin synthetase (EC 2.7.8.-) |
| AP53_854 | 0.189379963 | 6.218076921 | 1.32881775 | 0.272584924 | 0.9997444 | hypothetical protein |

| | | | | | | |
|---|---|---|---|---|---|---|
| AP53_194 | -0.166802646 | 6.239699763 | 1.327577677 | 0.272797 | 0.9997444 | Sensory transduction protein kinas |
| AP53_rna6 | 0.443501293 | 4.56855931 | 1.322230871 | 0.273714013 | 0.9997444 | tRNA-Asp-GTC |
| AP53_805 | 0.215471697 | 4.94977484 | 1.321741466 | 0.273798162 | 0.9997444 | FIG01115418: hypothetical protei |
| AP53_1672 | -0.219382922 | 7.82243254 | 1.319787176 | 0.274134538 | 0.9997444 | PTS system, maltose and glucose-sp |
| AP53_394 | 0.231199931 | 4.161062865 | 1.312321307 | 0.275424838 | 0.9997444 | Competence-stimulating peptide A |
| AP53_381 | 0.202838498 | 5.405779303 | 1.309943315 | 0.275837578 | 0.9997444 | hypothetical protein |
| AP53_1625 | 0.30496597 | 6.69419007 | 1.308614503 | 0.276068587 | 0.9997444 | DNA-damage-inducible protein J |
| AP53_1049 | -0.349319476 | 4.90069802 | 1.306490318 | 0.276438423 | 0.9997444 | hypothetical phage protein |
| AP53_988 | 0.258119099 | 9.577451512 | 1.305773188 | 0.276563436 | 0.9997444 | ABC-type polar amino acid transpo |
| AP53_821 | -0.418802655 | 9.889874738 | 1.304811138 | 0.276731266 | 0.9997444 | PTS system, mannose-specific IIC cc |
| AP53_1296 | -0.16251253 | 6.172112517 | 1.30084953 | 0.277423855 | 0.9997444 | Multiple sugar ABC transporter, me |
| AP53_1726 | -0.542149526 | 9.766519192 | 1.298457558 | 0.277843193 | 0.9997444 | Preprotein translocase subunit Sec |
| AP53_rna43 | 0.533945759 | 4.568139016 | 1.296761345 | 0.278141088 | 0.9997444 | tRNA-Pseudo-TGA |
| AP53_765 | 0.218424994 | 8.431233271 | 1.295164012 | 0.278422021 | 0.9997444 | Exfoliative toxin A |
| AP53_556 | -0.160645508 | 9.196164744 | 1.293936641 | 0.278638154 | 0.9997444 | Substrate-specific component Que |
| AP53_1380 | 0.183904227 | 8.52007152 | 1.288791966 | 0.27954663 | 0.9997444 | Methionyl-tRNA formyltransferase |
| AP53_rna17 | 2.195293191 | -0.659752514 | 1.287646544 | 0.279749453 | 0.9997444 | 5S RNA |
| AP53_rna37 | 1.459633973 | 5.803484641 | 1.295057363 | 0.280254134 | 0.9997444 | Large Subunit Ribosomal RNA; lsuR |
| AP53_449 | -0.514373147 | 1.82602402 | 1.284622479 | 0.280285914 | 0.9997444 | hypothetical protein |
| AP53_986 | 0.172661173 | 7.078026064 | 1.28097299 | 0.280935222 | 0.9997444 | CAMP factor |
| AP53_1490 | 0.21860912 | 13.61212773 | 1.277540439 | 0.281547838 | 0.9997444 | Chaperone protein DnaJ |
| AP53_1465 | -0.106413288 | 8.952617724 | 1.277198298 | 0.281609002 | 0.9997444 | Hypothetical protein in cluster wit |
| AP53_387 | -0.506246049 | 1.849198738 | 1.276770798 | 0.281685451 | 0.9997444 | hypothetical protein |
| AP53_1550 | -0.155646737 | 7.688059374 | 1.275501838 | 0.281912549 | 0.9997444 | Phosphatidylglycerophosphatase B |
| AP53_rna70 | 1.468716954 | 5.783091751 | 1.279686838 | 0.282968988 | 0.9997444 | Large Subunit Ribosomal RNA; lsuR |
| AP53_209 | -0.130596572 | 7.368054621 | 1.267899349 | 0.28327845 | 0.9997444 | Putative deoxyribonuclease YcfH |
| AP53_1037 | -0.258403407 | 5.399904477 | 1.265372098 | 0.283734543 | 0.9997444 | FIG01114622: hypothetical protei |
| AP53_rna77 | 2.102421942 | -1.139188048 | 1.256533034 | 0.285337782 | 0.9997444 | 5S RNA |
| AP53_1735 | -0.222851818 | 7.533997899 | 1.250929986 | 0.286360601 | 0.9997444 | hypothetical protein |
| AP53_1187 | 0.310988619 | 2.279544779 | 1.239267377 | 0.288506011 | 0.9997444 | Hyaluronate lyase (phage associate |
| AP53_1823 | -0.13305456 | 5.856257941 | 1.238187529 | 0.288705787 | 0.9997444 | Phosphohydrolase (MutT/nudix fan |
| AP53_294 | 0.141654955 | 7.702865212 | 1.237434906 | 0.28884514 | 0.9997444 | tRNA (cytidine(34)-2'-O)-methyltra |
| AP53_1257 | -0.133431592 | 7.218499909 | 1.234295699 | 0.289427398 | 0.9997444 | LacI family regulatory protein |
| AP53_923 | 0.174628705 | 8.605489428 | 1.231785951 | 0.289894083 | 0.9997444 | Signal recognition particle associat |
| AP53_rna3 | 1.449953603 | 5.832630068 | 1.238276483 | 0.290469009 | 0.9997444 | Large Subunit Ribosomal RNA; lsuR |

| | | | | | | |
|---|---|---|---|---|---|---|
| AP53_1452 | 0.132369997 | 8.900423862 | 1.226310201 | 0.290915948 | 0.9997444 | putative esterase |
| AP53_1772 | 0.315973418 | 11.24275464 | 1.225960497 | 0.290981379 | 0.9997444 | Ortholog of S. aureus MRSA252 (BX |
| AP53_206 | -0.159137974 | 6.775523524 | 1.22128519 | 0.291858132 | 0.9997444 | N-acetylneuraminate lyase (EC 4.1.3 |
| AP53_1057 | -0.409054516 | 4.900415044 | 1.218958375 | 0.292295854 | 0.9997444 | FIG01115240: hypothetical protein |
| AP53_195 | -0.162380519 | 6.691933789 | 1.209387154 | 0.294106094 | 0.9997444 | Response regulator of the competen |
| AP53_1131 | -0.125900572 | 8.424826186 | 1.208736665 | 0.294229692 | 0.9997444 | Glutaredoxin-like protein NrdH, red |
| AP53_555 | -0.172349025 | 8.73793431 | 1.199236269 | 0.296043183 | 0.9997444 | Endonuclease/Exonuclease/phosph |
| AP53_290 | -0.126478471 | 7.280403496 | 1.195225125 | 0.296813564 | 0.9997444 | Segregation and condensation prot |
| AP53_1124 | -0.225230379 | 4.615390786 | 1.194403982 | 0.29697162 | 0.9997444 | FIG01113965: hypothetical protein |
| AP53_1427 | 0.20965904 | 11.35827444 | 1.185937593 | 0.298608175 | 0.9997444 | FIG01116327: hypothetical protein |
| AP53_1117 | -0.188688936 | 8.568234233 | 1.185129988 | 0.298764948 | 0.9997444 | S-adenosylmethionine synthetase ( |
| AP53_1722 | 0.16766792 | 9.069333087 | 1.18404329 | 0.29897608 | 0.9997444 | Deoxyribonucleoside regulator Dec |
| AP53_1245 | -0.110346904 | 8.936590676 | 1.179373372 | 0.299885786 | 0.9997444 | UDP-N-acetylglucosamine--N-acety |
| AP53_1359 | 0.388978284 | 10.7346197 | 1.179156577 | 0.299928112 | 0.9997444 | Phage antirepressor protein |
| AP53_1138 | 0.177930595 | 6.670704766 | 1.178889091 | 0.299980347 | 0.9997444 | Membrane-bound protease, CAAX f |
| AP53_1462 | -0.116944147 | 7.438063323 | 1.176350836 | 0.300476658 | 0.9997444 | ABC transporter, permease protein |
| AP53_1361 | 0.296995054 | 9.39071655 | 1.171486281 | 0.301431075 | 0.9997444 | putative cI-like repressor - phage as |
| AP53_318 | 0.12158819 | 10.427855 | 1.170038183 | 0.301716015 | 0.9997444 | rRNA small subunit methyltransfera |
| AP53_1568 | 0.334996615 | 4.251335018 | 1.169508073 | 0.301820419 | 0.9997444 | Transcriptional regulator, Cro/CI fa |
| AP53_1441 | -0.111579588 | 8.92053845 | 1.167653655 | 0.302186044 | 0.9997444 | Tagatose 1,6-diphosphate aldolase |
| AP53_1311 | 0.225936756 | 8.270877675 | 1.165248071 | 0.30266127 | 0.9997444 | Phage protein |
| AP53_1235 | 0.18263001 | 10.02292846 | 1.164190229 | 0.302870582 | 0.9997444 | FIG01114027: hypothetical protein |
| AP53_1593 | -0.342604443 | 8.927438765 | 1.164093573 | 0.302889718 | 0.9997444 | LSU ribosomal protein L28p |
| AP53_905 | 0.197384247 | 5.511271964 | 1.151440821 | 0.30540944 | 0.9997444 | Methylmalonyl-CoA decarboxylase, |
| AP53_1547 | -0.113762578 | 8.074757721 | 1.14874366 | 0.305950395 | 0.9997444 | A/G-specific adenine glycosylase (EC |
| AP53_1003 | 0.238681923 | 6.570567463 | 1.148692361 | 0.305960698 | 0.9997444 | Streptococcal pyrogenic exotoxin K |
| AP53_1590 | 0.151445781 | 9.872711188 | 1.148516528 | 0.305996012 | 0.9997444 | Putative stomatin/prohibitin-famil |
| AP53_1821 | -0.408402576 | 5.51795796 | 1.146106799 | 0.306480569 | 0.9997444 | Transcriptional regulator |
| AP53_845 | 0.182219091 | 5.452419827 | 1.14257091 | 0.307193549 | 0.9997444 | Two-component response regulato |
| AP53_320 | 0.135650676 | 8.206756372 | 1.138522812 | 0.308012701 | 0.9997444 | Cytoplasmic copper homeostasis pr |
| AP53_1126 | 0.132791841 | 8.692747408 | 1.138206568 | 0.308076825 | 0.9997444 | ATP-dependent RNA helicase YfmL |
| AP53_1228 | -0.193038846 | 10.22004434 | 1.137783046 | 0.308162731 | 0.9997444 | putative phosphomannomutase |
| AP53_1703 | 0.240240831 | 8.116455482 | 1.136850836 | 0.308351937 | 0.9997444 | periplasmic component of efflux sy |
| AP53_935 | -0.224874056 | 9.235748517 | 1.134646818 | 0.308799933 | 0.9997444 | Formate--tetrahydrofolate ligase (E |
| AP53_794 | -0.287126988 | 9.90566459 | 1.124182887 | 0.310939528 | 0.9997444 | Acetoin dehydrogenase E1 compon |

| | | | | | | |
|---|---|---|---|---|---|---|
| AP53_1710 | 0.295558323 | 4.837104462 | 1.124099988 | 0.310956563 | 0.9997444 | FIG01114305: hypothetical protei |
| AP53_472 | 0.189208954 | 11.25387969 | 1.122629337 | 0.311258983 | 0.9997444 | Phage shock protein C, putative; str |
| AP53_1365 | 0.260370995 | 9.690534392 | 1.121581148 | 0.311474785 | 0.9997444 | Regulatory protein recX |
| AP53_32 | -0.290980775 | 5.687921236 | 1.120199327 | 0.311759601 | 0.9997444 | hypothetical protein |
| AP53_876 | 0.15790252 | 7.431415743 | 1.115876653 | 0.312652971 | 0.9997444 | Thymidine kinase (EC 2.7.1.21) |
| AP53_256 | 0.144548784 | 8.209398764 | 1.114167381 | 0.313007234 | 0.9997444 | Methionine ABC transporter substr |
| AP53_456 | 0.174470225 | 10.88979319 | 1.112602529 | 0.313332066 | 0.9997444 | hypothetical protein |
| AP53_350 | -0.121192358 | 8.629141679 | 1.100079401 | 0.315949021 | 0.9997444 | FIG01115635: hypothetical protei |
| AP53_pseudo656 | 0.189466499 | 8.518667932 | 1.091362574 | 0.317789039 | 0.9997444 | Lipoteichoic acid synthase LtaS Typ |
| AP53_188 | -0.253616565 | 7.499966239 | 1.091072095 | 0.317850619 | 0.9997444 | Deoxyuridine 5'-triphosphate nucle |
| AP53_654 | 0.146918287 | 8.648977839 | 1.089178498 | 0.318252472 | 0.9997444 | Pathogenicity island SaPIn2 |
| AP53_105 | -0.1413556 | 7.031738335 | 1.078771463 | 0.320474062 | 0.9997444 | Fibronectin-binding protein/major |
| AP53_1374 | 0.133140179 | 8.160834778 | 1.074253665 | 0.321445404 | 0.9997444 | Two component transcriptional reg |
| AP53_894 | 0.259654883 | 9.692295089 | 1.071895296 | 0.32195414 | 0.9997444 | Hemolysin III |
| AP53_1601 | -0.107619705 | 7.763040086 | 1.063529229 | 0.323768193 | 0.9997444 | Hypothetical protein ywlG |
| AP53_232 | -0.154928177 | 8.703000013 | 1.062777689 | 0.323931872 | 0.9997444 | Iron-sulfur cluster assembly protei |
| AP53_1561 | -0.181449734 | 8.081135354 | 1.059612068 | 0.324622626 | 0.9997444 | Beta-lactamase, class C |
| AP53_110 | 0.155460799 | 8.013028847 | 1.054018817 | 0.325848297 | 0.9997444 | FIG01114239: hypothetical protei |
| AP53_203 | -0.158582646 | 6.238333279 | 1.053275047 | 0.326011785 | 0.9997444 | ABC transporter, predicted N-acety |
| AP53_304 | 0.152384413 | 7.635473271 | 1.051222696 | 0.326463524 | 0.9997444 | ABC-type Fe3+-siderophore transpc |
| AP53_433 | -0.135129652 | 8.225934195 | 1.049973894 | 0.326738838 | 0.9997444 | Two-component response regulato |
| AP53_664 | -0.26090946 | 7.83188895 | 1.048961151 | 0.326962355 | 0.9997444 | FIG01114555: hypothetical protei |
| AP53_1062 | 0.130791554 | 7.882999782 | 1.04727945 | 0.327334003 | 0.9997444 | lipoprotein, putative |
| AP53_1734 | -0.222664829 | 7.484039976 | 1.046217915 | 0.327568912 | 0.9997444 | Transcriptional regulator, MutR fan |
| AP53_501 | 0.130371634 | 8.033709296 | 1.044226426 | 0.328010268 | 0.9997444 | Promiscuous sugar phosphatase Yic |
| AP53_569 | -0.130890131 | 8.07311659 | 1.043965841 | 0.328068083 | 0.9997444 | Competence-associated EpuA prote |
| AP53_248 | 0.150657244 | 8.921296728 | 1.038606016 | 0.329260514 | 0.9997444 | FIG004454: RNA binding protein |
| AP53_1141 | -0.306391223 | 5.045558232 | 1.035851362 | 0.329875798 | 0.9997444 | FIG01114322: hypothetical protei |
| AP53_906 | 0.143443408 | 5.05535495 | 1.033811213 | 0.330332562 | 0.9997444 | 2-(5''-triphosphoribosyl)-3'-dephos |
| AP53_1596 | 0.149099657 | 7.534487352 | 1.032783769 | 0.330562941 | 0.9997444 | hypothetical protein |
| AP53_892 | 0.187071114 | 12.79974523 | 1.03074696 | 0.331020333 | 0.9997444 | FIG01115412: hypothetical protei |
| AP53_843 | -0.132866936 | 7.471071188 | 1.030107187 | 0.331164191 | 0.9997444 | Spermidine Putrescine ABC transpc |
| AP53_112 | -0.208137054 | 5.848035247 | 1.028553143 | 0.331514009 | 0.9997444 | Transcriptional regulators, LysR fan |
| AP53_551 | -0.232812313 | 7.020067052 | 1.022172068 | 0.332956031 | 0.9997444 | Export ABC transporter ATP-binding |
| AP53_391 | -0.195941867 | 5.127292272 | 1.020210423 | 0.333401161 | 0.9997444 | Histidine kinase of the competence |

| | | | | | | |
|---|---|---|---|---|---|---|
| AP53_363 | 0.11578301 | 9.754538294 | 1.018146257 | 0.333870489 | 0.9997444 | Ribosome recycling factor |
| AP53_1739 | -0.380739515 | 9.058686756 | 1.015952275 | 0.334370385 | 0.9997444 | Transcriptional regulator CtsR |
| AP53_1342 | -0.280324956 | 4.625304636 | 1.013492913 | 0.334932042 | 0.9997444 | Phage protein |
| AP53_1081 | -0.364301904 | 4.303329124 | 1.013475941 | 0.334935923 | 0.9997444 | FIG01114324: hypothetical protei |
| AP53_698 | 0.203753284 | 8.420522513 | 1.010568685 | 0.335601651 | 0.9997444 | FIG01115236: hypothetical protei |
| AP53_1548 | 0.105653398 | 7.216243169 | 1.008012218 | 0.336188644 | 0.9997444 | Transcriptional regulator, XRE fami |
| AP53_653 | 0.185678632 | 7.969931946 | 1.007431029 | 0.3363223 | 0.9997444 | Phage protein |
| AP53_1586 | 0.24228012 | 7.962072421 | 0.996549031 | 0.338839198 | 0.9997444 | Transcriptional regulator, MerR fan |
| AP53_974 | 0.132353824 | 5.52041317 | 0.995586496 | 0.339063144 | 0.9997444 | ABC transporter ATP-binding protei |
| AP53_1580 | -0.14845667 | 8.126588309 | 0.990858778 | 0.340166253 | 0.9997444 | TsaD/Kae1/Qri7 protein, required f |
| AP53_543 | -0.113521717 | 9.723083223 | 0.980100629 | 0.342696072 | 0.9997444 | Septation ring formation regulator |
| AP53_723 | 0.202259813 | 7.192492431 | 0.973535124 | 0.344253538 | 0.9997444 | GTP pyrophosphokinase (EC 2.7.6.5 |
| AP53_971 | 0.192747029 | 7.836608819 | 0.972216632 | 0.344567562 | 0.9997444 | membrane protein |
| AP53_825 | -0.178689017 | 5.835413574 | 0.970160964 | 0.345057998 | 0.9997444 | Ferric iron ABC transporter, iron-bi |
| AP53_1376 | 0.134366286 | 6.677968156 | 0.960251334 | 0.347436666 | 0.9997444 | Transporter associated with VraSR |
| AP53_1814 | -0.253898033 | 9.414836082 | 0.958052453 | 0.347967746 | 0.9997444 | LSU ribosomal protein L32p |
| AP53_1508 | 0.106350036 | 8.636400623 | 0.957014102 | 0.348218948 | 0.9997444 | Hydrolase (HAD superfamily) |
| AP53_1317 | -0.189466477 | 3.704008296 | 0.946982616 | 0.35065963 | 0.9997444 | Hyaluronate lyase (phage associate |
| AP53_1730 | 0.10789457 | 8.408576889 | 0.946719491 | 0.350723989 | 0.9997444 | Similar to ribosomal large subunit |
| AP53_251 | -0.162880824 | 7.811421738 | 0.946136555 | 0.350866632 | 0.9997444 | Iojap protein |
| AP53_1214 | 0.189268316 | 7.586571696 | 0.945862667 | 0.350933682 | 0.9997444 | Phage protein |
| AP53_424 | 0.210207243 | 6.560053996 | 0.940626456 | 0.352219193 | 0.9997444 | hypothetical protein |
| AP53_1381 | 0.189847636 | 7.592901113 | 0.940443945 | 0.352264126 | 0.9997444 | Helicase PriA essential for oriC/Dna |
| AP53_966 | -0.270179085 | 5.433533808 | 0.940396724 | 0.352275753 | 0.9997444 | FIG006438: hypothetical protein |
| AP53_1077 | -0.239609672 | 6.908046039 | 0.938535779 | 0.352734409 | 0.9997444 | FIG01114114: hypothetical protei |
| AP53_1551 | -0.114481785 | 7.86464506 | 0.931828417 | 0.354394892 | 0.9997444 | Recombination inhibitory protein |
| AP53_586 | 0.119975757 | 10.17786669 | 0.93065424 | 0.354686764 | 0.9997444 | RNA polymerase sigma factor RpoD |
| AP53_1704 | 0.189937348 | 8.338143977 | 0.92639766 | 0.355747841 | 0.9997444 | FIG01115531: hypothetical protei |
| AP53_1808 | -0.192452504 | 7.402905307 | 0.92271787 | 0.35666894 | 0.9997444 | bacteriocin immunity protein |
| AP53_1540 | -0.094706462 | 9.211085047 | 0.920086562 | 0.357329765 | 0.9997444 | Excinuclease ABC subunit A |
| AP53_253 | 0.109736189 | 7.304285585 | 0.915856094 | 0.358396025 | 0.9997444 | FIG007079: UPF0348 protein famil |
| AP53_1664 | 0.128683862 | 7.418852833 | 0.913840679 | 0.358905663 | 0.9997444 | Leucine rich protein |
| AP53_1145 | 0.22199497 | 7.188093254 | 0.913458404 | 0.35900245 | 0.9997444 | Major facilitator:Oxalate:Formate |
| AP53_120 | -0.141886059 | 5.734036467 | 0.913012766 | 0.35911533 | 0.9997444 | V-type ATP synthase subunit I (EC 3. |
| AP53_755 | 0.115843936 | 8.12430739 | 0.90582638 | 0.360942963 | 0.9997444 | Acyl-phosphate:glycerol-3-phospha |

| | | | | | | |
|---|---|---|---|---|---|---|
| AP53_375 | -0.3867498 | 2.887972822 | 0.90240096 | 0.361819 | 0.9997444 | FIG01115781: hypothetical protei |
| AP53_1575 | -0.129228155 | 8.329307005 | 0.900060792 | 0.362419315 | 0.9997444 | Deoxyribose-phosphate aldolase (E |
| AP53_pseudo268 | -0.110244922 | 7.889228161 | 0.891789797 | 0.36455303 | 0.9997444 | Transmembrane histidine kinase Cs |
| AP53_1249 | -0.310310793 | 7.710650513 | 0.889570028 | 0.365128879 | 0.9997444 | Rhodanese-like domain protein |
| AP53_1208 | 0.328597381 | 2.736754577 | 0.886508906 | 0.36592523 | 0.9997444 | FIG01114174: hypothetical protei |
| AP53_1199 | -0.349865638 | 2.890927429 | 0.88562939 | 0.366154518 | 0.9997444 | FIG01115109: hypothetical protei |
| AP53_124 | -0.173438592 | 5.968741084 | 0.878041398 | 0.368141686 | 0.9997444 | V-type ATP synthase subunit F (EC 3 |
| AP53_1518 | -0.114374119 | 6.978776583 | 0.875626221 | 0.368777583 | 0.9997444 | Heme ABC transporter (Streptococ |
| AP53_1375 | 0.090100697 | 7.123257277 | 0.872745672 | 0.369538173 | 0.9997444 | Sensor histidine kinase VraS |
| AP53_103 | -0.109389196 | 6.880115778 | 0.871728568 | 0.369807297 | 0.9997444 | putative collagen binding protein |
| AP53_844 | -0.129507306 | 7.79716985 | 0.869270897 | 0.370458814 | 0.9997444 | ABC transporter, periplasmic sperm |
| AP53_1420 | -0.261147825 | 7.217537269 | 0.869038384 | 0.370520541 | 0.9997444 | Transaldolase (EC 2.2.1.2) |
| AP53_1437 | -0.13964345 | 7.885640041 | 0.869018151 | 0.370525913 | 0.9997444 | Transcriptional regulator, TetR fam |
| AP53_1513 | -0.104365371 | 7.803939915 | 0.86666047 | 0.371152723 | 0.9997444 | Duplicated ATPase component BL0 |
| AP53_493 | -0.085053362 | 8.217555346 | 0.860095932 | 0.372906418 | 0.9997444 | Phosphoenolpyruvate carboxylase |
| AP53_1207 | 0.324552816 | 3.359093794 | 0.852837353 | 0.37486015 | 0.9997444 | hypothetical protein |
| AP53_453 | 0.676383215 | 1.390259302 | 0.852221225 | 0.375026702 | 0.9997444 | FIG01115104: hypothetical protei |
| AP53_rna73 | 0.22198985 | 5.936613833 | 0.85150138 | 0.375221432 | 0.9997444 | tRNA-Ser-GGA |
| AP53_1677 | -0.118332093 | 7.899458382 | 0.851302153 | 0.375275353 | 0.9997444 | Para-aminobenzoate synthase, ami |
| AP53_279 | 0.130586762 | 10.25874936 | 0.848492183 | 0.376037133 | 0.9997444 | Inner membrane protein translocas |
| AP53_1536 | 0.126833376 | 11.00225356 | 0.83912935 | 0.378592401 | 0.9997444 | Alkaline shock protein |
| AP53_1350 | -0.193395791 | 3.748987503 | 0.838327088 | 0.378812579 | 0.9997444 | FIG01116817: hypothetical protei |
| AP53_rna33 | -0.51267931 | 4.405899646 | 0.834810333 | 0.37978004 | 0.9997444 | tRNA-Ile-GAT |
| AP53_791 | 0.09411235 | 8.265255128 | 0.832981908 | 0.380284528 | 0.9997444 | Tributyrin esterase |
| AP53_1125 | 0.151934265 | 7.154776504 | 0.832501567 | 0.380417231 | 0.9997444 | Uridine kinase (EC 2.7.1.48) [C1] |
| AP53_434 | -0.095647354 | 8.537490947 | 0.82678995 | 0.38200059 | 0.9997444 | Two-component sensor kinase SA1 |
| AP53_989 | 0.185040606 | 8.859544528 | 0.826348366 | 0.382123423 | 0.9997444 | ABC transporter membrane-spanni |
| AP53_1017 | -0.337664917 | 6.274401588 | 0.82571138 | 0.382300717 | 0.9997444 | Phage major tail protein |
| AP53_846 | 0.126200674 | 5.626682958 | 0.82287298 | 0.383092267 | 0.9997444 | Two-component sensor histidine ki |
| AP53_167 | -0.19165347 | 6.615908403 | 0.822277079 | 0.383258764 | 0.9997444 | FIG01116967: hypothetical protei |
| AP53_355 | -0.082592224 | 7.784997873 | 0.820580436 | 0.383733423 | 0.9997444 | Manganese ABC transporter, ATP-bi |
| AP53_1674 | -0.174861914 | 7.214747101 | 0.820217115 | 0.383835183 | 0.9997444 | Ribosomal protein L11 methyltrans |
| AP53_604 | 0.335954775 | 6.321387552 | 0.819576211 | 0.384014792 | 0.9997444 | Phage protein |
| AP53_1616 | -0.322653759 | 4.775695083 | 0.813145611 | 0.385824064 | 0.9997444 | Lantibiotic salivaricin A |
| AP53_1203 | -0.190015686 | 3.052125253 | 0.811654808 | 0.386245373 | 0.9997444 | Phage protein |

| | | | | | | |
|---|---|---|---|---|---|---|
| AP53_1227 | 0.076133104 | 7.515049606 | 0.807702586 | 0.387365716 | 0.9997444 | Methylenetetrahydrofolate dehydr |
| AP53_200 | -0.268687892 | 10.29106243 | 0.804108747 | 0.388388807 | 0.9997444 | LSU ribosomal protein L34p |
| AP53_1656 | -0.11637629 | 9.178346442 | 0.801978498 | 0.388997208 | 0.9997444 | Intramembrane protease RasP/YluC |
| AP53_475 | -0.10080638 | 9.041570762 | 0.798291575 | 0.39005367 | 0.9997444 | General stress protein |
| AP53_494 | -0.120592353 | 7.071482449 | 0.797725126 | 0.390216373 | 0.9997444 | Cell division protein FtsW |
| AP53_751 | -0.100782146 | 9.44213199 | 0.797464841 | 0.390291171 | 0.9997444 | Cystine ABC transporter, periplasm |
| AP53_207 | -0.113354247 | 7.12594599 | 0.79684528 | 0.390469303 | 0.9997444 | N-acetylmannosamine kinase (EC 2. |
| AP53_1859 | -0.117975395 | 8.358397793 | 0.791071124 | 0.392135496 | 0.9997444 | ABC transporter ATP-binding protei |
| AP53_1255 | -0.133877831 | 8.539361175 | 0.790903786 | 0.392183947 | 0.9997444 | Hypothetical DUF1027 domain pro |
| AP53_823 | -0.158856891 | 5.578408491 | 0.785637794 | 0.393713392 | 0.9997444 | Two-component sensor kinase, asso |
| AP53_376 | -0.177097204 | 3.814968251 | 0.784721701 | 0.393980403 | 0.9997444 | Competence-stimulating peptide A |
| AP53_335 | -0.20532635 | 8.142871266 | 0.779332576 | 0.395556847 | 0.9997444 | Ribonucleotide reductase of class Il |
| AP53_1483 | -0.187572906 | 9.419940924 | 0.77593818 | 0.396554814 | 0.9997444 | 3-oxoacyl-[acyl-carrier protein] red |
| AP53_573 | 0.163529538 | 6.896267721 | 0.77372722 | 0.39720695 | 0.9997444 | Phenylalanyl-tRNA synthetase alpha |
| AP53_1776 | 0.102251147 | 10.17093042 | 0.771185436 | 0.397958725 | 0.9997444 | RecA protein |
| AP53_441 | -0.197057114 | 4.005282735 | 0.768488842 | 0.398758711 | 0.9997444 | putative L-glutamate ligase |
| AP53_497 | 0.14505683 | 10.59102922 | 0.76432625 | 0.399998533 | 0.9997444 | Triosephosphate isomerase (EC 5.3. |
| AP53_1418 | -0.175966475 | 7.937633942 | 0.763199737 | 0.400335097 | 0.9997444 | FIG01113961: hypothetical protei |
| AP53_507 | 0.154956679 | 4.440330087 | 0.761693622 | 0.400785763 | 0.9997444 | Heparinase II/III-like |
| AP53_1306 | -0.137502486 | 6.047640193 | 0.761283555 | 0.400908602 | 0.9997444 | RNA methyltransferase, TrmA family |
| AP53_898 | 0.295510212 | 6.544989312 | 0.760870686 | 0.40103234 | 0.9997444 | Rossmann fold nucleotide-binding |
| AP53_1171 | 0.163196108 | 6.530184668 | 0.758190827 | 0.401836952 | 0.9997444 | hypothetical protein |
| AP53_900 | 0.115779419 | 5.716788735 | 0.757632204 | 0.402004993 | 0.9997444 | Mobile element protein |
| AP53_146 | 0.164240974 | 6.035456131 | 0.757610098 | 0.402011645 | 0.9997444 | L-ribulose-5-phosphate 4-epimeras |
| AP53_942 | -0.243444063 | 9.17685393 | 0.755271409 | 0.402716365 | 0.9997444 | Protein:protein lipoyl transferase |
| AP53_1757 | -0.239434879 | 8.184491381 | 0.755243549 | 0.402724772 | 0.9997444 | Trehalose-6-phosphate hydrolase (E |
| AP53_581 | -0.085880682 | 7.651479409 | 0.754126234 | 0.403062147 | 0.9997444 | ATP-dependent nuclease, subunit A |
| AP53_300 | -0.289837789 | 8.145165313 | 0.753694317 | 0.403192684 | 0.9997444 | Pyruvate formate-lyase activating e |
| AP53_1281 | 0.13726885 | 9.007721286 | 0.751343855 | 0.403904217 | 0.9997444 | hypothetical protein |
| AP53_1855 | 0.159526216 | 11.18573207 | 0.750407668 | 0.404188167 | 0.9997444 | Inosine-5'-monophosphate dehydr |
| AP53_367 | 0.14025664 | 10.43533034 | 0.749074828 | 0.404592965 | 0.9997444 | Group B streptococcal surface imm |
| AP53_1582 | -0.12510623 | 8.371673484 | 0.741380993 | 0.406942148 | 0.9997444 | Inactive homolog of metal-depende |
| AP53_706 | 0.208515231 | 8.158050813 | 0.738495424 | 0.407828739 | 0.9997444 | Transcriptional repressor of the fru |
| AP53_71 | 0.263805648 | 5.566105766 | 0.730923235 | 0.410169813 | 0.9997444 | hypothetical protein |
| AP53_679 | 0.117070788 | 7.309242498 | 0.73090946 | 0.410174091 | 0.9997444 | Capsule biosynthesis protein capA |

| | | | | | | |
|---|---|---|---|---|---|---|
| AP53_640 | -0.157061581 | 4.209067475 | 0.728530094 | 0.410914105 | 0.9997444 | hypothetical phage protein |
| AP53_1351 | 0.248747637 | 3.956639772 | 0.72675727 | 0.411466849 | 0.9997444 | hypothetical phage protein |
| AP53_1314 | -0.199723554 | 4.001388929 | 0.722721195 | 0.412729639 | 0.9997444 | Phage protein |
| AP53_1301 | -0.15702943 | 8.23204417 | 0.72181365 | 0.413014433 | 0.9997444 | Hyaluronoglucosaminidase (EC 3.2. |
| AP53_170 | 0.211788056 | 9.354608329 | 0.719961225 | 0.413596703 | 0.9997444 | Substrate-specific component BioY |
| AP53_1392 | 0.1037836 | 8.160192567 | 0.717469557 | 0.414381961 | 0.9997444 | S-ribosylhomocysteine lyase (EC 4.4 |
| AP53_1491 | 0.202784209 | 14.69533013 | 0.714128805 | 0.415438529 | 0.9997444 | Chaperone protein DnaK |
| AP53_963 | 0.122199902 | 7.67750797 | 0.712700828 | 0.415891456 | 0.9997444 | Phosphate ABC transporter, peripla |
| AP53_637 | 0.154549609 | 4.800871695 | 0.712033818 | 0.416103288 | 0.9997444 | hypothetical phage protein |
| AP53_743 | -0.19232132 | 9.690537483 | 0.709349408 | 0.416957552 | 0.9997444 | Purine nucleoside phosphorylase (E |
| AP53_1175 | 0.087046199 | 8.466828996 | 0.706161433 | 0.417975699 | 0.9997444 | Dihydroorotate dehydrogenase, cat |
| AP53_1326 | -0.375286918 | 4.142163145 | 0.704026838 | 0.418659643 | 0.9997444 | Phage minor capsid protein |
| AP53_rna15 | 0.345280083 | 3.977905366 | 0.700520055 | 0.419787131 | 0.9997444 | tRNA-Ala-TGC |
| AP53_527 | -0.103894208 | 8.774876702 | 0.699584048 | 0.420088893 | 0.9997444 | Asparaginyl-tRNA synthetase (EC 6.1 |
| AP53_1061 | 0.163150382 | 5.182023961 | 0.699036518 | 0.420265573 | 0.9997444 | Phage integrase: site-specific recom |
| AP53_980 | -0.311925119 | 11.32503678 | 0.697852556 | 0.420648028 | 0.9997444 | FIG01114232: hypothetical protein |
| AP53_831 | 0.29159249 | 9.968418741 | 0.692555835 | 0.422365862 | 0.9997444 | LSU ribosomal protein L10p (P0) |
| AP53_1283 | -0.10350295 | 8.006384195 | 0.684407485 | 0.425030589 | 0.9997444 | FIG01114159: hypothetical protein |
| AP53_523 | -0.235530806 | 9.522155055 | 0.681612666 | 0.425950796 | 0.9997444 | 2,3-butanediol dehydrogenase, S-al |
| AP53_262 | -0.111186312 | 7.796758407 | 0.672009167 | 0.429137393 | 0.9997444 | Potassium uptake protein TrkH |
| AP53_1056 | -0.104333809 | 7.167086984 | 0.668953354 | 0.430159443 | 0.9997444 | hypothetical protein |
| AP53_513 | 0.113504807 | 5.840755779 | 0.667508219 | 0.430644157 | 0.9997444 | 2-deoxy-D-gluconate 3-dehydrogen |
| AP53_1709 | 0.211574202 | 6.130877228 | 0.664470344 | 0.431665984 | 0.9997444 | Strepotococcal cysteine protease (S |
| AP53_770 | -0.108855808 | 8.138427926 | 0.662595498 | 0.432298575 | 0.9997444 | Ribonuclease Z (EC 3.1.26.11) |
| AP53_724 | 0.118029385 | 7.381698237 | 0.658908598 | 0.433546971 | 0.9997444 | Regulation of D-alanyl-lipoteichoic |
| AP53_1285 | -0.171167596 | 8.215176469 | 0.65584972 | 0.434587172 | 0.9997444 | 2-keto-3-deoxy-D-arabino-heptulos |
| AP53_rna47 | 0.365898919 | 4.404252428 | 0.652802738 | 0.435627374 | 0.9997444 | tRNA-Trp-CCA |
| AP53_1083 | -0.288783514 | 4.977482168 | 0.652395563 | 0.435766686 | 0.9997444 | Putative uncharacterized protein Sl |
| AP53_243 | 0.253915756 | 4.137115508 | 0.650739554 | 0.436334026 | 0.9997444 | Transposase |
| AP53_86 | 0.273071104 | 3.208787068 | 0.649120935 | 0.436889723 | 0.9997444 | Late competence protein ComGD, a |
| AP53_1344 | -0.326718855 | 4.348525858 | 0.647807781 | 0.437341399 | 0.9997444 | FIG01115146: hypothetical protein |
| AP53_996 | 0.132115034 | 8.831870744 | 0.647189437 | 0.43755435 | 0.9997444 | Transcriptional regulator, GntR fam |
| AP53_1572 | 0.106847279 | 6.581863799 | 0.647000742 | 0.437619368 | 0.9997444 | DNA polymerase III epsilon subunit |
| AP53_398 | 0.316650401 | 2.694450071 | 0.643189252 | 0.438936081 | 0.9997444 | unknown |
| AP53_887 | 0.1467648 | 10.0984361 | 0.642877983 | 0.439043898 | 0.9997444 | L-lactate dehydrogenase (EC 1.1.1.2 |

| | | | | | | |
|---|---|---|---|---|---|---|
| AP53_1101 | 0.104637923 | 6.992606094 | 0.639804571 | 0.440110796 | 0.9997444 | Ribosomal small subunit pseudouri |
| AP53_1573 | 0.118594176 | 8.771126769 | 0.639680158 | 0.440154074 | 0.9997444 | Transcriptional regulator OrfX |
| AP53_1848 | 0.079056222 | 8.29535345 | 0.639623855 | 0.440173662 | 0.9997444 | Hyaluronan synthase (EC 2.4.1.212 |
| AP53_1675 | -0.129796746 | 5.85093154 | 0.63932502 | 0.44027765 | 0.9997444 | FIG01115320: hypothetical protei |
| AP53_1428 | -0.12510013 | 7.56840856 | 0.630769241 | 0.443272084 | 0.9997444 | FIG01114118: hypothetical protei |
| AP53_949 | -0.209776466 | 7.838170168 | 0.630593322 | 0.443334005 | 0.9997444 | Putative deoxyribose-specific ABC t |
| AP53_1758 | -0.251983588 | 9.430769949 | 0.630525495 | 0.443357883 | 0.9997444 | PTS system, trehalose-specific IIB co |
| AP53_pseudo388 | 0.122091838 | 6.228121309 | 0.628329273 | 0.444132194 | 0.9997444 | Mobile element protein |
| AP53_244 | 0.197265445 | 3.470264022 | 0.626529691 | 0.444768327 | 0.9997444 | Competence-specific sigma factor C |
| AP53_869 | -0.101703866 | 8.672479792 | 0.619444904 | 0.447287417 | 0.9997444 | L-proline glycine betaine ABC trans |
| AP53_1322 | -0.200620485 | 2.744344257 | 0.619199258 | 0.447375183 | 0.9997444 | Phage protein |
| AP53_786 | 0.231180918 | 3.925446239 | 0.617357747 | 0.448034039 | 0.9997444 | hypothetical protein |
| AP53_1588 | 0.157202637 | 10.15300196 | 0.616282669 | 0.448419425 | 0.9997444 | Phosphoglycerate kinase (EC 2.7.2. |
| AP53_891 | 0.12508123 | 11.92681912 | 0.61254334 | 0.449764167 | 0.9997444 | FIG01114048: hypothetical protei |
| AP53_1282 | -0.28747807 | 8.051671386 | 0.611727701 | 0.450058378 | 0.9997444 | hypothetical protein |
| AP53_190 | -0.0883549 | 8.865796182 | 0.610815237 | 0.450387895 | 0.9997444 | Carbonic anhydrase (EC 4.2.1.1) |
| AP53_100 | -0.108011873 | 9.169923012 | 0.610307441 | 0.450571448 | 0.9997444 | tRNA dihydrouridine synthase B (EC |
| AP53_588 | 0.10363105 | 8.92811689 | 0.608636559 | 0.451176303 | 0.9997444 | dTDP-4-dehydrorhamnose reducta |
| AP53_1413 | -0.273530809 | 6.936292989 | 0.60822709 | 0.451324735 | 0.9997444 | FIG01115263: hypothetical protei |
| AP53_924 | 0.119330986 | 8.901812604 | 0.605659295 | 0.452257422 | 0.9997444 | Transcriptional regulator in cluster |
| AP53_1608 | -0.093622188 | 7.521305155 | 0.605130741 | 0.452449805 | 0.9997444 | Type I restriction-modification syst |
| AP53_rna42 | 0.540778132 | 3.837079763 | 0.604887542 | 0.45253837 | 0.9997444 | tRNA-Glu-TTC |
| AP53_535 | -0.166816237 | 10.82592105 | 0.602027937 | 0.453581922 | 0.9997444 | LSU ribosomal protein L31p @ LSU |
| AP53_632 | -0.162938781 | 4.285425534 | 0.600586853 | 0.454109339 | 0.9997444 | hypothetical protein |
| AP53_413 | -0.100871403 | 9.859277469 | 0.598289005 | 0.454952445 | 0.9997444 | Glutaminyl-peptide cyclotransfera |
| AP53_926 | 0.114497801 | 8.964821659 | 0.598191274 | 0.454988362 | 0.9997444 | GMP synthase [glutamine-hydrolyzi |
| AP53_rna8 | -0.329671223 | 4.747622072 | 0.597985718 | 0.45506392 | 0.9997444 | tRNA-Leu-TAG |
| AP53_1555 | -0.090469879 | 8.924918181 | 0.59571993 | 0.455898175 | 0.9997444 | Putative signal peptidase I (EC 3.4.2 |
| AP53_pseudo1695 | 0.198010177 | 11.9771486 | 0.594632298 | 0.456299548 | 0.9997444 | Antiphagocytic M protein/pseudog |
| AP53_933 | 0.176882154 | 9.329273809 | 0.59425963 | 0.456437211 | 0.9997444 | Anaerobic ribonucleoside-triphosp |
| AP53_799 | 0.131685063 | 7.171421152 | 0.592798712 | 0.456977547 | 0.9997444 | Lipoate-protein ligase A |
| AP53_690 | -0.116958958 | 8.924203573 | 0.586440099 | 0.459341918 | 0.9997444 | Carbamoyl-phosphate synthase larg |
| AP53_1602 | -0.180768768 | 5.230470574 | 0.586152171 | 0.459449468 | 0.9997444 | hypothetical protein |
| AP53_520 | -0.112333741 | 8.789098109 | 0.585578514 | 0.459663872 | 0.9997444 | Cell division transporter, ATP-bindi |
| AP53_63 | -0.125326234 | 10.43719894 | 0.584494623 | 0.460069437 | 0.9997444 | Preprotein translocase secY subuni |

| | | | | | | |
|---|---|---|---|---|---|---|
| AP53_2 | -0.102375078 | 8.274963851 | 0.581232272 | 0.461293777 | 0.9997444 | DNA polymerase III beta subunit (EC |
| AP53_1742 | -0.240006626 | 10.55339526 | 0.577686641 | 0.462630687 | 0.9997444 | Alkyl hydroperoxide reductase prot |
| AP53_1844 | -0.086834414 | 8.19988329 | 0.57557603 | 0.463429627 | 0.9997444 | CDP-diacylglycerol--glycerol-3-pho |
| AP53_1670 | -0.157184805 | 6.467137102 | 0.574217208 | 0.463945227 | 0.9997444 | hypothetical protein |
| AP53_1706 | 0.129775254 | 12.43279509 | 0.573753201 | 0.464121515 | 0.9997444 | FIG01114845: hypothetical protein |
| AP53_1607 | -0.115119262 | 5.26947702 | 0.573512645 | 0.464212954 | 0.9997444 | Glycerate kinase (EC 2.7.1.31) |
| AP53_1528 | 0.09344548 | 5.725136531 | 0.570250809 | 0.465455844 | 0.9997444 | Mobile element protein |
| AP53_696 | 0.150347718 | 9.102386373 | 0.565476732 | 0.467285188 | 0.9997444 | KH domain RNA binding protein Ylc |
| AP53_1719 | 0.127995547 | 6.55302028 | 0.565151054 | 0.467410429 | 0.9997444 | PTS system, cellobiose-specific IIC c |
| AP53_830 | 0.129453745 | 7.795508083 | 0.56059987 | 0.469166608 | 0.9997444 | GTPase and tRNA-U34 5-formylatio |
| AP53_1565 | 0.23617365 | 7.738902507 | 0.560476033 | 0.469214551 | 0.9997444 | transporter |
| AP53_237 | -0.092704883 | 6.217745756 | 0.559961058 | 0.469414011 | 0.9997444 | D-alanyl-D-alanine carboxypeptidas |
| AP53_314 | 0.127335281 | 9.769757424 | 0.558741962 | 0.469886767 | 0.9997444 | DNA polymerase III delta prime sub |
| AP53_633 | 0.162133088 | 4.286017489 | 0.557270183 | 0.4704586 | 0.9997444 | Single-stranded DNA-binding prote |
| AP53_488 | -0.234710659 | 7.678716534 | 0.555098903 | 0.47130439 | 0.9997444 | Phage lysin, glycosyl hydrolase, fam |
| AP53_605 | 0.26559284 | 6.486805845 | 0.552897939 | 0.472164406 | 0.9997444 | FIG01116522: hypothetical protein |
| AP53_1431 | -0.078700641 | 9.994209097 | 0.551539471 | 0.472696567 | 0.9997444 | INTEGRAL MEMBRANE PROTEIN (Rh |
| AP53_506 | 0.096858448 | 7.599458418 | 0.551149118 | 0.472849673 | 0.9997444 | Transcriptional regulator RegR, rpr |
| AP53_907 | -0.143059806 | 4.789085088 | 0.550612311 | 0.473060361 | 0.9997444 | Transcriptional regulator, GntR fam |
| AP53_1111 | -0.089258774 | 8.445947905 | 0.5491309 | 0.473642629 | 0.9997444 | Inner membrane protein YihY, form |
| AP53_1270 | -0.16452373 | 8.361294977 | 0.549036592 | 0.473679738 | 0.9997444 | Putative uncharacterized protein gl |
| AP53_787 | 0.098239034 | 5.790644875 | 0.546503647 | 0.474678307 | 0.9997444 | ABC transporter substrate-binding |
| AP53_1526 | -0.134143873 | 8.884422822 | 0.544917956 | 0.475305286 | 0.9997444 | Holo-[acyl-carrier protein] synthase |
| AP53_1463 | -0.092780685 | 7.910418677 | 0.543827948 | 0.475737102 | 0.9997444 | ABC transporter, ATP-binding prote |
| AP53_525 | -0.087654723 | 6.856350395 | 0.543275541 | 0.475956201 | 0.9997444 | DinG family ATP-dependent helicas |
| AP53_730 | 0.119772594 | 7.593321494 | 0.54319643 | 0.475987593 | 0.9997444 | Hydroxymethylglutaryl-CoA reduct |
| AP53_104 | -0.087203494 | 7.441903889 | 0.542106872 | 0.476420301 | 0.9997444 | Signal peptidase I (EC 3.4.21.89) |
| AP53_631 | 0.164423168 | 4.708619 | 0.542047509 | 0.476443896 | 0.9997444 | putaive immunity repressor protein |
| AP53_650 | 0.165477536 | 4.173284571 | 0.539743076 | 0.477361402 | 0.9997444 | conserved domain protein |
| AP53_862 | -0.149368072 | 5.314573916 | 0.538724015 | 0.477768114 | 0.9997444 | Adenylate cyclase (EC 4.6.1.1) |
| AP53_537 | -0.152344632 | 8.498753398 | 0.538419848 | 0.477889625 | 0.9997444 | Flavodoxin |
| AP53_1861 | 0.085534912 | 7.569565934 | 0.53831019 | 0.477933445 | 0.9997444 | Mobile element protein |
| AP53_1569 | 0.126922592 | 7.211031447 | 0.537897349 | 0.478098482 | 0.9997444 | putative repressor - phage associate |
| AP53_499 | 0.081767546 | 7.651709629 | 0.537675162 | 0.478187343 | 0.9997444 | tRNA-dependent lipid II-Ala--L-alani |
| AP53_919 | 0.096638511 | 9.876042395 | 0.537247891 | 0.478358307 | 0.9997444 | FIG01114660: hypothetical protein |

| | | | | | | |
|---|---|---|---|---|---|---|
| AP53_rna12 | 0.446571805 | 3.70589102 | 0.535719522 | 0.478970723 | 0.9997444 | tRNA-Arg-ACG |
| AP53_853 | 0.130640301 | 6.647686072 | 0.535448069 | 0.479079636 | 0.9997444 | Lipase/Acylhydrolase family protei |
| AP53_583 | -0.271185067 | 11.23882585 | 0.535272848 | 0.479149961 | 0.9997444 | SSU ribosomal protein S21p |
| AP53_1667 | 0.101060817 | 10.53558133 | 0.53382428 | 0.479732035 | 0.9997444 | GTP pyrophosphokinase (EC 2.7.6.5 |
| AP53_217 | -0.098591453 | 8.032206253 | 0.533365585 | 0.479916607 | 0.9997444 | PurR: transcription regulator assoc |
| AP53_454 | 0.103364513 | 9.919725063 | 0.533220303 | 0.479975092 | 0.9997444 | Plasmid stabilization system antito |
| AP53_59 | -0.152854943 | 11.0043086 | 0.531557695 | 0.480645277 | 0.9997444 | LSU ribosomal protein L18p (L5e) |
| AP53_1008 | -0.221352916 | 5.425898908 | 0.53127879 | 0.480757862 | 0.9997444 | Phage protein |
| AP53_606 | 0.185696407 | 5.885791463 | 0.529522608 | 0.481467825 | 0.9997444 | Phage hyaluronidase |
| AP53_1434 | -0.064377171 | 7.360167075 | 0.528703198 | 0.481799709 | 0.9997444 | Sodium-dependent phosphate tran |
| AP53_1192 | 0.290068824 | 2.102695241 | 0.528518779 | 0.481874459 | 0.9997444 | conserved hypothetical protein - pl |
| AP53_734 | 0.231534801 | 8.466451951 | 0.528005655 | 0.482082548 | 0.9997444 | FIG01114189: hypothetical protei |
| AP53_659 | 0.108229993 | 7.44072087 | 0.526326822 | 0.482764466 | 0.9997444 | Heteropolysaccharide repeat unit e |
| AP53_266 | -0.109019245 | 9.303906861 | 0.525238994 | 0.483207225 | 0.9997444 | COG1399 protein in cluster with ri |
| AP53_332 | -0.160305995 | 4.479779567 | 0.524693816 | 0.483429386 | 0.9997444 | FIG01115797: hypothetical protei |
| AP53_336 | -0.095429708 | 10.01169484 | 0.523259074 | 0.484014899 | 0.9997444 | C3 family ADP-ribosyltransferase (E |
| AP53_1328 | -0.209832305 | 3.983716311 | 0.522628349 | 0.484272688 | 0.9997444 | Phage major capsid protein |
| AP53_1741 | -0.278335473 | 10.54648531 | 0.519053884 | 0.485738189 | 0.9997444 | Cold shock protein CspA |
| AP53_1330 | 0.23385672 | 3.811752405 | 0.5187946 | 0.485844796 | 0.9997444 | Phage protein |
| AP53_1243 | -0.170233233 | 9.14538567 | 0.518015737 | 0.486165277 | 0.9997444 | Cell division protein FtsA |
| AP53_142 | 0.145471429 | 4.788570738 | 0.51680684 | 0.486663442 | 0.9997444 | Ascorbate-specific PTS system, EIIB |
| AP53_274 | 0.092153892 | 10.45166169 | 0.515124251 | 0.4873583 | 0.9997444 | FIG01114242: hypothetical protei |
| AP53_278 | 0.075416669 | 9.52934866 | 0.513868063 | 0.487878205 | 0.9997444 | Transcription elongation factor Gre |
| AP53_1378 | 0.084951843 | 10.25019188 | 0.512567648 | 0.488417442 | 0.9997444 | Protein serine/threonine phosphat |
| AP53_851 | -0.248812011 | 6.138214621 | 0.510204005 | 0.489400256 | 0.9997444 | Class B acid phosphatase precursor |
| AP53_1438 | 0.08853379 | 8.510849309 | 0.50804413 | 0.490301394 | 0.9997444 | Hydrolase (HAD superfamily) |
| AP53_407 | 0.092982467 | 7.994504104 | 0.506855588 | 0.490798526 | 0.9997444 | Dephospho-CoA kinase (EC 2.7.1.24 |
| AP53_785 | 0.078678211 | 6.737096775 | 0.505987902 | 0.491162015 | 0.9997444 | Fibronectin/fibrinogen-binding pro |
| AP53_11 | -0.084051885 | 7.488083093 | 0.505005283 | 0.491574227 | 0.9997444 | FIG01114299: hypothetical protei |
| AP53_393 | 0.143406548 | 3.336583387 | 0.504375807 | 0.491838616 | 0.9997444 | Competence-stimulating peptide A |
| AP53_1258 | -0.076878722 | 8.322350417 | 0.503666657 | 0.492136772 | 0.9997444 | Lon-like protease with PDZ domain |
| AP53_430 | 0.117178231 | 7.043124354 | 0.501485734 | 0.493055728 | 0.9997444 | 3-ketoacyl-CoA thiolase (EC 2.3.1.1 |
| AP53_1223 | -0.119491618 | 8.566335184 | 0.500946837 | 0.493283267 | 0.9997444 | RNA binding methyltransferase FtsJ |
| AP53_715 | 0.259755063 | 3.336580757 | 0.50033047 | 0.493543745 | 0.9997444 | Nucleoside diphosphate kinase (EC |
| AP53_1450 | 0.072210244 | 7.186068963 | 0.499552456 | 0.493872884 | 0.9997444 | Copper-translocating P-type ATPase |

| | | | | | | |
|---|---|---|---|---|---|---|
| AP53_rna45 | 0.370018015 | 3.076635878 | 0.497225072 | 0.494859815 | 0.9997444 | tRNA-Phe-GAA |
| AP53_838 | -0.157991104 | 8.42697884 | 0.497149061 | 0.494892107 | 0.9997444 | Dihydroneopterin aldolase (EC 4.1. |
| AP53_101 | -0.124109903 | 8.357710187 | 0.495357349 | 0.495654366 | 0.9997444 | Chaperonin (heat shock protein 33) |
| AP53_1799 | -0.174073476 | 4.830050309 | 0.494790226 | 0.495896077 | 0.9997444 | FIG01115461: hypothetical protei |
| AP53_1466 | -0.118952061 | 9.729073719 | 0.493925982 | 0.496264825 | 0.9997444 | Cell envelope-associated transcript |
| AP53_444 | -0.190215668 | 3.147686211 | 0.4938415 | 0.496300898 | 0.9997444 | FIG01116163: hypothetical protei |
| AP53_1458 | 0.08496975 | 9.111977327 | 0.492649387 | 0.496810408 | 0.9997444 | FIG000325: clustered with transcri |
| AP53_797 | -0.154126632 | 11.03110948 | 0.488537524 | 0.498575008 | 0.9997444 | Dihydrolipoamide dehydrogenase |
| AP53_1494 | 0.077525276 | 9.042313129 | 0.487561457 | 0.498995532 | 0.9997444 | putative N-acetyl-muramidase |
| AP53_528 | 0.075566044 | 8.886217142 | 0.487015077 | 0.49923121 | 0.9997444 | FIG000506: Predicted P-loop-cont |
| AP53_308 | -0.09391402 | 7.965217081 | 0.485941701 | 0.499694784 | 0.9997444 | Membrane protein involved in the |
| AP53_1030 | -0.177578895 | 4.945597315 | 0.485827955 | 0.499743955 | 0.9997444 | Phage endonuclease |
| AP53_1180 | -0.158061298 | 6.123507441 | 0.482320945 | 0.501264242 | 0.9997444 | Streptococcal pyrogenic exotoxin |
| AP53_1109 | -0.076013914 | 6.796823817 | 0.482304103 | 0.501271563 | 0.9997444 | Shikimate kinase I (EC 2.7.1.71) |
| AP53_362 | 0.088527918 | 9.437692988 | 0.480110925 | 0.502226559 | 0.9997444 | Uridylate kinase (EC 2.7.4.-) |
| AP53_826 | -0.245978653 | 8.443236576 | 0.47879118 | 0.502802807 | 0.9997444 | Succinate-semialdehyde dehydroge |
| AP53_874 | 0.108929612 | 7.794643657 | 0.478281272 | 0.50302577 | 0.9997444 | Hypothetical similar to thiamin bic |
| AP53_1137 | 0.11370124 | 5.163380194 | 0.478103636 | 0.503103485 | 0.9997444 | Mobile element protein |
| AP53_1345 | 0.17786093 | 4.364938321 | 0.477933646 | 0.503177876 | 0.9997444 | FIG01114622: hypothetical protei |
| AP53_rna74 | 0.247728022 | 3.94592579 | 0.477507668 | 0.503364378 | 0.9997444 | tRNA-Leu-AAG |
| AP53_246 | 0.106898853 | 8.990207353 | 0.476111253 | 0.503976633 | 0.9997444 | FIG001553: Hydrolase, HAD subfar |
| AP53_386 | 0.251167186 | 3.007733876 | 0.472990467 | 0.505349808 | 0.9997444 | Mobile element protein |
| AP53_967 | 0.201010159 | 9.252986656 | 0.471529602 | 0.505994929 | 0.9997444 | Transcriptional regulator SpxA1 |
| AP53_904 | 0.194185607 | 5.773598337 | 0.469370919 | 0.506950944 | 0.9997444 | Biotin carboxyl carrier protein of o |
| AP53_1346 | 0.1928736 | 4.250518521 | 0.468338979 | 0.507409117 | 0.9997444 | Phage Holliday junction resolvase |
| AP53_317 | -0.203756661 | 9.054935529 | 0.46754527 | 0.50776203 | 0.9997444 | DNA replication intiation control p |
| AP53_1409 | 0.071302246 | 9.200307258 | 0.467130128 | 0.507946794 | 0.9997444 | Phospho-N-acetylmuramoyl-penta |
| AP53_913 | -0.133281697 | 7.533091996 | 0.466354286 | 0.508292422 | 0.9997444 | Citrate lyase gamma chain, acyl car |
| AP53_1070 | 0.132340921 | 6.090033599 | 0.46564381 | 0.508609305 | 0.9997444 | Mobile element protein |
| AP53_657 | 0.108415446 | 7.849295204 | 0.464954153 | 0.508917246 | 0.9997444 | Glycosyl transferase, group 2 family |
| AP53_608 | 0.199299526 | 5.182205261 | 0.464269671 | 0.509223213 | 0.9997444 | Phage endopeptidase |
| AP53_1273 | -0.118428873 | 6.981510129 | 0.463702292 | 0.509477088 | 0.9997444 | Two-component response regulato |
| AP53_175 | -0.239619622 | 5.691972231 | 0.462209751 | 0.510146031 | 0.9997444 | hypothetical protein |
| AP53_1671 | -0.137209669 | 6.337641092 | 0.45987768 | 0.511194457 | 0.9997444 | Exodeoxyribonuclease III (EC 3.1.1 |
| AP53_252 | 0.091624846 | 7.357726558 | 0.458429684 | 0.511847414 | 0.9997444 | FIG145533: Methyltransferase (EC |

| | | | | | | |
|---|---|---|---|---|---|---|
| AP53_463 | -0.124823253 | 5.479056278 | 0.45566025 | 0.513100517 | 0.9997444 | Multidrug resistance protein B |
| AP53_1160 | 0.10943999 | 7.302703675 | 0.452783551 | 0.514408123 | 0.9997444 | tRNA (adenine37-N(6))-methyltrans |
| AP53_1313 | -0.274623314 | 3.823947758 | 0.45149573 | 0.514995488 | 0.9997444 | Phage holin |
| AP53_502 | -0.102711984 | 6.893721111 | 0.450936876 | 0.515250761 | 0.9997444 | FIG01114293: hypothetical protei |
| AP53_865 | 0.094115811 | 7.424202608 | 0.449511819 | 0.515902752 | 0.9997444 | Similar to ribosomal large subunit |
| AP53_1727 | 0.07367276 | 9.545110216 | 0.446137114 | 0.517452826 | 0.9997444 | Multimodular transpeptidase-trans |
| AP53_1788 | 0.175169879 | 7.228241139 | 0.444034649 | 0.518422886 | 0.9997444 | FIG01118151: hypothetical protei |
| AP53_798 | 0.08519492 | 6.426757126 | 0.443069581 | 0.518869288 | 0.9997444 | Hyaluronate lyase precursor (EC 4.2 |
| AP53_198 | 0.099627216 | 10.34227161 | 0.442522025 | 0.519122882 | 0.9997444 | RNA-binding protein Jag |
| AP53_1650 | 0.199080241 | 6.99247406 | 0.442205475 | 0.519269593 | 0.9997444 | FIG01118158: hypothetical protei |
| AP53_425 | 0.142763065 | 7.428724349 | 0.439698201 | 0.520434358 | 0.9997444 | Threonyl-tRNA synthetase (EC 6.1.1 |
| AP53_1146 | 0.133196449 | 9.620017711 | 0.439039981 | 0.520740941 | 0.9997444 | Oligoendopeptidase F |
| AP53_837 | 0.131868005 | 9.548222719 | 0.438958255 | 0.520779031 | 0.9997444 | Dihydropteroate synthase (EC 2.5.1 |
| AP53_1707 | 0.134518893 | 11.49237008 | 0.433698929 | 0.523241149 | 0.9997444 | Peptidylproline cis-trans-isomerase |
| AP53_1837 | -0.0952912 | 8.456360843 | 0.43332989 | 0.523414727 | 0.9997444 | tRNA-specific 2-thiouridylase Mnm |
| AP53_1316 | -0.178545504 | 3.674597187 | 0.432343628 | 0.523879143 | 0.9997444 | Phage hyaluronidase |
| AP53_1422 | -0.127207171 | 6.883149016 | 0.431778334 | 0.524145679 | 0.9997444 | NADH peroxidase (EC 1.11.1.1) |
| AP53_465 | -0.133933046 | 6.15883924 | 0.42718655 | 0.52632014 | 0.9997444 | PTS system, beta-glucoside-specific |
| AP53_199 | 0.103135647 | 6.130953473 | 0.426106993 | 0.526833826 | 0.9997444 | FIG01115631: hypothetical protei |
| AP53_836 | 0.07975021 | 9.379810407 | 0.425021611 | 0.527351234 | 0.9997444 | GTP cyclohydrolase I (EC 3.5.4.16) t |
| AP53_1368 | -0.137675555 | 3.964798628 | 0.424822865 | 0.527446081 | 0.9997444 | Competence protein F homolog, ph |
| AP53_1642 | -0.09990651 | 8.132237723 | 0.424446211 | 0.527625918 | 0.9997444 | FIG01114029: hypothetical protei |
| AP53_21 | 0.160447523 | 6.210596514 | 0.424318044 | 0.527687139 | 0.9997444 | Phosphoribosylaminoimidazole-su |
| AP53_1046 | -0.189273035 | 5.864546982 | 0.42426264 | 0.527713607 | 0.9997444 | hypothetical phage protein |
| AP53_1204 | -0.146872632 | 2.871581077 | 0.419750378 | 0.529877719 | 0.9997444 | Recombinational DNA repair protei |
| AP53_951 | -0.110592021 | 9.597594622 | 0.419722336 | 0.529891221 | 0.9997444 | Cytidine deaminase (EC 3.5.4.5) |
| AP53_1201 | -0.174652526 | 3.887389286 | 0.418107141 | 0.530669991 | 0.9997444 | conserved hypothetical protein - ph |
| AP53_117 | 0.078235225 | 7.004833486 | 0.417267187 | 0.531075833 | 0.9997444 | Endoribonuclease L-PSP |
| AP53_1640 | -0.08343264 | 6.301335828 | 0.4166745 | 0.531362556 | 0.9997444 | FIG01114632: hypothetical protei |
| AP53_505 | -0.100662036 | 9.691599483 | 0.416563091 | 0.531416485 | 0.9997444 | Calcium-transporting ATPase (EC 3. |
| AP53_518 | 0.065654731 | 7.32366515 | 0.415326657 | 0.532015692 | 0.9997444 | Epoxyqueuosine (oQ) reductase Qu |
| AP53_553 | -0.177672236 | 7.009642797 | 0.414125242 | 0.532599156 | 0.9997444 | Streptolysin S export transmembra |
| AP53_76 | 0.082815288 | 7.413343077 | 0.41381262 | 0.532751179 | 0.9997444 | Zinc ABC transporter, inner membr |
| AP53_910 | -0.161554436 | 7.052291359 | 0.413603166 | 0.53285308 | 0.9997444 | Biotin carboxyl carrier protein of o |
| AP53_435 | -0.106637673 | 7.890446052 | 0.412853477 | 0.53321811 | 0.9997444 | Zn-dependent hydrolase YycJ/WalJ, |

| | | | | | | |
|---|---|---|---|---|---|---|
| AP53_rna13 | 0.262690356 | 3.975155994 | 0.411584329 | 0.533837154 | 0.9997444 | tRNA-Pro-TGG |
| AP53_1 | -0.092913485 | 7.944552078 | 0.41025721 | 0.534485938 | 0.9997444 | Chromosomal replication initiator |
| AP53_130 | -0.077494191 | 7.733056354 | 0.4096847 | 0.534766282 | 0.9997444 | FIG013576: hypothetical protein |
| AP53_60 | -0.130734456 | 10.31908623 | 0.408569154 | 0.535313344 | 0.9997444 | SSU ribosomal protein S5p (S2e) |
| AP53_587 | 0.072238094 | 9.35155574 | 0.407418449 | 0.535878767 | 0.9997444 | probably aromatic ring hydroxylati |
| AP53_197 | 0.084298745 | 9.919655201 | 0.406476092 | 0.536342664 | 0.9997444 | Inner membrane protein translocas |
| AP53_901 | 0.072691631 | 7.690586902 | 0.405894172 | 0.53662951 | 0.9997444 | tRNA:m(5)U-54 MTase gid |
| AP53_422 | 0.087332506 | 10.12176022 | 0.405675596 | 0.536737329 | 0.9997444 | Glycosyltransferase LafB, responsib |
| AP53_1532 | 0.168309929 | 11.03730632 | 0.404084275 | 0.537523544 | 0.9997444 | PTS system, sucrose-specific IIB con |
| AP53_1041 | -0.124654088 | 5.849702813 | 0.402919359 | 0.538100487 | 0.9997444 | FIG01115822: hypothetical protei |
| AP53_1687 | -0.071861491 | 6.531062586 | 0.402704318 | 0.538207119 | 0.9997444 | Streptococcal histidine triad protei |
| AP53_651 | -0.208023287 | 3.86697987 | 0.402120324 | 0.538496907 | 0.9997444 | hypothetical phage protein |
| AP53_rna62 | 1.160213364 | 0.675720235 | 0.399861375 | 0.539620667 | 0.9997444 | tRNA-Glu-TTC |
| AP53_810 | -0.097662333 | 7.328800618 | 0.399693958 | 0.539704131 | 0.9997444 | Translation elongation factor LepA |
| AP53_1765 | 0.100984149 | 10.01501353 | 0.399620737 | 0.539740643 | 0.9997444 | Ribonucleotide reductase of class II |
| AP53_211 | 0.078942692 | 8.865840499 | 0.399050151 | 0.540025327 | 0.9997444 | SSU rRNA (adenine(1518)-N(6)/ader |
| AP53_1538 | 0.110638381 | 8.55751471 | 0.398221024 | 0.540439521 | 0.9997444 | dCMP deaminase (EC 3.5.4.12); Lat |
| AP53_220 | -0.132756172 | 11.41052102 | 0.397905925 | 0.54059709 | 0.9997444 | SSU ribosomal protein S7p (S5e) |
| AP53_242 | 0.088998736 | 9.151222099 | 0.396988635 | 0.541056298 | 0.9997444 | Oligopeptide transport ATP-binding |
| AP53_827 | -0.131612689 | 8.427106352 | 0.395870953 | 0.541616842 | 0.9997444 | Excinuclease ABC subunit C |
| AP53_881 | 0.093343528 | 9.178826799 | 0.395100532 | 0.542003879 | 0.9997444 | Serine hydroxymethyltransferase (E |
| AP53_1191 | 0.170306991 | 2.921683598 | 0.394589058 | 0.542261123 | 0.9997444 | Phage protein |
| AP53_1252 | 0.08786102 | 12.92724852 | 0.394027478 | 0.542543841 | 0.9997444 | Non-specific DNA-binding protein D |
| AP53_172 | -0.117496831 | 8.017024394 | 0.393018382 | 0.543052569 | 0.9997444 | tRNA-specific adenosine-34 deamir |
| AP53_1785 | -0.204323003 | 5.049146662 | 0.392175843 | 0.543478036 | 0.9997444 | conserved hypothetical protein - pl |
| AP53_1198 | -0.140430705 | 3.798169381 | 0.3920979 | 0.543517428 | 0.9997444 | FIG040428: Phage-associated prote |
| AP53_612 | -0.206967027 | 4.914644896 | 0.391411474 | 0.543864587 | 0.9997444 | Phage base plate |
| AP53_1327 | -0.194356138 | 3.739674086 | 0.39105445 | 0.544045321 | 0.9997444 | Phage protein |
| AP53_1585 | 0.159594455 | 8.253436029 | 0.390675568 | 0.544237248 | 0.9997444 | Glutamine synthetase type I (EC 6.3 |
| AP53_1629 | -0.275269548 | 2.025165839 | 0.390353557 | 0.544400469 | 0.9997444 | Protein of unknown function DUF2 |
| AP53_1298 | -0.114339272 | 7.18348429 | 0.390178281 | 0.544489353 | 0.9997444 | N-acetylmannosamine kinase (EC 2. |
| AP53_1000 | -0.220363999 | 7.176395449 | 0.389513004 | 0.544826977 | 0.9997444 | Paratox |
| AP53_835 | 0.175027281 | 10.22822728 | 0.389387048 | 0.544890945 | 0.9997444 | Dihydrofolate synthase (EC 6.3.2.1 |
| AP53_rna81 | 0.247229171 | 8.333760293 | 0.389243519 | 0.544963855 | 0.9997444 | tRNA-Lys-CTT |
| AP53_549 | -0.139446315 | 6.651482013 | 0.389103911 | 0.545034792 | 0.9997444 | Streptolysin S self-immunity protei |

| | | | | | | |
|---|---|---|---|---|---|---|
| AP53_226 | 0.054451775 | 7.602630791 | 0.388056906 | 0.545567359 | 0.9997444 | FIG01114729: hypothetical protei |
| AP53_334 | -0.153936475 | 7.109344875 | 0.387667765 | 0.545765557 | 0.9997444 | Ribonucleotide reduction protein I |
| AP53_260 | -0.167075944 | 7.925695479 | 0.387432466 | 0.545885468 | 0.9997444 | Dicarboxylate/amino acid:cation (N |
| AP53_784 | 0.103337248 | 7.389123837 | 0.381919736 | 0.548709575 | 0.9997444 | TPR-repeat-containing protein |
| AP53_396 | -0.315628468 | 2.364449789 | 0.381790378 | 0.548776186 | 0.9997444 | FIG01116077: hypothetical protei |
| AP53_1261 | 0.1046579 | 5.821106996 | 0.381552646 | 0.548898644 | 0.9997444 | Aspartate--ammonia ligase (EC 6.3.1 |
| AP53_168 | -0.119519703 | 8.972750008 | 0.380228861 | 0.549581519 | 0.9997444 | tRNA-guanine transglycosylase (EC 2 |
| AP53_3 | 0.173044427 | 5.923167754 | 0.379743825 | 0.549832141 | 0.9997444 | FIG01114317: hypothetical protei |
| AP53_1411 | 0.08804165 | 8.762450511 | 0.379551455 | 0.549931602 | 0.9997444 | Cell division protein FtsL |
| AP53_790 | -0.087366659 | 9.013399893 | 0.379217625 | 0.550104287 | 0.9997444 | Ribonuclease J2 (endoribonuclease |
| AP53_66 | -0.1614865 | 10.13338686 | 0.378610175 | 0.550418783 | 0.9997444 | LSU ribosomal protein L36p |
| AP53_832 | 0.160373075 | 9.839306377 | 0.37747801 | 0.551005883 | 0.9997444 | LSU ribosomal protein L7/L12 (P1/I |
| AP53_1232 | 0.248695155 | 8.0946323 | 0.376469079 | 0.551530117 | 0.9997444 | FIG01114188: hypothetical protei |
| AP53_481 | -0.152971803 | 8.287145628 | 0.375868261 | 0.551842764 | 0.9997444 | bacterial seryl-tRNA synthetase rela |
| AP53_287 | -0.09902868 | 7.802514992 | 0.375469445 | 0.552050489 | 0.9997444 | FIG009886: phosphoesterase |
| AP53_973 | 0.075048813 | 5.879572272 | 0.374405758 | 0.552605266 | 0.9997444 | ABC transporter permease protein |
| AP53_1167 | 0.068496651 | 8.0253434 | 0.374074704 | 0.552778155 | 0.9997444 | D-alanine--D-alanine ligase A (EC 6.3 |
| AP53_733 | 0.134641214 | 7.69791382 | 0.374053519 | 0.552789222 | 0.9997444 | Dihydrofolate reductase (EC 1.5.1.3 |
| AP53_1843 | 0.05724054 | 8.406362663 | 0.373535361 | 0.553060049 | 0.9997444 | ATPase component of general energ |
| AP53_10 | -0.121401124 | 6.92771867 | 0.373248273 | 0.553210214 | 0.9997444 | FIG01114659: hypothetical protei |
| AP53_1215 | 0.125139587 | 6.297480204 | 0.373112799 | 0.553281104 | 0.9997444 | Phage integrase |
| AP53_pseudo928 | 0.08653619 | 5.682119527 | 0.372938488 | 0.553372342 | 0.9997444 | UDP-N-acetylmuramoylpentapepti |
| AP53_625 | 0.195005242 | 3.57983179 | 0.372719425 | 0.553487047 | 0.9997444 | hypothetical phage protein |
| AP53_415 | 0.093698981 | 9.242783866 | 0.371600407 | 0.554073714 | 0.9997444 | FIG001614: Membrane protein |
| AP53_1763 | 0.133990712 | 7.269390876 | 0.369061738 | 0.555409231 | 0.9997444 | Glyoxalase family protein |
| AP53_89 | -0.13213685 | 3.062870992 | 0.368222528 | 0.555852118 | 0.9997444 | Late competence protein ComGG, I |
| AP53_pseudo812 | 0.132276039 | 8.76666172 | 0.367272512 | 0.556354327 | 0.9997444 | Nucleoside diphosphate kinase (EC |
| AP53_1678 | -0.097848436 | 8.21689848 | 0.367169435 | 0.556408871 | 0.9997444 | Anthranilate synthase, amidotransf |
| AP53_9 | 0.13030774 | 7.0436227 | 0.366596257 | 0.556712367 | 0.9997444 | Cell division protein DivIC (FtsB), st |
| AP53_37 | -0.173720182 | 8.621551496 | 0.36645373 | 0.556787886 | 0.9997444 | MORN motif family protein |
| AP53_1668 | 0.093992771 | 13.35226329 | 0.364441972 | 0.557856002 | 0.9997444 | Collagen-like surface protein |
| AP53_692 | -0.152841633 | 8.311700924 | 0.363621131 | 0.558292989 | 0.9997444 | ABC transporter ATP-binding protei |
| AP53_1360 | 0.273857416 | 9.222909666 | 0.362647279 | 0.55881232 | 0.9997444 | FIG01117062: hypothetical protei |
| AP53_872 | 0.121337457 | 7.873853789 | 0.36237654 | 0.558956869 | 0.9997444 | Xanthine phosphoribosyltransferas |
| AP53_1825 | 0.085049166 | 6.740475722 | 0.360757636 | 0.559822775 | 0.9997444 | FIG04612: Integral membrane prot |

| | | | | | | |
|---|---|---|---|---|---|---|
| AP53_442 | 0.11228836 | 3.886485751 | 0.359233992 | 0.560640181 | 0.9997444 | FIG01115625: hypothetical protei |
| AP53_1850 | 0.132518222 | 8.012850345 | 0.358555502 | 0.561004948 | 0.9997444 | FIG01117167: hypothetical protei |
| AP53_1579 | 0.100209495 | 8.787446596 | 0.358351579 | 0.561114673 | 0.9997444 | SSU ribosomal protein S14p (S29e) |
| AP53_590 | 0.071165846 | 7.752761326 | 0.357721094 | 0.561454191 | 0.9997444 | Alpha-L-Rha alpha-1,3-L-rhamnosyl |
| AP53_649 | -0.157469074 | 6.231483278 | 0.357565008 | 0.561538307 | 0.9997444 | hypothetical phage protein |
| AP53_1544 | 0.099580949 | 11.31501403 | 0.357301401 | 0.561680425 | 0.9997444 | Single-stranded DNA-binding prote |
| AP53_257 | 0.112769175 | 8.245360951 | 0.354606877 | 0.563137274 | 0.9997444 | Methionine ABC transporter ATP-bi |
| AP53_1840 | -0.122086472 | 8.984295955 | 0.35454749 | 0.563169469 | 0.9997444 | Immunodominant antigen A |
| AP53_1501 | -0.073960634 | 8.011588379 | 0.353662735 | 0.563649543 | 0.9997444 | Aspartyl-tRNA(Asn) amidotransfera |
| AP53_526 | -0.063008379 | 8.990784056 | 0.353294801 | 0.56384943 | 0.9997444 | Aspartate aminotransferase (EC 2.6. |
| AP53_849 | -0.19777092 | 6.509506297 | 0.351827426 | 0.564648031 | 0.9997444 | Alcohol dehydrogenase, zinc-conta |
| AP53_1159 | 0.076796629 | 8.174832113 | 0.351041973 | 0.565076444 | 0.9997444 | 1-acyl-sn-glycerol-3-phosphate acyl |
| AP53_1189 | 0.155860755 | 1.973860912 | 0.350949418 | 0.565126969 | 0.9997444 | FIG022125: Phage-associated prote |
| AP53_1348 | 0.135998747 | 4.605248262 | 0.350842303 | 0.565185455 | 0.9997444 | Single-stranded DNA-binding prote |
| AP53_955 | 0.096510429 | 6.856014716 | 0.349575231 | 0.565878215 | 0.9997444 | Two component system sensor hist |
| AP53_1681 | -0.082967048 | 8.463565769 | 0.349536841 | 0.565899231 | 0.9997444 | Dipeptide-binding ABC transporter |
| AP53_1034 | -0.110134303 | 6.332954571 | 0.349519725 | 0.565908601 | 0.9997444 | putative methyltransferase |
| AP53_1010 | -0.167612905 | 5.822470884 | 0.349316601 | 0.566019829 | 0.9997444 | Phage hyaluronidase |
| AP53_897 | 0.098456017 | 8.720799367 | 0.348185356 | 0.566640095 | 0.9997444 | Ribonuclease HII (EC 3.1.26.4) |
| AP53_466 | -0.109804809 | 6.612090354 | 0.346817275 | 0.567392062 | 0.9997444 | 6-phospho-beta-glucosidase (EC 3.2 |
| AP53_1464 | -0.066419971 | 8.759238143 | 0.345392584 | 0.568177302 | 0.9997444 | Histidine triad (HIT) nucleotide-bin |
| AP53_162 | 0.084590917 | 5.480491043 | 0.344516118 | 0.568661478 | 0.9997444 | Quinolinate phosphoribosyltransfe |
| AP53_1170 | 0.081900722 | 8.008830292 | 0.342461092 | 0.569800014 | 0.9997444 | Substrate-specific component NiaX |
| AP53_1107 | -0.074624348 | 7.126780371 | 0.342400248 | 0.569833794 | 0.9997444 | 23S rRNA (Uracil-5-) -methyltransfe |
| AP53_685 | 0.073943938 | 8.867203535 | 0.341942027 | 0.570088325 | 0.9997444 | Ribosomal large subunit pseudouri |
| AP53_298 | 0.069360393 | 7.251107526 | 0.340762583 | 0.570744547 | 0.9997444 | SAM-dependent methyltransferase, |
| AP53_328 | 0.141911846 | 8.781683546 | 0.340036745 | 0.571149159 | 0.9997444 | FIG01115281: hypothetical protei |
| AP53_324 | -0.219746614 | 10.21515832 | 0.339305988 | 0.571557106 | 0.9997444 | Lactate 2-monooxygenase (EC 1.13. |
| AP53_1286 | -0.135459046 | 9.381978509 | 0.339270896 | 0.571576711 | 0.9997444 | 3-dehydroquinate synthase (EC 4.2. |
| AP53_1516 | -0.053641719 | 6.984875302 | 0.339015543 | 0.571719413 | 0.9997444 | Putative ABC transporter ATP-bindi |
| AP53_574 | 0.108250088 | 7.228015948 | 0.338800522 | 0.571839633 | 0.9997444 | Phenylalanyl-tRNA synthetase beta |
| AP53_1798 | -0.172406668 | 4.356196758 | 0.338603743 | 0.571949699 | 0.9997444 | Phage protein |
| AP53_1529 | -0.069701581 | 8.227547356 | 0.336670347 | 0.573033434 | 0.9997444 | Mannose-6-phosphate isomerase (E |
| AP53_1713 | -0.234402223 | 10.57102058 | 0.335209841 | 0.573854899 | 0.9997444 | Streptodornase B; Mitogenic factor |
| AP53_1193 | 0.207788793 | 2.503623599 | 0.334612284 | 0.574191695 | 0.9997444 | Phage major tail protein |

| | | | | | | |
|---|---|---|---|---|---|---|
| AP53_1847 | -0.109052517 | 5.744415718 | 0.333668974 | 0.574724194 | 0.9997444 | FIG001621: Zinc protease |
| AP53_591 | 0.07974234 | 6.796622764 | 0.332986951 | 0.575109831 | 0.9997444 | rhamnose-containing polysacharid |
| AP53_1278 | -0.090400976 | 9.295328339 | 0.332041682 | 0.575645199 | 0.9997444 | Oxidoreductase, Gfo/Idh/MocA fam |
| AP53_1456 | 0.068990184 | 9.922923369 | 0.331328016 | 0.576050078 | 0.9997444 | COG2740: Predicted nucleic-acid-b |
| AP53_rna18 | 0.285701116 | 4.238330892 | 0.331325251 | 0.576051647 | 0.9997444 | tRNA-Val-TAC |
| AP53_1801 | -0.123588639 | 3.605031207 | 0.331314669 | 0.576057656 | 0.9997444 | FIG01116340: hypothetical protei |
| AP53_416 | 0.067150535 | 8.589937643 | 0.331044812 | 0.576210909 | 0.9997444 | hypothetical protein |
| AP53_1714 | -0.175932286 | 3.071270221 | 0.328787261 | 0.5774963 | 0.9997444 | FIG01115319: hypothetical protei |
| AP53_1060 | -0.142840536 | 5.333577603 | 0.326727994 | 0.57867399 | 0.9997444 | hypothetical protein |
| AP53_85 | 0.12919731 | 3.168246607 | 0.326532512 | 0.578786045 | 0.9997444 | Late competence protein ComGC, a |
| AP53_496 | -0.100841592 | 12.88864028 | 0.325604762 | 0.57931847 | 0.9997444 | Translation elongation factor Tu |
| AP53_1048 | -0.143503552 | 6.144025653 | 0.325204226 | 0.579548648 | 0.9997444 | hypothetical protein, phage associa |
| AP53_151 | 0.088150595 | 9.943785209 | 0.325110013 | 0.579602817 | 0.9997444 | DNA polymerase I (EC 2.7.7.7) |
| AP53_985 | -0.0724932 | 8.1689722 | 0.324776897 | 0.579794434 | 0.9997444 | Cobalt-zinc-cadmium resistance pr |
| AP53_457 | 0.29669797 | 2.788920823 | 0.324514519 | 0.579945452 | 0.9997444 | FIG01117988: hypothetical protei |
| AP53_747 | 0.159160876 | 9.584863311 | 0.323936916 | 0.580278197 | 0.9997444 | FIG01114562: hypothetical protei |
| AP53_570 | 0.118735208 | 9.326678034 | 0.322587569 | 0.581057081 | 0.9997444 | DNA-entry nuclease (Competence-s |
| AP53_155 | 0.124303458 | 9.707554236 | 0.321670043 | 0.581587953 | 0.9997444 | Co-activator of prophage gene expr |
| AP53_783 | 0.085145149 | 7.122457783 | 0.320534057 | 0.582246633 | 0.9997444 | Hypothetical protein SPy1011 |
| AP53_464 | -0.136001643 | 5.259576695 | 0.320049678 | 0.582527966 | 0.9997444 | Beta-glucoside bgl operon antiterm |
| AP53_1242 | -0.141323481 | 9.804031108 | 0.318754185 | 0.583281806 | 0.9997444 | Cell division protein FtsZ (EC 3.4.24 |
| AP53_181 | -0.127697196 | 6.519016854 | 0.318727453 | 0.583297384 | 0.9997444 | Integral membrane protein (Rhomb |
| AP53_969 | 0.090252357 | 8.451923842 | 0.318439525 | 0.583465215 | 0.9997444 | tRNA pseudouridine synthase B (EC |
| AP53_1651 | 0.112995851 | 8.844024322 | 0.31690161 | 0.584363378 | 0.9997444 | Peptide deformylase (EC 3.5.1.88) |
| AP53_760 | 0.165583664 | 10.99692506 | 0.316536198 | 0.584577211 | 0.9997444 | SSU ribosomal protein S1p |
| AP53_291 | -0.063462435 | 7.932082255 | 0.315004547 | 0.585475305 | 0.9997444 | Segregation and condensation prot |
| AP53_31 | -0.089803101 | 7.975795889 | 0.313476589 | 0.58637414 | 0.9997444 | FIG01114657: hypothetical protei |
| AP53_736 | 0.124912378 | 8.860886175 | 0.312517442 | 0.586939858 | 0.9997444 | GTP-binding protein EngB |
| AP53_306 | 0.099866614 | 6.313976703 | 0.31185049 | 0.587333917 | 0.9997444 | Ferrichrome transport ATP-binding |
| AP53_678 | -0.057486139 | 8.376474821 | 0.309145379 | 0.588937962 | 0.9997444 | tRNA S(4)U 4-thiouridine synthase ( |
| AP53_1657 | -0.073046203 | 8.21286656 | 0.308689851 | 0.589208991 | 0.9997444 | Phosphatidate cytidylyltransferase |
| AP53_296 | 0.103638988 | 7.707430947 | 0.308638426 | 0.589239604 | 0.9997444 | Membrane-associated phospholipi |
| AP53_968 | 0.053976264 | 7.568667338 | 0.307433421 | 0.589957918 | 0.9997444 | Riboflavin kinase (EC 2.7.1.26) / FM |
| AP53_337 | 0.072186992 | 5.666366932 | 0.306682407 | 0.590406548 | 0.9997444 | Mobile element protein |
| AP53_1028 | -0.21299565 | 5.282694774 | 0.306401985 | 0.590574249 | 0.9997444 | Phage terminase, large subunit |

| | | | | | | |
|---|---|---|---|---|---|---|
| AP53_1834 | -0.102037493 | 7.62930424 | 0.306360567 | 0.590599027 | 0.9997444 | Phosphoesterase, DHH family prote |
| AP53_920 | 0.093860635 | 9.684761164 | 0.304257236 | 0.591860249 | 0.9997444 | Tyrosine recombinase XerC |
| AP53_1826 | -0.10256981 | 6.765204913 | 0.304247446 | 0.591866133 | 0.9997444 | FIG015389: hypothetical membrar |
| AP53_621 | 0.129187169 | 3.43401959 | 0.304225146 | 0.591879536 | 0.9997444 | Portal protein, phage associated |
| AP53_1862 | 0.155350506 | 8.378678594 | 0.303966208 | 0.592035211 | 0.9997444 | LSU m3Psi1915 methyltransferase I |
| AP53_1172 | 0.088909994 | 7.299821759 | 0.302044058 | 0.593193577 | 0.9997444 | Transcriptional repressor for NAD b |
| AP53_693 | 0.089469551 | 9.317769129 | 0.300865544 | 0.593906206 | 0.9997444 | ABC transporter permease protein |
| AP53_1691 | 0.134446836 | 10.28433427 | 0.300520319 | 0.594115306 | 0.9997444 | C5a peptidase (EC 3.4.21.-) |
| AP53_1688 | -0.074445479 | 5.913434124 | 0.300201307 | 0.594308671 | 0.9997444 | Laminin-binding surface protein |
| AP53_1451 | -0.081253882 | 6.309742493 | 0.299503869 | 0.594731885 | 0.9997444 | Negative transcriptional regulator- |
| AP53_1140 | -0.1577365 | 6.040327791 | 0.299248372 | 0.594887086 | 0.9997444 | Transcriptional regulator, Cro/CI fa |
| AP53_7 | 0.078157446 | 6.414484418 | 0.299195908 | 0.594918966 | 0.9997444 | Transcription-repair coupling facto |
| AP53_1103 | 0.097826812 | 7.380225059 | 0.298979706 | 0.595050381 | 0.9997444 | Transporter, MFS superfamily |
| AP53_814 | 0.077958421 | 5.701081 | 0.297889539 | 0.595713976 | 0.9997444 | Mobile element protein |
| AP53_618 | -0.190914671 | 4.72583804 | 0.296362433 | 0.596646231 | 0.9997444 | hypothetical phage protein |
| AP53_1835 | -0.061501958 | 7.517769788 | 0.295584244 | 0.597122509 | 0.9997444 | tRNA uridine 5-carboxymethylamir |
| AP53_628 | 0.15333104 | 2.940810817 | 0.294152976 | 0.598000646 | 0.9997444 | Phage integrase: site-specific recom |
| AP53_899 | 0.067387922 | 8.193042399 | 0.293312963 | 0.598517331 | 0.9997444 | DNA topoisomerase I (EC 5.99.1.2) |
| AP53_1197 | 0.151748056 | 3.318117855 | 0.293023406 | 0.59869566 | 0.9997444 | FIG048485: Phage protein |
| AP53_34 | -0.193930618 | 5.755964412 | 0.292458941 | 0.599043629 | 0.9997444 | Transcriptional regulator |
| AP53_219 | -0.115263948 | 11.68278344 | 0.291391324 | 0.599702974 | 0.9997444 | SSU ribosomal protein S12p (S23e) |
| AP53_855 | -0.101146447 | 6.633769712 | 0.290850041 | 0.600037867 | 0.9997444 | Permease |
| AP53_1609 | -0.059622045 | 7.566110226 | 0.288701887 | 0.601370969 | 0.9997444 | Type I restriction-modification syst |
| AP53_177 | 0.083817262 | 8.91860337 | 0.288524255 | 0.601481493 | 0.9997444 | Glucose-6-phosphate isomerase (EC |
| AP53_1013 | -0.228635386 | 5.972803166 | 0.287992732 | 0.601812479 | 0.9997444 | Phage-related protein |
| AP53_1398 | -0.065453181 | 9.98185083 | 0.287567402 | 0.602077623 | 0.9997444 | Multimodular transpeptidase-trans |
| AP53_281 | 0.061143714 | 8.123607792 | 0.287348921 | 0.602213921 | 0.9997444 | FIG011178: rRNA methylase |
| AP53_622 | -0.120360833 | 3.759174187 | 0.286030242 | 0.603038007 | 0.9997444 | Phage terminase, large subunit |
| AP53_297 | 0.080783585 | 7.089865324 | 0.285525483 | 0.603354102 | 0.9997444 | COG1242: Predicted Fe-S oxidored |
| AP53_rna21 | -0.26159738 | 4.774862161 | 0.285085789 | 0.603629749 | 0.9997444 | tRNA-Leu-TAG |
| AP53_954 | -0.152704992 | 9.975002689 | 0.284855887 | 0.603773987 | 0.9997444 | SSU ribosomal protein S20p |
| AP53_719 | 0.054869249 | 8.308501386 | 0.28402352 | 0.604296837 | 0.9997444 | COG0488: ATPase components of A |
| AP53_1597 | 0.058467492 | 8.736347993 | 0.283832828 | 0.60441676 | 0.9997444 | CTP synthase (EC 6.3.4.2) |
| AP53_1430 | 0.102162192 | 7.962175931 | 0.283680766 | 0.604512427 | 0.9997444 | Glycyl-tRNA synthetase alpha chain |
| AP53_1810 | -0.089324056 | 7.450918334 | 0.282260277 | 0.605407717 | 0.9997444 | membrane protein, putative |

| | | | | | | |
|---|---|---|---|---|---|---|
| AP53_956 | 0.147613735 | 7.171513726 | 0.281942547 | 0.605608373 | 0.9997444 | Two component system response re |
| AP53_1708 | 0.127309829 | 6.4287874 | 0.281676746 | 0.605776347 | 0.9997444 | Streptopain inhibitor |
| AP53_1702 | 0.088163476 | 7.891787285 | 0.280825718 | 0.606314851 | 0.9997444 | ABC transporter ATP-binding protei |
| AP53_1084 | -0.164829638 | 5.287488837 | 0.280347284 | 0.606618054 | 0.9997444 | PTS system, cellobiose-specific IIA c |
| AP53_636 | -0.104133533 | 4.112039212 | 0.279779752 | 0.606978158 | 0.9997444 | Phage protein |
| AP53_697 | 0.072864782 | 7.56475385 | 0.279646452 | 0.607062807 | 0.9997444 | Cell surface protein |
| AP53_213 | 0.062015295 | 9.447110586 | 0.27714187 | 0.608658161 | 0.9997444 | Ribulose-phosphate 3-epimerase (E |
| AP53_1455 | 0.080315365 | 9.896842533 | 0.276715304 | 0.608930803 | 0.9997444 | ribosomal protein L7Ae family prot |
| AP53_704 | -0.079621085 | 8.096405396 | 0.27649093 | 0.609074321 | 0.9997444 | Membrane protein |
| AP53_541 | 0.062095303 | 8.691959385 | 0.275808686 | 0.609511174 | 0.9997444 | hydrolase, haloacid dehalogenase-l |
| AP53_1460 | 0.074881148 | 9.238176012 | 0.275417376 | 0.609762052 | 0.9997444 | tRNA (guanine46-N7-)-methyltransf |
| AP53_111 | -0.123494421 | 6.855954303 | 0.274595544 | 0.6102897 | 0.9997444 | Short chain fatty acids transporter |
| AP53_1039 | -0.098759458 | 5.831925291 | 0.274264431 | 0.610502575 | 0.9997444 | putative replication protein |
| AP53_118 | 0.070003526 | 7.703854256 | 0.273890445 | 0.610743213 | 0.9997444 | Transcriptional regulator pfoR |
| AP53_35 | -0.097616026 | 8.844644952 | 0.273764685 | 0.61082418 | 0.9997444 | Holliday junction DNA helicase Ruv |
| AP53_1355 | 0.18652733 | 4.089876018 | 0.273638712 | 0.610905308 | 0.9997444 | hypothetical protein, phage associa |
| AP53_1841 | -0.082868349 | 7.681483501 | 0.273395049 | 0.611062299 | 0.9997444 | Transmembrane component of gen |
| AP53_647 | 0.185362212 | 3.469571357 | 0.273258643 | 0.611150224 | 0.9997444 | hypothetical phage protein |
| AP53_1442 | -0.073429672 | 8.913391452 | 0.272841475 | 0.611419298 | 0.9997444 | Tagatose-6-phosphate kinase (EC 2.7 |
| AP53_1827 | 0.128116141 | 7.738498587 | 0.272286142 | 0.611777902 | 0.9997444 | Phage infection protein |
| AP53_rna61 | 0.384986769 | 1.540564622 | 0.271936863 | 0.612003689 | 0.9997444 | tRNA-Thr-GGT |
| AP53_1649 | 0.114933299 | 7.311415974 | 0.271652812 | 0.612187448 | 0.9997444 | FIG01114948: hypothetical protei |
| AP53_716 | 0.174753736 | 9.254855066 | 0.271628585 | 0.612203126 | 0.9997444 | Hypothetical protein SP1558 |
| AP53_1755 | -0.07345526 | 11.0368012 | 0.271181175 | 0.612492834 | 0.9997444 | Translation elongation factor Ts |
| AP53_1815 | -0.141216771 | 9.372363644 | 0.269093607 | 0.61384866 | 0.9997444 | LSU ribosomal protein L33p @ LSU |
| AP53_29 | 0.190197549 | 9.432280448 | 0.26891658 | 0.613963946 | 0.9997444 | Phosphoribosylaminoimidazole car |
| AP53_1104 | 0.067785178 | 8.137301301 | 0.268822632 | 0.614025148 | 0.9997444 | ATP-dependent Zn protease |
| AP53_1628 | 0.217417762 | 2.437120727 | 0.268768804 | 0.61406022 | 0.9997444 | Integrase |
| AP53_173 | 0.095182952 | 6.451482905 | 0.266964049 | 0.615238745 | 0.9997444 | S-layer homology domain / putative |
| AP53_1096 | 0.084182152 | 5.522855 | 0.266960718 | 0.615240925 | 0.9997444 | Quinolinate phosphoribosyltransfe |
| AP53_1817 | 0.085233951 | 5.415938694 | 0.266629196 | 0.61545797 | 0.9997444 | Cadmium efflux system accessory p |
| AP53_1004 | -0.058432114 | 9.335315697 | 0.264988126 | 0.616534924 | 0.9997444 | Phage lysin |
| AP53_rna64 | 0.257607848 | 4.164097717 | 0.263830816 | 0.617296983 | 0.9997444 | tRNA-Thr-TGT |
| AP53_738 | 0.141346435 | 13.6746115 | 0.263570534 | 0.617468666 | 0.9997444 | putative ATP-dependent Clp protei |
| AP53_1024 | -0.23901674 | 7.609041708 | 0.263543303 | 0.617486634 | 0.9997444 | Prophage Clp protease-like protein |

| | | | | | | |
|---|---|---|---|---|---|---|
| AP53_51 | 0.127097117 | 11.07336914 | 0.263285193 | 0.617657004 | 0.9997444 | LSU ribosomal protein L29p (L35e) |
| AP53_1509 | 0.074178427 | 7.486170921 | 0.263163437 | 0.617737407 | 0.9997444 | L-asparaginase (EC 3.5.1.1) |
| AP53_1025 | -0.248662416 | 6.219577531 | 0.262302105 | 0.618306887 | 0.9997444 | Phage portal protein |
| AP53_655 | -0.06356809 | 9.319195173 | 0.26054673 | 0.619471183 | 0.9997444 | Phage integrase: site-specific recom |
| AP53_136 | 0.054726504 | 9.356913988 | 0.260502043 | 0.619500887 | 0.9997444 | Thiol-activated cytolysin |
| AP53_1440 | -0.178419191 | 6.64335366 | 0.25971356 | 0.620025557 | 0.9997444 | hypothetical protein |
| AP53_852 | 0.084268004 | 6.142123985 | 0.259566888 | 0.620123267 | 0.9997444 | Voltage-gated chloride channel fam |
| AP53_1830 | 0.072263067 | 5.675547829 | 0.259010562 | 0.620494201 | 0.9997444 | Mobile element protein |
| AP53_781 | -0.057136686 | 9.416215669 | 0.258458817 | 0.620862583 | 0.9997444 | dTDP-glucose 4,6-dehydratase (EC 4 |
| AP53_993 | 0.094629339 | 11.49556352 | 0.257500973 | 0.621503295 | 0.9997444 | Pyruvate kinase (EC 2.7.1.40) |
| AP53_1190 | 0.145272521 | 2.186103157 | 0.257293451 | 0.621642309 | 0.9997444 | Phage minor tail protein |
| AP53_548 | -0.123597557 | 7.536666595 | 0.256087974 | 0.622451244 | 0.9997444 | Streptolysin S biosynthesis protein |
| AP53_288 | -0.098126633 | 7.785351794 | 0.255673595 | 0.622729872 | 0.9997444 | FIG042801: CBS domain containing |
| AP53_114 | 0.105517382 | 5.371535414 | 0.254288937 | 0.623663003 | 0.9997444 | Acetyl-CoA:acetoacetyl-CoA transfe |
| AP53_1545 | 0.080801107 | 11.4512735 | 0.254261764 | 0.623681347 | 0.9997444 | SSU ribosomal protein S6p |
| AP53_1690 | 0.16025135 | 11.27051386 | 0.254036302 | 0.623833604 | 0.9997444 | Fibronectin-binding protein |
| AP53_345 | -0.128241101 | 8.13500415 | 0.253600684 | 0.624128024 | 0.9997444 | 3-oxoacyl-[acyl-carrier protein] red |
| AP53_1539 | 0.054671204 | 8.072907953 | 0.253355406 | 0.62429394 | 0.9997444 | Aminopeptidase YpdF (MP-, MA-, M |
| AP53_1753 | -0.110430811 | 5.306778081 | 0.251226394 | 0.625738395 | 0.9997444 | Transcriptional regulator, LuxR fam |
| AP53_rna52 | 0.293524575 | 3.930072791 | 0.251180343 | 0.625769724 | 0.9997444 | tRNA-Ala-TGC |
| AP53_1510 | 0.097821227 | 6.923368282 | 0.249667248 | 0.626801145 | 0.9997444 | ABC transporter permease protein |
| AP53_rna46 | 0.290576401 | 3.464757328 | 0.249041089 | 0.627229132 | 0.9997444 | tRNA-Tyr-GTA |
| AP53_1792 | -0.190566763 | 4.794705572 | 0.248808521 | 0.627388269 | 0.9997444 | phage protein |
| AP53_rna79 | 0.238013493 | 3.888713338 | 0.248039744 | 0.627914981 | 0.9997444 | tRNA-Ala-TGC |
| AP53_410 | -0.096486586 | 9.975201927 | 0.246602888 | 0.628902185 | 0.9997444 | Preprotein translocase subunit Sec( |
| AP53_1047 | -0.127784075 | 6.075203301 | 0.245875914 | 0.629403043 | 0.9997444 | hypothetical phage protein |
| AP53_644 | 0.083583427 | 4.75123839 | 0.245326361 | 0.629782284 | 0.9997444 | Replicative DNA helicase (DnaB) (EC |
| AP53_138 | 0.135911873 | 10.18656462 | 0.245284527 | 0.629811175 | 0.9997444 | FIG01114903: hypothetical protein |
| AP53_57 | 0.106139092 | 10.59405612 | 0.245056016 | 0.629969043 | 0.9997444 | SSU ribosomal protein S8p (S15Ae) |
| AP53_1211 | -0.129448685 | 1.877716067 | 0.244765682 | 0.630169757 | 0.9997444 | Phage excisionase |
| AP53_1421 | -0.113649602 | 6.927983678 | 0.244428813 | 0.630402828 | 0.9997444 | Trans-acting positive regulator |
| AP53_385 | 0.097626303 | 4.362309074 | 0.2443571 | 0.630452471 | 0.9997444 | Mobile element protein |
| AP53_1566 | -0.098772926 | 7.086791417 | 0.243807864 | 0.630832978 | 0.9997444 | Transcriptional regulator, CrP/Fnr f |
| AP53_540 | -0.098298285 | 12.75185578 | 0.242738012 | 0.631575719 | 0.9997444 | LSU ribosomal protein L19p |
| AP53_768 | -0.049048105 | 8.452878255 | 0.242508496 | 0.631735328 | 0.9997444 | GTP-binding protein HflX |

| | | | | | | |
|---|---|---|---|---|---|---|
| AP53_412 | -0.141358833 | 9.840928338 | 0.24237999 | 0.631824734 | 0.9997444 | tmRNA-binding protein SmpB |
| AP53_231 | -0.088372846 | 8.46742801 | 0.242133108 | 0.631996583 | 0.9997444 | Iron-sulfur cluster assembly ATPase |
| AP53_202 | -0.085934481 | 7.286735702 | 0.241922066 | 0.632143572 | 0.9997444 | ABC transporter, predicted N-acety |
| AP53_1436 | -0.14089412 | 7.919112504 | 0.240867063 | 0.632879583 | 0.9997444 | DegV family protein |
| AP53_rna84 | -0.258523796 | 1.78152497 | 0.240704253 | 0.632993347 | 0.9997444 | tRNA-Glu-TTC |
| AP53_1118 | 0.097464593 | 4.120227588 | 0.240012226 | 0.633477438 | 0.9997444 | internalin, putative |
| AP53_1498 | -0.106208686 | 6.613320693 | 0.238482933 | 0.634550337 | 0.9997444 | hypothetical protein |
| AP53_594 | 0.068899782 | 6.746826299 | 0.238453208 | 0.634571234 | 0.9997444 | Alpha-L-Rha alpha-1,2-L-rhamnosyl |
| AP53_102 | -0.083942838 | 6.187645192 | 0.238391854 | 0.634614371 | 0.9997444 | Transcriptional regulator, RofA-like |
| AP53_503 | -0.059383673 | 7.641910473 | 0.237967846 | 0.634912675 | 0.9997444 | Deoxyguanosinetriphosphate triph |
| AP53_303 | 0.109493211 | 6.727022064 | 0.237773858 | 0.635049263 | 0.9997444 | ABC-type Fe3+-siderophore transpo |
| AP53_1012 | -0.196226983 | 5.87453472 | 0.236496015 | 0.635950751 | 0.9997444 | Phage hyaluronidase; Phage endop |
| AP53_965 | 0.072685159 | 6.971739361 | 0.236479079 | 0.63596272 | 0.9997444 | Inositol-1-monophosphatase (EC 3. |
| AP53_rna67 | 0.192817141 | 4.349876004 | 0.236327152 | 0.636070109 | 0.9997444 | tRNA-Asp-GTC |
| AP53_1217 | -0.052849731 | 8.364502351 | 0.235450099 | 0.636690893 | 0.9997444 | YfaA |
| AP53_1209 | 0.181959807 | 2.866562843 | 0.235395719 | 0.636729431 | 0.9997444 | DNA replication protein dnaD |
| AP53_1619 | -0.10420119 | 6.430822598 | 0.235152384 | 0.636901947 | 0.9997444 | PTS system, lactose-specific IIA com |
| AP53_479 | 0.055575928 | 8.07023155 | 0.234834507 | 0.637127478 | 0.9997444 | peptidase, U32 family large subunit |
| AP53_883 | -0.047678883 | 8.535576241 | 0.23452315 | 0.637348568 | 0.9997444 | Pneumococcal vaccine antigen A ho |
| AP53_1839 | -0.061553504 | 6.918112071 | 0.233578645 | 0.638020372 | 0.9997444 | L-serine dehydratase, alpha subunit |
| AP53_1599 | 0.099829392 | 9.99443516 | 0.233538908 | 0.638048673 | 0.9997444 | Cell division trigger factor (EC 5.2.1 |
| AP53_153 | 0.064546629 | 10.55554949 | 0.232107531 | 0.639070128 | 0.9997444 | Peroxide stress regulator PerR, FUR |
| AP53_369 | 0.116975064 | 6.804805955 | 0.231884108 | 0.639229921 | 0.9997444 | Phosphate starvation-inducible pro |
| AP53_1169 | -0.092942398 | 6.773293348 | 0.231838615 | 0.639262469 | 0.9997444 | Formate/nitrite family of transport |
| AP53_478 | 0.045699488 | 6.86776325 | 0.230470893 | 0.640242886 | 0.9997444 | PROTEASE (EC 3.4.-.-) |
| AP53_961 | -0.060148758 | 7.148994965 | 0.230354996 | 0.64032613 | 0.9997444 | Phosphate transport system perme |
| AP53_1108 | 0.055400641 | 8.456743553 | 0.230068916 | 0.640531721 | 0.9997444 | Cell envelope-associated transcript |
| AP53_645 | -0.145228525 | 4.180054042 | 0.229633287 | 0.640845093 | 0.9997444 | hypothetical protein |
| AP53_950 | -0.100304572 | 10.0211407 | 0.228646628 | 0.641556219 | 0.9997444 | Nucleoside-binding protein |
| AP53_1324 | 0.150210885 | 3.836125107 | 0.228521531 | 0.641646519 | 0.9997444 | Phage minor capsid protein |
| AP53_741 | -0.090387328 | 9.877237239 | 0.227892985 | 0.642100689 | 0.9997444 | Arsenate reductase (EC 1.20.4.1) |
| AP53_1621 | 0.09890737 | 6.750819631 | 0.227422741 | 0.642440984 | 0.9997444 | Tagatose-6-phosphate kinase (EC 2. |
| AP53_1673 | -0.099564553 | 7.054821001 | 0.226819484 | 0.642878175 | 0.9997444 | Ribosomal RNA small subunit meth |
| AP53_pseudo1337 | -0.161102261 | 2.222517796 | 0.226638543 | 0.643009447 | 0.9997444 | DNA modification methylase |
| AP53_778 | 0.059284926 | 7.90192138 | 0.226270727 | 0.643276497 | 0.9997444 | Glycine/D-amino acid oxidases fam |

| | | | | | | |
|---|---|---|---|---|---|---|
| AP53_295 | 0.099269765 | 6.996968242 | 0.226140687 | 0.643370975 | 0.9997444 | Substrate-specific component RibU |
| AP53_576 | 0.092553336 | 6.351748009 | 0.225542041 | 0.643806349 | 0.9997444 | FIG01115289: hypothetical protein |
| AP53_509 | 0.075646521 | 5.783510146 | 0.224622165 | 0.644476741 | 0.9997444 | PTS system, N-acetylgalactosamine- |
| AP53_1831 | -0.137100599 | 5.756421826 | 0.224378611 | 0.644654525 | 0.9997444 | FIG01114855: hypothetical protein |
| AP53_592 | 0.087067533 | 7.379795712 | 0.222598797 | 0.645957352 | 0.9997444 | ABC-transporter (ATP-binding prote |
| AP53_225 | 0.056084591 | 8.009070346 | 0.222571067 | 0.645977701 | 0.9997444 | Amino acid ABC transporter, glutan |
| AP53_353 | 0.067034631 | 9.770017578 | 0.221713833 | 0.646607547 | 0.9997444 | Mn-dependent transcriptional regu |
| AP53_1221 | -0.045638774 | 8.478727955 | 0.221108315 | 0.647053355 | 0.9997444 | DNA repair protein RecN |
| AP53_547 | -0.098747176 | 7.169960578 | 0.220496645 | 0.647504459 | 0.9997444 | Streptolysin S biosynthesis protein |
| AP53_rna32 | -0.26431447 | 4.160629324 | 0.220246648 | 0.647689053 | 0.9997444 | tRNA-Gly-TCC |
| AP53_990 | 0.11450587 | 8.554446163 | 0.219549373 | 0.648204596 | 0.9997444 | Alkylphosphonate utilization operc |
| AP53_1659 | 0.141132193 | 9.560678367 | 0.218624023 | 0.648890331 | 0.9997444 | Preprotein translocase subunit Yaj( |
| AP53_1846 | -0.066722548 | 6.160194467 | 0.217216115 | 0.649937111 | 0.9997444 | FIG009210: peptidase, M16 family |
| AP53_1113 | 0.05063179 | 7.915725624 | 0.215408357 | 0.651287325 | 0.9997444 | FIG009439: Cytosolic protein cont |
| AP53_630 | 0.103526531 | 4.836682553 | 0.215139227 | 0.651488934 | 0.9997444 | B-cell receptor associated protein-r |
| AP53_1497 | -0.088916054 | 7.204863828 | 0.214793792 | 0.651747931 | 0.9997444 | Integral membrane protein |
| AP53_1064 | 0.093671793 | 6.36318039 | 0.213514099 | 0.652709644 | 0.9997444 | 4-alpha-glucanotransferase (amylor |
| AP53_909 | -0.079641906 | 5.601753535 | 0.212661909 | 0.653352045 | 0.9997444 | FIG01114846: hypothetical protein |
| AP53_rna20 | -0.278113331 | 4.503413543 | 0.212106802 | 0.653771348 | 0.9997444 | tRNA-Lys-TTT |
| AP53_1595 | 0.053368325 | 9.256043471 | 0.211604433 | 0.654151395 | 0.9997444 | Hydrolases of the alpha/beta superf |
| AP53_998 | -0.053873596 | 8.269286534 | 0.211545898 | 0.654195713 | 0.9997444 | ABC transporter, ATP-binding prote |
| AP53_850 | -0.058993617 | 5.641731126 | 0.211268076 | 0.654406162 | 0.9997444 | Mobile element protein |
| AP53_780 | 0.067585602 | 9.426762818 | 0.211031361 | 0.654585606 | 0.9997444 | dTDP-4-dehydrorhamnose 3,5-epin |
| AP53_447 | 0.110138876 | 4.439548682 | 0.209614614 | 0.65566217 | 0.9997444 | ABC transporter membrane-spanni |
| AP53_1349 | 0.148738762 | 3.309771167 | 0.20924744 | 0.655941906 | 0.9997444 | hypothetical protein (phage associa |
| AP53_228 | -0.068156815 | 8.054936901 | 0.207492342 | 0.657283213 | 0.9997444 | Negative regulator of genetic comp |
| AP53_1684 | -0.071029829 | 6.496705515 | 0.207351234 | 0.657391354 | 0.9997444 | Dipeptide transport ATP-binding pr |
| AP53_1120 | -0.045962663 | 8.887645674 | 0.20668652 | 0.657901373 | 0.9997444 | FIG01114335: hypothetical protein |
| AP53_867 | -0.067296795 | 6.75101341 | 0.20641311 | 0.658111445 | 0.9997444 | Short chain dehydrogenase |
| AP53_222 | 0.070306239 | 12.83297456 | 0.205631586 | 0.658712858 | 0.9997444 | NAD-dependent glyceraldehyde-3-p |
| AP53_1086 | 0.104353377 | 4.62286242 | 0.205475064 | 0.658833475 | 0.9997444 | Transcriptional antiterminator of li |
| AP53_1147 | 0.143844822 | 9.234701475 | 0.204462738 | 0.659614939 | 0.9997444 | Competence protein CoiA |
| AP53_1795 | -0.126603362 | 5.452817499 | 0.202415697 | 0.661202381 | 0.9997444 | Phage protein |
| AP53_740 | -0.088823659 | 10.38253532 | 0.201019922 | 0.662290386 | 0.9997444 | Phosphopentomutase (EC 5.4.2.7) |
| AP53_1166 | -0.051296239 | 8.162139025 | 0.200136356 | 0.662981497 | 0.9997444 | UDP-N-acetylmuramoylalanyl-D-glu |

| | | | | | | |
|---|---|---|---|---|---|---|
| AP53_888 | 0.042693673 | 9.182311722 | 0.200132376 | 0.662984614 | 0.9997444 | DNA gyrase subunit A (EC 5.99.1.3) |
| AP53_1335 | 0.163864531 | 3.024041073 | 0.199493381 | 0.663485588 | 0.9997444 | Phage protein |
| AP53_1652 | 0.062369886 | 8.385416906 | 0.199324949 | 0.663617801 | 0.9997444 | FIG01114677: hypothetical protei |
| AP53_486 | 0.109919838 | 8.007340446 | 0.199215766 | 0.663703542 | 0.9997444 | Transcriptional regulator |
| AP53_683 | -0.071447437 | 7.300666495 | 0.198677814 | 0.664126409 | 0.9997444 | Cysteine and methionine metabolis |
| AP53_1624 | 0.097560707 | 6.419915055 | 0.198059877 | 0.664613009 | 0.9997444 | Lactose phosphotransferase system |
| AP53_157 | 0.115483748 | 5.443569593 | 0.198001161 | 0.664659293 | 0.9997444 | FIG01115469: hypothetical protei |
| AP53_65 | -0.135837349 | 8.071274492 | 0.197829042 | 0.664795018 | 0.9997444 | Translation initiation factor 1 |
| AP53_1194 | 0.234387454 | 1.506943889 | 0.197777245 | 0.664835877 | 0.9997444 | FIG018217: Phage-associated prote |
| AP53_1631 | -0.089608466 | 11.02108569 | 0.19750552 | 0.665050325 | 0.9997444 | LSU ribosomal protein L13p (L13Ae |
| AP53_1238 | -0.076481101 | 9.054908391 | 0.197311975 | 0.665203182 | 0.9997444 | FIG001583: hypothetical protein, |
| AP53_662 | 0.070402017 | 8.236886967 | 0.197011937 | 0.665440324 | 0.9997444 | Pore forming protein ebsA |
| AP53_1425 | 0.059153561 | 7.44292707 | 0.195605784 | 0.666554633 | 0.9997444 | Glycerol kinase (EC 2.7.1.30) |
| AP53_1637 | 0.064409648 | 6.216054756 | 0.195491088 | 0.666645737 | 0.9997444 | Hypothetical protein, Spy1939 hor |
| AP53_879 | 0.039999353 | 8.401266095 | 0.195055891 | 0.666991714 | 0.9997444 | TsaC protein (YrdC domain) require |
| AP53_224 | 0.061662785 | 8.276251153 | 0.19466423 | 0.667303479 | 0.9997444 | Amino acid transport ATP-binding |
| AP53_1866 | 0.113869379 | 4.323233914 | 0.194367475 | 0.667539951 | 0.9997444 | paratox? |
| AP53_908 | -0.08924969 | 5.781226351 | 0.194127394 | 0.667731421 | 0.9997444 | Mg2+/citrate complex transporter |
| AP53_1072 | -0.067231525 | 9.306028122 | 0.193349354 | 0.66835291 | 0.9997444 | putative esterase |
| AP53_634 | -0.131454041 | 4.245122489 | 0.190821546 | 0.670382566 | 0.9997444 | hypothetical phage protein |
| AP53_344 | -0.086364946 | 7.517459375 | 0.190487078 | 0.670652331 | 0.9997444 | FIG01114304: hypothetical protei |
| AP53_1176 | 0.114856127 | 10.08705089 | 0.189532495 | 0.671423822 | 0.9997444 | FIG01114217: hypothetical protei |
| AP53_1417 | -0.056081005 | 8.884941693 | 0.188982853 | 0.671869102 | 0.9997444 | Methionine ABC transporter ATP-bi |
| AP53_1226 | 0.039273869 | 8.30245655 | 0.18831995 | 0.672407175 | 0.9997444 | Exodeoxyribonuclease VII large sub |
| AP53_1162 | 0.119008856 | 5.587893052 | 0.188213375 | 0.672493787 | 0.9997444 | Kup system potassium uptake prote |
| AP53_414 | 0.066752623 | 9.780415718 | 0.187753464 | 0.672867888 | 0.9997444 | Pyrrolidone-carboxylate peptidase |
| AP53_482 | -0.065930109 | 7.245107736 | 0.187474798 | 0.673094829 | 0.9997444 | FIG01116574: hypothetical protei |
| AP53_1253 | -0.070450854 | 4.832420676 | 0.187473357 | 0.673096003 | 0.9997444 | Late competence protein ComC, pr |
| AP53_1325 | -0.154242297 | 2.984739862 | 0.187199117 | 0.673319538 | 0.9997444 | Phage minor capsid protein #Fam0 |
| AP53_1005 | -0.063765359 | 7.959082217 | 0.186973269 | 0.673503777 | 0.9997444 | Phage protein |
| AP53_53 | 0.111165072 | 11.43545424 | 0.186593742 | 0.673813681 | 0.9997444 | LSU ribosomal protein L14p (L23e) |
| AP53_rna58 | 0.194222792 | 3.880953751 | 0.186101539 | 0.674216156 | 0.9997444 | tRNA-Tyr-GTA |
| AP53_959 | -0.059585105 | 7.296947342 | 0.184776523 | 0.675302803 | 0.9997444 | Phosphate transport ATP-binding p |
| AP53_201 | -0.078259161 | 6.70652531 | 0.183999087 | 0.675942552 | 0.9997444 | N-acetylmannosamine-6-phosphat |
| AP53_1819 | 0.099603854 | 6.604633592 | 0.183886669 | 0.676035194 | 0.9997444 | FtsK/SpoIIIE family |

| | | | | | | |
|---|---|---|---|---|---|---|
| AP53_275 | -0.061250462 | 9.150603593 | 0.183751711 | 0.676146457 | 0.9997444 | UDP-N-acetylmuramate--alanine lig |
| AP53_1157 | 0.087782058 | 3.708622363 | 0.183050625 | 0.676725233 | 0.9997444 | Late competence protein ComEC, D |
| AP53_365 | -0.127707358 | 7.934240173 | 0.182943881 | 0.676813471 | 0.9997444 | Peptide methionine sulfoxide redu |
| AP53_373 | -0.061892929 | 9.046666375 | 0.182358994 | 0.677297504 | 0.9997444 | GTP-binding protein Era |
| AP53_321 | -0.055739042 | 9.194140231 | 0.182229202 | 0.677405042 | 0.9997444 | Arsenate reductase (EC 1.20.4.1) |
| AP53_1026 | -0.15268863 | 5.864797043 | 0.181834785 | 0.677732112 | 0.9997444 | Phage protein |
| AP53_1655 | -0.070835926 | 9.654728026 | 0.181429081 | 0.678068984 | 0.9997444 | Prolyl-tRNA synthetase (EC 6.1.1.15 |
| AP53_159 | 0.089171623 | 5.702756779 | 0.181320349 | 0.678159345 | 0.9997444 | Mobile element protein |
| AP53_409 | -0.053482608 | 9.444312881 | 0.181319923 | 0.678159699 | 0.9997444 | Multidrug resistance efflux pump P |
| AP53_577 | 0.129285876 | 8.271785237 | 0.180962922 | 0.67845661 | 0.9997444 | ABC transporter permease protein |
| AP53_184 | 0.05260621 | 8.108886386 | 0.180639188 | 0.678726157 | 0.9997444 | Transcriptional regulator, MarR fan |
| AP53_1496 | 0.045595829 | 8.657112087 | 0.180631993 | 0.67873215 | 0.9997444 | Phosphoglycerate mutase family |
| AP53_322 | -0.103102987 | 3.828451019 | 0.178193349 | 0.680771973 | 0.9997444 | FIG01115198: hypothetical protei |
| AP53_1352 | 0.187476473 | 4.312188956 | 0.178146821 | 0.680811053 | 0.9997444 | hypothetical phage protein |
| AP53_1589 | 0.064774749 | 9.549365128 | 0.178087697 | 0.680860722 | 0.9997444 | Acid phosphatase (EC 3.1.3.2) |
| AP53_842 | -0.058067052 | 7.789003183 | 0.177495541 | 0.681358718 | 0.9997444 | Spermidine Putrescine ABC transpo |
| AP53_1154 | 0.088430028 | 6.87505983 | 0.177232639 | 0.681580131 | 0.9997444 | FIG01116792: hypothetical protei |
| AP53_174 | -0.073249225 | 6.388664455 | 0.177005994 | 0.681771167 | 0.9997444 | Streptococcal pyrogenic exotoxin C |
| AP53_1543 | -0.065306037 | 10.58250761 | 0.176225736 | 0.682429949 | 0.9997444 | SSU ribosomal protein S18p @ SSU |
| AP53_1320 | -0.142422488 | 3.6637731 | 0.175508265 | 0.683037252 | 0.9997444 | Phage tail length tape-measure prot |
| AP53_1564 | 0.063966564 | 8.421111977 | 0.175195756 | 0.683302235 | 0.9997444 | Aquaporin (Major Intrinsic Protein |
| AP53_315 | 0.106886559 | 8.989167006 | 0.175065284 | 0.683412949 | 0.9997444 | hypothetical protein |
| AP53_1557 | -0.042459647 | 9.093260226 | 0.174774035 | 0.683660268 | 0.9997444 | Putative membrane protein precur |
| AP53_212 | 0.042474888 | 9.368271815 | 0.174026432 | 0.68429623 | 0.9997444 | Ribosome small subunit-stimulated |
| AP53_1760 | 0.119534496 | 4.176515525 | 0.173820347 | 0.684471825 | 0.9997444 | FIG01115695: hypothetical protei |
| AP53_902 | 0.075299789 | 4.822374628 | 0.172960764 | 0.685205568 | 0.9997444 | Oxaloacetate decarboxylase alpha c |
| AP53_1693 | -0.083945776 | 13.096781 | 0.172766243 | 0.685371911 | 0.9997444 | Antiphagocytic M protein |
| AP53_1546 | 0.083769751 | 9.595490849 | 0.172574405 | 0.685536069 | 0.9997444 | hypothetical protein |
| AP53_1105 | -0.10439804 | 7.325119416 | 0.172329206 | 0.685746046 | 0.9997444 | Phenylacetic acid degradation prot |
| AP53_326 | 0.058717807 | 9.854876452 | 0.172220477 | 0.685839213 | 0.9997444 | Serine endopeptidase ScpC (EC 3.4. |
| AP53_1469 | -0.14985697 | 8.997062334 | 0.171354985 | 0.686582076 | 0.9997444 | Guanine-hypoxanthine permease |
| AP53_992 | 0.122966478 | 9.403799049 | 0.171210579 | 0.686706238 | 0.9997444 | Signal peptidase I (EC 3.4.21.89) |
| AP53_1771 | 0.367487514 | 2.941398101 | 0.170279555 | 0.687508232 | 0.9997444 | hypothetical protein |
| AP53_1080 | 0.071719279 | 7.621737499 | 0.169634374 | 0.68806552 | 0.9997444 | Glutamate transport ATP-binding p |
| AP53_1091 | 0.07346421 | 8.27044064 | 0.16948415 | 0.688195458 | 0.9997444 | GTP-binding protein Obg |

| | | | | | | |
|---|---|---|---|---|---|---|
| AP53_616 | -0.146801997 | 5.136891904 | 0.169284313 | 0.688368416 | 0.9997444 | hypothetical phage protein |
| AP53_rna39 | -0.233444755 | 3.543319887 | 0.169213088 | 0.688430091 | 0.9997444 | tRNA-Val-TAC |
| AP53_426 | -0.077508977 | 3.992919516 | 0.168849529 | 0.688745136 | 0.9997444 | Daunorubicin resistance ATP-bindi |
| AP53_90 | 0.048907902 | 7.485381372 | 0.16884656 | 0.688747711 | 0.9997444 | Adenine-specific methyltransferase |
| AP53_1574 | -0.057611841 | 6.894513115 | 0.168557865 | 0.688998171 | 0.9997444 | NAD(FAD)-utilizing dehydrogenases |
| AP53_476 | -0.076769439 | 9.659574946 | 0.168529863 | 0.689022478 | 0.9997444 | FIG01115301: hypothetical protei |
| AP53_129 | -0.055426863 | 8.445112756 | 0.168359845 | 0.689170111 | 0.9997444 | putative toxic anion resistance pro |
| AP53_1824 | -0.089496896 | 3.817303732 | 0.166726529 | 0.69059289 | 0.9997444 | FIG014387: Transcriptional regulat |
| AP53_550 | -0.085537906 | 7.078859446 | 0.165747946 | 0.691449272 | 0.9997444 | Streptolysin S biosynthesis protein |
| AP53_1195 | -0.169995491 | 3.18282213 | 0.16557972 | 0.691596789 | 0.9997444 | FIG033577: Phage protein |
| AP53_1339 | -0.15430411 | 4.132509886 | 0.163818183 | 0.693146817 | 0.9997444 | hypothetical protein |
| AP53_1783 | 0.083032835 | 8.552077433 | 0.163549547 | 0.693384057 | 0.9997444 | hypothetical protein |
| AP53_680 | -0.080084338 | 10.64839791 | 0.16338969 | 0.69352534 | 0.9997444 | LSU ribosomal protein L21p |
| AP53_22 | 0.114857353 | 8.533996792 | 0.163227475 | 0.693668791 | 0.9997444 | Phosphoribosylformylglycinamidir |
| AP53_538 | -0.064074983 | 8.313456641 | 0.163080315 | 0.693799 | 0.9997444 | Chorismate mutase I (EC 5.4.99.5) |
| AP53_557 | -0.051058191 | 7.132158754 | 0.162917943 | 0.69394275 | 0.9997444 | DNA ligase (EC 6.5.1.2) |
| AP53_1006 | -0.168645335 | 6.091945859 | 0.162674391 | 0.694158526 | 0.9997444 | Phage lysin |
| AP53_1470 | -0.146300468 | 8.386196245 | 0.162673436 | 0.694159372 | 0.9997444 | Hydrolase (HAD superfamily) |
| AP53_806 | -0.077190492 | 6.347657363 | 0.162365202 | 0.694432727 | 0.9997444 | Hypothetical radical SAM family en |
| AP53_445 | 0.129044066 | 3.478242748 | 0.160890726 | 0.695744586 | 0.9997444 | Dolichyl-phosphate mannoosyltrar |
| AP53_rna85 | 0.118186684 | 4.683211769 | 0.160340807 | 0.69623566 | 0.9997444 | tRNA-Arg-TCT |
| AP53_52 | 0.101064067 | 11.32480098 | 0.159848947 | 0.696675724 | 0.9997444 | SSU ribosomal protein S17p (S11e) |
| AP53_684 | -0.054081746 | 7.260847375 | 0.159705484 | 0.696804228 | 0.9997444 | Lipoprotein signal peptidase (EC 3.4 |
| AP53_880 | 0.041591187 | 8.829390239 | 0.159419165 | 0.697060897 | 0.9997444 | Phosphinothricin N-acetyltransfera |
| AP53_417 | 0.080892855 | 7.960783991 | 0.158722501 | 0.697686545 | 0.9997444 | Bactoprenol glucosyl transferase (E |
| AP53_508 | 0.054306675 | 5.896771962 | 0.158588632 | 0.697806952 | 0.9997444 | PTS system, N-acetylgalactosamine- |
| AP53_1774 | -0.127320417 | 10.47176881 | 0.158189577 | 0.69816623 | 0.9997444 | Hypothetical protein possible func |
| AP53_504 | 0.065270206 | 7.698002855 | 0.157932086 | 0.698398337 | 0.9997444 | FIG01114378: hypothetical protei |
| AP53_135 | 0.049768451 | 8.773173766 | 0.157420573 | 0.698860079 | 0.9997444 | Streptococcal NAD glycohydrolase |
| AP53_rna11 | 0.169879621 | 4.042386343 | 0.156741952 | 0.699474026 | 0.9997444 | tRNA-Leu-TAA |
| AP53_1820 | 0.110530699 | 6.814986037 | 0.156459011 | 0.699730461 | 0.9997444 | FIG01114548: hypothetical protei |
| AP53_rna27 | -0.139282372 | 4.911695727 | 0.155816282 | 0.700313986 | 0.9997444 | tRNA-Met-CAT |
| AP53_1100 | 0.078545641 | 6.426479153 | 0.155540974 | 0.700564364 | 0.9997444 | Mobile element protein |
| AP53_1762 | 0.103268464 | 5.868500859 | 0.155309719 | 0.700774878 | 0.9997444 | Transcriptional regulator, HxlR fam |
| AP53_461 | -0.060541857 | 7.991869669 | 0.155132919 | 0.700935945 | 0.9997444 | Hydrolase (HAD superfamily) |

| | | | | | | |
|---|---|---|---|---|---|---|
| AP53_1797 | -0.111356408 | 3.912813951 | 0.155090106 | 0.700974963 | 0.9997444 | FIG01114328: hypothetical protei |
| AP53_1023 | -0.161430265 | 7.650861882 | 0.154901929 | 0.70114654 | 0.9997444 | Phage capsid protein |
| AP53_748 | 0.085932944 | 8.60819477 | 0.154599396 | 0.701422639 | 0.9997444 | Orotidine 5'-phosphate decarboxyl |
| AP53_420 | 0.048476469 | 9.55445663 | 0.153884367 | 0.702076446 | 0.9997444 | Proline dipeptidase (EC 3.4.13.9) |
| AP53_152 | 0.05355201 | 9.753289459 | 0.153183457 | 0.702719063 | 0.9997444 | Succinyl-CoA synthetase, alpha sub |
| AP53_663 | -0.073403771 | 7.573759383 | 0.153145375 | 0.702754027 | 0.9997444 | Ferredoxin |
| AP53_1151 | -0.050098789 | 7.822189144 | 0.152288369 | 0.703542202 | 0.9997444 | FIG01115059: hypothetical protei |
| AP53_284 | 0.085226167 | 9.147368134 | 0.152220496 | 0.703604734 | 0.9997444 | FIG027054: hypothetical protein |
| AP53_rna68 | 0.129205425 | 4.055443708 | 0.152220401 | 0.703604822 | 0.9997444 | tRNA-Val-TAC |
| AP53_1250 | 0.063485428 | 8.688805681 | 0.152118573 | 0.703698667 | 0.9997444 | Glucokinase (EC 2.7.1.2) |
| AP53_1042 | -0.075325494 | 6.407160573 | 0.15147627 | 0.704291462 | 0.9997444 | FIG01117510: hypothetical protei |
| AP53_675 | -0.086796603 | 8.237706878 | 0.151164765 | 0.704579484 | 0.9997444 | Dihydrofolate synthase (EC 6.3.2.1 |
| AP53_695 | -0.075100884 | 8.830345711 | 0.150265366 | 0.705413017 | 0.9997444 | SSU ribosomal protein S16p |
| AP53_818 | 0.038914511 | 9.40215327 | 0.150164483 | 0.705506693 | 0.9997444 | Putative membrane protein YeiH |
| AP53_1423 | 0.083941758 | 8.381643768 | 0.149690626 | 0.705947186 | 0.9997444 | Glycerol uptake facilitator protein |
| AP53_204 | -0.048981358 | 6.544829362 | 0.148853255 | 0.706727581 | 0.9997444 | ABC transporter, predicted N-acety |
| AP53_1807 | 0.058740748 | 9.245643543 | 0.148742702 | 0.706830802 | 0.9997444 | Arginyl-tRNA synthetase (EC 6.1.1.1 |
| AP53_510 | 0.05771348 | 5.849857875 | 0.148631427 | 0.706934741 | 0.9997444 | PTS system, N-acetylgalactosamine- |
| AP53_544 | -0.06749271 | 8.092722271 | 0.147850137 | 0.707665804 | 0.9997444 | FIG01114091: hypothetical protei |
| AP53_643 | -0.100379086 | 4.218836449 | 0.147811137 | 0.707702356 | 0.9997444 | replication initiator protein A |
| AP53_1505 | 0.055406687 | 10.10113094 | 0.147163641 | 0.708310017 | 0.9997444 | GTP-sensing transcriptional pleiotr |
| AP53_1610 | -0.041459791 | 7.599596944 | 0.146775429 | 0.708675087 | 0.9997444 | Type I restriction-modification syst |
| AP53_889 | 0.045741489 | 8.84098912 | 0.146729606 | 0.708718215 | 0.9997444 | Sortase A, LPXTG specific |
| AP53_918 | 0.092851362 | 9.936688656 | 0.1460752 | 0.709334992 | 0.9997444 | [Citrate [pro-3S]-lyase] ligase (EC 6.2 |
| AP53_1372 | 0.07405507 | 9.148815649 | 0.144981637 | 0.71036925 | 0.9997444 | General stress protein 13 |
| AP53_62 | -0.06354542 | 10.56937826 | 0.144932333 | 0.710415986 | 0.9997444 | LSU ribosomal protein L15p (L27Ae |
| AP53_1132 | -0.093066912 | 9.079262359 | 0.144875751 | 0.710469632 | 0.9997444 | Ribonucleotide reductase of class I |
| AP53_438 | 0.075463657 | 5.192022396 | 0.144457188 | 0.710866857 | 0.9997444 | Positive transcriptional regulator, I |
| AP53_578 | 0.092680797 | 8.24742685 | 0.143268425 | 0.711998654 | 0.9997444 | ABC transporter ATP-binding protei |
| AP53_732 | 0.068629621 | 6.437125728 | 0.142911608 | 0.712339428 | 0.9997444 | Thymidylate synthase (EC 2.1.1.45) |
| AP53_161 | 0.053519119 | 6.18135813 | 0.142794279 | 0.71245159 | 0.9997444 | Malonate permease |
| AP53_1127 | 0.051700171 | 9.163542651 | 0.142422925 | 0.712806938 | 0.9997444 | Peptidoglycan N-acetylglucosamin |
| AP53_1791 | -0.138251785 | 5.1115042 | 0.142394949 | 0.71283373 | 0.9997444 | FIG01115767: hypothetical protei |
| AP53_1852 | 0.039461164 | 7.54075747 | 0.141679025 | 0.713520385 | 0.9997444 | FIG002958: hypothetical protein |
| AP53_1156 | -0.037746388 | 7.151800104 | 0.141348034 | 0.713838521 | 0.9997444 | DNA polymerase III delta subunit (E |

| | | | | | | |
|---|---|---|---|---|---|---|
| AP53_316 | 0.089549937 | 9.497923771 | 0.141142959 | 0.714035846 | 0.9997444 | hypothetical |
| AP53_1759 | 0.068388567 | 6.652835082 | 0.141054151 | 0.714121349 | 0.9997444 | Trehalose operon transcriptional re |
| AP53_1251 | -0.090238871 | 6.28830736 | 0.140677673 | 0.714484162 | 0.9997444 | FIG01114576: hypothetical protei |
| AP53_545 | 0.078519798 | 12.02450849 | 0.140597868 | 0.714561142 | 0.9997444 | Enolase (EC 4.2.1.11) |
| AP53_360 | -0.064631414 | 11.50774728 | 0.139951094 | 0.715185951 | 0.9997444 | LSU ribosomal protein L11p (L12e) |
| AP53_107 | 0.042470221 | 8.10523054 | 0.139644681 | 0.715482537 | 0.9997444 | FIG01115182: hypothetical protei |
| AP53_1682 | -0.05607859 | 5.782192653 | 0.13962548 | 0.715501135 | 0.9997444 | Dipeptide transport system permea |
| AP53_250 | 0.032252536 | 8.649442822 | 0.139077549 | 0.716032471 | 0.9997444 | Hydrolase (HAD superfamily), YqeK |
| AP53_16 | -0.088146255 | 9.407115863 | 0.139021015 | 0.71608736 | 0.9997444 | Secreted antigen GbpB/SagA/PcsB, |
| AP53_929 | -0.050334294 | 6.614404055 | 0.138822643 | 0.716280066 | 0.9997444 | ABC transporter ATP-binding protei |
| AP53_rna76 | -0.435775186 | 0.798034033 | 0.138815167 | 0.716287332 | 0.9997444 | tRNA-Asn-GTT |
| AP53_833 | 0.075062062 | 6.204504065 | 0.138661926 | 0.716436308 | 0.9997444 | D-alanyl-D-alanine carboxypeptidas |
| AP53_979 | 0.093611725 | 13.36863302 | 0.13825609 | 0.716831308 | 0.9997444 | General stress protein, Gls24 family |
| AP53_208 | 0.046737524 | 7.245340904 | 0.135804778 | 0.719231441 | 0.9997444 | Sialic acid utilization regulator, Rpi |
| AP53_864 | 0.04444361 | 7.658762239 | 0.135330046 | 0.719699126 | 0.9997444 | NAD kinase (EC 2.7.1.23) |
| AP53_795 | -0.083690377 | 10.58190416 | 0.135074471 | 0.719951296 | 0.9997444 | Acetoin dehydrogenase E1 compon |
| AP53_1429 | 0.06436838 | 7.988638528 | 0.134691763 | 0.720329417 | 0.9997444 | Glycyl-tRNA synthetase beta chain ( |
| AP53_761 | -0.065681288 | 8.406277453 | 0.134107679 | 0.720907687 | 0.9997444 | FIG015049: hypothetical protein |
| AP53_459 | 0.087404389 | 4.324628536 | 0.134083546 | 0.720931611 | 0.9997444 | Mobile element protein |
| AP53_958 | -0.06038049 | 7.312010699 | 0.133786978 | 0.721225809 | 0.9997444 | Phosphate transport system regulat |
| AP53_462 | 0.057342347 | 8.914955832 | 0.132460209 | 0.722546562 | 0.9997444 | Signal recognition particle receptor |
| AP53_1482 | -0.103140794 | 10.11432222 | 0.132343386 | 0.722663216 | 0.9997444 | 3-oxoacyl-[acyl-carrier-protein] syn |
| AP53_1400 | 0.037581605 | 7.909196084 | 0.13213568 | 0.722870769 | 0.9997444 | NAD synthetase (EC 6.3.1.5) |
| AP53_1011 | -0.090411534 | 5.389861213 | 0.131890932 | 0.723115575 | 0.9997444 | Hyaluronate lyase (phage associated |
| AP53_276 | -0.067824807 | 8.425597474 | 0.131522215 | 0.723484869 | 0.9997444 | putative arylalkylamine n-acetyltra |
| AP53_82 | 0.068140682 | 8.333549952 | 0.131315805 | 0.723691861 | 0.9997444 | DNA binding protein, FIG046916 |
| AP53_660 | 0.059366086 | 7.324931732 | 0.131038505 | 0.723970234 | 0.9997444 | membrane protein, related to Actir |
| AP53_1594 | 0.077964898 | 11.13498404 | 0.131033117 | 0.723975646 | 0.9997444 | Fructose-bisphosphate aldolase cla |
| AP53_861 | -0.053437595 | 8.722673962 | 0.130163871 | 0.724850461 | 0.9997444 | Ribose-phosphate pyrophosphokin |
| AP53_rna59 | 0.177858473 | 3.594400094 | 0.129348361 | 0.725674228 | 0.9997444 | tRNA-Gln-TTG |
| AP53_30 | 0.136696684 | 10.05230692 | 0.128573675 | 0.726459502 | 0.9997444 | Phosphoribosylaminoimidazole car |
| AP53_1636 | 0.078766788 | 8.608762784 | 0.127496142 | 0.727556253 | 0.9997444 | 23S rRNA (guanosine-2'-O-) -methyl |
| AP53_18 | -0.081388466 | 4.922451675 | 0.127436828 | 0.727616778 | 0.9997444 | DNA recombination and repair prot |
| AP53_rna5 | -0.184544653 | 4.152167762 | 0.127354971 | 0.727700332 | 0.9997444 | tRNA-Val-TAC |
| AP53_38 | -0.049192747 | 8.772914564 | 0.126091614 | 0.728993772 | 0.9997444 | Acyltransferase family |

| | | | | | | |
|---|---|---|---|---|---|---|
| AP53_1058 | -0.074892473 | 4.464736719 | 0.126039597 | 0.729047184 | 0.9997444 | hypothetical protein |
| AP53_403 | 0.097374956 | 4.275269101 | 0.125983262 | 0.729105046 | 0.9997444 | hypothetical protein |
| AP53_1486 | -0.078749503 | 9.62811879 | 0.125853376 | 0.729238506 | 0.9997444 | Acyl carrier protein |
| AP53_735 | 0.083854246 | 8.638929863 | 0.125396582 | 0.729708494 | 0.9997444 | ATP-dependent Clp protease ATP-bi |
| AP53_1134 | -0.074685156 | 8.551652088 | 0.125198679 | 0.729912414 | 0.9997444 | Ribonucleotide reductase of class Il |
| AP53_1308 | -0.070818843 | 7.466925505 | 0.124457251 | 0.730678016 | 0.9997444 | Streptococcal extracellular nucleas |
| AP53_638 | -0.080393148 | 3.705966894 | 0.124448893 | 0.730686662 | 0.9997444 | FIG01116946: hypothetical protei |
| AP53_333 | -0.090333222 | 7.235476923 | 0.123500044 | 0.731670267 | 0.9997444 | Ribonucleotide reductase of class Il |
| AP53_1495 | 0.060265897 | 9.023869357 | 0.123305363 | 0.731872608 | 0.9997444 | D-alanyl-D-alanine carboxypeptidas |
| AP53_1832 | 0.03472122 | 8.819021249 | 0.123296985 | 0.73188132 | 0.9997444 | Replicative DNA helicase (DnaB) (EC |
| AP53_467 | -0.047071515 | 8.671118333 | 0.122316566 | 0.732903113 | 0.9997444 | ABC transporter, substrate-binding |
| AP53_829 | -0.076810739 | 10.09032713 | 0.122072346 | 0.733158361 | 0.9997444 | Acetylornithine deacetylase/Succir |
| AP53_521 | -0.045033185 | 8.920122241 | 0.12203121 | 0.733201383 | 0.9997444 | Cell division protein FtsX |
| AP53_1416 | -0.073422173 | 9.717240124 | 0.121934756 | 0.73330229 | 0.9997444 | ABC transporter permease protein |
| AP53_1858 | -0.06525118 | 7.645699048 | 0.121406536 | 0.733855703 | 0.9997444 | hypothetical protein BH3604 |
| AP53_789 | -0.078689104 | 5.972167453 | 0.119533033 | 0.735829612 | 0.9997444 | ABC transporter ATP-binding protei |
| AP53_106 | -0.042051683 | 7.294570993 | 0.119370995 | 0.736001152 | 0.9997444 | NPQTN specific sortase B |
| AP53_81 | -0.101689739 | 9.263004421 | 0.119246851 | 0.736132665 | 0.9997444 | DNA-directed RNA polymerase beta |
| AP53_1002 | 0.110855081 | 9.186568837 | 0.119149156 | 0.736236213 | 0.9997444 | Streptococcal phospholipase A2; _ |
| AP53_1512 | -0.036814531 | 8.214941548 | 0.118824996 | 0.736580139 | 0.9997444 | ATP-dependent DNA helicase RecG ( |
| AP53_500 | 0.033543364 | 8.026905762 | 0.118178937 | 0.737267172 | 0.9997444 | tRNA-dependent lipid II--L-alanine l |
| AP53_313 | 0.070834751 | 9.55902715 | 0.118146546 | 0.737301673 | 0.9997444 | Thymidylate kinase (EC 2.7.4.9) |
| AP53_1393 | 0.037004972 | 10.5883986 | 0.118015694 | 0.737441103 | 0.9997444 | Hypothetical protein SPy1643 |
| AP53_1723 | 0.046945617 | 8.835087518 | 0.117267536 | 0.738239984 | 0.9997444 | Transcriptional regulator, DeoR fam |
| AP53_443 | 0.081601967 | 4.271592966 | 0.116961649 | 0.738567436 | 0.9997444 | Archaeal S-adenosylmethionine syr |
| AP53_406 | -0.045790884 | 6.982701596 | 0.116780164 | 0.738761944 | 0.9997444 | Formamidopyrimidine-DNA glycos |
| AP53_109 | -0.099558856 | 6.207061618 | 0.116460248 | 0.73910523 | 0.9997444 | Fibronectin-binding protein |
| AP53_432 | -0.037688652 | 7.462376129 | 0.116374166 | 0.739197692 | 0.9997444 | FIG01114264: hypothetical protei |
| AP53_661 | 0.055458888 | 8.138897903 | 0.115934106 | 0.739670964 | 0.9997444 | Tripeptide aminopeptidase (EC 3.4. |
| AP53_1043 | -0.060441248 | 6.060397034 | 0.115719483 | 0.739902152 | 0.9997444 | FIG01115915: hypothetical protei |
| AP53_348 | 0.059787795 | 8.534545207 | 0.115639753 | 0.739988098 | 0.9997444 | N-acetylglucosamine-1-phosphate |
| AP53_809 | -0.043712534 | 7.618084778 | 0.115549759 | 0.740085146 | 0.9997444 | COG4478, integral membrane prot |
| AP53_1419 | -0.05250102 | 9.811227587 | 0.115081931 | 0.740590336 | 0.9997444 | Transketolase (EC 2.2.1.1) |
| AP53_1233 | 0.073919028 | 12.28677147 | 0.114974533 | 0.740706475 | 0.9997444 | ATP-dependent Clp protease, ATP-b |
| AP53_1300 | -0.063202746 | 9.157013082 | 0.114851341 | 0.740839767 | 0.9997444 | Beta-glucosidase (EC 3.2.1.21) |

| | | | | | | |
|---|---|---|---|---|---|---|
| AP53_1174 | 0.16389184 | 3.571063544 | 0.114765981 | 0.740932172 | 0.9997444 | FIG01117945: hypothetical protei |
| AP53_140 | 0.058938023 | 9.852723686 | 0.114676981 | 0.74102856 | 0.9997444 | Leucyl-tRNA synthetase (EC 6.1.1.4) |
| AP53_519 | 0.03136856 | 8.866446243 | 0.114464445 | 0.741258906 | 0.9997444 | Peptide chain release factor 2; prog |
| AP53_514 | 0.042997023 | 6.573820684 | 0.114399056 | 0.741329824 | 0.9997444 | predicted 4-deoxy-L-threo-5-hexos |
| AP53_1136 | -0.097861496 | 5.191885985 | 0.112862579 | 0.743002791 | 0.9997444 | Mobile element protein |
| AP53_480 | 0.076282613 | 5.721513167 | 0.112227191 | 0.743698355 | 0.9997444 | FIG01114033: hypothetical protei |
| AP53_1435 | 0.04765707 | 8.804048446 | 0.111487816 | 0.744510539 | 0.9997444 | FIG01116130: hypothetical protei |
| AP53_343 | -0.036850516 | 6.997685949 | 0.111388613 | 0.74461974 | 0.9997444 | FIG01115464: hypothetical protei |
| AP53_1648 | -0.071363494 | 11.11452243 | 0.111278911 | 0.744740561 | 0.9997444 | SSU ribosomal protein S15p (S13e) |
| AP53_566 | 0.087773287 | 10.8465092 | 0.111227451 | 0.744797261 | 0.9997444 | ATP synthase epsilon chain (EC 3.6. |
| AP53_28 | 0.120567312 | 9.347749304 | 0.111163421 | 0.74486783 | 0.9997444 | Phosphoribosylamine--glycine ligas |
| AP53_1521 | 0.052175363 | 7.131119651 | 0.111143888 | 0.744889363 | 0.9997444 | Cell surface protein Shp, transfers h |
| AP53_1404 | -0.058261326 | 6.130712776 | 0.11057094 | 0.745521903 | 0.9997444 | Hypothetical protein SPy1656 |
| AP53_1179 | 0.034544851 | 7.530511432 | 0.110395325 | 0.745716152 | 0.9997444 | Streptococcal extracellular nucleas |
| AP53_pseudo399 | -0.069822364 | 4.327962628 | 0.109548924 | 0.74665479 | 0.9997444 | FIG01114499: hypothetical protei |
| AP53_983 | -0.048658026 | 8.231438178 | 0.108687866 | 0.747613852 | 0.9997444 | ATP-dependent DNA helicase UvrD/ |
| AP53_976 | 0.032428973 | 9.357629102 | 0.108488434 | 0.747836587 | 0.9997444 | Transcriptional regulator, GntR fam |
| AP53_1685 | 0.052395089 | 6.368726164 | 0.107920135 | 0.748472549 | 0.9997444 | Dipeptide transport ATP-binding pr |
| AP53_401 | -0.068429964 | 4.320376361 | 0.107545816 | 0.748892457 | 0.9997444 | FIG01114077: hypothetical protei |
| AP53_627 | -0.108422111 | 3.230233061 | 0.106675607 | 0.749871811 | 0.9997444 | hypothetical phage protein |
| AP53_1315 | -0.086149372 | 3.418809219 | 0.106638798 | 0.749913335 | 0.9997444 | FIG01116522: hypothetical protei |
| AP53_1053 | 0.084748392 | 7.111292246 | 0.1064075 | 0.750174442 | 0.9997444 | conserved hypothetical protein - ph |
| AP53_1731 | 0.0568599 | 8.042599518 | 0.106288804 | 0.750308558 | 0.9997444 | Hypothetical protein SPy2065 |
| AP53_558 | 0.055256254 | 7.727189147 | 0.106123332 | 0.750495667 | 0.9997444 | Transcription regulator [contains d |
| AP53_1530 | 0.083337087 | 7.232502317 | 0.10599407 | 0.750641944 | 0.9997444 | Fructokinase (EC 2.7.1.4) |
| AP53_804 | 0.06830239 | 8.79923924 | 0.105896745 | 0.750752146 | 0.9997444 | Phosphoglucosamine mutase (EC 5. |
| AP53_1280 | 0.049396407 | 9.75997479 | 0.105820512 | 0.750838504 | 0.9997444 | Valyl-tRNA synthetase (EC 6.1.1.9) |
| AP53_668 | -0.056888748 | 10.80527704 | 0.105762676 | 0.750904045 | 0.9997444 | LSU ribosomal protein L20p |
| AP53_1302 | -0.059716145 | 7.945785161 | 0.104828718 | 0.751965202 | 0.9997444 | Transcriptional repressor of arabin |
| AP53_84 | 0.064367017 | 3.193047419 | 0.104393903 | 0.752461032 | 0.9997444 | Late competence protein ComGB, a |
| AP53_305 | 0.066885147 | 6.571725935 | 0.104263346 | 0.752610133 | 0.9997444 | Ferrichrome-binding periplasmic p |
| AP53_1341 | -0.125747571 | 4.226506109 | 0.103579281 | 0.753393064 | 0.9997444 | Phage protein |
| AP53_12 | -0.077316474 | 7.130205953 | 0.103562302 | 0.753412534 | 0.9997444 | tRNA(Ile)-lysidine synthetase |
| AP53_43 | 0.097713418 | 10.88048974 | 0.102387901 | 0.754763524 | 0.9997444 | LSU ribosomal protein L3p (L3e) |
| AP53_1358 | 0.071030274 | 13.07681823 | 0.102251865 | 0.75492057 | 0.9997444 | hypothetical protein - phage associ |

| | | | | | | |
|---|---|---|---|---|---|---|
| AP53_484 | -0.040576084 | 6.791743853 | 0.101525233 | 0.755761396 | 0.9997444 | 2-haloalkanoic acid dehalogenase ( |
| AP53_1230 | 0.074996475 | 7.713718669 | 0.101221929 | 0.756113354 | 0.9997444 | Glutamate transport ATP-binding p |
| AP53_255 | -0.048527178 | 9.625319323 | 0.100965251 | 0.756411663 | 0.9997444 | Cysteine ABC transporter, substrate |
| AP53_1570 | 0.048826794 | 7.812768783 | 0.10094451 | 0.756435787 | 0.9997444 | Transcriptional regulatory protein |
| AP53_515 | 0.043230735 | 6.975175126 | 0.100726955 | 0.756688986 | 0.9997444 | 2-dehydro-3-deoxygluconate kinas |
| AP53_1065 | 0.060736352 | 5.632716854 | 0.100269681 | 0.757222169 | 0.9997444 | Maltose operon transcriptional rep |
| AP53_674 | 0.033634866 | 8.27484944 | 0.100167428 | 0.757341581 | 0.9997444 | Glutathione reductase (EC 1.8.1.7) |
| AP53_1523 | 0.059202709 | 4.841995829 | 0.099934111 | 0.757614304 | 0.9997444 | hypothetical protein |
| AP53_658 | 0.066352723 | 6.883404077 | 0.099588462 | 0.758018979 | 0.9997444 | Membrane protein |
| AP53_1531 | 0.058450956 | 9.761708956 | 0.098586314 | 0.759196676 | 0.9997444 | Secreted Endo-beta-N-acetylglucos |
| AP53_873 | 0.087647012 | 8.200934288 | 0.097963488 | 0.759931945 | 0.9997444 | Xanthine permease |
| AP53_815 | 0.049679438 | 8.798029072 | 0.097940044 | 0.759959672 | 0.9997444 | Translation elongation factor LepA |
| AP53_1554 | -0.05397394 | 8.751635964 | 0.097821073 | 0.760100434 | 0.9997444 | Ribonuclease HIII (EC 3.1.26.4) |
| AP53_858 | 0.03578346 | 9.390154202 | 0.097813554 | 0.760109333 | 0.9997444 | Redox-sensitive transcriptional reg |
| AP53_1075 | -0.059746919 | 9.355194238 | 0.097699822 | 0.76024399 | 0.9997444 | D-alanyl transfer protein DltB |
| AP53_1260 | 0.043015963 | 8.863973916 | 0.09756566 | 0.760402949 | 0.9997444 | 16S rRNA (guanine(966)-N(2))-meth |
| AP53_931 | 0.050875312 | 6.177726661 | 0.097334741 | 0.760676829 | 0.9997444 | Pyridoxal kinase (EC 2.7.1.35) |
| AP53_731 | 0.052173578 | 7.025910352 | 0.096942117 | 0.761143325 | 0.9997444 | Hydroxymethylglutaryl-CoA syntha |
| AP53_293 | -0.038214047 | 7.345830115 | 0.096933879 | 0.761153124 | 0.9997444 | Protein YidD |
| AP53_615 | -0.125303632 | 5.69586088 | 0.096658143 | 0.76148138 | 0.9997444 | hypothetical phage protein |
| AP53_1303 | 0.067384069 | 8.524657579 | 0.096258179 | 0.761958443 | 0.9997444 | hypothetical protein |
| AP53_1074 | -0.06487214 | 9.483726773 | 0.096253252 | 0.761964327 | 0.9997444 | D-alanine--poly(phosphoribitol) lig |
| AP53_356 | -0.046643585 | 8.106574505 | 0.096219539 | 0.76200459 | 0.9997444 | Manganese ABC transporter, inner |
| AP53_752 | -0.034627178 | 9.431559898 | 0.095226448 | 0.763194133 | 0.9997444 | Cystine transport system permease |
| AP53_249 | 0.058283684 | 8.951446759 | 0.094729736 | 0.763791671 | 0.9997444 | Nicotinate-nucleotide adenylyltran |
| AP53_263 | 0.03009639 | 6.84634355 | 0.094433278 | 0.764149129 | 0.9997444 | rRNA small subunit 7-methylguano |
| AP53_771 | -0.050229491 | 9.030765767 | 0.094317215 | 0.764289242 | 0.9997444 | Short chain dehydrogenase |
| AP53_448 | -0.140032551 | 1.098269493 | 0.094125188 | 0.764521268 | 0.9997444 | FIG01115329: hypothetical protei |
| AP53_1863 | 0.035688488 | 10.1139576 | 0.093651907 | 0.765094249 | 0.9997444 | Serine protease, DegP/HtrA, do-like |
| AP53_338 | -0.089895214 | 12.83272165 | 0.092624818 | 0.766343209 | 0.9997444 | FIG01114323: hypothetical protei |
| AP53_1237 | -0.043127672 | 9.540523851 | 0.092573405 | 0.766405929 | 0.9997444 | Cell division initiation protein Div |
| AP53_1246 | -0.0308088 | 8.444518672 | 0.092483384 | 0.766515792 | 0.9997444 | UDP-N-acetylmuramoylalanine--D- |
| AP53_1184 | -0.086681585 | 3.96316391 | 0.092160505 | 0.766910323 | 0.9997444 | Phage protein |
| AP53_1307 | -0.093526326 | 4.02732108 | 0.092023462 | 0.767078009 | 0.9997444 | Paratox |
| AP53_1210 | -0.158376215 | 1.936449278 | 0.091921445 | 0.767202924 | 0.9997444 | Phage protein |

| | | | | | | |
|---|---|---|---|---|---|---|
| AP53_1514 | -0.036258612 | 7.701027762 | 0.091879458 | 0.767254358 | 0.9997444 | Transmembrane component BL069 |
| AP53_1079 | 0.057142419 | 6.775970913 | 0.091496951 | 0.767723524 | 0.9997444 | Glutamine ABC transporter, glutam |
| AP53_635 | -0.067826452 | 3.98579451 | 0.091491037 | 0.767730786 | 0.9997444 | conserved hypothetical protein - ph |
| AP53_1481 | -0.077390293 | 10.0964202 | 0.091455536 | 0.767774385 | 0.9997444 | Biotin carboxyl carrier protein of ac |
| AP53_991 | 0.056048997 | 11.37384296 | 0.091108065 | 0.768201619 | 0.9997444 | Glucosamine--fructose-6-phosphat |
| AP53_1696 | 0.037702254 | 8.493555226 | 0.090652139 | 0.768763563 | 0.9997444 | M protein trans-acting positive regu |
| AP53_1654 | -0.042264476 | 7.992457834 | 0.090167999 | 0.769361981 | 0.9997444 | DNA polymerase III alpha subunit (E |
| AP53_1489 | -0.072702297 | 10.06214315 | 0.090057092 | 0.769499314 | 0.9997444 | Trans-2,cis-3-Decenoyl-ACP isomera |
| AP53_1152 | 0.040143861 | 7.02683456 | 0.089235988 | 0.770518971 | 0.9997444 | FIG01113992: hypothetical protein |
| AP53_1088 | 0.055852469 | 4.336772951 | 0.088651228 | 0.771248272 | 0.9997444 | Beta-glucosidase (EC 3.2.1.21) |
| AP53_977 | 0.059810416 | 13.251696 | 0.088393478 | 0.771570569 | 0.9997444 | General stress protein, Gls24 family |
| AP53_1666 | -0.064937898 | 9.740172386 | 0.088178979 | 0.771839177 | 0.9997444 | D-tyrosyl-tRNA(Tyr) deacylase |
| AP53_33 | 0.04231312 | 9.672083697 | 0.087071467 | 0.773231792 | 0.9997444 | Adenylosuccinate lyase (EC 4.3.2.2) |
| AP53_610 | 0.084043803 | 4.371625699 | 0.086550001 | 0.773890854 | 0.9997444 | Phage tail length tape-measure prot |
| AP53_1857 | -0.096454498 | 7.917900341 | 0.085912201 | 0.774699903 | 0.9997444 | FIG01116071: hypothetical protein |
| AP53_1082 | -0.071109169 | 5.178836739 | 0.085710677 | 0.774956218 | 0.9997444 | PTS system, cellobiose-specific IIC c |
| AP53_166 | -0.030862202 | 7.535923888 | 0.085160807 | 0.775657267 | 0.9997444 | membrane protein |
| AP53_307 | -0.051610624 | 7.624681867 | 0.084545026 | 0.77644529 | 0.9997444 | UDP-N-acetylmuramoylalanyl-D-glu |
| AP53_1021 | -0.144068484 | 6.68383023 | 0.084288494 | 0.7767745 | 0.9997444 | DNA packaging protein, phage assoc |
| AP53_25 | 0.076529974 | 9.107144995 | 0.084285824 | 0.776777929 | 0.9997444 | Phosphoribosylglycinamide formyl |
| AP53_1836 | 0.049050033 | 7.000800069 | 0.084261176 | 0.77680959 | 0.9997444 | MutT/nudix family protein |
| AP53_437 | 0.057551232 | 6.32848962 | 0.084242381 | 0.776833735 | 0.9997444 | Chromosome partition protein smc |
| AP53_rna30 | 0.099040364 | 3.832576393 | 0.083918008 | 0.777250916 | 0.9997444 | tRNA-Met-CAT |
| AP53_1576 | -0.038573718 | 8.093332469 | 0.083346278 | 0.777988371 | 0.9997444 | Nucleoside permease NupC |
| AP53_534 | -0.065092872 | 7.793605183 | 0.083120519 | 0.778280329 | 0.9997444 | Galactosamine-6-phosphate isomer |
| AP53_1076 | -0.054648229 | 8.966213956 | 0.083013435 | 0.778418965 | 0.9997444 | D-alanine--poly(phosphoribitol) lig |
| AP53_745 | -0.085659903 | 5.133878848 | 0.082914238 | 0.778547476 | 0.9997444 | hypothetical protein |
| AP53_358 | -0.029586272 | 7.203562096 | 0.082132237 | 0.779563518 | 0.9997444 | Cell division protein FtsK |
| AP53_840 | -0.057165906 | 8.092291269 | 0.082027468 | 0.779700042 | 0.9997444 | UDP-N-acetylenolpyruvoylglucosar |
| AP53_1200 | 0.061661031 | 3.627071698 | 0.080967555 | 0.78108658 | 0.9997444 | hypothetical protein |
| AP53_816 | 0.054441814 | 13.52738783 | 0.080838267 | 0.781256384 | 0.9997444 | Collagen-like surface protein |
| AP53_271 | -0.028744851 | 9.333956541 | 0.080204418 | 0.782091009 | 0.9997444 | Helicase loader DnaI |
| AP53_1323 | -0.100686813 | 2.327562017 | 0.079910136 | 0.782479725 | 0.9997444 | Phage major tail shaft protein |
| AP53_1453 | -0.054042831 | 10.39467819 | 0.079709119 | 0.782745695 | 0.9997444 | Ribosome-binding factor A |
| AP53_1811 | -0.031178426 | 7.932330972 | 0.079615016 | 0.78287033 | 0.9997444 | Integral membrane protein |

| | | | | | | |
|---|---|---|---|---|---|---|
| AP53_238 | 0.028111454 | 9.458554412 | 0.07945848 | 0.783077832 | 0.9997444 | Oligopeptide ABC transporter, peri |
| AP53_359 | -0.081098089 | 7.623428763 | 0.078757056 | 0.784010364 | 0.9997444 | Integral membrane protein |
| AP53_pseudo595 | 0.042844311 | 6.325910274 | 0.078751606 | 0.784017627 | 0.9997444 | Lipoteichoic acid synthase LtaS Typ |
| AP53_rna22 | 0.146189518 | 4.187408477 | 0.0779409 | 0.78510112 | 0.9997444 | tRNA-Thr-TGT |
| AP53_1007 | -0.109959675 | 5.788601678 | 0.077749982 | 0.785357166 | 0.9997444 | Phage holin |
| AP53_1558 | -0.034882808 | 7.796647122 | 0.077747104 | 0.785361029 | 0.9997444 | DNA polymerase IV (EC 2.7.7.7) |
| AP53_1822 | -0.026223429 | 7.315602649 | 0.077594247 | 0.78556628 | 0.9997444 | FIG028593: membrane protein |
| AP53_620 | -0.098385162 | 4.71110218 | 0.077187092 | 0.786114065 | 0.9997444 | Prophage Clp protease-like protein |
| AP53_1144 | 0.048149552 | 6.785620853 | 0.077119817 | 0.786204728 | 0.9997444 | O-methyltransferase family protein |
| AP53_945 | -0.033131422 | 7.848707534 | 0.077113569 | 0.78621315 | 0.9997444 | Substrate-specific component PanT |
| AP53_rna44 | 0.124398845 | 3.465197914 | 0.076874506 | 0.786535681 | 0.9997444 | tRNA-Met-CAT |
| AP53_1517 | -0.031011301 | 7.018966461 | 0.076792167 | 0.786646895 | 0.9997444 | Putative ABC transporter (ATP-bind |
| AP53_999 | -0.040131845 | 7.754957157 | 0.076434526 | 0.787130701 | 0.9997444 | Immunoreactive protein Se23.5 (Fr |
| AP53_1321 | -0.101506689 | 2.997535762 | 0.076432422 | 0.787133552 | 0.9997444 | Phage protein |
| AP53_273 | 0.031665342 | 7.785493392 | 0.076014638 | 0.787700282 | 0.9997444 | SWF/SNF family helicase |
| AP53_1754 | -0.0359885 | 11.52845548 | 0.075971791 | 0.787758499 | 0.9997444 | SSU ribosomal protein S2p (SAe) |
| AP53_6 | -0.054897095 | 5.955464852 | 0.075188859 | 0.78882544 | 0.9997444 | Peptidyl-tRNA hydrolase (EC 3.1.1.2 |
| AP53_1504 | -0.041895069 | 6.423323182 | 0.074920225 | 0.789192905 | 0.9997444 | Nicotinamidase (EC 3.5.1.19) |
| AP53_1630 | -0.066772384 | 11.41737697 | 0.074389479 | 0.789921011 | 0.9997444 | SSU ribosomal protein S9p (S16e) |
| AP53_1778 | -0.061288064 | 9.281653972 | 0.074358242 | 0.789963951 | 0.9997444 | DNA-3-methyladenine glycosylase ( |
| AP53_593 | -0.03685522 | 6.727169012 | 0.074350222 | 0.789974978 | 0.9997444 | putative glycosyltransferase - possil |
| AP53_439 | -0.083092289 | 2.385210709 | 0.073787129 | 0.790750756 | 0.9997444 | Shikimate 5-dehydrogenase I alpha |
| AP53_1485 | -0.067339136 | 9.463774976 | 0.073528479 | 0.791108171 | 0.9997444 | Enoyl-[acyl-carrier-protein] reducta |
| AP53_436 | 0.080161806 | 5.781994632 | 0.073439813 | 0.791230852 | 0.9997444 | Ribonuclease III (EC 3.1.26.3) |
| AP53_1377 | -0.033057479 | 10.53840346 | 0.073431174 | 0.791242808 | 0.9997444 | Serine/threonine protein kinase Prl |
| AP53_289 | -0.032032046 | 7.004993998 | 0.073231978 | 0.791518727 | 0.9997444 | Tyrosine recombinase XerD |
| AP53_749 | 0.070953204 | 8.761878754 | 0.072919645 | 0.791952176 | 0.9997444 | Orotate phosphoribosyltransferase |
| AP53_774 | 0.027591893 | 8.855491958 | 0.072895196 | 0.791986149 | 0.9997444 | Chromosome replication initiation |
| AP53_672 | 0.025023064 | 8.457220619 | 0.072816532 | 0.792095495 | 0.9997444 | Chorismate synthase (EC 4.2.3.5) |
| AP53_1669 | -0.029374152 | 7.375156326 | 0.072646805 | 0.792331638 | 0.9997444 | Ribonucleotide reduction protein I |
| AP53_8 | 0.062205297 | 7.292519567 | 0.071233659 | 0.794309428 | 0.9997444 | Ribosome-associated heat shock pr |
| AP53_754 | 0.051677823 | 9.268599668 | 0.071190137 | 0.794370674 | 0.9997444 | Dihydroorotase (EC 3.5.2.3) |
| AP53_641 | -0.054292908 | 4.573462804 | 0.070177443 | 0.79580151 | 0.9997444 | hypothetical phage protein |
| AP53_875 | -0.074170883 | 7.024084202 | 0.070069827 | 0.795954211 | 0.9997444 | 4-oxalocrotonate tautomerase (EC |
| AP53_491 | 0.032082403 | 7.708053578 | 0.069344032 | 0.79698739 | 0.9997444 | Glutathione peroxidase family prot |

| | | | | | | |
|---|---|---|---|---|---|---|
| AP53_1818 | -0.04893049 | 5.452001863 | 0.06928639 | 0.797069692 | 0.9997444 | FIG01115119: hypothetical protei |
| AP53_487 | 0.050019787 | 9.432426044 | 0.06906079 | 0.797392164 | 0.9997444 | Substrate-specific component ThiT |
| AP53_254 | -0.052140455 | 8.141883276 | 0.069005527 | 0.797471243 | 0.9997444 | FIG007491: hypothetical protein Y |
| AP53_1397 | -0.037085399 | 8.742721169 | 0.068600715 | 0.798051552 | 0.9997444 | RecU Holliday junction resolvase |
| AP53_210 | -0.027708039 | 8.282924267 | 0.067811802 | 0.799187789 | 0.9997444 | Ribonuclease M5 (EC 3.1.26.8) |
| AP53_722 | 0.058487997 | 7.447617739 | 0.067444721 | 0.799718898 | 0.9997444 | 5'-nucleotidase (EC 3.1.3.5) |
| AP53_866 | 0.070521829 | 10.61348664 | 0.066987409 | 0.800382731 | 0.9997444 | Phosphate acetyltransferase (EC 2.3 |
| AP53_339 | 0.025552121 | 6.952917524 | 0.066567799 | 0.800993978 | 0.9997444 | Mobile element protein |
| AP53_1845 | -0.027437929 | 8.383367852 | 0.066191033 | 0.801544575 | 0.9997444 | Transcriptional regulator in cluster |
| AP53_1660 | 0.087298195 | 8.781260688 | 0.066104533 | 0.801671222 | 0.9997444 | putative transport accessory protei |
| AP53_1045 | -0.04779005 | 6.293677744 | 0.065987596 | 0.801842573 | 0.9997444 | hypothetical phage protein |
| AP53_247 | 0.038346024 | 8.868964733 | 0.065503206 | 0.802554099 | 0.9997444 | GTP-binding protein YqeH, require |
| AP53_286 | -0.032996922 | 8.349778934 | 0.065249605 | 0.80292774 | 0.9997444 | unknown domain / Nucleoside 5-tr |
| AP53_1410 | 0.036436031 | 9.086127775 | 0.064534545 | 0.803985475 | 0.9997444 | Cell division protein FtsI [Peptidogl |
| AP53_rna41 | 0.183483104 | 4.30478976 | 0.063848179 | 0.805006677 | 0.9997444 | tRNA-Ile-GAT |
| AP53_341 | -0.06733767 | 3.9403568 | 0.063764942 | 0.805130918 | 0.9997444 | hypothetical protein |
| AP53_1073 | -0.043108428 | 9.28714035 | 0.063601354 | 0.805375347 | 0.9997444 | Poly(glycerophosphate chain) D-ala |
| AP53_572 | 0.048116442 | 8.917857401 | 0.063597505 | 0.805381101 | 0.9997444 | DNA-entry nuclease (Competence-s |
| AP53_1533 | 0.034242696 | 8.145137342 | 0.063359063 | 0.805737988 | 0.9997444 | Sucrose-6-phosphate hydrolase (EC |
| AP53_726 | -0.035298075 | 7.212955973 | 0.062978473 | 0.806309124 | 0.9997444 | Mevalonate kinase (EC 2.7.1.36) |
| AP53_1556 | 0.059683766 | 8.243803113 | 0.062964162 | 0.806330637 | 0.9997444 | RecD-like DNA helicase YrrC |
| AP53_156 | 0.083774607 | 10.26429078 | 0.062839365 | 0.806518337 | 0.9997444 | Co-activator of prophage gene expr |
| AP53_582 | -0.02437761 | 8.124503157 | 0.062651309 | 0.806801562 | 0.9997444 | Amino acid ABC transporter, aminc |
| AP53_1519 | 0.030495239 | 7.460775577 | 0.06159251 | 0.808404711 | 0.9997444 | Heme ABC transporter (Streptococ |
| AP53_943 | 0.035241561 | 7.179684337 | 0.061368442 | 0.808745857 | 0.9997444 | Phosphopantothenoylcysteine synt |
| AP53_580 | -0.028733483 | 7.556711597 | 0.060895735 | 0.809467745 | 0.9997444 | ATP-dependent nuclease, subunit B |
| AP53_1542 | -0.030829025 | 9.030240073 | 0.060491005 | 0.810088199 | 0.9997444 | Integral membrane protein |
| AP53_1718 | 0.033841476 | 6.728872894 | 0.06038142 | 0.810256573 | 0.9997444 | Pyruvate formate-lyase (EC 2.3.1.54 |
| AP53_rna63 | -0.105038801 | 2.769722909 | 0.060308972 | 0.810367977 | 0.9997444 | tRNA-Ile-GAT |
| AP53_80 | -0.076658511 | 8.787198023 | 0.060164231 | 0.810590762 | 0.9997444 | DNA-directed RNA polymerase beta |
| AP53_1164 | -0.042147011 | 9.395850668 | 0.060095285 | 0.810696982 | 0.9997444 | Peptide chain release factor 3 |
| AP53_739 | -0.055927832 | 10.05085262 | 0.059995433 | 0.810850934 | 0.9997444 | Ribose 5-phosphate isomerase A (E |
| AP53_1790 | 0.084570743 | 4.848679332 | 0.059895334 | 0.811005403 | 0.9997444 | FIG01118442: hypothetical protei |
| AP53_703 | 0.055809071 | 6.57625008 | 0.05981036 | 0.81113664 | 0.9997444 | Thioredoxin reductase (EC 1.8.1.9) |
| AP53_1388 | -0.05372578 | 7.381478696 | 0.059669888 | 0.811353808 | 0.9997444 | Butyrate-acetoacetate CoA-transfer |

| | | | | | | |
|---|---|---|---|---|---|---|
| AP53_1094 | 0.092838475 | 4.265203367 | 0.059468843 | 0.811665094 | 0.9997444 | Mobile element protein |
| AP53_176 | 0.062297804 | 5.921850499 | 0.059413232 | 0.811751297 | 0.9997444 | hypothetical cytosolic protein |
| AP53_1743 | -0.080429126 | 10.51698796 | 0.059317284 | 0.811900129 | 0.9997444 | Alkyl hydroperoxide reductase prot |
| AP53_5 | -0.030036752 | 8.003714405 | 0.059078276 | 0.81227143 | 0.9997444 | GTP-binding and nucleic acid-bindi |
| AP53_1071 | 0.0507744 | 7.377320043 | 0.058832256 | 0.81265446 | 0.9997444 | Transposase (maltose area) |
| AP53_607 | -0.060206438 | 5.266162481 | 0.05826977 | 0.813533408 | 0.9997444 | Hyaluronate lyase (phage associatec |
| AP53_1591 | 0.029738745 | 10.34080977 | 0.058162539 | 0.81370148 | 0.9997444 | Dihydroxyacetone kinase family prc |
| AP53_1809 | -0.028108132 | 7.846217249 | 0.057819024 | 0.81424101 | 0.9997444 | Transporter |
| AP53_584 | -0.072474535 | 10.15155159 | 0.05742979 | 0.81485441 | 0.9997444 | Large-conductance mechanosensiti |
| AP53_753 | 0.035904739 | 8.807008081 | 0.057382347 | 0.814929327 | 0.9997444 | Uracil-DNA glycosylase, family 1 |
| AP53_759 | 0.076021166 | 8.73137515 | 0.057162914 | 0.81527626 | 0.9997444 | FIG01114490: hypothetical protei |
| AP53_1274 | 0.053716885 | 6.536350653 | 0.056684465 | 0.81603517 | 0.9997444 | Peptide methionine sulfoxide redu |
| AP53_762 | 0.065268262 | 9.11980704 | 0.056464561 | 0.81638512 | 0.9997444 | Rhodanese domain protein UPF017 |
| AP53_1812 | -0.038274407 | 8.060183411 | 0.056282115 | 0.816676008 | 0.9997444 | Aspartyl-tRNA synthetase (EC 6.1.1. |
| AP53_1604 | -0.033147439 | 7.401009618 | 0.056164266 | 0.81686417 | 0.9997444 | Novel pyridoxal kinase, thiD family |
| AP53_1106 | -0.025855203 | 7.140266528 | 0.055986695 | 0.817148083 | 0.9997444 | Uridine phosphorylase (EC 2.4.2.3) |
| AP53_1287 | 0.03323192 | 8.933424055 | 0.055564229 | 0.817825476 | 0.9997444 | Uncharacterized conserved protein |
| AP53_67 | -0.066325643 | 10.20321776 | 0.05492959 | 0.818848225 | 0.9997444 | SSU ribosomal protein S13p (S18e) |
| AP53_382 | 0.078965722 | 5.415796632 | 0.054899725 | 0.818896508 | 0.9997444 | FIG01119121: hypothetical protei |
| AP53_149 | -0.059532043 | 11.85960658 | 0.05461702 | 0.819354246 | 0.9997444 | Glycine betaine ABC transport syste |
| AP53_564 | 0.064931268 | 10.08214382 | 0.054605623 | 0.819372726 | 0.9997444 | ATP synthase gamma chain (EC 3.6.: |
| AP53_1168 | 0.021343556 | 7.247266685 | 0.054476282 | 0.819582589 | 0.9997444 | Recombination protein RecR |
| AP53_1715 | 0.034966661 | 6.006493938 | 0.054469879 | 0.819592985 | 0.9997444 | Low temperature requirement C pr |
| AP53_1015 | -0.071490369 | 6.161086585 | 0.054125131 | 0.820153683 | 0.9997444 | Phage protein |
| AP53_571 | 0.061872475 | 7.058893138 | 0.053704944 | 0.820839637 | 0.9997444 | hypothetical protein |
| AP53_330 | 0.050619734 | 10.15185276 | 0.053661844 | 0.820910159 | 0.9997444 | Methionyl-tRNA synthetase (EC 6.1. |
| AP53_1402 | 0.028547122 | 7.975794128 | 0.053551301 | 0.821091169 | 0.9997444 | Histidine transport protein (permea |
| AP53_1099 | 0.041830487 | 5.804455639 | 0.052936812 | 0.822100992 | 0.9997444 | Mobile element protein |
| AP53_361 | -0.04110487 | 12.21068719 | 0.052509209 | 0.822807357 | 0.9997444 | LSU ribosomal protein L1p (L10Ae) |
| AP53_856 | 0.038346982 | 6.912105337 | 0.051865746 | 0.823876057 | 0.9997444 | DNA repair protein RadC |
| AP53_340 | 0.028755694 | 6.586406523 | 0.05172656 | 0.824108145 | 0.9997444 | hypothetical protein |
| AP53_61 | -0.048976257 | 10.31748478 | 0.051654097 | 0.824229104 | 0.9997444 | LSU ribosomal protein L30p (L7e) |
| AP53_960 | 0.031490711 | 7.539641314 | 0.051492146 | 0.824499768 | 0.9997444 | Phosphate transport ATP-binding p |
| AP53_1219 | -0.022020874 | 8.732265628 | 0.051458275 | 0.824556432 | 0.9997444 | Hypothetical protein DUF194, Deg' |
| AP53_205 | -0.029229692 | 6.799939506 | 0.051281294 | 0.824852833 | 0.9997444 | FIG01114161: hypothetical protei |

| | | | | | | |
|---|---|---|---|---|---|---|
| AP53_97 | 0.031578918 | 7.423308075 | 0.050747946 | 0.825749341 | 0.9997444 | Phenylalanyl-tRNA synthetase dom |
| AP53_1087 | 0.045436689 | 4.395822584 | 0.050578913 | 0.826034505 | 0.9997444 | Outer surface protein of unknown f |
| AP53_758 | 0.034444681 | 7.399991744 | 0.050576622 | 0.826038373 | 0.9997444 | Branched-chain amino acid aminot |
| AP53_261 | -0.024755573 | 8.013127346 | 0.050503555 | 0.826161799 | 0.9997444 | Trk system potassium uptake protei |
| AP53_1038 | -0.042484187 | 5.76057865 | 0.050273803 | 0.826550512 | 0.9997444 | hypothetical protein |
| AP53_1052 | -0.039143723 | 7.631196824 | 0.050222662 | 0.826637164 | 0.9997444 | hypothetical protein |
| AP53_1020 | -0.102673176 | 6.170653395 | 0.050160586 | 0.826742406 | 0.9997444 | Phage protein |
| AP53_585 | 0.025187963 | 9.096517146 | 0.050078094 | 0.826882368 | 0.9997444 | DNA primase (EC 2.7.7.-) |
| AP53_1635 | -0.034803737 | 7.260475475 | 0.04987708 | 0.82722393 | 0.9997444 | Hypothetical protein DUF901, simi |
| AP53_1414 | -0.042205256 | 8.473384018 | 0.049856543 | 0.827258868 | 0.9997444 | Gamma-glutamyl phosphate reduct |
| AP53_1477 | -0.057830317 | 10.2050706 | 0.049703134 | 0.827520086 | 0.9997444 | Acetyl-coenzyme A carboxyl transfe |
| AP53_pseudo1615 | 0.060976367 | 3.079259341 | 0.049511407 | 0.827847149 | 0.9997444 | Lanthionine biosynthesis protein La |
| AP53_1537 | -0.036931904 | 10.26151985 | 0.048709222 | 0.829222865 | 0.9997444 | Translation elongation factor P |
| AP53_1044 | -0.043037954 | 6.201115305 | 0.048541202 | 0.829512521 | 0.9997444 | hypothetical phage protein |
| AP53_808 | 0.03979197 | 7.800966263 | 0.048410378 | 0.829738417 | 0.9997444 | Hypothetical NagD-like phosphatas |
| AP53_536 | -0.02464825 | 8.741025202 | 0.047965537 | 0.83050895 | 0.9997444 | FIG146085: 3'-to-5' oligoribonucle |
| AP53_912 | 0.050753897 | 6.658422814 | 0.047634232 | 0.831085265 | 0.9997444 | FIG01114213: hypothetical protein |
| AP53_371 | -0.032406231 | 8.04646574 | 0.047389303 | 0.831512682 | 0.9997444 | Metal-dependent hydrolase YbeY, in |
| AP53_50 | 0.047500995 | 11.20930675 | 0.047228515 | 0.8317939 | 0.9997444 | LSU ribosomal protein L16p (L10e) |
| AP53_579 | -0.030492742 | 7.606143427 | 0.046915543 | 0.832342734 | 0.9997444 | YpfJ protein, zinc metalloprotease s |
| AP53_1761 | -0.074661051 | 4.562062021 | 0.046595648 | 0.832905699 | 0.9997444 | FIG01114717: hypothetical protein |
| AP53_868 | 0.033093225 | 8.622066787 | 0.045980173 | 0.833994579 | 0.9997444 | Na+ driven multidrug efflux pump |
| AP53_948 | -0.053656141 | 8.363698522 | 0.04595276 | 0.834043255 | 0.9997444 | Putative deoxyribose-specific ABC t |
| AP53_1683 | -0.035047679 | 6.01827946 | 0.045640472 | 0.834598845 | 0.9997444 | Dipeptide transport system permea |
| AP53_772 | -0.040527235 | 8.155570193 | 0.045550426 | 0.834759417 | 0.9997444 | Single-stranded-DNA-specific exonu |
| AP53_839 | 0.029330605 | 9.240711618 | 0.045372487 | 0.83507721 | 0.9997444 | 2-amino-4-hydroxy-6-hydroxymeth |
| AP53_777 | 0.019677253 | 8.043231212 | 0.044876394 | 0.835966684 | 0.9997444 | FIG137478: Hypothetical protein |
| AP53_397 | -0.08573465 | 3.124222425 | 0.043444331 | 0.838563622 | 0.9997444 | hypothetical protein |
| AP53_1256 | 0.030686737 | 7.810463382 | 0.043286733 | 0.838852136 | 0.9997444 | FIG01114963: hypothetical protein |
| AP53_pseudo73 | 0.033607748 | 6.753476038 | 0.043251538 | 0.838916641 | 0.9997444 | 4-diphosphocytidyl-2-C-methyl-D-e |
| AP53_841 | -0.032982545 | 8.131143388 | 0.04280108 | 0.839744692 | 0.9997444 | Putrescine transport ATP-binding p |
| AP53_1488 | -0.053270647 | 10.09266506 | 0.042742509 | 0.839852694 | 0.9997444 | Transcriptional regulator of fatty ac |
| AP53_411 | -0.031607136 | 10.31818213 | 0.042604224 | 0.840107992 | 0.9997444 | 3'-to-5' exoribonuclease RNase R |
| AP53_1294 | -0.040604279 | 5.511894944 | 0.042274242 | 0.840718954 | 0.9997444 | Integral membrane protein |
| AP53_309 | -0.03721429 | 8.569034772 | 0.041959996 | 0.841303102 | 0.9997444 | Uracil phosphoribosyltransferase (E |

| | | | | | | |
|---|---|---|---|---|---|---|
| AP53_1401 | 0.018050797 | 7.825276973 | 0.041525909 | 0.842113794 | 0.9997444 | Nicotinate phosphoribosyltransfera |
| AP53_623 | -0.055750718 | 3.912249943 | 0.041472784 | 0.842213313 | 0.9997444 | Phage protein |
| AP53_1178 | -0.073849816 | 6.83398811 | 0.041357569 | 0.842429374 | 0.9997444 | Paratox |
| AP53_1813 | -0.034193972 | 6.865511878 | 0.041052515 | 0.843002961 | 0.9997444 | Histidyl-tRNA synthetase (EC 6.1.1. |
| AP53_1793 | -0.065707866 | 5.339455552 | 0.040902095 | 0.843286611 | 0.9997444 | FIG01114419: hypothetical protei |
| AP53_236 | -0.037522804 | 8.985063389 | 0.04054333 | 0.84396535 | 0.9997444 | D-alanyl-D-alanine carboxypeptidas |
| AP53_546 | -0.047721705 | 7.330619277 | 0.040487068 | 0.844072076 | 0.9997444 | Streptolysin S biosynthesis protein |
| AP53_981 | 0.044092908 | 12.5374196 | 0.040440326 | 0.8441608 | 0.9997444 | FIG01114374: hypothetical protei |
| AP53_1387 | 0.053439209 | 7.519770716 | 0.040045084 | 0.844913191 | 0.9997444 | Butyrate-acetoacetate CoA-transfer |
| AP53_27 | 0.084784322 | 7.905731302 | 0.039934509 | 0.845124377 | 0.9997444 | putative choline binding protein |
| AP53_1838 | 0.034437766 | 6.845123703 | 0.039422691 | 0.846105888 | 0.9997444 | L-serine dehydratase, beta subunit |
| AP53_483 | 0.026022731 | 9.888218567 | 0.039390254 | 0.846168315 | 0.9997444 | Lysyl-tRNA synthetase (class II) (EC 6 |
| AP53_1740 | 0.041332554 | 6.04185596 | 0.039318234 | 0.84630702 | 0.9997444 | hypothetical protein |
| AP53_374 | -0.035135648 | 8.82937416 | 0.039124411 | 0.846680966 | 0.9997444 | MutT/nudix family protein |
| AP53_682 | -0.062821874 | 10.79805948 | 0.038985855 | 0.846948875 | 0.9997444 | LSU ribosomal protein L27p |
| AP53_1318 | -0.060777242 | 4.204982717 | 0.038926216 | 0.847064345 | 0.9997444 | Phage endopeptidase |
| AP53_108 | -0.024292171 | 7.882968078 | 0.038524655 | 0.847844237 | 0.9997444 | Multiple sugar metabolism regulato |
| AP53_1459 | -0.028460026 | 5.692326177 | 0.038521845 | 0.847849711 | 0.9997444 | Mobile element protein |
| AP53_331 | -0.048774165 | 5.571994503 | 0.038368515 | 0.848148629 | 0.9997444 | hypothetical protein |
| AP53_45 | 0.051467398 | 11.20939483 | 0.038034898 | 0.848801173 | 0.9997444 | LSU ribosomal protein L23p (L23Ae |
| AP53_1329 | -0.054259249 | 3.992006818 | 0.037727303 | 0.849405468 | 0.9997444 | Phage protein |
| AP53_17 | 0.034413658 | 9.016055634 | 0.037458259 | 0.849936129 | 0.9997444 | Ribose-phosphate pyrophosphokin |
| AP53_rna24 | 0.095815226 | 3.822543586 | 0.037435655 | 0.849980804 | 0.9997444 | tRNA-Leu-TAA |
| AP53_596 | 0.069210013 | 8.004957284 | 0.03705607 | 0.850733119 | 0.9997444 | putative repressor C1 |
| AP53_14 | 0.027603616 | 10.75548072 | 0.036787015 | 0.8512688 | 0.9997444 | Cell division protein FtsH (EC 3.4.24 |
| AP53_531 | 0.019899127 | 7.100928641 | 0.036262166 | 0.852319656 | 0.9997444 | FIG01117156: hypothetical protei |
| AP53_1775 | 0.042099863 | 11.20879042 | 0.036178635 | 0.852487629 | 0.9997444 | Transcriptional regulator SpxA2 |
| AP53_164 | -0.061624012 | 4.340799239 | 0.035908074 | 0.853033091 | 0.9997444 | Mobile element protein |
| AP53_1001 | 0.057354401 | 7.743620698 | 0.03557743 | 0.853702591 | 0.9997444 | Phage protein |
| AP53_1736 | 0.024917129 | 13.53520806 | 0.035457429 | 0.853946371 | 0.9997444 | Heat shock protein 60 family chape |
| AP53_1698 | 0.037515946 | 9.395822895 | 0.034982206 | 0.854916014 | 0.9997444 | Immunogenic secreted protein |
| AP53_1475 | -0.026191232 | 5.649180463 | 0.034937535 | 0.855007511 | 0.9997444 | Mobile element protein |
| AP53_1816 | 0.034430605 | 5.463694811 | 0.034838404 | 0.855210772 | 0.9997444 | Cadmium resistance protein |
| AP53_269 | -0.022061521 | 8.07559697 | 0.034548422 | 0.85580709 | 0.9997444 | Ribonucleotide reductase transcrip |
| AP53_1305 | 0.024243656 | 7.406840544 | 0.034400593 | 0.856112085 | 0.9997444 | Putative two-component responsib |

| | | | | | | |
|---|---|---|---|---|---|---|
| AP53_957 | 0.040908379 | 8.083285336 | 0.033678111 | 0.857612548 | 0.9997444 | Lysyl aminopeptidase (EC 3.4.11.15 |
| AP53_364 | 0.021058078 | 9.106172219 | 0.033627313 | 0.857718672 | 0.9997444 | S1 RNA binding domain |
| AP53_737 | 0.020676611 | 7.671548301 | 0.033156373 | 0.858706505 | 0.9997444 | Hypothetical protein SPy0887 |
| AP53_1391 | 0.033041853 | 7.33462873 | 0.033005891 | 0.859023684 | 0.9997444 | D-beta-hydroxybutyrate permease |
| AP53_1369 | -0.03567768 | 4.662961336 | 0.032389149 | 0.86033151 | 0.9997444 | ComF operon protein A, DNA transp |
| AP53_245 | 0.027495503 | 5.525607065 | 0.032142902 | 0.860857286 | 0.9997444 | Mobile element protein |
| AP53_721 | 0.044252672 | 7.332076412 | 0.031906509 | 0.861363986 | 0.9997444 | Peptide deformylase (EC 3.5.1.88) |
| AP53_619 | -0.071488448 | 5.091305753 | 0.031714868 | 0.861776188 | 0.9997444 | Phage major capsid protein |
| AP53_1158 | 0.044989408 | 3.822656869 | 0.031642913 | 0.861931288 | 0.9997444 | Late competence protein ComEA, D |
| AP53_68 | 0.052560412 | 10.91520246 | 0.031541937 | 0.862149254 | 0.9997444 | SSU ribosomal protein S11p (S14e) |
| AP53_26 | 0.073048811 | 9.770366314 | 0.031434877 | 0.862380746 | 0.9997444 | IMP cyclohydrolase (EC 3.5.4.10) / I |
| AP53_750 | -0.04087367 | 8.941525709 | 0.031389261 | 0.862479503 | 0.9997444 | 6-aminohexanoate-cyclic-dimer hy |
| AP53_1461 | 0.017806103 | 9.01395109 | 0.030847968 | 0.863657102 | 0.9997444 | aminoglycoside phosphotransferas |
| AP53_1578 | 0.032654968 | 8.095803619 | 0.030637705 | 0.864117414 | 0.9997444 | Predicted regulator for deoxynucle |
| AP53_1319 | -0.048868993 | 3.772270914 | 0.030611516 | 0.86417486 | 0.9997444 | Phage-related protein |
| AP53_1165 | 0.018776768 | 9.470830918 | 0.030252425 | 0.864965128 | 0.9997444 | membrane protein, putative |
| AP53_rna66 | 0.075869139 | 4.587209041 | 0.030212873 | 0.865052467 | 0.9997444 | tRNA-Lys-TTT |
| AP53_1163 | -0.031201016 | 10.45687235 | 0.030087924 | 0.865328769 | 0.9997444 | Cold-shock DEAD-box protein A |
| AP53_681 | -0.032936297 | 9.560920647 | 0.029916376 | 0.865709085 | 0.9997444 | FIG139598: Potential ribosomal pr |
| AP53_258 | -0.035071741 | 7.671083701 | 0.02975633 | 0.866064918 | 0.9997444 | Methionine ABC transporter perme |
| AP53_rna25 | 0.117845992 | 3.774093468 | 0.029689747 | 0.866213243 | 0.9997444 | tRNA-Arg-ACG |
| AP53_91 | -0.034966445 | 8.79430569 | 0.029510546 | 0.866613301 | 0.9997444 | Acetate kinase (EC 2.7.2.1) |
| AP53_69 | 0.048505364 | 10.49884439 | 0.02948747 | 0.866664909 | 0.9997444 | DNA-directed RNA polymerase alph |
| AP53_727 | 0.02583157 | 7.583116513 | 0.029367169 | 0.866934287 | 0.9997444 | Diphosphomevalonate decarboxyla |
| AP53_530 | 0.025362533 | 9.094709856 | 0.029363006 | 0.866943619 | 0.9997444 | FIG001886: Cytoplasmic hypotheti |
| AP53_311 | 0.037238072 | 12.01622451 | 0.028969766 | 0.867828225 | 0.9997444 | ATP-dependent Clp protease protec |
| AP53_94 | 0.019943201 | 8.902392534 | 0.02892727 | 0.867924191 | 0.9997444 | Glutamyl aminopeptidase (EC 3.4.1 |
| AP53_946 | 0.02792751 | 8.691793309 | 0.028805271 | 0.868200098 | 0.9997444 | Phosphoglucomutase (EC 5.4.2.2) |
| AP53_1236 | 0.028056248 | 7.985314977 | 0.028789506 | 0.868235795 | 0.9997444 | Isoleucyl-tRNA synthetase (EC 6.1.1 |
| AP53_1764 | -0.044012233 | 9.072809522 | 0.028730493 | 0.86836951 | 0.9997444 | UPF0246 protein YaaA |
| AP53_372 | 0.02275351 | 8.032045335 | 0.02850969 | 0.868871079 | 0.9997444 | Diacylglycerol kinase (EC 2.7.1.107 |
| AP53_1385 | -0.045482878 | 6.421605107 | 0.028313264 | 0.869318959 | 0.9997444 | Transcriptional regulators, LysR fan |
| AP53_944 | -0.021157704 | 7.544989533 | 0.028226664 | 0.869516927 | 0.9997444 | Phosphopantothenoylcysteine dec |
| AP53_756 | 0.022387492 | 8.863382525 | 0.027868701 | 0.870338576 | 0.9997444 | Topoisomerase IV subunit B (EC 5.9 |
| AP53_744 | 0.01917703 | 8.440934461 | 0.027759814 | 0.870589587 | 0.9997444 | Histidine protein kinase |

| | | | | | | |
|---|---|---|---|---|---|---|
| AP53_589 | 0.025118624 | 6.987027995 | 0.027650058 | 0.870843115 | 0.9997444 | Alpha-D-GlcNAc alpha-1,2-L-rhamn |
| AP53_1333 | 0.045144589 | 3.062741465 | 0.027624541 | 0.87090213 | 0.9997444 | Phage minor capsid protein #Fam0 |
| AP53_1093 | -0.063231915 | 2.754249981 | 0.027512368 | 0.871161902 | 0.9997444 | Mobile element protein |
| AP53_1128 | -0.052493409 | 10.47072346 | 0.027356871 | 0.871522904 | 0.9997444 | Non-phosphorylating glyceraldehy |
| AP53_392 | -0.135498972 | 0.610601094 | 0.027125426 | 0.872062193 | 0.9997444 | FIG01116220: hypothetical protei |
| AP53_614 | 0.069731671 | 5.801452265 | 0.026966055 | 0.87243492 | 0.9997444 | hypothetical phage protein |
| AP53_133 | 0.036108121 | 10.50319653 | 0.026922687 | 0.872536541 | 0.9997444 | Transcription antitermination prot |
| AP53_39 | 0.027102067 | 10.17853263 | 0.026654548 | 0.87316674 | 0.9997444 | Alcohol dehydrogenase (EC 1.1.1.1 |
| AP53_962 | 0.022344056 | 7.398175823 | 0.026389976 | 0.873791759 | 0.9997444 | Phosphate transport system perme |
| AP53_1334 | -0.048265929 | 3.009464722 | 0.026356391 | 0.873871332 | 0.9997444 | Phage terminase large subunit |
| AP53_1804 | -0.020289756 | 6.605851342 | 0.02617685 | 0.874297594 | 0.9997444 | DNA mismatch repair protein MutS |
| AP53_532 | -0.027254617 | 8.611940454 | 0.025903066 | 0.874950509 | 0.9997444 | Candidate zinc-binding lipoprotein |
| AP53_1700 | 0.027424915 | 8.287535281 | 0.025723198 | 0.875381387 | 0.9997444 | Two-component system response r |
| AP53_113 | -0.041259967 | 5.138902606 | 0.025602596 | 0.87567116 | 0.9997444 | 3-ketoacyl-CoA thiolase (EC 2.3.1.1 |
| AP53_718 | 0.015502215 | 7.903787725 | 0.025485399 | 0.875953425 | 0.9997444 | tRNA nucleotidyltransferase (EC 2.7 |
| AP53_1293 | -0.023351117 | 5.977682409 | 0.025466549 | 0.875998887 | 0.9997444 | Two-component sensor kinase YesM |
| AP53_1433 | 0.019072487 | 10.47960649 | 0.025419228 | 0.876113089 | 0.9997444 | N-acetylglucosamine-6-phosphate |
| AP53_1787 | -0.049942261 | 5.888319301 | 0.025375384 | 0.876219 | 0.9997444 | Phage antirepressor protein |
| AP53_802 | 0.021863225 | 6.417002228 | 0.025328663 | 0.876331961 | 0.9997444 | Hypothetical protein YbbP, contair |
| AP53_568 | 0.019044487 | 9.895918157 | 0.025241548 | 0.876542877 | 0.9997444 | UDP-N-acetylglucosamine 1-carbox |
| AP53_775 | 0.022169621 | 9.194484888 | 0.025015709 | 0.877091405 | 0.9997444 | Endonuclease III (EC 4.2.99.18) |
| AP53_15 | 0.018341911 | 7.727893238 | 0.024926237 | 0.877309421 | 0.9997444 | Cationic amino acid transporter - A |
| AP53_1865 | -0.066804316 | 11.85379858 | 0.024721639 | 0.877809479 | 0.9997444 | Streptolysin S biosynthesis protein |
| AP53_1095 | -0.044281534 | 3.813457285 | 0.02471928 | 0.877815257 | 0.9997444 | Mobile element protein |
| AP53_994 | 0.024658848 | 10.55661971 | 0.024557942 | 0.878211103 | 0.9997444 | 6-phosphofructokinase (EC 2.7.1.1 |
| AP53_229 | -0.01986772 | 8.192570383 | 0.024487841 | 0.878383512 | 0.9997444 | Undecaprenyl-phosphate N-acetylg |
| AP53_788 | -0.02593607 | 5.911765568 | 0.024390622 | 0.878623038 | 0.9997444 | ABC transporter permease protein |
| AP53_1864 | 0.039102424 | 9.373845942 | 0.024169895 | 0.879168686 | 0.9997444 | Chromosome (plasmid) partitionin |
| AP53_1130 | 0.039188601 | 12.05479986 | 0.024036525 | 0.879499623 | 0.9997444 | Phosphocarrier protein of PTS syste |
| AP53_56 | 0.032949777 | 10.64711432 | 0.023987634 | 0.879621175 | 0.9997444 | SSU ribosomal protein S14p (S29e) |
| AP53_1354 | -0.056388175 | 3.37901892 | 0.023981577 | 0.879636243 | 0.9997444 | putative phage replication protein |
| AP53_1078 | -0.026932181 | 7.501289471 | 0.023922624 | 0.879782998 | 0.9997444 | Excinuclease ABC subunit B |
| AP53_1022 | -0.089193601 | 6.923782671 | 0.023895589 | 0.879850358 | 0.9997444 | conserved hypothetical protein - pl |
| AP53_186 | -0.030945518 | 8.362564137 | 0.023798132 | 0.880093515 | 0.9997444 | Lipid A export ATP-binding/permea |
| AP53_559 | 0.037637371 | 9.645722823 | 0.023574861 | 0.880652505 | 0.9997444 | ATP synthase C chain (EC 3.6.3.14) |

| | | | | | | |
|---|---|---|---|---|---|---|
| AP53_1622 | -0.028494384 | 7.523407045 | 0.023562914 | 0.880682493 | 0.9997444 | Galactose-6-phosphate isomerase, |
| AP53_471 | -0.025434294 | 8.036414201 | 0.023476926 | 0.880898556 | 0.9997444 | Putative metallopeptidase (Zinc) Sp |
| AP53_947 | -0.03941192 | 8.593368411 | 0.023358146 | 0.881197686 | 0.9997444 | Nucleoside ABC transporter, perme |
| AP53_686 | 0.019859116 | 8.354929584 | 0.02299119 | 0.882126765 | 0.9997444 | Uracil phosphoribosyltransferase (E |
| AP53_402 | 0.027791858 | 4.590131084 | 0.022859426 | 0.882462226 | 0.9997444 | hypothetical protein |
| AP53_1366 | -0.017671475 | 9.455194799 | 0.022719173 | 0.882820389 | 0.9997444 | Cysteinyl-tRNA synthetase related p |
| AP53_1289 | 0.019717705 | 8.056980692 | 0.02267866 | 0.882924056 | 0.9997444 | SAM-dependent methyltransferase |
| AP53_rna26 | 0.092944473 | 4.466309467 | 0.022444359 | 0.883525478 | 0.9997444 | tRNA-Pro-TGG |
| AP53_565 | 0.044386426 | 10.93748296 | 0.022370522 | 0.883715674 | 0.9997444 | ATP synthase beta chain (EC 3.6.3.1 |
| AP53_1090 | 0.052045644 | 7.046293108 | 0.021781915 | 0.885243504 | 0.9997444 | FIG01114098: hypothetical protei |
| AP53_rna71 | 0.076449463 | 3.871305666 | 0.021715904 | 0.885416156 | 0.9997444 | tRNA-Ala-TGC |
| AP53_886 | 0.027051389 | 11.36905956 | 0.021638387 | 0.885619248 | 0.9997444 | NADH peroxidase (EC 1.11.1.1) |
| AP53_629 | -0.047486594 | 3.905552694 | 0.02150843 | 0.885960568 | 0.9997444 | putative transcriptional activator - |
| AP53_517 | -0.021844334 | 5.639080472 | 0.02149797 | 0.885988085 | 0.9997444 | Beta-phosphoglucomutase (EC 5.4. |
| AP53_98 | -0.042360924 | 4.172683808 | 0.021408319 | 0.88622422 | 0.9997444 | FIG107367: tRNA-binding protein |
| AP53_468 | -0.017716259 | 8.330445525 | 0.021376815 | 0.886307319 | 0.9997444 | possible permease |
| AP53_1500 | -0.023545094 | 8.961250047 | 0.021104941 | 0.88702707 | 0.9997444 | Aspartyl-tRNA(Asn) amidotransfera |
| AP53_1524 | 0.026226426 | 10.72683241 | 0.021097449 | 0.88704697 | 0.9997444 | Immunogenic secreted protein |
| AP53_1613 | 0.042405823 | 3.795444941 | 0.020979986 | 0.887359463 | 0.9997444 | Salivaricin A modification enzyme |
| AP53_rna60 | 0.045337535 | 4.476833431 | 0.020842188 | 0.887727195 | 0.9997444 | tRNA-Arg-CCG |
| AP53_1220 | -0.014446737 | 8.806395054 | 0.020774508 | 0.887908261 | 0.9997444 | FIG01115348: hypothetical protei |
| AP53_1016 | -0.062585766 | 6.137775317 | 0.02071367 | 0.888071283 | 0.9997444 | Phage protein |
| AP53_560 | 0.040341361 | 9.652494913 | 0.0204701 | 0.888726415 | 0.9997444 | ATP synthase A chain (EC 3.6.3.14) |
| AP53_1276 | -0.028394631 | 5.528557162 | 0.020418495 | 0.88886573 | 0.9997444 | Cytochrome c-type biogenesis prot |
| AP53_83 | 0.032751847 | 3.433511599 | 0.020375607 | 0.888981647 | 0.9997444 | Late competence protein ComGA, a |
| AP53_1089 | -0.043606744 | 6.462081772 | 0.020201783 | 0.889452741 | 0.9997444 | Ribosyl nicotinamide transporter, |
| AP53_54 | 0.028069462 | 11.32762119 | 0.02015719 | 0.889573927 | 0.9997444 | LSU ribosomal protein L24p (L26e) |
| AP53_1600 | 0.019857285 | 6.597198535 | 0.020079815 | 0.889784533 | 0.9997444 | Potassium efflux system KefA protei |
| AP53_1472 | 0.03573005 | 9.358208714 | 0.020011591 | 0.889970573 | 0.9997444 | PTS system, mannose-specific IIC cc |
| AP53_1050 | 0.023911116 | 5.944521376 | 0.019995421 | 0.890014714 | 0.9997444 | hypothetical protein |
| AP53_1291 | -0.02055012 | 6.437852788 | 0.019887585 | 0.890309557 | 0.9997444 | Beta-galactosidase (EC 3.2.1.23) |
| AP53_529 | 0.016849318 | 9.140066361 | 0.019848711 | 0.890416044 | 0.9997444 | FIG002813: LPPG:FO 2-phospho-L- |
| AP53_713 | 0.034236129 | 2.030550548 | 0.019731457 | 0.890737889 | 0.9997444 | Immunoglobulin G-endopeptidase |
| AP53_1407 | -0.024239617 | 6.653605219 | 0.019671389 | 0.890903143 | 0.9997444 | FIG01116065: hypothetical protei |
| AP53_848 | 0.056809931 | 7.052381792 | 0.019324906 | 0.891861439 | 0.9997444 | NAD-dependent malic enzyme (EC 1 |

| | | | | | | |
|---|---|---|---|---|---|---|
| AP53_779 | 0.022912798 | 9.525736977 | 0.019276255 | 0.891996696 | 0.9997444 | Glucose-1-phosphate thymidylyltra |
| AP53_132 | -0.018628945 | 10.33594975 | 0.019215785 | 0.892165055 | 0.9997444 | Nucleoside-binding protein |
| AP53_1432 | -0.021851915 | 10.71659941 | 0.019205598 | 0.892193446 | 0.9997444 | oxidoreductase of aldo/keto reduct |
| AP53_rna28 | -0.068420844 | 4.380833458 | 0.019116256 | 0.892442752 | 0.9997444 | tRNA-Met-CAT |
| AP53_pseudo1389 | -0.030353865 | 7.021878509 | 0.019048859 | 0.892631218 | 0.9997444 | D-beta-hydroxybutyrate dehydroge |
| AP53_1121 | -0.020320277 | 8.876121787 | 0.018535351 | 0.894078474 | 0.9997444 | DNA polymerase III subunits gamma |
| AP53_240 | 0.015561981 | 8.982834402 | 0.018016992 | 0.895560299 | 0.9997444 | Oligopeptide transport system perr |
| AP53_666 | -0.021936565 | 10.2229329 | 0.018002882 | 0.895600935 | 0.9997444 | Translation initiation factor 3 |
| AP53_613 | -0.050837253 | 5.300012505 | 0.017685722 | 0.896518703 | 0.9997444 | Phage major tail protein |
| AP53_1114 | 0.015669259 | 7.467225765 | 0.017571494 | 0.896851296 | 0.9997444 | Ribosomal-protein-S5p-alanine ace |
| AP53_1729 | 0.013563338 | 8.363552599 | 0.017439975 | 0.897235604 | 0.9997444 | FIG01115477: hypothetical protei |
| AP53_1115 | -0.035726441 | 5.058997186 | 0.017353446 | 0.897489253 | 0.9997444 | Protein G-related alpha 2 macroglo |
| AP53_460 | 0.024845336 | 7.599552137 | 0.017206316 | 0.897922028 | 0.9997444 | Hydrolase (HAD superfamily) |
| AP53_rna65 | 0.06858614 | 4.639511609 | 0.017180149 | 0.897999197 | 0.9997444 | tRNA-Leu-TAG |
| AP53_1535 | 0.031037766 | 10.61596999 | 0.017151079 | 0.898084994 | 0.9997444 | Transcription termination protein |
| AP53_1063 | 0.021429202 | 6.562298696 | 0.017041334 | 0.898409566 | 0.9997444 | Maltodextrin phosphorylase (EC 2. |
| AP53_1161 | 0.030578255 | 7.283232404 | 0.01696945 | 0.898622742 | 0.9997444 | COG2827: putative endonuclease c |
| AP53_179 | 0.02319893 | 9.11682714 | 0.016964378 | 0.8986378 | 0.9997444 | FIG01118080: hypothetical protei |
| AP53_116 | -0.02801454 | 7.765070502 | 0.016850605 | 0.898976192 | 0.9997444 | YheO-like PAS domain |
| AP53_1143 | -0.023420194 | 9.577025249 | 0.016840193 | 0.899007217 | 0.9997444 | Foldase protein prsA 1 precursor (E |
| AP53_878 | -0.012332486 | 7.663320523 | 0.016389056 | 0.900361042 | 0.9997444 | Protein-N(5)-glutamine methyltran |
| AP53_847 | 0.048707426 | 6.940372562 | 0.016284944 | 0.900676158 | 0.9997444 | Malate Na(+) symporter |
| AP53_1571 | -0.025833775 | 6.358480657 | 0.016280097 | 0.900690854 | 0.9997444 | Transcriptional regulator, MerR fan |
| AP53_1699 | 0.031047776 | 8.424003252 | 0.016199405 | 0.900935826 | 0.9997444 | Two-component system histidine k |
| AP53_1721 | -0.028362662 | 6.061442966 | 0.015975692 | 0.901618265 | 0.9997444 | PTS system, cellobiose-specific IIA c |
| AP53_1031 | 0.023081747 | 6.564176908 | 0.015827832 | 0.902071984 | 0.9997444 | FIG01114733: hypothetical protei |
| AP53_1097 | -0.021638743 | 6.178719169 | 0.015696097 | 0.902478039 | 0.9997444 | Malonate permease |
| AP53_1367 | -0.036335879 | 10.32687169 | 0.015661679 | 0.902584415 | 0.9997444 | Ribosomal subunit interface protei |
| AP53_1408 | -0.021686897 | 8.416879746 | 0.015617235 | 0.902721954 | 0.9997444 | ATP-dependent RNA helicase YqfR |
| AP53_1769 | 0.018169351 | 9.866497896 | 0.015589869 | 0.90280674 | 0.9997444 | Ribonucleotide reductase of class II |
| AP53_259 | -0.028752879 | 7.002333297 | 0.015506987 | 0.90306399 | 0.9997444 | Branched-chain amino acid transpo |
| AP53_115 | -0.026317216 | 5.049902357 | 0.015208689 | 0.90399569 | 0.9997444 | Butyrate-acetoacetate CoA-transfer |
| AP53_1856 | 0.014131765 | 7.180055165 | 0.015018907 | 0.904593295 | 0.9997444 | Tryptophanyl-tRNA synthetase (EC 6 |
| AP53_694 | 0.013577168 | 8.383306483 | 0.01486546 | 0.905079297 | 0.9997444 | Glycerophosphoryl diester phosph |
| AP53_921 | -0.019315869 | 7.962211143 | 0.014739351 | 0.90548062 | 0.9997444 | Pleiotropic regulator of exopolysac |

| | | | | | | |
|---|---|---|---|---|---|---|
| AP53_458 | 0.019840036 | 3.931775774 | 0.014611902 | 0.90588799 | 0.9997444 | Mobile element protein |
| AP53_1474 | 0.024603976 | 6.336304883 | 0.014581988 | 0.905983865 | 0.9997444 | Putative regulator of the mannose ( |
| AP53_319 | 0.026757549 | 9.304296752 | 0.014578442 | 0.905995236 | 0.9997444 | FIG01114628: hypothetical protei |
| AP53_1638 | -0.017288564 | 7.493116441 | 0.014553788 | 0.90607434 | 0.9997444 | COG1939: Ribonuclease III family p |
| AP53_1503 | 0.021368852 | 7.92742632 | 0.014505222 | 0.90623036 | 0.9997444 | Pyruvate,phosphate dikinase (EC 2. |
| AP53_516 | -0.013443634 | 6.978985836 | 0.014496822 | 0.906257375 | 0.9997444 | 4-hydroxy-2-oxoglutarate aldolase |
| AP53_13 | -0.021205862 | 9.357275859 | 0.014469902 | 0.906343996 | 0.9997444 | Hypoxanthine-guanine phosphorib |
| AP53_346 | -0.030016726 | 9.215258084 | 0.014460893 | 0.906373003 | 0.9997444 | NAD-dependent oxidoreductase |
| AP53_215 | 0.015953506 | 9.515396733 | 0.014447657 | 0.906415638 | 0.9997444 | DNA recombination protein RmuC |
| AP53_1563 | 0.031158941 | 8.225798925 | 0.014246245 | 0.907066866 | 0.9997444 | Integral membrane protein |
| AP53_1781 | 0.022629858 | 9.060982847 | 0.014194376 | 0.90723533 | 0.9997444 | DNA mismatch repair protein MutL |
| AP53_1110 | -0.014683092 | 7.404757561 | 0.014166611 | 0.907325634 | 0.9997444 | 5-Enolpyruvylshikimate-3-phospha |
| AP53_807 | -0.018900782 | 6.845634405 | 0.01410299 | 0.907532901 | 0.9997444 | Acyl-ACP thioesterase |
| AP53_44 | 0.0287197 | 11.31807185 | 0.014014786 | 0.907821044 | 0.9997444 | LSU ribosomal protein L4p (L1e) |
| AP53_470 | -0.023589725 | 7.956823872 | 0.013943742 | 0.908053796 | 0.9997444 | Transcription accessory protein (S1 |
| AP53_917 | 0.034162291 | 8.589404302 | 0.013889239 | 0.908232766 | 0.9997444 | Oxaloacetate decarboxylase alpha c |
| AP53_47 | 0.02184825 | 10.83978006 | 0.013796154 | 0.908539255 | 0.9997444 | SSU ribosomal protein S19p (S15e) |
| AP53_1803 | -0.030126486 | 6.483353547 | 0.013697014 | 0.908866834 | 0.9997444 | hypothetical protein |
| AP53_1454 | 0.022297896 | 11.049852 | 0.013527302 | 0.909430408 | 0.9997444 | Translation initiation factor 2 |
| AP53_575 | -0.016783493 | 5.702429986 | 0.013484781 | 0.90957217 | 0.9997444 | Salt-stress induced protein |
| AP53_1484 | 0.029393224 | 9.517232347 | 0.013362197 | 0.909982132 | 0.9997444 | Malonyl CoA-acyl carrier protein tr |
| AP53_1036 | -0.032491485 | 5.48739783 | 0.013233826 | 0.910413497 | 0.9997444 | conserved hypothetical protein - pl |
| AP53_533 | -0.030923895 | 7.028844904 | 0.013194612 | 0.910545691 | 0.9997444 | Transcriptional regulator, GntR fam |
| AP53_283 | 0.018001127 | 9.767281081 | 0.013108843 | 0.910835524 | 0.9997444 | FIG005935: membrane protein |
| AP53_709 | 0.021119783 | 5.106968656 | 0.012928531 | 0.911447987 | 0.9997444 | Peptidoglycan hydrolase, Autolysin |
| AP53_473 | 0.022916769 | 9.506031141 | 0.012891563 | 0.91157409 | 0.9997444 | HPr kinase/phosphorylase (EC 2.7.1 |
| AP53_1612 | 0.021504525 | 4.415013354 | 0.01286916 | 0.9116506 | 0.9997444 | FIG00525014: hypothetical protei |
| AP53_1240 | 0.022669038 | 10.00904625 | 0.012712436 | 0.912187732 | 0.9997444 | FIG001960: FtsZ-interacting protei |
| AP53_70 | -0.028143224 | 10.44975777 | 0.01270468 | 0.912214398 | 0.9997444 | LSU ribosomal protein L17p |
| AP53_1277 | -0.024085278 | 10.15144344 | 0.012360669 | 0.913405677 | 0.9997444 | Magnesium and cobalt transport pr |
| AP53_139 | -0.025456355 | 6.850934411 | 0.012278016 | 0.913694392 | 0.9997444 | Cystathionine beta-lyase (EC 4.4.1.8 |
| AP53_267 | -0.026603182 | 10.33651236 | 0.012249544 | 0.913794074 | 0.9997444 | Response regulator CsrR |
| AP53_489 | 0.035686444 | 7.91146 | 0.012190104 | 0.914002559 | 0.9997444 | Substrate-specific component CbrT |
| AP53_1216 | 0.022949458 | 11.02528837 | 0.012158837 | 0.914112433 | 0.9997444 | DNA-binding protein HBsu |
| AP53_144 | 0.020524349 | 5.193900191 | 0.012148871 | 0.914147485 | 0.9997444 | 3-keto-L-gulonate 6-phosphate dec |

| | | | | | | |
|---|---|---|---|---|---|---|
| AP53_984 | 0.024121857 | 8.429015997 | 0.012139092 | 0.914181892 | 0.9997444 | Sodium/glycine symporter GlyP |
| AP53_1720 | -0.018502207 | 6.068932166 | 0.012120987 | 0.914245633 | 0.9997444 | PTS system, cellobiose-specific IIB c |
| AP53_1525 | 0.023994965 | 9.540654293 | 0.012098214 | 0.914325877 | 0.9997444 | Alanine racemase (EC 5.1.1.1) |
| AP53_742 | -0.026589136 | 10.24984754 | 0.012017888 | 0.914609524 | 0.9997444 | Purine nucleoside phosphorylase (E |
| AP53_64 | 0.030722514 | 7.801527973 | 0.011806478 | 0.915360686 | 0.9997444 | Adenylate kinase (EC 2.7.4.3) |
| AP53_1506 | -0.011037778 | 9.598567028 | 0.01170929 | 0.915708288 | 0.9997444 | Aspartate aminotransferase (EC 2.6. |
| AP53_792 | 0.012783975 | 7.475883425 | 0.011706229 | 0.915719261 | 0.9997444 | FIG01115226: hypothetical protei |
| AP53_1476 | -0.013909962 | 7.639481239 | 0.011643682 | 0.915943768 | 0.9997444 | Seryl-tRNA synthetase (EC 6.1.1.11) |
| AP53_1399 | -0.017035516 | 10.44428353 | 0.011551772 | 0.916274784 | 0.9997444 | Aminopeptidase C (EC 3.4.22.40) |
| AP53_1766 | -0.030779981 | 8.876366489 | 0.011417074 | 0.916762324 | 0.9997444 | Acetyltransferase (EC 2.3.1.-) |
| AP53_1202 | 0.024369342 | 3.55737465 | 0.011174345 | 0.917648284 | 0.9997444 | Phage protein |
| AP53_512 | -0.016885581 | 5.679056086 | 0.011101487 | 0.917916115 | 0.9997444 | PTS system, N-acetylgalactosamine- |
| AP53_1129 | -0.026688231 | 10.5572049 | 0.011052808 | 0.918095557 | 0.9997444 | Phosphoenolpyruvate-protein pho |
| AP53_92 | -0.017055276 | 6.356531205 | 0.01090226 | 0.918653053 | 0.9997444 | Hypothetical protein SPy0110 |
| AP53_1014 | -0.040588713 | 6.659339264 | 0.010692056 | 0.919438023 | 0.9997444 | Phage tail length tape-measure prot |
| AP53_79 | -0.014882973 | 8.410774121 | 0.010527664 | 0.920057372 | 0.9997444 | Multimodular transpeptidase-trans |
| AP53_1716 | 0.011020239 | 7.147578025 | 0.010169532 | 0.921423798 | 0.9997444 | Glycerol dehydrogenase (EC 1.1.1.6 |
| AP53_1424 | 0.013902199 | 7.751467306 | 0.010011983 | 0.922032616 | 0.9997444 | Alpha-glycerophosphate oxidase (E |
| AP53_1384 | -0.011760586 | 10.00184868 | 0.010008052 | 0.922047865 | 0.9997444 | Hydrolase (HAD superfamily) |
| AP53_1779 | 0.029413325 | 8.691103742 | 0.009992865 | 0.922106821 | 0.9997444 | Holliday junction DNA helicase Ruv |
| AP53_1842 | -0.014580142 | 6.89698383 | 0.009921565 | 0.922384202 | 0.9997444 | ATPase component of general energ |
| AP53_915 | 0.028633389 | 8.680433803 | 0.009904864 | 0.922449322 | 0.9997444 | Citrate lyase alpha chain (EC 4.1.3.6 |
| AP53_769 | -0.015163345 | 8.463737777 | 0.009807905 | 0.922828474 | 0.9997444 | Cystathionine beta-lyase (EC 4.4.1.8 |
| AP53_1295 | -0.017809104 | 7.199351983 | 0.00966853 | 0.923376826 | 0.9997444 | Multiple sugar ABC transporter, sub |
| AP53_264 | 0.01900884 | 9.718548387 | 0.009366842 | 0.924577624 | 0.9997444 | LemA protein |
| AP53_1142 | -0.01926986 | 8.075777564 | 0.009146105 | 0.925468638 | 0.9997444 | Alanyl-tRNA synthetase (EC 6.1.1.7) |
| AP53_1373 | -0.018989508 | 9.092152825 | 0.009140724 | 0.925490493 | 0.9997444 | Streptococcal lipoprotein rotamase |
| AP53_404 | 0.029107992 | 3.441721999 | 0.009118834 | 0.925579471 | 0.9997444 | hypothetical protein |
| AP53_49 | -0.022106683 | 11.30618214 | 0.0090678 | 0.925787325 | 0.9997444 | SSU ribosomal protein S3p (S3e) |
| AP53_617 | 0.03680643 | 4.683457916 | 0.008917889 | 0.926401336 | 0.9997444 | hypothetical phage protein |
| AP53_677 | -0.012352786 | 8.111374097 | 0.008917379 | 0.926403436 | 0.9997444 | Cysteine desulfurase (EC 2.8.1.7) |
| AP53_221 | -0.016476897 | 11.87240178 | 0.008840651 | 0.926719727 | 0.9997444 | Translation elongation factor G |
| AP53_1051 | 0.031621209 | 8.48216266 | 0.008727627 | 0.927188173 | 0.9997444 | hypothetical protein |
| AP53_1439 | 0.014362241 | 8.381408097 | 0.008675266 | 0.927406227 | 0.9997444 | INTEGRAL MEMBRANE PROTEIN (Rh |
| AP53_1487 | -0.022086125 | 10.25795846 | 0.008582325 | 0.927794921 | 0.9997444 | 3-oxoacyl-[acyl-carrier-protein] syn |

| | | | | | | |
|---|---|---|---|---|---|---|
| AP53_46 | -0.022544898 | 11.29190986 | 0.008424111 | 0.928461514 | 0.9997444 | LSU ribosomal protein L2p (L8e) |
| AP53_390 | -0.016750082 | 5.938200776 | 0.008391777 | 0.928598521 | 0.9997444 | Response regulator of the compete |
| AP53_1018 | -0.035858473 | 5.920351528 | 0.008152872 | 0.929619181 | 0.9997444 | Phage protein |
| AP53_431 | 0.019393397 | 6.495646796 | 0.008048633 | 0.930069238 | 0.9997444 | Long-chain-fatty-acid--CoA ligase (E |
| AP53_717 | 0.023544185 | 8.491106844 | 0.008014719 | 0.930216294 | 0.9997444 | DegV family protein |
| AP53_1415 | -0.011741271 | 8.159083617 | 0.007993108 | 0.93031017 | 0.9997444 | Glutamate 5-kinase (EC 2.7.2.11) |
| AP53_1522 | 0.015180147 | 8.075835223 | 0.007990098 | 0.930323255 | 0.9997444 | Streptococcal cell surface hemopro |
| AP53_1279 | -0.012549811 | 6.54295096 | 0.007960037 | 0.93045407 | 0.9997444 | FIG01114815: hypothetical protei |
| AP53_871 | 0.016401499 | 7.585096391 | 0.007744501 | 0.931399411 | 0.9997444 | GMP reductase (EC 1.7.1.7) |
| AP53_1786 | 0.021142644 | 6.068040946 | 0.007637976 | 0.93187153 | 0.9997444 | kilA protein, putative phage-relate |
| AP53_1789 | 0.011685383 | 5.668436909 | 0.007523109 | 0.932384357 | 0.9997444 | Mobile element protein |
| AP53_418 | 0.009316912 | 8.8980063 | 0.007466405 | 0.932638964 | 0.9997444 | Lactoylglutathione lyase (EC 4.4.1. |
| AP53_1248 | 0.019754972 | 9.88017853 | 0.007401756 | 0.932930445 | 0.9997444 | GTP-binding protein TypA/BipA |
| AP53_446 | 0.016665288 | 3.672188595 | 0.007374302 | 0.933054612 | 0.9997444 | UDP-glucose dehydrogenase (EC 1.1 |
| AP53_725 | 0.014940681 | 7.957002543 | 0.00736978 | 0.933075083 | 0.9997444 | Regulation of D-alanyl-lipoteichoic |
| AP53_1371 | -0.019118368 | 10.82174338 | 0.007360681 | 0.9331163 | 0.9997444 | Cysteine synthase (EC 2.5.1.47) |
| AP53_925 | 0.012231328 | 5.40964687 | 0.007253165 | 0.933605283 | 0.9997444 | FIG01114068: hypothetical protei |
| AP53_1794 | 0.023549475 | 5.449043881 | 0.007211537 | 0.933795589 | 0.9997444 | FIG01114073: hypothetical protei |
| AP53_688 | 0.012636868 | 9.040885876 | 0.007188522 | 0.933901042 | 0.9997444 | Aspartate carbamoyltransferase (EC |
| AP53_1112 | -0.015739309 | 9.61875054 | 0.00718344 | 0.933924348 | 0.9997444 | Methionine aminopeptidase (EC 3. |
| AP53_1205 | -0.025345712 | 2.924279 | 0.007073939 | 0.934428596 | 0.9997444 | FIG01114509: hypothetical protei |
| AP53_1767 | -0.013244964 | 9.051988918 | 0.007019789 | 0.934679406 | 0.9997444 | Putative oxidoreductase |
| AP53_1244 | -0.011341159 | 8.999923432 | 0.006963953 | 0.934939053 | 0.9997444 | Cell division protein FtsQ |
| AP53_1275 | 0.013854873 | 6.163761658 | 0.006884641 | 0.93530967 | 0.9997444 | Thiol:disulfide oxidoreductase asso |
| AP53_301 | 0.016380226 | 9.702029137 | 0.006728559 | 0.936045367 | 0.9997444 | Manganese-dependent inorganic p |
| AP53_1343 | -0.020486637 | 4.452770568 | 0.006708724 | 0.936139471 | 0.9997444 | Phage protein |
| AP53_42 | -0.019596118 | 10.79703521 | 0.006626866 | 0.936529334 | 0.9997444 | SSU ribosomal protein S10p (S20e) |
| AP53_1623 | -0.014040101 | 6.765951862 | 0.006615494 | 0.936583683 | 0.9997444 | Galactose-6-phosphate isomerase, |
| AP53_395 | -0.020310134 | 4.13609729 | 0.0065331 | 0.936978896 | 0.9997444 | Bacteriocin-like peptide M BlpM |
| AP53_782 | -0.016294255 | 6.754178647 | 0.006285509 | 0.938181844 | 0.9997444 | Mutator mutT protein (7,8-dihydr |
| AP53_1782 | -0.014220878 | 8.412433501 | 0.006231647 | 0.938446683 | 0.9997444 | Phage integrase: site-specific recom |
| AP53_88 | 0.019821546 | 3.287412157 | 0.006191411 | 0.938645272 | 0.9997444 | Late competence protein ComGF, a |
| AP53_1634 | 0.021264635 | 8.437611102 | 0.006143747 | 0.938881369 | 0.9997444 | DegV family protein in cluster with |
| AP53_1527 | -0.008998841 | 10.56430151 | 0.006140706 | 0.938896465 | 0.9997444 | Protein export cytoplasm protein S |
| AP53_511 | 0.012007135 | 6.045520278 | 0.006130625 | 0.938946526 | 0.9997444 | Unsaturated glucuronyl hydrolase ( |

| | | | | | | |
|---|---|---|---|---|---|---|
| AP53_241 | 0.012184169 | 9.196140432 | 0.006018549 | 0.939505947 | 0.9997444 | Oligopeptide transport ATP-binding |
| AP53_773 | 0.011289893 | 9.386672227 | 0.005937962 | 0.939911432 | 0.9997444 | Adenine phosphoribosyltransferase |
| AP53_626 | 0.025292841 | 3.253685203 | 0.00592869 | 0.939958264 | 0.9997444 | hypothetical phage protein |
| AP53_1499 | -0.011781938 | 9.780226003 | 0.005880669 | 0.940201402 | 0.9997444 | Aspartyl-tRNA(Asn) amidotransferas |
| AP53_24 | 0.02643492 | 8.609949346 | 0.00578879 | 0.940669397 | 0.9997444 | Phosphoribosylformylglycinamidin |
| AP53_rna19 | 0.042608266 | 4.531004466 | 0.005742151 | 0.940908385 | 0.9997444 | tRNA-Asp-GTC |
| AP53_1480 | -0.015378078 | 9.729546962 | 0.005700717 | 0.94112153 | 0.9997444 | (3R)-hydroxymyristoyl-[acyl carrier |
| AP53_378 | -0.014735536 | 4.547407381 | 0.005600525 | 0.941640167 | 0.9997444 | Peptide ABC transporter ATP bindin |
| AP53_1123 | 0.009990465 | 5.808376608 | 0.005593387 | 0.941677294 | 0.9997444 | FIG01115656: hypothetical protein |
| AP53_642 | -0.018471052 | 5.138889041 | 0.005581887 | 0.941737159 | 0.9997444 | Helicase loader DnaI |
| AP53_377 | -0.010733841 | 4.518264802 | 0.005433069 | 0.942517527 | 0.9997444 | Peptide ABC transporter ATP bindin |
| AP53_702 | 0.015271651 | 6.892190454 | 0.00540611 | 0.94266004 | 0.9997444 | tRNA (Guanine37-N1) -methyltrans |
| AP53_1382 | 0.012431408 | 9.307646867 | 0.005394128 | 0.942723491 | 0.9997444 | DNA-directed RNA polymerase ome |
| AP53_1055 | 0.008048904 | 7.699856721 | 0.005141165 | 0.944080076 | 0.9997444 | hypothetical phage protein |
| AP53_1340 | 0.014807218 | 4.087828755 | 0.005066553 | 0.944486586 | 0.9997444 | conserved hypothetical protein - ph |
| AP53_285 | -0.012690381 | 7.505391079 | 0.005057764 | 0.944534667 | 0.9997444 | Glutamate racemase (EC 5.1.1.3) |
| AP53_1092 | -0.017744649 | 6.259530038 | 0.004942498 | 0.945169189 | 0.9997444 | FIG01114947: hypothetical protein |
| AP53_701 | -0.01499249 | 7.052130267 | 0.004881603 | 0.945507411 | 0.9997444 | 16S rRNA processing protein RimM |
| AP53_1119 | 0.012376833 | 7.650473294 | 0.004873536 | 0.945552377 | 0.9997444 | Biotin operon repressor / Biotin-pro |
| AP53_712 | -0.044270214 | 1.394023947 | 0.004828105 | 0.945806305 | 0.9997444 | hypothetical protein |
| AP53_1304 | 0.014167873 | 9.344842128 | 0.004703801 | 0.94650728 | 0.9997444 | Alpha-mannosidase (EC 3.2.1.24) |
| AP53_265 | 0.017106323 | 9.95527985 | 0.004698495 | 0.946537409 | 0.9997444 | Heat shock protein HtpX (EC 3.4.24 |
| AP53_227 | 0.008167986 | 7.690531152 | 0.00464886 | 0.946820073 | 0.9997444 | Undecaprenyl-diphosphatase (EC 3. |
| AP53_707 | 0.010129404 | 8.129301303 | 0.00464129 | 0.946863312 | 0.9997444 | 1-phosphofructokinase (EC 2.7.1.5 |
| AP53_1029 | -0.024323204 | 5.505463083 | 0.004639482 | 0.946873646 | 0.9997444 | Phage terminase, small subunit |
| AP53_1502 | 0.00862302 | 5.633125602 | 0.004601658 | 0.947090289 | 0.9997444 | Mobile element protein |
| AP53_1426 | 0.013586257 | 6.450441917 | 0.00457699 | 0.947232058 | 0.9997444 | FIG01116532: hypothetical protein |
| AP53_1732 | 0.009060502 | 8.474699799 | 0.004546112 | 0.947410063 | 0.9997444 | Dipeptidase (EC 3.4.-.-) |
| AP53_1479 | 0.014762091 | 10.1080263 | 0.004492357 | 0.947721398 | 0.9997444 | Biotin carboxylase of acetyl-CoA car |
| AP53_863 | -0.015071746 | 7.861865308 | 0.004490125 | 0.947734364 | 0.9997444 | GTP pyrophosphokinase (EC 2.7.6.5 |
| AP53_599 | 0.035667037 | 9.392532048 | 0.004477243 | 0.94780927 | 0.9997444 | Phage protein |
| AP53_1412 | -0.00791578 | 8.859933775 | 0.004413562 | 0.948181163 | 0.9997444 | rRNA small subunit methyltransfera |
| AP53_171 | 0.01341002 | 7.61453829 | 0.004379219 | 0.948382847 | 0.9997444 | Metal-dependent hydrolase (EC 3.-. |
| AP53_rna40 | 0.03900585 | 3.861167023 | 0.004306883 | 0.948810258 | 0.9997444 | tRNA-Gly-TCC |
| AP53_1155 | -0.01957747 | 11.46796564 | 0.004289712 | 0.948912244 | 0.9997444 | Manganese superoxide dismutase (E |

| | | | | | | |
|---|---|---|---|---|---|---|
| AP53_351 | 0.011761756 | 9.231868952 | 0.004211689 | 0.949378261 | 0.9997444 | 5'-methylthioadenosine nucleosida |
| AP53_1353 | 0.019059757 | 4.458065608 | 0.004150002 | 0.949749788 | 0.9997444 | DNA replication protein DnaC |
| AP53_rna10 | 0.050722438 | 2.562764711 | 0.004124292 | 0.94990545 | 0.9997444 | tRNA-Gly-GCC |
| AP53_143 | 0.010079931 | 4.941427565 | 0.004111659 | 0.949982114 | 0.9997444 | Ascorbate-specific PTS system, EIIA |
| AP53_978 | -0.014936995 | 13.63851091 | 0.004083189 | 0.950155328 | 0.9997444 | FIG01114697: hypothetical protei |
| AP53_728 | -0.009195414 | 7.770431655 | 0.004035977 | 0.950443904 | 0.9997444 | Phosphomevalonate kinase (EC 2.7. |
| AP53_1135 | -0.014131233 | 7.158557208 | 0.004016888 | 0.950561065 | 0.9997444 | Chloride channel protein |
| AP53_1297 | 0.010267812 | 6.253431008 | 0.003980945 | 0.950782432 | 0.9997444 | Multiple sugar ABC transporter, me |
| AP53_1692 | -0.009735204 | 11.16790356 | 0.003879701 | 0.951411425 | 0.9997444 | Antiphagocytic M protein |
| AP53_539 | 0.014175396 | 8.780216336 | 0.003805443 | 0.951878022 | 0.9997444 | Chloride channel protein |
| AP53_1800 | 0.017586547 | 4.960088655 | 0.003753498 | 0.952207135 | 0.9997444 | FIG01115581: hypothetical protei |
| AP53_55 | -0.010940166 | 10.94485035 | 0.003734927 | 0.952325354 | 0.9997444 | LSU ribosomal protein L5p (L11e) |
| AP53_903 | -0.011676525 | 4.168785317 | 0.00373346 | 0.952334704 | 0.9997444 | FIG01114041: hypothetical protei |
| AP53_214 | -0.006261972 | 9.089887731 | 0.003623484 | 0.953041056 | 0.9997444 | Thiamin pyrophosphokinase (EC 2.7 |
| AP53_932 | 0.012707551 | 5.974249375 | 0.003515184 | 0.95374723 | 0.9997444 | Predicted transcriptional regulator |
| AP53_1288 | -0.006311882 | 9.249318484 | 0.003496395 | 0.953870857 | 0.9997444 | hypothetical protein |
| AP53_23 | 0.017040373 | 8.50615647 | 0.003448495 | 0.954187524 | 0.9997444 | Amidophosphoribosyltransferase (E |
| AP53_366 | 0.011020225 | 8.651241561 | 0.003413195 | 0.954422312 | 0.9997444 | FIG01114318: hypothetical protei |
| AP53_329 | 0.016487697 | 7.559046869 | 0.003340959 | 0.954906603 | 0.9997444 | Membrane protein, putative |
| AP53_1386 | 0.014616817 | 7.247876238 | 0.003333174 | 0.954959105 | 0.9997444 | 3-ketoacyl-CoA thiolase (EC 2.3.1.1 |
| AP53_834 | 0.010102395 | 7.083534567 | 0.003187216 | 0.955955138 | 0.9997444 | Polysaccharide deacetylase |
| AP53_427 | -0.00930509 | 4.410117983 | 0.003151896 | 0.956199584 | 0.9997444 | Daunorubicin resistance transmem |
| AP53_561 | -0.012787361 | 9.705734837 | 0.003148701 | 0.956221769 | 0.9997444 | ATP synthase B chain (EC 3.6.3.14) |
| AP53_379 | -0.017482417 | 4.715037934 | 0.003114395 | 0.956460634 | 0.9997444 | Bacteriocin BlpU |
| AP53_1229 | 0.006161295 | 6.707222003 | 0.00308262 | 0.956683066 | 0.9997444 | Deoxyribodipyrimidine photolyase |
| AP53_239 | -0.008882851 | 8.619236094 | 0.003071175 | 0.956763464 | 0.9997444 | Oligopeptide transport system perr |
| AP53_1271 | 0.010076807 | 7.72486972 | 0.002957742 | 0.95756857 | 0.9997444 | Hypothetical protein SpyM3_1200 |
| AP53_1122 | -0.011541041 | 8.656673592 | 0.002871082 | 0.958194142 | 0.9997444 | GAF domain-containing proteins |
| AP53_408 | 0.007232985 | 9.562030378 | 0.00283867 | 0.958430543 | 0.9997444 | ATPase |
| AP53_562 | -0.016382784 | 9.093359734 | 0.002735896 | 0.959189231 | 0.9997444 | ATP synthase delta chain (EC 3.6.3.1 |
| AP53_1473 | -0.012114497 | 9.677362925 | 0.002602269 | 0.960197384 | 0.9997444 | PTS system, mannose-specific IID cc |
| AP53_1394 | -0.010098632 | 10.01987976 | 0.002546235 | 0.960627844 | 0.9997444 | FIG001721: Predicted N6-adenine- |
| AP53_1520 | 0.006230138 | 7.600032679 | 0.002515928 | 0.960862651 | 0.9997444 | Heme transporter IsdDEF, lipoprote |
| AP53_609 | 0.013397482 | 4.442814656 | 0.002454775 | 0.961340788 | 0.9997444 | Phage tail fibers |
| AP53_1247 | -0.006344566 | 8.304777516 | 0.00238996 | 0.961854131 | 0.9997444 | FIG01114439: hypothetical protei |

| | | | | | | |
|---|---|---|---|---|---|---|
| AP53_803 | 0.008158667 | 6.424057096 | 0.002384974 | 0.961893905 | 0.9997444 | uncharacterized secreted protein, Y |
| AP53_485 | -0.007334464 | 6.997365124 | 0.00234939 | 0.962179004 | 0.9997444 | Phosphoglycerate mutase family 5 |
| AP53_1515 | 0.004480076 | 7.717556279 | 0.002248942 | 0.962995677 | 0.9997444 | Substrate-specific component BL06 |
| AP53_rna9 | 0.019693501 | 4.351643606 | 0.002220618 | 0.96322925 | 0.9997444 | tRNA-Thr-TGT |
| AP53_178 | -0.017476668 | 11.23122741 | 0.002217984 | 0.963251046 | 0.9997444 | Transcriptional regulator, RofA-like |
| AP53_1770 | -0.006138275 | 7.59429478 | 0.002174591 | 0.963612023 | 0.9997444 | Competence-induced protein Ccs4 |
| AP53_995 | -0.003913197 | 7.959449408 | 0.002104201 | 0.964205337 | 0.9997444 | DNA polymerase III alpha subunit (E |
| AP53_524 | 0.006172243 | 5.69811416 | 0.002101979 | 0.964224224 | 0.9997444 | Mobile element protein |
| AP53_474 | 0.006866721 | 9.129672143 | 0.002071481 | 0.964484517 | 0.9997444 | Prolipoprotein diacylglyceryl trans |
| AP53_165 | -0.013996929 | 2.954387678 | 0.002067875 | 0.964515414 | 0.9997444 | Mobile element protein |
| AP53_796 | -0.00948714 | 10.79042999 | 0.002018878 | 0.96493802 | 0.9997444 | Dihydrolipoamide acetyltransferase |
| AP53_624 | 0.014902956 | 4.183410825 | 0.001947037 | 0.965567053 | 0.9997444 | Phage-associated homing endonucl |
| AP53_492 | 0.007387648 | 7.705728591 | 0.001913809 | 0.965861929 | 0.9997444 | Oligoendopeptidase F |
| AP53_710 | 0.007081856 | 7.45928674 | 0.001880714 | 0.96615818 | 0.9997444 | Peptidoglycan hydrolase, Autolysin |
| AP53_1069 | 0.010699759 | 5.517887881 | 0.001839541 | 0.966530415 | 0.9997444 | Mobile element protein |
| AP53_rna7 | -0.019298937 | 4.436492387 | 0.001795378 | 0.966934357 | 0.9997444 | tRNA-Lys-TTT |
| AP53_639 | 0.008538895 | 4.770258326 | 0.001793121 | 0.966955133 | 0.9997444 | conserved hypothetical protein - ph |
| AP53_1777 | 0.007022535 | 7.689044846 | 0.001776078 | 0.96711245 | 0.9997444 | Molybdopterin binding motif, CinA |
| AP53_1756 | 0.009155598 | 9.032149695 | 0.001740745 | 0.967441011 | 0.9997444 | Neutral endopeptidase O (EC 3.4.24 |
| AP53_1606 | -0.008697475 | 4.367122512 | 0.001728004 | 0.967560313 | 0.9997444 | Competence-specific sigma factor C |
| AP53_1805 | -0.006264912 | 7.418596014 | 0.00171157 | 0.96771484 | 0.9997444 | biofilm formation protein YmcA |
| AP53_916 | -0.01143481 | 8.567138716 | 0.001709535 | 0.967734027 | 0.9997444 | Apo-citrate lyase phosphoribosyl-d |
| AP53_729 | -0.005647922 | 7.986503413 | 0.001674478 | 0.968066377 | 0.9997444 | Isopentenyl-diphosphate delta-isom |
| AP53_370 | 0.007132936 | 7.3926236 | 0.00164225 | 0.968374987 | 0.9997444 | Hypothetical protein VC0266 (suga |
| AP53_757 | -0.00712322 | 8.988796479 | 0.001634608 | 0.968448611 | 0.9997444 | Topoisomerase IV subunit A (EC 5.9 |
| AP53_48 | -0.008687386 | 11.44172855 | 0.001601561 | 0.968768988 | 0.9997444 | LSU ribosomal protein L22p (L17e) |
| AP53_rna31 | 0.02045153 | 3.575514669 | 0.001564966 | 0.969127654 | 0.9997444 | tRNA-Phe-GAA |
| AP53_1829 | 0.009240041 | 9.962085059 | 0.001497523 | 0.96979984 | 0.9997444 | SSU ribosomal protein S4p (S9e) |
| AP53_1332 | 0.006712532 | 4.927605505 | 0.001487796 | 0.969898028 | 0.9997444 | Phage minor capsid protein |
| AP53_1806 | 0.005517045 | 6.705292417 | 0.001452689 | 0.970255114 | 0.9997444 | Arginine pathway regulatory protei |
| AP53_982 | -0.006369827 | 11.67228753 | 0.00131442 | 0.971705377 | 0.9997444 | FIG01114177: hypothetical protein |
| AP53_911 | -0.007430446 | 7.402583754 | 0.001294912 | 0.971916031 | 0.9997444 | Oxaloacetate decarboxylase beta ch |
| AP53_187 | 0.007720977 | 7.024537248 | 0.001289074 | 0.971979377 | 0.9997444 | FIG053235: Diacylglucosamine hyd |
| AP53_1737 | -0.006955361 | 13.40341086 | 0.001279625 | 0.972082213 | 0.9997444 | Heat shock protein 60 family co-cha |
| AP53_323 | -0.006360579 | 7.876094137 | 0.001273862 | 0.972145121 | 0.9997444 | Exodeoxyribonuclease III (EC 3.1.11 |

| | | | | | | |
|---|---|---|---|---|---|---|
| AP53_567 | -0.005515806 | 7.522620469 | 0.001270723 | 0.972179451 | 0.9997444 | FIG01114865: hypothetical protei |
| AP53_964 | -0.004057354 | 6.85891838 | 0.001244706 | 0.972465594 | 0.9997444 | tRNA and rRNA cytosine-C5-methyl |
| AP53_1309 | 0.011897882 | 6.652699164 | 0.001236541 | 0.97255601 | 0.9997444 | FIG01117556: hypothetical protei |
| AP53_1009 | -0.011823843 | 5.668230454 | 0.001236374 | 0.972557867 | 0.9997444 | FIG01116522: hypothetical protei |
| AP53_163 | 0.008786093 | 3.782831697 | 0.001221102 | 0.972727803 | 0.9997444 | Mobile element protein |
| AP53_884 | 0.006103376 | 8.408772709 | 0.001216207 | 0.972782492 | 0.9997444 | ABC transporter, ATP-binding/perm |
| AP53_997 | -0.005050291 | 8.557511037 | 0.001207897 | 0.9728756 | 0.9997444 | ABC-type multidrug transport syste |
| AP53_676 | -0.008103867 | 8.098908033 | 0.001154632 | 0.973480138 | 0.9997444 | Histone acetyltransferase Gcn5 |
| AP53_667 | -0.007216815 | 10.64619676 | 0.001096619 | 0.974154683 | 0.9997444 | LSU ribosomal protein L35p |
| AP53_669 | -0.004711472 | 8.124166179 | 0.0010452 | 0.974767645 | 0.9997444 | Lipoteichoic acid synthase LtaS Typ |
| AP53_1150 | -0.004629836 | 7.68102015 | 0.001031239 | 0.974936664 | 0.9997444 | S-adenosylmethionine:tRNA ribosy |
| AP53_1019 | 0.013159625 | 7.874398624 | 0.00102143 | 0.975056109 | 0.9997444 | Phage protein |
| AP53_691 | -0.005465446 | 8.599600017 | 0.001017978 | 0.975098273 | 0.9997444 | periplasmic component of efflux sy |
| AP53_1059 | 0.003039832 | 6.906797958 | 0.001010944 | 0.975184428 | 0.9997444 | Pleiotropic regulator of exopolysac |
| AP53_419 | -0.003635421 | 9.639050574 | 0.001004793 | 0.975260009 | 0.9997444 | Oxygen-insensitive NAD(P)H nitrore |
| AP53_1290 | -0.00345853 | 6.889385727 | 0.001001378 | 0.975302067 | 0.9997444 | Shikimate/quinate 5-dehydrogenas |
| AP53_1347 | 0.009779157 | 4.277213214 | 0.000967626 | 0.975721714 | 0.9997444 | Single-stranded DNA-binding prote |
| AP53_490 | -0.007647762 | 8.144957934 | 0.000935436 | 0.976128831 | 0.9997444 | Additional lipoprotein component |
| AP53_312 | 0.007024516 | 9.453151372 | 0.000910871 | 0.976444248 | 0.9997444 | hypothetical protein |
| AP53_885 | 0.004109786 | 7.980501176 | 0.000882033 | 0.976820005 | 0.9997444 | ABC transporter, ATP-binding/perm |
| AP53_218 | -0.00336757 | 10.12084754 | 0.000825352 | 0.977576932 | 0.9997444 | Surface exclusion protein |
| AP53_1598 | 0.00925038 | 8.199124008 | 0.000807957 | 0.977814418 | 0.9997444 | DNA-directed RNA polymerase delt |
| AP53_1098 | -0.006730483 | 4.219661802 | 0.000792632 | 0.978025763 | 0.9997444 | Mobile element protein |
| AP53_93 | 0.004057432 | 8.777350987 | 0.000761012 | 0.978468408 | 0.9997444 | Pyrroline-5-carboxylate reductase ( |
| AP53_58 | 0.004960125 | 10.54885701 | 0.000715969 | 0.979115159 | 0.9997444 | LSU ribosomal protein L6p (L9e) |
| AP53_882 | -0.004737002 | 8.88071783 | 0.000704179 | 0.979287791 | 0.9997444 | Pseudouridylate synthases, 23S RN |
| AP53_137 | -0.008287326 | 6.003092304 | 0.000701252 | 0.979330867 | 0.9997444 | FIG01114743: hypothetical protei |
| AP53_423 | -0.003777042 | 10.05280575 | 0.000671836 | 0.979768917 | 0.9997444 | Glycosyltransferase LafA, responsib |
| AP53_469 | -0.005252022 | 9.808365257 | 0.000669053 | 0.979810851 | 0.9997444 | FIG01116444: hypothetical protei |
| AP53_1784 | 0.006389204 | 7.840220968 | 0.000666785 | 0.979845096 | 0.9997444 | Phage transcriptional repressor |
| AP53_1860 | 0.005040883 | 6.937017296 | 0.000661471 | 0.97992555 | 0.9997444 | ABC transporter, permease protein |
| AP53_1185 | -0.009479527 | 1.757692781 | 0.000617975 | 0.980596638 | 0.9997444 | FIG01116522: hypothetical protei |
| AP53_687 | 0.004111075 | 8.680041329 | 0.000611881 | 0.980692519 | 0.9997444 | Uracil permease |
| AP53_563 | -0.006936443 | 10.59655174 | 0.000581322 | 0.981180717 | 0.9997444 | ATP synthase alpha chain (EC 3.6.3. |
| AP53_1567 | 0.004897003 | 8.47442752 | 0.000513954 | 0.982304541 | 0.9997444 | Xaa-Pro dipeptidyl-peptidase (EC 3. |

| | | | | | | |
|---|---|---|---|---|---|---|
| AP53_720 | 0.007698411 | 6.455922938 | 0.000460501 | 0.983249818 | 0.9997444 | FIG01114610: hypothetical protei |
| AP53_542 | 0.001980571 | 9.748023883 | 0.000455787 | 0.983335759 | 0.9997444 | DNA gyrase subunit B (EC 5.99.1.3) |
| AP53_1639 | -0.003312977 | 7.323090863 | 0.000440537 | 0.983616865 | 0.9997444 | Cysteinyl-tRNA synthetase (EC 6.1.1 |
| AP53_1611 | 0.004726268 | 4.559297688 | 0.000425338 | 0.983901932 | 0.9997444 | Transcriptional regulator DegU, Lux |
| AP53_1587 | 0.003668865 | 5.648149901 | 0.000420923 | 0.983985684 | 0.9997444 | FIG01114549: hypothetical protei |
| AP53_1796 | -0.006046861 | 4.779736854 | 0.000415265 | 0.984093653 | 0.9997444 | DNA primase, phage associated |
| AP53_914 | -0.005536632 | 8.560464416 | 0.000373098 | 0.984922739 | 0.9997444 | Citrate lyase beta chain (EC 4.1.3.6) |
| AP53_1231 | 0.004714876 | 6.946548077 | 0.000353563 | 0.985322711 | 0.9997444 | ABC transporter membrane-spanni |
| AP53_1534 | 0.003115596 | 8.554004643 | 0.000305466 | 0.986357376 | 0.9997444 | Sucrose operon repressor ScrR, LacI |
| AP53_1241 | 0.004137275 | 9.750516897 | 0.00028332 | 0.986861161 | 0.9997444 | Hypothetical protein YggS, proline |
| AP53_648 | 0.004898787 | 7.381473531 | 0.000281105 | 0.986912637 | 0.9997444 | Phage antirepressor protein |
| AP53_1752 | 0.002661366 | 6.705251976 | 0.000239832 | 0.987911438 | 0.9997444 | Formiminoglutamase (EC 3.5.3.8) |
| AP53_302 | -0.001905018 | 9.263777797 | 0.000239009 | 0.98793219 | 0.9997444 | FIG01114725: hypothetical protei |
| AP53_746 | 0.002614037 | 8.06732286 | 0.000230628 | 0.988145642 | 0.9997444 | Methionine biosynthesis and trans |
| AP53_1853 | -0.00173789 | 7.660663908 | 0.000224305 | 0.988309258 | 0.9997444 | DNA recombination and repair pro |
| AP53_1717 | 0.002243453 | 6.665222329 | 0.000215138 | 0.988550611 | 0.9997444 | Transaldolase (EC 2.2.1.2) |
| AP53_1780 | -0.003131459 | 8.841926028 | 0.000195465 | 0.98908662 | 0.9997444 | Multidrug resistance protein B |
| AP53_1292 | -0.002419368 | 5.910146344 | 0.000193221 | 0.989149431 | 0.9997444 | DNA-binding response regulator, Ar |
| AP53_75 | -0.001898253 | 7.726886869 | 0.000181342 | 0.989488233 | 0.9997444 | Zinc ABC transporter, ATP-binding |
| AP53_20 | 0.002940773 | 7.070122534 | 0.000167299 | 0.989903423 | 0.9997444 | Acyl carrier protein |
| AP53_1541 | -0.001733439 | 7.953481663 | 0.000165437 | 0.98995976 | 0.9997444 | Magnesium and cobalt transport pr |
| AP53_183 | -0.001871056 | 9.708970245 | 0.000161584 | 0.99007736 | 0.9997444 | Glycerol-3-phosphate dehydrogena |
| AP53_1218 | -0.001330678 | 8.972107627 | 0.000146827 | 0.990541297 | 0.9997444 | FIG006988: Lipase/Acylhydrolase v |
| AP53_150 | -0.003590776 | 12.31151895 | 0.000135113 | 0.990926431 | 0.9997444 | Glycine betaine ABC transport syste |
| AP53_1471 | 0.002229926 | 8.431843594 | 0.00013455 | 0.990945355 | 0.9997444 | PTS system, mannose-specific IIB co |
| AP53_689 | 0.002774976 | 9.352042618 | 0.000123984 | 0.991308134 | 0.9997444 | Carbamoyl-phosphate synthase sma |
| AP53_1272 | -0.002124299 | 7.465339886 | 0.000119378 | 0.991471087 | 0.9997444 | Two-component sensor kinase yesM |
| AP53_1643 | 0.00214926 | 8.456063552 | 9.26E-05 | 0.992490014 | 0.9997444 | Polyribonucleotide nucleotidyltrar |
| AP53_1833 | 0.001389762 | 8.861494486 | 9.01E-05 | 0.992588943 | 0.9997444 | LSU ribosomal protein L9p |
| AP53_1577 | 0.002029156 | 7.414925905 | 7.81E-05 | 0.993101772 | 0.9997444 | Uridine phosphorylase (EC 2.4.2.3) |
| AP53_185 | -0.001894876 | 8.235030862 | 6.47E-05 | 0.993721698 | 0.9997444 | Lipid A export ATP-binding/permea |
| AP53_611 | -0.002656916 | 4.925121625 | 6.41E-05 | 0.993752091 | 0.9997444 | hypothetical protein |
| AP53_368 | -0.001781912 | 9.824903804 | 6.17E-05 | 0.993870076 | 0.9997444 | 67kDa Myosin-crossreactive strept |
| AP53_191 | -0.001040868 | 8.470630369 | 6.02E-05 | 0.993941173 | 0.9997444 | DUF124 domain-containing protei |
| AP53_192 | 0.001091559 | 9.179226164 | 5.21E-05 | 0.994365931 | 0.9997444 | Glutamyl-tRNA synthetase (EC 6.1.1 |

| | | | | | | |
|---|---|---|---|---|---|---|
| AP53_rna36 | -0.003154842 | 4.138285412 | 4.55E-05 | 0.994736986 | 0.9997444 | tRNA-Ala-TGC |
| AP53_131 | 0.000708672 | 10.18789975 | 4.16E-05 | 0.994967602 | 0.9997444 | Adenylosuccinate synthetase (EC 6. |
| AP53_277 | -0.000596867 | 9.564369913 | 4.11E-05 | 0.994995655 | 0.9997444 | FIG004453: protein YceG like |
| AP53_828 | 0.001199084 | 9.672546088 | 3.87E-05 | 0.99514137 | 0.9997444 | Oxygen-insensitive NAD(P)H nitrore |
| AP53_1478 | 0.001192874 | 10.06615235 | 3.62E-05 | 0.995304017 | 0.9997444 | Acetyl-coenzyme A carboxyl transfe |
| AP53_857 | 0.000749225 | 8.139744154 | 3.12E-05 | 0.995639359 | 0.9997444 | Glutamine amidotransferase, class I |
| AP53_1738 | 0.00106246 | 10.31251749 | 2.84E-05 | 0.995840561 | 0.9997444 | ATP-dependent Clp protease ATP-bi |
| AP53_19 | -0.000846067 | 7.876750887 | 2.75E-05 | 0.995904194 | 0.9997444 | Phosphate:acyl-ACP acyltransferase |
| AP53_1356 | -0.001340405 | 3.936521777 | 2.29E-05 | 0.996266221 | 0.9997444 | phage protein |
| AP53_952 | -0.001541106 | 7.544188929 | 2.08E-05 | 0.996441714 | 0.9997444 | Ribosomal RNA small subunit meth |
| AP53_705 | 0.000558925 | 8.668212891 | 1.21E-05 | 0.997289514 | 0.9997444 | 2-dehydropantoate 2-reductase (EC |
| AP53_160 | 0.000975079 | 4.973827245 | 1.05E-05 | 0.997471408 | 0.9997444 | Mobile element protein |
| AP53_87 | 0.000621749 | 3.194816257 | 4.24E-06 | 0.998392713 | 0.9997444 | Late competence protein ComGE, F |
| AP53_1728 | 0.000261936 | 7.033408301 | 3.90E-06 | 0.998458155 | 0.9997444 | Hypothetical protein SPy2060 |
| AP53_74 | -0.000165842 | 6.906016983 | 1.48E-06 | 0.999051322 | 0.9997444 | Transcriptional repressor AdcR for Z |
| AP53_342 | -0.000214621 | 9.409220889 | 9.75E-07 | 0.999229333 | 0.9997444 | FIG01114229: hypothetical protei |
| AP53_rna69 | 0 | -2.070645075 | 0 | 1 | 1 | 5S RNA |

lactonase UlaG (EC 3.1.1.-) (L-ascorbate utilization protein G)
rter, substrate binding periplasmic protein MalE

447

.2.1.85)
ated

.6.3.14)

.2.1.70)

o/CI family
mily

essor protein
n

family (EC 2.7.1.69)
omponent (EC 2.7.1.69)

: Ffh SRP54 (TC 3.A.5.1.1)

nponent (EC 2.7.1.69) / PTS system, lactose-specific IIC component (EC 2.7.1.69)
n
cetyltransferase (EC 2.3.1.-)

o-8-oxoguanine-triphosphatase) (EC 3.6.1.-)
rter, permease protein MalF

h PTS regulation domain, SPy0181 ortholog
em specificity subunit
.6.3.14)
(EC 3.1.2.6)

xyvinyltransferase (EC 2.5.1.7)
regulon ComD
n
3)
n

n
P-binding protein
C 2.1.3.3)
n ArgR, repressor of arg regulon

ates the switch between cylindrical and septal cell wall synthesis by re-localization of PBP1
porter protein
C 2.1.2.5) @ Glutamate formyltransferase
n

component
ransporting ATPase (EC 3.6.3.3) (EC 3.6.3.5); Copper-translocating P-type ATPase (EC 3.6.3.4)
LacA subunit (EC 5.3.1.26)
C 6.3.4.3)
n
protein, GcvH-like
ve
binase

n
eaminase (EC 4.3.1.4)
n
4)
1.14.-.-)
n SufB
nyltransferase (EC 2.1.1.176)
.6.3.14)
ily
e (EC 5.1.3.-)
n
se (EC 2.5.1.31)
yltransferase N (EC 2.1.1.-)
MutR family

clustered with an amidotransferase
.1)

e YggF
n

J of predicted pyridoxine ECF transporter
em, permease component
.6.3.14)
ing protein)
iacylglycerol kinase catalytic domain]

ohydrolase (NADGH) (EC 3.2.2.5)

TE) family transporter

ated
nzyme (EC 1.97.1.4)
mponent (EC 2.7.1.69) / PTS system, fructose-specific IIB component (EC 2.7.1.69) / PTS system, fructose-specific IIC component (EC 2.7.1.69)
ly scaffold protein for SUF system, SufE2
RNA; SSU rRNA
n

ufS subfamily
: (EC 3.1.26.5)
n
protein

xidoreductase in cluster with COG2110
(EC 5.2.1.8)
ly IIB component

transferase (EC 2.7.7.9)
e protein
n
4.2.1.70)

vith transport systems (COG1739)
RNA; SSU rRNA

RNA; SSU rRNA
.6.3.14)

o/CI family

ssor HrcA

transferase (EC 2.7.7.9)
.6.3.14)
n
0)
(EC 4.1.2.40)
n
RNA; SSU rRNA
operon
n
nyl-diaminopimelate desuccinylase and related deacylases
k protein
omponent (EC 2.7.1.69)

luronic acid synthesis (EC 1.1.1.22)

ulator, ArpU family
C 4.2.1.10)
.)
em, ATPase component
dine synthase B (EC 4.2.1.70)
o cobyric acid synthase

n

₹NA; SSU rRNA
ase (EC 3.5.99.6)
drolase (EC 3.5.2.12)
asmic-binding protein SitA
in
₹NA; SSU rRNA
sine phosphatase (EC 3.1.3.48)

omponent (EC 2.7.1.69)
ntial for anaerobic expression of the ADI pathway, Crp/Fnr family
NusA
utative), YggT family
5) / Deoxyguanosine kinase (EC 2.7.1.113)
anine amidase (EC 3.5.1.28)
ntaining CBS domains

ADP-ribose binding module

n
ntial role in cell wall biosynthesis

n
)
ulator (Mga)

Z (SmeZ)

ily

in

n
M3_1856/SPs1852

component (EC 2.7.1.69)
BC transporter protein ProX (TC 3.A.1.12.1) / Osmotic adaptation
omponent (EC 2.7.1.69)
ne permease (SagH)

protein (class beta)
otein J
acid-binding protein

nily
repressor
LacB subunit (EC 5.3.1.26)

5' exonuclease)

hage associated

rter, permease protein MalG

sferase (EC 2.7.7.3)

n

GTPase RbgA (B. subtilis YlqF)

n
r, associated with ferric iron transporter, SPy1062 homolog
ily

NA; LSU rRNA
.6.3.14)

n ArgR, repressor of arg regulon

e of SIR2 family
EC 2.5.1.90) / Dimethylallyltransferase (EC 2.5.1.1) / (2E,6E)-farnesyl diphosphate synthase (EC 2.5.1.10) / Geranylgeranyl diphosphate synthase (EC 2.
n
NA; LSU rRNA
ml
J2 of predicted pyridoxin-related ECF transporter
6.1.13)
)
nily
ctase MsrB (EC 1.8.4.12)

n
ated
otein CzcD
NA; LSU rRNA
idine synthase A (EC 4.2.1.70)
n

tor
unit (EC 3.1.11.6)
eptidase
.5.1.75)

e (EC 2.7.3.-)

n
ecific IIC component (EC 2.7.1.69) / PTS system, maltose and glucose-specific IIB component (EC 2.7.1.69) / PTS system, maltose and glucose-specific
BC transporter ATP-binding protein ComA

rt system, ATPase component
omponent (EC 2.7.1.69)
embrane-spanning permease protein MsmG
E (TC 3.A.5.1.1)

T (COG4708) of predicted queuosine-regulated ECF transporter
(EC 2.1.2.9)

NA; LSU rRNA

h Ecs transporter (in Streptococci)

(EC 3.1.3.27)
NA; LSU rRNA

n

d)
nily protein)
nsferase (EC 2.1.1.207)

ed protein
NA; LSU rRNA

571856) SAR1694
3.3)
n
nce regulon ComE
quired for reduction of Ribonucleotide reductase class Ib
atase family protein
ein A
n
n
EC 2.5.1.6)
R (transcriptional repressor)
lmuramyl-(pentapeptide) pyrophosphoryl-undecaprenol N-acetylglucosamine transferase (EC 2.4.1.227)

amily
EscB
sociated
ase I
mily
(EC 4.1.2.40)

n

, beta chain (EC 4.1.1.41); Oxaloacetate decarboxylase beta chain (EC 4.1.1.3)
C 3.2.2.-)
K (SpeK)
y membrane protease subunit YbbK

r, malate (EC 2.7.3.-)
rotein CutC

stem
C 6.3.4.3)
ent alpha-subunit (EC 1.2.4.-)

n
ress-responsive transcriptional regulator

ate-binding protein

n
e IIc (The first GCT-Ala is overlapped with the last codon of the former half interrupted protein)
eotidohydrolase (EC 3.6.1.23)

pilin backbone protein T-antigen
gulator VraR

SufD

n
lneuraminate transport system permease protein 1
ort system, permease component
r SA14-24
n

nily
dA, haloacid dehalogenase-like phosphatase family
ein

n
phocoenzyme-A synthase (EC 2.7.8.25)

n
orter permease component potC (TC_3.A.1.11.1)
nily
g protein; Streptolysin S export protein (SagG)
regulon ComD

n
n
ly

nily
in YvcR
or threonylcarbamoyladenosine t(6)A37 formation in tRNA
EzrA
5)

nding protein

d)
pseudouridine synthase D, Bacillus subtilis YhcT type

A-independent DNA replication

n
MutS2

n

ly

Antiporter
6.3.14)
ate O-acyltransferase PlsY

n
C 4.1.2.4)
rS / deletion of T at 1407 causing a early stop codon at 469th aa)

n
n
.6.3.14)
cus), ATP-binding protein

nidine putrescine-binding protein PotD (TC 3.A.1.11.1)

ily
693 of energizing module of predicted ECF transporter
(EC 4.1.1.31)

n

nase component (EC 2.6.1.85) / Aminodeoxychorismate lyase (EC 4.1.3.38)
se component YidC, OxaA protein

n

4-24
ng permease - glutamine transport

nase, malate (EC 2.7.3.-)
n
nding protein SitB
sferase (EC 2.1.1.-)

ogenase (NADP+) (EC 1.5.1.5) / Methenyltetrahydrofolate cyclohydrolase (EC 3.5.4.9)

:, implicated in cell division based on FtsL cleavage

ic cystine-binding protein FliY
7.1.60)
in uup
otein
ociated with ferric iron transporter, SPy1061 homolog
BC transporter ATP-binding protein ComA
o (aerobic), alpha subunit (EC 1.17.4.1)
uctase (EC 1.1.1.100)
a chain (EC 6.1.1.20)

1.1)
n

y
protein Smf possibly involved in DNA uptake

e (EC 5.1.3.4)

EC 3.2.1.93)

nzyme (EC 1.97.1.4)

ogenase (EC 1.1.1.205)
unogenic protein
ent proteases, putative molecular chaperone
ctose operon, DeoR family

.1.35)
of biotin ECF transporter
4.1.21) / Autoinducer-2 production protein LuxS

asmic phosphate-binding protein PstS (TC 3.A.1.7.1)

EC 2.4.2.1)
talytic subunit (EC 1.3.3.1)

1.1.22)
abinase
n

n
lcohol forming, (R)-acetoin-specific (EC 1.1.1.4) / Acetoin (diacetyl) reductase (EC 1.1.1.5)

nase (EC 1.1.1.125)
Streptopain) (EC 3.4.22.10) / Streptococcal pyrogenic exotoxin B (SpeB)

acid biosynthesis, DltR
sonate-7-phosphate synthase I beta (EC 2.5.1.54)

Ps0857

access of DNA to ComEA, FIG038316
n
nily
(EC 2.7.7.7)

27)

idine synthase A (EC 4.2.1.70)

)
n
n
ransporter, ATP-binding protein
omponent (EC 2.7.1.69) / PTS system, trehalose-specific IIC component (EC 2.7.1.69) / PTS system, trehalose-specific IIA component (EC 2.7.1.69)

ComX
port system permease protein ProV (TC 3.A.1.12.1)

3)
n

C 1.-.-.-)
se (EC 1.1.1.133)
n
with beta-lactamase, GntR family
em, restriction subunit R (EC 3.1.21.3)

ribosomal protein L31p, zinc-independent

se (EC 2.3.2.5)
ing], amidotransferase subunit (EC 6.3.5.2) / GMP synthase [glutamine-hydrolyzing], ATP pyrophosphatase subunit (EC 6.3.5.2)

21.89)
gene causing stop codon at 85th aa
phate reductase (EC 1.17.4.2)

ge chain (EC 6.3.5.5)

ng protein FtsE (TC 3.A.5.1.1)
t (TC 3.A.5.1.1)

: 2.7.7.7)
:ein C (EC 1.6.4.-)
sphate 3-phosphatidyltransferase (EC 2.7.8.5)

n

qC
:omponent (EC 2.7.1.69)
n enzyme TrmE

se (EC 3.4.16.4)
unit (EC 2.7.7.7)
in
ily 25
n
omboid family)
essor of hyaluronate and KDG utilization
ily
nerly thought to be RNase BN
bs2121
protein
e (EC 2.7.8.7)
ein EcsA
e YoaA
:ase (EC 1.1.1.34)

n - phage associated

ed
ine ligase @ tRNA-dependent lipid II-Ser--L-alanine ligase
n

ı

ɩ), (p)ppGpp synthetase I
iated with purine metabolism
xin protein

sporter
ıage associated
n
ɛxport protein
bosomal protein L32p, Firmicutes subfamily
n
C 2.4.2.-)

component (EC 2.7.1.69)
n
ɛA
ase PrpC, regulation of stationary phase
(EC 3.1.3.2)

4)
ɔtein
n
BC transporter permease protein ComB

6) @ Acetyl-CoA acetyltransferase (EC 2.3.1.9)
like
2.7.4.6)
ɛ (EC 3.6.3.4)

2.25)
)
n
ional attenuator LytR-CpsA-Psr, subfamily F2 (as in PMID19099556)
n
ption termination protein NusA
of acetoin dehydrogenase (EC 1.8.1.4)

aining kinase
export of O-antigen, teichoic acid lipoteichoic acids

C (SpeC); _Toximoron (Superantigen)

enase [NAD] (EC 1.2.1.24); Succinate-semialdehyde dehydrogenase [NADP+] (EC 1.2.1.16)
osynthesis lipoprotein ApbE

n

nily IIIA

xaloacetate decarboxylase; Biotin carboxyl carrier protein

rotein YabA
peptide-transferase (EC 2.7.8.13)
rier protein (EC 4.1.3.6)

/ protein

r yesN, associated with MetSO reductase

1.2)
2.1.1.-)

sferase TrmN6 (EC 2.1.1.223)

n
oseudouridine synthase D, Bacillus subtilis YjbO type
sglycosylase (EC 2.4.1.129) (EC 3.4.-.-)
n
.2.1)

n
.3)

.15)
e (EC 5.2.1.8); Proteinase maturation protein (RopA)
A

IIB component (EC 2.7.1.69) / PTS system, beta-glucoside-specific IIC component (EC 2.7.1.69) / PTS system, beta-glucoside-specific IIA component (
n
type 1
hosphoribosyltransferase domain; protein YhgH required for utilization of DNA as sole source of carbon and energy
n
ccinocarboxamide synthase (EC 6.3.2.6)

in RecT (prophage associated)

hage associated

n
6.3.8)
eG
ne permease (SagI)
ane permease protein ZnuB
xaloacetate decarboxylase; Biotin carboxyl carrier protein
, required for cell wall metabolism and coordination of cell division with DNA replication

protein DnaA

ng enzyme, evidenced by COGnitor; PaaD-like protein (DUF59) involved in Fe-S cluster assembly
se component YidC, short form OxaI-like

le for the formation of Gal-Glc-DAG
nponent (EC 2.7.1.69) / PTS system, sucrose-specific IIC component (EC 2.7.1.69) / PTS system, sucrose-specific IIA component (EC 2.7.1.69)
n
in

I (anaerobic), activating protein (EC 1.97.1.4)
nine(1519)-N(6))-dimethyltransferase (EC 2.1.1.182)
e competence protein ComEB

g protein OppF (TC 3.A.1.5.1)

C 2.1.2.1)

Dps / Iron-binding ferritin-like antioxidant protein / Ferroxidase (EC 1.16.3.1)
nase (EC 3.5.4.-)
nage associated
ein

.1.2)
08
7.1.60)

2) / Folylpolyglutamate synthase (EC 6.3.2.17)

n (SagE)

n
NrdI
Na+ or H+) symporter

n
1.1)
2.4.2.29)
n

e in RNA processing)

P2)
n
ated

3.2.4)
3)
gizing module of ECF transporters
n

de-lysine N(6)-alanyltransferase (EC 2.3.2.10) (EC 2.3.2.-)

FIG068335
2.7.4.6)
ferase component (EC 4.1.3.27) @ Para-aminobenzoate synthase, amidotransferase component (EC 2.6.1.85) # TrpAb@PabAb
abilizes FtsL against RasP cleavage

in
n
se (EC 2.4.2.22)
tein (putative)

n
n
@ SSU ribosomal protein S14p (S29e), zinc-independent
transferase (EC 2.4.1.-)

in
nding protein

se subunit C (EC 6.3.5.6) @ Glutamyl-tRNA(Gln) amidotransferase subunit C (EC 6.3.5.7)
.1.1)
ining
ltransferase (EC 2.3.1.51)
ein
in
idine kinase CiaH (EC 2.7.3.-)
, periplasmic substrate-binding component (TC 3.A.1.5.2)

2.1.86)
ding protein, similarity with At5g48545 and yeast YDL125C (HNT1)
erase [decarboxylating] (EC 2.4.2.19)
of predicted niacin ECF transporter
rase RumA (EC 2.1.1.-) ## LSU rRNA m(5)U1939
dine synthase D (EC 4.2.1.70)
MraW methylase family (EC 2.1.1.-)
n
.12.4)
.3.4)
ng protein, spy1790 homolog
chain (EC 6.1.1.20)

C 5.3.1.8)
1

e translocation permease
nily
binding protein implicated in transcription termination

n

n

access of DNA to ComEA, FIG007487

ated

otein
n
n
pecific nuclease) (EC 3.1.30.-)
ession IbrA

inator, BglG family
4.-)
boid family)
4.2.1.70)

tein B
n

protein FhuC (TC 3.A.1.14.3)
(former ThiI)
(EC 2.7.7.41)
d phosphatase
IN adenylyltransferase (EC 2.7.7.2)

ein

ne associated protein

RlmH
iosynthesis in gram-positives

copper transport operon
mily
r

nomethyl modification enzyme GidA
nbinase

em, specificity subunit S (EC 3.1.21.3)
C 5.3.1.9)

sglycosylase (EC 2.4.1.129) (EC 3.4.-.-) / Penicillin-binding protein 1A/1B (PBP1)

uctase

BC transporters with duplicated ATPase domains

(EC 6.1.1.14)

egulator CiaR

in
component (EC 2.7.1.69)

C 5.1.3.1)
ein

ike family
ferase (EC 2.1.1.33)

B
ated
eral energizing module of ECF transporters

7.1.144)

n

ribosomal protein L33p, zinc-independent
rboxylase catalytic subunit (EC 4.1.1.21)

e murein endopeptidase
erase [decarboxylating] (EC 2.4.2.19)
rotein

nase (ATP-binding subunit)

ıbinase

ıily protein

ı.2.1.46)

D (SagD)
g protein
erase, alpha subunit (EC 2.8.3.8)

uctase (EC 1.1.1.100)
IS-, AP-, NP- specific)
ıily

G (TC 3.A.5.1.1)

: 3.6.4.12)
n

family

protein SufC
lneuraminate-binding protein

transferase/alpha-L-Rha alpha-1,3-L- rhamnosyltransferase (EC 2.4.1.-)
: Protein (RALP)
osphohydrolase (EC 3.1.5.1)
ort system, permease 2 component
eptidase
1.3.25)

nponent (EC 2.7.1.69)
: [C1]
omolog
: (EC 4.3.1.17)
.8)
family
otein PhoH, predicted ATPase
ers

ase protein PstA (TC 3.A.1.7.1)
ional attenuator LytR-CpsA-Psr, subfamily F1 (as in PMID19099556)

7.1.144)
yltransferase E (EC 2.1.1.-)

ily

J of riboflavin ECF transporter
n
-specific IIC component (EC 2.7.1.69)
n
ein) -possibly involved in cell wall localization and side chain formation of rhamnose-glucose polysaccharide
nine-binding protein/permease protein
ılator MntR

C (SagC)

ɔn protein PhnA
C (TC 3.A.5.1.1)

aining multiple CBS domains
related protein

maltase) (EC 2.4.1.25)
n

family
ein

nerase (EC 5.1.3.13)
ng permease - macrolide efflux
ated)
etence MecA
rotein DppD (TC 3.A.1.5.2)
n

phosphate dehydrogenase (EC 1.2.1.12) / Plasmin(ogen) receptor
chenan operon, BglG family

utamyl-2,6-diaminopimelate--D-alanyl-D-alanine ligase (EC 6.3.2.10)

n

sm regulator CmbR, LysR family
repressor
n

ein
e)
contains S4-like RNA binding domain

molog
d for threonylcarbamoyladenosine t(6)A37 modification in tRNA
protein

n
n
nding protein
unit (EC 3.1.11.6)
ein
(EC 3.4.19.3)
n
ocessing protease
017

rotein PstB (TC 3.A.1.7.1)
e 2-epimerase (EC 5.1.3.9)

gase (EC 6.3.2.8)
DNA transport
ctase MsrA (EC 1.8.4.11)

i), bacterial type

mrA

nily

n

orter permease component PotB (TC 3.A.1.11.1)
n
G (SpeG)
ribosomal protein S18p, zinc-independent
tein
Family)

sor SPs0273
d GTPase EngC
n
chain (EC 4.1.1.3)

tein PaaD, thioesterase
21.-)

rotein

ng protein drrA
(EC 2.1.1.72)
;
n
tein
tor, PadR family
(SagF)

ne synthase, synthetase subunit (EC 6.3.5.3) / Phosphoribosylformylglycinamidine synthase, glutamine amidotransferase subunit (EC 6.3.5.3)

zyme in heat shock gene cluster, similarity with CPO of BS HemN-type
nsferase, involved in cell wall biogenesis

4.23.36)
ase (EC 2.3.1.-)
C 2.4.1.-)
-specific IID component (EC 2.7.1.69)
tionally linked with Alanyl-tRNA synthetase
n
inhibitor

n

ily

n

ase (EC 4.1.1.23)

unit-related enzymes

n

n
2) / Folylpolyglutamate synthase (EC 6.3.2.17)

lneuraminate transport system permease protein 2
.9)
-specific IIB component (EC 2.7.1.69)
n

opic repressor codY
em, DNA-methyltransferase subunit M (EC 2.1.1.72)

2.1.22)

e)
o (aerobic), alpha subunit (EC 1.17.4.1)
MutR family
in

e deacetylase (EC 3.5.1.-)
n

C 2.7.7.7)

epressor
n

n
ase protein DppB (TC 3.A.1.5.2)

putative peptidoglycan hydrolase
in

se (EC 3.4.16.4)
/
R family

ent beta-subunit (EC 1.2.4.-)
(EC 6.1.1.14)

tory protein PhoU
r protein FtsY (=alpha subunit) (TC 3.A.5.1.1)
thase, KASII (EC 2.3.1.41)

d)
nsferase

nobacillus protein (1944168)
ss II (EC 4.1.2.13)
ase (EC 2.7.6.1)

rboxylase ATPase subunit (EC 4.1.1.21)
transferase rlmB (EC 2.1.1.-) ## LSU rRNA Gm2251
tein RecO

nding subunit ClpX
(aerobic), beta subunit (EC 1.17.4.1)
e 3; Mitogenic factor 3
n
(aerobic), beta subunit (EC 1.17.4.1)
se (EC 3.4.16.4)
3.6.4.12)
protein
yl-diaminopimelate desuccinylase and related deacylases

in

' subunit (EC 2.7.7.6)
Toximoron (Other)
(EC 3.6.1.-)
igase @ tRNA-dependent lipid II--L-serine ligase

ily
thetase (EC 2.5.1.6)
ylase (EC 3.2.2.23)

n
11.4)
n
uridyltransferase (EC 2.7.7.23) / Glucosamine-1-phosphate N-acetyltransferase (EC 2.3.1.157)
ein

inding subunit ClpE

n
)
;rammed frameshift-containing
ulose-uronate ketol-isomerase (EC 5.3.1.17)

n
n
n

3.14)
;e (EC 6.3.4.13)
eme from hemoglobin to apo-SiaA/HtsA

;e 2; Mitogenic factor 2
n
'PcrA
nily
rotein DppF (TC 3.A.1.5.2)
n

n
nage associated

iacylglycerol kinase catalytic domain]

.4.2.10)

oside utilization operon, GntR family
access of DNA to ComEA
rotein precursor (TC 3.A.1.14.3)

ated

EC 3.8.1.2)
rotein
e-binding protein

e (EC 2.7.1.45)
ressor MalR, LacI family

aminidase (EndoS)

ulator (AT-rich DNA-binding protein)

yltransferase (EC 2.1.1.171) ## SSU rRNA m(2)G966

se (EC 2.3.3.10)

ase subunit 2 (EC 6.1.1.13)
membrane permease protein SitD
protein
nsferase (EC 2.7.7.18)
sine (m7G) methyltransferase GidB

n
e (EC 3.4.21.-)
n
VA
glutamate ligase (EC 6.3.2.9)

)4 of energizing module of predicted ECF transporter
nine-binding protein/permease protein
hage associated
cetyl-CoA carboxylase
e aminotransferase [isomerizing] (EC 2.6.1.16)
ulator (Mga)
EC 2.7.7.7)
ase
n

/

)
tein
n
component (EC 2.7.1.69)

utamate--L-lysine ligase (EC 6.3.2.7)
ciated
transferase (EC 2.1.2.2)

c

rase (EC 5.3.1.-)
ase subunit 1 (EC 6.1.1.13)

nine reductase (EC 1.1.1.158)

plasmic oligopeptide-binding protein OppA (TC 3.A.1.5.1)

e lic/interrupted by large phage

[C1]
[illegible] of predicted pantothenate ECF transporter

ing protein), spy1791 homolog
[illegible]agment)

[illegible]29)

EC 3.2.2.20)
[illegible]bly involved in cell wall localization and side chain formation of rhamnose-glucose polysaccharide
(EC 1.1.1.25)
[illegible]ase [FMN] (EC 1.3.1.9)

[illegible]C, regulator of stationary phase

(EC 2.4.2.10)
protein dnaD

[illegible]rdI
[illegible]otein implicated in the recycling of the 50S subunit (S4 paralog)

[illegible]5.3.2.-); Xylose transport system permease protein xylH
[illegible]ein

n
of thiamin ECF transporter
eeN

.1.8)

with unspecified monosaccharide ABC transport system
n

d for biogenesis of 30S ribosome subunit
iphosphatase RdgB (dHAPTP, dITP, XTP-specific) (EC 3.6.1.15)
ycan synthetase] (EC 2.4.1.129)

anine transfer protein DltD
pecific nuclease) (EC 3.1.30.-)
3.2.1.26)

ession IbrB
acid-binding protein
cus), permease protein
thetase (EC 6.3.2.5)

4)

subunit (EC 2.7.7.6)

C 5.3.1.6)
n

rase subunit B (EC 2.8.3.9)

:ein F (EC 1.6.4.-)
ng protein YchF

d)
otein

ive channel

n
ctase MsrA (EC 1.8.4.11) / Peptide methionine sulfoxide reductase MsrB (EC 1.8.4.12)
76, Firmicutes subgroup
12)
(EC 2.7.1.35)

, contains double-stranded beta-helix domain

n
em, ATP-binding protein OpuAA (EC 3.6.3.32)
3.14)

otein

.1.10)
ase)

rotein PstB (TC 3.A.1.7.1)
V family
n

ain protein (Bsu YtpR)
unction, cellobiose operon
ransferase (EC 2.6.1.42)
in TrkA

lar to C-terminal domain of ribosome protection-type Tc-resistance proteins
tase (EC 1.2.1.41)
erase alpha chain (EC 6.4.1.2)
anM/pseudogene causing stop codon at 177th aa

se
ase A, Bacillus type
n
nvolved in rRNA and/or ribosome maturation and assembly

superfamily
n

ransporter, permease protein
ase protein DppC (TC 3.A.1.5.2)
uclease RecJ (EC 3.1.-.-)
nyldihydropteridine pyrophosphokinase (EC 2.7.6.3)

n
erythritol kinase (EC 2.7.1.148)
protein PotA (TC 3.A.1.11.1)
cid biosynthesis FabT

EC 2.4.2.9)

ase (EC 2.4.2.11)

21)
n
se (EC 3.4.16.4)
B (SagB)
n
rase subunit A (EC 2.8.3.9)

(EC 4.3.1.17)
5.1.1.6)

or

e)

ase (EC 2.7.6.1)

4.-)
n

erone GroEL

ptional regulator NrdR
ble histidine kinase

;)

porter ATPase

DNA receptor

Phosphoribosylaminoimidazolecarboxamide formyltransferase (EC 2.1.2.3)
drolase (EC 3.5.2.12)
e family protein
oside utilization, GntR family

rotein
ease protein

a subunit (EC 2.7.7.6)
ase (EC 4.1.1.33)
ical protein
olytic subunit (EC 3.4.21.92)
.1.7)

.5)

)
nily
arboxylase (EC 4.1.1.36)
9.1.-)

osyltransferase (EC 2.4.1.-)
015

de-3-phosphate dehydrogenase (NADP) (EC 1.2.1.9)
n

ein NusG
); Acetaldehyde dehydrogenase (EC 1.2.1.10)
ase protein PstC (TC 3.A.1.7.1)

[illegible]
ZinT
egulator
6) @ Acetyl-CoA acetyltransferase (EC 2.3.1.9)
.7.21) (EC 2.7.7.25)
M (EC 2.7.3.-)
deacetylase (EC 3.5.1.25)

ns nucleotide-binding domain of DisA bacterial checkpoint controller
yvinyltransferase (EC 2.5.1.7)

PC Superfamily
B (SagA)

1)
lucosaminyl 1-phosphate transferase (EC 2.7.8.-)

g protein ParB / Stage 0 sporulation protein J
em
@ SSU ribosomal protein S14p (S29e), zinc-dependent

nage associated
se protein MsbA

LacB subunit (EC 5.3.1.26)
orT family
ase protein 2
C 2.4.2.9) / Pyrimidine operon regulatory protein PyrR

protein
HI0095 (UbiE paralog)

4)
n

phage associated
2.6) (EC 2.7.1.41)

se subunit A (EC 6.3.5.6) @ Glutamyl-tRNA(Gln) amidotransferase subunit A (EC 6.3.5.7)

n

ein CcdA homolog, associated with MetSO reductase
access of DNA to ComEA
PnuC-like

in / Small-conductance mechanosensitive channel
omponent (EC 2.7.1.69)

lactate transferase like, CofD-like
(IdeS) / Mac/ Secreted immunoglobulin binding protein (Sib38)
n
.1.1.38)

ansferase (EC 2.7.7.24)

ase family, subgroup 1

enase (EC 1.1.1.30)
a and tau (EC 2.7.7.7)
nease protein OppC (TC 3.A.1.5.1)

etyltransferase
n
bulin-binding protein (GRAB)

NusB
4.1.1)
containing a URI domain
n

C 5.2.1.8)
sferase PrmC, methylates polypeptide chain release factors RF1 and RF2

nily
inase
component (EC 2.7.1.69)
n

n

I (anaerobic), large subunit (EC 1.17.4.2)
rt system carrier protein
rase subunit B (EC 2.8.3.9)
5.1.1.2)
odiesterase (EC 3.1.4.46)
charide synthesis, competence and biofilm formation Ftr, XRE family

ɔperon, ManO
n
ɔrotein
7.9.1)
(EC 4.1.3.16) @ 2-dehydro-3-deoxyphosphogluconate aldolase (EC 4.1.2.14)
osyltransferase (EC 2.4.2.8)

ite synthase (EC 2.5.1.19)

RNA-binding domain)
chain (EC 4.1.1.3)

ansacylase (EC 2.3.1.39)
nage associated
nily

n1 (EC 3.5.1.28)
..-) (EC 2.7.4.-)
n
n related to cell division

rotein CorA
3)

of predicted cobalamin ECF transporter

arboxylase

:omponent (EC 2.7.1.69)

:C 2.4.2.1); N-Ribosylnicotinamide phosphorylase (EC 2.4.2.1) ## possible

.1.1)
n
)

- and galactosamine-specific IIA component (EC 2.7.1.69)
sphotransferase of PTS system (EC 2.7.3.9)

tein
sglycosylase (EC 2.4.1.129) (EC 3.4.-.-)
;)
C 1.1.3.21)

'A
;izing module of ECF transporters
;)
3)
ɔstrate-binding protein

)
e A; Peptidyl-prolyl cis-trans isomerase (EC 5.2.1.8)

omboid family)
thase, KASIII (EC 2.3.1.41)

nce regulon ComE

C 6.2.1.3)

otein receptor Shr
n

d DNA binding protein

5)

1.1.22)
acid biosynthesis, sensor histidine kinase

n
n
2.1.3.2)
4.11.18)
n

ciated with MetSO reductase
yrophosphatase (EC 3.6.1.1)

LacA subunit (EC 5.3.1.26)

o-8-oxoguanine-triphosphatase) (EC 3.6.1.-)
nbinase
ccess of DNA to ComEA, FIG012620
TrmH family tRNA/rRNA methyltransferase YacO
SecA ATPase RNA helicase (TC 3.A.5.1.1)
(EC 3.2.1.-)

g protein OppD (TC 3.A.1.5.1)
e (EC 2.4.2.7)

se subunit B (EC 6.3.5.6) @ Glutamyl-tRNA(Gln) amidotransferase subunit B (EC 6.3.5.7)
ne cyclo-ligase (EC 6.3.3.1)

protein] dehydratase (EC 4.2.1.-)
ng/permease protein
n

ng/permease protein
ferase (EC 2.1.1.31)
ega subunit (EC 2.7.7.6)

nage associated

n

otein ligase (EC 6.3.4.15)

.-)
.6.1.27)
6)

n

rboxylase (EC 6.3.4.14)
5)

ase H
-.-)

C 1.15.1.1)

ase (EC 3.2.2.16) / S-adenosylhomocysteine nucleosidase (EC 3.2.2.9)

component (EC 2.7.1.-)
n
.4.2)

embrane-spanning permease protein MsmF

n

n
7.6.2)
of pyridoxine metabolism

EC 2.4.2.14)
n

6) @ Acetyl-CoA acetyltransferase (EC 2.3.1.9)

brane protein

(EC 4.1.99.3)
mease protein OppB (TC 3.A.1.5.1)

14)
omponent (EC 2.7.1.69)
specific DNA methylase
ein IsdE

n

/BBR Bacillus subtilis homolog

595 of predicted ECF transporter

: Protein (RALP)

EC 2.7.7.7)

ferase (EC 2.4.99.-)

e component (E2) of acetoin dehydrogenase complex (EC 2.3.1.-)
lease

a2 (EC 3.5.1.28)

nage associated
A N-terminal domain / C-terminal domain of CinA type S
4.-)
ComX

ephospho-CoA transferase (EC 2.7.7.61)
nerase, FMN-dependent (EC 5.3.3.2)
ır utilization related?)
9.1.-)

n ArgR, repressor of arg regulon
n
nain (EC 4.1.1.3)
drolase like
aperone GroES
1.2)

n
ases
n
n

ıease protein
:m, ATPase component

e IIc
ltransferase-isomerase (EC 5.-.-.-)

stem
charide synthesis, competence and biofilm formation Ftr, XRE family
:ductase (EC 1.-.-.-) / Dihydropteridine reductase (EC 1.5.1.34)
e I beta (EC 1.1.1.282)
in
of predicted cobalamin ECF transporter

ıease protein

a subunit (EC 2.7.7.6)

(EC 1.5.1.2)

A-specific
n
le for the formation of Glc-DAG
n

n

14)
4.14.11)

n

..16)
‹R family
n

)
ng permease - glutamine transport
l family
synthase co-transcribed bacterial homolog PROSC

n
port regulator MtaR, LysR family
tein RecF

raC family
protein ZnuC

rotein CorA
ase [NAD(P)+] (EC 1.1.1.94)
with GDSL-like motif
em, permease protein OpuAB / Glycine betaine ABC transport system, glycine betaine-binding protein OpuAC
mponent (EC 2.7.1.69) / PTS system, mannose-specific IIA component (EC 2.7.1.69)
all chain (EC 6.3.5.5)
M (EC 2.7.3.-), associated with MetSO reductase
nsferase (EC 2.7.7.8)

se protein MsbA

coccal antigen
n
1.17) @ Glutamyl-tRNA(Gln) synthetase (EC 6.1.1.24)

3.4.4)

:ductase (EC 1.-.-.-) / Dihydropteridine reductase (EC 1.5.1.34)
:rase beta chain (EC 6.4.1.2)
l
nding subunit ClpA
: PlsX

yltransferase C (EC 2.1.1.52) ## SSU rRNA m2G1207
: 1.1.1.169)

:IG075573

Zn(2+)-responsive expression
n

.5.1.29)

IIA component (EC 2.7.1.69)

EC 2.7.1.69)

| genes | logFC.8hr | logCPM.8hr | statistic.8hr | Pvalue.8hr | FDR.8hr | function |
|---|---|---|---|---|---|---|
| AP53_1665 | -10.45441011 | 6.355545902 | 199.516795 | 1.40E-08 | 2.71E-05 | Streptokinase |
| AP53_1749 | 2.227699898 | 6.229693034 | 38.1354637 | 5.92E-05 | 0.014447645 | Conserved protein |
| AP53_1750 | 2.120873836 | 6.210996007 | 36.73900683 | 7.00E-05 | 0.014447645 | putative cationic amino acid transp |
| AP53_915 | 1.788913635 | 8.680433803 | 36.50579652 | 7.20E-05 | 0.014447645 | Citrate lyase alpha chain (EC 4.1.3.6 |
| AP53_911 | 1.25975678 | 7.402583754 | 36.17867595 | 7.49E-05 | 0.014447645 | Oxaloacetate decarboxylase beta ch |
| AP53_914 | 1.758293536 | 8.560464416 | 35.58486634 | 8.06E-05 | 0.014447645 | Citrate lyase beta chain (EC 4.1.3.6) |
| AP53_908 | 1.222705622 | 5.781226351 | 35.44132977 | 8.21E-05 | 0.014447645 | Mg2+/citrate complex transporter |
| AP53_1751 | 2.242674357 | 6.490343622 | 35.4372308 | 8.21E-05 | 0.014447645 | Histidine ammonia-lyase (EC 4.3.1.3 |
| AP53_916 | 1.683172292 | 8.567138716 | 35.23918442 | 8.41E-05 | 0.014447645 | Apo-citrate lyase phosphoribosyl-d |
| AP53_1747 | 2.63406484 | 6.135558656 | 35.18796204 | 8.47E-05 | 0.014447645 | Formiminotetrahydrofolate cyclod |
| AP53_1748 | 2.478199958 | 6.709134603 | 34.83070309 | 8.86E-05 | 0.014447645 | Formate--tetrahydrofolate ligase (E |
| AP53_rna15 | 2.581074746 | 3.977905366 | 34.7651952 | 8.93E-05 | 0.014447645 | tRNA-Ala-TGC |
| AP53_1746 | 2.357355009 | 6.287957149 | 33.76044722 | 0.000101574 | 0.015165737 | Glutamate formiminotransferase (E |
| AP53_rna2 | 2.527912505 | 3.836238549 | 31.81249904 | 0.00013145 | 0.018164871 | tRNA-Ala-TGC |
| AP53_1082 | 1.349123726 | 5.178836739 | 29.85357857 | 0.000172487 | 0.018164871 | PTS system, cellobiose-specific IIC c |
| AP53_rna36 | 2.72559185 | 4.138285412 | 29.81930883 | 0.000173329 | 0.018164871 | tRNA-Ala-TGC |
| AP53_1758 | 1.767935056 | 9.430769949 | 29.35713283 | 0.000185175 | 0.018164871 | PTS system, trehalose-specific IIB co |
| AP53_697 | 0.751036314 | 7.56475385 | 29.34802494 | 0.000185417 | 0.018164871 | Cell surface protein |
| AP53_917 | 1.600879377 | 8.589404302 | 29.14712232 | 0.00019087 | 0.018164871 | Oxaloacetate decarboxylase alpha c |
| AP53_1084 | 1.71472175 | 5.287488837 | 28.75001194 | 0.000202214 | 0.018164871 | PTS system, cellobiose-specific IIA c |
| AP53_rna52 | 3.472056368 | 3.930072791 | 28.63230639 | 0.000205727 | 0.018164871 | tRNA-Ala-TGC |
| AP53_1744 | 2.040740013 | 5.749783198 | 28.62699506 | 0.000205887 | 0.018164871 | Imidazolonepropionase (EC 3.5.2.7 |
| AP53_1002 | 1.747149398 | 9.186568837 | 27.99233592 | 0.000226136 | 0.01836129 | Streptococcal phospholipase A2; _ |
| AP53_1732 | 0.711938302 | 8.474699799 | 27.79692891 | 0.000232836 | 0.01836129 | Dipeptidase (EC 3.4.-.-) |
| AP53_1645 | 2.090128056 | 6.523461692 | 27.54085828 | 0.000241973 | 0.01836129 | Transport protein SgaT, putative |
| AP53_rna13 | 2.249087349 | 3.975155994 | 27.43286538 | 0.000245952 | 0.01836129 | tRNA-Pro-TGG |
| AP53_rna79 | 2.63069565 | 3.888713338 | 26.99482848 | 0.000262904 | 0.018478361 | tRNA-Ala-TGC |
| AP53_rna60 | 1.672603001 | 4.476833431 | 26.90470174 | 0.000266561 | 0.018478361 | tRNA-Arg-CCG |
| AP53_1826 | -0.967332992 | 6.765204913 | 26.49663431 | 0.000283883 | 0.019000595 | FIG015389: hypothetical membran |
| AP53_1745 | 1.980903668 | 5.866330514 | 25.80994424 | 0.00031612 | 0.020241717 | Urocanate hydratase (EC 4.2.1.49) |
| AP53_164 | 1.682066749 | 4.340799239 | 25.58797411 | 0.000327449 | 0.020241717 | Mobile element protein |
| AP53_rna71 | 2.800443438 | 3.871305666 | 25.46911508 | 0.000333712 | 0.020241717 | tRNA-Ala-TGC |
| AP53_1039 | 0.934605469 | 5.831925291 | 24.0330863 | 0.000421718 | 0.023296182 | putative replication protein |
| AP53_698 | 1.000616896 | 8.420522513 | 23.88932275 | 0.000431951 | 0.023296182 | FIG01115236: hypothetical protei |

| | | | | | | |
|---|---|---|---|---|---|---|
| AP53_111 | -1.165473192 | 6.855954303 | 23.83574312 | 0.00043584 | 0.023296182 | Short chain fatty acids transporter |
| AP53_1865 | 2.157061623 | 11.85379858 | 23.70913888 | 0.000445193 | 0.023296182 | Streptolysin S biosynthesis protein |
| AP53_912 | 1.142603105 | 6.658422814 | 23.58826775 | 0.000454343 | 0.023296182 | FIG01114213: hypothetical protei |
| AP53_1086 | 1.13156748 | 4.62286242 | 23.56562824 | 0.000456082 | 0.023296182 | Transcriptional antiterminator of li |
| AP53_17 | 0.859944297 | 9.016055634 | 23.05227059 | 0.000497687 | 0.024291864 | Ribose-phosphate pyrophosphokin |
| AP53_484 | 0.613293782 | 6.791743853 | 22.95827401 | 0.000505781 | 0.024291864 | 2-haloalkanoic acid dehalogenase ( |
| AP53_109 | 1.426264754 | 6.207061618 | 22.8694196 | 0.000513575 | 0.024291864 | Fibronectin-binding protein |
| AP53_949 | 1.279571136 | 7.838170168 | 22.73500475 | 0.000525635 | 0.024291864 | Putative deoxyribose-specific ABC t |
| AP53_80 | 1.51874692 | 8.787198023 | 22.53927758 | 0.000543799 | 0.024546847 | DNA-directed RNA polymerase beta |
| AP53_905 | 0.873158893 | 5.511271964 | 22.23396569 | 0.000573631 | 0.024786532 | Methylmalonyl-CoA decarboxylase, |
| AP53_904 | 1.351014989 | 5.773598337 | 21.97553833 | 0.000600396 | 0.024786532 | Biotin carboxyl carrier protein of o |
| AP53_849 | 1.595203434 | 6.509506297 | 21.92305311 | 0.00060601 | 0.024786532 | Alcohol dehydrogenase, zinc-conta |
| AP53_820 | 1.623828773 | 9.906872654 | 21.84393046 | 0.000614591 | 0.024786532 | PTS system, mannose/fructose fami |
| AP53_rna6 | 1.868985507 | 4.56855931 | 21.71636768 | 0.000628728 | 0.024786532 | tRNA-Asp-GTC |
| AP53_rna30 | 1.640194372 | 3.832576393 | 21.70040263 | 0.000630524 | 0.024786532 | tRNA-Met-CAT |
| AP53_733 | 1.035893178 | 7.69791382 | 21.6301564 | 0.000638499 | 0.024786532 | Dihydrofolate reductase (EC 1.5.1.3 |
| AP53_942 | 1.309784658 | 9.17685393 | 21.1740101 | 0.000693276 | 0.026385285 | Protein:protein lipoyl transferase |
| AP53_rna48 | 2.114629358 | 5.123122914 | 21.06010203 | 0.000707807 | 0.026420274 | tRNA-His-GTG |
| AP53_1031 | 0.843099018 | 6.564176908 | 20.75603145 | 0.000748389 | 0.027104845 | FIG01114733: hypothetical protei |
| AP53_1045 | 0.854494798 | 6.293677744 | 20.71497302 | 0.000754076 | 0.027104845 | hypothetical phage protein |
| AP53_760 | 1.357052298 | 10.99692506 | 20.52300328 | 0.000781352 | 0.02757461 | SSU ribosomal protein S1p |
| AP53_1644 | 1.388037674 | 6.060227028 | 20.35479145 | 0.000806211 | 0.027943856 | Transaldolase (EC 2.2.1.2) |
| AP53_rna47 | 2.117452058 | 4.404252428 | 19.99541437 | 0.000862514 | 0.029370868 | tRNA-Trp-CCA |
| AP53_rna7 | 2.122379596 | 4.436492387 | 19.66583615 | 0.00091826 | 0.030730058 | tRNA-Lys-TTT |
| AP53_1042 | 0.861179377 | 6.407160573 | 19.43459839 | 0.000959914 | 0.031579554 | FIG01117510: hypothetical protei |
| AP53_rna49 | 2.165360222 | 5.221521578 | 19.32049031 | 0.000981288 | 0.031744655 | tRNA-Gln-TTG |
| AP53_16 | 1.046391028 | 9.407115863 | 19.17091249 | 0.00101016 | 0.032142961 | Secreted antigen GbpB/SagA/PcsB, |
| AP53_1125 | 0.727252637 | 7.154776504 | 18.83477087 | 0.001078778 | 0.033424585 | Uridine kinase (EC 2.7.1.48) [C1] |
| AP53_965 | 0.650291253 | 6.971739361 | 18.7454619 | 0.001097923 | 0.033424585 | Inositol-1-monophosphatase (EC 3. |
| AP53_948 | 1.091731604 | 8.363698522 | 18.57691515 | 0.001135156 | 0.033424585 | Putative deoxyribose-specific ABC t |
| AP53_957 | 0.969482011 | 8.083285336 | 18.53555333 | 0.001144519 | 0.033424585 | Lysyl aminopeptidase (EC 3.4.11.15 |
| AP53_489 | 1.420597345 | 7.91146 | 18.52396631 | 0.001147158 | 0.033424585 | Substrate-specific component CbrT |
| AP53_369 | 1.057821449 | 6.804805955 | 18.48835024 | 0.001155316 | 0.033424585 | Phosphate starvation-inducible pro |
| AP53_576 | 0.841377214 | 6.351748009 | 18.37544757 | 0.001181632 | 0.033424585 | FIG01115289: hypothetical protei |
| AP53_1001 | 1.317248476 | 7.743620698 | 18.23523858 | 0.001215302 | 0.033424585 | Phage protein |

| | | | | | | |
|---|---|---|---|---|---|---|
| AP53_1757 | 1.188145419 | 8.184491381 | 18.16431556 | 0.001232763 | 0.033424585 | Trehalose-6-phosphate hydrolase (E |
| AP53_821 | 1.594823025 | 9.889874738 | 18.08248283 | 0.00125328 | 0.033424585 | PTS system, mannose-specific IIC cc |
| AP53_rna44 | 1.979203063 | 3.465197914 | 17.94619974 | 0.001288348 | 0.033424585 | tRNA-Met-CAT |
| AP53_1052 | 0.738983859 | 7.631196824 | 17.66703015 | 0.001363862 | 0.033424585 | hypothetical protein |
| AP53_1303 | -0.920117591 | 8.524657579 | 17.63733729 | 0.001372197 | 0.033424585 | hypothetical protein |
| AP53_1083 | 1.528191407 | 4.977482168 | 17.58251797 | 0.001387743 | 0.033424585 | Putative uncharacterized protein S |
| AP53_913 | 0.822225127 | 7.533091996 | 17.52493724 | 0.001404296 | 0.033424585 | Citrate lyase gamma chain, acyl car |
| AP53_1742 | 1.342987286 | 10.55339526 | 17.48811131 | 0.001415005 | 0.033424585 | Alkyl hydroperoxide reductase prot |
| AP53_847 | 1.599644669 | 6.940372562 | 17.37178872 | 0.001449469 | 0.033424585 | Malate Na(+) symporter |
| AP53_1044 | 0.81962626 | 6.201115305 | 17.32404178 | 0.001463901 | 0.033424585 | hypothetical phage protein |
| AP53_1308 | 0.839869686 | 7.466925505 | 17.30022683 | 0.001471163 | 0.033424585 | Streptococcal extracellular nucleas |
| AP53_rna12 | 2.698441107 | 3.70589102 | 17.2924753 | 0.001473535 | 0.033424585 | tRNA-Arg-ACG |
| AP53_1311 | 0.873467749 | 8.270877675 | 17.174172 | 0.001510314 | 0.033424585 | Phage protein |
| AP53_1408 | 0.722969383 | 8.416879746 | 17.12595534 | 0.001525614 | 0.033424585 | ATP-dependent RNA helicase YqfR |
| AP53_171 | 0.84228986 | 7.61453829 | 16.9963088 | 0.001567668 | 0.033424585 | Metal-dependent hydrolase (EC 3.-. |
| AP53_952 | 1.418565789 | 7.544188929 | 16.96724118 | 0.001577283 | 0.033424585 | Ribosomal RNA small subunit meth |
| AP53_1088 | 0.776613884 | 4.336772951 | 16.92351143 | 0.00159188 | 0.033424585 | Beta-glucosidase (EC 3.2.1.21) |
| AP53_1094 | 1.59704497 | 4.265203367 | 16.88725125 | 0.001604104 | 0.033424585 | Mobile element protein |
| AP53_1848 | -0.406922989 | 8.29535345 | 16.81829886 | 0.001627657 | 0.033424585 | Hyaluronan synthase (EC 2.4.1.212 |
| AP53_rna24 | 2.118868955 | 3.822543586 | 16.7926616 | 0.001636518 | 0.033424585 | tRNA-Leu-TAA |
| AP53_rna74 | 1.499667396 | 3.94592579 | 16.79047552 | 0.001637276 | 0.033424585 | tRNA-Leu-AAG |
| AP53_614 | 1.804044602 | 5.801452265 | 16.73143035 | 0.001657912 | 0.033424585 | hypothetical phage protein |
| AP53_1091 | 0.733927006 | 8.27044064 | 16.7192181 | 0.001662219 | 0.033424585 | GTP-binding protein Obg |
| AP53_467 | -0.55150708 | 8.671118333 | 16.67091475 | 0.001679381 | 0.033424585 | ABC transporter, substrate-binding |
| AP53_rna23 | 2.306278026 | 2.598266444 | 16.61772821 | 0.001698522 | 0.033424585 | tRNA-Gly-GCC |
| AP53_1585 | 1.051858392 | 8.253436029 | 16.5989606 | 0.001705337 | 0.033424585 | Glutamine synthetase type I (EC 6.3 |
| AP53_684 | -0.553487108 | 7.260847375 | 16.58946989 | 0.001708795 | 0.033424585 | Lipoprotein signal peptidase (EC 3.4 |
| AP53_588 | 0.541875174 | 8.92811689 | 16.54610906 | 0.001724703 | 0.033424585 | dTDP-4-dehydrorhamnose reducta: |
| AP53_941 | 1.051247662 | 8.638612252 | 16.39892822 | 0.001780016 | 0.033424585 | Probable NADH-dependent flavin o |
| AP53_112 | -0.836693311 | 5.848035247 | 16.39355308 | 0.001782076 | 0.033424585 | Transcriptional regulators, LysR fan |
| AP53_1752 | 0.697765692 | 6.705251976 | 16.3736834 | 0.001789713 | 0.033424585 | Formiminoglutamase (EC 3.5.3.8) |
| AP53_rna81 | 1.641856464 | 8.333760293 | 16.32895564 | 0.001807046 | 0.033424585 | tRNA-Lys-CTT |
| AP53_734 | 1.307949967 | 8.466451951 | 16.30290022 | 0.001817234 | 0.033424585 | FIG01114189: hypothetical protei |
| AP53_902 | 0.732533201 | 4.822374628 | 16.18738473 | 0.001863228 | 0.033424585 | Oxaloacetate decarboxylase alpha c |
| AP53_1594 | 0.872818613 | 11.13498404 | 16.17412721 | 0.001868594 | 0.033424585 | Fructose-bisphosphate aldolase cla |

| | | | | | | |
|---|---|---|---|---|---|---|
| AP53_830 | 0.697809715 | 7.795508083 | 16.11992101 | 0.001890724 | 0.033424585 | GTPase and tRNA-U34 5-formylatio |
| AP53_1729 | 0.413120951 | 8.363552599 | 16.08685557 | 0.001904375 | 0.033424585 | FIG01115477: hypothetical protei |
| AP53_1743 | 1.346996039 | 10.51698796 | 16.07136855 | 0.001910809 | 0.033424585 | Alkyl hydroperoxide reductase prot |
| AP53_1489 | 0.980278314 | 10.06214315 | 16.05530196 | 0.001917511 | 0.033424585 | Trans-2,cis-3-Decenoyl-ACP isomera |
| AP53_1128 | 1.289709096 | 10.47072346 | 15.98530304 | 0.001947034 | 0.033424585 | Non-phosphorylating glyceraldehy |
| AP53_947 | 1.041836674 | 8.593368411 | 15.97527603 | 0.001951307 | 0.033424585 | Nucleoside ABC transporter, perme |
| AP53_257 | -0.76101857 | 8.245360951 | 15.96645643 | 0.001955074 | 0.033424585 | Methionine ABC transporter ATP-bi |
| AP53_165 | 1.239867051 | 2.954387678 | 15.89630038 | 0.001985351 | 0.033424585 | Mobile element protein |
| AP53_762 | 1.107058187 | 9.11980704 | 15.84864235 | 0.002006233 | 0.033424585 | Rhodanese domain protein UPF017 |
| AP53_725 | 0.696321544 | 7.957002543 | 15.84794829 | 0.002006539 | 0.033424585 | Regulation of D-alanyl-lipoteichoic |
| AP53_1241 | 0.987644996 | 9.750516897 | 15.83490905 | 0.002012299 | 0.033424585 | Hypothetical protein YggS, proline |
| AP53_742 | 0.973111338 | 10.24984754 | 15.80001242 | 0.00202781 | 0.033424585 | Purine nucleoside phosphorylase (E |
| AP53_1053 | 1.044453591 | 7.111292246 | 15.79765232 | 0.002028864 | 0.033424585 | conserved hypothetical protein - pl |
| AP53_1599 | 0.82641789 | 9.99443516 | 15.79065097 | 0.002031994 | 0.033424585 | Cell division trigger factor (EC 5.2.1 |
| AP53_1242 | 0.999998776 | 9.804031108 | 15.64445516 | 0.002098689 | 0.033679589 | Cell division protein FtsZ (EC 3.4.24 |
| AP53_1719 | -0.675849188 | 6.55302028 | 15.63478782 | 0.002103189 | 0.033679589 | PTS system, cellobiose-specific IIC c |
| AP53_309 | 0.722027905 | 8.569034772 | 15.61288669 | 0.002113428 | 0.033679589 | Uracil phosphoribosyltransferase (E |
| AP53_550 | 0.833112509 | 7.078859446 | 15.56510934 | 0.002135967 | 0.033679589 | Streptolysin S biosynthesis protein |
| AP53_1248 | 0.913308431 | 9.88017853 | 15.55314632 | 0.002141655 | 0.033679589 | GTP-binding protein TypA/BipA |
| AP53_1721 | -0.892177595 | 6.061442966 | 15.53230426 | 0.002151607 | 0.033679589 | PTS system, cellobiose-specific IIA c |
| AP53_1550 | -0.543738968 | 7.688059374 | 15.44712849 | 0.00219285 | 0.03403125 | Phosphatidylglycerophosphatase B |
| AP53_1808 | -0.790452882 | 7.402905307 | 15.37808381 | 0.002226969 | 0.03403125 | bacteriocin immunity protein |
| AP53_146 | 0.743032738 | 6.035456131 | 15.3447735 | 0.002243653 | 0.03403125 | L-ribulose-5-phosphate 4-epimeras |
| AP53_115 | -0.842509661 | 5.049902357 | 15.32545537 | 0.002253396 | 0.03403125 | Butyrate-acetoacetate CoA-transfer |
| AP53_rna75 | 7.254038285 | -0.117416991 | 17.10911612 | 0.002301853 | 0.03403125 | tRNA-Arg-ACG |
| AP53_368 | 0.891030825 | 9.824903804 | 15.1841073 | 0.002326226 | 0.03403125 | 67kDa Myosin-crossreactive strept |
| AP53_551 | 0.901847986 | 7.020067052 | 15.18303758 | 0.002326788 | 0.03403125 | Export ABC transporter ATP-binding |
| AP53_rna11 | 1.720065373 | 4.042386343 | 15.15109139 | 0.002343636 | 0.03403125 | tRNA-Leu-TAA |
| AP53_1243 | 0.928333836 | 9.14538567 | 15.13944476 | 0.002349814 | 0.03403125 | Cell division protein FtsA |
| AP53_853 | 0.696696831 | 6.647686072 | 15.05148754 | 0.002397097 | 0.03403125 | Lipase/Acylhydrolase family protei |
| AP53_1713 | 1.607327868 | 10.57102058 | 15.04679199 | 0.002399653 | 0.03403125 | Streptodornase B; Mitogenic factor |
| AP53_703 | 0.893976656 | 6.57625008 | 15.04276329 | 0.002401848 | 0.03403125 | Thioredoxin reductase (EC 1.8.1.9) |
| AP53_1087 | 0.788592469 | 4.395822584 | 15.03539273 | 0.00240587 | 0.03403125 | Outer surface protein of unknown f |
| AP53_105 | -0.529474966 | 7.031738335 | 15.01046574 | 0.002419532 | 0.03403125 | Fibronectin-binding protein/major |
| AP53_1586 | 0.945814536 | 7.962072421 | 14.92524449 | 0.002466939 | 0.034282862 | Transcriptional regulator, MerR fan |

| | | | | | | |
|---|---|---|---|---|---|---|
| AP53_1019 | 1.636868472 | 7.874398624 | 14.91493681 | 0.002472746 | 0.034282862 | Phage protein |
| AP53_558 | 0.656380508 | 7.727189147 | 14.81319177 | 0.002530949 | 0.034778781 | Transcription regulator [contains d |
| AP53_889 | -0.46043907 | 8.84098912 | 14.79015264 | 0.002544352 | 0.034778781 | Sortase A, LPXTG specific |
| AP53_1051 | 1.321601857 | 8.48216266 | 14.75203555 | 0.002566711 | 0.03483906 | hypothetical protein |
| AP53_560 | 1.092723966 | 9.652494913 | 14.65647724 | 0.002623791 | 0.035325693 | ATP synthase A chain (EC 3.6.3.14) |
| AP53_436 | 1.147249512 | 5.781994632 | 14.61073574 | 0.002651644 | 0.035325693 | Ribonuclease III (EC 3.1.26.3) |
| AP53_1514 | -0.458489728 | 7.701027762 | 14.59256082 | 0.002662807 | 0.035325693 | Transmembrane component BL069 |
| AP53_1815 | 1.050353948 | 9.372363644 | 14.56830653 | 0.002677792 | 0.035325693 | LSU ribosomal protein L33p @ LSU |
| AP53_815 | 0.60653289 | 8.798029072 | 14.48987195 | 0.002726935 | 0.035325693 | Translation elongation factor LepA |
| AP53_1636 | 0.845837255 | 8.608762784 | 14.48856299 | 0.002727764 | 0.035325693 | 23S rRNA (guanosine-2'-O-) -methyl |
| AP53_623 | 1.063541524 | 3.912249943 | 14.4539583 | 0.002749792 | 0.035325693 | Phage protein |
| AP53_865 | 0.534677636 | 7.424202608 | 14.43288468 | 0.002763309 | 0.035325693 | Similar to ribosomal large subunit p |
| AP53_rna43 | 1.829035706 | 4.568139016 | 14.40029537 | 0.002784367 | 0.035325693 | tRNA-Pseudo-TGA |
| AP53_736 | 0.854413837 | 8.860886175 | 14.3999983 | 0.00278456 | 0.035325693 | GTP-binding protein EngB |
| AP53_590 | 0.451525519 | 7.752761326 | 14.32949204 | 0.002830775 | 0.035678355 | Alpha-L-Rha alpha-1,3-L-rhamnosyl |
| AP53_1304 | -0.785142366 | 9.344842128 | 14.27574534 | 0.002866614 | 0.035678355 | Alpha-mannosidase (EC 3.2.1.24) |
| AP53_1487 | 0.908078229 | 10.25795846 | 14.26826319 | 0.002871645 | 0.035678355 | 3-oxoacyl-[acyl-carrier-protein] syn |
| AP53_1659 | 1.153197424 | 9.560678367 | 14.21685412 | 0.002906499 | 0.035678355 | Preprotein translocase subunit Yaj( |
| AP53_735 | 0.899837641 | 8.638929863 | 14.1902202 | 0.002924753 | 0.035678355 | ATP-dependent Clp protease ATP-bi |
| AP53_216 | -0.403915541 | 8.772890398 | 14.1770108 | 0.002933856 | 0.035678355 | 3'->5' exoribonuclease Bsu YhaM |
| AP53_946 | 0.620914089 | 8.691793309 | 14.10691595 | 0.002982725 | 0.035678355 | Phosphoglucomutase (EC 5.4.2.2) |
| AP53_1324 | 1.205670115 | 3.836125107 | 14.10320385 | 0.00298534 | 0.035678355 | Phage minor capsid protein |
| AP53_783 | 0.567249883 | 7.122457783 | 14.08318326 | 0.002999488 | 0.035678355 | Hypothetical protein SPy1011 |
| AP53_1647 | 1.004929474 | 5.062154392 | 14.07688121 | 0.003003959 | 0.035678355 | Transcription antiterminator, BglG |
| AP53_577 | 1.15461435 | 8.271785237 | 14.06198691 | 0.003014554 | 0.035678355 | ABC transporter permease protein |
| AP53_rna63 | 1.677059743 | 2.769722909 | 14.01043077 | 0.003051574 | 0.035897609 | tRNA-Ile-GAT |
| AP53_1702 | -0.623670694 | 7.891787285 | 13.93419668 | 0.003107302 | 0.035978315 | ABC transporter ATP-binding protei |
| AP53_38 | -0.518568795 | 8.772914564 | 13.93244902 | 0.003108594 | 0.035978315 | Acyltransferase family |
| AP53_81 | 1.113479164 | 9.263004421 | 13.92508554 | 0.003114043 | 0.035978315 | DNA-directed RNA polymerase beta |
| AP53_103 | -0.437263039 | 6.880115778 | 13.83925976 | 0.003178391 | 0.036504484 | putative collagen binding protein |
| AP53_822 | 1.410649243 | 9.76644687 | 13.75559963 | 0.003242633 | 0.037005635 | PTS system, mannose-specific IID co |
| AP53_1038 | 0.707114421 | 5.76057865 | 13.72282744 | 0.003268215 | 0.037005635 | hypothetical protein |
| AP53_rna50 | 2.096024096 | 3.729627845 | 13.70882117 | 0.003279222 | 0.037005635 | tRNA-Leu-CAA |
| AP53_559 | 0.914043531 | 9.645722823 | 13.65677947 | 0.003320502 | 0.037096293 | ATP synthase C chain (EC 3.6.3.14) |
| AP53_861 | 0.549000536 | 8.722673962 | 13.65055582 | 0.003325479 | 0.037096293 | Ribose-phosphate pyrophosphokin |

| | | | | | | |
|---|---|---|---|---|---|---|
| AP53_804 | 0.779707352 | 8.79923924 | 13.61399673 | 0.003354897 | 0.037210603 | Phosphoglucosamine mutase (EC 5. |
| AP53_433 | 0.487473372 | 8.225934195 | 13.56797663 | 0.003392367 | 0.037412406 | Two-component response regulato |
| AP53_615 | 1.521743195 | 5.69586088 | 13.48005562 | 0.003465334 | 0.03800121 | hypothetical phage protein |
| AP53_1700 | -0.627583938 | 8.287535281 | 13.348264 | 0.003578214 | 0.039018619 | Two-component system response r |
| AP53_1240 | 0.73678771 | 10.00904625 | 13.27970118 | 0.003638654 | 0.039387053 | FIG001960: FtsZ-interacting protei |
| AP53_886 | 0.671891991 | 11.36905956 | 13.22967419 | 0.003683516 | 0.039387053 | NADH peroxidase (EC 1.11.1.1) |
| AP53_490 | 0.918229089 | 8.144957934 | 13.22094944 | 0.003691406 | 0.039387053 | Additional lipoprotein component |
| AP53_348 | 0.641782119 | 8.534545207 | 13.18508779 | 0.00372405 | 0.039387053 | N-acetylglucosamine-1-phosphate |
| AP53_75 | 0.513227421 | 7.726886869 | 13.15808501 | 0.003748855 | 0.039387053 | Zinc ABC transporter, ATP-binding |
| AP53_286 | 0.469905178 | 8.349778934 | 13.15443726 | 0.003752221 | 0.039387053 | unknown domain / Nucleoside 5-tr |
| AP53_1203 | -0.776848533 | 3.052125253 | 13.09903123 | 0.003803786 | 0.039387053 | Phage protein |
| AP53_1553 | -0.62260861 | 7.983589658 | 13.08448904 | 0.003817458 | 0.039387053 | FIG01114225: hypothetical protei |
| AP53_rna45 | 1.95227481 | 3.076635878 | 13.03078886 | 0.003868452 | 0.039387053 | tRNA-Phe-GAA |
| AP53_835 | 1.022273699 | 10.22822728 | 13.02012921 | 0.00387867 | 0.039387053 | Dihydrofolate synthase (EC 6.3.2.1 |
| AP53_1634 | 0.988126908 | 8.437611102 | 13.01886325 | 0.003879886 | 0.039387053 | DegV family protein in cluster with |
| AP53_882 | -0.646509174 | 8.88071783 | 12.98990796 | 0.003907813 | 0.039387053 | Pseudouridylate synthases, 23S RN |
| AP53_831 | 1.283369374 | 9.968418741 | 12.98798716 | 0.003909674 | 0.039387053 | LSU ribosomal protein L10p (P0) |
| AP53_307 | 0.642634395 | 7.624681867 | 12.98295291 | 0.003914556 | 0.039387053 | UDP-N-acetylmuramoylalanyl-D-gl |
| AP53_564 | 1.010614346 | 10.08214382 | 12.95645834 | 0.003940371 | 0.039387053 | ATP synthase gamma chain (EC 3.6. |
| AP53_1197 | -1.025494215 | 3.318117855 | 12.94360544 | 0.003952967 | 0.039387053 | FIG048485: Phage protein |
| AP53_1175 | 0.372567418 | 8.466828996 | 12.89977172 | 0.003996282 | 0.039387053 | Dihydroorotate dehydrogenase, ca |
| AP53_114 | -0.755463606 | 5.371535414 | 12.8670978 | 0.004028934 | 0.039387053 | Acetyl-CoA:acetoacetyl-CoA transfe |
| AP53_1578 | -0.671342723 | 8.095803619 | 12.81112251 | 0.004085609 | 0.039387053 | Predicted regulator for deoxynucle |
| AP53_300 | 1.20891156 | 8.145165313 | 12.7980653 | 0.004098964 | 0.039387053 | Pyruvate formate-lyase activating e |
| AP53_rna8 | 1.563556892 | 4.747622072 | 12.78773022 | 0.004109572 | 0.039387053 | tRNA-Leu-TAG |
| AP53_rna9 | 1.530928592 | 4.351643606 | 12.7823354 | 0.004115122 | 0.039387053 | tRNA-Thr-TGT |
| AP53_566 | 0.949412874 | 10.8465092 | 12.7820319 | 0.004115434 | 0.039387053 | ATP synthase epsilon chain (EC 3.6. |
| AP53_64 | 1.023687657 | 7.801527973 | 12.75085571 | 0.004147684 | 0.039387053 | Adenylate kinase (EC 2.7.4.3) |
| AP53_753 | -0.536874714 | 8.807008081 | 12.74692421 | 0.004151773 | 0.039387053 | Uracil-DNA glycosylase, family 1 |
| AP53_301 | 0.71643194 | 9.702029137 | 12.7391615 | 0.004159858 | 0.039387053 | Manganese-dependent inorganic p |
| AP53_rna83 | 2.434411983 | 3.752216362 | 12.73913151 | 0.00415989 | 0.039387053 | tRNA-Asn-GTT |
| AP53_1522 | -0.608232301 | 8.075835223 | 12.71569431 | 0.004184416 | 0.03941425 | Streptococcal cell surface hemopr |
| AP53_710 | 0.584153517 | 7.45928674 | 12.69768872 | 0.004203374 | 0.03941425 | Peptidoglycan hydrolase, Autolysi |
| AP53_866 | 0.976189 | 10.61348664 | 12.60046192 | 0.004307518 | 0.039906806 | Phosphate acetyltransferase (EC 2.3 |
| AP53_555 | 0.55973509 | 8.73793431 | 12.58022449 | 0.004329576 | 0.039906806 | Endonuclease/Exonuclease/phosph |

| | | | | | | |
|---|---|---|---|---|---|---|
| AP53_267 | 0.858533551 | 10.33651236 | 12.57161975 | 0.004338996 | 0.039906806 | Response regulator CsrR |
| AP53_351 | 0.645007568 | 9.231868952 | 12.55891726 | 0.004352946 | 0.039906806 | 5'-methylthioadenosine nucleosida |
| AP53_372 | 0.479139497 | 8.032045335 | 12.55368865 | 0.004358703 | 0.039906806 | Diacylglycerol kinase (EC 2.7.1.107 |
| AP53_69 | 1.008839071 | 10.49884439 | 12.49463523 | 0.004424358 | 0.040317738 | DNA-directed RNA polymerase alph |
| AP53_799 | 0.605911891 | 7.171421152 | 12.42032883 | 0.004508638 | 0.040893771 | Lipoate-protein ligase A |
| AP53_1218 | 0.386871885 | 8.972107627 | 12.37126138 | 0.004565333 | 0.041113826 | FIG006988: Lipase/Acylhydrolase w |
| AP53_273 | -0.404849462 | 7.785493392 | 12.36274105 | 0.004575263 | 0.041113826 | SWF/SNF family helicase |
| AP53_1130 | 0.895065872 | 12.05479986 | 12.34013216 | 0.004601738 | 0.041121993 | Phosphocarrier protein of PTS syste |
| AP53_176 | -0.905322766 | 5.921850499 | 12.29635021 | 0.004653523 | 0.041121993 | hypothetical cytosolic protein |
| AP53_113 | -0.912903946 | 5.138902606 | 12.29133627 | 0.004659497 | 0.041121993 | 3-ketoacyl-CoA thiolase (EC 2.3.1.1 |
| AP53_1105 | 0.888722953 | 7.325119416 | 12.27697795 | 0.004676656 | 0.041121993 | Phenylacetic acid degradation prot |
| AP53_1460 | 0.501114367 | 9.238176012 | 12.26865315 | 0.004686638 | 0.041121993 | tRNA (guanine46-N7-)-methyltrans |
| AP53_1015 | 1.093634646 | 6.161086585 | 12.23825319 | 0.004723307 | 0.041121993 | Phage protein |
| AP53_926 | 0.519280827 | 8.964821659 | 12.23412145 | 0.004728316 | 0.041121993 | GMP synthase [glutamine-hydrolyzi |
| AP53_681 | 0.667928894 | 9.560920647 | 12.188635 | 0.004783887 | 0.041121993 | FIG139598: Potential ribosomal pr |
| AP53_832 | 0.91984238 | 9.839306377 | 12.18514007 | 0.004788189 | 0.041121993 | LSU ribosomal protein L7/L12 (P1/ |
| AP53_909 | 0.604784885 | 5.601753535 | 12.16997515 | 0.004806907 | 0.041121993 | FIG01114846: hypothetical protei |
| AP53_rna27 | 1.251316852 | 4.911695727 | 12.16810761 | 0.004809218 | 0.041121993 | tRNA-Met-CAT |
| AP53_732 | 0.634784149 | 6.437125728 | 12.09694059 | 0.004898275 | 0.04169979 | Thymidylate synthase (EC 2.1.1.45) |
| AP53_719 | 0.357757599 | 8.308501386 | 12.04640133 | 0.004962704 | 0.042063793 | COG0488: ATPase components of A |
| AP53_rna25 | 2.501161498 | 3.774093468 | 11.97740666 | 0.005052286 | 0.042202251 | tRNA-Arg-ACG |
| AP53_rna46 | 2.098978021 | 3.464757328 | 11.9710494 | 0.005060636 | 0.042202251 | tRNA-Tyr-GTA |
| AP53_1488 | 0.898859445 | 10.09266506 | 11.96982231 | 0.005062249 | 0.042202251 | Transcriptional regulator of fatty ac |
| AP53_1812 | 0.560442285 | 8.060183411 | 11.9669646 | 0.00506601 | 0.042202251 | Aspartyl-tRNA synthetase (EC 6.1.1. |
| AP53_1474 | 0.707465082 | 6.336304883 | 11.93255419 | 0.005111547 | 0.042399631 | Putative regulator of the mannose |
| AP53_1559 | 1.01266939 | 11.45855556 | 11.91283187 | 0.005137866 | 0.042436584 | Pyruvate formate-lyase (EC 2.3.1.54 |
| AP53_1850 | -0.769996928 | 8.012850345 | 11.89645944 | 0.005159836 | 0.042437461 | FIG01117167: hypothetical protei |
| AP53_848 | 1.401282975 | 7.052381792 | 11.87786262 | 0.005184925 | 0.042457101 | NAD-dependent malic enzyme (EC 1 |
| AP53_1029 | 1.254994276 | 5.505463083 | 11.86234481 | 0.005205971 | 0.042457101 | Phage terminase, small subunit |
| AP53_692 | -0.878645874 | 8.311700924 | 11.83014775 | 0.005249962 | 0.042636717 | ABC transporter ATP-binding protei |
| AP53_963 | 0.499256779 | 7.67750797 | 11.80333679 | 0.005286929 | 0.042731997 | Phosphate ABC transporter, peripla |
| AP53_578 | 0.846844624 | 8.24742685 | 11.78978983 | 0.005305725 | 0.042731997 | ABC transporter ATP-binding protei |
| AP53_1845 | -0.365771619 | 8.383367852 | 11.71143404 | 0.005416003 | 0.043439924 | Transcriptional regulator in cluster |
| AP53_1579 | 0.574225267 | 8.787446596 | 11.69095859 | 0.005445265 | 0.043494898 | SSU ribosomal protein S14p (S29e) |
| AP53_826 | 1.229941131 | 8.443236576 | 11.66043482 | 0.005489237 | 0.043666427 | Succinate-semialdehyde dehydroge |

| | | | | | | |
|---|---|---|---|---|---|---|
| AP53_716 | 1.157243019 | 9.254855066 | 11.61579768 | 0.005554297 | 0.044003638 | Hypothetical protein SP1558 |
| AP53_rna26 | 2.208598411 | 4.466309467 | 11.59907063 | 0.005578913 | 0.044018981 | tRNA-Pro-TGG |
| AP53_383 | 0.941480644 | 6.861804942 | 11.5769015 | 0.005611735 | 0.044098697 | putative role in replication |
| AP53_68 | 1.015167993 | 10.91520246 | 11.52534964 | 0.005688948 | 0.044525195 | SSU ribosomal protein S11p (S14e) |
| AP53_1526 | -0.618580059 | 8.884422822 | 11.49647707 | 0.005732742 | 0.044526071 | Holo-[acyl-carrier protein] synthase |
| AP53_1127 | -0.465221492 | 9.163542651 | 11.48072661 | 0.0057568 | 0.044526071 | Peptidoglycan N-acetylglucosamin |
| AP53_1651 | 0.684043456 | 8.844024322 | 11.48002161 | 0.005757879 | 0.044526071 | Peptide deformylase (EC 3.5.1.88) |
| AP53_562 | 1.070948624 | 9.093359734 | 11.41182612 | 0.005863456 | 0.045127668 | ATP synthase delta chain (EC 3.6.3. |
| AP53_747 | 0.951711814 | 9.584863311 | 11.39149906 | 0.005895368 | 0.045127668 | FIG01114562: hypothetical protei |
| AP53_1764 | -0.883023155 | 9.072809522 | 11.38512043 | 0.005905424 | 0.045127668 | UPF0246 protein YaaA |
| AP53_275 | -0.481064028 | 9.150603593 | 11.27723773 | 0.006078618 | 0.046269011 | UDP-N-acetylmuramate--alanine lig |
| AP53_1419 | 0.519741593 | 9.811227587 | 11.21861607 | 0.006175247 | 0.046646034 | Transketolase (EC 2.2.1.1) |
| AP53_1720 | -0.564487891 | 6.068932166 | 11.20477407 | 0.006198329 | 0.046646034 | PTS system, cellobiose-specific IIB c |
| AP53_387 | 1.537363617 | 1.849198738 | 11.19424749 | 0.00621595 | 0.046646034 | hypothetical protein |
| AP53_1637 | 0.489008167 | 6.216054756 | 11.18928479 | 0.006224278 | 0.046646034 | Hypothetical protein, Spy1939 hor |
| AP53_259 | -0.777870015 | 7.002333297 | 11.15436622 | 0.006283249 | 0.046748067 | Branched-chain amino acid transpo |
| AP53_97 | 0.46978645 | 7.423308075 | 11.15271085 | 0.006286062 | 0.046748067 | Phenylalanyl-tRNA synthetase dom |
| AP53_312 | -0.780787641 | 9.453151372 | 11.11570553 | 0.006349314 | 0.047038239 | hypothetical protein |
| AP53_1423 | 0.725013647 | 8.381643768 | 11.07989381 | 0.006411245 | 0.047230408 | Glycerol uptake facilitator protein |
| AP53_592 | 0.615939139 | 7.379795712 | 11.04345717 | 0.006474992 | 0.047230408 | ABC-transporter (ATP-binding prote |
| AP53_1663 | 0.712608285 | 6.881238382 | 11.03868183 | 0.006483402 | 0.047230408 | Multiple sugar ABC transporter, AT |
| AP53_784 | 0.557657713 | 7.389123837 | 11.02911574 | 0.006500288 | 0.047230408 | TPR-repeat-containing protein |
| AP53_730 | 0.542044484 | 7.593321494 | 11.02494059 | 0.006507674 | 0.047230408 | Hydroxymethylglutaryl-CoA reduct |
| AP53_1338 | 0.823830661 | 3.122952067 | 11.01728047 | 0.006521252 | 0.047230408 | FIG01114733: hypothetical protei |
| AP53_311 | -0.72857498 | 12.01622451 | 10.97391734 | 0.006598748 | 0.047456799 | ATP-dependent Clp protease protec |
| AP53_1510 | -0.651971561 | 6.923368282 | 10.97209613 | 0.006602026 | 0.047456799 | ABC transporter permease protein |
| AP53_910 | 0.836791089 | 7.052291359 | 10.95737135 | 0.006628605 | 0.047456799 | Biotin carboxyl carrier protein of o |
| AP53_rna67 | 1.343134034 | 4.349876004 | 10.94209967 | 0.006656304 | 0.047456799 | tRNA-Asp-GTC |
| AP53_rna58 | 1.522721353 | 3.880953751 | 10.92289652 | 0.006691329 | 0.047456799 | tRNA-Tyr-GTA |
| AP53_1660 | 1.136917409 | 8.781260688 | 10.91131513 | 0.006712558 | 0.047456799 | putative transport accessory protei |
| AP53_43 | 1.018354106 | 10.88048974 | 10.89588231 | 0.006740971 | 0.047456799 | LSU ribosomal protein L3p (L3e) |
| AP53_1643 | 0.743001203 | 8.456063552 | 10.89201807 | 0.006748107 | 0.047456799 | Polyribonucleotide nucleotidyltra |
| AP53_1756 | 0.726536534 | 9.032149695 | 10.82116164 | 0.006880566 | 0.048135285 | Neutral endopeptidase O (EC 3.4.2 |
| AP53_1122 | 0.71200303 | 8.656673592 | 10.81396928 | 0.006894183 | 0.048135285 | GAF domain-containing proteins |
| AP53_1468 | -0.984923808 | 8.166070866 | 10.78266197 | 0.006953828 | 0.048377704 | ATPase YjeE, predicted to have esse |

| | | | | | | |
|---|---|---|---|---|---|---|
| AP53_1014 | 1.315726708 | 6.659339264 | 10.76205254 | 0.006993425 | 0.048479419 | Phage tail length tape-measure pro |
| AP53_1112 | 0.610042007 | 9.61875054 | 10.7079942 | 0.007098563 | 0.049033134 | Methionine aminopeptidase (EC 3.4 |
| AP53_1026 | 1.192268763 | 5.864797043 | 10.69001582 | 0.007133943 | 0.049102778 | Phage protein |
| AP53_431 | -0.710307167 | 6.495646796 | 10.66848878 | 0.007176582 | 0.049150597 | Long-chain-fatty-acid--CoA ligase (E |
| AP53_939 | 0.797720837 | 9.212780987 | 10.64108317 | 0.007231302 | 0.049150597 | Biotinyl-lipoyl attachment domain |
| AP53_1016 | 1.455355128 | 6.137775317 | 10.64032505 | 0.007232822 | 0.049150597 | Phage protein |
| AP53_1024 | 1.563860192 | 7.609041708 | 10.63566436 | 0.00724218 | 0.049150597 | Prophage Clp protease-like protein |
| AP53_935 | 0.69008319 | 9.235748517 | 10.58636729 | 0.007342036 | 0.049654676 | Formate--tetrahydrofolate ligase (E |
| AP53_1342 | -0.915464754 | 4.625304636 | 10.55849456 | 0.007399217 | 0.049665347 | Phage protein |
| AP53_702 | 0.679078875 | 6.892190454 | 10.55126549 | 0.007414133 | 0.049665347 | tRNA (Guanine37-N1) -methyltrans |
| AP53_829 | 0.717601964 | 10.09032713 | 10.54689387 | 0.007423171 | 0.049665347 | Acetylornithine deacetylase/Succir |
| AP53_345 | 0.832472169 | 8.13500415 | 10.52748527 | 0.007463453 | 0.049665347 | 3-oxoacyl-[acyl-carrier protein] red |
| AP53_rna84 | 1.83888244 | 1.78152497 | 10.52359809 | 0.007471551 | 0.049665347 | tRNA-Glu-TTC |
| AP53_589 | 0.489956123 | 6.987027995 | 10.46257997 | 0.007600055 | 0.050077363 | Alpha-D-GlcNAc alpha-1,2-L-rhamn |
| AP53_1471 | 0.625155797 | 8.431843594 | 10.4510463 | 0.007624638 | 0.050077363 | PTS system, mannose-specific IIB cc |
| AP53_840 | 0.648334596 | 8.092291269 | 10.43945606 | 0.007649437 | 0.050077363 | UDP-N-acetylenolpyruvoylglucosar |
| AP53_371 | 0.482509681 | 8.04646574 | 10.43367623 | 0.007661839 | 0.050077363 | Metal-dependent hydrolase YbeY, ir |
| AP53_1047 | 0.839972932 | 6.075203301 | 10.43335317 | 0.007662533 | 0.050077363 | hypothetical phage protein |
| AP53_284 | 0.707984507 | 9.147368134 | 10.39994578 | 0.007734694 | 0.050314953 | FIG027054: hypothetical protein |
| AP53_1309 | 1.100746891 | 6.652699164 | 10.39257117 | 0.007750732 | 0.050314953 | FIG01117556: hypothetical protei |
| AP53_rna19 | 1.884274832 | 4.531004466 | 10.37802856 | 0.007782474 | 0.050352605 | tRNA-Asp-GTC |
| AP53_563 | 0.93409076 | 10.59655174 | 10.35592445 | 0.007831015 | 0.050428044 | ATP synthase alpha chain (EC 3.6.3. |
| AP53_649 | -0.853522511 | 6.231483278 | 10.34909081 | 0.007846094 | 0.050428044 | hypothetical phage protein |
| AP53_473 | 0.650970464 | 9.506031141 | 10.31548582 | 0.007920751 | 0.050739861 | HPr kinase/phosphorylase (EC 2.7.1 |
| AP53_1129 | 0.819463319 | 10.5572049 | 10.28015064 | 0.008000161 | 0.050839803 | Phosphoenolpyruvate-protein pho |
| AP53_1810 | -0.540950517 | 7.450918334 | 10.27439306 | 0.008013189 | 0.050839803 | membrane protein, putative |
| AP53_1722 | 0.494272198 | 9.069333087 | 10.25528055 | 0.008056618 | 0.050839803 | Deoxyribonucleoside regulator Dec |
| AP53_156 | 1.080643583 | 10.26429078 | 10.22773576 | 0.008119699 | 0.050839803 | Co-activator of prophage gene expr |
| AP53_rna66 | 1.43247101 | 4.587209041 | 10.22552079 | 0.008124797 | 0.050839803 | tRNA-Lys-TTT |
| AP53_738 | -0.88654946 | 13.6746115 | 10.21382199 | 0.008151785 | 0.050839803 | putative ATP-dependent Clp protei |
| AP53_1025 | 1.596908294 | 6.219577531 | 10.20623051 | 0.008169355 | 0.050839803 | Phage portal protein |
| AP53_932 | -0.688461526 | 5.974249375 | 10.18210801 | 0.008225484 | 0.050839803 | Predicted transcriptional regulator |
| AP53_1237 | 0.453314677 | 9.540523851 | 10.17362315 | 0.008245335 | 0.050839803 | Cell division initiation protein DivI |
| AP53_701 | 0.688771736 | 7.052130267 | 10.17031159 | 0.008253098 | 0.050839803 | 16S rRNA processing protein RimM |
| AP53_863 | 0.720379896 | 7.861865308 | 10.15337992 | 0.008292926 | 0.050839803 | GTP pyrophosphokinase (EC 2.7.6.5 |

| | | | | | | |
|---|---|---|---|---|---|---|
| AP53_1626 | 1.05423917 | 7.678876515 | 10.13979161 | 0.008325054 | 0.050839803 | Toxin to DNA-damage-inducible pro |
| AP53_1676 | 0.865738436 | 7.594097753 | 10.12487164 | 0.008360501 | 0.050839803 | 6-aminohexanoate-cyclic-dimer hy |
| AP53_370 | 0.562511778 | 7.3926236 | 10.11488467 | 0.008384328 | 0.050839803 | Hypothetical protein VC0266 (suga |
| AP53_rna28 | 1.611778253 | 4.380833458 | 10.10979409 | 0.008396504 | 0.050839803 | tRNA-Met-CAT |
| AP53_1814 | 0.830379843 | 9.414836082 | 10.10832501 | 0.008400021 | 0.050839803 | LSU ribosomal protein L32p |
| AP53_1784 | -0.791759439 | 7.840220968 | 10.09968369 | 0.008420749 | 0.050839803 | Phage transcriptional repressor |
| AP53_1233 | -0.696082928 | 12.28677147 | 10.09932617 | 0.008421607 | 0.050839803 | ATP-dependent Clp protease, ATP-b |
| AP53_1673 | 0.668194148 | 7.054821001 | 10.09416747 | 0.008434012 | 0.050839803 | Ribosomal RNA small subunit meth |
| AP53_969 | 0.508659906 | 8.451923842 | 10.0628847 | 0.008509697 | 0.050906987 | tRNA pseudouridine synthase B (EC |
| AP53_rna29 | 1.385314138 | 4.81700164 | 10.06090607 | 0.008514511 | 0.050906987 | tRNA-Pseudo-TGA |
| AP53_1380 | 0.514469155 | 8.52007152 | 10.04280669 | 0.008558698 | 0.050906987 | Methionyl-tRNA formyltransferase ( |
| AP53_116 | -0.687956173 | 7.765070502 | 10.03979448 | 0.008566078 | 0.050906987 | YheO-like PAS domain |
| AP53_1813 | 0.537598267 | 6.865511878 | 10.03563017 | 0.008576293 | 0.050906987 | Histidyl-tRNA synthetase (EC 6.1.1.: |
| AP53_936 | 0.819162471 | 8.458939196 | 10.01598754 | 0.008624673 | 0.050960992 | Lipoate-protein ligase A |
| AP53_1420 | 0.891180258 | 7.217537269 | 10.006494 | 0.008648171 | 0.050960992 | Transaldolase (EC 2.2.1.2) |
| AP53_1840 | 0.651471478 | 8.984295955 | 10.00005331 | 0.008664156 | 0.050960992 | Immunodominant antigen A |
| AP53_1328 | 0.93463596 | 3.983716311 | 9.967235844 | 0.008746152 | 0.05128786 | Phage major capsid protein |
| AP53_874 | -0.498318638 | 7.794643657 | 9.955489028 | 0.008775726 | 0.051306275 | Hypothetical similar to thiamin bic |
| AP53_792 | -0.373338213 | 7.475883425 | 9.935105453 | 0.008827324 | 0.051452961 | FIG01115226: hypothetical protei |
| AP53_1155 | 0.946448118 | 11.46796564 | 9.849915654 | 0.009046901 | 0.052574955 | Manganese superoxide dismutase (I |
| AP53_731 | 0.527970503 | 7.025910352 | 9.818647321 | 0.009129116 | 0.052894368 | Hydroxymethylglutaryl-CoA syntha |
| AP53_1238 | 0.540826758 | 9.054908391 | 9.801108214 | 0.009175619 | 0.053005585 | FIG001583: hypothetical protein, ( |
| AP53_265 | 0.783737518 | 9.95527985 | 9.744416064 | 0.009327867 | 0.053710261 | Heat shock protein HtpX (EC 3.4.24 |
| AP53_384 | 1.014841011 | 5.312484241 | 9.735182615 | 0.009352946 | 0.053710261 | Mobile element protein |
| AP53_437 | 0.621633677 | 6.32848962 | 9.71773081 | 0.009400566 | 0.05382448 | Chromosome partition protein smc |
| AP53_1864 | -0.78742707 | 9.373845942 | 9.681154582 | 0.009501306 | 0.054029757 | Chromosome (plasmid) partitionin |
| AP53_680 | 0.618664666 | 10.64839791 | 9.67555666 | 0.009516837 | 0.054029757 | LSU ribosomal protein L21p |
| AP53_1704 | -0.616148765 | 8.338143977 | 9.674444449 | 0.009519926 | 0.054029757 | FIG01115531: hypothetical protei |
| AP53_1588 | 0.624278923 | 10.15300196 | 9.64141471 | 0.009612214 | 0.054054439 | Phosphoglycerate kinase (EC 2.7.2.: |
| AP53_1455 | -0.475144814 | 9.896842533 | 9.64025815 | 0.009615465 | 0.054054439 | ribosomal protein L7Ae family prot |
| AP53_1699 | -0.762056942 | 8.424003252 | 9.634275796 | 0.0096323 | 0.054054439 | Two-component system histidine k |
| AP53_319 | -0.690993367 | 9.304296752 | 9.628743983 | 0.009647898 | 0.054054439 | FIG01114628: hypothetical protei |
| AP53_1605 | -0.381348642 | 7.544310561 | 9.593276184 | 0.009748624 | 0.054054439 | tRNA pseudouridine synthase A (EC |
| AP53_1753 | 0.683430539 | 5.306778081 | 9.571416435 | 0.009811326 | 0.054054439 | Transcriptional regulator, LuxR fam |
| AP53_533 | -0.840080278 | 7.028844904 | 9.562693794 | 0.009836478 | 0.054054439 | Transcriptional regulator, GntR fam |

| | | | | | | |
|---|---|---|---|---|---|---|
| AP53_989 | -0.631364043 | 8.859544528 | 9.547148309 | 0.009881496 | 0.054054439 | ABC transporter membrane-spanni |
| AP53_819 | 1.334984827 | 8.051631786 | 9.547029624 | 0.009881841 | 0.054054439 | PTS system, IIA component, putativ |
| AP53_1041 | 0.609573572 | 5.849702813 | 9.514161688 | 0.009977832 | 0.054054439 | FIG01115822: hypothetical protei |
| AP53_1838 | -0.538373309 | 6.845123703 | 9.512790329 | 0.009981861 | 0.054054439 | L-serine dehydratase, beta subunit |
| AP53_1633 | 0.548488323 | 6.935103688 | 9.510258326 | 0.009989305 | 0.054054439 | hypothetical protein |
| AP53_1078 | 0.539240649 | 7.501289471 | 9.509468884 | 0.009991628 | 0.054054439 | Excinuclease ABC subunit B |
| AP53_95 | -0.626011684 | 6.349589747 | 9.496369493 | 0.010030256 | 0.054054439 | FIG014356: hypothetical protein |
| AP53_1464 | 0.348840445 | 8.759238143 | 9.489136676 | 0.01005166 | 0.054054439 | Histidine triad (HIT) nucleotide-bin |
| AP53_938 | 0.755084394 | 8.540954766 | 9.485920628 | 0.010061195 | 0.054054439 | COG2110, Macro domain, possibly |
| AP53_561 | 0.705514953 | 9.705734837 | 9.478991511 | 0.010081775 | 0.054054439 | ATP synthase B chain (EC 3.6.3.14) |
| AP53_1688 | 0.419500619 | 5.913434124 | 9.476108655 | 0.010090352 | 0.054054439 | Laminin-binding surface protein |
| AP53_741 | 0.58428332 | 9.877237239 | 9.459634066 | 0.010139532 | 0.054054439 | Arsenate reductase (EC 1.20.4.1) |
| AP53_1250 | 0.501401744 | 8.688805681 | 9.441711253 | 0.010193359 | 0.054054439 | Glucokinase (EC 2.7.1.2) |
| AP53_740 | 0.610736067 | 10.38253532 | 9.433242312 | 0.010218912 | 0.054054439 | Phosphopentomutase (EC 5.4.2.7) |
| AP53_1703 | -0.695445374 | 8.116455482 | 9.432047915 | 0.010222521 | 0.054054439 | periplasmic component of efflux sy |
| AP53_1566 | -0.617764283 | 7.086791417 | 9.429313877 | 0.01023079 | 0.054054439 | Transcriptional regulator, CrP/Fnr f |
| AP53_51 | 0.765049854 | 11.07336914 | 9.429131123 | 0.010231343 | 0.054054439 | LSU ribosomal protein L29p (L35e) |
| AP53_565 | 0.918938624 | 10.93748296 | 9.425556619 | 0.010242167 | 0.054054439 | ATP synthase beta chain (EC 3.6.3.1 |
| AP53_288 | 0.598056909 | 7.785351794 | 9.423518856 | 0.010248343 | 0.054054439 | FIG042801: CBS domain containin |
| AP53_1093 | 1.173706484 | 2.754249981 | 9.402348502 | 0.01031277 | 0.054192742 | Mobile element protein |
| AP53_1028 | 1.205014571 | 5.282694774 | 9.396538444 | 0.010330536 | 0.054192742 | Phage terminase, large subunit |
| AP53_1844 | -0.351257253 | 8.19988329 | 9.382925098 | 0.010372304 | 0.054192742 | CDP-diacylglycerol--glycerol-3-pho |
| AP53_1486 | 0.683308319 | 9.62811879 | 9.378396274 | 0.010386244 | 0.054192742 | Acyl carrier protein |
| AP53_193 | -0.411697314 | 6.180584908 | 9.362605883 | 0.010435021 | 0.054291259 | Histidine kinase of the competence |
| AP53_705 | 0.493932203 | 8.668212891 | 9.354209986 | 0.010461067 | 0.054291259 | 2-dehydropantoate 2-reductase (EC |
| AP53_359 | -0.890248975 | 7.623428763 | 9.320081092 | 0.010567739 | 0.054373006 | Integral membrane protein |
| AP53_413 | -0.398260465 | 9.859277469 | 9.298330893 | 0.010636394 | 0.054373006 | Glutaminyl-peptide cyclotransfera |
| AP53_1085 | 0.826670492 | 4.943668624 | 9.292026885 | 0.010656392 | 0.054373006 | PTS system, cellobiose-specific IIB c |
| AP53_488 | -0.97040347 | 7.678716534 | 9.291740424 | 0.010657301 | 0.054373006 | Phage lysin, glycosyl hydrolase, fam |
| AP53_1357 | -1.030910289 | 4.280988049 | 9.289953048 | 0.01066298 | 0.054373006 | conserved hypothetical protein - pl |
| AP53_1517 | -0.341823305 | 7.018966461 | 9.289032475 | 0.010665906 | 0.054373006 | Putative ABC transporter (ATP-bind |
| AP53_613 | 1.193403837 | 5.300012505 | 9.262835357 | 0.010749576 | 0.054373006 | Phage major tail protein |
| AP53_1422 | 0.58976073 | 6.883149016 | 9.250778437 | 0.010788346 | 0.054373006 | NADH peroxidase (EC 1.11.1.1) |
| AP53_1373 | 0.60639563 | 9.092152825 | 9.248140526 | 0.010796851 | 0.054373006 | Streptococcal lipoprotein rotamas |
| AP53_1521 | -0.477305583 | 7.131119651 | 9.239259306 | 0.010825542 | 0.054373006 | Cell surface protein Shp, transfers h |

| | | | | | | |
|---|---|---|---|---|---|---|
| AP53_1421 | 0.70062713 | 6.927983678 | 9.237416285 | 0.010831507 | 0.054373006 | Trans-acting positive regulator |
| AP53_816 | 0.583755552 | 13.52738783 | 9.231628281 | 0.010850267 | 0.054373006 | Collagen-like surface protein |
| AP53_1598 | 0.998287466 | 8.199124008 | 9.223817009 | 0.010875645 | 0.054373006 | DNA-directed RNA polymerase delt |
| AP53_841 | 0.485836035 | 8.131143388 | 9.223706055 | 0.010876006 | 0.054373006 | Putrescine transport ATP-binding p |
| AP53_1074 | 0.637633246 | 9.483726773 | 9.217257232 | 0.010897011 | 0.054373006 | D-alanine--poly(phosphoribitol) lig |
| AP53_163 | 0.769509694 | 3.782831697 | 9.207507957 | 0.010928859 | 0.05439209 | Mobile element protein |
| AP53_1013 | 1.321037223 | 5.972803166 | 9.19796933 | 0.010960125 | 0.054408192 | Phage-related protein |
| AP53_1440 | -1.074783364 | 6.64335366 | 9.181733677 | 0.011013588 | 0.054534118 | hypothetical protein |
| AP53_1833 | 0.443755719 | 8.861494486 | 9.149885773 | 0.011119362 | 0.054917767 | LSU ribosomal protein L9p |
| AP53_1516 | -0.2788911 | 6.984875302 | 9.119216842 | 0.011222362 | 0.055237338 | Putative ABC transporter ATP-bindi |
| AP53_317 | -0.906514412 | 9.054935529 | 9.11295334 | 0.011243537 | 0.055237338 | DNA replication intiation control p |
| AP53_1068 | -0.599823105 | 6.318886915 | 9.105309821 | 0.011269441 | 0.055237338 | Maltose/maltodextrin ABC transpo |
| AP53_452 | 1.365591311 | 1.143102125 | 9.07765131 | 0.011363771 | 0.055518715 | FIG01116748: hypothetical protei |
| AP53_1135 | -0.674543703 | 7.158557208 | 9.059526143 | 0.011426096 | 0.055518715 | Chloride channel protein |
| AP53_532 | -0.511308642 | 8.611940454 | 9.059075275 | 0.011427652 | 0.055518715 | Candidate zinc-binding lipoprotein |
| AP53_1406 | -0.396640839 | 9.002072289 | 9.055133923 | 0.01144126 | 0.055518715 | Cysteine ABC transporter, permease |
| AP53_91 | 0.613747388 | 8.79430569 | 9.0207266 | 0.01156088 | 0.055959274 | Acetate kinase (EC 2.7.2.1) |
| AP53_1817 | -0.496740275 | 5.415938694 | 8.998149166 | 0.01164018 | 0.056202959 | Cadmium efflux system accessory p |
| AP53_1679 | -0.399463517 | 7.966748178 | 8.964991153 | 0.011757816 | 0.056630076 | Chromosome segregation helicase |
| AP53_1159 | 0.388397661 | 8.174832113 | 8.946717561 | 0.011823249 | 0.056804272 | 1-acyl-sn-glycerol-3-phosphate acyl |
| AP53_987 | -0.559558438 | 9.004300497 | 8.933351966 | 0.011871381 | 0.056894695 | Amino acid ABC transporter, aminc |
| AP53_540 | -0.597716662 | 12.75185578 | 8.913162888 | 0.011944529 | 0.056950287 | LSU ribosomal protein L19p |
| AP53_1133 | -0.533851942 | 9.591269898 | 8.910386367 | 0.01195463 | 0.056950287 | FIG01116145: hypothetical protei |
| AP53_1782 | -0.539484809 | 8.412433501 | 8.905891976 | 0.011971003 | 0.056950287 | Phage integrase: site-specific recom |
| AP53_523 | 0.85603995 | 9.522155055 | 8.890956986 | 0.012025602 | 0.057070156 | 2,3-butanediol dehydrogenase, S-al |
| AP53_1144 | 0.519568261 | 6.785620853 | 8.877297506 | 0.012075797 | 0.057168591 | O-methyltransferase family protein |
| AP53_937 | 0.715230238 | 8.459560788 | 8.867275126 | 0.012112785 | 0.057204176 | NAD-dependent protein deacetylas |
| AP53_1360 | -1.3772932 | 9.222909666 | 8.846503097 | 0.012189874 | 0.05733666 | FIG01117062: hypothetical protei |
| AP53_1670 | 0.619378163 | 6.467137102 | 8.843808346 | 0.012199918 | 0.05733666 | hypothetical protein |
| AP53_460 | 0.565562598 | 7.599552137 | 8.813295468 | 0.012314328 | 0.057734569 | Hydrolase (HAD superfamily) |
| AP53_285 | 0.531081977 | 7.505391079 | 8.797280249 | 0.012374889 | 0.057878696 | Glutamate racemase (EC 5.1.1.3) |
| AP53_1405 | -0.319840584 | 9.099489907 | 8.782739839 | 0.012430178 | 0.057997539 | ABC-type multidrug transport syste |
| AP53_685 | -0.375217228 | 8.867203535 | 8.774234845 | 0.012462655 | 0.058009623 | Ribosomal large subunit pseudouri |
| AP53_534 | -0.670434924 | 7.793605183 | 8.721111874 | 0.0126678 | 0.058728633 | Galactosamine-6-phosphate isomer |
| AP53_1023 | 1.238344471 | 7.650861882 | 8.71033579 | 0.012709902 | 0.058728633 | Phage capsid protein |

| | | | | | | |
|---|---|---|---|---|---|---|
| AP53_466 | -0.552895151 | 6.612090354 | 8.706120468 | 0.012726416 | 0.058728633 | 6-phospho-beta-glucosidase (EC 3.2 |
| AP53_1485 | 0.736854339 | 9.463774976 | 8.695854927 | 0.01276674 | 0.058728633 | Enoyl-[acyl-carrier-protein] reducta |
| AP53_pseudo1389 | -0.651828414 | 7.021878509 | 8.688297722 | 0.012796523 | 0.058728633 | D-beta-hydroxybutyrate dehydroge |
| AP53_254 | 0.587993145 | 8.141883276 | 8.687754562 | 0.012798667 | 0.058728633 | FIG007491: hypothetical protein Y |
| AP53_1809 | -0.344732456 | 7.846217249 | 8.663519046 | 0.012894755 | 0.059029997 | Transporter |
| AP53_1436 | 0.851366353 | 7.919112504 | 8.617464664 | 0.013079722 | 0.059500298 | DegV family protein |
| AP53_894 | -0.739664249 | 9.692295089 | 8.612772755 | 0.013098742 | 0.059500298 | Hemolysin III |
| AP53_1839 | -0.375221489 | 6.918112071 | 8.612089867 | 0.013101513 | 0.059500298 | L-serine dehydratase, alpha subunit |
| AP53_239 | -0.471514961 | 8.619236094 | 8.607512111 | 0.013120107 | 0.059500298 | Oligopeptide transport system perr |
| AP53_1730 | 0.325521493 | 8.408576889 | 8.589632778 | 0.01319303 | 0.059691543 | Similar to ribosomal large subunit p |
| AP53_412 | -0.845750445 | 9.840928338 | 8.554273937 | 0.013338669 | 0.060210131 | tmRNA-binding protein SmpB |
| AP53_52 | 0.742687697 | 11.32480098 | 8.537744273 | 0.013407407 | 0.060358265 | SSU ribosomal protein S17p (S11e) |
| AP53_316 | -0.699511754 | 9.497923771 | 8.531453338 | 0.013433679 | 0.060358265 | hypothetical |
| AP53_954 | 0.840024709 | 9.975002689 | 8.496488186 | 0.013580815 | 0.060878435 | SSU ribosomal protein S20p |
| AP53_1370 | -0.455306151 | 7.087991316 | 8.454648414 | 0.013759405 | 0.061444685 | FIG000605: protein co-occurring w |
| AP53_876 | 0.435746057 | 7.431415743 | 8.452082694 | 0.013770447 | 0.061444685 | Thymidine kinase (EC 2.7.1.21) |
| AP53_390 | -0.532460077 | 5.938200776 | 8.405508179 | 0.013972733 | 0.06186366 | Response regulator of the compete |
| AP53_108 | -0.359611641 | 7.882968078 | 8.404374746 | 0.0139777 | 0.06186366 | Multiple sugar metabolism regulato |
| AP53_782 | 0.598742399 | 6.754178647 | 8.401459648 | 0.013990483 | 0.06186366 | Mutator mutT protein (7,8-dihydro |
| AP53_779 | 0.479536508 | 9.525736977 | 8.401152206 | 0.013991832 | 0.06186366 | Glucose-1-phosphate thymidylyltra |
| AP53_808 | 0.525497797 | 7.800966263 | 8.386718771 | 0.014055339 | 0.062003213 | Hypothetical NagD-like phosphatas |
| AP53_1484 | 0.740029261 | 9.517232347 | 8.364282828 | 0.014154739 | 0.062300109 | Malonyl CoA-acyl carrier protein tra |
| AP53_42 | 0.698981045 | 10.79703521 | 8.347743766 | 0.014228548 | 0.062388817 | SSU ribosomal protein S10p (S20e) |
| AP53_464 | -0.699705258 | 5.259576695 | 8.345370135 | 0.014239179 | 0.062388817 | Beta-glucoside bgl operon antiterm |
| AP53_1391 | -0.527039212 | 7.33462873 | 8.331878357 | 0.014299782 | 0.062513235 | D-beta-hydroxybutyrate permease |
| AP53_992 | -0.863080389 | 9.403799049 | 8.31034417 | 0.014397147 | 0.062797441 | Signal peptidase I (EC 3.4.21.89) |
| AP53_618 | 1.035849112 | 4.72583804 | 8.277488091 | 0.014547225 | 0.063137528 | hypothetical phage protein |
| AP53_212 | -0.293086511 | 9.368271815 | 8.269553768 | 0.014583746 | 0.063137528 | Ribosome small subunit-stimulated |
| AP53_1666 | -0.630981066 | 9.740172386 | 8.259967879 | 0.014628014 | 0.063137528 | D-tyrosyl-tRNA(Tyr) deacylase |
| AP53_364 | 0.330323902 | 9.106172219 | 8.25192577 | 0.014665276 | 0.063137528 | S1 RNA binding domain |
| AP53_240 | -0.333483122 | 8.982834402 | 8.251857858 | 0.014665591 | 0.063137528 | Oligopeptide transport system perr |
| AP53_1143 | 0.520115622 | 9.577025249 | 8.25084627 | 0.014670286 | 0.063137528 | Foldase protein prsA 1 precursor (E |
| AP53_780 | 0.42283927 | 9.426762818 | 8.225470195 | 0.014788653 | 0.063400282 | dTDP-4-dehydrorhamnose 3,5-epir |
| AP53_1121 | 0.428652337 | 8.876121787 | 8.213276005 | 0.014845936 | 0.063400282 | DNA polymerase III subunits gamma |
| AP53_447 | 0.691513191 | 4.439548682 | 8.210135966 | 0.014860729 | 0.063400282 | ABC transporter membrane-spanni |

| | | | | | | |
|---|---|---|---|---|---|---|
| AP53_690 | -0.438415017 | 8.924203573 | 8.20493498 | 0.014885271 | 0.063400282 | Carbamoyl-phosphate synthase larg |
| AP53_1608 | -0.345571019 | 7.521305155 | 8.202948592 | 0.014894657 | 0.063400282 | Type I restriction-modification syst |
| AP53_120 | -0.425832576 | 5.734036467 | 8.16139849 | 0.015092605 | 0.063972575 | V-type ATP synthase subunit I (EC 3. |
| AP53_346 | 0.716934275 | 9.215258084 | 8.160894905 | 0.015095023 | 0.063972575 | NAD-dependent oxidoreductase |
| AP53_1350 | -0.608518186 | 3.748987503 | 8.149655975 | 0.015149113 | 0.063974845 | FIG01116817: hypothetical protei |
| AP53_rna39 | 1.671419448 | 3.543319887 | 8.140620185 | 0.015192767 | 0.063974845 | tRNA-Val-TAC |
| AP53_1018 | 1.154849124 | 5.920351528 | 8.134668159 | 0.015221604 | 0.063974845 | Phage protein |
| AP53_1646 | 1.058041632 | 4.87587575 | 8.12223752 | 0.015282041 | 0.063974845 | PTS system IIB component |
| AP53_1470 | -1.045527544 | 8.386196245 | 8.120557454 | 0.015290231 | 0.063974845 | Hydrolase (HAD superfamily) |
| AP53_827 | 0.597866845 | 8.427106352 | 8.119924694 | 0.015293317 | 0.063974845 | Excinuclease ABC subunit C |
| AP53_571 | -0.765522796 | 7.058893138 | 8.110919898 | 0.015337314 | 0.064020918 | hypothetical protein |
| AP53_934 | 0.545964607 | 8.425851427 | 8.094482568 | 0.015418015 | 0.064219674 | Cardiolipin synthetase (EC 2.7.8.-) |
| AP53_276 | -0.532935997 | 8.425597474 | 8.071373204 | 0.01553233 | 0.064342688 | putative arylalkylamine n-acetyltra |
| AP53_260 | -0.768105356 | 7.925695479 | 8.066390985 | 0.015557107 | 0.064342688 | Dicarboxylate/amino acid:cation (N |
| AP53_739 | 0.650905014 | 10.05085262 | 8.058041808 | 0.015598734 | 0.064342688 | Ribose 5-phosphate isomerase A (EC |
| AP53_1301 | -0.525754211 | 8.23204417 | 8.05486805 | 0.015614593 | 0.064342688 | Hyaluronoglucosaminidase (EC 3.2. |
| AP53_487 | -0.541595685 | 9.432426044 | 8.047764729 | 0.015650156 | 0.064342688 | Substrate-specific component ThiT |
| AP53_549 | 0.635460125 | 6.651482013 | 8.040420517 | 0.015687026 | 0.064342688 | Streptolysin S self-immunity protei |
| AP53_362 | 0.362705126 | 9.437692988 | 8.036093636 | 0.015708797 | 0.064342688 | Uridylate kinase (EC 2.7.4.-) |
| AP53_557 | 0.359758606 | 7.132158754 | 8.035310111 | 0.015712743 | 0.064342688 | DNA ligase (EC 6.5.1.2) |
| AP53_rna73 | 0.683124704 | 5.936613833 | 7.974887802 | 0.016020638 | 0.065465387 | tRNA-Ser-GGA |
| AP53_1582 | 0.410637434 | 8.371673484 | 7.957553395 | 0.016110291 | 0.065693435 | Inactive homolog of metal-depende |
| AP53_pseudo812 | 0.616830634 | 8.76666172 | 7.931902244 | 0.016244056 | 0.066090068 | Nucleoside diphosphate kinase (EC |
| AP53_229 | -0.358161274 | 8.192570383 | 7.925878768 | 0.016275658 | 0.066090068 | Undecaprenyl-phosphate N-acetylg |
| AP53_1150 | 0.406988262 | 7.68102015 | 7.905725978 | 0.016381923 | 0.0663827 | S-adenosylmethionine:tRNA ribosy |
| AP53_1310 | 0.43310417 | 8.517374264 | 7.885000621 | 0.01649207 | 0.066554419 | Phage protein |
| AP53_249 | 0.533179283 | 8.951446759 | 7.884849267 | 0.016492878 | 0.066554419 | Nicotinate-nucleotide adenylyltran |
| AP53_1046 | 0.822280272 | 5.864546982 | 7.870147345 | 0.016571552 | 0.066733157 | hypothetical phage protein |
| AP53_320 | 0.357101416 | 8.206756372 | 7.861662166 | 0.016617161 | 0.066778282 | Cytoplasmic copper homeostasis pr |
| AP53_852 | 0.46526738 | 6.142123985 | 7.84676134 | 0.016697619 | 0.06696297 | Voltage-gated chloride channel fam |
| AP53_168 | 0.54445613 | 8.972750008 | 7.834257119 | 0.016765493 | 0.067096541 | tRNA-guanine transglycosylase (EC 2 |
| AP53_1027 | 1.017679747 | 4.361305901 | 7.820131688 | 0.016842564 | 0.067266288 | hypothetical phage protein |
| AP53_104 | -0.33147362 | 7.441903889 | 7.804075836 | 0.016930679 | 0.067479361 | Signal peptidase I (EC 3.4.21.89) |
| AP53_224 | -0.391241804 | 8.276251153 | 7.797426504 | 0.016967332 | 0.067486868 | Amino acid transport ATP-binding p |
| AP53_247 | 0.418593066 | 8.868964733 | 7.774346415 | 0.01709529 | 0.067819464 | GTP-binding protein YqeH, required |

| | | | | | | |
|---|---|---|---|---|---|---|
| AP53_1652 | 0.39011985 | 8.385416906 | 7.769763685 | 0.017120833 | 0.067819464 | FIG01114677: hypothetical protei |
| AP53_1298 | -0.511120177 | 7.18348429 | 7.759539512 | 0.017177985 | 0.067839848 | N-acetylmannosamine kinase (EC 2. |
| AP53_828 | 0.538141247 | 9.672546088 | 7.756346416 | 0.017195881 | 0.067839848 | Oxygen-insensitive NAD(P)H nitrore |
| AP53_553 | 0.770234952 | 7.009642797 | 7.74716755 | 0.017247447 | 0.067860855 | Streptolysin S export transmembra |
| AP53_717 | 0.735716075 | 8.491106844 | 7.74296307 | 0.017271129 | 0.067860855 | DegV family protein |
| AP53_35 | -0.520055509 | 8.844644952 | 7.722806795 | 0.0173852 | 0.068120145 | Holliday junction DNA helicase Ruv |
| AP53_rna64 | 1.418785612 | 4.164097717 | 7.715353753 | 0.017427606 | 0.068120145 | tRNA-Thr-TGT |
| AP53_504 | 0.457517218 | 7.698002855 | 7.712757536 | 0.017442407 | 0.068120145 | FIG01114378: hypothetical protei |
| AP53_374 | 0.494371302 | 8.82937416 | 7.701964001 | 0.017504101 | 0.068223815 | MutT/nudix family protein |
| AP53_266 | 0.417866292 | 9.303906861 | 7.68669383 | 0.017591826 | 0.068428327 | COG1399 protein in cluster with ri |
| AP53_1178 | -1.017609575 | 6.83398811 | 7.671967343 | 0.017676925 | 0.068621823 | Paratox |
| AP53_743 | 0.633650073 | 9.690537483 | 7.643287128 | 0.017844068 | 0.068910562 | Purine nucleoside phosphorylase (E |
| AP53_1307 | -0.86211553 | 4.02732108 | 7.640661856 | 0.017859461 | 0.068910562 | Paratox |
| AP53_5 | 0.342177448 | 8.003714405 | 7.637898318 | 0.017875683 | 0.068910562 | GTP-binding and nucleic acid-bindi |
| AP53_1006 | 1.174665849 | 6.091945859 | 7.634897838 | 0.017893314 | 0.068910562 | Phage lysin |
| AP53_172 | 0.519177917 | 8.017024394 | 7.622433886 | 0.017966779 | 0.06894411 | tRNA-specific adenosine-34 deamir |
| AP53_1012 | 1.13483125 | 5.87453472 | 7.621370217 | 0.017973065 | 0.06894411 | Phage hyaluronidase; Phage endop |
| AP53_pseudo388 | 0.425830541 | 6.228121309 | 7.598831668 | 0.018106881 | 0.06931516 | Mobile element protein |
| AP53_994 | 0.434316698 | 10.55661971 | 7.587869349 | 0.018172396 | 0.06931516 | 6-phosphofructokinase (EC 2.7.1.1 |
| AP53_594 | 0.389861739 | 6.746826299 | 7.584605394 | 0.018191957 | 0.06931516 | Alpha-L-Rha alpha-1,2-L-rhamnosyl |
| AP53_202 | 0.482259684 | 7.286735702 | 7.581158913 | 0.018212639 | 0.06931516 | ABC transporter, predicted N-acety |
| AP53_1022 | 1.648192068 | 6.923782671 | 7.571552745 | 0.018270433 | 0.06932965 | conserved hypothetical protein - pl |
| AP53_1141 | -0.835990965 | 5.045558232 | 7.568659356 | 0.018287883 | 0.06932965 | FIG01114322: hypothetical protei |
| AP53_98 | -0.805058591 | 4.172683808 | 7.554696519 | 0.018372374 | 0.069357692 | FIG107367: tRNA-binding protein |
| AP53_1165 | -0.297059128 | 9.470830918 | 7.554063353 | 0.018376217 | 0.069357692 | membrane protein, putative |
| AP53_597 | 0.865473736 | 8.519968898 | 7.54973998 | 0.018402479 | 0.069357692 | hypothetical protein - phage associ |
| AP53_1472 | 0.697422252 | 9.358208714 | 7.537342263 | 0.018478036 | 0.069453677 | PTS system, mannose-specific IIC co |
| AP53_1299 | -0.527808272 | 6.93179985 | 7.533829687 | 0.018499511 | 0.069453677 | FIG01114249: hypothetical protei |
| AP53_520 | -0.402999707 | 8.789098109 | 7.505445524 | 0.018674136 | 0.069973933 | Cell division transporter, ATP-bindi |
| AP53_1860 | -0.538434828 | 6.937017296 | 7.486281897 | 0.018793145 | 0.07018563 | ABC transporter, permease protein |
| AP53_493 | -0.251241095 | 8.217555346 | 7.484709315 | 0.018802951 | 0.07018563 | Phosphoenolpyruvate carboxylase |
| AP53_70 | 0.686138504 | 10.44975777 | 7.474916357 | 0.018864153 | 0.070278929 | LSU ribosomal protein L17p |
| AP53_883 | -0.26932757 | 8.535576241 | 7.462203004 | 0.018943962 | 0.070441054 | Pneumococcal vaccine antigen A ho |
| AP53_1020 | 1.275827942 | 6.170653395 | 7.422706044 | 0.019194485 | 0.070839811 | Phage protein |
| AP53_1263 | 1.033341034 | 10.24910459 | 7.420628404 | 0.019207772 | 0.070839811 | Acetylornithine deacetylase/Succir |

| | | | | | | |
|---|---|---|---|---|---|---|
| AP53_314 | -0.464768407 | 9.769757424 | 7.414823788 | 0.019244952 | 0.070839811 | DNA polymerase III delta prime sub |
| AP53_967 | 0.801403302 | 9.252986656 | 7.411862529 | 0.019263953 | 0.070839811 | Transcriptional regulator SpxA1 |
| AP53_867 | 0.403908035 | 6.75101341 | 7.409083505 | 0.019281805 | 0.070839811 | Short chain dehydrogenase |
| AP53_682 | 0.872323755 | 10.79805948 | 7.405067117 | 0.01930764 | 0.070839811 | LSU ribosomal protein L27p |
| AP53_931 | -0.445148589 | 6.177726661 | 7.400408132 | 0.01933766 | 0.070839811 | Pyridoxal kinase (EC 2.7.1.35) |
| AP53_1519 | -0.334839191 | 7.460775577 | 7.398081941 | 0.01935267 | 0.070839811 | Heme ABC transporter (Streptococ |
| AP53_1374 | 0.349636713 | 8.160834778 | 7.392387358 | 0.019389473 | 0.070839811 | Two component transcriptional re |
| AP53_1268 | 0.501076751 | 5.460772353 | 7.383948799 | 0.019444163 | 0.070839811 | Transcriptional regulator ArcR esse |
| AP53_1371 | -0.607602482 | 10.82174338 | 7.377503556 | 0.019486059 | 0.070839811 | Cysteine synthase (EC 2.5.1.47) |
| AP53_471 | 0.452109814 | 8.036414201 | 7.377027139 | 0.01948916 | 0.070839811 | Putative metallopeptidase (Zinc) Sp |
| AP53_704 | 0.412018557 | 8.096405396 | 7.357495714 | 0.019616801 | 0.070869013 | Membrane protein |
| AP53_940 | 0.621054851 | 8.107856263 | 7.352502148 | 0.019649594 | 0.070869013 | Luciferase-like monooxygenase (EC |
| AP53_457 | 1.416428389 | 2.788920823 | 7.351601769 | 0.019655514 | 0.070869013 | FIG01117988: hypothetical protei |
| AP53_997 | -0.394728456 | 8.557511037 | 7.34848606 | 0.019676016 | 0.070869013 | ABC-type multidrug transport syste |
| AP53_213 | -0.319692315 | 9.447110586 | 7.347918716 | 0.019679752 | 0.070869013 | Ribulose-phosphate 3-epimerase (E |
| AP53_pseudo73 | 0.439325159 | 6.753476038 | 7.333903851 | 0.019772308 | 0.071070464 | 4-diphosphocytidyl-2-C-methyl-D- |
| AP53_340 | 0.343144358 | 6.586406523 | 7.325996864 | 0.019824756 | 0.071127268 | hypothetical protein |
| AP53_696 | 0.542156952 | 9.102386373 | 7.305513538 | 0.019961398 | 0.071397717 | KH domain RNA binding protein Yl |
| AP53_669 | -0.394852484 | 8.124166179 | 7.301235904 | 0.019990075 | 0.071397717 | Lipoteichoic acid synthase LtaS Typ |
| AP53_674 | 0.287313949 | 8.27484944 | 7.292817178 | 0.020046657 | 0.071397717 | Glutathione reductase (EC 1.8.1.7) |
| AP53_424 | 0.58840973 | 6.560053996 | 7.289086569 | 0.020071792 | 0.071397717 | hypothetical protein |
| AP53_823 | 0.485784565 | 5.578408491 | 7.287268246 | 0.020084056 | 0.071397717 | Two-component sensor kinase, ass |
| AP53_289 | 0.320103686 | 7.004993998 | 7.277958041 | 0.020146993 | 0.071490517 | Tyrosine recombinase XerD |
| AP53_408 | 0.366396926 | 9.562030378 | 7.262761085 | 0.020250229 | 0.071630904 | ATPase |
| AP53_1737 | -0.525311388 | 13.40341086 | 7.259314878 | 0.020273727 | 0.071630904 | Heat shock protein 60 family co-ch |
| AP53_1367 | 0.786889612 | 10.32687169 | 7.255176025 | 0.020301992 | 0.071630904 | Ribosomal subunit interface protei |
| AP53_132 | -0.362383846 | 10.33594975 | 7.250472087 | 0.020334172 | 0.071630904 | Nucleoside-binding protein |
| AP53_365 | -0.80682873 | 7.934240173 | 7.201778377 | 0.020670893 | 0.0725711 | Peptide methionine sulfoxide redu |
| AP53_1491 | -0.646344749 | 14.69533013 | 7.201069104 | 0.020675847 | 0.0725711 | Chaperone protein DnaK |
| AP53_1058 | 0.566652246 | 4.464736719 | 7.133583699 | 0.021153712 | 0.07411436 | hypothetical protein |
| AP53_231 | 0.480590597 | 8.46742801 | 7.118913955 | 0.021259325 | 0.074350181 | Iron-sulfur cluster assembly ATPase |
| AP53_754 | -0.51802682 | 9.268599668 | 7.11304072 | 0.021301785 | 0.074364685 | Dihydroorotase (EC 3.5.2.3) |
| AP53_1829 | 0.638744872 | 9.962085059 | 7.091586202 | 0.02145775 | 0.074686166 | SSU ribosomal protein S4p (S9e) |
| AP53_1411 | -0.381154274 | 8.762450511 | 7.089795459 | 0.02147083 | 0.074686166 | Cell division protein FtsL |
| AP53_1359 | -0.958543953 | 10.7346197 | 7.052642552 | 0.021744349 | 0.075502293 | Phage antirepressor protein |

| | | | | | | |
|---|---|---|---|---|---|---|
| AP53_1837 | 0.385016166 | 8.456360843 | 7.040073888 | 0.021837819 | 0.075691442 | tRNA-specific 2-thiouridylase Mnm |
| AP53_13 | -0.468139138 | 9.357275859 | 7.019285925 | 0.021993469 | 0.076095051 | Hypoxanthine-guanine phosphorib |
| AP53_1772 | 0.758841397 | 11.24275464 | 7.005539551 | 0.022097122 | 0.076317641 | Ortholog of S. aureus MRSA252 (BX |
| AP53_1738 | -0.528203328 | 10.31251749 | 6.978281188 | 0.02230439 | 0.076896664 | ATP-dependent Clp protease ATP-bi |
| AP53_1552 | -0.252643253 | 7.991986028 | 6.945492985 | 0.022556789 | 0.07762895 | Colicin V production protein |
| AP53_rna21 | 1.315951216 | 4.774862161 | 6.934444606 | 0.022642605 | 0.077786364 | tRNA-Leu-TAG |
| AP53_724 | 0.383728299 | 7.381698237 | 6.926473379 | 0.022704761 | 0.077862088 | Regulation of D-alanyl-lipoteichoic |
| AP53_1315 | 0.71225182 | 3.418809219 | 6.914874595 | 0.022795569 | 0.07789282 | FIG01116522: hypothetical protei |
| AP53_242 | -0.371903282 | 9.151222099 | 6.911538943 | 0.022821764 | 0.07789282 | Oligopeptide transport ATP-bindin |
| AP53_691 | -0.451291773 | 8.599600017 | 6.909967918 | 0.022834114 | 0.07789282 | periplasmic component of efflux sy |
| AP53_1698 | -0.527731275 | 9.395822895 | 6.882095405 | 0.023054552 | 0.078400637 | Immunogenic secreted protein |
| AP53_751 | -0.296298581 | 9.44213199 | 6.876667148 | 0.023097778 | 0.078400637 | Cystine ABC transporter, periplasm |
| AP53_1739 | -0.999197568 | 9.058686756 | 6.875867393 | 0.023104155 | 0.078400637 | Transcriptional regulator CtsR |
| AP53_192 | 0.397137572 | 9.179226164 | 6.868175661 | 0.023165591 | 0.078471924 | Glutamyl-tRNA synthetase (EC 6.1.1 |
| AP53_44 | 0.638170624 | 11.31807185 | 6.862045573 | 0.023214694 | 0.078501256 | LSU ribosomal protein L4p (L1e) |
| AP53_1009 | 0.891510582 | 5.668230454 | 6.842515939 | 0.02337196 | 0.078843198 | FIG01116522: hypothetical protei |
| AP53_837 | 0.521606782 | 9.548222719 | 6.830535768 | 0.023469062 | 0.078843198 | Dihydropteroate synthase (EC 2.5.1 |
| AP53_1842 | -0.383279298 | 6.89698383 | 6.825676928 | 0.023508581 | 0.078843198 | ATPase component of general energ |
| AP53_170 | -0.654536185 | 9.354608329 | 6.824546796 | 0.023517784 | 0.078843198 | Substrate-specific component BioY |
| AP53_1180 | -0.596676679 | 6.123507441 | 6.824408143 | 0.023518914 | 0.078843198 | Streptococcal pyrogenic exotoxin ( |
| AP53_1671 | -0.528905838 | 6.337641092 | 6.814193193 | 0.0236023 | 0.078986318 | Exodeoxyribonuclease III (EC 3.1.11 |
| AP53_646 | 0.670577845 | 3.949294806 | 6.802086659 | 0.023701585 | 0.07918206 | hypothetical phage protein |
| AP53_1804 | 0.327283014 | 6.605851342 | 6.793521587 | 0.023772128 | 0.079281274 | DNA mismatch repair protein MutS |
| AP53_895 | -0.647165489 | 9.171990467 | 6.783467303 | 0.023855257 | 0.079422048 | FIG01115071: hypothetical protei |
| AP53_795 | 0.594776479 | 10.58190416 | 6.774272338 | 0.023931585 | 0.079539738 | Acetoin dehydrogenase E1 compon |
| AP53_465 | -0.535133494 | 6.15883924 | 6.757665006 | 0.024070184 | 0.079863637 | PTS system, beta-glucoside-specific |
| AP53_1520 | -0.323176135 | 7.600032679 | 6.747148693 | 0.024158446 | 0.079964811 | Heme transporter IsdDEF, lipoprote |
| AP53_353 | -0.370247664 | 9.770017578 | 6.744222547 | 0.024183073 | 0.079964811 | Mn-dependent transcriptional regu |
| AP53_1066 | -0.379074687 | 7.930079305 | 6.732102019 | 0.024285403 | 0.080166609 | Maltose/maltodextrin ABC transpo |
| AP53_726 | 0.365661248 | 7.212955973 | 6.71498917 | 0.024430763 | 0.080509526 | Mevalonate kinase (EC 2.7.1.36) |
| AP53_617 | 1.034663332 | 4.683457916 | 6.680624863 | 0.024725813 | 0.081216052 | hypothetical phage protein |
| AP53_93 | -0.380699981 | 8.777350987 | 6.680274371 | 0.024728844 | 0.081216052 | Pyrroline-5-carboxylate reductase ( |
| AP53_1063 | -0.424948232 | 6.562298696 | 6.666585811 | 0.024847576 | 0.081380169 | Maltodextrin phosphorylase (EC 2.4 |
| AP53_1095 | 0.733104394 | 3.813457285 | 6.664851309 | 0.024862669 | 0.081380169 | Mobile element protein |
| AP53_1076 | 0.48997187 | 8.966213956 | 6.629638897 | 0.025171461 | 0.081688268 | D-alanine--poly(phosphoribitol) lig |

| | | | | | | |
|---|---|---|---|---|---|---|
| AP53_1007 | 1.03110626 | 5.788601678 | 6.627612009 | 0.025189374 | 0.081688268 | Phage holin |
| AP53_1388 | -0.568832971 | 7.381478696 | 6.626408913 | 0.025200015 | 0.081688268 | Butyrate-acetoacetate CoA-transfer |
| AP53_752 | -0.289108988 | 9.431559898 | 6.625605261 | 0.025207125 | 0.081688268 | Cystine transport system permease |
| AP53_750 | -0.59615642 | 8.941525709 | 6.625516673 | 0.025207909 | 0.081688268 | 6-aminohexanoate-cyclic-dimer hy |
| AP53_334 | -0.640147171 | 7.109344875 | 6.625358254 | 0.025209311 | 0.081688268 | Ribonucleotide reduction protein I |
| AP53_1678 | -0.41610229 | 8.21689848 | 6.61524995 | 0.025298961 | 0.081842139 | Anthranilate synthase, amidotrans |
| AP53_1403 | 0.294948496 | 8.232437424 | 6.603491026 | 0.02540373 | 0.081910793 | Thioredoxin reductase (EC 1.8.1.9) |
| AP53_421 | 0.267685173 | 8.761816951 | 6.599001301 | 0.025443869 | 0.081910793 | Catabolite control protein A |
| AP53_1418 | -0.519355439 | 7.937633942 | 6.598675544 | 0.025446784 | 0.081910793 | FIG01113961: hypothetical protei |
| AP53_67 | 0.73063814 | 10.20321776 | 6.588953731 | 0.025533971 | 0.082055362 | SSU ribosomal protein S13p (S18e) |
| AP53_264 | 0.504770802 | 9.718548387 | 6.569326872 | 0.025711079 | 0.082418991 | LemA protein |
| AP53_958 | 0.424414396 | 7.312010699 | 6.567013666 | 0.02573205 | 0.082418991 | Phosphate transport system regula |
| AP53_rna22 | 1.369747299 | 4.187408477 | 6.559676743 | 0.025798698 | 0.082481312 | tRNA-Thr-TGT |
| AP53_1134 | -0.542575108 | 8.551652088 | 6.551136336 | 0.025876539 | 0.082481312 | Ribonucleotide reductase of class I |
| AP53_1329 | 0.726536078 | 3.992006818 | 6.550867942 | 0.02587899 | 0.082481312 | Phage protein |
| AP53_102 | -0.440399436 | 6.187645192 | 6.529648601 | 0.026073632 | 0.082965444 | Transcriptional regulator, RofA-like |
| AP53_964 | 0.294187391 | 6.85891838 | 6.514369872 | 0.026214867 | 0.083217138 | tRNA and rRNA cytosine-C5-methyl |
| AP53_1561 | 0.450758262 | 8.081135354 | 6.509255704 | 0.026262346 | 0.083217138 | Beta-lactamase, class C |
| AP53_36 | -0.301406846 | 9.058116692 | 6.507211496 | 0.026281353 | 0.083217138 | Low molecular weight protein tyro |
| AP53_1851 | -0.345120015 | 9.03965394 | 6.486422783 | 0.02647558 | 0.083612578 | UTP--glucose-1-phosphate uridylyl |
| AP53_366 | -0.481584587 | 8.651241561 | 6.484631806 | 0.026492393 | 0.083612578 | FIG01114318: hypothetical protei |
| AP53_1140 | -0.738025113 | 6.040327791 | 6.459356267 | 0.02673104 | 0.084228812 | Transcriptional regulator, Cro/CI fa |
| AP53_1161 | -0.598892564 | 7.283232404 | 6.453977286 | 0.026782158 | 0.08425311 | COG2827: putative endonuclease c |
| AP53_344 | 0.505014059 | 7.517459375 | 6.439810271 | 0.026917353 | 0.084541395 | FIG01114304: hypothetical protei |
| AP53_45 | 0.671002012 | 11.20939483 | 6.405367361 | 0.027249459 | 0.085446204 | LSU ribosomal protein L23p (L23Ae |
| AP53_1518 | -0.309646791 | 6.978776583 | 6.400654282 | 0.027295284 | 0.085451848 | Heme ABC transporter (Streptococ |
| AP53_1705 | -0.338013937 | 7.659559924 | 6.363096521 | 0.027663768 | 0.086190942 | FIG01116435: hypothetical protei |
| AP53_195 | -0.373595624 | 6.691933789 | 6.359865478 | 0.027695745 | 0.086190942 | Response regulator of the compete |
| AP53_492 | 0.427911794 | 7.705728591 | 6.357763143 | 0.027716576 | 0.086190942 | Oligoendopeptidase F |
| AP53_1414 | -0.477805307 | 8.473384018 | 6.356017647 | 0.027733885 | 0.086190942 | Gamma-glutamyl phosphate reduc |
| AP53_1775 | -0.559677829 | 11.20879042 | 6.354051746 | 0.027753395 | 0.086190942 | Transcriptional regulator SpxA2 |
| AP53_463 | -0.467637556 | 5.479056278 | 6.348348463 | 0.027810088 | 0.086229044 | Multidrug resistance protein B |
| AP53_1825 | -0.357145541 | 6.740475722 | 6.333803107 | 0.027955306 | 0.086541066 | FIG04612: Integral membrane prot |
| AP53_1563 | -0.658860267 | 8.225798925 | 6.305336481 | 0.028242144 | 0.087289812 | Integral membrane protein |
| AP53_474 | 0.379125539 | 9.129672143 | 6.297285955 | 0.028323902 | 0.087403328 | Prolipoprotein diacylglyceryl trans |

| | | | | | | |
|---|---|---|---|---|---|---|
| AP53_335 | -0.583785536 | 8.142871266 | 6.253649665 | 0.028772014 | 0.088564526 | Ribonucleotide reductase of class Il |
| AP53_1320 | 0.867777358 | 3.6637731 | 6.251774911 | 0.028791456 | 0.088564526 | Phage tail length tape-measure pro |
| AP53_178 | -0.934421379 | 11.23122741 | 6.232343843 | 0.02899389 | 0.089046109 | Transcriptional regulator, RofA-like |
| AP53_1170 | -0.349860082 | 8.008830292 | 6.224619679 | 0.029074834 | 0.089153639 | Substrate-specific component NiaX |
| AP53_1285 | -0.528160297 | 8.215176469 | 6.198951841 | 0.029345766 | 0.089824265 | 2-keto-3-deoxy-D-arabino-heptulos |
| AP53_1556 | -0.594406525 | 8.243803113 | 6.195155522 | 0.029386094 | 0.089824265 | RecD-like DNA helicase YrrC |
| AP53_86 | -0.855172819 | 3.208787068 | 6.177081744 | 0.029579 | 0.090271759 | Late competence protein ComGD, a |
| AP53_1696 | 0.31150494 | 8.493555226 | 6.170422292 | 0.02965046 | 0.090347791 | M protein trans-acting positive regu |
| AP53_806 | 0.477215097 | 6.347657363 | 6.15908982 | 0.029772541 | 0.090497748 | Hypothetical radical SAM family en |
| AP53_759 | 0.794200619 | 8.73137515 | 6.157143201 | 0.029793571 | 0.090497748 | FIG01114490: hypothetical protei |
| AP53_1232 | 1.015361344 | 8.0946323 | 6.152893472 | 0.029839546 | 0.090497748 | FIG01114188: hypothetical protei |
| AP53_148 | 0.705937247 | 6.799100627 | 6.146693541 | 0.029906771 | 0.090560129 | Probable L-ascorbate-6-phosphate |
| AP53_1003 | -0.553298694 | 6.570567463 | 6.137240529 | 0.030009618 | 0.09061215 | Streptococcal pyrogenic exotoxin k |
| AP53_619 | 1.018182488 | 5.091305753 | 6.136534455 | 0.030017317 | 0.09061215 | Phage major capsid protein |
| AP53_1340 | -0.51874312 | 4.087828755 | 6.123299666 | 0.030162067 | 0.090818917 | conserved hypothetical protein - pl |
| AP53_1017 | 0.932117477 | 6.274401588 | 6.121720603 | 0.030179393 | 0.090818917 | Phage major tail protein |
| AP53_1089 | -0.76396464 | 6.462081772 | 6.10965545 | 0.03031217 | 0.090859417 | Ribosyl nicotinamide transporter, I |
| AP53_884 | -0.433428372 | 8.408772709 | 6.097168381 | 0.030450329 | 0.090859417 | ABC transporter, ATP-binding/perm |
| AP53_1382 | -0.418753339 | 9.307646867 | 6.096130606 | 0.030461845 | 0.090859417 | DNA-directed RNA polymerase ome |
| AP53_573 | 0.46177954 | 6.896267721 | 6.094751501 | 0.030477157 | 0.090859417 | Phenylalanyl-tRNA synthetase alpha |
| AP53_241 | -0.388274879 | 9.196140432 | 6.091406378 | 0.030514335 | 0.090859417 | Oligopeptide transport ATP-binding |
| AP53_1314 | 0.588254178 | 4.001388929 | 6.086462357 | 0.030569383 | 0.090859417 | Phage protein |
| AP53_461 | 0.380190777 | 7.991869669 | 6.085587516 | 0.030579136 | 0.090859417 | Hydrolase (HAD superfamily) |
| AP53_1107 | 0.315867611 | 7.126780371 | 6.082789845 | 0.03061035 | 0.090859417 | 23S rRNA (Uracil-5-) -methyltransfe |
| AP53_620 | 0.893330883 | 4.71110218 | 6.081930719 | 0.030619943 | 0.090859417 | Prophage Clp protease-like protein |
| AP53_591 | 0.341419317 | 6.796622764 | 6.078261331 | 0.030660957 | 0.090859417 | rhamnose-containing polysacharid |
| AP53_689 | -0.616689419 | 9.352042618 | 6.072724288 | 0.030722971 | 0.090904402 | Carbamoyl-phosphate synthase sma |
| AP53_453 | 1.858026478 | 1.390259302 | 6.053456178 | 0.030939946 | 0.091407054 | FIG01115104: hypothetical protei |
| AP53_315 | -0.630684217 | 8.989167006 | 6.04585834 | 0.031026008 | 0.091499474 | hypothetical protein |
| AP53_1343 | -0.620201846 | 4.452770568 | 6.042379475 | 0.031065509 | 0.091499474 | Phage protein |
| AP53_1305 | -0.32167722 | 7.406840544 | 6.038148599 | 0.031113631 | 0.09150236 | Putative two-component responsib |
| AP53_1567 | -0.532377564 | 8.47442752 | 6.033803547 | 0.031163144 | 0.091509323 | Xaa-Pro dipeptidyl-peptidase (EC 3. |
| AP53_313 | -0.506922564 | 9.55902715 | 6.020399748 | 0.031316481 | 0.091820678 | Thymidylate kinase (EC 2.7.4.9) |
| AP53_1064 | -0.498447628 | 6.36318039 | 6.013439047 | 0.031396466 | 0.091916351 | 4-alpha-glucanotransferase (amylor |
| AP53_258 | -0.498865957 | 7.671083701 | 5.994842533 | 0.031611363 | 0.092245885 | Methionine ABC transporter perme |

| | | | | | | |
|---|---|---|---|---|---|---|
| AP53_1600 | 0.343994739 | 6.597198535 | 5.990255948 | 0.031664635 | 0.092245885 | Potassium efflux system KefA protei |
| AP53_rna65 | 1.306650594 | 4.639511609 | 5.984639665 | 0.031730014 | 0.092245885 | tRNA-Leu-TAG |
| AP53_1043 | 0.436659245 | 6.060397034 | 5.984600155 | 0.031730474 | 0.092245885 | FIG01115915: hypothetical protei |
| AP53_956 | 0.683981816 | 7.171513726 | 5.98321263 | 0.031746652 | 0.092245885 | Two component system response re |
| AP53_1081 | 0.888687125 | 4.303329124 | 5.976919593 | 0.031820147 | 0.092321233 | FIG01114324: hypothetical protei |
| AP53_1385 | -0.663132772 | 6.421605107 | 5.957234681 | 0.032051363 | 0.092754486 | Transcriptional regulators, LysR fam |
| AP53_778 | -0.304681573 | 7.90192138 | 5.956074874 | 0.032065049 | 0.092754486 | Glycine/D-amino acid oxidases fam |
| AP53_1260 | 0.336091478 | 8.863973916 | 5.941722846 | 0.032234983 | 0.093107296 | 16S rRNA (guanine(966)-N(2))-meth |
| AP53_1261 | 0.413738713 | 5.821106996 | 5.930638803 | 0.032366963 | 0.093349591 | Aspartate--ammonia ligase (EC 6.3.: |
| AP53_723 | 0.500294218 | 7.192492431 | 5.918220882 | 0.032515595 | 0.093608634 | GTP pyrophosphokinase (EC 2.7.6.5 |
| AP53_1827 | -0.598971579 | 7.738498587 | 5.914486593 | 0.032560452 | 0.093608634 | Phage infection protein |
| AP53_1220 | 0.243804121 | 8.806395054 | 5.907195343 | 0.032648248 | 0.093608634 | FIG01115348: hypothetical protei |
| AP53_794 | 0.660373339 | 9.90566459 | 5.902015004 | 0.032710799 | 0.093608634 | Acetoin dehydrogenase E1 compon |
| AP53_1326 | 1.10963022 | 4.142163145 | 5.901721655 | 0.032714346 | 0.093608634 | Phage minor capsid protein |
| AP53_1341 | -0.964493942 | 4.226506109 | 5.899093202 | 0.032746143 | 0.093608634 | Phage protein |
| AP53_1437 | -0.363825903 | 7.885640041 | 5.871643371 | 0.033080432 | 0.094425175 | Transcriptional regulator, TetR fam |
| AP53_1410 | -0.347603426 | 9.086127775 | 5.854723145 | 0.03328853 | 0.094879643 | Cell division protein FtsI [Peptidogl |
| AP53_1106 | 0.264761429 | 7.140266528 | 5.846139154 | 0.033394703 | 0.095042695 | Uridine phosphorylase (EC 2.4.2.3) |
| AP53_462 | 0.379449728 | 8.914955832 | 5.776258363 | 0.034274323 | 0.09739107 | Signal recognition particle receptor |
| AP53_1504 | -0.368385388 | 6.423323182 | 5.772673564 | 0.034320191 | 0.09739107 | Nicotinamidase (EC 3.5.1.19) |
| AP53_430 | -0.398223207 | 7.043124354 | 5.760694953 | 0.034473996 | 0.09768471 | 3-ketoacyl-CoA thiolase (EC 2.3.1.1 |
| AP53_rna20 | 1.488292025 | 4.503413543 | 5.754060007 | 0.034559544 | 0.097755369 | tRNA-Lys-TTT |
| AP53_1821 | -0.920850994 | 5.51795796 | 5.750955549 | 0.034599659 | 0.097755369 | Transcriptional regulator |
| AP53_1090 | 0.850850866 | 7.046293108 | 5.745233142 | 0.034673749 | 0.097822308 | FIG01114098: hypothetical protei |
| AP53_1439 | -0.369867839 | 8.381408097 | 5.730932673 | 0.034859737 | 0.098032247 | INTEGRAL MEMBRANE PROTEIN (Rh |
| AP53_552 | 0.634358409 | 6.275588373 | 5.727912612 | 0.034899168 | 0.098032247 | Streptolysin S export transmembra |
| AP53_1452 | -0.286360603 | 8.900423862 | 5.72588744 | 0.03492564 | 0.098032247 | putative esterase |
| AP53_287 | 0.387559156 | 7.802514992 | 5.724011014 | 0.034950188 | 0.098032247 | FIG009886: phosphoesterase |
| AP53_234 | -0.276661498 | 8.561751269 | 5.719386507 | 0.035010777 | 0.098060489 | Putative iron-sulfur cluster assembl |
| AP53_1580 | 0.356825173 | 8.126588309 | 5.707709294 | 0.035164332 | 0.098348657 | TsaD/Kae1/Qri7 protein, required f |
| AP53_117 | -0.289483849 | 7.004833486 | 5.680599777 | 0.035523948 | 0.099078715 | Endoribonuclease L-PSP |
| AP53_548 | 0.582712958 | 7.536666595 | 5.678077913 | 0.035557626 | 0.099078715 | Streptolysin S biosynthesis protein |
| AP53_1781 | -0.453564783 | 9.060982847 | 5.676516374 | 0.035578498 | 0.099078715 | DNA mismatch repair protein MutL |
| AP53_1783 | -0.489855306 | 8.552077433 | 5.663853015 | 0.035748307 | 0.09919063 | hypothetical protein |
| AP53_1117 | 0.413091954 | 8.568234233 | 5.654749497 | 0.035870981 | 0.09919063 | S-adenosylmethionine synthetase ( |

| | | | | | | |
|---|---|---|---|---|---|---|
| AP53_1495 | -0.408719749 | 9.023869357 | 5.652933878 | 0.035895508 | 0.09919063 | D-alanyl-D-alanine carboxypeptidas |
| AP53_1482 | 0.677589214 | 10.11432222 | 5.652528216 | 0.035900991 | 0.09919063 | 3-oxoacyl-[acyl-carrier-protein] syn |
| AP53_1394 | -0.476923661 | 10.01987976 | 5.651446035 | 0.035915622 | 0.09919063 | FIG001721: Predicted N6-adenine- |
| AP53_451 | 1.953738063 | 0.454461434 | 5.650730287 | 0.035925303 | 0.09919063 | FIG01114600: hypothetical protei |
| AP53_1543 | -0.369728012 | 10.58250761 | 5.631582905 | 0.036185451 | 0.099766988 | SSU ribosomal protein S18p @ SSU |
| AP53_248 | 0.351144273 | 8.921296728 | 5.626570371 | 0.036253927 | 0.099814004 | FIG004454: RNA binding protein |
| AP53_1509 | -0.343396945 | 7.486170921 | 5.620602555 | 0.036335657 | 0.099897323 | L-asparaginase (EC 3.5.1.1) |
| AP53_303 | -0.534273799 | 6.727022064 | 5.616828562 | 0.036387456 | 0.099898236 | ABC-type Fe3+-siderophore transpo |
| AP53_1862 | 0.669729434 | 8.378678594 | 5.609001818 | 0.036495164 | 0.099927292 | LSU m3Psi1915 methyltransferase |
| AP53_1289 | 0.310642217 | 8.056980692 | 5.608578232 | 0.036501005 | 0.099927292 | SAM-dependent methyltransferase |
| AP53_951 | -0.404771699 | 9.597594622 | 5.602786561 | 0.036580969 | 0.100005155 | Cytidine deaminase (EC 3.5.4.5) |
| AP53_541 | 0.279380383 | 8.691959385 | 5.57270037 | 0.036999764 | 0.100891883 | hydrolase, haloacid dehalogenase-l |
| AP53_rna31 | 1.248430439 | 3.575514669 | 5.572021181 | 0.037009284 | 0.100891883 | tRNA-Phe-GAA |
| AP53_1132 | -0.578950435 | 9.079262359 | 5.560914317 | 0.037165393 | 0.101132934 | Ribonucleotide reductase of class II |
| AP53_1524 | -0.426459997 | 10.72683241 | 5.557785433 | 0.037209512 | 0.101132934 | Immunogenic secreted protein |
| AP53_1544 | 0.39322338 | 11.31501403 | 5.55463373 | 0.037254017 | 0.101132934 | Single-stranded DNA-binding prote |
| AP53_1176 | -0.623530176 | 10.08705089 | 5.545248136 | 0.03738693 | 0.101352 | FIG01114217: hypothetical protei |
| AP53_1416 | -0.495867644 | 9.717240124 | 5.534403509 | 0.037541215 | 0.101625526 | ABC transporter permease protein |
| AP53_1554 | -0.406771009 | 8.751635964 | 5.530807517 | 0.037592544 | 0.101625526 | Ribonuclease HIII (EC 3.1.26.4) |
| AP53_41 | -0.297328176 | 7.04652198 | 5.522039212 | 0.037718055 | 0.101823012 | Multi antimicrobial extrusion (MAT |
| AP53_1164 | -0.404491992 | 9.395850668 | 5.514235739 | 0.03783018 | 0.101983859 | Peptide chain release factor 3 |
| AP53_1231 | -0.589778953 | 6.946548077 | 5.479121674 | 0.038339701 | 0.103196512 | ABC transporter membrane-spanni |
| AP53_984 | -0.513922034 | 8.429015997 | 5.475935721 | 0.038386338 | 0.103196512 | Sodium/glycine symporter GlyP |
| AP53_1257 | -0.280581202 | 7.218499909 | 5.441321307 | 0.038897454 | 0.104352722 | LacI family regulatory protein |
| AP53_1147 | -0.746188569 | 9.234701475 | 5.439542386 | 0.038923942 | 0.104352722 | Competence protein CoiA |
| AP53_825 | 0.423832435 | 5.835413574 | 5.425684566 | 0.039131027 | 0.104763204 | Ferric iron ABC transporter, iron-bi |
| AP53_1718 | -0.320899683 | 6.728872894 | 5.410394601 | 0.039361053 | 0.10523389 | Pyruvate formate-lyase (EC 2.3.1.54 |
| AP53_648 | -0.683577767 | 7.381473531 | 5.405893899 | 0.039429073 | 0.105270743 | Phage antirepressor protein |
| AP53_1707 | -0.474856667 | 11.49237008 | 5.380899752 | 0.039809394 | 0.10602992 | Peptidylproline cis-trans-isomerase |
| AP53_622 | 0.52962245 | 3.759174187 | 5.38003215 | 0.039822675 | 0.10602992 | Phage terminase, large subunit |
| AP53_1381 | -0.454749065 | 7.592901113 | 5.371098554 | 0.039959737 | 0.106159473 | Helicase PriA essential for oriC/Dna |
| AP53_1454 | -0.44503415 | 11.049852 | 5.366848814 | 0.040025137 | 0.106159473 | Translation initiation factor 2 |
| AP53_1142 | 0.469083597 | 8.075777564 | 5.3661819 | 0.040035412 | 0.106159473 | Alanyl-tRNA synthetase (EC 6.1.1.7) |
| AP53_1302 | -0.427630847 | 7.945785161 | 5.354820495 | 0.040210942 | 0.106378896 | Transcriptional repressor of arabino |
| AP53_1356 | -0.654743589 | 3.936521777 | 5.35039346 | 0.040279589 | 0.106378896 | phage protein |

| | | | | | | |
|---|---|---|---|---|---|---|
| AP53_rna40 | 1.409594704 | 3.861167023 | 5.350200729 | 0.040282581 | 0.106378896 | tRNA-Gly-TCC |
| AP53_1849 | -0.303976201 | 8.259439319 | 5.342444788 | 0.040403198 | 0.106475996 | UDP-glucose dehydrogenase in hyal |
| AP53_993 | 0.431632679 | 11.49556352 | 5.337847811 | 0.040474893 | 0.106475996 | Pyruvate kinase (EC 2.7.1.40) |
| AP53_985 | -0.294241403 | 8.1689722 | 5.331658534 | 0.040571664 | 0.106475996 | Cobalt-zinc-cadmium resistance pr |
| AP53_1768 | -1.083115234 | 7.359813173 | 5.330423047 | 0.040591014 | 0.106475996 | FIG01115181: hypothetical protei |
| AP53_1331 | 0.521938983 | 7.374790994 | 5.330256074 | 0.040593631 | 0.106475996 | hypothetical protein, phage associ |
| AP53_1538 | 0.405055073 | 8.55751471 | 5.325171985 | 0.040673382 | 0.106541207 | dCMP deaminase (EC 3.5.4.12); Lat |
| AP53_1604 | 0.323286315 | 7.401009618 | 5.317808962 | 0.040789217 | 0.106633019 | Novel pyridoxal kinase, thiD family |
| AP53_18 | -0.528163945 | 4.922451675 | 5.315963804 | 0.040818307 | 0.106633019 | DNA recombination and repair pro |
| AP53_640 | -0.426294535 | 4.209067475 | 5.306361913 | 0.04097009 | 0.106885679 | hypothetical phage protein |
| AP53_1492 | -0.462150794 | 14.03692929 | 5.292792929 | 0.041185744 | 0.107304067 | Heat shock protein GrpE |
| AP53_94 | -0.269707755 | 8.902392534 | 5.282003749 | 0.041358194 | 0.107608921 | Glutamyl aminopeptidase (EC 3.4.1 |
| AP53_1435 | -0.327983859 | 8.804048446 | 5.266405935 | 0.041609043 | 0.108116671 | FIG01116130: hypothetical protei |
| AP53_333 | -0.593131728 | 7.235476923 | 5.262079691 | 0.041678944 | 0.108153515 | Ribonucleotide reductase of class I |
| AP53_1625 | 0.612355405 | 6.69419007 | 5.253143673 | 0.041823773 | 0.108384438 | DNA-damage-inducible protein J |
| AP53_695 | 0.445020457 | 8.830345711 | 5.249677766 | 0.04188011 | 0.108385724 | SSU ribosomal protein S16p |
| AP53_834 | 0.409394627 | 7.083534567 | 5.212618432 | 0.042488238 | 0.109704801 | Polysaccharide deacetylase |
| AP53_1798 | -0.679601576 | 4.356196758 | 5.210713994 | 0.042519775 | 0.109704801 | Phage protein |
| AP53_236 | -0.426097042 | 8.985063389 | 5.206452162 | 0.042590453 | 0.109704801 | D-alanyl-D-alanine carboxypeptida |
| AP53_1770 | -0.300703495 | 7.59429478 | 5.204921118 | 0.042615879 | 0.109704801 | Competence-induced protein Ccs4 |
| AP53_687 | -0.379517495 | 8.680041329 | 5.194248773 | 0.042793617 | 0.110016439 | Uracil permease |
| AP53_856 | -0.383859942 | 6.912105337 | 5.182748265 | 0.042986149 | 0.110255978 | DNA repair protein RadC |
| AP53_1828 | -0.559748206 | 5.847888352 | 5.178775376 | 0.043052902 | 0.110255978 | Transcriptional regulator, TetR fam |
| AP53_1229 | -0.252861012 | 6.707222003 | 5.174634111 | 0.043122618 | 0.110255978 | Deoxyribodipyrimidine photolyase |
| AP53_1306 | 0.359091956 | 6.047640193 | 5.171515718 | 0.043175203 | 0.110255978 | RNA methyltransferase, TrmA famil |
| AP53_1674 | 0.440797117 | 7.214747101 | 5.169107692 | 0.043215863 | 0.110255978 | Ribosomal protein L11 methyltran |
| AP53_185 | -0.537247908 | 8.235030862 | 5.167313357 | 0.04324619 | 0.110255978 | Lipid A export ATP-binding/permea |
| AP53_422 | -0.311918941 | 10.12176022 | 5.165053762 | 0.043284418 | 0.110255978 | Glycosyltransferase LafB, responsib |
| AP53_1715 | 0.340165743 | 6.006493938 | 5.142393791 | 0.043670038 | 0.111092455 | Low temperature requirement C pr |
| AP53_749 | -0.597849892 | 8.761878754 | 5.136493338 | 0.043771128 | 0.111110702 | Orotate phosphoribosyltransferase |
| AP53_1525 | -0.4956632 | 9.540654293 | 5.135294706 | 0.043791698 | 0.111110702 | Alanine racemase (EC 5.1.1.1) |
| AP53_329 | -0.649586875 | 7.559046869 | 5.131672805 | 0.043853926 | 0.11112333 | Membrane protein, putative |
| AP53_1114 | -0.268091807 | 7.467225765 | 5.125508464 | 0.043960081 | 0.111215045 | Ribosomal-protein-S5p-alanine ace |
| AP53_361 | 0.406333248 | 12.21068719 | 5.114459066 | 0.044151136 | 0.111215045 | LSU ribosomal protein L1p (L10Ae) |
| AP53_1119 | 0.402284278 | 7.650473294 | 5.11263381 | 0.044182792 | 0.111215045 | Biotin operon repressor / Biotin-pr |

| | | | | | | |
|---|---|---|---|---|---|---|
| AP53_328 | -0.552205374 | 8.781683546 | 5.110724413 | 0.044215938 | 0.111215045 | FIG01115281: hypothetical protei |
| AP53_339 | 0.22430208 | 6.952917524 | 5.110572761 | 0.044218571 | 0.111215045 | Mobile element protein |
| AP53_854 | 0.372273877 | 6.218076921 | 5.109689931 | 0.044233908 | 0.111215045 | hypothetical protein |
| AP53_574 | 0.42154569 | 7.228015948 | 5.084658986 | 0.04467142 | 0.112169762 | Phenylalanyl-tRNA synthetase beta |
| AP53_1574 | -0.316686174 | 6.894513115 | 5.073287495 | 0.044871903 | 0.112527603 | NAD(FAD)-utilizing dehydrogenases |
| AP53_633 | -0.493036681 | 4.286017489 | 5.066930406 | 0.044984454 | 0.112574753 | Single-stranded DNA-binding prote |
| AP53_486 | -0.556772616 | 8.007340446 | 5.065676062 | 0.045006702 | 0.112574753 | Transcriptional regulator |
| AP53_1438 | -0.279338087 | 8.510849309 | 5.045687304 | 0.045363034 | 0.113151293 | Hydrolase (HAD superfamily) |
| AP53_1239 | 0.460213191 | 8.396129883 | 5.045058646 | 0.045374296 | 0.113151293 | Cell division protein YlmG/Ycf19 (p |
| AP53_1467 | -0.291339158 | 9.113828949 | 5.042950594 | 0.045412085 | 0.113151293 | Acetyltransferase (EC 2.3.1.-) |
| AP53_1413 | -0.792486945 | 6.936292989 | 5.036224505 | 0.04553291 | 0.113306897 | FIG01115263: hypothetical protei |
| AP53_1000 | -0.797191305 | 7.176395449 | 5.029734155 | 0.045649869 | 0.113452491 | Paratox |
| AP53_180 | -0.884199598 | 2.720633585 | 5.006512231 | 0.046071315 | 0.114353482 | hypothetical protein |
| AP53_1226 | 0.202347851 | 8.30245655 | 4.995358603 | 0.046275407 | 0.114713364 | Exodeoxyribonuclease VII large sub |
| AP53_173 | -0.412986924 | 6.451482905 | 4.990893008 | 0.046357425 | 0.114770103 | S-layer homology domain / putativ |
| AP53_175 | -0.793298301 | 5.691972231 | 4.983327888 | 0.046496771 | 0.114968449 | hypothetical protein |
| AP53_871 | -0.415955031 | 7.585096391 | 4.95367392 | 0.047047883 | 0.11618313 | GMP reductase (EC 1.7.1.7) |
| AP53_121 | -0.304926485 | 6.021714868 | 4.933968773 | 0.04741846 | 0.116949468 | V-type ATP synthase subunit K (EC 3 |
| AP53_347 | 0.256137343 | 7.310843593 | 4.924798529 | 0.047592118 | 0.117228808 | Glycerol-3-phosphate transporter |
| AP53_1101 | 0.290602073 | 6.992606094 | 4.911544426 | 0.047844471 | 0.117701037 | Ribosomal small subunit pseudouri |
| AP53_545 | 0.464520155 | 12.02450849 | 4.89920011 | 0.048080957 | 0.118046099 | Enolase (EC 4.2.1.11) |
| AP53_1535 | 0.525868789 | 10.61596999 | 4.897877916 | 0.04810637 | 0.118046099 | Transcription termination protein |
| AP53_1137 | -0.364628876 | 5.163380194 | 4.888196027 | 0.048292957 | 0.118218507 | Mobile element protein |
| AP53_991 | -0.411146712 | 11.37384296 | 4.886218883 | 0.048331168 | 0.118218507 | Glucosamine--fructose-6-phosphat |
| AP53_1021 | 1.120575115 | 6.68383023 | 4.884761957 | 0.048359348 | 0.118218507 | DNA packaging protein, phage asso |
| AP53_50 | 0.483703054 | 11.20930675 | 4.872664439 | 0.048594108 | 0.118642974 | LSU ribosomal protein L16p (L10e) |
| AP53_1253 | -0.359702341 | 4.832420676 | 4.867354022 | 0.048697595 | 0.118746271 | Late competence protein ComC, pr |
| AP53_737 | 0.250707827 | 7.671548301 | 4.858079723 | 0.048878967 | 0.118935751 | Hypothetical protein SPy0887 |
| AP53_149 | -0.5631674 | 11.85960658 | 4.857116459 | 0.048897851 | 0.118935751 | Glycine betaine ABC transport syste |
| AP53_1213 | -0.423591066 | 7.298142231 | 4.837916486 | 0.049276111 | 0.119705795 | Phage CI-like repressor |
| AP53_1498 | -0.47895302 | 6.613320693 | 4.815835035 | 0.049715518 | 0.120622275 | hypothetical protein |
| AP53_923 | -0.345169728 | 8.605489428 | 4.803171612 | 0.049969649 | 0.121087501 | Signal recognition particle associat |
| AP53_rna42 | 1.554088092 | 3.837079763 | 4.796596453 | 0.050102219 | 0.121257365 | tRNA-Glu-TTC |
| AP53_1427 | 0.421350248 | 11.35827444 | 4.777228441 | 0.050495199 | 0.122056266 | FIG01116327: hypothetical protei |
| AP53_851 | 0.76242361 | 6.138214621 | 4.751178516 | 0.05102965 | 0.123194715 | Class B acid phosphatase precursor |

| | | | | | | |
|---|---|---|---|---|---|---|
| AP53_rna41 | 1.628155076 | 4.30478976 | 4.740384045 | 0.051253116 | 0.123580496 | tRNA-Ile-GAT |
| AP53_1219 | 0.211068582 | 8.732265628 | 4.719857151 | 0.051681337 | 0.124458405 | Hypothetical protein DUF194, Deg' |
| AP53_1766 | -0.627677514 | 8.876366489 | 4.707396273 | 0.0519434 | 0.124934496 | Acetyltransferase (EC 2.3.1.-) |
| AP53_729 | 0.29938152 | 7.986503413 | 4.685203447 | 0.052414126 | 0.125910666 | Isopentenyl-diphosphate delta-isor |
| AP53_1779 | -0.638222483 | 8.691103742 | 4.664277597 | 0.052862714 | 0.126831308 | Holliday junction DNA helicase Ruv |
| AP53_129 | -0.291733801 | 8.445112756 | 4.654535585 | 0.053073138 | 0.127178964 | putative toxic anion resistance pro |
| AP53_292 | 0.224426738 | 7.715056802 | 4.641275955 | 0.053361173 | 0.127711512 | Ribosomal large subunit pseudouri |
| AP53_1551 | -0.255557433 | 7.86464506 | 4.630156487 | 0.053604177 | 0.128097904 | Recombination inhibitory protein |
| AP53_955 | 0.351917125 | 6.856014716 | 4.62785645 | 0.053654609 | 0.128097904 | Two component system sensor hist |
| AP53_270 | -0.234470627 | 8.047303966 | 4.616499928 | 0.053904462 | 0.128536316 | Helicase loader DnaB |
| AP53_971 | -0.421016517 | 7.836608819 | 4.613482817 | 0.053971078 | 0.128537254 | membrane protein |
| AP53_748 | -0.469507616 | 8.60819477 | 4.592170557 | 0.054444488 | 0.129505822 | Orotidine 5'-phosphate decarboxyl |
| AP53_1208 | -0.761090172 | 2.736754577 | 4.579929893 | 0.054718663 | 0.129918557 | FIG01114174: hypothetical protei |
| AP53_1767 | -0.338739701 | 9.051988918 | 4.578452407 | 0.05475187 | 0.129918557 | Putative oxidoreductase |
| AP53_186 | -0.429382188 | 8.362564137 | 4.564877854 | 0.055058106 | 0.130485695 | Lipid A export ATP-binding/permea |
| AP53_1654 | 0.300900957 | 7.992457834 | 4.551758926 | 0.055356037 | 0.13103179 | DNA polymerase III alpha subunit (E |
| AP53_305 | -0.443059033 | 6.571725935 | 4.544669359 | 0.055517854 | 0.131254756 | Ferrichrome-binding periplasmic p |
| AP53_1769 | -0.310241785 | 9.866497896 | 4.536082057 | 0.055714625 | 0.131433553 | Ribonucleotide reductase of class II |
| AP53_773 | 0.312408733 | 9.386672227 | 4.533888501 | 0.055765023 | 0.131433553 | Adenine phosphoribosyltransferase |
| AP53_417 | -0.433490378 | 7.960783991 | 4.532514337 | 0.055796624 | 0.131433553 | Bactoprenol glucosyl transferase (E |
| AP53_1215 | -0.436607516 | 6.297480204 | 4.525839761 | 0.055950424 | 0.131636088 | Phage integrase |
| AP53_1791 | -0.784739642 | 5.1115042 | 4.518404636 | 0.056122354 | 0.13185044 | FIG01115767: hypothetical protei |
| AP53_1577 | 0.489285971 | 7.414925905 | 4.515353574 | 0.056193092 | 0.13185044 | Uridine phosphorylase (EC 2.4.2.3) |
| AP53_324 | 0.805810461 | 10.21515832 | 4.512282084 | 0.056264413 | 0.13185044 | Lactate 2-monooxygenase (EC 1.13. |
| AP53_1731 | -0.371039291 | 8.042599518 | 4.506059333 | 0.056409244 | 0.13185044 | Hypothetical protein SPy2065 |
| AP53_135 | -0.266466351 | 8.773173766 | 4.504114153 | 0.056454609 | 0.13185044 | Streptococcal NAD glycohydrolase |
| AP53_1173 | 0.282539456 | 9.926256259 | 4.503752891 | 0.05646304 | 0.13185044 | Phosphoglycerate mutase (EC 5.4.2 |
| AP53_903 | 0.406406944 | 4.168785317 | 4.498683285 | 0.056581503 | 0.13185044 | FIG01114041: hypothetical protei |
| AP53_998 | -0.248439393 | 8.269286534 | 4.492903584 | 0.056716926 | 0.13185044 | ABC transporter, ATP-binding prote |
| AP53_839 | 0.291958905 | 9.240711618 | 4.484694246 | 0.056909954 | 0.13185044 | 2-amino-4-hydroxy-6-hydroxymeth |
| AP53_1198 | -0.47881117 | 3.798169381 | 4.482279983 | 0.056966873 | 0.13185044 | FIG040428: Phage-associated prote |
| AP53_1363 | -0.501072747 | 9.67730007 | 4.482053036 | 0.056972227 | 0.13185044 | Chromosome segregation ATPase |
| AP53_377 | -0.309732293 | 4.518264802 | 4.481834672 | 0.056977379 | 0.13185044 | Peptide ABC transporter ATP bindir |
| AP53_1072 | 0.324129987 | 9.306028122 | 4.479888564 | 0.057023321 | 0.13185044 | putative esterase |
| AP53_rna32 | 1.218069989 | 4.160629324 | 4.476516836 | 0.057103024 | 0.13185044 | tRNA-Gly-TCC |

| | | | | | | |
|---|---|---|---|---|---|---|
| AP53_803 | 0.354118461 | 6.424057096 | 4.472486787 | 0.057198466 | 0.13185044 | uncharacterized secreted protein, Y |
| AP53_1479 | 0.466946393 | 10.1080263 | 4.470546448 | 0.057244488 | 0.13185044 | Biotin carboxylase of acetyl-CoA ca |
| AP53_1846 | 0.303246889 | 6.160194467 | 4.467629243 | 0.057313763 | 0.13185044 | FIG009210: peptidase, M16 family |
| AP53_1794 | -0.588610811 | 5.449043881 | 4.467108056 | 0.05732615 | 0.13185044 | FIG01114073: hypothetical protei |
| AP53_1648 | -0.453131358 | 11.11452243 | 4.466854191 | 0.057332185 | 0.13185044 | SSU ribosomal protein S15p (S13e) |
| AP53_1325 | 0.765297393 | 2.984739862 | 4.416639792 | 0.058541196 | 0.134471552 | Phage minor capsid protein #Fam0 |
| AP53_1067 | -0.326737078 | 5.709482512 | 4.408992646 | 0.058728019 | 0.134741235 | Maltose/maltodextrin ABC transpo |
| AP53_796 | 0.443056784 | 10.79042999 | 4.385249475 | 0.059312702 | 0.135922024 | Dihydrolipoamide acetyltransferas |
| AP53_1795 | -0.591290428 | 5.452817499 | 4.381929073 | 0.05939503 | 0.13595018 | Phage protein |
| AP53_1216 | 0.434877308 | 11.02528837 | 4.347775816 | 0.060249946 | 0.137713069 | DNA-binding protein HBsu |
| AP53_197 | -0.275839254 | 9.919655201 | 4.345513023 | 0.060307114 | 0.137713069 | Inner membrane protein translocas |
| AP53_1534 | -0.371397731 | 8.554004643 | 4.330233176 | 0.060694876 | 0.138435669 | Sucrose operon repressor ScrR, Lac |
| AP53_586 | 0.258781043 | 10.17786669 | 4.325552142 | 0.060814273 | 0.138545193 | RNA polymerase sigma factor RpoD |
| AP53_1555 | -0.243804805 | 8.924918181 | 4.318008826 | 0.061007279 | 0.138821956 | Putative signal peptidase I (EC 3.4.2 |
| AP53_996 | 0.341463509 | 8.831870744 | 4.312868002 | 0.06113924 | 0.138959326 | Transcriptional regulator, GntR fam |
| AP53_187 | -0.446853281 | 7.024537248 | 4.288436072 | 0.061771149 | 0.140231345 | FIG053235: Diacylglucosamine hyd |
| AP53_712 | 1.346809121 | 1.394023947 | 4.269804302 | 0.06225838 | 0.141172332 | hypothetical protein |
| AP53_1139 | -0.244891643 | 6.556102348 | 4.26600259 | 0.06235837 | 0.141234068 | FIG01114175: hypothetical protei |
| AP53_1154 | 0.433363228 | 6.87505983 | 4.249785038 | 0.062787111 | 0.142039373 | FIG01116792: hypothetical protei |
| AP53_1477 | 0.536080284 | 10.2050706 | 4.244056919 | 0.0629394 | 0.14212098 | Acetyl-coenzyme A carboxyl transfe |
| AP53_676 | -0.493310817 | 8.098908033 | 4.242922036 | 0.062969625 | 0.14212098 | Histone acetyltransferase Gcn5 |
| AP53_662 | 0.326710292 | 8.236886967 | 4.238120377 | 0.063097704 | 0.14224465 | Pore forming protein ebsA |
| AP53_1281 | -0.326222615 | 9.007721286 | 4.234209806 | 0.063202248 | 0.142315039 | hypothetical protein |
| AP53_1469 | -0.748445954 | 8.997062334 | 4.223406797 | 0.063492149 | 0.142802156 | Guanine-hypoxanthine permease |
| AP53_145 | 0.36001937 | 5.127435458 | 4.217578133 | 0.063649233 | 0.14281999 | L-xylulose 5-phosphate 3-epimeras |
| AP53_144 | 0.383130017 | 5.193900191 | 4.217324371 | 0.063656083 | 0.14281999 | 3-keto-L-gulonate 6-phosphate dec |
| AP53_1286 | -0.478591308 | 9.381978509 | 4.214927693 | 0.06372082 | 0.14281999 | 3-dehydroquinate synthase (EC 4.2. |
| AP53_1290 | 0.224218824 | 6.889385727 | 4.200521786 | 0.064111631 | 0.143530191 | Shikimate/quinate 5-dehydrogenas |
| AP53_933 | 0.470087721 | 9.329273809 | 4.181907184 | 0.064620944 | 0.144503747 | Anaerobic ribonucleoside-triphosp |
| AP53_468 | -0.248107716 | 8.330445525 | 4.177890323 | 0.064731494 | 0.144584384 | possible permease |
| AP53_503 | -0.248841443 | 7.641910473 | 4.160778585 | 0.06520502 | 0.145428727 | Deoxyguanosinetriphosphate triph |
| AP53_1362 | -0.448432256 | 9.411929701 | 4.158824513 | 0.065259362 | 0.145428727 | Phage protein |
| AP53_1131 | -0.232984798 | 8.424826186 | 4.127721224 | 0.066131798 | 0.147143125 | Glutaredoxin-like protein NrdH, red |
| AP53_929 | -0.2751415 | 6.614404055 | 4.126007008 | 0.066180293 | 0.147143125 | ABC transporter ATP-binding protei |
| AP53_1035 | -0.645923301 | 4.328280742 | 4.117372368 | 0.066425223 | 0.147518716 | FIG01118889: hypothetical protei |

| | | | | | | |
|---|---|---|---|---|---|---|
| AP53_435 | 0.337009711 | 7.890446052 | 4.112870167 | 0.06655337 | 0.147576557 | Zn-dependent hydrolase YycJ/WalJ, |
| AP53_877 | 0.197628005 | 8.076302094 | 4.111117785 | 0.06660333 | 0.147576557 | Peptide chain release factor 1 |
| AP53_893 | -0.406093754 | 10.31994222 | 4.098458754 | 0.066965595 | 0.148210057 | Transcription regulator [contains d |
| AP53_294 | -0.257639368 | 7.702865212 | 4.087051423 | 0.067294097 | 0.148767474 | tRNA (cytidine(34)-2'-O)-methyltra |
| AP53_1505 | 0.291840365 | 10.10113094 | 4.074716768 | 0.067651515 | 0.149387475 | GTP-sensing transcriptional pleiotr |
| AP53_1201 | 0.543409634 | 3.887389286 | 4.022379734 | 0.069194043 | 0.152479468 | conserved hypothetical protein - pl |
| AP53_124 | -0.372493741 | 5.968741084 | 4.021883664 | 0.069208867 | 0.152479468 | V-type ATP synthase subunit F (EC 3 |
| AP53_1483 | 0.427841915 | 9.419940924 | 4.016092835 | 0.069382207 | 0.152688054 | 3-oxoacyl-[acyl-carrier protein] red |
| AP53_1777 | -0.334426981 | 7.689044846 | 4.012491596 | 0.069490271 | 0.15275268 | Molybdopterin binding motif, CinA |
| AP53_1456 | -0.239825211 | 9.922923369 | 3.998629279 | 0.069908166 | 0.153497456 | COG2740: Predicted nucleic-acid-b |
| AP53_1677 | -0.256616876 | 7.899458382 | 3.995341 | 0.070007744 | 0.153542409 | Para-aminobenzoate synthase, ami |
| AP53_411 | -0.306173851 | 10.31818213 | 3.989735664 | 0.070177888 | 0.153741853 | 3'-to-5' exoribonuclease RNase R |
| AP53_1778 | -0.449101475 | 9.281653972 | 3.976832334 | 0.070571476 | 0.154429802 | DNA-3-methyladenine glycosylase ( |
| AP53_14 | 0.287095743 | 10.75548072 | 3.972732788 | 0.070697087 | 0.154530456 | Cell division protein FtsH (EC 3.4.2 |
| AP53_896 | -0.390449797 | 9.148982694 | 3.967958238 | 0.070843724 | 0.154676792 | 50S ribosomal subunit maturation |
| AP53_rna18 | 1.004727048 | 4.238330892 | 3.963355391 | 0.070985439 | 0.154812064 | tRNA-Val-TAC |
| AP53_1591 | 0.245369986 | 10.34080977 | 3.95456416 | 0.071257071 | 0.155140508 | Dihydroxyacetone kinase family pro |
| AP53_556 | 0.280950376 | 9.196164744 | 3.953310956 | 0.071295895 | 0.155140508 | Substrate-specific component Que |
| AP53_37 | -0.571830608 | 8.621551496 | 3.945921782 | 0.07152534 | 0.155465494 | MORN motif family protein |
| AP53_1330 | -0.648198552 | 3.811752405 | 3.926729792 | 0.072125504 | 0.156477816 | Phage protein |
| AP53_770 | -0.265467368 | 8.138427926 | 3.92587695 | 0.072152316 | 0.156477816 | Ribonuclease Z (EC 3.1.26.11) |
| AP53_1687 | 0.224457571 | 6.531062586 | 3.913094519 | 0.072555638 | 0.15717689 | Streptococcal histidine triad protei |
| AP53_1387 | -0.530119098 | 7.519770716 | 3.908168425 | 0.072711805 | 0.15733959 | Butyrate-acetoacetate CoA-transfer |
| AP53_79 | 0.286802938 | 8.410774121 | 3.896633019 | 0.073079108 | 0.157958294 | Multimodular transpeptidase-trans |
| AP53_953 | 0.404576307 | 5.365556629 | 3.884606352 | 0.073464469 | 0.15861461 | Pantothenate kinase (EC 2.7.1.33) |
| AP53_665 | -0.233725194 | 8.344185197 | 3.880114414 | 0.073609037 | 0.158750157 | Cytidylate kinase (EC 2.7.4.14) |
| AP53_707 | -0.292976941 | 8.129301303 | 3.867503825 | 0.074016757 | 0.159452302 | 1-phosphofructokinase (EC 2.7.1.5 |
| AP53_787 | 0.261448485 | 5.790644875 | 3.858952132 | 0.074294816 | 0.159655329 | ABC transporter substrate-binding |
| AP53_pseudo1695 | -0.50550646 | 11.9771486 | 3.858793176 | 0.074299997 | 0.159655329 | Antiphagocytic M protein/pseudog |
| AP53_885 | -0.272277829 | 7.980501176 | 3.857021637 | 0.074357763 | 0.159655329 | ABC transporter, ATP-binding/perm |
| AP53_1275 | -0.327047634 | 6.163761658 | 3.823403475 | 0.075464458 | 0.1618525 | Thiol:disulfide oxidoreductase asso |
| AP53_1661 | -0.363108077 | 5.406063771 | 3.816217225 | 0.075703632 | 0.162181772 | Pullulanase (EC 3.2.1.41) |
| AP53_706 | 0.474887945 | 8.158050813 | 3.813775954 | 0.075785094 | 0.162181772 | Transcriptional repressor of the fru |
| AP53_1344 | -0.799509482 | 4.348525858 | 3.810011514 | 0.075910918 | 0.16220349 | FIG01115146: hypothetical protei |
| AP53_1642 | 0.299998633 | 8.132237723 | 3.80664274 | 0.076023734 | 0.16220349 | FIG01114029: hypothetical protei |

| | | | | | | |
|---|---|---|---|---|---|---|
| AP53_1780 | -0.43787619 | 8.841926028 | 3.805980253 | 0.076045943 | 0.16220349 | Multidrug resistance protein B |
| AP53_1065 | 0.374372106 | 5.632716854 | 3.795786482 | 0.076388693 | 0.162755712 | Maltose operon transcriptional rep |
| AP53_683 | -0.312348485 | 7.300666495 | 3.783961513 | 0.076788659 | 0.163303309 | Cysteine and methionine metabolis |
| AP53_49 | 0.452545724 | 11.30618214 | 3.783215661 | 0.076813973 | 0.163303309 | SSU ribosomal protein S3p (S3e) |
| AP53_1276 | -0.386884066 | 5.528557162 | 3.771652715 | 0.077207715 | 0.163846739 | Cytochrome c-type biogenesis prot |
| AP53_1571 | -0.393864133 | 6.358480657 | 3.769400647 | 0.07728469 | 0.163846739 | Transcriptional regulator, MerR fan |
| AP53_230 | -0.465315298 | 5.128972314 | 3.767597633 | 0.077346384 | 0.163846739 | hypothetical cytosolic protein |
| AP53_198 | -0.29081311 | 10.34227161 | 3.76471596 | 0.077445112 | 0.163846739 | RNA-binding protein Jag |
| AP53_181 | -0.43986789 | 6.519016854 | 3.762447468 | 0.07752294 | 0.163846739 | Integral membrane protein (Rhomb |
| AP53_1252 | 0.271622091 | 12.92724852 | 3.760900438 | 0.077576071 | 0.163846739 | Non-specific DNA-binding protein D |
| AP53_1333 | 0.535381798 | 3.062741465 | 3.75473569 | 0.077788235 | 0.164116266 | Phage minor capsid protein #Fam0( |
| AP53_6 | -0.38863212 | 5.955464852 | 3.752054172 | 0.077880743 | 0.164133032 | Peptidyl-tRNA hydrolase (EC 3.1.1.2 |
| AP53_567 | 0.300335101 | 7.522620469 | 3.745031043 | 0.078123668 | 0.164466421 | FIG01114865: hypothetical protei |
| AP53_1160 | -0.315033516 | 7.302703675 | 3.738015281 | 0.078367266 | 0.164800501 | tRNA (adenine37-N(6))-methyltrans |
| AP53_769 | -0.295745351 | 8.463737777 | 3.716766567 | 0.079110746 | 0.166183937 | Cystathionine beta-lyase (EC 4.4.1.8 |
| AP53_1820 | -0.540400791 | 6.814986037 | 3.714149479 | 0.079202912 | 0.166197679 | FIG01114548: hypothetical protei |
| AP53_438 | -0.382680112 | 5.192022396 | 3.705771314 | 0.079498854 | 0.166638526 | Positive transcriptional regulator, I |
| AP53_1210 | -1.0421452 | 1.936449278 | 3.693091942 | 0.079949307 | 0.167401948 | Phage protein |
| AP53_496 | 0.339665808 | 12.88864028 | 3.686013912 | 0.080202125 | 0.167689092 | Translation elongation factor Tu |
| AP53_1103 | -0.344088649 | 7.380225059 | 3.684418929 | 0.08025923 | 0.167689092 | Transporter, MFS superfamily |
| AP53_96 | -0.453569421 | 6.056549802 | 3.666969002 | 0.080887262 | 0.168819544 | FIG009688: Thioredoxin |
| AP53_244 | 0.478613994 | 3.470264022 | 3.661511343 | 0.081084922 | 0.169050304 | Competence-specific sigma factor C |
| AP53_rna85 | 0.566255599 | 4.683211769 | 3.653931518 | 0.081360422 | 0.169354618 | tRNA-Arg-TCT |
| AP53_897 | -0.31939886 | 8.720799367 | 3.652697363 | 0.081405388 | 0.169354618 | Ribonuclease HII (EC 3.1.26.4) |
| AP53_253 | -0.219016904 | 7.304285585 | 3.639964088 | 0.081871103 | 0.170141124 | FIG007079: UPF0348 protein famil |
| AP53_1494 | -0.211602015 | 9.042313129 | 3.631005314 | 0.082200725 | 0.17064343 | putative N-acetyl-muramidase |
| AP53_150 | -0.589299443 | 12.31151895 | 3.614089003 | 0.08282758 | 0.171704324 | Glycine betaine ABC transport syste |
| AP53_1431 | -0.201480584 | 9.994209097 | 3.612448246 | 0.082888692 | 0.171704324 | INTEGRAL MEMBRANE PROTEIN (Rh |
| AP53_1396 | 0.510301022 | 4.564997894 | 3.609326599 | 0.083005114 | 0.171762182 | FIG005686: hypothetical protein |
| AP53_857 | -0.254865803 | 8.139744154 | 3.602204646 | 0.08327148 | 0.172129863 | Glutamine amidotransferase, class I |
| AP53_1251 | -0.456845578 | 6.28830736 | 3.587164215 | 0.083837456 | 0.173115427 | FIG01114576: hypothetical protei |
| AP53_1062 | -0.2422801 | 7.882999782 | 3.582320842 | 0.084020717 | 0.173309471 | lipoprotein, putative |
| AP53_824 | 0.335283979 | 5.45028148 | 3.573901025 | 0.084340475 | 0.173784355 | Two-component response regulato |
| AP53_1832 | 0.187005541 | 8.819021249 | 3.571437413 | 0.084434318 | 0.173793225 | Replicative DNA helicase (DnaB) (EC |
| AP53_786 | 0.560529489 | 3.925446239 | 3.566050781 | 0.084639949 | 0.17403193 | hypothetical protein |

| | | | | | | |
|---|---|---|---|---|---|---|
| AP53_887 | 0.346055074 | 10.0984361 | 3.563415376 | 0.084740778 | 0.174054869 | L-lactate dehydrogenase (EC 1.1.1.2 |
| AP53_1541 | -0.254584457 | 7.953481663 | 3.556044797 | 0.085023556 | 0.174209861 | Magnesium and cobalt transport pr |
| AP53_rna34 | 1.029017349 | 3.37489484 | 3.554712979 | 0.085074776 | 0.174209861 | tRNA-Pseudo-GCT |
| AP53_652 | 0.376565356 | 8.254094096 | 3.554434354 | 0.085085496 | 0.174209861 | Phage transcriptional regulator, Cr |
| AP53_1077 | 0.467488753 | 6.908046039 | 3.546351691 | 0.085397206 | 0.174560515 | FIG01114114: hypothetical protei |
| AP53_1759 | -0.343397032 | 6.652835082 | 3.543201168 | 0.085519086 | 0.174560515 | Trehalose operon transcriptional re |
| AP53_76 | 0.242775181 | 7.413343077 | 3.543008203 | 0.085526558 | 0.174560515 | Zinc ABC transporter, inner membr |
| AP53_679 | 0.257800444 | 7.309242498 | 3.535310056 | 0.085825299 | 0.174986245 | Capsule biosynthesis protein capA |
| AP53_1214 | -0.365658936 | 7.586571696 | 3.526754638 | 0.086158813 | 0.175481906 | Phage protein |
| AP53_1230 | -0.443365598 | 7.713718669 | 3.523321959 | 0.086293076 | 0.175571133 | Glutamate transport ATP-binding p |
| AP53_pseudo268 | 0.219274537 | 7.889228161 | 3.52097407 | 0.086385058 | 0.175574238 | Transmembrane histidine kinase Cs |
| AP53_155 | 0.411526511 | 9.707554236 | 3.516138177 | 0.08657489 | 0.175776005 | Co-activator of prophage gene expr |
| AP53_1126 | 0.232826357 | 8.692747408 | 3.497573973 | 0.087308402 | 0.177069159 | ATP-dependent RNA helicase YfmL |
| AP53_1819 | -0.435294856 | 6.604633592 | 3.495413626 | 0.087394258 | 0.177069159 | FtsK/SpoIIIE family |
| AP53_101 | 0.330211435 | 8.357710187 | 3.490496055 | 0.087590077 | 0.177280854 | Chaperonin (heat shock protein 33) |
| AP53_1318 | 0.582778224 | 4.204982717 | 3.487855471 | 0.087695448 | 0.177296772 | Phage endopeptidase |
| AP53_962 | 0.257277773 | 7.398175823 | 3.484302673 | 0.087837467 | 0.177296772 | Phosphate transport system perme |
| AP53_1153 | 0.22654406 | 7.176369711 | 3.483440585 | 0.08787197 | 0.177296772 | FIG01114764: hypothetical protei |
| AP53_199 | 0.294764975 | 6.130953473 | 3.478101692 | 0.088086022 | 0.177441733 | FIG01115631: hypothetical protei |
| AP53_310 | 0.281263526 | 6.753458318 | 3.475460711 | 0.088192143 | 0.177441733 | hypothetical protein |
| AP53_758 | -0.286764431 | 7.399991744 | 3.473746198 | 0.088261121 | 0.177441733 | Branched-chain amino acid aminot |
| AP53_139 | -0.428765028 | 6.850934411 | 3.470471091 | 0.088393069 | 0.177441733 | Cystathionine beta-lyase (EC 4.4.1.8 |
| AP53_658 | -0.392477916 | 6.883404077 | 3.468550379 | 0.088470563 | 0.177441733 | Membrane protein |
| AP53_1386 | -0.47344187 | 7.247876238 | 3.468011458 | 0.088492322 | 0.177441733 | 3-ketoacyl-CoA thiolase (EC 2.3.1.1 |
| AP53_1461 | 0.188719183 | 9.01395109 | 3.4603945 | 0.088800559 | 0.177697908 | aminoglycoside phosphotransferas |
| AP53_1222 | -0.287495232 | 7.686486073 | 3.458013722 | 0.088897174 | 0.177697908 | Arginine pathway regulatory protei |
| AP53_675 | -0.416168227 | 8.237706878 | 3.457409499 | 0.088921714 | 0.177697908 | Dihydrofolate synthase (EC 6.3.2.12 |
| AP53_1466 | -0.314959972 | 9.729073719 | 3.455820814 | 0.088986278 | 0.177697908 | Cell envelope-associated transcript |
| AP53_330 | -0.406744353 | 10.15185276 | 3.452842909 | 0.089107456 | 0.177757011 | Methionyl-tRNA synthetase (EC 6.1. |
| AP53_429 | 0.623154929 | 2.085177102 | 3.438823609 | 0.089680669 | 0.178716815 | Hypothetical protein |
| AP53_1867 | -0.362942532 | 7.390902448 | 3.434858871 | 0.089843599 | 0.178799988 | small RNA fasX |
| AP53_1313 | 0.772320211 | 3.823947758 | 3.433327202 | 0.08990664 | 0.178799988 | Phage holin |
| AP53_580 | -0.215956957 | 7.556711597 | 3.426059532 | 0.09020651 | 0.179040798 | ATP-dependent nuclease, subunit B |
| AP53_547 | 0.388922285 | 7.169960578 | 3.42590924 | 0.090212724 | 0.179040798 | Streptolysin S biosynthesis protein |
| AP53_354 | 0.296613384 | 9.752665909 | 3.423692234 | 0.090304452 | 0.179040798 | Manganese ABC transporter, peripla |

| | | | | | | |
|---|---|---|---|---|---|---|
| AP53_1415 | -0.243076593 | 8.159083617 | 3.417681163 | 0.090553737 | 0.179331668 | Glutamate 5-kinase (EC 2.7.2.11) |
| AP53_1149 | -0.361958097 | 11.11120345 | 3.415703363 | 0.090635943 | 0.179331668 | Glucosamine-6-phosphate deamina |
| AP53_2 | -0.248092583 | 8.274963851 | 3.406121269 | 0.091035523 | 0.179905636 | DNA polymerase III beta subunit (EC |
| AP53_1071 | 0.387618491 | 7.377320043 | 3.404307398 | 0.091111407 | 0.179905636 | Transposase (maltose area) |
| AP53_864 | 0.222712505 | 7.658762239 | 3.392518715 | 0.091606491 | 0.180620048 | NAD kinase (EC 2.7.1.23) |
| AP53_191 | -0.247168603 | 8.470630369 | 3.391265329 | 0.091659324 | 0.180620048 | DUF124 domain-containing protei |
| AP53_772 | -0.350080969 | 8.155570193 | 3.381578275 | 0.09206892 | 0.181243179 | Single-stranded-DNA-specific exonu |
| AP53_1641 | 0.267145578 | 8.263963792 | 3.374743581 | 0.092359268 | 0.181630537 | Serine acetyltransferase (EC 2.3.1.3 |
| AP53_1441 | -0.189386462 | 8.92053845 | 3.360579415 | 0.092964584 | 0.182463858 | Tagatose 1,6-diphosphate aldolase |
| AP53_1327 | 0.579446899 | 3.739674086 | 3.360257722 | 0.092978388 | 0.182463858 | Phage protein |
| AP53_rna68 | 0.615901533 | 4.055443708 | 3.35709531 | 0.093114228 | 0.182463858 | tRNA-Val-TAC |
| AP53_rna55 | 2.333848592 | 0.474471177 | 3.355566308 | 0.093179993 | 0.182463858 | tRNA-Asn-GTT |
| AP53_1690 | -0.584132262 | 11.27051386 | 3.353869606 | 0.093253038 | 0.182463858 | Fibronectin-binding protein |
| AP53_1818 | -0.340348618 | 5.452001863 | 3.339149266 | 0.093889742 | 0.183382288 | FIG01115119: hypothetical protei |
| AP53_445 | 0.588547107 | 3.478242748 | 3.338110409 | 0.093934878 | 0.183382288 | Dolichyl-phosphate mannoosyltran |
| AP53_1691 | -0.448627687 | 10.28433427 | 3.332565182 | 0.09417626 | 0.183382288 | C5a peptidase (EC 3.4.21.-) |
| AP53_1807 | -0.278325703 | 9.245643543 | 3.332429071 | 0.094182194 | 0.183382288 | Arginyl-tRNA synthetase (EC 6.1.1.1 |
| AP53_1247 | 0.237170278 | 8.304777516 | 3.332139575 | 0.094194818 | 0.183382288 | FIG01114439: hypothetical protei |
| AP53_1725 | -0.499222009 | 6.193449723 | 3.326890816 | 0.094424055 | 0.183511904 | FIG01114281: hypothetical protei |
| AP53_46 | 0.44897214 | 11.29190986 | 3.326286663 | 0.094450485 | 0.183511904 | LSU ribosomal protein L2p (L8e) |
| AP53_836 | 0.222885497 | 9.379810407 | 3.314655866 | 0.094961089 | 0.184319473 | GTP cyclohydrolase I (EC 3.5.4.16) t |
| AP53_1270 | 0.403856297 | 8.361294977 | 3.298295818 | 0.095685089 | 0.185478052 | Putative uncharacterized protein gl |
| AP53_1465 | 0.171077746 | 8.952617724 | 3.296856668 | 0.095749103 | 0.185478052 | Hypothetical protein in cluster wit |
| AP53_645 | -0.555172676 | 4.180054042 | 3.289410015 | 0.096081178 | 0.185882346 | hypothetical protein |
| AP53_745 | -0.54158578 | 5.133878848 | 3.287885366 | 0.096149343 | 0.185882346 | hypothetical protein |
| AP53_943 | -0.258319997 | 7.179684337 | 3.279535896 | 0.096523698 | 0.186420397 | Phosphopantothenoylcysteine syn |
| AP53_1255 | -0.27257073 | 8.539361175 | 3.271315194 | 0.09689404 | 0.186949634 | Hypothetical DUF1027 domain pro |
| AP53_746 | -0.311553227 | 8.06732286 | 3.26520612 | 0.09717039 | 0.187195373 | Methionine biosynthesis and trans |
| AP53_875 | -0.508291025 | 7.024084202 | 3.263075718 | 0.09726699 | 0.187195373 | 4-oxalocrotonate tautomerase (EC |
| AP53_881 | -0.268528639 | 9.178826799 | 3.26211188 | 0.097310732 | 0.187195373 | Serine hydroxymethyltransferase (E |
| AP53_130 | -0.21881956 | 7.733056354 | 3.259375058 | 0.097435072 | 0.187230455 | FIG013576: hypothetical protein |
| AP53_1847 | 0.341201356 | 5.744415718 | 3.253389065 | 0.097707714 | 0.187230455 | FIG001621: Zinc protease |
| AP53_1048 | 0.455775945 | 6.144025653 | 3.253286217 | 0.097712407 | 0.187230455 | hypothetical protein, phage associa |
| AP53_57 | 0.387387539 | 10.59405612 | 3.253233502 | 0.097714812 | 0.187230455 | SSU ribosomal protein S8p (S15Ae) |
| AP53_755 | -0.21956997 | 8.12430739 | 3.250339465 | 0.097846976 | 0.187298797 | Acyl-phosphate:glycerol-3-phospha |

| | | | | | | |
|---|---|---|---|---|---|---|
| AP53_583 | 0.669913025 | 11.23882585 | 3.24164145 | 0.098245524 | 0.187876416 | SSU ribosomal protein S21p |
| AP53_771 | -0.29430975 | 9.030765767 | 3.229307953 | 0.098814098 | 0.188456308 | Short chain dehydrogenase |
| AP53_477 | -0.216708336 | 8.971203091 | 3.225810681 | 0.098976062 | 0.188456308 | FIG01114801: hypothetical protei |
| AP53_1274 | -0.40587197 | 6.536350653 | 3.222909456 | 0.099110671 | 0.188456308 | Peptide methionine sulfoxide redu |
| AP53_988 | -0.405838661 | 9.577451512 | 3.220688068 | 0.09921389 | 0.188456308 | ABC-type polar amino acid transpo |
| AP53_1365 | -0.44162866 | 9.690534392 | 3.217923764 | 0.099342522 | 0.188456308 | Regulatory protein recX |
| AP53_637 | -0.330106771 | 4.800871695 | 3.215763037 | 0.099443212 | 0.188456308 | hypothetical phage protein |
| AP53_521 | -0.231386614 | 8.920122241 | 3.21521895 | 0.099468586 | 0.188456308 | Cell division protein FtsX |
| AP53_1010 | 0.512262402 | 5.822470884 | 3.215020746 | 0.099477832 | 0.188456308 | Phage hyaluronidase |
| AP53_575 | 0.259350245 | 5.702429986 | 3.214413123 | 0.099506182 | 0.188456308 | Salt-stress induced protein |
| AP53_1855 | -0.330432436 | 11.18573207 | 3.214123708 | 0.099519689 | 0.188456308 | Inosine-5'-monophosphate dehydr |
| AP53_444 | 0.48626695 | 3.147686211 | 3.210537105 | 0.099687261 | 0.188589644 | FIG01116163: hypothetical protei |
| AP53_1113 | -0.195470051 | 7.915725624 | 3.205125353 | 0.09994077 | 0.188885136 | FIG009439: Cytosolic protein cont |
| AP53_502 | -0.274027556 | 6.893721111 | 3.188589521 | 0.100720331 | 0.19017331 | FIG01114293: hypothetical protei |
| AP53_215 | -0.236840563 | 9.515396733 | 3.179648732 | 0.101144967 | 0.190789487 | DNA recombination protein RmuC |
| AP53_1262 | 0.650999232 | 10.79217666 | 3.16485044 | 0.101852681 | 0.191729872 | Carbamate kinase (EC 2.7.2.2) |
| AP53_1741 | 0.689985025 | 10.54648531 | 3.163641507 | 0.101910767 | 0.191729872 | Cold shock protein CspA |
| AP53_525 | 0.2118455 | 6.856350395 | 3.163036761 | 0.101939839 | 0.191729872 | DinG family ATP-dependent helicas |
| AP53_713 | 0.439873533 | 2.030550548 | 3.14823843 | 0.102654451 | 0.192887019 | Immunoglobulin G-endopeptidase |
| AP53_343 | -0.196292132 | 6.997685949 | 3.145625713 | 0.102781263 | 0.192938522 | FIG01115464: hypothetical protei |
| AP53_360 | -0.305939569 | 11.50774728 | 3.129959343 | 0.103545735 | 0.194082804 | LSU ribosomal protein L11p (L12e) |
| AP53_256 | -0.242222874 | 8.209398764 | 3.127226889 | 0.103679791 | 0.194082804 | Methionine ABC transporter substr |
| AP53_727 | 0.267193113 | 7.583116513 | 3.127002431 | 0.103690813 | 0.194082804 | Diphosphomevalonate decarboxyla |
| AP53_950 | 0.371044965 | 10.0211407 | 3.120336262 | 0.104018806 | 0.194387272 | Nucleoside-binding protein |
| AP53_1866 | -0.45839259 | 4.323233914 | 3.119627099 | 0.104053774 | 0.194387272 | paratox? |
| AP53_639 | -0.357369481 | 4.770258326 | 3.116974299 | 0.104184711 | 0.194444735 | conserved hypothetical protein - pl |
| AP53_918 | -0.42941748 | 9.936688656 | 3.113581323 | 0.10435248 | 0.194570762 | [Citrate [pro-3S]-lyase] ligase (EC 6.2 |
| AP53_539 | -0.40564307 | 8.780216336 | 3.105342406 | 0.104761255 | 0.195145485 | Chloride channel protein |
| AP53_297 | -0.266448848 | 7.089865324 | 3.093704802 | 0.105342045 | 0.196039223 | COG1242: Predicted Fe-S oxidored |
| AP53_1060 | -0.439637167 | 5.333577603 | 3.078316396 | 0.106116167 | 0.197290691 | hypothetical protein |
| AP53_1271 | -0.325054352 | 7.72486972 | 3.06818698 | 0.106629587 | 0.198055529 | Hypothetical protein SpyM3_1200 |
| AP53_898 | -0.595625532 | 6.544989312 | 3.064185726 | 0.106833243 | 0.198133339 | Rossmann fold nucleotide-binding |
| AP53_1692 | -0.273809021 | 11.16790356 | 3.063354058 | 0.106875634 | 0.198133339 | Antiphagocytic M protein |
| AP53_159 | -0.367250803 | 5.702756779 | 3.058668649 | 0.107114844 | 0.198387321 | Mobile element protein |
| AP53_1763 | -0.386138647 | 7.269390876 | 3.050770364 | 0.107519592 | 0.198947119 | Glyoxalase family protein |

| | | | | | | |
|---|---|---|---|---|---|---|
| AP53_718 | 0.16918417 | 7.903787725 | 3.032114251 | 0.108483189 | 0.200538923 | tRNA nucleotidyltransferase (EC 2.7 |
| AP53_1171 | -0.326321851 | 6.530184668 | 3.022950221 | 0.108960443 | 0.201229514 | hypothetical protein |
| AP53_1473 | 0.413680756 | 9.677362925 | 3.019755648 | 0.109127426 | 0.201346324 | PTS system, mannose-specific IID co |
| AP53_570 | -0.363219315 | 9.326678034 | 3.010693341 | 0.109602853 | 0.201896213 | DNA-entry nuclease (Competence-s |
| AP53_476 | 0.324780467 | 9.659574946 | 3.010110997 | 0.109633492 | 0.201896213 | FIG01115301: hypothetical protei |
| AP53_434 | 0.182601412 | 8.537490947 | 3.007218964 | 0.109785809 | 0.201985076 | Two-component sensor kinase SA14 |
| AP53_536 | 0.194863096 | 8.741025202 | 2.993493087 | 0.110512315 | 0.202979101 | FIG146085: 3'-to-5' oligoribonucle |
| AP53_630 | 0.388014657 | 4.836682553 | 2.993061709 | 0.110535244 | 0.202979101 | B-cell receptor associated protein-r |
| AP53_469 | 0.35055635 | 9.808365257 | 2.969968099 | 0.111771441 | 0.205055168 | FIG01116444: hypothetical protei |
| AP53_299 | 0.174971372 | 8.536673103 | 2.954867941 | 0.112589056 | 0.205902522 | Hemolysins and related proteins co |
| AP53_797 | 0.379442211 | 11.03110948 | 2.953386944 | 0.112669646 | 0.205902522 | Dihydrolipoamide dehydrogenase o |
| AP53_53 | 0.443146859 | 11.43545424 | 2.953063369 | 0.112687263 | 0.205902522 | LSU ribosomal protein L14p (L23e) |
| AP53_169 | -0.212825171 | 9.092234106 | 2.951160816 | 0.112790919 | 0.205902522 | Zinc finger protein |
| AP53_720 | -0.618737388 | 6.455922938 | 2.949951146 | 0.112856886 | 0.205902522 | FIG01114610: hypothetical protei |
| AP53_880 | -0.17907812 | 8.829390239 | 2.949714429 | 0.112869801 | 0.205902522 | Phosphinothricin N-acetyltransfera |
| AP53_1287 | 0.241946011 | 8.933424055 | 2.942036416 | 0.113289689 | 0.206474448 | Uncharacterized conserved protein |
| AP53_pseudo1337 | 0.593367514 | 2.222517796 | 2.931923259 | 0.113845721 | 0.207103231 | DNA modification methylase |
| AP53_1256 | 0.252692617 | 7.810463382 | 2.930595472 | 0.113918976 | 0.207103231 | FIG01114963: hypothetical protei |
| AP53_1111 | 0.20631698 | 8.445947905 | 2.92994669 | 0.113954792 | 0.207103231 | Inner membrane protein YihY, form |
| AP53_217 | -0.231236158 | 8.032206253 | 2.927090534 | 0.114112629 | 0.207196083 | PurR: transcription regulator assoc |
| AP53_1334 | 0.517863197 | 3.009464722 | 2.922347444 | 0.114375345 | 0.207479013 | Phage terminase large subunit |
| AP53_279 | -0.242152994 | 10.25874936 | 2.914789667 | 0.114795522 | 0.208009902 | Inner membrane protein translocas |
| AP53_1332 | 0.297423125 | 4.927605505 | 2.913232427 | 0.114882336 | 0.208009902 | Phage minor capsid protein |
| AP53_510 | -0.254496161 | 5.849857875 | 2.889081492 | 0.116239253 | 0.210270633 | PTS system, N-acetylgalactosamine- |
| AP53_479 | 0.194966707 | 8.07023155 | 2.88334485 | 0.116564497 | 0.210662652 | peptidase, U32 family large subunit |
| AP53_147 | -0.203128599 | 5.637764626 | 2.866258608 | 0.117539963 | 0.212227971 | Transcriptional antiterminator with |
| AP53_529 | 0.201769356 | 9.140066361 | 2.842523804 | 0.118911964 | 0.21450569 | FIG002813: LPPG:FO 2-phospho-L- |
| AP53_873 | -0.473121118 | 8.200934288 | 2.839223482 | 0.11910432 | 0.21465319 | Xanthine permease |
| AP53_123 | -0.243694186 | 5.959595513 | 2.829148933 | 0.119693914 | 0.215515665 | V-type ATP synthase subunit C (EC 3 |
| AP53_1372 | 0.326606648 | 9.148815649 | 2.813353839 | 0.120625649 | 0.216830271 | General stress protein 13 |
| AP53_1793 | -0.547132447 | 5.339455552 | 2.812008973 | 0.120705399 | 0.216830271 | FIG01114419: hypothetical protei |
| AP53_373 | 0.243291938 | 9.046666375 | 2.811103062 | 0.120759157 | 0.216830271 | GTP-binding protein Era |
| AP53_77 | -0.34894723 | 6.050714383 | 2.800729315 | 0.12137688 | 0.217738007 | HIT family protein |
| AP53_1858 | -0.313777599 | 7.645699048 | 2.797099805 | 0.121593937 | 0.217925977 | hypothetical protein BH3604 |
| AP53_579 | 0.235397165 | 7.606143427 | 2.789683523 | 0.122038969 | 0.218521807 | YpfJ protein, zinc metalloprotease s |

| | | | | | | |
|---|---|---|---|---|---|---|
| AP53_291 | 0.188766152 | 7.932082255 | 2.780217008 | 0.122609993 | 0.219341932 | Segregation and condensation prot |
| AP53_85 | -0.379251425 | 3.168246607 | 2.776173181 | 0.122854936 | 0.219577745 | Late competence protein ComGC, a |
| AP53_89 | -0.366109464 | 3.062870992 | 2.771776793 | 0.12312193 | 0.219728752 | Late competence protein ComGG, I |
| AP53_1533 | -0.226478758 | 8.145137342 | 2.770927639 | 0.123173583 | 0.219728752 | Sucrose-6-phosphate hydrolase (EC |
| AP53_625 | 0.537960778 | 3.57983179 | 2.769195403 | 0.123279037 | 0.219728752 | hypothetical phage protein |
| AP53_1118 | -0.332336145 | 4.120227588 | 2.765495069 | 0.123504682 | 0.219889122 | internalin, putative |
| AP53_133 | -0.366040506 | 10.50319653 | 2.759279485 | 0.123884871 | 0.219889122 | Transcription antitermination prot |
| AP53_401 | -0.347527541 | 4.320376361 | 2.757375891 | 0.124001601 | 0.219889122 | FIG01114077: hypothetical protei |
| AP53_1146 | -0.334146022 | 9.620017711 | 2.756944317 | 0.124028084 | 0.219889122 | Oligoendopeptidase F |
| AP53_544 | -0.291877291 | 8.092722271 | 2.756919054 | 0.124029635 | 0.219889122 | FIG01114091: hypothetical protei |
| AP53_1836 | 0.28123514 | 7.000800069 | 2.756607929 | 0.124048732 | 0.219889122 | MutT/nudix family protein |
| AP53_478 | -0.157809986 | 6.86776325 | 2.737849954 | 0.125206934 | 0.221739653 | PROTEASE (EC 3.4.-.-) |
| AP53_378 | -0.326669858 | 4.547407381 | 2.73203352 | 0.125568816 | 0.222058518 | Peptide ABC transporter ATP bindir |
| AP53_1069 | -0.413032791 | 5.517887881 | 2.731280019 | 0.125615792 | 0.222058518 | Mobile element protein |
| AP53_395 | -0.417333003 | 4.13609729 | 2.726805195 | 0.125895225 | 0.222349984 | Bacteriocin-like peptide M BlpM |
| AP53_515 | -0.225089019 | 6.975175126 | 2.724709063 | 0.126026387 | 0.222379289 | 2-dehydro-3-deoxygluconate kinas |
| AP53_974 | 0.219223764 | 5.52041317 | 2.717333225 | 0.126489282 | 0.222993367 | ABC transporter ATP-binding protei |
| AP53_1136 | -0.481373352 | 5.191885985 | 2.712702805 | 0.126780968 | 0.223304773 | Mobile element protein |
| AP53_1458 | -0.199461807 | 9.111977327 | 2.710572982 | 0.126915416 | 0.223338915 | FIG000325: clustered with transcri |
| AP53_1824 | -0.362350232 | 3.817303732 | 2.702167032 | 0.127447801 | 0.224072629 | FIG014387: Transcriptional regulat |
| AP53_593 | 0.221766021 | 6.727169012 | 2.681672845 | 0.128757572 | 0.226170541 | putative glycosyltransferase - possil |
| AP53_1040 | 0.40712227 | 5.15098903 | 2.666305421 | 0.12975079 | 0.227572586 | Phage protein |
| AP53_1186 | 0.424172155 | 2.275852303 | 2.665698105 | 0.129790238 | 0.227572586 | Phage hyaluronidase |
| AP53_211 | 0.203573608 | 8.865840499 | 2.650294063 | 0.130795878 | 0.22912888 | SSU rRNA (adenine(1518)-N(6)/ader |
| AP53_397 | -0.67703649 | 3.124222425 | 2.646001728 | 0.131077841 | 0.229415771 | hypothetical protein |
| AP53_415 | -0.250103713 | 9.242783866 | 2.642834436 | 0.13128639 | 0.229573769 | FIG001614: Membrane protein |
| AP53_1354 | -0.598488933 | 3.37901892 | 2.626045193 | 0.132398861 | 0.231121096 | putative phage replication protein |
| AP53_1590 | -0.229067729 | 9.872711188 | 2.625886893 | 0.132409407 | 0.231121096 | Putative stomatin/prohibitin-famil |
| AP53_1279 | 0.228001708 | 6.54295096 | 2.620073555 | 0.132797403 | 0.23114273 | FIG01114815: hypothetical protei |
| AP53_879 | 0.146714928 | 8.401266095 | 2.618866516 | 0.132878142 | 0.23114273 | TsaC protein (YrdC domain) require |
| AP53_833 | -0.326961353 | 6.204504065 | 2.618644326 | 0.132893012 | 0.23114273 | D-alanyl-D-alanine carboxypeptida: |
| AP53_1347 | -0.51304991 | 4.277213214 | 2.618567724 | 0.132898138 | 0.23114273 | Single-stranded DNA-binding prote |
| AP53_1693 | -0.326452019 | 13.096781 | 2.607446493 | 0.1336451 | 0.232233787 | Antiphagocytic M protein |
| AP53_1157 | -0.332644102 | 3.708622363 | 2.59513156 | 0.134478411 | 0.233374791 | Late competence protein ComEC, D |
| AP53_1099 | -0.293444888 | 5.804455639 | 2.594192967 | 0.13454219 | 0.233374791 | Mobile element protein |

| | | | | | | |
|---|---|---|---|---|---|---|
| AP53_396 | -0.844266601 | 2.364449789 | 2.584961021 | 0.135171553 | 0.234257129 | FIG01116077: hypothetical protei |
| AP53_1167 | 0.179866224 | 8.0253434 | 2.573608024 | 0.135950603 | 0.235335889 | D-alanine--D-alanine ligase A (EC 6.3 |
| AP53_423 | -0.233837346 | 10.05280575 | 2.572361118 | 0.136036511 | 0.235335889 | Glycosyltransferase LafA, responsib |
| AP53_394 | -0.324699803 | 4.161062865 | 2.566637229 | 0.136431747 | 0.235809457 | Competence-stimulating peptide A |
| AP53_924 | -0.245640133 | 8.901812604 | 2.563669012 | 0.136637273 | 0.235954578 | Transcriptional regulator in cluster |
| AP53_482 | -0.243974363 | 7.245107736 | 2.558818809 | 0.136973951 | 0.236217922 | FIG01116574: hypothetical protei |
| AP53_842 | 0.220859802 | 7.789003183 | 2.557967278 | 0.137033168 | 0.236217922 | Spermidine Putrescine ABC transpo |
| AP53_1166 | -0.183516125 | 8.162139025 | 2.555388533 | 0.137212696 | 0.23629675 | UDP-N-acetylmuramoylalanyl-D-glu |
| AP53_1110 | -0.19738309 | 7.404757561 | 2.55381517 | 0.137322377 | 0.23629675 | 5-Enolpyruvylshikimate-3-phospha |
| AP53_1463 | 0.201353186 | 7.910418677 | 2.550623383 | 0.137545219 | 0.236445761 | ABC transporter, ATP-binding prote |
| AP53_1 | -0.231893592 | 7.944552078 | 2.54908759 | 0.137652607 | 0.236445761 | Chromosomal replication initiator |
| AP53_1212 | 0.433810255 | 4.201745088 | 2.544911094 | 0.137945175 | 0.236738802 | Cro-like phage transcriptional repre |
| AP53_1630 | 0.390576134 | 11.41737697 | 2.536814323 | 0.138514601 | 0.237506043 | SSU ribosomal protein S9p (S16e) |
| AP53_1163 | 0.286050413 | 10.45687235 | 2.523425775 | 0.139462704 | 0.238920661 | Cold-shock DEAD-box protein A |
| AP53_1244 | -0.215829719 | 8.999923432 | 2.516942484 | 0.139924758 | 0.239500843 | Cell division protein FtsQ |
| AP53_667 | -0.346103859 | 10.64619676 | 2.514903044 | 0.140070505 | 0.239539075 | LSU ribosomal protein L35p |
| AP53_66 | 0.416414113 | 10.13338686 | 2.506099627 | 0.140701838 | 0.240406925 | LSU ribosomal protein L36p |
| AP53_1711 | -0.561281572 | 3.491323419 | 2.500363098 | 0.141115164 | 0.240901085 | hypothetical protein |
| AP53_601 | -0.495542248 | 8.445616324 | 2.490286222 | 0.14184494 | 0.241879104 | Phage lysin |
| AP53_1321 | 0.591633202 | 2.997535762 | 2.489015575 | 0.1419373 | 0.241879104 | Phage protein |
| AP53_1714 | -0.486211401 | 3.071270221 | 2.483547617 | 0.142335618 | 0.242345117 | FIG01115319: hypothetical protei |
| AP53_807 | 0.250598746 | 6.845634405 | 2.467317342 | 0.143526276 | 0.244158195 | Acyl-ACP thioesterase |
| AP53_1560 | 0.271451632 | 7.297212437 | 2.454302764 | 0.144490141 | 0.24558263 | hypothetical protein |
| AP53_166 | -0.16590199 | 7.535923888 | 2.451360975 | 0.144709145 | 0.245739676 | membrane protein |
| AP53_1264 | 0.517840458 | 10.19077399 | 2.447499724 | 0.144997237 | 0.246013668 | Arginine/ornithine antiporter ArcD |
| AP53_480 | 0.355544621 | 5.721513167 | 2.430008139 | 0.146311433 | 0.24802663 | FIG01114033: hypothetical protei |
| AP53_781 | 0.175195086 | 9.416215669 | 2.427741884 | 0.146482806 | 0.248100459 | dTDP-glucose 4,6-dehydratase (EC 4 |
| AP53_188 | -0.378871914 | 7.499966239 | 2.422772713 | 0.146859462 | 0.248521548 | Deoxyuridine 5'-triphosphate nucle |
| AP53_990 | -0.379570187 | 8.554446163 | 2.404043634 | 0.148290178 | 0.250724072 | Alkylphosphonate utilization opero |
| AP53_642 | -0.384418313 | 5.138889041 | 2.400314896 | 0.148577122 | 0.250990596 | Helicase loader DnaI |
| AP53_1806 | 0.224360357 | 6.705292417 | 2.398297836 | 0.148732639 | 0.251034828 | Arginine pathway regulatory protei |
| AP53_1102 | 0.34611014 | 6.954638951 | 2.374804934 | 0.150559252 | 0.253748012 | Dihydrolipoamide acyltransferases |
| AP53_232 | 0.231783212 | 8.703000013 | 2.374264541 | 0.150601602 | 0.253748012 | Iron-sulfur cluster assembly protein |
| AP53_802 | 0.211991352 | 6.417002228 | 2.372320698 | 0.150754066 | 0.253784598 | Hypothetical protein YbbP, contain |
| AP53_222 | 0.238731287 | 12.83297456 | 2.36831515 | 0.151068856 | 0.253854211 | NAD-dependent glyceraldehyde-3-p |

| | | | | | | |
|---|---|---|---|---|---|---|
| AP53_296 | 0.28750205 | 7.707430947 | 2.367892427 | 0.151102126 | 0.253854211 | Membrane-associated phospholipi |
| AP53_15 | -0.17879299 | 7.727893238 | 2.366804745 | 0.151187773 | 0.253854211 | Cationic amino acid transporter - A |
| AP53_511 | -0.235768287 | 6.045520278 | 2.361315889 | 0.151620924 | 0.254253855 | Unsaturated glucuronyl hydrolase ( |
| AP53_40 | 0.509539584 | 10.27616672 | 2.360470583 | 0.151687771 | 0.254253855 | Alcohol dehydrogenase (EC 1.1.1.1) |
| AP53_1816 | -0.28325957 | 5.463694811 | 2.350348319 | 0.152491159 | 0.255379931 | Cadmium resistance protein |
| AP53_137 | -0.480345945 | 6.003092304 | 2.344015828 | 0.152996509 | 0.256005366 | FIG01114743: hypothetical protei |
| AP53_721 | -0.379692411 | 7.332076412 | 2.337718817 | 0.153501141 | 0.256628522 | Peptide deformylase (EC 3.5.1.88) |
| AP53_1348 | -0.352715819 | 4.605248262 | 2.335195098 | 0.153703981 | 0.256737244 | Single-stranded DNA-binding prote |
| AP53_1225 | -0.282396081 | 7.639382578 | 2.333620459 | 0.153830714 | 0.256737244 | Exodeoxyribonuclease VII small sub |
| AP53_1070 | -0.295708087 | 6.090033599 | 2.322068409 | 0.154764537 | 0.258073855 | Mobile element protein |
| AP53_1792 | -0.584655775 | 4.794705572 | 2.318889265 | 0.155022792 | 0.258282609 | phage protein |
| AP53_1672 | -0.289520703 | 7.82243254 | 2.303374792 | 0.156290996 | 0.260172233 | PTS system, maltose and glucose-sp |
| AP53_1428 | -0.239783191 | 7.56840856 | 2.297048111 | 0.156811951 | 0.260815764 | FIG01114118: hypothetical protei |
| AP53_1610 | -0.164073797 | 7.599596944 | 2.295232438 | 0.156961866 | 0.260841595 | Type I restriction-modification syst |
| AP53_572 | -0.288851151 | 8.917857401 | 2.287282772 | 0.157620404 | 0.261334317 | DNA-entry nuclease (Competence-s |
| AP53_1568 | -0.470225581 | 4.251335018 | 2.284914789 | 0.157817244 | 0.261334317 | Transcriptional regulator, Cro/CI fa |
| AP53_1669 | 0.164998941 | 7.375156326 | 2.283141751 | 0.157964835 | 0.261334317 | Ribonucleotide reduction protein I |
| AP53_rna69 | -5.616224634 | -2.070645075 | 2.680785734 | 0.158003365 | 0.261334317 | 5S RNA |
| AP53_1148 | -0.26487057 | 11.24128382 | 2.281793564 | 0.158077178 | 0.261334317 | Ribosomal small subunit pseudouri |
| AP53_688 | -0.225280822 | 9.040885876 | 2.280484329 | 0.158186372 | 0.261334317 | Aspartate carbamoyltransferase (EC |
| AP53_125 | -0.272116107 | 5.993230058 | 2.280310994 | 0.158200836 | 0.261334317 | hypothetical protein |
| AP53_872 | -0.304479528 | 7.873853789 | 2.274405788 | 0.158694604 | 0.261927064 | Xanthine phosphoribosyltransferas |
| AP53_1653 | 0.198117211 | 6.900789998 | 2.267870051 | 0.159243389 | 0.262494315 | Transcriptional regulator, MarR fan |
| AP53_196 | -0.141648313 | 8.712936276 | 2.267093428 | 0.15930876 | 0.262494315 | Ribonuclease P protein component |
| AP53_1569 | -0.260043334 | 7.211031447 | 2.24953502 | 0.1607959 | 0.264719967 | putative repressor - phage associate |
| AP53_24 | -0.522330532 | 8.609949346 | 2.24590758 | 0.161105338 | 0.265004628 | Phosphoribosylformylglycinamidir |
| AP53_1497 | -0.286961013 | 7.204863828 | 2.224556572 | 0.162942182 | 0.267799133 | Integral membrane protein |
| AP53_631 | 0.334017601 | 4.708619 | 2.221349818 | 0.163220369 | 0.268029388 | putaive immunity repressor protei |
| AP53_651 | 0.490510593 | 3.86697987 | 2.208133513 | 0.16437332 | 0.269694517 | hypothetical phage protein |
| AP53_rna76 | 1.75613531 | 0.798034033 | 2.191236783 | 0.165862562 | 0.271803542 | tRNA-Asn-GTT |
| AP53_425 | 0.320656469 | 7.428724349 | 2.190069306 | 0.165966098 | 0.271803542 | Threonyl-tRNA synthetase (EC 6.1.1 |
| AP53_218 | -0.173497699 | 10.12084754 | 2.188799244 | 0.166078825 | 0.271803542 | Surface exclusion protein |
| AP53_1573 | -0.218960274 | 8.771126769 | 2.176712782 | 0.167156512 | 0.273158065 | Transcriptional regulator OrfX |
| AP53_838 | 0.33107238 | 8.42697884 | 2.176361865 | 0.167187935 | 0.273158065 | Dihydroneopterin aldolase (EC 4.1. |
| AP53_rna5 | 0.771947528 | 4.152167762 | 2.173775538 | 0.167419763 | 0.273306778 | tRNA-Val-TAC |

| | | | | | | |
|---|---|---|---|---|---|---|
| AP53_210 | 0.156722179 | 8.282924267 | 2.166251563 | 0.168096529 | 0.274180977 | Ribonuclease M5 (EC 3.1.26.8) |
| AP53_920 | -0.249982957 | 9.684761164 | 2.156294419 | 0.168997556 | 0.27541919 | Tyrosine recombinase XerC |
| AP53_1152 | 0.196675654 | 7.02683456 | 2.144656266 | 0.170058567 | 0.276915838 | FIG01113992: hypothetical protei |
| AP53_1728 | 0.19450339 | 7.033408301 | 2.142822045 | 0.170226566 | 0.276957053 | Hypothetical protein SPy2060 |
| AP53_1462 | 0.158092044 | 7.438063323 | 2.139739968 | 0.170509338 | 0.277032262 | ABC transporter, permease protein |
| AP53_1097 | -0.252858064 | 6.178719169 | 2.139207574 | 0.170558245 | 0.277032262 | Malonate permease |
| AP53_1480 | 0.298272409 | 9.729546962 | 2.137352563 | 0.17072879 | 0.27707741 | (3R)-hydroxymyristoyl-[acyl carrier |
| AP53_1831 | -0.423898511 | 5.756421826 | 2.13112455 | 0.171302985 | 0.27777702 | FIG01114855: hypothetical protei |
| AP53_280 | 0.14524879 | 8.991995718 | 2.125565923 | 0.171817561 | 0.27837887 | Predicted nucleoside phosphatase |
| AP53_1453 | -0.278972252 | 10.39467819 | 2.120990615 | 0.172242601 | 0.278513842 | Ribosome-binding factor A |
| AP53_728 | 0.21118173 | 7.770431655 | 2.120059277 | 0.172329286 | 0.278513842 | Phosphomevalonate kinase (EC 2.7. |
| AP53_569 | -0.186696016 | 8.07311659 | 2.120037255 | 0.172331336 | 0.278513842 | Competence-associated EpuA prote |
| AP53_1536 | 0.201522324 | 11.00225356 | 2.117437055 | 0.172573654 | 0.27867343 | Alkaline shock protein |
| AP53_1490 | -0.281347589 | 13.61212773 | 2.114775159 | 0.172822175 | 0.278842761 | Chaperone protein DnaJ |
| AP53_1366 | -0.16987451 | 9.455194799 | 2.098059788 | 0.174393328 | 0.281144062 | Cysteinyl-tRNA synthetase related p |
| AP53_23 | -0.42082094 | 8.50615647 | 2.093650169 | 0.174810869 | 0.281583317 | Amidophosphoribosyltransferase (I |
| AP53_1079 | 0.273286318 | 6.775970913 | 2.074011034 | 0.176686185 | 0.284368064 | Glutamine ABC transporter, glutam |
| AP53_1288 | -0.153662383 | 9.249318484 | 2.070060576 | 0.177066536 | 0.284656154 | hypothetical protein |
| AP53_1478 | 0.285637133 | 10.06615235 | 2.069107119 | 0.177158493 | 0.284656154 | Acetyl-coenzyme A carboxyl transfe |
| AP53_174 | -0.250579733 | 6.388664455 | 2.065688676 | 0.177488694 | 0.284950832 | Streptococcal pyrogenic exotoxin ( |
| AP53_1223 | -0.242204925 | 8.566335184 | 2.054951267 | 0.17853104 | 0.286387395 | RNA binding methyltransferase FtsJ |
| AP53_39 | 0.237689914 | 10.17853263 | 2.04804573 | 0.179205577 | 0.28723206 | Alcohol dehydrogenase (EC 1.1.1.1) |
| AP53_677 | -0.187346046 | 8.111374097 | 2.045881176 | 0.179417689 | 0.287334764 | Cysteine desulfurase (EC 2.8.1.7) |
| AP53_405 | -0.180363246 | 5.795434844 | 2.042568442 | 0.179742943 | 0.287618344 | Positive transcriptional regulator, I |
| AP53_1273 | -0.24866734 | 6.981510129 | 2.038153519 | 0.180177597 | 0.287797421 | Two-component response regulato |
| AP53_599 | -0.765519664 | 9.392532048 | 2.03803637 | 0.180189149 | 0.287797421 | Phage protein |
| AP53_1657 | 0.187780922 | 8.21286656 | 2.036916037 | 0.180299672 | 0.287797421 | Phosphatidate cytidylyltransferase |
| AP53_308 | 0.19220598 | 7.965217081 | 2.031351506 | 0.180849924 | 0.288264954 | Membrane protein involved in the ( |
| AP53_167 | -0.302430518 | 6.615908403 | 2.030951116 | 0.1808896 | 0.288264954 | FIG01116967: hypothetical protei |
| AP53_263 | -0.139420523 | 6.84634355 | 2.017448448 | 0.18223425 | 0.29012289 | rRNA small subunit 7-methylguano |
| AP53_363 | 0.162995821 | 9.754538294 | 2.016248015 | 0.182354418 | 0.29012289 | Ribosome recycling factor |
| AP53_1075 | 0.271337672 | 9.355194238 | 2.009508397 | 0.183030989 | 0.290920044 | D-alanyl transfer protein DltB |
| AP53_29 | -0.521007534 | 9.432280448 | 2.00827432 | 0.183155226 | 0.290920044 | Phosphoribosylaminoimidazole ca |
| AP53_1709 | -0.3671482 | 6.130877228 | 2.004202275 | 0.183565941 | 0.290981521 | Strepotococcal cysteine protease (S |
| AP53_818 | 0.142190367 | 9.40215327 | 2.004060076 | 0.183580305 | 0.290981521 | Putative membrane protein YeiH |

| | | | | | | |
|---|---|---|---|---|---|---|
| AP53_1319 | 0.400609255 | 3.772270914 | 2.003432961 | 0.18364367 | 0.290981521 | Phage-related protein |
| AP53_1717 | -0.216253226 | 6.665222329 | 1.994394367 | 0.184560093 | 0.292195057 | Transaldolase (EC 2.2.1.2) |
| AP53_711 | 0.499803124 | 2.784300559 | 1.983899134 | 0.185631649 | 0.293565232 | Mobile element protein |
| AP53_817 | 0.153737702 | 9.179772291 | 1.982958994 | 0.185728029 | 0.293565232 | Peptide methionine sulfoxide redu |
| AP53_295 | -0.294251659 | 6.996968242 | 1.978902358 | 0.186144649 | 0.29376665 | Substrate-specific component RibU |
| AP53_1417 | -0.181570845 | 8.884941693 | 1.978771039 | 0.186158155 | 0.29376665 | Methionine ABC transporter ATP-bi |
| AP53_1754 | -0.183465212 | 11.52845548 | 1.972875595 | 0.186765841 | 0.294486188 | SSU ribosomal protein S2p (SAe) |
| AP53_1853 | -0.16285342 | 7.660663908 | 1.963868352 | 0.187699258 | 0.295717743 | DNA recombination and repair pro |
| AP53_238 | 0.139484123 | 9.458554412 | 1.955212315 | 0.18860199 | 0.296898996 | Oligopeptide ABC transporter, peri |
| AP53_757 | -0.24592388 | 8.988796479 | 1.943963625 | 0.189783547 | 0.298516909 | Topoisomerase IV subunit A (EC 5.9 |
| AP53_765 | -0.267508924 | 8.431233271 | 1.939682212 | 0.19023579 | 0.298823724 | Exfoliative toxin A |
| AP53_709 | -0.259321314 | 5.106968656 | 1.939202839 | 0.190286514 | 0.298823724 | Peptidoglycan hydrolase, Autolysir |
| AP53_1565 | 0.441908943 | 7.738902507 | 1.930642738 | 0.191195235 | 0.300008044 | transporter |
| AP53_235 | -0.187820395 | 8.612244144 | 1.922880717 | 0.192024117 | 0.301065275 | Iron-sulfur cluster assembly proteir |
| AP53_983 | -0.204751454 | 8.231438178 | 1.920268878 | 0.192304077 | 0.301260867 | ATP-dependent DNA helicase UvrD/ |
| AP53_414 | 0.213154226 | 9.780415718 | 1.911751843 | 0.193220709 | 0.302452739 | Pyrrolidone-carboxylate peptidase |
| AP53_1272 | -0.269059491 | 7.465339886 | 1.909077865 | 0.193509664 | 0.302660966 | Two-component sensor kinase yesN |
| AP53_27 | -0.58612997 | 7.905731302 | 1.892588762 | 0.195304001 | 0.305066283 | putative choline binding protein |
| AP53_513 | 0.191289815 | 5.840755779 | 1.892060265 | 0.19536187 | 0.305066283 | 2-deoxy-D-gluconate 3-dehydroger |
| AP53_298 | -0.163477167 | 7.251107526 | 1.890401722 | 0.19554362 | 0.305104635 | SAM-dependent methyltransferase, |
| AP53_1481 | 0.352003057 | 10.0964202 | 1.884681036 | 0.196172211 | 0.30567714 | Biotin carboxyl carrier protein of ac |
| AP53_1629 | 0.596855831 | 2.025165839 | 1.88419706 | 0.196225511 | 0.30567714 | Protein of unknown function DUF2 |
| AP53_1432 | 0.215936457 | 10.71659941 | 1.874054978 | 0.197346817 | 0.307177364 | oxidoreductase of aldo/keto reduct |
| AP53_1424 | 0.189936284 | 7.751467306 | 1.869970082 | 0.197800805 | 0.307637309 | Alpha-glycerophosphate oxidase (E |
| AP53_1316 | 0.375735437 | 3.674597187 | 1.863808551 | 0.198488168 | 0.308243898 | Phage hyaluronidase |
| AP53_632 | 0.288715214 | 4.285425534 | 1.863627298 | 0.198508435 | 0.308243898 | hypothetical protein |
| AP53_rna61 | 1.041340158 | 1.540564622 | 1.861064248 | 0.198795319 | 0.308442617 | tRNA-Thr-GGT |
| AP53_1689 | -0.418779466 | 2.901079833 | 1.855958672 | 0.199368405 | 0.309084723 | hypothetical protein |
| AP53_582 | -0.132476599 | 8.124503157 | 1.845649561 | 0.200532171 | 0.310640817 | Amino acid ABC transporter, aminc |
| AP53_664 | -0.346116537 | 7.83188895 | 1.84416841 | 0.200700103 | 0.31065303 | FIG01114555: hypothetical protei |
| AP53_272 | -0.130840754 | 9.628509917 | 1.842037233 | 0.200942057 | 0.310779708 | GTP-binding protein EngA |
| AP53_1662 | 0.230199453 | 4.998466761 | 1.839989097 | 0.201174943 | 0.31089217 | Glucan 1,6-alpha-glucosidase (EC 3. |
| AP53_208 | -0.172160555 | 7.245340904 | 1.83780643 | 0.201423516 | 0.311028675 | Sialic acid utilization regulator, Rpi |
| AP53_980 | -0.506609434 | 11.32503678 | 1.835073265 | 0.201735348 | 0.311262569 | FIG01114232: hypothetical protei |
| AP53_1724 | -0.240478759 | 8.526789921 | 1.83317213 | 0.201952626 | 0.311350315 | Pyruvate formate-lyase activating e |

| | | | | | | |
|---|---|---|---|---|---|---|
| AP53_855 | 0.253609341 | 6.633769712 | 1.82237615 | 0.203192306 | 0.31301291 | Permease |
| AP53_722 | -0.304102381 | 7.447617739 | 1.815350269 | 0.204004433 | 0.314014753 | 5'-nucleotidase (EC 3.1.3.5) |
| AP53_1049 | 0.413235433 | 4.90069802 | 1.811329744 | 0.204471082 | 0.314483653 | hypothetical phage protein |
| AP53_700 | 0.265565729 | 4.841515719 | 1.801791482 | 0.205583771 | 0.315944655 | Transcriptional regulator, TetR fam |
| AP53_612 | 0.450070976 | 4.914644896 | 1.791617036 | 0.206779448 | 0.317452846 | Phage base plate |
| AP53_481 | 0.334225312 | 8.287145628 | 1.79066118 | 0.206892246 | 0.317452846 | bacterial seryl-tRNA synthetase rela |
| AP53_878 | -0.128848402 | 7.663320523 | 1.783014495 | 0.207797531 | 0.318590054 | Protein-N(5)-glutamine methyltran |
| AP53_1050 | 0.22613143 | 5.944521376 | 1.777360763 | 0.208470221 | 0.319369139 | hypothetical protein |
| AP53_1790 | -0.460358937 | 4.848679332 | 1.764652309 | 0.209992778 | 0.321447936 | FIG01118442: hypothetical protei |
| AP53_54 | 0.262540715 | 11.32762119 | 1.760317246 | 0.210515493 | 0.321810907 | LSU ribosomal protein L24p (L26e) |
| AP53_406 | -0.17817454 | 6.982701596 | 1.759936455 | 0.21056149 | 0.321810907 | Formamidopyrimidine-DNA glycos |
| AP53_327 | -0.170933865 | 8.892621413 | 1.757174926 | 0.210895459 | 0.321962121 | FIG01118451: hypothetical protei |
| AP53_214 | -0.137852145 | 9.089887731 | 1.754827631 | 0.211179882 | 0.321962121 | Thiamin pyrophosphokinase (EC 2. |
| AP53_221 | 0.232230054 | 11.87240178 | 1.754049453 | 0.211274285 | 0.321962121 | Translation elongation factor G |
| AP53_92 | -0.216362238 | 6.356531205 | 1.753494853 | 0.2113416 | 0.321962121 | Hypothetical protein SPy0110 |
| AP53_152 | -0.181212627 | 9.753289459 | 1.752274646 | 0.211489801 | 0.321962121 | Succinyl-CoA synthetase, alpha sub |
| AP53_1532 | -0.350599817 | 11.03730632 | 1.750296594 | 0.211730339 | 0.322075696 | PTS system, sucrose-specific IIB con |
| AP53_243 | -0.418765913 | 4.137115508 | 1.748812062 | 0.2119111 | 0.322098234 | Transposase |
| AP53_106 | -0.161071961 | 7.294570993 | 1.747190375 | 0.212108793 | 0.322146453 | NPQTN specific sortase B |
| AP53_1843 | -0.123828302 | 8.406362663 | 1.745794309 | 0.212279177 | 0.322153152 | ATPase component of general energ |
| AP53_1726 | -0.626727376 | 9.766519192 | 1.73179885 | 0.213997273 | 0.324203129 | Preprotein translocase subunit Sec |
| AP53_1207 | 0.465900342 | 3.359093794 | 1.731200873 | 0.21407109 | 0.324203129 | hypothetical protein |
| AP53_306 | -0.235602558 | 6.313976703 | 1.730715159 | 0.214131072 | 0.324203129 | Ferrichrome transport ATP-binding |
| AP53_99 | -0.176691135 | 8.792586277 | 1.727490017 | 0.214529921 | 0.32455384 | Deoxyadenosine kinase (EC 2.7.1.7 |
| AP53_323 | 0.233937827 | 7.876094137 | 1.717235201 | 0.215804655 | 0.326133923 | Exodeoxyribonuclease III (EC 3.1.1 |
| AP53_1349 | -0.431273366 | 3.309771167 | 1.716388089 | 0.215910402 | 0.326133923 | hypothetical protein (phage associa |
| AP53_8 | 0.304812982 | 7.292519567 | 1.70783927 | 0.216981421 | 0.327496842 | Ribosome-associated heat shock pr |
| AP53_767 | 0.152354924 | 6.583723387 | 1.705256833 | 0.217306337 | 0.3277324 | tRNA dimethylallyltransferase (EC 2 |
| AP53_1618 | 0.190235913 | 7.218910504 | 1.701364026 | 0.217797336 | 0.32821788 | PTS system, lactose-specific IIB com |
| AP53_342 | 0.283033632 | 9.409220889 | 1.691640848 | 0.219030147 | 0.32981964 | FIG01114229: hypothetical protei |
| AP53_1369 | -0.258096908 | 4.662961336 | 1.679122778 | 0.220630967 | 0.33197264 | ComF operon protein A, DNA trans |
| AP53_1108 | -0.149638821 | 8.456743553 | 1.676941576 | 0.220911484 | 0.332137251 | Cell envelope-associated transcript |
| AP53_1205 | -0.395740075 | 2.924279 | 1.67381693 | 0.22131416 | 0.332485127 | FIG01114509: hypothetical protei |
| AP53_1765 | -0.206460553 | 10.01501353 | 1.669411528 | 0.221883543 | 0.333082719 | Ribonucleotide reductase of class II |
| AP53_rna38 | 1.886278072 | -0.468952949 | 1.70241447 | 0.222962739 | 0.334444108 | 5S RNA |

| | | | | | | |
|---|---|---|---|---|---|---|
| AP53_616 | 0.464180496 | 5.136891904 | 1.655083243 | 0.223748918 | 0.335207393 | hypothetical phage protein |
| AP53_1221 | 0.124909325 | 8.478727955 | 1.654563353 | 0.223816992 | 0.335207393 | DNA repair protein RecN |
| AP53_219 | -0.274702081 | 11.68278344 | 1.652649978 | 0.224067766 | 0.33527207 | SSU ribosomal protein S12p (S23e) |
| AP53_1195 | 0.539283904 | 3.18282213 | 1.650790092 | 0.224311887 | 0.33527207 | FIG033577: Phage protein |
| AP53_1234 | -0.193883123 | 9.054293806 | 1.650284037 | 0.224378371 | 0.33527207 | Mutator mutT protein (7,8-dihydro |
| AP53_1800 | -0.368552858 | 4.960088655 | 1.640960077 | 0.225608025 | 0.336756818 | FIG01115581: hypothetical protei |
| AP53_74 | 0.175186617 | 6.906016983 | 1.640121765 | 0.225719021 | 0.336756818 | Transcriptional repressor AdcR for |
| AP53_194 | -0.185358602 | 6.239699763 | 1.635036481 | 0.226393893 | 0.33750426 | Sensory transduction protein kinas |
| AP53_495 | -0.236414292 | 6.404686299 | 1.628835355 | 0.227220487 | 0.338476566 | FIG01114557: hypothetical protei |
| AP53_1355 | -0.457566826 | 4.089876018 | 1.626939337 | 0.227474023 | 0.338594386 | hypothetical protein, phage associa |
| AP53_979 | 0.32118476 | 13.36863302 | 1.624526227 | 0.22779725 | 0.33881568 | General stress protein, Gls24 family |
| AP53_1531 | -0.237117536 | 9.761708956 | 1.62147431 | 0.228206919 | 0.339165107 | Secreted Endo-beta-N-acetylglucos |
| AP53_1073 | 0.217201541 | 9.28714035 | 1.610814783 | 0.22964551 | 0.341042031 | Poly(glycerophosphate chain) D-ala |
| AP53_634 | 0.38434679 | 4.245122489 | 1.609428058 | 0.229833547 | 0.341060333 | hypothetical phage protein |
| AP53_982 | 0.222869707 | 11.67228753 | 1.607316739 | 0.230120233 | 0.341154008 | FIG01114177: hypothetical protei |
| AP53_302 | -0.156206991 | 9.263777797 | 1.606375475 | 0.230248197 | 0.341154008 | FIG01114725: hypothetical protei |
| AP53_3 | 0.356140438 | 5.923167754 | 1.604872306 | 0.230452748 | 0.341196631 | FIG01114317: hypothetical protei |
| AP53_1548 | -0.133281998 | 7.216243169 | 1.602885717 | 0.230723454 | 0.341337061 | Transcriptional regulator, XRE fami |
| AP53_800 | 0.117148674 | 8.470305784 | 1.596105228 | 0.231650614 | 0.342043486 | Putative amidotransferase similar t |
| AP53_1120 | 0.127772649 | 8.887645674 | 1.595691953 | 0.231707285 | 0.342043486 | FIG01114335: hypothetical protei |
| AP53_546 | 0.299187111 | 7.330619277 | 1.595529144 | 0.231729616 | 0.342043486 | Streptolysin S biosynthesis protein |
| AP53_rna10 | 1.030495849 | 2.562764711 | 1.577024178 | 0.234286599 | 0.345385666 | tRNA-Gly-GCC |
| AP53_1621 | 0.26075501 | 6.750819631 | 1.576570363 | 0.23434978 | 0.345385666 | Tagatose-6-phosphate kinase (EC 2. |
| AP53_1449 | -0.443863027 | 5.031016365 | 1.575197715 | 0.234541019 | 0.345405249 | Copper chaperone |
| AP53_1395 | -0.449185074 | 7.290890495 | 1.572779094 | 0.234878493 | 0.345533897 | Cell division protein GpsB, coordin |
| AP53_1008 | 0.383582729 | 5.425898908 | 1.570902478 | 0.235140788 | 0.345533897 | Phage protein |
| AP53_1109 | -0.137338832 | 6.796823817 | 1.570747754 | 0.235162431 | 0.345533897 | Shikimate kinase I (EC 2.7.1.71) |
| AP53_26 | -0.516588895 | 9.770366314 | 1.563666071 | 0.236155902 | 0.346473981 | IMP cyclohydrolase (EC 3.5.4.10) / |
| AP53_1145 | -0.290741687 | 7.188093254 | 1.56364237 | 0.236159236 | 0.346473981 | Major facilitator:Oxalate:Formate |
| AP53_131 | -0.137427309 | 10.18789975 | 1.561808127 | 0.236417478 | 0.34659088 | Adenylosuccinate synthetase (EC 6. |
| AP53_1184 | 0.3602765 | 3.96316391 | 1.55849093 | 0.236885467 | 0.346874745 | Phage protein |
| AP53_71 | 0.386332843 | 5.566105766 | 1.557903094 | 0.236968528 | 0.346874745 | hypothetical protein |
| AP53_246 | 0.193277531 | 8.990207353 | 1.555703113 | 0.237279731 | 0.347068544 | FIG001553: Hydrolase, HAD subfam |
| AP53_48 | 0.270440232 | 11.44172855 | 1.549124089 | 0.238213658 | 0.348156428 | LSU ribosomal protein L22p (L17e) |
| AP53_269 | -0.147846373 | 8.07559697 | 1.547940333 | 0.238382222 | 0.348156428 | Ribonucleotide reductase transcrip |

| | | | | | | |
|---|---|---|---|---|---|---|
| AP53_1277 | -0.269430287 | 10.15144344 | 1.544469267 | 0.238877416 | 0.348617341 | Magnesium and cobalt transport pr |
| AP53_321 | 0.162088799 | 9.194140231 | 1.538550495 | 0.239724997 | 0.34959145 | Arsenate reductase (EC 1.20.4.1) |
| AP53_1537 | -0.207484792 | 10.26151985 | 1.535816866 | 0.240117823 | 0.349901422 | Translation elongation factor P |
| AP53_1624 | 0.271654504 | 6.419915055 | 1.532393265 | 0.240611018 | 0.350357079 | Lactose phosphotransferase system |
| AP53_535 | 0.265021068 | 10.82592105 | 1.517636295 | 0.24275253 | 0.353210391 | LSU ribosomal protein L31p @ LSU |
| AP53_325 | 0.532689196 | 6.18675495 | 1.515304825 | 0.243093213 | 0.353441144 | hypothetical protein |
| AP53_1098 | -0.295232651 | 4.219661802 | 1.507669868 | 0.244213378 | 0.354785433 | Mobile element protein |
| AP53_1716 | 0.134066612 | 7.147578025 | 1.506515325 | 0.244383371 | 0.354785433 | Glycerol dehydrogenase (EC 1.1.1.6 |
| AP53_891 | -0.195900526 | 11.92681912 | 1.501769303 | 0.245083849 | 0.355536436 | FIG01114048: hypothetical protei |
| AP53_118 | -0.163985307 | 7.703854256 | 1.496469262 | 0.2458693 | 0.356147856 | Transcriptional regulator pfoR |
| AP53_764 | -0.183885748 | 7.227939036 | 1.496449085 | 0.245872296 | 0.356147856 | Glutathione S-transferase domain p |
| AP53_pseudo603 | -0.386287728 | 7.457202431 | 1.489280448 | 0.246940097 | 0.357427837 | Phage holin |
| AP53_858 | 0.139198005 | 9.390154202 | 1.479502032 | 0.248406757 | 0.359195777 | Redox-sensitive transcriptional reg |
| AP53_1493 | 0.250415302 | 13.43323349 | 1.478672973 | 0.248531648 | 0.359195777 | Heat-inducible transcription repres |
| AP53_206 | -0.175051875 | 6.775523524 | 1.47595106 | 0.248942279 | 0.35952155 | N-acetylneuraminate lyase (EC 4.1.3 |
| AP53_1425 | 0.161705862 | 7.44292707 | 1.463218285 | 0.250875396 | 0.361837723 | Glycerol kinase (EC 2.7.1.30) |
| AP53_1401 | 0.107266312 | 7.825276973 | 1.4625519 | 0.250977127 | 0.361837723 | Nicotinate phosphoribosyltransfera |
| AP53_358 | 0.124967559 | 7.203562096 | 1.461712738 | 0.251105313 | 0.361837723 | Cell division protein FtsK |
| AP53_1228 | 0.218512351 | 10.22004434 | 1.457765979 | 0.251709392 | 0.362439117 | putative phosphomannomutase |
| AP53_1011 | 0.302401606 | 5.389861213 | 1.454934907 | 0.252143921 | 0.362795663 | Hyaluronate lyase (phage associated |
| AP53_1852 | 0.126418557 | 7.54075747 | 1.450064838 | 0.252893788 | 0.363605068 | FIG002958: hypothetical protein |
| AP53_986 | -0.183441108 | 7.078026064 | 1.445923168 | 0.253533881 | 0.364138483 | CAMP factor |
| AP53_1433 | -0.143838846 | 10.47960649 | 1.445237941 | 0.253639994 | 0.364138483 | N-acetylglucosamine-6-phosphate |
| AP53_1506 | -0.122475175 | 9.598567028 | 1.440774424 | 0.254332686 | 0.364863077 | Aspartate aminotransferase (EC 2.6. |
| AP53_1138 | 0.196563127 | 6.670704766 | 1.4382122 | 0.254731477 | 0.365165286 | Membrane-bound protease, CAAX f |
| AP53_pseudo595 | -0.182558348 | 6.325910274 | 1.423153271 | 0.257092543 | 0.368246594 | Lipoteichoic acid synthase LtaS Typ |
| AP53_1061 | -0.233012375 | 5.182023961 | 1.422090012 | 0.257260372 | 0.368246594 | Phage integrase: site-specific recom |
| AP53_1631 | 0.24030081 | 11.02108569 | 1.418140136 | 0.257885145 | 0.368720257 | LSU ribosomal protein L13p (L13Ae |
| AP53_1515 | -0.112600079 | 7.717556279 | 1.417597078 | 0.257971205 | 0.368720257 | Substrate-specific component BL0 |
| AP53_975 | -0.201886378 | 5.218803085 | 1.411413973 | 0.258953821 | 0.369852366 | Transcriptional regulator, TetR fam |
| AP53_1335 | 0.441332826 | 3.024041073 | 1.409664583 | 0.259232758 | 0.369978517 | Phage protein |
| AP53_1542 | -0.148577509 | 9.030240073 | 1.402562368 | 0.260369409 | 0.371196161 | Integral membrane protein |
| AP53_659 | -0.17675589 | 7.44072087 | 1.401945818 | 0.260468403 | 0.371196161 | Heteropolysaccharide repeat unit e |
| AP53_1322 | 0.308829751 | 2.744344257 | 1.394639792 | 0.261645388 | 0.372510847 | Phage protein |
| AP53_161 | -0.167300565 | 6.18135813 | 1.393839506 | 0.261774753 | 0.372510847 | Malonate permease |

| | | | | | | |
|---|---|---|---|---|---|---|
| AP53_1796 | -0.350599356 | 4.779736854 | 1.389204664 | 0.262525689 | 0.37330576 | DNA primase, phage associated |
| AP53_1191 | -0.320990656 | 2.921683598 | 1.386473519 | 0.262969567 | 0.373663198 | Phage protein |
| AP53_380 | -0.291457603 | 5.255508454 | 1.38204484 | 0.263691518 | 0.37441495 | hypothetical protein |
| AP53_153 | -0.157262151 | 10.55554949 | 1.377149118 | 0.264492763 | 0.375278109 | Peroxide stress regulator PerR, FUR |
| AP53_605 | -0.420056146 | 6.486805845 | 1.373970116 | 0.26501483 | 0.375744182 | FIG01116522: hypothetical protei |
| AP53_1384 | 0.137721465 | 10.00184868 | 1.371580105 | 0.265408255 | 0.375888213 | Hydrolase (HAD superfamily) |
| AP53_122 | -0.183787845 | 6.032187503 | 1.370517413 | 0.265583444 | 0.375888213 | V-type ATP synthase subunit E (EC 3 |
| AP53_184 | -0.14491384 | 8.108886386 | 1.369748321 | 0.265710331 | 0.375888213 | Transcriptional regulator, MarR fan |
| AP53_763 | 0.14356482 | 7.618129502 | 1.368653838 | 0.265891044 | 0.375888213 | hypothetical protein |
| AP53_28 | -0.423356162 | 9.347749304 | 1.365821552 | 0.266359474 | 0.376276375 | Phosphoribosylamine--glycine ligas |
| AP53_rna82 | 0.68146958 | 2.664243267 | 1.364357798 | 0.266602007 | 0.376302446 | tRNA-Cys-GCA |
| AP53_506 | -0.152304349 | 7.599458418 | 1.363371077 | 0.26676567 | 0.376302446 | Transcriptional regulator RegR, rpr |
| AP53_530 | -0.172733866 | 9.094709856 | 1.360347843 | 0.267267978 | 0.376737215 | FIG001886: Cytoplasmic hypotheti |
| AP53_1622 | 0.216028686 | 7.523407045 | 1.34822783 | 0.269294785 | 0.379318707 | Galactose-6-phosphate isomerase, |
| AP53_543 | -0.133032507 | 9.723083223 | 1.345500748 | 0.269753736 | 0.379689632 | Septation ring formation regulator |
| AP53_443 | 0.277007501 | 4.271592966 | 1.34085221 | 0.270538537 | 0.380518333 | Archaeal S-adenosylmethionine syr |
| AP53_1755 | -0.16309951 | 11.0368012 | 1.336221934 | 0.271323378 | 0.381345892 | Translation elongation factor Ts |
| AP53_1650 | -0.346053615 | 6.99247406 | 1.332660638 | 0.271929157 | 0.381920762 | FIG01118158: hypothetical protei |
| AP53_56 | 0.244794 | 10.64711432 | 1.321580715 | 0.273825809 | 0.3841799 | SSU ribosomal protein S14p (S29e) |
| AP53_1799 | -0.285029917 | 4.830050309 | 1.320954603 | 0.27393353 | 0.3841799 | FIG01115461: hypothetical protei |
| AP53_352 | -0.167525352 | 8.762418006 | 1.309699065 | 0.27588002 | 0.38663041 | FIG01116268: hypothetical protei |
| AP53_1616 | 0.411607722 | 4.775695083 | 1.305994492 | 0.276524849 | 0.387254496 | Lantibiotic salivaricin A |
| AP53_1383 | 0.130801902 | 9.539422849 | 1.304454439 | 0.276793528 | 0.387274397 | Guanylate kinase (EC 2.7.4.8) |
| AP53_293 | 0.140329514 | 7.345830115 | 1.303626583 | 0.276938106 | 0.387274397 | Protein YidD |
| AP53_227 | 0.136683167 | 7.690531152 | 1.299522887 | 0.277656321 | 0.387999222 | Undecaprenyl-diphosphatase (EC 3. |
| AP53_715 | 0.420403717 | 3.336580757 | 1.295896799 | 0.278293093 | 0.388609275 | Nucleoside diphosphate kinase (EC |
| AP53_1283 | -0.142378058 | 8.006384195 | 1.292557617 | 0.278881269 | 0.389150641 | FIG01114159: hypothetical protei |
| AP53_151 | -0.175609297 | 9.943785209 | 1.289268629 | 0.279462286 | 0.389681248 | DNA polymerase I (EC 2.7.7.7) |
| AP53_140 | -0.197421443 | 9.852723686 | 1.285548076 | 0.280121564 | 0.390063429 | Leucyl-tRNA synthetase (EC 6.1.1.4 |
| AP53_970 | -0.162089575 | 6.591033471 | 1.284649448 | 0.280281123 | 0.390063429 | FIG01115860: hypothetical protei |
| AP53_182 | 0.126461973 | 8.834217995 | 1.284322258 | 0.28033925 | 0.390063429 | UTP--glucose-1-phosphate uridylylt |
| AP53_119 | -0.168653995 | 5.4753407 | 1.281827234 | 0.28078305 | 0.390401075 | V-type ATP synthase subunit G (EC 3 |
| AP53_1861 | -0.131455567 | 7.569565934 | 1.272313384 | 0.282484287 | 0.39237571 | Mobile element protein |
| AP53_1681 | 0.15847926 | 8.463565769 | 1.271006195 | 0.282719151 | 0.39237571 | Dipeptide-binding ABC transporter |
| AP53_899 | -0.140443823 | 8.193042399 | 1.270502654 | 0.282809695 | 0.39237571 | DNA topoisomerase I (EC 5.99.1.2) |

| | | | | | | |
|---|---|---|---|---|---|---|
| AP53_959 | 0.156504051 | 7.296947342 | 1.268945538 | 0.283089942 | 0.392483984 | Phosphate transport ATP-binding p |
| AP53_1296 | -0.159207037 | 6.172112517 | 1.250652598 | 0.28641137 | 0.396805474 | Multiple sugar ABC transporter, me |
| AP53_661 | 0.182112848 | 8.138897903 | 1.249167222 | 0.286683443 | 0.396899118 | Tripeptide aminopeptidase (EC 3.4. |
| AP53_1269 | 0.143310226 | 7.264681156 | 1.23483594 | 0.289327078 | 0.400273598 | Arginine pathway regulatory protei |
| AP53_1771 | 0.994224979 | 2.941398101 | 1.229402356 | 0.290338284 | 0.401386473 | hypothetical protein |
| AP53_rna77 | 2.099656569 | -1.139188048 | 1.227551057 | 0.290683943 | 0.401578316 | 5S RNA |
| AP53_1162 | -0.30345581 | 5.587893052 | 1.220516707 | 0.292002598 | 0.402846528 | Kup system potassium uptake prote |
| AP53_663 | -0.207359861 | 7.573759383 | 1.22043994 | 0.292017035 | 0.402846528 | Ferredoxin |
| AP53_624 | 0.375680701 | 4.183410825 | 1.213512826 | 0.293323871 | 0.404122914 | Phage-associated homing endonucl |
| AP53_868 | -0.170112869 | 8.622066787 | 1.213328938 | 0.293358673 | 0.404122914 | Na+ driven multidrug efflux pump |
| AP53_1399 | 0.173764377 | 10.44428353 | 1.200861616 | 0.295731817 | 0.406951548 | Aminopeptidase C (EC 3.4.22.40) |
| AP53_1706 | -0.187739594 | 12.43279509 | 1.200341832 | 0.295831342 | 0.406951548 | FIG01114845: hypothetical protei |
| AP53_1364 | -0.207745142 | 8.777863786 | 1.191079932 | 0.297612653 | 0.409112011 | Phage integrase: site-specific recom |
| AP53_338 | -0.321974008 | 12.83272165 | 1.185926761 | 0.298610277 | 0.409802998 | FIG01114323: hypothetical protei |
| AP53_12 | 0.261565707 | 7.130205953 | 1.185404802 | 0.298711588 | 0.409802998 | tRNA(Ile)-lysidine synthetase |
| AP53_209 | -0.126383576 | 7.368054621 | 1.184679585 | 0.29885243 | 0.409802998 | Putative deoxyribonuclease YcfH |
| AP53_756 | -0.146032407 | 8.863382525 | 1.184126854 | 0.298959837 | 0.409802998 | Topoisomerase IV subunit B (EC 5.9 |
| AP53_1774 | -0.348137265 | 10.47176881 | 1.180242217 | 0.299716239 | 0.410549908 | Hypothetical protein possible func |
| AP53_981 | 0.237956179 | 12.5374196 | 1.176552429 | 0.300437198 | 0.41124725 | FIG01114374: hypothetical protei |
| AP53_1115 | -0.293166554 | 5.058997186 | 1.162315616 | 0.303242009 | 0.414794038 | Protein G-related alpha 2 macroglo |
| AP53_1701 | -0.112143723 | 8.428105587 | 1.160484606 | 0.303605418 | 0.414890399 | ABC transporter permease protein |
| AP53_379 | -0.340519604 | 4.715037934 | 1.159807526 | 0.303739957 | 0.414890399 | Bacteriocin BlpU |
| AP53_1236 | 0.178030858 | 7.985314977 | 1.152087089 | 0.305280023 | 0.416403445 | Isoleucyl-tRNA synthetase (EC 6.1.1 |
| AP53_350 | -0.124114907 | 8.629141679 | 1.151575487 | 0.305382466 | 0.416403445 | FIG01115635: hypothetical protei |
| AP53_472 | 0.191614811 | 11.25387969 | 1.151032518 | 0.305491244 | 0.416403445 | Phage shock protein C, putative; str |
| AP53_1856 | 0.122410785 | 7.180055165 | 1.123664903 | 0.311045989 | 0.423677379 | Tryptophanyl-tRNA synthetase (EC 6 |
| AP53_809 | 0.136357444 | 7.618084778 | 1.121403801 | 0.311511318 | 0.424013653 | COG4478, integral membrane prot |
| AP53_1762 | -0.27565939 | 5.868500859 | 1.104984524 | 0.314920296 | 0.428353395 | Transcriptional regulator, HxlR fam |
| AP53_1400 | 0.108425335 | 7.909196084 | 1.097734004 | 0.316442605 | 0.430122617 | NAD synthetase (EC 6.3.1.5) |
| AP53_65 | 0.319994064 | 8.071274492 | 1.090633116 | 0.317943713 | 0.431860565 | Translation initiation factor 1 |
| AP53_250 | -0.090006309 | 8.649442822 | 1.082101531 | 0.319760781 | 0.434024948 | Hydrolase (HAD superfamily), YqeK |
| AP53_1377 | -0.126756533 | 10.53840346 | 1.079241054 | 0.320373341 | 0.434552519 | Serine/threonine protein kinase Prk |
| AP53_1547 | -0.110126929 | 8.074757721 | 1.073787387 | 0.321545895 | 0.43561236 | A/G-specific adenine glycosylase (EC |
| AP53_483 | 0.13590792 | 9.888218567 | 1.073519917 | 0.321603561 | 0.43561236 | Lysyl-tRNA synthetase (class II) (EC 6 |
| AP53_134 | 0.107488637 | 8.862852606 | 1.07061295 | 0.322231248 | 0.436158195 | Nicotine adenine dinucleotide glyc |

| | | | | | | |
|---|---|---|---|---|---|---|
| AP53_427 | -0.171476198 | 4.410117983 | 1.064553115 | 0.323545391 | 0.43763178 | Daunorubicin resistance transmem |
| AP53_1789 | -0.139077551 | 5.668436909 | 1.063314236 | 0.323815005 | 0.437691451 | Mobile element protein |
| AP53_1375 | -0.099364573 | 7.123257277 | 1.060319649 | 0.324468045 | 0.438268945 | Sensor histidine kinase VraS |
| AP53_1339 | 0.395216267 | 4.132509886 | 1.047567853 | 0.327270224 | 0.441746527 | hypothetical protein |
| AP53_1407 | 0.176818215 | 6.653605219 | 1.044195957 | 0.328017027 | 0.442431374 | FIG01116065: hypothetical protei |
| AP53_1282 | -0.375957774 | 8.051671386 | 1.043220733 | 0.328233477 | 0.442431374 | hypothetical protein |
| AP53_160 | -0.307900981 | 4.973827245 | 1.040455669 | 0.328848304 | 0.442952504 | Mobile element protein |
| AP53_1581 | -0.250549128 | 6.347730772 | 1.03797453 | 0.329401417 | 0.443119319 | Ribosomal-protein-S18p-alanine ac |
| AP53_1291 | 0.148344651 | 6.437852788 | 1.03785214 | 0.329428736 | 0.443119319 | Beta-galactosidase (EC 3.2.1.23) |
| AP53_1592 | 0.118386387 | 9.927094547 | 1.034204849 | 0.330244361 | 0.443650139 | FIG001802: Putative alkaline-shocl |
| AP53_382 | 0.34385342 | 5.415796632 | 1.034043544 | 0.3302805 | 0.443650139 | FIG01119121: hypothetical protei |
| AP53_788 | 0.168136181 | 5.911765568 | 1.022447581 | 0.332893582 | 0.44658684 | ABC transporter permease protein |
| AP53_1442 | -0.142193757 | 8.913391452 | 1.022300479 | 0.332926923 | 0.44658684 | Tagatose-6-phosphate kinase (EC 2. |
| AP53_1187 | 0.283246595 | 2.279544779 | 1.021118326 | 0.333195036 | 0.446637821 | Hyaluronate lyase (phage associate |
| AP53_1512 | -0.107885227 | 8.214941548 | 1.018761769 | 0.33373044 | 0.44704678 | ATP-dependent DNA helicase RecG |
| AP53_432 | 0.111448069 | 7.462376129 | 1.016589863 | 0.334224999 | 0.447379856 | FIG01114264: hypothetical protei |
| AP53_862 | -0.205399614 | 5.314573916 | 1.015086294 | 0.334567997 | 0.447379856 | Adenylate cyclase (EC 4.6.1.1) |
| AP53_628 | 0.28925723 | 2.940810817 | 1.014637147 | 0.334670557 | 0.447379856 | Phage integrase: site-specific recom |
| AP53_860 | -0.144723974 | 8.307023261 | 1.000384715 | 0.337948918 | 0.451451376 | Cysteine desulfurase (EC 2.8.1.7) |
| AP53_22 | -0.284139124 | 8.533996792 | 0.996523194 | 0.338845206 | 0.452337377 | Phosphoribosylformylglycinamidir |
| AP53_416 | -0.116347592 | 8.589937643 | 0.992288207 | 0.339832175 | 0.453343128 | hypothetical protein |
| AP53_1188 | 0.289473311 | 2.169798669 | 0.991196697 | 0.340087233 | 0.453371786 | Phage hyaluronidase; Phage endope |
| AP53_1773 | 0.176540057 | 11.49108682 | 0.989859162 | 0.340400163 | 0.4534775 | Putative Holliday junction resolvas |
| AP53_440 | 0.309803935 | 2.590914792 | 0.980064076 | 0.342704714 | 0.456234466 | FIG01114171: hypothetical protei |
| AP53_1174 | -0.479938055 | 3.571063544 | 0.977898003 | 0.343217424 | 0.456603852 | FIG01117945: hypothetical protei |
| AP53_1530 | -0.252730214 | 7.232502317 | 0.973906067 | 0.344165267 | 0.457551221 | Fructokinase (EC 2.7.1.4) |
| AP53_538 | -0.156559287 | 8.313456641 | 0.972873946 | 0.344410958 | 0.457564455 | Chorismate mutase I (EC 5.4.99.5) |
| AP53_426 | -0.186846546 | 3.992919516 | 0.969996827 | 0.345097202 | 0.458162564 | Daunorubicin resistance ATP-bindi |
| AP53_pseudo1615 | 0.270484605 | 3.079259341 | 0.96786171 | 0.345607764 | 0.458526774 | Lanthionine biosynthesis protein L |
| AP53_1352 | -0.437374462 | 4.312188956 | 0.959509982 | 0.347615586 | 0.460875582 | hypothetical phage protein |
| AP53_1740 | -0.203982393 | 6.04185596 | 0.953886695 | 0.348977158 | 0.462364958 | hypothetical protein |
| AP53_1733 | 0.149727135 | 7.51982531 | 0.948142121 | 0.350376232 | 0.463901955 | transcriptional regulator, MutR fan |
| AP53_30 | -0.370907426 | 10.05230692 | 0.944272632 | 0.351323308 | 0.464728852 | Phosphoribosylaminoimidazole car |
| AP53_1570 | -0.149395156 | 7.812768783 | 0.943635441 | 0.351479626 | 0.464728852 | Transcriptional regulatory protein |
| AP53_47 | 0.1806414 | 10.83978006 | 0.941901163 | 0.351905607 | 0.464975346 | SSU ribosomal protein S19p (S15e) |

| | | | | | | |
|---|---|---|---|---|---|---|
| AP53_966 | 0.269070939 | 5.433533808 | 0.930156067 | 0.354810705 | 0.468494952 | FIG006438: hypothetical protein |
| AP53_485 | -0.145398694 | 6.997365124 | 0.920130392 | 0.357318742 | 0.470935088 | Phosphoglycerate mutase family 5 |
| AP53_609 | -0.261476614 | 4.442814656 | 0.920103638 | 0.35732547 | 0.470935088 | Phage tail fibers |
| AP53_906 | 0.135640143 | 5.05535495 | 0.919860602 | 0.357386597 | 0.470935088 | 2-(5''-triphosphoribosyl)-3'-dephos |
| AP53_455 | -0.160803851 | 10.67283373 | 0.91785296 | 0.357892139 | 0.471281304 | Plasmid stabilization system toxin p |
| AP53_643 | -0.250744248 | 4.218836449 | 0.912569362 | 0.359227696 | 0.472719293 | replication initiator protein A |
| AP53_62 | 0.158161819 | 10.56937826 | 0.896981556 | 0.363211494 | 0.477637879 | LSU ribosomal protein L15p (L27Ae |
| AP53_995 | -0.080773799 | 7.959449408 | 0.895251 | 0.363657843 | 0.477901065 | DNA polymerase III alpha subunit (E |
| AP53_505 | -0.147217966 | 9.691599483 | 0.890487891 | 0.364890603 | 0.479196658 | Calcium-transporting ATPase (EC 3. |
| AP53_1760 | -0.271711876 | 4.176515525 | 0.888587676 | 0.365384154 | 0.479469858 | FIG01115695: hypothetical protei |
| AP53_978 | 0.22035349 | 13.63851091 | 0.887785994 | 0.365592679 | 0.479469858 | FIG01114697: hypothetical protei |
| AP53_1030 | 0.241148299 | 4.945597315 | 0.882727766 | 0.366912497 | 0.480875866 | Phage endonuclease |
| AP53_pseudo928 | 0.13302459 | 5.682119527 | 0.881222477 | 0.367306646 | 0.481067612 | UDP-N-acetylmuramoylpentapepti |
| AP53_1558 | 0.11745757 | 7.796647122 | 0.879554746 | 0.367744073 | 0.481315743 | DNA polymerase IV (EC 2.7.7.7) |
| AP53_1457 | -0.107912581 | 10.3336213 | 0.87521402 | 0.368886277 | 0.482485353 | Transcription termination protein |
| AP53_456 | -0.15445262 | 10.88979319 | 0.872088515 | 0.369712022 | 0.483239754 | hypothetical protein |
| AP53_rna33 | 0.528658784 | 4.405899646 | 0.868157701 | 0.370754486 | 0.484276216 | tRNA-Ile-GAT |
| AP53_1227 | 0.078648903 | 7.515049606 | 0.860640333 | 0.372760507 | 0.486569028 | Methylenetetrahydrofolate dehydr |
| AP53_491 | 0.113034792 | 7.708053578 | 0.85687827 | 0.373770583 | 0.487559611 | Glutathione peroxidase family prot |
| AP53_251 | -0.154317281 | 7.811421738 | 0.848499737 | 0.376035082 | 0.490184079 | Iojap protein |
| AP53_972 | -0.104642065 | 8.01282363 | 0.844878223 | 0.377020327 | 0.490866786 | Type I restriction-modification syst |
| AP53_1177 | -0.175964791 | 10.37121398 | 0.844715846 | 0.377064594 | 0.490866786 | Lead, cadmium, zinc and mercury t |
| AP53_568 | 0.109831438 | 9.895918157 | 0.838976371 | 0.378634371 | 0.492381603 | UDP-N-acetylglucosamine 1-carbox |
| AP53_653 | 0.169368034 | 7.969931946 | 0.83860762 | 0.378735566 | 0.492381603 | Phage protein |
| AP53_450 | 0.293311769 | 1.699702403 | 0.836869365 | 0.379213143 | 0.492672497 | FIG01116026: hypothetical protei |
| AP53_1675 | 0.148663271 | 5.85093154 | 0.832720639 | 0.3803567 | 0.493827661 | FIG01115320: hypothetical protei |
| AP53_1353 | -0.270477785 | 4.458065608 | 0.830122781 | 0.381075452 | 0.494430116 | DNA replication protein DnaC |
| AP53_357 | -0.109995194 | 9.9521086 | 0.829178586 | 0.381337197 | 0.494439212 | Peptidyl-prolyl cis-trans isomerase |
| AP53_604 | -0.337667267 | 6.321387552 | 0.824659486 | 0.382593769 | 0.49573732 | Phage protein |
| AP53_1100 | 0.180791866 | 6.426479153 | 0.821620317 | 0.383442396 | 0.496505464 | Mobile element protein |
| AP53_rna59 | 0.45048792 | 3.594400094 | 0.818110108 | 0.384426139 | 0.497447424 | tRNA-Gln-TTG |
| AP53_1345 | -0.23365931 | 4.364938321 | 0.816313509 | 0.384931136 | 0.497471452 | FIG01114622: hypothetical protei |
| AP53_rna72 | 1.100576378 | 5.80827402 | 0.82142675 | 0.384957301 | 0.497471452 | Small Subunit Ribosomal RNA; ssuR |
| AP53_1446 | 0.09061526 | 8.608058006 | 0.81448988 | 0.385444771 | 0.497769993 | PTS system, galactose-specific IIB co |
| AP53_1736 | -0.119101041 | 13.53520806 | 0.809902765 | 0.386741413 | 0.499112422 | Heat shock protein 60 family chape |

| | | | | | | |
|---|---|---|---|---|---|---|
| AP53_945 | 0.107179792 | 7.848707534 | 0.80584612 | 0.387893695 | 0.500266885 | Substrate-specific component PanT |
| AP53_205 | -0.115296649 | 6.799939506 | 0.796926081 | 0.390446065 | 0.503026193 | FIG01114161: hypothetical protei |
| AP53_1346 | -0.252960285 | 4.250518521 | 0.796559518 | 0.390551506 | 0.503026193 | Phage Holliday junction resolvase |
| AP53_1445 | 0.103385161 | 8.853964934 | 0.793706232 | 0.39137375 | 0.50375096 | PTS system, galactose-specific IIC cc |
| AP53_960 | 0.123762548 | 7.539641314 | 0.791531831 | 0.392002152 | 0.50406351 | Phosphate transport ATP-binding p |
| AP53_1602 | 0.210510857 | 5.230470574 | 0.791069519 | 0.392135961 | 0.50406351 | hypothetical protein |
| AP53_673 | 0.110113718 | 9.971696134 | 0.786936421 | 0.393335366 | 0.505270645 | FIG01116936: hypothetical protei |
| AP53_1859 | -0.11739651 | 8.358397793 | 0.78163355 | 0.394882565 | 0.506922658 | ABC transporter ATP-binding protei |
| AP53_183 | 0.130032917 | 9.708970245 | 0.779682044 | 0.395454324 | 0.507321112 | Glycerol-3-phosphate dehydrogena |
| AP53_641 | -0.181363932 | 4.573462804 | 0.775257638 | 0.396755366 | 0.508654007 | hypothetical phage protein |
| AP53_1712 | -0.187690163 | 5.106717319 | 0.772068097 | 0.397697413 | 0.5095252 | RopB; Rgg-like transcription regula |
| AP53_rna80 | 1.138484328 | 5.803466699 | 0.775549483 | 0.398086313 | 0.509687027 | Small Subunit Ribosomal RNA; ssuF |
| AP53_rna35 | 1.125931694 | 5.821934084 | 0.76701861 | 0.40060455 | 0.512573125 | Small Subunit Ribosomal RNA; ssuF |
| AP53_1005 | -0.12868283 | 7.959082217 | 0.761223478 | 0.400926604 | 0.512647258 | Phage protein |
| AP53_1096 | -0.142066068 | 5.522855 | 0.759171733 | 0.401542148 | 0.512820277 | Quinolinate phosphoribosyltransfe |
| AP53_rna51 | 1.094990253 | 5.804361528 | 0.763702028 | 0.401590325 | 0.512820277 | Small Subunit Ribosomal RNA; ssuF |
| AP53_rna14 | 1.118960403 | 5.834101851 | 0.760277886 | 0.402612075 | 0.513787007 | Small Subunit Ribosomal RNA; ssuF |
| AP53_1834 | 0.158903119 | 7.62930424 | 0.742032296 | 0.406742455 | 0.518716889 | Phosphoesterase, DHH family prote |
| AP53_1635 | 0.133898818 | 7.260475475 | 0.736648521 | 0.408397796 | 0.520485963 | Hypothetical protein DUF901, simi |
| AP53_584 | 0.259559809 | 10.15155159 | 0.73570592 | 0.408688707 | 0.520514948 | Large-conductance mechanosensiti |
| AP53_356 | 0.128986461 | 8.106574505 | 0.734118066 | 0.409179499 | 0.5207983 | Manganese ABC transporter, inner r |
| AP53_654 | 0.120441822 | 8.648977839 | 0.73240308 | 0.40971063 | 0.520836642 | Pathogenicity island SaPIn2 |
| AP53_844 | -0.119052389 | 7.79716985 | 0.732288058 | 0.409746291 | 0.520836642 | ABC transporter, periplasmic sperm |
| AP53_1196 | -0.317681043 | 3.612372154 | 0.7192886 | 0.413808451 | 0.525655892 | FIG042840: Phage protein |
| AP53_1448 | 0.098920794 | 8.01963308 | 0.715757835 | 0.414922788 | 0.526726704 | Lactose phosphotransferase system |
| AP53_744 | 0.097181361 | 8.440934461 | 0.712567691 | 0.415933725 | 0.527664941 | Histidine protein kinase |
| AP53_1378 | 0.100040178 | 10.25019188 | 0.710678172 | 0.416534349 | 0.528081758 | Protein serine/threonine phosphat |
| AP53_1761 | -0.291440495 | 4.562062021 | 0.707103223 | 0.417674506 | 0.529181604 | FIG01114717: hypothetical protei |
| AP53_1607 | 0.12774246 | 5.26947702 | 0.704006538 | 0.418666155 | 0.530091981 | Glycerate kinase (EC 2.7.1.31) |
| AP53_494 | 0.113330429 | 7.071482449 | 0.701240407 | 0.419555131 | 0.530828335 | Cell division protein FtsW |
| AP53_410 | -0.162674369 | 9.975201927 | 0.700496596 | 0.41979469 | 0.530828335 | Preprotein translocase subunit Sec |
| AP53_1500 | 0.135421386 | 8.961250047 | 0.697027642 | 0.420914828 | 0.53189823 | Aspartyl-tRNA(Asn) amidotransfera |
| AP53_977 | 0.167791399 | 13.251696 | 0.695358797 | 0.421455409 | 0.532234839 | General stress protein, Gls24 family |
| AP53_1557 | 0.0845366 | 9.093260226 | 0.692941363 | 0.422240449 | 0.532879526 | Putative membrane protein precur |
| AP53_403 | 0.227023872 | 4.275269101 | 0.681854719 | 0.425870972 | 0.537112123 | hypothetical protein |

| | | | | | | |
|---|---|---|---|---|---|---|
| AP53_1192 | -0.336990062 | 2.102695241 | 0.680231602 | 0.426406702 | 0.537280469 | conserved hypothetical protein - ph |
| AP53_1686 | -0.437338658 | 7.500481105 | 0.67977361 | 0.426558064 | 0.537280469 | FIG01115696: hypothetical protein |
| AP53_1199 | -0.307958191 | 2.890927429 | 0.674411299 | 0.428336726 | 0.53896251 | FIG01115109: hypothetical protein |
| AP53_968 | -0.079967118 | 7.568667338 | 0.674074574 | 0.428448816 | 0.53896251 | Riboflavin kinase (EC 2.7.1.26) / FM |
| AP53_587 | -0.09279989 | 9.35155574 | 0.672398427 | 0.429007482 | 0.539036682 | probably aromatic ring hydroxylati |
| AP53_1426 | 0.164633519 | 6.450441917 | 0.672231448 | 0.429063202 | 0.539036682 | FIG01116532: hypothetical protein |
| AP53_1179 | 0.085202035 | 7.530511432 | 0.670671204 | 0.429584405 | 0.539342387 | Streptococcal extracellular nucleas |
| AP53_1667 | -0.113023661 | 10.53558133 | 0.667666091 | 0.430591162 | 0.540256913 | GTP pyrophosphokinase (EC 2.7.6.5 |
| AP53_1546 | 0.163959609 | 9.595490849 | 0.660948972 | 0.432855371 | 0.542536021 | hypothetical protein |
| AP53_190 | -0.091928176 | 8.865796182 | 0.660620272 | 0.432966665 | 0.542536021 | Carbonic anhydrase (EC 4.2.1.1) |
| AP53_554 | 0.249743615 | 2.74527286 | 0.659491038 | 0.433349366 | 0.542665238 | Asparagine-rich ORFan next to Sag c |
| AP53_1735 | -0.160961608 | 7.533997899 | 0.654929267 | 0.434900976 | 0.544057335 | hypothetical protein |
| AP53_127 | -0.10825882 | 6.639887807 | 0.654575652 | 0.43502163 | 0.544057335 | V-type ATP synthase subunit B (EC 3 |
| AP53_32 | -0.221625593 | 5.687921236 | 0.649517447 | 0.436753487 | 0.545520325 | hypothetical protein |
| AP53_rna1 | 1.039018507 | 5.84424604 | 0.65313167 | 0.436805277 | 0.545520325 | Small Subunit Ribosomal RNA; ssuR |
| AP53_1620 | 0.140720893 | 6.896683767 | 0.648699573 | 0.437034573 | 0.545520325 | Tagatose 1,6-diphosphate aldolase |
| AP53_428 | -0.125694991 | 4.324315563 | 0.643137425 | 0.43895403 | 0.547460392 | ABC-type multidrug transport syste |
| AP53_1697 | 0.107757237 | 8.455477189 | 0.642527669 | 0.439165291 | 0.547460392 | Hypothetical mga-associated prote |
| AP53_282 | -0.097041866 | 8.804343559 | 0.641749982 | 0.439434977 | 0.547460392 | FIG005986: HD family hydrolase |
| AP53_810 | -0.123259507 | 7.328800618 | 0.634911653 | 0.441818115 | 0.550076306 | Translation elongation factor LepA |
| AP53_1404 | -0.138947539 | 6.130712776 | 0.628126448 | 0.444203815 | 0.552692055 | Hypothetical protein SPy1656 |
| AP53_1655 | -0.13168644 | 9.654728026 | 0.626568793 | 0.444754489 | 0.552764426 | Prolyl-tRNA synthetase (EC 6.1.1.15 |
| AP53_1246 | 0.080232355 | 8.444518672 | 0.626351079 | 0.444831547 | 0.552764426 | UDP-N-acetylmuramoylalanine--D-g |
| AP53_608 | -0.232127575 | 5.182205261 | 0.624458043 | 0.445502498 | 0.553243985 | Phage endopeptidase |
| AP53_1863 | -0.091930448 | 10.1139576 | 0.621052146 | 0.446713875 | 0.554393626 | Serine protease, DegP/HtrA, do-like |
| AP53_1788 | -0.206194556 | 7.228241139 | 0.614007636 | 0.449236778 | 0.557061965 | FIG01118151: hypothetical protein |
| AP53_rna56 | 0.83607418 | 0.251749348 | 0.613448778 | 0.449437938 | 0.557061965 | tRNA-Arg-ACG |
| AP53_600 | 0.205305473 | 10.95078582 | 0.609064847 | 0.451021135 | 0.55780093 | Phage protein |
| AP53_1158 | -0.19844632 | 3.822656869 | 0.609059854 | 0.451022943 | 0.55780093 | Late competence protein ComEA, D |
| AP53_237 | -0.096832469 | 6.217745756 | 0.608762788 | 0.451130561 | 0.55780093 | D-alanyl-D-alanine carboxypeptidas |
| AP53_693 | 0.127229722 | 9.317769129 | 0.608616291 | 0.451183648 | 0.55780093 | ABC transporter permease protein |
| AP53_678 | -0.080647524 | 8.376474821 | 0.607318915 | 0.451654241 | 0.558027295 | tRNA S(4)U 4-thiouridine synthase ( |
| AP53_1785 | -0.254832232 | 5.049146662 | 0.605994187 | 0.452135599 | 0.558266665 | conserved hypothetical protein - ph |
| AP53_1429 | 0.13643293 | 7.988638528 | 0.602455608 | 0.453425597 | 0.559503549 | Glycyl-tRNA synthetase beta chain ( |
| AP53_158 | -0.195443421 | 4.055778573 | 0.59438946 | 0.456389244 | 0.562802747 | Mobile element protein |

| | | | | | | |
|---|---|---|---|---|---|---|
| AP53_1447 | -0.092965958 | 8.138222723 | 0.588462562 | 0.458587663 | 0.564707079 | PTS system, galactose-specific IIA co |
| AP53_1056 | -0.097954845 | 7.167086984 | 0.588129106 | 0.458711878 | 0.564707079 | hypothetical protein |
| AP53_671 | 0.074445275 | 8.008832027 | 0.587875683 | 0.458806318 | 0.564707079 | 3-dehydroquinate dehydratase I (EC |
| AP53_pseudo399 | -0.161546715 | 4.327962628 | 0.583330017 | 0.460505876 | 0.566439737 | FIG01114499: hypothetical protei |
| AP53_1300 | -0.142062221 | 9.157013082 | 0.580635345 | 0.461518396 | 0.567145615 | Beta-glucosidase (EC 3.2.1.21) |
| AP53_650 | -0.172489295 | 4.173284571 | 0.5796537 | 0.461888183 | 0.567145615 | conserved domain protein |
| AP53_657 | -0.1210563 | 7.849295204 | 0.578874799 | 0.462181953 | 0.567145615 | Glycosyl transferase, group 2 family |
| AP53_1092 | -0.192598023 | 6.259530038 | 0.578698438 | 0.462248512 | 0.567145615 | FIG01114947: hypothetical protei |
| AP53_393 | -0.15479003 | 3.336583387 | 0.577274177 | 0.462786635 | 0.567447163 | Competence-stimulating peptide A |
| AP53_790 | 0.107680547 | 9.013399893 | 0.575512222 | 0.463453817 | 0.567906476 | Ribonuclease J2 (endoribonuclease |
| AP53_1280 | -0.115004658 | 9.75997479 | 0.573264356 | 0.464307364 | 0.568280144 | Valyl-tRNA synthetase (EC 6.1.1.9) |
| AP53_204 | -0.096104356 | 6.544829362 | 0.572133558 | 0.464737753 | 0.568280144 | ABC transporter, predicted N-acety |
| AP53_1649 | -0.166835844 | 7.311415974 | 0.571751823 | 0.464883196 | 0.568280144 | FIG01114948: hypothetical protei |
| AP53_1507 | 0.125729585 | 8.566607648 | 0.571629377 | 0.464929865 | 0.568280144 | Universal stress protein family |
| AP53_448 | -0.338908255 | 1.098269493 | 0.569214107 | 0.465852053 | 0.568993674 | FIG01115329: hypothetical protei |
| AP53_1685 | 0.120513218 | 6.368726164 | 0.568260651 | 0.466216954 | 0.568993674 | Dipeptide transport ATP-binding pr |
| AP53_1595 | 0.087480234 | 9.256043471 | 0.567800947 | 0.466393063 | 0.568993674 | Hydrolases of the alpha/beta superf |
| AP53_811 | 0.238750935 | 11.91569231 | 0.557438864 | 0.470393002 | 0.573513076 | Collagen-like surface protein |
| AP53_409 | -0.093337361 | 9.444312881 | 0.551959045 | 0.472532095 | 0.575759445 | Multidrug resistance efflux pump P |
| AP53_1444 | 0.073168924 | 9.09676772 | 0.548329774 | 0.473958025 | 0.577134584 | Galactose-6-phosphate isomerase, |
| AP53_rna70 | 0.934680411 | 5.783091751 | 0.543252797 | 0.477128827 | 0.580631382 | Large Subunit Ribosomal RNA; lsuR |
| AP53_1194 | 0.396031925 | 1.506943889 | 0.538355696 | 0.477915259 | 0.581224009 | FIG018217: Phage-associated prote |
| AP53_611 | 0.245469115 | 4.925121625 | 0.536037615 | 0.478843152 | 0.581987825 | hypothetical protein |
| AP53_rna62 | 1.26682465 | 0.675720235 | 0.532087421 | 0.480431572 | 0.583552992 | tRNA-Glu-TTC |
| AP53_944 | -0.091620482 | 7.544989533 | 0.527718095 | 0.482199231 | 0.585333776 | Phosphopantothenoylcysteine dec |
| AP53_84 | 0.144005547 | 3.193047419 | 0.521563031 | 0.484708649 | 0.58801218 | Late competence protein ComGB, a |
| AP53_976 | -0.070707392 | 9.357629102 | 0.515559105 | 0.487178552 | 0.590639331 | Transcriptional regulator, GntR fam |
| AP53_rna3 | 0.908223393 | 5.832630068 | 0.50912315 | 0.490972259 | 0.594867138 | Large Subunit Ribosomal RNA; lsuR |
| AP53_1123 | -0.09509933 | 5.808376608 | 0.505205418 | 0.491490221 | 0.595123218 | FIG01115656: hypothetical protei |
| AP53_177 | 0.110417068 | 8.91860337 | 0.499533906 | 0.493880736 | 0.597359177 | Glucose-6-phosphate isomerase (EC |
| AP53_1619 | 0.151795539 | 6.430822598 | 0.49804793 | 0.494510481 | 0.597359177 | PTS system, lactose-specific IIA com |
| AP53_785 | -0.078134273 | 6.737096775 | 0.497978674 | 0.494539866 | 0.597359177 | Fibronectin/fibrinogen-binding pro |
| AP53_1664 | -0.09507131 | 7.418852833 | 0.497912726 | 0.49456785 | 0.597359177 | Leucine rich protein |
| AP53_598 | 0.205223162 | 9.372826546 | 0.496833257 | 0.49502631 | 0.597541087 | conserved hypothetical protein |
| AP53_1185 | 0.265744342 | 1.757692781 | 0.490638246 | 0.49767209 | 0.600361421 | FIG01116522: hypothetical protei |

| | | | | | | |
|---|---|---|---|---|---|---|
| AP53_1172 | -0.113025515 | 7.299821759 | 0.487930937 | 0.498836273 | 0.601392053 | Transcriptional repressor for NAD b |
| AP53_1857 | 0.230139653 | 7.917900341 | 0.486838981 | 0.49930721 | 0.601586154 | FIG01116071: hypothetical protei |
| AP53_rna78 | 0.87387889 | 5.81260375 | 0.482856966 | 0.502119315 | 0.604599001 | Large Subunit Ribosomal RNA; lsuR |
| AP53_907 | -0.133942787 | 4.789085088 | 0.479241019 | 0.502606257 | 0.604810133 | Transcriptional regulator, GntR fam |
| AP53_rna53 | 0.862978759 | 5.811675989 | 0.478216977 | 0.504135872 | 0.606274924 | Large Subunit Ribosomal RNA; lsuR |
| AP53_1508 | -0.074265006 | 8.636400623 | 0.466676917 | 0.508148642 | 0.6107223 | Hydrolase (HAD superfamily) |
| AP53_537 | -0.141640925 | 8.498753398 | 0.464968723 | 0.508910737 | 0.611076159 | Flavodoxin |
| AP53_200 | -0.204216849 | 10.29106243 | 0.464606247 | 0.50907272 | 0.611076159 | LSU ribosomal protein L34p |
| AP53_1475 | -0.095452307 | 5.649180463 | 0.463273934 | 0.50966891 | 0.611413692 | Mobile element protein |
| AP53_890 | -0.096960352 | 6.723217123 | 0.460293572 | 0.511007197 | 0.612306604 | FIG01280259: hypothetical protei |
| AP53_791 | -0.06994565 | 8.265255128 | 0.459517748 | 0.511356622 | 0.612306604 | Tributyrin esterase |
| AP53_768 | -0.067567104 | 8.452878255 | 0.459511117 | 0.511359611 | 0.612306604 | GTP-binding protein HflX |
| AP53_rna16 | 0.859008902 | 5.830256733 | 0.45702803 | 0.513535148 | 0.614532505 | Large Subunit Ribosomal RNA; lsuR |
| AP53_rna37 | 0.842600518 | 5.803484641 | 0.45348421 | 0.515138595 | 0.616071481 | Large Subunit Ribosomal RNA; lsuR |
| AP53_610 | 0.193088457 | 4.371625699 | 0.447260185 | 0.516936022 | 0.617840406 | Phage tail length tape-measure pro |
| AP53_793 | 0.070700697 | 7.110767067 | 0.446119744 | 0.517460826 | 0.618087055 | ABC transporter ATP-binding protei |
| AP53_585 | -0.074805183 | 9.096517146 | 0.441598254 | 0.519551236 | 0.619834025 | DNA primase (EC 2.7.7.-) |
| AP53_1217 | -0.072397452 | 8.364502351 | 0.44157493 | 0.51956206 | 0.619834025 | YfaA |
| AP53_277 | -0.061542247 | 9.564369913 | 0.436836713 | 0.521769616 | 0.622085273 | FIG004453: protein YceG like |
| AP53_626 | 0.218941483 | 3.253685203 | 0.434848807 | 0.522700991 | 0.622813152 | hypothetical phage protein |
| AP53_1312 | -0.179297552 | 6.203251073 | 0.433354 | 0.523403383 | 0.623267465 | Phage lysin |
| AP53_1211 | 0.171303947 | 1.877716067 | 0.425573249 | 0.527088146 | 0.627270442 | Phage excisionase |
| AP53_531 | -0.067805438 | 7.100928641 | 0.419312385 | 0.530088675 | 0.630237128 | FIG01117156: hypothetical protei |
| AP53_499 | 0.072270265 | 7.651709629 | 0.418745272 | 0.530362057 | 0.630237128 | tRNA-dependent lipid II-Ala--L-alani |
| AP53_1841 | 0.102473868 | 7.681483501 | 0.418345129 | 0.530555109 | 0.630237128 | Transmembrane component of gen |
| AP53_336 | 0.085089173 | 10.01169484 | 0.41595536 | 0.531710847 | 0.631223703 | C3 family ADP-ribosyltransferase (E |
| AP53_398 | -0.25828522 | 2.694450071 | 0.414540989 | 0.532397113 | 0.631652076 | unknown |
| AP53_629 | 0.208349715 | 3.905552694 | 0.41101879 | 0.534113443 | 0.633301279 | putative transcriptional activator - |
| AP53_337 | -0.083432233 | 5.666366932 | 0.40903037 | 0.535087035 | 0.634068336 | Mobile element protein |
| AP53_900 | 0.084625649 | 5.716788735 | 0.404214085 | 0.537459327 | 0.636357337 | Mobile element protein |
| AP53_1854 | 0.066006868 | 9.006424022 | 0.403779424 | 0.537674411 | 0.636357337 | Putative sugar uptake protein SpyM |
| AP53_201 | 0.11573359 | 6.70652531 | 0.401996256 | 0.538558511 | 0.637015277 | N-acetylmannosamine-6-phosphate |
| AP53_226 | 0.055406413 | 7.602630791 | 0.401325563 | 0.538891765 | 0.637021264 | FIG01114729: hypothetical protei |
| AP53_1628 | -0.266267819 | 2.437120727 | 0.394344243 | 0.542384336 | 0.640759584 | Integrase |
| AP53_1502 | -0.07957217 | 5.633125602 | 0.390987495 | 0.544079228 | 0.642370913 | Mobile element protein |

| | | | | | | |
|---|---|---|---|---|---|---|
| AP53_367 | -0.100561785 | 10.43533034 | 0.385092711 | 0.547080626 | 0.645521882 | Group B streptococcal surface imm |
| AP53_1822 | -0.058011845 | 7.315602649 | 0.37934919 | 0.550036218 | 0.648615005 | FIG028593: membrane protein |
| AP53_1368 | -0.129568867 | 3.964798628 | 0.371092168 | 0.554340574 | 0.653293901 | Competence protein F homolog, ph |
| AP53_1501 | -0.074865792 | 8.011588379 | 0.361283035 | 0.55954146 | 0.659023043 | Aspartyl-tRNA(Asn) amidotransfera |
| AP53_459 | -0.143718135 | 4.324628536 | 0.360262074 | 0.560088373 | 0.659267151 | Mobile element protein |
| AP53_31 | 0.095604538 | 7.975795889 | 0.355316806 | 0.5627527 | 0.662001813 | FIG01114657: hypothetical protei |
| AP53_1295 | -0.10746492 | 7.199351983 | 0.353205071 | 0.5638982 | 0.66294755 | Multiple sugar ABC transporter, sub |
| AP53_391 | -0.11519155 | 5.127292272 | 0.351902481 | 0.564607128 | 0.663379199 | Histidine kinase of the competence |
| AP53_1523 | -0.111104956 | 4.841995829 | 0.350723177 | 0.565250513 | 0.663733361 | hypothetical protein |
| AP53_694 | 0.065414898 | 8.383306483 | 0.345408634 | 0.568168443 | 0.666553151 | Glycerophosphoryl diester phosph |
| AP53_497 | 0.097480718 | 10.59102922 | 0.345100205 | 0.568338724 | 0.666553151 | Triosephosphate isomerase (EC 5.3. |
| AP53_1054 | 0.187994168 | 4.635450138 | 0.338869389 | 0.571801123 | 0.670208925 | Phage transcriptional regulator, Cr |
| AP53_203 | -0.089345194 | 6.238333279 | 0.334245069 | 0.574398867 | 0.672847436 | ABC transporter, predicted N-acety |
| AP53_1412 | -0.068758904 | 8.859933775 | 0.332755924 | 0.575240582 | 0.673427002 | rRNA small subunit methyltransfera |
| AP53_1034 | 0.107516635 | 6.332954571 | 0.332017997 | 0.575658627 | 0.673510184 | putative methyltransferase |
| AP53_55 | 0.102352904 | 10.94485035 | 0.326703003 | 0.578688313 | 0.676296584 | LSU ribosomal protein L5p (L11e) |
| AP53_526 | 0.060578154 | 8.990784056 | 0.326299949 | 0.578919415 | 0.676296584 | Aspartate aminotransferase (EC 2.6. |
| AP53_789 | 0.130061565 | 5.972167453 | 0.326010524 | 0.579085483 | 0.676296584 | ABC transporter ATP-binding protei |
| AP53_925 | 0.08191201 | 5.40964687 | 0.324651812 | 0.579866419 | 0.67666654 | FIG01114068: hypothetical protei |
| AP53_672 | 0.052818967 | 8.457220619 | 0.324247031 | 0.580099497 | 0.67666654 | Chorismate synthase (EC 4.2.3.5) |
| AP53_1037 | 0.13052423 | 5.399904477 | 0.321661424 | 0.581592945 | 0.678001145 | FIG01114622: hypothetical protei |
| AP53_507 | -0.100384795 | 4.440330087 | 0.318664982 | 0.583333789 | 0.67962238 | Heparinase II/III-like |
| AP53_1202 | 0.130649351 | 3.55737465 | 0.316553914 | 0.58456684 | 0.680650412 | Phage protein |
| AP53_1459 | -0.080556339 | 5.692326177 | 0.308281348 | 0.589452268 | 0.685927369 | Mobile element protein |
| AP53_59 | 0.116296321 | 11.0043086 | 0.307620781 | 0.589846109 | 0.685974414 | LSU ribosomal protein L18p (L5e) |
| AP53_859 | 0.100999606 | 7.980890273 | 0.303901849 | 0.592073919 | 0.687813652 | FIG01114333: hypothetical protei |
| AP53_1627 | 0.211875325 | 2.557416665 | 0.303798099 | 0.592136328 | 0.687813652 | hypothetical protein |
| AP53_1576 | 0.073374096 | 8.093332469 | 0.301243036 | 0.593677742 | 0.689191685 | Nucleoside permease NupC |
| AP53_699 | 0.11190331 | 5.236852529 | 0.299826418 | 0.594536077 | 0.689775569 | Cobalt-zinc-cadmium resistance pr |
| AP53_1503 | -0.097009486 | 7.92742632 | 0.29911768 | 0.594966508 | 0.6898626 | Pyruvate,phosphate dikinase (EC 2. |
| AP53_1294 | -0.107892469 | 5.511894944 | 0.298530013 | 0.59532392 | 0.689864913 | Integral membrane protein |
| AP53_508 | -0.074034798 | 5.896771962 | 0.295313927 | 0.597288144 | 0.691728095 | PTS system, N-acetylgalactosamine- |
| AP53_1200 | 0.117943276 | 3.627071698 | 0.293732839 | 0.598258947 | 0.692439247 | hypothetical protein |
| AP53_921 | -0.086069247 | 7.962211143 | 0.292579889 | 0.598969033 | 0.692847969 | Pleiotropic regulator of exopolysac |
| AP53_1708 | -0.129023103 | 6.4287874 | 0.290288376 | 0.600385802 | 0.694073164 | Streptopain inhibitor |

| | | | | | | |
|---|---|---|---|---|---|---|
| AP53_1434 | -0.047522546 | 7.360167075 | 0.287558126 | 0.602083409 | 0.695309085 | Sodium-dependent phosphate tran |
| AP53_1245 | -0.054492574 | 8.936590676 | 0.287417165 | 0.60217134 | 0.695309085 | UDP-N-acetylglucosamine--N-acety |
| AP53_1593 | -0.169698605 | 8.927438765 | 0.285520171 | 0.603357431 | 0.696264431 | LSU ribosomal protein L28p |
| AP53_670 | 0.056967367 | 8.14152311 | 0.283417114 | 0.604678378 | 0.697374172 | LSU m5C1962 methyltransferase Rl |
| AP53_1193 | -0.191198686 | 2.503623599 | 0.280709314 | 0.60638859 | 0.698931267 | Phage major tail protein |
| AP53_1584 | -0.049112234 | 9.405547829 | 0.276221963 | 0.609246462 | 0.701808536 | Ribonuclease J1 (endonuclease and |
| AP53_1317 | 0.102845695 | 3.704008296 | 0.272953561 | 0.611346977 | 0.703810487 | Hyaluronate lyase (phage associate |
| AP53_1190 | 0.150080698 | 2.186103157 | 0.271476476 | 0.612301586 | 0.704219908 | Phage minor tail protein |
| AP53_660 | -0.085435184 | 7.324931732 | 0.271280897 | 0.612428235 | 0.704219908 | membrane protein, related to Actir |
| AP53_814 | -0.074341955 | 5.701081 | 0.270564209 | 0.612892838 | 0.704336884 | Mobile element protein |
| AP53_973 | 0.063747997 | 5.879572272 | 0.269159039 | 0.613806061 | 0.704968973 | ABC transporter permease protein |
| AP53_1575 | -0.070273139 | 8.329307005 | 0.265893672 | 0.615940131 | 0.707001653 | Deoxyribose-phosphate aldolase (E |
| AP53_1059 | 0.049249434 | 6.906797958 | 0.264733601 | 0.616702339 | 0.707122246 | Pleiotropic regulator of exopolysac |
| AP53_25 | -0.135645142 | 9.107144995 | 0.264624992 | 0.616773808 | 0.707122246 | Phosphoribosylglycinamide formyl |
| AP53_1727 | -0.056349279 | 9.545110216 | 0.260901096 | 0.619235737 | 0.709486186 | Multimodular transpeptidase-trans |
| AP53_1376 | -0.070003825 | 6.677968156 | 0.260402961 | 0.619566762 | 0.709486186 | Transporter associated with VraSR |
| AP53_110 | 0.077039194 | 8.013028847 | 0.258914236 | 0.620558479 | 0.710202835 | FIG01114239: hypothetical protei |
| AP53_1036 | -0.141795159 | 5.48739783 | 0.250762335 | 0.626054271 | 0.716070324 | conserved hypothetical protein - pl |
| AP53_606 | -0.127362237 | 5.885791463 | 0.247740409 | 0.628120342 | 0.718010356 | Phage hyaluronidase |
| AP53_708 | 0.086543783 | 9.580698848 | 0.245891357 | 0.629392393 | 0.718526496 | PTS system, fructose-specific IIA cor |
| AP53_1209 | 0.186882742 | 2.866562843 | 0.245794146 | 0.629459436 | 0.718526496 | DNA replication protein dnaD |
| AP53_1124 | -0.102351458 | 4.615390786 | 0.245471003 | 0.629682416 | 0.718526496 | FIG01113965: hypothetical protei |
| AP53_524 | -0.066527932 | 5.69811416 | 0.243836281 | 0.630813278 | 0.719393991 | Mobile element protein |
| AP53_1513 | -0.055203987 | 7.803939915 | 0.242026094 | 0.632071107 | 0.720256386 | Duplicated ATPase component BL0 |
| AP53_627 | 0.166417792 | 3.230233061 | 0.241574198 | 0.632386035 | 0.720256386 | hypothetical phage protein |
| AP53_9 | 0.105472093 | 7.0436227 | 0.240139461 | 0.633388368 | 0.720256386 | Cell division protein DivIC (FtsB), st |
| AP53_278 | -0.051561255 | 9.52934866 | 0.240095701 | 0.633418999 | 0.720256386 | Transcription elongation factor Gre |
| AP53_1527 | -0.056272033 | 10.56430151 | 0.24008733 | 0.633424859 | 0.720256386 | Protein export cytoplasm protein S |
| AP53_375 | -0.200730224 | 2.887972822 | 0.2391853 | 0.634057047 | 0.720553119 | FIG01115781: hypothetical protei |
| AP53_542 | 0.04512833 | 9.748023883 | 0.236599381 | 0.635877716 | 0.722199325 | DNA gyrase subunit B (EC 5.99.1.3) |
| AP53_777 | -0.045006443 | 8.043231212 | 0.234355681 | 0.637467561 | 0.7235816 | FIG137478: Hypothetical protein |
| AP53_136 | 0.051661766 | 9.356913988 | 0.232052252 | 0.639109654 | 0.72465437 | Thiol-activated cytolysin |
| AP53_100 | 0.066546241 | 9.169923012 | 0.231379024 | 0.639591512 | 0.72465437 | tRNA dihydrouridine synthase B (EC |
| AP53_446 | 0.093286777 | 3.672188595 | 0.230519873 | 0.640207713 | 0.72465437 | UDP-glucose dehydrogenase (EC 1.1 |
| AP53_1169 | -0.092682332 | 6.773293348 | 0.230190074 | 0.640444632 | 0.72465437 | Formate/nitrite family of transport |

| | | | | | | |
|---|---|---|---|---|---|---|
| AP53_1587 | -0.085867582 | 5.648149901 | 0.230020732 | 0.640566365 | 0.72465437 | FIG01114549: hypothetical protei |
| AP53_846 | -0.066717391 | 5.626682958 | 0.229900672 | 0.640652704 | 0.72465437 | Two-component sensor histidine ki |
| AP53_1393 | 0.051130933 | 10.5883986 | 0.225305594 | 0.643978506 | 0.727992009 | Hypothetical protein SPy1643 |
| AP53_228 | 0.070760095 | 8.054936901 | 0.223399199 | 0.645370661 | 0.729118259 | Negative regulator of genetic comp |
| AP53_1323 | 0.172823757 | 2.327562017 | 0.222782075 | 0.645822894 | 0.729118259 | Phage major tail shaft protein |
| AP53_rna17 | 0.757511737 | -0.659752514 | 0.222402136 | 0.646101703 | 0.729118259 | 5S RNA |
| AP53_11 | 0.055621599 | 7.488083093 | 0.220933598 | 0.64718213 | 0.72991314 | FIG01114299: hypothetical protei |
| AP53_1835 | -0.053149965 | 7.517769788 | 0.219879767 | 0.647960187 | 0.730366274 | tRNA uridine 5-carboxymethylamir |
| AP53_290 | 0.053803501 | 7.280403496 | 0.215845968 | 0.650959836 | 0.733024604 | Segregation and condensation prot |
| AP53_621 | 0.111135457 | 3.43401959 | 0.215365972 | 0.651319066 | 0.733024604 | Portal protein, phage associated |
| AP53_1358 | -0.103048976 | 13.07681823 | 0.215189127 | 0.651451541 | 0.733024604 | hypothetical protein - phage associ |
| AP53_766 | -0.100160744 | 9.494367576 | 0.214108132 | 0.652262777 | 0.733512196 | FIG01115306: hypothetical protei |
| AP53_1181 | -0.115637994 | 3.955616634 | 0.213565216 | 0.65267116 | 0.733546452 | Phage lysin, N-acetylmuramoyl-L-al |
| AP53_1254 | 0.037257431 | 8.539635484 | 0.211359473 | 0.65433691 | 0.734993022 | Ribosomal RNA large subunit methy |
| AP53_381 | 0.081619154 | 5.405779303 | 0.21083071 | 0.654737807 | 0.735017978 | hypothetical protein |
| AP53_82 | 0.086025259 | 8.333549952 | 0.20932486 | 0.655882898 | 0.735877864 | DNA binding protein, FIG046916 |
| AP53_714 | -0.571942751 | -0.182630266 | 0.207799524 | 0.657047955 | 0.736729861 | Phage protein |
| AP53_512 | -0.073027339 | 5.679056086 | 0.207338126 | 0.657401401 | 0.736729861 | PTS system, N-acetylgalactosamine- |
| AP53_1511 | 0.061048693 | 7.467094329 | 0.201999022 | 0.6615267 | 0.740925173 | putative ABC transporter (ATP-bind |
| AP53_154 | 0.060079211 | 9.987487702 | 0.198935721 | 0.66392359 | 0.743180904 | M protein trans-acting positive regu |
| AP53_801 | 0.038805652 | 7.615810176 | 0.196497249 | 0.66584763 | 0.744801196 | proposed amino acid ligase found c |
| AP53_1545 | 0.070972351 | 11.4512735 | 0.196130161 | 0.666138525 | 0.744801196 | SSU ribosomal protein S6p |
| AP53_1640 | -0.057178268 | 6.301335828 | 0.195090783 | 0.666963958 | 0.745294785 | FIG01114632: hypothetical protei |
| AP53_1249 | -0.144885897 | 7.710650513 | 0.194254162 | 0.667630303 | 0.745610137 | Rhodanese-like domain protein |
| AP53_1004 | 0.049792784 | 9.335315697 | 0.192506432 | 0.669027931 | 0.746741353 | Phage lysin |
| AP53_225 | 0.051424655 | 8.009070346 | 0.186627452 | 0.67378614 | 0.751620056 | Amino acid ABC transporter, glutan |
| AP53_475 | -0.048524742 | 9.041570762 | 0.18487158 | 0.675224691 | 0.752792145 | General stress protein |
| AP53_245 | -0.065468279 | 5.525607065 | 0.181912305 | 0.677667795 | 0.755082199 | Mobile element protein |
| AP53_1379 | 0.046851884 | 9.287515947 | 0.179533244 | 0.679649163 | 0.756426644 | 16S rRNA (cytosine(967)-C(5))-metl |
| AP53_1801 | -0.091357525 | 3.605031207 | 0.179527672 | 0.679653821 | 0.756426644 | FIG01116340: hypothetical protei |
| AP53_1409 | 0.04410001 | 9.200307258 | 0.17867715 | 0.680365977 | 0.756785308 | Phospho-N-acetylmuramoyl-penta |
| AP53_636 | -0.083203411 | 4.112039212 | 0.176905051 | 0.681856297 | 0.758008632 | Phage protein |
| AP53_892 | 0.076911822 | 12.79974523 | 0.174334268 | 0.684034168 | 0.759994459 | FIG01115412: hypothetical protei |
| AP53_666 | -0.068020681 | 10.2229329 | 0.172983377 | 0.685186238 | 0.760838952 | Translation initiation factor 3 |
| AP53_501 | 0.052451613 | 8.033709296 | 0.168475184 | 0.689069948 | 0.764713991 | Promiscuous sugar phosphatase Yic |

| | | | | | | |
|---|---|---|---|---|---|---|
| AP53_1259 | 0.049765109 | 8.599922287 | 0.164164526 | 0.692841288 | 0.768348718 | Phosphopantetheine adenylyltrans |
| AP53_33 | 0.057961722 | 9.672083697 | 0.163401506 | 0.693514895 | 0.768348718 | Adenylosuccinate lyase (EC 4.3.2.2) |
| AP53_1235 | -0.068393214 | 10.02292846 | 0.163381376 | 0.693532691 | 0.768348718 | FIG01114027: hypothetical protei |
| AP53_776 | -0.048400451 | 7.944744549 | 0.162666956 | 0.694165115 | 0.768526805 | Putative tRNA-m1A22 methylase |
| AP53_1265 | 0.110047594 | 12.06436896 | 0.162020087 | 0.694739151 | 0.768526805 | Ornithine carbamoyltransferase (EC |
| AP53_470 | -0.080509017 | 7.956823872 | 0.161860171 | 0.694881269 | 0.768526805 | Transcription accessory protein (S1 |
| AP53_1080 | 0.069964708 | 7.621737499 | 0.16061448 | 0.695991148 | 0.769315956 | Glutamate transport ATP-binding p |
| AP53_331 | -0.099277767 | 5.571994503 | 0.158195987 | 0.698160455 | 0.771274584 | hypothetical protein |
| AP53_1204 | -0.090986081 | 2.871581077 | 0.156400522 | 0.699783504 | 0.772240458 | Recombinational DNA repair protei |
| AP53_1284 | -0.081202931 | 6.949306293 | 0.156348732 | 0.699830482 | 0.772240458 | Aldose 1-epimerase (EC 5.1.3.3) |
| AP53_304 | -0.058510214 | 7.635473271 | 0.154901349 | 0.701147069 | 0.773152398 | ABC-type Fe3+-siderophore transpo |
| AP53_1057 | -0.146398383 | 4.900415044 | 0.154565528 | 0.701453567 | 0.773152398 | FIG01115240: hypothetical protei |
| AP53_850 | -0.050335001 | 5.641731126 | 0.153644563 | 0.702296114 | 0.773641746 | Mobile element protein |
| AP53_1776 | -0.045567649 | 10.17093042 | 0.153155465 | 0.702744762 | 0.773696871 | RecA protein |
| AP53_922 | 0.042946607 | 9.687420644 | 0.152700532 | 0.703162821 | 0.773718274 | Signal recognition particle, subunit |
| AP53_88 | -0.098181733 | 3.287412157 | 0.150016073 | 0.705644566 | 0.776009124 | Late competence protein ComGF, a |
| AP53_138 | -0.105918618 | 10.18656462 | 0.148895555 | 0.706688098 | 0.776664519 | FIG01114903: hypothetical protei |
| AP53_1402 | -0.047624726 | 7.975794128 | 0.148517927 | 0.707040807 | 0.776664519 | Histidine transport protein (perme |
| AP53_1156 | -0.038469699 | 7.151800104 | 0.146438692 | 0.708992205 | 0.778367573 | DNA polymerase III delta subunit (E |
| AP53_rna54 | 0.577749387 | -0.548503406 | 0.139849506 | 0.716714284 | 0.786400467 | 5S RNA |
| AP53_774 | -0.037226898 | 8.855491958 | 0.132531417 | 0.722475486 | 0.792273965 | Chromosome replication initiation |
| AP53_638 | 0.082774658 | 3.705966894 | 0.131016623 | 0.723992215 | 0.793488926 | FIG01116946: hypothetical protei |
| AP53_1830 | -0.051019243 | 5.675547829 | 0.128927862 | 0.726100141 | 0.795350098 | Mobile element protein |
| AP53_1033 | -0.134888863 | 5.62939968 | 0.127314288 | 0.72774187 | 0.796698799 | hypothetical protein |
| AP53_281 | 0.040629744 | 8.123607792 | 0.126678906 | 0.728391584 | 0.796960577 | FIG011178: rRNA methylase |
| AP53_pseudo656 | -0.064339794 | 8.518667932 | 0.125923831 | 0.729166102 | 0.797358538 | Lipoteichoic acid synthase LtaS Typ |
| AP53_1617 | 0.046275647 | 6.251830487 | 0.125238201 | 0.729871676 | 0.797680699 | 6-phospho-beta-galactosidase (EC 3 |
| AP53_1603 | 0.049611234 | 6.695972091 | 0.121628514 | 0.733622972 | 0.801329313 | Substrate-specific component PdxU |
| AP53_1668 | -0.054035349 | 13.35226329 | 0.120455657 | 0.734855376 | 0.802224007 | Collagen-like surface protein |
| AP53_442 | 0.065055179 | 3.886485751 | 0.11976312 | 0.735586257 | 0.802570503 | FIG01115625: hypothetical protei |
| AP53_1151 | 0.044088805 | 7.822189144 | 0.117685104 | 0.737793753 | 0.804526784 | FIG01115059: hypothetical protei |
| AP53_1638 | -0.048918561 | 7.493116441 | 0.116012191 | 0.739586913 | 0.806029308 | COG1939: Ribonuclease III family p |
| AP53_404 | -0.1029976 | 3.441721999 | 0.113815866 | 0.741963327 | 0.808165441 | hypothetical protein |
| AP53_1623 | 0.058248017 | 6.765951862 | 0.113166054 | 0.742671349 | 0.808482944 | Galactose-6-phosphate isomerase, |
| AP53_223 | 0.090979496 | 4.257604621 | 0.111332941 | 0.744681047 | 0.810216319 | FIG01114598: hypothetical protei |

| | | | | | | |
|---|---|---|---|---|---|---|
| AP53_441 | -0.074356309 | 4.005282735 | 0.108899458 | 0.747377785 | 0.812694835 | putative L-glutamate ligase |
| AP53_1802 | 0.100073737 | 3.801703709 | 0.104786004 | 0.75201386 | 0.81727822 | Phage encoded transcriptional regu |
| AP53_870 | -0.037495715 | 8.825899477 | 0.102992386 | 0.754067077 | 0.818452775 | L-proline glycine betaine binding A |
| AP53_407 | 0.041887561 | 7.994504104 | 0.102699427 | 0.75440432 | 0.818452775 | Dephospho-CoA kinase (EC 2.7.1.24 |
| AP53_1597 | 0.035164466 | 8.736347993 | 0.102486735 | 0.754649498 | 0.818452775 | CTP synthase (EC 6.3.4.2) |
| AP53_220 | 0.067333817 | 11.41052102 | 0.102372514 | 0.754781281 | 0.818452775 | SSU ribosomal protein S7p (S5e) |
| AP53_1476 | 0.041216522 | 7.639481239 | 0.101617702 | 0.75565421 | 0.818941832 | Seryl-tRNA synthetase (EC 6.1.1.11) |
| AP53_386 | -0.116376409 | 3.007733876 | 0.100848663 | 0.756547299 | 0.819452181 | Mobile element protein |
| AP53_668 | -0.055330189 | 10.80527704 | 0.100007638 | 0.757528321 | 0.82005715 | LSU ribosomal protein L20p |
| AP53_1589 | 0.048277726 | 9.549365128 | 0.098938311 | 0.758782268 | 0.820666473 | Acid phosphatase (EC 3.1.3.2) |
| AP53_1206 | 0.122215698 | 3.161078979 | 0.098528677 | 0.75926461 | 0.820666473 | Phage protein |
| AP53_rna57 | -0.095932187 | 5.964443146 | 0.098317239 | 0.759514013 | 0.820666473 | tRNA-Arg-CCT |
| AP53_655 | -0.039014887 | 9.319195173 | 0.09808998 | 0.759782407 | 0.820666473 | Phage integrase: site-specific recom |
| AP53_517 | -0.045854905 | 5.639080472 | 0.094362917 | 0.764234058 | 0.825015743 | Beta-phosphoglucomutase (EC 5.4. |
| AP53_1267 | -0.083919704 | 11.23135134 | 0.093313056 | 0.765505462 | 0.825928906 | Arginine deiminase (EC 3.5.3.6) |
| AP53_4 | 0.061554615 | 4.818853348 | 0.089037195 | 0.770766607 | 0.831143324 | hypothetical protein |
| AP53_pseudo1183 | -0.089942829 | 4.144345891 | 0.086832196 | 0.77353393 | 0.833664274 | Phage holin |
| AP53_87 | -0.08728257 | 3.194816257 | 0.082304706 | 0.77933898 | 0.839454473 | Late competence protein ComGE, F |
| AP53_255 | 0.043151039 | 9.625319323 | 0.079793071 | 0.782634571 | 0.842536718 | Cysteine ABC transporter, substrate |
| AP53_1656 | -0.03663164 | 9.178346442 | 0.079429331 | 0.783116495 | 0.842588202 | Intramembrane protease RasP/YluC |
| AP53_1451 | 0.041705776 | 6.309742493 | 0.078804664 | 0.783946927 | 0.843014397 | Negative transcriptional regulator- |
| AP53_1823 | -0.033497236 | 5.856257941 | 0.078435232 | 0.784439732 | 0.843077253 | Phosphohydrolase (MutT/nudix fan |
| AP53_1549 | -0.057066929 | 8.063844327 | 0.076298825 | 0.787314594 | 0.845522924 | Thioredoxin |
| AP53_1639 | 0.043666962 | 7.323090863 | 0.076098397 | 0.787586526 | 0.845522924 | Cysteinyl-tRNA synthetase (EC 6.1.1 |
| AP53_1609 | -0.03014995 | 7.566110226 | 0.07384024 | 0.790677449 | 0.848332143 | Type I restriction-modification syst |
| AP53_1496 | 0.029095594 | 8.657112087 | 0.073550749 | 0.791077372 | 0.848332143 | Phosphoglycerate mutase family |
| AP53_21 | -0.066438291 | 6.210596514 | 0.072657905 | 0.792316187 | 0.849080934 | Phosphoribosylaminoimidazole-su |
| AP53_1658 | 0.028574395 | 8.238727787 | 0.072417953 | 0.792650516 | 0.849080934 | Undecaprenyl diphosphate synthas |
| AP53_1278 | -0.042101647 | 9.295328339 | 0.071993049 | 0.793244011 | 0.849248001 | Oxidoreductase, Gfo/Idh/MocA fan |
| AP53_644 | 0.045242276 | 4.75123839 | 0.07144629 | 0.794010495 | 0.849599984 | Replicative DNA helicase (DnaB) (EC |
| AP53_90 | 0.03175186 | 7.485381372 | 0.070979909 | 0.7946668 | 0.849833751 | Adenine-specific methyltransferase |
| AP53_1540 | -0.026003287 | 9.211085047 | 0.069335022 | 0.797000251 | 0.85185985 | Excinuclease ABC subunit A |
| AP53_961 | -0.032732326 | 7.148994965 | 0.067904647 | 0.799053701 | 0.853584609 | Phosphate transport system perme |
| AP53_919 | -0.034225026 | 9.876042395 | 0.067406208 | 0.79977471 | 0.85388488 | FIG01114660: hypothetical protei |
| AP53_1443 | 0.027196119 | 8.795064872 | 0.065052218 | 0.803219097 | 0.857090856 | Galactose-6-phosphate isomerase, |

| | | | | | | |
|---|---|---|---|---|---|---|
| AP53_1055 | 0.027799512 | 7.699856721 | 0.0611647 | 0.809056634 | 0.862845564 | hypothetical phage protein |
| AP53_332 | 0.054807448 | 4.479779567 | 0.060842104 | 0.809549835 | 0.862897436 | FIG01115797: hypothetical protei |
| AP53_7 | -0.034923035 | 6.414484418 | 0.059703864 | 0.811301255 | 0.864289647 | Transcription-repair coupling facto |
| AP53_596 | -0.087346185 | 8.004957284 | 0.059094219 | 0.812246638 | 0.86482212 | putative repressor C1 |
| AP53_1292 | -0.041926191 | 5.910146344 | 0.058054052 | 0.813871688 | 0.865625319 | DNA-binding response regulator, Ar |
| AP53_1351 | -0.070546149 | 3.956639772 | 0.057889213 | 0.814130632 | 0.865625319 | hypothetical phage protein |
| AP53_162 | -0.034653088 | 5.480491043 | 0.057756805 | 0.814338914 | 0.865625319 | Quinolinate phosphoribosyltransfe |
| AP53_142 | -0.048346308 | 4.788570738 | 0.056693525 | 0.816020769 | 0.866906644 | Ascorbate-specific PTS system, EIIB |
| AP53_1734 | -0.051642864 | 7.484039976 | 0.056431633 | 0.816437582 | 0.866906644 | Transcriptional regulator, MutR far |
| AP53_798 | -0.03027551 | 6.426757126 | 0.05582199 | 0.817411853 | 0.867466597 | Hyaluronate lyase precursor (EC 4.2 |
| AP53_1682 | 0.035365756 | 5.782192653 | 0.055244456 | 0.818340026 | 0.867803589 | Dipeptide transport system permea |
| AP53_1168 | 0.021472575 | 7.247266685 | 0.054940016 | 0.818831372 | 0.867803589 | Recombination protein RecR |
| AP53_1116 | 0.030848434 | 7.551347705 | 0.0543817 | 0.819736219 | 0.867803589 | UDP-N-acetylglucosamine 1-carbox |
| AP53_189 | -0.020870642 | 8.5345132 | 0.054373208 | 0.819750021 | 0.867803589 | DNA repair protein RadA |
| AP53_143 | -0.036682127 | 4.941427565 | 0.054241104 | 0.819964854 | 0.867803589 | Ascorbate-specific PTS system, EIIA |
| AP53_1529 | 0.027905075 | 8.227547356 | 0.053895537 | 0.820528145 | 0.867926501 | Mannose-6-phosphate isomerase (E |
| AP53_pseudo1680 | -0.04926358 | 5.028817522 | 0.053278141 | 0.821539307 | 0.868076996 | Streptococcal mitogenic exotoxin Z |
| AP53_869 | 0.02984159 | 8.672479792 | 0.053262572 | 0.821564885 | 0.868076996 | L-proline glycine betaine ABC transp |
| AP53_1398 | 0.027679816 | 9.98185083 | 0.051426955 | 0.824608847 | 0.870819245 | Multimodular transpeptidase-trans |
| AP53_262 | 0.030228222 | 7.796758407 | 0.049543319 | 0.827792665 | 0.873706124 | Potassium uptake protein TrkH |
| AP53_528 | -0.023596546 | 8.886217142 | 0.047434862 | 0.831433091 | 0.876655538 | FIG000506: Predicted P-loop-conta |
| AP53_1499 | 0.033458888 | 9.780226003 | 0.047402061 | 0.831490389 | 0.876655538 | Aspartyl-tRNA(Asn) amidotransferas |
| AP53_1293 | -0.031488138 | 5.977682409 | 0.046340593 | 0.833356013 | 0.878145505 | Two-component sensor kinase YesM |
| AP53_514 | -0.026836564 | 6.573820684 | 0.044519472 | 0.836609824 | 0.881095859 | predicted 4-deoxy-L-threo-5-hexosu |
| AP53_1612 | -0.039461481 | 4.415013354 | 0.043203433 | 0.839004855 | 0.883139058 | FIG00525014: hypothetical protei |
| AP53_607 | -0.051813821 | 5.266162481 | 0.042667222 | 0.839991633 | 0.883605961 | Hyaluronate lyase (phage associated |
| AP53_179 | 0.036711952 | 9.11682714 | 0.042468552 | 0.840358889 | 0.883605961 | FIG01118080: hypothetical protei |
| AP53_1032 | 0.054179041 | 5.685132842 | 0.042011921 | 0.841206421 | 0.884018226 | hypothetical protein - phage associ |
| AP53_349 | 0.022512118 | 8.400918934 | 0.041154399 | 0.842811145 | 0.885225342 | ADP-ribose pyrophosphatase (EC 3. |
| AP53_1805 | -0.030523455 | 7.418596014 | 0.040634337 | 0.843792881 | 0.885777167 | biofilm formation protein YmcA |
| AP53_60 | 0.040978289 | 10.31908623 | 0.04012434 | 0.844762009 | 0.886315167 | SSU ribosomal protein S5p (S2e) |
| AP53_141 | -0.03007674 | 5.279381164 | 0.039845605 | 0.845294395 | 0.886394609 | Ascorbate-specific PTS system, EIIC |
| AP53_61 | 0.042514083 | 10.31748478 | 0.038890023 | 0.847134465 | 0.88784449 | LSU ribosomal protein L30p (L7e) |
| AP53_419 | -0.022319668 | 9.639050574 | 0.037855499 | 0.849153307 | 0.889480069 | Oxygen-insensitive NAD(P)H nitrore |
| AP53_1572 | 0.025341466 | 6.581863799 | 0.036374116 | 0.852094848 | 0.892079881 | DNA polymerase III epsilon subunit |

| | | | | | | |
|---|---|---|---|---|---|---|
| AP53_1564 | 0.029049644 | 8.421111977 | 0.03601172 | 0.852823885 | 0.892361812 | Aquaporin (Major Intrinsic Protein |
| AP53_449 | 0.08280991 | 1.82602402 | 0.034953806 | 0.854974177 | 0.894129783 | hypothetical protein |
| AP53_439 | -0.056113334 | 2.385210709 | 0.03386946 | 0.857213541 | 0.895988951 | Shikimate 5-dehydrogenase I alpha |
| AP53_1297 | -0.028665118 | 6.253431008 | 0.031040352 | 0.86323735 | 0.901799622 | Multiple sugar ABC transporter, me |
| AP53_1632 | 0.022746626 | 7.214749221 | 0.030678987 | 0.86402691 | 0.902138909 | Transcriptional regulator, Cro/CI fa |
| AP53_805 | 0.032701637 | 4.94977484 | 0.030170248 | 0.865146659 | 0.9028224 | FIG01115418: hypothetical protei |
| AP53_1189 | 0.045610015 | 1.973860912 | 0.029622001 | 0.866364336 | 0.903607295 | FIG022125: Phage-associated prote |
| AP53_1611 | -0.039052113 | 4.559297688 | 0.029037453 | 0.867675521 | 0.904488822 | Transcriptional regulator DegU, Lu |
| AP53_63 | -0.02702966 | 10.43719894 | 0.027179064 | 0.871937002 | 0.908443221 | Preprotein translocase secY subuni |
| AP53_1430 | 0.031013061 | 7.962175931 | 0.025998197 | 0.87472324 | 0.910857193 | Glycyl-tRNA synthetase alpha chain |
| AP53_83 | -0.036657583 | 3.433511599 | 0.025479922 | 0.875966632 | 0.911662859 | Late competence protein ComGA, a |
| AP53_761 | 0.028340633 | 8.406277453 | 0.024946236 | 0.877260653 | 0.912520326 | FIG015049: hypothetical protein |
| AP53_516 | 0.017390656 | 6.978985836 | 0.024280842 | 0.878894101 | 0.913729754 | 4-hydroxy-2-oxoglutarate aldolase |
| AP53_843 | -0.020125255 | 7.471071188 | 0.023547284 | 0.880721737 | 0.915139663 | Spermidine Putrescine ABC transpo |
| AP53_rna4 | 0.178714372 | -0.831064152 | 0.019561744 | 0.891205457 | 0.925537609 | 5S RNA |
| AP53_647 | 0.047808345 | 3.469571357 | 0.017924323 | 0.89582749 | 0.929637797 | hypothetical phage protein |
| AP53_1583 | -0.015662166 | 8.319343603 | 0.017826106 | 0.896111447 | 0.929637797 | Protein of unknown function DUF1 |
| AP53_376 | 0.026584802 | 3.814968251 | 0.017516228 | 0.897012609 | 0.929871239 | Competence-stimulating peptide A |
| AP53_274 | 0.01685177 | 10.45166169 | 0.017227988 | 0.897858164 | 0.929871239 | FIG01114242: hypothetical protei |
| AP53_498 | 0.168316782 | -1.006521509 | 0.017266322 | 0.898223167 | 0.929871239 | hypothetical protein |
| AP53_775 | 0.018333098 | 9.194484888 | 0.017094315 | 0.898252742 | 0.929871239 | Endonuclease III (EC 4.2.99.18) |
| AP53_402 | 0.023668364 | 4.590131084 | 0.01656661 | 0.899825983 | 0.931003322 | hypothetical protein |
| AP53_522 | 0.015648703 | 7.975341605 | 0.016387199 | 0.900366656 | 0.931066424 | Hydroxyacylglutathione hydrolase |
| AP53_930 | -0.020924413 | 6.355157634 | 0.015486228 | 0.903128532 | 0.933373528 | Substrate-specific component Pdx |
| AP53_1266 | 0.036722876 | 10.8204009 | 0.015347997 | 0.903559433 | 0.933373528 | Acetyltransferase (EC 2.3.1.-) |
| AP53_458 | -0.020126972 | 3.931775774 | 0.014923692 | 0.904894562 | 0.934255503 | Mobile element protein |
| AP53_1606 | -0.025265496 | 4.367122512 | 0.014514403 | 0.906200845 | 0.935106773 | Competence-specific sigma factor ( |
| AP53_128 | -0.018520281 | 6.379202322 | 0.013856582 | 0.908340174 | 0.936816301 | V-type ATP synthase subunit D (EC 3 |
| AP53_1797 | 0.032746177 | 3.912813951 | 0.013413809 | 0.909809294 | 0.937496047 | FIG01114328: hypothetical protei |
| AP53_635 | -0.026067938 | 3.98579451 | 0.013341404 | 0.910051861 | 0.937496047 | conserved hypothetical protein - p |
| AP53_1450 | -0.011756398 | 7.186068963 | 0.01321594 | 0.910473768 | 0.937496047 | Copper-translocating P-type ATPas |
| AP53_1723 | 0.015660119 | 8.835087518 | 0.013052014 | 0.911028091 | 0.937496047 | Transcriptional regulator, DeoR fan |
| AP53_1528 | -0.014087245 | 5.725136531 | 0.012938434 | 0.911414241 | 0.937496047 | Mobile element protein |
| AP53_355 | 0.010178643 | 7.784997873 | 0.012424437 | 0.913183597 | 0.938168742 | Manganese ABC transporter, ATP-bi |
| AP53_1562 | -0.011697687 | 7.62799416 | 0.012330228 | 0.913511896 | 0.938168742 | C3-degrading proteinase |

| | | | | | | |
|---|---|---|---|---|---|---|
| AP53_271 | -0.011271795 | 9.333956541 | 0.012328408 | 0.913518249 | 0.938168742 | Helicase loader DnaI |
| AP53_999 | -0.015757055 | 7.754957157 | 0.011767226 | 0.915500901 | 0.939707693 | Immunoreactive protein Se23.5 (Fr |
| AP53_283 | 0.016462722 | 9.767281081 | 0.010956907 | 0.918450241 | 0.941908876 | FIG005935: membrane protein |
| AP53_261 | 0.011515801 | 8.013127346 | 0.010912255 | 0.918615921 | 0.941908876 | Trk system potassium uptake protei |
| AP53_1392 | -0.011764795 | 8.160192567 | 0.00920412 | 0.925233418 | 0.948193276 | S-ribosylhomocysteine lyase (EC 4.4 |
| AP53_518 | 0.009303455 | 7.32366515 | 0.008331566 | 0.928854361 | 0.951247707 | Epoxyqueuosine (oQ) reductase Qu |
| AP53_454 | -0.012856373 | 9.919725063 | 0.008251959 | 0.929194051 | 0.951247707 | Plasmid stabilization system antito |
| AP53_901 | -0.009820647 | 7.690586902 | 0.007395107 | 0.932960493 | 0.954600062 | tRNA:m(5)U-54 MTase gid |
| AP53_326 | -0.0112867 | 9.854876452 | 0.006363707 | 0.937799373 | 0.959045618 | Serine endopeptidase ScpC (EC 3.4. |
| AP53_233 | 0.009561648 | 8.635923669 | 0.00542775 | 0.942545616 | 0.962555833 | Cysteine desulfurase (EC 2.8.1.7), Su |
| AP53_34 | -0.026411713 | 5.755964412 | 0.005422044 | 0.942575762 | 0.962555833 | Transcriptional regulator |
| AP53_1601 | 0.007674883 | 7.763040086 | 0.005394873 | 0.942719546 | 0.962555833 | Hypothetical protein ywlG |
| AP53_1811 | -0.007822396 | 7.932330972 | 0.005003189 | 0.944834174 | 0.964207746 | Integral membrane protein |
| AP53_500 | 0.006563409 | 8.026905762 | 0.004510435 | 0.947616489 | 0.966538942 | tRNA-dependent lipid II--L-alanine l |
| AP53_1361 | -0.017973362 | 9.39071655 | 0.004300924 | 0.948845625 | 0.967284327 | putative cI-like repressor - phage as |
| AP53_207 | 0.008107211 | 7.12594599 | 0.004077831 | 0.950187995 | 0.967894841 | N-acetylmannosamine kinase (EC 2. |
| AP53_322 | -0.01556321 | 3.828451019 | 0.004036318 | 0.950441817 | 0.967894841 | FIG01115198: hypothetical protei |
| AP53_1258 | 0.006460478 | 8.322350417 | 0.00355377 | 0.953494392 | 0.970494292 | Lon-like protease with PDZ domain |
| AP53_888 | 0.005463172 | 9.182311722 | 0.003276418 | 0.955343763 | 0.971867004 | DNA gyrase subunit A (EC 5.99.1.3) |
| AP53_519 | 0.004965215 | 8.866446243 | 0.002864556 | 0.958241633 | 0.974304353 | Peptide chain release factor 2; prog |
| AP53_1613 | -0.015289033 | 3.795444941 | 0.002721974 | 0.959293095 | 0.974862773 | Salivaricin A modification enzyme |
| AP53_318 | -0.005753892 | 10.427855 | 0.002621555 | 0.960050305 | 0.975121739 | rRNA small subunit methyltransfera |
| AP53_1787 | -0.015209424 | 5.888319301 | 0.002352416 | 0.962154676 | 0.976748027 | Phage antirepressor protein |
| AP53_385 | -0.009042574 | 4.362309074 | 0.002086479 | 0.964356269 | 0.978379598 | Mobile element protein |
| AP53_1104 | -0.005909716 | 8.137301301 | 0.002038289 | 0.96476999 | 0.978379598 | ATP-dependent Zn protease |
| AP53_107 | 0.005016526 | 8.10523054 | 0.001946952 | 0.965567806 | 0.978677343 | FIG01115182: hypothetical protei |
| AP53_509 | 0.006700717 | 5.783510146 | 0.001765237 | 0.967212908 | 0.979833118 | PTS system, N-acetylgalactosamine- |
| AP53_1539 | 0.004315895 | 8.072907953 | 0.001578603 | 0.968993511 | 0.981124885 | Aminopeptidase YpdF (MP-, MA-, M |
| AP53_527 | 0.004620964 | 8.774876702 | 0.001382503 | 0.970982193 | 0.981521881 | Asparaginyl-tRNA synthetase (EC 6.1 |
| AP53_1803 | -0.00958732 | 6.483353547 | 0.001381954 | 0.970987953 | 0.981521881 | hypothetical protein |
| AP53_1683 | -0.006084856 | 6.01827946 | 0.001370368 | 0.971109761 | 0.981521881 | Dipeptide transport system permea |
| AP53_341 | 0.009627212 | 3.9403568 | 0.001300176 | 0.971859025 | 0.981521881 | hypothetical protein |
| AP53_1786 | 0.008693392 | 6.068040946 | 0.001291421 | 0.971953896 | 0.981521881 | kilA protein, putative phage-relatec |
| AP53_1684 | 0.005495143 | 6.496705515 | 0.001235973 | 0.972562313 | 0.981521881 | Dipeptide transport ATP-binding pr |
| AP53_10 | -0.006894068 | 6.92771867 | 0.001203468 | 0.972925347 | 0.981521881 | FIG01114659: hypothetical protei |

| | | | | | | |
|---|---|---|---|---|---|---|
| AP53_686 | -0.004167509 | 8.354929584 | 0.00101133 | 0.975179692 | 0.982826543 | Uracil phosphoribosyltransferase (E |
| AP53_581 | -0.003142339 | 7.651479409 | 0.001007129 | 0.975231284 | 0.982826543 | ATP-dependent nuclease, subunit A |
| AP53_1596 | 0.00407941 | 7.534487352 | 0.000771477 | 0.978320909 | 0.985428585 | hypothetical protein |
| AP53_1710 | 0.007083215 | 4.837104462 | 0.000650825 | 0.980087706 | 0.986696182 | FIG01114305: hypothetical protei |
| AP53_126 | -0.003524891 | 6.282394857 | 0.000591457 | 0.981017409 | 0.987120161 | V-type ATP synthase subunit A (EC 3 |
| AP53_78 | -0.002783063 | 7.419845793 | 0.000534154 | 0.981960204 | 0.987498585 | Tyrosyl-tRNA synthetase (EC 6.1.1.1 |
| AP53_19 | 0.003633954 | 7.876750887 | 0.000507787 | 0.982411009 | 0.987498585 | Phosphate:acyl-ACP acyltransferase |
| AP53_845 | 0.002776601 | 5.452419827 | 0.000265765 | 0.987274697 | 0.991873803 | Two-component response regulato |
| AP53_1397 | 0.002112271 | 8.742721169 | 0.000222379 | 0.988359553 | 0.992450022 | RecU Holliday junction resolvase |
| AP53_418 | -0.001463153 | 8.8980063 | 0.000183964 | 0.98941252 | 0.992993641 | Lactoylglutathione lyase (EC 4.4.1.5 |
| AP53_252 | 0.001381551 | 7.357726558 | 0.00010412 | 0.992034736 | 0.995110812 | FIG145533: Methyltransferase (EC 2 |
| AP53_392 | 0.005442859 | 0.610601094 | 4.51E-05 | 0.994760439 | 0.996998671 | FIG01116220: hypothetical protei |
| AP53_157 | 0.001680262 | 5.443569593 | 4.19E-05 | 0.994944063 | 0.996998671 | FIG01115469: hypothetical protei |
| AP53_1224 | 0.000459666 | 8.260536247 | 1.77E-05 | 0.99671392 | 0.998256821 | Octaprenyl diphosphate synthase (E |
| AP53_420 | 0.000347689 | 9.55445663 | 7.91E-06 | 0.997804481 | 0.998833676 | Proline dipeptidase (EC 3.4.13.9) |
| AP53_20 | 0.000404652 | 7.070122534 | 3.17E-06 | 0.998609787 | 0.999120058 | Acyl carrier protein |
| AP53_58 | -0.000209035 | 10.54885701 | 1.27E-06 | 0.999120058 | 0.999120058 | LSU ribosomal protein L6p (L9e) |

porter protein
5)
hain (EC 4.1.1.3)
)

3)
ephospho-CoA transferase (EC 2.7.7.61)
eaminase (EC 4.3.1.4)
C 6.3.4.3)

C 2.1.2.5) @ Glutamate formyltransferase

component (EC 2.7.1.69)

omponent (EC 2.7.1.69) / PTS system, trehalose-specific IIC component (EC 2.7.1.69) / PTS system, trehalose-specific IIA component (EC 2.7.1.69)

chain (EC 4.1.1.3)
component (EC 2.7.1.69)

)
Toximoron (Other)

ne associated protein

n

B (SagA)
n
chenan operon, BglG family
ase (EC 2.7.6.1)
EC 3.8.1.2)

ransporter, ATP-binding protein
subunit (EC 2.7.7.6)
, beta chain (EC 4.1.1.41); Oxaloacetate decarboxylase beta chain (EC 4.1.1.3)
xaloacetate decarboxylase; Biotin carboxyl carrier protein
ining
ly IIB component

3)

n

n

putative peptidoglycan hydrolase

1.3.25)
ransporter, permease protein
5)
of predicted cobalamin ECF transporter
otein PhoH, predicted ATPase
n

:C 3.2.1.93)
omponent (EC 2.7.1.69)

Ps0857
rier protein (EC 4.1.3.6)
tein C (EC 1.6.4.-)

se 3; Mitogenic factor 3

-.-)
yltransferase C (EC 2.1.1.52) ## SSU rRNA m2G1207

)

protein

.1.2)
4.23.36)
se (EC 1.1.1.133)
xidoreductase in cluster with COG2110
nily

n
chain (EC 4.1.1.3)
ss II (EC 4.1.2.13)

n enzyme TrmE
n
tein F (EC 1.6.4.-)
ase
de-3-phosphate dehydrogenase (NADP) (EC 1.2.1.9)
ease protein 2
nding protein

76, Firmicutes subgroup
acid biosynthesis, sensor histidine kinase
synthase co-transcribed bacterial homolog PROSC
EC 2.4.2.1); N-Ribosylnicotinamide phosphorylase (EC 2.4.2.1) ## possible
hage associated
.8)
4.-)
component (EC 2.7.1.69)
EC 2.4.2.9)
(SagF)

component (EC 2.7.1.69)
(EC 3.1.3.27)

e (EC 5.1.3.4)
rase subunit B (EC 2.8.3.9)

ococcal antigen
g protein; Streptolysin S export protein (SagG)

n
1

unction, cellobiose operon
pilin backbone protein T-antigen
nily

iacylglycerol kinase catalytic domain]

)4 of energizing module of predicted ECF transporter
ribosomal protein L33p, zinc-independent

transferase rlmB (EC 2.1.1.-) ## LSU rRNA Gm2251

oseudouridine synthase D, Bacillus subtilis YjbO type

transferase (EC 2.4.1.-)

thase, KASIII (EC 2.3.1.41)
C (TC 3.A.5.1.1)
nding subunit ClpX

family (EC 2.7.1.69)

in

' subunit (EC 2.7.7.6)

omponent (EC 2.7.1.69)

ase (EC 2.7.6.1)

.4.2.10)
r SA14-24

egulator
n related to cell division

of predicted cobalamin ECF transporter
uridyltransferase (EC 2.7.7.23) / Glucosamine-1-phosphate N-acetyltransferase (EC 2.3.1.157)
protein ZnuC
iphosphatase RdgB (dHAPTP, dITP, XTP-specific) (EC 3.6.1.15)

n

2) / Folylpolyglutamate synthase (EC 6.3.2.17)
TrmH family tRNA/rRNA methyltransferase YacO
A-specific

utamate--L-lysine ligase (EC 6.3.2.7)
3.14)

talytic subunit (EC 1.3.3.1)
erase, alpha subunit (EC 2.8.3.8)
oside utilization, GntR family
nzyme (EC 1.97.1.4)

3.14)

yrophosphatase (EC 3.6.1.1)

otein receptor Shr
a2 (EC 3.5.1.28)
3.1.8)
hatase family protein

ase (EC 3.2.2.16) / S-adenosylhomocysteine nucleosidase (EC 3.2.2.9)
)
a subunit (EC 2.7.7.6)

vith GDSL-like motif

em

6) @ Acetyl-CoA acetyltransferase (EC 2.3.1.9)
tein PaaD, thioesterase
ferase (EC 2.1.1.33)

ing], amidotransferase subunit (EC 6.3.5.2) / GMP synthase [glutamine-hydrolyzing], ATP pyrophosphatase subunit (EC 6.3.5.2)
rotein
P2)
n

BC transporters with duplicated ATPase domains

cid biosynthesis FabT
12)
operon, ManO
4)
n
.1.1.38)

in
asmic phosphate-binding protein PstS (TC 3.A.1.7.1)
in
with unspecified monosaccharide ABC transport system
@ SSU ribosomal protein S14p (S29e), zinc-independent
enase [NAD] (EC 1.2.1.24); Succinate-semialdehyde dehydrogenase [NADP+] (EC 1.2.1.16)

e (EC 2.7.8.7)
e deacetylase (EC 3.5.1.-)

14)
n

gase (EC 6.3.2.8)

component (EC 2.7.1.69)

nolog
ort system carrier protein
ain protein (Bsu YtpR)

ein) -possibly involved in cell wall localization and side chain formation of rhamnose-glucose polysaccharide
P-binding protein

tase (EC 1.1.1.34)
n
olytic subunit (EC 3.4.21.92)

xaloacetate decarboxylase; Biotin carboxyl carrier protein

n

nsferase (EC 2.7.7.8)
4.-)

ntial role in cell wall biosynthesis

tein
4.11.18)

C 6.2.1.3)
protein, GcvH-like

C 6.3.4.3)

ferase (EC 2.1.1.31)
nyl-diaminopimelate desuccinylase and related deacylases
uctase (EC 1.1.1.100)

osyltransferase (EC 2.4.1.-)
omponent (EC 2.7.1.69) / PTS system, mannose-specific IIA component (EC 2.7.1.69)
mine reductase (EC 1.1.1.158)
nvolved in rRNA and/or ribosome maturation and assembly

n

14)

..-) (EC 2.7.4.-)
sphotransferase of PTS system (EC 2.7.3.9)

oR (transcriptional repressor)
ession IbrB

nase (ATP-binding subunit)

of pyridoxine metabolism
VA

5)

otein J
drolase (EC 3.5.2.12)
r utilization related?)

inding subunit ClpE
yltransferase E (EC 2.1.1.-)
4.2.1.70)

(EC 2.1.2.9)

21)

osynthesis lipoprotein ApbE
n
EC 1.15.1.1)
se (EC 2.3.3.10)
contains S4-like RNA binding domain
.-)

c
g protein ParB / Stage 0 sporulation protein J

n
3)
tein
inase
n
4.2.1.70)
ily
ily

ng permease - glutamine transport
/e
n
(EC 4.3.1.17)

ding protein, similarity with At5g48545 and yeast YDL125C (HNT1)
ADP-ribose binding module

stem
family

4)
g protein

sphate 3-phosphatidyltransferase (EC 2.7.8.5)

regulon ComD
C 1.1.1.169)

se (EC 2.3.2.5)
component (EC 2.7.1.69)
ily 25
nage associated
ing protein), spy1791 homolog

e A; Peptidyl-prolyl cis-trans isomerase (EC 5.2.1.8)
eme from hemoglobin to apo-SiaA/HtsA

a subunit (EC 2.7.7.6)
rotein PotA (TC 3.A.1.11.1)
ase subunit 2 (EC 6.1.1.13)

ng protein, spy1790 homolog
rotein YabA
rter, permease protein MalG
n

ZinT
protein

rotein

ltransferase (EC 2.3.1.51)
acid-binding protein

n
binase
lcohol forming, (R)-acetoin-specific (EC 1.1.1.4) / Acetoin (diacetyl) reductase (EC 1.1.1.5)
[C1]
e of SIR2 family
n

m, ATPase component
dine synthase D (EC 4.2.1.70)
rase (EC 5.3.1.-)

2.1.86)
ase [FMN] (EC 1.3.1.9)
enase (EC 1.1.1.30)
eeN

: (EC 4.3.1.17)
nease protein OppB (TC 3.A.1.5.1)
pseudouridine synthase D, Bacillus subtilis YhcT type

vith transport systems (COG1739)

nce regulon ComE
or
o-8-oxoguanine-triphosphatase) (EC 3.6.1.-)
ansferase (EC 2.7.7.24)
se
ansacylase (EC 2.3.1.39)

ninator, BglG family

d GTPase EngC

nease protein OppC (TC 3.A.1.5.1)
C 5.2.1.8)
nerase (EC 5.1.3.13)
a and tau (EC 2.7.7.7)
ng permease - macrolide efflux

ge chain (EC 6.3.5.5)

em, restriction subunit R (EC 3.1.21.3)

6.3.14)

n

nsferase

Na+ or H+) symporter

C 5.3.1.6)

.1.35)

of thiamin ECF transporter

n (SagE)

ent proteases, putative molecular chaperone

2.7.4.6)

glucosaminyl 1-phosphate transferase (EC 2.7.8.-)

ltransferase-isomerase (EC 5.-.-.-)

nsferase (EC 2.7.7.18)

rotein CutC

ily protein

2.4.2.29)

protein

d for biogenesis of 30S ribosome subunit

n
7.1.60)
eductase (EC 1.-.-.-) / Dihydropteridine reductase (EC 1.5.1.34)
ne permease (SagI)

'B

n

bosomal protein L32p, Firmicutes subfamily

EC 2.4.2.1)

ng protein YchF

ase (EC 3.5.4.-)
eptidase

1)
transferase/alpha-L-Rha alpha-1,3-L- rhamnosyltransferase (EC 2.4.1.-)
lneuraminate-binding protein
nage associated
n

ated
omponent (EC 2.7.1.69)
n
ng protein FtsE (TC 3.A.5.1.1)

(EC 4.1.1.31)

omolog

nyl-diaminopimelate desuccinylase and related deacylases

unit (EC 2.7.7.7)

cus), permease protein
gulator VraR
ntial for anaerobic expression of the ADI pathway, Crp/Fnr family

orT family

1.14.-.-)
n
em, ATPase component
C 5.1.3.1)
erythritol kinase (EC 2.7.1.148)

qC
e IIc

ociated with ferric iron transporter, SPy1061 homolog

aperone GroES
n

ctase MsrA (EC 1.8.4.11)

protein SufC

A
osyltransferase (EC 2.4.2.8)
571856) SAR1694
nding subunit ClpA

acid biosynthesis, DltR
n
g protein OppF (TC 3.A.1.5.1)
stem

ic cystine-binding protein FliY

1.17) @ Glutamyl-tRNA(Gln) synthetase (EC 6.1.1.24)

n
..15)
gizing module of ECF transporters
of biotin ECF transporter
C (SpeC); _Toximoron (Superantigen)
1.2)

;
n
ent beta-subunit (EC 1.2.4.-)
IIB component (EC 2.7.1.69) / PTS system, beta-glucoside-specific IIC component (EC 2.7.1.69) / PTS system, beta-glucoside-specific IIA component (EC 2.7
ein IsdE
ulator MntR
rter, substrate binding periplasmic protein MalE

(EC 1.5.1.2)
4.1.1)

ase subunit 1 (EC 6.1.1.13)

rase subunit B (EC 2.8.3.9)
protein
drolase (EC 3.5.2.12)
NrdI
ferase component (EC 4.1.3.27) @ Para-aminobenzoate synthase, amidotransferase component (EC 2.6.1.85) # TrpAb@PabAb

n

tory protein PhoU

(aerobic), beta subunit (EC 1.17.4.1)

Protein (RALP)
ases

sine phosphatase (EC 3.1.3.48)
transferase (EC 2.7.7.9)
n
mily
containing a URI domain
n
e)
cus), ATP-binding protein
n
nce regulon ComE

tase (EC 1.2.1.41)

tein (putative)

ferase (EC 2.4.99.-)

ɔ (aerobic), alpha subunit (EC 1.17.4.1)
tein
: Protein (RALP)
[illegible] of predicted niacin ECF transporter
sonate-7-phosphate synthase I beta (EC 2.5.1.54)

access of DNA to ComEA, FIG038316
ulator (Mga)
zyme in heat shock gene cluster, similarity with CPO of BS HemN-type
n
n
lactonase UlaG (EC 3.1.1.-) (L-ascorbate utilization protein G)
( (SpeK)

nage associated

PnuC-like
nease protein
ga subunit (EC 2.7.7.6)
a chain (EC 6.1.1.20)
g protein OppD (TC 3.A.1.5.1)

rase RumA (EC 2.1.1.-) ## LSU rRNA m(5)U1939

e translocation permease
all chain (EC 6.3.5.5)
n

ɔle histidine kinase
4.14.11)

maltase) (EC 2.4.1.25)
ease protein

in / Small-conductance mechanosensitive channel

n
egulator CiaR
n
nily
ily
nyltransferase (EC 2.1.1.171) ## SSU rRNA m(2)G966
1.1)
5)

n
ent alpha-subunit (EC 1.2.4.-)

ily
ycan synthetase] (EC 2.4.1.129)

r protein FtsY (=alpha subunit) (TC 3.A.5.1.1)

6) @ Acetyl-CoA acetyltransferase (EC 2.3.1.9)

n
omboid family)
ne permease (SagH)

ly scaffold protein for SUF system, SufE2
or threonylcarbamoyladenosine t(6)A37 formation in tRNA

D (SagD)

EC 2.5.1.6)

se (EC 3.4.16.4)
thase, KASII (EC 2.3.1.41)
specific DNA methylase
n
ribosomal protein S18p, zinc-independent

rt system, permease 2 component
RlmH
HI0095 (UbiE paralog)

ike family

(aerobic), alpha subunit (EC 1.17.4.1)

in
n

E) family transporter

ng permease - glutamine transport

nding protein
4)

(EC 5.2.1.8); Proteinase maturation protein (RopA)

A-independent DNA replication

)
oside utilization operon, GntR family

luronic acid synthesis (EC 1.1.1.22)

otein
n
ated
e competence protein ComEB
(EC 2.7.1.35)
tein RecO

.1.7)
n
o (aerobic), beta subunit (EC 1.17.4.1)

se (EC 3.4.16.4)

ily
e (EC 4.1.99.3)
y
sferase (EC 2.1.1.-)
se protein MsbA
le for the formation of Gal-Glc-DAG
otein
(EC 2.4.2.10)

tyltransferase

otein ligase (EC 6.3.4.15)

n

chain (EC 6.1.1.20)
;
in

utative), YggT family

n

unit (EC 3.1.11.6)
e murein endopeptidase

.6.3.14)

idine synthase A (EC 4.2.1.70)

NusB

e aminotransferase [isomerizing] (EC 2.6.1.16)
ciated

ocessing protease

em, ATP-binding protein OpuAA (EC 3.6.3.32)

ed protein

n
(EC 3.1.3.2)

V family

nerase, FMN-dependent (EC 5.3.3.2)
'A
tein
dine synthase B (EC 4.2.1.70)
MutS2
idine kinase CiaH (EC 2.7.3.-)

ase (EC 4.1.1.23)
n

se protein MsbA
C 2.7.7.7)
rotein precursor (TC 3.A.1.14.3)
I (anaerobic), large subunit (EC 1.17.4.2)
e (EC 2.4.2.7)
C 2.4.1.-)

n

.12.4)

inhibitor
.1)
n
ein
hyldihydropteridine pyrophosphokinase (EC 2.7.6.3)
ein

ng/permease protein

'BBR Bacillus subtilis homolog
rboxylase (EC 6.3.4.14)

n

017
rter, permease protein MalF
e component (E2) of acetoin dehydrogenase complex (EC 2.3.1.-)

se component YidC, short form OxaI-like
I family

21.89)
nily
drolase like

n
n
erase alpha chain (EC 6.4.1.2)

e (EC 5.1.3.-)
arboxylase
.3.4)
se I beta (EC 1.1.1.282)
hate reductase (EC 1.17.4.2)

osphohydrolase (EC 3.1.5.1)

quired for reduction of Ribonucleotide reductase class Ib
in
n

, required for cell wall metabolism and coordination of cell division with DNA replication

iacylglycerol kinase catalytic domain]
ısferase (EC 2.1.1.207)
opic repressor codY
ıage associated
.6.3.14)
uctase (EC 1.1.1.100)
ı N-terminal domain / C-terminal domain of CinA type S
ɔinding protein implicated in transcription termination
nase component (EC 2.6.1.85) / Aminodeoxychorismate lyase (EC 4.1.3.38)

EC 3.2.2.20)
4.-)
GTPase RbgA (B. subtilis YlqF)

ɔtein
T (COG4708) of predicted queuosine-regulated ECF transporter

in
rase subunit A (EC 2.8.3.9)
sglycosylase (EC 2.4.1.129) (EC 3.4.-.-)

6)
protein
gene causing stop codon at 85th aa
ıease protein
ciated with MetSO reductase

ctose operon, DeoR family
n
n

ressor MalR, LacI family
sm regulator CmbR, LysR family

ein CcdA homolog, associated with MetSO reductase
nily

oid family)
Dps / Iron-binding ferritin-like antioxidant protein / Ferroxidase (EC 1.16.3.1)
015
29)
n
sferase TrmN6 (EC 2.1.1.223)
3)
n
MutR family

ComX

ly

em, permease protein OpuAB / Glycine betaine ABC transport system, glycine betaine-binding protein OpuAC
omboid family)

l
n

r, associated with ferric iron transporter, SPy1062 homolog
3.6.4.12)

:7)
rotein CorA

o/CI family
n
epressor
ane permease protein ZnuB

rotein
rS / deletion of T at 1407 causing a early stop codon at 469th aa)
ession IbrA

)

ase protein PstC (TC 3.A.1.7.1)
n
n

ransferase (EC 2.6.1.42)
3)

6) @ Acetyl-CoA acetyltransferase (EC 2.3.1.9)
e family protein
n ArgR, repressor of arg regulon
2) / Folylpolyglutamate synthase (EC 6.3.2.17)
ional attenuator LytR-CpsA-Psr, subfamily F2 (as in PMID19099556)
.1.10)

C (SagC)
asmic-binding protein SitA

ase (EC 3.5.99.6)
C 2.7.7.7)

n
uclease RecJ (EC 3.1.-.-)
0)
(EC 4.1.2.40)

n
nsferase, involved in cell wall biogenesis

.9)
n
n

type 1
bs2121
h Ecs transporter (in Streptococci)

thetase (EC 6.3.2.5)
otein
port regulator MtaR, LysR family
5.3.2.-); Xylose transport system permease protein xylH
C 2.1.2.1)

ated

ate O-acyltransferase PlsY

n
ctase MsrA (EC 1.8.4.11) / Peptide methionine sulfoxide reductase MsrB (EC 1.8.4.12)
rt system, ATPase component

ogenase (EC 1.1.1.205)
n
aining multiple CBS domains
n

e YoaA
(IdeS) / Mac/ Secreted immunoglobulin binding protein (Sib38)
n

ate-binding protein
ase (EC 4.1.1.33)

hage associated
2.1.22)

uctase

protein Smf possibly involved in DNA uptake

'.7.21) (EC 2.7.7.25)

omponent (EC 2.7.1.69)
pecific nuclease) (EC 3.1.30.-)
n
4-24
ase A, Bacillus type
related protein
n
ntaining CBS domains
of acetoin dehydrogenase (EC 1.8.1.4)

n
ase (EC 2.3.1.-)
, contains double-stranded beta-helix domain

n
nerly thought to be RNase BN
iated with purine metabolism

se component YidC, OxaA protein

-specific IIB component (EC 2.7.1.69)
: [C1]
h PTS regulation domain, SPy0181 ortholog
lactate transferase like, CofD-like

.6.3.14)

n

superfamily

:ein B
access of DNA to ComEA, FIG007487
FIG068335
3.2.1.26)

:ein NusG
n

n

ng/permease protein

e (EC 2.7.1.45)
in YvcR

ption termination protein NusA
tor, PadR family
oly involved in cell wall localization and side chain formation of rhamnose-glucose polysaccharide

nine(1519)-N(6))-dimethyltransferase (EC 2.1.1.182)

y membrane protease subunit YbbK
n
d for threonylcarbamoyladenosine t(6)A37 modification in tRNA
se (EC 3.4.16.4)
in

DNA transport

n
3.2.4)
le for the formation of Glc-DAG
BC transporter ATP-binding protein ComA
with beta-lactamase, GntR family
n
orter permease component PotB (TC 3.A.1.11.1)
utamyl-2,6-diaminopimelate--D-alanyl-D-alanine ligase (EC 6.3.2.10)
ate synthase (EC 2.5.1.19)
ein EcsA
protein DnaA
essor protein

n

n
4.2.1.46)
eotidohydrolase (EC 3.6.1.23)
on protein PhnA

n ArgR, repressor of arg regulon

n SufD
ns nucleotide-binding domain of DisA bacterial checkpoint controller
phosphate dehydrogenase (EC 1.2.1.12) / Plasmin(ogen) receptor

d phosphatase
PC Superfamily
(EC 3.2.1.-)
)

n

in
ɔunit (EC 3.1.11.6)

ecific IIC component (EC 2.7.1.69) / PTS system, maltose and glucose-specific IIB component (EC 2.7.1.69) / PTS system, maltose and glucose-specific IIA co
n
em, DNA-methyltransferase subunit M (EC 2.1.1.72)
pecific nuclease) (EC 3.1.30.-)
mily
NrdI

idine synthase A (EC 4.2.1.70)
2.1.3.2)

se (EC 2.4.2.22)
nily
(EC 3.1.26.5)
ed
ne cyclo-ligase (EC 6.3.3.1)

n - phage associated

.3)

2.25)

n

EscB

protein] dehydratase (EC 4.2.1.-)
n

.4.2)
ein

protein
EC 2.4.2.14)
nine-binding protein/permease protein

erase beta chain (EC 6.4.1.2)
G (SpeG)
like
); Acetaldehyde dehydrogenase (EC 1.2.1.10)

MutR family
r yesN, associated with MetSO reductase

(EC 2.7.7.41)
export of O-antigen, teichoic acid lipoteichoic acids
n
sine (m7G) methyltransferase GidB

rboxylase catalytic subunit (EC 4.1.1.21)
Streptopain) (EC 3.4.22.10) / Streptococcal pyrogenic exotoxin B (SpeB)

ctase MsrB (EC 1.8.4.12)
J of riboflavin ECF transporter
nding protein

tein RecF
plasmic oligopeptide-binding protein OppA (TC 3.A.1.5.1)
9.1.-)

ı1 (EC 3.5.1.28)

ı SufB
'PcrA
(EC 3.4.19.3)
/I (EC 2.7.3.-), associated with MetSO reductase

ıase (EC 1.1.1.125)
MraW methylase family (EC 2.1.1.-)
cetyl-CoA carboxylase
08
tase family, subgroup 1
C 1.1.3.21)

acid-binding protein
n

.2.1.70)
R family
n
nzyme (EC 1.97.1.4)

ily

ited
sferase PrmC, methylates polypeptide chain release factors RF1 and RF2

n

ylase (EC 3.2.2.23)
n
7.6.2)

unit-related enzymes
nponent (EC 2.7.1.69) / PTS system, sucrose-specific IIC component (EC 2.7.1.69) / PTS system, sucrose-specific IIA component (EC 2.7.1.69)

gizing module of ECF transporters
E (TC 3.A.5.1.1)

; protein FhuC (TC 3.A.1.14.3)
5) / Deoxyguanosine kinase (EC 2.7.1.113)
1.2)
ated)
otein implicated in the recycling of the 50S subunit (S4 paralog)
.5.1.75)
nponent (EC 2.7.1.69) / PTS system, lactose-specific IIC component (EC 2.7.1.69)
n
porter ATPase
ional attenuator LytR-CpsA-Psr, subfamily F1 (as in PMID19099556)
n
I (anaerobic), activating protein (EC 1.97.1.4)

›-8-oxoguanine-triphosphatase) (EC 3.6.1.-)
n
Zn(2+)-responsive expression
e (EC 2.7.3.-)
n
ated
/
aminidase (EndoS)
anine transfer protein DltD

n
n
n
ly
o cobyric acid synthase
n
B (SagB)

7.1.144)

ates the switch between cylindrical and septal cell wall synthesis by re-localization of PBP1

Phosphoribosylaminoimidazolecarboxamide formyltransferase (EC 2.1.2.3)
Antiporter
3.4.4)

nily IIIA

›tional regulator NrdR

rotein CorA

repressor
ribosomal protein L31p, zinc-independent

;)
n

protein (class beta)

ulator (AT-rich DNA-binding protein)
ssor HrcA
3.3)

ase (EC 2.4.2.11)

d)

deacetylase (EC 3.5.1.25)
.1.1)
family
e IIc/interrupted by large phage
nbinase
e)
595 of predicted ECF transporter
ily

export protein

family
n

.6.3.14)
nily

se (EC 6.3.4.13)

essor of hyaluronate and KDG utilization
ical protein
LacB subunit (EC 5.3.1.26)
EzrA
thetase (EC 2.5.1.6)

n
@ SSU ribosomal protein S14p (S29e), zinc-dependent
n
n

.6.1.27)
2.7.4.6)
n

)
n
transferase (EC 2.7.7.9)
.6.3.14)

, periplasmic substrate-binding component (TC 3.A.1.5.2)

rotein PstB (TC 3.A.1.7.1)
embrane-spanning permease protein MsmG
11.4)
n ArgR, repressor of arg regulon

ein

lease

n
binase
n

9.1.-)
tionally linked with Alanyl-tRNA synthetase
n
bulin-binding protein (GRAB)

.5)
n
ress-responsive transcriptional regulator
5.1.1.2)
ein
ily

C, regulator of stationary phase
C 3.2.2.-)
5.1.1.6)
ohydrolase (NADGH) (EC 3.2.2.5)

brane protein

n

cetyltransferase (EC 2.3.1.-)

k protein
n

7.1.144)
d)
(EC 3.6.1.-)
n

abinase

ne synthase, synthetase subunit (EC 6.3.5.3) / Phosphoribosylformylglycinamidine synthase, glutamine amidotransferase subunit (EC 6.3.5.3)

eptidase
e YggF
n
n

ng protein drrA
anM/pseudogene causing stop codon at 177th aa

nily
rboxylase ATPase subunit (EC 4.1.1.21)

phocoenzyme-A synthase (EC 2.7.8.25)
protein

e)
EC 2.7.7.7)
6.3.8)
n
n

de-lysine N(6)-alanyltransferase (EC 2.3.2.10) (EC 2.3.2.-)

NusA

ogenase (NADP+) (EC 1.5.1.5) / Methenyltetrahydrofolate cyclohydrolase (EC 3.5.4.9)
tein

em specificity subunit
ransporting ATPase (EC 3.6.3.3) (EC 3.6.3.5); Copper-translocating P-type ATPase (EC 3.6.3.4)
xyvinyltransferase (EC 2.5.1.7)

n
n

(EC 5.2.1.8)

n
RNA; SSU rRNA
omponent (EC 2.7.1.69)
erone GroEL

Γ of predicted pantothenate ECF transporter
n

omponent (EC 2.7.1.69)
rotein PstB (TC 3.A.1.7.1)

n
in uup
ase [NAD(P)+] (EC 1.1.1.94)

tor
RNA; SSU rRNA
RNA; SSU rRNA

erase [decarboxylating] (EC 2.4.2.19)
RNA; SSU rRNA
RNA; SSU rRNA
ein
lar to C-terminal domain of ribosome protection-type Tc-resistance proteins
ive channel
membrane permease protein SitD

nidine putrescine-binding protein PotD (TC 3.A.1.11.1)

repressor

ase PrpC, regulation of stationary phase
n

G (TC 3.A.5.1.1)
se subunit A (EC 6.3.5.6) @ Glutamyl-tRNA(Gln) amidotransferase subunit A (EC 6.3.5.7)
/
sor SPs0273

ıage associated
n
n
IN adenylyltransferase (EC 2.7.7.2)
ng enzyme, evidenced by COGnitor; PaaD-like protein (DUF59) involved in Fe-S cluster assembly
n
se 2; Mitogenic factor 2
5), (p)ppGpp synthetase I

operon

.6.3.14)

RNA; SSU rRNA
(EC 4.1.2.40)
em, permease component
in

5), bacterial type
glutamate ligase (EC 6.3.2.9)

e (EC 3.4.21.-)
n

DNA receptor
se (EC 3.4.16.4)

(former ThiI)
ıage associated
(EC 6.1.1.14)

omponent (EC 2.7.1.69)

C 4.2.1.10)
n

/ protein
n
BC transporter permease protein ComB
: in RNA processing)

lneuraminate transport system permease protein 2
n

n
rotein DppF (TC 3.A.1.5.2)
family

mrA
LacA subunit (EC 5.3.1.26)
NA; LSU rRNA
ein

arboxylase (EC 4.1.1.36)
access of DNA to ComEA
nily
NA; LSU rRNA
n
C 5.3.1.9)
nponent (EC 2.7.1.69)
otein

n

iosynthesis in gram-positives
n
NA; LSU rRNA
nily
NA; LSU rRNA

n

NA; LSU rRNA
NA; LSU rRNA
tein
in

n
ine ligase @ tRNA-dependent lipid II-Ser--L-alanine ligase
eral energizing module of ECF transporters
C 2.4.2.-)

phage associated

M3_1856/SPs1852
e 2-epimerase (EC 5.1.3.9)
n

unogenic protein

osphoribosyltransferase domain; protein YhgH required for utilization of DNA as sole source of carbon and energy
se subunit C (EC 6.3.5.6) @ Glutamyl-tRNA(Gln) amidotransferase subunit C (EC 6.3.5.7)

n
strate-binding protein
regulon ComD

odiesterase (EC 3.1.4.46)
1.1)
o/CI family
lneuraminate transport system permease protein 1
ase H

.1.1)
in
n

n

n

otein CzcD
7.9.1)

-specific IID component (EC 2.7.1.69)

charide synthesis, competence and biofilm formation Ftr, XRE family

sporter
lmuramyl-(pentapeptide) pyrophosphoryl-undecaprenol N-acetylglucosamine transferase (EC 2.4.1.227)

mI

5' exonuclease)
d)

nobacillus protein (1944168)

C 4.1.2.4)
ccharide synthesis, competence and biofilm formation Ftr, XRE family
transferase (EC 2.1.2.2)
sglycosylase (EC 2.4.1.129) (EC 3.4.-.-)

n
nage associated

mponent (EC 2.7.1.69) / PTS system, fructose-specific IIB component (EC 2.7.1.69) / PTS system, fructose-specific IIC component (EC 2.7.1.69)

n

693 of energizing module of predicted ECF transporter

abilizes FtsL against RasP cleavage
eA
SecA ATPase RNA helicase (TC 3.A.5.1.1)
n

C 1.-.-.-)
1.1.22)
ers

n
nase, malate (EC 2.7.3.-)

etence MecA

n
nomethyl modification enzyme GidA
tein A

ated
n
anine amidase (EC 3.5.1.28)
yltransferase N (EC 2.1.1.-)

- and galactosamine-specific IIA component (EC 2.7.1.69)
ing protein)
ulator (Mga)
clustered with an amidotransferase

n

nine-binding protein/permease protein

nyltransferase (EC 2.1.1.176)
n
peptide-transferase (EC 2.7.8.13)

n

dA, haloacid dehalogenase-like phosphatase family

ferase (EC 2.7.7.3)
)
n

C 2.1.3.3)
RNA-binding domain)
rotein

in RecT (prophage associated)

rt system, permease component
n

: Ffh SRP54 (TC 3.A.5.1.1)
ccess of DNA to ComEA, FIG012620
n
ase)
C 2.7.7.7)

protein dnaD
n

e IIc (The first GCT-Ala is overlapped with the last codon of the former half interrupted protein)
.2.1.85)
J2 of predicted pyridoxin-related ECF transporter

n
n
rotein

LacA subunit (EC 5.3.1.26)
n

ılator, ArpU family
BC transporter protein ProX (TC 3.A.1.12.1) / Osmotic adaptation
4)

)

ıbinase
2.6) (EC 2.7.1.41)

FIG075573
e-binding protein
C, implicated in cell division based on FtsL cleavage
copper transport operon
nily protein)

..16)
em, specificity subunit S (EC 3.1.21.3)

ccinocarboxamide synthase (EC 6.3.2.6)
se (EC 2.5.1.31)
nily
: 3.6.4.12)
(EC 2.1.1.72)

ase protein PstA (TC 3.A.1.7.1)
n
LacB subunit (EC 5.3.1.26)

n
r

raC family

erase [decarboxylating] (EC 2.4.2.19)
component (EC 2.7.1.69)
nily
.2.1)
ase protein DppB (TC 3.A.1.5.2)

yvinyltransferase (EC 2.5.1.7)

component (EC 2.7.1.-)
C 5.3.1.8)
Z (SmeZ)
port system permease protein ProV (TC 3.A.1.12.1)
sglycosylase (EC 2.4.1.129) (EC 3.4.-.-) / Penicillin-binding protein 1A/1B (PBP1)

aining kinase
se subunit B (EC 6.3.5.6) @ Glutamyl-tRNA(Gln) amidotransferase subunit B (EC 6.3.5.7)
M (EC 2.7.3.-)
ulose-uronate ketol-isomerase (EC 5.3.1.17)
n
d)
n
ated
6.1.13)

component

eductase (EC 1.-.-.-) / Dihydropteridine reductase (EC 1.5.1.34)
(EC 2.7.7.7)

Family)

(EC 1.1.1.25)
embrane-spanning permease protein MsmF
mily
n
ein
R family
t (TC 3.A.5.1.1)
(EC 6.1.1.14)
access of DNA to ComEA

(EC 4.1.3.16) @ 2-dehydro-3-deoxyphosphogluconate aldolase (EC 4.1.2.14)
rter permease component potC (TC_3.A.1.11.1)

447
BC transporter ATP-binding protein ComA
n

(EC 3.1.2.6)
J of predicted pyridoxine ECF transporter

ComX
.6.3.14)
n
hage associated
e (EC 3.6.3.4)
nily

nding protein SitB

ragment)

in TrkA
4.1.21) / Autoinducer-2 production protein LuxS
eG
xin protein

21.-)
ufS subfamily

igase @ tRNA-dependent lipid II--L-serine ligase
sociated
7.1.60)
n

grammed frameshift-containing

ase I

n
-specific IIC component (EC 2.7.1.69)
IS-, AP-, NP- specific)
1.1.22)

ase protein DppC (TC 3.A.1.5.2)

d DNA binding protein
rotein DppD (TC 3.A.1.5.2)
n

EC 2.4.2.9) / Pyrimidine operon regulatory protein PyrR

n
.6.3.14)
)
e PlsX
r, malate (EC 2.7.3.-)

5)
2.1.1.-)
n
n
EC 2.5.1.90) / Dimethylallyltransferase (EC 2.5.1.1) / (2E,6E)-farnesyl diphosphate synthase (EC 2.5.1.10) / Geranylgeranyl diphosphate synthase (EC 2.5.1.2

.1.69)

mponent (EC 2.7.1.69)

9)

| genes | logFC.wildSK | logCPM.wildSK | statistic.wildSK | Pvalue.wildSK | FDR.wildSK | function |
|---|---|---|---|---|---|---|
| AP53_832 | -2.089060634 | 9.839306377 | 59.07930293 | 7.64E-06 | 0.01160193 | LSU ribosomal protein |
| AP53_618 | -2.351559093 | 4.72583804 | 40.14312293 | 4.70E-05 | 0.01160193 | hypothetical phage pro |
| AP53_318 | 0.691370775 | 10.427855 | 37.49123272 | 6.39E-05 | 0.01160193 | rRNA small subunit met |
| AP53_848 | 2.596198738 | 7.052381792 | 37.04244937 | 6.74E-05 | 0.01160193 | NAD-dependent malic e |
| AP53_612 | -2.097297492 | 4.914644896 | 36.71078552 | 7.02E-05 | 0.01160193 | Phage base plate |
| AP53_1483 | -1.298695612 | 9.419940924 | 35.99534239 | 7.66E-05 | 0.01160193 | 3-oxoacyl-[acyl-carrier |
| AP53_1015 | -1.907591787 | 6.161086585 | 35.80120002 | 7.85E-05 | 0.01160193 | Phage protein |
| AP53_613 | -2.421455977 | 5.300012505 | 35.728621 | 7.92E-05 | 0.01160193 | Phage major tail protei |
| AP53_1867 | -1.178475919 | 7.390902448 | 35.70730704 | 7.94E-05 | 0.01160193 | small RNA fasX |
| AP53_766 | -1.311417871 | 9.494367576 | 35.57816062 | 8.07E-05 | 0.01160193 | FIG01115306: hypothe |
| AP53_1107 | -0.765794255 | 7.126780371 | 35.54017675 | 8.10E-05 | 0.01160193 | 23S rRNA (Uracil-5-) -m |
| AP53_66 | -1.594970325 | 10.13338686 | 35.18885335 | 8.47E-05 | 0.01160193 | LSU ribosomal protein |
| AP53_1494 | 0.656859383 | 9.042313129 | 34.7266374 | 8.98E-05 | 0.01160193 | putative N-acetyl-mura |
| AP53_425 | -1.275265181 | 7.428724349 | 33.84349927 | 0.000100489 | 0.01160193 | Threonyl-tRNA syntheta |
| AP53_1023 | -2.53158498 | 7.650861882 | 33.77062399 | 0.00010144 | 0.01160193 | Phage capsid protein |
| AP53_620 | -2.106397546 | 4.71110218 | 32.17618244 | 0.00012516 | 0.01160193 | Prophage Clp protease- |
| AP53_1482 | -1.63912747 | 10.11432222 | 31.75140974 | 0.000132543 | 0.01160193 | 3-oxoacyl-[acyl-carrier- |
| AP53_492 | -0.958863589 | 7.705728591 | 31.56666409 | 0.000135914 | 0.01160193 | Oligoendopeptidase F |
| AP53_831 | -2.040127454 | 9.968418741 | 31.41428641 | 0.00013877 | 0.01160193 | LSU ribosomal protein |
| AP53_1617 | -0.73404363 | 6.251830487 | 31.29055107 | 0.000141141 | 0.01160193 | 6-phospho-beta-galact |
| AP53_1017 | -2.171511076 | 6.274401588 | 31.23472212 | 0.000142226 | 0.01160193 | Phage major tail protei |
| AP53_1643 | -1.266604552 | 8.456063552 | 31.08429749 | 0.0001452 | 0.01160193 | Polyribonucleotide nuc |
| AP53_619 | -2.362818649 | 5.091305753 | 30.95787487 | 0.000147756 | 0.01160193 | Phage major capsid pro |
| AP53_1471 | -1.079416794 | 8.431843594 | 30.74989955 | 0.000152077 | 0.01160193 | PTS system, mannose-s |
| AP53_1237 | -0.790373219 | 9.540523851 | 30.69189175 | 0.000153309 | 0.01160193 | Cell division initiation |
| AP53_1481 | -1.439289384 | 10.0964202 | 30.38587407 | 0.000160004 | 0.01160193 | Biotin carboxyl carrier |
| AP53_1267 | -1.547189377 | 11.23135134 | 30.32452781 | 0.000161387 | 0.01160193 | Arginine deiminase (EC |
| AP53_1476 | -0.698875033 | 7.639481239 | 29.04338183 | 0.00019376 | 0.012277899 | Seryl-tRNA synthetase ( |
| AP53_1142 | -1.098166697 | 8.075777564 | 28.93523859 | 0.000196827 | 0.012277899 | Alanyl-tRNA synthetase |
| AP53_1565 | -1.746253124 | 7.738902507 | 28.79794957 | 0.000200803 | 0.012277899 | transporter |
| AP53_617 | -2.198443828 | 4.683457916 | 28.67320863 | 0.000204498 | 0.012277899 | hypothetical phage pro |
| AP53_1478 | -1.073559275 | 10.06615235 | 28.64412453 | 0.000205371 | 0.012277899 | Acetyl-coenzyme A carb |
| AP53_1315 | -1.451882888 | 3.418809219 | 28.46899624 | 0.000210721 | 0.012277899 | FIG01116522: hypothe |
| AP53_1428 | -0.842654228 | 7.56840856 | 28.16305029 | 0.000220467 | 0.012277899 | FIG01114118: hypothe |

| | | | | | | |
|---|---|---|---|---|---|---|
| AP53_1480 | -1.08953404 | 9.729546962 | 27.94096279 | 0.000227875 | 0.012277899 | (3R)-hydroxymyristoyl- |
| AP53_847 | 2.070087404 | 6.940372562 | 27.82595937 | 0.000231826 | 0.012277899 | Malate Na(+) symporter |
| AP53_1631 | -1.067177392 | 11.02108569 | 27.39454824 | 0.000247383 | 0.012277899 | LSU ribosomal protein I |
| AP53_614 | -2.346098416 | 5.801452265 | 27.37104908 | 0.000248265 | 0.012277899 | hypothetical phage pro |
| AP53_1719 | 0.895780852 | 6.55302028 | 27.31394973 | 0.000250424 | 0.012277899 | PTS system, cellobiose-s |
| AP53_1236 | -0.869376709 | 7.985314977 | 27.17088205 | 0.000255931 | 0.012277899 | Isoleucyl-tRNA syntheta |
| AP53_665 | 0.619413901 | 8.344185197 | 27.08392148 | 0.000259348 | 0.012277899 | Cytidylate kinase (EC 2.7 |
| AP53_70 | -1.313524919 | 10.44975777 | 26.75696575 | 0.000272684 | 0.012340526 | LSU ribosomal protein I |
| AP53_47 | -0.970677613 | 10.83978006 | 26.72626769 | 0.000273978 | 0.012340526 | SSU ribosomal protein S |
| AP53_81 | -1.540936202 | 9.263004421 | 26.1368386 | 0.000300265 | 0.012340526 | DNA-directed RNA poly |
| AP53_694 | 0.569302757 | 8.383306483 | 25.99247977 | 0.000307148 | 0.012340526 | Glycerophosphoryl dies |
| AP53_508 | 0.694116505 | 5.896771962 | 25.71251818 | 0.000321035 | 0.012340526 | PTS system, N-acetylgal |
| AP53_559 | -1.26110618 | 9.645722823 | 25.64099106 | 0.000324701 | 0.012340526 | ATP synthase C chain (E |
| AP53_1130 | -1.297309908 | 12.05479986 | 25.49585968 | 0.000332291 | 0.012340526 | Phosphocarrier protein |
| AP53_1314 | -1.204161637 | 4.001388929 | 25.27401535 | 0.0003443 | 0.012340526 | Phage protein |
| AP53_1675 | -0.822163185 | 5.85093154 | 25.25278428 | 0.000345476 | 0.012340526 | FIG01115320: hypothe |
| AP53_48 | -1.101568739 | 11.44172855 | 25.14380548 | 0.000351587 | 0.012340526 | LSU ribosomal protein I |
| AP53_1408 | -0.877255497 | 8.416879746 | 25.12287251 | 0.000352775 | 0.012340526 | ATP-dependent RNA hel |
| AP53_1736 | 0.66546485 | 13.53520806 | 25.07048996 | 0.00035577 | 0.012340526 | Heat shock protein 60 f |
| AP53_1848 | -0.496154399 | 8.29535345 | 25.0418727 | 0.000357418 | 0.012340526 | Hyaluronan synthase (E |
| AP53_1729 | -0.515213609 | 8.363552599 | 25.0001687 | 0.000359837 | 0.012340526 | FIG01115477: hypothe |
| AP53_1756 | -1.109078557 | 9.032149695 | 24.90300647 | 0.000365546 | 0.012340526 | Neutral endopeptidase |
| AP53_1024 | -2.451910027 | 7.609041708 | 24.75514432 | 0.000374442 | 0.012340526 | Prophage Clp protease- |
| AP53_433 | -0.659251525 | 8.225934195 | 24.7509337 | 0.000374699 | 0.012340526 | Two-component respor |
| AP53_887 | -0.918146128 | 10.0984361 | 24.74418056 | 0.000375111 | 0.012340526 | L-lactate dehydrogenas |
| AP53_1490 | 0.966495229 | 13.61212773 | 24.54169685 | 0.000387736 | 0.012543256 | Chaperone protein Dna |
| AP53_64 | -1.425752426 | 7.801527973 | 24.3449569 | 0.000400484 | 0.012638048 | Adenylate kinase (EC 2.7 |
| AP53_1011 | -1.248900162 | 5.389861213 | 24.29666553 | 0.000403688 | 0.012638048 | Hyaluronate lyase (phag |
| AP53_1028 | -1.96425251 | 5.282694774 | 24.07486553 | 0.000418798 | 0.012902972 | Phage terminase, large s |
| AP53_1265 | -1.359013982 | 12.06436896 | 23.85695817 | 0.000434295 | 0.013171364 | Ornithine carbamoyltra |
| AP53_1861 | 0.569645631 | 7.569565934 | 23.75421499 | 0.000441836 | 0.013193902 | Mobile element protein |
| AP53_344 | -0.963598294 | 7.517459375 | 23.21398607 | 0.000484116 | 0.013742853 | FIG01114304: hypothe |
| AP53_1039 | -0.915034324 | 5.831925291 | 23.08355517 | 0.000495027 | 0.013742853 | putative replication pro |
| AP53_1642 | -0.740866736 | 8.132237723 | 23.05479001 | 0.000497472 | 0.013742853 | FIG01114029: hypothe |
| AP53_1479 | -1.069534077 | 10.1080263 | 23.0481573 | 0.000498038 | 0.013742853 | Biotin carboxylase of ac |

| | | | | | | |
|---|---|---|---|---|---|---|
| AP53_1012 | -2.013973894 | 5.87453472 | 22.9775353 | 0.00050411 | 0.013742853 | Phage hyaluronidase; P |
| AP53_1150 | -0.694803786 | 7.68102015 | 22.9491367 | 0.000506576 | 0.013742853 | S-adenosylmethionine: |
| AP53_1010 | -1.385498299 | 5.822470884 | 22.91246656 | 0.000509781 | 0.013742853 | Phage hyaluronidase |
| AP53_1018 | -1.970100941 | 5.920351528 | 22.74238174 | 0.000524965 | 0.01378326 | Phage protein |
| AP53_1436 | -1.396420797 | 7.919112504 | 22.71864624 | 0.000527126 | 0.01378326 | DegV family protein |
| AP53_615 | -1.998274709 | 5.69586088 | 22.65920781 | 0.000532583 | 0.01378326 | hypothetical phage pro |
| AP53_1008 | -1.473799839 | 5.425898908 | 22.4939344 | 0.000548112 | 0.013998494 | Phage protein |
| AP53_67 | -1.356067839 | 10.20321776 | 22.15223557 | 0.000581941 | 0.014669439 | SSU ribosomal protein S |
| AP53_573 | -0.881496698 | 6.896267721 | 22.05061528 | 0.000592472 | 0.014743425 | Phenylalanyl-tRNA synt |
| AP53_69 | -1.344710733 | 10.49884439 | 21.87694725 | 0.000610993 | 0.014809411 | DNA-directed RNA poly |
| AP53_1021 | -2.457936072 | 6.68383023 | 21.85031065 | 0.000613893 | 0.014809411 | DNA packaging protein, |
| AP53_1681 | -0.657772914 | 8.463565769 | 21.75931502 | 0.000623926 | 0.014809411 | Dipeptide-binding ABC |
| AP53_1671 | 0.948958403 | 6.337641092 | 21.68444211 | 0.000632326 | 0.014809411 | Exodeoxyribonuclease I |
| AP53_1531 | 0.870149095 | 9.761708956 | 21.53662364 | 0.000649303 | 0.014809411 | Secreted Endo-beta-N-a |
| AP53_496 | -0.823975696 | 12.88864028 | 21.45318217 | 0.000659124 | 0.014809411 | Translation elongation |
| AP53_1485 | -1.164662262 | 9.463774976 | 21.40358891 | 0.000665043 | 0.014809411 | Enoyl-[acyl-carrier-prot |
| AP53_1295 | 0.8413297 | 7.199351983 | 21.34469009 | 0.000672155 | 0.014809411 | Multiple sugar ABC tran |
| AP53_1533 | 0.630552546 | 8.145137342 | 21.33937074 | 0.000672802 | 0.014809411 | Sucrose-6-phosphate hy |
| AP53_1013 | -2.050763176 | 5.972803166 | 21.32414075 | 0.000674658 | 0.014809411 | Phage-related protein |
| AP53_1462 | -0.498737322 | 7.438063323 | 21.26002873 | 0.000682536 | 0.014809411 | ABC transporter, perme |
| AP53_1651 | -0.930194844 | 8.844024322 | 21.08572867 | 0.000704507 | 0.014809411 | Peptide deformylase (EC |
| AP53_1143 | -0.832329247 | 9.577025249 | 20.96738649 | 0.0007199 | 0.014809411 | Foldase protein prsA 1 p |
| AP53_1484 | -1.177064292 | 9.517232347 | 20.8383511 | 0.000737138 | 0.014809411 | Malonyl CoA-acyl carrie |
| AP53_54 | -0.909812436 | 11.32762119 | 20.83179087 | 0.000738027 | 0.014809411 | LSU ribosomal protein L |
| AP53_922 | 0.50268573 | 9.687420644 | 20.81647278 | 0.000740109 | 0.014809411 | Signal recognition parti |
| AP53_80 | -1.455196393 | 8.787198023 | 20.77161948 | 0.000746243 | 0.014809411 | DNA-directed RNA poly |
| AP53_621 | -1.088851949 | 3.43401959 | 20.74230601 | 0.000750284 | 0.014809411 | Portal protein, phage as |
| AP53_1489 | -1.116118129 | 10.06214315 | 20.70588061 | 0.000755342 | 0.014809411 | Trans-2,cis-3-Decenoyl- |
| AP53_939 | -1.117286398 | 9.212780987 | 20.64554821 | 0.000763809 | 0.014809411 | Biotinyl-lipoyl attachm |
| AP53_169 | 0.564295351 | 9.092234106 | 20.63926137 | 0.000764697 | 0.014809411 | Zinc finger protein |
| AP53_1328 | -1.350553728 | 3.983716311 | 20.63855654 | 0.000764797 | 0.014809411 | Phage major capsid pro |
| AP53_1672 | 0.870260047 | 7.82243254 | 20.54011083 | 0.000778874 | 0.014809411 | PTS system, maltose and |
| AP53_1043 | -0.811095866 | 6.060397034 | 20.51620149 | 0.000782339 | 0.014809411 | FIG01115915: hypothe |
| AP53_50 | -0.998871216 | 11.20930675 | 20.4763596 | 0.000788153 | 0.014809411 | LSU ribosomal protein L |
| AP53_1022 | -2.804011931 | 6.923782671 | 20.36515173 | 0.000804653 | 0.014809411 | conserved hypothetical |

| | | | | | | |
|---|---|---|---|---|---|---|
| AP53_1304 | 0.939731691 | 9.344842128 | 20.35295899 | 0.000806487 | 0.014809411 | Alpha-mannosidase (EC |
| AP53_1709 | 1.183626746 | 6.130877228 | 20.31799343 | 0.000811773 | 0.014809411 | Strepotococcal cystein |
| AP53_616 | -1.653781621 | 5.136891904 | 20.25363769 | 0.000821609 | 0.014809411 | hypothetical phage pro |
| AP53_1045 | -0.843980628 | 6.293677744 | 20.23802269 | 0.000824017 | 0.014809411 | hypothetical phage pro |
| AP53_68 | -1.35349463 | 10.91520246 | 20.18481409 | 0.000832284 | 0.014820757 | SSU ribosomal protein |
| AP53_109 | -1.333148951 | 6.207061618 | 20.09121809 | 0.000847064 | 0.014913079 | Fibronectin-binding pr |
| AP53_1300 | 0.839727503 | 9.157013082 | 20.01367795 | 0.000859543 | 0.014913079 | Beta-glucosidase (EC 3. |
| AP53_1016 | -2.027085879 | 6.137775317 | 20.00767829 | 0.000860518 | 0.014913079 | Phage protein |
| AP53_1026 | -1.64511035 | 5.864797043 | 19.94702616 | 0.000870443 | 0.014913455 | Phage protein |
| AP53_1009 | -1.53796783 | 5.668230454 | 19.87228006 | 0.000882862 | 0.014913455 | FIG01116522: hypothe |
| AP53_1042 | -0.870321835 | 6.407160573 | 19.867939 | 0.00088359 | 0.014913455 | FIG01117510: hypothe |
| AP53_1004 | 0.505171052 | 9.335315697 | 19.71524823 | 0.000909638 | 0.015110515 | Phage lysin |
| AP53_561 | -1.02244563 | 9.705734837 | 19.70835659 | 0.000910835 | 0.015110515 | ATP synthase B chain (E |
| AP53_1464 | -0.501835315 | 8.759238143 | 19.6042131 | 0.000929149 | 0.015283714 | Histidine triad (HIT) nu |
| AP53_1492 | 0.892281647 | 14.03692929 | 19.50599968 | 0.000946819 | 0.015443497 | Heat shock protein Grp |
| AP53_1041 | -0.873343446 | 5.849702813 | 19.4302056 | 0.000960727 | 0.015539757 | FIG01115822: hypothe |
| AP53_1025 | -2.243515612 | 6.219577531 | 19.38277056 | 0.000969553 | 0.015552918 | Phage portal protein |
| AP53_58 | -0.820751842 | 10.54885701 | 19.33938692 | 0.00097771 | 0.015555201 | LSU ribosomal protein |
| AP53_256 | 0.601852391 | 8.209398764 | 19.19933552 | 0.001004597 | 0.015853037 | Methionine ABC transp |
| AP53_53 | -1.140021011 | 11.43545424 | 19.13426808 | 0.001017385 | 0.015925362 | LSU ribosomal protein |
| AP53_622 | -0.997235125 | 3.759174187 | 19.03569092 | 0.001037126 | 0.016053992 | Phage terminase, large |
| AP53_506 | 0.570251913 | 7.599458418 | 19.00666953 | 0.001043024 | 0.016053992 | Transcriptional regulat |
| AP53_565 | -1.313390906 | 10.93748296 | 18.93956791 | 0.001056812 | 0.016053992 | ATP synthase beta chair |
| AP53_1030 | -1.125009633 | 4.945597315 | 18.93052298 | 0.001058687 | 0.016053992 | Phage endonuclease |
| AP53_574 | -0.811659544 | 7.228015948 | 18.73525835 | 0.001100135 | 0.016553198 | Phenylalanyl-tRNA synt |
| AP53_1594 | -0.937794501 | 11.13498404 | 18.63135277 | 0.00112297 | 0.016766801 | Fructose-bisphosphate |
| AP53_623 | -1.203855467 | 3.912249943 | 18.52527434 | 0.00114686 | 0.016992789 | Phage protein |
| AP53_1296 | 0.613354492 | 6.172112517 | 18.44025126 | 0.001166442 | 0.017084913 | Multiple sugar ABC trar |
| AP53_1472 | -1.097473236 | 9.358208714 | 18.42206232 | 0.001170682 | 0.017084913 | PTS system, mannose-s |
| AP53_1463 | -0.540509635 | 7.910418677 | 18.32646246 | 0.001193269 | 0.017284584 | ABC transporter, ATP-bi |
| AP53_563 | -1.243533937 | 10.59655174 | 18.12456239 | 0.00124268 | 0.017866979 | ATP synthase alpha chai |
| AP53_460 | -0.811975101 | 7.599552137 | 18.08611206 | 0.001252361 | 0.017873777 | Hydrolase (HAD superfa |
| AP53_1020 | -2.022952936 | 6.170653395 | 17.94443157 | 0.001288811 | 0.018259719 | Phage protein |
| AP53_820 | -1.460681215 | 9.906872654 | 17.83832939 | 0.001316927 | 0.018522868 | PTS system, mannose/fr |
| AP53_1486 | -0.940625613 | 9.62811879 | 17.63672877 | 0.001372368 | 0.018821016 | Acyl carrier protein |

| | | | | | | |
|---|---|---|---|---|---|---|
| AP53_1234 | 0.635900488 | 9.054293806 | 17.62593246 | 0.001375414 | 0.018821016 | Mutator mutT protein ( |
| AP53_57 | -0.906483178 | 10.59405612 | 17.58610052 | 0.001386721 | 0.018821016 | SSU ribosomal protein S |
| AP53_1708 | 1.013978708 | 6.4287874 | 17.56514223 | 0.001392714 | 0.018821016 | Streptopain inhibitor |
| AP53_1144 | -0.731641579 | 6.785620853 | 17.5581901 | 0.001394708 | 0.018821016 | O-methyltransferase far |
| AP53_587 | 0.475048159 | 9.35155574 | 17.54636526 | 0.001398109 | 0.018821016 | probably aromatic ring |
| AP53_1014 | -1.693518788 | 6.659339264 | 17.51905346 | 0.001406001 | 0.018821016 | Phage tail length tape-n |
| AP53_292 | -0.435478884 | 7.715056802 | 17.46078571 | 0.001423013 | 0.018918279 | Ribosomal large subuni |
| AP53_1019 | -1.772061705 | 7.874398624 | 17.36236233 | 0.001452305 | 0.019058727 | Phage protein |
| AP53_469 | -0.852292261 | 9.808365257 | 17.35933898 | 0.001453216 | 0.019058727 | FIG01116444: hypothe |
| AP53_314 | 0.710159622 | 9.769757424 | 17.21687863 | 0.001496914 | 0.019377937 | DNA polymerase III delt |
| AP53_510 | 0.622995778 | 5.849857875 | 17.21493079 | 0.001497522 | 0.019377937 | PTS system, N-acetylgal |
| AP53_1029 | -1.511407901 | 5.505463083 | 17.03079155 | 0.00155635 | 0.020005802 | Phage terminase, small |
| AP53_1814 | -1.078453136 | 9.414836082 | 16.90534407 | 0.001597991 | 0.02040592 | LSU ribosomal protein L |
| AP53_1079 | -0.781338311 | 6.775970913 | 16.843036 | 0.00161916 | 0.020541114 | Glutamine ABC transpo |
| AP53_56 | -0.877687271 | 10.64711432 | 16.75948781 | 0.001648068 | 0.020655887 | SSU ribosomal protein S |
| AP53_1473 | -0.98165485 | 9.677362925 | 16.75541946 | 0.001649491 | 0.020655887 | PTS system, mannose-sp |
| AP53_545 | -0.861385241 | 12.02450849 | 16.67526696 | 0.001677826 | 0.020876032 | Enolase (EC 4.2.1.11) |
| AP53_1363 | 0.969322523 | 9.67730007 | 16.55094362 | 0.001722921 | 0.021300568 | Chromosome segregatio |
| AP53_760 | -1.213126148 | 10.99692506 | 16.51492153 | 0.001736253 | 0.021326621 | SSU ribosomal protein S |
| AP53_1630 | -1.003931715 | 11.41737697 | 16.48611162 | 0.001747003 | 0.021326621 | SSU ribosomal protein S |
| AP53_1007 | -1.640800436 | 5.788601678 | 16.35703058 | 0.001796143 | 0.021681682 | Phage holin |
| AP53_62 | -0.678084985 | 10.56937826 | 16.35112784 | 0.001798429 | 0.021681682 | LSU ribosomal protein L |
| AP53_919 | 0.532773004 | 9.876042395 | 16.24306071 | 0.001840891 | 0.022056597 | FIG01114660: hypothe |
| AP53_704 | -0.611274026 | 8.096405396 | 16.15364557 | 0.00187692 | 0.022324894 | Membrane protein |
| AP53_1588 | -0.809008636 | 10.15300196 | 16.11252434 | 0.001893768 | 0.022324894 | Phosphoglycerate kinas |
| AP53_41 | 0.507967217 | 7.04652198 | 16.08327279 | 0.001905861 | 0.022324894 | Multi antimicrobial ext |
| AP53_348 | -0.708957315 | 8.534545207 | 16.07501885 | 0.00190929 | 0.022324894 | N-acetylglucosamine-1- |
| AP53_1038 | -0.76417141 | 5.76057865 | 16.02933197 | 0.001928402 | 0.022398814 | hypothetical protein |
| AP53_1216 | -0.838151834 | 11.02528837 | 15.99155696 | 0.001944375 | 0.022398814 | DNA-binding protein HI |
| AP53_1829 | -0.961708591 | 9.962085059 | 15.92168314 | 0.001974333 | 0.022398814 | SSU ribosomal protein S |
| AP53_1322 | -1.037858981 | 2.744344257 | 15.90863716 | 0.001979987 | 0.022398814 | Phage protein |
| AP53_611 | -1.352399614 | 4.925121625 | 15.903736 | 0.001982116 | 0.022398814 | hypothetical protein |
| AP53_1430 | -0.769450041 | 7.962175931 | 15.87132246 | 0.001996263 | 0.022398814 | Glycyl-tRNA synthetase |
| AP53_63 | -0.655818572 | 10.43719894 | 15.87103122 | 0.001996391 | 0.022398814 | Preprotein translocase |
| AP53_127 | -0.532384463 | 6.639887807 | 15.8063531 | 0.002024981 | 0.02258901 | V-type ATP synthase sub |

| | | | | | | |
|---|---|---|---|---|---|---|
| AP53_954 | -1.147208629 | 9.975002689 | 15.66870332 | 0.00208745 | 0.023152806 | SSU ribosomal protein S |
| AP53_491 | -0.483364743 | 7.708053578 | 15.63996506 | 0.002100778 | 0.023168238 | Glutathione peroxidase |
| AP53_1716 | 0.431105845 | 7.147578025 | 15.52366486 | 0.002155748 | 0.023640155 | Glycerol dehydrogenase |
| AP53_46 | -0.972313346 | 11.29190986 | 15.37834015 | 0.002226841 | 0.024282574 | LSU ribosomal protein L |
| AP53_1329 | -1.112888073 | 3.992006818 | 15.30545951 | 0.002263534 | 0.02443816 | Phage protein |
| AP53_1431 | 0.414936746 | 9.994209097 | 15.28363987 | 0.002274658 | 0.02443816 | INTEGRAL MEMBRANE P |
| AP53_36 | 0.462267021 | 9.058116692 | 15.27539007 | 0.00227888 | 0.02443816 | Low molecular weight p |
| AP53_1214 | 0.761808801 | 7.586571696 | 15.18260714 | 0.002327014 | 0.024486289 | Phage protein |
| AP53_1583 | -0.457106929 | 8.319343603 | 15.14926162 | 0.002344605 | 0.024486289 | Protein of unknown fur |
| AP53_1163 | -0.703602844 | 10.45687235 | 15.14774291 | 0.00234541 | 0.024486289 | Cold-shock DEAD-box p |
| AP53_731 | -0.655756916 | 7.025910352 | 15.13582201 | 0.002351739 | 0.024486289 | Hydroxymethylglutaryl |
| AP53_51 | -0.972515471 | 11.07336914 | 15.13272095 | 0.002353389 | 0.024486289 | LSU ribosomal protein L |
| AP53_1647 | -1.041121707 | 5.062154392 | 15.12208082 | 0.00235906 | 0.024486289 | Transcription antitermi |
| AP53_1433 | 0.465268656 | 10.47960649 | 15.06373449 | 0.002390447 | 0.024680089 | N-acetylglucosamine-6- |
| AP53_1618 | -0.565352731 | 7.218910504 | 14.98831129 | 0.002431752 | 0.024973709 | PTS system, lactose-spe |
| AP53_1557 | 0.392509263 | 9.093260226 | 14.89567536 | 0.002483643 | 0.025372369 | Putative membrane pro |
| AP53_111 | 0.91373951 | 6.855954303 | 14.84662584 | 0.002511647 | 0.025430398 | Short chain fatty acids t |
| AP53_1813 | -0.65346663 | 6.865511878 | 14.8211224 | 0.002526355 | 0.025430398 | Histidyl-tRNA synthetas |
| AP53_1076 | -0.734346163 | 8.966213956 | 14.81719596 | 0.002528628 | 0.025430398 | D-alanine--poly(phosph |
| AP53_560 | -1.095855601 | 9.652494913 | 14.742564 | 0.002572303 | 0.025680661 | ATP synthase A chain (E |
| AP53_59 | -0.809737333 | 11.0043086 | 14.72960997 | 0.002579974 | 0.025680661 | LSU ribosomal protein L |
| AP53_1564 | -0.587373239 | 8.421111977 | 14.65257002 | 0.002626157 | 0.025991865 | Aquaporin (Major Intrir |
| AP53_1291 | 0.558394798 | 6.437852788 | 14.61512663 | 0.002648955 | 0.025991865 | Beta-galactosidase (EC 3 |
| AP53_44 | -0.934768791 | 11.31807185 | 14.59163052 | 0.00266338 | 0.025991865 | LSU ribosomal protein L |
| AP53_564 | -1.072974955 | 10.08214382 | 14.57178498 | 0.002675637 | 0.025991865 | ATP synthase gamma ch |
| AP53_177 | -0.597941889 | 8.91860337 | 14.567659 | 0.002678193 | 0.025991865 | Glucose-6-phosphate is |
| AP53_49 | -0.89140104 | 11.30618214 | 14.5124051 | 0.002712709 | 0.026195858 | SSU ribosomal protein S |
| AP53_257 | 0.72373017 | 8.245360951 | 14.47330197 | 0.002737453 | 0.026303939 | Methionine ABC transp |
| AP53_61 | -0.82379875 | 10.31748478 | 14.41601517 | 0.002774186 | 0.026507592 | LSU ribosomal protein L |
| AP53_55 | -0.682453995 | 10.94485035 | 14.39784197 | 0.00278596 | 0.026507592 | LSU ribosomal protein L |
| AP53_1706 | 0.651538944 | 12.43279509 | 14.34586271 | 0.002819964 | 0.02655086 | FIG01114845: hypothe |
| AP53_1641 | -0.551465838 | 8.263963792 | 14.33663768 | 0.00282605 | 0.02655086 | Serine acetyltransferase |
| AP53_455 | 0.637650848 | 10.67283373 | 14.32447597 | 0.002834097 | 0.02655086 | Plasmid stabilization sy |
| AP53_1851 | -0.512592278 | 9.03965394 | 14.2930173 | 0.002855039 | 0.02655086 | UTP--glucose-1-phosph |
| AP53_1327 | -1.198851975 | 3.739674086 | 14.28724245 | 0.002858902 | 0.02655086 | Phage protein |

| | | | | | | |
|---|---|---|---|---|---|---|
| AP53_1562 | -0.397195648 | 7.62799416 | 14.20930911 | 0.002911657 | 0.026777921 | C3-degrading proteinas |
| AP53_254 | -0.752592275 | 8.141883276 | 14.19075293 | 0.002924386 | 0.026777921 | FIG007491: hypothetic |
| AP53_1491 | 0.910261719 | 14.69533013 | 14.16860048 | 0.002939669 | 0.026777921 | Chaperone protein Dna |
| AP53_1722 | 0.581030045 | 9.069333087 | 14.13733618 | 0.002961399 | 0.026777921 | Deoxyribonucleoside re |
| AP53_1735 | 0.751557666 | 7.533997899 | 14.12629412 | 0.002969119 | 0.026777921 | hypothetical protein |
| AP53_45 | -1.001329141 | 11.20939483 | 14.11450866 | 0.002977385 | 0.026777921 | LSU ribosomal protein |
| AP53_758 | -0.57791135 | 7.399991744 | 14.11089802 | 0.002979923 | 0.026777921 | Branched-chain amino |
| AP53_920 | 0.640377897 | 9.684761164 | 14.05450085 | 0.003019897 | 0.027012072 | Tyrosine recombinase X |
| AP53_rna6 | -1.486250228 | 4.56855931 | 14.03172242 | 0.003036221 | 0.02703351 | tRNA-Asp-GTC |
| AP53_516 | 0.416538055 | 6.978985836 | 13.88650989 | 0.003142773 | 0.027692283 | 4-hydroxy-2-oxoglutara |
| AP53_1697 | 0.502018679 | 8.455477189 | 13.87703506 | 0.003149877 | 0.027692283 | Hypothetical mga-asso |
| AP53_1317 | -0.731653097 | 3.704008296 | 13.87286394 | 0.003153011 | 0.027692283 | Hyaluronate lyase (phag |
| AP53_454 | 0.52795274 | 9.919725063 | 13.83926262 | 0.003178389 | 0.027789431 | Plasmid stabilization sy |
| AP53_1152 | 0.499785572 | 7.02683456 | 13.78344954 | 0.003221079 | 0.028036384 | FIG01113992: hypothe |
| AP53_1242 | -0.930272155 | 9.804031108 | 13.57645815 | 0.003385424 | 0.029335303 | Cell division protein Fts |
| AP53_rna7 | -1.739218332 | 4.436492387 | 13.52391893 | 0.003428702 | 0.029578269 | tRNA-Lys-TTT |
| AP53_1425 | 0.491655105 | 7.44292707 | 13.4641712 | 0.003478714 | 0.029876922 | Glycerol kinase (EC 2.7. |
| AP53_509 | 0.584936775 | 5.783510146 | 13.36081654 | 0.003567278 | 0.030502583 | PTS system, N-acetylgal |
| AP53_1073 | -0.624821376 | 9.28714035 | 13.25053362 | 0.003664731 | 0.031198435 | Poly(glycerophosphate |
| AP53_1865 | 1.581511407 | 11.85379858 | 13.2217375 | 0.003690693 | 0.031282248 | Streptolysin S biosynth |
| AP53_302 | 0.448156106 | 9.263777797 | 13.17867806 | 0.003729921 | 0.031477286 | FIG01114725: hypothe |
| AP53_101 | -0.640993328 | 8.357710187 | 13.09397589 | 0.003808532 | 0.032001562 | Chaperonin (heat shock |
| AP53_1488 | -0.937532516 | 10.09266506 | 13.00687309 | 0.003891422 | 0.032426301 | Transcriptional regulat |
| AP53_607 | -0.908579937 | 5.266162481 | 13.00576197 | 0.003892493 | 0.032426301 | Hyaluronate lyase (phag |
| AP53_1710 | 1.008616896 | 4.837104462 | 12.91043149 | 0.003985697 | 0.032952896 | FIG01114305: hypothe |
| AP53_693 | 0.587850349 | 9.317769129 | 12.90643626 | 0.00398966 | 0.032952896 | ABC transporter perme |
| AP53_1674 | -0.696506942 | 7.214747101 | 12.86588289 | 0.004030155 | 0.033146314 | Ribosomal protein L11 |
| AP53_355 | -0.326037719 | 7.784997873 | 12.74922284 | 0.004149382 | 0.033631048 | Manganese ABC transpo |
| AP53_rna75 | -6.716005924 | -0.117416991 | 14.12142103 | 0.004153834 | 0.033631048 | tRNA-Arg-ACG |
| AP53_566 | -0.947476147 | 10.8465092 | 12.73215371 | 0.004167174 | 0.033631048 | ATP synthase epsilon ch |
| AP53_60 | -0.733545146 | 10.31908623 | 12.72774499 | 0.004171784 | 0.033631048 | SSU ribosomal protein |
| AP53_122 | -0.559564145 | 6.032187503 | 12.68666832 | 0.004215028 | 0.033631048 | V-type ATP synthase sub |
| AP53_1320 | -1.238351895 | 3.6637731 | 12.681518 | 0.004220487 | 0.033631048 | Phage tail length tape-r |
| AP53_557 | -0.451802763 | 7.132158754 | 12.68007411 | 0.004222019 | 0.033631048 | DNA ligase (EC 6.5.1.2) |
| AP53_1054 | -1.15825678 | 4.635450138 | 12.67472087 | 0.004227705 | 0.033631048 | Phage transcriptional r |

| | | | | | | |
|---|---|---|---|---|---|---|
| AP53_1362 | 0.785156703 | 9.411929701 | 12.64839625 | 0.004255797 | 0.033716334 | Phage protein |
| AP53_1027 | -1.295836414 | 4.361305901 | 12.59870643 | 0.004309426 | 0.034002421 | hypothetical phage pro |
| AP53_500 | -0.345481956 | 8.026905762 | 12.49246632 | 0.004426791 | 0.034523817 | tRNA-dependent lipid I |
| AP53_52 | -0.900268555 | 11.32480098 | 12.48352714 | 0.004436838 | 0.034523817 | SSU ribosomal protein S |
| AP53_1646 | -1.316242271 | 4.87587575 | 12.47563007 | 0.004445735 | 0.034523817 | PTS system IIB compon |
| AP53_1405 | 0.381252149 | 9.099489907 | 12.47481638 | 0.004446653 | 0.034523817 | ABC-type multidrug tra |
| AP53_234 | -0.407540278 | 8.561751269 | 12.41331723 | 0.004516689 | 0.034927859 | Putative iron-sulfur clu |
| AP53_1837 | -0.511069491 | 8.456360843 | 12.38967983 | 0.004543953 | 0.034999259 | tRNA-specific 2-thiouri |
| AP53_15 | 0.409116442 | 7.727893238 | 12.36955738 | 0.004567317 | 0.035040167 | Cationic amino acid tra |
| AP53_1072 | -0.538663018 | 9.306028122 | 12.33479446 | 0.004608015 | 0.035213216 | putative esterase |
| AP53_490 | -0.884454121 | 8.144957934 | 12.29100507 | 0.004659892 | 0.035470001 | Additional lipoprotein |
| AP53_1508 | 0.38041251 | 8.636400623 | 12.21526177 | 0.004751264 | 0.036024234 | Hydrolase (HAD superfa |
| AP53_1127 | 0.479374343 | 9.163542651 | 12.19310534 | 0.004778392 | 0.036088943 | Peptidoglycan N-acetyl |
| AP53_1701 | 0.36333602 | 8.428105587 | 12.1587871 | 0.004820772 | 0.03610535 | ABC transporter perme |
| AP53_93 | 0.513990963 | 8.777350987 | 12.15492357 | 0.004825571 | 0.03610535 | Pyrroline-5-carboxylat |
| AP53_1047 | -0.906607293 | 6.075203301 | 12.14624743 | 0.004836368 | 0.03610535 | hypothetical phage pro |
| AP53_1432 | 0.549812884 | 10.71659941 | 12.08739248 | 0.004910371 | 0.036517359 | oxidoreductase of aldo, |
| AP53_1031 | -0.639455834 | 6.564176908 | 12.01746847 | 0.00500004 | 0.037042278 | FIG01114733: hypothe |
| AP53_1737 | 0.676071684 | 13.40341086 | 11.98132926 | 0.005047142 | 0.037207584 | Heat shock protein 60 f |
| AP53_852 | -0.574655867 | 6.142123985 | 11.9646116 | 0.005069108 | 0.037207584 | Voltage-gated chloride |
| AP53_456 | 0.57362083 | 10.88979319 | 11.95645943 | 0.005079861 | 0.037207584 | hypothetical protein |
| AP53_1213 | 0.666078013 | 7.298142231 | 11.91376587 | 0.005136616 | 0.037481846 | Phage CI-like repressor |
| AP53_703 | -0.792766496 | 6.57625008 | 11.88068188 | 0.005181112 | 0.03766494 | Thioredoxin reductase |
| AP53_977 | 0.695730835 | 13.251696 | 11.84752842 | 0.00522616 | 0.037774152 | General stress protein, |
| AP53_168 | -0.670138205 | 8.972750008 | 11.84102021 | 0.005235058 | 0.037774152 | tRNA-guanine transglyc |
| AP53_1765 | 0.550557971 | 10.01501353 | 11.81109237 | 0.005276204 | 0.037930043 | Ribonucleotide reducta |
| AP53_923 | 0.541457748 | 8.605489428 | 11.78539605 | 0.005311838 | 0.038045302 | Signal recognition parti |
| AP53_1534 | 0.611724671 | 8.554004643 | 11.69847665 | 0.005434499 | 0.038650205 | Sucrose operon repress |
| AP53_1301 | 0.634064336 | 8.23204417 | 11.69734484 | 0.005436119 | 0.038650205 | Hyaluronoglucosamini |
| AP53_1316 | -0.940009874 | 3.674597187 | 11.65705884 | 0.005494126 | 0.038843564 | Phage hyaluronidase |
| AP53_1235 | 0.579114846 | 10.02292846 | 11.63770105 | 0.005522259 | 0.038843564 | FIG01114027: hypothe |
| AP53_1590 | 0.483049708 | 9.872711188 | 11.63695219 | 0.005523351 | 0.038843564 | Putative stomatin/proh |
| AP53_1233 | 0.744832855 | 12.28677147 | 11.54808657 | 0.00565474 | 0.039583511 | ATP-dependent Clp pro |
| AP53_1733 | 0.523394715 | 7.51982531 | 11.52391691 | 0.005691112 | 0.039583511 | transcriptional regulat |
| AP53_123 | -0.491239551 | 5.959595513 | 11.50239189 | 0.005723738 | 0.039583511 | V-type ATP synthase sub |

| | | | | | | |
|---|---|---|---|---|---|---|
| AP53_562 | -1.074970399 | 9.093359734 | 11.50070577 | 0.005726303 | 0.039583511 | ATP synthase delta chai |
| AP53_1313 | -1.422276467 | 3.823947758 | 11.49632124 | 0.005732979 | 0.039583511 | Phage holin |
| AP53_1075 | -0.650675972 | 9.355194238 | 11.48456394 | 0.005750927 | 0.039583511 | D-alanyl transfer protei |
| AP53_986 | 0.517409315 | 7.078026064 | 11.45934662 | 0.005789648 | 0.039707882 | CAMP factor |
| AP53_946 | -0.558744158 | 8.691793309 | 11.44622915 | 0.005809912 | 0.039707882 | Phosphoglucomutase ( |
| AP53_94 | 0.396820752 | 8.902392534 | 11.42045917 | 0.005849965 | 0.039821049 | Glutamyl aminopeptid |
| AP53_233 | -0.438448889 | 8.635923669 | 11.38536108 | 0.005905044 | 0.039821049 | Cysteine desulfurase (E |
| AP53_1663 | -0.723168116 | 6.881238382 | 11.38451724 | 0.005906376 | 0.039821049 | Multiple sugar ABC trar |
| AP53_654 | 0.47584519 | 8.648977839 | 11.38315109 | 0.005908533 | 0.039821049 | Pathogenicity island Sa |
| AP53_1424 | 0.469164938 | 7.751467306 | 11.36453515 | 0.005938016 | 0.039881277 | Alpha-glycerophosphat |
| AP53_1006 | -1.437590813 | 6.091945859 | 11.31553225 | 0.006016461 | 0.040268795 | Phage lysin |
| AP53_1667 | 0.465341837 | 10.53558133 | 11.27264955 | 0.006086116 | 0.040595023 | GTP pyrophosphokinas |
| AP53_100 | -0.46383754 | 9.169923012 | 11.20397385 | 0.006199666 | 0.040839988 | tRNA dihydrouridine sy |
| AP53_rna63 | -1.488531179 | 2.769722909 | 11.18611571 | 0.006229603 | 0.040839988 | tRNA-Ile-GAT |
| AP53_1318 | -1.046440652 | 4.204982717 | 11.18289651 | 0.006235017 | 0.040839988 | Phage endopeptidase |
| AP53_821 | -1.242044354 | 9.889874738 | 11.17057245 | 0.006255798 | 0.040839988 | PTS system, mannose-s |
| AP53_258 | 0.681950798 | 7.671083701 | 11.16755855 | 0.006260892 | 0.040839988 | Methionine ABC transp |
| AP53_1620 | -0.584475682 | 6.896683767 | 11.16147994 | 0.006271182 | 0.040839988 | Tagatose 1,6-diphosph |
| AP53_372 | -0.451335417 | 8.032045335 | 11.15564889 | 0.006281072 | 0.040839988 | Diacylglycerol kinase (E |
| AP53_1548 | 0.351514221 | 7.216243169 | 11.13807994 | 0.00631098 | 0.040839988 | Transcriptional regulat |
| AP53_423 | 0.487416613 | 10.05280575 | 11.13736067 | 0.006312208 | 0.040839988 | Glycosyltransferase Laf |
| AP53_1050 | -0.565907765 | 5.944521376 | 11.10379651 | 0.00636983 | 0.041075881 | hypothetical protein |
| AP53_1040 | -0.832873072 | 5.15098903 | 11.09140435 | 0.006391262 | 0.041077612 | Phage protein |
| AP53_610 | -0.959338907 | 4.371625699 | 10.97795767 | 0.006591481 | 0.042093499 | Phage tail length tape- |
| AP53_1493 | 0.684862348 | 13.43323349 | 10.97133122 | 0.006603404 | 0.042093499 | Heat-inducible transcri |
| AP53_845 | 0.565774909 | 5.452419827 | 10.96524156 | 0.006614383 | 0.042093499 | Two-component respo |
| AP53_413 | 0.432173595 | 9.859277469 | 10.94608492 | 0.006649062 | 0.042175916 | Glutaminyl-peptide cyc |
| AP53_987 | 0.618983704 | 9.004300497 | 10.92280192 | 0.006691502 | 0.042306859 | Amino acid ABC transp |
| AP53_1177 | 0.633766543 | 10.37121398 | 10.87778252 | 0.006774475 | 0.042390858 | Lead, cadmium, zinc an |
| AP53_998 | 0.386697948 | 8.269286534 | 10.87451297 | 0.006780549 | 0.042390858 | ABC transporter, ATP-bi |
| AP53_1532 | 0.878821222 | 11.03730632 | 10.85818524 | 0.006810974 | 0.042390858 | PTS system, sucrose-spe |
| AP53_1840 | -0.678878968 | 8.984295955 | 10.85720255 | 0.006812811 | 0.042390858 | Immunodominant anti |
| AP53_1361 | 0.910367381 | 9.39071655 | 10.85357067 | 0.006819603 | 0.042390858 | putative cI-like repress |
| AP53_1176 | 0.875135414 | 10.08705089 | 10.8449126 | 0.006835826 | 0.042390858 | FIG01114217: hypothe |
| AP53_1326 | -1.51113602 | 4.142163145 | 10.82309582 | 0.00687691 | 0.042509816 | Phage minor capsid pr |

| | | | | | | |
|---|---|---|---|---|---|---|
| AP53_316 | 0.787689472 | 9.497923771 | 10.79390872 | 0.006932331 | 0.042674594 | hypothetical |
| AP53_1595 | -0.381662812 | 9.256043471 | 10.78594817 | 0.006947538 | 0.042674594 | Hydrolases of the alpha |
| AP53_856 | 0.551239316 | 6.912105337 | 10.67041623 | 0.007172752 | 0.043865831 | DNA repair protein Rad |
| AP53_1639 | -0.517372748 | 7.323090863 | 10.66341567 | 0.007186674 | 0.043865831 | Cysteinyl-tRNA syntheta |
| AP53_294 | 0.415277751 | 7.702865212 | 10.60749792 | 0.007299035 | 0.044369101 | tRNA (cytidine(34)-2'-O |
| AP53_960 | -0.453162225 | 7.539641314 | 10.59971242 | 0.007314844 | 0.044369101 | Phosphate transport AT |
| AP53_494 | -0.438978225 | 7.071482449 | 10.51545125 | 0.007488559 | 0.045162037 | Cell division protein Fts |
| AP53_1495 | 0.558020574 | 9.023869357 | 10.51162748 | 0.007496557 | 0.045162037 | D-alanyl-D-alanine carb |
| AP53_1248 | -0.748474566 | 9.88017853 | 10.50265159 | 0.007515373 | 0.045162037 | GTP-binding protein Ty |
| AP53_1429 | -0.568831425 | 7.988638528 | 10.43341659 | 0.007662397 | 0.045903433 | Glycyl-tRNA synthetase |
| AP53_567 | -0.501179974 | 7.522620469 | 10.41805463 | 0.007695479 | 0.045959769 | FIG01114865: hypothe |
| AP53_1827 | 0.792351727 | 7.738498587 | 10.30634121 | 0.007941212 | 0.047281879 | Phage infection protein |
| AP53_874 | 0.504462225 | 7.794643657 | 10.21588811 | 0.008147011 | 0.048323896 | Hypothetical similar to |
| AP53_1044 | -0.627245778 | 6.201115305 | 10.2076716 | 0.008166016 | 0.048323896 | hypothetical phage pro |
| AP53_rna84 | -1.79528695 | 1.78152497 | 10.16789401 | 0.008258771 | 0.048724239 | tRNA-Glu-TTC |
| AP53_1243 | -0.75784781 | 9.14538567 | 10.14818954 | 0.008305181 | 0.048846982 | Cell division protein Fts |
| AP53_1128 | -1.020549069 | 10.47072346 | 10.12782449 | 0.008353471 | 0.048846982 | Non-phosphorylating g |
| AP53_1117 | -0.553282853 | 8.568234233 | 10.12676183 | 0.008356 | 0.048846982 | S-adenosylmethionine s |
| AP53_600 | 0.841998925 | 10.95078582 | 10.10975308 | 0.008396602 | 0.048846982 | Phage protein |
| AP53_1487 | -0.762397751 | 10.25795846 | 10.10607793 | 0.008405405 | 0.048846982 | 3-oxoacyl-[acyl-carrier- |
| AP53_1364 | 0.606466656 | 8.777863786 | 10.08780747 | 0.008449334 | 0.048925356 | Phage integrase: site-sp |
| AP53_816 | -0.610159431 | 13.52738783 | 10.07953884 | 0.008469304 | 0.048925356 | Collagen-like surface pr |
| AP53_167 | -0.67356139 | 6.615908403 | 10.05458628 | 0.008529909 | 0.049039555 | FIG01116967: hypothe |
| AP53_437 | -0.631853945 | 6.32848962 | 10.05061436 | 0.008539603 | 0.049039555 | Chromosome partition |
| AP53_755 | 0.385559256 | 8.12430739 | 10.01052097 | 0.008638194 | 0.049459396 | Acyl-phosphate:glycero |
| AP53_349 | -0.35070337 | 8.400918934 | 9.978507892 | 0.008717885 | 0.049768869 | ADP-ribose pyrophosph |
| AP53_1266 | -0.938120955 | 10.8204009 | 9.848378495 | 0.009050922 | 0.051518592 | Acetyltransferase (EC 2. |
| AP53_546 | 0.745558071 | 7.330619277 | 9.803123265 | 0.009170262 | 0.052045259 | Streptolysin S biosynth |
| AP53_1477 | -0.816011983 | 10.2050706 | 9.764025 | 0.00927487 | 0.052397654 | Acetyl-coenzyme A carb |
| AP53_1720 | 0.525901106 | 6.068932166 | 9.75145172 | 0.00930881 | 0.052397654 | PTS system, cellobiose-s |
| AP53_489 | -1.021403486 | 7.91146 | 9.749778318 | 0.009313339 | 0.052397654 | Substrate-specific comp |
| AP53_1319 | -0.882342267 | 3.772270914 | 9.719105042 | 0.009396806 | 0.052615241 | Phage-related protein |
| AP53_804 | -0.657252166 | 8.79923924 | 9.710724419 | 0.009419765 | 0.052615241 | Phosphoglucosamine m |
| AP53_1682 | -0.46899823 | 5.782192653 | 9.705782319 | 0.009433335 | 0.052615241 | Dipeptide transport sys |
| AP53_501 | -0.398072833 | 8.033709296 | 9.693509076 | 0.009467136 | 0.052652469 | Promiscuous sugar pho |

| | | | | | | |
|---|---|---|---|---|---|---|
| AP53_1685 | -0.497988978 | 6.368726164 | 9.683100064 | 0.009495916 | 0.052661635 | Dipeptide transport ATI |
| AP53_235 | -0.421263411 | 8.612244144 | 9.66300724 | 0.009551763 | 0.052688575 | Iron-sulfur cluster asser |
| AP53_547 | 0.655449896 | 7.169960578 | 9.661824216 | 0.009555064 | 0.052688575 | Streptolysin S biosynth |
| AP53_424 | -0.675626586 | 6.560053996 | 9.606606941 | 0.00971062 | 0.053394658 | hypothetical protein |
| AP53_121 | -0.424383457 | 6.021714868 | 9.57932624 | 0.009788582 | 0.053671295 | V-type ATP synthase sub |
| AP53_438 | 0.615236 | 5.192022396 | 9.550211265 | 0.009872607 | 0.05397952 | Positive transcriptional |
| AP53_1644 | -0.940151743 | 6.060227028 | 9.52029604 | 0.009959833 | 0.054271826 | Transaldolase (EC 2.2.1. |
| AP53_1849 | -0.405360898 | 8.259439319 | 9.512746607 | 0.00998199 | 0.054271826 | UDP-glucose dehydroge |
| AP53_783 | -0.46403484 | 7.122457783 | 9.455957784 | 0.010150546 | 0.054976019 | Hypothetical protein SI |
| AP53_705 | -0.49629733 | 8.668212891 | 9.450089169 | 0.010168156 | 0.054976019 | 2-dehydropantoate 2-r |
| AP53_212 | 0.312446081 | 9.368271815 | 9.39984144 | 0.010320432 | 0.055644328 | Ribosome small subuni |
| AP53_1293 | 0.448551188 | 5.977682409 | 9.370273703 | 0.010411301 | 0.055978769 | Two-component sensor |
| AP53_535 | -0.659173826 | 10.82592105 | 9.323382343 | 0.010557364 | 0.056472253 | LSU ribosomal protein I |
| AP53_65 | -0.940382659 | 8.071274492 | 9.302141431 | 0.010624328 | 0.056472253 | Translation initiation fa |
| AP53_rna34 | -1.675969156 | 3.37489484 | 9.296010466 | 0.01064375 | 0.056472253 | tRNA-Pseudo-GCT |
| AP53_484 | -0.389175713 | 6.791743853 | 9.293338916 | 0.010652226 | 0.056472253 | 2-haloalkanoic acid deh |
| AP53_1126 | 0.37978735 | 8.692747408 | 9.2885969 | 0.010667291 | 0.056472253 | ATP-dependent RNA hel |
| AP53_511 | 0.468044756 | 6.045520278 | 9.284683534 | 0.010679742 | 0.056472253 | Unsaturated glucurony |
| AP53_170 | 0.764035618 | 9.354608329 | 9.274180262 | 0.010713247 | 0.056472253 | Substrate-specific comp |
| AP53_1080 | -0.532123133 | 7.621737499 | 9.267120239 | 0.010735838 | 0.056472253 | Glutamate transport AT |
| AP53_819 | -1.311667629 | 8.051631786 | 9.234532511 | 0.010840849 | 0.056739892 | PTS system, IIA compor |
| AP53_38 | 0.421590445 | 8.772914564 | 9.233195728 | 0.010845183 | 0.056739892 | Acyltransferase family |
| AP53_1262 | -1.118435527 | 10.79217666 | 9.19573779 | 0.010967455 | 0.057225352 | Carbamate kinase (EC 2. |
| AP53_978 | 0.711963156 | 13.63851091 | 9.184216307 | 0.011005393 | 0.05726935 | FIG01114697: hypothe |
| AP53_1702 | 0.504304109 | 7.891787285 | 9.150305291 | 0.011117961 | 0.057700436 | ABC transporter ATP-bi |
| AP53_726 | -0.425570254 | 7.212955973 | 9.104070403 | 0.011273649 | 0.058352405 | Mevalonate kinase (EC 2 |
| AP53_1687 | -0.342033123 | 6.531062586 | 9.091145523 | 0.011317633 | 0.05836351 | Streptococcal histidine |
| AP53_1623 | -0.522326766 | 6.765951862 | 9.085785832 | 0.011335932 | 0.05836351 | Galactose-6-phosphate |
| AP53_371 | -0.448602389 | 8.04646574 | 9.033069472 | 0.011517799 | 0.059142986 | Metal-dependent hydro |
| AP53_rna68 | -1.010357923 | 4.055443708 | 9.011865258 | 0.011591928 | 0.059172684 | tRNA-Val-TAC |
| AP53_963 | -0.435685711 | 7.67750797 | 9.009426017 | 0.011600492 | 0.059172684 | Phosphate ABC transpo |
| AP53_1723 | 0.412007756 | 8.835087518 | 9.005286983 | 0.01161504 | 0.059172684 | Transcriptional regulat |
| AP53_738 | 0.83068672 | 13.6746115 | 8.984078332 | 0.011689928 | 0.059344013 | putative ATP-dependen |
| AP53_1298 | 0.549525427 | 7.18348429 | 8.978472499 | 0.011709818 | 0.059344013 | N-acetylmannosamine I |
| AP53_1046 | -0.877872029 | 5.864546982 | 8.969366818 | 0.011742212 | 0.05935321 | hypothetical phage pro |

| | | | | | | |
|---|---|---|---|---|---|---|
| AP53_1119 | -0.532434301 | 7.650473294 | 8.950172646 | 0.011810844 | 0.05954506 | Biotin operon repressor |
| AP53_1070 | 0.58008938 | 6.090033599 | 8.899240475 | 0.011995283 | 0.06031825 | Mobile element protein |
| AP53_357 | 0.360507995 | 9.9521086 | 8.88727414 | 0.012039111 | 0.060382209 | Peptidyl-prolyl cis-tran |
| AP53_77 | 0.621932174 | 6.050714383 | 8.858357173 | 0.01214581 | 0.060760356 | HIT family protein |
| AP53_429 | -0.991400685 | 2.085177102 | 8.816620469 | 0.012301799 | 0.061382501 | Hypothetical protein |
| AP53_1338 | -0.733246963 | 3.122952067 | 8.806310759 | 0.012340697 | 0.061418701 | FIG01114733: hypothe |
| AP53_924 | 0.455464749 | 8.901812604 | 8.791879013 | 0.012395393 | 0.061525096 | Transcriptional regulat |
| AP53_730 | -0.483256564 | 7.593321494 | 8.783975051 | 0.01242547 | 0.061525096 | Hydroxymethylglutaryl |
| AP53_1603 | -0.420341161 | 6.695972091 | 8.727842504 | 0.012641588 | 0.061989704 | Substrate-specific comp |
| AP53_935 | -0.625585373 | 9.235748517 | 8.717376565 | 0.012682375 | 0.061989704 | Formate--tetrahydrofol |
| AP53_1503 | 0.524878513 | 7.92742632 | 8.712544951 | 0.012701257 | 0.061989704 | Pyruvate,phosphate dil |
| AP53_1215 | 0.606346528 | 6.297480204 | 8.707919573 | 0.012719365 | 0.061989704 | Phage integrase |
| AP53_313 | 0.609723076 | 9.55902715 | 8.692848421 | 0.012778579 | 0.061989704 | Thymidylate kinase (EC |
| AP53_1600 | -0.414199987 | 6.597198535 | 8.690710749 | 0.012787004 | 0.061989704 | Potassium efflux system |
| AP53_1402 | -0.364123287 | 7.975794128 | 8.678969207 | 0.0128334 | 0.061989704 | Histidine transport pro |
| AP53_321 | -0.385188035 | 9.194140231 | 8.673461378 | 0.012855232 | 0.061989704 | Arsenate reductase (EC |
| AP53_92 | 0.481754491 | 6.356531205 | 8.668149753 | 0.012876329 | 0.061989704 | Hypothetical protein SI |
| AP53_1366 | 0.345519852 | 9.455194799 | 8.666478151 | 0.012882977 | 0.061989704 | Cysteinyl-tRNA syntheta |
| AP53_1173 | -0.392063809 | 9.926256259 | 8.662142611 | 0.012900238 | 0.061989704 | Phosphoglycerate muta |
| AP53_1074 | -0.617866694 | 9.483726773 | 8.661563511 | 0.012902545 | 0.061989704 | D-alanine--poly(phosph |
| AP53_1645 | -1.139319401 | 6.523461692 | 8.621647162 | 0.013062794 | 0.06260465 | Transport protein SgaT, |
| AP53_1453 | 0.563351861 | 10.39467819 | 8.609032873 | 0.013113926 | 0.062694905 | Ribosome-binding facto |
| AP53_1048 | -0.741263836 | 6.144025653 | 8.567891283 | 0.013282356 | 0.063250337 | hypothetical protein, p |
| AP53_1609 | 0.324938371 | 7.566110226 | 8.564760079 | 0.013295279 | 0.063250337 | Type I restriction-modi |
| AP53_434 | -0.307462029 | 8.537490947 | 8.523919286 | 0.013465222 | 0.06390219 | Two-component sensor |
| AP53_823 | -0.524563431 | 5.578408491 | 8.507081875 | 0.013536035 | 0.064081569 | Two-component sensor |
| AP53_1375 | 0.280674982 | 7.123257277 | 8.459327637 | 0.013739294 | 0.064885572 | Sensor histidine kinase |
| AP53_1812 | -0.469284541 | 8.060183411 | 8.412753595 | 0.013941034 | 0.06567851 | Aspartyl-tRNA syntheta |
| AP53_311 | 0.636515863 | 12.01622451 | 8.397139341 | 0.014009455 | 0.065833174 | ATP-dependent Clp pro |
| AP53_1854 | 0.301111869 | 9.006424022 | 8.389812751 | 0.014041697 | 0.065833174 | Putative sugar uptake p |
| AP53_1660 | -0.992706338 | 8.781260688 | 8.370140155 | 0.014128708 | 0.065842128 | putative transport acce |
| AP53_989 | 0.590513562 | 8.859544528 | 8.366431913 | 0.014145182 | 0.065842128 | ABC transporter membr |
| AP53_1264 | -0.963417645 | 10.19077399 | 8.366389028 | 0.014145372 | 0.065842128 | Arginine/ornithine anti |
| AP53_937 | -0.693852062 | 8.459560788 | 8.356064018 | 0.01419136 | 0.065898158 | NAD-dependent protein |
| AP53_33 | 0.414823692 | 9.672083697 | 8.34164751 | 0.01425587 | 0.066029646 | Adenylosuccinate lyase |

| | | | | | | |
|---|---|---|---|---|---|---|
| AP53_1339 | -1.11898239 | 4.132509886 | 8.329380979 | 0.014311033 | 0.066029646 | hypothetical protein |
| AP53_1692 | -0.451955953 | 11.16790356 | 8.327008527 | 0.014321732 | 0.066029646 | Antiphagocytic M prote |
| AP53_299 | 0.293237059 | 8.536673103 | 8.2906891 | 0.014486704 | 0.066631972 | Hemolysins and related |
| AP53_198 | 0.431305613 | 10.34227161 | 8.265288834 | 0.014603422 | 0.067010027 | RNA-binding protein Ja |
| AP53_1496 | 0.308225654 | 8.657112087 | 8.240792216 | 0.014717048 | 0.067372146 | Phosphoglycerate muta |
| AP53_817 | 0.313191596 | 9.179772291 | 8.217680089 | 0.014825217 | 0.067707638 | Peptide methionine sul |
| AP53_373 | -0.415787597 | 9.046666375 | 8.197650419 | 0.014919725 | 0.067979311 | GTP-binding protein Era |
| AP53_1597 | -0.313452219 | 8.736347993 | 8.136719929 | 0.015211656 | 0.069147131 | CTP synthase (EC 6.3.4.: |
| AP53_627 | -0.960775566 | 3.230233061 | 8.101973525 | 0.015381175 | 0.069754348 | hypothetical phage pro |
| AP53_991 | 0.528251219 | 11.37384296 | 8.049033515 | 0.015643796 | 0.070779974 | Glucosamine--fructose- |
| AP53_1184 | -0.818367683 | 3.96316391 | 8.03716958 | 0.01570338 | 0.070884326 | Phage protein |
| AP53_1835 | -0.320886342 | 7.517769788 | 8.020827667 | 0.015785895 | 0.071091468 | tRNA uridine 5-carboxy |
| AP53_840 | -0.566325828 | 8.092291269 | 7.987362641 | 0.015956486 | 0.071693379 | UDP-N-acetylenolpyruv |
| AP53_733 | -0.624818444 | 7.69791382 | 7.973060652 | 0.01603006 | 0.071857614 | Dihydrofolate reductas |
| AP53_128 | -0.444230919 | 6.379202322 | 7.963275408 | 0.016080631 | 0.071918215 | V-type ATP synthase sub |
| AP53_461 | -0.433423376 | 7.991869669 | 7.911897547 | 0.016349293 | 0.072951674 | Hydrolase (HAD superfa |
| AP53_959 | -0.390163434 | 7.296947342 | 7.886239618 | 0.016485461 | 0.073390548 | Phosphate transport AT |
| AP53_1808 | 0.562116591 | 7.402905307 | 7.844677162 | 0.016708909 | 0.074215086 | bacteriocin immunity p |
| AP53_1738 | 0.558331382 | 10.31251749 | 7.793465101 | 0.016989213 | 0.075287814 | ATP-dependent Clp pro |
| AP53_rna66 | -1.240359779 | 4.587209041 | 7.748891469 | 0.017237749 | 0.076215193 | tRNA-Lys-TTT |
| AP53_585 | 0.313124174 | 9.096517146 | 7.725524501 | 0.017369768 | 0.076624361 | DNA primase (EC 2.7.7.- |
| AP53_1052 | -0.486520094 | 7.631196824 | 7.707875006 | 0.017470283 | 0.076785241 | hypothetical protein |
| AP53_1324 | -0.885431359 | 3.836125107 | 7.705238616 | 0.017485356 | 0.076785241 | Phage minor capsid pro |
| AP53_286 | -0.358179822 | 8.349778934 | 7.661754289 | 0.01773623 | 0.077711112 | unknown domain / Nuc |
| AP53_841 | -0.44180889 | 8.131143388 | 7.6406686 | 0.017859422 | 0.077782807 | Putrescine transport AT |
| AP53_1334 | -0.834295549 | 3.009464722 | 7.635304264 | 0.017890925 | 0.077782807 | Phage terminase large s |
| AP53_1423 | 0.60127878 | 8.381643768 | 7.634100885 | 0.017898001 | 0.077782807 | Glycerol uptake facilita |
| AP53_272 | 0.266428905 | 9.628509917 | 7.631571138 | 0.017912888 | 0.077782807 | GTP-binding protein En |
| AP53_1855 | 0.508631341 | 11.18573207 | 7.593523253 | 0.018138571 | 0.078586978 | Inosine-5'-monophosph |
| AP53_184 | 0.340003309 | 8.108886386 | 7.531584267 | 0.018513254 | 0.080031684 | Transcriptional regulat |
| AP53_1616 | -0.990982535 | 4.775695083 | 7.504275846 | 0.018681374 | 0.080578992 | Lantibiotic salivaricin A |
| AP53_1393 | 0.294895434 | 10.5883986 | 7.482222795 | 0.018818468 | 0.080990348 | Hypothetical protein SP |
| AP53_rna52 | -1.675747713 | 3.930072791 | 7.439493223 | 0.019087527 | 0.081966571 | tRNA-Ala-TGC |
| AP53_422 | 0.374190279 | 10.12176022 | 7.428402174 | 0.019158113 | 0.082088073 | Glycosyltransferase Laf |
| AP53_1659 | -0.824844446 | 9.560678367 | 7.364681623 | 0.019569724 | 0.083456495 | Preprotein translocase |

| | | | | | | |
|---|---|---|---|---|---|---|
| AP53_1226 | 0.245652722 | 8.30245655 | 7.360251355 | 0.019598732 | 0.083456495 | Exodeoxyribonuclease V |
| AP53_843 | -0.355628749 | 7.471071188 | 7.355330291 | 0.019631013 | 0.083456495 | Spermidine Putrescine |
| AP53_1003 | 0.605468111 | 6.570567463 | 7.352521254 | 0.019649468 | 0.083456495 | Streptococcal pyrogeni |
| AP53_277 | 0.252200582 | 9.564369913 | 7.328986335 | 0.019804907 | 0.083667863 | FIG004453: protein Yce |
| AP53_370 | -0.478058987 | 7.3926236 | 7.326896894 | 0.019818778 | 0.083667863 | Hypothetical protein V( |
| AP53_43 | -0.832207315 | 10.88048974 | 7.325425793 | 0.019828551 | 0.083667863 | LSU ribosomal protein I |
| AP53_735 | -0.641864532 | 8.638929863 | 7.277854431 | 0.020147694 | 0.084830097 | ATP-dependent Clp pro |
| AP53_842 | -0.372198981 | 7.789003183 | 7.263911444 | 0.020242392 | 0.085044336 | Spermidine Putrescine |
| AP53_309 | -0.490711015 | 8.569034772 | 7.253487405 | 0.020313537 | 0.085158909 | Uracil phosphoribosylt |
| AP53_197 | 0.356242635 | 9.919655201 | 7.242344446 | 0.020389918 | 0.085294894 | Inner membrane protei |
| AP53_806 | -0.516438009 | 6.347657363 | 7.218023301 | 0.020557824 | 0.08564448 | Hypothetical radical SA |
| AP53_240 | 0.311620803 | 8.982834402 | 7.211908828 | 0.020600296 | 0.08564448 | Oligopeptide transport |
| AP53_1325 | -0.974613074 | 2.984739862 | 7.204865694 | 0.020649347 | 0.08564448 | Phage minor capsid pro |
| AP53_1305 | 0.351161087 | 7.406840544 | 7.204774609 | 0.020649983 | 0.08564448 | Putative two-compone |
| AP53_586 | 0.33350278 | 10.17786669 | 7.177593027 | 0.020840606 | 0.086250783 | RNA polymerase sigma |
| AP53_668 | -0.46930864 | 10.80527704 | 7.165765299 | 0.020924209 | 0.086412531 | LSU ribosomal protein I |
| AP53_1773 | 0.475036449 | 11.49108682 | 7.139249686 | 0.021113088 | 0.08679818 | Putative Holliday junct |
| AP53_961 | -0.335448649 | 7.148994965 | 7.138666733 | 0.021117264 | 0.08679818 | Phosphate transport sy |
| AP53_369 | -0.65277273 | 6.804805955 | 7.13385763 | 0.021151746 | 0.08679818 | Phosphate starvation-ir |
| AP53_1704 | 0.527575052 | 8.338143977 | 7.116025645 | 0.021280193 | 0.087141045 | FIG01115531: hypothe |
| AP53_1684 | -0.416705224 | 6.496705515 | 7.10255996 | 0.021377806 | 0.087356467 | Dipeptide transport ATI |
| AP53_938 | -0.651932648 | 8.540954766 | 7.094910642 | 0.021433494 | 0.087400025 | COG2110, Macro doma |
| AP53_119 | -0.395960771 | 5.4753407 | 7.077440033 | 0.021561332 | 0.087736994 | V-type ATP synthase sub |
| AP53_42 | -0.642463421 | 10.79703521 | 7.063551567 | 0.021663608 | 0.087816971 | SSU ribosomal protein S |
| AP53_1828 | 0.653701396 | 5.847888352 | 7.059261676 | 0.021695316 | 0.087816971 | Transcriptional regulat |
| AP53_680 | -0.527763782 | 10.64839791 | 7.056370626 | 0.021716716 | 0.087816971 | LSU ribosomal protein I |
| AP53_773 | -0.389624884 | 9.386672227 | 7.047616438 | 0.02178167 | 0.08789651 | Adenine phosphoribosy |
| AP53_1297 | 0.431785701 | 6.253431008 | 7.02007907 | 0.021987506 | 0.088520582 | Multiple sugar ABC trar |
| AP53_1864 | 0.668961706 | 9.373845942 | 7.014759912 | 0.022027533 | 0.088520582 | Chromosome (plasmid) |
| AP53_1769 | 0.385406289 | 9.866497896 | 6.994789878 | 0.022178585 | 0.088702884 | Ribonucleotide reducta |
| AP53_pseudo73 | -0.428531759 | 6.753476038 | 6.990893142 | 0.022208204 | 0.088702884 | 4-diphosphocytidyl-2-C |
| AP53_1539 | 0.287371484 | 8.072907953 | 6.990657615 | 0.022209996 | 0.088702884 | Aminopeptidase YpdF ( |
| AP53_148 | -0.751685428 | 6.799100627 | 6.976738099 | 0.022316193 | 0.088944006 | Probable L-ascorbate-6 |
| AP53_1240 | -0.531718334 | 10.00904625 | 6.952373101 | 0.022503546 | 0.08950693 | FIG001960: FtsZ-intera |
| AP53_1129 | -0.671958092 | 10.5572049 | 6.942640015 | 0.022578912 | 0.089623042 | Phosphoenolpyruvate-j |

| | | | | | | |
|---|---|---|---|---|---|---|
| AP53_1669 | -0.287181083 | 7.375156326 | 6.923052299 | 0.0227315 | 0.089655407 | Ribonucleotide reducti |
| AP53_340 | -0.333298126 | 6.586406523 | 6.922672686 | 0.02273447 | 0.089655407 | hypothetical protein |
| AP53_1303 | 0.573070321 | 8.524657579 | 6.921625337 | 0.022742664 | 0.089655407 | hypothetical protein |
| AP53_114 | 0.551384533 | 5.371535414 | 6.91239597 | 0.02281503 | 0.089655407 | Acetyl-CoA:acetoacetyl |
| AP53_662 | 0.417520862 | 8.236886967 | 6.909408897 | 0.02283851 | 0.089655407 | Pore forming protein el |
| AP53_rna8 | -1.136965208 | 4.747622072 | 6.901750857 | 0.022898838 | 0.089655407 | tRNA-Leu-TAG |
| AP53_892 | 0.48498334 | 12.79974523 | 6.900286253 | 0.022910398 | 0.089655407 | FIG01115412: hypothe |
| AP53_1536 | 0.363228242 | 11.00225356 | 6.866398481 | 0.023179814 | 0.090527201 | Alkaline shock protein |
| AP53_728 | -0.379638482 | 7.770431655 | 6.850126322 | 0.023310525 | 0.090854878 | Phosphomevalonate ki |
| AP53_298 | 0.310513659 | 7.251107526 | 6.819929953 | 0.023555427 | 0.091625417 | SAM-dependent methyl |
| AP53_752 | 0.292970315 | 9.431559898 | 6.806185752 | 0.023667913 | 0.091846441 | Cystine transport syster |
| AP53_1333 | -0.717600649 | 3.062741465 | 6.795722713 | 0.023753976 | 0.091846441 | Phage minor capsid prc |
| AP53_1679 | 0.347421857 | 7.966748178 | 6.79569482 | 0.023754206 | 0.091846441 | Chromosome segregati |
| AP53_112 | 0.535525488 | 5.848035247 | 6.789867623 | 0.023802299 | 0.091849429 | Transcriptional regulat |
| AP53_rna33 | -1.49522724 | 4.405899646 | 6.783042476 | 0.023858777 | 0.091884696 | tRNA-Ile-GAT |
| AP53_1064 | 0.528757111 | 6.36318039 | 6.772830975 | 0.023943576 | 0.092005941 | 4-alpha-glucanotransfe |
| AP53_1817 | 0.429934423 | 5.415938694 | 6.766048444 | 0.024000099 | 0.092005941 | Cadmium efflux system |
| AP53_1049 | -0.800095645 | 4.90069802 | 6.762173448 | 0.024032464 | 0.092005941 | hypothetical phage pro |
| AP53_1713 | 1.062184058 | 10.57102058 | 6.745795523 | 0.02416983 | 0.09231048 | Streptodornase B; Mito |
| AP53_918 | 0.633163379 | 9.936688656 | 6.741367887 | 0.024207127 | 0.09231048 | [Citrate [pro-3S]-lyase] l |
| AP53_1321 | -0.967562804 | 2.997535762 | 6.69124324 | 0.024634192 | 0.093754837 | Phage protein |
| AP53_242 | 0.365309996 | 9.151222099 | 6.672841005 | 0.024793235 | 0.094175479 | Oligopeptide transport |
| AP53_1632 | 0.335484103 | 7.214749221 | 6.662265517 | 0.02488519 | 0.094340144 | Transcriptional regulat |
| AP53_805 | -0.485450256 | 4.94977484 | 6.65484835 | 0.024949926 | 0.094401183 | FIG01115418: hypothe |
| AP53_1153 | 0.312919703 | 7.176369711 | 6.639273233 | 0.02508652 | 0.094733337 | FIG01114764: hypothe |
| AP53_957 | -0.57642639 | 8.083285336 | 6.630202682 | 0.025166481 | 0.094850755 | Lysyl aminopeptidase (l |
| AP53_691 | 0.441491121 | 8.599600017 | 6.620124661 | 0.02525568 | 0.09500247 | periplasmic componen |
| AP53_326 | 0.364194923 | 9.854876452 | 6.609032799 | 0.02535429 | 0.095188929 | Serine endopeptidase S |
| AP53_1666 | 0.562465122 | 9.740172386 | 6.576522167 | 0.025645981 | 0.096098164 | D-tyrosyl-tRNA(Tyr) dea |
| AP53_rna67 | -1.028840881 | 4.349876004 | 6.511874096 | 0.026238024 | 0.098127178 | tRNA-Asp-GTC |
| AP53_350 | -0.294474918 | 8.629141679 | 6.482539556 | 0.026512051 | 0.098771815 | FIG01115635: hypothe |
| AP53_1770 | 0.335407854 | 7.59429478 | 6.482527243 | 0.026512167 | 0.098771815 | Competence-induced p |
| AP53_1238 | -0.43882754 | 9.054908391 | 6.467984428 | 0.026649287 | 0.099092462 | FIG001583: hypothetic |
| AP53_319 | 0.563781795 | 9.304296752 | 6.433677775 | 0.026976128 | 0.099837797 | FIG01114628: hypothe |
| AP53_rna19 | -1.466019389 | 4.531004466 | 6.430811537 | 0.027003651 | 0.099837797 | tRNA-Asp-GTC |

| | | | | | | |
|---|---|---|---|---|---|---|
| AP53_846 | 0.353029471 | 5.626682958 | 6.427761539 | 0.027032976 | 0.099837797 | Two-component sensor |
| AP53_1501 | -0.315684034 | 8.011588379 | 6.425423934 | 0.027055477 | 0.099837797 | Aspartyl-tRNA(Asn) ami |
| AP53_186 | 0.507184115 | 8.362564137 | 6.365113825 | 0.027643825 | 0.101695878 | Lipid A export ATP-binc |
| AP53_171 | -0.513070174 | 7.61453829 | 6.363093895 | 0.027663794 | 0.101695878 | Metal-dependent hydro |
| AP53_632 | -0.531714186 | 4.285425534 | 6.335556612 | 0.027937751 | 0.102508837 | hypothetical protein |
| AP53_936 | -0.647744297 | 8.458939196 | 6.296182067 | 0.028335135 | 0.103770748 | Lipoate-protein ligase A |
| AP53_182 | -0.279763343 | 8.834217995 | 6.282849563 | 0.028471222 | 0.104072772 | UTP--glucose-1-phosph |
| AP53_497 | -0.416315548 | 10.59102922 | 6.27541582 | 0.02854744 | 0.104155226 | Triosephosphate isome |
| AP53_1455 | 0.382649166 | 9.896842533 | 6.264189897 | 0.028663001 | 0.104380647 | ribosomal protein L7Ae |
| AP53_729 | -0.345613396 | 7.986503413 | 6.248013024 | 0.028830515 | 0.104794063 | Isopentenyl-diphospha |
| AP53_594 | -0.352773608 | 6.746826299 | 6.223388728 | 0.029087759 | 0.105531477 | Alpha-L-Rha alpha-1,2-l |
| AP53_221 | -0.437620566 | 11.87240178 | 6.212195537 | 0.029205599 | 0.105761322 | Translation elongation |
| AP53_1673 | -0.522220526 | 7.054821001 | 6.193348638 | 0.029405311 | 0.106170051 | Ribosomal RNA small su |
| AP53_436 | -0.741415684 | 5.781994632 | 6.191229791 | 0.029427866 | 0.106170051 | Ribonuclease III (EC 3.1 |
| AP53_1589 | 0.382026319 | 9.549365128 | 6.17797536 | 0.029569427 | 0.106482852 | Acid phosphatase (EC 3. |
| AP53_462 | -0.391445814 | 8.914955832 | 6.149746292 | 0.029873648 | 0.10713527 | Signal recognition parti |
| AP53_1292 | 0.432107656 | 5.910146344 | 6.147239706 | 0.029900842 | 0.10713527 | DNA-binding response r |
| AP53_1654 | -0.349463505 | 7.992457834 | 6.143116138 | 0.029945643 | 0.10713527 | DNA polymerase III alph |
| AP53_1139 | 0.293669072 | 6.556102348 | 6.140750439 | 0.029971382 | 0.10713527 | FIG01114175: hypothe |
| AP53_818 | 0.248340575 | 9.40215327 | 6.108689113 | 0.030322835 | 0.108192322 | Putative membrane pro |
| AP53_353 | 0.352082154 | 9.770017578 | 6.102216072 | 0.03039439 | 0.108248644 | Mn-dependent transcri |
| AP53_1601 | -0.257677181 | 7.763040086 | 6.08557935 | 0.030579227 | 0.10826161 | Hypothetical protein yv |
| AP53_274 | 0.316985825 | 10.45166169 | 6.083893737 | 0.030598029 | 0.10826161 | FIG01114242: hypothe |
| AP53_558 | -0.419467205 | 7.727189147 | 6.083387748 | 0.030603676 | 0.10826161 | Transcription regulator |
| AP53_rna47 | -1.139663454 | 4.404252428 | 6.081824006 | 0.030621135 | 0.10826161 | tRNA-Trp-CCA |
| AP53_794 | -0.669357745 | 9.90566459 | 6.063702777 | 0.030824333 | 0.108781873 | Acetoin dehydrogenase |
| AP53_702 | -0.511138888 | 6.892190454 | 6.007865936 | 0.031460684 | 0.110674515 | tRNA (Guanine37-N1) -r |
| AP53_1448 | 0.286674355 | 8.01963308 | 6.004512061 | 0.031499405 | 0.110674515 | Lactose phosphotransfe |
| AP53_1705 | 0.327993772 | 7.659559924 | 6.00171964 | 0.031531688 | 0.110674515 | FIG01116435: hypothe |
| AP53_1057 | -0.91377995 | 4.900415044 | 5.983950094 | 0.031738052 | 0.111197761 | FIG01115240: hypothe |
| AP53_75 | -0.344994225 | 7.726886869 | 5.96722986 | 0.03193371 | 0.11157419 | Zinc ABC transporter, A |
| AP53_213 | 0.287891511 | 9.447110586 | 5.963610922 | 0.031976249 | 0.11157419 | Ribulose-phosphate 3-e |
| AP53_727 | -0.368590179 | 7.583116513 | 5.953458759 | 0.032095945 | 0.11157419 | Diphosphomevalonate |
| AP53_1783 | 0.502046105 | 8.552077433 | 5.952944541 | 0.032102022 | 0.11157419 | hypothetical protein |
| AP53_1409 | 0.254585898 | 9.200307258 | 5.948469492 | 0.032154966 | 0.11157419 | Phospho-N-acetylmura |

| | | | | | | |
|---|---|---|---|---|---|---|
| AP53_1302 | 0.450406554 | 7.945785161 | 5.945480268 | 0.03219039 | 0.11157419 | Transcriptional represso |
| AP53_1509 | 0.351613831 | 7.486170921 | 5.901930242 | 0.032711824 | 0.113179412 | L-asparaginase (EC 3.5.1 |
| AP53_732 | -0.442047868 | 6.437125728 | 5.894689549 | 0.032799497 | 0.113280826 | Thymidylate synthase (E |
| AP53_rna23 | -1.346584344 | 2.598266444 | 5.874985875 | 0.033039509 | 0.11390708 | tRNA-Gly-GCC |
| AP53_498 | 5.250123094 | -1.006521509 | 6.216011819 | 0.033167929 | 0.114147076 | hypothetical protein |
| AP53_1805 | 0.36611166 | 7.418596014 | 5.834507689 | 0.033539217 | 0.115220568 | biofilm formation prote |
| AP53_rna72 | 3.212133731 | 5.80827402 | 5.938042382 | 0.033830227 | 0.116014965 | Small Subunit Ribosom |
| AP53_1638 | -0.344474782 | 7.493116441 | 5.756025533 | 0.034534175 | 0.118220165 | COG1939: Ribonucleas |
| AP53_161 | 0.339774012 | 6.18135813 | 5.745147354 | 0.034674862 | 0.118492793 | Malonate permease |
| AP53_701 | -0.516066266 | 7.052130267 | 5.738653577 | 0.034759173 | 0.118572152 | 16S rRNA processing pr |
| AP53_487 | 0.455948135 | 9.432426044 | 5.716064291 | 0.035054382 | 0.119369396 | Substrate-specific comp |
| AP53_153 | 0.318989693 | 10.55554949 | 5.658043711 | 0.035826532 | 0.121770836 | Peroxide stress regulato |
| AP53_1239 | -0.486952433 | 8.396129883 | 5.650594024 | 0.035927146 | 0.121770836 | Cell division protein Ylm |
| AP53_1299 | 0.456218988 | 6.93179985 | 5.649067538 | 0.035947805 | 0.121770836 | FIG01114249: hypothe |
| AP53_822 | -0.890157621 | 9.76644687 | 5.601902109 | 0.036593199 | 0.123741113 | PTS system, mannose-sp |
| AP53_pseudo656 | 0.429180574 | 8.518667932 | 5.584683246 | 0.036832277 | 0.124244437 | Lipoteichoic acid synth |
| AP53_pseudo1695 | 0.608323211 | 11.9771486 | 5.575777515 | 0.036956667 | 0.124244437 | Antiphagocytic M prote |
| AP53_799 | -0.404901467 | 7.171421152 | 5.573657423 | 0.036986353 | 0.124244437 | Lipoate-protein ligase A |
| AP53_888 | 0.225394146 | 9.182311722 | 5.572820101 | 0.036998086 | 0.124244437 | DNA gyrase subunit A (E |
| AP53_736 | -0.528659437 | 8.860886175 | 5.562210611 | 0.037147132 | 0.124301675 | GTP-binding protein En |
| AP53_1734 | 0.513854848 | 7.484039976 | 5.560178631 | 0.037175761 | 0.124301675 | Transcriptional regulato |
| AP53_1116 | -0.311757524 | 7.551347705 | 5.556612131 | 0.037226073 | 0.124301675 | UDP-N-acetylglucosami |
| AP53_825 | -0.428460153 | 5.835413574 | 5.553411728 | 0.03727129 | 0.124301675 | Ferric iron ABC transpo |
| AP53_1005 | 0.347391655 | 7.959082217 | 5.5405289 | 0.037453976 | 0.124520093 | Phage protein |
| AP53_1518 | 0.287692004 | 6.978776583 | 5.538202564 | 0.03748708 | 0.124520093 | Heme ABC transporter ( |
| AP53_129 | 0.318003719 | 8.445112756 | 5.533514246 | 0.037553901 | 0.124520093 | putative toxic anion res |
| AP53_1172 | 0.380858079 | 7.299821759 | 5.530748129 | 0.037593392 | 0.124520093 | Transcriptional represso |
| AP53_rna30 | -0.814226535 | 3.832576393 | 5.515408049 | 0.03781331 | 0.125035152 | tRNA-Met-CAT |
| AP53_1406 | 0.308686373 | 9.002072289 | 5.495163568 | 0.038105909 | 0.125636093 | Cysteine ABC transporte |
| AP53_1504 | 0.35907281 | 6.423323182 | 5.493883447 | 0.038124502 | 0.125636093 | Nicotinamidase (EC 3.5 |
| AP53_253 | 0.267840053 | 7.304285585 | 5.450225193 | 0.038765201 | 0.127530942 | FIG007079: UPF0348 p |
| AP53_1108 | 0.269560683 | 8.456743553 | 5.440096332 | 0.038915691 | 0.127601262 | Cell envelope-associate |
| AP53_rna20 | -1.443669783 | 4.503413543 | 5.439937522 | 0.038918056 | 0.127601262 | tRNA-Lys-TTT |
| AP53_381 | -0.414314423 | 5.405779303 | 5.434905205 | 0.038993091 | 0.127631684 | hypothetical protein |
| AP53_762 | -0.642822337 | 9.11980704 | 5.427453226 | 0.039104523 | 0.127780942 | Rhodanese domain pro |

| | | | | | | |
|---|---|---|---|---|---|---|
| AP53_1077 | -0.576089329 | 6.908046039 | 5.385478413 | 0.039739394 | 0.12963725 | FIG01114114: hypothe |
| AP53_191 | 0.311246126 | 8.470630369 | 5.378025005 | 0.03985342 | 0.129791087 | DUF124 domain-contai |
| AP53_1071 | -0.486819995 | 7.377320043 | 5.370576801 | 0.039967759 | 0.129945428 | Transposase (maltose ar |
| AP53_1688 | -0.315080316 | 5.913434124 | 5.360417222 | 0.040124359 | 0.130236422 | Laminin-binding surfac |
| AP53_126 | -0.335047275 | 6.282394857 | 5.344409827 | 0.040372597 | 0.13082339 | V-type ATP synthase sub |
| AP53_279 | 0.327831174 | 10.25874936 | 5.338000484 | 0.040472509 | 0.130928567 | Inner membrane protei |
| AP53_rna15 | -0.971550201 | 3.977905366 | 5.325519335 | 0.040667927 | 0.131341842 | tRNA-Ala-TGC |
| AP53_312 | 0.538138312 | 9.453151372 | 5.316724915 | 0.040806304 | 0.131569828 | hypothetical protein |
| AP53_1411 | 0.329593784 | 8.762450511 | 5.309441146 | 0.040921341 | 0.131721928 | Cell division protein Fts |
| AP53_1381 | 0.451516245 | 7.592901113 | 5.304041216 | 0.041006877 | 0.131778722 | Helicase PriA essential f |
| AP53_276 | 0.431124672 | 8.425597474 | 5.298445474 | 0.041095742 | 0.131846008 | putative arylalkylamine |
| AP53_rna36 | -1.099681307 | 4.138285412 | 5.292220017 | 0.04119488 | 0.13186349 | tRNA-Ala-TGC |
| AP53_214 | 0.239373408 | 9.089887731 | 5.289461529 | 0.0412389 | 0.13186349 | Thiamin pyrophosphok |
| AP53_1678 | 0.37145748 | 8.21689848 | 5.281620152 | 0.041364342 | 0.13186349 | Anthranilate synthase, a |
| AP53_237 | -0.28499497 | 6.217745756 | 5.279566584 | 0.041397269 | 0.13186349 | D-alanyl-D-alanine carb |
| AP53_1711 | 0.815963512 | 3.491323419 | 5.276850406 | 0.04144087 | 0.13186349 | hypothetical protein |
| AP53_pseudo928 | 0.325351754 | 5.682119527 | 5.263670986 | 0.041653216 | 0.132167352 | UDP-N-acetylmuramoy |
| AP53_1465 | -0.21611268 | 8.952617724 | 5.262475046 | 0.04167255 | 0.132167352 | Hypothetical protein ir |
| AP53_1545 | -0.366794737 | 11.4512735 | 5.225705588 | 0.042272276 | 0.133836919 | SSU ribosomal protein S |
| AP53_1379 | 0.25259907 | 9.287515947 | 5.21317799 | 0.042478977 | 0.133836919 | 16S rRNA (cytosine(967 |
| AP53_477 | -0.275373717 | 8.971203091 | 5.211868503 | 0.042500654 | 0.133836919 | FIG01114801: hypothe |
| AP53_16 | -0.541235508 | 9.407115863 | 5.208854006 | 0.042550604 | 0.133836919 | Secreted antigen GbpB/ |
| AP53_1535 | 0.54194571 | 10.61596999 | 5.199417559 | 0.042707424 | 0.133836919 | Transcription terminati |
| AP53_rna51 | 3.118147833 | 5.804361528 | 5.302345444 | 0.042732629 | 0.133836919 | Small Subunit Ribosom |
| AP53_1331 | 0.515455305 | 7.374790994 | 5.19049739 | 0.042856306 | 0.133836919 | hypothetical protein, p |
| AP53_548 | 0.557219721 | 7.536666595 | 5.186127343 | 0.042929472 | 0.133836919 | Streptolysin S biosynth |
| AP53_1113 | 0.24854465 | 7.915725624 | 5.185482585 | 0.042940279 | 0.133836919 | FIG009439: Cytosolic p |
| AP53_74 | -0.311319442 | 6.906016983 | 5.185460059 | 0.042940657 | 0.133836919 | Transcriptional repress |
| AP53_472 | -0.407161019 | 11.25387969 | 5.184458921 | 0.042957445 | 0.133836919 | Phage shock protein C, |
| AP53_110 | 0.344860107 | 8.013028847 | 5.177833349 | 0.043068749 | 0.133968656 | FIG01114239: hypothe |
| AP53_666 | -0.372251762 | 10.2229329 | 5.168940286 | 0.043218691 | 0.134177774 | Translation initiation fa |
| AP53_1281 | 0.360305761 | 9.007721286 | 5.165655549 | 0.043274233 | 0.134177774 | hypothetical protein |
| AP53_780 | -0.334378151 | 9.426762818 | 5.152206323 | 0.043502545 | 0.134191445 | dTDP-4-dehydrorhamn |
| AP53_1230 | 0.536159578 | 7.713718669 | 5.149551066 | 0.043547792 | 0.134191445 | Glutamate transport AT |
| AP53_968 | 0.220974068 | 7.568667338 | 5.14858406 | 0.043564284 | 0.134191445 | Riboflavin kinase (EC 2.7 |

| | | | | | | |
|---|---|---|---|---|---|---|
| AP53_135 | 0.284771115 | 8.773173766 | 5.146718142 | 0.043596129 | 0.134191445 | Streptococcal NAD glyc |
| AP53_222 | -0.35197097 | 12.83297456 | 5.141269418 | 0.04368928 | 0.134191445 | NAD-dependent glycera |
| AP53_1078 | -0.395766108 | 7.501289471 | 5.141025608 | 0.043693453 | 0.134191445 | Excinuclease ABC subur |
| AP53_139 | 0.521456992 | 6.850934411 | 5.133006447 | 0.043831001 | 0.13440122 | Cystathionine beta-lyas |
| AP53_974 | -0.300775222 | 5.52041317 | 5.122646709 | 0.044009467 | 0.134520441 | ABC transporter ATP-bi |
| AP53_1676 | -0.613143436 | 7.594097753 | 5.117784069 | 0.044093538 | 0.134520441 | 6-aminohexanoate-cycl |
| AP53_282 | -0.273982564 | 8.804343559 | 5.114946165 | 0.044142692 | 0.134520441 | FIG005986: HD family I |
| AP53_1547 | -0.240223606 | 8.074757721 | 5.114691872 | 0.0441471 | 0.134520441 | A/G-specific adenine gly |
| AP53_1742 | -0.71577811 | 10.55339526 | 5.090948513 | 0.044560997 | 0.135528297 | Alkyl hydroperoxide red |
| AP53_540 | 0.450890636 | 12.75185578 | 5.087727429 | 0.044617508 | 0.135528297 | LSU ribosomal protein I |
| AP53_rna32 | -1.296603262 | 4.160629324 | 5.077412038 | 0.044799061 | 0.135758948 | tRNA-Gly-TCC |
| AP53_1764 | 0.586816986 | 9.072809522 | 5.075470846 | 0.044833326 | 0.135758948 | UPF0246 protein YaaA |
| AP53_1841 | 0.35660527 | 7.681483501 | 5.055596638 | 0.045185961 | 0.136613628 | Transmembrane compo |
| AP53_199 | 0.354635839 | 6.130953473 | 5.028649185 | 0.045669456 | 0.137730916 | FIG01115631: hypothe |
| AP53_379 | -0.707562576 | 4.715037934 | 5.027100652 | 0.045697429 | 0.137730916 | Bacteriocin BlpU |
| AP53_553 | 0.618822616 | 7.009642797 | 5.006404336 | 0.046073284 | 0.138471269 | Streptolysin S export tr |
| AP53_rna1 | 3.144772848 | 5.84424604 | 5.10192424 | 0.046114564 | 0.138471269 | Small Subunit Ribosom |
| AP53_1712 | 0.478328087 | 5.106717319 | 5.001817392 | 0.04615709 | 0.138471269 | RopB; Rgg-like transcrip |
| AP53_949 | -0.592262179 | 7.838170168 | 4.987567368 | 0.046418619 | 0.138860174 | Putative deoxyribose-sp |
| AP53_681 | -0.426095336 | 9.560920647 | 4.98696002 | 0.046429806 | 0.138860174 | FIG139598: Potential r |
| AP53_207 | 0.283244064 | 7.12594599 | 4.973729967 | 0.046674289 | 0.139376608 | N-acetylmannosamine I |
| AP53_624 | -0.760353414 | 4.183410825 | 4.969359584 | 0.046755392 | 0.139404327 | Phage-associated homi |
| AP53_1519 | 0.273410474 | 7.460775577 | 4.945522005 | 0.047200763 | 0.140516382 | Heme ABC transporter ( |
| AP53_pseudo1337 | -0.764715869 | 2.222517796 | 4.938714546 | 0.047328889 | 0.140682043 | DNA modification meth |
| AP53_1068 | 0.440292971 | 6.318886915 | 4.931404288 | 0.047466947 | 0.140876673 | Maltose/maltodextrin A |
| AP53_rna80 | 3.127137676 | 5.803466699 | 5.011930719 | 0.04773888 | 0.14146743 | Small Subunit Ribosom |
| AP53_13 | 0.39121543 | 9.357275859 | 4.910991019 | 0.047855043 | 0.141595485 | Hypoxanthine-guanine |
| AP53_1520 | 0.275294713 | 7.600032679 | 4.907096831 | 0.047929513 | 0.141599978 | Heme transporter IsdDI |
| AP53_317 | 0.661743138 | 9.054935529 | 4.896844697 | 0.048126241 | 0.141965096 | DNA replication intiatic |
| AP53_rna54 | 6.214022266 | -0.548503406 | 5.145269426 | 0.048264541 | 0.142015362 | 5S RNA |
| AP53_1538 | 0.388116362 | 8.55751471 | 4.888369174 | 0.048289613 | 0.142015362 | dCMP deaminase (EC 3. |
| AP53_154 | 0.297501151 | 9.987487702 | 4.870106514 | 0.048643923 | 0.142754295 | M protein trans-acting |
| AP53_1549 | -0.456081208 | 8.063844327 | 4.865132011 | 0.048740975 | 0.142754295 | Thioredoxin |
| AP53_97 | -0.309715608 | 7.423308075 | 4.864080836 | 0.048761514 | 0.142754295 | Phenylalanyl-tRNA synt |
| AP53_753 | 0.329812523 | 8.807008081 | 4.832501122 | 0.049383437 | 0.144159566 | Uracil-DNA glycosylase, |

| | | | | | | |
|---|---|---|---|---|---|---|
| AP53_1135 | 0.490970011 | 7.158557208 | 4.832167291 | 0.049390063 | 0.144159566 | Chloride channel prote |
| AP53_rna14 | 3.066373665 | 5.834101851 | 4.917016454 | 0.049528296 | 0.14434598 | Small Subunit Ribosom |
| AP53_958 | -0.36312763 | 7.312010699 | 4.818737182 | 0.049657497 | 0.144505551 | Phosphate transport sy |
| AP53_216 | 0.234773065 | 8.772890398 | 4.804421285 | 0.0499445 | 0.145123167 | 3'->5' exoribonuclease I |
| AP53_232 | -0.329435061 | 8.703000013 | 4.794978239 | 0.050134911 | 0.145458687 | Iron-sulfur cluster asser |
| AP53_rna21 | -1.087388349 | 4.774862161 | 4.787997233 | 0.05027624 | 0.145651019 | tRNA-Leu-TAG |
| AP53_20 | 0.496293657 | 7.070122534 | 4.74885823 | 0.051077585 | 0.147631429 | Acyl carrier protein |
| AP53_653 | 0.403522831 | 7.969931946 | 4.747195772 | 0.051111963 | 0.147631429 | Phage protein |
| AP53_1778 | 0.490512998 | 9.281653972 | 4.742385753 | 0.051211588 | 0.147699394 | DNA-3-methyladenine g |
| AP53_1718 | 0.299522717 | 6.728872894 | 4.72397925 | 0.051594998 | 0.148584407 | Pyruvate formate-lyase |
| AP53_345 | -0.554953485 | 8.13500415 | 4.715171019 | 0.051779702 | 0.148655408 | 3-oxoacyl-[acyl-carrier p |
| AP53_542 | 0.201421873 | 9.748023883 | 4.710438819 | 0.051879264 | 0.148655408 | DNA gyrase subunit B (E |
| AP53_515 | 0.295758952 | 6.975175126 | 4.708692352 | 0.051916067 | 0.148655408 | 2-dehydro-3-deoxygluc |
| AP53_115 | 0.463630893 | 5.049902357 | 4.70822123 | 0.051926 | 0.148655408 | Butyrate-acetoacetate ( |
| AP53_rna35 | 3.024797545 | 5.821934084 | 4.787196954 | 0.052110381 | 0.148963548 | Small Subunit Ribosom |
| AP53_893 | 0.434108923 | 10.31994222 | 4.682149487 | 0.052479306 | 0.149797548 | Transcription regulator |
| AP53_rna2 | -0.934448478 | 3.836238549 | 4.677805082 | 0.052572197 | 0.14984234 | tRNA-Ala-TGC |
| AP53_231 | -0.388814435 | 8.46742801 | 4.669901808 | 0.052741693 | 0.150105023 | Iron-sulfur cluster asser |
| AP53_979 | 0.544695892 | 13.36863302 | 4.654277172 | 0.053078734 | 0.150843077 | General stress protein, ( |
| AP53_1524 | 0.388695939 | 10.72683241 | 4.620571172 | 0.05381473 | 0.152528684 | Immunogenic secreted |
| AP53_659 | 0.320896389 | 7.44072087 | 4.619921779 | 0.053829031 | 0.152528684 | Heteropolysaccharide r |
| AP53_1721 | 0.481537894 | 6.061442966 | 4.588967851 | 0.054516063 | 0.154142725 | PTS system, cellobiose-s |
| AP53_rna13 | -0.892107159 | 3.975155994 | 4.587117329 | 0.054557471 | 0.154142725 | tRNA-Pro-TGG |
| AP53_1401 | -0.18965115 | 7.825276973 | 4.576539548 | 0.054794898 | 0.154588514 | Nicotinate phosphoribo |
| AP53_1698 | 0.429113522 | 9.395822895 | 4.561103528 | 0.055143622 | 0.155346545 | Immunogenic secreted |
| AP53_1842 | 0.312585934 | 6.89698383 | 4.555022539 | 0.055281738 | 0.155509933 | ATPase component of g |
| AP53_193 | -0.286156681 | 6.180584908 | 4.547608949 | 0.05545069 | 0.155647623 | Histidine kinase of the c |
| AP53_rna78 | 2.902862272 | 5.81260375 | 4.625821976 | 0.055553594 | 0.155647623 | Large Subunit Ribosom |
| AP53_241 | 0.334835712 | 9.196140432 | 4.536004621 | 0.055716403 | 0.155647623 | Oligopeptide transport |
| AP53_698 | -0.432629121 | 8.420522513 | 4.535536259 | 0.05572716 | 0.155647623 | FIG01115236: hypothe |
| AP53_695 | -0.413360872 | 8.830345711 | 4.534360854 | 0.055754166 | 0.155647623 | SSU ribosomal protein S |
| AP53_1846 | -0.30520731 | 6.160194467 | 4.531853806 | 0.055811822 | 0.155647623 | FIG009210: peptidase, |
| AP53_266 | -0.320120748 | 9.303906861 | 4.519115621 | 0.056105886 | 0.15624322 | COG1399 protein in clu |
| AP53_1775 | 0.471221832 | 11.20879042 | 4.512953236 | 0.056248819 | 0.156292574 | Transcriptional regulato |
| AP53_981 | 0.466709011 | 12.5374196 | 4.511411351 | 0.056284652 | 0.156292574 | FIG01114374: hypothe |

| | | | | | | |
|---|---|---|---|---|---|---|
| AP53_155 | 0.466211551 | 9.707554236 | 4.5074958 | 0.056375771 | 0.156321959 | Co-activator of prophag |
| AP53_1648 | -0.454630821 | 11.11452243 | 4.49772856 | 0.056603846 | 0.156730477 | SSU ribosomal protein S |
| AP53_rna16 | 2.912965507 | 5.830256733 | 4.557792653 | 0.05708824 | 0.157846544 | Large Subunit Ribosom |
| AP53_1323 | -0.765439541 | 2.327562017 | 4.47055939 | 0.05724418 | 0.158052567 | Phage major tail shaft p |
| AP53_1365 | 0.520624999 | 9.690534392 | 4.466649272 | 0.057337057 | 0.158084131 | Regulatory protein recX |
| AP53_1637 | -0.308185001 | 6.216054756 | 4.460543993 | 0.057482441 | 0.158171755 | Hypothetical protein, S |
| AP53_rna56 | 2.69435991 | 0.251749348 | 4.458474701 | 0.057531818 | 0.158171755 | tRNA-Arg-ACG |
| AP53_225 | -0.25025437 | 8.009070346 | 4.421649337 | 0.058419202 | 0.160252162 | Amino acid ABC transpo |
| AP53_1253 | 0.342193522 | 4.832420676 | 4.420233556 | 0.058453648 | 0.160252162 | Late competence prote |
| AP53_262 | -0.28511705 | 7.796758407 | 4.409248302 | 0.058721762 | 0.16076014 | Potassium uptake prote |
| AP53_1815 | -0.573107038 | 9.372363644 | 4.402538992 | 0.058886247 | 0.160983389 | LSU ribosomal protein L |
| AP53_150 | 0.650116829 | 12.31151895 | 4.39219918 | 0.059140835 | 0.161451984 | Glycine betaine ABC tra |
| AP53_894 | 0.526013853 | 9.692295089 | 4.381154387 | 0.059414258 | 0.161957973 | Hemolysin III |
| AP53_rna53 | 2.816424597 | 5.811675989 | 4.455573847 | 0.059493063 | 0.161957973 | Large Subunit Ribosom |
| AP53_1294 | 0.412400259 | 5.511894944 | 4.352012523 | 0.060143086 | 0.163498222 | Integral membrane pro |
| AP53_rna37 | 2.814571317 | 5.803484641 | 4.424553571 | 0.060247299 | 0.163552458 | Large Subunit Ribosom |
| AP53_921 | 0.331410446 | 7.962211143 | 4.331983595 | 0.060650302 | 0.164333123 | Pleiotropic regulator of |
| AP53_1662 | -0.352829453 | 4.998466761 | 4.32986727 | 0.060704199 | 0.164333123 | Glucan 1,6-alpha-gluco |
| AP53_660 | 0.341391833 | 7.324931732 | 4.32619917 | 0.060797753 | 0.164357156 | membrane protein, rela |
| AP53_696 | -0.415488432 | 9.102386373 | 4.302362626 | 0.061409985 | 0.165723144 | KH domain RNA binding |
| AP53_1396 | -0.556240789 | 4.564997894 | 4.299894312 | 0.061473809 | 0.165723144 | FIG005686: hypothetic |
| AP53_941 | -0.533570545 | 8.638612252 | 4.291555165 | 0.061690036 | 0.165743161 | Probable NADH-depend |
| AP53_824 | -0.366838179 | 5.45028148 | 4.285264805 | 0.061853751 | 0.165743161 | Two-component respon |
| AP53_1724 | 0.36775027 | 8.526789921 | 4.283409741 | 0.061902132 | 0.165743161 | Pyruvate formate-lyase |
| AP53_1435 | 0.295612963 | 8.804048446 | 4.283044733 | 0.061911658 | 0.165743161 | FIG01116130: hypothe |
| AP53_879 | -0.187519779 | 8.401266095 | 4.281183173 | 0.061960264 | 0.165743161 | TsaC protein (YrdC dom |
| AP53_1168 | -0.18936399 | 7.247266685 | 4.279908294 | 0.061993578 | 0.165743161 | Recombination protein |
| AP53_754 | 0.401127978 | 9.268599668 | 4.27653999 | 0.062081701 | 0.165750456 | Dihydroorotase (EC 3.5 |
| AP53_289 | -0.244961637 | 7.004993998 | 4.27197947 | 0.062201258 | 0.165841542 | Tyrosine recombinase X |
| AP53_1106 | -0.225998725 | 7.140266528 | 4.267937204 | 0.062307463 | 0.165896825 | Uridine phosphorylase |
| AP53_1822 | 0.194370747 | 7.315602649 | 4.261213609 | 0.062484604 | 0.166140571 | FIG028593: membrane |
| AP53_1125 | -0.34397342 | 7.154776504 | 4.24821929 | 0.062828694 | 0.166826941 | Uridine kinase (EC 2.7.1 |
| AP53_366 | 0.389085772 | 8.651241561 | 4.243505774 | 0.062954076 | 0.166931506 | FIG01114318: hypothe |
| AP53_1069 | 0.513643404 | 5.517887881 | 4.223383142 | 0.063492785 | 0.168130281 | Mobile element protein |
| AP53_1622 | -0.381929683 | 7.523407045 | 4.214567208 | 0.063730564 | 0.168530006 | Galactose-6-phosphate |

| | | | | | | |
|---|---|---|---|---|---|---|
| AP53_1034 | -0.382967975 | 6.332954571 | 4.210091751 | 0.063851688 | 0.16862058 | putative methyltransfe |
| AP53_1180 | 0.467338451 | 6.123507441 | 4.206541562 | 0.06394797 | 0.168645393 | Streptococcal pyrogeni |
| AP53_1826 | 0.381434156 | 6.765204913 | 4.201323273 | 0.064089811 | 0.168790127 | FIG015389: hypothetic |
| AP53_994 | -0.321704387 | 10.55661971 | 4.170629316 | 0.064931912 | 0.170776208 | 6-phosphofructokinase |
| AP53_1416 | 0.429379067 | 9.717240124 | 4.156273193 | 0.065330396 | 0.171579196 | ABC transporter perme |
| AP53_79 | -0.295952186 | 8.410774121 | 4.152396767 | 0.065438504 | 0.171579196 | Multimodular transpep |
| AP53_rna49 | -0.97360873 | 5.221521578 | 4.15010884 | 0.065502413 | 0.171579196 | tRNA-Gln-TTG |
| AP53_rna3 | 2.789344526 | 5.832630068 | 4.216999838 | 0.065607569 | 0.171623035 | Large Subunit Ribosom |
| AP53_192 | -0.307798115 | 9.179226164 | 4.132276558 | 0.066003139 | 0.172425428 | Glutamyl-tRNA synthet |
| AP53_rna70 | 2.760255887 | 5.783091751 | 4.172711632 | 0.066826207 | 0.174176204 | Large Subunit Ribosom |
| AP53_802 | -0.278636585 | 6.417002228 | 4.102391551 | 0.066852793 | 0.174176204 | Hypothetical protein Y |
| AP53_1626 | 0.666497412 | 7.678876515 | 4.0902351 | 0.067202218 | 0.174851884 | Toxin to DNA-damage-i |
| AP53_1752 | 0.347529128 | 6.705251976 | 4.083510818 | 0.067396456 | 0.174959388 | Formiminoglutamase ( |
| AP53_1703 | 0.455894781 | 8.116455482 | 4.082565649 | 0.067423813 | 0.174959388 | periplasmic componen |
| AP53_1171 | 0.378875656 | 6.530184668 | 4.078225365 | 0.067549613 | 0.1750518 | hypothetical protein |
| AP53_877 | -0.196290252 | 8.076302094 | 4.060183642 | 0.06807561 | 0.176179678 | Peptide chain release fa |
| AP53_1530 | 0.514250698 | 7.232502317 | 4.020009606 | 0.069264906 | 0.178812106 | Fructokinase (EC 2.7.1. |
| AP53_940 | -0.45826834 | 8.107856263 | 4.019604609 | 0.069277024 | 0.178812106 | Luciferase-like monoox |
| AP53_1653 | -0.263350877 | 6.900789998 | 4.013343604 | 0.069464686 | 0.179058374 | Transcriptional regulat |
| AP53_598 | 0.584802752 | 9.372826546 | 4.009563564 | 0.069578286 | 0.179113333 | conserved hypothetical |
| AP53_1456 | 0.23982786 | 9.922923369 | 3.999768011 | 0.069873723 | 0.179635623 | COG2740: Predicted nu |
| AP53_315 | 0.511144648 | 8.989167006 | 3.984968045 | 0.070323001 | 0.180410157 | hypothetical protein |
| AP53_133 | 0.439940108 | 10.50319653 | 3.981714674 | 0.070422235 | 0.180410157 | Transcription antitermi |
| AP53_1377 | 0.243529168 | 10.53840346 | 3.980679752 | 0.070453838 | 0.180410157 | Serine/threonine prote |
| AP53_844 | -0.276419042 | 7.79716985 | 3.952389954 | 0.071324445 | 0.182398877 | ABC transporter, peripl |
| AP53_1403 | -0.227911157 | 8.232437424 | 3.948913053 | 0.071432349 | 0.182434459 | Thioredoxin reductase |
| AP53_rna38 | 5.233671714 | -0.468952949 | 4.105964033 | 0.071988219 | 0.183612528 | 5S RNA |
| AP53_782 | -0.408431166 | 6.754178647 | 3.928078225 | 0.072083136 | 0.183613343 | Mutator mutT protein ( |
| AP53_1092 | -0.501690124 | 6.259530038 | 3.924647528 | 0.072190989 | 0.183647063 | FIG01114947: hypothe |
| AP53_858 | 0.226599408 | 9.390154202 | 3.91869231 | 0.072378674 | 0.183835565 | Redox-sensitive transcr |
| AP53_1640 | -0.256042772 | 6.301335828 | 3.916291335 | 0.072454512 | 0.183835565 | FIG01114632: hypothe |
| AP53_1859 | -0.262602368 | 8.358397793 | 3.913202102 | 0.072552232 | 0.183843188 | ABC transporter ATP-bi |
| AP53_685 | 0.250160737 | 8.867203535 | 3.909459125 | 0.072670847 | 0.183903669 | Ribosomal large subuni |
| AP53_rna71 | -1.048379055 | 3.871305666 | 3.900777582 | 0.072946879 | 0.184361839 | tRNA-Ala-TGC |
| AP53_1781 | 0.375053293 | 9.060982847 | 3.888927614 | 0.073325721 | 0.185078316 | DNA mismatch repair p |

| | | | | | | |
|---|---|---|---|---|---|---|
| AP53_82 | 0.370857204 | 8.333549952 | 3.88087928 | 0.073584396 | 0.185421429 | DNA binding protein, Fl |
| AP53_1610 | 0.213183946 | 7.599596944 | 3.878759372 | 0.073652716 | 0.185421429 | Type I restriction-modi |
| AP53_1522 | 0.334151094 | 8.075835223 | 3.864593569 | 0.074111241 | 0.186334092 | Streptococcal cell surfa |
| AP53_1510 | 0.384528051 | 6.923368282 | 3.849741525 | 0.074595729 | 0.187309587 | ABC transporter perme |
| AP53_719 | 0.201874995 | 8.308501386 | 3.841947058 | 0.074851541 | 0.187658529 | COG0488: ATPase comp |
| AP53_583 | -0.729651939 | 11.23882585 | 3.839621956 | 0.074928058 | 0.187658529 | SSU ribosomal protein S |
| AP53_rna44 | -0.892291072 | 3.465197914 | 3.807559893 | 0.075992999 | 0.189935107 | tRNA-Met-CAT |
| AP53_1359 | 0.701201738 | 10.7346197 | 3.806373607 | 0.076032755 | 0.189935107 | Phage antirepressor pro |
| AP53_1683 | -0.320080617 | 6.01827946 | 3.794077818 | 0.076446329 | 0.190425516 | Dipeptide transport sys |
| AP53_539 | 0.448297281 | 8.780216336 | 3.792398156 | 0.076503038 | 0.190425516 | Chloride channel prote |
| AP53_1546 | 0.393215381 | 9.595490849 | 3.791795695 | 0.076523391 | 0.190425516 | hypothetical protein |
| AP53_246 | 0.301416334 | 8.990207353 | 3.779602747 | 0.076936736 | 0.191208968 | FIG001553: Hydrolase, |
| AP53_1700 | 0.332385387 | 8.287535281 | 3.771507911 | 0.077212662 | 0.191649331 | Two-component systen |
| AP53_387 | -0.884407873 | 1.849198738 | 3.767300269 | 0.077356565 | 0.191761293 | hypothetical protein |
| AP53_646 | -0.496667507 | 3.949294806 | 3.757281325 | 0.077700539 | 0.192123882 | hypothetical phage pro |
| AP53_552 | 0.513470476 | 6.275588373 | 3.757273839 | 0.077700797 | 0.192123882 | Streptolysin S export tr |
| AP53_1037 | -0.445298121 | 5.399904477 | 3.742909922 | 0.078197219 | 0.193105346 | FIG01114622: hypothe |
| AP53_518 | 0.196278245 | 7.32366515 | 3.709424183 | 0.079369658 | 0.195751595 | Epoxyqueuosine (oQ) re |
| AP53_976 | 0.189366961 | 9.357629102 | 3.69687114 | 0.079814719 | 0.196599455 | Transcriptional regulat |
| AP53_405 | -0.241749489 | 5.795434844 | 3.679598976 | 0.080432104 | 0.197869092 | Positive transcriptional |
| AP53_1097 | 0.331333416 | 6.178719169 | 3.67446726 | 0.080616661 | 0.198067806 | Malonate permease |
| AP53_684 | 0.259397443 | 7.260847375 | 3.671684204 | 0.080716968 | 0.198067806 | Lipoprotein signal pept |
| AP53_185 | 0.451800036 | 8.235030862 | 3.664931325 | 0.080960991 | 0.198415763 | Lipid A export ATP-binc |
| AP53_1197 | 0.536229049 | 3.318117855 | 3.653890771 | 0.081361906 | 0.199146861 | FIG048485: Phage prot |
| AP53_1825 | 0.270471244 | 6.740475722 | 3.644447534 | 0.081706751 | 0.199739047 | FIG04612: Integral mer |
| AP53_1500 | -0.309284656 | 8.961250047 | 3.632934183 | 0.082129618 | 0.200410848 | Aspartyl-tRNA(Asn) ami |
| AP53_1686 | -1.015430969 | 7.500481105 | 3.631348563 | 0.082188066 | 0.200410848 | FIG01115696: hypothe |
| AP53_1204 | -0.434009716 | 2.871581077 | 3.622838204 | 0.082502638 | 0.200925497 | Recombinational DNA r |
| AP53_1181 | 0.476327879 | 3.955616634 | 3.615410447 | 0.082778402 | 0.201344459 | Phage lysin, N-acetylmu |
| AP53_942 | -0.532417224 | 9.17685393 | 3.594518014 | 0.083560142 | 0.202973689 | Protein:protein lipoyl t |
| AP53_rna27 | -0.672994358 | 4.911695727 | 3.590914454 | 0.083695892 | 0.202973689 | tRNA-Met-CAT |
| AP53_869 | -0.244984312 | 8.672479792 | 3.589163786 | 0.08376194 | 0.202973689 | L-proline glycine betair |
| AP53_219 | -0.404176803 | 11.68278344 | 3.571986437 | 0.084413394 | 0.204297253 | SSU ribosomal protein S |
| AP53_1784 | 0.467983535 | 7.840220968 | 3.564769831 | 0.084688939 | 0.204373784 | Phage transcriptional re |
| AP53_1521 | 0.295519037 | 7.131119651 | 3.561024106 | 0.084832395 | 0.204373784 | Cell surface protein Shp |

| | | | | | | |
|---|---|---|---|---|---|---|
| AP53_205 | 0.243618765 | 6.799939506 | 3.560107443 | 0.084867547 | 0.204373784 | FIG01114161: hypothe |
| AP53_499 | -0.210512646 | 7.651709629 | 3.556773985 | 0.084995529 | 0.204373784 | tRNA-dependent lipid II |
| AP53_988 | 0.426361297 | 9.577451512 | 3.554698628 | 0.085075328 | 0.204373784 | ABC-type polar amino a |
| AP53_634 | -0.570469013 | 4.245122489 | 3.554039177 | 0.085100704 | 0.204373784 | hypothetical phage pro |
| AP53_1154 | 0.396293759 | 6.87505983 | 3.551926397 | 0.085182067 | 0.204373784 | FIG01116792: hypothe |
| AP53_853 | -0.336916583 | 6.647686072 | 3.546724011 | 0.085382817 | 0.204602527 | Lipase/Acylhydrolase fa |
| AP53_39 | 0.312681807 | 10.17853263 | 3.541291432 | 0.08559307 | 0.204853451 | Alcohol dehydrogenase |
| AP53_172 | -0.352520811 | 8.017024394 | 3.526697635 | 0.086161041 | 0.205958843 | tRNA-specific adenosin |
| AP53_649 | 0.495037101 | 6.231483278 | 3.523512532 | 0.086285616 | 0.206002928 | hypothetical phage pro |
| AP53_7 | 0.268056331 | 6.414484418 | 3.515457549 | 0.086601649 | 0.20650344 | Transcription-repair co |
| AP53_596 | 0.675979815 | 8.004957284 | 3.508059125 | 0.086893176 | 0.206944362 | putative repressor C1 |
| AP53_1850 | -0.41427469 | 8.012850345 | 3.48853004 | 0.087668515 | 0.208446314 | FIG01117167: hypothe |
| AP53_1422 | 0.361588201 | 6.883149016 | 3.486775005 | 0.087738608 | 0.208446314 | NADH peroxidase (EC 1. |
| AP53_1517 | 0.208835771 | 7.018966461 | 3.48091369 | 0.087973201 | 0.208748146 | Putative ABC transport |
| AP53_1511 | 0.253337133 | 7.467094329 | 3.476573804 | 0.088147397 | 0.208906102 | putative ABC transport |
| AP53_1426 | 0.374193527 | 6.450441917 | 3.464875281 | 0.088619075 | 0.209612485 | FIG01116532: hypothe |
| AP53_1762 | 0.488520667 | 5.868500859 | 3.462989381 | 0.088695403 | 0.209612485 | Transcriptional regulat |
| AP53_630 | -0.416729734 | 4.836682553 | 3.461162276 | 0.08876943 | 0.209612485 | B-cell receptor associat |
| AP53_1161 | 0.436366478 | 7.283232404 | 3.44610401 | 0.089382428 | 0.210803514 | COG2827: putative enc |
| AP53_19 | 0.298981723 | 7.876750887 | 3.433236061 | 0.089910393 | 0.211619477 | Phosphate:acyl-ACP acy |
| AP53_1376 | 0.254153951 | 6.677968156 | 3.432360535 | 0.089946455 | 0.211619477 | Transporter associated |
| AP53_1274 | 0.418289366 | 6.536350653 | 3.427820426 | 0.090133741 | 0.211803379 | Peptide methionine sul |
| AP53_118 | -0.247006675 | 7.703854256 | 3.402005174 | 0.091207833 | 0.213990784 | Transcriptional regulat |
| AP53_1741 | -0.715511186 | 10.54648531 | 3.400162663 | 0.091285095 | 0.213990784 | Cold shock protein Csp |
| AP53_1527 | 0.210656268 | 10.56430151 | 3.362179702 | 0.09289595 | 0.217504268 | Protein export cytoplas |
| AP53_rna39 | -1.058860067 | 3.543319887 | 3.353800383 | 0.09325602 | 0.217825943 | tRNA-Val-TAC |
| AP53_1593 | -0.58292774 | 8.927438765 | 3.353759433 | 0.093257784 | 0.217825943 | LSU ribosomal protein |
| AP53_692 | 0.464505314 | 8.311700924 | 3.348746876 | 0.093474008 | 0.218068568 | ABC transporter ATP-bi |
| AP53_pseudo595 | -0.27961497 | 6.325910274 | 3.345123654 | 0.093630685 | 0.218171861 | Lipoteichoic acid synth |
| AP53_992 | 0.54285919 | 9.403799049 | 3.319094288 | 0.094765837 | 0.220552145 | Signal peptidase I (EC 3. |
| AP53_1370 | -0.283959945 | 7.087991316 | 3.312602996 | 0.095051565 | 0.220952201 | FIG000605: protein co- |
| AP53_1241 | -0.447480089 | 9.750516897 | 3.299800729 | 0.095618207 | 0.221837738 | Hypothetical protein Y |
| AP53_628 | -0.518626438 | 2.940810817 | 3.297785317 | 0.09570779 | 0.221837738 | Phage integrase: site-sp |
| AP53_973 | -0.22293834 | 5.879572272 | 3.296266117 | 0.095775386 | 0.221837738 | ABC transporter perme |
| AP53_571 | 0.484359497 | 7.058893138 | 3.279700851 | 0.096516285 | 0.223287377 | hypothetical protein |

| | | | | | | |
|---|---|---|---|---|---|---|
| AP53_1085 | -0.487880407 | 4.943668624 | 3.266847553 | 0.097096043 | 0.224361213 | PTS system, cellobiose-: |
| AP53_273 | 0.206694172 | 7.785493392 | 3.236522794 | 0.098481002 | 0.227290874 | SWF/SNF family helicas |
| AP53_581 | -0.177299784 | 7.651479409 | 3.210514985 | 0.099688296 | 0.2296099 | ATP-dependent nucleas |
| AP53_102 | 0.30817108 | 6.187645192 | 3.209786331 | 0.099722383 | 0.2296099 | Transcriptional regulat |
| AP53_1526 | 0.325351071 | 8.884422822 | 3.201149887 | 0.100127505 | 0.229988432 | Holo-[acyl-carrier prote |
| AP53_157 | 0.465047836 | 5.443569593 | 3.201125689 | 0.100128643 | 0.229988432 | FIG01115469: hypothe |
| AP53_570 | 0.374338801 | 9.326678034 | 3.198711519 | 0.100242253 | 0.229988432 | DNA-entry nuclease (Co |
| AP53_1275 | 0.298537027 | 6.163761658 | 3.192214514 | 0.100548794 | 0.230381677 | Thiol:disulfide oxidore |
| AP53_304 | 0.265586354 | 7.635473271 | 3.190052961 | 0.100651037 | 0.230381677 | ABC-type Fe3+-siderop |
| AP53_1571 | 0.36152201 | 6.358480657 | 3.182298625 | 0.101018882 | 0.230951296 | Transcriptional regulat |
| AP53_1438 | 0.221529551 | 8.510849309 | 3.178280057 | 0.101210167 | 0.23096539 | Hydrolase (HAD superfa |
| AP53_1105 | -0.449548791 | 7.325119416 | 3.177171086 | 0.101263033 | 0.23096539 | Phenylacetic acid degra |
| AP53_1063 | 0.29254734 | 6.562298696 | 3.172029564 | 0.101508584 | 0.23125371 | Maltodextrin phosphor |
| AP53_1604 | -0.249249929 | 7.401009618 | 3.167841442 | 0.101709146 | 0.231438983 | Novel pyridoxal kinase, |
| AP53_1780 | 0.398626385 | 8.841926028 | 3.158533446 | 0.102156651 | 0.231785301 | Multidrug resistance pr |
| AP53_rna24 | -0.89507208 | 3.822543586 | 3.15829522 | 0.102168136 | 0.231785301 | tRNA-Leu-TAA |
| AP53_1310 | 0.2738001 | 8.517374264 | 3.15722844 | 0.102219587 | 0.231785301 | Phage protein |
| AP53_pseudo388 | -0.273797938 | 6.228121309 | 3.15109478 | 0.102516036 | 0.231937044 | Mobile element protein |
| AP53_275 | 0.253766842 | 9.150603593 | 3.150899453 | 0.102525494 | 0.231937044 | UDP-N-acetylmuramate |
| AP53_625 | -0.570842797 | 3.57983179 | 3.136943109 | 0.103204081 | 0.233200374 | hypothetical phage pro |
| AP53_744 | 0.203803562 | 8.440934461 | 3.133038577 | 0.103394923 | 0.233359937 | Histidine protein kinase |
| AP53_1572 | 0.234799078 | 6.581863799 | 3.121674175 | 0.103952874 | 0.234346723 | DNA polymerase III epsi |
| AP53_1457 | 0.203677414 | 10.3336213 | 3.116471237 | 0.104209565 | 0.234465091 | Transcription terminati |
| AP53_287 | -0.285488302 | 7.802514992 | 3.113227684 | 0.104369985 | 0.234465091 | FIG009886: phosphoes |
| AP53_40 | 0.585668211 | 10.27616672 | 3.112828416 | 0.104389753 | 0.234465091 | Alcohol dehydrogenase |
| AP53_1104 | -0.230808467 | 8.137301301 | 3.110834077 | 0.104488565 | 0.234465091 | ATP-dependent Zn prot |
| AP53_1857 | -0.582172502 | 7.917900341 | 3.104602702 | 0.104798053 | 0.234888014 | FIG01116071: hypothe |
| AP53_420 | -0.217767899 | 9.55445663 | 3.101762741 | 0.104939479 | 0.234933713 | Proline dipeptidase (EC |
| AP53_609 | -0.476764906 | 4.442814656 | 3.080869783 | 0.105987231 | 0.236841628 | Phage tail fibers |
| AP53_113 | 0.4530866 | 5.138902606 | 3.07990865 | 0.106035742 | 0.236841628 | 3-ketoacyl-CoA thiolase |
| AP53_1725 | 0.479165046 | 6.193449723 | 3.071701913 | 0.10645108 | 0.237262521 | FIG01114281: hypothe |
| AP53_rna18 | -0.880516312 | 4.238330892 | 3.066947194 | 0.106692638 | 0.237262521 | tRNA-Val-TAC |
| AP53_1819 | 0.407170167 | 6.604633592 | 3.065179419 | 0.106782621 | 0.237262521 | FtsK/SpoIIIE family |
| AP53_1798 | 0.518769035 | 4.356196758 | 3.064674312 | 0.106808349 | 0.237262521 | Phage protein |
| AP53_183 | -0.257859453 | 9.708970245 | 3.064144088 | 0.106835365 | 0.237262521 | Glycerol-3-phosphate c |

| | | | | | | |
|---|---|---|---|---|---|---|
| AP53_215 | 0.232295965 | 9.515396733 | 3.060055296 | 0.107043981 | 0.237454133 | DNA recombination pro |
| AP53_531 | -0.182705013 | 7.100928641 | 3.051199502 | 0.107497552 | 0.238188068 | FIG01117156: hypothe |
| AP53_1605 | -0.214386203 | 7.544310561 | 3.048702433 | 0.107625876 | 0.238200485 | tRNA pseudouridine syn |
| AP53_1252 | 0.244095614 | 12.92724852 | 3.038117803 | 0.108171937 | 0.238682764 | Non-specific DNA-bindi |
| AP53_1096 | 0.284102046 | 5.522855 | 3.036925602 | 0.108233658 | 0.238682764 | Quinolinate phosphoril |
| AP53_1454 | 0.334411856 | 11.049852 | 3.035971009 | 0.108283109 | 0.238682764 | Translation initiation fa |
| AP53_948 | -0.437128286 | 8.363698522 | 3.03495718 | 0.10833566 | 0.238682764 | Putative deoxyribose-sp |
| AP53_1032 | -0.460151527 | 5.685132842 | 3.027632828 | 0.108716252 | 0.239249711 | hypothetical protein - p |
| AP53_1091 | -0.31052093 | 8.27044064 | 3.019034055 | 0.109165188 | 0.239965606 | GTP-binding protein Ob |
| AP53_502 | -0.265678707 | 6.893721111 | 3.010151924 | 0.109631339 | 0.240717679 | FIG01114293: hypothe |
| AP53_1542 | -0.217294249 | 9.030240073 | 3.001348204 | 0.110095818 | 0.241464386 | Integral membrane pro |
| AP53_324 | 0.655180615 | 10.21515832 | 2.994858569 | 0.110439773 | 0.241945373 | Lactate 2-monooxygena |
| AP53_667 | -0.376791253 | 10.64619676 | 2.980433034 | 0.111209136 | 0.243356182 | LSU ribosomal protein L |
| AP53_1658 | -0.182410119 | 8.238727787 | 2.95306057 | 0.112687416 | 0.246265477 | Undecaprenyl diphosph |
| AP53_414 | -0.264858123 | 9.780415718 | 2.951134014 | 0.11279238 | 0.246265477 | Pyrrolidone-carboxylat |
| AP53_538 | 0.272380389 | 8.313456641 | 2.944082857 | 0.113177586 | 0.246828869 | Chorismate mutase I (EC |
| AP53_1821 | 0.654711154 | 5.51795796 | 2.940853468 | 0.113354554 | 0.246937362 | Transcriptional regulato |
| AP53_rna55 | -2.147979269 | 0.474471177 | 2.901042084 | 0.115564767 | 0.251469969 | tRNA-Asn-GTT |
| AP53_1759 | 0.310202022 | 6.652835082 | 2.897898699 | 0.115741556 | 0.251572632 | Trehalose operon trans |
| AP53_1084 | -0.53168175 | 5.287488837 | 2.893168217 | 0.116008242 | 0.251870244 | PTS system, cellobiose-s |
| AP53_658 | 0.357746299 | 6.883404077 | 2.889803767 | 0.116198383 | 0.252001186 | Membrane protein |
| AP53_162 | 0.244872307 | 5.480491043 | 2.88423397 | 0.116514013 | 0.252403682 | Quinolinate phosphoril |
| AP53_1145 | 0.393913274 | 7.188093254 | 2.87060029 | 0.117291131 | 0.253524947 | Major facilitator:Oxalat |
| AP53_rna65 | -0.897287653 | 4.639511609 | 2.869714262 | 0.117341858 | 0.253524947 | tRNA-Leu-TAG |
| AP53_1602 | -0.400694982 | 5.230470574 | 2.868023713 | 0.117438722 | 0.253524947 | hypothetical protein |
| AP53_1751 | 0.614512096 | 6.490343622 | 2.863998918 | 0.117669732 | 0.253524947 | Histidine ammonia-lyas |
| AP53_125 | -0.304473568 | 5.993230058 | 2.863481303 | 0.117699482 | 0.253524947 | hypothetical protein |
| AP53_793 | 0.179023444 | 7.110767067 | 2.86146776 | 0.117815302 | 0.253524947 | ABC transporter ATP-bin |
| AP53_431 | 0.365869831 | 6.495646796 | 2.8587784 | 0.117970218 | 0.253577179 | Long-chain-fatty-acid--C |
| AP53_748 | 0.369407796 | 8.60819477 | 2.850289475 | 0.118460877 | 0.2542904 | Orotidine 5'-phosphate |
| AP53_962 | -0.232408325 | 7.398175823 | 2.848144882 | 0.118585237 | 0.2542904 | Phosphate transport sy |
| AP53_1138 | 0.27655039 | 6.670704766 | 2.845094609 | 0.118762396 | 0.2542904 | Membrane-bound prot |
| AP53_1820 | 0.472015939 | 6.814986037 | 2.843999751 | 0.118826065 | 0.2542904 | FIG01114548: hypothe |
| AP53_1277 | 0.365326725 | 10.15144344 | 2.83659625 | 0.119257724 | 0.254933086 | Magnesium and cobalt |
| AP53_1217 | 0.182867656 | 8.364502351 | 2.817795985 | 0.120362699 | 0.2570121 | YfaA |

| | | | | | | |
|---|---|---|---|---|---|---|
| AP53_527 | -0.208464943 | 8.774876702 | 2.813828265 | 0.120597532 | 0.257230559 | Asparaginyl-tRNA synth |
| AP53_807 | -0.266702592 | 6.845634405 | 2.7997434 | 0.121435793 | 0.25873422 | Acyl-ACP thioesterase |
| AP53_124 | -0.309549209 | 5.968741084 | 2.790976642 | 0.121961226 | 0.258887978 | V-type ATP synthase sub |
| AP53_697 | -0.230404766 | 7.56475385 | 2.789228823 | 0.12206632 | 0.258887978 | Cell surface protein |
| AP53_rna31 | -0.876578226 | 3.575514669 | 2.787914569 | 0.12214542 | 0.258887978 | tRNA-Phe-GAA |
| AP53_1271 | 0.309558751 | 7.72486972 | 2.787228464 | 0.122186739 | 0.258887978 | Hypothetical protein Sp |
| AP53_470 | -0.334037059 | 7.956823872 | 2.786781764 | 0.12221365 | 0.258887978 | Transcription accessory |
| AP53_1058 | -0.353023047 | 4.464736719 | 2.784495873 | 0.122351477 | 0.258887978 | hypothetical protein |
| AP53_rna60 | -0.527133141 | 4.476833431 | 2.782468393 | 0.122473886 | 0.258887978 | tRNA-Arg-CCG |
| AP53_734 | -0.53289746 | 8.466451951 | 2.778860238 | 0.122692107 | 0.258887978 | FIG01114189: hypothe |
| AP53_29 | 0.61336761 | 9.432280448 | 2.778591714 | 0.122708367 | 0.258887978 | Phosphoribosylaminoi |
| AP53_792 | 0.196784679 | 7.475883425 | 2.77213903 | 0.123099904 | 0.259432045 | FIG01115226: hypothe |
| AP53_104 | 0.197098889 | 7.441903889 | 2.769027615 | 0.123289258 | 0.259549294 | Signal peptidase I (EC 3. |
| AP53_784 | -0.278257032 | 7.389123837 | 2.760282849 | 0.123823399 | 0.260363943 | TPR-repeat-containing |
| AP53_1287 | 0.234256678 | 8.933424055 | 2.758306713 | 0.123944505 | 0.260363943 | Uncharacterized conser |
| AP53_rna5 | -0.867981784 | 4.152167762 | 2.755156112 | 0.124137893 | 0.260488271 | tRNA-Val-TAC |
| AP53_rna69 | 5.616224634 | -2.070645075 | 3.299425981 | 0.124514312 | 0.260995983 | 5S RNA |
| AP53_572 | 0.316244037 | 8.917857401 | 2.742365961 | 0.124926862 | 0.261379659 | DNA-entry nuclease (Co |
| AP53_964 | -0.190619256 | 6.85891838 | 2.741723173 | 0.124966679 | 0.261379659 | tRNA and rRNA cytosine |
| AP53_448 | 0.742380206 | 1.098269493 | 2.738009576 | 0.125197022 | 0.261579568 | FIG01115329: hypothe |
| AP53_310 | -0.249262126 | 6.753458318 | 2.734706652 | 0.125402339 | 0.261726817 | hypothetical protein |
| AP53_672 | 0.152868025 | 8.457220619 | 2.716329616 | 0.126552432 | 0.263843469 | Chorismate synthase (E |
| AP53_239 | 0.26389125 | 8.619236094 | 2.707331781 | 0.127120365 | 0.26458511 | Oligopeptide transport |
| AP53_839 | -0.226518197 | 9.240711618 | 2.70244213 | 0.127430334 | 0.26458511 | 2-amino-4-hydroxy-6-h |
| AP53_1131 | -0.188274268 | 8.424826186 | 2.701097335 | 0.12751575 | 0.26458511 | Glutaredoxin-like prote |
| AP53_1414 | 0.310740007 | 8.473384018 | 2.698519508 | 0.127679686 | 0.26458511 | Gamma-glutamyl phosp |
| AP53_332 | -0.364354769 | 4.479779567 | 2.697835111 | 0.127723254 | 0.26458511 | FIG01115797: hypothe |
| AP53_307 | -0.291915927 | 7.624681867 | 2.69695215 | 0.12777949 | 0.26458511 | UDP-N-acetylmuramoy |
| AP53_149 | 0.418732178 | 11.85960658 | 2.69295295 | 0.12803459 | 0.26458511 | Glycine betaine ABC tra |
| AP53_814 | 0.234470541 | 5.701081 | 2.692326673 | 0.128074597 | 0.26458511 | Mobile element protein |
| AP53_1101 | -0.214877951 | 6.992606094 | 2.691381814 | 0.128134984 | 0.26458511 | Ribosomal small subun |
| AP53_593 | -0.221082841 | 6.727169012 | 2.669850628 | 0.129520807 | 0.267162472 | putative glycosyltransfe |
| AP53_346 | -0.40825067 | 9.215258084 | 2.66472186 | 0.129853683 | 0.267474861 | NAD-dependent oxidor |
| AP53_1421 | 0.375293497 | 6.927983678 | 2.66327384 | 0.129947859 | 0.267474861 | Trans-acting positive re |
| AP53_652 | 0.325741268 | 8.254094096 | 2.660501791 | 0.130128387 | 0.26756271 | Phage transcriptional r |

| | | | | | | |
|---|---|---|---|---|---|---|
| AP53_1368 | -0.34524263 | 3.964798628 | 2.656896311 | 0.130363662 | 0.267657199 | Competence protein F l |
| AP53_1809 | 0.190537555 | 7.846217249 | 2.655573223 | 0.130450134 | 0.267657199 | Transporter |
| AP53_rna48 | -0.726229472 | 5.123122914 | 2.648008417 | 0.130945927 | 0.268390755 | tRNA-His-GTG |
| AP53_1155 | -0.48619058 | 11.46796564 | 2.63293943 | 0.131940615 | 0.270144233 | Manganese superoxide |
| AP53_178 | 0.602978611 | 11.23122741 | 2.621501139 | 0.13270199 | 0.271416821 | Transcriptional regulat |
| AP53_1112 | -0.300036545 | 9.61875054 | 2.604937134 | 0.133814372 | 0.273357933 | Methionine aminopept |
| AP53_854 | -0.265392478 | 6.218076921 | 2.603185182 | 0.133932712 | 0.273357933 | hypothetical protein |
| AP53_137 | 0.50463477 | 6.003092304 | 2.59026149 | 0.134809756 | 0.274858967 | FIG01114743: hypothe |
| AP53_rna9 | -0.676613048 | 4.351643606 | 2.575568713 | 0.135815656 | 0.276619296 | tRNA-Thr-TGT |
| AP53_238 | 0.159754777 | 9.458554412 | 2.564881718 | 0.136553255 | 0.277749345 | Oligopeptide ABC trans |
| AP53_140 | 0.27885362 | 9.852723686 | 2.563388955 | 0.136656684 | 0.277749345 | Leucyl-tRNA synthetase |
| AP53_1383 | 0.182865484 | 9.539422849 | 2.549052471 | 0.137655063 | 0.279485856 | Guanylate kinase (EC 2. |
| AP53_797 | -0.351982528 | 11.03110948 | 2.542637313 | 0.138104786 | 0.280051263 | Dihydrolipoamide dehy |
| AP53_291 | -0.180333549 | 7.932082255 | 2.540451409 | 0.138258448 | 0.280051263 | Segregation and conder |
| AP53_724 | -0.23197774 | 7.381698237 | 2.538917731 | 0.138366389 | 0.280051263 | Regulation of D-alanyl-l |
| AP53_267 | -0.383168196 | 10.33651236 | 2.533246305 | 0.138766472 | 0.28056846 | Response regulator Csrl |
| AP53_1556 | 0.377856207 | 8.243803113 | 2.517637852 | 0.139875108 | 0.282515697 | RecD-like DNA helicase |
| AP53_1061 | 0.309582847 | 5.182023961 | 2.513769391 | 0.140151604 | 0.282779899 | Phage integrase: site-sp |
| AP53_1289 | -0.207472725 | 8.056980692 | 2.506571533 | 0.140667904 | 0.283526897 | SAM-dependent methyl |
| AP53_1794 | 0.439011284 | 5.449043881 | 2.499434398 | 0.141182223 | 0.284268355 | FIG01114073: hypothe |
| AP53_1103 | 0.282504735 | 7.380225059 | 2.49031514 | 0.141842839 | 0.28530254 | Transporter, MFS super |
| AP53_1810 | 0.265110612 | 7.450918334 | 2.485059487 | 0.142225343 | 0.285775767 | membrane protein, put |
| AP53_1358 | 0.350048351 | 13.07681823 | 2.477566935 | 0.142772904 | 0.286579325 | hypothetical protein - p |
| AP53_rna79 | -0.763712573 | 3.888713338 | 2.474405114 | 0.143004772 | 0.286748205 | tRNA-Ala-TGC |
| AP53_1561 | -0.277264001 | 8.081135354 | 2.470540963 | 0.143288791 | 0.286752395 | Beta-lactamase, class C |
| AP53_1795 | 0.442700509 | 5.452817499 | 2.47035699 | 0.14330233 | 0.286752395 | Phage protein |
| AP53_1776 | 0.18267325 | 10.17093042 | 2.460166455 | 0.144054863 | 0.287816014 | RecA protein |
| AP53_5 | -0.193932529 | 8.003714405 | 2.459146868 | 0.144130431 | 0.287816014 | GTP-binding and nuclei |
| AP53_737 | -0.17785417 | 7.671548301 | 2.4493081 | 0.144862221 | 0.288980033 | Hypothetical protein SI |
| AP53_663 | 0.293303378 | 7.573759383 | 2.443038642 | 0.145330987 | 0.2896175 | Ferredoxin |
| AP53_495 | -0.288957241 | 6.404686299 | 2.439124038 | 0.145624655 | 0.289905082 | FIG01114557: hypothe |
| AP53_411 | 0.239036138 | 10.31818213 | 2.434130911 | 0.146000323 | 0.29035515 | 3'-to-5' exoribonucleas |
| AP53_1585 | -0.39858656 | 8.253436029 | 2.426373586 | 0.146586399 | 0.291076296 | Glutamine synthetase t |
| AP53_1730 | -0.172782483 | 8.408576889 | 2.425060226 | 0.14668592 | 0.291076296 | Similar to ribosomal lar |
| AP53_1102 | -0.349282342 | 6.954638951 | 2.422912831 | 0.146848824 | 0.291076296 | Dihydrolipoamide acyl |

| | | | | | | |
|---|---|---|---|---|---|---|
| AP53_1160 | 0.253207041 | 7.302703675 | 2.421412382 | 0.146962787 | 0.291076296 | tRNA (adenine37-N(6))- |
| AP53_740 | -0.30775138 | 10.38253532 | 2.408627435 | 0.147938394 | 0.292709911 | Phosphopentomutase ( |
| AP53_1802 | 0.480632347 | 3.801703709 | 2.405990929 | 0.148140603 | 0.292811518 | Phage encoded transcri |
| AP53_rna10 | -1.265880245 | 2.562764711 | 2.398492463 | 0.148717624 | 0.293653009 | tRNA-Gly-GCC |
| AP53_1753 | 0.340755005 | 5.306778081 | 2.391132211 | 0.149286783 | 0.294477283 | Transcriptional regulat |
| AP53_374 | -0.274654485 | 8.82937416 | 2.3858806 | 0.149694574 | 0.294981897 | MutT/nudix family prot |
| AP53_1860 | 0.30252603 | 6.937017296 | 2.379690728 | 0.150177038 | 0.295632486 | ABC transporter, perme |
| AP53_891 | 0.24617892 | 11.92681912 | 2.370632596 | 0.15088663 | 0.296436357 | FIG01114048: hypothe |
| AP53_354 | -0.246716925 | 9.752665909 | 2.370579012 | 0.15089084 | 0.296436357 | Manganese ABC transpo |
| AP53_934 | -0.29448907 | 8.425851427 | 2.365793626 | 0.151267447 | 0.296875748 | Cardiolipin synthetase |
| AP53_1053 | -0.399905015 | 7.111292246 | 2.355855447 | 0.152053399 | 0.298116815 | conserved hypothetical |
| AP53_1779 | 0.451969794 | 8.691103742 | 2.35096322 | 0.152442201 | 0.29857751 | Holliday junction DNA |
| AP53_1728 | -0.2028161 | 7.033408301 | 2.334180648 | 0.153785612 | 0.300905115 | Hypothetical protein SI |
| AP53_483 | -0.200324214 | 9.888218567 | 2.331974296 | 0.153963344 | 0.300909115 | Lysyl-tRNA synthetase ( |
| AP53_889 | 0.182327127 | 8.84098912 | 2.329892447 | 0.154131287 | 0.300909115 | Sortase A, LPXTG specifi |
| AP53_1224 | 0.166610341 | 8.260536247 | 2.328388676 | 0.15425274 | 0.300909115 | Octaprenyl diphosphat |
| AP53_31 | 0.244075886 | 7.975795889 | 2.314502554 | 0.155380044 | 0.302467546 | FIG01114657: hypothe |
| AP53_1611 | 0.348552641 | 4.559297688 | 2.311763541 | 0.155603639 | 0.302467546 | Transcriptional regulat |
| AP53_471 | -0.252558003 | 8.036414201 | 2.309851881 | 0.155759937 | 0.302467546 | Putative metallopeptid |
| AP53_1816 | 0.280498258 | 5.463694811 | 2.309726314 | 0.155770211 | 0.302467546 | Cadmium resistance pr |
| AP53_1228 | 0.275066648 | 10.22004434 | 2.30874983 | 0.155850132 | 0.302467546 | putative phosphomanr |
| AP53_778 | 0.189343996 | 7.90192138 | 2.30655088 | 0.1560303 | 0.302467546 | Glycine/D-amino acid c |
| AP53_323 | -0.270808695 | 7.876094137 | 2.303389334 | 0.156289801 | 0.302467546 | Exodeoxyribonuclease |
| AP53_rna40 | -0.9156264 | 3.861167023 | 2.302389297 | 0.156371999 | 0.302467546 | tRNA-Gly-TCC |
| AP53_107 | 0.172419934 | 8.10523054 | 2.300316845 | 0.156542519 | 0.302467546 | FIG01115182: hypothe |
| AP53_1507 | -0.25216664 | 8.566607648 | 2.299498291 | 0.156609935 | 0.302467546 | Universal stress protein |
| AP53_1447 | 0.183656447 | 8.138222723 | 2.297438238 | 0.156779763 | 0.302494552 | PTS system, galactose-s |
| AP53_664 | 0.385900785 | 7.83188895 | 2.294226062 | 0.157045039 | 0.302705482 | FIG01114555: hypothe |
| AP53_980 | 0.566142016 | 11.32503678 | 2.289014797 | 0.157476626 | 0.303236242 | FIG01114232: hypothe |
| AP53_1055 | -0.169881193 | 7.699856721 | 2.286907625 | 0.157651568 | 0.303272243 | hypothetical phage pro |
| AP53_878 | -0.145662266 | 7.663320523 | 2.283703516 | 0.157918054 | 0.303484101 | Protein-N(5)-glutamine |
| AP53_4 | -0.31114173 | 4.818853348 | 2.280513885 | 0.158183906 | 0.303694324 | hypothetical protein |
| AP53_1650 | 0.452296184 | 6.99247406 | 2.276183251 | 0.158545774 | 0.304088288 | FIG01118158: hypothe |
| AP53_37 | 0.432729541 | 8.621551496 | 2.268366585 | 0.159201611 | 0.305044746 | MORN motif family pro |
| AP53_337 | 0.19601883 | 5.666366932 | 2.259802267 | 0.159924157 | 0.306127011 | Mobile element protein |

| | | | | | | |
|---|---|---|---|---|---|---|
| AP53_1505 | -0.216821712 | 10.10113094 | 2.251231242 | 0.160651464 | 0.307216248 | GTP-sensing transcripti |
| AP53_99 | -0.201505924 | 8.792586277 | 2.249029207 | 0.160839002 | 0.307272149 | Deoxyadenosine kinase |
| AP53_rna11 | -0.649579056 | 4.042386343 | 2.24550391 | 0.16113982 | 0.30754414 | tRNA-Leu-TAA |
| AP53_993 | -0.279497891 | 11.49556352 | 2.243102523 | 0.161345144 | 0.307633522 | Pyruvate kinase (EC 2.7. |
| AP53_356 | -0.225021626 | 8.106574505 | 2.235641613 | 0.161985207 | 0.308300814 | Manganese ABC transpo |
| AP53_430 | 0.247510697 | 7.043124354 | 2.2353209 | 0.162012793 | 0.308300814 | 3-ketoacyl-CoA thiolase |
| AP53_330 | 0.326641015 | 10.15185276 | 2.229927357 | 0.162477621 | 0.308827024 | Methionyl-tRNA synthe |
| AP53_1374 | 0.191737645 | 8.160834778 | 2.226726546 | 0.16275428 | 0.308827024 | Two component transc |
| AP53_283 | -0.234759873 | 9.767281081 | 2.226583725 | 0.162766638 | 0.308827024 | FIG005935: membrane |
| AP53_606 | -0.38145974 | 5.885791463 | 2.224609642 | 0.162937583 | 0.308849462 | Phage hyaluronidase |
| AP53_514 | 0.189325517 | 6.573820684 | 2.216599738 | 0.163633558 | 0.309866083 | predicted 4-deoxy-L-thr |
| AP53_1380 | 0.241024233 | 8.52007152 | 2.212350602 | 0.164004305 | 0.310071307 | Methionyl-tRNA formyl |
| AP53_1717 | 0.227549745 | 6.665222329 | 2.211697005 | 0.164061429 | 0.310071307 | Transaldolase (EC 2.2.1. |
| AP53_1649 | 0.327647256 | 7.311415974 | 2.204230509 | 0.164715797 | 0.310870793 | FIG01114948: hypothe |
| AP53_1391 | 0.270103139 | 7.33462873 | 2.203218315 | 0.164804764 | 0.310870793 | D-beta-hydroxybutyrat |
| AP53_1427 | 0.285710158 | 11.35827444 | 2.200674594 | 0.165028614 | 0.310913719 | FIG01116327: hypothe |
| AP53_975 | -0.251475302 | 5.218803085 | 2.199321071 | 0.165147885 | 0.310913719 | Transcriptional regulat |
| AP53_1578 | 0.276627999 | 8.095803619 | 2.195865639 | 0.165452873 | 0.311186072 | Predicted regulator for |
| AP53_1459 | 0.214170295 | 5.692326177 | 2.180437574 | 0.166823439 | 0.313460112 | Mobile element protein |
| AP53_504 | -0.242516099 | 7.698002855 | 2.175046951 | 0.167305747 | 0.314062336 | FIG01114378: hypothe |
| AP53_rna4 | 2.195933896 | -0.831064152 | 2.17130484 | 0.167641611 | 0.314145678 | 5S RNA |
| AP53_402 | 0.270870766 | 4.590131084 | 2.169980727 | 0.167760661 | 0.314145678 | hypothetical protein |
| AP53_1570 | 0.226434655 | 7.812768783 | 2.169146878 | 0.167835687 | 0.314145678 | Transcriptional regulat |
| AP53_711 | -0.519882641 | 2.784300559 | 2.165259864 | 0.168185991 | 0.314498082 | Mobile element protein |
| AP53_226 | 0.128332746 | 7.602630791 | 2.154533489 | 0.169157547 | 0.316010394 | FIG01114729: hypothe |
| AP53_800 | 0.136000931 | 8.470305784 | 2.151835495 | 0.169403054 | 0.316164739 | Putative amidotransfer |
| AP53_rna50 | -0.80871894 | 3.729627845 | 2.146795538 | 0.169862897 | 0.316659534 | tRNA-Leu-CAA |
| AP53_236 | 0.273123218 | 8.985063389 | 2.145356872 | 0.169994454 | 0.316659534 | D-alanyl-D-alanine carb |
| AP53_1146 | 0.294392047 | 9.620017711 | 2.14182533 | 0.170317946 | 0.316957942 | Oligoendopeptidase F |
| AP53_6 | 0.292816887 | 5.955464852 | 2.137172624 | 0.170745345 | 0.317448961 | Peptidyl-tRNA hydrolas |
| AP53_1767 | 0.231115823 | 9.051988918 | 2.13537234 | 0.170911089 | 0.317453038 | Putative oxidoreductas |
| AP53_635 | -0.328149887 | 3.98579451 | 2.128758333 | 0.171521789 | 0.318282785 | conserved hypothetical |
| AP53_1782 | 0.262458925 | 8.412433501 | 2.120269978 | 0.17230967 | 0.319439417 | Phage integrase: site-sp |
| AP53_984 | 0.31877828 | 8.429015997 | 2.116384085 | 0.172671907 | 0.319805507 | Sodium/glycine sympor |
| AP53_1633 | 0.257954006 | 6.935103688 | 2.11079055 | 0.173195048 | 0.319918337 | hypothetical protein |

| | | | | | | |
|---|---|---|---|---|---|---|
| AP53_678 | -0.150256431 | 8.376474821 | 2.110340113 | 0.173237265 | 0.319918337 | tRNA S(4)U 4-thiouridir |
| AP53_764 | 0.218134798 | 7.227939036 | 2.108779918 | 0.173383593 | 0.319918337 | Glutathione S-transfera |
| AP53_1330 | 0.47195488 | 3.811752405 | 2.108689127 | 0.173392113 | 0.319918337 | Phage protein |
| AP53_383 | -0.397412047 | 6.861804942 | 2.091027055 | 0.175059859 | 0.322688686 | putative role in replicat |
| AP53_1818 | 0.268655577 | 5.452001863 | 2.087404747 | 0.175404445 | 0.323017104 | FIG01115119: hypothe |
| AP53_252 | 0.195234268 | 7.357726558 | 2.080012941 | 0.176110333 | 0.323828934 | FIG145533: Methyltrar |
| AP53_439 | 0.439254121 | 2.385210709 | 2.077987832 | 0.176304361 | 0.323828934 | Shikimate 5-dehydroge |
| AP53_1162 | 0.39584023 | 5.587893052 | 2.077555773 | 0.176345793 | 0.323828934 | Kup system potassium u |
| AP53_156 | 0.482186959 | 10.26429078 | 2.072787891 | 0.176803837 | 0.32410266 | Co-activator of prophag |
| AP53_1382 | 0.243725061 | 9.307646867 | 2.071128406 | 0.176963621 | 0.32410266 | DNA-directed RNA poly |
| AP53_795 | -0.328035171 | 10.58190416 | 2.070794586 | 0.176995785 | 0.32410266 | Acetoin dehydrogenase |
| AP53_1677 | 0.184059397 | 7.899458382 | 2.059696554 | 0.178069415 | 0.325715133 | Para-aminobenzoate sy |
| AP53_1263 | -0.540135627 | 10.24910459 | 2.058229265 | 0.178211989 | 0.325715133 | Acetylornithine deacety |
| AP53_467 | 0.192984512 | 8.671118333 | 2.054830856 | 0.178542773 | 0.326012722 | ABC transporter, substr |
| AP53_1836 | -0.241544051 | 7.000800069 | 2.037682845 | 0.180224015 | 0.328773321 | MutT/nudix family prot |
| AP53_1574 | 0.200187196 | 6.894513115 | 2.034637333 | 0.180524741 | 0.329012697 | NAD(FAD)-utilizing dehy |
| AP53_1309 | 0.482643468 | 6.652699164 | 2.028827164 | 0.18110026 | 0.329536405 | FIG01117556: hypothe |
| AP53_1786 | 0.344832242 | 6.068040946 | 2.028309558 | 0.181151645 | 0.329536405 | kilA protein, putative p |
| AP53_280 | 0.141073613 | 8.991995718 | 2.005677847 | 0.183416974 | 0.333344894 | Predicted nucleoside pl |
| AP53_440 | 0.444421936 | 2.590914792 | 2.003804379 | 0.183606138 | 0.333376533 | FIG01114171: hypothe |
| AP53_1838 | -0.245646455 | 6.845123703 | 2.000502012 | 0.183940192 | 0.333670947 | L-serine dehydratase, b |
| AP53_1525 | 0.307741942 | 9.540654293 | 1.986481194 | 0.185367276 | 0.335945736 | Alanine racemase (EC 5. |
| AP53_713 | -0.346637965 | 2.030550548 | 1.982047956 | 0.185821488 | 0.33615038 | Immunoglobulin G-enc |
| AP53_166 | -0.148975523 | 7.535923888 | 1.981998509 | 0.185826563 | 0.33615038 | membrane protein |
| AP53_720 | 0.505233789 | 6.455922938 | 1.975522333 | 0.186492705 | 0.336990275 | FIG01114610: hypothe |
| AP53_rna73 | -0.33864884 | 5.936613833 | 1.974112837 | 0.186638096 | 0.336990275 | tRNA-Ser-GGA |
| AP53_770 | -0.187838919 | 8.138427926 | 1.971142421 | 0.18694498 | 0.337164016 | Ribonuclease Z (EC 3.1.: |
| AP53_850 | 0.180152774 | 5.641731126 | 1.969820783 | 0.187081734 | 0.337164016 | Mobile element proteir |
| AP53_519 | -0.129862174 | 8.866446243 | 1.960409079 | 0.188059349 | 0.338348573 | Peptide chain release fa |
| AP53_857 | 0.187830605 | 8.139744154 | 1.960137656 | 0.18808764 | 0.338348573 | Glutamine amidotransf |
| AP53_997 | 0.2031162 | 8.557511037 | 1.952470395 | 0.188889117 | 0.339475719 | ABC-type multidrug tra |
| AP53_786 | -0.412634324 | 3.925446239 | 1.945413838 | 0.189630679 | 0.340493198 | hypothetical protein |
| AP53_9 | 0.300078024 | 7.0436227 | 1.941691867 | 0.190023338 | 0.340882901 | Cell division protein Div |
| AP53_1441 | 0.143818294 | 8.92053845 | 1.939839207 | 0.190219182 | 0.340919144 | Tagatose 1,6-diphospha |
| AP53_836 | -0.17026341 | 9.379810407 | 1.935767296 | 0.190650547 | 0.341377041 | GTP cyclohydrolase I (E |

| | | | | | | |
|---|---|---|---|---|---|---|
| AP53_765 | 0.26660858 | 8.431233271 | 1.928448687 | 0.191429058 | 0.342455117 | Exfoliative toxin A |
| AP53_661 | 0.22617551 | 8.138897903 | 1.926578527 | 0.191628657 | 0.342496523 | Tripeptide aminopeptic |
| AP53_718 | 0.134746292 | 7.903787725 | 1.924734041 | 0.191825782 | 0.342533434 | tRNA nucleotidyltransf |
| AP53_1229 | 0.153877144 | 6.707222003 | 1.921971882 | 0.192121474 | 0.342543785 | Deoxyribodipyrimidine |
| AP53_1774 | 0.444496535 | 10.47176881 | 1.921383521 | 0.192184535 | 0.342543785 | Hypothetical protein p |
| AP53_838 | -0.30976599 | 8.42697884 | 1.907386112 | 0.193692769 | 0.344528336 | Dihydroneopterin aldo |
| AP53_rna57 | 0.42307063 | 5.964443146 | 1.906944457 | 0.193740608 | 0.344528336 | tRNA-Arg-CCT |
| AP53_339 | -0.1368042 | 6.952917524 | 1.905204444 | 0.193929232 | 0.344528336 | Mobile element protein |
| AP53_1134 | -0.291595176 | 8.551652088 | 1.90447875 | 0.194007971 | 0.344528336 | Ribonucleotide reducta |
| AP53_895 | 0.341347872 | 9.171990467 | 1.90012129 | 0.194481637 | 0.344756548 | FIG01115071: hypothe |
| AP53_907 | -0.266029169 | 4.789085088 | 1.899236948 | 0.194577951 | 0.344756548 | Transcriptional regulat |
| AP53_1373 | -0.27392979 | 9.092152825 | 1.898398417 | 0.194669333 | 0.344756548 | Streptococcal lipoprot |
| AP53_12 | 0.331014922 | 7.130205953 | 1.896558564 | 0.194870034 | 0.34479739 | tRNA(Ile)-lysidine synth |
| AP53_1699 | 0.335390859 | 8.424003252 | 1.886714937 | 0.195948424 | 0.346029338 | Two-component systen |
| AP53_1306 | -0.216548626 | 6.047640193 | 1.885726289 | 0.196057161 | 0.346029338 | RNA methyltransferase, |
| AP53_683 | 0.220124782 | 7.300666495 | 1.885326831 | 0.196101119 | 0.346029338 | Cysteine and methionir |
| AP53_551 | 0.315409213 | 7.020067052 | 1.87834236 | 0.196871788 | 0.347073697 | Export ABC transporter |
| AP53_549 | 0.305958407 | 6.651482013 | 1.872782695 | 0.19748807 | 0.34755776 | Streptolysin S self-immu |
| AP53_1750 | 0.462412352 | 6.210996007 | 1.872634904 | 0.197504487 | 0.34755776 | putative cationic aminc |
| AP53_365 | 0.408617066 | 7.934240173 | 1.869106545 | 0.197896951 | 0.347932954 | Peptide methionine sul |
| AP53_1506 | 0.139378773 | 9.598567028 | 1.86637272 | 0.198201738 | 0.348148651 | Aspartate aminotransfe |
| AP53_1170 | 0.191176562 | 8.008830292 | 1.864790943 | 0.198378366 | 0.348148651 | Substrate-specific comp |
| AP53_367 | 0.221145664 | 10.43533034 | 1.86112053 | 0.198789013 | 0.348554178 | Group B streptococcal : |
| AP53_641 | -0.279555288 | 4.573462804 | 1.85324757 | 0.199673596 | 0.349789214 | hypothetical phage pro |
| AP53_76 | -0.175251031 | 7.413343077 | 1.849767011 | 0.200066301 | 0.350161128 | Zinc ABC transporter, ir |
| AP53_449 | 0.604408787 | 1.82602402 | 1.846123781 | 0.200478443 | 0.350566358 | hypothetical protein |
| AP53_550 | 0.285327535 | 7.078859446 | 1.843617627 | 0.200762597 | 0.350747255 | Streptolysin S biosynth |
| AP53_1862 | 0.381758541 | 8.378678594 | 1.831624397 | 0.202129739 | 0.3525414 | LSU m3Psi1915 methyl |
| AP53_673 | 0.168000466 | 9.971696134 | 1.83142304 | 0.202152797 | 0.3525414 | FIG01116936: hypothe |
| AP53_1001 | 0.411615742 | 7.743620698 | 1.828130249 | 0.202530338 | 0.352882753 | Phage protein |
| AP53_250 | 0.116738589 | 8.649442822 | 1.821452648 | 0.203298813 | 0.353904031 | Hydrolase (HAD superfa |
| AP53_1385 | 0.364249664 | 6.421605107 | 1.812581117 | 0.204325689 | 0.355372906 | Transcriptional regulat |
| AP53_303 | 0.302133754 | 6.727022064 | 1.808019621 | 0.204856329 | 0.355951904 | ABC-type Fe3+-siderop |
| AP53_651 | -0.442523393 | 3.86697987 | 1.806569821 | 0.205025362 | 0.355951904 | hypothetical phage pro |
| AP53_1398 | 0.163105646 | 9.98185083 | 1.785019813 | 0.207559621 | 0.359742721 | Multimodular transpep |

| | | | | | | |
|---|---|---|---|---|---|---|
| AP53_195 | -0.19734667 | 6.691933789 | 1.78485197 | 0.20757952 | 0.359742721 | Response regulator of tl |
| AP53_pseudo1389 | 0.293460751 | 7.021878509 | 1.778721584 | 0.208308047 | 0.360602624 | D-beta-hydroxybutyrat |
| AP53_1830 | 0.189370317 | 5.675547829 | 1.777553296 | 0.208447266 | 0.360602624 | Mobile element proteir |
| AP53_1636 | -0.293986532 | 8.608762784 | 1.77164801 | 0.209152844 | 0.361212121 | 23S rRNA (guanosine-2' |
| AP53_809 | -0.171269949 | 7.618084778 | 1.771489894 | 0.20917178 | 0.361212121 | COG4478, integral men |
| AP53_1443 | 0.141834313 | 8.795064872 | 1.769433855 | 0.209418207 | 0.361316212 | Galactose-6-phosphate |
| AP53_rna12 | -0.827079963 | 3.70589102 | 1.767223815 | 0.209683517 | 0.361452671 | tRNA-Arg-ACG |
| AP53_83 | 0.304266755 | 3.433511599 | 1.759070545 | 0.210666135 | 0.362824284 | Late competence prote |
| AP53_1839 | -0.168659435 | 6.918112071 | 1.751271332 | 0.211611763 | 0.364129815 | L-serine dehydratase, al |
| AP53_220 | -0.277922087 | 11.41052102 | 1.741726417 | 0.212776675 | 0.365736345 | SSU ribosomal protein S |
| AP53_1417 | 0.170228826 | 8.884941693 | 1.740538534 | 0.212922241 | 0.365736345 | Methionine ABC transp |
| AP53_947 | -0.340220325 | 8.593368411 | 1.735215191 | 0.213576193 | 0.36653527 | Nucleoside ABC transpc |
| AP53_1790 | 0.45479894 | 4.848679332 | 1.727767315 | 0.214495589 | 0.367787932 | FIG01118442: hypothe |
| AP53_1763 | 0.289381268 | 7.269390876 | 1.719366376 | 0.215538915 | 0.369250693 | Glyoxalase family prote |
| AP53_158 | 0.331474774 | 4.055778573 | 1.714414374 | 0.216157053 | 0.369983105 | Mobile element proteir |
| AP53_507 | 0.232262345 | 4.440330087 | 1.709543401 | 0.216767364 | 0.37070084 | Heparinase II/III-like |
| AP53_525 | -0.155251356 | 6.856350395 | 1.702286109 | 0.217680901 | 0.371935413 | DinG family ATP-depen |
| AP53_803 | -0.217829923 | 6.424057096 | 1.696680741 | 0.218389985 | 0.372155626 | uncharacterized secrete |
| AP53_967 | 0.38167068 | 9.252986656 | 1.69657341 | 0.218403592 | 0.372155626 | Transcriptional regulat |
| AP53_468 | -0.157852805 | 8.330445525 | 1.695394905 | 0.218553075 | 0.372155626 | possible permease |
| AP53_1567 | 0.281429597 | 8.47442752 | 1.695208591 | 0.21857672 | 0.372155626 | Xaa-Pro dipeptidyl-pep |
| AP53_1251 | 0.312723996 | 6.28830736 | 1.687770937 | 0.219523387 | 0.373439872 | FIG01114576: hypothe |
| AP53_1110 | 0.160219933 | 7.404757561 | 1.686194847 | 0.219724689 | 0.373455009 | 5-Enolpyruvylshikimate |
| AP53_rna41 | -0.956730799 | 4.30478976 | 1.684076752 | 0.219995602 | 0.373588332 | tRNA-Ile-GAT |
| AP53_779 | -0.214172695 | 9.525736977 | 1.682191378 | 0.220237122 | 0.37367155 | Glucose-1-phosphate tl |
| AP53_916 | 0.358543502 | 8.567138716 | 1.677590817 | 0.220827938 | 0.374083085 | Apo-citrate lyase phosp |
| AP53_749 | 0.340639023 | 8.761878754 | 1.677301763 | 0.220865129 | 0.374083085 | Orotate phosphoribosy |
| AP53_899 | -0.160715579 | 8.193042399 | 1.666407719 | 0.222272889 | 0.376139213 | DNA topoisomerase I (E |
| AP53_747 | 0.359892622 | 9.584863311 | 1.652961829 | 0.224026868 | 0.378777135 | FIG01114562: hypothe |
| AP53_990 | 0.314098188 | 8.554446163 | 1.649382775 | 0.224496841 | 0.3792414 | Alkylphosphonate utili: |
| AP53_1625 | 0.341449027 | 6.69419007 | 1.639116919 | 0.225852163 | 0.381199172 | DNA-damage-inducible |
| AP53_206 | 0.183306329 | 6.775523524 | 1.621398681 | 0.228217083 | 0.384856089 | N-acetylneuraminate ly |
| AP53_1137 | 0.208538198 | 5.163380194 | 1.607068801 | 0.230153931 | 0.3877854 | Mobile element proteir |
| AP53_1449 | -0.44473254 | 5.031016365 | 1.593047506 | 0.232070352 | 0.390675241 | Copper chaperone |
| AP53_1357 | 0.419700107 | 4.280988049 | 1.583393085 | 0.233402318 | 0.392577035 | conserved hypothetical |

| | | | | | | |
|---|---|---|---|---|---|---|
| AP53_1568 | 0.389683802 | 4.251335018 | 1.580410685 | 0.233815848 | 0.392932088 | Transcriptional regulat |
| AP53_982 | -0.220830968 | 11.67228753 | 1.578175067 | 0.234126475 | 0.393113743 | FIG01114177: hypothe |
| AP53_28 | 0.454313196 | 9.347749304 | 1.572643831 | 0.234897385 | 0.393772677 | Phosphoribosylamine-- |
| AP53_1777 | 0.209024085 | 7.689044846 | 1.5724486 | 0.234924658 | 0.393772677 | Molybdopterin binding |
| AP53_rna25 | -0.870035367 | 3.774093468 | 1.560573889 | 0.236591459 | 0.39622435 | tRNA-Arg-ACG |
| AP53_690 | 0.190570601 | 8.924203573 | 1.556540517 | 0.23716121 | 0.396836127 | Carbamoyl-phosphate s |
| AP53_rna26 | -0.780861055 | 4.466309467 | 1.547597466 | 0.238431075 | 0.398617327 | tRNA-Pro-TGG |
| AP53_956 | -0.34569204 | 7.171513726 | 1.539469787 | 0.239593088 | 0.400215304 | Two component system |
| AP53_1191 | 0.33590455 | 2.921683598 | 1.53390312 | 0.240393345 | 0.401108109 | Phage protein |
| AP53_917 | 0.359253087 | 8.589404302 | 1.532879598 | 0.240540875 | 0.401108109 | Oxaloacetate decarbox |
| AP53_1439 | 0.190681856 | 8.381408097 | 1.528189984 | 0.241218397 | 0.401892625 | INTEGRAL MEMBRANE P |
| AP53_582 | -0.120049221 | 8.124503157 | 1.518218642 | 0.242667536 | 0.403960281 | Amino acid ABC transp |
| AP53_rna22 | -0.649767854 | 4.187408477 | 1.51181064 | 0.243605003 | 0.404826747 | tRNA-Thr-TGT |
| AP53_1335 | -0.455006263 | 3.024041073 | 1.511000137 | 0.243723923 | 0.404826747 | Phage protein |
| AP53_1268 | -0.225901839 | 5.460772353 | 1.507948733 | 0.244172342 | 0.404826747 | Transcriptional regulat |
| AP53_203 | 0.189583528 | 6.238333279 | 1.505948367 | 0.244466908 | 0.404826747 | ABC transporter, predic |
| AP53_84 | 0.244500904 | 3.193047419 | 1.505853921 | 0.244480828 | 0.404826747 | Late competence prote |
| AP53_1148 | 0.215066588 | 11.24128382 | 1.505247155 | 0.24457028 | 0.404826747 | Ribosomal small subun |
| AP53_626 | -0.40544457 | 3.253685203 | 1.504720184 | 0.244648003 | 0.404826747 | hypothetical phage pro |
| AP53_1670 | -0.254023895 | 6.467137102 | 1.497304244 | 0.245745333 | 0.406054669 | hypothetical protein |
| AP53_1843 | 0.114604512 | 8.406362663 | 1.496878924 | 0.245808468 | 0.406054669 | ATPase component of g |
| AP53_1276 | 0.242747246 | 5.528557162 | 1.491340722 | 0.24663257 | 0.407069574 | Cytochrome c-type bio |
| AP53_1772 | 0.3475216 | 11.24275464 | 1.48234637 | 0.247978925 | 0.408944005 | Ortholog of S. aureus M |
| AP53_1378 | 0.144066261 | 10.25019188 | 1.473668943 | 0.249287269 | 0.410233836 | Protein serine/threonir |
| AP53_30 | 0.463488512 | 10.05230692 | 1.472510628 | 0.24946262 | 0.410233836 | Phosphoribosylaminoi |
| AP53_873 | 0.340034797 | 8.200934288 | 1.471676087 | 0.24958906 | 0.410233836 | Xanthine permease |
| AP53_1095 | -0.342722147 | 3.813457285 | 1.469423212 | 0.249930823 | 0.410233836 | Mobile element proteir |
| AP53_790 | -0.171935862 | 9.013399893 | 1.467675024 | 0.250196461 | 0.410233836 | Ribonuclease J2 (endor |
| AP53_524 | 0.163076225 | 5.69811416 | 1.46667947 | 0.250347907 | 0.410233836 | Mobile element proteir |
| AP53_1690 | 0.385468405 | 11.27051386 | 1.466266845 | 0.250410713 | 0.410233836 | Fibronectin-binding pr |
| AP53_1852 | -0.127013803 | 7.54075747 | 1.46599647 | 0.250451878 | 0.410233836 | FIG002958: hypothetic |
| AP53_1800 | 0.347543942 | 4.960088655 | 1.463919971 | 0.250768337 | 0.410249845 | FIG01115581: hypothe |
| AP53_401 | 0.252344998 | 4.320376361 | 1.462964004 | 0.250914208 | 0.410249845 | FIG01114077: hypothe |
| AP53_699 | 0.247112306 | 5.236852529 | 1.46177544 | 0.251095732 | 0.410249845 | Cobalt-zinc-cadmium r |
| AP53_85 | 0.272768503 | 3.168246607 | 1.45434379 | 0.252234777 | 0.411764257 | Late competence prote |

| | | | | | | |
|---|---|---|---|---|---|---|
| AP53_1803 | -0.310726585 | 6.483353547 | 1.452578393 | 0.252506387 | 0.411861258 | hypothetical protein |
| AP53_1360 | 0.548626063 | 9.222909666 | 1.449114182 | 0.253040517 | 0.412385931 | FIG01117062: hypothe |
| AP53_1599 | -0.24860844 | 9.99443516 | 1.446468121 | 0.253449533 | 0.412705993 | Cell division trigger fact |
| AP53_474 | 0.181190515 | 9.129672143 | 1.441414475 | 0.254233199 | 0.413635071 | Prolipoprotein diacylgl |
| AP53_1151 | -0.153937653 | 7.822189144 | 1.436294271 | 0.255030545 | 0.414507944 | FIG01115059: hypothe |
| AP53_130 | 0.14498232 | 7.733056354 | 1.433901328 | 0.255404351 | 0.414507944 | FIG013576: hypothetic |
| AP53_1404 | 0.209844208 | 6.130712776 | 1.433862925 | 0.255410356 | 0.414507944 | Hypothetical protein SI |
| AP53_163 | -0.301983781 | 3.782831697 | 1.431106684 | 0.255841851 | 0.414861347 | Mobile element proteir |
| AP53_rna81 | -0.474430627 | 8.333760293 | 1.425576794 | 0.256710559 | 0.41588879 | tRNA-Lys-CTT |
| AP53_774 | -0.122005529 | 8.855491958 | 1.424348895 | 0.256903998 | 0.41588879 | Chromosome replicatic |
| AP53_1225 | 0.220219939 | 7.639382578 | 1.42243689 | 0.257205603 | 0.416030062 | Exodeoxyribonuclease \ |
| AP53_rna46 | -0.703872866 | 3.464757328 | 1.416283767 | 0.25817949 | 0.417011227 | tRNA-Tyr-GTA |
| AP53_1823 | 0.14223292 | 5.856257941 | 1.415503436 | 0.258303356 | 0.417011227 | Phosphohydrolase (Mu |
| AP53_1114 | 0.140616597 | 7.467225765 | 1.414537935 | 0.258456726 | 0.417011227 | Ribosomal-protein-S5p |
| AP53_532 | 0.201158273 | 8.611940454 | 1.410156452 | 0.259154289 | 0.417789431 | Candidate zinc-binding |
| AP53_929 | -0.159827713 | 6.614404055 | 1.398023195 | 0.261099429 | 0.420250894 | ABC transporter ATP-bi |
| AP53_138 | -0.324697391 | 10.18656462 | 1.397204385 | 0.261231413 | 0.420250894 | FIG01114903: hypothe |
| AP53_rna17 | 2.2863868 | -0.659752514 | 1.396588979 | 0.26133067 | 0.420250894 | 5S RNA |
| AP53_1174 | 0.569932213 | 3.571063544 | 1.385009535 | 0.263207922 | 0.422919352 | FIG01117945: hypothe |
| AP53_517 | -0.175302034 | 5.639080472 | 1.382203477 | 0.263665611 | 0.423304343 | Beta-phosphoglucomu |
| AP53_pseudo603 | 0.371152283 | 7.457202431 | 1.377675312 | 0.264406485 | 0.424142965 | Phage holin |
| AP53_750 | 0.270492647 | 8.941525709 | 1.373013157 | 0.265172261 | 0.425020115 | 6-aminohexanoate-cycl |
| AP53_1269 | -0.150488473 | 7.264681156 | 1.363934904 | 0.266672134 | 0.426557375 | Arginine pathway regul |
| AP53_1766 | 0.336586247 | 8.876366489 | 1.362590151 | 0.266895296 | 0.426557375 | Acetyltransferase (EC 2. |
| AP53_1257 | 0.140144219 | 7.218499909 | 1.362232693 | 0.266954659 | 0.426557375 | LacI family regulatory p |
| AP53_1202 | -0.270004725 | 3.55737465 | 1.361897072 | 0.267010413 | 0.426557375 | Phage protein |
| AP53_1415 | 0.153056987 | 8.159083617 | 1.35775106 | 0.267700467 | 0.427308064 | Glutamate 5-kinase (EC |
| AP53_835 | -0.326660389 | 10.22822728 | 1.353860422 | 0.268350243 | 0.42799328 | Dihydrofolate synthase |
| AP53_743 | -0.264635905 | 9.690537483 | 1.341996809 | 0.270345006 | 0.430546066 | Purine nucleoside phos |
| AP53_742 | -0.28118892 | 10.24984754 | 1.341704196 | 0.270394464 | 0.430546066 | Purine nucleoside phos |
| AP53_263 | -0.113129674 | 6.84634355 | 1.332551952 | 0.271947673 | 0.43266429 | rRNA small subunit 7-m |
| AP53_608 | -0.337386922 | 5.182205261 | 1.324680447 | 0.273293368 | 0.434449163 | Phage endopeptidase |
| AP53_376 | -0.229948976 | 3.814968251 | 1.319552543 | 0.274174962 | 0.435146055 | Competence-stimulatir |
| AP53_861 | -0.170219604 | 8.722673962 | 1.31952257 | 0.274180126 | 0.435146055 | Ribose-phosphate pyro |
| AP53_86 | 0.388813683 | 3.208787068 | 1.31396363 | 0.275140284 | 0.436128252 | Late competence prote |

| | | | | | | |
|---|---|---|---|---|---|---|
| AP53_543 | 0.13141227 | 9.723083223 | 1.313339463 | 0.275248381 | 0.436128252 | Septation ring formatic |
| AP53_363 | -0.131116313 | 9.754538294 | 1.305275777 | 0.276650192 | 0.437991861 | Ribosome recycling fac |
| AP53_1789 | 0.15365127 | 5.668436909 | 1.300157181 | 0.277545141 | 0.439050627 | Mobile element proteir |
| AP53_1475 | 0.159415275 | 5.649180463 | 1.293921011 | 0.278640908 | 0.440425083 | Mobile element proteir |
| AP53_965 | -0.169767153 | 6.971739361 | 1.287389361 | 0.279795022 | 0.441889452 | Inositol-1-monophospl |
| AP53_1627 | 0.43593665 | 2.557416665 | 1.279810597 | 0.28114247 | 0.443651761 | hypothetical protein |
| AP53_1284 | -0.231952991 | 6.949306293 | 1.278546926 | 0.281368016 | 0.443651761 | Aldose 1-epimerase (EC |
| AP53_26 | 0.464754874 | 9.770366314 | 1.26729546 | 0.283387341 | 0.446473076 | IMP cyclohydrolase (EC |
| AP53_320 | -0.142699537 | 8.206756372 | 1.258737844 | 0.284936695 | 0.448549979 | Cytoplasmic copper ho |
| AP53_1178 | 0.407575015 | 6.83398811 | 1.257320419 | 0.285194455 | 0.448591927 | Paratox |
| AP53_906 | -0.158259016 | 5.05535495 | 1.25552153 | 0.285522053 | 0.448676878 | 2-(5"-triphosphoribosy |
| AP53_451 | -0.92996463 | 0.454461434 | 1.254486508 | 0.285710779 | 0.448676878 | FIG01114600: hypothe |
| AP53_855 | -0.209060313 | 6.633769712 | 1.240709049 | 0.288239595 | 0.452282178 | Permease |
| AP53_870 | -0.129868496 | 8.825899477 | 1.23620621 | 0.289072843 | 0.453202526 | L-proline glycine betair |
| AP53_264 | -0.218400706 | 9.718548387 | 1.235018911 | 0.289293112 | 0.453202526 | LemA protein |
| AP53_1133 | 0.197917151 | 9.591269898 | 1.231369306 | 0.289971659 | 0.453539636 | FIG01116145: hypothe |
| AP53_1544 | -0.184914613 | 11.31501403 | 1.231348011 | 0.289975625 | 0.453539636 | Single-stranded DNA-bi |
| AP53_528 | -0.119965181 | 8.886217142 | 1.226720703 | 0.290839167 | 0.454524012 | FIG000506: Predicted I |
| AP53_343 | -0.121897463 | 6.997685949 | 1.217510899 | 0.292568617 | 0.456858959 | FIG01115464: hypothe |
| AP53_1563 | 0.288006888 | 8.225798925 | 1.215450777 | 0.292957443 | 0.457098389 | Integral membrane pro |
| AP53_406 | -0.147427593 | 6.982701596 | 1.20897388 | 0.294184611 | 0.458644442 | Formamidopyrimidine- |
| AP53_rna58 | -0.497744011 | 3.880953751 | 1.20336478 | 0.295253188 | 0.459627177 | tRNA-Tyr-GTA |
| AP53_601 | 0.343786206 | 8.445616324 | 1.203084505 | 0.295306725 | 0.459627177 | Phage lysin |
| AP53_1395 | -0.39153684 | 7.290890495 | 1.201940681 | 0.295525356 | 0.459627177 | Cell division protein Gp |
| AP53_331 | -0.272999741 | 5.571994503 | 1.198697277 | 0.296146538 | 0.459817584 | hypothetical protein |
| AP53_830 | -0.189422665 | 7.795508083 | 1.198192898 | 0.296243303 | 0.459817584 | GTPase and tRNA-U34 5 |
| AP53_428 | 0.171211942 | 4.324315563 | 1.197592874 | 0.296358474 | 0.459817584 | ABC-type multidrug tra |
| AP53_1460 | -0.15601228 | 9.238176012 | 1.194656471 | 0.296923008 | 0.460217259 | tRNA (guanine46-N7-)-r |
| AP53_141 | 0.164529745 | 5.279381164 | 1.193787938 | 0.297090276 | 0.460217259 | Ascorbate-specific PTS s |
| AP53_759 | -0.34452292 | 8.73137515 | 1.170363677 | 0.301651935 | 0.46691101 | FIG01114490: hypothe |
| AP53_288 | -0.209883427 | 7.785351794 | 1.167999738 | 0.302117761 | 0.467259422 | FIG042801: CBS domai |
| AP53_1537 | 0.180344822 | 10.26151985 | 1.160805963 | 0.303541593 | 0.469087764 | Translation elongation |
| AP53_995 | 0.091747363 | 7.959449408 | 1.156377804 | 0.304422756 | 0.470075234 | DNA polymerase III alpl |
| AP53_577 | -0.326651247 | 8.271785237 | 1.151455784 | 0.305406443 | 0.471219321 | ABC transporter perme |
| AP53_329 | 0.303971395 | 7.559046869 | 1.134018033 | 0.308927911 | 0.476274088 | Membrane protein, put |

| | | | | | | |
|---|---|---|---|---|---|---|
| AP53_868 | 0.164284065 | 8.622066787 | 1.132558826 | 0.309225198 | 0.476354055 | Na+ driven multidrug e |
| AP53_1060 | 0.265563569 | 5.333577603 | 1.129500223 | 0.309849652 | 0.476758813 | hypothetical protein |
| AP53_933 | 0.243821751 | 9.329273809 | 1.128462156 | 0.310061994 | 0.476758813 | Anaerobic ribonucleosi |
| AP53_568 | -0.1273219 | 9.895918157 | 1.127666827 | 0.310224823 | 0.476758813 | UDP-N-acetylglucosami |
| AP53_1807 | 0.161270966 | 9.245643543 | 1.120632554 | 0.311670266 | 0.478601254 | Arginyl-tRNA synthetas |
| AP53_1586 | -0.256794733 | 7.962072421 | 1.117372305 | 0.312343452 | 0.478617095 | Transcriptional regulat |
| AP53_1558 | -0.132286731 | 7.796647122 | 1.117051866 | 0.312409728 | 0.478617095 | DNA polymerase IV (EC |
| AP53_432 | 0.116770126 | 7.462376129 | 1.116926558 | 0.312435652 | 0.478617095 | FIG01114264: hypothe |
| AP53_1467 | 0.136891257 | 9.113828949 | 1.11558582 | 0.312713209 | 0.478617095 | Acetyltransferase (EC 2. |
| AP53_871 | 0.196812807 | 7.585096391 | 1.114456459 | 0.312947279 | 0.478617095 | GMP reductase (EC 1.7. |
| AP53_1223 | 0.178152473 | 8.566335184 | 1.113430781 | 0.313160077 | 0.478617095 | RNA binding methyltra |
| AP53_421 | -0.109733591 | 8.761816951 | 1.110749431 | 0.313717353 | 0.479091568 | Catabolite control prot |
| AP53_1866 | 0.271773557 | 4.323233914 | 1.106058929 | 0.314695605 | 0.480125478 | paratox? |
| AP53_71 | -0.324958631 | 5.566105766 | 1.105133653 | 0.314889095 | 0.480125478 | hypothetical protein |
| AP53_1231 | 0.263535213 | 6.946548077 | 1.103607131 | 0.315208686 | 0.480235525 | ABC transporter memb |
| AP53_479 | -0.120324466 | 8.07023155 | 1.099718884 | 0.31602482 | 0.480904298 | peptidase, U32 family l |
| AP53_745 | 0.31197424 | 5.133878848 | 1.099156249 | 0.316143165 | 0.480904298 | hypothetical protein |
| AP53_1147 | 0.333607117 | 9.234701475 | 1.097837066 | 0.316420893 | 0.480949846 | Competence protein C |
| AP53_657 | 0.166392774 | 7.849295204 | 1.094669987 | 0.317089084 | 0.481302244 | Glycosyl transferase, gr |
| AP53_526 | -0.110917092 | 8.990784056 | 1.094388021 | 0.317148671 | 0.481302244 | Aspartate aminotransfe |
| AP53_103 | 0.122356611 | 6.880115778 | 1.090996567 | 0.317866633 | 0.482014949 | putative collagen bindi |
| AP53_682 | -0.331982771 | 10.79805948 | 1.086283351 | 0.318868278 | 0.483156384 | LSU ribosomal protein |
| AP53_325 | 0.446858564 | 6.18675495 | 1.067085049 | 0.322995377 | 0.488210864 | hypothetical protein |
| AP53_1584 | 0.09648401 | 9.405547829 | 1.066311435 | 0.323163287 | 0.488210864 | Ribonuclease J1 (endor |
| AP53_1664 | 0.139031643 | 7.418852833 | 1.066228448 | 0.323181307 | 0.488210864 | Leucine rich protein |
| AP53_245 | 0.158388513 | 5.525607065 | 1.066095485 | 0.323210181 | 0.488210864 | Mobile element protei |
| AP53_1206 | -0.398908227 | 3.161078979 | 1.05979677 | 0.324582265 | 0.489902159 | Phage protein |
| AP53_394 | 0.207766216 | 4.161062865 | 1.058597645 | 0.324844427 | 0.489916886 | Competence-stimulati |
| AP53_1796 | 0.304798695 | 4.779736854 | 1.054223229 | 0.325803386 | 0.490981656 | DNA primase, phage ass |
| AP53_1136 | 0.298968168 | 5.191885985 | 1.052940439 | 0.326085373 | 0.491025375 | Mobile element protei |
| AP53_636 | -0.201823722 | 4.112039212 | 1.048022042 | 0.327169819 | 0.492276449 | Phage protein |
| AP53_1528 | 0.126517602 | 5.725136531 | 1.04479814 | 0.327883476 | 0.492748798 | Mobile element protei |
| AP53_208 | 0.129634987 | 7.245340904 | 1.044311157 | 0.327991472 | 0.492748798 | Sialic acid utilization re |
| AP53_229 | 0.129648078 | 8.192570383 | 1.042455577 | 0.328403447 | 0.49298615 | Undecaprenyl-phospha |
| AP53_404 | 0.310443583 | 3.441721999 | 1.037591329 | 0.329486963 | 0.494092671 | hypothetical protein |

| | | | | | | |
|---|---|---|---|---|---|---|
| AP53_966 | -0.283805663 | 5.433533808 | 1.036816606 | 0.329660011 | 0.494092671 | FIG006438: hypothetic |
| AP53_1452 | 0.121574852 | 8.900423862 | 1.034335691 | 0.330215051 | 0.494092671 | putative esterase |
| AP53_446 | 0.197309056 | 3.672188595 | 1.033592963 | 0.33038148 | 0.494092671 | UDP-glucose dehydroge |
| AP53_900 | 0.135255547 | 5.716788735 | 1.033450856 | 0.330413337 | 0.494092671 | Mobile element protein |
| AP53_10 | 0.201564878 | 6.92771867 | 1.029223357 | 0.331363077 | 0.494508281 | FIG01114659: hypothe |
| AP53_8 | 0.236541455 | 7.292519567 | 1.029171815 | 0.331374681 | 0.494508281 | Ribosome-associated h |
| AP53_926 | -0.150209201 | 8.964821659 | 1.028812563 | 0.331455576 | 0.494508281 | GMP synthase [glutami |
| AP53_512 | 0.162349409 | 5.679056086 | 1.025882643 | 0.332116396 | 0.495113613 | PTS system, N-acetylgal |
| AP53_196 | 0.095177278 | 8.712936276 | 1.024626795 | 0.332400227 | 0.495156439 | Ribonuclease P protein |
| AP53_1442 | 0.141790701 | 8.913391452 | 1.017136255 | 0.334100481 | 0.497307542 | Tagatose-6-phosphate k |
| AP53_796 | -0.212910353 | 10.79042999 | 1.015780762 | 0.33440951 | 0.497386099 | Dihydrolipoamide acet |
| AP53_721 | 0.248885441 | 7.332076412 | 1.008319062 | 0.33611811 | 0.499500669 | Peptide deformylase (E |
| AP53_359 | 0.290042451 | 7.623428763 | 1.00649908 | 0.336536782 | 0.499500669 | Integral membrane pro |
| AP53_1394 | 0.200827995 | 10.01987976 | 1.006210518 | 0.336603233 | 0.499500669 | FIG001721: Predicted I |
| AP53_1307 | 0.308419952 | 4.02732108 | 1.001487456 | 0.337693598 | 0.500735885 | Paratox |
| AP53_pseudo1680 | 0.213034945 | 5.028817522 | 0.997476296 | 0.338623662 | 0.501322372 | Streptococcal mitogeni |
| AP53_pseudo1183 | 0.304591738 | 4.144345891 | 0.997145586 | 0.33870051 | 0.501322372 | Phage holin |
| AP53_308 | -0.134366514 | 7.965217081 | 0.99410359 | 0.339408585 | 0.501322372 | Membrane protein invo |
| AP53_746 | 0.171633546 | 8.06732286 | 0.993744994 | 0.339492196 | 0.501322372 | Methionine biosynthes |
| AP53_35 | 0.185929736 | 8.844644952 | 0.992883157 | 0.339693268 | 0.501322372 | Holliday junction DNA I |
| AP53_1000 | 0.351669866 | 7.176395449 | 0.992015626 | 0.339895844 | 0.501322372 | Paratox |
| AP53_1582 | -0.144746553 | 8.371673484 | 0.992010317 | 0.339897084 | 0.501322372 | Inactive homolog of me |
| AP53_1167 | -0.111447014 | 8.0253434 | 0.989380439 | 0.340512268 | 0.501848376 | D-alanine--D-alanine lig |
| AP53_865 | 0.139459692 | 7.424202608 | 0.986475186 | 0.341193761 | 0.502471237 | Similar to ribosomal lar |
| AP53_1175 | 0.102483512 | 8.466828996 | 0.978573073 | 0.343057515 | 0.504363624 | Dihydroorotate dehydr |
| AP53_599 | 0.528676733 | 9.392532048 | 0.978551393 | 0.343062648 | 0.504363624 | Phage protein |
| AP53_301 | -0.197555303 | 9.702029137 | 0.977725544 | 0.343258294 | 0.504363624 | Manganese-dependent |
| AP53_1668 | -0.153805923 | 13.35226329 | 0.9755515 | 0.343774112 | 0.504739449 | Collagen-like surface pr |
| AP53_1788 | 0.259555823 | 7.228241139 | 0.974070555 | 0.344126135 | 0.504832968 | FIG01118151: hypothe |
| AP53_293 | -0.121139729 | 7.345830115 | 0.973096385 | 0.344357986 | 0.504832968 | Protein YidD |
| AP53_785 | 0.108964707 | 6.737096775 | 0.97006613 | 0.345080648 | 0.505510595 | Fibronectin/fibrinogen |
| AP53_534 | 0.222154942 | 7.793605183 | 0.96768423 | 0.345650254 | 0.505963155 | Galactosamine-6-phos |
| AP53_rna28 | -0.488166478 | 4.380833458 | 0.962805994 | 0.346821154 | 0.507294544 | tRNA-Met-CAT |
| AP53_382 | -0.330919687 | 5.415796632 | 0.959923811 | 0.347515695 | 0.507927683 | FIG01119121: hypothe |
| AP53_450 | 0.31422235 | 1.699702403 | 0.957722094 | 0.348047639 | 0.508322398 | FIG01116026: hypothe |

| | | | | | | |
|---|---|---|---|---|---|---|
| AP53_181 | 0.220805297 | 6.519016854 | 0.952945705 | 0.349205768 | 0.509516687 | Integral membrane pro |
| AP53_1121 | -0.145699556 | 8.876121787 | 0.952186518 | 0.349390371 | 0.509516687 | DNA polymerase III subu |
| AP53_1399 | -0.15428246 | 10.44428353 | 0.946950139 | 0.350667573 | 0.510658225 | Aminopeptidase C (EC 3 |
| AP53_1246 | -0.098604003 | 8.444518672 | 0.946820268 | 0.350699337 | 0.510658225 | UDP-N-acetylmuramoy |
| AP53_1256 | 0.143380704 | 7.810463382 | 0.944587583 | 0.351246081 | 0.510967616 | FIG01114963: hypothe |
| AP53_872 | 0.19587348 | 7.873853789 | 0.943754403 | 0.351450434 | 0.510967616 | Xanthine phosphoribos |
| AP53_120 | -0.144212067 | 5.734036467 | 0.94273149 | 0.351701563 | 0.510967616 | V-type ATP synthase sub |
| AP53_1196 | -0.360481858 | 3.612372154 | 0.940269485 | 0.352307084 | 0.511248913 | FIG042840: Phage prot |
| AP53_687 | 0.161027509 | 8.680041329 | 0.938307239 | 0.352790797 | 0.511248913 | Uracil permease |
| AP53_1261 | -0.164250461 | 5.821106996 | 0.938093573 | 0.352843527 | 0.511248913 | Aspartate--ammonia lig |
| AP53_1863 | -0.112945775 | 10.1139576 | 0.937667303 | 0.35294876 | 0.511248913 | Serine protease, DegP/H |
| AP53_270 | 0.105184395 | 8.047303966 | 0.931233412 | 0.354542751 | 0.513174855 | Helicase loader DnaB |
| AP53_364 | -0.11018698 | 9.106172219 | 0.920210761 | 0.357298532 | 0.516096234 | S1 RNA binding domair |
| AP53_908 | -0.19436952 | 5.781226351 | 0.919175151 | 0.35755908 | 0.516096234 | Mg2+/citrate complex t |
| AP53_503 | -0.116751652 | 7.641910473 | 0.919151962 | 0.357564917 | 0.516096234 | Deoxyguanosinetripho |
| AP53_1847 | -0.181082195 | 5.744415718 | 0.918914727 | 0.357624644 | 0.516096234 | FIG001621: Zinc protea |
| AP53_876 | -0.142285215 | 7.431415743 | 0.904782212 | 0.361209668 | 0.520683851 | Thymidine kinase (EC 2. |
| AP53_1689 | 0.289987129 | 2.901079833 | 0.903379767 | 0.36156835 | 0.520683851 | hypothetical protein |
| AP53_151 | 0.146895873 | 9.943785209 | 0.902534641 | 0.361784752 | 0.520683851 | DNA polymerase I (EC 2. |
| AP53_1607 | -0.144452063 | 5.26947702 | 0.902176101 | 0.361876617 | 0.520683851 | Glycerate kinase (EC 2.7 |
| AP53_1035 | -0.299291357 | 4.328280742 | 0.900847853 | 0.362217247 | 0.520787909 | FIG01118889: hypothe |
| AP53_1591 | 0.116910389 | 10.34080977 | 0.898640094 | 0.36278449 | 0.52121739 | Dihydroxyacetone kina |
| AP53_1272 | 0.183998401 | 7.465339886 | 0.895165619 | 0.363679885 | 0.521602667 | Two-component sensor |
| AP53_1164 | 0.162642658 | 9.395850668 | 0.894512114 | 0.363848668 | 0.521602667 | Peptide chain release fa |
| AP53_1844 | 0.108246582 | 8.19988329 | 0.894472738 | 0.363858841 | 0.521602667 | CDP-diacylglycerol--gly |
| AP53_1193 | 0.338550692 | 2.503623599 | 0.887026788 | 0.36579032 | 0.52398451 | Phage major tail protei |
| AP53_338 | 0.278024408 | 12.83272165 | 0.884760181 | 0.366381331 | 0.524444073 | FIG01114323: hypothe |
| AP53_1179 | 0.097679231 | 7.530511432 | 0.882317513 | 0.367019856 | 0.524970922 | Streptococcal extracell |
| AP53_1086 | -0.215389126 | 4.62286242 | 0.871448731 | 0.369881393 | 0.528674362 | Transcriptional antiterr |
| AP53_1715 | 0.139704985 | 6.006493938 | 0.86910066 | 0.370504007 | 0.529174597 | Low temperature requi |
| AP53_1613 | 0.272414725 | 3.795444941 | 0.865707932 | 0.371406416 | 0.530073422 | Salivaricin A modificati |
| AP53_285 | -0.165626313 | 7.505391079 | 0.860188439 | 0.372881618 | 0.531787818 | Glutamate racemase (E( |
| AP53_575 | 0.133833852 | 5.702429986 | 0.857204558 | 0.373682814 | 0.532539165 | Salt-stress induced prot |
| AP53_1714 | 0.282818367 | 3.071270221 | 0.853738979 | 0.374616625 | 0.53347826 | FIG01115319: hypothe |
| AP53_1731 | 0.160250489 | 8.042599518 | 0.843828424 | 0.377306662 | 0.536915125 | Hypothetical protein SI |

| | | | | | | |
|---|---|---|---|---|---|---|
| AP53_1696 | -0.114288426 | 8.493555226 | 0.832392183 | 0.38044746 | 0.540987926 | M protein trans-acting |
| AP53_290 | -0.105475621 | 7.280403496 | 0.830969432 | 0.380840981 | 0.541151057 | Segregation and conde |
| AP53_204 | 0.115378912 | 6.544829362 | 0.825838497 | 0.382265327 | 0.542393293 | ABC transporter, predic |
| AP53_412 | 0.260863864 | 9.840928338 | 0.824748136 | 0.382569058 | 0.542393293 | tmRNA-binding protein |
| AP53_1369 | -0.180243747 | 4.662961336 | 0.824682154 | 0.38258745 | 0.542393293 | ComF operon protein A |
| AP53_1598 | -0.295795107 | 8.199124008 | 0.823801766 | 0.382832979 | 0.542393293 | DNA-directed RNA poly |
| AP53_pseudo268 | -0.104669259 | 7.889228161 | 0.803610883 | 0.388530867 | 0.550064487 | Transmembrane histidi |
| AP53_132 | 0.120308489 | 10.33594975 | 0.801221243 | 0.389213834 | 0.550629776 | Nucleoside-binding pro |
| AP53_644 | -0.151133987 | 4.75123839 | 0.800032363 | 0.389554309 | 0.55066266 | Replicative DNA helicas |
| AP53_676 | -0.213371204 | 8.098908033 | 0.799159832 | 0.389804479 | 0.55066266 | Histone acetyltransfera |
| AP53_25 | 0.235339356 | 9.107144995 | 0.796276236 | 0.390633021 | 0.551237984 | Phosphoribosylglycina |
| AP53_1122 | -0.192181869 | 8.656673592 | 0.795106994 | 0.390969752 | 0.551237984 | GAF domain-containing |
| AP53_829 | -0.196068071 | 10.09032713 | 0.794780942 | 0.391063732 | 0.551237984 | Acetylornithine deacet |
| AP53_179 | 0.158355457 | 9.11682714 | 0.790060938 | 0.392428124 | 0.552569649 | FIG01118080: hypothe |
| AP53_768 | -0.088489406 | 8.452878255 | 0.789015891 | 0.392731206 | 0.552569649 | GTP-binding protein Hf |
| AP53_24 | 0.308760678 | 8.609949346 | 0.788417975 | 0.392904775 | 0.552569649 | Phosphoribosylformylg |
| AP53_1797 | 0.250744555 | 3.912813951 | 0.787583595 | 0.393147184 | 0.552569649 | FIG01114328: hypothe |
| AP53_808 | -0.160490193 | 7.800966263 | 0.786336915 | 0.39350981 | 0.552679119 | Hypothetical NagD-like |
| AP53_597 | 0.276449155 | 8.519968898 | 0.779549063 | 0.395493331 | 0.554951125 | hypothetical protein - |
| AP53_1410 | 0.126600041 | 9.086127775 | 0.77884721 | 0.395699308 | 0.554951125 | Cell division protein Ft |
| AP53_106 | 0.107266798 | 7.294570993 | 0.776619021 | 0.396354331 | 0.555382784 | NPQTN specific sortase |
| AP53_108 | 0.109026767 | 7.882968078 | 0.775854857 | 0.39657936 | 0.555382784 | Multiple sugar metabol |
| AP53_1727 | 0.097097911 | 9.545110216 | 0.774829576 | 0.396881595 | 0.555405318 | Multimodular transpep |
| AP53_147 | -0.105337039 | 5.637764626 | 0.773584153 | 0.397249205 | 0.555519242 | Transcriptional antiter |
| AP53_1227 | 0.074295669 | 7.515049606 | 0.768860299 | 0.398648364 | 0.557074496 | Methylenetetrahydrofo |
| AP53_pseudo812 | 0.191294709 | 8.76666172 | 0.767750419 | 0.398978212 | 0.557134324 | Nucleoside diphosphat |
| AP53_1118 | 0.173963834 | 4.120227588 | 0.763949011 | 0.40011119 | 0.558314752 | internalin, putative |
| AP53_1466 | 0.147555087 | 9.729073719 | 0.759965126 | 0.401303947 | 0.55928228 | Cell envelope-associate |
| AP53_944 | -0.109814102 | 7.544989533 | 0.759708923 | 0.401380843 | 0.55928228 | Phosphopantothenoyl |
| AP53_447 | 0.209430932 | 4.439548682 | 0.757238093 | 0.402123613 | 0.559915303 | ABC transporter memb |
| AP53_295 | 0.180948082 | 6.996968242 | 0.750916791 | 0.404033709 | 0.561970738 | Substrate-specific com |
| AP53_rna74 | -0.312218225 | 3.94592579 | 0.750438367 | 0.404178851 | 0.561970738 | tRNA-Leu-AAG |
| AP53_200 | -0.259306504 | 10.29106243 | 0.74897726 | 0.404622622 | 0.562185046 | LSU ribosomal protein |
| AP53_1499 | -0.132620547 | 9.780226003 | 0.744728375 | 0.405917456 | 0.563580674 | Aspartyl-tRNA(Asn) ami |
| AP53_751 | -0.097055175 | 9.44213199 | 0.739512631 | 0.407515856 | 0.565155919 | Cystine ABC transporte |

| | | | | | | |
|---|---|---|---|---|---|---|
| AP53_1612 | 0.16300491 | 4.415013354 | 0.739127231 | 0.407634357 | 0.565155919 | FIG00525014: hypothe |
| AP53_1367 | -0.249580873 | 10.32687169 | 0.737900338 | 0.408011957 | 0.565275666 | Ribosomal subunit inte |
| AP53_1580 | -0.127655815 | 8.126588309 | 0.732528142 | 0.409671862 | 0.567170531 | TsaD/Kae1/Qri7 protein |
| AP53_712 | -0.555348298 | 1.394023947 | 0.729681716 | 0.410555673 | 0.567547012 | hypothetical protein |
| AP53_890 | 0.12196941 | 6.723217123 | 0.729560205 | 0.410593469 | 0.567547012 | FIG01280259: hypothe |
| AP53_1249 | 0.280472086 | 7.710650513 | 0.727957125 | 0.411092621 | 0.567547012 | Rhodanese-like domain |
| AP53_1856 | -0.098433603 | 7.180055165 | 0.727890475 | 0.411113395 | 0.567547012 | Tryptophanyl-tRNA syn |
| AP53_18 | 0.193360458 | 4.922451675 | 0.719401001 | 0.413773054 | 0.570654724 | DNA recombination and |
| AP53_1550 | 0.116811759 | 7.688059374 | 0.718831252 | 0.413952525 | 0.570654724 | Phosphatidylglyceroph |
| AP53_rna83 | -0.553201184 | 3.752216362 | 0.706673133 | 0.41781201 | 0.575566438 | tRNA-Asn-GTT |
| AP53_297 | 0.12695695 | 7.089865324 | 0.704866894 | 0.418390268 | 0.575954263 | COG1242: Predicted Fe |
| AP53_91 | -0.170519755 | 8.79430569 | 0.701142085 | 0.419586785 | 0.577192026 | Acetate kinase (EC 2.7.2 |
| AP53_1497 | -0.159480323 | 7.204863828 | 0.690269294 | 0.423110909 | 0.581627674 | Integral membrane pro |
| AP53_1247 | -0.107703574 | 8.304777516 | 0.688236362 | 0.423775093 | 0.58173216 | FIG01114439: hypothe |
| AP53_251 | 0.138873835 | 7.811421738 | 0.688202 | 0.423786334 | 0.58173216 | Iojap protein |
| AP53_769 | -0.126601815 | 8.463737777 | 0.683204588 | 0.425426256 | 0.583276414 | Cystathionine beta-lyas |
| AP53_530 | 0.122274253 | 9.094709856 | 0.682251946 | 0.425740027 | 0.583276414 | FIG001886: Cytoplasm |
| AP53_386 | 0.301621872 | 3.007733876 | 0.680734352 | 0.426240647 | 0.583276414 | Mobile element protein |
| AP53_837 | -0.164208163 | 9.548222719 | 0.680309929 | 0.426380824 | 0.583276414 | Dihydropteroate synth |
| AP53_1132 | -0.201748374 | 9.079262359 | 0.680210047 | 0.426413824 | 0.583276414 | Ribonucleotide reducta |
| AP53_23 | 0.239053399 | 8.50615647 | 0.678008519 | 0.427142222 | 0.583861305 | Amidophosphoribosylt |
| AP53_864 | 0.099385453 | 7.658762239 | 0.676475696 | 0.427650558 | 0.584144781 | NAD kinase (EC 2.7.1.23 |
| AP53_176 | 0.209999318 | 5.921850499 | 0.674602818 | 0.428272994 | 0.584499002 | hypothetical cytosolic |
| AP53_1446 | -0.082391932 | 8.608058006 | 0.673884382 | 0.428512149 | 0.584499002 | PTS system, galactose-s |
| AP53_417 | 0.166306098 | 7.960783991 | 0.670516928 | 0.429635997 | 0.585620414 | Bactoprenol glucosyl tr |
| AP53_592 | -0.151002535 | 7.379795712 | 0.668405153 | 0.430343211 | 0.586172753 | ABC-transporter (ATP-b |
| AP53_1791 | 0.299300751 | 5.1115042 | 0.667343078 | 0.430699603 | 0.586246795 | FIG01115767: hypothe |
| AP53_271 | 0.082745961 | 9.333956541 | 0.664524816 | 0.431647627 | 0.587125468 | Helicase loader DnaI |
| AP53_1185 | 0.304772719 | 1.757692781 | 0.643698062 | 0.438759933 | 0.596025619 | FIG01116522: hypothe |
| AP53_95 | -0.162025363 | 6.349589747 | 0.643235569 | 0.438920041 | 0.596025619 | FIG014356: hypothetic |
| AP53_1082 | -0.194941764 | 5.178836739 | 0.642681074 | 0.439112125 | 0.596025619 | PTS system, cellobiose- |
| AP53_631 | -0.179005047 | 4.708619 | 0.64036227 | 0.439916882 | 0.596700677 | putaive immunity repre |
| AP53_580 | -0.093132238 | 7.556711597 | 0.639274683 | 0.44029517 | 0.596796736 | ATP-dependent nucleas |
| AP53_710 | -0.130370414 | 7.45928674 | 0.636524054 | 0.441254289 | 0.597483209 | Peptidoglycan hydrolas |
| AP53_249 | 0.151025178 | 8.951446759 | 0.635769971 | 0.44151783 | 0.597483209 | Nicotinate-nucleotide a |

| | | | | | | |
|---|---|---|---|---|---|---|
| AP53_1726 | 0.378484879 | 9.766519192 | 0.635014863 | 0.441781989 | 0.597483209 | Preprotein translocase |
| AP53_1208 | 0.278196099 | 2.736754577 | 0.633628543 | 0.442267641 | 0.597483209 | FIG01114174: hypothe |
| AP53_1187 | 0.222970978 | 2.279544779 | 0.633420103 | 0.442340738 | 0.597483209 | Hyaluronate lyase (phag |
| AP53_689 | 0.197944047 | 9.352042618 | 0.630394246 | 0.443404094 | 0.598462647 | Carbamoyl-phosphate s |
| AP53_351 | -0.143857939 | 9.231868952 | 0.629603972 | 0.443682508 | 0.598462647 | 5'-methylthioadenosin |
| AP53_1569 | 0.137082738 | 7.211031447 | 0.627182151 | 0.444537516 | 0.599199527 | putative repressor - pha |
| AP53_136 | 0.084815377 | 9.356913988 | 0.625593496 | 0.445099858 | 0.599397152 | Thiol-activated cytolysi |
| AP53_142 | -0.160193411 | 4.788570738 | 0.62502395 | 0.445301748 | 0.599397152 | Ascorbate-specific PTS s |
| AP53_2 | 0.106078645 | 8.274963851 | 0.624133731 | 0.445617612 | 0.599406643 | DNA polymerase III beta |
| AP53_194 | -0.114226302 | 6.239699763 | 0.622600688 | 0.44616243 | 0.599723876 | Sensory transduction p |
| AP53_772 | -0.149645158 | 8.155570193 | 0.620477768 | 0.446918701 | 0.600324705 | Single-stranded-DNA-sp |
| AP53_1188 | 0.228517841 | 2.169798669 | 0.618052843 | 0.447785159 | 0.60107261 | Phage hyaluronidase; P |
| AP53_306 | 0.140487888 | 6.313976703 | 0.616863435 | 0.448211168 | 0.601132547 | Ferrichrome transport |
| AP53_187 | 0.168847586 | 7.024537248 | 0.616199629 | 0.448449216 | 0.601132547 | FIG053235: Diacylgluc |
| AP53_1513 | -0.087897407 | 7.803939915 | 0.614576139 | 0.4490323 | 0.601498754 | Duplicated ATPase com |
| AP53_578 | -0.19177802 | 8.24742685 | 0.612481764 | 0.449786367 | 0.60178911 | ABC transporter ATP-bi |
| AP53_1793 | 0.254216684 | 5.339455552 | 0.611952897 | 0.449977115 | 0.60178911 | FIG01114419: hypothe |
| AP53_1450 | 0.079873402 | 7.186068963 | 0.610910022 | 0.450353646 | 0.60178911 | Copper-translocating P |
| AP53_1165 | 0.084347111 | 9.470830918 | 0.610351872 | 0.450555383 | 0.60178911 | membrane protein, put |
| AP53_1785 | -0.254896371 | 5.049146662 | 0.609506947 | 0.450861056 | 0.60178911 | conserved hypothetical |
| AP53_105 | 0.106070587 | 7.031738335 | 0.607785485 | 0.451484909 | 0.60178911 | Fibronectin-binding pro |
| AP53_576 | -0.151903708 | 6.351748009 | 0.606607069 | 0.451912794 | 0.60178911 | FIG01115289: hypothe |
| AP53_1787 | 0.244208961 | 5.888319301 | 0.606392235 | 0.451990873 | 0.60178911 | Phage antirepressor pro |
| AP53_1470 | 0.282526649 | 8.386196245 | 0.606258678 | 0.452039424 | 0.60178911 | Hydrolase (HAD superfa |
| AP53_1553 | 0.132730516 | 7.983589658 | 0.600493316 | 0.454143608 | 0.604175972 | FIG01114225: hypothe |
| AP53_426 | -0.145499399 | 3.992919516 | 0.593578236 | 0.456689099 | 0.607146261 | Daunorubicin resistanc |
| AP53_1308 | 0.153895365 | 7.466925505 | 0.587734615 | 0.458858902 | 0.609613367 | Streptococcal extracell |
| AP53_255 | -0.117002889 | 9.625319323 | 0.586730265 | 0.459233576 | 0.609693824 | Cysteine ABC transport |
| AP53_523 | 0.216614773 | 9.522155055 | 0.576794351 | 0.462968165 | 0.614231858 | 2,3-butanediol dehydro |
| AP53_457 | 0.393677458 | 2.788920823 | 0.570306651 | 0.465434518 | 0.61708224 | FIG01117988: hypothe |
| AP53_116 | 0.162350444 | 7.765070502 | 0.565721668 | 0.467191035 | 0.618988259 | YheO-like PAS domain |
| AP53_1124 | -0.154871202 | 4.615390786 | 0.564785578 | 0.467551042 | 0.619042682 | FIG01113965: hypothe |
| AP53_1665 | 0.330005999 | 6.355545902 | 0.563927729 | 0.467881373 | 0.619057768 | Streptokinase |
| AP53_1100 | -0.149197001 | 6.426479153 | 0.560413577 | 0.469238734 | 0.620430778 | Mobile element protein |
| AP53_1629 | 0.326346274 | 2.025165839 | 0.558740546 | 0.469887317 | 0.620865407 | Protein of unknown fur |

| | | | | | | |
|---|---|---|---|---|---|---|
| AP53_174 | 0.12962017 | 6.388664455 | 0.554267115 | 0.471629091 | 0.622440564 | Streptococcal pyrogeni |
| AP53_rna61 | -0.565538665 | 1.540564622 | 0.554032366 | 0.471720798 | 0.622440564 | tRNA-Thr-GGT |
| AP53_1099 | 0.133627934 | 5.804455639 | 0.539964381 | 0.477273157 | 0.629339129 | Mobile element proteir |
| AP53_1739 | 0.276185599 | 9.058686756 | 0.535600948 | 0.479018292 | 0.631211476 | Transcriptional regulat |
| AP53_335 | 0.169394235 | 8.142871266 | 0.530830787 | 0.480938801 | 0.63331222 | Ribonucleotide reducta |
| AP53_1288 | 0.077643236 | 9.249318484 | 0.52897922 | 0.481687868 | 0.633868577 | hypothetical protein |
| AP53_645 | -0.22016812 | 4.180054042 | 0.526575955 | 0.482663165 | 0.634721683 | hypothetical protein |
| AP53_1444 | 0.071510666 | 9.09676772 | 0.523949046 | 0.48373317 | 0.635698092 | Galactose-6-phosphate |
| AP53_159 | 0.150836736 | 5.702756779 | 0.518529355 | 0.485953895 | 0.637587708 | Mobile element proteir |
| AP53_90 | -0.085676884 | 7.485381372 | 0.517663975 | 0.48631014 | 0.637587708 | Adenine-specific methy |
| AP53_27 | 0.30537653 | 7.905731302 | 0.517299225 | 0.486460431 | 0.637587708 | putative choline bindir |
| AP53_152 | 0.098289837 | 9.753289459 | 0.515936919 | 0.487022475 | 0.637587708 | Succinyl-CoA synthetas |
| AP53_1036 | -0.203043984 | 5.48739783 | 0.51577823 | 0.48708802 | 0.637587708 | conserved hypothetical |
| AP53_1761 | 0.248227393 | 4.562062021 | 0.515201911 | 0.48732619 | 0.637587708 | FIG01114717: hypothe |
| AP53_1514 | 0.085770956 | 7.701027762 | 0.514060422 | 0.487798529 | 0.637587708 | Transmembrane compo |
| AP53_725 | -0.124847849 | 7.957002543 | 0.514059424 | 0.487798942 | 0.637587708 | Regulation of D-alanyl-l |
| AP53_1502 | 0.090891048 | 5.633125602 | 0.511073319 | 0.489038386 | 0.638465543 | Mobile element proteir |
| AP53_1806 | 0.103477069 | 6.705292417 | 0.510856927 | 0.48912842 | 0.638465543 | Arginine pathway regul |
| AP53_347 | -0.082326015 | 7.310843593 | 0.509880132 | 0.489535195 | 0.638567079 | Glycerol-3-phosphate t |
| AP53_445 | 0.229490599 | 3.478242748 | 0.508242429 | 0.490218538 | 0.639029001 | Dolichyl-phosphate ma |
| AP53_1219 | -0.068903884 | 8.732265628 | 0.503640036 | 0.492147971 | 0.641113564 | Hypothetical protein D |
| AP53_14 | 0.101899995 | 10.75548072 | 0.50121041 | 0.493171955 | 0.64201661 | Cell division protein Fts |
| AP53_358 | -0.072910393 | 7.203562096 | 0.498413013 | 0.494355629 | 0.643126191 | Cell division protein Fts |
| AP53_1413 | 0.246565536 | 6.936292989 | 0.495053421 | 0.495783876 | 0.644552246 | FIG01115263: hypothe |
| AP53_880 | -0.072899595 | 8.829390239 | 0.489575473 | 0.498128518 | 0.646834424 | Phosphinothricin N-ace |
| AP53_1592 | 0.081406788 | 9.927094547 | 0.489166408 | 0.498304398 | 0.646834424 | FIG001802: Putative al |
| AP53_886 | -0.128565139 | 11.36905956 | 0.48857517 | 0.498558802 | 0.646834424 | NADH peroxidase (EC 1. |
| AP53_544 | 0.122572873 | 8.092722271 | 0.487600032 | 0.498978901 | 0.646834424 | FIG01114091: hypothe |
| AP53_1523 | 0.130678706 | 4.841995829 | 0.486598995 | 0.499410817 | 0.646834424 | hypothetical protein |
| AP53_648 | 0.203790579 | 7.381473531 | 0.48617631 | 0.499593394 | 0.646834424 | Phage antirepressor prc |
| AP53_723 | -0.142939228 | 7.192492431 | 0.48553174 | 0.499872044 | 0.646834424 | GTP pyrophosphokinas |
| AP53_1109 | 0.075818114 | 6.796823817 | 0.479859712 | 0.502336156 | 0.649406894 | Shikimate kinase I (EC 2 |
| AP53_1260 | 0.095366069 | 8.863973916 | 0.479417492 | 0.502529188 | 0.649406894 | 16S rRNA (guanine(966 |
| AP53_1245 | -0.070056342 | 8.936590676 | 0.475377289 | 0.504298977 | 0.650951137 | UDP-N-acetylglucosami |
| AP53_1186 | 0.179088422 | 2.275852303 | 0.47515903 | 0.504394905 | 0.650951137 | Phage hyaluronidase |

| | | | | | | |
|---|---|---|---|---|---|---|
| AP53_739 | -0.157144168 | 10.05085262 | 0.473388489 | 0.505174298 | 0.651523796 | Ribose 5-phosphate iso |
| AP53_827 | 0.143515685 | 8.427106352 | 0.470822861 | 0.506307563 | 0.652485385 | Excinuclease ABC subur |
| AP53_1858 | 0.128414172 | 7.645699048 | 0.470180081 | 0.506592208 | 0.652485385 | hypothetical protein BI |
| AP53_1279 | -0.096376009 | 6.54295096 | 0.469029128 | 0.507102614 | 0.652709664 | FIG01114815: hypothe |
| AP53_482 | 0.103770275 | 7.245107736 | 0.464438144 | 0.509147873 | 0.654626732 | FIG01116574: hypothe |
| AP53_943 | -0.096973656 | 7.179684337 | 0.464172791 | 0.509266546 | 0.654626732 | Phosphopantothenoylc |
| AP53_1461 | -0.068844731 | 9.01395109 | 0.460978918 | 0.510698886 | 0.656033446 | aminoglycoside phospl |
| AP53_971 | 0.132171546 | 7.836608819 | 0.457126021 | 0.512436592 | 0.657830307 | membrane protein |
| AP53_930 | 0.113532841 | 6.355157634 | 0.456359139 | 0.512783757 | 0.657840893 | Substrate-specific comp |
| AP53_1149 | 0.131448424 | 11.11120345 | 0.45157301 | 0.514960207 | 0.660045718 | Glucosamine-6-phosph |
| AP53_1254 | 0.054412241 | 8.539635484 | 0.451086233 | 0.515182515 | 0.660045718 | Ribosomal RNA large su |
| AP53_533 | 0.180658363 | 7.028844904 | 0.450167717 | 0.515602478 | 0.660148027 | Transcriptional regulat |
| AP53_3 | 0.18663974 | 5.923167754 | 0.441543376 | 0.519576704 | 0.664567422 | FIG01114317: hypothe |
| AP53_1412 | 0.079150538 | 8.859933775 | 0.441193834 | 0.519738973 | 0.664567422 | rRNA small subunit met |
| AP53_415 | 0.101964058 | 9.242783866 | 0.4399862 | 0.520300321 | 0.664847217 | FIG001614: Membrane |
| AP53_1290 | 0.072417711 | 6.889385727 | 0.438885785 | 0.52081281 | 0.665064253 | Shikimate/quinate 5-de |
| AP53_913 | -0.128629746 | 7.533091996 | 0.434174331 | 0.523017702 | 0.667440736 | Citrate lyase gamma cha |
| AP53_352 | 0.096085359 | 8.762418006 | 0.431351765 | 0.524346974 | 0.668669597 | FIG01116268: hypothe |
| AP53_898 | 0.222302529 | 6.544989312 | 0.430668607 | 0.524669653 | 0.668669597 | Rossmann fold nucleoti |
| AP53_22 | 0.186249258 | 8.533996792 | 0.428981939 | 0.525467917 | 0.669247524 | Phosphoribosylformylg |
| AP53_1573 | 0.096544979 | 8.771126769 | 0.423887035 | 0.527893115 | 0.671545864 | Transcriptional regulat |
| AP53_1543 | -0.101275253 | 10.58250761 | 0.423737848 | 0.527964445 | 0.671545864 | SSU ribosomal protein S |
| AP53_248 | -0.096115285 | 8.921296728 | 0.422632682 | 0.528493422 | 0.671778475 | FIG004454: RNA bindir |
| AP53_811 | 0.207205163 | 11.91569231 | 0.419986413 | 0.5297641 | 0.672952958 | Collagen-like surface pr |
| AP53_362 | -0.082470777 | 9.437692988 | 0.416562778 | 0.531416636 | 0.674426926 | Uridylate kinase (EC 2.7 |
| AP53_1189 | 0.170081883 | 1.973860912 | 0.416144153 | 0.53161937 | 0.674426926 | FIG022125: Phage-asso |
| AP53_591 | -0.088378736 | 6.796622764 | 0.408516685 | 0.535339101 | 0.678497934 | rhamnose-containing p |
| AP53_1387 | 0.170650944 | 7.519770716 | 0.408133093 | 0.535527478 | 0.678497934 | Butyrate-acetoacetate C |
| AP53_247 | -0.095596116 | 8.868964733 | 0.406890158 | 0.536138736 | 0.67882928 | GTP-binding protein Yq |
| AP53_17 | -0.112293644 | 9.016055634 | 0.398597298 | 0.540251476 | 0.683590687 | Ribose-phosphate pyro |
| AP53_11 | 0.074115438 | 7.488083093 | 0.392712071 | 0.543207175 | 0.686882819 | FIG01114299: hypothe |
| AP53_828 | -0.119973956 | 9.672546088 | 0.387635921 | 0.545781782 | 0.689689088 | Oxygen-insensitive NAD |
| AP53_1559 | -0.180558456 | 11.45855556 | 0.38612301 | 0.546553717 | 0.690028514 | Pyruvate formate-lyase |
| AP53_341 | 0.165557614 | 3.9403568 | 0.385716707 | 0.546761388 | 0.690028514 | hypothetical protein |
| AP53_1388 | 0.135822909 | 7.381478696 | 0.381309891 | 0.549023745 | 0.691753562 | Butyrate-acetoacetate C |

| | | | | | | |
|---|---|---|---|---|---|---|
| AP53_1169 | 0.119165827 | 6.773293348 | 0.381192114 | 0.54908446 | 0.691753562 | Formate/nitrite family ( |
| AP53_78 | -0.074257145 | 7.419845793 | 0.380973011 | 0.549197444 | 0.691753562 | Tyrosyl-tRNA synthetase |
| AP53_409 | 0.077406025 | 9.444312881 | 0.379786967 | 0.54980984 | 0.69207581 | Multidrug resistance ef |
| AP53_637 | -0.112701311 | 4.800871695 | 0.377852645 | 0.550811475 | 0.69288728 | hypothetical phage pro |
| AP53_1560 | -0.105814179 | 7.297212437 | 0.373801215 | 0.552921061 | 0.695090531 | hypothetical protein |
| AP53_951 | 0.104079923 | 9.597594622 | 0.371778039 | 0.553980505 | 0.695832057 | Cytidine deaminase (EC |
| AP53_1083 | -0.217737695 | 4.977482168 | 0.370935187 | 0.554423051 | 0.695832057 | Putative uncharacterize |
| AP53_708 | -0.106198665 | 9.580698848 | 0.37037179 | 0.554719258 | 0.695832057 | PTS system, fructose-sp |
| AP53_305 | 0.125938252 | 6.571725935 | 0.36948348 | 0.555186925 | 0.695832057 | Ferrichrome-binding pe |
| AP53_418 | -0.065525668 | 8.8980063 | 0.369184163 | 0.555344681 | 0.695832057 | Lactoylglutathione lyas |
| AP53_1286 | 0.141184237 | 9.381978509 | 0.368582861 | 0.55566187 | 0.695832057 | 3-dehydroquinate synth |
| AP53_1757 | 0.166075152 | 8.184491381 | 0.363852778 | 0.5581696 | 0.69817061 | Trehalose-6-phosphate |
| AP53_397 | 0.247627976 | 3.124222425 | 0.363132464 | 0.558553463 | 0.69817061 | hypothetical protein |
| AP53_1371 | -0.134292509 | 10.82174338 | 0.363029404 | 0.558608428 | 0.69817061 | Cysteine synthase (EC 2. |
| AP53_218 | 0.070142488 | 10.12084754 | 0.358026242 | 0.561289816 | 0.701070485 | Surface exclusion prote |
| AP53_1392 | -0.072923933 | 8.160192567 | 0.354034932 | 0.563447486 | 0.703312907 | S-ribosylhomocysteine |
| AP53_rna45 | -0.313899839 | 3.076635878 | 0.350165573 | 0.565555237 | 0.705318747 | tRNA-Phe-GAA |
| AP53_671 | 0.057394974 | 8.008832027 | 0.349752503 | 0.56578119 | 0.705318747 | 3-dehydroquinate dehy |
| AP53_541 | 0.069719475 | 8.691959385 | 0.347623347 | 0.566948759 | 0.705891805 | hydrolase, haloacid deh |
| AP53_1400 | -0.060972934 | 7.909196084 | 0.347587925 | 0.566968225 | 0.705891805 | NAD synthetase (EC 6.3. |
| AP53_134 | 0.060895951 | 8.862852606 | 0.343824083 | 0.569044365 | 0.708022508 | Nicotine adenine dinuc |
| AP53_361 | 0.105074353 | 12.21068719 | 0.34305315 | 0.569471523 | 0.708100081 | LSU ribosomal protein L |
| AP53_188 | -0.141704793 | 7.499966239 | 0.340890566 | 0.570673265 | 0.709140081 | Deoxyuridine 5'-tripho: |
| AP53_378 | 0.114782665 | 4.547407381 | 0.33972277 | 0.571324363 | 0.709178508 | Peptide ABC transporte |
| AP53_1087 | -0.117894494 | 4.395822584 | 0.339524742 | 0.571434923 | 0.709178508 | Outer surface protein o |
| AP53_970 | 0.082990705 | 6.591033471 | 0.337550933 | 0.572539313 | 0.710095084 | FIG01115860: hypothe |
| AP53_390 | 0.105939099 | 5.938200776 | 0.335603219 | 0.573633404 | 0.710997725 | Response regulator of tl |
| AP53_1273 | 0.100592139 | 6.981510129 | 0.334671399 | 0.574158358 | 0.711194239 | Two-component respor |
| AP53_1232 | -0.23231323 | 8.0946323 | 0.328022809 | 0.577932907 | 0.71541312 | FIG01114188: hypothe |
| AP53_375 | -0.232593964 | 2.887972822 | 0.327115282 | 0.57845212 | 0.715599468 | FIG01115781: hypothe |
| AP53_rna62 | 1.056296458 | 0.675720235 | 0.325192869 | 0.579555177 | 0.716507388 | tRNA-Glu-TTC |
| AP53_1355 | 0.203012936 | 4.089876018 | 0.323742295 | 0.580390404 | 0.717083242 | hypothetical protein, p |
| AP53_427 | 0.094122267 | 4.410117983 | 0.322354329 | 0.581191935 | 0.71761676 | Daunorubicin resistanc |
| AP53_650 | -0.127093905 | 4.173284571 | 0.317250181 | 0.584159553 | 0.720442502 | conserved domain prot |
| AP53_1332 | 0.098012514 | 4.927605505 | 0.317065353 | 0.584267611 | 0.720442502 | Phage minor capsid pro |

| | | | | | | |
|---|---|---|---|---|---|---|
| AP53_21 | 0.13859342 | 6.210596514 | 0.316507531 | 0.584593993 | 0.720442502 | Phosphoribosylaminoi |
| AP53_1120 | -0.056642898 | 8.887645674 | 0.313835702 | 0.586162627 | 0.721917296 | FIG01114335: hypothe |
| AP53_260 | -0.149949364 | 7.925695479 | 0.312023467 | 0.587231662 | 0.722775305 | Dicarboxylate/amino a |
| AP53_643 | -0.144778016 | 4.218836449 | 0.306936125 | 0.590254904 | 0.726035975 | replication initiator pro |
| AP53_1832 | -0.054576042 | 8.819021249 | 0.304522252 | 0.59170102 | 0.727147844 | Replicative DNA helicas |
| AP53_1804 | 0.069121167 | 6.605851342 | 0.30372372 | 0.592181078 | 0.727147844 | DNA mismatch repair p |
| AP53_584 | 0.166656481 | 10.15155159 | 0.30355484 | 0.592282711 | 0.727147844 | Large-conductance mec |
| AP53_201 | 0.100143359 | 6.70652531 | 0.301319763 | 0.593631329 | 0.728342863 | N-acetylmannosamine- |
| AP53_rna76 | -0.64389365 | 0.798034033 | 0.293261962 | 0.598548732 | 0.733912249 | tRNA-Asn-GTT |
| AP53_1255 | 0.08092361 | 8.539361175 | 0.289076966 | 0.601137736 | 0.736621431 | Hypothetical DUF1027 |
| AP53_1747 | -0.225539161 | 6.135558656 | 0.288214337 | 0.601674434 | 0.736813928 | Formiminotetrahydrof |
| AP53_909 | -0.092054654 | 5.601753535 | 0.28391895 | 0.604362593 | 0.739639214 | FIG01114846: hypothe |
| AP53_1792 | 0.203129398 | 4.794705572 | 0.283068589 | 0.604897905 | 0.739827872 | phage protein |
| AP53_791 | 0.054191391 | 8.265255128 | 0.276124731 | 0.609308718 | 0.744753288 | Tributyrin esterase |
| AP53_905 | 0.096458646 | 5.511271964 | 0.274839428 | 0.610133011 | 0.745291488 | Methylmalonyl-CoA de |
| AP53_1529 | -0.062627838 | 8.227547356 | 0.271735196 | 0.612134139 | 0.747265637 | Mannose-6-phosphate i |
| AP53_1280 | 0.078714887 | 9.75997479 | 0.268678859 | 0.614118835 | 0.748226143 | Valyl-tRNA synthetase ( |
| AP53_1707 | -0.105829535 | 11.49237008 | 0.268455902 | 0.614264184 | 0.748226143 | Peptidylproline cis-trar |
| AP53_882 | 0.092428233 | 8.88071783 | 0.268037181 | 0.614537363 | 0.748226143 | Pseudouridylate syntha |
| AP53_1244 | -0.070363026 | 8.999923432 | 0.267976335 | 0.614577083 | 0.748226143 | Cell division protein Fts |
| AP53_1474 | -0.105447165 | 6.336304883 | 0.267560983 | 0.614848376 | 0.748226143 | Putative regulator of th |
| AP53_688 | 0.076796989 | 9.040885876 | 0.265439761 | 0.616238112 | 0.748905194 | Aspartate carbamoyltra |
| AP53_556 | 0.072727869 | 9.196164744 | 0.26533736 | 0.616305381 | 0.748905194 | Substrate-specific comp |
| AP53_1088 | -0.096692575 | 4.336772951 | 0.264944085 | 0.616563885 | 0.748905194 | Beta-glucosidase (EC 3.2 |
| AP53_1205 | -0.154831346 | 2.924279 | 0.262604433 | 0.618106863 | 0.750309832 | FIG01114509: hypothe |
| AP53_536 | -0.057276883 | 8.741025202 | 0.258931412 | 0.620547016 | 0.752705843 | FIG146085: 3'-to-5' olig |
| AP53_465 | -0.104171544 | 6.15883924 | 0.258384489 | 0.620912247 | 0.752705843 | PTS system, beta-glucos |
| AP53_392 | 0.412456158 | 0.610601094 | 0.257888217 | 0.621244081 | 0.752705843 | FIG01116220: hypothe |
| AP53_639 | 0.101968947 | 4.770258326 | 0.255596004 | 0.622782076 | 0.754098571 | conserved hypothetical |
| AP53_393 | -0.102217366 | 3.336583387 | 0.254875934 | 0.623267028 | 0.754112261 | Competence-stimulatir |
| AP53_1833 | -0.073851876 | 8.861494486 | 0.254426111 | 0.623570417 | 0.754112261 | LSU ribosomal protein |
| AP53_1250 | 0.081953984 | 8.688805681 | 0.253437257 | 0.624238561 | 0.754450217 | Glucokinase (EC 2.7.1.2 |
| AP53_284 | -0.109686418 | 9.147368134 | 0.252016288 | 0.625201579 | 0.754848328 | FIG027054: hypothetic |
| AP53_380 | -0.124035593 | 5.255508454 | 0.251803986 | 0.625345756 | 0.754848328 | hypothetical protein |
| AP53_360 | 0.085408534 | 11.50774728 | 0.244379835 | 0.630436732 | 0.760520631 | LSU ribosomal protein |

| | | | | | | |
|---|---|---|---|---|---|---|
| AP53_217 | 0.066598721 | 8.032206253 | 0.243420884 | 0.631101401 | 0.760849577 | PurR: transcription regu |
| AP53_98 | 0.142417056 | 4.172683808 | 0.242005006 | 0.632085796 | 0.761563333 | FIG107367: tRNA-bindi |
| AP53_1157 | 0.100572959 | 3.708622363 | 0.239939045 | 0.633528682 | 0.762672011 | Late competence prote |
| AP53_408 | -0.066428813 | 9.562030378 | 0.239379012 | 0.633921158 | 0.762672011 | ATPase |
| AP53_554 | 0.150112694 | 2.74527286 | 0.239003328 | 0.634184763 | 0.762672011 | Asparagine-rich ORFan |
| AP53_1758 | 0.154761221 | 9.430769949 | 0.23811287 | 0.634810608 | 0.762951945 | PTS system, trehalose-s |
| AP53_rna64 | -0.24450873 | 4.164097717 | 0.235922033 | 0.636356675 | 0.764336823 | tRNA-Thr-TGT |
| AP53_1123 | -0.06457997 | 5.808376608 | 0.233539745 | 0.638048077 | 0.765447857 | FIG01115656: hypothe |
| AP53_709 | 0.089776998 | 5.106968656 | 0.233508415 | 0.638070393 | 0.765447857 | Peptidoglycan hydrolas |
| AP53_775 | -0.066964426 | 9.194484888 | 0.228168382 | 0.641901599 | 0.769568254 | Endonuclease III (EC 4.2 |
| AP53_722 | 0.10730334 | 7.447617739 | 0.226916738 | 0.642807645 | 0.76994969 | 5'-nucleotidase (EC 3.1. |
| AP53_925 | 0.068332929 | 5.40964687 | 0.226287456 | 0.643264344 | 0.76994969 | FIG01114068: hypothe |
| AP53_1190 | 0.136442335 | 2.186103157 | 0.226006866 | 0.643468236 | 0.76994969 | Phage minor tail protei |
| AP53_955 | -0.077528308 | 6.856014716 | 0.225315485 | 0.643971302 | 0.76994969 | Two component system |
| AP53_rna43 | -0.222918794 | 4.568139016 | 0.224997303 | 0.644203141 | 0.76994969 | tRNA-Pseudo-TGA |
| AP53_1207 | -0.166787006 | 3.359093794 | 0.223337095 | 0.645416136 | 0.770924751 | hypothetical protein |
| AP53_1619 | -0.101012063 | 6.430822598 | 0.220893177 | 0.64721193 | 0.772594315 | PTS system, lactose-spe |
| AP53_952 | -0.158479367 | 7.544188929 | 0.219341044 | 0.648358823 | 0.773487692 | Ribosomal RNA small su |
| AP53_590 | -0.055656151 | 7.752761326 | 0.218629196 | 0.648886493 | 0.773641697 | Alpha-L-Rha alpha-1,3-l |
| AP53_714 | 0.566124765 | -0.182630266 | 0.217602638 | 0.649649316 | 0.77364664 | Phage protein |
| AP53_1089 | 0.143116023 | 6.462081772 | 0.217550925 | 0.649687802 | 0.77364664 | Ribosyl nicotinamide tr |
| AP53_342 | -0.100861784 | 9.409220889 | 0.215174551 | 0.651462464 | 0.774747303 | FIG01114229: hypothe |
| AP53_278 | 0.048797831 | 9.52934866 | 0.215128436 | 0.651497021 | 0.774747303 | Transcription elongatic |
| AP53_776 | -0.055420268 | 7.944744549 | 0.213592238 | 0.652650819 | 0.774747303 | Putative tRNA-m1A22 r |
| AP53_1194 | -0.246951334 | 1.506943889 | 0.213396741 | 0.652798017 | 0.774747303 | FIG018217: Phage-asso |
| AP53_1440 | 0.161621867 | 6.64335366 | 0.213268038 | 0.652894968 | 0.774747303 | hypothetical protein |
| AP53_915 | 0.132845169 | 8.680433803 | 0.213119374 | 0.653007 | 0.774747303 | Citrate lyase alpha chai |
| AP53_rna82 | 0.268167095 | 2.664243267 | 0.211385862 | 0.654316919 | 0.775827208 | tRNA-Cys-GCA |
| AP53_505 | 0.071500094 | 9.691599483 | 0.210191688 | 0.655223124 | 0.775997578 | Calcium-transporting A |
| AP53_336 | 0.060454404 | 10.01169484 | 0.210018048 | 0.655355155 | 0.775997578 | C3 family ADP-ribosyltr |
| AP53_788 | -0.076061495 | 5.911765568 | 0.209617485 | 0.655659984 | 0.775997578 | ABC transporter perme |
| AP53_rna59 | -0.226373214 | 3.594400094 | 0.20794266 | 0.656938405 | 0.777036834 | tRNA-Gln-TTG |
| AP53_1624 | 0.099195767 | 6.419915055 | 0.204674667 | 0.659451146 | 0.779533906 | Lactose phosphotransfe |
| AP53_1033 | -0.169971063 | 5.62939968 | 0.202694119 | 0.660985898 | 0.780872567 | hypothetical protein |
| AP53_333 | -0.114774658 | 7.235476923 | 0.199244246 | 0.663681174 | 0.78357978 | Ribonucleotide reducta |

| | | | | | | |
|---|---|---|---|---|---|---|
| AP53_904 | -0.125758985 | 5.773598337 | 0.196530094 | 0.665821619 | 0.785186673 | Biotin carboxyl carrier p |
| AP53_32 | 0.121720732 | 5.687921236 | 0.196492686 | 0.665851243 | 0.785186673 | hypothetical protein |
| AP53_243 | 0.138722112 | 4.137115508 | 0.194008081 | 0.667826628 | 0.787037939 | Transposase |
| AP53_1515 | 0.041502071 | 7.717556279 | 0.192937693 | 0.66868235 | 0.787568229 | Substrate-specific comp |
| AP53_932 | 0.093944427 | 5.974249375 | 0.192056005 | 0.669389368 | 0.787575573 | Predicted transcriptior |
| AP53_453 | -0.332102491 | 1.390259302 | 0.191918137 | 0.6695001 | 0.787575573 | FIG01115104: hypothe |
| AP53_1051 | -0.148055734 | 8.48216266 | 0.19111686 | 0.670144616 | 0.787627327 | hypothetical protein |
| AP53_931 | 0.071252736 | 6.177726661 | 0.190854918 | 0.670355664 | 0.787627327 | Pyridoxal kinase (EC 2.7 |
| AP53_897 | 0.07277 | 8.720799367 | 0.190186066 | 0.670895356 | 0.787784565 | Ribonuclease HII (EC 3.1 |
| AP53_670 | 0.046536352 | 8.14152311 | 0.189299867 | 0.671612186 | 0.78780481 | LSU m5C1962 methyltr |
| AP53_1498 | 0.094579379 | 6.613320693 | 0.189161428 | 0.671724349 | 0.78780481 | hypothetical protein |
| AP53_466 | -0.080814953 | 6.612090354 | 0.187822475 | 0.672811718 | 0.788495368 | 6-phospho-beta-glucos |
| AP53_767 | -0.050449982 | 6.583723387 | 0.187293429 | 0.673242642 | 0.788495368 | tRNA dimethylallyltran |
| AP53_1199 | -0.15999298 | 2.890927429 | 0.185770563 | 0.674487154 | 0.788495368 | FIG01115109: hypothe |
| AP53_463 | 0.079621806 | 5.479056278 | 0.185481323 | 0.674724216 | 0.788495368 | Multidrug resistance pr |
| AP53_513 | 0.059823465 | 5.840755779 | 0.185340177 | 0.67483998 | 0.788495368 | 2-deoxy-D-gluconate 3- |
| AP53_1066 | 0.062688499 | 7.930079305 | 0.184987133 | 0.675129769 | 0.788495368 | Maltose/maltodextrin A |
| AP53_1062 | 0.054970114 | 7.882999782 | 0.184954252 | 0.675156776 | 0.788495368 | lipoprotein, putative |
| AP53_1799 | 0.105533542 | 4.830050309 | 0.182074171 | 0.677533551 | 0.790637467 | FIG01115461: hypothe |
| AP53_1056 | -0.054386453 | 7.167086984 | 0.181746189 | 0.677805639 | 0.790637467 | hypothetical protein |
| AP53_1420 | 0.118279691 | 7.217537269 | 0.178662986 | 0.680377854 | 0.79316121 | Transaldolase (EC 2.2.1. |
| AP53_368 | 0.095522933 | 9.824903804 | 0.177170643 | 0.681632373 | 0.79414672 | 67kDa Myosin-crossrea |
| AP53_1156 | -0.042143757 | 7.151800104 | 0.176111893 | 0.682526213 | 0.794711086 | DNA polymerase III delt |
| AP53_700 | -0.082752523 | 4.841515719 | 0.175510412 | 0.683035432 | 0.794827202 | Transcriptional regulat |
| AP53_950 | -0.087712591 | 10.0211407 | 0.174840055 | 0.683604185 | 0.795012416 | Nucleoside-binding pro |
| AP53_281 | -0.047178672 | 8.123607792 | 0.170993706 | 0.686892824 | 0.798358665 | FIG011178: rRNA meth |
| AP53_1352 | 0.182076613 | 4.312188956 | 0.167857366 | 0.68960695 | 0.801033567 | hypothetical phage pro |
| AP53_419 | -0.046679325 | 9.639050574 | 0.165633056 | 0.691550009 | 0.802810148 | Oxygen-insensitive NAD |
| AP53_1628 | 0.169888036 | 2.437120727 | 0.163471929 | 0.693452647 | 0.804504493 | Integrase |
| AP53_999 | 0.058615407 | 7.754957157 | 0.163035693 | 0.693838496 | 0.804504493 | Immunoreactive protei |
| AP53_1652 | -0.056303113 | 8.385416906 | 0.162355199 | 0.694441603 | 0.804723076 | FIG01114677: hypothe |
| AP53_395 | 0.100630251 | 4.13609729 | 0.160328294 | 0.696246845 | 0.806333607 | Bacteriocin-like peptid |
| AP53_588 | 0.052924138 | 8.92811689 | 0.158725539 | 0.697683814 | 0.80751597 | dTDP-4-dehydrorhamn |
| AP53_160 | 0.119679197 | 4.973827245 | 0.158043102 | 0.698298238 | 0.807745459 | Mobile element protein |
| AP53_902 | -0.071525339 | 4.822374628 | 0.155764672 | 0.700360902 | 0.808923081 | Oxaloacetate decarboxy |

| | | | | | | |
|---|---|---|---|---|---|---|
| AP53_910 | -0.099141891 | 7.052291359 | 0.15576071 | 0.700364504 | 0.808923081 | Biotin carboxyl carrier |
| AP53_1656 | -0.051247243 | 9.178346442 | 0.155538557 | 0.700566563 | 0.808923081 | Intramembrane proteas |
| AP53_1195 | -0.163810885 | 3.18282213 | 0.153376576 | 0.702541834 | 0.810261916 | FIG033577: Phage prot |
| AP53_1340 | 0.081492726 | 4.087828755 | 0.15335574 | 0.70256095 | 0.810261916 | conserved hypothetical |
| AP53_1755 | 0.055093567 | 11.0368012 | 0.152558084 | 0.703293872 | 0.810365451 | Translation elongation |
| AP53_1 | 0.056615663 | 7.944552078 | 0.15234969 | 0.703485721 | 0.810365451 | Chromosomal replicati |
| AP53_1201 | -0.105036009 | 3.887389286 | 0.151142251 | 0.704600314 | 0.810747392 | conserved hypothetical |
| AP53_1587 | 0.069519625 | 5.648149901 | 0.15108566 | 0.704652679 | 0.810747392 | FIG01114549: hypothe |
| AP53_655 | -0.048078082 | 9.319195173 | 0.14903007 | 0.706562585 | 0.812442797 | Phage integrase: site-sp |
| AP53_798 | -0.049355138 | 6.426757126 | 0.148600796 | 0.706963362 | 0.812442797 | Hyaluronate lyase precu |
| AP53_763 | -0.047142992 | 7.618129502 | 0.147809462 | 0.707703926 | 0.812812615 | hypothetical protein |
| AP53_1743 | -0.126194704 | 10.51698796 | 0.145989159 | 0.709416205 | 0.81429737 | Alkyl hydroperoxide re |
| AP53_875 | 0.106729757 | 7.024084202 | 0.145100729 | 0.710256398 | 0.814779946 | 4-oxalocrotonate taut |
| AP53_1356 | 0.10645055 | 3.936521777 | 0.144247763 | 0.711065855 | 0.815226712 | phage protein |
| AP53_296 | -0.070762993 | 7.707430947 | 0.143774562 | 0.711516111 | 0.815261377 | Membrane-associated |
| AP53_1353 | 0.111450072 | 4.458065608 | 0.141813952 | 0.713390822 | 0.81692719 | DNA replication protei |
| AP53_851 | 0.130215038 | 6.138214621 | 0.139993526 | 0.715144908 | 0.818452988 | Class B acid phosphatas |
| AP53_87 | -0.112592821 | 3.194816257 | 0.138607238 | 0.716489498 | 0.819508613 | Late competence prote |
| AP53_269 | -0.043640418 | 8.07559697 | 0.135138145 | 0.719888445 | 0.822907109 | Ribonucleotide reduct |
| AP53_385 | 0.072530691 | 4.362309074 | 0.13471272 | 0.720308696 | 0.822907109 | Mobile element protei |
| AP53_1824 | 0.079873684 | 3.817303732 | 0.132853298 | 0.722154471 | 0.824530487 | FIG014387: Transcripti |
| AP53_781 | -0.040757327 | 9.416215669 | 0.131505639 | 0.723501485 | 0.825582824 | dTDP-glucose 4,6-dehy |
| AP53_1634 | -0.098240814 | 8.437611102 | 0.131032482 | 0.723976283 | 0.825639228 | DegV family protein in |
| AP53_1811 | -0.03993717 | 7.932330972 | 0.130580762 | 0.724430485 | 0.825672091 | Integral membrane pro |
| AP53_860 | 0.051292769 | 8.307023261 | 0.125853687 | 0.729238187 | 0.83066392 | Cysteine desulfurase (E |
| AP53_833 | 0.071077037 | 6.204504065 | 0.124283875 | 0.730857418 | 0.831598552 | D-alanyl-D-alanine carb |
| AP53_881 | -0.052344413 | 9.178826799 | 0.124227702 | 0.730915574 | 0.831598552 | Serine hydroxymethyltr |
| AP53_815 | -0.055403216 | 8.798029072 | 0.121762647 | 0.733482459 | 0.833554579 | Translation elongation |
| AP53_1657 | -0.045876888 | 8.21286656 | 0.121741406 | 0.733504705 | 0.833554579 | Phosphatidate cytidyly |
| AP53_pseudo399 | 0.073484502 | 4.327962628 | 0.121342278 | 0.733923119 | 0.833554579 | FIG01114499: hypothe |
| AP53_210 | -0.036139871 | 8.282924267 | 0.115327591 | 0.740324914 | 0.840333718 | Ribonuclease M5 (EC 3. |
| AP53_1345 | -0.085065408 | 4.364938321 | 0.10904267 | 0.747218152 | 0.847662438 | FIG01114622: hypothe |
| AP53_741 | -0.062045968 | 9.877237239 | 0.107387025 | 0.749070833 | 0.849267808 | Arsenate reductase (EC |
| AP53_1748 | 0.130452429 | 6.709134603 | 0.1063636 | 0.750224036 | 0.85001971 | Formate--tetrahydrofol |
| AP53_1745 | -0.122845926 | 5.866330514 | 0.106022394 | 0.750609883 | 0.85001971 | Urocanate hydratase (E |

| | | | | | | |
|---|---|---|---|---|---|---|
| AP53_1115 | 0.08795965 | 5.058997186 | 0.105170053 | 0.751576772 | 0.850195873 | Protein G-related alpha |
| AP53_459 | 0.077432642 | 4.324628536 | 0.105113145 | 0.751641483 | 0.850195873 | Mobile element proteir |
| AP53_1458 | 0.038982626 | 9.111977327 | 0.10368725 | 0.753269301 | 0.851540893 | FIG000325: clustered v |
| AP53_190 | -0.036294998 | 8.865796182 | 0.103074359 | 0.753972807 | 0.851840058 | Carbonic anhydrase (EC |
| AP53_475 | -0.036118897 | 9.041570762 | 0.10249419 | 0.754640899 | 0.852098886 | General stress protein |
| AP53_1407 | -0.055142559 | 6.653605219 | 0.101737874 | 0.755514993 | 0.852589884 | FIG01116065: hypothe |
| AP53_629 | -0.102498448 | 3.905552694 | 0.099985469 | 0.757554241 | 0.854394412 | putative transcriptiona |
| AP53_117 | -0.038197396 | 7.004833486 | 0.099386842 | 0.758255388 | 0.854688564 | Endoribonuclease L-PSI |
| AP53_1098 | -0.074929212 | 4.219661802 | 0.098059617 | 0.759818292 | 0.855953166 | Mobile element proteir |
| AP53_674 | 0.033008536 | 8.27484944 | 0.096444151 | 0.761736487 | 0.857616311 | Glutathione reductase ( |
| AP53_1434 | 0.027403216 | 7.360167075 | 0.095789181 | 0.762519245 | 0.857999916 | Sodium-dependent pho |
| AP53_810 | -0.047443203 | 7.328800618 | 0.094310879 | 0.764296893 | 0.859501894 | Translation elongation |
| AP53_520 | 0.044725637 | 8.789098109 | 0.092849969 | 0.766068772 | 0.860995649 | Cell division transporte |
| AP53_1354 | -0.110789006 | 3.37901892 | 0.092313562 | 0.766723205 | 0.861232489 | putative phage replicat |
| AP53_165 | -0.09312054 | 2.954387678 | 0.091027629 | 0.768300646 | 0.862111326 | Mobile element proteir |
| AP53_647 | -0.106948622 | 3.469571357 | 0.090467815 | 0.768991188 | 0.862111326 | hypothetical phage pro |
| AP53_969 | 0.048108687 | 8.451923842 | 0.090460389 | 0.769000363 | 0.862111326 | tRNA pseudouridine syı |
| AP53_1166 | -0.03444696 | 8.162139025 | 0.090232434 | 0.769282235 | 0.862111326 | UDP-N-acetylmuramoy |
| AP53_1347 | -0.09425601 | 4.277213214 | 0.089698664 | 0.769943785 | 0.86235481 | Single-stranded DNA-bi |
| AP53_173 | -0.05483377 | 6.451482905 | 0.088542007 | 0.771384782 | 0.863238187 | S-layer homology doma |
| AP53_529 | -0.035535986 | 9.140066361 | 0.088269259 | 0.771726079 | 0.863238187 | FIG002813: LPPG:FO 2- |
| AP53_983 | 0.043784109 | 8.231438178 | 0.087997501 | 0.772066714 | 0.863238187 | ATP-dependent DNA he |
| AP53_rna42 | -0.20610561 | 3.837079763 | 0.087140049 | 0.773145275 | 0.863484499 | tRNA-Glu-TTC |
| AP53_633 | -0.064099182 | 4.286017489 | 0.086893458 | 0.773456528 | 0.863484499 | Single-stranded DNA-bi |
| AP53_1285 | -0.06224287 | 8.215176469 | 0.086762827 | 0.77362161 | 0.863484499 | 2-keto-3-deoxy-D-arabi |
| AP53_1749 | 0.100664466 | 6.229693034 | 0.084530459 | 0.776463969 | 0.866158945 | Conserved protein |
| AP53_164 | -0.094240044 | 4.340799239 | 0.083777614 | 0.777431752 | 0.866740396 | Mobile element proteir |
| AP53_476 | 0.05383086 | 9.659574946 | 0.08286815 | 0.778607213 | 0.867552584 | FIG01115301: hypothe |
| AP53_996 | 0.047069384 | 8.831870744 | 0.082150009 | 0.779540369 | 0.868094008 | Transcriptional regulat |
| AP53_1551 | 0.033857321 | 7.86464506 | 0.08152411 | 0.780357293 | 0.868505451 | Recombination inhibit |
| AP53_1283 | -0.035529505 | 8.006384195 | 0.080649261 | 0.781504885 | 0.868969675 | FIG01114159: hypothe |
| AP53_884 | -0.049671838 | 8.408772709 | 0.080523974 | 0.781669785 | 0.868969675 | ABC transporter, ATP-bi |
| AP53_679 | 0.038665661 | 7.309242498 | 0.079687317 | 0.782774564 | 0.869235784 | Capsule biosynthesis pr |
| AP53_224 | 0.039420393 | 8.276251153 | 0.079538276 | 0.782972028 | 0.869235784 | Amino acid transport A |
| AP53_488 | 0.088663208 | 7.678716534 | 0.079326732 | 0.783252647 | 0.869235784 | Phage lysin, glycosyl hy |

| | | | | | | |
|---|---|---|---|---|---|---|
| AP53_1732 | 0.037346574 | 8.474699799 | 0.077220474 | 0.786069095 | 0.871862922 | Dipeptidase (EC 3.4.-.-) |
| AP53_1218 | 0.030449993 | 8.972107627 | 0.076872471 | 0.786538429 | 0.87188526 | FIG006988: Lipase/Acy |
| AP53_1845 | 0.029052956 | 8.383367852 | 0.074202807 | 0.790177765 | 0.87514071 | Transcriptional regulat |
| AP53_1220 | 0.027278571 | 8.806395054 | 0.074058163 | 0.790376952 | 0.87514071 | FIG01115348: hypothe |
| AP53_1211 | 0.069777042 | 1.877716067 | 0.071294224 | 0.794224233 | 0.878899223 | Phage excisionase |
| AP53_912 | 0.061760708 | 6.658422814 | 0.070495748 | 0.795350586 | 0.879644153 | FIG01114213: hypothe |
| AP53_642 | 0.064617285 | 5.138889041 | 0.068288186 | 0.798500831 | 0.882392772 | Helicase loader DnaI |
| AP53_1002 | 0.083824853 | 9.186568837 | 0.068118637 | 0.798745029 | 0.882392772 | Streptococcal phospho |
| AP53_1693 | -0.052497546 | 13.096781 | 0.067573201 | 0.799532832 | 0.882640513 | Antiphagocytic M prote |
| AP53_pseudo1615 | 0.071186617 | 3.079259341 | 0.067334445 | 0.799878754 | 0.882640513 | Lanthionine biosynthes |
| AP53_1576 | 0.034195622 | 8.093332469 | 0.065491443 | 0.802571412 | 0.884786941 | Nucleoside permease N |
| AP53_1566 | 0.051143282 | 7.086791417 | 0.065379949 | 0.802735602 | 0.884786941 | Transcriptional regulat |
| AP53_202 | -0.044437642 | 7.286735702 | 0.064674432 | 0.803778058 | 0.885433151 | ABC transporter, predic |
| AP53_863 | -0.056908125 | 7.861865308 | 0.063977633 | 0.804813622 | 0.886071039 | GTP pyrophosphokinas |
| AP53_1384 | -0.029495985 | 10.00184868 | 0.062947417 | 0.806355809 | 0.886192473 | Hydrolase (HAD superfa |
| AP53_1691 | 0.06142062 | 10.28433427 | 0.062732662 | 0.806678983 | 0.886192473 | C5a peptidase (EC 3.4.2 |
| AP53_1081 | -0.090389541 | 4.303329124 | 0.062627177 | 0.806837938 | 0.886192473 | FIG01114324: hypothe |
| AP53_261 | -0.02744767 | 8.013127346 | 0.062064729 | 0.807687912 | 0.886192473 | Trk system potassium u |
| AP53_144 | 0.046309809 | 5.193900191 | 0.061814704 | 0.80806707 | 0.886192473 | 3-keto-L-gulonate 6-ph |
| AP53_1344 | 0.100562505 | 4.348525858 | 0.061621534 | 0.80836057 | 0.886192473 | FIG01115146: hypothe |
| AP53_480 | 0.056530585 | 5.721513167 | 0.061602156 | 0.80839004 | 0.886192473 | FIG01114033: hypothe |
| AP53_1258 | 0.026824485 | 8.322350417 | 0.061321516 | 0.808817386 | 0.886192473 | Lon-like protease with I |
| AP53_1445 | 0.028693968 | 8.853964934 | 0.061180178 | 0.809033005 | 0.886192473 | PTS system, galactose-s |
| AP53_787 | -0.032712695 | 5.790644875 | 0.060550911 | 0.809996224 | 0.886747135 | ABC transporter substra |
| AP53_396 | -0.123220157 | 2.364449789 | 0.058361058 | 0.813390455 | 0.889961033 | FIG01116077: hypothe |
| AP53_444 | -0.06425052 | 3.147686211 | 0.056367533 | 0.816539756 | 0.892903474 | FIG01116163: hypothe |
| AP53_327 | -0.030056047 | 8.892621413 | 0.05440803 | 0.819693437 | 0.895847389 | FIG01118451: hypothe |
| AP53_1418 | -0.046637866 | 7.937633942 | 0.05364129 | 0.8209438 | 0.896403803 | FIG01113961: hypothe |
| AP53_1093 | 0.088178963 | 2.754249981 | 0.053529919 | 0.821126204 | 0.896403803 | Mobile element protein |
| AP53_1094 | -0.086899649 | 4.265203367 | 0.051900257 | 0.823818561 | 0.898837452 | Mobile element protein |
| AP53_1270 | -0.049907896 | 8.361294977 | 0.05054447 | 0.826092673 | 0.900812291 | Putative uncharacterize |
| AP53_1740 | -0.046174798 | 6.04185596 | 0.04903805 | 0.828657505 | 0.902290936 | hypothetical protein |
| AP53_605 | -0.079017431 | 6.486805845 | 0.048938341 | 0.828828725 | 0.902290936 | FIG01116522: hypothe |
| AP53_1575 | -0.030126978 | 8.329307005 | 0.04892989 | 0.828843246 | 0.902290936 | Deoxyribose-phosphate |
| AP53_1350 | 0.045798663 | 3.748987503 | 0.047099351 | 0.832020172 | 0.904876701 | FIG01116817: hypothe |

| | | | | | | |
|---|---|---|---|---|---|---|
| AP53_1198 | -0.048432024 | 3.798169381 | 0.046628054 | 0.832848578 | 0.904876701 | FIG040428: Phage-asso |
| AP53_1349 | -0.070332592 | 3.309771167 | 0.046570606 | 0.832949854 | 0.904876701 | hypothetical protein (p |
| AP53_1437 | -0.032153681 | 7.885640041 | 0.0460876 | 0.833803974 | 0.904876701 | Transcriptional regulat |
| AP53_1346 | -0.060442503 | 4.250518521 | 0.045877173 | 0.834177549 | 0.904876701 | Phage Holliday junctior |
| AP53_34 | 0.076685797 | 5.755964412 | 0.045781889 | 0.834347004 | 0.904876701 | Transcriptional regulat |
| AP53_801 | 0.018668875 | 7.615810176 | 0.04553951 | 0.834778894 | 0.904876701 | proposed amino acid li |
| AP53_777 | -0.019748938 | 8.043231212 | 0.045186049 | 0.835410884 | 0.904876701 | FIG137478: Hypothetic |
| AP53_896 | 0.041476129 | 9.148982694 | 0.044950065 | 0.83583427 | 0.904876701 | 50S ribosomal subunit |
| AP53_175 | -0.07446261 | 5.691972231 | 0.044685691 | 0.836309977 | 0.904876701 | hypothetical protein |
| AP53_244 | 0.052763317 | 3.470264022 | 0.044665357 | 0.836346626 | 0.904876701 | Competence-specific si |
| AP53_1468 | -0.062389764 | 8.166070866 | 0.044243566 | 0.837108833 | 0.905196794 | ATPase YjeE, predicted t |
| AP53_1111 | -0.025044433 | 8.445947905 | 0.04323175 | 0.838952922 | 0.90644906 | Inner membrane protei |
| AP53_1596 | -0.030465878 | 7.534487352 | 0.04309662 | 0.839200908 | 0.90644906 | hypothetical protein |
| AP53_384 | 0.066724986 | 5.312484241 | 0.042810819 | 0.839726741 | 0.906512572 | Mobile element proteir |
| AP53_493 | 0.01868306 | 8.217555346 | 0.04150426 | 0.842154341 | 0.908385187 | Phosphoenolpyruvate c |
| AP53_1192 | -0.081711438 | 2.102695241 | 0.041349622 | 0.842444289 | 0.908385187 | conserved hypothetical |
| AP53_901 | 0.023143373 | 7.690586902 | 0.041125573 | 0.84286539 | 0.908385187 | tRNA:m(5)U-54 MTase g |
| AP53_1065 | 0.038454999 | 5.632716854 | 0.040173228 | 0.84466883 | 0.90982364 | Maltose operon transcr |
| AP53_1555 | 0.023091565 | 8.924918181 | 0.038814706 | 0.847280493 | 0.912130581 | Putative signal peptida: |
| AP53_1209 | 0.073491221 | 2.866562843 | 0.03823897 | 0.848401663 | 0.912831279 | DNA replication proteir |
| AP53_1577 | -0.044279953 | 7.414925905 | 0.037160677 | 0.850525396 | 0.914609304 | Uridine phosphorylase |
| AP53_398 | -0.075210219 | 2.694450071 | 0.035960831 | 0.852926563 | 0.916683532 | unknown |
| AP53_300 | 0.062787729 | 8.145165313 | 0.035449312 | 0.853962875 | 0.917289397 | Pyruvate formate-lyase |
| AP53_522 | -0.022532852 | 7.975341605 | 0.034016564 | 0.856907586 | 0.919684497 | Hydroxyacylglutathion |
| AP53_211 | 0.023012219 | 8.865840499 | 0.033904661 | 0.857140265 | 0.919684497 | SSU rRNA (adenine(151 |
| AP53_328 | 0.044609566 | 8.781683546 | 0.033603002 | 0.85776949 | 0.919851149 | FIG01115281: hypothe |
| AP53_209 | -0.020989765 | 7.368054621 | 0.032759904 | 0.85954378 | 0.92124488 | Putative deoxyribonucl |
| AP53_1348 | -0.041413724 | 4.605248262 | 0.032478015 | 0.860142277 | 0.921377572 | Single-stranded DNA-bi |
| AP53_1419 | 0.027771275 | 9.811227587 | 0.032200727 | 0.860733634 | 0.92149083 | Transketolase (EC 2.2.1. |
| AP53_1621 | 0.037102315 | 6.750819631 | 0.031984076 | 0.861197509 | 0.92149083 | Tagatose-6-phosphate k |
| AP53_403 | 0.048780632 | 4.275269101 | 0.031569134 | 0.862090511 | 0.921653471 | hypothetical protein |
| AP53_416 | -0.020707535 | 8.589937643 | 0.031472579 | 0.862299177 | 0.921653471 | hypothetical protein |
| AP53_377 | -0.025575222 | 4.518264802 | 0.030811657 | 0.863736479 | 0.922226465 | Peptide ABC transporte |
| AP53_1540 | -0.017307925 | 9.211085047 | 0.030733344 | 0.863907836 | 0.922226465 | Excinuclease ABC subur |
| AP53_145 | 0.030604117 | 5.127435458 | 0.030572424 | 0.86426066 | 0.922226465 | L-xylulose 5-phosphate |

| | | | | | | |
|---|---|---|---|---|---|---|
| AP53_1768 | 0.079007561 | 7.359813173 | 0.029170079 | 0.867376846 | 0.925043108 | FIG01115181: hypothe |
| AP53_1655 | -0.02792655 | 9.654728026 | 0.028199695 | 0.869578643 | 0.926462767 | Prolyl-tRNA synthetase |
| AP53_1312 | 0.045650825 | 6.203251073 | 0.028163016 | 0.869662628 | 0.926462767 | Phage lysin |
| AP53_569 | 0.021406662 | 8.07311659 | 0.027933922 | 0.870188468 | 0.926514436 | Competence-associatec |
| AP53_1606 | 0.034492639 | 4.367122512 | 0.027158451 | 0.871985097 | 0.927820415 | Competence-specific si |
| AP53_1581 | -0.040333876 | 6.347730772 | 0.026993319 | 0.872371075 | 0.927820415 | Ribosomal-protein-S18 |
| AP53_442 | 0.030668839 | 3.886485751 | 0.026743049 | 0.87295838 | 0.927936592 | FIG01115625: hypothe |
| AP53_867 | 0.023801395 | 6.75101341 | 0.025817328 | 0.875155703 | 0.92976312 | Short chain dehydroger |
| AP53_985 | -0.020119816 | 8.1689722 | 0.025015426 | 0.877092092 | 0.930735956 | Cobalt-zinc-cadmium r |
| AP53_859 | 0.028912014 | 7.980890273 | 0.024929923 | 0.87730043 | 0.930735956 | FIG01114333: hypothe |
| AP53_189 | 0.014101877 | 8.5345132 | 0.024844089 | 0.877509943 | 0.930735956 | DNA repair protein Rad |
| AP53_1351 | -0.045537593 | 3.956639772 | 0.024279779 | 0.87889673 | 0.931314726 | hypothetical phage pro |
| AP53_1554 | 0.026828653 | 8.751635964 | 0.024168589 | 0.879171922 | 0.931314726 | Ribonuclease HIII (EC 3. |
| AP53_1140 | 0.044691732 | 6.040327791 | 0.024038365 | 0.87949505 | 0.931314726 | Transcriptional regulat |
| AP53_1200 | -0.032925678 | 3.627071698 | 0.023018837 | 0.882056505 | 0.933517816 | hypothetical protein |
| AP53_675 | 0.033395051 | 8.237706878 | 0.022379442 | 0.88369268 | 0.934739778 | Dihydrofolate synthase |
| AP53_407 | -0.019160068 | 7.994504104 | 0.021512423 | 0.885950065 | 0.936617143 | Dephospho-CoA kinase |
| AP53_707 | -0.021650788 | 8.129301303 | 0.021196501 | 0.886784151 | 0.936983247 | 1-phosphofructokinase |
| AP53_1635 | -0.022596862 | 7.260475475 | 0.02101665 | 0.887261828 | 0.936983247 | Hypothetical protein D |
| AP53_1853 | -0.016697205 | 7.660663908 | 0.02069661 | 0.888117041 | 0.937376388 | DNA recombination an |
| AP53_391 | 0.027371983 | 5.127292272 | 0.019932387 | 0.890186959 | 0.939050483 | Histidine kinase of the c |
| AP53_717 | -0.03694823 | 8.491106844 | 0.019730446 | 0.890740668 | 0.939124191 | DegV family protein |
| AP53_1090 | 0.049048525 | 7.046293108 | 0.019333602 | 0.89183728 | 0.939769903 | FIG01114098: hypothe |
| AP53_604 | -0.050526989 | 6.321387552 | 0.018554283 | 0.894024756 | 0.94109715 | Phage protein |
| AP53_756 | 0.018261418 | 8.863382525 | 0.018539558 | 0.894066535 | 0.94109715 | Topoisomerase IV subu |
| AP53_1579 | 0.022352452 | 8.787446596 | 0.017827355 | 0.896107832 | 0.942302803 | SSU ribosomal protein |
| AP53_rna77 | 0.290965373 | -1.139188048 | 0.017801441 | 0.896182882 | 0.942302803 | 5S RNA |
| AP53_1159 | -0.016896921 | 8.174832113 | 0.016986826 | 0.898571168 | 0.943329501 | 1-acyl-sn-glycerol-3-ph |
| AP53_1801 | -0.027925088 | 3.605031207 | 0.016911407 | 0.898795206 | 0.943329501 | FIG01116340: hypothe |
| AP53_1341 | -0.050724568 | 4.226506109 | 0.016851308 | 0.898974096 | 0.943329501 | Phage protein |
| AP53_638 | -0.029422301 | 3.705966894 | 0.016653248 | 0.899565956 | 0.943329501 | FIG01116946: hypothe |
| AP53_686 | 0.01689988 | 8.354929584 | 0.016645448 | 0.899589339 | 0.943329501 | Uracil phosphoribosylt |
| AP53_89 | -0.027268277 | 3.062870992 | 0.015675996 | 0.902540153 | 0.944828688 | Late competence prote |
| AP53_1771 | 0.111680845 | 2.941398101 | 0.015621624 | 0.902708362 | 0.944828688 | hypothetical protein |
| AP53_1834 | 0.023010276 | 7.62930424 | 0.015582428 | 0.902829808 | 0.944828688 | Phosphoesterase, DHH |

| | | | | | | |
|---|---|---|---|---|---|---|
| AP53_1141 | -0.037480155 | 5.045558232 | 0.015538497 | 0.902966108 | 0.944828688 | FIG01114322: hypothe |
| AP53_1451 | 0.017898171 | 6.309742493 | 0.014532512 | 0.906142658 | 0.947641648 | Negative transcriptiona |
| AP53_677 | -0.015678288 | 8.111374097 | 0.014360622 | 0.906696479 | 0.947710213 | Cysteine desulfurase (EC |
| AP53_458 | -0.019207763 | 3.931775774 | 0.013669785 | 0.908957017 | 0.949561663 | Mobile element proteir |
| AP53_1342 | 0.030973548 | 4.625304636 | 0.012404546 | 0.913252809 | 0.953084784 | Phage protein |
| AP53_1212 | -0.030201219 | 4.201745088 | 0.012387679 | 0.91331154 | 0.953084784 | Cro-like phage transcrip |
| AP53_771 | -0.017775412 | 9.030765767 | 0.011810878 | 0.915344981 | 0.954693502 | Short chain dehydroger |
| AP53_1221 | 0.01046014 | 8.478727955 | 0.011613593 | 0.91605199 | 0.954917783 | DNA repair protein Recl |
| AP53_826 | 0.037406233 | 8.443236576 | 0.011089322 | 0.917960921 | 0.956394067 | Succinate-semialdehyd |
| AP53_1278 | 0.01635966 | 9.295328339 | 0.010875507 | 0.918752532 | 0.956705292 | Oxidoreductase, Gfo/Id |
| AP53_1067 | -0.015930643 | 5.709482512 | 0.010535176 | 0.920028966 | 0.957520763 | Maltose/maltodextrin / |
| AP53_716 | -0.033583092 | 9.254855066 | 0.010033413 | 0.921949519 | 0.959005367 | Hypothetical protein SI |
| AP53_1516 | -0.008994222 | 6.984875302 | 0.009528478 | 0.923931882 | 0.959740678 | Putative ABC transport |
| AP53_1760 | 0.027939831 | 4.176515525 | 0.009480249 | 0.924123976 | 0.959740678 | FIG01115695: hypothe |
| AP53_146 | 0.018176261 | 6.035456131 | 0.009274322 | 0.924949781 | 0.959740678 | L-ribulose-5-phosphate |
| AP53_1469 | 0.034681203 | 8.997062334 | 0.009181969 | 0.925323141 | 0.959740678 | Guanine-hypoxanthine |
| AP53_96 | -0.022553009 | 6.056549802 | 0.009131848 | 0.92552656 | 0.959740678 | FIG009688: Thioredoxi |
| AP53_485 | -0.014398273 | 6.997365124 | 0.009049266 | 0.925862958 | 0.959740678 | Phosphoglycerate muta |
| AP53_1744 | 0.034932455 | 5.749783198 | 0.008987003 | 0.926117614 | 0.959740678 | Imidazolonepropionase |
| AP53_486 | 0.022595953 | 8.007340446 | 0.008416531 | 0.928493608 | 0.961689484 | Transcriptional regulat |
| AP53_589 | -0.013659577 | 6.987027995 | 0.00817092 | 0.92954155 | 0.962261413 | Alpha-D-GlcNAc alpha-1 |
| AP53_521 | 0.011539718 | 8.920122241 | 0.008012574 | 0.930225607 | 0.962456238 | Cell division protein Fts |
| AP53_1397 | 0.01257512 | 8.742721169 | 0.00788692 | 0.930773298 | 0.962509841 | RecU Holliday junction |
| AP53_866 | 0.023935596 | 10.61348664 | 0.007716936 | 0.931521265 | 0.962770381 | Phosphate acetyltransf |
| AP53_715 | -0.031359669 | 3.336580757 | 0.007250637 | 0.933616827 | 0.964422705 | Nucleoside diphosphat |
| AP53_1552 | 0.007954659 | 7.991986028 | 0.00690653 | 0.935207171 | 0.965056972 | Colicin V production pr |
| AP53_757 | -0.014543379 | 8.988796479 | 0.006812719 | 0.935647622 | 0.965056972 | Topoisomerase IV subu |
| AP53_1386 | 0.020835861 | 7.247876238 | 0.006770017 | 0.935849123 | 0.965056972 | 3-ketoacyl-CoA thiolase |
| AP53_rna29 | -0.034839591 | 4.81700164 | 0.006560206 | 0.936848603 | 0.965056972 | tRNA-Pseudo-TGA |
| AP53_1831 | 0.023375233 | 5.756421826 | 0.006525518 | 0.937015392 | 0.965056972 | FIG01114855: hypothe |
| AP53_885 | -0.01114512 | 7.980501176 | 0.00648433 | 0.937214009 | 0.965056972 | ABC transporter, ATP-bi |
| AP53_579 | -0.011191683 | 7.606143427 | 0.006317772 | 0.938023759 | 0.965312674 | YpfJ protein, zinc metal |
| AP53_911 | -0.01630278 | 7.402583754 | 0.006229556 | 0.938456989 | 0.965312674 | Oxaloacetate decarbox |
| AP53_464 | -0.018651555 | 5.259576695 | 0.006021142 | 0.939492945 | 0.965393028 | Beta-glucoside bgl oper |
| AP53_849 | 0.025841482 | 6.509506297 | 0.006013784 | 0.939529845 | 0.965393028 | Alcohol dehydrogenase |

| | | | | | | |
|---|---|---|---|---|---|---|
| AP53_435 | -0.012556996 | 7.890446052 | 0.005725235 | 0.94099531 | 0.966387246 | Zn-dependent hydrolas |
| AP53_669 | -0.010824692 | 8.124166179 | 0.005515638 | 0.942083245 | 0.966992903 | Lipoteichoic acid synth |
| AP53_443 | 0.017534107 | 4.271592966 | 0.005391202 | 0.942738998 | 0.967154543 | Archaeal S-adenosylme |
| AP53_410 | 0.013975392 | 9.975201927 | 0.005174475 | 0.943899553 | 0.967802185 | Preprotein translocase |
| AP53_452 | 0.032670462 | 1.143102125 | 0.005088351 | 0.944367511 | 0.967802185 | FIG01116748: hypothe |
| AP53_706 | 0.01674023 | 8.158050813 | 0.004760365 | 0.946187164 | 0.969155296 | Transcriptional repress |
| AP53_228 | 0.010225132 | 8.054936901 | 0.00466987 | 0.94670024 | 0.969169391 | Negative regulator of ge |
| AP53_88 | -0.016080445 | 3.287412157 | 0.004063609 | 0.950274801 | 0.972091248 | Late competence prote |
| AP53_1608 | -0.007497498 | 7.521305155 | 0.003880951 | 0.951403612 | 0.972091248 | Type I restriction-modi |
| AP53_473 | 0.012517188 | 9.506031141 | 0.003845576 | 0.951625285 | 0.972091248 | HPr kinase/phosphoryla |
| AP53_1754 | -0.008086158 | 11.52845548 | 0.003835423 | 0.9516891 | 0.972091248 | SSU ribosomal protein S |
| AP53_537 | 0.01263479 | 8.498753398 | 0.003705339 | 0.952514314 | 0.972091248 | Flavodoxin |
| AP53_1222 | -0.009383402 | 7.686486073 | 0.003698316 | 0.952559275 | 0.972091248 | Arginine pathway regul |
| AP53_1372 | 0.011193731 | 9.148815649 | 0.003312154 | 0.955101181 | 0.974132484 | General stress protein 1 |
| AP53_761 | 0.01019933 | 8.406277453 | 0.003233739 | 0.955635224 | 0.974132484 | FIG015049: hypothetic |
| AP53_555 | 0.008859947 | 8.73793431 | 0.003171302 | 0.956065112 | 0.974132484 | Endonuclease/Exonucle |
| AP53_945 | 0.006031306 | 7.848707534 | 0.002554914 | 0.960560866 | 0.978199706 | Substrate-specific comp |
| AP53_883 | 0.004723272 | 8.535576241 | 0.002301346 | 0.96256739 | 0.978825287 | Pneumococcal vaccine |
| AP53_322 | 0.011687458 | 3.828451019 | 0.002289525 | 0.962663571 | 0.978825287 | FIG01115198: hypothe |
| AP53_131 | -0.005244717 | 10.18789975 | 0.002276168 | 0.962772544 | 0.978825287 | Adenylosuccinate synth |
| AP53_223 | -0.012835229 | 4.257604621 | 0.002225084 | 0.963192322 | 0.978825287 | FIG01114598: hypothe |
| AP53_914 | -0.013284798 | 8.560464416 | 0.002147255 | 0.963841274 | 0.97895169 | Citrate lyase beta chain |
| AP53_972 | 0.005200297 | 8.01282363 | 0.002090096 | 0.964325415 | 0.97895169 | Type I restriction-modi |
| AP53_1541 | -0.005838404 | 7.953481663 | 0.001876147 | 0.966199265 | 0.979967079 | Magnesium and cobalt |
| AP53_1746 | -0.016717466 | 6.287957149 | 0.001861056 | 0.966335389 | 0.979967079 | Glutamate formiminotr |
| AP53_862 | -0.008600892 | 5.314573916 | 0.001787102 | 0.967010607 | 0.980139733 | Adenylate cyclase (EC 4. |
| AP53_1259 | -0.004879879 | 8.599922287 | 0.001579731 | 0.968982448 | 0.981625747 | Phosphopantetheine ad |
| AP53_1059 | -0.003520568 | 6.906797958 | 0.001355252 | 0.971269464 | 0.983429333 | Pleiotropic regulator of |
| AP53_1661 | -0.00669604 | 5.406063771 | 0.001305758 | 0.971798721 | 0.983452199 | Pullulanase (EC 3.2.1.4 |
| AP53_rna85 | 0.010224652 | 4.683211769 | 0.001198311 | 0.972983395 | 0.984005231 | tRNA-Arg-TCT |
| AP53_834 | 0.006109867 | 7.083534567 | 0.001165199 | 0.973359116 | 0.984005231 | Polysaccharide deacety |
| AP53_1203 | -0.006739478 | 3.052125253 | 0.001022322 | 0.975045225 | 0.98496726 | Phage protein |
| AP53_1311 | -0.00650682 | 8.270877675 | 0.000966612 | 0.975734444 | 0.98496726 | Phage protein |
| AP53_334 | 0.007653365 | 7.109344875 | 0.000958765 | 0.975833097 | 0.98496726 | Ribonucleotide reducti |
| AP53_1282 | 0.010596495 | 8.051671386 | 0.000832805 | 0.977475953 | 0.986112695 | hypothetical protein |

| | | | | | | |
|---|---|---|---|---|---|---|
| AP53_143 | -0.004178949 | 4.941427565 | 0.000706094 | 0.97925966 | 0.987029622 | Ascorbate-specific PTS s |
| AP53_1158 | 0.006680846 | 3.822656869 | 0.000696441 | 0.979401882 | 0.987029622 | Late competence prote |
| AP53_230 | 0.005821721 | 5.128972314 | 0.000595096 | 0.980959125 | 0.98804575 | hypothetical cytosolic |
| AP53_789 | 0.005415547 | 5.972167453 | 0.000566128 | 0.981428236 | 0.98804575 | ABC transporter ATP-bi |
| AP53_481 | 0.005374782 | 8.287145628 | 0.000464256 | 0.983181691 | 0.989297907 | bacterial seryl-tRNA syr |
| AP53_441 | -0.004124005 | 4.005282735 | 0.000337006 | 0.985670438 | 0.991288249 | putative L-glutamate lig |
| AP53_478 | 0.001621911 | 6.86776325 | 0.000290117 | 0.986704529 | 0.99181434 | PROTEASE (EC 3.4.-.-) |
| AP53_227 | 0.001891693 | 7.690531152 | 0.000249253 | 0.987676314 | 0.992081047 | Undecaprenyl-diphospl |
| AP53_953 | -0.003151926 | 5.365556629 | 0.000236641 | 0.987992098 | 0.992081047 | Pantothenate kinase (E( |
| AP53_903 | -0.002233462 | 4.168785317 | 0.000136427 | 0.990882401 | 0.994468842 | FIG01114041: hypothe |
| AP53_1343 | 0.001800061 | 4.452770568 | 5.17E-05 | 0.994384553 | 0.997467916 | Phage protein |
| AP53_1512 | -0.000604863 | 8.214941548 | 3.21E-05 | 0.9955793 | 0.998150528 | ATP-dependent DNA he |
| AP53_640 | -0.000743765 | 4.209067475 | 1.63E-05 | 0.996844573 | 0.998879979 | hypothetical phage pro |
| AP53_180 | -0.001109353 | 2.720633585 | 8.25E-06 | 0.997757287 | 0.998879979 | hypothetical protein |
| AP53_1210 | 0.001430835 | 1.936449278 | 7.58E-06 | 0.997850736 | 0.998879979 | Phage protein |
| AP53_259 | -5.43E-05 | 7.002333297 | 5.53E-08 | 0.999816452 | 0.999921318 | Branched-chain amino |
| AP53_265 | -2.52E-05 | 9.95527985 | 1.02E-08 | 0.999921318 | 0.999921318 | Heat shock protein Htp |

L7/L12 (P1/P2)
:tein
:hyltransferase I
enzyme (EC 1.1.1.38)

protein] reductase (EC 1.1.1.100)

n

etical protein
ethyltransferase RumA (EC 2.1.1.-) ## LSU rRNA m(5)U1939
L36p
midase
ase (EC 6.1.1.3)

like protein
protein] synthase, KASII (EC 2.3.1.41)

L10p (P0)
osidase (EC 3.2.1.85)
n
cleotidyltransferase (EC 2.7.7.8)
:tein
pecific IIB component (EC 2.7.1.69) / PTS system, mannose-specific IIA component (EC 2.7.1.69)
protein DivIVA
protein of acetyl-CoA carboxylase
3.5.3.6)
EC 6.1.1.11)
(EC 6.1.1.7)

:tein
boxyl transferase beta chain (EC 6.4.1.2)
etical protein
etical protein

[acyl carrier protein] dehydratase (EC 4.2.1.-)
-
L13p (L13Ae)
tein
specific IIC component (EC 2.7.1.69)
ase (EC 6.1.1.5)
7.4.14)
L17p
S19p (S15e)
merase beta' subunit (EC 2.7.7.6)
ster phosphodiesterase (EC 3.1.4.46)
actosamine-specific IID component (EC 2.7.1.69)
C 3.6.3.14)
of PTS system

etical protein
L22p (L17e)
licase YqfR
amily chaperone GroEL
C 2.4.1.212)
etical protein
O (EC 3.4.24.-)
like protein
nse regulator SA14-24
e (EC 1.1.1.27)
J
7.4.3)
ge associated)
subunit
ansferase (EC 2.1.3.3)
n
etical protein
otein
etical protein
cetyl-CoA carboxylase (EC 6.3.4.14)

hage endopeptidase
tRNA ribosyltransferase-isomerase (EC 5.-.-.-)

tein

S13p (S18e)
hetase alpha chain (EC 6.1.1.20)
merase alpha subunit (EC 2.7.7.6)
, phage associated
transporter, periplasmic substrate-binding component (TC 3.A.1.5.2)
III (EC 3.1.11.2)
cetylglucosaminidase (EndoS)
factor Tu
ein] reductase [FMN] (EC 1.3.1.9)
nsporter, substrate-binding protein
ydrolase (EC 3.2.1.26)

ease protein EscB
C 3.5.1.88)
precursor (EC 5.2.1.8)
er protein transacylase (EC 2.3.1.39)
L24p (L26e)
icle, subunit Ffh SRP54 (TC 3.A.5.1.1)
merase beta subunit (EC 2.7.7.6)
ssociated
-ACP isomerase
ent domain protein, GcvH-like

tein
d glucose-specific IIC component (EC 2.7.1.69) / PTS system, maltose and glucose-specific IIB component (EC 2.7.1.69) / PTS system, maltose and gluco
etical protein
L16p (L10e)
l protein - phage associated

3.2.1.24)
e protease (Streptopain) (EC 3.4.22.10) / Streptococcal pyrogenic exotoxin B (SpeB)
tein
tein
S11p (S14e)
otein
2.1.21)

etical protein
etical protein

C 3.6.3.14)
cleotide-binding protein, similarity with At5g48545 and yeast YDL125C (HNT1)
E
etical protein

L6p (L9e)
orter substrate-binding protein
L14p (L23e)
subunit
or RegR, rpressor of hyaluronate and KDG utilization
n (EC 3.6.3.14)

thetase beta chain (EC 6.1.1.20)
aldolase class II (EC 4.1.2.13)

nsporter, membrane-spanning permease protein MsmG
pecific IIC component (EC 2.7.1.69)
inding protein EcsA
in (EC 3.6.3.14)
amily)

ructose family IIB component

(7,8-dihydro-8-oxoguanine-triphosphatase) (EC 3.6.1.-)
S8p (S15Ae)

mily protein [C1]
hydroxylating enzyme, evidenced by COGnitor; PaaD-like protein (DUF59) involved in Fe-S cluster assembly
neasure protein
t pseudouridine synthase B (EC 4.2.1.70)

etical protein
a prime subunit (EC 2.7.7.7)
actosamine-specific IIB component (EC 2.7.1.69)
subunit
L32p
rter, glutamine-binding protein/permease protein
S14p (S29e) @ SSU ribosomal protein S14p (S29e), zinc-dependent
pecific IID component (EC 2.7.1.69)

on ATPase
S1p
S9p (S16e)

L15p (L27Ae)
etical protein

se (EC 2.7.2.3)
rusion (MATE) family transporter
-phosphate uridyltransferase (EC 2.7.7.23) / Glucosamine-1-phosphate N-acetyltransferase (EC 2.3.1.157)

Bsu
S4p (S9e)

alpha chain (EC 6.1.1.14)
secY subunit (TC 3.A.5.1.1)
bunit B (EC 3.6.3.14)

S20p
e family protein
e (EC 1.1.1.6)
L2p (L8e)

PROTEIN (Rhomboid family)
protein tyrosine phosphatase (EC 3.1.3.48)

nction DUF1447
protein A
-CoA synthase (EC 2.3.3.10)
L29p (L35e)
inator, BglG family (EC 2.7.1.69)
-phosphate deacetylase (EC 3.5.1.25)
cific IIB component (EC 2.7.1.69) / PTS system, lactose-specific IIC component (EC 2.7.1.69)
otein precursor SPs0273
transporter
se (EC 6.1.1.21)
horibitol) ligase subunit 1 (EC 6.1.1.13)
C 3.6.3.14)
L18p (L5e)
nsic Protein Family)
3.2.1.23)
L4p (L1e)
hain (EC 3.6.3.14)
somerase (EC 5.3.1.9)
S3p (S3e)
orter ATP-binding protein
L30p (L7e)
L5p (L11e)
etical protein
e (EC 2.3.1.30)
ystem toxin protein
ate uridylyltransferase (EC 2.7.7.9)

;e
:al protein YeeN
ıK
egulator DeoR (transcriptional repressor)

L23p (L23Ae)
acid aminotransferase (EC 2.6.1.42)
(erC

ate aldolase (EC 4.1.3.16) @ 2-dehydro-3-deoxyphosphogluconate aldolase (EC 4.1.2.14)
:iated protein
ge associated)
'stem antitoxin protein
etical protein
sZ (EC 3.4.24.-)

1.30)
actosamine-specific IIC component (EC 2.7.1.69)
chain) D-alanine transfer protein DltD
esis protein B (SagA)
etical protein
( protein 33)
or of fatty acid biosynthesis FabT
ge associated)
etical protein
ase protein
methyltransferase (EC 2.1.1.-)
orter, ATP-binding protein SitB

ıain (EC 3.6.3.14)
S5p (S2e)
ounit E (EC 3.6.3.14)
neasure protein

egulator, Cro/CI family

tein
I--L-alanine ligase @ tRNA-dependent lipid II--L-serine ligase
S17p (S11e)
ent
nsport system, ATPase component
ster assembly scaffold protein for SUF system, SufE2
dylase MnmA
nsporter - APC Superfamily

component of predicted cobalamin ECF transporter
mily)
glucosamine deacetylase (EC 3.5.1.-)
ase protein
e reductase (EC 1.5.1.2)
tein
/keto reductase family, subgroup 1
etical protein
amily co-chaperone GroES
channel family protein

(EC 1.8.1.9)
Gls24 family
osylase (EC 2.4.2.29)
ase of class III (anaerobic), activating protein (EC 1.97.1.4)
icle associated protein
or ScrR, LacI family
dase (EC 3.2.1.35)

etical protein
ibitin-family membrane protease subunit YbbK
tease, ATP-binding subunit ClpE
or, MutR family
bunit C (EC 3.6.3.14)

n (EC 3.6.3.14)

n DltB

EC 5.4.2.2)
ase (EC 3.4.11.7)
C 2.8.1.7), SufS subfamily
nsporter, ATP-binding protein
PIn2
e oxidase (EC 1.1.3.21)

e (EC 2.7.6.5), (p)ppGpp synthetase I
nthase B (EC 1.-.-.-)

pecific IIC component (EC 2.7.1.69)
orter permease protein
ate aldolase (EC 4.1.2.40)
C 2.7.1.107)
or, XRE family
A, responsible for the formation of Glc-DAG

measure protein
ption repressor HrcA
nse regulator, malate (EC 2.7.3.-)
clotransferase (EC 2.3.2.5)
orter, amino acid-binding protein
d mercury transporting ATPase (EC 3.6.3.3) (EC 3.6.3.5); Copper-translocating P-type ATPase (EC 3.6.3.4)
inding protein
ecific IIB component (EC 2.7.1.69) / PTS system, sucrose-specific IIC component (EC 2.7.1.69) / PTS system, sucrose-specific IIA component (EC 2.7.1.69
gen A
or - phage associated
etical protein
otein

/beta superfamily
C
ase (EC 6.1.1.16)
)-methyltransferase (EC 2.1.1.207)
TP-binding protein PstB (TC 3.A.1.7.1)
sW
oxypeptidase (EC 3.4.16.4)
pA/BipA
beta chain (EC 6.1.1.14)
etical protein
n
thiamin biosynthesis lipoprotein ApbE
tein

sA
lyceraldehyde-3-phosphate dehydrogenase (NADP) (EC 1.2.1.9)
synthetase (EC 2.5.1.6)

protein] synthase, KASIII (EC 2.3.1.41)
ecific recombinase
otein
etical protein
protein smc
ol-3-phosphate O-acyltransferase PlsY
natase (EC 3.6.1.13)
3.1.-)
esis protein B (SagB)
oxyl transferase alpha chain (EC 6.4.1.2)
specific IIB component (EC 2.7.1.69)
ponent CbrT of predicted cobalamin ECF transporter

nutase (EC 5.4.2.10)
stem permease protein DppB (TC 3.A.1.5.2)
sphatase YidA, haloacid dehalogenase-like phosphatase family

P-binding protein DppF (TC 3.A.1.5.2)
nbly protein SufB
esis protein C (SagC)

bunit K (EC 3.6.3.14)
l regulator, MutR family
.2)
enase in hyaluronic acid synthesis (EC 1.1.1.22)
Py1011
eductase (EC 1.1.1.169)
t-stimulated GTPase EngC
r kinase YesM (EC 2.7.3.-)
L31p @ LSU ribosomal protein L31p, zinc-independent
actor 1

halogenase (EC 3.8.1.2)
licase YfmL
l hydrolase (EC 3.2.1.-)
ponent BioY of biotin ECF transporter
TP-binding protein
nent, putative

.7.2.2)
etical protein
nding protein
2.7.1.36)
e triad protein
isomerase, LacA subunit (EC 5.3.1.26)
plase YbeY, involved in rRNA and/or ribosome maturation and assembly

rter, periplasmic phosphate-binding protein PstS (TC 3.A.1.7.1)
or, DeoR family
t Clp proteinase (ATP-binding subunit)
kinase (EC 2.7.1.60)
tein

r / Biotin-protein ligase (EC 6.3.4.15)
n
s isomerase (EC 5.2.1.8)

etical protein
or in cluster with beta-lactamase, GntR family
-CoA reductase (EC 1.1.1.34)
ponent PdxU2 of predicted pyridoxin-related ECF transporter
ate ligase (EC 6.3.4.3)
kinase (EC 2.7.9.1)

2.7.4.9)
n KefA protein / Small-conductance mechanosensitive channel
tein (permease)
1.20.4.1)
Py0110
ase related protein
ase (EC 5.4.2.1)
noribitol) ligase subunit 2 (EC 6.1.1.13)
putative
or A
hage associated
fication system, specificity subunit S (EC 3.1.21.3)
r kinase SA14-24
r kinase, associated with ferric iron transporter, SPy1061 homolog
VraS
se (EC 6.1.1.12)
tease proteolytic subunit (EC 3.4.21.92)
rotein SpyM3_1856/SPs1852
essory protein
rane-spanning permease - glutamine transport
porter ArcD
n deacetylase of SIR2 family
(EC 4.3.2.2)

ein
proteins containing CBS domains
g
ase family
foxide reductase MsrB (EC 1.8.4.12)
a
2)
tein
-6-phosphate aminotransferase [isomerizing] (EC 2.6.1.16)

methylaminomethyl modification enzyme GidA
oylglucosamine reductase (EC 1.1.1.158)
e (EC 1.5.1.3)
unit D (EC 3.6.3.14)
mily)
P-binding protein PstB (TC 3.A.1.7.1)
rotein
tease ATP-binding subunit ClpA

-)

tein
leoside 5-triphosphatase RdgB (dHAPTP, dITP, XTP-specific) (EC 3.6.1.15)
P-binding protein PotA (TC 3.A.1.11.1)
ubunit
tor protein
gA
ate dehydrogenase (EC 1.1.1.205)
or, MarR family
[illegible]
y1643

3, responsible for the formation of Gal-Glc-DAG
subunit YajC (TC 3.A.5.1.1)

VII large subunit (EC 3.1.11.6)
ABC transporter permease component potC (TC_3.A.1.11.1)
c exotoxin K (SpeK)
eG like
C0266 (sugar utilization related?)
L3p (L3e)
tease ATP-binding subunit ClpX
ABC transporter permease component PotB (TC 3.A.1.11.1)
ransferase (EC 2.4.2.9)
n translocase component YidC, short form OxaI-like
M family enzyme in heat shock gene cluster, similarity with CPO of BS HemN-type
system permease protein OppC (TC 3.A.1.5.1)
tein #Fam0017
nt responsible histidine kinase
factor RpoD
L20p
ion resolvase YggF
stem permease protein PstA (TC 3.A.1.7.1)
nducible protein PhoH, predicted ATPase
etical protein
P-binding protein DppD (TC 3.A.1.5.2)
ain, possibly ADP-ribose binding module
bunit G (EC 3.6.3.14)
S10p (S20e)
or, TetR family
L21p
yltransferase (EC 2.4.2.7)
nsporter, membrane-spanning permease protein MsmF
partitioning protein ParB / Stage 0 sporulation protein J
ase of class III (anaerobic), large subunit (EC 1.17.4.2)
C-methyl-D-erythritol kinase (EC 2.7.1.148)
MP-, MA-, MS-, AP-, NP- specific)
-phosphate lactonase UlaG (EC 3.1.1.-) (L-ascorbate utilization protein G)
cting protein related to cell division
protein phosphotransferase of PTS system (EC 2.7.3.9)

on protein NrdI

-CoA transferase, alpha subunit (EC 2.8.3.8)
osA

etical protein

nase (EC 2.7.4.2)
ltransferase, MraW methylase family (EC 2.1.1.-)
n permease protein
otein #Fam0015
on helicase
ors, LysR family

rase (amylomaltase) (EC 2.4.1.25)
accessory protein
otein
genic factor 1
igase (EC 6.2.1.22)

ATP-binding protein OppF (TC 3.A.1.5.1)
or, Cro/CI family
etical protein
etical protein
EC 3.4.11.15)
t of efflux system
cpC (EC 3.4.21.-)
cylase

etical protein
rotein Ccs4
cal protein, contains S4-like RNA binding domain
etical protein

histidine kinase, malate (EC 2.7.3.-)
dotransferase subunit C (EC 6.3.5.6) @ Glutamyl-tRNA(Gln) amidotransferase subunit C (EC 6.3.5.7)
ling/permease protein MsbA
olase (EC 3.-.-.-)

[illegible]
ate uridylyltransferase (EC 2.7.7.9)
rase (EC 5.3.1.1)
e family protein
te delta-isomerase, FMN-dependent (EC 5.3.3.2)
L-rhamnosyltransferase/alpha-L-Rha alpha-1,3-L- rhamnosyltransferase (EC 2.4.1.-)
factor G
ubunit methyltransferase E (EC 2.1.1.-)
.26.3)
.1.3.2)
icle receptor protein FtsY (=alpha subunit) (TC 3.A.5.1.1)
regulator, AraC family
ha subunit (EC 2.7.7.7)
etical protein
otein YeiH
ptional regulator MntR
wlG
etical protein
[contains diacylglycerol kinase catalytic domain]

e E1 component alpha-subunit (EC 1.2.4.-)
methyltransferase (EC 2.1.1.31)
erase system repressor
etical protein
etical protein
TP-binding protein ZnuC
epimerase (EC 5.1.3.1)
decarboxylase (EC 4.1.1.33)

moyl-pentapeptide-transferase (EC 2.7.8.13)

or of arabinoside utilization operon, GntR family
1.1)
EC 2.1.1.45)

ein YmcA
al RNA; ssuRNA; SSU rRNA
e III family protein

otein RimM
ponent ThiT of thiamin ECF transporter
or PerR, FUR family
mG/Ycf19 (putative), YggT family
etical protein
pecific IID component (EC 2.7.1.69)
ase LtaS Type Iic (The first GCT-Ala is overlapped with the last codon of the former half interrupted protein)
ein/pseudogene causing stop codon at 85th aa
A
C 5.99.1.3)
gB
or, MutR family
ine 1-carboxyvinyltransferase (EC 2.5.1.7)
rter, iron-binding protein

(Streptococcus), ATP-binding protein
sistance protein
or for NAD biosynthesis in gram-positives

er, permease protein
.1.19)
protein family
d transcriptional attenuator LytR-CpsA-Psr, subfamily F1 (as in PMID19099556)

tein UPF0176, Firmicutes subgroup

etical protein
ining protein
rea)
e protein
ounit A (EC 3.6.3.14)
n translocase component YidC, OxaA protein

sL
for oriC/DnaA-independent DNA replication
e n-acetyltransferase

kinase (EC 2.7.6.2)
amidotransferase component (EC 4.1.3.27) @ Para-aminobenzoate synthase, amidotransferase component (EC 2.6.1.85) # TrpAb@PabAb
oxypeptidase (EC 3.4.16.4)

lpentapeptide-lysine N(6)-alanyltransferase (EC 2.3.2.10) (EC 2.3.2.-)
n cluster with Ecs transporter (in Streptococci)
S6p
')-C(5))-methyltransferase (EC 2.1.1.176)
etical protein
/SagA/PcsB, putative peptidoglycan hydrolase
ion protein NusB
al RNA; ssuRNA; SSU rRNA
hage associated
esis protein D (SagD)
protein containing multiple CBS domains
or AdcR for Zn(2+)-responsive expression
putative; stress-responsive transcriptional regulator
etical protein
actor 3

ose 3,5-epimerase (EC 5.1.3.13)
TP-binding protein
7.1.26) / FMN adenylyltransferase (EC 2.7.7.2)

:ohydrolase inhibitor
aldehyde-3-phosphate dehydrogenase (EC 1.2.1.12) / Plasmin(ogen) receptor
nit B
se (EC 4.4.1.8)
nding protein YvcR
lic-dimer hydrolase (EC 3.5.2.12)
hydrolase
ycosylase (EC 3.2.2.-)
ductase protein C (EC 1.6.4.-)
L19p

onent of general energizing module of ECF transporters
etical protein

ansmembrane permease (SagI)
al RNA; ssuRNA; SSU rRNA
otion regulator
pecific ABC transporter, ATP-binding protein
ibosomal protein
kinase (EC 2.7.1.60)
ng endonuclease
(Streptococcus), permease protein
nylase
ABC transporter, permease protein MalG
al RNA; ssuRNA; SSU rRNA
phosphoribosyltransferase (EC 2.4.2.8)
EF, lipoprotein IsdE
on control protein YabA

5.4.12); Late competence protein ComEB
positive regulator (Mga)

thetase domain protein (Bsu YtpR)
family 1

in
al RNA; ssuRNA; SSU rRNA
stem regulatory protein PhoU
3su YhaM
nbly protein SufD

glycosylase (EC 3.2.2.20)
(EC 2.3.1.54)
protein] reductase (EC 1.1.1.100)
C 5.99.1.3)
conate kinase (EC 2.7.1.45)
CoA-transferase subunit B (EC 2.8.3.9)
al RNA; ssuRNA; SSU rRNA
[contains diacylglycerol kinase catalytic domain]

nbly ATPase protein SufC
Gls24 family
protein
epeat unit export protein
specific IIA component (EC 2.7.1.69)

osyltransferase (EC 2.4.2.11)
protein
eneral energizing module of ECF transporters
competence regulon ComD
al RNA; lsuRNA; LSU rRNA
ATP-binding protein OppD (TC 3.A.1.5.1)
etical protein
S16p
M16 family
ster with ribosomal protein L32p, Firmicutes subfamily
or SpxA2
etical protein

ge gene expression IbrA
S15p (S13e)
al RNA; lsuRNA; LSU rRNA
rotein
(
py1939 homolog

orter, glutamine-binding protein/permease protein
in ComC, processing protease
ein TrkH
L33p @ LSU ribosomal protein L33p, zinc-independent
nsport system, permease protein OpuAB / Glycine betaine ABC transport system, glycine betaine-binding protein OpuAC

al RNA; lsuRNA; LSU rRNA
tein
al RNA; lsuRNA; LSU rRNA
f exopolysaccharide synthesis, competence and biofilm formation Ftr, XRE family
sidase (EC 3.2.1.70)
ated to Actinobacillus protein (1944168)
g protein YlqC
cal protein
dent flavin oxidoreductase in cluster with COG2110
nse regulator, associated with ferric iron transporter, SPy1062 homolog
activating enzyme (EC 1.97.1.4)
etical protein
ain) required for threonylcarbamoyladenosine t(6)A37 modification in tRNA
RecR
.2.3)
XerD
(EC 2.4.2.3)
protein
.48) [C1]
etical protein
n
isomerase, LacB subunit (EC 5.3.1.26)

rase
c exotoxin C (SpeC); _Toximoron (Superantigen)
cal membrane associated protein
e (EC 2.7.1.11)
ase protein
otidase-transglycosylase (EC 2.4.1.129) (EC 3.4.-.-)

al RNA; lsuRNA; LSU rRNA
ase (EC 6.1.1.17) @ Glutamyl-tRNA(Gln) synthetase (EC 6.1.1.24)
al RNA; lsuRNA; LSU rRNA
obP, contains nucleotide-binding domain of DisA bacterial checkpoint controller
nducible protein J
EC 3.5.3.8)
t of efflux system

ictor 1
4)
ygenase (EC 1.14.-.-)
or, MarR family
l protein
ucleic-acid-binding protein implicated in transcription termination

ination protein NusG
in kinase PrkC, regulator of stationary phase
asmic spermidine putrescine-binding protein PotD (TC 3.A.1.11.1)
(EC 1.8.1.9)

(7,8-dihydro-8-oxoguanine-triphosphatase) (EC 3.6.1.-)
etical protein
iptional regulator (AT-rich DNA-binding protein)
etical protein
nding protein uup
t pseudouridine synthase D (EC 4.2.1.70)

rotein MutL

IG046916
fication system, DNA-methyltransferase subunit M (EC 2.1.1.72)
ice hemoprotein receptor Shr
ase protein
ponents of ABC transporters with duplicated ATPase domains
S21p

otein
stem permease protein DppC (TC 3.A.1.5.2)
in

HAD subfamily IIIA
n response regulator

otein
ansmembrane permease (SagH)
etical protein
eductase QueG
or, GntR family
l regulator, MutR family

tidase (EC 3.4.23.36)
ling/permease protein MsbA
tein
mbrane protein (putative)
dotransferase subunit A (EC 6.3.5.6) @ Glutamyl-tRNA(Gln) amidotransferase subunit A (EC 6.3.5.7)
etical protein
repair protein RecT (prophage associated)
uramoyl-L-alanine amidase (EC 3.5.1.28)
transferase

ne ABC transport system permease protein ProV (TC 3.A.1.12.1)
S12p (S23e)
epressor
o, transfers heme from hemoglobin to apo-SiaA/HtsA

ɛtical protein
l-Ala--L-alanine ligase @ tRNA-dependent lipid II-Ser--L-alanine ligase
ıcid transport system, ATPase component
ɔtein
ɛtical protein
ımily protein
ɛ (EC 1.1.1.1); Acetaldehyde dehydrogenase (EC 1.2.1.10)
e-34 deaminase (EC 3.5.4.-)
ɔtein
upling factor

ɛtical protein
11.1.1)
er (ATP-binding protein), spy1791 homolog
er (ATP-binding protein)
ɛtical protein
or, HxlR family
ed protein-related protein
lonuclease containing a URI domain
yltransferase PlsX
with VraSR
foxide reductase MsrA (EC 1.8.4.11) / Peptide methionine sulfoxide reductase MsrB (EC 1.8.4.12)
or pfoR
A
sm protein SecA ATPase RNA helicase (TC 3.A.5.1.1)

L28p
nding protein
ase LtaS Type Iic/interrupted by large phage
4.21.89)
-occurring with transport systems (COG1739)
ggS, proline synthase co-transcribed bacterial homolog PROSC
ecific recombinase
ase protein

specific IIB component (EC 2.7.1.69)
e
se, subunit A
or, RofA-like Protein (RALP)
ein] synthase (EC 2.7.8.7)
etical protein
mpetence-specific nuclease) (EC 3.1.30.-)
ductase associated with MetSO reductase
nore transport system, permease component
or, MerR family
mily)
adation protein PaaD, thioesterase
rylase (EC 2.4.1.1)
thiD family (EC 2.7.1.35)
otein B

n
e--alanine ligase (EC 6.3.2.8)
tein
e
lon subunit (EC 2.7.7.7)
ion protein NusA
terase
e (EC 1.1.1.1)
ease
etical protein
3.4.13.9)

e (EC 2.3.1.16) @ Acetyl-CoA acetyltransferase (EC 2.3.1.9)
etical protein

lehydrogenase [NAD(P)+] (EC 1.1.1.94)

otein RmuC
etical protein
nthase A (EC 4.2.1.70)
ng protein Dps / Iron-binding ferritin-like antioxidant protein / Ferroxidase (EC 1.16.3.1)
bosyltransferase [decarboxylating] (EC 2.4.2.19)
actor 2
pecific ABC transporter, permease protein
phage associated
og
etical protein
tein
ase (EC 1.13.12.4)
L35p
nate synthase (EC 2.5.1.31)
e peptidase (EC 3.4.19.3)
C 5.4.99.5)
or

criptional repressor
specific IIA component (EC 2.7.1.69)

bosyltransferase [decarboxylating] (EC 2.4.2.19)
te:Formate Antiporter

se (EC 4.3.1.3)

nding protein
CoA ligase (EC 6.2.1.3)
e decarboxylase (EC 4.1.1.23)
stem permease protein PstC (TC 3.A.1.7.1)
ease, CAAX family
etical protein
transport protein CorA

etase (EC 6.1.1.22)

ɔunit F (EC 3.6.3.14)

ɔyM3_1200
/ protein (S1 RNA-binding domain)

ɛtical protein
midazole carboxylase catalytic subunit (EC 4.1.1.21)
ɛtical protein
4.21.89)
protein
rved protein, contains double-stranded beta-helix domain

ɔmpetence-specific nuclease) (EC 3.1.30.-)
e-C5-methylases
ɛtical protein

C 4.2.3.5)
system permease protein OppB (TC 3.A.1.5.1)
ydroxymethyldihydropteridine pyrophosphokinase (EC 2.7.6.3)
ein NrdH, required for reduction of Ribonucleotide reductase class Ib
ɔhate reductase (EC 1.2.1.41)
ɛtical protein
lalanyl-D-glutamate--L-lysine ligase (EC 6.3.2.7)
nsport system, ATP-binding protein OpuAA (EC 3.6.3.32)
n
it pseudouridine synthase A (EC 4.2.1.70)
erase - possibly involved in cell wall localization and side chain formation of rhamnose-glucose polysaccharide
eductase
gulator
egulator, Cro/CI family

ıomolog, phosphoribosyltransferase domain; protein YhgH required for utilization of DNA as sole source of carbon and energy

dismutase (EC 1.15.1.1)
or, RofA-like Protein (RALP)
:idase (EC 3.4.11.18)

:tical protein

porter, periplasmic oligopeptide-binding protein OppA (TC 3.A.1.5.1)
: (EC 6.1.1.4)
7.4.8)
ydrogenase of acetoin dehydrogenase (EC 1.8.1.4)
ısation protein B
ipoteichoic acid biosynthesis, DltR
R
YrrC
ecific recombinase
ltransferase HI0095 (UbiE paralog)
:tical protein
family
:ative
ɔhage associated

c acid-binding protein YchF
Py0887

:tical protein
e RNase R
ype I (EC 6.3.1.2)
rge subunit pseudouridine synthase D, Bacillus subtilis YhcT type
transferases

methyltransferase TrmN6 (EC 2.1.1.223)
(EC 5.4.2.7)
ptional regulator, ArpU family

or, LuxR family
tein
ease protein
etical protein
orter, periplasmic-binding protein SitA
(EC 2.7.8.-)
l protein - phage associated
helicase RuvA
Py2060
class II) (EC 6.1.1.6)
ic
e synthase (EC 2.5.1.90) / Dimethylallyltransferase (EC 2.5.1.1) / (2E,6E)-farnesyl diphosphate synthase (EC 2.5.1.10) / Geranylgeranyl diphosphate synt
etical protein
or DegU, LuxR family
ase (Zinc) SprT family
otein
nomutase
oxidases family
III (EC 3.1.11.2)

etical protein
family
pecific IIA component (EC 2.7.1.69)
etical protein
etical protein
tein
methyltransferase PrmC, methylates polypeptide chain release factors RF1 and RF2

etical protein
tein
n

onal pleiotropic repressor codY
(EC 2.7.1.76) / Deoxyguanosine kinase (EC 2.7.1.113)

.1.40)
orter, inner membrane permease protein SitD
e (EC 2.3.1.16) @ Acetyl-CoA acetyltransferase (EC 2.3.1.9)
etase (EC 6.1.1.10)
riptional regulator VraR
e protein

reo-5-hexosulose-uronate ketol-isomerase (EC 5.3.1.17)
transferase (EC 2.1.2.9)
.2)
etical protein
e permease
etical protein
or, TetR family
deoxynucleoside utilization, GntR family
n
etical protein

ory protein
n
etical protein
ase similar to cobyric acid synthase

oxypeptidase (EC 3.4.16.4)

se (EC 3.1.1.29)
se
l protein - phage associated
ecific recombinase
rter GlyP

ne synthase (former ThiI)
ıse domain protein (class beta)

tion
etical protein
ısferase (EC 2.1.1.-)
nase I alpha (EC 1.1.1.25)
uptake protein
ge gene expression IbrB
merase omega subunit (EC 2.7.7.6)
E1 component beta-subunit (EC 1.2.4.-)
nthase, aminase component (EC 2.6.1.85) / Aminodeoxychorismate lyase (EC 4.1.3.38)
ylase/Succinyl-diaminopimelate desuccinylase and related deacylases
rate-binding protein
tein
ydrogenases
etical protein
hage-related DNA binding protein
hosphatase
etical protein
eta subunit (EC 4.3.1.17)
1.1.1)
lopeptidase (IdeS) / Mac/ Secreted immunoglobulin binding protein (Sib38)

etical protein

26.11)
n
ıctor 2; programmed frameshift-containing
ferase, class I
nsport system, ATPase component

vIC (FtsB), stabilizes FtsL against RasP cleavage
ate aldolase (EC 4.1.2.40)
C 3.5.4.16) type 1

dase (EC 3.4.11.4)
erase (EC 2.7.7.21) (EC 2.7.7.25)
e photolyase (EC 4.1.99.3)
ossible functionally linked with Alanyl-tRNA synthetase
lase (EC 4.1.2.25)

n
ase of class Ib (aerobic), beta subunit (EC 1.17.4.1)
etical protein
or, GntR family
ein rotamase A; Peptidyl-prolyl cis-trans isomerase (EC 5.2.1.8)
etase
n histidine kinase
TrmA family
ne metabolism regulator CmbR, LysR family
ATP-binding protein; Streptolysin S export protein (SagG)
unity protein (SagE)
o acid transporter protein
foxide reductase MsrA (EC 1.8.4.11)
erase (EC 2.6.1.1)
ponent NiaX of predicted niacin ECF transporter
surface immunogenic protein
tein
nner membrane permease protein ZnuB

esis protein (SagF)
transferase RlmH
etical protein

amily), YqeK
ors, LysR family
hore transport system, permease 2 component
tein
ptidase-transglycosylase (EC 2.4.1.129) (EC 3.4.-.-) / Penicillin-binding protein 1A/1B (PBP1)

he competence regulon ComE
e dehydrogenase (EC 1.1.1.30)
ı
-O-) -methyltransferase rlmB (EC 2.1.1.-) ## LSU rRNA Gm2251
nbrane protein
isomerase, LacB subunit (EC 5.3.1.26)

in ComGA, access of DNA to ComEA
pha subunit (EC 4.3.1.17)
S7p (S5e)
orter ATP-binding protein
rter, permease protein 2
tical protein
in
ı

dent helicase YoaA
ed protein, YBBR Bacillus subtilis homolog
or SpxA1

tidase (EC 3.4.14.11)
tical protein
e-3-phosphate synthase (EC 2.5.1.19)

nymidylyltransferase (EC 2.7.7.24)
horibosyl-dephospho-CoA transferase (EC 2.7.7.61)
ltransferase (EC 2.4.2.10)
C 5.99.1.2)
tical protein
zation operon protein PhnA
protein J
ase (EC 4.1.3.3)
ı

l protein - phage associated

or, Cro/CI family
etical protein
-glycine ligase (EC 6.3.4.13)
; motif, CinA N-terminal domain / C-terminal domain of CinA type S

synthase large chain (EC 6.3.5.5)

n response regulator CiaR

ylase alpha chain (EC 4.1.1.3)
PROTEIN (Rhomboid family)
orter, amino acid-binding protein

or ArcR essential for anaerobic expression of the ADI pathway, Crp/Fnr family
cted N-acetylneuraminate transport system permease protein 1
in ComGB, access of DNA to ComEA
it pseudouridine synthase A (EC 4.2.1.70)
tein

eneral energizing module of ECF transporters
genesis protein CcdA homolog, associated with MetSO reductase
IRSA252 (BX571856) SAR1694
ne phosphatase PrpC, regulation of stationary phase
midazole carboxylase ATPase subunit (EC 4.1.1.21)

n
ibonuclease in RNA processing)
n
otein
cal protein
etical protein
etical protein
esistance protein CzcD
in ComGC, access of DNA to ComEA, FIG007487

etical protein
tor (EC 5.2.1.8)
yceryl transferase (EC 2.4.99.-)
etical protein
cal protein
Py1656
n

on initiation protein dnaD
VII small subunit (EC 3.1.11.6)

tT/nudix family protein)
-alanine acetyltransferase
lipoprotein ZinT
nding protein
etical protein

etical protein
tase (EC 5.4.2.6) (EC 2.7.1.41)

lic-dimer hydrolase (EC 3.5.2.12)
atory protein ArgR, repressor of arg regulon
3.1.-)
rotein

2.7.2.11)
(EC 6.3.2.12) / Folylpolyglutamate synthase (EC 6.3.2.17)
sphorylase (EC 2.4.2.1)
sphorylase (EC 2.4.2.1); N-Ribosylnicotinamide phosphorylase (EC 2.4.2.1) ## possible
ethylguanosine (m7G) methyltransferase GidB

ng peptide ABC transporter ATP-binding protein ComA
phosphokinase (EC 2.7.6.1)
in ComGD, access of DNA to ComEA, FIG038316

ɔn regulator EzrA
tor
ı
ı
ıatase (EC 3.1.3.25)

5.1.3.3)
3.5.4.10) / Phosphoribosylaminoimidazolecarboxamide formyltransferase (EC 2.1.2.3)
meostasis protein CutC

l)-3'-dephosphocoenzyme-A synthase (EC 2.7.8.25)
etical protein

ıe binding ABC transporter protein ProX (TC 3.A.1.12.1) / Osmotic adaptation

etical protein
nding protein
P-loop-containing kinase
etical protein
tein
-DNA glycosylase (EC 3.2.2.23)

ɔsB, coordinates the switch between cylindrical and septal cell wall synthesis by re-localization of PBP1

5-formylation enzyme TrmE
nsport system, permease component
nethyltransferase (EC 2.1.1.33)
system, EIIC component
etical protein
n containing protein
factor P
ıa subunit (EC 2.7.7.7)
ase protein
tative

fflux pump

de-triphosphate reductase (EC 1.17.4.2)
ine 1-carboxyvinyltransferase (EC 2.5.1.7)
e (EC 6.1.1.19)
or, MerR family
2.7.7.7)
etical protein
3.1.-)
1.7)
nsferase FtsJ like
ein A

rane-spanning permease - glutamine transport
arge subunit [C1]

piA
oup 2 family protein
erase (EC 2.6.1.1)
ng protein
L27p

nuclease and 5' exonuclease)

n

ng peptide ABC transporter ATP-binding protein ComA
sociated
n

n
egulator, RpiR family
ate N-acetylglucosaminyl 1-phosphate transferase (EC 2.7.8.-)

:al protein

enase (EC 1.1.1.22)
n
etical protein
eat shock protein implicated in the recycling of the 50S subunit (S4 paralog)
ne-hydrolyzing], amidotransferase subunit (EC 6.3.5.2) / GMP synthase [glutamine-hydrolyzing], ATP pyrophosphatase subunit (EC 6.3.5.2)
actosamine- and galactosamine-specific IIA component (EC 2.7.1.69)
component (EC 3.1.26.5)
kinase (EC 2.7.1.144)
yltransferase component (E2) of acetoin dehydrogenase complex (EC 2.3.1.-)
C 3.5.1.88)
tein
N6-adenine-specific DNA methylase

ic exotoxin Z (SmeZ)

olved in the export of O-antigen, teichoic acid lipoteichoic acids
is and transport regulator MtaR, LysR family
helicase RuvB

etal-dependent proteases, putative molecular chaperone
gase A (EC 6.3.2.4)
rge subunit pseudouridine synthase D, Bacillus subtilis YjbO type
rogenase, catalytic subunit (EC 1.3.3.1)

inorganic pyrophosphatase (EC 3.6.1.1)
otein
etical protein

-binding protein
phate isomerase (EC 5.3.1.-)

etical protein
etical protein

tein (Rhomboid family)
units gamma and tau (EC 2.7.7.7)
3.4.22.40)
lalanine--D-glutamate ligase (EC 6.3.2.9)
etical protein
syltransferase (EC 2.4.2.22)
bunit I (EC 3.6.3.14)
tein

gase (EC 6.3.1.1)
HtrA, do-like (EC 3.4.21.-)

n
transporter
sphate triphosphohydrolase (EC 3.1.5.1)
ase
7.1.21)

.7.7.7)
7.1.31)
etical protein
se family protein
r kinase yesM (EC 2.7.3.-), associated with MetSO reductase
actor 3
cerol-3-phosphate 3-phosphatidyltransferase (EC 2.7.8.5)
n
etical protein
ular nuclease 2; Mitogenic factor 2
minator of lichenan operon, BglG family
rement C protein
on enzyme
C 5.1.1.3)
tein
etical protein
Py2065

positive regulator (Mga)
nsation protein A
cted N-acetylneuraminate transport system permease protein 2
n SmpB
, DNA transporter ATPase
merase delta subunit (EC 2.7.7.6)
ne kinase CsrS / deletion of T at 1407 causing a early stop codon at 469th aa)
otein
se (DnaB) (EC 3.6.4.12)
se Gcn5
mide formyltransferase (EC 2.1.2.2)
g proteins
ylase/Succinyl-diaminopimelate desuccinylase and related deacylases
etical protein
lX
glycinamidine cyclo-ligase (EC 6.3.3.1)
etical protein
phosphatase
phage associated
sI [Peptidoglycan synthetase] (EC 2.4.1.129)
B
ism regulator
ptidase-transglycosylase (EC 2.4.1.129) (EC 3.4.-.-)
minator with PTS regulation domain, SPy0181 ortholog
plate dehydrogenase (NADP+) (EC 1.5.1.5) / Methenyltetrahydrofolate cyclohydrolase (EC 3.5.4.9)
e kinase (EC 2.7.4.6)

d transcriptional attenuator LytR-CpsA-Psr, subfamily F2 (as in PMID19099556)
cysteine decarboxylase (EC 4.1.1.36)
rane-spanning permease - macrolide efflux
ponent RibU of riboflavin ECF transporter

L34p
dotransferase subunit B (EC 6.3.5.6) @ Glutamyl-tRNA(Gln) amidotransferase subunit B (EC 6.3.5.7)
r, periplasmic cystine-binding protein FliY

:tical protein
:rface protein
ı, required for threonylcarbamoyladenosine t(6)A37 formation in tRNA

:tical protein
 protein
thetase (EC 6.1.1.2)
d repair protein RecO
osphatase B (EC 3.1.3.27)

:-S oxidoreductase
:.1)
tein
:tical protein

:e (EC 4.4.1.8)
ic hypothetical protein
ı
ase (EC 2.5.1.15)
ase of class Ib (aerobic), alpha subunit (EC 1.17.4.1)
:ransferase (EC 2.4.2.14)
3)
protein
pecific IIB component (EC 2.7.1.69)
ansferase (EC 2.4.1.-)
inding protein) -possibly involved in cell wall localization and side chain formation of rhamnose-glucose polysaccharide
:tical protein

:tical protein
:al protein
specific IIC component (EC 2.7.1.69)
:ssor protein - phage associated
:e, subunit B
:e, Autolysin2 (EC 3.5.1.28)
adenylyltransferase (EC 2.7.7.18)

subunit SecE (TC 3.A.5.1.1)
etical protein
ge associated)
synthase small chain (EC 6.3.5.5)
e nucleosidase (EC 3.2.2.16) / S-adenosylhomocysteine nucleosidase (EC 3.2.2.9)
age associated
n
system, EIIB component (EC 2.7.1.69)
a subunit (EC 2.7.7.7)
rotein kinase (EC 2.7.3.-)
pecific exonuclease RecJ (EC 3.1.-.-)
hage endopeptidase
ATP-binding protein FhuC (TC 3.A.1.14.3)
osamine hydrolase like
ponent BL0693 of energizing module of predicted ECF transporter
nding protein
etical protein
-type ATPase (EC 3.6.3.4)
tative
l protein - phage associated
otein/major pilin backbone protein T-antigen
etical protein
otein
amily)
etical protein
e ATP-binding protein drrA
ular nuclease 3; Mitogenic factor 3
er, substrate-binding protein
ogenase, S-alcohol forming, (R)-acetoin-specific (EC 1.1.1.4) / Acetoin (diacetyl) reductase (EC 1.1.1.5)
etical protein

etical protein

n
nction DUF208

c exotoxin G (SpeG)

n
or CtsR
ase of class Ib (aerobic), alpha subunit (EC 1.17.4.1)

isomerase, LacA subunit (EC 5.3.1.26)
n
ltransferase (EC 2.1.1.72)
g protein
e, alpha subunit-related enzymes
l protein - phage associated
etical protein
onent BL0694 of energizing module of predicted ECF transporter
ipoteichoic acid biosynthesis, sensor histidine kinase
n
atory protein ArgR, repressor of arg regulon
ransporter
nnoosyltransferase, involved in cell wall biogenesis
UF194, DegV family
sH (EC 3.4.24.-)
sK
etical protein
etyltransferase (EC 2.3.1.-)
kaline-shock protein
11.1.1)
etical protein

otein
e (EC 2.7.6.5)
.7.1.71)
)-N(2))-methyltransferase (EC 2.1.1.171) ## SSU rRNA m(2)G966
ine--N-acetylmuramyl-(pentapeptide) pyrophosphoryl-undecaprenol N-acetylglucosamine transferase (EC 2.4.1.227)

merase A (EC 5.3.1.6)
nit C
H3604
etical protein
etical protein
cysteine synthetase (EC 6.3.2.5)
notransferase family protein

ponent PdxU of predicted pyridoxine ECF transporter
ate deaminase (EC 3.5.99.6)
bunit methyltransferase N (EC 2.1.1.-)
or, GntR family
etical protein
thyltransferase H
protein
ehydrogenase I beta (EC 1.1.1.282)
ain, acyl carrier protein (EC 4.1.3.6)
etical protein
ide-binding protein Smf possibly involved in DNA uptake
glycinamidine synthase, synthetase subunit (EC 6.3.5.3) / Phosphoribosylformylglycinamidine synthase, glutamine amidotransferase subunit (EC 6.3.5
or OrfX
S18p @ SSU ribosomal protein S18p, zinc-independent
ng protein
otein
.4.-)
ciated protein
olysacharide translocation permease
CoA-transferase subunit A (EC 2.8.3.9)
eH, required for biogenesis of 30S ribosome subunit
phosphokinase (EC 2.7.6.1)
etical protein
(P)H nitroreductase (EC 1.-.-.-) / Dihydropteridine reductase (EC 1.5.1.34)
(EC 2.3.1.54)

CoA-transferase subunit B (EC 2.8.3.9)

of transporters
e (EC 6.1.1.1)
flux pump PmrA
tein

3.5.4.5)
ed protein SPs0857
ecific IIA component (EC 2.7.1.69) / PTS system, fructose-specific IIB component (EC 2.7.1.69) / PTS system, fructose-specific IIC component (EC 2.7.1.
eriplasmic protein precursor (TC 3.A.1.14.3)
se (EC 4.4.1.5)
hase (EC 4.2.3.4)
hydrolase (EC 3.2.1.93)

.5.1.47)
ein
lyase (EC 4.4.1.21) / Autoinducer-2 production protein LuxS

dratase I (EC 4.2.1.10)
alogenase-like family
.1.5)
leotide glycohydrolase (NADGH) (EC 3.2.2.5)
L1p (L10Ae)
sphate nucleotidohydrolase (EC 3.6.1.23)
r ATP binding/permease protein
f unknown function, cellobiose operon
etical protein
he competence regulon ComE
nse regulator yesN, associated with MetSO reductase
etical protein
etical protein

hage associated
e transmembrane protein
tein
otein

midazole-succinocarboxamide synthase (EC 6.3.2.6)
etical protein
cid:cation (Na+ or H+) symporter
otein A
se (DnaB) (EC 3.6.4.12)
rotein MutS
chanosensitive channel
6-phosphate 2-epimerase (EC 5.1.3.9)

domain protein
olate cyclodeaminase (EC 4.3.1.4)
etical protein

carboxylase, beta chain (EC 4.1.1.41); Oxaloacetate decarboxylase beta chain (EC 4.1.1.3)
isomerase (EC 5.3.1.8)
EC 6.1.1.9)
ns-isomerase (EC 5.2.1.8); Proteinase maturation protein (RopA)
ases, 23S RNA-specific
sQ
e mannose operon, ManO
ansferase (EC 2.1.3.2)
ponent QueT (COG4708) of predicted queuosine-regulated ECF transporter
2.1.21)
etical protein
goribonuclease A, Bacillus type
side-specific IIB component (EC 2.7.1.69) / PTS system, beta-glucoside-specific IIC component (EC 2.7.1.69) / PTS system, beta-glucoside-specific IIA cor
etical protein
l protein - phage associated
ng peptide ABC transporter permease protein ComB
L9p
2)
cal protein

L11p (L12e)

ulator associated with purine metabolism
ing protein
in ComEC, DNA transport

next to Sag operon
pecific IIB component (EC 2.7.1.69) / PTS system, trehalose-specific IIC component (EC 2.7.1.69) / PTS system, trehalose-specific IIA component (EC 2.7

etical protein
se, Autolysin1 (EC 3.5.1.28)
.99.18)
3.5)
etical protein
n
sensor histidine kinase CiaH (EC 2.7.3.-)

cific IIA component (EC 2.7.1.69)
ubunit methyltransferase C (EC 2.1.1.52) ## SSU rRNA m2G1207
-rhamnosyltransferase (EC 2.4.1.-)

ansporter, PnuC-like
etical protein
on factor GreA
nethylase
ciated protein

n (EC 4.1.3.6)

TPase (EC 3.6.3.8)
ansferase (EC 2.4.2.-)
ase protein

erase system repressor

ase of class Ib (aerobic), beta subunit (EC 1.17.4.1)

protein of oxaloacetate decarboxylase; Biotin carboxyl carrier protein

ponent BL0695 of predicted ECF transporter
nal regulator of pyridoxine metabolism
etical protein

7.1.35)
1.26.4)
ransferase RlmI

idase (EC 3.2.1.86)
sferase (EC 2.5.1.75)
etical protein
rotein B
-dehydrogenase (EC 1.1.1.125)
ABC transporter, substrate binding periplasmic protein MalE

etical protein

.2)
ctive streptococcal antigen
a subunit (EC 2.7.7.7)
or, TetR family
otein
ylase
tein
D(P)H nitroreductase (EC 1.-.-.-) / Dihydropteridine reductase (EC 1.5.1.34)

n Se23.5 (Fragment)
etical protein
e M BlpM
ose reductase (EC 1.1.1.133)
n
ylase alpha chain (EC 4.1.1.3)

protein of oxaloacetate decarboxylase; Biotin carboxyl carrier protein
se RasP/YluC, implicated in cell division based on FtsL cleavage
ein
l protein - phage associated
factor Ts
on initiator protein DnaA
l protein - phage associated
etical protein
ecific recombinase
ursor (EC 4.2.2.1)

ductase protein F (EC 1.6.4.-)
omerase (EC 5.3.2.-); Xylose transport system permease protein xylH

phospholipid phosphatase
n DnaC
se precursor (EC 3.1.3.2)
in ComGE, FIG075573
ase transcriptional regulator NrdR
n
onal regulator, PadR family
dratase (EC 4.2.1.46)
cluster with TrmH family tRNA/rRNA methyltransferase YacO
tein
C 2.8.1.7)
oxypeptidase (EC 3.4.16.4)
ransferase (EC 2.1.2.1)
factor LepA
ltransferase (EC 2.7.7.41)
etical protein
1.26.8)
etical protein
1.20.4.1)
ate ligase (EC 6.3.4.3)
C 4.2.1.49)

ı 2 macroglobulin-binding protein (GRAB)
ı
vith transcription termination protein NusA
: 4.2.1.1)

etical protein
l activator - phage associated
ɔ

ı
(EC 1.8.1.7)
ɔsphate transporter
factor LepA
er, ATP-binding protein FtsE (TC 3.A.5.1.1)
ion protein
ı
tein
nthase B (EC 4.2.1.70)
lalanyl-D-glutamyl-2,6-diaminopimelate--D-alanyl-D-alanine ligase (EC 6.3.2.10)
nding protein
ain / putative murein endopeptidase
-phospho-L-lactate transferase like, CofD-like
licase UvrD/PcrA

nding protein
no-heptulosonate-7-phosphate synthase I beta (EC 2.5.1.54)

ı
etical protein
or, GntR family
ory protein MutS2
etical protein
inding/permease protein
otein capA
TP-binding protein
drolase, family 25

lhydrolase with GDSL-like motif
or in cluster with unspecified monosaccharide ABC transport system
etical protein

etical protein

lipase A2; _Toximoron (Other)
ein
sis protein LanM/pseudogene causing stop codon at 177th aa
upC
or, CrP/Fnr family
cted N-acetylneuraminate-binding protein
e (EC 2.7.6.5)
mily)
1.-)
etical protein
ptake protein TrkA
osphate decarboxylase
etical protein
etical protein
PDZ domain
pecific IIC component (EC 2.7.1.69)
ate-binding protein
etical protein
etical protein
etical protein
etical protein
n
n
ed protein gbs2121

etical protein
e aldolase (EC 4.1.2.4)
etical protein

ciated protein
hage associated)
or, TetR family
n resolvase
or
gase found clustered with an amidotransferase
cal protein
maturation GTPase RbgA (B. subtilis YlqF)

gma factor ComX
to have essential role in cell wall biosynthesis
n YihY, formerly thought to be RNase BN

n
carboxylase (EC 4.1.1.31)
l protein - phage associated
gid
iptional repressor MalR, LacI family
se I (EC 3.4.21.89)
n dnaD
(EC 2.4.2.3)

activating enzyme (EC 1.97.1.4)
e hydrolase (EC 3.1.2.6)
8)-N(6)/adenine(1519)-N(6))-dimethyltransferase (EC 2.1.1.182)
etical protein
lease YcfH
nding protein
.1)
kinase (EC 2.7.1.144)

er ATP binding/permease protein
nit A
3-epimerase (EC 5.1.3.-)

etical protein
(EC 6.1.1.15), bacterial type

d EpuA protein
gma factor ComX
p-alanine acetyltransferase (EC 2.3.1.-)
etical protein
nase
esistance protein
etical protein
A
tein
1.26.4)
or, Cro/CI family

(EC 6.3.2.12) / Folylpolyglutamate synthase (EC 6.3.2.17)
(EC 2.7.1.24)
e (EC 2.7.1.56)
UF901, similar to C-terminal domain of ribosome protection-type Tc-resistance proteins
d repair protein RecF
competence regulon ComD

etical protein

nit B (EC 5.99.1.-)
S14p (S29e) @ SSU ribosomal protein S14p (S29e), zinc-independent

osphate acyltransferase (EC 2.3.1.51)
etical protein

etical protein
ransferase (EC 2.4.2.9) / Pyrimidine operon regulatory protein PyrR
in ComGG, FIG068335

family protein

etical protein
al regulator-copper transport operon
C 2.8.1.7)
n

otional repressor protein
nase
N
e dehydrogenase [NAD] (EC 1.2.1.24); Succinate-semialdehyde dehydrogenase [NADP+] (EC 1.2.1.16)
h/MocA family
ABC transporter, permease protein MalF
P1558
er ATP-binding protein, spy1790 homolog
etical protein
e 4-epimerase (EC 5.1.3.4)
permease
n
ase family 5
e (EC 3.5.2.7)
or
1,2-L-rhamnosyltransferase (EC 2.4.1.-)
sX
resolvase
erase (EC 2.3.1.8)
e kinase (EC 2.7.4.6)
rotein
nit A (EC 5.99.1.-)
e (EC 2.3.1.16) @ Acetyl-CoA acetyltransferase (EC 2.3.1.9)

etical protein
inding/permease protein
loprotease superfamily
ylase beta chain (EC 4.1.1.3)
ron antiterminator, BglG family
e, zinc-containing

e YycJ/WalJ, required for cell wall metabolism and coordination of cell division with DNA replication
ase LtaS Type IIc
thionine synthetase (EC 2.5.1.6)
subunit SecG (TC 3.A.5.1.1)
etical protein
or of the fructose operon, DeoR family
enetic competence MecA
in ComGF, access of DNA to ComEA, FIG012620
fication system, restriction subunit R (EC 3.1.21.3)
ase (EC 2.7.1.-) (EC 2.7.4.-)
S2p (SAe)

atory protein ArgR, repressor of arg regulon
.3
cal protein
ease/phosphatase family protein
ponent PanT of predicted pantothenate ECF transporter
antigen A homolog
etical protein
netase (EC 6.3.4.4)
etical protein
(EC 4.1.3.6)
fication system specificity subunit
transport protein CorA
ransferase (EC 2.1.2.5) @ Glutamate formyltransferase
.6.1.1)
denylyltransferase (EC 2.7.7.3)
f exopolysaccharide synthesis, competence and biofilm formation Ftr, XRE family
1)

ylase

on protein NrdI

system, EIIA component (EC 2.7.1.-)
in ComEA, DNA receptor
protein
nding protein
thetase related
gase

hatase (EC 3.6.1.27)
C 2.7.1.33)
etical protein

licase RecG (EC 3.6.1.-)
tein

acid transport system carrier protein
X (EC 3.4.24.-)

se-specific IIA component (EC 2.7.1.69)

)

hase (EC 2.5.1.29)

.3)

59)

mponent (EC 2.7.1.69)

.1.69)

| genes | logFC.deltaSK | logCPM.deltaSK | statistic.deltaSK | Pvalue.deltaSK | FDR.deltaSK |
|---|---|---|---|---|---|
| AP53_1848 | -0.98213361 | 8.29535345 | 96.61224912 | 6.51E-07 | 0.001005338 |
| AP53_1867 | -1.891152939 | 7.390902448 | 88.23668512 | 1.04E-06 | 0.001005338 |
| AP53_848 | 3.940671781 | 7.052381792 | 76.21508327 | 2.17E-06 | 0.001113893 |
| AP53_847 | 3.621024648 | 6.940372562 | 75.39172206 | 2.30E-06 | 0.001113893 |
| AP53_1865 | 3.805377346 | 11.85379858 | 64.15092953 | 5.11E-06 | 0.001985184 |
| AP53_1851 | -1.054016136 | 9.03965394 | 59.40793183 | 7.44E-06 | 0.002406919 |
| AP53_916 | 2.053150604 | 8.567138716 | 51.17974511 | 1.52E-05 | 0.004219385 |
| AP53_1849 | -0.887717984 | 8.259439319 | 45.01805953 | 2.78E-05 | 0.006212426 |
| AP53_1713 | 2.90391415 | 10.57102058 | 44.66442051 | 2.88E-05 | 0.006212426 |
| AP53_1758 | 2.174679866 | 9.430769949 | 43.21311123 | 3.35E-05 | 0.006512044 |
| AP53_917 | 1.925970174 | 8.589404302 | 41.33716271 | 4.11E-05 | 0.007205485 |
| AP53_915 | 1.893125414 | 8.680433803 | 40.61566708 | 4.45E-05 | 0.007205485 |
| AP53_551 | 1.450069512 | 7.020067052 | 38.26644477 | 5.83E-05 | 0.008705873 |
| AP53_1428 | -0.957337289 | 7.56840856 | 36.12159123 | 7.54E-05 | 0.009381703 |
| AP53_1752 | 1.042633454 | 6.705251976 | 36.11853636 | 7.55E-05 | 0.009381703 |
| AP53_911 | 1.250884446 | 7.402583754 | 35.70595721 | 7.94E-05 | 0.009381703 |
| AP53_914 | 1.750545371 | 8.560464416 | 35.29940218 | 8.35E-05 | 0.009381703 |
| AP53_1722 | 0.907634323 | 9.069333087 | 34.18337895 | 9.62E-05 | 0.009381703 |
| AP53_1850 | -1.316789839 | 8.012850345 | 34.11094003 | 9.71E-05 | 0.009381703 |
| AP53_1751 | 2.185303365 | 6.490343622 | 33.81265084 | 0.00010089 | 0.009381703 |
| AP53_758 | -0.899120461 | 7.399991744 | 33.76589109 | 0.000101502 | 0.009381703 |
| AP53_1750 | 1.995401029 | 6.210996007 | 32.83344535 | 0.000114667 | 0.010100806 |
| AP53_550 | 1.203977951 | 7.078859446 | 32.02184593 | 0.000127785 | 0.010100806 |
| AP53_1757 | 1.59365545 | 8.184491381 | 31.99443287 | 0.000128258 | 0.010100806 |
| AP53_1423 | 1.242350669 | 8.381643768 | 31.88893399 | 0.000130098 | 0.010100806 |
| AP53_553 | 1.566729804 | 7.009642797 | 30.89208587 | 0.000149107 | 0.010727151 |
| AP53_694 | 0.621140487 | 8.383306483 | 30.88670082 | 0.000149218 | 0.010727151 |
| AP53_1422 | 1.078556103 | 6.883149016 | 30.42144851 | 0.000159208 | 0.010881088 |
| AP53_552 | 1.48011463 | 6.275588373 | 30.09450072 | 0.0001667 | 0.010881088 |
| AP53_1732 | 0.740224374 | 8.474699799 | 30.03207205 | 0.000168178 | 0.010881088 |
| AP53_908 | 1.117585792 | 5.781226351 | 29.77385231 | 0.000174454 | 0.010923078 |
| AP53_1004 | 0.61339595 | 9.335315697 | 28.98897878 | 0.000195295 | 0.011474647 |
| AP53_547 | 1.143119356 | 7.169960578 | 28.8865807 | 0.000198225 | 0.011474647 |
| AP53_1001 | 1.671509817 | 7.743620698 | 28.79128298 | 0.000200998 | 0.011474647 |

| | | | | | |
|---|---|---|---|---|---|
| AP53_849 | 1.818815836 | 6.509506297 | 28.11506799 | 0.000222043 | 0.012313892 |
| AP53_766 | -1.157092892 | 9.494367576 | 27.87960639 | 0.000229973 | 0.012399366 |
| AP53_1308 | 1.064583895 | 7.466925505 | 27.56431354 | 0.000241119 | 0.012648951 |
| AP53_1002 | 1.72011917 | 9.186568837 | 27.18312939 | 0.000255454 | 0.013048311 |
| AP53_1421 | 1.189570229 | 6.927983678 | 26.13316103 | 0.000300438 | 0.014310489 |
| AP53_1557 | 0.519505511 | 9.093260226 | 26.02521774 | 0.000305571 | 0.014310489 |
| AP53_548 | 1.263530236 | 7.536666595 | 26.02478528 | 0.000305592 | 0.014310489 |
| AP53_1753 | 1.134616356 | 5.306778081 | 25.94085267 | 0.000309655 | 0.014310489 |
| AP53_1716 | 0.554152218 | 7.147578025 | 25.5686841 | 0.000328456 | 0.014501791 |
| AP53_425 | -1.097371778 | 7.428724349 | 25.27496489 | 0.000344248 | 0.014501791 |
| AP53_832 | -1.329591328 | 9.839306377 | 25.06860212 | 0.000355878 | 0.014501791 |
| AP53_318 | 0.564028693 | 10.427855 | 25.02858801 | 0.000358186 | 0.014501791 |
| AP53_1082 | 1.225291131 | 5.178836739 | 24.80399959 | 0.000371475 | 0.014501791 |
| AP53_164 | 1.649450717 | 4.340799239 | 24.69912558 | 0.000377878 | 0.014501791 |
| AP53_1291 | 0.727289568 | 6.437852788 | 24.67039817 | 0.000379655 | 0.014501791 |
| AP53_1476 | -0.64374855 | 7.639481239 | 24.65469876 | 0.00038063 | 0.014501791 |
| AP53_1432 | 0.787601256 | 10.71659941 | 24.64816805 | 0.000381036 | 0.014501791 |
| AP53_912 | 1.153609916 | 6.658422814 | 24.04560506 | 0.000420841 | 0.015708693 |
| AP53_719 | 0.504763345 | 8.308501386 | 23.92036781 | 0.000429717 | 0.015737378 |
| AP53_1152 | 0.656317364 | 7.02683456 | 23.67321423 | 0.00044789 | 0.016099145 |
| AP53_1749 | 1.715539596 | 6.229693034 | 23.45156135 | 0.000464961 | 0.016408896 |
| AP53_549 | 1.080864847 | 6.651482013 | 22.90232402 | 0.000510672 | 0.017700259 |
| AP53_1565 | -1.54051783 | 7.738902507 | 22.71394299 | 0.000527555 | 0.017964646 |
| AP53_1748 | 1.929166499 | 6.709134603 | 22.01800148 | 0.000595899 | 0.01982005 |
| AP53_555 | 0.740944062 | 8.73793431 | 21.94187778 | 0.000603989 | 0.01982005 |
| AP53_1838 | -0.81845753 | 6.845123703 | 21.86147793 | 0.000612675 | 0.01982005 |
| AP53_1424 | 0.645199023 | 7.751467306 | 21.39971012 | 0.000665509 | 0.020898149 |
| AP53_621 | -1.106903661 | 3.43401959 | 21.38287108 | 0.000667535 | 0.020898149 |
| AP53_300 | 1.561537078 | 8.145165313 | 20.96538734 | 0.000720164 | 0.022187902 |
| AP53_546 | 1.092466887 | 7.330619277 | 20.78120136 | 0.000744927 | 0.022285841 |
| AP53_1692 | -0.716029769 | 11.16790356 | 20.77116175 | 0.000746306 | 0.022285841 |
| AP53_1309 | 1.571492477 | 6.652699164 | 20.67776167 | 0.000759274 | 0.02232957 |
| AP53_1448 | 0.531134365 | 8.01963308 | 20.51047321 | 0.000783172 | 0.02268861 |
| AP53_523 | 1.30818553 | 9.522155055 | 20.37747338 | 0.000802805 | 0.022791766 |
| AP53_1226 | 0.408726704 | 8.30245655 | 20.32825586 | 0.000810217 | 0.022791766 |

| | | | | | |
|---|---|---|---|---|---|
| AP53_1420 | 1.270607774 | 7.217537269 | 20.0138557 | 0.000859514 | 0.023833103 |
| AP53_234 | -0.513442507 | 8.561751269 | 19.63852954 | 0.000923067 | 0.024492137 |
| AP53_1559 | 1.308185602 | 11.45855556 | 19.62398056 | 0.00092564 | 0.024492137 |
| AP53_1425 | 0.594207406 | 7.44292707 | 19.61351548 | 0.000927496 | 0.024492137 |
| AP53_58 | -0.825921002 | 10.54885701 | 19.57840274 | 0.000933755 | 0.024492137 |
| AP53_40 | 1.48843863 | 10.27616672 | 19.40226468 | 0.000965914 | 0.024997866 |
| AP53_1083 | 1.599237226 | 4.977482168 | 19.19023928 | 0.001006374 | 0.025702254 |
| AP53_1430 | -0.840599172 | 7.962175931 | 18.89066266 | 0.001066996 | 0.026114206 |
| AP53_324 | 1.68073769 | 10.21515832 | 18.84708734 | 0.001076168 | 0.026114206 |
| AP53_47 | -0.811884462 | 10.83978006 | 18.79915799 | 0.001086364 | 0.026114206 |
| AP53_156 | 1.479055935 | 10.26429078 | 18.79730246 | 0.001086761 | 0.026114206 |
| AP53_1535 | 1.036776733 | 10.61596999 | 18.74326771 | 0.001098398 | 0.026114206 |
| AP53_1236 | -0.719402099 | 7.985314977 | 18.68647411 | 0.001110787 | 0.026114206 |
| AP53_368 | 0.98833567 | 9.824903804 | 18.61658659 | 0.00112626 | 0.026114206 |
| AP53_1296 | 0.616659985 | 6.172112517 | 18.59925721 | 0.001130136 | 0.026114206 |
| AP53_1744 | 1.602123668 | 5.749783198 | 18.1782076 | 0.00122932 | 0.027787402 |
| AP53_1084 | 1.347869638 | 5.287488837 | 18.17070551 | 0.001231178 | 0.027787402 |
| AP53_612 | -1.440259489 | 4.914644896 | 18.02890304 | 0.001266931 | 0.028265669 |
| AP53_662 | 0.673829136 | 8.236886967 | 17.89843592 | 0.001300912 | 0.028693969 |
| AP53_1746 | 1.668747513 | 6.287957149 | 17.77065454 | 0.001335239 | 0.029041575 |
| AP53_913 | 0.826877078 | 7.533091996 | 17.72922921 | 0.001346595 | 0.029041575 |
| AP53_905 | 0.772233293 | 5.511271964 | 17.46368418 | 0.001422161 | 0.030327479 |
| AP53_826 | 1.513326017 | 8.443236576 | 17.39856256 | 0.001441449 | 0.030327479 |
| AP53_1672 | 0.800122266 | 7.82243254 | 17.3597424 | 0.001453094 | 0.030327479 |
| AP53_827 | 0.872995219 | 8.427106352 | 17.1735203 | 0.00151052 | 0.031190632 |
| AP53_508 | 0.565775032 | 5.896771962 | 17.11365026 | 0.001529548 | 0.03125108 |
| AP53_1295 | 0.751673884 | 7.199351983 | 17.05293313 | 0.001549134 | 0.031321561 |
| AP53_865 | 0.580021516 | 7.424202608 | 16.97348826 | 0.001575211 | 0.031520451 |
| AP53_747 | 1.152443559 | 9.584863311 | 16.56927669 | 0.001716182 | 0.033990911 |
| AP53_1735 | 0.813447876 | 7.533997899 | 16.47840725 | 0.001749891 | 0.034015399 |
| AP53_1300 | 0.760868028 | 9.157013082 | 16.46173799 | 0.001756159 | 0.034015399 |
| AP53_1564 | -0.622290158 | 8.421111977 | 16.42519715 | 0.001769992 | 0.034015399 |
| AP53_978 | 0.947253641 | 13.63851091 | 16.1348351 | 0.001884605 | 0.035862916 |
| AP53_1747 | 1.720313425 | 6.135558656 | 16.07621468 | 0.001908793 | 0.035970557 |
| AP53_516 | 0.447372345 | 6.978985836 | 15.98825567 | 0.001945778 | 0.036314954 |

| | | | | | |
|---|---|---|---|---|---|
| AP53_17 | 0.713236994 | 9.016055634 | 15.9343129 | 0.001968878 | 0.036396118 |
| AP53_977 | 0.803711817 | 13.251696 | 15.76047378 | 0.002045554 | 0.037456795 |
| AP53_1478 | -0.789115017 | 10.06615235 | 15.63484187 | 0.002103164 | 0.038151794 |
| AP53_1626 | 1.311057157 | 7.678876515 | 15.5005905 | 0.002166855 | 0.038943207 |
| AP53_1736 | 0.52144668 | 13.53520806 | 15.4449954 | 0.002193895 | 0.039067437 |
| AP53_887 | -0.718855853 | 10.0984361 | 15.26595975 | 0.002283719 | 0.040009597 |
| AP53_1481 | -1.009896034 | 10.0964202 | 15.25757728 | 0.002288029 | 0.040009597 |
| AP53_1648 | -0.836398685 | 11.11452243 | 15.07366212 | 0.002385072 | 0.041034778 |
| AP53_1743 | 1.301230461 | 10.51698796 | 15.03265245 | 0.002407368 | 0.041034778 |
| AP53_1538 | 0.682533054 | 8.55751471 | 15.02769454 | 0.00241008 | 0.041034778 |
| AP53_1419 | 0.600013889 | 9.811227587 | 14.92568899 | 0.002466688 | 0.04140287 |
| AP53_975 | -0.657835206 | 5.218803085 | 14.91207793 | 0.00247436 | 0.04140287 |
| AP53_1126 | 0.479821865 | 8.692747408 | 14.80087159 | 0.002538106 | 0.042106523 |
| AP53_66 | -1.017069711 | 10.13338686 | 14.71495968 | 0.002588682 | 0.042581621 |
| AP53_693 | 0.62561052 | 9.317769129 | 14.60241209 | 0.002656749 | 0.043334039 |
| AP53_500 | -0.372461911 | 8.026905762 | 14.50343155 | 0.002718364 | 0.043825493 |
| AP53_1218 | 0.418652556 | 8.972107627 | 14.48182399 | 0.002732037 | 0.043825493 |
| AP53_586 | 0.472308067 | 10.17786669 | 14.36377211 | 0.002808193 | 0.044307971 |
| AP53_121 | -0.520569958 | 6.021714868 | 14.34685392 | 0.002819311 | 0.044307971 |
| AP53_1480 | -0.775883553 | 9.729546962 | 14.32976145 | 0.002830597 | 0.044307971 |
| AP53_1228 | 0.686617845 | 10.22004434 | 14.27165425 | 0.002869363 | 0.044409042 |
| AP53_1645 | 1.471518043 | 6.523461692 | 14.22403108 | 0.002901604 | 0.044409042 |
| AP53_1839 | -0.482327419 | 6.918112071 | 14.21803643 | 0.002905692 | 0.044409042 |
| AP53_48 | -0.822441122 | 11.44172855 | 14.16108426 | 0.002944876 | 0.044656287 |
| AP53_165 | 1.16074344 | 2.954387678 | 14.04265766 | 0.003028371 | 0.045566424 |
| AP53_1175 | 0.388004731 | 8.466828996 | 13.99070716 | 0.003065878 | 0.045775923 |
| AP53_501 | -0.475992854 | 8.033709296 | 13.83553329 | 0.003181221 | 0.047135493 |
| AP53_347 | 0.429015383 | 7.310843593 | 13.78284133 | 0.003221548 | 0.047371392 |
| AP53_588 | 0.491168263 | 8.92811689 | 13.61159663 | 0.003356839 | 0.048989662 |
| AP53_167 | -0.784338438 | 6.615908403 | 13.55190411 | 0.003405569 | 0.049119702 |
| AP53_952 | 1.261627527 | 7.544188929 | 13.53881868 | 0.003416363 | 0.049119702 |
| AP53_1745 | 1.406434394 | 5.866330514 | 13.49808078 | 0.003450225 | 0.049218942 |
| AP53_1094 | 1.417306846 | 4.265203367 | 13.46978351 | 0.00347398 | 0.049218942 |
| AP53_474 | 0.553449333 | 9.129672143 | 13.37634972 | 0.003553799 | 0.049984956 |
| AP53_177 | -0.571342083 | 8.91860337 | 13.30628422 | 0.003615078 | 0.05048106 |

| | | | | | |
|---|---|---|---|---|---|
| AP53_1631 | -0.737268116 | 11.02108569 | 13.22565143 | 0.003687151 | 0.05108937 |
| AP53_556 | 0.514323753 | 9.196164744 | 13.1990686 | 0.003711283 | 0.05108937 |
| AP53_942 | 1.020811498 | 9.17685393 | 13.03182861 | 0.003867457 | 0.052848179 |
| AP53_904 | 1.031070397 | 5.773598337 | 13.00471398 | 0.003893503 | 0.052848179 |
| AP53_118 | -0.480995508 | 7.703854256 | 12.83991954 | 0.004056335 | 0.054676019 |
| AP53_1134 | -0.759485128 | 8.551652088 | 12.77316736 | 0.004124574 | 0.055212402 |
| AP53_607 | -0.90018732 | 5.266162481 | 12.74298247 | 0.004155876 | 0.05525038 |
| AP53_1402 | -0.440295135 | 7.975794128 | 12.66294692 | 0.004240242 | 0.055988506 |
| AP53_910 | 0.899203635 | 7.052291359 | 12.6310061 | 0.004274475 | 0.056059168 |
| AP53_805 | -0.668220315 | 4.94977484 | 12.54853045 | 0.004364392 | 0.056854258 |
| AP53_1380 | 0.571589161 | 8.52007152 | 12.38334213 | 0.004551297 | 0.058579803 |
| AP53_1267 | -0.975400105 | 11.23135134 | 12.37824477 | 0.004557213 | 0.058579803 |
| AP53_1154 | 0.741226959 | 6.87505983 | 12.33800093 | 0.004604243 | 0.058794972 |
| AP53_1086 | 0.811824977 | 4.62286242 | 12.31180572 | 0.004635164 | 0.058802962 |
| AP53_818 | 0.351616431 | 9.40215327 | 12.22901395 | 0.004734518 | 0.059673374 |
| AP53_1707 | -0.715205096 | 11.49237008 | 12.13863397 | 0.004845867 | 0.060682756 |
| AP53_rna60 | 1.100132325 | 4.476833431 | 12.04489847 | 0.004964635 | 0.06177152 |
| AP53_941 | 0.893949909 | 8.638612252 | 11.93436688 | 0.005109137 | 0.063058175 |
| AP53_1093 | 1.325117361 | 2.754249981 | 11.90799766 | 0.005144341 | 0.063058175 |
| AP53_199 | 0.546265167 | 6.130953473 | 11.89224601 | 0.005165507 | 0.063058175 |
| AP53_155 | 0.753434605 | 9.707554236 | 11.69257616 | 0.005442947 | 0.065102553 |
| AP53_909 | 0.592372137 | 5.601753535 | 11.68637404 | 0.005451843 | 0.065102553 |
| AP53_rna36 | 1.629065385 | 4.138285412 | 11.65998922 | 0.005489882 | 0.065102553 |
| AP53_1841 | 0.541947487 | 7.681483501 | 11.62622236 | 0.005539022 | 0.065102553 |
| AP53_1331 | 0.772404956 | 7.374790994 | 11.5962498 | 0.005583076 | 0.065102553 |
| AP53_1011 | -0.856087022 | 5.389861213 | 11.59284631 | 0.005588105 | 0.065102553 |
| AP53_325 | 1.502128437 | 6.18675495 | 11.5827374 | 0.005603073 | 0.065102553 |
| AP53_1605 | -0.418963623 | 7.544310561 | 11.5781953 | 0.005609813 | 0.065102553 |
| AP53_54 | -0.675341182 | 11.32762119 | 11.56138638 | 0.005634842 | 0.065102553 |
| AP53_63 | -0.557521998 | 10.43719894 | 11.49500956 | 0.005734978 | 0.065867412 |
| AP53_866 | 0.929602768 | 10.61348664 | 11.44670064 | 0.005809182 | 0.066327188 |
| AP53_61 | -0.732308409 | 10.31748478 | 11.4219723 | 0.005847604 | 0.06637544 |
| AP53_949 | 0.897085423 | 7.838170168 | 11.36850394 | 0.005931716 | 0.066938722 |
| AP53_953 | 0.693761561 | 5.365556629 | 11.34546638 | 0.005968397 | 0.066963344 |
| AP53_rna79 | 1.628969583 | 3.888713338 | 11.24641802 | 0.006129195 | 0.068372229 |

| | | | | | |
|---|---|---|---|---|---|
| AP53_117 | -0.40591647 | 7.004833486 | 11.16166413 | 0.00627087 | 0.068929159 |
| AP53_193 | -0.448815271 | 6.180584908 | 11.12555808 | 0.0063324 | 0.068929159 |
| AP53_1675 | -0.543703168 | 5.85093154 | 11.11347493 | 0.006353151 | 0.068929159 |
| AP53_867 | 0.495006225 | 6.75101341 | 11.11248292 | 0.006354858 | 0.068929159 |
| AP53_1051 | 1.141924914 | 8.48216266 | 11.11142351 | 0.006356682 | 0.068929159 |
| AP53_1644 | 1.014033483 | 6.060227028 | 11.06768484 | 0.006432522 | 0.069106321 |
| AP53_967 | 0.982063823 | 9.252986656 | 11.06098868 | 0.006444227 | 0.069106321 |
| AP53_1804 | 0.416693937 | 6.605851342 | 11.00002058 | 0.006551968 | 0.069416662 |
| AP53_1088 | 0.62406884 | 4.336772951 | 10.99995848 | 0.006552079 | 0.069416662 |
| AP53_148 | -0.943323524 | 6.799100627 | 10.97601086 | 0.006594981 | 0.069416662 |
| AP53_122 | -0.520573665 | 6.032187503 | 10.96199464 | 0.006620246 | 0.069416662 |
| AP53_1494 | 0.367732092 | 9.042313129 | 10.94128787 | 0.00665778 | 0.069416662 |
| AP53_380 | -0.822938262 | 5.255508454 | 10.92485527 | 0.006687746 | 0.069416662 |
| AP53_1370 | -0.516528753 | 7.087991316 | 10.87639531 | 0.006777051 | 0.069481311 |
| AP53_pseudo595 | -0.505017629 | 6.325910274 | 10.85749437 | 0.006812265 | 0.069481311 |
| AP53_491 | -0.402412354 | 7.708053578 | 10.84935409 | 0.006827498 | 0.069481311 |
| AP53_249 | 0.625920778 | 8.951446759 | 10.84420107 | 0.006837162 | 0.069481311 |
| AP53_299 | 0.334987613 | 8.536673103 | 10.8129275 | 0.006896158 | 0.069715847 |
| AP53_633 | -0.719268951 | 4.286017489 | 10.76020751 | 0.006996983 | 0.070368618 |
| AP53_611 | -1.104273583 | 4.925121625 | 10.72020763 | 0.007074646 | 0.070592506 |
| AP53_447 | 0.790805247 | 4.439548682 | 10.71134955 | 0.007091983 | 0.070592506 |
| AP53_667 | -0.715678297 | 10.64619676 | 10.67205243 | 0.007169502 | 0.071000022 |
| AP53_1697 | 0.437690333 | 8.455477189 | 10.55966886 | 0.007396797 | 0.072691896 |
| AP53_697 | 0.447766765 | 7.56475385 | 10.52065851 | 0.007477683 | 0.072691896 |
| AP53_637 | -0.597357691 | 4.800871695 | 10.50558721 | 0.007509213 | 0.072691896 |
| AP53_379 | -1.030599763 | 4.715037934 | 10.4972051 | 0.007526817 | 0.072691896 |
| AP53_851 | 1.141450659 | 6.138214621 | 10.49683275 | 0.0075276 | 0.072691896 |
| AP53_1371 | -0.722776623 | 10.82174338 | 10.4087795 | 0.007715535 | 0.07337028 |
| AP53_902 | 0.585708074 | 4.822374628 | 10.40516385 | 0.00772337 | 0.07337028 |
| AP53_509 | 0.51599097 | 5.783510146 | 10.40050318 | 0.007733484 | 0.07337028 |
| AP53_741 | 0.61262468 | 9.877237239 | 10.39334342 | 0.007749051 | 0.07337028 |
| AP53_259 | -0.749171445 | 7.002333297 | 10.35855133 | 0.007825228 | 0.073641856 |
| AP53_1633 | 0.570650785 | 6.935103688 | 10.29393787 | 0.007969065 | 0.073641856 |
| AP53_1445 | 0.371560401 | 8.853964934 | 10.22425712 | 0.008127707 | 0.073641856 |
| AP53_468 | -0.388244262 | 8.330445525 | 10.21763854 | 0.008142969 | 0.073641856 |

| | | | | | |
|---|---|---|---|---|---|
| AP53_1483 | -0.683280791 | 9.419940924 | 10.19857669 | 0.008187115 | 0.073641856 |
| AP53_1609 | 0.354410465 | 7.566110226 | 10.17048063 | 0.008252702 | 0.073641856 |
| AP53_725 | 0.556533013 | 7.957002543 | 10.16541492 | 0.008264593 | 0.073641856 |
| AP53_654 | 0.449368726 | 8.648977839 | 10.15473088 | 0.00828974 | 0.073641856 |
| AP53_1250 | 0.5198703 | 8.688805681 | 10.14818715 | 0.008305186 | 0.073641856 |
| AP53_1733 | 0.490691302 | 7.51982531 | 10.1335393 | 0.008339886 | 0.073641856 |
| AP53_207 | 0.404705522 | 7.12594599 | 10.11961628 | 0.008373029 | 0.073641856 |
| AP53_123 | -0.461253912 | 5.959595513 | 10.11745288 | 0.008378193 | 0.073641856 |
| AP53_1310 | 0.490795915 | 8.517374264 | 10.11625826 | 0.008381046 | 0.073641856 |
| AP53_1081 | 1.162599488 | 4.303329124 | 10.11320028 | 0.008388354 | 0.073641856 |
| AP53_1634 | 0.868621459 | 8.437611102 | 10.10815721 | 0.008400423 | 0.073641856 |
| AP53_492 | -0.538339442 | 7.705728591 | 10.0772679 | 0.008474799 | 0.073641856 |
| AP53_1374 | 0.408234179 | 8.160834778 | 10.07085482 | 0.008490338 | 0.073641856 |
| AP53_127 | -0.424689161 | 6.639887807 | 10.05626224 | 0.008525822 | 0.073641856 |
| AP53_rna71 | 1.67561492 | 3.871305666 | 10.04375414 | 0.008556378 | 0.073641856 |
| AP53_55 | -0.569160926 | 10.94485035 | 10.04171021 | 0.008561383 | 0.073641856 |
| AP53_1444 | 0.31282008 | 9.09676772 | 10.00305869 | 0.008656693 | 0.073641856 |
| AP53_473 | 0.640570883 | 9.506031141 | 9.99236435 | 0.008683286 | 0.073641856 |
| AP53_1862 | 0.896137469 | 8.378678594 | 9.972181173 | 0.008733737 | 0.073641856 |
| AP53_1536 | 0.437917191 | 11.00225356 | 9.968575099 | 0.008742788 | 0.073641856 |
| AP53_146 | 0.596968025 | 6.035456131 | 9.951654276 | 0.008785405 | 0.073641856 |
| AP53_41 | 0.39980937 | 7.04652198 | 9.950136376 | 0.00878924 | 0.073641856 |
| AP53_618 | -1.12479531 | 4.72583804 | 9.942160233 | 0.008809425 | 0.073641856 |
| AP53_815 | 0.501450236 | 8.798029072 | 9.930096728 | 0.008840058 | 0.073641856 |
| AP53_1493 | 0.649956165 | 13.43323349 | 9.890547813 | 0.008941374 | 0.073736571 |
| AP53_39 | 0.523269654 | 10.17853263 | 9.882608489 | 0.008961878 | 0.073736571 |
| AP53_1708 | 0.757645777 | 6.4287874 | 9.881249034 | 0.008965395 | 0.073736571 |
| AP53_1287 | 0.44297077 | 8.933424055 | 9.834385757 | 0.009087624 | 0.07442649 |
| AP53_1734 | 0.684876812 | 7.484039976 | 9.816125944 | 0.009135784 | 0.074506537 |
| AP53_573 | -0.583246696 | 6.896267721 | 9.749075042 | 0.009315243 | 0.075652242 |
| AP53_53 | -0.808039224 | 11.43545424 | 9.733327837 | 0.009357993 | 0.075682772 |
| AP53_56 | -0.665843048 | 10.64711432 | 9.707082886 | 0.009429762 | 0.075946754 |
| AP53_619 | -1.273147712 | 5.091305753 | 9.67088072 | 0.009529833 | 0.07643556 |
| AP53_620 | -1.114681501 | 4.71110218 | 9.579291595 | 0.009788682 | 0.078188608 |
| AP53_817 | 0.337705402 | 9.179772291 | 9.550576154 | 0.009871548 | 0.078527358 |

| | | | | | |
|---|---|---|---|---|---|
| AP53_1087 | 0.625261286 | 4.395822584 | 9.51732386 | 0.009968549 | 0.078840395 |
| AP53_pseudo812 | 0.675849304 | 8.76666172 | 9.509295883 | 0.009992137 | 0.078840395 |
| AP53_1531 | 0.574580602 | 9.761708956 | 9.46595447 | 0.010120631 | 0.079530949 |
| AP53_361 | 0.552512471 | 12.21068719 | 9.430044242 | 0.010228581 | 0.08005514 |
| AP53_882 | -0.549343939 | 8.88071783 | 9.400326139 | 0.01031895 | 0.080438081 |
| AP53_1597 | -0.336755245 | 8.736347993 | 9.385172355 | 0.010365396 | 0.080476932 |
| AP53_265 | 0.76660604 | 9.95527985 | 9.32868927 | 0.010540712 | 0.080583195 |
| AP53_576 | 0.59692017 | 6.351748009 | 9.32263893 | 0.0105597 | 0.080583195 |
| AP53_1080 | -0.533877704 | 7.621737499 | 9.322601426 | 0.010559817 | 0.080583195 |
| AP53_1311 | 0.641024173 | 8.270877675 | 9.320063266 | 0.010567795 | 0.080583195 |
| AP53_1252 | 0.427856685 | 12.92724852 | 9.311364465 | 0.01059519 | 0.080583195 |
| AP53_979 | 0.772268926 | 13.36863302 | 9.300924208 | 0.01062818 | 0.080583195 |
| AP53_919 | 0.401909467 | 9.876042395 | 9.264373809 | 0.010744641 | 0.0810315 |
| AP53_173 | -0.563003646 | 6.451482905 | 9.256228091 | 0.010770802 | 0.0810315 |
| AP53_rna15 | 1.264244462 | 3.977905366 | 9.228911482 | 0.010859085 | 0.081380251 |
| AP53_613 | -1.177214886 | 5.300012505 | 9.171945306 | 0.01104597 | 0.082462414 |
| AP53_1357 | -1.022724896 | 4.280988049 | 9.152836188 | 0.011109513 | 0.082603496 |
| AP53_617 | -1.200586926 | 4.683457916 | 9.13995676 | 0.011152585 | 0.082603496 |
| AP53_1861 | 0.352655152 | 7.569565934 | 9.128062302 | 0.01119254 | 0.082603496 |
| AP53_1142 | -0.60981324 | 8.075777564 | 9.071229252 | 0.011385808 | 0.083698372 |
| AP53_381 | -0.535533767 | 5.405779303 | 9.055767649 | 0.011439071 | 0.083698372 |
| AP53_1482 | -0.858397462 | 10.11432222 | 9.031437555 | 0.011523484 | 0.083698372 |
| AP53_863 | 0.678543517 | 7.861865308 | 9.022977169 | 0.011553011 | 0.083698372 |
| AP53_1293 | 0.440414167 | 5.977682409 | 9.021979525 | 0.011556499 | 0.083698372 |
| AP53_981 | 0.660572282 | 12.5374196 | 8.998798135 | 0.011637892 | 0.083974527 |
| AP53_33 | 0.430472294 | 9.672083697 | 8.979072073 | 0.011707689 | 0.084165275 |
| AP53_1595 | -0.347550904 | 9.256043471 | 8.946871899 | 0.011822694 | 0.084286437 |
| AP53_896 | -0.587152865 | 9.148982694 | 8.941891285 | 0.011840603 | 0.084286437 |
| AP53_202 | 0.523756523 | 7.286735702 | 8.937944689 | 0.011854816 | 0.084286437 |
| AP53_589 | 0.451177922 | 6.987027995 | 8.884473937 | 0.012049395 | 0.085357209 |
| AP53_1079 | -0.565194412 | 6.775970913 | 8.864206757 | 0.012124136 | 0.085574357 |
| AP53_1639 | -0.470392809 | 7.323090863 | 8.813883606 | 0.012312111 | 0.086549531 |
| AP53_718 | 0.288428246 | 7.903787725 | 8.803459979 | 0.012351479 | 0.086549531 |
| AP53_1715 | 0.444904067 | 6.006493938 | 8.783598488 | 0.012426905 | 0.086660031 |
| AP53_831 | -1.04835057 | 9.968418741 | 8.771772943 | 0.012472074 | 0.086660031 |

| | | | | | |
|---|---|---|---|---|---|
| AP53_1681 | -0.416326606 | 8.463565769 | 8.757609575 | 0.012526429 | 0.086660031 |
| AP53_1579 | 0.496368225 | 8.787446596 | 8.752570334 | 0.012545837 | 0.086660031 |
| AP53_899 | -0.368547323 | 8.193042399 | 8.73554792 | 0.012611658 | 0.086805773 |
| AP53_224 | -0.413484195 | 8.276251153 | 8.70864817 | 0.01271651 | 0.087218186 |
| AP53_742 | 0.718511554 | 10.24984754 | 8.687489001 | 0.012799715 | 0.087479741 |
| AP53_610 | -0.850294253 | 4.371625699 | 8.653932107 | 0.012933001 | 0.0878572 |
| AP53_176 | -0.757621252 | 5.921850499 | 8.650813615 | 0.012945471 | 0.0878572 |
| AP53_1107 | -0.375302296 | 7.126780371 | 8.609101601 | 0.013113647 | 0.088688464 |
| AP53_922 | 0.321331661 | 9.687420644 | 8.530532945 | 0.013437528 | 0.090563336 |
| AP53_861 | 0.432218527 | 8.722673962 | 8.48315361 | 0.013637432 | 0.091219464 |
| AP53_676 | -0.698578155 | 8.098908033 | 8.474470832 | 0.013674449 | 0.091219464 |
| AP53_969 | 0.466516236 | 8.451923842 | 8.474137996 | 0.01367587 | 0.091219464 |
| AP53_57 | -0.62523473 | 10.59405612 | 8.437410855 | 0.013833793 | 0.091746313 |
| AP53_454 | 0.411731855 | 9.919725063 | 8.433810237 | 0.013849392 | 0.091746313 |
| AP53_1845 | -0.309280734 | 8.383367852 | 8.380598099 | 0.014082374 | 0.092972406 |
| AP53_457 | 1.513407876 | 2.788920823 | 8.36171206 | 0.014166182 | 0.093208673 |
| AP53_616 | -1.042799128 | 5.136891904 | 8.303291577 | 0.014429205 | 0.094355278 |
| AP53_858 | 0.330013953 | 9.390154202 | 8.300847042 | 0.014440337 | 0.094355278 |
| AP53_263 | -0.282646586 | 6.84634355 | 8.290781375 | 0.014486282 | 0.094355278 |
| AP53_120 | -0.428158585 | 5.734036467 | 8.254373616 | 0.014653922 | 0.095127968 |
| AP53_1393 | 0.309021395 | 10.5883986 | 8.214140949 | 0.014841864 | 0.095798539 |
| AP53_235 | -0.38845885 | 8.612244144 | 8.211154283 | 0.01485593 | 0.095798539 |
| AP53_947 | 0.74102827 | 8.593368411 | 8.171815463 | 0.015042685 | 0.096681628 |
| AP53_327 | -0.367329685 | 8.892621413 | 8.100954102 | 0.015386182 | 0.097956099 |
| AP53_502 | -0.436994278 | 6.893721111 | 8.100084714 | 0.015390454 | 0.097956099 |
| AP53_1220 | 0.285529429 | 8.806395054 | 8.099692849 | 0.01539238 | 0.097956099 |
| AP53_929 | -0.384634919 | 6.614404055 | 8.058144481 | 0.015598222 | 0.098941661 |
| AP53_1008 | -0.868864194 | 5.425898908 | 8.024283141 | 0.015768405 | 0.099635133 |
| AP53_575 | 0.40996759 | 5.702429986 | 8.016031261 | 0.015810212 | 0.099635133 |
| AP53_1474 | 0.577413942 | 6.336304883 | 7.980798103 | 0.015990206 | 0.100119347 |
| AP53_1429 | -0.496766874 | 7.988638528 | 7.967070732 | 0.016060994 | 0.100119347 |
| AP53_1443 | 0.30119823 | 8.795064872 | 7.963355637 | 0.016080216 | 0.100119347 |
| AP53_rna82 | 1.696755941 | 2.664243267 | 7.947554436 | 0.016162277 | 0.100119347 |
| AP53_932 | -0.60722465 | 5.974249375 | 7.937957599 | 0.01621236 | 0.100119347 |
| AP53_1260 | 0.388441584 | 8.863973916 | 7.931831456 | 0.016244427 | 0.100119347 |

| | | | | | |
|---|---|---|---|---|---|
| AP53_948 | 0.708259458 | 8.363698522 | 7.93112746 | 0.016248117 | 0.100119347 |
| AP53_880 | -0.293568902 | 8.829390239 | 7.921162959 | 0.016300451 | 0.100123971 |
| AP53_1268 | 0.518297954 | 5.460772353 | 7.901894828 | 0.016402218 | 0.100431248 |
| AP53_716 | 0.948906191 | 9.254855066 | 7.877964907 | 0.016529662 | 0.100724903 |
| AP53_1514 | -0.33646016 | 7.701027762 | 7.873426745 | 0.016553964 | 0.100724903 |
| AP53_886 | 0.516275463 | 11.36905956 | 7.840011432 | 0.016734217 | 0.10139103 |
| AP53_1132 | -0.687631897 | 9.079262359 | 7.825586295 | 0.016812753 | 0.10139103 |
| AP53_466 | -0.523905295 | 6.612090354 | 7.82423166 | 0.01682015 | 0.10139103 |
| AP53_1479 | -0.617349775 | 10.1080263 | 7.794070534 | 0.016985867 | 0.101745999 |
| AP53_972 | -0.317865604 | 8.01282363 | 7.788224419 | 0.017018212 | 0.101745999 |
| AP53_497 | -0.463891661 | 10.59102922 | 7.784961724 | 0.017036296 | 0.101745999 |
| AP53_884 | -0.489203586 | 8.408772709 | 7.76207455 | 0.017163794 | 0.10201149 |
| AP53_1617 | -0.364312145 | 6.251830487 | 7.755648929 | 0.017199793 | 0.10201149 |
| AP53_881 | -0.41421658 | 9.178826799 | 7.748772498 | 0.017238418 | 0.10201149 |
| AP53_260 | -0.750978777 | 7.925695479 | 7.716493694 | 0.017421112 | 0.102673304 |
| AP53_12 | 0.669897103 | 7.130205953 | 7.71036699 | 0.017456049 | 0.102673304 |
| AP53_1723 | 0.380722258 | 8.835087518 | 7.690062419 | 0.017572429 | 0.103045575 |
| AP53_1203 | -0.593572324 | 3.052125253 | 7.672199571 | 0.017675579 | 0.103338251 |
| AP53_1265 | -0.758824061 | 12.06436896 | 7.619218288 | 0.017985791 | 0.104674825 |
| AP53_195 | -0.408561775 | 6.691933789 | 7.606384315 | 0.018061908 | 0.104674825 |
| AP53_1835 | -0.312534348 | 7.517769788 | 7.598438873 | 0.018109224 | 0.104674825 |
| AP53_1772 | 0.790389578 | 11.24275464 | 7.593060175 | 0.018141338 | 0.104674825 |
| AP53_1431 | 0.292156803 | 9.994209097 | 7.587628831 | 0.018173836 | 0.104674825 |
| AP53_609 | -0.751639002 | 4.442814656 | 7.569871708 | 0.018280569 | 0.104978059 |
| AP53_1030 | -0.706282438 | 4.945597315 | 7.560501626 | 0.01833719 | 0.104992585 |
| AP53_1671 | 0.557262234 | 6.337641092 | 7.540413588 | 0.018459284 | 0.105380795 |
| AP53_669 | -0.400965704 | 8.124166179 | 7.530421521 | 0.018520376 | 0.105393968 |
| AP53_1290 | 0.300095065 | 6.889385727 | 7.520645572 | 0.018580379 | 0.105393968 |
| AP53_60 | -0.561832402 | 10.31908623 | 7.497895353 | 0.018720916 | 0.105393968 |
| AP53_943 | -0.390535214 | 7.179684337 | 7.490172394 | 0.018768911 | 0.105393968 |
| AP53_587 | 0.310010174 | 9.35155574 | 7.489152664 | 0.018775259 | 0.105393968 |
| AP53_124 | -0.508604358 | 5.968741084 | 7.487206141 | 0.018787384 | 0.105393968 |
| AP53_421 | 0.284965044 | 8.761816951 | 7.478362607 | 0.018842589 | 0.105399033 |
| AP53_710 | 0.446701247 | 7.45928674 | 7.45263728 | 0.019004277 | 0.105513315 |
| AP53_62 | -0.456377746 | 10.56937826 | 7.44309041 | 0.019064701 | 0.105513315 |

| | | | | | |
|---|---|---|---|---|---|
| AP53_1742 | 0.867215802 | 10.55339526 | 7.440335433 | 0.01908218 | 0.105513315 |
| AP53_1254 | 0.221113347 | 8.539635484 | 7.436727586 | 0.019105099 | 0.105513315 |
| AP53_590 | 0.324703522 | 7.752761326 | 7.430240791 | 0.01914639 | 0.105513315 |
| AP53_965 | 0.407838942 | 6.971739361 | 7.42123987 | 0.01920386 | 0.105513315 |
| AP53_1224 | 0.297608852 | 8.260536247 | 7.412878811 | 0.01925743 | 0.105513315 |
| AP53_1023 | -1.131810244 | 7.650861882 | 7.396486569 | 0.019362972 | 0.105513315 |
| AP53_878 | -0.262178183 | 7.663320523 | 7.380357526 | 0.019467494 | 0.105513315 |
| AP53_1496 | 0.291725419 | 8.657112087 | 7.380344447 | 0.019467579 | 0.105513315 |
| AP53_238 | 0.271127447 | 9.458554412 | 7.379483884 | 0.019473175 | 0.105513315 |
| AP53_1533 | 0.369831092 | 8.145137342 | 7.369329693 | 0.019539345 | 0.105513315 |
| AP53_1153 | 0.329560545 | 7.176369711 | 7.364368495 | 0.019571773 | 0.105513315 |
| AP53_829 | 0.598344632 | 10.09032713 | 7.35614853 | 0.019625641 | 0.105513315 |
| AP53_1629 | 1.198471654 | 2.025165839 | 7.348120406 | 0.019678424 | 0.105513315 |
| AP53_889 | -0.323853432 | 8.84098912 | 7.332197654 | 0.019783612 | 0.105785097 |
| AP53_606 | -0.694518384 | 5.885791463 | 7.317903153 | 0.019878615 | 0.106001075 |
| AP53_674 | 0.286687619 | 8.27484944 | 7.263170248 | 0.020247441 | 0.107329931 |
| AP53_351 | 0.489387873 | 9.231868952 | 7.257085882 | 0.020288943 | 0.107329931 |
| AP53_1591 | 0.33254163 | 10.34080977 | 7.256388492 | 0.020293707 | 0.107329931 |
| AP53_1685 | -0.42987085 | 6.368726164 | 7.220951826 | 0.020537519 | 0.108099759 |
| AP53_1582 | 0.390997112 | 8.371673484 | 7.219057665 | 0.02055065 | 0.108099759 |
| AP53_166 | -0.284015312 | 7.535923888 | 7.182437578 | 0.020806479 | 0.109149662 |
| AP53_31 | 0.429483525 | 7.975795889 | 7.142384875 | 0.02109065 | 0.110342185 |
| AP53_574 | -0.498363943 | 7.228015948 | 7.126510746 | 0.021204555 | 0.110380736 |
| AP53_1542 | -0.335042733 | 9.030240073 | 7.121900083 | 0.021237776 | 0.110380736 |
| AP53_668 | -0.46775008 | 10.80527704 | 7.117626777 | 0.021268622 | 0.110380736 |
| AP53_1092 | -0.676543498 | 6.259530038 | 7.095447294 | 0.021429582 | 0.110919514 |
| AP53_328 | -0.649507654 | 8.781683546 | 7.05640214 | 0.021716483 | 0.111640149 |
| AP53_1263 | 1.006968169 | 10.24910459 | 7.054491784 | 0.021730637 | 0.111640149 |
| AP53_rna75 | 3.036448967 | -0.117416991 | 7.5318185 | 0.021791306 | 0.111640149 |
| AP53_rna13 | 1.094289834 | 3.975155994 | 7.044007053 | 0.021808518 | 0.111640149 |
| AP53_769 | -0.407183822 | 8.463737777 | 7.037584147 | 0.021856392 | 0.111640149 |
| AP53_554 | 0.822918264 | 2.74527286 | 7.014387779 | 0.022030336 | 0.112233288 |
| AP53_272 | 0.254653486 | 9.628509917 | 6.96992039 | 0.022368429 | 0.113657384 |
| AP53_542 | 0.244569632 | 9.748023883 | 6.940711162 | 0.022593883 | 0.114503204 |
| AP53_449 | 1.201591844 | 1.82602402 | 6.929277949 | 0.022682869 | 0.114562193 |

| | | | | | |
|---|---|---|---|---|---|
| AP53_219 | -0.563614935 | 11.68278344 | 6.924066767 | 0.022723567 | 0.114562193 |
| AP53_822 | 0.989789056 | 9.76644687 | 6.904370515 | 0.02287818 | 0.115042868 |
| AP53_pseudo928 | 0.371840153 | 5.682119527 | 6.866804728 | 0.023176562 | 0.116242135 |
| AP53_1638 | -0.37610478 | 7.493116441 | 6.849444996 | 0.023316017 | 0.116640179 |
| AP53_95 | -0.529975624 | 6.349589747 | 6.821322204 | 0.023544068 | 0.117402683 |
| AP53_rna74 | 0.939721149 | 3.94592579 | 6.812105393 | 0.023619387 | 0.117402683 |
| AP53_1426 | 0.525240789 | 6.450441917 | 6.804695643 | 0.023680147 | 0.117402683 |
| AP53_608 | -0.768814023 | 5.182205261 | 6.801016921 | 0.023710382 | 0.117402683 |
| AP53_1543 | -0.405697227 | 10.58250761 | 6.777189519 | 0.023907338 | 0.118076699 |
| AP53_486 | -0.644096501 | 8.007340446 | 6.768656226 | 0.023978348 | 0.118126838 |
| AP53_1623 | -0.450038648 | 6.765951862 | 6.746038911 | 0.024167782 | 0.118596006 |
| AP53_931 | -0.424771165 | 6.177726661 | 6.742713537 | 0.024195785 | 0.118596006 |
| AP53_960 | -0.360890388 | 7.539641314 | 6.728818216 | 0.024313216 | 0.118871415 |
| AP53_435 | 0.431090388 | 7.890446052 | 6.721008938 | 0.024379512 | 0.11889606 |
| AP53_751 | -0.29257161 | 9.44213199 | 6.705703937 | 0.024510069 | 0.119233192 |
| AP53_1826 | -0.483329025 | 6.765204913 | 6.695149075 | 0.024600593 | 0.119374375 |
| AP53_531 | -0.270409577 | 7.100928641 | 6.668304435 | 0.024832631 | 0.120199844 |
| AP53_1427 | 0.497401366 | 11.35827444 | 6.648236235 | 0.025007806 | 0.120475941 |
| AP53_600 | 0.680941984 | 10.95078582 | 6.64401538 | 0.025044837 | 0.120475941 |
| AP53_1710 | 0.720141789 | 4.837104462 | 6.639588555 | 0.025083746 | 0.120475941 |
| AP53_1131 | -0.295358495 | 8.424826186 | 6.631675418 | 0.025153477 | 0.120475941 |
| AP53_387 | 1.159201793 | 1.849198738 | 6.613764964 | 0.025312163 | 0.120475941 |
| AP53_59 | -0.540586069 | 11.0043086 | 6.611401942 | 0.025333189 | 0.120475941 |
| AP53_770 | -0.344450479 | 8.138427926 | 6.606283619 | 0.025378802 | 0.120475941 |
| AP53_350 | -0.297397467 | 8.629141679 | 6.605451845 | 0.025386224 | 0.120475941 |
| AP53_50 | -0.562669158 | 11.20930675 | 6.584394392 | 0.025574983 | 0.121075714 |
| AP53_717 | 0.67522366 | 8.491106844 | 6.53517983 | 0.026022727 | 0.122867758 |
| AP53_355 | -0.233266852 | 7.784997873 | 6.52520667 | 0.026114599 | 0.122867758 |
| AP53_666 | -0.418335878 | 10.2229329 | 6.522086332 | 0.026143423 | 0.122867758 |
| AP53_1560 | 0.441262948 | 7.297212437 | 6.46948659 | 0.026635084 | 0.124876082 |
| AP53_343 | -0.281339079 | 6.997685949 | 6.462147757 | 0.026704557 | 0.124900109 |
| AP53_1501 | -0.316589192 | 8.011588379 | 6.454463653 | 0.026777531 | 0.124940356 |
| AP53_1516 | -0.234243604 | 6.984875302 | 6.438069405 | 0.026934022 | 0.125369154 |
| AP53_rna52 | 1.50278408 | 3.930072791 | 6.404739888 | 0.027255555 | 0.126562276 |
| AP53_1580 | 0.377626028 | 8.126588309 | 6.392379647 | 0.027375961 | 0.126817994 |

| | | | | | |
|---|---|---|---|---|---|
| AP53_1433 | 0.302357323 | 10.47960649 | 6.376374188 | 0.027532823 | 0.127240974 |
| AP53_1833 | 0.368514081 | 8.861494486 | 6.319510649 | 0.028098883 | 0.128832758 |
| AP53_1149 | -0.492761141 | 11.11120345 | 6.316761808 | 0.028126598 | 0.128832758 |
| AP53_1670 | 0.522539073 | 6.467137102 | 6.31305608 | 0.028164013 | 0.128832758 |
| AP53_301 | 0.502496411 | 9.702029137 | 6.300575455 | 0.028290461 | 0.128832758 |
| AP53_503 | -0.306209422 | 7.641910473 | 6.299907997 | 0.028297243 | 0.128832758 |
| AP53_1682 | -0.377553884 | 5.782192653 | 6.291515863 | 0.028382675 | 0.128832758 |
| AP53_14 | 0.361392121 | 10.75548072 | 6.289043155 | 0.028407906 | 0.128832758 |
| AP53_rna2 | 1.064427142 | 3.836238549 | 6.284450133 | 0.028454844 | 0.128832758 |
| AP53_1085 | 0.677694653 | 4.943668624 | 6.282516865 | 0.028474628 | 0.128832758 |
| AP53_16 | 0.593301775 | 9.407115863 | 6.255649354 | 0.028751294 | 0.129520154 |
| AP53_288 | 0.486300115 | 7.785351794 | 6.250007635 | 0.028809797 | 0.129520154 |
| AP53_246 | 0.387795012 | 8.990207353 | 6.248376144 | 0.028826742 | 0.129520154 |
| AP53_1062 | -0.31810154 | 7.882999782 | 6.173232221 | 0.029620283 | 0.132778218 |
| AP53_1452 | -0.297155748 | 8.900423862 | 6.165965611 | 0.029698399 | 0.13282164 |
| AP53_1256 | 0.365386584 | 7.810463382 | 6.119453049 | 0.030204294 | 0.134670421 |
| AP53_477 | -0.298461964 | 8.971203091 | 6.115247511 | 0.030250543 | 0.134670421 |
| AP53_1010 | -0.705622992 | 5.822470884 | 6.107564202 | 0.030335256 | 0.134738516 |
| AP53_934 | 0.472122956 | 8.425851427 | 6.064439394 | 0.030816042 | 0.1365615 |
| AP53_1122 | 0.531362202 | 8.656673592 | 6.052454499 | 0.030951275 | 0.136615229 |
| AP53_506 | 0.321089115 | 7.599458418 | 6.04405284 | 0.031046501 | 0.136615229 |
| AP53_rna55 | 3.404495856 | 0.474471177 | 6.042985015 | 0.031058629 | 0.136615229 |
| AP53_971 | -0.481592 | 7.836608819 | 6.032859766 | 0.031173911 | 0.136615229 |
| AP53_465 | -0.505371992 | 6.15883924 | 6.032318728 | 0.031180086 | 0.136615229 |
| AP53_961 | -0.308032218 | 7.148994965 | 6.011935694 | 0.031413773 | 0.137150908 |
| AP53_743 | 0.561335488 | 9.690537483 | 6.009341557 | 0.031443665 | 0.137150908 |
| AP53_1315 | -0.653481696 | 3.418809219 | 5.987858405 | 0.031692525 | 0.137857202 |
| AP53_744 | 0.281807893 | 8.440934461 | 5.983127888 | 0.03174764 | 0.137857202 |
| AP53_469 | -0.496483889 | 9.808365257 | 5.946365635 | 0.032179893 | 0.139422259 |
| AP53_476 | 0.455380766 | 9.659574946 | 5.90579328 | 0.032665164 | 0.141209538 |
| AP53_1625 | 0.648838461 | 6.69419007 | 5.894590229 | 0.032800702 | 0.141480361 |
| AP53_541 | 0.287004555 | 8.691959385 | 5.881596236 | 0.032958753 | 0.141846875 |
| AP53_895 | -0.601311937 | 9.171990467 | 5.863201833 | 0.033184057 | 0.142500564 |
| AP53_772 | -0.459198892 | 8.155570193 | 5.811078014 | 0.033832603 | 0.144418927 |
| AP53_1297 | 0.39285277 | 6.253431008 | 5.808085307 | 0.033870299 | 0.144418927 |

| | | | | | |
|---|---|---|---|---|---|
| AP53_rna45 | 1.268356956 | 3.076635878 | 5.803421089 | 0.03392915 | 0.144418927 |
| AP53_1490 | 0.46653852 | 13.61212773 | 5.798934673 | 0.033985873 | 0.144418927 |
| AP53_464 | -0.58235517 | 5.259576695 | 5.797596963 | 0.034002808 | 0.144418927 |
| AP53_1017 | -0.901728682 | 6.274401588 | 5.782382656 | 0.034196131 | 0.144438584 |
| AP53_432 | 0.265906847 | 7.462376129 | 5.780155662 | 0.03422454 | 0.144438584 |
| AP53_776 | -0.28851133 | 7.944744549 | 5.775990977 | 0.034277742 | 0.144438584 |
| AP53_580 | -0.280355712 | 7.556711597 | 5.773852872 | 0.034305094 | 0.144438584 |
| AP53_739 | 0.549688678 | 10.05085262 | 5.761253024 | 0.034466812 | 0.14447852 |
| AP53_1173 | -0.319441478 | 9.926256259 | 5.756736979 | 0.034524998 | 0.14447852 |
| AP53_1015 | -0.742466772 | 6.161086585 | 5.755738045 | 0.034537884 | 0.14447852 |
| AP53_1360 | -1.102524553 | 9.222909666 | 5.740305314 | 0.034737704 | 0.145001902 |
| AP53_70 | -0.599243191 | 10.44975777 | 5.717253966 | 0.03503876 | 0.145944706 |
| AP53_1090 | 0.847853747 | 7.046293108 | 5.708762642 | 0.035150447 | 0.146016479 |
| AP53_333 | -0.617573164 | 7.235476923 | 5.704517343 | 0.035206446 | 0.146016479 |
| AP53_302 | 0.293854133 | 9.263777797 | 5.676149372 | 0.035583405 | 0.146977295 |
| AP53_rna28 | 1.19203262 | 4.380833458 | 5.675689491 | 0.035589556 | 0.146977295 |
| AP53_1589 | 0.365529296 | 9.549365128 | 5.656217872 | 0.03585116 | 0.147743315 |
| AP53_1343 | -0.597915149 | 4.452770568 | 5.623031102 | 0.036302371 | 0.14928581 |
| AP53_1471 | -0.456490923 | 8.431843594 | 5.606591083 | 0.036528417 | 0.149897795 |
| AP53_1021 | -1.193292472 | 6.68383023 | 5.59601706 | 0.0366747 | 0.150180577 |
| AP53_1545 | -0.376623494 | 11.4512735 | 5.508541067 | 0.037912256 | 0.154836323 |
| AP53_1643 | -0.525752609 | 8.456063552 | 5.499404458 | 0.038044389 | 0.154836323 |
| AP53_91 | 0.478194078 | 8.79430569 | 5.494157721 | 0.038120517 | 0.154836323 |
| AP53_284 | 0.513071922 | 9.147368134 | 5.489305525 | 0.038191084 | 0.154836323 |
| AP53_1546 | 0.473405239 | 9.595490849 | 5.487971713 | 0.038210509 | 0.154836323 |
| AP53_940 | 0.535483108 | 8.107856263 | 5.479608149 | 0.038332586 | 0.155007395 |
| AP53_582 | -0.22814821 | 8.124503157 | 5.473002168 | 0.03842934 | 0.155075571 |
| AP53_933 | 0.537027318 | 9.329273809 | 5.450760769 | 0.038757263 | 0.156074374 |
| AP53_1473 | -0.555859597 | 9.677362925 | 5.441748886 | 0.03889109 | 0.15628904 |
| AP53_1270 | 0.518472131 | 8.361294977 | 5.42513187 | 0.039139314 | 0.156359351 |
| AP53_237 | -0.289122555 | 6.217745756 | 5.424458138 | 0.039149418 | 0.156359351 |
| AP53_1566 | -0.467848075 | 7.086791417 | 5.424402349 | 0.039150255 | 0.156359351 |
| AP53_1104 | -0.30450336 | 8.137301301 | 5.406752641 | 0.039416084 | 0.156989776 |
| AP53_1709 | 0.604904345 | 6.130877228 | 5.403199139 | 0.039469866 | 0.156989776 |
| AP53_533 | -0.62849802 | 7.028844904 | 5.382248339 | 0.039788761 | 0.15793453 |

| | | | | | |
|---|---|---|---|---|---|
| AP53_rna12 | 1.424789339 | 3.70589102 | 5.352268213 | 0.040250501 | 0.159441271 |
| AP53_467 | -0.311451053 | 8.671118333 | 5.337438591 | 0.040481283 | 0.160028858 |
| AP53_1599 | 0.477980058 | 9.99443516 | 5.330800788 | 0.040585097 | 0.160113157 |
| AP53_rna81 | 0.920196666 | 8.333760293 | 5.316704733 | 0.040806622 | 0.160660556 |
| AP53_116 | -0.497591189 | 7.765070502 | 5.274258545 | 0.041482527 | 0.162577808 |
| AP53_864 | 0.277654347 | 7.658762239 | 5.271221386 | 0.041531405 | 0.162577808 |
| AP53_1348 | -0.53012829 | 4.605248262 | 5.270385434 | 0.04154487 | 0.162577808 |
| AP53_578 | 0.562385807 | 8.24742685 | 5.244929588 | 0.041957437 | 0.163861942 |
| AP53_577 | 0.698677227 | 8.271785237 | 5.239457318 | 0.04204677 | 0.163881086 |
| AP53_1503 | 0.406500174 | 7.92742632 | 5.231442083 | 0.04217803 | 0.16406324 |
| AP53_1578 | -0.427369692 | 8.095803619 | 5.21740114 | 0.042409161 | 0.164331902 |
| AP53_1208 | -0.811491453 | 2.736754577 | 5.216961293 | 0.042416426 | 0.164331902 |
| AP53_rna43 | 1.072171153 | 4.568139016 | 5.180285662 | 0.043027512 | 0.166367331 |
| AP53_1159 | 0.294704112 | 8.174832113 | 5.160002196 | 0.043370027 | 0.167358296 |
| AP53_1676 | 0.614638771 | 7.594097753 | 5.147002853 | 0.043591268 | 0.167380689 |
| AP53_1854 | 0.235462242 | 9.006424022 | 5.132754841 | 0.043835325 | 0.167380689 |
| AP53_545 | -0.475384884 | 12.02450849 | 5.130655433 | 0.043871425 | 0.167380689 |
| AP53_845 | 0.38633242 | 5.452419827 | 5.125255961 | 0.043964435 | 0.167380689 |
| AP53_1815 | 0.618463681 | 9.372363644 | 5.124357534 | 0.043979935 | 0.167380689 |
| AP53_472 | -0.404755162 | 11.25387969 | 5.123590811 | 0.043993167 | 0.167380689 |
| AP53_307 | 0.402329092 | 7.624681867 | 5.119814424 | 0.044058411 | 0.167380689 |
| AP53_1201 | 0.613026151 | 3.887389286 | 5.116015866 | 0.044124156 | 0.167380689 |
| AP53_834 | 0.405402099 | 7.083534567 | 5.114412752 | 0.044151938 | 0.167380689 |
| AP53_rna24 | 1.127981648 | 3.822543586 | 5.102855751 | 0.044352847 | 0.167814572 |
| AP53_1197 | -0.641013222 | 3.318117855 | 5.094375735 | 0.044500966 | 0.168047421 |
| AP53_1805 | 0.341853117 | 7.418596014 | 5.081425193 | 0.044728323 | 0.168481478 |
| AP53_444 | 0.612232098 | 3.147686211 | 5.077953346 | 0.044789512 | 0.168481478 |
| AP53_1292 | 0.392600833 | 5.910146344 | 5.070576563 | 0.044919858 | 0.16864496 |
| AP53_1111 | 0.270531321 | 8.445947905 | 5.035236014 | 0.0455507 | 0.170626647 |
| AP53_1341 | -0.889470939 | 4.226506109 | 5.031195219 | 0.045623509 | 0.170626647 |
| AP53_1317 | -0.439340925 | 3.704008296 | 5.009318374 | 0.046020139 | 0.171779019 |
| AP53_390 | -0.409770896 | 5.938200776 | 4.989377872 | 0.046385292 | 0.172809698 |
| AP53_1332 | 0.388723107 | 4.927605505 | 4.971628356 | 0.046713268 | 0.173698188 |
| AP53_1630 | -0.546583196 | 11.41737697 | 4.954708228 | 0.047028528 | 0.174190608 |
| AP53_1547 | -0.236587957 | 8.074757721 | 4.954483215 | 0.047032738 | 0.174190608 |

| | | | | | |
|---|---|---|---|---|---|
| AP53_1468 | -0.663833021 | 8.166070866 | 4.945768585 | 0.04719613 | 0.174190608 |
| AP53_1806 | 0.322320381 | 6.705292417 | 4.945314223 | 0.047204668 | 0.174190608 |
| AP53_377 | -0.324573674 | 4.518264802 | 4.925839574 | 0.047572365 | 0.175022809 |
| AP53_1642 | -0.340961593 | 8.132237723 | 4.9238285 | 0.047610532 | 0.175022809 |
| AP53_883 | -0.216925415 | 8.535576241 | 4.843954945 | 0.049156767 | 0.180365377 |
| AP53_144 | 0.408915477 | 5.193900191 | 4.804826951 | 0.04993634 | 0.182880068 |
| AP53_830 | 0.378933305 | 7.795508083 | 4.791586876 | 0.050203508 | 0.18351226 |
| AP53_1539 | 0.237016176 | 8.072907953 | 4.754594882 | 0.05095917 | 0.185434472 |
| AP53_707 | -0.324757132 | 8.129301303 | 4.751881561 | 0.051015136 | 0.185434472 |
| AP53_rna25 | 1.51328014 | 3.774093468 | 4.751840927 | 0.051015975 | 0.185434472 |
| AP53_1409 | 0.227383663 | 9.200307258 | 4.745284179 | 0.051151528 | 0.185579655 |
| AP53_1684 | -0.340180252 | 6.496705515 | 4.733004571 | 0.051406567 | 0.186120627 |
| AP53_1241 | 0.536027633 | 9.750516897 | 4.728887757 | 0.051492414 | 0.186120627 |
| AP53_903 | 0.415850007 | 4.168785317 | 4.715634996 | 0.051769953 | 0.186775982 |
| AP53_496 | -0.383468296 | 12.88864028 | 4.695284873 | 0.052199654 | 0.187736972 |
| AP53_20 | 0.493757535 | 7.070122534 | 4.693866701 | 0.05222976 | 0.187736972 |
| AP53_1361 | 0.595398965 | 9.39071655 | 4.685077776 | 0.052416806 | 0.188061037 |
| AP53_661 | 0.352829469 | 8.138897903 | 4.680113384 | 0.052522816 | 0.188093703 |
| AP53_828 | 0.416968206 | 9.672546088 | 4.668086903 | 0.052780709 | 0.188669164 |
| AP53_1340 | -0.452057611 | 4.087828755 | 4.658178244 | 0.05299434 | 0.189084583 |
| AP53_1105 | 0.543572202 | 7.325119416 | 4.645206017 | 0.053275604 | 0.189608394 |
| AP53_408 | 0.292735128 | 9.562030378 | 4.641237105 | 0.053362019 | 0.189608394 |
| AP53_1342 | -0.60416625 | 4.625304636 | 4.633342772 | 0.053534407 | 0.189608394 |
| AP53_86 | -0.73943024 | 3.208787068 | 4.632905276 | 0.053543981 | 0.189608394 |
| AP53_1608 | -0.259446329 | 7.521305155 | 4.628997768 | 0.053629577 | 0.189608394 |
| AP53_420 | -0.26589668 | 9.55445663 | 4.621175236 | 0.053801431 | 0.189826562 |
| AP53_1053 | 0.559800184 | 7.111292246 | 4.617296973 | 0.053886881 | 0.189826562 |
| AP53_1050 | -0.363687451 | 5.944521376 | 4.601433009 | 0.054238125 | 0.190604331 |
| AP53_1163 | -0.386351415 | 10.45687235 | 4.598468595 | 0.054304068 | 0.190604331 |
| AP53_1165 | -0.231488785 | 9.470830918 | 4.590637493 | 0.054478735 | 0.190872247 |
| AP53_225 | -0.254914305 | 8.009070346 | 4.585065982 | 0.054603418 | 0.190964386 |
| AP53_1636 | 0.473083935 | 8.608762784 | 4.579856421 | 0.054720314 | 0.191029009 |
| AP53_445 | 0.68899364 | 3.478242748 | 4.566953437 | 0.055011148 | 0.191699529 |
| AP53_rna44 | 0.962513146 | 3.465197914 | 4.554043228 | 0.05530402 | 0.192374738 |
| AP53_1073 | -0.364511407 | 9.28714035 | 4.532131268 | 0.055805437 | 0.193771652 |

| | | | | | |
|---|---|---|---|---|---|
| AP53_936 | 0.548303901 | 8.458939196 | 4.523563615 | 0.05600299 | 0.193776739 |
| AP53_605 | -0.764666418 | 6.486805845 | 4.523408726 | 0.056006569 | 0.193776739 |
| AP53_1703 | -0.479791423 | 8.116455482 | 4.510635197 | 0.0563027 | 0.193845166 |
| AP53_1773 | 0.37723967 | 11.49108682 | 4.50978176 | 0.056322552 | 0.193845166 |
| AP53_rna26 | 1.334792883 | 4.466309467 | 4.505938005 | 0.056412072 | 0.193845166 |
| AP53_98 | -0.62028061 | 4.172683808 | 4.503725171 | 0.056463687 | 0.193845166 |
| AP53_1691 | -0.521653902 | 10.28433427 | 4.499818048 | 0.05655496 | 0.193845166 |
| AP53_1598 | 0.693241978 | 8.199124008 | 4.496801196 | 0.056625558 | 0.193845166 |
| AP53_rna30 | 0.726927472 | 3.832576393 | 4.490130437 | 0.056782043 | 0.194038636 |
| AP53_478 | -0.201887562 | 6.86776325 | 4.482467981 | 0.056962438 | 0.194312992 |
| AP53_285 | 0.378146046 | 7.505391079 | 4.476592987 | 0.057101223 | 0.19444469 |
| AP53_1406 | -0.278170877 | 9.002072289 | 4.460807609 | 0.057476154 | 0.195378661 |
| AP53_652 | 0.421400456 | 8.254094096 | 4.448833624 | 0.057762551 | 0.195725359 |
| AP53_1447 | 0.255797738 | 8.138222723 | 4.448113916 | 0.05777982 | 0.195725359 |
| AP53_1398 | 0.256238644 | 9.98185083 | 4.401216677 | 0.058918731 | 0.19923564 |
| AP53_821 | 0.771581326 | 9.889874738 | 4.392232049 | 0.059140024 | 0.199636149 |
| AP53_653 | 0.387212234 | 7.969931946 | 4.371879029 | 0.059645062 | 0.200991434 |
| AP53_100 | -0.289279426 | 9.169923012 | 4.367673683 | 0.059750065 | 0.20099632 |
| AP53_rna11 | 0.900606696 | 4.042386343 | 4.361315737 | 0.059909243 | 0.201183114 |
| AP53_406 | -0.279811249 | 6.982701596 | 4.340422326 | 0.060435967 | 0.202129244 |
| AP53_267 | 0.501968538 | 10.33651236 | 4.339447538 | 0.060460679 | 0.202129244 |
| AP53_1440 | -0.734742305 | 6.64335366 | 4.337763638 | 0.060503396 | 0.202129244 |
| AP53_326 | 0.294190416 | 9.854876452 | 4.315659353 | 0.061067545 | 0.203663411 |
| AP53_484 | 0.264694153 | 6.791743853 | 4.310466666 | 0.061200999 | 0.203758386 |
| AP53_885 | -0.287532735 | 7.980501176 | 4.302081445 | 0.061417251 | 0.203976501 |
| AP53_510 | 0.310786137 | 5.849857875 | 4.299783012 | 0.061476689 | 0.203976501 |
| AP53_1237 | -0.293930869 | 9.540523851 | 4.290361731 | 0.061721056 | 0.20412744 |
| AP53_1462 | -0.223701132 | 7.438063323 | 4.289921156 | 0.061732513 | 0.20412744 |
| AP53_105 | -0.282048779 | 7.031738335 | 4.274014869 | 0.062147864 | 0.205151368 |
| AP53_1168 | -0.18923497 | 7.247266685 | 4.269341796 | 0.062270534 | 0.205207313 |
| AP53_897 | -0.345084877 | 8.720799367 | 4.26326623 | 0.062430461 | 0.205226262 |
| AP53_82 | 0.38874178 | 8.333549952 | 4.261094456 | 0.062487749 | 0.205226262 |
| AP53_1863 | -0.240564711 | 10.1139576 | 4.249109586 | 0.062805045 | 0.20591992 |
| AP53_645 | -0.630112271 | 4.180054042 | 4.237879841 | 0.063104128 | 0.206485256 |
| AP53_138 | -0.566527882 | 10.18656462 | 4.234658843 | 0.063190233 | 0.206485256 |

| | | | | | |
|---|---|---|---|---|---|
| AP53_277 | 0.191255202 | 9.564369913 | 4.215801369 | 0.063697212 | 0.207484789 |
| AP53_650 | -0.465060736 | 4.173284571 | 4.21533141 | 0.06370991 | 0.207484789 |
| AP53_8 | 0.479149141 | 7.292519567 | 4.20819677 | 0.063903058 | 0.20776522 |
| AP53_592 | 0.377869071 | 7.379795712 | 4.181070486 | 0.064643952 | 0.209739838 |
| AP53_1216 | -0.426223984 | 11.02528837 | 4.178071101 | 0.064726514 | 0.209739838 |
| AP53_1322 | -0.528408745 | 2.744344257 | 4.170480102 | 0.064936039 | 0.209795506 |
| AP53_488 | -0.647029604 | 7.678716534 | 4.169618745 | 0.064959866 | 0.209795506 |
| AP53_627 | -0.685935664 | 3.230233061 | 4.147410812 | 0.065577876 | 0.211439629 |
| AP53_46 | -0.500796308 | 11.29190986 | 4.135372096 | 0.065915884 | 0.212176999 |
| AP53_892 | 0.374824048 | 12.79974523 | 4.129347207 | 0.066085839 | 0.212371878 |
| AP53_1121 | 0.303273058 | 8.876121787 | 4.120155575 | 0.066346154 | 0.212856009 |
| AP53_1458 | -0.24544893 | 9.111977327 | 4.103582799 | 0.066818671 | 0.21401822 |
| AP53_rna27 | 0.717604865 | 4.911695727 | 4.098128305 | 0.066975084 | 0.214165795 |
| AP53_528 | -0.219127771 | 8.886217142 | 4.08853073 | 0.067251386 | 0.214405546 |
| AP53_1369 | -0.402662975 | 4.662961336 | 4.087003921 | 0.067295469 | 0.214405546 |
| AP53_1244 | -0.274851586 | 8.999923432 | 4.080285727 | 0.067489859 | 0.214405546 |
| AP53_1823 | 0.241790244 | 5.856257941 | 4.080215132 | 0.067491906 | 0.214405546 |
| AP53_1561 | 0.354943994 | 8.081135354 | 4.0445441 | 0.068535616 | 0.217365408 |
| AP53_384 | 0.646926728 | 5.312484241 | 4.011398629 | 0.069523109 | 0.22013761 |
| AP53_273 | -0.229820632 | 7.785493392 | 3.992027307 | 0.070108267 | 0.221628903 |
| AP53_970 | -0.285369715 | 6.591033471 | 3.980647128 | 0.070454834 | 0.222362331 |
| AP53_78 | -0.239509075 | 7.419845793 | 3.956370126 | 0.071201166 | 0.224353026 |
| AP53_rna9 | 0.834622043 | 4.351643606 | 3.944286225 | 0.071576249 | 0.225169366 |
| AP53_1285 | -0.419235571 | 8.215176469 | 3.913776461 | 0.072534051 | 0.227813258 |
| AP53_1349 | -0.650344721 | 3.309771167 | 3.906283094 | 0.072771682 | 0.227998032 |
| AP53_740 | 0.391808345 | 10.38253532 | 3.899799169 | 0.072978068 | 0.227998032 |
| AP53_641 | -0.406626313 | 4.573462804 | 3.89724063 | 0.073059704 | 0.227998032 |
| AP53_1859 | -0.262023483 | 8.358397793 | 3.891639594 | 0.073238808 | 0.227998032 |
| AP53_888 | 0.188163645 | 9.182311722 | 3.883783343 | 0.073490931 | 0.227998032 |
| AP53_499 | -0.220009927 | 7.651709629 | 3.882298556 | 0.0735387 | 0.227998032 |
| AP53_481 | 0.492571897 | 8.287145628 | 3.879189445 | 0.07363885 | 0.227998032 |
| AP53_1367 | 0.573644618 | 10.32687169 | 3.878215088 | 0.07367027 | 0.227998032 |
| AP53_67 | -0.559104056 | 10.20321776 | 3.87781815 | 0.073683074 | 0.227998032 |
| AP53_393 | -0.400413944 | 3.336583387 | 3.875202368 | 0.073767524 | 0.227998032 |
| AP53_196 | -0.185025739 | 8.712936276 | 3.868067574 | 0.073998471 | 0.228348223 |

| | | | | | |
|---|---|---|---|---|---|
| AP53_1550 | -0.271280473 | 7.688059374 | 3.860131655 | 0.074256388 | 0.228539421 |
| AP53_898 | -0.668833215 | 6.544989312 | 3.858918358 | 0.074295917 | 0.228539421 |
| AP53_1150 | -0.283185687 | 7.68102015 | 3.843903763 | 0.074787222 | 0.229686705 |
| AP53_1091 | 0.349941866 | 8.27044064 | 3.836305579 | 0.075037361 | 0.230090867 |
| AP53_1463 | -0.246375763 | 7.910418677 | 3.823234381 | 0.075470075 | 0.231052706 |
| AP53_470 | -0.390956351 | 7.956823872 | 3.81017179 | 0.075905555 | 0.232019973 |
| AP53_665 | 0.231552978 | 8.344185197 | 3.806118418 | 0.076041311 | 0.232069472 |
| AP53_1706 | 0.334024096 | 12.43279509 | 3.79380683 | 0.076455475 | 0.232494465 |
| AP53_1018 | -0.779393343 | 5.920351528 | 3.79139789 | 0.076536834 | 0.232494465 |
| AP53_672 | 0.180663928 | 8.457220619 | 3.791306915 | 0.076539909 | 0.232494465 |
| AP53_1347 | -0.617085077 | 4.277213214 | 3.787335512 | 0.076674277 | 0.232538706 |
| AP53_1652 | 0.271446851 | 8.385416906 | 3.769331898 | 0.077287041 | 0.233822505 |
| AP53_604 | -0.724149031 | 6.321387552 | 3.76782765 | 0.07733851 | 0.233822505 |
| AP53_820 | 0.659638668 | 9.906872654 | 3.759074402 | 0.077638841 | 0.234335617 |
| AP53_1668 | -0.301834043 | 13.35226329 | 3.751885644 | 0.077886562 | 0.234335617 |
| AP53_1235 | 0.328091622 | 10.02292846 | 3.751670381 | 0.077893994 | 0.234335617 |
| AP53_1647 | 0.513956301 | 5.062154392 | 3.748859123 | 0.077991143 | 0.234335617 |
| AP53_1356 | -0.546952635 | 3.936521777 | 3.74332072 | 0.078182968 | 0.234548903 |
| AP53_1418 | -0.39002683 | 7.937633942 | 3.729971068 | 0.078647717 | 0.235579043 |
| AP53_1585 | 0.493677376 | 8.253436029 | 3.723255178 | 0.078882801 | 0.235919131 |
| AP53_334 | -0.478557331 | 7.109344875 | 3.715149602 | 0.079167675 | 0.236406858 |
| AP53_1577 | 0.442976863 | 7.414925905 | 3.706613282 | 0.079469052 | 0.236918708 |
| AP53_pseudo268 | 0.2248502 | 7.889228161 | 3.703390363 | 0.079583203 | 0.236918708 |
| AP53_1345 | -0.496585648 | 4.364938321 | 3.68513405 | 0.08023362 | 0.238489215 |
| AP53_1541 | -0.258689423 | 7.953481663 | 3.672654435 | 0.080681981 | 0.239455238 |
| AP53_1005 | 0.282474184 | 7.959082217 | 3.660984275 | 0.081104042 | 0.240340374 |
| AP53_rna72 | 2.442278639 | 5.80827402 | 3.713146225 | 0.08127706 | 0.240412162 |
| AP53_844 | -0.265964124 | 7.79716985 | 3.653504233 | 0.081375987 | 0.240412162 |
| AP53_973 | -0.234239156 | 5.879572272 | 3.635768876 | 0.082025256 | 0.241962039 |
| AP53_1303 | -0.414431339 | 8.524657579 | 3.622996141 | 0.082496787 | 0.242960709 |
| AP53_429 | 0.637879401 | 2.085177102 | 3.619830906 | 0.082614152 | 0.242960709 |
| AP53_rna48 | 0.843778728 | 5.123122914 | 3.611455087 | 0.08292571 | 0.243508024 |
| AP53_985 | -0.241868019 | 8.1689722 | 3.605135448 | 0.083161739 | 0.243694905 |
| AP53_247 | 0.284650926 | 8.868964733 | 3.603032739 | 0.083240455 | 0.243694905 |
| AP53_1460 | 0.270220939 | 9.238176012 | 3.581327182 | 0.084058376 | 0.245349606 |

| | | | | | |
|---|---|---|---|---|---|
| AP53_585 | 0.213131029 | 9.096517146 | 3.581324735 | 0.084058469 | 0.245349606 |
| AP53_682 | 0.603162858 | 10.79805948 | 3.568265229 | 0.08455534 | 0.246429301 |
| AP53_rna46 | 1.104528753 | 3.464757328 | 3.562515131 | 0.084775255 | 0.246699805 |
| AP53_1554 | -0.325968415 | 8.751635964 | 3.556094656 | 0.08502164 | 0.247046411 |
| AP53_894 | -0.47330528 | 9.692295089 | 3.548704219 | 0.085306336 | 0.247395686 |
| AP53_1700 | -0.322623466 | 8.287535281 | 3.546363247 | 0.085396759 | 0.247395686 |
| AP53_274 | 0.241683703 | 10.45166169 | 3.539338099 | 0.085668824 | 0.24781399 |
| AP53_19 | 0.303461744 | 7.876750887 | 3.533131814 | 0.085910062 | 0.248142011 |
| AP53_1009 | -0.634633405 | 5.668230454 | 3.520421081 | 0.086406739 | 0.24898229 |
| AP53_344 | -0.37221929 | 7.517459375 | 3.519126364 | 0.086457529 | 0.24898229 |
| AP53_859 | 0.343872493 | 7.980890273 | 3.514341434 | 0.086645551 | 0.249154096 |
| AP53_rna51 | 2.471198418 | 5.804361528 | 3.561328552 | 0.086884252 | 0.24919182 |
| AP53_1141 | -0.567079897 | 5.045558232 | 3.504719857 | 0.087025152 | 0.24919182 |
| AP53_1089 | -0.577241873 | 6.462081772 | 3.504248018 | 0.08704382 | 0.24919182 |
| AP53_1699 | -0.45771386 | 8.424003252 | 3.499680881 | 0.08722477 | 0.24934209 |
| AP53_1221 | 0.181008239 | 8.478727955 | 3.473808635 | 0.088258608 | 0.251926409 |
| AP53_1510 | -0.365264737 | 6.923368282 | 3.461112017 | 0.088771467 | 0.253018234 |
| AP53_1497 | -0.357525283 | 7.204863828 | 3.45253799 | 0.089119875 | 0.253638824 |
| AP53_843 | -0.242887069 | 7.471071188 | 3.430009226 | 0.09004339 | 0.255686994 |
| AP53_835 | 0.520586029 | 10.22822728 | 3.428565071 | 0.09010299 | 0.255686994 |
| AP53_1140 | -0.53559688 | 6.040327791 | 3.417922515 | 0.090543712 | 0.256192435 |
| AP53_305 | -0.384005929 | 6.571725935 | 3.417889444 | 0.090545085 | 0.256192435 |
| AP53_1586 | 0.446739683 | 7.962072421 | 3.380281392 | 0.092123927 | 0.259644083 |
| AP53_rna58 | 0.83075455 | 3.880953751 | 3.380220848 | 0.092126496 | 0.259644083 |
| AP53_1386 | -0.467222825 | 7.247876238 | 3.379283214 | 0.092166292 | 0.259644083 |
| AP53_1508 | 0.199797468 | 8.636400623 | 3.374748012 | 0.09235908 | 0.259810107 |
| AP53_1065 | 0.352090753 | 5.632716854 | 3.36045046 | 0.092970117 | 0.261150502 |
| AP53_1484 | -0.466428255 | 9.517232347 | 3.348217833 | 0.093496865 | 0.262084553 |
| AP53_233 | -0.237477888 | 8.635923669 | 3.346463899 | 0.093572692 | 0.262084553 |
| AP53_774 | -0.18682432 | 8.855491958 | 3.336885371 | 0.093988138 | 0.26219452 |
| AP53_950 | 0.383636946 | 10.0211407 | 3.335292821 | 0.094057431 | 0.26219452 |
| AP53_108 | -0.226292704 | 7.882968078 | 3.332989307 | 0.094157771 | 0.26219452 |
| AP53_336 | 0.240973286 | 10.01169484 | 3.332672288 | 0.09417159 | 0.26219452 |
| AP53_522 | 0.223150125 | 7.975341605 | 3.328434115 | 0.094356581 | 0.26219452 |
| AP53_1022 | -1.066626262 | 6.923782671 | 3.324295636 | 0.094537653 | 0.26219452 |

| | | | | | |
|---|---|---|---|---|---|
| AP53_15 | 0.211981542 | 7.727893238 | 3.323841949 | 0.094557529 | 0.26219452 |
| AP53_163 | 0.45873982 | 3.782831697 | 3.314296781 | 0.094976907 | 0.262694706 |
| AP53_428 | 0.285692737 | 4.324315563 | 3.310904786 | 0.095126491 | 0.262694706 |
| AP53_rna64 | 0.916669034 | 4.164097717 | 3.310509638 | 0.095143935 | 0.262694706 |
| AP53_1667 | 0.251257359 | 10.53558133 | 3.296041517 | 0.095785384 | 0.264090101 |
| AP53_579 | 0.254698224 | 7.606143427 | 3.266489826 | 0.09711224 | 0.267368591 |
| AP53_11 | 0.213788922 | 7.488083093 | 3.260944092 | 0.097363764 | 0.267681395 |
| AP53_1138 | 0.295182922 | 6.670704766 | 3.240699842 | 0.09828879 | 0.269842349 |
| AP53_1359 | -0.646320499 | 10.7346197 | 3.237635415 | 0.098429758 | 0.269847684 |
| AP53_808 | 0.325215634 | 7.800966263 | 3.225919378 | 0.098971023 | 0.270820125 |
| AP53_698 | 0.364234491 | 8.420522513 | 3.223925179 | 0.099063518 | 0.270820125 |
| AP53_1410 | -0.257439416 | 9.086127775 | 3.214952159 | 0.099481031 | 0.271579018 |
| AP53_49 | -0.416748633 | 11.30618214 | 3.210984958 | 0.099666318 | 0.271702701 |
| AP53_359 | -0.519108436 | 7.623428763 | 3.198489936 | 0.100252689 | 0.272917909 |
| AP53_1651 | -0.359147239 | 8.844024322 | 3.19124353 | 0.100594706 | 0.273299424 |
| AP53_1346 | -0.506276388 | 4.250518521 | 3.189558562 | 0.10067444 | 0.273299424 |
| AP53_1756 | -0.39169762 | 9.032149695 | 3.173622698 | 0.101432419 | 0.274972523 |
| AP53_601 | -0.559432138 | 8.445616324 | 3.17059618 | 0.101577171 | 0.274980877 |
| AP53_1306 | 0.280045816 | 6.047640193 | 3.150711447 | 0.102534599 | 0.276550901 |
| AP53_1640 | -0.2297884 | 6.301335828 | 3.150572353 | 0.102541335 | 0.276550901 |
| AP53_349 | -0.196632388 | 8.400918934 | 3.139503294 | 0.103079184 | 0.276550901 |
| AP53_452 | 0.780121849 | 1.143102125 | 3.139196055 | 0.103094163 | 0.276550901 |
| AP53_175 | -0.628141289 | 5.691972231 | 3.137761377 | 0.103164142 | 0.276550901 |
| AP53_1683 | -0.291117794 | 6.01827946 | 3.135831673 | 0.10325836 | 0.276550901 |
| AP53_684 | -0.240007919 | 7.260847375 | 3.132430678 | 0.103424675 | 0.276550901 |
| AP53_1354 | -0.652889764 | 3.37901892 | 3.129690053 | 0.103558937 | 0.276550901 |
| AP53_282 | -0.214334853 | 8.804343559 | 3.129446562 | 0.103570876 | 0.276550901 |
| AP53_1312 | -0.481991289 | 6.203251073 | 3.123185862 | 0.103878441 | 0.276550901 |
| AP53_1373 | 0.351455348 | 9.092152825 | 3.122934741 | 0.103890801 | 0.276550901 |
| AP53_rna35 | 2.380400895 | 5.821934084 | 3.166328409 | 0.103896662 | 0.276550901 |
| AP53_1453 | 0.33842244 | 10.39467819 | 3.118515594 | 0.104108611 | 0.276550901 |
| AP53_rna8 | 0.756262907 | 4.747622072 | 3.117640025 | 0.104151833 | 0.276550901 |
| AP53_423 | 0.257356308 | 10.05280575 | 3.114620728 | 0.10430105 | 0.276568768 |
| AP53_1043 | -0.313995373 | 6.060397034 | 3.109527644 | 0.104553356 | 0.276859569 |
| AP53_1314 | -0.416183905 | 4.001388929 | 3.098280569 | 0.10511321 | 0.277962862 |

| | | | | | |
|---|---|---|---|---|---|
| AP53_110 | 0.266438503 | 8.013028847 | 3.092520835 | 0.105401356 | 0.278345621 |
| AP53_405 | -0.221174343 | 5.795434844 | 3.072402232 | 0.106415559 | 0.280186469 |
| AP53_rna29 | 0.75406239 | 4.81700164 | 3.070951558 | 0.106489156 | 0.280186469 |
| AP53_1112 | 0.325744771 | 9.61875054 | 3.07011776 | 0.106531486 | 0.280186469 |
| AP53_103 | -0.205517231 | 6.880115778 | 3.064793691 | 0.106802268 | 0.280518541 |
| AP53_416 | -0.204205662 | 8.589937643 | 3.056080112 | 0.107247285 | 0.280870825 |
| AP53_1057 | -0.651123817 | 4.900415044 | 3.046570372 | 0.107735593 | 0.280870825 |
| AP53_rna1 | 2.352773556 | 5.84424604 | 3.088101944 | 0.107745506 | 0.280870825 |
| AP53_1294 | 0.345112068 | 5.511894944 | 3.044340747 | 0.10785048 | 0.280870825 |
| AP53_142 | -0.354011148 | 4.788570738 | 3.040614342 | 0.108042831 | 0.280870825 |
| AP53_1147 | -0.556426274 | 9.234701475 | 3.039122192 | 0.108119973 | 0.280870825 |
| AP53_1192 | -0.708770324 | 2.102695241 | 3.037969995 | 0.108179586 | 0.280870825 |
| AP53_1130 | -0.441432637 | 12.05479986 | 3.037890268 | 0.108183713 | 0.280870825 |
| AP53_1584 | 0.162887691 | 9.405547829 | 3.036827641 | 0.108238731 | 0.280870825 |
| AP53_920 | 0.296534304 | 9.684761164 | 3.031863333 | 0.108496222 | 0.281163106 |
| AP53_128 | -0.273528126 | 6.379202322 | 3.021658699 | 0.109027913 | 0.281873291 |
| AP53_1663 | 0.371064689 | 6.881238382 | 3.021031229 | 0.109060712 | 0.281873291 |
| AP53_819 | 0.740693497 | 8.051631786 | 3.014474308 | 0.109404184 | 0.28213826 |
| AP53_rna80 | 2.346743169 | 5.803466699 | 3.053209897 | 0.109519797 | 0.28213826 |
| AP53_1350 | -0.369323733 | 3.748987503 | 3.010760825 | 0.109599304 | 0.28213826 |
| AP53_364 | 0.199078844 | 9.106172219 | 3.002262043 | 0.11004749 | 0.282916793 |
| AP53_92 | 0.28244753 | 6.356531205 | 2.983341469 | 0.111053484 | 0.285125414 |
| AP53_1822 | 0.16258233 | 7.315602649 | 2.977310173 | 0.111376566 | 0.285577165 |
| AP53_1472 | -0.435781034 | 9.358208714 | 2.963665061 | 0.112111825 | 0.286988303 |
| AP53_1205 | -0.525225709 | 2.924279 | 2.956447943 | 0.112503157 | 0.286988303 |
| AP53_1388 | -0.379284282 | 7.381478696 | 2.955685415 | 0.112544603 | 0.286988303 |
| AP53_926 | 0.254573825 | 8.964821659 | 2.953688051 | 0.112653255 | 0.286988303 |
| AP53_394 | -0.348133517 | 4.161062865 | 2.953450318 | 0.112666196 | 0.286988303 |
| AP53_1170 | -0.240584242 | 8.008830292 | 2.947301776 | 0.113001533 | 0.287224633 |
| AP53_938 | 0.419014229 | 8.540954766 | 2.946324804 | 0.11305493 | 0.287224633 |
| AP53_1198 | -0.386812489 | 3.798169381 | 2.930882124 | 0.113903157 | 0.289001342 |
| AP53_417 | -0.348077135 | 7.960783991 | 2.926429863 | 0.114149178 | 0.28924746 |
| AP53_1437 | -0.256336134 | 7.885640041 | 2.91861339 | 0.114582701 | 0.289967435 |
| AP53_1588 | -0.341932349 | 10.15300196 | 2.909558728 | 0.115087465 | 0.290865585 |
| AP53_781 | 0.191574444 | 9.416215669 | 2.902846575 | 0.115463431 | 0.291436307 |

| | | | | | |
|---|---|---|---|---|---|
| AP53_rna14 | 2.279677778 | 5.834101851 | 2.936718043 | 0.115715528 | 0.291501116 |
| AP53_734 | 0.543517706 | 8.466451951 | 2.897047808 | 0.115789469 | 0.291501116 |
| AP53_1522 | -0.289261354 | 8.075835223 | 2.890845691 | 0.116139457 | 0.29200348 |
| AP53_1721 | -0.382277039 | 6.061442966 | 2.884128348 | 0.116520009 | 0.292581289 |
| AP53_1035 | -0.539242238 | 4.328280742 | 2.876627988 | 0.116946758 | 0.29327346 |
| AP53_534 | -0.38318711 | 7.793605183 | 2.86447069 | 0.117642624 | 0.294547717 |
| AP53_1470 | -0.616700427 | 8.386196245 | 2.862456936 | 0.117758387 | 0.294547717 |
| AP53_1219 | 0.164185571 | 8.732265628 | 2.857714002 | 0.118031601 | 0.294851142 |
| AP53_689 | -0.421520348 | 9.352042618 | 2.848356147 | 0.118572979 | 0.295750237 |
| AP53_rna3 | 2.247614316 | 5.832630068 | 2.883434286 | 0.118696257 | 0.295750237 |
| AP53_243 | -0.533959558 | 4.137115508 | 2.84322815 | 0.118870962 | 0.295805817 |
| AP53_519 | -0.156265519 | 8.866446243 | 2.837012576 | 0.119233398 | 0.296327818 |
| AP53_rna78 | 2.212563362 | 5.81260375 | 2.86637066 | 0.119671208 | 0.29674376 |
| AP53_1012 | -0.682915661 | 5.87453472 | 2.82893413 | 0.119706525 | 0.29674376 |
| AP53_rna70 | 2.226219344 | 5.783091751 | 2.857350583 | 0.120190661 | 0.297275092 |
| AP53_rna37 | 2.197537861 | 5.803484641 | 2.856371285 | 0.120247228 | 0.297275092 |
| AP53_431 | -0.363830732 | 6.495646796 | 2.817497743 | 0.120380331 | 0.297275092 |
| AP53_890 | -0.239632063 | 6.723217123 | 2.811414083 | 0.120740697 | 0.297786141 |
| AP53_678 | -0.173417816 | 8.376474821 | 2.80805763 | 0.120940094 | 0.297899394 |
| AP53_520 | -0.245940328 | 8.789098109 | 2.800810268 | 0.121372044 | 0.298330473 |
| AP53_203 | 0.25882098 | 6.238333279 | 2.799965798 | 0.1214225 | 0.298330473 |
| AP53_rna83 | 1.085397364 | 3.752216362 | 2.795653347 | 0.121680576 | 0.298586597 |
| AP53_1075 | -0.319591381 | 9.355194238 | 2.788059584 | 0.122136689 | 0.299248969 |
| AP53_rna53 | 2.175735348 | 5.811675989 | 2.821608866 | 0.122277148 | 0.299248969 |
| AP53_532 | -0.282895752 | 8.611940454 | 2.782598183 | 0.122466045 | 0.299248969 |
| AP53_1052 | 0.291607488 | 7.631196824 | 2.780924558 | 0.122567198 | 0.299248969 |
| AP53_921 | 0.264657068 | 7.962211143 | 2.76151966 | 0.123747677 | 0.301440386 |
| AP53_132 | -0.223446412 | 10.33594975 | 2.761067276 | 0.123775367 | 0.301440386 |
| AP53_rna16 | 2.206616247 | 5.830256733 | 2.792542219 | 0.124007771 | 0.301627925 |
| AP53_648 | -0.484685975 | 7.381473531 | 2.729281728 | 0.12574048 | 0.305459665 |
| AP53_1204 | -0.378123166 | 2.871581077 | 2.719154031 | 0.126374814 | 0.306345448 |
| AP53_1366 | 0.193316816 | 9.455194799 | 2.716134795 | 0.126564695 | 0.306345448 |
| AP53_354 | 0.26411931 | 9.752665909 | 2.715914027 | 0.126578593 | 0.306345448 |
| AP53_115 | -0.352561553 | 5.049902357 | 2.708958243 | 0.127017468 | 0.307024789 |
| AP53_584 | 0.498690825 | 10.15155159 | 2.705896268 | 0.127211268 | 0.307110785 |

| | | | | | |
|---|---|---|---|---|---|
| AP53_229 | -0.208645476 | 8.192570383 | 2.694112749 | 0.127960543 | 0.308535919 |
| AP53_998 | 0.192132151 | 8.269286534 | 2.686788596 | 0.128429053 | 0.309281378 |
| AP53_833 | -0.330946379 | 6.204504065 | 2.683741262 | 0.128624617 | 0.309368503 |
| AP53_1137 | -0.269791919 | 5.163380194 | 2.679620659 | 0.128889655 | 0.309622301 |
| AP53_1646 | 0.601774807 | 4.87587575 | 2.676998787 | 0.12905865 | 0.309645043 |
| AP53_1657 | 0.214950237 | 8.21286656 | 2.669050561 | 0.129572664 | 0.310494494 |
| AP53_1210 | -0.88233815 | 1.936449278 | 2.655395283 | 0.130461769 | 0.312239573 |
| AP53_398 | -0.65014584 | 2.694450071 | 2.639572193 | 0.131501628 | 0.313993008 |
| AP53_1573 | -0.241009471 | 8.771126769 | 2.63716802 | 0.131660535 | 0.313993008 |
| AP53_84 | 0.324139433 | 3.193047419 | 2.636878218 | 0.131679706 | 0.313993008 |
| AP53_352 | 0.237670649 | 8.762418006 | 2.633583799 | 0.131897887 | 0.314127359 |
| AP53_753 | -0.24296693 | 8.807008081 | 2.621925803 | 0.132673624 | 0.315587628 |
| AP53_113 | -0.418557379 | 5.138902606 | 2.611835936 | 0.133349653 | 0.316490355 |
| AP53_430 | -0.267890741 | 7.043124354 | 2.611395764 | 0.133379243 | 0.316490355 |
| AP53_974 | -0.213905283 | 5.52041317 | 2.593054431 | 0.134619607 | 0.318748343 |
| AP53_201 | 0.29413611 | 6.70652531 | 2.59247146 | 0.134659269 | 0.318748343 |
| AP53_493 | -0.147504674 | 8.217555346 | 2.582158313 | 0.135363353 | 0.320024687 |
| AP53_310 | 0.241568984 | 6.753458318 | 2.566477566 | 0.136442792 | 0.322012839 |
| AP53_1155 | 0.479835008 | 11.46796564 | 2.565129478 | 0.136536098 | 0.322012839 |
| AP53_792 | -0.189337508 | 7.475883425 | 2.559977761 | 0.136893407 | 0.322463718 |
| AP53_771 | -0.261855671 | 9.030765767 | 2.557573517 | 0.137060562 | 0.322466123 |
| AP53_1143 | -0.288793431 | 9.577025249 | 2.555025988 | 0.13723796 | 0.322492591 |
| AP53_708 | -0.278045989 | 9.580698848 | 2.535422355 | 0.138612793 | 0.325329421 |
| AP53_1391 | -0.289977926 | 7.33462873 | 2.530804576 | 0.138939169 | 0.325388945 |
| AP53_345 | 0.405759785 | 8.13500415 | 2.53032057 | 0.138973434 | 0.325388945 |
| AP53_10 | 0.316071934 | 6.92771867 | 2.522737698 | 0.139511651 | 0.32625556 |
| AP53_1368 | -0.337135942 | 3.964798628 | 2.517084242 | 0.139914635 | 0.326804219 |
| AP53_65 | -0.484551247 | 8.071274492 | 2.500326117 | 0.141117834 | 0.329218408 |
| AP53_957 | 0.352147242 | 8.083285336 | 2.489068259 | 0.141933469 | 0.330723725 |
| AP53_685 | -0.199000429 | 8.867203535 | 2.472835836 | 0.143120029 | 0.332915287 |
| AP53_415 | -0.241838636 | 9.242783866 | 2.471516301 | 0.143217035 | 0.332915287 |
| AP53_rna68 | -0.523661814 | 4.055443708 | 2.466421081 | 0.143592392 | 0.333289998 |
| AP53_256 | 0.215080732 | 8.209398764 | 2.464670343 | 0.143721653 | 0.333289998 |
| AP53_559 | -0.384700021 | 9.645722823 | 2.454576336 | 0.144469796 | 0.334625148 |
| AP53_1662 | 0.265199402 | 4.998466761 | 2.442756655 | 0.145352116 | 0.336267529 |

| | | | | | |
|---|---|---|---|---|---|
| AP53_996 | 0.256417859 | 8.831870744 | 2.435207741 | 0.145919201 | 0.337177583 |
| AP53_171 | 0.315809666 | 7.61453829 | 2.42294428 | 0.146846437 | 0.338916688 |
| AP53_pseudo603 | -0.492204542 | 7.457202431 | 2.415217762 | 0.147434471 | 0.339869725 |
| AP53_1469 | -0.563907782 | 8.997062334 | 2.40867082 | 0.147935069 | 0.340619181 |
| AP53_1157 | -0.319853201 | 3.708622363 | 2.403030986 | 0.148368037 | 0.340908919 |
| AP53_rna85 | 0.458293567 | 4.683211769 | 2.402457177 | 0.148412178 | 0.340908919 |
| AP53_412 | -0.443527748 | 9.840928338 | 2.376172191 | 0.150452168 | 0.344782042 |
| AP53_1387 | -0.412907363 | 7.519770716 | 2.376154211 | 0.150453575 | 0.344782042 |
| AP53_1095 | 0.43466378 | 3.813457285 | 2.370491766 | 0.150897696 | 0.345222461 |
| AP53_1319 | -0.432864019 | 3.772270914 | 2.36760315 | 0.151124899 | 0.345222461 |
| AP53_1834 | 0.283950888 | 7.62930424 | 2.366911856 | 0.151179336 | 0.345222461 |
| AP53_795 | 0.350431685 | 10.58190416 | 2.362646884 | 0.151515745 | 0.345281099 |
| AP53_1392 | -0.188472329 | 8.160192567 | 2.362076697 | 0.151560792 | 0.345281099 |
| AP53_1563 | -0.402012319 | 8.225798925 | 2.359362227 | 0.151775477 | 0.345364832 |
| AP53_622 | -0.347251842 | 3.759174187 | 2.351358906 | 0.152410708 | 0.346404197 |
| AP53_1731 | -0.267648702 | 8.042599518 | 2.347870074 | 0.152688677 | 0.346630084 |
| AP53_1853 | -0.177812734 | 7.660663908 | 2.341968483 | 0.153160348 | 0.347294668 |
| AP53_64 | -0.432787283 | 7.801527973 | 2.329282759 | 0.154180514 | 0.349199974 |
| AP53_1765 | 0.24311327 | 10.01501353 | 2.313674409 | 0.155447605 | 0.351659442 |
| AP53_1185 | 0.579996589 | 1.757692781 | 2.311354132 | 0.155637096 | 0.351678234 |
| AP53_439 | 0.466233075 | 2.385210709 | 2.305473782 | 0.156118647 | 0.352297209 |
| AP53_303 | -0.341633256 | 6.727022064 | 2.303581255 | 0.156274032 | 0.352297209 |
| AP53_494 | -0.205055443 | 7.071482449 | 2.298519241 | 0.156690617 | 0.352826552 |
| AP53_172 | 0.284153937 | 8.017024394 | 2.293943962 | 0.157068363 | 0.353267315 |
| AP53_1098 | -0.36343138 | 4.219661802 | 2.286563248 | 0.157680182 | 0.354232909 |
| AP53_283 | -0.236298278 | 9.767281081 | 2.255413296 | 0.160296063 | 0.359693247 |
| AP53_99 | -0.201643007 | 8.792586277 | 2.249919149 | 0.160763176 | 0.359940124 |
| AP53_362 | 0.191706431 | 9.437692988 | 2.249757285 | 0.160776964 | 0.359940124 |
| AP53_1217 | 0.163319934 | 8.364502351 | 2.245786682 | 0.161115664 | 0.360202368 |
| AP53_1144 | -0.26022287 | 6.785620853 | 2.244036424 | 0.161265254 | 0.360202368 |
| AP53_1521 | -0.23396191 | 7.131119651 | 2.22645851 | 0.162777475 | 0.363162159 |
| AP53_992 | -0.443187677 | 9.403799049 | 2.214317516 | 0.163832554 | 0.365096426 |
| AP53_pseudo1389 | -0.328013798 | 7.021878509 | 2.212044436 | 0.16403106 | 0.365119597 |
| AP53_rna39 | 0.846004137 | 3.543319887 | 2.197211223 | 0.165334021 | 0.367598322 |
| AP53_1339 | -0.569462013 | 4.132509886 | 2.193861772 | 0.165630071 | 0.367733928 |

| | | | | | |
|---|---|---|---|---|---|
| AP53_1307 | -0.460169252 | 4.02732108 | 2.192237012 | 0.165773924 | 0.367733928 |
| AP53_456 | 0.244697985 | 10.88979319 | 2.187220907 | 0.166219051 | 0.368300432 |
| AP53_226 | 0.129287384 | 7.602630791 | 2.18500306 | 0.16641635 | 0.368317144 |
| AP53_309 | 0.26853118 | 8.569034772 | 2.180908033 | 0.166781431 | 0.368440132 |
| AP53_114 | -0.309596455 | 5.371535414 | 2.178026094 | 0.167038978 | 0.368440132 |
| AP53_565 | -0.438838708 | 10.93748296 | 2.177999253 | 0.167041379 | 0.368440132 |
| AP53_480 | 0.335792593 | 5.721513167 | 2.169276269 | 0.167824042 | 0.369746272 |
| AP53_850 | 0.18881139 | 5.641731126 | 2.159681523 | 0.168690361 | 0.37123355 |
| AP53_1178 | -0.536184744 | 6.83398811 | 2.157338049 | 0.168902827 | 0.371234965 |
| AP53_779 | 0.242451015 | 9.525736977 | 2.155458021 | 0.169073524 | 0.371234965 |
| AP53_1686 | -0.77931642 | 7.500481105 | 2.145113849 | 0.17001669 | 0.372884062 |
| AP53_640 | -0.269976719 | 4.209067475 | 2.132026855 | 0.171219643 | 0.375062324 |
| AP53_1693 | -0.295003789 | 13.096781 | 2.130114584 | 0.171396333 | 0.375062324 |
| AP53_1782 | -0.262805006 | 8.412433501 | 2.122074383 | 0.172141798 | 0.37589753 |
| AP53_159 | -0.305585691 | 5.702756779 | 2.119958608 | 0.172338659 | 0.37589753 |
| AP53_677 | -0.190671548 | 8.111374097 | 2.119740271 | 0.172358991 | 0.37589753 |
| AP53_297 | -0.220275483 | 7.089865324 | 2.115895814 | 0.172717492 | 0.376256624 |
| AP53_1148 | -0.254674046 | 11.24128382 | 2.109759624 | 0.173291689 | 0.376929176 |
| AP53_1166 | -0.166666846 | 8.162139025 | 2.108449441 | 0.173414608 | 0.376929176 |
| AP53_762 | 0.398967589 | 9.11980704 | 2.103282945 | 0.173900414 | 0.377390729 |
| AP53_1485 | -0.360468787 | 9.463774976 | 2.100351763 | 0.174176811 | 0.377390729 |
| AP53_402 | 0.266747273 | 4.590131084 | 2.099997564 | 0.174210249 | 0.377390729 |
| AP53_711 | 0.512121309 | 2.784300559 | 2.088469253 | 0.175303089 | 0.378134289 |
| AP53_45 | -0.381794527 | 11.20939483 | 2.086777552 | 0.175464198 | 0.378134289 |
| AP53_1299 | 0.277106925 | 6.93179985 | 2.079318344 | 0.176176852 | 0.378134289 |
| AP53_119 | -0.214753218 | 5.4753407 | 2.079167246 | 0.176191326 | 0.378134289 |
| AP53_681 | 0.274769855 | 9.560920647 | 2.078644011 | 0.176241461 | 0.378134289 |
| AP53_1725 | -0.39423683 | 6.193449723 | 2.078604732 | 0.176245226 | 0.378134289 |
| AP53_5 | 0.17828167 | 8.003714405 | 2.078416489 | 0.176263268 | 0.378134289 |
| AP53_1569 | -0.249883188 | 7.211031447 | 2.078206325 | 0.176283414 | 0.378134289 |
| AP53_596 | 0.519423617 | 8.004957284 | 2.076770239 | 0.176421152 | 0.378134289 |
| AP53_536 | 0.162234464 | 8.741025202 | 2.075931826 | 0.176501631 | 0.378134289 |
| AP53_1243 | 0.340719259 | 9.14538567 | 2.070948038 | 0.176980999 | 0.37874324 |
| AP53_757 | -0.253344039 | 8.988796479 | 2.063220963 | 0.177727554 | 0.379922007 |
| AP53_511 | 0.220269334 | 6.045520278 | 2.059383586 | 0.178099813 | 0.380093627 |

| | | | | | |
|---|---|---|---|---|---|
| AP53_1301 | 0.265339555 | 8.23204417 | 2.05723789 | 0.178308403 | 0.380093627 |
| AP53_374 | 0.254852466 | 8.82937416 | 2.055142238 | 0.178512432 | 0.380093627 |
| AP53_614 | -0.611785485 | 5.801452265 | 2.054334708 | 0.178591132 | 0.380093627 |
| AP53_189 | 0.128073872 | 8.5345132 | 2.046863317 | 0.179321405 | 0.381085363 |
| AP53_460 | -0.271257839 | 7.599552137 | 2.043800065 | 0.17962193 | 0.381085363 |
| AP53_944 | -0.18027688 | 7.544989533 | 2.04355384 | 0.179646114 | 0.381085363 |
| AP53_269 | -0.169425269 | 8.07559697 | 2.033179162 | 0.180668956 | 0.382519452 |
| AP53_1179 | 0.148336416 | 7.530511432 | 2.032700793 | 0.1807163 | 0.382519452 |
| AP53_1847 | 0.269171678 | 5.744415718 | 2.028076413 | 0.181174797 | 0.383072202 |
| AP53_1527 | 0.163383076 | 10.56430151 | 2.02294227 | 0.181685599 | 0.383658154 |
| AP53_1705 | -0.190369971 | 7.659559924 | 2.021321436 | 0.181847244 | 0.383658154 |
| AP53_1603 | -0.20204788 | 6.695972091 | 2.018686382 | 0.182110434 | 0.383796258 |
| AP53_463 | -0.263192497 | 5.479056278 | 2.016140612 | 0.182365175 | 0.383916273 |
| AP53_169 | 0.175755865 | 9.092234106 | 2.012703131 | 0.182709875 | 0.384225208 |
| AP53_280 | 0.140740802 | 8.991995718 | 1.996044597 | 0.184392335 | 0.387011796 |
| AP53_1028 | -0.546242288 | 5.282694774 | 1.995637007 | 0.184433751 | 0.387011796 |
| AP53_1316 | -0.385728933 | 3.674597187 | 1.989026993 | 0.185107094 | 0.387881123 |
| AP53_1704 | -0.278511061 | 8.338143977 | 1.987646749 | 0.185248097 | 0.387881123 |
| AP53_446 | 0.273930546 | 3.672188595 | 1.984117003 | 0.185609323 | 0.387881123 |
| AP53_1211 | 0.370529675 | 1.877716067 | 1.983745648 | 0.185647379 | 0.387881123 |
| AP53_1446 | 0.141183458 | 8.608058006 | 1.97701114 | 0.186339292 | 0.388908135 |
| AP53_1436 | -0.404160324 | 7.919112504 | 1.971677732 | 0.186889627 | 0.389637773 |
| AP53_778 | -0.174622503 | 7.90192138 | 1.958844299 | 0.188222528 | 0.39199563 |
| AP53_1020 | -0.644451818 | 6.170653395 | 1.954211469 | 0.18870673 | 0.392582812 |
| AP53_1740 | -0.291489745 | 6.04185596 | 1.947492973 | 0.189411794 | 0.393627721 |
| AP53_210 | 0.148290347 | 8.282924267 | 1.940101698 | 0.190191419 | 0.394445466 |
| AP53_1797 | 0.39484714 | 3.912813951 | 1.939909718 | 0.190211724 | 0.394445466 |
| AP53_1829 | -0.33220376 | 9.962085059 | 1.930725227 | 0.191186451 | 0.396043651 |
| AP53_1125 | 0.231344953 | 7.154776504 | 1.928244297 | 0.191450859 | 0.396168569 |
| AP53_1024 | -0.649033095 | 7.609041708 | 1.922516694 | 0.192063104 | 0.397012231 |
| AP53_800 | 0.12813032 | 8.470305784 | 1.909677433 | 0.193444825 | 0.399442985 |
| AP53_266 | 0.206764789 | 9.303906861 | 1.88753106 | 0.19585872 | 0.40297716 |
| AP53_893 | -0.275282833 | 10.31994222 | 1.886662116 | 0.195954231 | 0.40297716 |
| AP53_518 | 0.139926969 | 7.32366515 | 1.884010775 | 0.196246032 | 0.40297716 |
| AP53_455 | 0.230422901 | 10.67283373 | 1.883501848 | 0.196302108 | 0.40297716 |

| | | | | | |
|---|---|---|---|---|---|
| AP53_1786 | 0.33238299 | 6.068040946 | 1.882676996 | 0.196393039 | 0.40297716 |
| AP53_154 | 0.18479226 | 9.987487702 | 1.880915065 | 0.196587456 | 0.40297716 |
| AP53_877 | -0.133551183 | 8.076302094 | 1.880197061 | 0.196666755 | 0.40297716 |
| AP53_424 | -0.297424099 | 6.560053996 | 1.878144747 | 0.19689365 | 0.40297716 |
| AP53_935 | 0.289371873 | 9.235748517 | 1.876959 | 0.197024897 | 0.40297716 |
| AP53_495 | -0.253408146 | 6.404686299 | 1.871920658 | 0.197583852 | 0.403695006 |
| AP53_1058 | 0.288521672 | 4.464736719 | 1.86402773 | 0.198463663 | 0.405066215 |
| AP53_1844 | -0.156176257 | 8.19988329 | 1.858243211 | 0.199111705 | 0.405704298 |
| AP53_1601 | -0.142382592 | 7.763040086 | 1.857507451 | 0.199194331 | 0.405704298 |
| AP53_264 | 0.267361256 | 9.718548387 | 1.850161439 | 0.200021748 | 0.406773995 |
| AP53_69 | -0.384377026 | 10.49884439 | 1.84591805 | 0.200501749 | 0.406773995 |
| AP53_1532 | 0.359911477 | 11.03730632 | 1.844050647 | 0.200713463 | 0.406773995 |
| AP53_1327 | -0.425048939 | 3.739674086 | 1.842687357 | 0.200868208 | 0.406773995 |
| AP53_1323 | -0.49192897 | 2.327562017 | 1.842096191 | 0.200935358 | 0.406773995 |
| AP53_223 | -0.369370533 | 4.257604621 | 1.839522495 | 0.201228049 | 0.406773995 |
| AP53_346 | 0.338700331 | 9.215258084 | 1.836742489 | 0.201544828 | 0.406773995 |
| AP53_1372 | 0.263745309 | 9.148815649 | 1.836330183 | 0.201591866 | 0.406773995 |
| AP53_521 | -0.174813711 | 8.920122241 | 1.836209307 | 0.201605658 | 0.406773995 |
| AP53_471 | 0.224986106 | 8.036414201 | 1.834030665 | 0.201854467 | 0.406853085 |
| AP53_1249 | 0.445896982 | 7.710650513 | 1.831041629 | 0.202196481 | 0.407119678 |
| AP53_157 | 0.35124435 | 5.443569593 | 1.827211675 | 0.202635824 | 0.407581487 |
| AP53_rna47 | 0.611889685 | 4.404252428 | 1.822459834 | 0.203182659 | 0.408258324 |
| AP53_51 | -0.334562734 | 11.07336914 | 1.820630673 | 0.203393671 | 0.408259685 |
| AP53_426 | -0.254836968 | 3.992919516 | 1.8071827 | 0.204953884 | 0.410966414 |
| AP53_628 | -0.382700248 | 2.940810817 | 1.802944201 | 0.205448879 | 0.411533822 |
| AP53_1378 | 0.159154596 | 10.25019188 | 1.798219648 | 0.206002488 | 0.41221735 |
| AP53_244 | 0.334111866 | 3.470264022 | 1.795418078 | 0.206331693 | 0.412305906 |
| AP53_44 | -0.325317867 | 11.31807185 | 1.794229378 | 0.206471582 | 0.412305906 |
| AP53_687 | -0.222601061 | 8.680041329 | 1.790227179 | 0.206943488 | 0.412823547 |
| AP53_1568 | -0.415538395 | 4.251335018 | 1.786970293 | 0.207328559 | 0.413162232 |
| AP53_1055 | -0.150130585 | 7.699856721 | 1.784700972 | 0.207597424 | 0.413162232 |
| AP53_517 | -0.199312606 | 5.639080472 | 1.783399313 | 0.207751849 | 0.413162232 |
| AP53_561 | -0.304143317 | 9.705734837 | 1.777157946 | 0.208494406 | 0.414214577 |
| AP53_1784 | -0.330165109 | 7.840220968 | 1.772425392 | 0.209059782 | 0.414913125 |
| AP53_1534 | 0.237211344 | 8.554004643 | 1.767987287 | 0.209591814 | 0.415178732 |

| | | | | | |
|---|---|---|---|---|---|
| AP53_907 | -0.25691215 | 4.789085088 | 1.764261994 | 0.210039772 | 0.415178732 |
| AP53_1394 | -0.265997035 | 10.01987976 | 1.7633871 | 0.210145158 | 0.415178732 |
| AP53_1611 | 0.304774259 | 4.559297688 | 1.762894746 | 0.210204496 | 0.415178732 |
| AP53_rna76 | 1.548016846 | 0.798034033 | 1.76240859 | 0.210263109 | 0.415178732 |
| AP53_187 | -0.285726671 | 7.024537248 | 1.757171746 | 0.210895844 | 0.415449117 |
| AP53_1266 | -0.393412563 | 10.8204009 | 1.756382911 | 0.210991372 | 0.415449117 |
| AP53_714 | -1.594012689 | -0.182630266 | 1.755963724 | 0.211042158 | 0.415449117 |
| AP53_675 | -0.295976573 | 8.237706878 | 1.751730299 | 0.21155596 | 0.41603862 |
| AP53_68 | -0.390887049 | 10.91520246 | 1.739000104 | 0.21311096 | 0.418672442 |
| AP53_1231 | -0.330958616 | 6.946548077 | 1.732444422 | 0.21391762 | 0.419184127 |
| AP53_1232 | 0.53435296 | 8.0946323 | 1.731240911 | 0.214066146 | 0.419184127 |
| AP53_3 | 0.369735751 | 5.923167754 | 1.730078854 | 0.214209686 | 0.419184127 |
| AP53_1688 | 0.178865782 | 5.913434124 | 1.729871859 | 0.214235267 | 0.419184127 |
| AP53_712 | 0.835731038 | 1.394023947 | 1.727589856 | 0.214517559 | 0.419313779 |
| AP53_679 | 0.179395316 | 7.309242498 | 1.714353914 | 0.216164615 | 0.422108166 |
| AP53_rna49 | 0.617675136 | 5.221521578 | 1.712173944 | 0.216437486 | 0.422216242 |
| AP53_639 | -0.263939428 | 4.770258326 | 1.702483546 | 0.217655981 | 0.424166927 |
| AP53_131 | -0.143380698 | 10.18789975 | 1.700095191 | 0.217957691 | 0.424328865 |
| AP53_1195 | 0.54546851 | 3.18282213 | 1.69277292 | 0.218886137 | 0.425486155 |
| AP53_1318 | -0.402885186 | 4.204982717 | 1.691951979 | 0.218990556 | 0.425486155 |
| AP53_331 | -0.323503343 | 5.571994503 | 1.679033053 | 0.220642496 | 0.428267086 |
| AP53_1184 | -0.371409598 | 3.96316391 | 1.672521916 | 0.221481335 | 0.429465805 |
| AP53_1593 | -0.410021902 | 8.927438765 | 1.663057368 | 0.222708223 | 0.431413833 |
| AP53_798 | -0.164825568 | 6.426757126 | 1.654924167 | 0.223769744 | 0.433037959 |
| AP53_937 | 0.307337052 | 8.459560788 | 1.652508047 | 0.224086383 | 0.433218794 |
| AP53_rna54 | 2.240444526 | -0.548503406 | 1.690299026 | 0.224487911 | 0.433563219 |
| AP53_853 | 0.229139948 | 6.647686072 | 1.645357271 | 0.225027005 | 0.433569123 |
| AP53_997 | -0.186561965 | 8.557511037 | 1.645228249 | 0.225044025 | 0.433569123 |
| AP53_571 | -0.343035775 | 7.058893138 | 1.641648733 | 0.225516899 | 0.433569123 |
| AP53_1118 | -0.255836904 | 4.120227588 | 1.641048566 | 0.225596313 | 0.433569123 |
| AP53_321 | -0.167360193 | 9.194140231 | 1.640961487 | 0.225607838 | 0.433569123 |
| AP53_9 | 0.275242376 | 7.0436227 | 1.633044937 | 0.226658924 | 0.435158231 |
| AP53_1498 | -0.278164955 | 6.613320693 | 1.628841438 | 0.227219674 | 0.435803743 |
| AP53_1696 | 0.15951426 | 8.493555226 | 1.621117897 | 0.228254826 | 0.437356976 |
| AP53_635 | -0.286391373 | 3.98579451 | 1.614494943 | 0.229147479 | 0.438634376 |

| | | | | | |
|---|---|---|---|---|---|
| AP53_1123 | -0.169669765 | 5.808376608 | 1.608642639 | 0.22994014 | 0.439718041 |
| AP53_35 | -0.236509747 | 8.844644952 | 1.603693911 | 0.230613273 | 0.440361732 |
| AP53_329 | -0.362103178 | 7.559046869 | 1.602834123 | 0.23073049 | 0.440361732 |
| AP53_1007 | -0.499734502 | 5.788601678 | 1.592960437 | 0.232082319 | 0.442234341 |
| AP53_871 | -0.235543723 | 7.585096391 | 1.591872486 | 0.232231919 | 0.442234341 |
| AP53_1760 | -0.363306541 | 4.176515525 | 1.590011874 | 0.232488064 | 0.442234341 |
| AP53_567 | -0.195329066 | 7.522620469 | 1.589032738 | 0.23262301 | 0.442234341 |
| AP53_1330 | -0.410100391 | 3.811752405 | 1.577169582 | 0.234266361 | 0.444915726 |
| AP53_733 | 0.27643352 | 7.69791382 | 1.574544596 | 0.234632086 | 0.444915726 |
| AP53_959 | -0.174074278 | 7.296947342 | 1.57237057 | 0.234935559 | 0.444915726 |
| AP53_1439 | -0.193548224 | 8.381408097 | 1.572264743 | 0.234950345 | 0.444915726 |
| AP53_591 | 0.17329824 | 6.796622764 | 1.57049317 | 0.235198049 | 0.444950695 |
| AP53_529 | 0.149384053 | 9.140066361 | 1.559002047 | 0.236813278 | 0.447570177 |
| AP53_749 | -0.328164073 | 8.761878754 | 1.554929663 | 0.237389271 | 0.447918024 |
| AP53_1702 | -0.207530061 | 7.891787285 | 1.552243414 | 0.23777024 | 0.447918024 |
| AP53_1624 | 0.273289564 | 6.419915055 | 1.551459109 | 0.237881626 | 0.447918024 |
| AP53_1351 | -0.364831378 | 3.956639772 | 1.551186258 | 0.237920393 | 0.447918024 |
| AP53_287 | 0.201099534 | 7.802514992 | 1.545901616 | 0.238672905 | 0.448741891 |
| AP53_396 | -0.651858289 | 2.364449789 | 1.544868511 | 0.238820388 | 0.448741891 |
| AP53_239 | -0.19874086 | 8.619236094 | 1.534368082 | 0.240326365 | 0.451134888 |
| AP53_750 | -0.284790103 | 8.941525709 | 1.519843039 | 0.242430665 | 0.454645333 |
| AP53_1355 | -0.44108122 | 4.089876018 | 1.51389586 | 0.243299408 | 0.455442891 |
| AP53_158 | -0.311724471 | 4.055778573 | 1.513719505 | 0.243325234 | 0.455442891 |
| AP53_1377 | 0.149830113 | 10.53840346 | 1.5076558 | 0.244215448 | 0.456668771 |
| AP53_1860 | -0.240949681 | 6.937017296 | 1.504916669 | 0.24461902 | 0.456794493 |
| AP53_597 | 0.383681794 | 8.519968898 | 1.502771951 | 0.244935641 | 0.456794493 |
| AP53_945 | 0.14634252 | 7.848707534 | 1.502412903 | 0.2449887 | 0.456794493 |
| AP53_1553 | -0.20939342 | 7.983589658 | 1.488875798 | 0.247000558 | 0.460103727 |
| AP53_205 | 0.157551807 | 6.799939506 | 1.486998891 | 0.247281259 | 0.460184969 |
| AP53_642 | -0.301329977 | 5.138889041 | 1.47682619 | 0.248810156 | 0.462586698 |
| AP53_525 | 0.144248868 | 6.856350395 | 1.46845511 | 0.25007788 | 0.464498722 |
| AP53_1825 | -0.171723462 | 6.740475722 | 1.466842422 | 0.25032311 | 0.46450971 |
| AP53_1549 | -0.249964842 | 8.063844327 | 1.463316674 | 0.250860381 | 0.46479207 |
| AP53_1114 | -0.143144469 | 7.467225765 | 1.462702093 | 0.250954193 | 0.46479207 |
| AP53_75 | 0.17013145 | 7.726886869 | 1.45461576 | 0.252192969 | 0.466641137 |

| | | | | | |
|---|---|---|---|---|---|
| AP53_1102 | 0.270322957 | 6.954638951 | 1.451245309 | 0.252711748 | 0.467143486 |
| AP53_1044 | 0.235418435 | 6.201115305 | 1.449727744 | 0.252945803 | 0.467143486 |
| AP53_660 | 0.196590563 | 7.324931732 | 1.434749985 | 0.255271695 | 0.470990837 |
| AP53_1596 | -0.175486125 | 7.534487352 | 1.428148695 | 0.256306033 | 0.472450152 |
| AP53_1754 | -0.15556287 | 11.52845548 | 1.418707529 | 0.25779527 | 0.474744421 |
| AP53_440 | 0.371930503 | 2.590914792 | 1.413888644 | 0.258559936 | 0.475293637 |
| AP53_89 | -0.26124089 | 3.062870992 | 1.413742051 | 0.258583246 | 0.475293637 |
| AP53_107 | 0.134966239 | 8.10523054 | 1.408798972 | 0.259370929 | 0.47629042 |
| AP53_369 | 0.288073654 | 6.804805955 | 1.404008744 | 0.260137378 | 0.476375104 |
| AP53_1239 | 0.242261055 | 8.396129883 | 1.403251701 | 0.26025879 | 0.476375104 |
| AP53_448 | 0.543504503 | 1.098269493 | 1.402973024 | 0.260303502 | 0.476375104 |
| AP53_222 | -0.183545922 | 12.83297456 | 1.40064195 | 0.260677923 | 0.476375104 |
| AP53_1567 | -0.25584497 | 8.47442752 | 1.399123648 | 0.260922191 | 0.476375104 |
| AP53_1329 | -0.332092746 | 3.992006818 | 1.398763168 | 0.260980231 | 0.476375104 |
| AP53_453 | 0.849540772 | 1.390259302 | 1.397802218 | 0.261135039 | 0.476375104 |
| AP53_211 | 0.147643135 | 8.865840499 | 1.394942384 | 0.261596497 | 0.476768827 |
| AP53_664 | 0.300693708 | 7.83188895 | 1.390792392 | 0.262268115 | 0.477544476 |
| AP53_1438 | -0.146342326 | 8.510849309 | 1.386386324 | 0.262983755 | 0.478398752 |
| AP53_624 | -0.399575669 | 4.183410825 | 1.384838322 | 0.263235817 | 0.478408915 |
| AP53_906 | -0.16606228 | 5.05535495 | 1.381549159 | 0.263772491 | 0.47866557 |
| AP53_395 | -0.296392618 | 4.13609729 | 1.377692809 | 0.264403617 | 0.47866557 |
| AP53_759 | 0.373656534 | 8.73137515 | 1.37720039 | 0.264484354 | 0.47866557 |
| AP53_1620 | -0.204826888 | 6.896683767 | 1.376555723 | 0.264590105 | 0.47866557 |
| AP53_1673 | 0.245538174 | 7.054821001 | 1.376434023 | 0.264610075 | 0.47866557 |
| AP53_1013 | -0.501090567 | 5.972803166 | 1.373234201 | 0.265135885 | 0.479170162 |
| AP53_852 | -0.193656492 | 6.142123985 | 1.367641667 | 0.266058317 | 0.480389947 |
| AP53_706 | 0.283112943 | 8.158050813 | 1.360331611 | 0.267270678 | 0.481718442 |
| AP53_1016 | -0.509144984 | 6.137775317 | 1.360212776 | 0.267290449 | 0.481718442 |
| AP53_875 | -0.327390385 | 7.024084202 | 1.357388245 | 0.267760969 | 0.482118777 |
| AP53_rna42 | 0.80720435 | 3.837079763 | 1.349541755 | 0.269074043 | 0.48336186 |
| AP53_275 | -0.166046725 | 9.150603593 | 1.348740273 | 0.269208664 | 0.48336186 |
| AP53_206 | 0.167392428 | 6.775523524 | 1.3486416 | 0.269225244 | 0.48336186 |
| AP53_1164 | -0.199702324 | 9.395850668 | 1.347319891 | 0.269447467 | 0.48336186 |
| AP53_1556 | -0.276234083 | 8.243803113 | 1.344721223 | 0.269885123 | 0.483672736 |
| AP53_1866 | -0.300488412 | 4.323233914 | 1.343334091 | 0.270119137 | 0.483672736 |

| | | | | | |
|---|---|---|---|---|---|
| AP53_1120 | 0.117092415 | 8.887645674 | 1.34039891 | 0.270615234 | 0.484114442 |
| AP53_1807 | -0.175795485 | 9.245643543 | 1.330703236 | 0.272262905 | 0.486246336 |
| AP53_1572 | 0.153293265 | 6.581863799 | 1.330439193 | 0.272307969 | 0.486246336 |
| AP53_184 | 0.142483259 | 8.108886386 | 1.323766185 | 0.273450263 | 0.487599073 |
| AP53_846 | 0.160111407 | 5.626682958 | 1.323080868 | 0.27356795 | 0.487599073 |
| AP53_rna50 | 0.620185387 | 3.729627845 | 1.31480431 | 0.274994782 | 0.489692542 |
| AP53_1269 | 0.147652694 | 7.264681156 | 1.311406837 | 0.275583459 | 0.490291011 |
| AP53_765 | -0.219325338 | 8.431233271 | 1.30464395 | 0.276760446 | 0.49193409 |
| AP53_872 | -0.229943505 | 7.873853789 | 1.29835008 | 0.277862055 | 0.493440302 |
| AP53_796 | 0.239633571 | 10.79042999 | 1.286498774 | 0.279952897 | 0.49639463 |
| AP53_538 | 0.179896085 | 8.313456641 | 1.283815463 | 0.280429317 | 0.49639463 |
| AP53_1401 | -0.100435635 | 7.825276973 | 1.283420426 | 0.28049955 | 0.49639463 |
| AP53_837 | 0.225530614 | 9.548222719 | 1.282980832 | 0.280577734 | 0.49639463 |
| AP53_722 | -0.255287039 | 7.447617739 | 1.280302434 | 0.281054752 | 0.49639463 |
| AP53_1455 | -0.172811013 | 9.896842533 | 1.280057066 | 0.281098508 | 0.49639463 |
| AP53_rna38 | 1.604160295 | -0.468952949 | 1.306190406 | 0.281315864 | 0.49639463 |
| AP53_1690 | -0.358915207 | 11.27051386 | 1.271528062 | 0.282625354 | 0.498252327 |
| AP53_308 | 0.151753486 | 7.965217081 | 1.267189228 | 0.283406503 | 0.498339703 |
| AP53_1729 | -0.115655997 | 8.363552599 | 1.266418657 | 0.283545547 | 0.498339703 |
| AP53_643 | -0.295143177 | 4.218836449 | 1.265791244 | 0.28365883 | 0.498339703 |
| AP53_1803 | -0.290187419 | 6.483353547 | 1.265552836 | 0.283701892 | 0.498339703 |
| AP53_489 | 0.363507415 | 7.91146 | 1.261302972 | 0.284471046 | 0.49923897 |
| AP53_644 | -0.189475138 | 4.75123839 | 1.255177977 | 0.285584677 | 0.500329203 |
| AP53_286 | 0.144722278 | 8.349778934 | 1.253830269 | 0.285830527 | 0.500329203 |
| AP53_1583 | -0.131332398 | 8.319343603 | 1.253638262 | 0.285865578 | 0.500329203 |
| AP53_756 | -0.150158481 | 8.863382525 | 1.252177049 | 0.286132512 | 0.500345231 |
| AP53_1459 | 0.162073981 | 5.692326177 | 1.246920339 | 0.287095685 | 0.50157761 |
| AP53_951 | -0.190099755 | 9.597594622 | 1.23885857 | 0.288581619 | 0.503720254 |
| AP53_1555 | -0.130243362 | 8.924918181 | 1.23334588 | 0.289603892 | 0.505050454 |
| AP53_989 | -0.225891087 | 8.859544528 | 1.230077371 | 0.290212393 | 0.5055104 |
| AP53_18 | -0.25341502 | 4.922451675 | 1.227831375 | 0.290631567 | 0.5055104 |
| AP53_1810 | -0.186515849 | 7.450918334 | 1.226679776 | 0.290846821 | 0.5055104 |
| AP53_1099 | -0.20164744 | 5.804455639 | 1.226253417 | 0.290926571 | 0.5055104 |
| AP53_1382 | -0.187459686 | 9.307646867 | 1.224953951 | 0.291169824 | 0.5055104 |
| AP53_1172 | 0.17892257 | 7.299821759 | 1.221603063 | 0.291798405 | 0.506033028 |

| | | | | | |
|---|---|---|---|---|---|
| AP53_81 | -0.3257673 | 9.263004421 | 1.220571641 | 0.291992268 | 0.506033028 |
| AP53_1036 | -0.312347658 | 5.48739783 | 1.216445313 | 0.292769645 | 0.506927636 |
| AP53_436 | 0.325672021 | 5.781994632 | 1.209429258 | 0.294098096 | 0.508773979 |
| AP53_728 | -0.159261339 | 7.770431655 | 1.207940806 | 0.294381012 | 0.508809923 |
| AP53_188 | -0.266960142 | 7.499966239 | 1.204627079 | 0.295012236 | 0.509447287 |
| AP53_rna22 | 0.573789927 | 4.187408477 | 1.195428955 | 0.296774349 | 0.511769236 |
| AP53_1576 | 0.146143437 | 8.093332469 | 1.194858271 | 0.296884163 | 0.511769236 |
| AP53_1379 | 0.12052813 | 9.287515947 | 1.187912661 | 0.298225262 | 0.513372474 |
| AP53_306 | -0.194981284 | 6.313976703 | 1.186069742 | 0.298582534 | 0.513372474 |
| AP53_955 | 0.177878388 | 6.856014716 | 1.185940102 | 0.298607688 | 0.513372474 |
| AP53_227 | 0.130406874 | 7.690531152 | 1.183446668 | 0.299092087 | 0.513750212 |
| AP53_7 | 0.15497585 | 6.414484418 | 1.175008861 | 0.300739525 | 0.51612327 |
| AP53_688 | -0.161120702 | 9.040885876 | 1.16715278 | 0.302284905 | 0.518317139 |
| AP53_221 | -0.188913614 | 11.87240178 | 1.161232547 | 0.303456897 | 0.519475151 |
| AP53_1809 | -0.126086769 | 7.846217249 | 1.161037949 | 0.303495529 | 0.519475151 |
| AP53_768 | -0.107008405 | 8.452878255 | 1.152749653 | 0.305147422 | 0.521442016 |
| AP53_731 | -0.179959991 | 7.025910352 | 1.150163483 | 0.30566546 | 0.521442016 |
| AP53_1396 | 0.287067289 | 4.564997894 | 1.149323659 | 0.305833954 | 0.521442016 |
| AP53_258 | 0.218156583 | 7.671083701 | 1.149078106 | 0.305883244 | 0.521442016 |
| AP53_312 | -0.249673844 | 9.453151372 | 1.148557064 | 0.30598787 | 0.521442016 |
| AP53_558 | 0.181657049 | 7.727189147 | 1.145535055 | 0.306595697 | 0.522019516 |
| AP53_1548 | 0.112578826 | 7.216243169 | 1.143255677 | 0.307055289 | 0.522343835 |
| AP53_791 | -0.109866609 | 8.265255128 | 1.133963918 | 0.308938929 | 0.52508797 |
| AP53_1214 | 0.206881549 | 7.586571696 | 1.129512162 | 0.309847211 | 0.526170985 |
| AP53_262 | -0.143702516 | 7.796758407 | 1.120187004 | 0.311762142 | 0.528577021 |
| AP53_101 | -0.18667199 | 8.357710187 | 1.118616601 | 0.312086282 | 0.528577021 |
| AP53_986 | 0.161307034 | 7.078026064 | 1.117345151 | 0.312349067 | 0.528577021 |
| AP53_535 | -0.227336521 | 10.82592105 | 1.117324463 | 0.312353345 | 0.528577021 |
| AP53_784 | 0.176063433 | 7.389123837 | 1.10742437 | 0.31441038 | 0.531096729 |
| AP53_988 | -0.237596463 | 9.577451512 | 1.106221685 | 0.314661588 | 0.531096729 |
| AP53_1160 | -0.171266465 | 7.302703675 | 1.106214053 | 0.314663183 | 0.531096729 |
| AP53_563 | -0.302506734 | 10.59655174 | 1.103392602 | 0.315253636 | 0.531631024 |
| AP53_1552 | -0.100535333 | 7.991986028 | 1.100933748 | 0.315769499 | 0.532038712 |
| AP53_1247 | 0.13581127 | 8.304777516 | 1.093973757 | 0.317236245 | 0.534046446 |
| AP53_1763 | -0.23074809 | 7.269390876 | 1.091359409 | 0.31778971 | 0.534514581 |

| | | | | | |
|---|---|---|---|---|---|
| AP53_180 | -0.411563256 | 2.720633585 | 1.086887526 | 0.318739628 | 0.535648153 |
| AP53_1689 | 0.32171193 | 2.901079833 | 1.083467703 | 0.319468813 | 0.53640914 |
| AP53_1238 | 0.178480319 | 9.054908391 | 1.073563869 | 0.321594084 | 0.538850443 |
| AP53_85 | -0.235680233 | 3.168246607 | 1.073450926 | 0.321618437 | 0.538850443 |
| AP53_214 | 0.107783235 | 9.089887731 | 1.072814936 | 0.321755622 | 0.538850443 |
| AP53_413 | 0.134784532 | 9.859277469 | 1.067753867 | 0.322850314 | 0.540217637 |
| AP53_824 | 0.18266641 | 5.45028148 | 1.063773819 | 0.323714949 | 0.541197861 |
| AP53_692 | -0.261298927 | 8.311700924 | 1.059634712 | 0.324617678 | 0.542240028 |
| AP53_794 | 0.278142582 | 9.90566459 | 1.054785288 | 0.325679943 | 0.543546664 |
| AP53_793 | 0.108576412 | 7.110767067 | 1.052377211 | 0.326209295 | 0.543962406 |
| AP53_709 | -0.190664099 | 5.106968656 | 1.049243861 | 0.326899938 | 0.544260298 |
| AP53_1279 | 0.14417551 | 6.54295096 | 1.049022805 | 0.326948742 | 0.544260298 |
| AP53_52 | -0.258644925 | 11.32480098 | 1.045652387 | 0.327694158 | 0.545007736 |
| AP53_pseudo1615 | 0.280694855 | 3.079259341 | 1.044456124 | 0.327959318 | 0.545007736 |
| AP53_83 | 0.234857326 | 3.433511599 | 1.042893647 | 0.32830612 | 0.545117347 |
| AP53_1384 | 0.119986066 | 10.00184868 | 1.041243397 | 0.328672979 | 0.545260045 |
| AP53_764 | -0.153131569 | 7.227939036 | 1.038313475 | 0.329325778 | 0.54587646 |
| AP53_295 | -0.212573342 | 6.996968242 | 1.033754301 | 0.330345317 | 0.546860977 |
| AP53_1258 | 0.110163685 | 8.322350417 | 1.0331392 | 0.330483218 | 0.546860977 |
| AP53_1128 | 0.321653436 | 10.47072346 | 1.025023618 | 0.332310504 | 0.549416259 |
| AP53_1461 | 0.102068349 | 9.01395109 | 1.012985272 | 0.335048146 | 0.552831981 |
| AP53_811 | -0.321965085 | 11.91569231 | 1.01286656 | 0.335075305 | 0.552831981 |
| AP53_485 | -0.152462502 | 6.997365124 | 1.012186583 | 0.335230934 | 0.552831981 |
| AP53_782 | 0.206605487 | 6.754178647 | 1.008584383 | 0.336057139 | 0.553724029 |
| AP53_231 | 0.180149007 | 8.46742801 | 1.005039311 | 0.336873138 | 0.554597762 |
| AP53_984 | -0.219265611 | 8.429015997 | 1.000740862 | 0.337866427 | 0.55576164 |
| AP53_1076 | -0.189726065 | 8.966213956 | 0.999113734 | 0.33824354 | 0.555910847 |
| AP53_378 | -0.197151657 | 4.547407381 | 0.996385153 | 0.338877311 | 0.556149012 |
| AP53_rna21 | 0.490160247 | 4.774862161 | 0.996023233 | 0.338961505 | 0.556149012 |
| AP53_152 | -0.1364748 | 9.753289459 | 0.994202979 | 0.339385416 | 0.556374233 |
| AP53_228 | 0.149142041 | 8.054936901 | 0.992232089 | 0.339845282 | 0.556657968 |
| AP53_74 | -0.135966984 | 6.906016983 | 0.989946572 | 0.3403797 | 0.557047881 |
| AP53_980 | 0.3714577 | 11.32503678 | 0.988761412 | 0.340657308 | 0.557047881 |
| AP53_1352 | -0.442774323 | 4.312188956 | 0.984310418 | 0.341702855 | 0.558287241 |
| AP53_513 | 0.137608474 | 5.840755779 | 0.980098917 | 0.342696477 | 0.559439749 |

| | | | | | |
|---|---|---|---|---|---|
| AP53_930 | -0.166130291 | 6.355157634 | 0.97640163 | 0.343572273 | 0.560172346 |
| AP53_1832 | 0.097708279 | 8.819021249 | 0.97576905 | 0.343722444 | 0.560172346 |
| AP53_271 | 0.100219017 | 9.333956541 | 0.974415624 | 0.344044065 | 0.560226115 |
| AP53_1381 | -0.193080455 | 7.592901113 | 0.971103312 | 0.344833048 | 0.56104019 |
| AP53_1031 | 0.180561436 | 6.564176908 | 0.967224549 | 0.345760341 | 0.56207774 |
| AP53_787 | 0.130496757 | 5.790644875 | 0.963244039 | 0.346715774 | 0.563159261 |
| AP53_1078 | 0.170406723 | 7.501289471 | 0.956162886 | 0.348425076 | 0.565462436 |
| AP53_1411 | -0.13960214 | 8.762450511 | 0.953300508 | 0.349119544 | 0.566116152 |
| AP53_543 | 0.111901479 | 9.723083223 | 0.95203841 | 0.349426401 | 0.566140772 |
| AP53_1066 | 0.142339994 | 7.930079305 | 0.950544047 | 0.349790245 | 0.566257602 |
| AP53_1525 | -0.211916223 | 9.540654293 | 0.942325178 | 0.351801388 | 0.568990162 |
| AP53_1764 | -0.252193936 | 9.072809522 | 0.941255086 | 0.352064495 | 0.568990162 |
| AP53_pseudo656 | 0.175374281 | 8.518667932 | 0.934916182 | 0.353629046 | 0.571043243 |
| AP53_179 | 0.17186848 | 9.11682714 | 0.930247336 | 0.354787993 | 0.572404336 |
| AP53_1234 | 0.145471586 | 9.054293806 | 0.929147878 | 0.355061731 | 0.572404336 |
| AP53_1072 | -0.147301507 | 9.306028122 | 0.927277126 | 0.355528223 | 0.572680731 |
| AP53_925 | 0.138013612 | 5.40964687 | 0.921736062 | 0.356915297 | 0.573665643 |
| AP53_126 | -0.138987878 | 6.282394857 | 0.919970235 | 0.357359021 | 0.573665643 |
| AP53_1701 | 0.099822484 | 8.428105587 | 0.919409096 | 0.357500197 | 0.573665643 |
| AP53_825 | 0.174061299 | 5.835413574 | 0.919080559 | 0.357582892 | 0.573665643 |
| AP53_1621 | 0.198949955 | 6.750819631 | 0.918943385 | 0.357617428 | 0.573665643 |
| AP53_581 | -0.094561441 | 7.651479409 | 0.912476792 | 0.359251162 | 0.575810491 |
| AP53_1139 | -0.113072454 | 6.556102348 | 0.91030363 | 0.35980269 | 0.576218664 |
| AP53_1451 | 0.140857829 | 6.309742493 | 0.898693625 | 0.36277072 | 0.580189752 |
| AP53_1186 | 0.244886604 | 2.275852303 | 0.898268309 | 0.362880143 | 0.580189752 |
| AP53_1229 | -0.105145163 | 6.707222003 | 0.895604228 | 0.363566671 | 0.580808977 |
| AP53_1375 | 0.091209712 | 7.123257277 | 0.893256977 | 0.364173164 | 0.581299433 |
| AP53_1791 | -0.347187106 | 5.1115042 | 0.891513868 | 0.364624537 | 0.581541682 |
| AP53_789 | 0.214166216 | 5.972167453 | 0.883614206 | 0.366680686 | 0.584340897 |
| AP53_514 | 0.11949193 | 6.573820684 | 0.882364002 | 0.367007688 | 0.584382217 |
| AP53_1635 | 0.146105693 | 7.260475475 | 0.877404736 | 0.368309154 | 0.585787539 |
| AP53_1063 | -0.153830093 | 6.562298696 | 0.8759068 | 0.368703624 | 0.585787539 |
| AP53_1385 | -0.25340023 | 6.421605107 | 0.87555766 | 0.368795658 | 0.585787539 |
| AP53_964 | 0.10762549 | 6.85891838 | 0.874311162 | 0.369124523 | 0.585830498 |
| AP53_1785 | -0.3054056 | 5.049146662 | 0.870722485 | 0.370073796 | 0.58685722 |

| | | | | | |
|---|---|---|---|---|---|
| AP53_1158 | -0.236754882 | 3.822656869 | 0.868279458 | 0.370722129 | 0.587405431 |
| AP53_987 | -0.173179678 | 9.004300497 | 0.860243134 | 0.372866956 | 0.590279334 |
| AP53_1500 | -0.150318176 | 8.961250047 | 0.859186103 | 0.373150463 | 0.590279334 |
| AP53_801 | -0.081066359 | 7.615810176 | 0.858077198 | 0.373448234 | 0.590279334 |
| AP53_1817 | -0.152039803 | 5.415938694 | 0.845921442 | 0.376736114 | 0.594991699 |
| AP53_194 | -0.132782259 | 6.239699763 | 0.839508657 | 0.37848837 | 0.596526173 |
| AP53_97 | 0.128491924 | 7.423308075 | 0.83905078 | 0.378613956 | 0.596526173 |
| AP53_966 | 0.25544436 | 5.433533808 | 0.838993399 | 0.378629699 | 0.596526173 |
| AP53_636 | -0.1808936 | 4.112039212 | 0.837655982 | 0.37899691 | 0.59662044 |
| AP53_94 | 0.107169797 | 8.902392534 | 0.834752145 | 0.379796079 | 0.597343396 |
| AP53_rna56 | 0.974748125 | 0.251749348 | 0.832787494 | 0.38033823 | 0.597343396 |
| AP53_1313 | -0.375332941 | 3.823947758 | 0.832638435 | 0.380379411 | 0.597343396 |
| AP53_1344 | -0.372228123 | 4.348525858 | 0.831486005 | 0.380698029 | 0.59735518 |
| AP53_504 | 0.149730913 | 7.698002855 | 0.830194998 | 0.381055444 | 0.59735518 |
| AP53_1517 | -0.101976233 | 7.018966461 | 0.828110125 | 0.381633722 | 0.59735518 |
| AP53_777 | -0.084432634 | 8.043231212 | 0.82504097 | 0.38248745 | 0.59735518 |
| AP53_1831 | -0.263422679 | 5.756421826 | 0.82463663 | 0.38260014 | 0.59735518 |
| AP53_341 | 0.242522496 | 3.9403568 | 0.823318585 | 0.382967834 | 0.59735518 |
| AP53_1240 | 0.182400337 | 10.00904625 | 0.822381992 | 0.383229443 | 0.59735518 |
| AP53_876 | 0.135558321 | 7.431415743 | 0.821938512 | 0.383353411 | 0.59735518 |
| AP53_1227 | 0.076811468 | 7.515049606 | 0.821252913 | 0.383545181 | 0.59735518 |
| AP53_1739 | -0.342272455 | 9.058686756 | 0.820223999 | 0.383833255 | 0.59735518 |
| AP53_289 | 0.107174095 | 7.004993998 | 0.818408136 | 0.384342466 | 0.59735518 |
| AP53_1679 | 0.120577005 | 7.966748178 | 0.818392681 | 0.384346805 | 0.59735518 |
| AP53_365 | -0.270504306 | 7.934240173 | 0.817817048 | 0.384508445 | 0.59735518 |
| AP53_1027 | 0.324578221 | 4.361305901 | 0.816135982 | 0.384981092 | 0.59735518 |
| AP53_1551 | -0.107218327 | 7.86464506 | 0.815823265 | 0.385069113 | 0.59735518 |
| AP53_1766 | -0.260311287 | 8.876366489 | 0.814605961 | 0.385412045 | 0.59735518 |
| AP53_360 | -0.15589962 | 11.50774728 | 0.813872937 | 0.385618774 | 0.59735518 |
| AP53_860 | 0.130348479 | 8.307023261 | 0.811351626 | 0.38633114 | 0.597981454 |
| AP53_736 | 0.200842022 | 8.860886175 | 0.807365706 | 0.387461442 | 0.599253023 |
| AP53_335 | -0.209064951 | 8.142871266 | 0.806280295 | 0.387770117 | 0.599253023 |
| AP53_1828 | -0.220004454 | 5.847888352 | 0.803610159 | 0.388531073 | 0.599607958 |
| AP53_1562 | -0.094393924 | 7.62799416 | 0.803306962 | 0.388617626 | 0.599607958 |
| AP53_1334 | -0.268166423 | 3.009464722 | 0.801492765 | 0.389136139 | 0.599931093 |

| | | | | | |
|---|---|---|---|---|---|
| AP53_968 | 0.087030687 | 7.568667338 | 0.798308823 | 0.390048717 | 0.600860762 |
| AP53_879 | -0.080804203 | 8.401266095 | 0.795164064 | 0.390953306 | 0.601731196 |
| AP53_443 | 0.212939642 | 4.271592966 | 0.794191286 | 0.391233781 | 0.601731196 |
| AP53_1103 | -0.159410726 | 7.380225059 | 0.79208702 | 0.391841554 | 0.602188802 |
| AP53_1787 | 0.278941798 | 5.888319301 | 0.789631387 | 0.392552657 | 0.602804357 |
| AP53_1034 | -0.165317036 | 6.332954571 | 0.785656348 | 0.393707987 | 0.604100556 |
| AP53_315 | -0.226426128 | 8.989167006 | 0.784357401 | 0.394086662 | 0.604203958 |
| AP53_rna7 | 0.4024602 | 4.436492387 | 0.777608859 | 0.39606314 | 0.606754976 |
| AP53_251 | 0.147437378 | 7.811421738 | 0.774204453 | 0.397066046 | 0.607811668 |
| AP53_93 | 0.12923355 | 8.777350987 | 0.771373931 | 0.397902899 | 0.608612708 |
| AP53_869 | -0.113438856 | 8.672479792 | 0.769811247 | 0.398366087 | 0.608841398 |
| AP53_1824 | -0.192979652 | 3.817303732 | 0.767411935 | 0.399078892 | 0.609282375 |
| AP53_976 | 0.086230596 | 9.357629102 | 0.766728072 | 0.399282422 | 0.609282375 |
| AP53_1526 | -0.159085115 | 8.884422822 | 0.765432887 | 0.399668337 | 0.609392178 |
| AP53_701 | 0.187697959 | 7.052130267 | 0.763708131 | 0.400183151 | 0.609698192 |
| AP53_162 | 0.125628302 | 5.480491043 | 0.758821503 | 0.401647368 | 0.611449053 |
| AP53_754 | -0.168576665 | 9.268599668 | 0.756700523 | 0.402285499 | 0.61194056 |
| AP53_615 | -0.351227882 | 5.69586088 | 0.755543416 | 0.402634302 | 0.611991527 |
| AP53_1286 | -0.201948025 | 9.381978509 | 0.753196164 | 0.403343321 | 0.612589504 |
| AP53_90 | -0.102832926 | 7.485381372 | 0.74499287 | 0.405836663 | 0.615691679 |
| AP53_680 | 0.170985222 | 10.64839791 | 0.744391905 | 0.406020272 | 0.615691679 |
| AP53_1146 | -0.172950425 | 9.620017711 | 0.739766699 | 0.407437766 | 0.617261848 |
| AP53_27 | -0.365537762 | 7.905731302 | 0.738940631 | 0.407691751 | 0.617261848 |
| AP53_1014 | -0.337203366 | 6.659339264 | 0.734503963 | 0.409060136 | 0.617734315 |
| AP53_1259 | -0.105244949 | 8.599922287 | 0.734429584 | 0.409083138 | 0.617734315 |
| AP53_690 | -0.130885458 | 8.924203573 | 0.733664881 | 0.409319745 | 0.617734315 |
| AP53_1477 | -0.222101382 | 10.2050706 | 0.732233978 | 0.40976306 | 0.617734315 |
| AP53_1492 | 0.171545745 | 14.03692929 | 0.731921646 | 0.409859926 | 0.617734315 |
| AP53_183 | -0.12595548 | 9.708970245 | 0.73174946 | 0.409913343 | 0.617734315 |
| AP53_1261 | 0.144830351 | 5.821106996 | 0.72990694 | 0.410485631 | 0.618116843 |
| AP53_218 | -0.099987641 | 10.12084754 | 0.727270433 | 0.41130673 | 0.61887315 |
| AP53_1669 | -0.09280799 | 7.375156326 | 0.723616672 | 0.412448937 | 0.619715603 |
| AP53_748 | -0.186032763 | 8.60819477 | 0.723437164 | 0.412505183 | 0.619715603 |
| AP53_663 | 0.159347287 | 7.573759383 | 0.720431279 | 0.413448829 | 0.620652882 |
| AP53_178 | -0.3139661 | 11.23122741 | 0.714298028 | 0.415384906 | 0.62307736 |

| | | | | | |
|---|---|---|---|---|---|
| AP53_1407 | 0.145915274 | 6.653605219 | 0.711650701 | 0.416225038 | 0.623496068 |
| AP53_727 | -0.127228637 | 7.583116513 | 0.710964112 | 0.416443369 | 0.623496068 |
| AP53_1242 | 0.211050102 | 9.804031108 | 0.710384832 | 0.416627717 | 0.623496068 |
| AP53_1206 | 0.327961994 | 3.161078979 | 0.707307548 | 0.417609207 | 0.624097642 |
| AP53_1338 | -0.205123259 | 3.122952067 | 0.706105912 | 0.417993465 | 0.624097642 |
| AP53_1665 | -0.816098115 | 6.355545902 | 0.706103308 | 0.417994299 | 0.624097642 |
| AP53_161 | 0.118954328 | 6.18135813 | 0.704553504 | 0.418490728 | 0.624358573 |
| AP53_342 | 0.182386469 | 9.409220889 | 0.703249693 | 0.41890909 | 0.624502722 |
| AP53_1003 | -0.186512506 | 6.570567463 | 0.700569209 | 0.419771294 | 0.624915399 |
| AP53_1026 | -0.300152956 | 5.864797043 | 0.699801675 | 0.420018701 | 0.624915399 |
| AP53_1610 | 0.09056994 | 7.599596944 | 0.699389136 | 0.420151775 | 0.624915399 |
| AP53_1298 | 0.152744523 | 7.18348429 | 0.694910184 | 0.421600915 | 0.62659064 |
| AP53_1333 | -0.227363441 | 3.062741465 | 0.692095006 | 0.42251585 | 0.62746998 |
| AP53_1771 | 0.73841831 | 2.941398101 | 0.688821688 | 0.423583688 | 0.628574877 |
| AP53_746 | -0.142533719 | 8.06732286 | 0.684382427 | 0.425038825 | 0.630156444 |
| AP53_397 | -0.343673864 | 3.124222425 | 0.683591925 | 0.425298785 | 0.630156444 |
| AP53_125 | -0.148482679 | 5.993230058 | 0.679662854 | 0.426594681 | 0.631497353 |
| AP53_797 | 0.181586314 | 11.03110948 | 0.677936061 | 0.42716623 | 0.631497353 |
| AP53_1225 | 0.152187376 | 7.639382578 | 0.677895057 | 0.427179817 | 0.631497353 |
| AP53_1161 | -0.193104341 | 7.283232404 | 0.674546981 | 0.428291574 | 0.632659014 |
| AP53_1025 | -0.397944902 | 6.219577531 | 0.668601291 | 0.430277447 | 0.635109145 |
| AP53_1054 | -0.262624027 | 4.635450138 | 0.664015876 | 0.431819188 | 0.636900489 |
| AP53_1737 | 0.157715657 | 13.40341086 | 0.657613394 | 0.433986922 | 0.639611705 |
| AP53_141 | -0.12188657 | 5.279381164 | 0.654828833 | 0.434935239 | 0.640522988 |
| AP53_rna33 | -0.453889146 | 4.405899646 | 0.647909851 | 0.437306263 | 0.643277477 |
| AP53_527 | -0.099949771 | 8.774876702 | 0.646880782 | 0.437660711 | 0.643277477 |
| AP53_1486 | -0.17856779 | 9.62811879 | 0.646477136 | 0.43779987 | 0.643277477 |
| AP53_1119 | -0.142526856 | 7.650473294 | 0.644909585 | 0.438340975 | 0.64358535 |
| AP53_1812 | 0.129432151 | 8.060183411 | 0.642905584 | 0.439034336 | 0.644116134 |
| AP53_838 | 0.179297494 | 8.42697884 | 0.639641794 | 0.440167421 | 0.645175714 |
| AP53_1515 | -0.075578085 | 7.717556279 | 0.638912284 | 0.44042134 | 0.645175714 |
| AP53_1637 | 0.116413519 | 6.216054756 | 0.637915818 | 0.440768566 | 0.645197425 |
| AP53_1511 | 0.108363114 | 7.467094329 | 0.636509892 | 0.441259236 | 0.645428921 |
| AP53_213 | -0.093816099 | 9.447110586 | 0.633913616 | 0.442167704 | 0.646270717 |
| AP53_1777 | -0.132425431 | 7.689044846 | 0.630186875 | 0.443477123 | 0.647696837 |

| | | | | | |
|---|---|---|---|---|---|
| AP53_1358 | 0.1759691 | 13.07681823 | 0.627228626 | 0.444521083 | 0.648240825 |
| AP53_257 | -0.150057575 | 8.245360951 | 0.626971702 | 0.444611942 | 0.648240825 |
| AP53_1449 | -0.279502551 | 5.031016365 | 0.625876827 | 0.44499948 | 0.648240825 |
| AP53_438 | 0.157092231 | 5.192022396 | 0.624830248 | 0.445370445 | 0.648240825 |
| AP53_358 | 0.081643438 | 7.203562096 | 0.624410265 | 0.445519454 | 0.648240825 |
| AP53_704 | -0.119634384 | 8.096405396 | 0.623222723 | 0.445941236 | 0.648368494 |
| AP53_873 | -0.220733333 | 8.200934288 | 0.619980261 | 0.44709624 | 0.649561229 |
| AP53_1070 | 0.152040372 | 6.090033599 | 0.614081111 | 0.449210342 | 0.652144558 |
| AP53_1581 | 0.192526791 | 6.347730772 | 0.612635205 | 0.44973105 | 0.652319205 |
| AP53_900 | 0.104101777 | 5.716788735 | 0.611881739 | 0.45000279 | 0.652319205 |
| AP53_1405 | -0.084167275 | 9.099489907 | 0.609289177 | 0.450939896 | 0.653189804 |
| AP53_651 | 0.256010486 | 3.86697987 | 0.606097609 | 0.452097988 | 0.654378967 |
| AP53_855 | 0.145695475 | 6.633769712 | 0.602447753 | 0.453428467 | 0.655427521 |
| AP53_1857 | -0.255578351 | 7.917900341 | 0.601267125 | 0.453860241 | 0.655427521 |
| AP53_252 | 0.104990973 | 7.357726558 | 0.601251599 | 0.453865924 | 0.655427521 |
| AP53_1413 | -0.272390601 | 6.936292989 | 0.600412786 | 0.454173115 | 0.655427521 |
| AP53_414 | -0.118456519 | 9.780415718 | 0.590954592 | 0.45766115 | 0.659970499 |
| AP53_803 | 0.12812987 | 6.424057096 | 0.587420776 | 0.458975925 | 0.660028398 |
| AP53_1604 | 0.107183824 | 7.401009618 | 0.586338478 | 0.459379872 | 0.660028398 |
| AP53_1464 | -0.086574899 | 8.759238143 | 0.586283826 | 0.459400285 | 0.660028398 |
| AP53_856 | 0.129032392 | 6.912105337 | 0.586258876 | 0.459409605 | 0.660028398 |
| AP53_rna40 | 0.454962455 | 3.861167023 | 0.583388638 | 0.460483891 | 0.660028398 |
| AP53_1724 | -0.135568204 | 8.526789921 | 0.583067535 | 0.460604338 | 0.660028398 |
| AP53_1408 | -0.132599217 | 8.416879746 | 0.58301259 | 0.460624954 | 0.660028398 |
| AP53_373 | -0.11060273 | 9.046666375 | 0.581840957 | 0.461064924 | 0.660028398 |
| AP53_763 | -0.093519331 | 7.618129502 | 0.581531486 | 0.461181255 | 0.660028398 |
| AP53_136 | 0.081750638 | 9.356913988 | 0.581073925 | 0.461353344 | 0.660028398 |
| AP53_649 | -0.201016336 | 6.231483278 | 0.580838842 | 0.461441801 | 0.660028398 |
| AP53_983 | -0.112309319 | 8.231438178 | 0.578132851 | 0.462462079 | 0.661000659 |
| AP53_1135 | -0.169442459 | 7.158557208 | 0.575629172 | 0.463409482 | 0.661867406 |
| AP53_723 | 0.155095178 | 7.192492431 | 0.572287981 | 0.464678938 | 0.663150966 |
| AP53_151 | -0.116864019 | 9.943785209 | 0.571181606 | 0.465100597 | 0.663150966 |
| AP53_1518 | 0.092419332 | 6.978776583 | 0.570572186 | 0.465333135 | 0.663150966 |
| AP53_22 | -0.21274722 | 8.533996792 | 0.559109819 | 0.469744029 | 0.668295592 |
| AP53_1653 | 0.098282744 | 6.900789998 | 0.558989152 | 0.469790843 | 0.668295592 |

| | | | | | |
|---|---|---|---|---|---|
| AP53_1519 | -0.091923956 | 7.460775577 | 0.558512005 | 0.469976034 | 0.668295592 |
| AP53_278 | -0.078180093 | 9.52934866 | 0.552012213 | 0.47251126 | 0.671408753 |
| AP53_1321 | -0.274422913 | 2.997535762 | 0.549778421 | 0.47338797 | 0.671711686 |
| AP53_1628 | -0.313797545 | 2.437120727 | 0.549705636 | 0.473416583 | 0.671711686 |
| AP53_1622 | -0.137406613 | 7.523407045 | 0.546623718 | 0.47463089 | 0.672942701 |
| AP53_1365 | -0.181374656 | 9.690534392 | 0.544473965 | 0.475481095 | 0.673656063 |
| AP53_1404 | 0.129157995 | 6.130712776 | 0.542540211 | 0.476248123 | 0.673780343 |
| AP53_958 | 0.121667256 | 7.312010699 | 0.542305469 | 0.476341379 | 0.673780343 |
| AP53_990 | -0.179977868 | 8.554446163 | 0.54162923 | 0.476610206 | 0.673780343 |
| AP53_735 | 0.174118864 | 8.638929863 | 0.540233222 | 0.477165995 | 0.674075106 |
| AP53_702 | 0.152668336 | 6.892190454 | 0.539351479 | 0.477517619 | 0.674081235 |
| AP53_1779 | -0.215666013 | 8.691103742 | 0.536167388 | 0.478791123 | 0.675120826 |
| AP53_1613 | 0.214719869 | 3.795444941 | 0.535771919 | 0.478949705 | 0.675120826 |
| AP53_363 | -0.083903502 | 9.754538294 | 0.534587517 | 0.479425193 | 0.675300653 |
| AP53_1590 | 0.102536198 | 9.872711188 | 0.52652757 | 0.482682836 | 0.679187782 |
| AP53_153 | 0.097180913 | 10.55554949 | 0.526031323 | 0.482884667 | 0.679187782 |
| AP53_348 | -0.126962992 | 8.534545207 | 0.520900969 | 0.484979932 | 0.681640874 |
| AP53_1619 | 0.154984666 | 6.430822598 | 0.519392533 | 0.485599013 | 0.682017137 |
| AP53_738 | -0.197209175 | 13.6746115 | 0.512870531 | 0.488291753 | 0.684876882 |
| AP53_1071 | -0.149975904 | 7.377320043 | 0.512157733 | 0.488587638 | 0.684876882 |
| AP53_1616 | -0.256721053 | 4.775695083 | 0.511902397 | 0.488693705 | 0.684876882 |
| AP53_487 | -0.135667337 | 9.432426044 | 0.507629484 | 0.490474727 | 0.686876945 |
| AP53_625 | -0.227887261 | 3.57983179 | 0.505437792 | 0.491392713 | 0.687400877 |
| AP53_96 | -0.168078638 | 6.056549802 | 0.505045976 | 0.491557144 | 0.687400877 |
| AP53_788 | 0.118010756 | 5.911765568 | 0.5041448 | 0.491935706 | 0.687434487 |
| AP53_1843 | -0.066464329 | 8.406362663 | 0.503130025 | 0.492362606 | 0.687434487 |
| AP53_1041 | -0.139115785 | 5.849702813 | 0.500610893 | 0.493425208 | 0.687434487 |
| AP53_418 | -0.076305733 | 8.8980063 | 0.500413976 | 0.493508441 | 0.687434487 |
| AP53_1328 | -0.206085464 | 3.983716311 | 0.500060598 | 0.49365787 | 0.687434487 |
| AP53_1202 | -0.163724716 | 3.55737465 | 0.499946419 | 0.493706169 | 0.687434487 |
| AP53_248 | 0.104371744 | 8.921296728 | 0.498436529 | 0.494345658 | 0.687652856 |
| AP53_721 | -0.175059643 | 7.332076412 | 0.497903997 | 0.494571554 | 0.687652856 |
| AP53_1487 | 0.167766603 | 10.25795846 | 0.494862482 | 0.495865269 | 0.68877012 |
| AP53_208 | -0.089263093 | 7.245340904 | 0.494347731 | 0.496084816 | 0.68877012 |
| AP53_1187 | 0.195228954 | 2.279544779 | 0.488531349 | 0.498577667 | 0.691655651 |

| | | | | | |
|---|---|---|---|---|---|
| AP53_1774 | 0.223679687 | 10.47176881 | 0.487839234 | 0.498875792 | 0.691655651 |
| AP53_1793 | -0.227207897 | 5.339455552 | 0.486881666 | 0.499288786 | 0.691734143 |
| AP53_840 | 0.139174675 | 8.092291269 | 0.485626631 | 0.499831004 | 0.691991426 |
| AP53_1530 | 0.178183397 | 7.232502317 | 0.483918188 | 0.50057081 | 0.692521698 |
| AP53_32 | 0.191075914 | 5.687921236 | 0.482399437 | 0.501230125 | 0.692570957 |
| AP53_319 | -0.153969122 | 9.304296752 | 0.482192596 | 0.501320039 | 0.692570957 |
| AP53_1454 | -0.13292019 | 11.049852 | 0.48046031 | 0.502074204 | 0.692773788 |
| AP53_314 | 0.118055933 | 9.769757424 | 0.480216091 | 0.50218069 | 0.692773788 |
| AP53_pseudo1183 | -0.210866901 | 4.144345891 | 0.478006015 | 0.503146203 | 0.69361277 |
| AP53_24 | -0.240004774 | 8.609949346 | 0.476078877 | 0.503990844 | 0.694104724 |
| AP53_483 | -0.090439025 | 9.888218567 | 0.475560977 | 0.50421827 | 0.694104724 |
| AP53_1177 | 0.131688008 | 10.37121398 | 0.473199915 | 0.505257437 | 0.6950423 |
| AP53_647 | -0.244502489 | 3.469571357 | 0.471130705 | 0.506171342 | 0.695449045 |
| AP53_1129 | 0.174193457 | 10.5572049 | 0.470543315 | 0.50643132 | 0.695449045 |
| AP53_956 | 0.190676041 | 7.171513726 | 0.470013257 | 0.506666131 | 0.695449045 |
| AP53_700 | -0.135237392 | 4.841515719 | 0.469157289 | 0.507045733 | 0.695449045 |
| AP53_23 | -0.198807914 | 8.50615647 | 0.468475362 | 0.507348521 | 0.695449045 |
| AP53_rna63 | 0.293567365 | 2.769722909 | 0.467678221 | 0.507702883 | 0.695449045 |
| AP53_732 | 0.12410666 | 6.437125728 | 0.46684059 | 0.50807573 | 0.695468964 |
| AP53_1768 | -0.31669159 | 7.359813173 | 0.463949467 | 0.50936646 | 0.696000831 |
| AP53_1040 | 0.169356663 | 5.15098903 | 0.463560558 | 0.509540543 | 0.696000831 |
| AP53_292 | -0.07080545 | 7.715056802 | 0.46298701 | 0.509797471 | 0.696000831 |
| AP53_34 | 0.244204701 | 5.755964412 | 0.462761402 | 0.5098986 | 0.696000831 |
| AP53_1304 | 0.140421451 | 9.344842128 | 0.461725924 | 0.510363222 | 0.696145477 |
| AP53_1698 | -0.136133699 | 9.395822895 | 0.460227101 | 0.511037118 | 0.696153977 |
| AP53_rna31 | 0.351400683 | 3.575514669 | 0.460116816 | 0.511086768 | 0.696153977 |
| AP53_rna62 | 1.162907745 | 0.675720235 | 0.458743063 | 0.51170597 | 0.696506581 |
| AP53_242 | -0.095592023 | 9.151222099 | 0.457730834 | 0.512163098 | 0.696506581 |
| AP53_842 | -0.093272127 | 7.789003183 | 0.457157938 | 0.512422152 | 0.696506581 |
| AP53_332 | -0.149241326 | 4.479779567 | 0.45218096 | 0.514682807 | 0.699089803 |
| AP53_rna41 | 0.487941173 | 4.30478976 | 0.450201397 | 0.515587068 | 0.699090376 |
| AP53_524 | 0.09037605 | 5.69811416 | 0.449913705 | 0.515718728 | 0.699090376 |
| AP53_1512 | -0.071675559 | 8.214941548 | 0.449815382 | 0.515763739 | 0.699090376 |
| AP53_823 | 0.120078025 | 5.578408491 | 0.448109137 | 0.516545992 | 0.699662087 |
| AP53_392 | 0.553397988 | 0.610601094 | 0.446222235 | 0.517413622 | 0.700081289 |

| | | | | | |
|---|---|---|---|---|---|
| AP53_1320 | -0.228152048 | 3.6637731 | 0.445867921 | 0.517576842 | 0.700081289 |
| AP53_rna19 | 0.375647178 | 4.531004466 | 0.444827555 | 0.518056651 | 0.700242312 |
| AP53_1594 | -0.142940786 | 11.13498404 | 0.440278431 | 0.520164379 | 0.70260199 |
| AP53_512 | 0.106207651 | 5.679056086 | 0.43843231 | 0.521024277 | 0.703274077 |
| AP53_87 | -0.200497139 | 3.194816257 | 0.435842949 | 0.522234829 | 0.703656489 |
| AP53_1414 | -0.124860044 | 8.473384018 | 0.435698222 | 0.522302644 | 0.703656489 |
| AP53_109 | 0.192674658 | 6.207061618 | 0.435500845 | 0.522395157 | 0.703656489 |
| AP53_598 | 0.191469694 | 9.372826546 | 0.432564777 | 0.52377494 | 0.705025769 |
| AP53_71 | -0.202431435 | 5.566105766 | 0.429851419 | 0.525056127 | 0.705354982 |
| AP53_1383 | 0.075035667 | 9.539422849 | 0.429430675 | 0.525255318 | 0.705354982 |
| AP53_rna23 | 0.344830444 | 2.598266444 | 0.428820235 | 0.525544568 | 0.705354982 |
| AP53_1741 | 0.252809311 | 10.54648531 | 0.428266193 | 0.525807352 | 0.705354982 |
| AP53_298 | 0.077676099 | 7.251107526 | 0.426801994 | 0.526503016 | 0.705354982 |
| AP53_1288 | -0.069707265 | 9.249318484 | 0.426172296 | 0.526802727 | 0.705354982 |
| AP53_1544 | 0.108727818 | 11.31501403 | 0.425918348 | 0.526923686 | 0.705354982 |
| AP53_874 | -0.102786025 | 7.794643657 | 0.425245226 | 0.527244557 | 0.705354982 |
| AP53_1465 | 0.061378354 | 8.952617724 | 0.424712778 | 0.527498631 | 0.705354982 |
| AP53_1281 | -0.103185704 | 9.007721286 | 0.424388474 | 0.527653495 | 0.705354982 |
| AP53_200 | -0.194835461 | 10.29106243 | 0.422950847 | 0.528341033 | 0.705359645 |
| AP53_rna65 | 0.3407768 | 4.639511609 | 0.422861573 | 0.528383784 | 0.705359645 |
| AP53_403 | 0.178429548 | 4.275269101 | 0.422034172 | 0.52878031 | 0.705403836 |
| AP53_1029 | -0.232090421 | 5.505463083 | 0.420904377 | 0.529322663 | 0.705642369 |
| AP53_1607 | 0.098409659 | 5.26947702 | 0.418191815 | 0.530629112 | 0.706437596 |
| AP53_1212 | -0.175003256 | 4.201745088 | 0.418072891 | 0.530686529 | 0.706437596 |
| AP53_290 | 0.074806351 | 7.280403496 | 0.41740118 | 0.531011052 | 0.706437596 |
| AP53_802 | -0.088508458 | 6.417002228 | 0.414370148 | 0.53248012 | 0.706836361 |
| AP53_1475 | 0.0901542 | 5.649180463 | 0.41320804 | 0.533045411 | 0.706836361 |
| AP53_147 | -0.077069114 | 5.637764626 | 0.412902644 | 0.533194156 | 0.706836361 |
| AP53_383 | 0.175991969 | 6.861804942 | 0.412808633 | 0.53323996 | 0.706836361 |
| AP53_459 | -0.153689882 | 4.324628536 | 0.412451687 | 0.533413941 | 0.706836361 |
| AP53_1600 | -0.090062533 | 6.597198535 | 0.412283901 | 0.53349576 | 0.706836361 |
| AP53_626 | -0.211795928 | 3.253685203 | 0.410815879 | 0.53421264 | 0.70705205 |
| AP53_1858 | -0.120112248 | 7.645699048 | 0.410459194 | 0.534387098 | 0.70705205 |
| AP53_1820 | -0.17891555 | 6.814986037 | 0.408825644 | 0.535187467 | 0.707376953 |
| AP53_1441 | 0.06601142 | 8.92053845 | 0.408470983 | 0.535361538 | 0.707376953 |

| | | | | | |
|---|---|---|---|---|---|
| AP53_1289 | 0.083451787 | 8.056980692 | 0.405831128 | 0.536660604 | 0.708591178 |
| AP53_1000 | -0.22515744 | 7.176395449 | 0.405121933 | 0.537010625 | 0.708591178 |
| AP53_291 | 0.071895038 | 7.932082255 | 0.403802884 | 0.537662798 | 0.708969763 |
| AP53_1403 | -0.072456491 | 8.232437424 | 0.399570819 | 0.539765537 | 0.711168488 |
| AP53_1248 | 0.145078892 | 9.88017853 | 0.398974616 | 0.540063036 | 0.711168488 |
| AP53_1188 | 0.182413955 | 2.169798669 | 0.396784447 | 0.541158619 | 0.712128055 |
| AP53_1664 | -0.084723529 | 7.418852833 | 0.395597686 | 0.541754062 | 0.712428614 |
| AP53_160 | -0.189196863 | 4.973827245 | 0.393305919 | 0.542907515 | 0.712850587 |
| AP53_1039 | 0.118330603 | 5.831925291 | 0.393149787 | 0.54298627 | 0.712850587 |
| AP53_1864 | -0.157567788 | 9.373845942 | 0.392054033 | 0.543539601 | 0.712850587 |
| AP53_145 | 0.109536594 | 5.127435458 | 0.392045365 | 0.543543982 | 0.712850587 |
| AP53_339 | 0.061945759 | 6.952917524 | 0.39059791 | 0.544276602 | 0.713329429 |
| AP53_1115 | -0.169480463 | 5.058997186 | 0.388894818 | 0.545141068 | 0.71375707 |
| AP53_1794 | -0.173149003 | 5.449043881 | 0.388506887 | 0.545338349 | 0.71375707 |
| AP53_1659 | 0.187220785 | 9.560678367 | 0.384568629 | 0.54734903 | 0.715526184 |
| AP53_320 | 0.078751203 | 8.206756372 | 0.383589921 | 0.547850953 | 0.715526184 |
| AP53_236 | -0.115451019 | 8.985063389 | 0.383422568 | 0.547936869 | 0.715526184 |
| AP53_1726 | 0.293907028 | 9.766519192 | 0.382979278 | 0.548164573 | 0.715526184 |
| AP53_1169 | 0.119425894 | 6.773293348 | 0.382042937 | 0.548646149 | 0.715673505 |
| AP53_1196 | -0.230815058 | 3.612372154 | 0.380184951 | 0.549604198 | 0.716441738 |
| AP53_357 | 0.073810848 | 9.9521086 | 0.373411744 | 0.553124699 | 0.720547007 |
| AP53_1632 | 0.078844301 | 7.214749221 | 0.368644557 | 0.555629308 | 0.72332427 |
| AP53_296 | 0.11310007 | 7.707430947 | 0.367263628 | 0.556359028 | 0.723788789 |
| AP53_129 | 0.08169678 | 8.445112756 | 0.36543036 | 0.557330722 | 0.724206172 |
| AP53_773 | -0.088506043 | 9.386672227 | 0.364708779 | 0.55771411 | 0.724206172 |
| AP53_1618 | -0.087963318 | 7.218910504 | 0.364548784 | 0.557799189 | 0.724206172 |
| AP53_1353 | -0.17808747 | 4.458065608 | 0.360271608 | 0.560083261 | 0.726358227 |
| AP53_1395 | -0.214640961 | 7.290890495 | 0.360044269 | 0.560205186 | 0.726358227 |
| AP53_353 | -0.085200141 | 9.770017578 | 0.3580227 | 0.561291724 | 0.727035363 |
| AP53_330 | -0.130723072 | 10.15185276 | 0.357679576 | 0.561476563 | 0.727035363 |
| AP53_1767 | -0.094378914 | 9.051988918 | 0.356118364 | 0.562319125 | 0.727640947 |
| AP53_1415 | -0.078278335 | 8.159083617 | 0.354773525 | 0.563046952 | 0.728097358 |
| AP53_1283 | -0.074404613 | 8.006384195 | 0.353147683 | 0.563929395 | 0.728752966 |
| AP53_281 | -0.067692641 | 8.123607792 | 0.351806655 | 0.564659352 | 0.729149678 |
| AP53_630 | -0.132241608 | 4.836682553 | 0.349945468 | 0.565675614 | 0.729149678 |

| | | | | | |
|---|---|---|---|---|---|
| AP53_311 | -0.129297189 | 12.01622451 | 0.349128071 | 0.566123105 | 0.729149678 |
| AP53_1106 | 0.064617906 | 7.140266528 | 0.349120623 | 0.566127186 | 0.729149678 |
| AP53_924 | 0.09049363 | 8.901812604 | 0.348210141 | 0.566626491 | 0.729149678 |
| AP53_1171 | -0.110642303 | 6.530184668 | 0.348070735 | 0.566703019 | 0.729149678 |
| AP53_1775 | -0.13055586 | 11.20879042 | 0.347774011 | 0.566865979 | 0.729149678 |
| AP53_1641 | -0.085262166 | 8.263963792 | 0.344564572 | 0.568634689 | 0.730915242 |
| AP53_699 | 0.119816894 | 5.236852529 | 0.343919176 | 0.568991721 | 0.730915242 |
| AP53_638 | 0.133745505 | 3.705966894 | 0.342350628 | 0.569861346 | 0.731548196 |
| AP53_404 | 0.178337992 | 3.441721999 | 0.340451365 | 0.570917961 | 0.732046601 |
| AP53_192 | 0.088247898 | 9.179226164 | 0.34029722 | 0.571003892 | 0.732046601 |
| AP53_1276 | -0.115742189 | 5.528557162 | 0.338285774 | 0.572127642 | 0.733003138 |
| AP53_761 | 0.10422125 | 8.406277453 | 0.33733175 | 0.572662221 | 0.73320407 |
| AP53_544 | -0.101811708 | 8.092722271 | 0.335890469 | 0.573471777 | 0.733559598 |
| AP53_1097 | 0.100114095 | 6.178719169 | 0.335492443 | 0.573695759 | 0.733559598 |
| AP53_1456 | -0.068987535 | 9.922923369 | 0.331215282 | 0.576114088 | 0.73616685 |
| AP53_814 | 0.082170165 | 5.701081 | 0.33051105 | 0.576514284 | 0.73619357 |
| AP53_999 | 0.082990197 | 7.754957157 | 0.326405391 | 0.578858938 | 0.738516831 |
| AP53_583 | 0.211446152 | 11.23882585 | 0.32557629 | 0.579334826 | 0.738516831 |
| AP53_1191 | -0.155393098 | 2.921683598 | 0.32511663 | 0.579599013 | 0.738516831 |
| AP53_419 | -0.065363572 | 9.639050574 | 0.324670672 | 0.579855565 | 0.738516831 |
| AP53_rna84 | 0.302119287 | 1.78152497 | 0.323961302 | 0.580264141 | 0.738552588 |
| AP53_1499 | -0.087379722 | 9.780226003 | 0.323299688 | 0.580645752 | 0.738554001 |
| AP53_1 | -0.082364443 | 7.944552078 | 0.321856027 | 0.581480263 | 0.7391311 |
| AP53_254 | -0.112458675 | 8.141883276 | 0.32052201 | 0.582253626 | 0.73962977 |
| AP53_1280 | -0.085686178 | 9.75997479 | 0.318290254 | 0.583552264 | 0.74078355 |
| AP53_1048 | -0.141984339 | 6.144025653 | 0.31765126 | 0.583925209 | 0.74078355 |
| AP53_1711 | -0.199303023 | 3.491323419 | 0.315871061 | 0.584966862 | 0.741620299 |
| AP53_1435 | -0.080027967 | 8.804048446 | 0.314119917 | 0.585995342 | 0.742439268 |
| AP53_994 | 0.087953464 | 10.55661971 | 0.312350802 | 0.587038262 | 0.743174063 |
| AP53_1108 | 0.064521221 | 8.456743553 | 0.311838395 | 0.587341068 | 0.743174063 |
| AP53_1759 | -0.101583578 | 6.652835082 | 0.31060226 | 0.588072924 | 0.743615339 |
| AP53_391 | 0.1081223 | 5.127292272 | 0.309903397 | 0.588487544 | 0.743655158 |
| AP53_1277 | 0.119981716 | 10.15144344 | 0.3066024 | 0.590454384 | 0.745655146 |
| AP53_695 | 0.106760469 | 8.830345711 | 0.303448005 | 0.592347024 | 0.747558891 |
| AP53_1819 | -0.127728543 | 6.604633592 | 0.301813495 | 0.593332851 | 0.74831648 |

| | | | | | |
|---|---|---|---|---|---|
| AP53_366 | -0.10351904 | 8.651241561 | 0.300827185 | 0.593929432 | 0.748582485 |
| AP53_111 | -0.128239261 | 6.855954303 | 0.294974899 | 0.597496023 | 0.752589085 |
| AP53_186 | 0.108747445 | 8.362564137 | 0.293586778 | 0.598348803 | 0.753174466 |
| AP53_1324 | 0.170027871 | 3.836125107 | 0.292227829 | 0.599186227 | 0.753476427 |
| AP53_954 | -0.154478928 | 9.975002689 | 0.291404787 | 0.59969465 | 0.753476427 |
| AP53_1719 | 0.091936117 | 6.55302028 | 0.291310004 | 0.599753261 | 0.753476427 |
| AP53_407 | -0.070254974 | 7.994504104 | 0.289056136 | 0.601150683 | 0.754126368 |
| AP53_1612 | 0.102038904 | 4.415013354 | 0.288795908 | 0.601312485 | 0.754126368 |
| AP53_204 | 0.068255914 | 6.544829362 | 0.288597079 | 0.601436176 | 0.754126368 |
| AP53_1655 | -0.088777064 | 9.654728026 | 0.284837022 | 0.603785826 | 0.756226968 |
| AP53_rna20 | 0.322735573 | 4.503413543 | 0.284669984 | 0.603890675 | 0.756226968 |
| AP53_1781 | -0.101141348 | 9.060982847 | 0.283295211 | 0.604755142 | 0.756821232 |
| AP53_253 | -0.06091304 | 7.304285585 | 0.28176969 | 0.605717598 | 0.75753728 |
| AP53_88 | -0.134083723 | 3.287412157 | 0.280444409 | 0.606556475 | 0.757825005 |
| AP53_1077 | 0.131009097 | 6.908046039 | 0.280173108 | 0.60672852 | 0.757825005 |
| AP53_rna17 | 0.848605346 | -0.659752514 | 0.279521609 | 0.607142109 | 0.757853912 |
| AP53_441 | 0.1185768 | 4.005282735 | 0.276728721 | 0.608922223 | 0.759587426 |
| AP53_1802 | 0.161093124 | 3.801703709 | 0.271652051 | 0.61218794 | 0.763170708 |
| AP53_1540 | 0.05139525 | 9.211085047 | 0.270862969 | 0.612699066 | 0.763184433 |
| AP53_901 | -0.059368906 | 7.690586902 | 0.270382789 | 0.613010571 | 0.763184433 |
| AP53_1442 | 0.073026616 | 8.913391452 | 0.2696899 | 0.613460694 | 0.763184433 |
| AP53_1495 | 0.089034928 | 9.023869357 | 0.268994041 | 0.613913495 | 0.763184433 |
| AP53_1176 | 0.136749111 | 10.08705089 | 0.268608174 | 0.614164907 | 0.763184433 |
| AP53_1507 | 0.085875201 | 8.566607648 | 0.266791605 | 0.61535162 | 0.764169863 |
| AP53_570 | -0.107615723 | 9.326678034 | 0.264894692 | 0.616596368 | 0.765226056 |
| AP53_139 | 0.118148319 | 6.850934411 | 0.263967793 | 0.617206675 | 0.76549403 |
| AP53_pseudo1680 | -0.109485194 | 5.028817522 | 0.263340563 | 0.617620447 | 0.765518064 |
| AP53_530 | -0.075822146 | 9.094709856 | 0.262247557 | 0.618342991 | 0.765924535 |
| AP53_1145 | -0.118823384 | 7.188093254 | 0.261491494 | 0.618843924 | 0.766056158 |
| AP53_599 | -0.272509968 | 9.392532048 | 0.260795168 | 0.619306096 | 0.766139663 |
| AP53_rna73 | 0.122486015 | 5.936613833 | 0.259271798 | 0.620319957 | 0.766905119 |
| AP53_857 | -0.067784423 | 8.139744154 | 0.255074141 | 0.623133453 | 0.769478575 |
| AP53_775 | -0.070800948 | 9.194484888 | 0.2549837 | 0.623194395 | 0.769478575 |
| AP53_854 | -0.082498564 | 6.218076921 | 0.252028172 | 0.62519351 | 0.771073957 |
| AP53_230 | -0.120147267 | 5.128972314 | 0.251899335 | 0.625280994 | 0.771073957 |

| | | | | | |
|---|---|---|---|---|---|
| AP53_1434 | 0.044257841 | 7.360167075 | 0.249443591 | 0.62695394 | 0.772646093 |
| AP53_1792 | -0.190959614 | 4.794705572 | 0.248442548 | 0.627638879 | 0.772999406 |
| AP53_646 | -0.126649751 | 3.949294806 | 0.247129783 | 0.628539756 | 0.773618051 |
| AP53_1190 | 0.141250513 | 2.186103157 | 0.241506076 | 0.632433542 | 0.777917303 |
| AP53_1457 | -0.056503873 | 10.3336213 | 0.240000499 | 0.633485649 | 0.778717952 |
| AP53_217 | -0.066045984 | 8.032206253 | 0.239001015 | 0.634186386 | 0.779085934 |
| AP53_482 | -0.07427398 | 7.245107736 | 0.237364703 | 0.635337582 | 0.77960923 |
| AP53_1363 | 0.115028833 | 9.67730007 | 0.237253973 | 0.635415663 | 0.77960923 |
| AP53_191 | 0.065118391 | 8.470630369 | 0.23557908 | 0.636599509 | 0.780568317 |
| AP53_1813 | -0.081674391 | 6.865511878 | 0.233671548 | 0.637954217 | 0.781249924 |
| AP53_745 | -0.143951638 | 5.133878848 | 0.233217906 | 0.638277405 | 0.781249924 |
| AP53_841 | 0.077009691 | 8.131143388 | 0.233098 | 0.638362895 | 0.781249924 |
| AP53_1335 | -0.177537968 | 3.024041073 | 0.23210963 | 0.639068627 | 0.781620797 |
| AP53_658 | -0.10108434 | 6.883404077 | 0.230579112 | 0.64016518 | 0.7824689 |
| AP53_629 | 0.153337861 | 3.905552694 | 0.223161721 | 0.645544595 | 0.788151562 |
| AP53_810 | -0.073040376 | 7.328800618 | 0.223049985 | 0.645626473 | 0.788151562 |
| AP53_891 | -0.074802836 | 11.92681912 | 0.219105212 | 0.648533521 | 0.791202743 |
| AP53_293 | 0.057403831 | 7.345830115 | 0.218378376 | 0.64907267 | 0.791363099 |
| AP53_1830 | 0.066088007 | 5.675547829 | 0.216339943 | 0.650590657 | 0.792277867 |
| AP53_1151 | -0.059750059 | 7.822189144 | 0.21627479 | 0.65063932 | 0.792277867 |
| AP53_1788 | -0.121808612 | 7.228241139 | 0.214421023 | 0.65202771 | 0.793470711 |
| AP53_1840 | 0.094678982 | 8.984295955 | 0.213138821 | 0.652992343 | 0.794146703 |
| AP53_1046 | 0.133681278 | 5.864546982 | 0.211345835 | 0.654347243 | 0.795232831 |
| AP53_737 | 0.052177045 | 7.671548301 | 0.210874183 | 0.654704824 | 0.795232831 |
| AP53_564 | -0.127291876 | 10.08214382 | 0.209763712 | 0.655548664 | 0.795637561 |
| AP53_1778 | 0.102699587 | 9.281653972 | 0.208676588 | 0.656377414 | 0.795637561 |
| AP53_918 | 0.110894537 | 9.936688656 | 0.208315151 | 0.656653534 | 0.795637561 |
| AP53_rna34 | 0.244590501 | 3.37489484 | 0.20827142 | 0.656686962 | 0.795637561 |
| AP53_1509 | -0.065961541 | 7.486170921 | 0.207747753 | 0.657087589 | 0.795637561 |
| AP53_112 | -0.093030769 | 5.848035247 | 0.204973771 | 0.659220153 | 0.797722143 |
| AP53_1846 | 0.064762127 | 6.160194467 | 0.204350616 | 0.659701637 | 0.7978074 |
| AP53_80 | 0.140209039 | 8.787198023 | 0.201113861 | 0.662217016 | 0.800350702 |
| AP53_1302 | 0.082491852 | 7.945785161 | 0.199777091 | 0.663263038 | 0.800845192 |
| AP53_673 | 0.055391339 | 9.971696134 | 0.199185854 | 0.663727037 | 0.800845192 |
| AP53_1059 | 0.042689034 | 6.906797958 | 0.199011607 | 0.663863944 | 0.800845192 |

| | | | | | |
|---|---|---|---|---|---|
| AP53_371 | 0.066313522 | 8.04646574 | 0.198231328 | 0.664477907 | 0.801087961 |
| AP53_1061 | -0.086579909 | 5.182023961 | 0.196501247 | 0.665844464 | 0.802237184 |
| AP53_462 | -0.069338433 | 8.914955832 | 0.193558602 | 0.668185617 | 0.804558487 |
| AP53_370 | 0.077319854 | 7.3926236 | 0.192705667 | 0.66886822 | 0.804881101 |
| AP53_1520 | -0.05411156 | 7.600032679 | 0.189456454 | 0.671485379 | 0.807529814 |
| AP53_1033 | 0.163843855 | 5.62939968 | 0.187685419 | 0.672923284 | 0.808757953 |
| AP53_785 | -0.047847778 | 6.737096775 | 0.186825619 | 0.673624296 | 0.80909948 |
| AP53_1156 | -0.042867067 | 7.151800104 | 0.181918203 | 0.677662902 | 0.812762435 |
| AP53_245 | 0.065424731 | 5.525607065 | 0.181674094 | 0.677865487 | 0.812762435 |
| AP53_1272 | -0.082936791 | 7.465339886 | 0.181596241 | 0.67793013 | 0.812762435 |
| AP53_816 | -0.080845692 | 13.52738783 | 0.17824892 | 0.680725306 | 0.815609764 |
| AP53_1574 | -0.058887138 | 6.894513115 | 0.175663709 | 0.682905551 | 0.816907052 |
| AP53_1068 | 0.083077769 | 6.318886915 | 0.175610429 | 0.682950685 | 0.816907052 |
| AP53_1674 | -0.08084791 | 7.214747101 | 0.174986319 | 0.683479979 | 0.816907052 |
| AP53_1762 | 0.109592813 | 5.868500859 | 0.174731764 | 0.683696183 | 0.816907052 |
| AP53_241 | -0.065623337 | 9.196140432 | 0.17447742 | 0.683912396 | 0.816907052 |
| AP53_1364 | 0.0788565 | 8.777863786 | 0.171705237 | 0.686281183 | 0.819232335 |
| AP53_1278 | 0.064658989 | 9.295328339 | 0.169802113 | 0.687920511 | 0.820356641 |
| AP53_755 | 0.05014535 | 8.12430739 | 0.169631135 | 0.688068321 | 0.820356641 |
| AP53_170 | -0.102288624 | 9.354608329 | 0.167995906 | 0.689486431 | 0.821273462 |
| AP53_427 | -0.068048841 | 4.410117983 | 0.167769363 | 0.689683536 | 0.821273462 |
| AP53_799 | 0.069325361 | 7.171421152 | 0.164162571 | 0.692843012 | 0.823065655 |
| AP53_322 | 0.099227235 | 3.828451019 | 0.164087513 | 0.692909193 | 0.823065655 |
| AP53_963 | -0.058628833 | 7.67750797 | 0.163901444 | 0.693073333 | 0.823065655 |
| AP53_1074 | 0.084638691 | 9.483726773 | 0.163784081 | 0.693176921 | 0.823065655 |
| AP53_1730 | 0.04484444 | 8.408576889 | 0.163478787 | 0.693446586 | 0.823065655 |
| AP53_181 | -0.091365396 | 6.519016854 | 0.162808666 | 0.69403954 | 0.823065655 |
| AP53_1687 | -0.045714061 | 6.531062586 | 0.162676264 | 0.694156866 | 0.823065655 |
| AP53_1718 | -0.055218441 | 6.728872894 | 0.160437345 | 0.696149381 | 0.824924266 |
| AP53_104 | -0.047171237 | 7.441903889 | 0.158298267 | 0.698068322 | 0.826693479 |
| AP53_657 | -0.063078973 | 7.849295204 | 0.157234357 | 0.699028395 | 0.827325679 |
| AP53_1769 | 0.056995153 | 9.866497896 | 0.153358046 | 0.702558834 | 0.830997378 |
| AP53_1255 | -0.057769289 | 8.539361175 | 0.147131249 | 0.708340456 | 0.837325716 |
| AP53_1842 | -0.056113221 | 6.89698383 | 0.146598366 | 0.708841781 | 0.837408337 |
| AP53_1852 | -0.04005641 | 7.54075747 | 0.145761198 | 0.709631506 | 0.837831359 |

| | | | | | |
|---|---|---|---|---|---|
| AP53_232 | 0.057276328 | 8.703000013 | 0.145197126 | 0.710165091 | 0.837951636 |
| AP53_632 | -0.080060192 | 4.285425534 | 0.144179357 | 0.711130891 | 0.838581446 |
| AP53_1056 | -0.048007489 | 7.167086984 | 0.141328067 | 0.713857727 | 0.841285882 |
| AP53_730 | -0.060984674 | 7.593321494 | 0.140681989 | 0.714479999 | 0.8415083 |
| AP53_1821 | 0.142262736 | 5.51795796 | 0.138940277 | 0.716165773 | 0.842795578 |
| AP53_939 | 0.090456923 | 9.212780987 | 0.138656719 | 0.716441372 | 0.842795578 |
| AP53_pseudo1695 | -0.095193426 | 11.9771486 | 0.137508251 | 0.717560928 | 0.843601309 |
| AP53_220 | -0.077832098 | 11.41052102 | 0.136786035 | 0.718267712 | 0.843921083 |
| AP53_304 | 0.054691727 | 7.635473271 | 0.135352226 | 0.719677255 | 0.84506567 |
| AP53_1397 | 0.051772791 | 8.742721169 | 0.133605569 | 0.721405952 | 0.846583405 |
| AP53_1376 | 0.04978384 | 6.677968156 | 0.131707779 | 0.723298941 | 0.84829199 |
| AP53_1069 | 0.089910853 | 5.517887881 | 0.129656305 | 0.725362819 | 0.849059589 |
| AP53_1019 | -0.148352858 | 7.874398624 | 0.129583762 | 0.72543614 | 0.849059589 |
| AP53_458 | -0.059174771 | 3.931775774 | 0.129227534 | 0.72579653 | 0.849059589 |
| AP53_498 | -0.442103857 | -1.006521509 | 0.1303382 | 0.726050596 | 0.849059589 |
| AP53_rna10 | -0.286106833 | 2.562764711 | 0.128887951 | 0.726140607 | 0.849059589 |
| AP53_21 | -0.088292395 | 6.210596514 | 0.128364806 | 0.726671686 | 0.849169021 |
| AP53_1677 | 0.045774614 | 7.899458382 | 0.127231696 | 0.727826221 | 0.850006435 |
| AP53_1096 | 0.057853827 | 5.522855 | 0.125912129 | 0.729178126 | 0.850460677 |
| AP53_185 | -0.083552996 | 8.235030862 | 0.125592931 | 0.729506355 | 0.850460677 |
| AP53_1524 | -0.063990483 | 10.72683241 | 0.125570306 | 0.729529638 | 0.850460677 |
| AP53_1592 | 0.041036423 | 9.927094547 | 0.124306817 | 0.73083367 | 0.850986051 |
| AP53_190 | -0.039868273 | 8.865796182 | 0.124284125 | 0.73085716 | 0.850986051 |
| AP53_715 | 0.129288984 | 3.336580757 | 0.12376346 | 0.731396773 | 0.851103799 |
| AP53_1417 | 0.044738986 | 8.884941693 | 0.120196722 | 0.735128368 | 0.85493359 |
| AP53_1513 | -0.038736024 | 7.803939915 | 0.119207586 | 0.736174277 | 0.855021925 |
| AP53_1783 | -0.070842036 | 8.552077433 | 0.118951936 | 0.736445395 | 0.855021925 |
| AP53_38 | -0.047785603 | 8.772914564 | 0.118876139 | 0.736525841 | 0.855021925 |
| AP53_752 | 0.038488505 | 9.431559898 | 0.117600358 | 0.737884244 | 0.855720501 |
| AP53_1042 | 0.066183036 | 6.407160573 | 0.116790301 | 0.738751075 | 0.855720501 |
| AP53_4 | 0.0704617 | 4.818853348 | 0.11675666 | 0.738787147 | 0.855720501 |
| AP53_437 | -0.067771501 | 6.32848962 | 0.116659774 | 0.738891066 | 0.855720501 |
| AP53_rna57 | -0.103911038 | 5.964443146 | 0.115393835 | 0.740253395 | 0.856787024 |
| AP53_720 | -0.12120201 | 6.455922938 | 0.113889288 | 0.741883471 | 0.858161989 |
| AP53_149 | -0.084903178 | 11.85960658 | 0.111072884 | 0.744967653 | 0.861216328 |

| | | | | | |
|---|---|---|---|---|---|
| AP53_1678 | 0.053203627 | 8.21689848 | 0.108440103 | 0.747890599 | 0.862919135 |
| AP53_rna32 | 0.185781197 | 4.160629324 | 0.108368598 | 0.747970536 | 0.862919135 |
| AP53_991 | 0.06105551 | 11.37384296 | 0.108102597 | 0.748268159 | 0.862919135 |
| AP53_1223 | 0.055439165 | 8.566335184 | 0.107753577 | 0.748659292 | 0.862919135 |
| AP53_356 | -0.04939158 | 8.106574505 | 0.107749839 | 0.748663484 | 0.862919135 |
| AP53_566 | -0.085836561 | 10.8465092 | 0.106362216 | 0.750225599 | 0.864206462 |
| AP53_rna18 | -0.161490381 | 4.238330892 | 0.105586604 | 0.751103697 | 0.864263662 |
| AP53_1253 | 0.052942035 | 4.832420676 | 0.105531878 | 0.751165789 | 0.864263662 |
| AP53_557 | -0.040985966 | 7.132158754 | 0.104771036 | 0.752030912 | 0.864338021 |
| AP53_143 | -0.050941006 | 4.941427565 | 0.104691959 | 0.752121029 | 0.864338021 |
| AP53_240 | -0.0374243 | 8.982834402 | 0.104129976 | 0.752762554 | 0.864563383 |
| AP53_1284 | -0.065507515 | 6.949306293 | 0.101799953 | 0.755443111 | 0.867128964 |
| AP53_862 | -0.064632434 | 5.314573916 | 0.100577544 | 0.756863052 | 0.867699864 |
| AP53_279 | -0.044908582 | 10.25874936 | 0.100361214 | 0.757115334 | 0.867699864 |
| AP53_42 | 0.076113741 | 10.79703521 | 0.099954611 | 0.757590327 | 0.867699864 |
| AP53_13 | -0.055717845 | 9.357275859 | 0.099836394 | 0.757728629 | 0.867699864 |
| AP53_1199 | -0.118085534 | 2.890927429 | 0.099143989 | 0.758540498 | 0.868117398 |
| AP53_1045 | 0.05830422 | 6.293677744 | 0.098099122 | 0.759771604 | 0.869013956 |
| AP53_1064 | -0.06336231 | 6.36318039 | 0.097558689 | 0.760411211 | 0.86923331 |
| AP53_1770 | 0.040842634 | 7.59429478 | 0.096127255 | 0.762114843 | 0.869928897 |
| AP53_1650 | -0.092837673 | 6.99247406 | 0.09608014 | 0.762171155 | 0.869928897 |
| AP53_337 | 0.040399606 | 5.666366932 | 0.095918658 | 0.762364273 | 0.869928897 |
| AP53_993 | 0.057505449 | 11.49556352 | 0.095098451 | 0.763347946 | 0.870391779 |
| AP53_623 | -0.084563226 | 3.912249943 | 0.094833443 | 0.76366677 | 0.870391779 |
| AP53_rna67 | 0.121476013 | 4.349876004 | 0.093697624 | 0.765038833 | 0.871443881 |
| AP53_568 | -0.036534949 | 9.895918157 | 0.092869232 | 0.76604531 | 0.872078561 |
| AP53_26 | -0.124882831 | 9.770366314 | 0.091824738 | 0.76732141 | 0.87301926 |
| AP53_1776 | 0.034854454 | 10.17093042 | 0.089609241 | 0.770054826 | 0.875339321 |
| AP53_409 | 0.037551273 | 9.444312881 | 0.08935337 | 0.77037289 | 0.875339321 |
| AP53_1727 | -0.032924128 | 9.545110216 | 0.089079808 | 0.770713498 | 0.875339321 |
| AP53_2 | -0.039638861 | 8.274963851 | 0.087039935 | 0.773271583 | 0.877731077 |
| AP53_729 | -0.040583955 | 7.986503413 | 0.086349344 | 0.774145034 | 0.878208949 |
| AP53_1529 | 0.034978818 | 8.227547356 | 0.084705053 | 0.7762402 | 0.880071395 |
| AP53_1136 | -0.084543688 | 5.191885985 | 0.083938367 | 0.777224706 | 0.880272482 |
| AP53_43 | 0.088433373 | 10.88048974 | 0.083860793 | 0.777324592 | 0.880272482 |

| | | | | | |
|---|---|---|---|---|---|
| AP53_1466 | -0.048452825 | 9.729073719 | 0.081937125 | 0.779817844 | 0.882581011 |
| AP53_1362 | 0.062564596 | 9.411929701 | 0.081247288 | 0.780719689 | 0.883086782 |
| AP53_786 | -0.083285753 | 3.925446239 | 0.079862607 | 0.782542579 | 0.884633166 |
| AP53_767 | -0.032764987 | 6.583723387 | 0.078981027 | 0.78371211 | 0.885305121 |
| AP53_102 | -0.048285518 | 6.187645192 | 0.078727915 | 0.784049203 | 0.885305121 |
| AP53_1117 | 0.048498036 | 8.568234233 | 0.078270321 | 0.784660122 | 0.885479824 |
| AP53_433 | -0.036648502 | 8.225934195 | 0.077160356 | 0.786150091 | 0.885864694 |
| AP53_434 | -0.029213262 | 8.537490947 | 0.077064957 | 0.786278692 | 0.885864694 |
| AP53_593 | 0.0375384 | 6.727169012 | 0.076996994 | 0.786370359 | 0.885864694 |
| AP53_1127 | -0.037547319 | 9.163542651 | 0.07508144 | 0.788972293 | 0.887186638 |
| AP53_1506 | 0.027941376 | 9.598567028 | 0.075009022 | 0.78907136 | 0.887186638 |
| AP53_174 | -0.047710338 | 6.388664455 | 0.074958281 | 0.789140805 | 0.887186638 |
| AP53_1467 | 0.035446629 | 9.113828949 | 0.07476027 | 0.789412046 | 0.887186638 |
| AP53_198 | 0.040865287 | 10.34227161 | 0.074456279 | 0.789829217 | 0.887186638 |
| AP53_410 | -0.05221239 | 9.975201927 | 0.072197989 | 0.792957522 | 0.889954046 |
| AP53_569 | -0.034399223 | 8.07311659 | 0.072017403 | 0.793209943 | 0.889954046 |
| AP53_29 | -0.097837474 | 9.432280448 | 0.071171265 | 0.794397238 | 0.890394186 |
| AP53_rna66 | 0.116242092 | 4.587209041 | 0.070764159 | 0.794971195 | 0.890394186 |
| AP53_1827 | 0.065264007 | 7.738498587 | 0.070607886 | 0.795191988 | 0.890394186 |
| AP53_1116 | -0.035095838 | 7.551347705 | 0.070434579 | 0.795437155 | 0.890394186 |
| AP53_1661 | 0.048701382 | 5.406063771 | 0.068744051 | 0.797845867 | 0.892061778 |
| AP53_839 | 0.036110102 | 9.240711618 | 0.0687439 | 0.797846083 | 0.892061778 |
| AP53_1489 | -0.063137518 | 10.06214315 | 0.067904422 | 0.799054026 | 0.892526422 |
| AP53_1790 | -0.09013074 | 4.848679332 | 0.067816287 | 0.799181309 | 0.892526422 |
| AP53_804 | 0.054152797 | 8.79923924 | 0.066538246 | 0.801037107 | 0.894084545 |
| AP53_1491 | 0.061132761 | 14.69533013 | 0.064949986 | 0.803370186 | 0.896173294 |
| AP53_1275 | -0.04236548 | 6.163761658 | 0.064263674 | 0.80438779 | 0.896793051 |
| AP53_868 | -0.03892203 | 8.622066787 | 0.063552142 | 0.805448944 | 0.897333917 |
| AP53_1067 | -0.039079102 | 5.709482512 | 0.063165715 | 0.806027908 | 0.897333917 |
| AP53_135 | -0.031463688 | 8.773173766 | 0.062875813 | 0.806463498 | 0.897333917 |
| AP53_1627 | -0.095761313 | 2.557416665 | 0.06270401 | 0.806722145 | 0.897333917 |
| AP53_28 | -0.089610278 | 9.347749304 | 0.061390697 | 0.808711944 | 0.899032006 |
| AP53_1037 | -0.056370484 | 5.399904477 | 0.060096756 | 0.810694715 | 0.900381013 |
| AP53_1755 | -0.034550684 | 11.0368012 | 0.05999398 | 0.810853174 | 0.900381013 |
| AP53_540 | -0.048527741 | 12.75185578 | 0.059165773 | 0.81213541 | 0.901289212 |

| | | | | | |
|---|---|---|---|---|---|
| AP53_rna4 | 0.307365165 | -0.831064152 | 0.058000581 | 0.813955642 | 0.902056712 |
| AP53_659 | 0.035910507 | 7.44072087 | 0.057884633 | 0.814137832 | 0.902056712 |
| AP53_713 | 0.05899944 | 2.030550548 | 0.057831614 | 0.814221205 | 0.902056712 |
| AP53_1047 | 0.061149714 | 6.075203301 | 0.056230158 | 0.81675894 | 0.904352026 |
| AP53_1100 | -0.046950776 | 6.426479153 | 0.055504962 | 0.817920725 | 0.905122079 |
| AP53_411 | -0.035530578 | 10.31818213 | 0.053825928 | 0.820641841 | 0.907265678 |
| AP53_724 | 0.033721174 | 7.381698237 | 0.053733673 | 0.820792648 | 0.907265678 |
| AP53_1399 | 0.036517433 | 10.44428353 | 0.053071598 | 0.821878963 | 0.907949384 |
| AP53_1213 | -0.043802321 | 7.298142231 | 0.051868464 | 0.823871528 | 0.909624816 |
| AP53_212 | -0.023115317 | 9.368271815 | 0.051520453 | 0.824452428 | 0.909624816 |
| AP53_1189 | 0.059831143 | 1.973860912 | 0.051182773 | 0.825018066 | 0.909624816 |
| AP53_1006 | -0.094279629 | 6.091945859 | 0.050758451 | 0.825731635 | 0.909624816 |
| AP53_594 | -0.031811651 | 6.746826299 | 0.050754226 | 0.825738756 | 0.909624816 |
| AP53_1251 | -0.053882711 | 6.28830736 | 0.050064537 | 0.826905381 | 0.910393275 |
| AP53_670 | -0.023767337 | 8.14152311 | 0.049357533 | 0.828110122 | 0.911044472 |
| AP53_836 | -0.027128123 | 9.379810407 | 0.049167437 | 0.828435596 | 0.911044472 |
| AP53_1101 | -0.028913801 | 6.992606094 | 0.04878774 | 0.829087685 | 0.911245298 |
| AP53_1656 | 0.028497407 | 9.178346442 | 0.048074445 | 0.830319957 | 0.912083213 |
| AP53_1215 | 0.044599424 | 6.297480204 | 0.047356138 | 0.831570646 | 0.912940398 |
| AP53_450 | 0.068666269 | 1.699702403 | 0.046545355 | 0.832994392 | 0.913986498 |
| AP53_1504 | 0.032582492 | 6.423323182 | 0.045243626 | 0.835307759 | 0.915119468 |
| AP53_1837 | -0.030762125 | 8.456360843 | 0.045120347 | 0.835528644 | 0.915119468 |
| AP53_1282 | -0.077883209 | 8.051671386 | 0.044870861 | 0.835976634 | 0.915119468 |
| AP53_1575 | 0.028828037 | 8.329307005 | 0.044753193 | 0.836188376 | 0.915119468 |
| AP53_1523 | -0.039628959 | 4.841995829 | 0.044644457 | 0.836384305 | 0.915119468 |
| AP53_1209 | 0.078414155 | 2.866562843 | 0.043496053 | 0.838469054 | 0.915837184 |
| AP53_150 | 0.064408163 | 12.31151895 | 0.04346595 | 0.838524087 | 0.915837184 |
| AP53_946 | 0.034242421 | 8.691793309 | 0.043279195 | 0.838865948 | 0.915837184 |
| AP53_1649 | 0.045878113 | 7.311415974 | 0.043245549 | 0.838927621 | 0.915837184 |
| AP53_6 | -0.040918138 | 5.955464852 | 0.041685491 | 0.84181525 | 0.918472963 |
| AP53_1816 | -0.037191917 | 5.463694811 | 0.040563479 | 0.843927149 | 0.919848703 |
| AP53_515 | 0.027439198 | 6.975175126 | 0.040512419 | 0.844023978 | 0.919848703 |
| AP53_1712 | 0.042674318 | 5.106717319 | 0.039919221 | 0.8451536 | 0.920562928 |
| AP53_703 | 0.045401089 | 6.57625008 | 0.039536953 | 0.845886195 | 0.920844142 |
| AP53_806 | 0.037967579 | 6.347657363 | 0.039228342 | 0.84648033 | 0.920974395 |

| | | | | | |
|---|---|---|---|---|---|
| AP53_382 | -0.066031989 | 5.415796632 | 0.038307671 | 0.848267416 | 0.922401711 |
| AP53_pseudo388 | 0.029940764 | 6.228121309 | 0.037759097 | 0.849342887 | 0.923054056 |
| AP53_655 | -0.023524879 | 9.319195173 | 0.035667253 | 0.853520395 | 0.927075034 |
| AP53_216 | -0.020062647 | 8.772890398 | 0.035077383 | 0.854721269 | 0.92786017 |
| AP53_1570 | 0.028212705 | 7.812768783 | 0.033662826 | 0.857644473 | 0.929632596 |
| AP53_1274 | -0.041299488 | 6.536350653 | 0.033454456 | 0.858080415 | 0.929632596 |
| AP53_422 | -0.025061168 | 10.12176022 | 0.033404303 | 0.858185554 | 0.929632596 |
| AP53_1110 | -0.022480065 | 7.404757561 | 0.0331458 | 0.858728766 | 0.929632596 |
| AP53_276 | -0.033986518 | 8.425597474 | 0.032987126 | 0.859063288 | 0.929632596 |
| AP53_634 | -0.054668181 | 4.245122489 | 0.032824276 | 0.859407481 | 0.929632596 |
| AP53_386 | -0.065921722 | 3.007733876 | 0.032414143 | 0.860278261 | 0.929632596 |
| AP53_rna77 | 0.288199999 | -1.139188048 | 0.032097995 | 0.860953393 | 0.929632596 |
| AP53_451 | -0.131249215 | 0.454461434 | 0.032087506 | 0.860975852 | 0.929632596 |
| AP53_1808 | -0.035883786 | 7.402905307 | 0.032009292 | 0.861143435 | 0.929632596 |
| AP53_870 | -0.020681039 | 8.825899477 | 0.031337547 | 0.862591552 | 0.930678267 |
| AP53_726 | -0.024610931 | 7.212955973 | 0.030559252 | 0.864289582 | 0.931815087 |
| AP53_995 | 0.014886761 | 7.959449408 | 0.030415637 | 0.864605343 | 0.931815087 |
| AP53_385 | -0.034138186 | 4.362309074 | 0.029776198 | 0.866020692 | 0.932638841 |
| AP53_133 | 0.037791481 | 10.50319653 | 0.029487393 | 0.866665081 | 0.932638841 |
| AP53_323 | -0.030510289 | 7.876094137 | 0.029271271 | 0.867149431 | 0.932638841 |
| AP53_rna5 | 0.088510397 | 4.152167762 | 0.029148737 | 0.867424862 | 0.932638841 |
| AP53_1273 | -0.029646329 | 6.981510129 | 0.028994612 | 0.867772152 | 0.932638841 |
| AP53_1660 | 0.056912876 | 8.781260688 | 0.028086676 | 0.869837605 | 0.933875825 |
| AP53_1193 | -0.060436787 | 2.503623599 | 0.028065858 | 0.869885364 | 0.933875825 |
| AP53_479 | 0.019066313 | 8.07023155 | 0.02761508 | 0.870924019 | 0.933935204 |
| AP53_255 | -0.025324672 | 9.625319323 | 0.027487764 | 0.871218952 | 0.933935204 |
| AP53_490 | 0.04142273 | 8.144957934 | 0.02741658 | 0.87138416 | 0.933935204 |
| AP53_790 | 0.023111344 | 9.013399893 | 0.026523868 | 0.873475054 | 0.935659537 |
| AP53_1780 | -0.036118346 | 8.841926028 | 0.025984075 | 0.874756951 | 0.936016708 |
| AP53_1194 | -0.085306863 | 1.506943889 | 0.025748307 | 0.875321146 | 0.936016708 |
| AP53_1558 | 0.020053646 | 7.796647122 | 0.025661553 | 0.875529415 | 0.936016708 |
| AP53_505 | 0.024944164 | 9.691599483 | 0.025575058 | 0.875737425 | 0.936016708 |
| AP53_rna6 | -0.060766013 | 4.56855931 | 0.024890117 | 0.877397548 | 0.936097306 |
| AP53_77 | -0.03267513 | 6.050714383 | 0.024602426 | 0.878101827 | 0.936097306 |
| AP53_338 | 0.045945615 | 12.83272165 | 0.024197969 | 0.879099146 | 0.936097306 |

| | | | | | |
|---|---|---|---|---|---|
| AP53_313 | 0.031965762 | 9.55902715 | 0.024054528 | 0.879454897 | 0.936097306 |
| AP53_1124 | -0.031992281 | 4.615390786 | 0.023996797 | 0.879598384 | 0.936097306 |
| AP53_215 | -0.020498104 | 9.515396733 | 0.023842399 | 0.879983005 | 0.936097306 |
| AP53_560 | -0.043472996 | 9.652494913 | 0.023764773 | 0.880176861 | 0.936097306 |
| AP53_1412 | 0.018307414 | 8.859933775 | 0.023593179 | 0.88060654 | 0.936097306 |
| AP53_1325 | -0.055073383 | 2.984739862 | 0.023589095 | 0.880616787 | 0.936097306 |
| AP53_1528 | 0.018984877 | 5.725136531 | 0.023507241 | 0.880822338 | 0.936097306 |
| AP53_1658 | 0.016175393 | 8.238727787 | 0.023213584 | 0.881562797 | 0.936097306 |
| AP53_1174 | -0.073897683 | 3.571063544 | 0.023198823 | 0.881600142 | 0.936097306 |
| AP53_1811 | -0.016581139 | 7.932330972 | 0.022488011 | 0.883413187 | 0.937415507 |
| AP53_340 | -0.018909462 | 6.586406523 | 0.022334915 | 0.88380751 | 0.937415507 |
| AP53_1738 | 0.029065595 | 10.31251749 | 0.021244047 | 0.886658215 | 0.939925503 |
| AP53_209 | -0.01677677 | 7.368054621 | 0.020883466 | 0.887616907 | 0.940428175 |
| AP53_475 | 0.016162741 | 9.041570762 | 0.020512717 | 0.888611503 | 0.940841782 |
| AP53_1222 | -0.022056161 | 7.686486073 | 0.020377426 | 0.888976727 | 0.940841782 |
| AP53_780 | 0.020875517 | 9.426762818 | 0.020128649 | 0.889651565 | 0.941042882 |
| AP53_pseudo73 | -0.022814348 | 6.753476038 | 0.019898018 | 0.890280994 | 0.941195757 |
| AP53_1245 | -0.014202012 | 8.936590676 | 0.019525927 | 0.891304401 | 0.941764748 |
| AP53_1271 | -0.025572407 | 7.72486972 | 0.019020209 | 0.892711437 | 0.94194856 |
| AP53_923 | 0.021659315 | 8.605489428 | 0.018950273 | 0.89290751 | 0.94194856 |
| AP53_317 | -0.041014613 | 9.054935529 | 0.018940751 | 0.892934235 | 0.94194856 |
| AP53_1505 | 0.019611966 | 10.10113094 | 0.01843584 | 0.894361288 | 0.942941478 |
| AP53_1800 | -0.038595463 | 4.960088655 | 0.018023993 | 0.895540142 | 0.943585263 |
| AP53_1796 | -0.039753799 | 4.779736854 | 0.01788393 | 0.895944173 | 0.943585263 |
| AP53_1109 | 0.014493196 | 6.796823817 | 0.017497297 | 0.897067924 | 0.944173939 |
| AP53_376 | -0.02626697 | 3.814968251 | 0.017137694 | 0.898124523 | 0.944173939 |
| AP53_1181 | 0.032671044 | 3.955616634 | 0.017048667 | 0.898387844 | 0.944173939 |
| AP53_507 | -0.023079128 | 4.440330087 | 0.016853976 | 0.898966148 | 0.944173939 |
| AP53_683 | -0.020776266 | 7.300666495 | 0.016764028 | 0.899234476 | 0.944173939 |
| AP53_140 | 0.022494154 | 9.852723686 | 0.016701396 | 0.899421748 | 0.944173939 |
| AP53_401 | -0.026752579 | 4.320376361 | 0.016352219 | 0.900472421 | 0.944264951 |
| AP53_1180 | 0.028723071 | 6.123507441 | 0.015900627 | 0.901848337 | 0.944264951 |
| AP53_294 | 0.015983429 | 7.702865212 | 0.015744111 | 0.902329843 | 0.944264951 |
| AP53_1060 | -0.031233062 | 5.333577603 | 0.015576302 | 0.902848802 | 0.944264951 |
| AP53_539 | 0.028478816 | 8.780216336 | 0.015349688 | 0.903554148 | 0.944264951 |

| | | | | | |
|---|---|---|---|---|---|
| AP53_rna61 | 0.090814725 | 1.540564622 | 0.015141558 | 0.904206645 | 0.944264951 |
| AP53_1246 | 0.012437151 | 8.444518672 | 0.015058257 | 0.904469073 | 0.944264951 |
| AP53_1049 | -0.037540737 | 4.90069802 | 0.01505332 | 0.904484648 | 0.944264951 |
| AP53_1818 | -0.022762551 | 5.452001863 | 0.014958139 | 0.904785457 | 0.944264951 |
| AP53_367 | -0.019672762 | 10.43533034 | 0.014740829 | 0.905475908 | 0.944264951 |
| AP53_270 | 0.013232239 | 8.047303966 | 0.014714737 | 0.905559155 | 0.944264951 |
| AP53_37 | 0.034619114 | 8.621551496 | 0.014544934 | 0.906102764 | 0.944264951 |
| AP53_783 | 0.018069895 | 7.122457783 | 0.014419277 | 0.906507119 | 0.944264951 |
| AP53_526 | 0.01266944 | 8.990784056 | 0.01427659 | 0.906968453 | 0.944264951 |
| AP53_1720 | -0.020084578 | 6.068932166 | 0.014255596 | 0.907036528 | 0.944264951 |
| AP53_76 | -0.015291138 | 7.413343077 | 0.014090341 | 0.907574166 | 0.944264951 |
| AP53_696 | -0.023679198 | 9.102386373 | 0.014027739 | 0.90777867 | 0.944264951 |
| AP53_30 | -0.044115598 | 10.05230692 | 0.013393266 | 0.909878047 | 0.945512113 |
| AP53_1233 | -0.0251691 | 12.28677147 | 0.013330527 | 0.910088356 | 0.945512113 |
| AP53_134 | 0.011944095 | 8.862852606 | 0.01322625 | 0.910439021 | 0.945512113 |
| AP53_375 | -0.046574389 | 2.887972822 | 0.012876078 | 0.911626969 | 0.946239543 |
| AP53_537 | 0.023338497 | 8.498753398 | 0.0126279 | 0.912478849 | 0.946555633 |
| AP53_1587 | -0.020016823 | 5.648149901 | 0.012504143 | 0.912906824 | 0.946555633 |
| AP53_572 | -0.020723556 | 8.917857401 | 0.011790854 | 0.915416467 | 0.948651021 |
| AP53_1200 | 0.023356567 | 3.627071698 | 0.011555439 | 0.916261553 | 0.949020104 |
| AP53_1262 | 0.03823772 | 10.79217666 | 0.011013071 | 0.918242333 | 0.950564463 |
| AP53_137 | 0.032576151 | 6.003092304 | 0.010816412 | 0.918972704 | 0.950813443 |
| AP53_1855 | 0.018672689 | 11.18573207 | 0.010285774 | 0.920977657 | 0.952380199 |
| AP53_1162 | -0.026624436 | 5.587893052 | 0.009407142 | 0.9244161 | 0.955332539 |
| AP53_106 | -0.011753481 | 7.294570993 | 0.009307273 | 0.924817023 | 0.955332539 |
| AP53_1400 | 0.009870796 | 7.909196084 | 0.009105373 | 0.925634237 | 0.955668114 |
| AP53_rna59 | 0.046256234 | 3.594400094 | 0.00871911 | 0.927223596 | 0.956800107 |
| AP53_1761 | 0.031447949 | 4.562062021 | 0.008235213 | 0.929265716 | 0.958397851 |
| AP53_1714 | -0.027460748 | 3.071270221 | 0.007907502 | 0.93068329 | 0.959182337 |
| AP53_442 | -0.016564342 | 3.886485751 | 0.007782888 | 0.931230083 | 0.959182337 |
| AP53_25 | 0.02316424 | 9.107144995 | 0.007719739 | 0.931508864 | 0.959182337 |
| AP53_1133 | 0.015478322 | 9.591269898 | 0.007530547 | 0.932351027 | 0.959191765 |
| AP53_pseudo399 | -0.01823985 | 4.327962628 | 0.007439617 | 0.932759586 | 0.959191765 |
| AP53_1606 | 0.017924618 | 4.367122512 | 0.007310779 | 0.933342807 | 0.959191765 |
| AP53_1856 | 0.009845418 | 7.180055165 | 0.007277407 | 0.933494717 | 0.959191765 |

| | | | | | |
|---|---|---|---|---|---|
| AP53_1264 | 0.027647498 | 10.19077399 | 0.007015836 | 0.934697757 | 0.95991976 |
| AP53_261 | 0.008823703 | 8.013127346 | 0.006408689 | 0.937580435 | 0.962371033 |
| AP53_1795 | -0.021986557 | 5.452817499 | 0.006089431 | 0.939151542 | 0.963474177 |
| AP53_1038 | -0.014572802 | 5.76057865 | 0.005904807 | 0.940079063 | 0.96391625 |
| AP53_1230 | 0.017797505 | 7.713718669 | 0.005694197 | 0.941155138 | 0.964510096 |
| AP53_760 | -0.021657514 | 10.99692506 | 0.005417637 | 0.942599057 | 0.96548009 |
| AP53_1207 | -0.02543948 | 3.359093794 | 0.005231683 | 0.943590873 | 0.965986226 |
| AP53_809 | 0.008800029 | 7.618084778 | 0.004676024 | 0.946665193 | 0.968622635 |
| AP53_1728 | -0.008574646 | 7.033408301 | 0.004172968 | 0.949611149 | 0.971124995 |
| AP53_250 | -0.005520256 | 8.649442822 | 0.004071381 | 0.950227343 | 0.971243429 |
| AP53_1717 | 0.009053065 | 6.665222329 | 0.003497393 | 0.953864282 | 0.973601593 |
| AP53_1537 | 0.009791934 | 10.26151985 | 0.003423428 | 0.954354128 | 0.973601593 |
| AP53_1326 | -0.026218882 | 4.142163145 | 0.003421619 | 0.954366177 | 0.973601593 |
| AP53_1257 | -0.007005391 | 7.218499909 | 0.003395439 | 0.954540871 | 0.973601593 |
| AP53_1488 | 0.014597576 | 10.09266506 | 0.00320904 | 0.955804774 | 0.974378711 |
| AP53_1836 | -0.009358943 | 7.000800069 | 0.003062562 | 0.956824062 | 0.974905777 |
| AP53_686 | -0.007126744 | 8.354929584 | 0.00295826 | 0.957564856 | 0.975148681 |
| AP53_1032 | -0.013208805 | 5.685132842 | 0.0024992 | 0.960992859 | 0.978126449 |
| AP53_982 | 0.008408566 | 11.67228753 | 0.002290311 | 0.962657167 | 0.979057326 |
| AP53_461 | 0.007309259 | 7.991869669 | 0.002258615 | 0.962916247 | 0.979057326 |
| AP53_1654 | -0.006298071 | 7.992457834 | 0.001999899 | 0.965103095 | 0.980767072 |
| AP53_631 | -0.009410614 | 4.708619 | 0.00177257 | 0.967144921 | 0.981732794 |
| AP53_1305 | 0.00524021 | 7.406840544 | 0.001604563 | 0.968739755 | 0.981732794 |
| AP53_1450 | -0.00409324 | 7.186068963 | 0.001602823 | 0.968756699 | 0.981732794 |
| AP53_1602 | -0.009415356 | 5.230470574 | 0.001586526 | 0.968915847 | 0.981732794 |
| AP53_79 | 0.005733726 | 8.410774121 | 0.001561159 | 0.969165213 | 0.981732794 |
| AP53_562 | 0.01236101 | 9.093359734 | 0.001556811 | 0.969208149 | 0.981732794 |
| AP53_1798 | 0.011574127 | 4.356196758 | 0.001518028 | 0.969593903 | 0.981732794 |
| AP53_372 | 0.00505057 | 8.032045335 | 0.001403277 | 0.970765101 | 0.982406184 |
| AP53_36 | 0.003991089 | 9.058116692 | 0.001142813 | 0.973616166 | 0.984777998 |
| AP53_1416 | 0.006933597 | 9.717240124 | 0.001087123 | 0.974266785 | 0.984831209 |
| AP53_1571 | -0.006508348 | 6.358480657 | 0.001031651 | 0.974931666 | 0.984831209 |
| AP53_168 | -0.006162372 | 8.972750008 | 0.001010415 | 0.975190925 | 0.984831209 |
| AP53_130 | 0.00365695 | 7.733056354 | 0.000910962 | 0.976443067 | 0.985127426 |
| AP53_pseudo1337 | -0.010246094 | 2.222517796 | 0.000901218 | 0.976569347 | 0.985127426 |

| | | | | | |
|---|---|---|---|---|---|
| AP53_197 | -0.003895363 | 9.919655201 | 0.000867867 | 0.97700685 | 0.985127426 |
| AP53_691 | -0.004335206 | 8.599600017 | 0.000639904 | 0.980255441 | 0.987889829 |
| AP53_1814 | 0.005824741 | 9.414836082 | 0.000504603 | 0.982466229 | 0.988554639 |
| AP53_1113 | 0.00244281 | 7.915725624 | 0.000500865 | 0.982531279 | 0.988554639 |
| AP53_1799 | -0.0054229 | 4.830050309 | 0.000478476 | 0.982926101 | 0.988554639 |
| AP53_1789 | 0.002888335 | 5.668436909 | 0.000458776 | 0.983281219 | 0.988554639 |
| AP53_1502 | 0.002695858 | 5.633125602 | 0.000448927 | 0.983461621 | 0.988554639 |
| AP53_1801 | 0.004306027 | 3.605031207 | 0.00039916 | 0.984405106 | 0.988990844 |
| AP53_962 | 0.002525392 | 7.398175823 | 0.00033661 | 0.98567886 | 0.98935455 |
| AP53_705 | -0.002924051 | 8.668212891 | 0.000329743 | 0.985825692 | 0.98935455 |
| AP53_807 | 0.002796936 | 6.845634405 | 0.000308209 | 0.986296267 | 0.98935455 |
| AP53_1666 | -0.003578045 | 9.740172386 | 0.000267627 | 0.987230192 | 0.989779857 |
| AP53_671 | 0.000728584 | 8.008832027 | 5.63E-05 | 0.994140662 | 0.996193611 |
| AP53_316 | -0.001372219 | 9.497923771 | 3.31E-05 | 0.995506484 | 0.997047516 |
| AP53_182 | 0.000509101 | 8.834217995 | 2.08E-05 | 0.996437415 | 0.9974652 |
| AP53_1167 | -7.74E-05 | 8.0253434 | 4.78E-07 | 0.999460459 | 0.999975645 |
| AP53_rna69 | 0 | -2.070645075 | 0 | 1 | 1 |

| function |
| --- |
| Hyaluronan synthase (EC 2.4.1.212) |
| small RNA fasX |
| NAD-dependent malic enzyme (EC 1.1.1.38) |
| Malate Na(+) symporter |
| Streptolysin S biosynthesis protein B (SagA) |
| UTP--glucose-1-phosphate uridylyltransferase (EC 2.7.7.9) |
| Apo-citrate lyase phosphoribosyl-dephospho-CoA transferase (EC 2.7.7.61) |
| UDP-glucose dehydrogenase in hyaluronic acid synthesis (EC 1.1.1.22) |
| Streptodornase B; Mitogenic factor 1 |
| PTS system, trehalose-specific IIB component (EC 2.7.1.69) / PTS system, trehalose-specific IIC component (EC 2.7.1.69) / PTS system, trehalose- |
| Oxaloacetate decarboxylase alpha chain (EC 4.1.1.3) |
| Citrate lyase alpha chain (EC 4.1.3.6) |
| Export ABC transporter ATP-binding protein; Streptolysin S export protein (SagG) |
| FIG01114118: hypothetical protein |
| Formiminoglutamase (EC 3.5.3.8) |
| Oxaloacetate decarboxylase beta chain (EC 4.1.1.3) |
| Citrate lyase beta chain (EC 4.1.3.6) |
| Deoxyribonucleoside regulator DeoR (transcriptional repressor) |
| FIG01117167: hypothetical protein |
| Histidine ammonia-lyase (EC 4.3.1.3) |
| Branched-chain amino acid aminotransferase (EC 2.6.1.42) |
| putative cationic amino acid transporter protein |
| Streptolysin S biosynthesis protein (SagF) |
| Trehalose-6-phosphate hydrolase (EC 3.2.1.93) |
| Glycerol uptake facilitator protein |
| Streptolysin S export transmembrane permease (SagI) |
| Glycerophosphoryl diester phosphodiesterase (EC 3.1.4.46) |
| NADH peroxidase (EC 1.11.1.1) |
| Streptolysin S export transmembrane permease (SagH) |
| Dipeptidase (EC 3.4.-.-) |
| Mg2+/citrate complex transporter |
| Phage lysin |
| Streptolysin S biosynthesis protein C (SagC) |
| Phage protein |

Alcohol dehydrogenase, zinc-containing
FIG01115306: hypothetical protein
Streptococcal extracellular nuclease 3; Mitogenic factor 3
Streptococcal phospholipase A2; _Toximoron (Other)
Trans-acting positive regulator
Putative membrane protein precursor SPs0273
Streptolysin S biosynthesis protein D (SagD)
Transcriptional regulator, LuxR family
Glycerol dehydrogenase (EC 1.1.1.6)
Threonyl-tRNA synthetase (EC 6.1.1.3)
LSU ribosomal protein L7/L12 (P1/P2)
rRNA small subunit methyltransferase I
PTS system, cellobiose-specific IIC component (EC 2.7.1.69)
Mobile element protein
Beta-galactosidase (EC 3.2.1.23)
Seryl-tRNA synthetase (EC 6.1.1.11)
oxidoreductase of aldo/keto reductase family, subgroup 1
FIG01114213: hypothetical protein
COG0488: ATPase components of ABC transporters with duplicated ATPase domains
FIG01113992: hypothetical protein
Conserved protein
Streptolysin S self-immunity protein (SagE)
transporter
Formate--tetrahydrofolate ligase (EC 6.3.4.3)
Endonuclease/Exonuclease/phosphatase family protein
L-serine dehydratase, beta subunit (EC 4.3.1.17)
Alpha-glycerophosphate oxidase (EC 1.1.3.21)
Portal protein, phage associated
Pyruvate formate-lyase activating enzyme (EC 1.97.1.4)
Streptolysin S biosynthesis protein B (SagB)
Antiphagocytic M protein
FIG01117556: hypothetical protein
Lactose phosphotransferase system repressor
2,3-butanediol dehydrogenase, S-alcohol forming, (R)-acetoin-specific (EC 1.1.1.4) / Acetoin (diacetyl) reductase (EC 1.1.1.5)
Exodeoxyribonuclease VII large subunit (EC 3.1.11.6)

Transaldolase (EC 2.2.1.2)
Putative iron-sulfur cluster assembly scaffold protein for SUF system, SufE2
Pyruvate formate-lyase (EC 2.3.1.54)
Glycerol kinase (EC 2.7.1.30)
LSU ribosomal protein L6p (L9e)
Alcohol dehydrogenase (EC 1.1.1.1)
Putative uncharacterized protein SPs0857
Glycyl-tRNA synthetase alpha chain (EC 6.1.1.14)
Lactate 2-monooxygenase (EC 1.13.12.4)
SSU ribosomal protein S19p (S15e)
Co-activator of prophage gene expression IbrB
Transcription termination protein NusB
Isoleucyl-tRNA synthetase (EC 6.1.1.5)
67kDa Myosin-crossreactive streptococcal antigen
Multiple sugar ABC transporter, membrane-spanning permease protein MsmG
Imidazolonepropionase (EC 3.5.2.7)
PTS system, cellobiose-specific IIA component (EC 2.7.1.69)
Phage base plate
Pore forming protein ebsA
Glutamate formiminotransferase (EC 2.1.2.5) @ Glutamate formyltransferase
Citrate lyase gamma chain, acyl carrier protein (EC 4.1.3.6)
Methylmalonyl-CoA decarboxylase, beta chain (EC 4.1.1.41); Oxaloacetate decarboxylase beta chain (EC 4.1.1.3)
Succinate-semialdehyde dehydrogenase [NAD] (EC 1.2.1.24); Succinate-semialdehyde dehydrogenase [NADP+] (EC 1.2.1.16)
PTS system, maltose and glucose-specific IIC component (EC 2.7.1.69) / PTS system, maltose and glucose-specific IIB component (EC 2.7.1.69) / [illegible]
Excinuclease ABC subunit C
PTS system, N-acetylgalactosamine-specific IID component (EC 2.7.1.69)
Multiple sugar ABC transporter, substrate-binding protein
Similar to ribosomal large subunit pseudouridine synthase D, Bacillus subtilis YjbO type
FIG01114562: hypothetical protein
hypothetical protein
Beta-glucosidase (EC 3.2.1.21)
Aquaporin (Major Intrinsic Protein Family)
FIG01114697: hypothetical protein
Formiminotetrahydrofolate cyclodeaminase (EC 4.3.1.4)
4-hydroxy-2-oxoglutarate aldolase (EC 4.1.3.16) @ 2-dehydro-3-deoxyphosphogluconate aldolase (EC 4.1.2.14)

Ribose-phosphate pyrophosphokinase (EC 2.7.6.1)
General stress protein, Gls24 family
Acetyl-coenzyme A carboxyl transferase beta chain (EC 6.4.1.2)
Toxin to DNA-damage-inducible protein J
Heat shock protein 60 family chaperone GroEL
L-lactate dehydrogenase (EC 1.1.1.27)
Biotin carboxyl carrier protein of acetyl-CoA carboxylase
SSU ribosomal protein S15p (S13e)
Alkyl hydroperoxide reductase protein F (EC 1.6.4.-)
dCMP deaminase (EC 3.5.4.12); Late competence protein ComEB
Transketolase (EC 2.2.1.1)
Transcriptional regulator, TetR family
ATP-dependent RNA helicase YfmL
LSU ribosomal protein L36p
ABC transporter permease protein
tRNA-dependent lipid II--L-alanine ligase @ tRNA-dependent lipid II--L-serine ligase
FIG006988: Lipase/Acylhydrolase with GDSL-like motif
RNA polymerase sigma factor RpoD
V-type ATP synthase subunit K (EC 3.6.3.14)
(3R)-hydroxymyristoyl-[acyl carrier protein] dehydratase (EC 4.2.1.-)
putative phosphomannomutase
Transport protein SgaT, putative
L-serine dehydratase, alpha subunit (EC 4.3.1.17)
LSU ribosomal protein L22p (L17e)
Mobile element protein
Dihydroorotate dehydrogenase, catalytic subunit (EC 1.3.3.1)
Promiscuous sugar phosphatase YidA, haloacid dehalogenase-like phosphatase family
Glycerol-3-phosphate transporter
dTDP-4-dehydrorhamnose reductase (EC 1.1.1.133)
FIG01116967: hypothetical protein
Ribosomal RNA small subunit methyltransferase C (EC 2.1.1.52) ## SSU rRNA m2G1207
Urocanate hydratase (EC 4.2.1.49)
Mobile element protein
Prolipoprotein diacylglyceryl transferase (EC 2.4.99.-)
Glucose-6-phosphate isomerase (EC 5.3.1.9)

LSU ribosomal protein L13p (L13Ae)
Substrate-specific component QueT (COG4708) of predicted queuosine-regulated ECF transporter
Protein:protein lipoyl transferase
Biotin carboxyl carrier protein of oxaloacetate decarboxylase; Biotin carboxyl carrier protein
Transcriptional regulator pfoR
Ribonucleotide reductase of class Ib (aerobic), beta subunit (EC 1.17.4.1)
Hyaluronate lyase (phage associated)
Histidine transport protein (permease)
Biotin carboxyl carrier protein of oxaloacetate decarboxylase; Biotin carboxyl carrier protein
FIG01115418: hypothetical protein
Methionyl-tRNA formyltransferase (EC 2.1.2.9)
Arginine deiminase (EC 3.5.3.6)
FIG01116792: hypothetical protein
Transcriptional antiterminator of lichenan operon, BglG family
Putative membrane protein YeiH
Peptidylproline cis-trans-isomerase (EC 5.2.1.8); Proteinase maturation protein (RopA)
tRNA-Arg-CCG
Probable NADH-dependent flavin oxidoreductase in cluster with COG2110
Mobile element protein
FIG01115631: hypothetical protein
Co-activator of prophage gene expression IbrA
FIG01114846: hypothetical protein
tRNA-Ala-TGC
Transmembrane component of general energizing module of ECF transporters
hypothetical protein, phage associated
Hyaluronate lyase (phage associated)
hypothetical protein
tRNA pseudouridine synthase A (EC 4.2.1.70)
LSU ribosomal protein L24p (L26e)
Preprotein translocase secY subunit (TC 3.A.5.1.1)
Phosphate acetyltransferase (EC 2.3.1.8)
LSU ribosomal protein L30p (L7e)
Putative deoxyribose-specific ABC transporter, ATP-binding protein
Pantothenate kinase (EC 2.7.1.33)
tRNA-Ala-TGC

Endoribonuclease L-PSP
Histidine kinase of the competence regulon ComD
FIG01115320: hypothetical protein
Short chain dehydrogenase
hypothetical protein
Transaldolase (EC 2.2.1.2)
Transcriptional regulator SpxA1
DNA mismatch repair protein MutS
Beta-glucosidase (EC 3.2.1.21)
Probable L-ascorbate-6-phosphate lactonase UlaG (EC 3.1.1.-) (L-ascorbate utilization protein G)
V-type ATP synthase subunit E (EC 3.6.3.14)
putative N-acetyl-muramidase
hypothetical protein
FIG000605: protein co-occurring with transport systems (COG1739)
Lipoteichoic acid synthase LtaS Type Iic/interrupted by large phage
Glutathione peroxidase family protein
Nicotinate-nucleotide adenylyltransferase (EC 2.7.7.18)
Hemolysins and related proteins containing CBS domains
Single-stranded DNA-binding protein
hypothetical protein
ABC transporter membrane-spanning permease - macrolide efflux
LSU ribosomal protein L35p
Hypothetical mga-associated protein
Cell surface protein
hypothetical phage protein
Bacteriocin BlpU
Class B acid phosphatase precursor (EC 3.1.3.2)
Cysteine synthase (EC 2.5.1.47)
Oxaloacetate decarboxylase alpha chain (EC 4.1.1.3)
PTS system, N-acetylgalactosamine-specific IIC component (EC 2.7.1.69)
Arsenate reductase (EC 1.20.4.1)
Branched-chain amino acid transport system carrier protein
hypothetical protein
PTS system, galactose-specific IIC component (EC 2.7.1.69)
possible permease

3-oxoacyl-[acyl-carrier protein] reductase (EC 1.1.1.100)
Type I restriction-modification system, specificity subunit S (EC 3.1.21.3)
Regulation of D-alanyl-lipoteichoic acid biosynthesis, sensor histidine kinase
Pathogenicity island SaPIn2
Glucokinase (EC 2.7.1.2)
transcriptional regulator, MutR family
N-acetylmannosamine kinase (EC 2.7.1.60)
V-type ATP synthase subunit C (EC 3.6.3.14)
Phage protein
FIG01114324: hypothetical protein
DegV family protein in cluster with TrmH family tRNA/rRNA methyltransferase YacO
Oligoendopeptidase F
Two component transcriptional regulator VraR
V-type ATP synthase subunit B (EC 3.6.3.14)
tRNA-Ala-TGC
LSU ribosomal protein L5p (L11e)
Galactose-6-phosphate isomerase, LacA subunit (EC 5.3.1.26)
HPr kinase/phosphorylase (EC 2.7.1.-) (EC 2.7.4.-)
LSU m3Psi1915 methyltransferase RlmH
Alkaline shock protein
L-ribulose-5-phosphate 4-epimerase (EC 5.1.3.4)
Multi antimicrobial extrusion (MATE) family transporter
hypothetical phage protein
Translation elongation factor LepA
Heat-inducible transcription repressor HrcA
Alcohol dehydrogenase (EC 1.1.1.1); Acetaldehyde dehydrogenase (EC 1.2.1.10)
Streptopain inhibitor
Uncharacterized conserved protein, contains double-stranded beta-helix domain
Transcriptional regulator, MutR family
Phenylalanyl-tRNA synthetase alpha chain (EC 6.1.1.20)
LSU ribosomal protein L14p (L23e)
SSU ribosomal protein S14p (S29e) @ SSU ribosomal protein S14p (S29e), zinc-dependent
Phage major capsid protein
Prophage Clp protease-like protein
Peptide methionine sulfoxide reductase MsrB (EC 1.8.4.12)

Outer surface protein of unknown function, cellobiose operon
Nucleoside diphosphate kinase (EC 2.7.4.6)
Secreted Endo-beta-N-acetylglucosaminidase (EndoS)
LSU ribosomal protein L1p (L10Ae)
Pseudouridylate synthases, 23S RNA-specific
CTP synthase (EC 6.3.4.2)
Heat shock protein HtpX (EC 3.4.24.-)
FIG01115289: hypothetical protein
Glutamate transport ATP-binding protein
Phage protein
Non-specific DNA-binding protein Dps / Iron-binding ferritin-like antioxidant protein / Ferroxidase (EC 1.16.3.1)
General stress protein, Gls24 family
FIG01114660: hypothetical protein
S-layer homology domain / putative murein endopeptidase
tRNA-Ala-TGC
Phage major tail protein
conserved hypothetical protein - phage associated
hypothetical phage protein
Mobile element protein
Alanyl-tRNA synthetase (EC 6.1.1.7)
hypothetical protein
3-oxoacyl-[acyl-carrier-protein] synthase, KASII (EC 2.3.1.41)
GTP pyrophosphokinase (EC 2.7.6.5)
Two-component sensor kinase YesM (EC 2.7.3.-)
FIG01114374: hypothetical protein
Adenylosuccinate lyase (EC 4.3.2.2)
Hydrolases of the alpha/beta superfamily
50S ribosomal subunit maturation GTPase RbgA (B. subtilis YlqF)
ABC transporter, predicted N-acetylneuraminate-binding protein
Alpha-D-GlcNAc alpha-1,2-L-rhamnosyltransferase (EC 2.4.1.-)
Glutamine ABC transporter, glutamine-binding protein/permease protein
Cysteinyl-tRNA synthetase (EC 6.1.1.16)
tRNA nucleotidyltransferase (EC 2.7.7.21) (EC 2.7.7.25)
Low temperature requirement C protein
LSU ribosomal protein L10p (P0)

Dipeptide-binding ABC transporter, periplasmic substrate-binding component (TC 3.A.1.5.2)
SSU ribosomal protein S14p (S29e) @ SSU ribosomal protein S14p (S29e), zinc-independent
DNA topoisomerase I (EC 5.99.1.2)
Amino acid transport ATP-binding protein
Purine nucleoside phosphorylase (EC 2.4.2.1); N-Ribosylnicotinamide phosphorylase (EC 2.4.2.1) ## possible
Phage tail length tape-measure protein
hypothetical cytosolic protein
23S rRNA (Uracil-5-) -methyltransferase RumA (EC 2.1.1.-) ## LSU rRNA m(5)U1939
Signal recognition particle, subunit Ffh SRP54 (TC 3.A.5.1.1)
Ribose-phosphate pyrophosphokinase (EC 2.7.6.1)
Histone acetyltransferase Gcn5
tRNA pseudouridine synthase B (EC 4.2.1.70)
SSU ribosomal protein S8p (S15Ae)
Plasmid stabilization system antitoxin protein
Transcriptional regulator in cluster with unspecified monosaccharide ABC transport system
FIG01117988: hypothetical protein
hypothetical phage protein
Redox-sensitive transcriptional regulator (AT-rich DNA-binding protein)
rRNA small subunit 7-methylguanosine (m7G) methyltransferase GidB
V-type ATP synthase subunit I (EC 3.6.3.14)
Hypothetical protein SPy1643
Iron-sulfur cluster assembly protein SufB
Nucleoside ABC transporter, permease protein 2
FIG01118451: hypothetical protein
FIG01114293: hypothetical protein
FIG01115348: hypothetical protein
ABC transporter ATP-binding protein
Phage protein
Salt-stress induced protein
Putative regulator of the mannose operon, ManO
Glycyl-tRNA synthetase beta chain (EC 6.1.1.14)
Galactose-6-phosphate isomerase, LacB subunit (EC 5.3.1.26)
tRNA-Cys-GCA
Predicted transcriptional regulator of pyridoxine metabolism
16S rRNA (guanine(966)-N(2))-methyltransferase (EC 2.1.1.171) ## SSU rRNA m(2)G966

Putative deoxyribose-specific ABC transporter, permease protein
Phosphinothricin N-acetyltransferase (EC 2.3.1.-)
Transcriptional regulator ArcR essential for anaerobic expression of the ADI pathway, Crp/Fnr family
Hypothetical protein SP1558
Transmembrane component BL0694 of energizing module of predicted ECF transporter
NADH peroxidase (EC 1.11.1.1)
Ribonucleotide reductase of class Ib (aerobic), alpha subunit (EC 1.17.4.1)
6-phospho-beta-glucosidase (EC 3.2.1.86)
Biotin carboxylase of acetyl-CoA carboxylase (EC 6.3.4.14)
Type I restriction-modification system specificity subunit
Triosephosphate isomerase (EC 5.3.1.1)
ABC transporter, ATP-binding/permease protein
6-phospho-beta-galactosidase (EC 3.2.1.85)
Serine hydroxymethyltransferase (EC 2.1.2.1)
Dicarboxylate/amino acid:cation (Na+ or H+) symporter
tRNA(Ile)-lysidine synthetase
Transcriptional regulator, DeoR family
Phage protein
Ornithine carbamoyltransferase (EC 2.1.3.3)
Response regulator of the competence regulon ComE
tRNA uridine 5-carboxymethylaminomethyl modification enzyme GidA
Ortholog of S. aureus MRSA252 (BX571856) SAR1694
INTEGRAL MEMBRANE PROTEIN (Rhomboid family)
Phage tail fibers
Phage endonuclease
Exodeoxyribonuclease III (EC 3.1.11.2)
Lipoteichoic acid synthase LtaS Type IIc
Shikimate/quinate 5-dehydrogenase I beta (EC 1.1.1.282)
SSU ribosomal protein S5p (S2e)
Phosphopantothenoylcysteine synthetase (EC 6.3.2.5)
probably aromatic ring hydroxylating enzyme, evidenced by COGnitor; PaaD-like protein (DUF59) involved in Fe-S cluster assembly
V-type ATP synthase subunit F (EC 3.6.3.14)
Catabolite control protein A
Peptidoglycan hydrolase, Autolysin2 (EC 3.5.1.28)
LSU ribosomal protein L15p (L27Ae)

Alkyl hydroperoxide reductase protein C (EC 1.6.4.-)
Ribosomal RNA large subunit methyltransferase N (EC 2.1.1.-)
Alpha-L-Rha alpha-1,3-L-rhamnosyltransferase (EC 2.4.1.-)
Inositol-1-monophosphatase (EC 3.1.3.25)
Octaprenyl diphosphate synthase (EC 2.5.1.90) / Dimethylallyltransferase (EC 2.5.1.1) / (2E,6E)-farnesyl diphosphate synthase (EC 2.5.1.10) / Ger
Phage capsid protein
Protein-N(5)-glutamine methyltransferase PrmC, methylates polypeptide chain release factors RF1 and RF2
Phosphoglycerate mutase family
Oligopeptide ABC transporter, periplasmic oligopeptide-binding protein OppA (TC 3.A.1.5.1)
Sucrose-6-phosphate hydrolase (EC 3.2.1.26)
FIG01114764: hypothetical protein
Acetylornithine deacetylase/Succinyl-diaminopimelate desuccinylase and related deacylases
Protein of unknown function DUF208
Sortase A, LPXTG specific
Phage hyaluronidase
Glutathione reductase (EC 1.8.1.7)
5'-methylthioadenosine nucleosidase (EC 3.2.2.16) / S-adenosylhomocysteine nucleosidase (EC 3.2.2.9)
Dihydroxyacetone kinase family protein
Dipeptide transport ATP-binding protein DppF (TC 3.A.1.5.2)
Inactive homolog of metal-dependent proteases, putative molecular chaperone
membrane protein
FIG01114657: hypothetical protein
Phenylalanyl-tRNA synthetase beta chain (EC 6.1.1.20)
Integral membrane protein
LSU ribosomal protein L20p
FIG01114947: hypothetical protein
FIG01115281: hypothetical protein
Acetylornithine deacetylase/Succinyl-diaminopimelate desuccinylase and related deacylases
tRNA-Arg-ACG
tRNA-Pro-TGG
Cystathionine beta-lyase (EC 4.4.1.8)
Asparagine-rich ORFan next to Sag operon
GTP-binding protein EngA
DNA gyrase subunit B (EC 5.99.1.3)
hypothetical protein

SSU ribosomal protein S12p (S23e)
PTS system, mannose-specific IID component (EC 2.7.1.69)
UDP-N-acetylmuramoylpentapeptide-lysine N(6)-alanyltransferase (EC 2.3.2.10) (EC 2.3.2.-)
COG1939: Ribonuclease III family protein
FIG014356: hypothetical protein
tRNA-Leu-AAG
FIG01116532: hypothetical protein
Phage endopeptidase
SSU ribosomal protein S18p @ SSU ribosomal protein S18p, zinc-independent
Transcriptional regulator
Galactose-6-phosphate isomerase, LacA subunit (EC 5.3.1.26)
Pyridoxal kinase (EC 2.7.1.35)
Phosphate transport ATP-binding protein PstB (TC 3.A.1.7.1)
Zn-dependent hydrolase YycJ/WalJ, required for cell wall metabolism and coordination of cell division with DNA replication
Cystine ABC transporter, periplasmic cystine-binding protein FliY
FIG015389: hypothetical membrane associated protein
FIG01117156: hypothetical protein
FIG01116327: hypothetical protein
Phage protein
FIG01114305: hypothetical protein
Glutaredoxin-like protein NrdH, required for reduction of Ribonucleotide reductase class Ib
hypothetical protein
LSU ribosomal protein L18p (L5e)
Ribonuclease Z (EC 3.1.26.11)
FIG01115635: hypothetical protein
LSU ribosomal protein L16p (L10e)
DegV family protein
Manganese ABC transporter, ATP-binding protein SitB
Translation initiation factor 3
hypothetical protein
FIG01115464: hypothetical protein
Aspartyl-tRNA(Asn) amidotransferase subunit C (EC 6.3.5.6) @ Glutamyl-tRNA(Gln) amidotransferase subunit C (EC 6.3.5.7)
Putative ABC transporter ATP-binding protein, spy1790 homolog
tRNA-Ala-TGC
TsaD/Kae1/Qri7 protein, required for threonylcarbamoyladenosine t(6)A37 formation in tRNA

N-acetylglucosamine-6-phosphate deacetylase (EC 3.5.1.25)
LSU ribosomal protein L9p
Glucosamine-6-phosphate deaminase (EC 3.5.99.6)
hypothetical protein
Manganese-dependent inorganic pyrophosphatase (EC 3.6.1.1)
Deoxyguanosinetriphosphate triphosphohydrolase (EC 3.1.5.1)
Dipeptide transport system permease protein DppB (TC 3.A.1.5.2)
Cell division protein FtsH (EC 3.4.24.-)
tRNA-Ala-TGC
PTS system, cellobiose-specific IIB component (EC 2.7.1.69)
Secreted antigen GbpB/SagA/PcsB, putative peptidoglycan hydrolase
FIG042801: CBS domain containing protein
FIG001553: Hydrolase, HAD subfamily IIIA
lipoprotein, putative
putative esterase
FIG01114963: hypothetical protein
FIG01114801: hypothetical protein
Phage hyaluronidase
Cardiolipin synthetase (EC 2.7.8.-)
GAF domain-containing proteins
Transcriptional regulator RegR, rpressor of hyaluronate and KDG utilization
tRNA-Asn-GTT
membrane protein
PTS system, beta-glucoside-specific IIB component (EC 2.7.1.69) / PTS system, beta-glucoside-specific IIC component (EC 2.7.1.69) / PTS system,
Phosphate transport system permease protein PstA (TC 3.A.1.7.1)
Purine nucleoside phosphorylase (EC 2.4.2.1)
FIG01116522: hypothetical protein
Histidine protein kinase
FIG01116444: hypothetical protein
FIG01115301: hypothetical protein
DNA-damage-inducible protein J
hydrolase, haloacid dehalogenase-like family
FIG01115071: hypothetical protein
Single-stranded-DNA-specific exonuclease RecJ (EC 3.1.-.-)
Multiple sugar ABC transporter, membrane-spanning permease protein MsmF

tRNA-Phe-GAA
Chaperone protein DnaJ
Beta-glucoside bgl operon antiterminator, BglG family
Phage major tail protein
FIG01114264: hypothetical protein
Putative tRNA-m1A22 methylase
ATP-dependent nuclease, subunit B
Ribose 5-phosphate isomerase A (EC 5.3.1.6)
Phosphoglycerate mutase (EC 5.4.2.1)
Phage protein
FIG01117062: hypothetical protein
LSU ribosomal protein L17p
FIG01114098: hypothetical protein
Ribonucleotide reductase of class Ib (aerobic), beta subunit (EC 1.17.4.1)
FIG01114725: hypothetical protein
tRNA-Met-CAT
Acid phosphatase (EC 3.1.3.2)
Phage protein
PTS system, mannose-specific IIB component (EC 2.7.1.69) / PTS system, mannose-specific IIA component (EC 2.7.1.69)
DNA packaging protein, phage associated
SSU ribosomal protein S6p
Polyribonucleotide nucleotidyltransferase (EC 2.7.7.8)
Acetate kinase (EC 2.7.2.1)
FIG027054: hypothetical protein
hypothetical protein
Luciferase-like monooxygenase (EC 1.14.-.-)
Amino acid ABC transporter, amino acid-binding protein
Anaerobic ribonucleoside-triphosphate reductase (EC 1.17.4.2)
PTS system, mannose-specific IID component (EC 2.7.1.69)
Putative uncharacterized protein gbs2121
D-alanyl-D-alanine carboxypeptidase (EC 3.4.16.4)
Transcriptional regulator, CrP/Fnr family
ATP-dependent Zn protease
Strepotococcal cysteine protease (Streptopain) (EC 3.4.22.10) / Streptococcal pyrogenic exotoxin B (SpeB)
Transcriptional regulator, GntR family

tRNA-Arg-ACG
ABC transporter, substrate-binding protein
Cell division trigger factor (EC 5.2.1.8)
tRNA-Lys-CTT
YheO-like PAS domain
NAD kinase (EC 2.7.1.23)
Single-stranded DNA-binding protein
ABC transporter ATP-binding protein
ABC transporter permease protein
Pyruvate,phosphate dikinase (EC 2.7.9.1)
Predicted regulator for deoxynucleoside utilization, GntR family
FIG01114174: hypothetical protein
tRNA-Pseudo-TGA
1-acyl-sn-glycerol-3-phosphate acyltransferase (EC 2.3.1.51)
6-aminohexanoate-cyclic-dimer hydrolase (EC 3.5.2.12)
Putative sugar uptake protein SpyM3_1856/SPs1852
Enolase (EC 4.2.1.11)
Two-component response regulator, malate (EC 2.7.3.-)
LSU ribosomal protein L33p @ LSU ribosomal protein L33p, zinc-independent
Phage shock protein C, putative; stress-responsive transcriptional regulator
UDP-N-acetylmuramoylalanyl-D-glutamate--L-lysine ligase (EC 6.3.2.7)
conserved hypothetical protein - phage associated
Polysaccharide deacetylase
tRNA-Leu-TAA
FIG048485: Phage protein
biofilm formation protein YmcA
FIG01116163: hypothetical protein
DNA-binding response regulator, AraC family
Inner membrane protein YihY, formerly thought to be RNase BN
Phage protein
Hyaluronate lyase (phage associated)
Response regulator of the competence regulon ComE
Phage minor capsid protein
SSU ribosomal protein S9p (S16e)
A/G-specific adenine glycosylase (EC 3.2.2.-)

ATPase YjeE, predicted to have essential role in cell wall biosynthesis
Arginine pathway regulatory protein ArgR, repressor of arg regulon
Peptide ABC transporter ATP binding/permease protein
FIG01114029: hypothetical protein
Pneumococcal vaccine antigen A homolog
3-keto-L-gulonate 6-phosphate decarboxylase
GTPase and tRNA-U34 5-formylation enzyme TrmE
Aminopeptidase YpdF (MP-, MA-, MS-, AP-, NP- specific)
1-phosphofructokinase (EC 2.7.1.56)
tRNA-Arg-ACG
Phospho-N-acetylmuramoyl-pentapeptide-transferase (EC 2.7.8.13)
Dipeptide transport ATP-binding protein DppD (TC 3.A.1.5.2)
Hypothetical protein YggS, proline synthase co-transcribed bacterial homolog PROSC
FIG01114041: hypothetical protein
Translation elongation factor Tu
Acyl carrier protein
putative cI-like repressor - phage associated
Tripeptide aminopeptidase (EC 3.4.11.4)
Oxygen-insensitive NAD(P)H nitroreductase (EC 1.-.-.-) / Dihydropteridine reductase (EC 1.5.1.34)
conserved hypothetical protein - phage associated
Phenylacetic acid degradation protein PaaD, thioesterase
ATPase
Phage protein
Late competence protein ComGD, access of DNA to ComEA, FIG038316
Type I restriction-modification system, restriction subunit R (EC 3.1.21.3)
Proline dipeptidase (EC 3.4.13.9)
conserved hypothetical protein - phage associated
hypothetical protein
Cold-shock DEAD-box protein A
membrane protein, putative
Amino acid ABC transporter, glutamine-binding protein/permease protein
23S rRNA (guanosine-2'-O-) -methyltransferase rlmB (EC 2.1.1.-) ## LSU rRNA Gm2251
Dolichyl-phosphate mannoosyltransferase, involved in cell wall biogenesis
tRNA-Met-CAT
Poly(glycerophosphate chain) D-alanine transfer protein DltD

Lipoate-protein ligase A
FIG01116522: hypothetical protein
periplasmic component of efflux system
Putative Holliday junction resolvase YggF
tRNA-Pro-TGG
FIG107367: tRNA-binding protein
C5a peptidase (EC 3.4.21.-)
DNA-directed RNA polymerase delta subunit (EC 2.7.7.6)
tRNA-Met-CAT
PROTEASE (EC 3.4.-.-)
Glutamate racemase (EC 5.1.1.3)
Cysteine ABC transporter, permease protein
Phage transcriptional regulator, Cro/CI family
PTS system, galactose-specific IIA component (EC 2.7.1.69)
Multimodular transpeptidase-transglycosylase (EC 2.4.1.129) (EC 3.4.-.-) / Penicillin-binding protein 1A/1B (PBP1)
PTS system, mannose-specific IIC component (EC 2.7.1.69)
Phage protein
tRNA dihydrouridine synthase B (EC 1.-.-.-)
tRNA-Leu-TAA
Formamidopyrimidine-DNA glycosylase (EC 3.2.2.23)
Response regulator CsrR
hypothetical protein
Serine endopeptidase ScpC (EC 3.4.21.-)
2-haloalkanoic acid dehalogenase (EC 3.8.1.2)
ABC transporter, ATP-binding/permease protein
PTS system, N-acetylgalactosamine-specific IIB component (EC 2.7.1.69)
Cell division initiation protein DivIVA
ABC transporter, permease protein EscB
Fibronectin-binding protein/major pilin backbone protein T-antigen
Recombination protein RecR
Ribonuclease HII (EC 3.1.26.4)
DNA binding protein, FIG046916
Serine protease, DegP/HtrA, do-like (EC 3.4.21.-)
hypothetical protein
FIG01114903: hypothetical protein

FIG004453: protein YceG like
conserved domain protein
Ribosome-associated heat shock protein implicated in the recycling of the 50S subunit (S4 paralog)
ABC-transporter (ATP-binding protein) -possibly involved in cell wall localization and side chain formation of rhamnose-glucose polysaccharide
DNA-binding protein HBsu
Phage protein
Phage lysin, glycosyl hydrolase, family 25
hypothetical phage protein
LSU ribosomal protein L2p (L8e)
FIG01115412: hypothetical protein
DNA polymerase III subunits gamma and tau (EC 2.7.7.7)
FIG000325: clustered with transcription termination protein NusA
tRNA-Met-CAT
FIG000506: Predicted P-loop-containing kinase
ComF operon protein A, DNA transporter ATPase
Cell division protein FtsQ
Phosphohydrolase (MutT/nudix family protein)
Beta-lactamase, class C
Mobile element protein
SWF/SNF family helicase
FIG01115860: hypothetical protein
Tyrosyl-tRNA synthetase (EC 6.1.1.1)
tRNA-Thr-TGT
2-keto-3-deoxy-D-arabino-heptulosonate-7-phosphate synthase I beta (EC 2.5.1.54)
hypothetical protein (phage associated)
Phosphopentomutase (EC 5.4.2.7)
hypothetical phage protein
ABC transporter ATP-binding protein uup
DNA gyrase subunit A (EC 5.99.1.3)
tRNA-dependent lipid II-Ala--L-alanine ligase @ tRNA-dependent lipid II-Ser--L-alanine ligase
bacterial seryl-tRNA synthetase related
Ribosomal subunit interface protein
SSU ribosomal protein S13p (S18e)
Competence-stimulating peptide ABC transporter permease protein ComB
Ribonuclease P protein component (EC 3.1.26.5)

Phosphatidylglycerophosphatase B (EC 3.1.3.27)
Rossmann fold nucleotide-binding protein Smf possibly involved in DNA uptake
S-adenosylmethionine:tRNA ribosyltransferase-isomerase (EC 5.-.-.-)
GTP-binding protein Obg
ABC transporter, ATP-binding protein EcsA
Transcription accessory protein (S1 RNA-binding domain)
Cytidylate kinase (EC 2.7.4.14)
FIG01114845: hypothetical protein
Phage protein
Chorismate synthase (EC 4.2.3.5)
Single-stranded DNA-binding protein
FIG01114677: hypothetical protein
Phage protein
PTS system, mannose/fructose family IIB component
Collagen-like surface protein
FIG01114027: hypothetical protein
Transcription antiterminator, BglG family (EC 2.7.1.69)
phage protein
FIG01113961: hypothetical protein
Glutamine synthetase type I (EC 6.3.1.2)
Ribonucleotide reduction protein NrdI
Uridine phosphorylase (EC 2.4.2.3)
Transmembrane histidine kinase CsrS / deletion of T at 1407 causing a early stop codon at 469th aa)
FIG01114622: hypothetical protein
Magnesium and cobalt transport protein CorA
Phage protein
Small Subunit Ribosomal RNA; ssuRNA; SSU rRNA
ABC transporter, periplasmic spermidine putrescine-binding protein PotD (TC 3.A.1.11.1)
ABC transporter permease protein
hypothetical protein
Hypothetical protein
tRNA-His-GTG
Cobalt-zinc-cadmium resistance protein
GTP-binding protein YqeH, required for biogenesis of 30S ribosome subunit
tRNA (guanine46-N7-)-methyltransferase (EC 2.1.1.33)

DNA primase (EC 2.7.7.-)
LSU ribosomal protein L27p
tRNA-Tyr-GTA
Ribonuclease HIII (EC 3.1.26.4)
Hemolysin III
Two-component system response regulator
FIG01114242: hypothetical protein
Phosphate:acyl-ACP acyltransferase PlsX
FIG01116522: hypothetical protein
FIG01114304: hypothetical protein
FIG01114333: hypothetical protein
Small Subunit Ribosomal RNA; ssuRNA; SSU rRNA
FIG01114322: hypothetical protein
Ribosyl nicotinamide transporter, PnuC-like
Two-component system histidine kinase
DNA repair protein RecN
ABC transporter permease protein
Integral membrane protein
Spermidine Putrescine ABC transporter permease component potC (TC_3.A.1.11.1)
Dihydrofolate synthase (EC 6.3.2.12) / Folylpolyglutamate synthase (EC 6.3.2.17)
Transcriptional regulator, Cro/CI family
Ferrichrome-binding periplasmic protein precursor (TC 3.A.1.14.3)
Transcriptional regulator, MerR family
tRNA-Tyr-GTA
3-ketoacyl-CoA thiolase (EC 2.3.1.16) @ Acetyl-CoA acetyltransferase (EC 2.3.1.9)
Hydrolase (HAD superfamily)
Maltose operon transcriptional repressor MalR, LacI family
Malonyl CoA-acyl carrier protein transacylase (EC 2.3.1.39)
Cysteine desulfurase (EC 2.8.1.7), SufS subfamily
Chromosome replication initiation protein dnaD
Nucleoside-binding protein
Multiple sugar metabolism regulator
C3 family ADP-ribosyltransferase (EC 2.4.2.-)
Hydroxyacylglutathione hydrolase (EC 3.1.2.6)
conserved hypothetical protein - phage associated

Cationic amino acid transporter - APC Superfamily
Mobile element protein
ABC-type multidrug transport system, permease component
tRNA-Thr-TGT
GTP pyrophosphokinase (EC 2.7.6.5), (p)ppGpp synthetase I
YpfJ protein, zinc metalloprotease superfamily
FIG01114299: hypothetical protein
Membrane-bound protease, CAAX family
Phage antirepressor protein
Hypothetical NagD-like phosphatase
FIG01115236: hypothetical protein
Cell division protein FtsI [Peptidoglycan synthetase] (EC 2.4.1.129)
SSU ribosomal protein S3p (S3e)
Integral membrane protein
Peptide deformylase (EC 3.5.1.88)
Phage Holliday junction resolvase
Neutral endopeptidase O (EC 3.4.24.-)
Phage lysin
RNA methyltransferase, TrmA family
FIG01114632: hypothetical protein
ADP-ribose pyrophosphatase (EC 3.6.1.13)
FIG01116748: hypothetical protein
hypothetical protein
Dipeptide transport system permease protein DppC (TC 3.A.1.5.2)
Lipoprotein signal peptidase (EC 3.4.23.36)
putative phage replication protein
FIG005986: HD family hydrolase
Phage lysin
Streptococcal lipoprotein rotamase A; Peptidyl-prolyl cis-trans isomerase (EC 5.2.1.8)
Small Subunit Ribosomal RNA; ssuRNA; SSU rRNA
Ribosome-binding factor A
tRNA-Leu-TAG
Glycosyltransferase LafA, responsible for the formation of Glc-DAG
FIG01115915: hypothetical protein
Phage protein

FIG01114239: hypothetical protein
Positive transcriptional regulator, MutR family
tRNA-Pseudo-TGA
Methionine aminopeptidase (EC 3.4.11.18)
putative collagen binding protein
hypothetical protein
FIG01115240: hypothetical protein
Small Subunit Ribosomal RNA; ssuRNA; SSU rRNA
Integral membrane protein
Ascorbate-specific PTS system, EIIB component (EC 2.7.1.69)
Competence protein CoiA
conserved hypothetical protein - phage associated
Phosphocarrier protein of PTS system
Ribonuclease J1 (endonuclease and 5' exonuclease)
Tyrosine recombinase XerC
V-type ATP synthase subunit D (EC 3.6.3.14)
Multiple sugar ABC transporter, ATP-binding protein
PTS system, IIA component, putative
Small Subunit Ribosomal RNA; ssuRNA; SSU rRNA
FIG01116817: hypothetical protein
S1 RNA binding domain
Hypothetical protein SPy0110
FIG028593: membrane protein
PTS system, mannose-specific IIC component (EC 2.7.1.69)
FIG01114509: hypothetical protein
Butyrate-acetoacetate CoA-transferase subunit B (EC 2.8.3.9)
GMP synthase [glutamine-hydrolyzing], amidotransferase subunit (EC 6.3.5.2) / GMP synthase [glutamine-hydrolyzing], ATP pyrophosphatase su
Competence-stimulating peptide ABC transporter ATP-binding protein ComA
Substrate-specific component NiaX of predicted niacin ECF transporter
COG2110, Macro domain, possibly ADP-ribose binding module
FIG040428: Phage-associated protein
Bactoprenol glucosyl transferase (EC 2.4.1.-)
Transcriptional regulator, TetR family
Phosphoglycerate kinase (EC 2.7.2.3)
dTDP-glucose 4,6-dehydratase (EC 4.2.1.46)

Small Subunit Ribosomal RNA; ssuRNA; SSU rRNA
FIG01114189: hypothetical protein
Streptococcal cell surface hemoprotein receptor Shr
PTS system, cellobiose-specific IIA component (EC 2.7.1.69)
FIG01118889: hypothetical protein
Galactosamine-6-phosphate isomerase (EC 5.3.1.-)
Hydrolase (HAD superfamily)
Hypothetical protein DUF194, DegV family
Carbamoyl-phosphate synthase small chain (EC 6.3.5.5)
Large Subunit Ribosomal RNA; lsuRNA; LSU rRNA
Transposase
Peptide chain release factor 2; programmed frameshift-containing
Large Subunit Ribosomal RNA; lsuRNA; LSU rRNA
Phage hyaluronidase; Phage endopeptidase
Large Subunit Ribosomal RNA; lsuRNA; LSU rRNA
Large Subunit Ribosomal RNA; lsuRNA; LSU rRNA
Long-chain-fatty-acid--CoA ligase (EC 6.2.1.3)
FIG01280259: hypothetical protein
tRNA S(4)U 4-thiouridine synthase (former ThiI)
Cell division transporter, ATP-binding protein FtsE (TC 3.A.5.1.1)
ABC transporter, predicted N-acetylneuraminate transport system permease protein 1
tRNA-Asn-GTT
D-alanyl transfer protein DltB
Large Subunit Ribosomal RNA; lsuRNA; LSU rRNA
Candidate zinc-binding lipoprotein ZinT
hypothetical protein
Pleiotropic regulator of exopolysaccharide synthesis, competence and biofilm formation Ftr, XRE family
Nucleoside-binding protein
Large Subunit Ribosomal RNA; lsuRNA; LSU rRNA
Phage antirepressor protein
Recombinational DNA repair protein RecT (prophage associated)
Cysteinyl-tRNA synthetase related protein
Manganese ABC transporter, periplasmic-binding protein SitA
Butyrate-acetoacetate CoA-transferase subunit B (EC 2.8.3.9)
Large-conductance mechanosensitive channel

Undecaprenyl-phosphate N-acetylglucosaminyl 1-phosphate transferase (EC 2.7.8.-)
ABC transporter, ATP-binding protein
D-alanyl-D-alanine carboxypeptidase (EC 3.4.16.4)
Mobile element protein
PTS system IIB component
Phosphatidate cytidylyltransferase (EC 2.7.7.41)
Phage protein
unknown
Transcriptional regulator OrfX
Late competence protein ComGB, access of DNA to ComEA
FIG01116268: hypothetical protein
Uracil-DNA glycosylase, family 1
3-ketoacyl-CoA thiolase (EC 2.3.1.16) @ Acetyl-CoA acetyltransferase (EC 2.3.1.9)
3-ketoacyl-CoA thiolase (EC 2.3.1.16) @ Acetyl-CoA acetyltransferase (EC 2.3.1.9)
ABC transporter ATP-binding protein YvcR
N-acetylmannosamine-6-phosphate 2-epimerase (EC 5.1.3.9)
Phosphoenolpyruvate carboxylase (EC 4.1.1.31)
hypothetical protein
Manganese superoxide dismutase (EC 1.15.1.1)
FIG01115226: hypothetical protein
Short chain dehydrogenase
Foldase protein prsA 1 precursor (EC 5.2.1.8)
PTS system, fructose-specific IIA component (EC 2.7.1.69) / PTS system, fructose-specific IIB component (EC 2.7.1.69) / PTS system, fructose-spe
D-beta-hydroxybutyrate permease
3-oxoacyl-[acyl-carrier protein] reductase (EC 1.1.1.100)
FIG01114659: hypothetical protein
Competence protein F homolog, phosphoribosyltransferase domain; protein YhgH required for utilization of DNA as sole source of carbon and e
Translation initiation factor 1
Lysyl aminopeptidase (EC 3.4.11.15)
Ribosomal large subunit pseudouridine synthase D (EC 4.2.1.70)
FIG001614: Membrane protein
tRNA-Val-TAC
Methionine ABC transporter substrate-binding protein
ATP synthase C chain (EC 3.6.3.14)
Glucan 1,6-alpha-glucosidase (EC 3.2.1.70)

Transcriptional regulator, GntR family
Metal-dependent hydrolase (EC 3.-.-.-)
Phage holin
Guanine-hypoxanthine permease
Late competence protein ComEC, DNA transport
tRNA-Arg-TCT
tmRNA-binding protein SmpB
Butyrate-acetoacetate CoA-transferase subunit A (EC 2.8.3.9)
Mobile element protein
Phage-related protein
Phosphoesterase, DHH family protein
Acetoin dehydrogenase E1 component beta-subunit (EC 1.2.4.-)
S-ribosylhomocysteine lyase (EC 4.4.1.21) / Autoinducer-2 production protein LuxS
Integral membrane protein
Phage terminase, large subunit
Hypothetical protein SPy2065
DNA recombination and repair protein RecF
Adenylate kinase (EC 2.7.4.3)
Ribonucleotide reductase of class III (anaerobic), activating protein (EC 1.97.1.4)
FIG01116522: hypothetical protein
Shikimate 5-dehydrogenase I alpha (EC 1.1.1.25)
ABC-type Fe3+-siderophore transport system, permease 2 component
Cell division protein FtsW
tRNA-specific adenosine-34 deaminase (EC 3.5.4.-)
Mobile element protein
FIG005935: membrane protein
Deoxyadenosine kinase (EC 2.7.1.76) / Deoxyguanosine kinase (EC 2.7.1.113)
Uridylate kinase (EC 2.7.4.-)
YfaA
O-methyltransferase family protein [C1]
Cell surface protein Shp, transfers heme from hemoglobin to apo-SiaA/HtsA
Signal peptidase I (EC 3.4.21.89)
D-beta-hydroxybutyrate dehydrogenase (EC 1.1.1.30)
tRNA-Val-TAC
hypothetical protein

Paratox
hypothetical protein
FIG01114729: hypothetical protein
Uracil phosphoribosyltransferase (EC 2.4.2.9)
Acetyl-CoA:acetoacetyl-CoA transferase, alpha subunit (EC 2.8.3.8)
ATP synthase beta chain (EC 3.6.3.14)
FIG01114033: hypothetical protein
Mobile element protein
Paratox
Glucose-1-phosphate thymidylyltransferase (EC 2.7.7.24)
FIG01115696: hypothetical protein
hypothetical phage protein
Antiphagocytic M protein
Phage integrase: site-specific recombinase
Mobile element protein
Cysteine desulfurase (EC 2.8.1.7)
COG1242: Predicted Fe-S oxidoreductase
Ribosomal small subunit pseudouridine synthase A (EC 4.2.1.70)
UDP-N-acetylmuramoylalanyl-D-glutamyl-2,6-diaminopimelate--D-alanyl-D-alanine ligase (EC 6.3.2.10)
Rhodanese domain protein UPF0176, Firmicutes subgroup
Enoyl-[acyl-carrier-protein] reductase [FMN] (EC 1.3.1.9)
hypothetical protein
Mobile element protein
LSU ribosomal protein L23p (L23Ae)
FIG01114249: hypothetical protein
V-type ATP synthase subunit G (EC 3.6.3.14)
FIG139598: Potential ribosomal protein
FIG01114281: hypothetical protein
GTP-binding and nucleic acid-binding protein YchF
putative repressor - phage associated
putative repressor C1
FIG146085: 3'-to-5' oligoribonuclease A, Bacillus type
Cell division protein FtsA
Topoisomerase IV subunit A (EC 5.99.1.-)
Unsaturated glucuronyl hydrolase (EC 3.2.1.-)

Hyaluronoglucosaminidase (EC 3.2.1.35)
MutT/nudix family protein
hypothetical phage protein
DNA repair protein RadA
Hydrolase (HAD superfamily)
Phosphopantothenoylcysteine decarboxylase (EC 4.1.1.36)
Ribonucleotide reductase transcriptional regulator NrdR
Streptococcal extracellular nuclease 2; Mitogenic factor 2
FIG001621: Zinc protease
Protein export cytoplasm protein SecA ATPase RNA helicase (TC 3.A.5.1.1)
FIG01116435: hypothetical protein
Substrate-specific component PdxU2 of predicted pyridoxin-related ECF transporter
Multidrug resistance protein B
Zinc finger protein
Predicted nucleoside phosphatase
Phage terminase, large subunit
Phage hyaluronidase
FIG01115531: hypothetical protein
UDP-glucose dehydrogenase (EC 1.1.1.22)
Phage excisionase
PTS system, galactose-specific IIB component (EC 2.7.1.69)
DegV family protein
Glycine/D-amino acid oxidases family
Phage protein
hypothetical protein
Ribonuclease M5 (EC 3.1.26.8)
FIG01114328: hypothetical protein
SSU ribosomal protein S4p (S9e)
Uridine kinase (EC 2.7.1.48) [C1]
Prophage Clp protease-like protein
Putative amidotransferase similar to cobyric acid synthase
COG1399 protein in cluster with ribosomal protein L32p, Firmicutes subfamily
Transcription regulator [contains diacylglycerol kinase catalytic domain]
Epoxyqueuosine (oQ) reductase QueG
Plasmid stabilization system toxin protein

kilA protein, putative phage-related DNA binding protein
M protein trans-acting positive regulator (Mga)
Peptide chain release factor 1
hypothetical protein
Formate--tetrahydrofolate ligase (EC 6.3.4.3)
FIG01114557: hypothetical protein
hypothetical protein
CDP-diacylglycerol--glycerol-3-phosphate 3-phosphatidyltransferase (EC 2.7.8.5)
Hypothetical protein ywlG
LemA protein
DNA-directed RNA polymerase alpha subunit (EC 2.7.7.6)
PTS system, sucrose-specific IIB component (EC 2.7.1.69) / PTS system, sucrose-specific IIC component (EC 2.7.1.69) / PTS system, sucrose-specif
Phage protein
Phage major tail shaft protein
FIG01114598: hypothetical protein
NAD-dependent oxidoreductase
General stress protein 13
Cell division protein FtsX
Putative metallopeptidase (Zinc) SprT family
Rhodanese-like domain protein
FIG01115469: hypothetical protein
tRNA-Trp-CCA
LSU ribosomal protein L29p (L35e)
Daunorubicin resistance ATP-binding protein drrA
Phage integrase: site-specific recombinase
Protein serine/threonine phosphatase PrpC, regulation of stationary phase
Competence-specific sigma factor ComX
LSU ribosomal protein L4p (L1e)
Uracil permease
Transcriptional regulator, Cro/CI family
hypothetical phage protein
Beta-phosphoglucomutase (EC 5.4.2.6) (EC 2.7.1.41)
ATP synthase B chain (EC 3.6.3.14)
Phage transcriptional repressor
Sucrose operon repressor ScrR, LacI family

Transcriptional regulator, GntR family
FIG001721: Predicted N6-adenine-specific DNA methylase
Transcriptional regulator DegU, LuxR family
tRNA-Asn-GTT
FIG053235: Diacylglucosamine hydrolase like
Acetyltransferase (EC 2.3.1.-)
Phage protein
Dihydrofolate synthase (EC 6.3.2.12) / Folylpolyglutamate synthase (EC 6.3.2.17)
SSU ribosomal protein S11p (S14e)
ABC transporter membrane-spanning permease - glutamine transport
FIG01114188: hypothetical protein
FIG01114317: hypothetical protein
Laminin-binding surface protein
hypothetical protein
Capsule biosynthesis protein capA
tRNA-Gln-TTG
conserved hypothetical protein - phage associated
Adenylosuccinate synthetase (EC 6.3.4.4)
FIG033577: Phage protein
Phage endopeptidase
hypothetical protein
Phage protein
LSU ribosomal protein L28p
Hyaluronate lyase precursor (EC 4.2.2.1)
NAD-dependent protein deacetylase of SIR2 family
5S RNA
Lipase/Acylhydrolase family protein
ABC-type multidrug transport system, ATPase component
hypothetical protein
internalin, putative
Arsenate reductase (EC 1.20.4.1)
Cell division protein DivIC (FtsB), stabilizes FtsL against RasP cleavage
hypothetical protein
M protein trans-acting positive regulator (Mga)
conserved hypothetical protein - phage associated

FIG01115656: hypothetical protein
Holliday junction DNA helicase RuvB
Membrane protein, putative
Phage holin
GMP reductase (EC 1.7.1.7)
FIG01115695: hypothetical protein
FIG01114865: hypothetical protein
Phage protein
Dihydrofolate reductase (EC 1.5.1.3)
Phosphate transport ATP-binding protein PstB (TC 3.A.1.7.1)
INTEGRAL MEMBRANE PROTEIN (Rhomboid family)
rhamnose-containing polysacharide translocation permease
FIG002813: LPPG:FO 2-phospho-L-lactate transferase like, CofD-like
Orotate phosphoribosyltransferase (EC 2.4.2.10)
ABC transporter ATP-binding protein
Lactose phosphotransferase system repressor
hypothetical phage protein
FIG009886: phosphoesterase
FIG01116077: hypothetical protein
Oligopeptide transport system permease protein OppB (TC 3.A.1.5.1)
6-aminohexanoate-cyclic-dimer hydrolase (EC 3.5.2.12)
hypothetical protein, phage associated
Mobile element protein
Serine/threonine protein kinase PrkC, regulator of stationary phase
ABC transporter, permease protein
hypothetical protein - phage associated
Substrate-specific component PanT of predicted pantothenate ECF transporter
FIG01114225: hypothetical protein
FIG01114161: hypothetical protein
Helicase loader DnaI
DinG family ATP-dependent helicase YoaA
FIG04612: Integral membrane protein (putative)
Thioredoxin
Ribosomal-protein-S5p-alanine acetyltransferase
Zinc ABC transporter, ATP-binding protein ZnuC

Dihydrolipoamide acyltransferases
hypothetical phage protein
membrane protein, related to Actinobacillus protein (1944168)
hypothetical protein
SSU ribosomal protein S2p (SAe)
FIG01114171: hypothetical protein
Late competence protein ComGG, FIG068335
FIG01115182: hypothetical protein
Phosphate starvation-inducible protein PhoH, predicted ATPase
Cell division protein YlmG/Ycf19 (putative), YggT family
FIG01115329: hypothetical protein
NAD-dependent glyceraldehyde-3-phosphate dehydrogenase (EC 1.2.1.12) / Plasmin(ogen) receptor
Xaa-Pro dipeptidyl-peptidase (EC 3.4.14.11)
Phage protein
FIG01115104: hypothetical protein
SSU rRNA (adenine(1518)-N(6)/adenine(1519)-N(6))-dimethyltransferase (EC 2.1.1.182)
FIG01114555: hypothetical protein
Hydrolase (HAD superfamily)
Phage-associated homing endonuclease
2-(5''-triphosphoribosyl)-3'-dephosphocoenzyme-A synthase (EC 2.7.8.25)
Bacteriocin-like peptide M BlpM
FIG01114490: hypothetical protein
Tagatose 1,6-diphosphate aldolase (EC 4.1.2.40)
Ribosomal RNA small subunit methyltransferase E (EC 2.1.1.-)
Phage-related protein
Voltage-gated chloride channel family protein
Transcriptional repressor of the fructose operon, DeoR family
Phage protein
4-oxalocrotonate tautomerase (EC 5.3.2.-); Xylose transport system permease protein xylH
tRNA-Glu-TTC
UDP-N-acetylmuramate--alanine ligase (EC 6.3.2.8)
N-acetylneuraminate lyase (EC 4.1.3.3)
Peptide chain release factor 3
RecD-like DNA helicase YrrC
paratox?

FIG01114335: hypothetical protein
Arginyl-tRNA synthetase (EC 6.1.1.19)
DNA polymerase III epsilon subunit (EC 2.7.7.7)
Transcriptional regulator, MarR family
Two-component sensor histidine kinase, malate (EC 2.7.3.-)
tRNA-Leu-CAA
Arginine pathway regulatory protein ArgR, repressor of arg regulon
Exfoliative toxin A
Xanthine phosphoribosyltransferase (EC 2.4.2.22)
Dihydrolipoamide acetyltransferase component (E2) of acetoin dehydrogenase complex (EC 2.3.1.-)
Chorismate mutase I (EC 5.4.99.5)
Nicotinate phosphoribosyltransferase (EC 2.4.2.11)
Dihydropteroate synthase (EC 2.5.1.15)
5'-nucleotidase (EC 3.1.3.5)
ribosomal protein L7Ae family protein
5S RNA
Fibronectin-binding protein
Membrane protein involved in the export of O-antigen, teichoic acid lipoteichoic acids
FIG01115477: hypothetical protein
replication initiator protein A
hypothetical protein
Substrate-specific component CbrT of predicted cobalamin ECF transporter
Replicative DNA helicase (DnaB) (EC 3.6.4.12)
unknown domain / Nucleoside 5-triphosphatase RdgB (dHAPTP, dITP, XTP-specific) (EC 3.6.1.15)
Protein of unknown function DUF1447
Topoisomerase IV subunit B (EC 5.99.1.-)
Mobile element protein
Cytidine deaminase (EC 3.5.4.5)
Putative signal peptidase I (EC 3.4.21.89)
ABC transporter membrane-spanning permease - glutamine transport
DNA recombination and repair protein RecO
membrane protein, putative
Mobile element protein
DNA-directed RNA polymerase omega subunit (EC 2.7.7.6)
Transcriptional repressor for NAD biosynthesis in gram-positives

DNA-directed RNA polymerase beta' subunit (EC 2.7.7.6)
conserved hypothetical protein - phage associated
Ribonuclease III (EC 3.1.26.3)
Phosphomevalonate kinase (EC 2.7.4.2)
Deoxyuridine 5'-triphosphate nucleotidohydrolase (EC 3.6.1.23)
tRNA-Thr-TGT
Nucleoside permease NupC
16S rRNA (cytosine(967)-C(5))-methyltransferase (EC 2.1.1.176)
Ferrichrome transport ATP-binding protein FhuC (TC 3.A.1.14.3)
Two component system sensor histidine kinase CiaH (EC 2.7.3.-)
Undecaprenyl-diphosphatase (EC 3.6.1.27)
Transcription-repair coupling factor
Aspartate carbamoyltransferase (EC 2.1.3.2)
Translation elongation factor G
Transporter
GTP-binding protein HflX
Hydroxymethylglutaryl-CoA synthase (EC 2.3.3.10)
FIG005686: hypothetical protein
Methionine ABC transporter permease protein
hypothetical protein
Transcription regulator [contains diacylglycerol kinase catalytic domain]
Transcriptional regulator, XRE family
Tributyrin esterase
Phage protein
Potassium uptake protein TrkH
Chaperonin (heat shock protein 33)
CAMP factor
LSU ribosomal protein L31p @ LSU ribosomal protein L31p, zinc-independent
TPR-repeat-containing protein
ABC-type polar amino acid transport system, ATPase component
tRNA (adenine37-N(6))-methyltransferase TrmN6 (EC 2.1.1.223)
ATP synthase alpha chain (EC 3.6.3.14)
Colicin V production protein
FIG01114439: hypothetical protein
Glyoxalase family protein

hypothetical protein
hypothetical protein
FIG001583: hypothetical protein, contains S4-like RNA binding domain
Late competence protein ComGC, access of DNA to ComEA, FIG007487
Thiamin pyrophosphokinase (EC 2.7.6.2)
Glutaminyl-peptide cyclotransferase (EC 2.3.2.5)
Two-component response regulator, associated with ferric iron transporter, SPy1062 homolog
ABC transporter ATP-binding protein
Acetoin dehydrogenase E1 component alpha-subunit (EC 1.2.4.-)
ABC transporter ATP-binding protein
Peptidoglycan hydrolase, Autolysin1 (EC 3.5.1.28)
FIG01114815: hypothetical protein
SSU ribosomal protein S17p (S11e)
Lanthionine biosynthesis protein LanM/pseudogene causing stop codon at 177th aa
Late competence protein ComGA, access of DNA to ComEA
Hydrolase (HAD superfamily)
Glutathione S-transferase domain protein (class beta)
Substrate-specific component RibU of riboflavin ECF transporter
Lon-like protease with PDZ domain
Non-phosphorylating glyceraldehyde-3-phosphate dehydrogenase (NADP) (EC 1.2.1.9)
aminoglycoside phosphotransferase family protein
Collagen-like surface protein
Phosphoglycerate mutase family 5
Mutator mutT protein (7,8-dihydro-8-oxoguanine-triphosphatase) (EC 3.6.1.-)
Iron-sulfur cluster assembly ATPase protein SufC
Sodium/glycine symporter GlyP
D-alanine--poly(phosphoribitol) ligase subunit 1 (EC 6.1.1.13)
Peptide ABC transporter ATP binding/permease protein
tRNA-Leu-TAG
Succinyl-CoA synthetase, alpha subunit-related enzymes
Negative regulator of genetic competence MecA
Transcriptional repressor AdcR for Zn(2+)-responsive expression
FIG01114232: hypothetical protein
hypothetical phage protein
2-deoxy-D-gluconate 3-dehydrogenase (EC 1.1.1.125)

Substrate-specific component PdxU of predicted pyridoxine ECF transporter
Replicative DNA helicase (DnaB) (EC 3.6.4.12)
Helicase loader DnaI
Helicase PriA essential for oriC/DnaA-independent DNA replication
FIG01114733: hypothetical protein
ABC transporter substrate-binding protein
Excinuclease ABC subunit B
Cell division protein FtsL
Septation ring formation regulator EzrA
Maltose/maltodextrin ABC transporter, substrate binding periplasmic protein MalE
Alanine racemase (EC 5.1.1.1)
UPF0246 protein YaaA
Lipoteichoic acid synthase LtaS Type IIc (The first GCT-Ala is overlapped with the last codon of the former half interrupted protein)
FIG01118080: hypothetical protein
Mutator mutT protein (7,8-dihydro-8-oxoguanine-triphosphatase) (EC 3.6.1.-)
putative esterase
FIG01114068: hypothetical protein
V-type ATP synthase subunit A (EC 3.6.3.14)
ABC transporter permease protein
Ferric iron ABC transporter, iron-binding protein
Tagatose-6-phosphate kinase (EC 2.7.1.144)
ATP-dependent nuclease, subunit A
FIG01114175: hypothetical protein
Negative transcriptional regulator-copper transport operon
Phage hyaluronidase
Deoxyribodipyrimidine photolyase (EC 4.1.99.3)
Sensor histidine kinase VraS
FIG01115767: hypothetical protein
ABC transporter ATP-binding protein
predicted 4-deoxy-L-threo-5-hexosulose-uronate ketol-isomerase (EC 5.3.1.17)
Hypothetical protein DUF901, similar to C-terminal domain of ribosome protection-type Tc-resistance proteins
Maltodextrin phosphorylase (EC 2.4.1.1)
Transcriptional regulators, LysR family
tRNA and rRNA cytosine-C5-methylases
conserved hypothetical protein - phage associated

Late competence protein ComEA, DNA receptor
Amino acid ABC transporter, amino acid-binding protein
Aspartyl-tRNA(Asn) amidotransferase subunit A (EC 6.3.5.6) @ Glutamyl-tRNA(Gln) amidotransferase subunit A (EC 6.3.5.7)
proposed amino acid ligase found clustered with an amidotransferase
Cadmium efflux system accessory protein
Sensory transduction protein kinase (EC 2.7.3.-)
Phenylalanyl-tRNA synthetase domain protein (Bsu YtpR)
FIG006438: hypothetical protein
Phage protein
Glutamyl aminopeptidase (EC 3.4.11.7)
tRNA-Arg-ACG
Phage holin
FIG01115146: hypothetical protein
FIG01114378: hypothetical protein
Putative ABC transporter (ATP-binding protein), spy1791 homolog
FIG137478: Hypothetical protein
FIG01114855: hypothetical protein
hypothetical protein
FIG001960: FtsZ-interacting protein related to cell division
Thymidine kinase (EC 2.7.1.21)
Methylenetetrahydrofolate dehydrogenase (NADP+) (EC 1.5.1.5) / Methenyltetrahydrofolate cyclohydrolase (EC 3.5.4.9)
Transcriptional regulator CtsR
Tyrosine recombinase XerD
Chromosome segregation helicase
Peptide methionine sulfoxide reductase MsrA (EC 1.8.4.11)
hypothetical phage protein
Recombination inhibitory protein MutS2
Acetyltransferase (EC 2.3.1.-)
LSU ribosomal protein L11p (L12e)
Cysteine desulfurase (EC 2.8.1.7)
GTP-binding protein EngB
Ribonucleotide reductase of class Ib (aerobic), alpha subunit (EC 1.17.4.1)
Transcriptional regulator, TetR family
C3-degrading proteinase
Phage terminase large subunit

Riboflavin kinase (EC 2.7.1.26) / FMN adenylyltransferase (EC 2.7.7.2)
TsaC protein (YrdC domain) required for threonylcarbamoyladenosine t(6)A37 modification in tRNA
Archaeal S-adenosylmethionine synthetase (EC 2.5.1.6)
Transporter, MFS superfamily
Phage antirepressor protein
putative methyltransferase
hypothetical protein
tRNA-Lys-TTT
Iojap protein
Pyrroline-5-carboxylate reductase (EC 1.5.1.2)
L-proline glycine betaine ABC transport system permease protein ProV (TC 3.A.1.12.1)
FIG014387: Transcriptional regulator, PadR family
Transcriptional regulator, GntR family
Holo-[acyl-carrier protein] synthase (EC 2.7.8.7)
16S rRNA processing protein RimM
Quinolinate phosphoribosyltransferase [decarboxylating] (EC 2.4.2.19)
Dihydroorotase (EC 3.5.2.3)
hypothetical phage protein
3-dehydroquinate synthase (EC 4.2.3.4)
Adenine-specific methyltransferase (EC 2.1.1.72)
LSU ribosomal protein L21p
Oligoendopeptidase F
putative choline binding protein
Phage tail length tape-measure protein
Phosphopantetheine adenylyltransferase (EC 2.7.7.3)
Carbamoyl-phosphate synthase large chain (EC 6.3.5.5)
Acetyl-coenzyme A carboxyl transferase alpha chain (EC 6.4.1.2)
Heat shock protein GrpE
Glycerol-3-phosphate dehydrogenase [NAD(P)+] (EC 1.1.1.94)
Aspartate--ammonia ligase (EC 6.3.1.1)
Surface exclusion protein
Ribonucleotide reduction protein NrdI
Orotidine 5'-phosphate decarboxylase (EC 4.1.1.23)
Ferredoxin
Transcriptional regulator, RofA-like Protein (RALP)

FIG01116065: hypothetical protein
Diphosphomevalonate decarboxylase (EC 4.1.1.33)
Cell division protein FtsZ (EC 3.4.24.-)
Phage protein
FIG01114733: hypothetical protein
Streptokinase
Malonate permease
FIG01114229: hypothetical protein
Streptococcal pyrogenic exotoxin K (SpeK)
Phage protein
Type I restriction-modification system, DNA-methyltransferase subunit M (EC 2.1.1.72)
N-acetylmannosamine kinase (EC 2.7.1.60)
Phage minor capsid protein #Fam0015
hypothetical protein
Methionine biosynthesis and transport regulator MtaR, LysR family
hypothetical protein
hypothetical protein
Dihydrolipoamide dehydrogenase of acetoin dehydrogenase (EC 1.8.1.4)
Exodeoxyribonuclease VII small subunit (EC 3.1.11.6)
COG2827: putative endonuclease containing a URI domain
Phage portal protein
Phage transcriptional regulator, Cro/CI family
Heat shock protein 60 family co-chaperone GroES
Ascorbate-specific PTS system, EIIC component
tRNA-Ile-GAT
Asparaginyl-tRNA synthetase (EC 6.1.1.22)
Acyl carrier protein
Biotin operon repressor / Biotin-protein ligase (EC 6.3.4.15)
Aspartyl-tRNA synthetase (EC 6.1.1.12)
Dihydroneopterin aldolase (EC 4.1.2.25)
Substrate-specific component BL0695 of predicted ECF transporter
Hypothetical protein, Spy1939 homolog
putative ABC transporter (ATP-binding protein)
Ribulose-phosphate 3-epimerase (EC 5.1.3.1)
Molybdopterin binding motif, CinA N-terminal domain / C-terminal domain of CinA type S

hypothetical protein - phage associated
Methionine ABC transporter ATP-binding protein
Copper chaperone
Positive transcriptional regulator, MutR family
Cell division protein FtsK
Membrane protein
Xanthine permease
Mobile element protein
Ribosomal-protein-S18p-alanine acetyltransferase (EC 2.3.1.-)
Mobile element protein
ABC-type multidrug transport system, ATPase component
hypothetical phage protein
Permease
FIG01116071: hypothetical protein
FIG145533: Methyltransferase (EC 2.1.1.-)
FIG01115263: hypothetical protein
Pyrrolidone-carboxylate peptidase (EC 3.4.19.3)
uncharacterized secreted protein, YBBR Bacillus subtilis homolog
Novel pyridoxal kinase, thiD family (EC 2.7.1.35)
Histidine triad (HIT) nucleotide-binding protein, similarity with At5g48545 and yeast YDL125C (HNT1)
DNA repair protein RadC
tRNA-Gly-TCC
Pyruvate formate-lyase activating enzyme (EC 1.97.1.4)
ATP-dependent RNA helicase YqfR
GTP-binding protein Era
hypothetical protein
Thiol-activated cytolysin
hypothetical phage protein
ATP-dependent DNA helicase UvrD/PcrA
Chloride channel protein
GTP pyrophosphokinase (EC 2.7.6.5)
DNA polymerase I (EC 2.7.7.7)
Heme ABC transporter (Streptococcus), ATP-binding protein
Phosphoribosylformylglycinamidine synthase, synthetase subunit (EC 6.3.5.3) / Phosphoribosylformylglycinamidine synthase, glutamine amid
Transcriptional regulator, MarR family

Heme ABC transporter (Streptococcus), permease protein
Transcription elongation factor GreA
Phage protein
Integrase
Galactose-6-phosphate isomerase, LacB subunit (EC 5.3.1.26)
Regulatory protein recX
Hypothetical protein SPy1656
Phosphate transport system regulatory protein PhoU
Alkylphosphonate utilization operon protein PhnA
ATP-dependent Clp protease ATP-binding subunit ClpX
tRNA (Guanine37-N1) -methyltransferase (EC 2.1.1.31)
Holliday junction DNA helicase RuvA
Salivaricin A modification enzyme
Ribosome recycling factor
Putative stomatin/prohibitin-family membrane protease subunit YbbK
Peroxide stress regulator PerR, FUR family
N-acetylglucosamine-1-phosphate uridyltransferase (EC 2.7.7.23) / Glucosamine-1-phosphate N-acetyltransferase (EC 2.3.1.157)
PTS system, lactose-specific IIA component (EC 2.7.1.69)
putative ATP-dependent Clp proteinase (ATP-binding subunit)
Transposase (maltose area)
Lantibiotic salivaricin A
Substrate-specific component ThiT of thiamin ECF transporter
hypothetical phage protein
FIG009688: Thioredoxin
ABC transporter permease protein
ATPase component of general energizing module of ECF transporters
FIG01115822: hypothetical protein
Lactoylglutathione lyase (EC 4.4.1.5)
Phage major capsid protein
Phage protein
FIG004454: RNA binding protein
Peptide deformylase (EC 3.5.1.88)
3-oxoacyl-[acyl-carrier-protein] synthase, KASIII (EC 2.3.1.41)
Sialic acid utilization regulator, RpiR family
Hyaluronate lyase (phage associated)

Hypothetical protein possible functionally linked with Alanyl-tRNA synthetase
FIG01114419: hypothetical protein
UDP-N-acetylenolpyruvoylglucosamine reductase (EC 1.1.1.158)
Fructokinase (EC 2.7.1.4)
hypothetical protein
FIG01114628: hypothetical protein
Translation initiation factor 2
DNA polymerase III delta prime subunit (EC 2.7.7.7)
Phage holin
Phosphoribosylformylglycinamidine cyclo-ligase (EC 6.3.3.1)
Lysyl-tRNA synthetase (class II) (EC 6.1.1.6)
Lead, cadmium, zinc and mercury transporting ATPase (EC 3.6.3.3) (EC 3.6.3.5); Copper-translocating P-type ATPase (EC 3.6.3.4)
hypothetical phage protein
Phosphoenolpyruvate-protein phosphotransferase of PTS system (EC 2.7.3.9)
Two component system response regulator CiaR
Transcriptional regulator, TetR family
Amidophosphoribosyltransferase (EC 2.4.2.14)
tRNA-Ile-GAT
Thymidylate synthase (EC 2.1.1.45)
FIG01115181: hypothetical protein
Phage protein
Ribosomal large subunit pseudouridine synthase B (EC 4.2.1.70)
Transcriptional regulator
Alpha-mannosidase (EC 3.2.1.24)
Immunogenic secreted protein
tRNA-Phe-GAA
tRNA-Glu-TTC
Oligopeptide transport ATP-binding protein OppF (TC 3.A.1.5.1)
Spermidine Putrescine ABC transporter permease component PotB (TC 3.A.1.11.1)
FIG01115797: hypothetical protein
tRNA-Ile-GAT
Mobile element protein
ATP-dependent DNA helicase RecG (EC 3.6.1.-)
Two-component sensor kinase, associated with ferric iron transporter, SPy1061 homolog
FIG01116220: hypothetical protein

Phage tail length tape-measure protein
tRNA-Asp-GTC
Fructose-bisphosphate aldolase class II (EC 4.1.2.13)
PTS system, N-acetylgalactosamine- and galactosamine-specific IIA component (EC 2.7.1.69)
Late competence protein ComGE, FIG075573
Gamma-glutamyl phosphate reductase (EC 1.2.1.41)
Fibronectin-binding protein
conserved hypothetical protein
hypothetical protein
Guanylate kinase (EC 2.7.4.8)
tRNA-Gly-GCC
Cold shock protein CspA
SAM-dependent methyltransferase, MraW methylase family (EC 2.1.1.-)
hypothetical protein
Single-stranded DNA-binding protein
Hypothetical similar to thiamin biosynthesis lipoprotein ApbE
Hypothetical protein in cluster with Ecs transporter (in Streptococci)
hypothetical protein
LSU ribosomal protein L34p
tRNA-Leu-TAG
hypothetical protein
Phage terminase, small subunit
Glycerate kinase (EC 2.7.1.31)
Cro-like phage transcriptional repressor protein
Segregation and condensation protein A
Hypothetical protein YbbP, contains nucleotide-binding domain of DisA bacterial checkpoint controller
Mobile element protein
Transcriptional antiterminator with PTS regulation domain, SPy0181 ortholog
putative role in replication
Mobile element protein
Potassium efflux system KefA protein / Small-conductance mechanosensitive channel
hypothetical phage protein
hypothetical protein BH3604
FIG01114548: hypothetical protein
Tagatose 1,6-diphosphate aldolase (EC 4.1.2.40)

SAM-dependent methyltransferase HI0095 (UbiE paralog)
Paratox
Segregation and condensation protein B
Thioredoxin reductase (EC 1.8.1.9)
GTP-binding protein TypA/BipA
Phage hyaluronidase; Phage endopeptidase
Leucine rich protein
Mobile element protein
putative replication protein
Chromosome (plasmid) partitioning protein ParB / Stage 0 sporulation protein J
L-xylulose 5-phosphate 3-epimerase (EC 5.1.3.-)
Mobile element protein
Protein G-related alpha 2 macroglobulin-binding protein (GRAB)
FIG01114073: hypothetical protein
Preprotein translocase subunit YajC (TC 3.A.5.1.1)
Cytoplasmic copper homeostasis protein CutC
D-alanyl-D-alanine carboxypeptidase (EC 3.4.16.4)
Preprotein translocase subunit SecE (TC 3.A.5.1.1)
Formate/nitrite family of transporters
FIG042840: Phage protein
Peptidyl-prolyl cis-trans isomerase (EC 5.2.1.8)
Transcriptional regulator, Cro/CI family
Membrane-associated phospholipid phosphatase
putative toxic anion resistance protein
Adenine phosphoribosyltransferase (EC 2.4.2.7)
PTS system, lactose-specific IIB component (EC 2.7.1.69) / PTS system, lactose-specific IIC component (EC 2.7.1.69)
DNA replication protein DnaC
Cell division protein GpsB, coordinates the switch between cylindrical and septal cell wall synthesis by re-localization of PBP1
Mn-dependent transcriptional regulator MntR
Methionyl-tRNA synthetase (EC 6.1.1.10)
Putative oxidoreductase
Glutamate 5-kinase (EC 2.7.2.11)
FIG01114159: hypothetical protein
FIG011178: rRNA methylase
B-cell receptor associated protein-related protein

ATP-dependent Clp protease proteolytic subunit (EC 3.4.21.92)
Uridine phosphorylase (EC 2.4.2.3)
Transcriptional regulator in cluster with beta-lactamase, GntR family
hypothetical protein
Transcriptional regulator SpxA2
Serine acetyltransferase (EC 2.3.1.30)
Cobalt-zinc-cadmium resistance protein CzcD
FIG01116946: hypothetical protein
hypothetical protein
Glutamyl-tRNA synthetase (EC 6.1.1.17) @ Glutamyl-tRNA(Gln) synthetase (EC 6.1.1.24)
Cytochrome c-type biogenesis protein CcdA homolog, associated with MetSO reductase
FIG015049: hypothetical protein
FIG01114091: hypothetical protein
Malonate permease
COG2740: Predicted nucleic-acid-binding protein implicated in transcription termination
Mobile element protein
Immunoreactive protein Se23.5 (Fragment)
SSU ribosomal protein S21p
Phage protein
Oxygen-insensitive NAD(P)H nitroreductase (EC 1.-.-.-) / Dihydropteridine reductase (EC 1.5.1.34)
tRNA-Glu-TTC
Aspartyl-tRNA(Asn) amidotransferase subunit B (EC 6.3.5.6) @ Glutamyl-tRNA(Gln) amidotransferase subunit B (EC 6.3.5.7)
Chromosomal replication initiator protein DnaA
FIG007491: hypothetical protein YeeN
Valyl-tRNA synthetase (EC 6.1.1.9)
hypothetical protein, phage associated
hypothetical protein
FIG01116130: hypothetical protein
6-phosphofructokinase (EC 2.7.1.11)
Cell envelope-associated transcriptional attenuator LytR-CpsA-Psr, subfamily F1 (as in PMID19099556)
Trehalose operon transcriptional repressor
Histidine kinase of the competence regulon ComD
Magnesium and cobalt transport protein CorA
SSU ribosomal protein S16p
FtsK/SpoIIIE family

FIG01114318: hypothetical protein
Short chain fatty acids transporter
Lipid A export ATP-binding/permease protein MsbA
Phage minor capsid protein
SSU ribosomal protein S20p
PTS system, cellobiose-specific IIC component (EC 2.7.1.69)
Dephospho-CoA kinase (EC 2.7.1.24)
FIG00525014: hypothetical protein
ABC transporter, predicted N-acetylneuraminate transport system permease protein 2
Prolyl-tRNA synthetase (EC 6.1.1.15), bacterial type
tRNA-Lys-TTT
DNA mismatch repair protein MutL
FIG007079: UPF0348 protein family
Late competence protein ComGF, access of DNA to ComEA, FIG012620
FIG01114114: hypothetical protein
5S RNA
putative L-glutamate ligase
Phage encoded transcriptional regulator, ArpU family
Excinuclease ABC subunit A
tRNA:m(5)U-54 MTase gid
Tagatose-6-phosphate kinase (EC 2.7.1.144)
D-alanyl-D-alanine carboxypeptidase (EC 3.4.16.4)
FIG01114217: hypothetical protein
Universal stress protein family
DNA-entry nuclease (Competence-specific nuclease) (EC 3.1.30.-)
Cystathionine beta-lyase (EC 4.4.1.8)
Streptococcal mitogenic exotoxin Z (SmeZ)
FIG001886: Cytoplasmic hypothetical protein
Major facilitator:Oxalate:Formate Antiporter
Phage protein
tRNA-Ser-GGA
Glutamine amidotransferase, class I
Endonuclease III (EC 4.2.99.18)
hypothetical protein
hypothetical cytosolic protein

Sodium-dependent phosphate transporter
phage protein
hypothetical phage protein
Phage minor tail protein
Transcription termination protein NusA
PurR: transcription regulator associated with purine metabolism
FIG01116574: hypothetical protein
Chromosome segregation ATPase
DUF124 domain-containing protein
Histidyl-tRNA synthetase (EC 6.1.1.21)
hypothetical protein
Putrescine transport ATP-binding protein PotA (TC 3.A.1.11.1)
Phage protein
Membrane protein
putative transcriptional activator - phage associated
Translation elongation factor LepA
FIG01114048: hypothetical protein
Protein YidD
Mobile element protein
FIG01115059: hypothetical protein
FIG01118151: hypothetical protein
Immunodominant antigen A
hypothetical phage protein
Hypothetical protein SPy0887
ATP synthase gamma chain (EC 3.6.3.14)
DNA-3-methyladenine glycosylase (EC 3.2.2.20)
[Citrate [pro-3S]-lyase] ligase (EC 6.2.1.22)
tRNA-Pseudo-GCT
L-asparaginase (EC 3.5.1.1)
Transcriptional regulators, LysR family
FIG009210: peptidase, M16 family
DNA-directed RNA polymerase beta subunit (EC 2.7.7.6)
Transcriptional repressor of arabinoside utilization operon, GntR family
FIG01116936: hypothetical protein
Pleiotropic regulator of exopolysaccharide synthesis, competence and biofilm formation Ftr, XRE family

Metal-dependent hydrolase YbeY, involved in rRNA and/or ribosome maturation and assembly
Phage integrase: site-specific recombinase
Signal recognition particle receptor protein FtsY (=alpha subunit) (TC 3.A.5.1.1)
Hypothetical protein VC0266 (sugar utilization related?)
Heme transporter IsdDEF, lipoprotein IsdE
hypothetical protein
Fibronectin/fibrinogen-binding protein
DNA polymerase III delta subunit (EC 2.7.7.7)
Mobile element protein
Two-component sensor kinase yesM (EC 2.7.3.-), associated with MetSO reductase
Collagen-like surface protein
NAD(FAD)-utilizing dehydrogenases
Maltose/maltodextrin ABC transporter, permease protein MalG
Ribosomal protein L11 methyltransferase (EC 2.1.1.-)
Transcriptional regulator, HxlR family
Oligopeptide transport ATP-binding protein OppD (TC 3.A.1.5.1)
Phage integrase: site-specific recombinase
Oxidoreductase, Gfo/Idh/MocA family
Acyl-phosphate:glycerol-3-phosphate O-acyltransferase PlsY
Substrate-specific component BioY of biotin ECF transporter
Daunorubicin resistance transmembrane protein
Lipoate-protein ligase A
FIG01115198: hypothetical protein
Phosphate ABC transporter, periplasmic phosphate-binding protein PstS (TC 3.A.1.7.1)
D-alanine--poly(phosphoribitol) ligase subunit 2 (EC 6.1.1.13)
Similar to ribosomal large subunit pseudouridine synthase D, Bacillus subtilis YhcT type
Integral membrane protein (Rhomboid family)
Streptococcal histidine triad protein
Pyruvate formate-lyase (EC 2.3.1.54)
Signal peptidase I (EC 3.4.21.89)
Glycosyl transferase, group 2 family protein
Ribonucleotide reductase of class III (anaerobic), large subunit (EC 1.17.4.2)
Hypothetical DUF1027 domain protein
ATPase component of general energizing module of ECF transporters
FIG002958: hypothetical protein

Iron-sulfur cluster assembly protein SufD
hypothetical protein
hypothetical protein
Hydroxymethylglutaryl-CoA reductase (EC 1.1.1.34)
Transcriptional regulator
Biotinyl-lipoyl attachment domain protein, GcvH-like
Antiphagocytic M protein/pseudogene causing stop codon at 85th aa
SSU ribosomal protein S7p (S5e)
ABC-type Fe3+-siderophore transport system, permease component
RecU Holliday junction resolvase
Transporter associated with VraSR
Mobile element protein
Phage protein
Mobile element protein
hypothetical protein
tRNA-Gly-GCC
Phosphoribosylaminoimidazole-succinocarboxamide synthase (EC 6.3.2.6)
Para-aminobenzoate synthase, aminase component (EC 2.6.1.85) / Aminodeoxychorismate lyase (EC 4.1.3.38)
Quinolinate phosphoribosyltransferase [decarboxylating] (EC 2.4.2.19)
Lipid A export ATP-binding/permease protein MsbA
Immunogenic secreted protein
FIG001802: Putative alkaline-shock protein
Carbonic anhydrase (EC 4.2.1.1)
Nucleoside diphosphate kinase (EC 2.7.4.6)
Methionine ABC transporter ATP-binding protein
Duplicated ATPase component BL0693 of energizing module of predicted ECF transporter
hypothetical protein
Acyltransferase family
Cystine transport system permease protein
FIG01117510: hypothetical protein
hypothetical protein
Chromosome partition protein smc
tRNA-Arg-CCT
FIG01114610: hypothetical protein
Glycine betaine ABC transport system, ATP-binding protein OpuAA (EC 3.6.3.32)

Anthranilate synthase, amidotransferase component (EC 4.1.3.27) @ Para-aminobenzoate synthase, amidotransferase component (EC 2.6.1.85
tRNA-Gly-TCC
Glucosamine--fructose-6-phosphate aminotransferase [isomerizing] (EC 2.6.1.16)
RNA binding methyltransferase FtsJ like
Manganese ABC transporter, inner membrane permease protein SitD
ATP synthase epsilon chain (EC 3.6.3.14)
tRNA-Val-TAC
Late competence protein ComC, processing protease
DNA ligase (EC 6.5.1.2)
Ascorbate-specific PTS system, EIIA component (EC 2.7.1.-)
Oligopeptide transport system permease protein OppC (TC 3.A.1.5.1)
Aldose 1-epimerase (EC 5.1.3.3)
Adenylate cyclase (EC 4.6.1.1)
Inner membrane protein translocase component YidC, OxaA protein
SSU ribosomal protein S10p (S20e)
Hypoxanthine-guanine phosphoribosyltransferase (EC 2.4.2.8)
FIG01115109: hypothetical protein
hypothetical phage protein
4-alpha-glucanotransferase (amylomaltase) (EC 2.4.1.25)
Competence-induced protein Ccs4
FIG01118158: hypothetical protein
Mobile element protein
Pyruvate kinase (EC 2.7.1.40)
Phage protein
tRNA-Asp-GTC
UDP-N-acetylglucosamine 1-carboxyvinyltransferase (EC 2.5.1.7)
IMP cyclohydrolase (EC 3.5.4.10) / Phosphoribosylaminoimidazolecarboxamide formyltransferase (EC 2.1.2.3)
RecA protein
Multidrug resistance efflux pump PmrA
Multimodular transpeptidase-transglycosylase (EC 2.4.1.129) (EC 3.4.-.-)
DNA polymerase III beta subunit (EC 2.7.7.7)
Isopentenyl-diphosphate delta-isomerase, FMN-dependent (EC 5.3.3.2)
Mannose-6-phosphate isomerase (EC 5.3.1.8)
Mobile element protein
LSU ribosomal protein L3p (L3e)

Cell envelope-associated transcriptional attenuator LytR-CpsA-Psr, subfamily F2 (as in PMID19099556)
Phage protein
hypothetical protein
tRNA dimethylallyltransferase (EC 2.5.1.75)
Transcriptional regulator, RofA-like Protein (RALP)
S-adenosylmethionine synthetase (EC 2.5.1.6)
Two-component response regulator SA14-24
Two-component sensor kinase SA14-24
putative glycosyltransferase - possibly involved in cell wall localization and side chain formation of rhamnose-glucose polysaccharide
Peptidoglycan N-acetylglucosamine deacetylase (EC 3.5.1.-)
Aspartate aminotransferase (EC 2.6.1.1)
Streptococcal pyrogenic exotoxin G (SpeG)
Acetyltransferase (EC 2.3.1.-)
RNA-binding protein Jag
Preprotein translocase subunit SecG (TC 3.A.5.1.1)
Competence-associated EpuA protein
Phosphoribosylaminoimidazole carboxylase catalytic subunit (EC 4.1.1.21)
tRNA-Lys-TTT
Phage infection protein
UDP-N-acetylglucosamine 1-carboxyvinyltransferase (EC 2.5.1.7)
Pullulanase (EC 3.2.1.41)
2-amino-4-hydroxy-6-hydroxymethyldihydropteridine pyrophosphokinase (EC 2.7.6.3)
Trans-2,cis-3-Decenoyl-ACP isomerase
FIG01118442: hypothetical protein
Phosphoglucosamine mutase (EC 5.4.2.10)
Chaperone protein DnaK
Thiol:disulfide oxidoreductase associated with MetSO reductase
Na+ driven multidrug efflux pump
Maltose/maltodextrin ABC transporter, permease protein MalF
Streptococcal NAD glycohydrolase inhibitor
hypothetical protein
Phosphoribosylamine--glycine ligase (EC 6.3.4.13)
FIG01114622: hypothetical protein
Translation elongation factor Ts
LSU ribosomal protein L19p

5S RNA
Heteropolysaccharide repeat unit export protein
Immunoglobulin G-endopeptidase (IdeS) / Mac/ Secreted immunoglobulin binding protein (Sib38)
hypothetical phage protein
Mobile element protein
3'-to-5' exoribonuclease RNase R
Regulation of D-alanyl-lipoteichoic acid biosynthesis, DltR
Aminopeptidase C (EC 3.4.22.40)
Phage CI-like repressor
Ribosome small subunit-stimulated GTPase EngC
FIG022125: Phage-associated protein
Phage lysin
Alpha-L-Rha alpha-1,2-L-rhamnosyltransferase/alpha-L-Rha alpha-1,3-L- rhamnosyltransferase (EC 2.4.1.-)
FIG01114576: hypothetical protein
LSU m5C1962 methyltransferase RlmI
GTP cyclohydrolase I (EC 3.5.4.16) type 1
Ribosomal small subunit pseudouridine synthase A (EC 4.2.1.70)
Intramembrane protease RasP/YluC, implicated in cell division based on FtsL cleavage
Phage integrase
FIG01116026: hypothetical protein
Nicotinamidase (EC 3.5.1.19)
tRNA-specific 2-thiouridylase MnmA
hypothetical protein
Deoxyribose-phosphate aldolase (EC 4.1.2.4)
hypothetical protein
DNA replication protein dnaD
Glycine betaine ABC transport system, permease protein OpuAB / Glycine betaine ABC transport system, glycine betaine-binding protein OpuAC
Phosphoglucomutase (EC 5.4.2.2)
FIG01114948: hypothetical protein
Peptidyl-tRNA hydrolase (EC 3.1.1.29)
Cadmium resistance protein
2-dehydro-3-deoxygluconate kinase (EC 2.7.1.45)
RopB; Rgg-like transcription regulator
Thioredoxin reductase (EC 1.8.1.9)
Hypothetical radical SAM family enzyme in heat shock gene cluster, similarity with CPO of BS HemN-type

FIG01119121: hypothetical protein
Mobile element protein
Phage integrase: site-specific recombinase
3'->5' exoribonuclease Bsu YhaM
Transcriptional regulatory protein
Peptide methionine sulfoxide reductase MsrA (EC 1.8.4.11) / Peptide methionine sulfoxide reductase MsrB (EC 1.8.4.12)
Glycosyltransferase LafB, responsible for the formation of Gal-Glc-DAG
5-Enolpyruvylshikimate-3-phosphate synthase (EC 2.5.1.19)
putative arylalkylamine n-acetyltransferase
hypothetical phage protein
Mobile element protein
5S RNA
FIG01114600: hypothetical protein
bacteriocin immunity protein
L-proline glycine betaine binding ABC transporter protein ProX (TC 3.A.1.12.1) / Osmotic adaptation
Mevalonate kinase (EC 2.7.1.36)
DNA polymerase III alpha subunit (EC 2.7.7.7)
Mobile element protein
Transcription antitermination protein NusG
Exodeoxyribonuclease III (EC 3.1.11.2)
tRNA-Val-TAC
Two-component response regulator yesN, associated with MetSO reductase
putative transport accessory protein
Phage major tail protein
peptidase, U32 family large subunit [C1]
Cysteine ABC transporter, substrate-binding protein
Additional lipoprotein component of predicted cobalamin ECF transporter
Ribonuclease J2 (endoribonuclease in RNA processing)
Multidrug resistance protein B
FIG018217: Phage-associated protein
DNA polymerase IV (EC 2.7.7.7)
Calcium-transporting ATPase (EC 3.6.3.8)
tRNA-Asp-GTC
HIT family protein
FIG01114323: hypothetical protein

Thymidylate kinase (EC 2.7.4.9)
FIG01113965: hypothetical protein
DNA recombination protein RmuC
ATP synthase A chain (EC 3.6.3.14)
rRNA small subunit methyltransferase H
Phage minor capsid protein #Fam0017
Mobile element protein
Undecaprenyl diphosphate synthase (EC 2.5.1.31)
FIG01117945: hypothetical protein
Integral membrane protein
hypothetical protein
ATP-dependent Clp protease ATP-binding subunit ClpA
Putative deoxyribonuclease YcfH
General stress protein
Arginine pathway regulatory protein ArgR, repressor of arg regulon
dTDP-4-dehydrorhamnose 3,5-epimerase (EC 5.1.3.13)
4-diphosphocytidyl-2-C-methyl-D-erythritol kinase (EC 2.7.1.148)
UDP-N-acetylglucosamine--N-acetylmuramyl-(pentapeptide) pyrophosphoryl-undecaprenol N-acetylglucosamine transferase (EC 2.4.1.227)
Hypothetical protein SpyM3_1200
Signal recognition particle associated protein
DNA replication intiation control protein YabA
GTP-sensing transcriptional pleiotropic repressor codY
FIG01115581: hypothetical protein
DNA primase, phage associated
Shikimate kinase I (EC 2.7.1.71)
Competence-stimulating peptide ABC transporter ATP-binding protein ComA
Phage lysin, N-acetylmuramoyl-L-alanine amidase (EC 3.5.1.28)
Heparinase II/III-like
Cysteine and methionine metabolism regulator CmbR, LysR family
Leucyl-tRNA synthetase (EC 6.1.1.4)
FIG01114077: hypothetical protein
Streptococcal pyrogenic exotoxin C (SpeC); _Toximoron (Superantigen)
tRNA (cytidine(34)-2'-O)-methyltransferase (EC 2.1.1.207)
hypothetical protein
Chloride channel protein

tRNA-Thr-GGT
UDP-N-acetylmuramoylalanine--D-glutamate ligase (EC 6.3.2.9)
hypothetical phage protein
FIG01115119: hypothetical protein
Group B streptococcal surface immunogenic protein
Helicase loader DnaB
MORN motif family protein
Hypothetical protein SPy1011
Aspartate aminotransferase (EC 2.6.1.1)
PTS system, cellobiose-specific IIB component (EC 2.7.1.69)
Zinc ABC transporter, inner membrane permease protein ZnuB
KH domain RNA binding protein YlqC
Phosphoribosylaminoimidazole carboxylase ATPase subunit (EC 4.1.1.21)
ATP-dependent Clp protease, ATP-binding subunit ClpE
Nicotine adenine dinucleotide glycohydrolase (NADGH) (EC 3.2.2.5)
FIG01115781: hypothetical protein
Flavodoxin
FIG01114549: hypothetical protein
DNA-entry nuclease (Competence-specific nuclease) (EC 3.1.30.-)
hypothetical protein
Carbamate kinase (EC 2.7.2.2)
FIG01114743: hypothetical protein
Inosine-5'-monophosphate dehydrogenase (EC 1.1.1.205)
Kup system potassium uptake protein
NPQTN specific sortase B
NAD synthetase (EC 6.3.1.5)
tRNA-Gln-TTG
FIG01114717: hypothetical protein
FIG01115319: hypothetical protein
FIG01115625: hypothetical protein
Phosphoribosylglycinamide formyltransferase (EC 2.1.2.2)
FIG01116145: hypothetical protein
FIG01114499: hypothetical protein
Competence-specific sigma factor ComX
Tryptophanyl-tRNA synthetase (EC 6.1.1.2)

Arginine/ornithine antiporter ArcD
Trk system potassium uptake protein TrkA
Phage protein
hypothetical protein
Glutamate transport ATP-binding protein
SSU ribosomal protein S1p
hypothetical protein
COG4478, integral membrane protein
Hypothetical protein SPy2060
Hydrolase (HAD superfamily), YqeK
Transaldolase (EC 2.2.1.2)
Translation elongation factor P
Phage minor capsid protein
LacI family regulatory protein
Transcriptional regulator of fatty acid biosynthesis FabT
MutT/nudix family protein
Uracil phosphoribosyltransferase (EC 2.4.2.9) / Pyrimidine operon regulatory protein PyrR
hypothetical protein - phage associated
FIG01114177: hypothetical protein
Hydrolase (HAD superfamily)
DNA polymerase III alpha subunit (EC 2.7.7.7)
putaive immunity repressor protein - phage associated
Putative two-component responsible histidine kinase
Copper-translocating P-type ATPase (EC 3.6.3.4)
hypothetical protein
Multimodular transpeptidase-transglycosylase (EC 2.4.1.129) (EC 3.4.-.-)
ATP synthase delta chain (EC 3.6.3.14)
Phage protein
Diacylglycerol kinase (EC 2.7.1.107)
Low molecular weight protein tyrosine phosphatase (EC 3.1.3.48)
ABC transporter permease protein
Transcriptional regulator, MerR family
tRNA-guanine transglycosylase (EC 2.4.2.29)
FIG013576: hypothetical protein
DNA modification methylase

Inner membrane protein translocase component YidC, short form OxaI-like
periplasmic component of efflux system
LSU ribosomal protein L32p
FIG009439: Cytosolic protein containing multiple CBS domains
FIG01115461: hypothetical protein
Mobile element protein
Mobile element protein
FIG01116340: hypothetical protein
Phosphate transport system permease protein PstC (TC 3.A.1.7.1)
2-dehydropantoate 2-reductase (EC 1.1.1.169)
Acyl-ACP thioesterase
D-tyrosyl-tRNA(Tyr) deacylase
3-dehydroquinate dehydratase I (EC 4.2.1.10)
hypothetical
UTP--glucose-1-phosphate uridylyltransferase (EC 2.7.7.9)
D-alanine--D-alanine ligase A (EC 6.3.2.4)
5S RNA

specific IIA component (EC 2.7.1.69)

PTS system, maltose and glucose-specific IIA component (EC 2.7.1.69)

ranylgeranyl diphosphate synthase (EC 2.5.1.29)

beta-glucoside-specific IIA component (EC 2.7.1.69)

ıbunit (EC 6.3.5.2)

cific IIC component (EC 2.7.1.69)

energy

fic IIA component (EC 2.7.1.69)

otransferase subunit (EC 6.3.5.3)

) # TrpAb@PabAb

| geneID | symbol | function. |
| --- | --- | --- |
| AP53_1 | dnaA | Chromosomal replication initiator protein DnaA |
| AP53_2 | dnaN | DNA polymerase III beta subunit (EC 2.7.7.7) |
| AP53_3 | - | FIG01114317: hypothetical protein |
| AP53_4 | - | hypothetical protein |
| AP53_5 | ychF | GTP-binding and nucleic acid-binding protein YchF |
| AP53_6 | pth | Peptidyl-tRNA hydrolase (EC 3.1.1.29) |
| AP53_7 | trcF | Transcription-repair coupling factor |
| AP53_8 | - | Ribosome-associated heat shock protein implicated in the recycling of the 50S subunit (S4 paralog) |
| AP53_9 | divIC | Cell division protein DivIC (FtsB), stabilizes FtsL against RasP cleavage |
| AP53_10 | - | FIG01114659: hypothetical protein |
| AP53_11 | - | FIG01114299: hypothetical protein |
| AP53_12 | tilS | tRNA(Ile)-lysidine synthetase |
| AP53_13 | hpt | Hypoxanthine-guanine phosphoribosyltransferase (EC 2.4.2.8) |
| AP53_14 | ftsH | Cell division protein FtsH (EC 3.4.24.-) |
| AP53_15 | - | Cationic amino acid transporter - APC Superfamily |
| AP53_rna1 | - | Small Subunit Ribosomal RNA; ssuRNA; SSU rRNA |
| AP53_rna2 | - | tRNA-Ala-TGC |
| AP53_rna3 | - | Large Subunit Ribosomal RNA; lsuRNA; LSU rRNA |
| AP53_rna4 | - | 5S RNA |
| AP53_rna5 | - | tRNA-Val-TAC |
| AP53_rna6 | - | tRNA-Asp-GTC |
| AP53_rna7 | - | tRNA-Lys-TTT |
| AP53_rna8 | - | tRNA-Leu-TAG |
| AP53_rna9 | - | tRNA-Thr-TGT |
| AP53_rna10 | - | tRNA-Gly-GCC |
| AP53_rna11 | - | tRNA-Leu-TAA |
| AP53_rna12 | - | tRNA-Arg-ACG |
| AP53_rna13 | - | tRNA-Pro-TGG |
| AP53_rna14 | - | Small Subunit Ribosomal RNA; ssuRNA; SSU rRNA |
| AP53_rna15 | - | tRNA-Ala-TGC |
| AP53_rna16 | - | Large Subunit Ribosomal RNA; lsuRNA; LSU rRNA |
| AP53_rna17 | - | 5S RNA |
| AP53_rna18 | - | tRNA-Val-TAC |
| AP53_rna19 | - | tRNA-Asp-GTC |

| | | |
|---|---|---|
| AP53_rna20 | - | tRNA-Lys-TTT |
| AP53_rna21 | - | tRNA-Leu-TAG |
| AP53_rna22 | - | tRNA-Thr-TGT |
| AP53_rna23 | - | tRNA-Gly-GCC |
| AP53_rna24 | - | tRNA-Leu-TAA |
| AP53_rna25 | - | tRNA-Arg-ACG |
| AP53_rna26 | - | tRNA-Pro-TGG |
| AP53_rna27 | - | tRNA-Met-CAT |
| AP53_rna28 | - | tRNA-Met-CAT |
| AP53_rna29 | - | tRNA-Pseudo-TGA |
| AP53_rna30 | - | tRNA-Met-CAT |
| AP53_rna31 | - | tRNA-Phe-GAA |
| AP53_rna32 | - | tRNA-Gly-TCC |
| AP53_rna33 | - | tRNA-Ile-GAT |
| AP53_rna34 | - | tRNA-Pseudo-GCT |
| AP53_16 | - | Secreted antigen GbpB/SagA/PcsB, putative peptidoglycan hydrolase |
| AP53_17 | prsA2 | Ribose-phosphate pyrophosphokinase (EC 2.7.6.1) |
| AP53_18 | recO | DNA recombination and repair protein RecO |
| AP53_19 | plsX | Phosphate:acyl-ACP acyltransferase PlsX |
| AP53_20 | acpP2 | Acyl carrier protein |
| AP53_21 | purC | Phosphoribosylaminoimidazole-succinocarboxamide synthase (EC 6.3.2.6) |
| AP53_22 | purL | Phosphoribosylformylglycinamidine synthase, synthetase subunit (EC 6.3.5.3) / Phosphoribosylformylglycir |
| AP53_23 | purF | Amidophosphoribosyltransferase (EC 2.4.2.14) |
| AP53_24 | purM | Phosphoribosylformylglycinamidine cyclo-ligase (EC 6.3.3.1) |
| AP53_25 | purN | Phosphoribosylglycinamide formyltransferase (EC 2.1.2.2) |
| AP53_26 | purH | IMP cyclohydrolase (EC 3.5.4.10) / Phosphoribosylaminoimidazolecarboxamide formyltransferase (EC 2.1.2. |
| AP53_27 | - | putative choline binding protein |
| AP53_28 | | Phosphoribosylamine--glycine ligase (EC 6.3.4.13) |
| AP53_29 | | Phosphoribosylaminoimidazole carboxylase catalytic subunit (EC 4.1.1.21) |
| AP53_30 | | Phosphoribosylaminoimidazole carboxylase ATPase subunit (EC 4.1.1.21) |
| AP53_31 | - | FIG01114657: hypothetical protein |
| AP53_32 | - | hypothetical protein |
| AP53_33 | purB | Adenylosuccinate lyase (EC 4.3.2.2) |
| AP53_34 | | Transcriptional regulator |
| AP53_35 | ruvB | Holliday junction DNA helicase RuvB |

| | | |
|---|---|---|
| AP53_36 | - | Low molecular weight protein tyrosine phosphatase (EC 3.1.3.48) |
| AP53_37 | - | MORN motif family protein |
| AP53_38 | - | Acyltransferase family |
| AP53_39 | adhE | Alcohol dehydrogenase (EC 1.1.1.1); Acetaldehyde dehydrogenase (EC 1.2.1.10) |
| AP53_40 | adhP | Alcohol dehydrogenase (EC 1.1.1.1) |
| AP53_41 | - | Multi antimicrobial extrusion (MATE) family transporter |
| AP53_42 | rpsJ | SSU ribosomal protein S10p (S20e) |
| AP53_43 | rplC | LSU ribosomal protein L3p (L3e) |
| AP53_44 | rplD | LSU ribosomal protein L4p (L1e) |
| AP53_45 | rplW | LSU ribosomal protein L23p (L23Ae) |
| AP53_46 | rplB | LSU ribosomal protein L2p (L8e) |
| AP53_47 | rpsS | SSU ribosomal protein S19p (S15e) |
| AP53_48 | rplV | LSU ribosomal protein L22p (L17e) |
| AP53_49 | rpsC | SSU ribosomal protein S3p (S3e) |
| AP53_50 | rplP | LSU ribosomal protein L16p (L10e) |
| AP53_51 | rpmC | LSU ribosomal protein L29p (L35e) |
| AP53_52 | rpsQ | SSU ribosomal protein S17p (S11e) |
| AP53_53 | rplN | LSU ribosomal protein L14p (L23e) |
| AP53_54 | rplX | LSU ribosomal protein L24p (L26e) |
| AP53_55 | rplE | LSU ribosomal protein L5p (L11e) |
| AP53_56 | rpsN | SSU ribosomal protein S14p (S29e) @ SSU ribosomal protein S14p (S29e), zinc-dependent |
| AP53_57 | rpsH | SSU ribosomal protein S8p (S15Ae) |
| AP53_58 | rplF | LSU ribosomal protein L6p (L9e) |
| AP53_59 | rplR | LSU ribosomal protein L18p (L5e) |
| AP53_60 | rpsE | SSU ribosomal protein S5p (S2e) |
| AP53_61 | rpmD | LSU ribosomal protein L30p (L7e) |
| AP53_62 | rplO | LSU ribosomal protein L15p (L27Ae) |
| AP53_63 | secY | Preprotein translocase secY subunit (TC 3.A.5.1.1) |
| AP53_64 | adk | Adenylate kinase (EC 2.7.4.3) |
| AP53_65 | infA | Translation initiation factor 1 |
| AP53_66 | rpmJ | LSU ribosomal protein L36p |
| AP53_67 | rpsM | SSU ribosomal protein S13p (S18e) |
| AP53_68 | rpsK | SSU ribosomal protein S11p (S14e) |
| AP53_69 | rpoA | DNA-directed RNA polymerase alpha subunit (EC 2.7.7.6) |
| AP53_70 | rplQ | LSU ribosomal protein L17p |

| | | |
|---|---|---|
| AP53_rna35 | - | Small Subunit Ribosomal RNA; ssuRNA; SSU rRNA |
| AP53_rna36 | - | tRNA-Ala-TGC |
| AP53_rna37 | - | Large Subunit Ribosomal RNA; lsuRNA; LSU rRNA |
| AP53_rna38 | - | 5S RNA |
| AP53_rna39 | - | tRNA-Val-TAC |
| AP53_rna40 | - | tRNA-Gly-TCC |
| AP53_rna41 | - | tRNA-Ile-GAT |
| AP53_rna42 | - | tRNA-Glu-TTC |
| AP53_rna43 | - | tRNA-Pseudo-TGA |
| AP53_rna44 | - | tRNA-Met-CAT |
| AP53_rna45 | - | tRNA-Phe-GAA |
| AP53_rna46 | - | tRNA-Tyr-GTA |
| AP53_rna47 | - | tRNA-Trp-CCA |
| AP53_rna48 | - | tRNA-His-GTG |
| AP53_rna49 | - | tRNA-Gln-TTG |
| AP53_rna50 | - | tRNA-Leu-CAA |
| AP53_71 | - | hypothetical protein |
| AP53_pseudo73 | -/pseudogene | 4-diphosphocytidyl-2-C-methyl-D-erythritol kinase (EC 2.7.1.148) |
| AP53_74 | adrR | Transcriptional repressor AdcR for Zn(2+)-responsive expression |
| AP53_75 | adrC | Zinc ABC transporter, ATP-binding protein ZnuC |
| AP53_76 | adcB | Zinc ABC transporter, inner membrane permease protein ZnuB |
| AP53_77 | - | HIT family protein |
| AP53_78 | tyrS | Tyrosyl-tRNA synthetase (EC 6.1.1.1) |
| AP53_79 | | Multimodular transpeptidase-transglycosylase (EC 2.4.1.129) (EC 3.4.-.-) |
| AP53_80 | | DNA-directed RNA polymerase beta subunit (EC 2.7.7.6) |
| AP53_81 | | DNA-directed RNA polymerase beta' subunit (EC 2.7.7.6) |
| AP53_82 | - | DNA binding protein, FIG046916 |
| AP53_83 | comGA | Late competence protein ComGA, access of DNA to ComEA |
| AP53_84 | comGB | Late competence protein ComGB, access of DNA to ComEA |
| AP53_85 | comGC | Late competence protein ComGC, access of DNA to ComEA, FIG007487 |
| AP53_86 | comGD | Late competence protein ComGD, access of DNA to ComEA, FIG038316 |
| AP53_87 | comGE | Late competence protein ComGE, FIG075573 |
| AP53_88 | comGF | Late competence protein ComGF, access of DNA to ComEA, FIG012620 |
| AP53_89 | comGF | Late competence protein ComGG, FIG068335 |
| AP53_90 | yujA | Adenine-specific methyltransferase (EC 2.1.1.72) |

| | | |
|---|---|---|
| AP53_91 | ackA | Acetate kinase (EC 2.7.2.1) |
| AP53_92 | - | Hypothetical protein SPy0110 |
| AP53_93 | proC | Pyrroline-5-carboxylate reductase (EC 1.5.1.2) |
| AP53_94 | pepA | Glutamyl aminopeptidase (EC 3.4.11.7) |
| AP53_95 | - | FIG014356: hypothetical protein |
| AP53_96 | trxA | FIG009688: Thioredoxin |
| AP53_97 | - | Phenylalanyl-tRNA synthetase domain protein (Bsu YtpR) |
| AP53_98 | - | FIG107367: tRNA-binding protein |
| AP53_99 | - | Deoxyadenosine kinase (EC 2.7.1.76) / Deoxyguanosine kinase (EC 2.7.1.113) |
| AP53_100 | - | tRNA dihydrouridine synthase B (EC 1.-.-.-) |
| AP53_101 | hslO | Chaperonin (heat shock protein 33) |
| AP53_102 | rofA | Transcriptional regulator, RofA-like Protein (RALP) |
| AP53_103 | cpa1 | putative collagen binding protein |
| AP53_104 | - | Signal peptidase I (EC 3.4.21.89) |
| AP53_105 | fctA | Fibronectin-binding protein/major pilin backbone protein T-antigen |
| AP53_106 | srtB | NPQTN specific sortase B |
| AP53_107 | | FIG01115182: hypothetical protein |
| AP53_108 | - | Multiple sugar metabolism regulator |
| AP53_109 | prtF2/fbaB | Fibronectin-binding protein |
| AP53_110 | | FIG01114239: hypothetical protein |
| AP53_111 | atoE | Short chain fatty acids transporter |
| AP53_112 | - | Transcriptional regulators, LysR family |
| AP53_113 | - | 3-ketoacyl-CoA thiolase (EC 2.3.1.16) @ Acetyl-CoA acetyltransferase (EC 2.3.1.9) |
| AP53_114 | atoD2 | Acetyl-CoA:acetoacetyl-CoA transferase, alpha subunit (EC 2.8.3.8) |
| AP53_115 | atoA1 | Butyrate-acetoacetate CoA-transferase subunit B (EC 2.8.3.9) |
| AP53_116 | - | YheO-like PAS domain |
| AP53_117 | - | Endoribonuclease L-PSP |
| AP53_118 | | Transcriptional regulator pfoR |
| AP53_119 | - | V-type ATP synthase subunit G (EC 3.6.3.14) |
| AP53_120 | ntpI | V-type ATP synthase subunit I (EC 3.6.3.14) |
| AP53_121 | ntpK | V-type ATP synthase subunit K (EC 3.6.3.14) |
| AP53_122 | ntpE | V-type ATP synthase subunit E (EC 3.6.3.14) |
| AP53_123 | ntpC | V-type ATP synthase subunit C (EC 3.6.3.14) |
| AP53_124 | ntpF | V-type ATP synthase subunit F (EC 3.6.3.14) |
| AP53_125 | - | hypothetical protein |

| | | |
|---|---|---|
| AP53_126 | ntpA | V-type ATP synthase subunit A (EC 3.6.3.14) |
| AP53_127 | ntpB | V-type ATP synthase subunit B (EC 3.6.3.14) |
| AP53_128 | ntpD | V-type ATP synthase subunit D (EC 3.6.3.14) |
| AP53_129 | telX | putative toxic anion resistance protein |
| AP53_130 | - | FIG013576: hypothetical protein |
| AP53_131 | purA | Adenylosuccinate synthetase (EC 6.3.4.4) |
| AP53_132 | - | Nucleoside-binding protein |
| AP53_133 | nusG | Transcription antitermination protein NusG |
| AP53_134 | nga | Nicotine adenine dinucleotide glycohydrolase (NADGH) (EC 3.2.2.5) |
| AP53_135 | ifs | Streptococcal NAD glycohydrolase inhibitor |
| AP53_136 | slo | Thiol-activated cytolysin |
| AP53_137 | hypo3 | FIG01114743: hypothetical protein |
| AP53_138 | hypo2 | FIG01114903: hypothetical protein |
| AP53_139 | - | Cystathionine beta-lyase (EC 4.4.1.8) |
| AP53_140 | leuS | Leucyl-tRNA synthetase (EC 6.1.1.4) |
| AP53_141 | | Ascorbate-specific PTS system, EIIC component |
| AP53_142 | | Ascorbate-specific PTS system, EIIB component (EC 2.7.1.69) |
| AP53_143 | | Ascorbate-specific PTS system, EIIA component (EC 2.7.1.-) |
| AP53_144 | | 3-keto-L-gulonate 6-phosphate decarboxylase |
| AP53_145 | | L-xylulose 5-phosphate 3-epimerase (EC 5.1.3.-) |
| AP53_146 | araD | L-ribulose-5-phosphate 4-epimerase (EC 5.1.3.4) |
| AP53_147 | - | Transcriptional antiterminator with PTS regulation domain, SPy0181 ortholog |
| AP53_148 | ulaG | Probable L-ascorbate-6-phosphate lactonase UlaG (EC 3.1.1.-) (L-ascorbate utilization protein G) |
| AP53_149 | opuA | Glycine betaine ABC transport system, ATP-binding protein OpuAA (EC 3.6.3.32) |
| AP53_150 | opuBC | Glycine betaine ABC transport system, permease protein OpuAB / Glycine betaine ABC transport system, glyc |
| AP53_151 | polA | DNA polymerase I (EC 2.7.7.7) |
| AP53_152 | - | Succinyl-CoA synthetase, alpha subunit-related enzymes |
| AP53_153 | perR | Peroxide stress regulator PerR, FUR family |
| AP53_154 | vlg | M protein trans-acting positive regulator (Mga) |
| AP53_155 | - | Co-activator of prophage gene expression IbrA |
| AP53_156 | - | Co-activator of prophage gene expression IbrB |
| AP53_157 | - | FIG01115469: hypothetical protein |
| AP53_158 | - | Mobile element protein |
| AP53_159 | - | Mobile element protein |
| AP53_160 | - | Mobile element protein |

| | | |
|---|---|---|
| AP53_161 | - | Malonate permease |
| AP53_162 | nadC | Quinolinate phosphoribosyltransferase [decarboxylating] (EC 2.4.2.19) |
| AP53_163 | - | Mobile element protein |
| AP53_164 | - | Mobile element protein |
| AP53_165 | - | Mobile element protein |
| AP53_166 | - | membrane protein |
| AP53_167 | - | FIG01116967: hypothetical protein |
| AP53_168 | tgt | tRNA-guanine transglycosylase (EC 2.4.2.29) |
| AP53_169 | hot13 | Zinc finger protein |
| AP53_170 | bioY | Substrate-specific component BioY of biotin ECF transporter |
| AP53_171 | - | Metal-dependent hydrolase (EC 3.-.-.-) |
| AP53_172 | tadA | tRNA-specific adenosine-34 deaminase (EC 3.5.4.-) |
| AP53_173 | - | S-layer homology domain / putative murein endopeptidase |
| AP53_174 | speG | Streptococcal pyrogenic exotoxin G (SpeG) |
| AP53_175 | - | hypothetical protein |
| AP53_176 | - | hypothetical cytosolic protein |
| AP53_177 | pgi | Glucose-6-phosphate isomerase (EC 5.3.1.9) |
| AP53_178 | - | Transcriptional regulator, RofA-like Protein (RALP) |
| AP53_179 | - | FIG01118080: hypothetical protein |
| AP53_180 | - | hypothetical protein |
| AP53_181 | - | Integral membrane protein (Rhomboid family) |
| AP53_182 | hasC2 | UTP--glucose-1-phosphate uridylyltransferase (EC 2.7.7.9) |
| AP53_183 | gpsA | Glycerol-3-phosphate dehydrogenase [NAD(P)+] (EC 1.1.1.94) |
| AP53_184 | | Transcriptional regulator, MarR family |
| AP53_185 | - | Lipid A export ATP-binding/permease protein MsbA |
| AP53_186 | - | Lipid A export ATP-binding/permease protein MsbA |
| AP53_187 | - | FIG053235: Diacylglucosamine hydrolase like |
| AP53_188 | dut | Deoxyuridine 5'-triphosphate nucleotidohydrolase (EC 3.6.1.23) |
| AP53_189 | radA | DNA repair protein RadA |
| AP53_190 | - | Carbonic anhydrase (EC 4.2.1.1) |
| AP53_191 | - | DUF124 domain-containing protein |
| AP53_192 | gltX | Glutamyl-tRNA synthetase (EC 6.1.1.17) @ Glutamyl-tRNA(Gln) synthetase (EC 6.1.1.24) |
| AP53_193 | | Histidine kinase of the competence regulon ComD |
| AP53_194 | | Sensory transduction protein kinase (EC 2.7.3.-) |
| AP53_195 | | Response regulator of the competence regulon ComE |

| | | |
|---|---|---|
| AP53_1867 | fasX | small RNA fasX |
| AP53_196 | rnpA | Ribonuclease P protein component (EC 3.1.26.5) |
| AP53_197 | - | Inner membrane protein translocase component YidC, short form OxaI-like |
| AP53_198 | - | RNA-binding protein Jag |
| AP53_199 | - | FIG01115631: hypothetical protein |
| AP53_200 | rpmH | LSU ribosomal protein L34p |
| AP53_201 | nanE | N-acetylmannosamine-6-phosphate 2-epimerase (EC 5.1.3.9) |
| AP53_202 | - | ABC transporter, predicted N-acetylneuraminate-binding protein |
| AP53_203 | - | ABC transporter, predicted N-acetylneuraminate transport system permease protein 1 |
| AP53_204 | - | ABC transporter, predicted N-acetylneuraminate transport system permease protein 2 |
| AP53_205 | - | FIG01114161: hypothetical protein |
| AP53_206 | nanH | N-acetylneuraminate lyase (EC 4.1.3.3) |
| AP53_207 | - | N-acetylmannosamine kinase (EC 2.7.1.60) |
| AP53_208 | - | Sialic acid utilization regulator, RpiR family |
| AP53_209 | tatD | Putative deoxyribonuclease YcfH |
| AP53_210 | - | Ribonuclease M5 (EC 3.1.26.8) |
| AP53_211 | ksgA | SSU rRNA (adenine(1518)-N(6)/adenine(1519)-N(6))-dimethyltransferase (EC 2.1.1.182) |
| AP53_212 | engC | Ribosome small subunit-stimulated GTPase EngC |
| AP53_213 | rpe | Ribulose-phosphate 3-epimerase (EC 5.1.3.1) |
| AP53_214 | thiN | Thiamin pyrophosphokinase (EC 2.7.6.2) |
| AP53_215 | - | DNA recombination protein RmuC |
| AP53_216 | cbf | 3'->5' exoribonuclease Bsu YhaM |
| AP53_217 | | PurR: transcription regulator associated with purine metabolism |
| AP53_218 | | Surface exclusion protein |
| AP53_219 | rpsL | SSU ribosomal protein S12p (S23e) |
| AP53_220 | rpsG | SSU ribosomal protein S7p (S5e) |
| AP53_221 | fusA | Translation elongation factor G |
| AP53_222 | plr | NAD-dependent glyceraldehyde-3-phosphate dehydrogenase (EC 1.2.1.12) / Plasmin(ogen) receptor |
| AP53_223 | - | FIG01114598: hypothetical protein |
| AP53_224 | - | Amino acid transport ATP-binding protein |
| AP53_225 | - | Amino acid ABC transporter, glutamine-binding protein/permease protein |
| AP53_226 | - | FIG01114729: hypothetical protein |
| AP53_227 | uppP | Undecaprenyl-diphosphatase (EC 3.6.1.27) |
| AP53_228 | mecA | Negative regulator of genetic competence MecA |
| AP53_229 | rgpG | Undecaprenyl-phosphate N-acetylglucosaminyl 1-phosphate transferase (EC 2.7.8.-) |

| | | |
|---|---|---|
| AP53_230 | - | hypothetical cytosolic protein |
| AP53_231 | sufC | Iron-sulfur cluster assembly ATPase protein SufC |
| AP53_232 | sufD | Iron-sulfur cluster assembly protein SufD |
| AP53_233 | sufS | Cysteine desulfurase (EC 2.8.1.7), SufS subfamily |
| AP53_234 | sufE2 | Putative iron-sulfur cluster assembly scaffold protein for SUF system, SufE2 |
| AP53_235 | sufB | Iron-sulfur cluster assembly protein SufB |
| AP53_236 | | D-alanyl-D-alanine carboxypeptidase (EC 3.4.16.4) |
| AP53_237 | | D-alanyl-D-alanine carboxypeptidase (EC 3.4.16.4) |
| AP53_238 | oppA | Oligopeptide ABC transporter, periplasmic oligopeptide-binding protein OppA (TC 3.A.1.5.1) |
| AP53_239 | oppB | Oligopeptide transport system permease protein OppB (TC 3.A.1.5.1) |
| AP53_240 | oppC | Oligopeptide transport system permease protein OppC (TC 3.A.1.5.1) |
| AP53_241 | oppD | Oligopeptide transport ATP-binding protein OppD (TC 3.A.1.5.1) |
| AP53_242 | oppF | Oligopeptide transport ATP-binding protein OppF (TC 3.A.1.5.1) |
| AP53_243 | - | Transposase |
| AP53_rna51 | - | Small Subunit Ribosomal RNA; ssuRNA; SSU rRNA |
| AP53_rna52 | - | tRNA-Ala-TGC |
| AP53_rna53 | - | Large Subunit Ribosomal RNA; lsuRNA; LSU rRNA |
| AP53_rna54 | - | 5S RNA |
| AP53_rna55 | - | tRNA-Asn-GTT |
| AP53_rna56 | - | tRNA-Arg-ACG |
| AP53_244 | comX2 | Competence-specific sigma factor ComX |
| AP53_245 | - | Mobile element protein |
| AP53_246 | - | FIG001553: Hydrolase, HAD subfamily IIIA |
| AP53_247 | - | GTP-binding protein YqeH, required for biogenesis of 30S ribosome subunit |
| AP53_248 | - | FIG004454: RNA binding protein |
| AP53_249 | nadD | Nicotinate-nucleotide adenylyltransferase (EC 2.7.7.18) |
| AP53_250 | - | Hydrolase (HAD superfamily), YqeK |
| AP53_251 | - | Iojap protein |
| AP53_252 | - | FIG145533: Methyltransferase (EC 2.1.1.-) |
| AP53_253 | - | FIG007079: UPF0348 protein family |
| AP53_254 | - | FIG007491: hypothetical protein YeeN |
| AP53_255 | - | Cysteine ABC transporter, substrate-binding protein |
| AP53_256 | - | Methionine ABC transporter substrate-binding protein |
| AP53_257 | - | Methionine ABC transporter ATP-binding protein |
| AP53_258 | - | Methionine ABC transporter permease protein |

| | | |
|---|---|---|
| AP53_259 | braB | Branched-chain amino acid transport system carrier protein |
| AP53_260 | - | Dicarboxylate/amino acid:cation (Na+ or H+) symporter |
| AP53_261 | trkA | Trk system potassium uptake protein TrkA |
| AP53_262 | trkH | Potassium uptake protein TrkH |
| AP53_263 | gidB | rRNA small subunit 7-methylguanosine (m7G) methyltransferase GidB |
| AP53_264 | lemA | LemA protein |
| AP53_265 | htpX | Heat shock protein HtpX (EC 3.4.24.-) |
| AP53_266 | - | COG1399 protein in cluster with ribosomal protein L32p, Firmicutes subfamily |
| AP53_267 | covR | Response regulator CsrR |
| AP53_pseudo268 | csrS/pseudogene | Transmembrane histidine kinase CsrS / deletion of T at 1407 causing a early stop codon at 469th aa) |
| AP53_269 | nrdR | Ribonucleotide reductase transcriptional regulator NrdR |
| AP53_270 | dnaB | Helicase loader DnaB |
| AP53_271 | dnaI | Helicase loader DnaI |
| AP53_272 | | GTP-binding protein EngA |
| AP53_273 | snf | SWF/SNF family helicase |
| AP53_274 | - | FIG01114242: hypothetical protein |
| AP53_275 | murC | UDP-N-acetylmuramate--alanine ligase (EC 6.3.2.8) |
| AP53_276 | - | putative arylalkylamine n-acetyltransferase |
| AP53_277 | - | FIG004453: protein YceG like |
| AP53_278 | greA | Transcription elongation factor GreA |
| AP53_279 | - | Inner membrane protein translocase component YidC, OxaA protein |
| AP53_280 | acyP | Predicted nucleoside phosphatase |
| AP53_281 | - | FIG011178: rRNA methylase |
| AP53_282 | - | FIG005986: HD family hydrolase |
| AP53_283 | - | FIG005935: membrane protein |
| AP53_284 | - | FIG027054: hypothetical protein |
| AP53_285 | glr | Glutamate racemase (EC 5.1.1.3) |
| AP53_286 | rdgB | unknown domain / Nucleoside 5-triphosphatase RdgB (dHAPTP, dITP, XTP-specific) (EC 3.6.1.15) |
| AP53_287 | - | FIG009886: phosphoesterase |
| AP53_288 | - | FIG042801: CBS domain containing protein |
| AP53_289 | xerD | Tyrosine recombinase XerD |
| AP53_290 | scpA | Segregation and condensation protein A |
| AP53_291 | scpB | Segregation and condensation protein B |
| AP53_292 | rluB | Ribosomal large subunit pseudouridine synthase B (EC 4.2.1.70) |
| AP53_293 | - | Protein YidD |

| | | |
|---|---|---|
| AP53_294 | - | tRNA (cytidine(34)-2'-O)-methyltransferase (EC 2.1.1.207) |
| AP53_295 | - | Substrate-specific component RibU of riboflavin ECF transporter |
| AP53_296 | - | Membrane-associated phospholipid phosphatase |
| AP53_297 | - | COG1242: Predicted Fe-S oxidoreductase |
| AP53_298 | - | SAM-dependent methyltransferase, MraW methylase family (EC 2.1.1.-) |
| AP53_299 | hlyX | Hemolysins and related proteins containing CBS domains |
| AP53_300 | pflC | Pyruvate formate-lyase activating enzyme (EC 1.97.1.4) |
| AP53_301 | ppaC | Manganese-dependent inorganic pyrophosphatase (EC 3.6.1.1) |
| AP53_302 | - | FIG01114725: hypothetical protein |
| AP53_303 | | ABC-type Fe3+-siderophore transport system, permease 2 component |
| AP53_304 | | ABC-type Fe3+-siderophore transport system, permease component |
| AP53_305 | | Ferrichrome-binding periplasmic protein precursor (TC 3.A.1.14.3) |
| AP53_306 | | Ferrichrome transport ATP-binding protein FhuC (TC 3.A.1.14.3) |
| AP53_307 | murE | UDP-N-acetylmuramoylalanyl-D-glutamate--L-lysine ligase (EC 6.3.2.7) |
| AP53_308 | - | Membrane protein involved in the export of O-antigen, teichoic acid lipoteichoic acids |
| AP53_309 | upp | Uracil phosphoribosyltransferase (EC 2.4.2.9) |
| AP53_310 | - | hypothetical protein |
| AP53_311 | clpP | ATP-dependent Clp protease proteolytic subunit (EC 3.4.21.92) |
| AP53_312 | - | hypothetical protein |
| AP53_313 | tmk | Thymidylate kinase (EC 2.7.4.9) |
| AP53_314 | dnaX | DNA polymerase III delta prime subunit (EC 2.7.7.7) |
| AP53_315 | - | hypothetical protein |
| AP53_316 | - | hypothetical |
| AP53_317 | - | DNA replication intiation control protein YabA |
| AP53_318 | - | rRNA small subunit methyltransferase I |
| AP53_319 | - | FIG01114628: hypothetical protein |
| AP53_320 | - | Cytoplasmic copper homeostasis protein CutC |
| AP53_321 | - | Arsenate reductase (EC 1.20.4.1) |
| AP53_322 | exoA | FIG01115198: hypothetical protein |
| AP53_323 | - | Exodeoxyribonuclease III (EC 3.1.11.2) |
| AP53_324 | lctO | Lactate 2-monooxygenase (EC 1.13.12.4) |
| AP53_325 | | hypothetical protein |
| AP53_326 | prtS | Serine endopeptidase ScpC (EC 3.4.21.-) |
| AP53_327 | - | FIG01118451: hypothetical protein |
| AP53_328 | - | FIG01115281: hypothetical protein |

| | | |
|---|---|---|
| AP53_329 | - | Membrane protein, putative |
| AP53_330 | metG | Methionyl-tRNA synthetase (EC 6.1.1.10) |
| AP53_331 | - | hypothetical protein |
| AP53_332 | - | FIG01115797: hypothetical protein |
| AP53_333 | nrdF | Ribonucleotide reductase of class Ib (aerobic), beta subunit (EC 1.17.4.1) |
| AP53_334 | nrdI | Ribonucleotide reduction protein NrdI |
| AP53_335 | nrdE1 | Ribonucleotide reductase of class Ib (aerobic), alpha subunit (EC 1.17.4.1) |
| AP53_336 | spyA | C3 family ADP-ribosyltransferase (EC 2.4.2.-) |
| AP53_337 | - | Mobile element protein |
| AP53_338 | - | FIG01114323: hypothetical protein |
| AP53_339 | - | Mobile element protein |
| AP53_340 | - | hypothetical protein |
| AP53_341 | - | hypothetical protein |
| AP53_342 | - | FIG01114229: hypothetical protein |
| AP53_343 | - | FIG01115464: hypothetical protein |
| AP53_344 | - | FIG01114304: hypothetical protein |
| AP53_345 | | 3-oxoacyl-[acyl-carrier protein] reductase (EC 1.1.1.100) |
| AP53_346 | | NAD-dependent oxidoreductase |
| AP53_347 | glpT | Glycerol-3-phosphate transporter |
| AP53_348 | glmU | N-acetylglucosamine-1-phosphate uridyltransferase (EC 2.7.7.23) / Glucosamine-1-phosphate N-acetyltransf |
| AP53_349 | MutT | ADP-ribose pyrophosphatase (EC 3.6.1.13) |
| AP53_350 | hypo20 | FIG01115635: hypothetical protein |
| AP53_351 | pfs | 5'-methylthioadenosine nucleosidase (EC 3.2.2.16) / S-adenosylhomocysteine nucleosidase (EC 3.2.2.9) |
| AP53_352 | - | FIG01116268: hypothetical protein |
| AP53_353 | scaR | Mn-dependent transcriptional regulator MntR |
| AP53_354 | mtsA | Manganese ABC transporter, periplasmic-binding protein SitA |
| AP53_355 | mtsB | Manganese ABC transporter, ATP-binding protein SitB |
| AP53_356 | mtsC | Manganese ABC transporter, inner membrane permease protein SitD |
| AP53_357 | ppi | Peptidyl-prolyl cis-trans isomerase (EC 5.2.1.8) |
| AP53_358 | ftsK | Cell division protein FtsK |
| AP53_359 | - | Integral membrane protein |
| AP53_360 | rplK | LSU ribosomal protein L11p (L12e) |
| AP53_361 | rplA | LSU ribosomal protein L1p (L10Ae) |
| AP53_362 | pyrH | Uridylate kinase (EC 2.7.4.-) |
| AP53_363 | frr | Ribosome recycling factor |

| | | |
|---|---|---|
| AP53_364 | - | S1 RNA binding domain |
| AP53_365 | | Peptide methionine sulfoxide reductase MsrA (EC 1.8.4.11) |
| AP53_366 | - | FIG01114318: hypothetical protein |
| AP53_367 | - | Group B streptococcal surface immunogenic protein |
| AP53_368 | - | 67kDa Myosin-crossreactive streptococcal antigen |
| AP53_369 | phoH | Phosphate starvation-inducible protein PhoH, predicted ATPase |
| AP53_370 | - | Hypothetical protein VC0266 (sugar utilization related?) |
| AP53_371 | ybeY | Metal-dependent hydrolase YbeY, involved in rRNA and/or ribosome maturation and assembly |
| AP53_372 | dgk | Diacylglycerol kinase (EC 2.7.1.107) |
| AP53_373 | era | GTP-binding protein Era |
| AP53_374 | - | MutT/nudix family protein |
| AP53_375 | - | FIG01115781: hypothetical protein |
| AP53_376 | - | Competence-stimulating peptide ABC transporter ATP-binding protein ComA |
| AP53_377 | - | Peptide ABC transporter ATP binding/permease protein |
| AP53_378 | - | Peptide ABC transporter ATP binding/permease protein |
| AP53_379 | - | Bacteriocin BlpU |
| AP53_380 | - | hypothetical protein |
| AP53_381 | - | hypothetical protein |
| AP53_382 | - | FIG01119121: hypothetical protein |
| AP53_383 | - | putative role in replication |
| AP53_384 | - | Mobile element protein |
| AP53_385 | - | Mobile element protein |
| AP53_386 | - | Mobile element protein |
| AP53_387 | - | hypothetical protein |
| AP53_pseudo388 | -/pseudogene | Mobile element protein |
| AP53_390 | - | Response regulator of the competence regulon ComE |
| AP53_391 | - | Histidine kinase of the competence regulon ComD |
| AP53_392 | - | FIG01116220: hypothetical protein |
| AP53_393 | - | Competence-stimulating peptide ABC transporter permease protein ComB |
| AP53_394 | | Competence-stimulating peptide ABC transporter ATP-binding protein ComA |
| AP53_395 | blpM | Bacteriocin-like peptide M BlpM |
| AP53_396 | - | FIG01116077: hypothetical protein |
| AP53_397 | silD | hypothetical protein |
| AP53_398 | - | unknown |
| AP53_pseudo399 | - | FIG01114499: hypothetical protein |

| | | |
|---|---|---|
| AP53_400 | - | FIG01114499: hypothetical protein |
| AP53_401 | - | FIG01114077: hypothetical protein |
| AP53_402 | - | hypothetical protein |
| AP53_403 | - | hypothetical protein |
| AP53_404 | - | hypothetical protein |
| AP53_405 | | Positive transcriptional regulator, MutR family |
| AP53_406 | fpg | Formamidopyrimidine-DNA glycosylase (EC 3.2.2.23) |
| AP53_407 | coaE | Dephospho-CoA kinase (EC 2.7.1.24) |
| AP53_408 | ATPase | ATPase |
| AP53_409 | pmrA | Multidrug resistance efflux pump PmrA |
| AP53_410 | secG | Preprotein translocase subunit SecG (TC 3.A.5.1.1) |
| AP53_411 | vacB | 3'-to-5' exoribonuclease RNase R |
| AP53_412 | smpB | tmRNA-binding protein SmpB |
| AP53_413 | - | Glutaminyl-peptide cyclotransferase (EC 2.3.2.5) |
| AP53_414 | pcp | Pyrrolidone-carboxylate peptidase (EC 3.4.19.3) |
| AP53_415 | - | FIG001614: Membrane protein |
| AP53_416 | - | hypothetical protein |
| AP53_417 | - | Bactoprenol glucosyl transferase (EC 2.4.1.-) |
| AP53_418 | gloA | Lactoylglutathione lyase (EC 4.4.1.5) |
| AP53_419 | yfkO | Oxygen-insensitive NAD(P)H nitroreductase (EC 1.-.-.-) / Dihydropteridine reductase (EC 1.5.1.34) |
| AP53_420 | pepQ | Proline dipeptidase (EC 3.4.13.9) |
| AP53_421 | ccpA | Catabolite control protein A |
| AP53_422 | - | Glycosyltransferase LafB, responsible for the formation of Gal-Glc-DAG |
| AP53_423 | - | Glycosyltransferase LafA, responsible for the formation of Glc-DAG |
| AP53_424 | - | hypothetical protein |
| AP53_425 | thrS | Threonyl-tRNA synthetase (EC 6.1.1.3) |
| AP53_426 | tagH | Daunorubicin resistance ATP-binding protein drrA |
| AP53_427 | - | Daunorubicin resistance transmembrane protein |
| AP53_428 | - | ABC-type multidrug transport system, permease component |
| AP53_429 | - | Hypothetical protein |
| AP53_430 | - | 3-ketoacyl-CoA thiolase (EC 2.3.1.16) @ Acetyl-CoA acetyltransferase (EC 2.3.1.9) |
| AP53_431 | - | Long-chain-fatty-acid--CoA ligase (EC 6.2.1.3) |
| AP53_432 | - | FIG01114264: hypothetical protein |
| AP53_433 | | Two-component response regulator SA14-24 |
| AP53_434 | | Two-component sensor kinase SA14-24 |

| | | |
|---|---|---|
| AP53_435 | | Zn-dependent hydrolase YycJ/WalJ, required for cell wall metabolism and coordination of cell division with |
| AP53_436 | rnc | Ribonuclease III (EC 3.1.26.3) |
| AP53_437 | smc | Chromosome partition protein smc |
| AP53_438 | mutR-Rgg3 | Positive transcriptional regulator, MutR family |
| AP53_439 | aroE | Shikimate 5-dehydrogenase I alpha (EC 1.1.1.25) |
| AP53_440 | - | FIG01114171: hypothetical protein |
| AP53_441 | - | putative L-glutamate ligase |
| AP53_442 | - | FIG01115625: hypothetical protein |
| AP53_443 | | Archaeal S-adenosylmethionine synthetase (EC 2.5.1.6) |
| AP53_444 | - | FIG01116163: hypothetical protein |
| AP53_445 | - | Dolichyl-phosphate mannoosyltransferase, involved in cell wall biogenesis |
| AP53_446 | - | UDP-glucose dehydrogenase (EC 1.1.1.22) |
| AP53_447 | mefE | ABC transporter membrane-spanning permease - macrolide efflux |
| AP53_448 | - | FIG01115329: hypothetical protein |
| AP53_449 | - | hypothetical protein |
| AP53_450 | - | FIG01116026: hypothetical protein |
| AP53_451 | - | FIG01114600: hypothetical protein |
| AP53_452 | - | FIG01116748: hypothetical protein |
| AP53_453 | - | FIG01115104: hypothetical protein |
| AP53_454 | - | Plasmid stabilization system antitoxin protein |
| AP53_455 | - | Plasmid stabilization system toxin protein |
| AP53_456 | - | hypothetical protein |
| AP53_457 | - | FIG01117988: hypothetical protein |
| AP53_458 | tnpA | Mobile element protein |
| AP53_459 | - | Mobile element protein |
| AP53_460 | - | Hydrolase (HAD superfamily) |
| AP53_461 | - | Hydrolase (HAD superfamily) |
| AP53_462 | ftsY | Signal recognition particle receptor protein FtsY (=alpha subunit) (TC 3.A.5.1.1) |
| AP53_463 | - | Multidrug resistance protein B |
| AP53_464 | bglG | Beta-glucoside bgl operon antiterminator, BglG family |
| AP53_465 | - | PTS system, beta-glucoside-specific IIB component (EC 2.7.1.69) / PTS system, beta-glucoside-specific IIC com |
| AP53_466 | bglA | 6-phospho-beta-glucosidase (EC 3.2.1.86) |
| AP53_467 | - | ABC transporter, substrate-binding protein |
| AP53_468 | - | possible permease |
| AP53_469 | - | FIG01116444: hypothetical protein |

| | | |
|---|---|---|
| AP53_470 | - | Transcription accessory protein (S1 RNA-binding domain) |
| AP53_471 | - | Putative metallopeptidase (Zinc) SprT family |
| AP53_472 | - | Phage shock protein C, putative; stress-responsive transcriptional regulator |
| AP53_473 | | HPr kinase/phosphorylase (EC 2.7.1.-) (EC 2.7.4.-) |
| AP53_474 | lgt | Prolipoprotein diacylglyceryl transferase (EC 2.4.99.-) |
| AP53_475 | - | General stress protein |
| AP53_476 | - | FIG01115301: hypothetical protein |
| AP53_477 | - | FIG01114801: hypothetical protein |
| AP53_478 | - | PROTEASE (EC 3.4.-.-) |
| AP53_479 | - | peptidase, U32 family large subunit [C1] |
| AP53_480 | - | FIG01114033: hypothetical protein |
| AP53_481 | - | bacterial seryl-tRNA synthetase related |
| AP53_482 | - | FIG01116574: hypothetical protein |
| AP53_483 | lysS | Lysyl-tRNA synthetase (class II) (EC 6.1.1.6) |
| AP53_484 | - | 2-haloalkanoic acid dehalogenase (EC 3.8.1.2) |
| AP53_485 | pgmA | Phosphoglycerate mutase family 5 |
| AP53_486 | ybaK | Transcriptional regulator |
| AP53_487 | - | Substrate-specific component ThiT of thiamin ECF transporter |
| AP53_488 | - | Phage lysin, glycosyl hydrolase, family 25 |
| AP53_489 | - | Substrate-specific component CbrT of predicted cobalamin ECF transporter |
| AP53_490 | - | Additional lipoprotein component of predicted cobalamin ECF transporter |
| AP53_491 | bsaA | Glutathione peroxidase family protein |
| AP53_492 | pepF | Oligoendopeptidase F |
| AP53_493 | ppc | Phosphoenolpyruvate carboxylase (EC 4.1.1.31) |
| AP53_494 | ftsW | Cell division protein FtsW |
| AP53_495 | - | FIG01114557: hypothetical protein |
| AP53_496 | tuf | Translation elongation factor Tu |
| AP53_497 | tpiA | Triosephosphate isomerase (EC 5.3.1.1) |
| AP53_498 | - | hypothetical protein |
| AP53_499 | murN | tRNA-dependent lipid II-Ala--L-alanine ligase @ tRNA-dependent lipid II-Ser--L-alanine ligase |
| AP53_500 | femB | tRNA-dependent lipid II--L-alanine ligase @ tRNA-dependent lipid II--L-serine ligase |
| AP53_501 | yidA | Promiscuous sugar phosphatase YidA, haloacid dehalogenase-like phosphatase family |
| AP53_502 | - | FIG01114293: hypothetical protein |
| AP53_503 | - | Deoxyguanosinetriphosphate triphosphohydrolase (EC 3.1.5.1) |
| AP53_504 | - | FIG01114378: hypothetical protein |

| | | |
|---|---|---|
| AP53_505 | pacL | Calcium-transporting ATPase (EC 3.6.3.8) |
| AP53_506 | regR | Transcriptional regulator RegR, rpressor of hyaluronate and KDG utilization |
| AP53_507 | - | Heparinase II/III-like |
| AP53_508 | agaD | PTS system, N-acetylgalactosamine-specific IID component (EC 2.7.1.69) |
| AP53_509 | agaW | PTS system, N-acetylgalactosamine-specific IIC component (EC 2.7.1.69) |
| AP53_510 | agaV | PTS system, N-acetylgalactosamine-specific IIB component (EC 2.7.1.69) |
| AP53_511 | ugl | Unsaturated glucuronyl hydrolase (EC 3.2.1.-) |
| AP53_512 | | PTS system, N-acetylgalactosamine- and galactosamine-specific IIA component (EC 2.7.1.69) |
| AP53_513 | idnO | 2-deoxy-D-gluconate 3-dehydrogenase (EC 1.1.1.125) |
| AP53_514 | - | predicted 4-deoxy-L-threo-5-hexosulose-uronate ketol-isomerase (EC 5.3.1.17) |
| AP53_515 | kdgK | 2-dehydro-3-deoxygluconate kinase (EC 2.7.1.45) |
| AP53_516 | kgdA | 4-hydroxy-2-oxoglutarate aldolase (EC 4.1.3.16) @ 2-dehydro-3-deoxyphosphogluconate aldolase (EC 4.1.2. |
| AP53_517 | - | Beta-phosphoglucomutase (EC 5.4.2.6) (EC 2.7.1.41) |
| AP53_518 | QueG | Epoxyqueuosine (oQ) reductase QueG |
| AP53_519 | prfB | Peptide chain release factor 2; programmed frameshift-containing |
| AP53_520 | ftsE | Cell division transporter, ATP-binding protein FtsE (TC 3.A.5.1.1) |
| AP53_521 | ftsX | Cell division protein FtsX |
| AP53_522 | - | Hydroxyacylglutathione hydrolase (EC 3.1.2.6) |
| AP53_523 | budC | 2,3-butanediol dehydrogenase, S-alcohol forming, (R)-acetoin-specific (EC 1.1.1.4) / Acetoin (diacetyl) reduc |
| AP53_524 | - | Mobile element protein |
| AP53_525 | dinG | DinG family ATP-dependent helicase YoaA |
| AP53_526 | | Aspartate aminotransferase (EC 2.6.1.1) |
| AP53_527 | | Asparaginyl-tRNA synthetase (EC 6.1.1.22) |
| AP53_528 | - | FIG000506: Predicted P-loop-containing kinase |
| AP53_529 | - | FIG002813: LPPG:FO 2-phospho-L-lactate transferase like, CofD-like |
| AP53_530 | - | FIG001886: Cytoplasmic hypothetical protein |
| AP53_531 | pepD | FIG01117156: hypothetical protein |
| AP53_532 | adcA | Candidate zinc-binding lipoprotein ZinT |
| AP53_533 | | Transcriptional regulator, GntR family |
| AP53_534 | | Galactosamine-6-phosphate isomerase (EC 5.3.1.-) |
| AP53_535 | rpmE2 | LSU ribosomal protein L31p @ LSU ribosomal protein L31p, zinc-independent |
| AP53_536 | - | FIG146085: 3'-to-5' oligoribonuclease A, Bacillus type |
| AP53_537 | flaV | Flavodoxin |
| AP53_538 | - | Chorismate mutase I (EC 5.4.99.5) |
| AP53_539 | - | Chloride channel protein |

| | | |
|---|---|---|
| AP53_540 | rplS | LSU ribosomal protein L19p |
| AP53_rna57 | - | tRNA-Arg-CCT |
| AP53_541 | - | hydrolase, haloacid dehalogenase-like family |
| AP53_542 | - | DNA gyrase subunit B (EC 5.99.1.3) |
| AP53_543 | - | Septation ring formation regulator EzrA |
| AP53_544 | - | FIG01114091: hypothetical protein |
| AP53_545 | eno | Enolase (EC 4.2.1.11) |
| AP53_1865 | sagA | Streptolysin S biosynthesis protein B (SagA) |
| AP53_546 | sagB | Streptolysin S biosynthesis protein B (SagB) |
| AP53_547 | sagC | Streptolysin S biosynthesis protein C (SagC) |
| AP53_548 | sagD | Streptolysin S biosynthesis protein D (SagD) |
| AP53_549 | sagE | Streptolysin S self-immunity protein (SagE) |
| AP53_550 | sagF | Streptolysin S biosynthesis protein (SagF) |
| AP53_551 | sagG | Export ABC transporter ATP-binding protein; Streptolysin S export protein (SagG) |
| AP53_552 | sagH | Streptolysin S export transmembrane permease (SagH) |
| AP53_553 | sagI | Streptolysin S export transmembrane permease (SagI) |
| AP53_554 | - | Asparagine-rich ORFan next to Sag operon |
| AP53_555 | - | Endonuclease/Exonuclease/phosphatase family protein |
| AP53_556 | - | Substrate-specific component QueT (COG4708) of predicted queuosine-regulated ECF transporter |
| AP53_557 | ligA | DNA ligase (EC 6.5.1.2) |
| AP53_558 | dagK | Transcription regulator [contains diacylglycerol kinase catalytic domain] |
| AP53_559 | atpE | ATP synthase C chain (EC 3.6.3.14) |
| AP53_560 | atpB | ATP synthase A chain (EC 3.6.3.14) |
| AP53_561 | atpF | ATP synthase B chain (EC 3.6.3.14) |
| AP53_562 | atpH | ATP synthase delta chain (EC 3.6.3.14) |
| AP53_563 | atpA | ATP synthase alpha chain (EC 3.6.3.14) |
| AP53_564 | atpG | ATP synthase gamma chain (EC 3.6.3.14) |
| AP53_565 | atpD | ATP synthase beta chain (EC 3.6.3.14) |
| AP53_566 | atpC | ATP synthase epsilon chain (EC 3.6.3.14) |
| AP53_567 | - | FIG01114865: hypothetical protein |
| AP53_568 | murA1 | UDP-N-acetylglucosamine 1-carboxyvinyltransferase (EC 2.5.1.7) |
| AP53_569 | epuA | Competence-associated EpuA protein |
| AP53_570 | - | DNA-entry nuclease (Competence-specific nuclease) (EC 3.1.30.-) |
| AP53_571 | endA | hypothetical protein |
| AP53_572 | endA | DNA-entry nuclease (Competence-specific nuclease) (EC 3.1.30.-) |

| | | |
|---|---|---|
| AP53_573 | | Phenylalanyl-tRNA synthetase alpha chain (EC 6.1.1.20) |
| AP53_574 | | Phenylalanyl-tRNA synthetase beta chain (EC 6.1.1.20) |
| AP53_575 | - | Salt-stress induced protein |
| AP53_576 | - | FIG01115289: hypothetical protein |
| AP53_577 | - | ABC transporter permease protein |
| AP53_578 | - | ABC transporter ATP-binding protein |
| AP53_579 | ypfJ | YpfJ protein, zinc metalloprotease superfamily |
| AP53_580 | addB | ATP-dependent nuclease, subunit B |
| AP53_581 | addA | ATP-dependent nuclease, subunit A |
| AP53_582 | - | Amino acid ABC transporter, amino acid-binding protein |
| AP53_583 | rpsU | SSU ribosomal protein S21p |
| AP53_584 | mscL | Large-conductance mechanosensitive channel |
| AP53_585 | dnaG | DNA primase (EC 2.7.7.-) |
| AP53_586 | rpoD | RNA polymerase sigma factor RpoD |
| AP53_587 | - | probably aromatic ring hydroxylating enzyme, evidenced by COGnitor; PaaD-like protein (DUF59) involved i |
| AP53_588 | rmlD | dTDP-4-dehydrorhamnose reductase (EC 1.1.1.133) |
| AP53_589 | rgpAc | Alpha-D-GlcNAc alpha-1,2-L-rhamnosyltransferase (EC 2.4.1.-) |
| AP53_590 | rgpBc | Alpha-L-Rha alpha-1,3-L-rhamnosyltransferase (EC 2.4.1.-) |
| AP53_591 | rgpCc | rhamnose-containing polysacharide translocation permease |
| AP53_592 | | ABC-transporter (ATP-binding protein) -possibly involved in cell wall localization and side chain formation o |
| AP53_593 | | putative glycosyltransferase - possibly involved in cell wall localization and side chain formation of rhamnos |
| AP53_594 | | Alpha-L-Rha alpha-1,2-L-rhamnosyltransferase/alpha-L-Rha alpha-1,3-L- rhamnosyltransferase (EC 2.4.1.-) |
| AP53_pseudo595 | ltaS/pseudogene | Lipoteichoic acid synthase LtaS Type Iic/interrupted by large phage |
| AP53_1866 | ptx | paratox? |
| AP53_596 | - | putative repressor C1 |
| AP53_597 | - | hypothetical protein - phage associated |
| AP53_598 | - | conserved hypothetical protein |
| AP53_599 | - | Phage protein |
| AP53_600 | - | Phage protein |
| AP53_601 | - | Phage lysin |
| AP53_pseudo603 | -/pseudogene | Phage holin |
| AP53_604 | - | Phage protein |
| AP53_605 | - | FIG01116522: hypothetical protein |
| AP53_606 | - | Phage hyaluronidase |
| AP53_607 | hylP | Hyaluronate lyase (phage associated) |

| | | |
|---|---|---|
| AP53_608 | - | Phage endopeptidase |
| AP53_609 | - | Phage tail fibers |
| AP53_610 | - | Phage tail length tape-measure protein |
| AP53_611 | - | hypothetical protein |
| AP53_612 | - | Phage base plate |
| AP53_613 | - | Phage major tail protein |
| AP53_614 | - | hypothetical phage protein |
| AP53_615 | - | hypothetical phage protein |
| AP53_616 | - | hypothetical phage protein |
| AP53_617 | - | hypothetical phage protein |
| AP53_618 | - | hypothetical phage protein |
| AP53_619 | - | Phage major capsid protein |
| AP53_620 | - | Prophage Clp protease-like protein |
| AP53_621 | - | Portal protein, phage associated |
| AP53_622 | - | Phage terminase, large subunit |
| AP53_623 | - | Phage protein |
| AP53_624 | - | Phage-associated homing endonuclease |
| AP53_625 | - | hypothetical phage protein |
| AP53_626 | - | hypothetical phage protein |
| AP53_627 | - | hypothetical phage protein |
| AP53_628 | - | Phage integrase: site-specific recombinase |
| AP53_629 | - | putative transcriptional activator - phage associated |
| AP53_630 | - | B-cell receptor associated protein-related protein |
| AP53_631 | - | putaive immunity repressor protein - phage associated |
| AP53_632 | - | hypothetical protein |
| AP53_633 | | Single-stranded DNA-binding protein |
| AP53_634 | - | hypothetical phage protein |
| AP53_635 | - | conserved hypothetical protein - phage associated |
| AP53_636 | - | Phage protein |
| AP53_637 | - | hypothetical phage protein |
| AP53_638 | - | FIG01116946: hypothetical protein |
| AP53_639 | - | conserved hypothetical protein - phage associated |
| AP53_640 | - | hypothetical phage protein |
| AP53_641 | - | hypothetical phage protein |
| AP53_642 | - | Helicase loader DnaI |

| | | |
|---|---|---|
| AP53_643 | - | replication initiator protein A |
| AP53_644 | dnaB | Replicative DNA helicase (DnaB) (EC 3.6.4.12) |
| AP53_645 | - | hypothetical protein |
| AP53_646 | - | hypothetical phage protein |
| AP53_647 | - | hypothetical phage protein |
| AP53_648 | - | Phage antirepressor protein |
| AP53_649 | - | hypothetical phage protein |
| AP53_650 | - | conserved domain protein |
| AP53_651 | - | hypothetical phage protein |
| AP53_652 | - | Phage transcriptional regulator, Cro/CI family |
| AP53_653 | - | Phage protein |
| AP53_654 | - | Pathogenicity island SaPIn2 |
| AP53_655 | - | Phage integrase: site-specific recombinase |
| AP53_pseudo656 | ltaS/pseudogene | Lipoteichoic acid synthase LtaS Type Iic (The first GCT-Ala is overlapped with the last codon of the former hal[illegible] |
| AP53_657 | - | Glycosyl transferase, group 2 family protein |
| AP53_658 | - | Membrane protein |
| AP53_659 | amrA | Heteropolysaccharide repeat unit export protein |
| AP53_660 | - | membrane protein, related to Actinobacillus protein (1944168) |
| AP53_661 | pepT | Tripeptide aminopeptidase (EC 3.4.11.4) |
| AP53_662 | ebsA | Pore forming protein ebsA |
| AP53_663 | fer | Ferredoxin |
| AP53_664 | - | FIG01114555: hypothetical protein |
| AP53_665 | | Cytidylate kinase (EC 2.7.4.14) |
| AP53_666 | infC | Translation initiation factor 3 |
| AP53_667 | rpmI | LSU ribosomal protein L35p |
| AP53_668 | rplT | LSU ribosomal protein L20p |
| AP53_669 | - | Lipoteichoic acid synthase LtaS Type IIc |
| AP53_670 | - | LSU m5C1962 methyltransferase RlmI |
| AP53_671 | | 3-dehydroquinate dehydratase I (EC 4.2.1.10) |
| AP53_672 | | Chorismate synthase (EC 4.2.3.5) |
| AP53_673 | - | FIG01116936: hypothetical protein |
| AP53_674 | gor | Glutathione reductase (EC 1.8.1.7) |
| AP53_675 | - | Dihydrofolate synthase (EC 6.3.2.12) / Folylpolyglutamate synthase (EC 6.3.2.17) |
| AP53_676 | - | Histone acetyltransferase Gcn5 |
| AP53_677 | nifS1 | Cysteine desulfurase (EC 2.8.1.7) |

| | | |
|---|---|---|
| AP53_678 | thiI | tRNA S(4)U 4-thiouridine synthase (former ThiI) |
| AP53_679 | capA | Capsule biosynthesis protein capA |
| AP53_680 | rplU | LSU ribosomal protein L21p |
| AP53_681 | - | FIG139598: Potential ribosomal protein |
| AP53_682 | rpmA | LSU ribosomal protein L27p |
| AP53_683 | - | Cysteine and methionine metabolism regulator CmbR, LysR family |
| AP53_684 | lsp | Lipoprotein signal peptidase (EC 3.4.23.36) |
| AP53_685 | - | Ribosomal large subunit pseudouridine synthase D (EC 4.2.1.70) |
| AP53_686 | pyrR | Uracil phosphoribosyltransferase (EC 2.4.2.9) / Pyrimidine operon regulatory protein PyrR |
| AP53_687 | pyrP | Uracil permease |
| AP53_688 | pyrB | Aspartate carbamoyltransferase (EC 2.1.3.2) |
| AP53_689 | carA | Carbamoyl-phosphate synthase small chain (EC 6.3.5.5) |
| AP53_690 | carB | Carbamoyl-phosphate synthase large chain (EC 6.3.5.5) |
| AP53_691 | arcA | periplasmic component of efflux system |
| AP53_692 | salX | ABC transporter ATP-binding protein |
| AP53_693 | salY | ABC transporter permease protein |
| AP53_694 | glpQ | Glycerophosphoryl diester phosphodiesterase (EC 3.1.4.46) |
| AP53_695 | rpsP | SSU ribosomal protein S16p |
| AP53_696 | - | KH domain RNA binding protein YlqC |
| AP53_697 | - | Cell surface protein |
| AP53_698 | - | FIG01115236: hypothetical protein |
| AP53_699 | czcD | Cobalt-zinc-cadmium resistance protein CzcD |
| AP53_700 | - | Transcriptional regulator, TetR family |
| AP53_701 | rimM | 16S rRNA processing protein RimM |
| AP53_702 | trmD | tRNA (Guanine37-N1) -methyltransferase (EC 2.1.1.31) |
| AP53_703 | trxB | Thioredoxin reductase (EC 1.8.1.9) |
| AP53_704 | - | Membrane protein |
| AP53_705 | apbA | 2-dehydropantoate 2-reductase (EC 1.1.1.169) |
| AP53_706 | | Transcriptional repressor of the fructose operon, DeoR family |
| AP53_707 | | 1-phosphofructokinase (EC 2.7.1.56) |
| AP53_708 | | PTS system, fructose-specific IIA component (EC 2.7.1.69) / PTS system, fructose-specific IIB component (EC 2 |
| AP53_709 | - | Peptidoglycan hydrolase, Autolysin1 (EC 3.5.1.28) |
| AP53_710 | - | Peptidoglycan hydrolase, Autolysin2 (EC 3.5.1.28) |
| AP53_711 | - | Mobile element protein |
| AP53_712 | - | hypothetical protein |

| | | |
|---|---|---|
| AP53_713 | mac;ideS | Immunoglobulin G-endopeptidase (IdeS) / Mac/ Secreted immunoglobulin binding protein (Sib38) |
| AP53_714 | - | Phage protein |
| AP53_715 | - | Nucleoside diphosphate kinase (EC 2.7.4.6) |
| AP53_716 | - | Hypothetical protein SP1558 |
| AP53_717 | - | DegV family protein |
| AP53_718 | | tRNA nucleotidyltransferase (EC 2.7.7.21) (EC 2.7.7.25) |
| AP53_719 | - | COG0488: ATPase components of ABC transporters with duplicated ATPase domains |
| AP53_720 | - | FIG01114610: hypothetical protein |
| AP53_721 | fms | Peptide deformylase (EC 3.5.1.88) |
| AP53_722 | - | 5'-nucleotidase (EC 3.1.3.5) |
| AP53_723 | - | GTP pyrophosphokinase (EC 2.7.6.5) |
| AP53_724 | sptR | Regulation of D-alanyl-lipoteichoic acid biosynthesis, DltR |
| AP53_725 | - | Regulation of D-alanyl-lipoteichoic acid biosynthesis, sensor histidine kinase |
| AP53_726 | | Mevalonate kinase (EC 2.7.1.36) |
| AP53_727 | | Diphosphomevalonate decarboxylase (EC 4.1.1.33) |
| AP53_728 | | Phosphomevalonate kinase (EC 2.7.4.2) |
| AP53_729 | fni | Isopentenyl-diphosphate delta-isomerase, FMN-dependent (EC 5.3.3.2) |
| AP53_730 | mvaA | Hydroxymethylglutaryl-CoA reductase (EC 1.1.1.34) |
| AP53_731 | mvaS | Hydroxymethylglutaryl-CoA synthase (EC 2.3.3.10) |
| AP53_732 | thyA | Thymidylate synthase (EC 2.1.1.45) |
| AP53_733 | dfr | Dihydrofolate reductase (EC 1.5.1.3) |
| AP53_734 | - | FIG01114189: hypothetical protein |
| AP53_735 | clpX | ATP-dependent Clp protease ATP-binding subunit ClpX |
| AP53_736 | engB | GTP-binding protein EngB |
| AP53_737 | vfr | Hypothetical protein SPy0887 |
| AP53_738 | clpL | putative ATP-dependent Clp proteinase (ATP-binding subunit) |
| AP53_739 | rpiA | Ribose 5-phosphate isomerase A (EC 5.3.1.6) |
| AP53_740 | deoB | Phosphopentomutase (EC 5.4.2.7) |
| AP53_741 | arsC | Arsenate reductase (EC 1.20.4.1) |
| AP53_742 | punA | Purine nucleoside phosphorylase (EC 2.4.2.1); N-Ribosylnicotinamide phosphorylase (EC 2.4.2.1) ## possible |
| AP53_743 | deoD | Purine nucleoside phosphorylase (EC 2.4.2.1) |
| AP53_744 | cpsX | Histidine protein kinase |
| AP53_745 | - | hypothetical protein |
| AP53_746 | | Methionine biosynthesis and transport regulator MtaR, LysR family |
| AP53_747 | | FIG01114562: hypothetical protein |

| | | |
|---|---|---|
| AP53_748 | pyrF | Orotidine 5'-phosphate decarboxylase (EC 4.1.1.23) |
| AP53_749 | pyrE | Orotate phosphoribosyltransferase (EC 2.4.2.10) |
| AP53_750 | | 6-aminohexanoate-cyclic-dimer hydrolase (EC 3.5.2.12) |
| AP53_751 | | Cystine ABC transporter, periplasmic cystine-binding protein FliY |
| AP53_752 | | Cystine transport system permease protein |
| AP53_753 | ung | Uracil-DNA glycosylase, family 1 |
| AP53_754 | pyrC | Dihydroorotase (EC 3.5.2.3) |
| AP53_755 | plsY | Acyl-phosphate:glycerol-3-phosphate O-acyltransferase PlsY |
| AP53_756 | parE | Topoisomerase IV subunit B (EC 5.99.1.-) |
| AP53_757 | parC | Topoisomerase IV subunit A (EC 5.99.1.-) |
| AP53_758 | bcaT | Branched-chain amino acid aminotransferase (EC 2.6.1.42) |
| AP53_759 | - | FIG01114490: hypothetical protein |
| AP53_rna58 | - | tRNA-Tyr-GTA |
| AP53_rna59 | - | tRNA-Gln-TTG |
| AP53_760 | rpsA | SSU ribosomal protein S1p |
| AP53_rna60 | - | tRNA-Arg-CCG |
| AP53_761 | - | FIG015049: hypothetical protein |
| AP53_762 | - | Rhodanese domain protein UPF0176, Firmicutes subgroup |
| AP53_763 | - | hypothetical protein |
| AP53_764 | - | Glutathione S-transferase domain protein (class beta) |
| AP53_765 | - | Exfoliative toxin A |
| AP53_766 | - | FIG01115306: hypothetical protein |
| AP53_767 | miaA | tRNA dimethylallyltransferase (EC 2.5.1.75) |
| AP53_768 | hflX | GTP-binding protein HflX |
| AP53_769 | - | Cystathionine beta-lyase (EC 4.4.1.8) |
| AP53_770 | elaC | Ribonuclease Z (EC 3.1.26.11) |
| AP53_771 | - | Short chain dehydrogenase |
| AP53_772 | recJ | Single-stranded-DNA-specific exonuclease RecJ (EC 3.1.-.-) |
| AP53_773 | apt | Adenine phosphoribosyltransferase (EC 2.4.2.7) |
| AP53_774 | dnaD | Chromosome replication initiation protein dnaD |
| AP53_775 | nth | Endonuclease III (EC 4.2.99.18) |
| AP53_776 | - | Putative tRNA-m1A22 methylase |
| AP53_777 | - | FIG137478: Hypothetical protein |
| AP53_778 | - | Glycine/D-amino acid oxidases family |
| AP53_779 | | Glucose-1-phosphate thymidylyltransferase (EC 2.7.7.24) |

| | | |
|---|---|---|
| AP53_780 | cpsFP | dTDP-4-dehydrorhamnose 3,5-epimerase (EC 5.1.3.13) |
| AP53_781 | rmlB | dTDP-glucose 4,6-dehydratase (EC 4.2.1.46) |
| AP53_782 | mutX | Mutator mutT protein (7,8-dihydro-8-oxoguanine-triphosphatase) (EC 3.6.1.-) |
| AP53_783 | - | Hypothetical protein SPy1011 |
| AP53_784 | - | TPR-repeat-containing protein |
| AP53_785 | fbp | Fibronectin/fibrinogen-binding protein |
| AP53_786 | - | hypothetical protein |
| AP53_787 | - | ABC transporter substrate-binding protein |
| AP53_788 | - | ABC transporter permease protein |
| AP53_789 | - | ABC transporter ATP-binding protein |
| AP53_790 | - | Ribonuclease J2 (endoribonuclease in RNA processing) |
| AP53_791 | estA | Tributyrin esterase |
| AP53_792 | - | FIG01115226: hypothetical protein |
| AP53_793 | - | ABC transporter ATP-binding protein |
| AP53_794 | | Acetoin dehydrogenase E1 component alpha-subunit (EC 1.2.4.-) |
| AP53_795 | | Acetoin dehydrogenase E1 component beta-subunit (EC 1.2.4.-) |
| AP53_796 | pdhC | Dihydrolipoamide acetyltransferase component (E2) of acetoin dehydrogenase complex (EC 2.3.1.-) |
| AP53_797 | acoL | Dihydrolipoamide dehydrogenase of acetoin dehydrogenase (EC 1.8.1.4) |
| AP53_798 | hylA | Hyaluronate lyase precursor (EC 4.2.2.1) |
| AP53_799 | - | Lipoate-protein ligase A |
| AP53_800 | cobQ | Putative amidotransferase similar to cobyric acid synthase |
| AP53_801 | - | proposed amino acid ligase found clustered with an amidotransferase |
| AP53_802 | - | Hypothetical protein YbbP, contains nucleotide-binding domain of DisA bacterial checkpoint controller |
| AP53_803 | - | uncharacterized secreted protein, YBBR Bacillus subtilis homolog |
| AP53_804 | | Phosphoglucosamine mutase (EC 5.4.2.10) |
| AP53_805 | - | FIG01115418: hypothetical protein |
| AP53_806 | hemN | Hypothetical radical SAM family enzyme in heat shock gene cluster, similarity with CPO of BS HemN-type |
| AP53_807 | - | Acyl-ACP thioesterase |
| AP53_808 | - | Hypothetical NagD-like phosphatase |
| AP53_809 | - | COG4478, integral membrane protein |
| AP53_810 | lepA | Translation elongation factor LepA |
| AP53_811 | - | Collagen-like surface protein |
| AP53_pseudo812 | - | Nucleoside diphosphate kinase (EC 2.7.4.6) |
| AP53_813 | ndk | FIG01117958: hypothetical protein |
| AP53_814 | - | Mobile element protein |

| | | |
|---|---|---|
| AP53_815 | lepA | Translation elongation factor LepA |
| AP53_816 | - | Collagen-like surface protein |
| AP53_817 | msrB | Peptide methionine sulfoxide reductase MsrB (EC 1.8.4.12) |
| AP53_818 | YeiH | Putative membrane protein YeiH |
| AP53_819 | ptsA | PTS system, IIA component, putative |
| AP53_820 | ptsB | PTS system, mannose/fructose family IIB component |
| AP53_821 | ptsC | PTS system, mannose-specific IIC component (EC 2.7.1.69) |
| AP53_822 | ptsD | PTS system, mannose-specific IID component (EC 2.7.1.69) |
| AP53_823 | - | Two-component sensor kinase, associated with ferric iron transporter, SPy1061 homolog |
| AP53_824 | - | Two-component response regulator, associated with ferric iron transporter, SPy1062 homolog |
| AP53_825 | - | Ferric iron ABC transporter, iron-binding protein |
| AP53_826 | gabD | Succinate-semialdehyde dehydrogenase [NAD] (EC 1.2.1.24); Succinate-semialdehyde dehydrogenase [NADP |
| AP53_827 | uvrC | Excinuclease ABC subunit C |
| AP53_828 | - | Oxygen-insensitive NAD(P)H nitroreductase (EC 1.-.-.-) / Dihydropteridine reductase (EC 1.5.1.34) |
| AP53_829 | pepV | Acetylornithine deacetylase/Succinyl-diaminopimelate desuccinylase and related deacylases |
| AP53_830 | trmE | GTPase and tRNA-U34 5-formylation enzyme TrmE |
| AP53_831 | rplJ | LSU ribosomal protein L10p (P0) |
| AP53_832 | rplL | LSU ribosomal protein L7/L12 (P1/P2) |
| AP53_833 | - | D-alanyl-D-alanine carboxypeptidase (EC 3.4.16.4) |
| AP53_834 | - | Polysaccharide deacetylase |
| AP53_835 | folC1 | Dihydrofolate synthase (EC 6.3.2.12) / Folylpolyglutamate synthase (EC 6.3.2.17) |
| AP53_836 | | GTP cyclohydrolase I (EC 3.5.4.16) type 1 |
| AP53_837 | | Dihydropteroate synthase (EC 2.5.1.15) |
| AP53_838 | | Dihydroneopterin aldolase (EC 4.1.2.25) |
| AP53_839 | | 2-amino-4-hydroxy-6-hydroxymethyldihydropteridine pyrophosphokinase (EC 2.7.6.3) |
| AP53_840 | murB | UDP-N-acetylenolpyruvoylglucosamine reductase (EC 1.1.1.158) |
| AP53_841 | potA | Putrescine transport ATP-binding protein PotA (TC 3.A.1.11.1) |
| AP53_842 | potB | Spermidine Putrescine ABC transporter permease component PotB (TC 3.A.1.11.1) |
| AP53_843 | potC | Spermidine Putrescine ABC transporter permease component potC (TC_3.A.1.11.1) |
| AP53_844 | potD | ABC transporter, periplasmic spermidine putrescine-binding protein PotD (TC 3.A.1.11.1) |
| AP53_845 | dpiA | Two-component response regulator, malate (EC 2.7.3.-) |
| AP53_846 | dpiB | Two-component sensor histidine kinase, malate (EC 2.7.3.-) |
| AP53_847 | | Malate Na(+) symporter |
| AP53_848 | | NAD-dependent malic enzyme (EC 1.1.1.38) |
| AP53_849 | adh | Alcohol dehydrogenase, zinc-containing |

| | | |
|---|---|---|
| AP53_850 | - | Mobile element protein |
| AP53_851 | | Class B acid phosphatase precursor (EC 3.1.3.2) |
| AP53_852 | - | Voltage-gated chloride channel family protein |
| AP53_853 | - | Lipase/Acylhydrolase family protein |
| AP53_854 | - | hypothetical protein |
| AP53_855 | - | Permease |
| AP53_856 | radC | DNA repair protein RadC |
| AP53_857 | - | Glutamine amidotransferase, class I |
| AP53_858 | rex | Redox-sensitive transcriptional regulator (AT-rich DNA-binding protein) |
| AP53_859 | - | FIG01114333: hypothetical protein |
| AP53_860 | | Cysteine desulfurase (EC 2.8.1.7) |
| AP53_861 | prs | Ribose-phosphate pyrophosphokinase (EC 2.7.6.1) |
| AP53_862 | - | Adenylate cyclase (EC 4.6.1.1) |
| AP53_863 | - | GTP pyrophosphokinase (EC 2.7.6.5) |
| AP53_864 | ppnK | NAD kinase (EC 2.7.1.23) |
| AP53_865 | rluD | Similar to ribosomal large subunit pseudouridine synthase D, Bacillus subtilis YjbO type |
| AP53_866 | eutD | Phosphate acetyltransferase (EC 2.3.1.8) |
| AP53_867 | ydfG | Short chain dehydrogenase |
| AP53_868 | - | Na+ driven multidrug efflux pump |
| AP53_869 | proV | L-proline glycine betaine ABC transport system permease protein ProV (TC 3.A.1.12.1) |
| AP53_870 | proX | L-proline glycine betaine binding ABC transporter protein ProX (TC 3.A.1.12.1) / Osmotic adaptation |
| AP53_871 | guaC | GMP reductase (EC 1.7.1.7) |
| AP53_872 | xpt | Xanthine phosphoribosyltransferase (EC 2.4.2.22) |
| AP53_873 | pbuX | Xanthine permease |
| AP53_874 | apbE | Hypothetical similar to thiamin biosynthesis lipoprotein ApbE |
| AP53_875 | - | 4-oxalocrotonate tautomerase (EC 5.3.2.-); Xylose transport system permease protein xylH |
| AP53_876 | tdk2 | Thymidine kinase (EC 2.7.1.21) |
| AP53_877 | | Peptide chain release factor 1 |
| AP53_878 | hemK | Protein-N(5)-glutamine methyltransferase PrmC, methylates polypeptide chain release factors RF1 and RF2 |
| AP53_879 | - | TsaC protein (YrdC domain) required for threonylcarbamoyladenosine t(6)A37 modification in tRNA |
| AP53_880 | pat | Phosphinothricin N-acetyltransferase (EC 2.3.1.-) |
| AP53_881 | glyA | Serine hydroxymethyltransferase (EC 2.1.2.1) |
| AP53_882 | rluA | Pseudouridylate synthases, 23S RNA-specific |
| AP53_883 | pvaA | Pneumococcal vaccine antigen A homolog |
| AP53_884 | - | ABC transporter, ATP-binding/permease protein |

| | | |
|---|---|---|
| AP53_885 | - | ABC transporter, ATP-binding/permease protein |
| AP53_886 | | NADH peroxidase (EC 1.11.1.1) |
| AP53_887 | ldh | L-lactate dehydrogenase (EC 1.1.1.27) |
| AP53_888 | gyrA | DNA gyrase subunit A (EC 5.99.1.3) |
| AP53_889 | srtA | Sortase A, LPXTG specific |
| AP53_890 | - | FIG01280259: hypothetical protein |
| AP53_891 | | FIG01114048: hypothetical protein |
| AP53_892 | | FIG01115412: hypothetical protein |
| AP53_893 | - | Transcription regulator [contains diacylglycerol kinase catalytic domain] |
| AP53_894 | - | Hemolysin III |
| AP53_895 | - | FIG01115071: hypothetical protein |
| AP53_896 | rbgA | 50S ribosomal subunit maturation GTPase RbgA (B. subtilis YlqF) |
| AP53_897 | rnhB | Ribonuclease HII (EC 3.1.26.4) |
| AP53_898 | smf | Rossmann fold nucleotide-binding protein Smf possibly involved in DNA uptake |
| AP53_899 | topA | DNA topoisomerase I (EC 5.99.1.2) |
| AP53_900 | - | Mobile element protein |
| AP53_901 | gid | tRNA:m(5)U-54 MTase gid |
| AP53_902 | oadA2 | Oxaloacetate decarboxylase alpha chain (EC 4.1.1.3) |
| AP53_903 | - | FIG01114041: hypothetical protein |
| AP53_904 | gcdC | Biotin carboxyl carrier protein of oxaloacetate decarboxylase; Biotin carboxyl carrier protein |
| AP53_905 | - | Methylmalonyl-CoA decarboxylase, beta chain (EC 4.1.1.41); Oxaloacetate decarboxylase beta chain (EC 4.1. |
| AP53_906 | citG | 2-(5''-triphosphoribosyl)-3'-dephosphocoenzyme-A synthase (EC 2.7.8.25) |
| AP53_907 | gntR | Transcriptional regulator, GntR family |
| AP53_908 | citN | Mg2+/citrate complex transporter |
| AP53_909 | - | FIG01114846: hypothetical protein |
| AP53_910 | - | Biotin carboxyl carrier protein of oxaloacetate decarboxylase; Biotin carboxyl carrier protein |
| AP53_911 | oadB | Oxaloacetate decarboxylase beta chain (EC 4.1.1.3) |
| AP53_912 | - | FIG01114213: hypothetical protein |
| AP53_913 | | Citrate lyase gamma chain, acyl carrier protein (EC 4.1.3.6) |
| AP53_914 | | Citrate lyase beta chain (EC 4.1.3.6) |
| AP53_915 | | Citrate lyase alpha chain (EC 4.1.3.6) |
| AP53_916 | | Apo-citrate lyase phosphoribosyl-dephospho-CoA transferase (EC 2.7.7.61) |
| AP53_917 | oadA1 | Oxaloacetate decarboxylase alpha chain (EC 4.1.1.3) |
| AP53_918 | citC | [Citrate [pro-3S]-lyase] ligase (EC 6.2.1.22) |
| AP53_919 | - | FIG01114660: hypothetical protein |

| | | |
|---|---|---|
| AP53_920 | xerS | Tyrosine recombinase XerC |
| AP53_921 | - | Pleiotropic regulator of exopolysaccharide synthesis, competence and biofilm formation Ftr, XRE family |
| AP53_922 | ffh | Signal recognition particle, subunit Ffh SRP54 (TC 3.A.5.1.1) |
| AP53_923 | ylxM | Signal recognition particle associated protein |
| AP53_924 | - | Transcriptional regulator in cluster with beta-lactamase, GntR family |
| AP53_925 | - | FIG01114068: hypothetical protein |
| AP53_926 | guaA | GMP synthase [glutamine-hydrolyzing], amidotransferase subunit (EC 6.3.5.2) / GMP synthase [glutamine-hy |
| AP53_927 | - | UDP-N-acetylmuramoylpentapeptide-lysine N(6)-alanyltransferase (EC 2.3.2.10) (EC 2.3.2.-) |
| AP53_pseudo928 | - | UDP-N-acetylmuramoylpentapeptide-lysine N(6)-alanyltransferase (EC 2.3.2.10) (EC 2.3.2.-) |
| AP53_929 | - | ABC transporter ATP-binding protein |
| AP53_930 | pdxK | Substrate-specific component PdxU of predicted pyridoxine ECF transporter |
| AP53_931 | pdxK | Pyridoxal kinase (EC 2.7.1.35) |
| AP53_932 | - | Predicted transcriptional regulator of pyridoxine metabolism |
| AP53_933 | rnhB | Anaerobic ribonucleoside-triphosphate reductase (EC 1.17.4.2) |
| AP53_934 | cls | Cardiolipin synthetase (EC 2.7.8.-) |
| AP53_935 | fhs1 | Formate--tetrahydrofolate ligase (EC 6.3.4.3) |
| AP53_936 | lplA | Lipoate-protein ligase A |
| AP53_937 | sir2 | NAD-dependent protein deacetylase of SIR2 family |
| AP53_938 | - | COG2110, Macro domain, possibly ADP-ribose binding module |
| AP53_939 | gcvH | Biotinyl-lipoyl attachment domain protein, GcvH-like |
| AP53_940 | - | Luciferase-like monooxygenase (EC 1.14.-.-) |
| AP53_941 | - | Probable NADH-dependent flavin oxidoreductase in cluster with COG2110 |
| AP53_942 | - | Protein:protein lipoyl transferase |
| AP53_943 | dpfB | Phosphopantothenoylcysteine synthetase (EC 6.3.2.5) |
| AP53_944 | dfp | Phosphopantothenoylcysteine decarboxylase (EC 4.1.1.36) |
| AP53_945 | - | Substrate-specific component PanT of predicted pantothenate ECF transporter |
| AP53_946 | pgmA | Phosphoglucomutase (EC 5.4.2.2) |
| AP53_947 | - | Nucleoside ABC transporter, permease protein 2 |
| AP53_948 | - | Putative deoxyribose-specific ABC transporter, permease protein |
| AP53_949 | - | Putative deoxyribose-specific ABC transporter, ATP-binding protein |
| AP53_950 | - | Nucleoside-binding protein |
| AP53_951 | | Cytidine deaminase (EC 3.5.4.5) |
| AP53_952 | - | Ribosomal RNA small subunit methyltransferase C (EC 2.1.1.52) ## SSU rRNA m2G1207 |
| AP53_953 | | Pantothenate kinase (EC 2.7.1.33) |
| AP53_954 | rpsT | SSU ribosomal protein S20p |

| | | |
|---|---|---|
| AP53_955 | ciaH | Two component system sensor histidine kinase CiaH (EC 2.7.3.-) |
| AP53_956 | ciaR | Two component system response regulator CiaR |
| AP53_957 | | Lysyl aminopeptidase (EC 3.4.11.15) |
| AP53_958 | phoU | Phosphate transport system regulatory protein PhoU |
| AP53_959 | pstB | Phosphate transport ATP-binding protein PstB (TC 3.A.1.7.1) |
| AP53_960 | pstB2 | Phosphate transport ATP-binding protein PstB (TC 3.A.1.7.1) |
| AP53_961 | pstA | Phosphate transport system permease protein PstA (TC 3.A.1.7.1) |
| AP53_962 | pstC | Phosphate transport system permease protein PstC (TC 3.A.1.7.1) |
| AP53_963 | pstS | Phosphate ABC transporter, periplasmic phosphate-binding protein PstS (TC 3.A.1.7.1) |
| AP53_964 | - | tRNA and rRNA cytosine-C5-methylases |
| AP53_965 | - | Inositol-1-monophosphatase (EC 3.1.3.25) |
| AP53_966 | - | FIG006438: hypothetical protein |
| AP53_967 | spxA | Transcriptional regulator SpxA1 |
| AP53_968 | ribF | Riboflavin kinase (EC 2.7.1.26) / FMN adenylyltransferase (EC 2.7.7.2) |
| AP53_969 | truB | tRNA pseudouridine synthase B (EC 4.2.1.70) |
| AP53_970 | - | FIG01115860: hypothetical protein |
| AP53_971 | - | membrane protein |
| AP53_972 | hsdS2 | Type I restriction-modification system specificity subunit |
| AP53_973 | - | ABC transporter permease protein |
| AP53_974 | - | ABC transporter ATP-binding protein YvcR |
| AP53_975 | - | Transcriptional regulator, TetR family |
| AP53_976 | - | Transcriptional regulator, GntR family |
| AP53_977 | - | General stress protein, Gls24 family |
| AP53_978 | | FIG01114697: hypothetical protein |
| AP53_979 | - | General stress protein, Gls24 family |
| AP53_980 | | FIG01114232: hypothetical protein |
| AP53_981 | hypo-high2 | FIG01114374: hypothetical protein |
| AP53_982 | hypo-high1 | FIG01114177: hypothetical protein |
| AP53_983 | pcrA | ATP-dependent DNA helicase UvrD/PcrA |
| AP53_984 | GlyP | Sodium/glycine symporter GlyP |
| AP53_985 | | Cobalt-zinc-cadmium resistance protein |
| AP53_986 | cfa | CAMP factor |
| AP53_987 | hisJ | Amino acid ABC transporter, amino acid-binding protein |
| AP53_988 | glnQ | ABC-type polar amino acid transport system, ATPase component |
| AP53_989 | hisM | ABC transporter membrane-spanning permease - glutamine transport |

| | | |
|---|---|---|
| AP53_990 | PhnA | Alkylphosphonate utilization operon protein PhnA |
| AP53_991 | glmS | Glucosamine--fructose-6-phosphate aminotransferase [isomerizing] (EC 2.6.1.16) |
| AP53_992 | - | Signal peptidase I (EC 3.4.21.89) |
| AP53_993 | pyk | Pyruvate kinase (EC 2.7.1.40) |
| AP53_994 | pfk | 6-phosphofructokinase (EC 2.7.1.11) |
| AP53_995 | dnaE | DNA polymerase III alpha subunit (EC 2.7.7.7) |
| AP53_996 | - | Transcriptional regulator, GntR family |
| AP53_997 | - | ABC-type multidrug transport system, ATPase component |
| AP53_998 | - | ABC transporter, ATP-binding protein |
| AP53_999 | - | Immunoreactive protein Se23.5 (Fragment) |
| AP53_1000 | prx | Paratox |
| AP53_1001 | - | Phage protein |
| AP53_1002 | slaA | Streptococcal phospholipase A2; _Toximoron (Other) |
| AP53_1003 | speK | Streptococcal pyrogenic exotoxin K (SpeK) |
| AP53_1004 | - | Phage lysin |
| AP53_1005 | - | Phage protein |
| AP53_1006 | - | Phage lysin |
| AP53_1007 | - | Phage holin |
| AP53_1008 | - | Phage protein |
| AP53_1009 | - | FIG01116522: hypothetical protein |
| AP53_1010 | - | Phage hyaluronidase |
| AP53_1011 | hylP | Hyaluronate lyase (phage associated) |
| AP53_1012 | - | Phage hyaluronidase; Phage endopeptidase |
| AP53_1013 | - | Phage-related protein |
| AP53_1014 | - | Phage tail length tape-measure protein |
| AP53_1015 | - | Phage protein |
| AP53_1016 | - | Phage protein |
| AP53_1017 | - | Phage major tail protein |
| AP53_1018 | - | Phage protein |
| AP53_1019 | - | Phage protein |
| AP53_1020 | - | Phage protein |
| AP53_1021 | - | DNA packaging protein, phage associated |
| AP53_1022 | - | conserved hypothetical protein - phage associated |
| AP53_1023 | - | Phage capsid protein |
| AP53_1024 | - | Prophage Clp protease-like protein |

| | | |
|---|---|---|
| AP53_1025 | - | Phage portal protein |
| AP53_1026 | - | Phage protein |
| AP53_1027 | - | hypothetical phage protein |
| AP53_1028 | - | Phage terminase, large subunit |
| AP53_1029 | - | Phage terminase, small subunit |
| AP53_1030 | - | Phage endonuclease |
| AP53_1031 | - | FIG01114733: hypothetical protein |
| AP53_1032 | - | hypothetical protein - phage associated |
| AP53_1033 | - | hypothetical protein |
| AP53_1034 | - | putative methyltransferase |
| AP53_1035 | - | FIG01118889: hypothetical protein |
| AP53_1036 | - | conserved hypothetical protein - phage associated |
| AP53_1037 | - | FIG01114622: hypothetical protein |
| AP53_1038 | - | hypothetical protein |
| AP53_1039 | - | putative replication protein |
| AP53_1040 | - | Phage protein |
| AP53_1041 | - | FIG01115822: hypothetical protein |
| AP53_1042 | - | FIG01117510: hypothetical protein |
| AP53_1043 | - | FIG01115915: hypothetical protein |
| AP53_1044 | - | hypothetical phage protein |
| AP53_1045 | - | hypothetical phage protein |
| AP53_1046 | - | hypothetical phage protein |
| AP53_1047 | - | hypothetical phage protein |
| AP53_1048 | - | hypothetical protein, phage associated |
| AP53_1049 | - | hypothetical phage protein |
| AP53_1050 | - | hypothetical protein |
| AP53_1051 | - | hypothetical protein |
| AP53_1052 | - | hypothetical protein |
| AP53_1053 | - | conserved hypothetical protein - phage associated |
| AP53_1054 | - | Phage transcriptional regulator, Cro/CI family |
| AP53_1055 | - | hypothetical phage protein |
| AP53_1056 | - | hypothetical protein |
| AP53_1057 | - | FIG01115240: hypothetical protein |
| AP53_1058 | - | hypothetical protein |
| AP53_1059 | - | Pleiotropic regulator of exopolysaccharide synthesis, competence and biofilm formation Ftr, XRE family |

| | | |
|---|---|---|
| AP53_1060 | - | hypothetical protein |
| AP53_1061 | int3 | Phage integrase: site-specific recombinase |
| AP53_1062 | - | lipoprotein, putative |
| AP53_1063 | glgP | Maltodextrin phosphorylase (EC 2.4.1.1) |
| AP53_1064 | malM | 4-alpha-glucanotransferase (amylomaltase) (EC 2.4.1.25) |
| AP53_1065 | malR | Maltose operon transcriptional repressor MalR, LacI family |
| AP53_1066 | malE | Maltose/maltodextrin ABC transporter, substrate binding periplasmic protein MalE |
| AP53_1067 | malF | Maltose/maltodextrin ABC transporter, permease protein MalF |
| AP53_1068 | malG | Maltose/maltodextrin ABC transporter, permease protein MalG |
| AP53_1069 | - | Mobile element protein |
| AP53_1070 | - | Mobile element protein |
| AP53_1071 | - | Transposase (maltose area) |
| AP53_1072 | | putative esterase |
| AP53_1073 | dltD | Poly(glycerophosphate chain) D-alanine transfer protein DltD |
| AP53_1074 | dltC | D-alanine--poly(phosphoribitol) ligase subunit 2 (EC 6.1.1.13) |
| AP53_1075 | dltB | D-alanyl transfer protein DltB |
| AP53_1076 | dltA | D-alanine--poly(phosphoribitol) ligase subunit 1 (EC 6.1.1.13) |
| AP53_1077 | - | FIG01114114: hypothetical protein |
| AP53_1078 | | Excinuclease ABC subunit B |
| AP53_1079 | glnPH | Glutamine ABC transporter, glutamine-binding protein/permease protein |
| AP53_1080 | | Glutamate transport ATP-binding protein |
| AP53_1081 | | FIG01114324: hypothetical protein |
| AP53_1082 | celB | PTS system, cellobiose-specific IIC component (EC 2.7.1.69) |
| AP53_1083 | | Putative uncharacterized protein SPs0857 |
| AP53_1084 | celC | PTS system, cellobiose-specific IIA component (EC 2.7.1.69) |
| AP53_1085 | celA | PTS system, cellobiose-specific IIB component (EC 2.7.1.69) |
| AP53_1086 | BglG | Transcriptional antiterminator of lichenan operon, BglG family |
| AP53_1087 | - | Outer surface protein of unknown function, cellobiose operon |
| AP53_1088 | bglA2 | Beta-glucosidase (EC 3.2.1.21) |
| AP53_1089 | pnuC | Ribosyl nicotinamide transporter, PnuC-like |
| AP53_1090 | - | FIG01114098: hypothetical protein |
| AP53_1091 | - | GTP-binding protein Obg |
| AP53_1092 | - | FIG01114947: hypothetical protein |
| AP53_1093 | - | Mobile element protein |
| AP53_1094 | - | Mobile element protein |

| | | |
|---|---|---|
| AP53_1095 | - | Mobile element protein |
| AP53_1096 | nadC | Quinolinate phosphoribosyltransferase [decarboxylating] (EC 2.4.2.19) |
| AP53_1097 | - | Malonate permease |
| AP53_1098 | - | Mobile element protein |
| AP53_1099 | - | Mobile element protein |
| AP53_1100 | - | Mobile element protein |
| AP53_1101 | | Ribosomal small subunit pseudouridine synthase A (EC 4.2.1.70) |
| AP53_1102 | - | Dihydrolipoamide acyltransferases |
| AP53_1103 | - | Transporter, MFS superfamily |
| AP53_1104 | - | ATP-dependent Zn protease |
| AP53_1105 | - | Phenylacetic acid degradation protein PaaD, thioesterase |
| AP53_1106 | - | Uridine phosphorylase (EC 2.4.2.3) |
| AP53_1107 | - | 23S rRNA (Uracil-5-) -methyltransferase RumA (EC 2.1.1.-) ## LSU rRNA m(5)U1939 |
| AP53_1108 | LytR-F1 | Cell envelope-associated transcriptional attenuator LytR-CpsA-Psr, subfamily F1 (as in PMID19099556) |
| AP53_1109 | | Shikimate kinase I (EC 2.7.1.71) |
| AP53_1110 | | 5-Enolpyruvylshikimate-3-phosphate synthase (EC 2.5.1.19) |
| AP53_1111 | YihY | Inner membrane protein YihY, formerly thought to be RNase BN |
| AP53_1112 | map | Methionine aminopeptidase (EC 3.4.11.18) |
| AP53_1113 | - | FIG009439: Cytosolic protein containing multiple CBS domains |
| AP53_1114 | | Ribosomal-protein-S5p-alanine acetyltransferase |
| AP53_1115 | graB | Protein G-related alpha 2 macroglobulin-binding protein (GRAB) |
| AP53_1116 | | UDP-N-acetylglucosamine 1-carboxyvinyltransferase (EC 2.5.1.7) |
| AP53_1117 | metK | S-adenosylmethionine synthetase (EC 2.5.1.6) |
| AP53_1118 | inlA | internalin, putative |
| AP53_1119 | birA | Biotin operon repressor / Biotin-protein ligase (EC 6.3.4.15) |
| AP53_1120 | - | FIG01114335: hypothetical protein |
| AP53_1121 | dnaX | DNA polymerase III subunits gamma and tau (EC 2.7.7.7) |
| AP53_1122 | - | GAF domain-containing proteins |
| AP53_1123 | - | FIG01115656: hypothetical protein |
| AP53_1124 | - | FIG01113965: hypothetical protein |
| AP53_1125 | udk | Uridine kinase (EC 2.7.1.48) [C1] |
| AP53_1126 | deaD2 | ATP-dependent RNA helicase YfmL |
| AP53_1127 | deadD2 | Peptidoglycan N-acetylglucosamine deacetylase (EC 3.5.1.-) |
| AP53_1128 | gapN | Non-phosphorylating glyceraldehyde-3-phosphate dehydrogenase (NADP) (EC 1.2.1.9) |
| AP53_1129 | ptsI | Phosphoenolpyruvate-protein phosphotransferase of PTS system (EC 2.7.3.9) |

| | | |
|---|---|---|
| AP53_1130 | | Phosphocarrier protein of PTS system |
| AP53_1131 | nrdH | Glutaredoxin-like protein NrdH, required for reduction of Ribonucleotide reductase class Ib |
| AP53_1132 | | Ribonucleotide reductase of class Ib (aerobic), alpha subunit (EC 1.17.4.1) |
| AP53_1133 | - | FIG01116145: hypothetical protein |
| AP53_1134 | | Ribonucleotide reductase of class Ib (aerobic), beta subunit (EC 1.17.4.1) |
| AP53_1135 | - | Chloride channel protein |
| AP53_1136 | - | Mobile element protein |
| AP53_1137 | - | Mobile element protein |
| AP53_1138 | - | Membrane-bound protease, CAAX family |
| AP53_1139 | - | FIG01114175: hypothetical protein |
| AP53_1140 | - | Transcriptional regulator, Cro/CI family |
| AP53_1141 | - | FIG01114322: hypothetical protein |
| AP53_1142 | | Alanyl-tRNA synthetase (EC 6.1.1.7) |
| AP53_1143 | prsA | Foldase protein prsA 1 precursor (EC 5.2.1.8) |
| AP53_1144 | - | O-methyltransferase family protein [C1] |
| AP53_1145 | - | Major facilitator:Oxalate:Formate Antiporter |
| AP53_1146 | pepB | Oligoendopeptidase F |
| AP53_1147 | CoiA | Competence protein CoiA |
| AP53_1148 | rsuA | Ribosomal small subunit pseudouridine synthase A (EC 4.2.1.70) |
| AP53_1149 | nagB | Glucosamine-6-phosphate deaminase (EC 3.5.99.6) |
| AP53_1150 | queA | S-adenosylmethionine:tRNA ribosyltransferase-isomerase (EC 5.-.-.-) |
| AP53_1151 | | FIG01115059: hypothetical protein |
| AP53_1152 | | FIG01113992: hypothetical protein |
| AP53_1153 | | FIG01114764: hypothetical protein |
| AP53_1154 | | FIG01116792: hypothetical protein |
| AP53_1155 | sodA | Manganese superoxide dismutase (EC 1.15.1.1) |
| AP53_1156 | holA | DNA polymerase III delta subunit (EC 2.7.7.7) |
| AP53_1157 | comEC | Late competence protein ComEC, DNA transport |
| AP53_1158 | comEA | Late competence protein ComEA, DNA receptor |
| AP53_1159 | plsC | 1-acyl-sn-glycerol-3-phosphate acyltransferase (EC 2.3.1.51) |
| AP53_1160 | - | tRNA (adenine37-N(6))-methyltransferase TrmN6 (EC 2.1.1.223) |
| AP53_1161 | - | COG2827: putative endonuclease containing a URI domain |
| AP53_1162 | - | Kup system potassium uptake protein |
| AP53_1163 | deaD | Cold-shock DEAD-box protein A |
| AP53_1164 | prfC | Peptide chain release factor 3 |

| | | |
|---|---|---|
| AP53_1165 | - | membrane protein, putative |
| AP53_1166 | | UDP-N-acetylmuramoylalanyl-D-glutamyl-2,6-diaminopimelate--D-alanyl-D-alanine ligase (EC 6.3.2.10) |
| AP53_1167 | ddl | D-alanine--D-alanine ligase A (EC 6.3.2.4) |
| AP53_1168 | recR | Recombination protein RecR |
| AP53_1169 | - | Formate/nitrite family of transporters |
| AP53_1170 | - | Substrate-specific component NiaX of predicted niacin ECF transporter |
| AP53_1171 | - | hypothetical protein |
| AP53_1172 | - | Transcriptional repressor for NAD biosynthesis in gram-positives |
| AP53_1173 | | Phosphoglycerate mutase (EC 5.4.2.1) |
| AP53_1174 | - | FIG01117945: hypothetical protein |
| AP53_1175 | pyrD | Dihydroorotate dehydrogenase, catalytic subunit (EC 1.3.3.1) |
| AP53_1176 | - | FIG01114217: hypothetical protein |
| AP53_1177 | | Lead, cadmium, zinc and mercury transporting ATPase (EC 3.6.3.3) (EC 3.6.3.5); Copper-translocating P-type |
| AP53_1178 | | Paratox |
| AP53_1179 | spd1 | Streptococcal extracellular nuclease 2; Mitogenic factor 2 |
| AP53_1180 | speC | Streptococcal pyrogenic exotoxin C (SpeC); _Toximoron (Superantigen) |
| AP53_1181 | - | Phage lysin, N-acetylmuramoyl-L-alanine amidase (EC 3.5.1.28) |
| AP53_pseudo1183 | -/pseudogene | Phage holin |
| AP53_1184 | - | Phage protein |
| AP53_1185 | - | FIG01116522: hypothetical protein |
| AP53_1186 | - | Phage hyaluronidase |
| AP53_1187 | | Hyaluronate lyase (phage associated) |
| AP53_1188 | pblB | Phage hyaluronidase; Phage endopeptidase |
| AP53_1189 | - | FIG022125: Phage-associated protein |
| AP53_1190 | - | Phage minor tail protein |
| AP53_1191 | - | Phage protein |
| AP53_1192 | - | conserved hypothetical protein - phage associated |
| AP53_1193 | - | Phage major tail protein |
| AP53_1194 | - | FIG018217: Phage-associated protein |
| AP53_1195 | - | FIG033577: Phage protein |
| AP53_1196 | - | FIG042840: Phage protein |
| AP53_1197 | - | FIG048485: Phage protein |
| AP53_1198 | - | FIG040428: Phage-associated protein |
| AP53_1199 | - | FIG01115109: hypothetical protein |
| AP53_1200 | pcfJ | hypothetical protein |

| | | |
|---|---|---|
| AP53_1201 | - | conserved hypothetical protein - phage associated |
| AP53_1202 | pi112 | Phage protein |
| AP53_1203 | - | Phage protein |
| AP53_1204 | - | Recombinational DNA repair protein RecT (prophage associated) |
| AP53_1205 | - | FIG01114509: hypothetical protein |
| AP53_1206 | - | Phage protein |
| AP53_1207 | - | hypothetical protein |
| AP53_1208 | - | FIG01114174: hypothetical protein |
| AP53_1209 | - | DNA replication protein dnaD |
| AP53_1210 | - | Phage protein |
| AP53_1211 | xis | Phage excisionase |
| AP53_1212 | - | Cro-like phage transcriptional repressor protein |
| AP53_1213 | - | Phage CI-like repressor |
| AP53_1214 | - | Phage protein |
| AP53_1215 | int4 | Phage integrase |
| AP53_1216 | HBsu | DNA-binding protein HBsu |
| AP53_1217 | yfaA | YfaA |
| AP53_1218 | - | FIG006988: Lipase/Acylhydrolase with GDSL-like motif |
| AP53_1219 | - | Hypothetical protein DUF194, DegV family |
| AP53_1220 | - | FIG01115348: hypothetical protein |
| AP53_1221 | recN | DNA repair protein RecN |
| AP53_1222 | argR | Arginine pathway regulatory protein ArgR, repressor of arg regulon |
| AP53_1223 | hlyA1 | RNA binding methyltransferase FtsJ like |
| AP53_1224 | fps | Octaprenyl diphosphate synthase (EC 2.5.1.90) / Dimethylallyltransferase (EC 2.5.1.1) / (2E,6E)-farnesyl diph |
| AP53_1225 | xseB | Exodeoxyribonuclease VII small subunit (EC 3.1.11.6) |
| AP53_1226 | xseA | Exodeoxyribonuclease VII large subunit (EC 3.1.11.6) |
| AP53_1227 | folD | Methylenetetrahydrofolate dehydrogenase (NADP+) (EC 1.5.1.5) / Methenyltetrahydrofolate cyclohydrolase |
| AP53_1228 | pgcA | putative phosphomannomutase |
| AP53_1229 | phr | Deoxyribodipyrimidine photolyase (EC 4.1.99.3) |
| AP53_1230 | | Glutamate transport ATP-binding protein |
| AP53_1231 | | ABC transporter membrane-spanning permease - glutamine transport |
| AP53_1232 | - | FIG01114188: hypothetical protein |
| AP53_1233 | clpE | ATP-dependent Clp protease, ATP-binding subunit ClpE |
| AP53_1234 | mutT | Mutator mutT protein (7,8-dihydro-8-oxoguanine-triphosphatase) (EC 3.6.1.-) |
| AP53_1235 | - | FIG01114027: hypothetical protein |

| | | |
|---|---|---|
| AP53_1236 | ileS | Isoleucyl-tRNA synthetase (EC 6.1.1.5) |
| AP53_1237 | divIVA | Cell division initiation protein DivIVA |
| AP53_1238 | - | FIG001583: hypothetical protein, contains S4-like RNA binding domain |
| AP53_1239 | ylmG | Cell division protein YlmG/Ycf19 (putative), YggT family |
| AP53_1240 | - | FIG001960: FtsZ-interacting protein related to cell division |
| AP53_1241 | - | Hypothetical protein YggS, proline synthase co-transcribed bacterial homolog PROSC |
| AP53_1242 | ftsZ | Cell division protein FtsZ (EC 3.4.24.-) |
| AP53_1243 | ftsA | Cell division protein FtsA |
| AP53_1244 | ftsQ | Cell division protein FtsQ |
| AP53_1245 | | UDP-N-acetylglucosamine--N-acetylmuramyl-(pentapeptide) pyrophosphoryl-undecaprenol N-acetylglucosa |
| AP53_1246 | | UDP-N-acetylmuramoylalanine--D-glutamate ligase (EC 6.3.2.9) |
| AP53_1247 | - | FIG01114439: hypothetical protein |
| AP53_1248 | typA | GTP-binding protein TypA/BipA |
| AP53_1249 | - | Rhodanese-like domain protein |
| AP53_1250 | glkA | Glucokinase (EC 2.7.1.2) |
| AP53_1251 | - | FIG01114576: hypothetical protein |
| AP53_1252 | dpr | Non-specific DNA-binding protein Dps / Iron-binding ferritin-like antioxidant protein / Ferroxidase (EC 1.16.[illegible] |
| AP53_1253 | - | Late competence protein ComC, processing protease |
| AP53_1254 | - | Ribosomal RNA large subunit methyltransferase N (EC 2.1.1.-) |
| AP53_1255 | - | Hypothetical DUF1027 domain protein |
| AP53_1256 | - | FIG01114963: hypothetical protein |
| AP53_1257 | - | LacI family regulatory protein |
| AP53_1258 | lon | Lon-like protease with PDZ domain |
| AP53_1259 | coaD | Phosphopantetheine adenylyltransferase (EC 2.7.7.3) |
| AP53_1260 | rsmD | 16S rRNA (guanine(966)-N(2))-methyltransferase (EC 2.1.1.171) ## SSU rRNA m(2)G966 |
| AP53_1261 | asnA | Aspartate--ammonia ligase (EC 6.3.1.1) |
| AP53_1262 | arcC | Carbamate kinase (EC 2.7.2.2) |
| AP53_1263 | arcE | Acetylornithine deacetylase/Succinyl-diaminopimelate desuccinylase and related deacylases |
| AP53_1264 | arcD | Arginine/ornithine antiporter ArcD |
| AP53_1265 | arcB | Ornithine carbamoyltransferase (EC 2.1.3.3) |
| AP53_1266 | - | Acetyltransferase (EC 2.3.1.-) |
| AP53_1267 | arcA | Arginine deiminase (EC 3.5.3.6) |
| AP53_1268 | arcR | Transcriptional regulator ArcR essential for anaerobic expression of the ADI pathway, Crp/Fnr family |
| AP53_1269 | argR | Arginine pathway regulatory protein ArgR, repressor of arg regulon |
| AP53_1270 | - | Putative uncharacterized protein gbs2121 |

| | | |
|---|---|---|
| AP53_1271 | - | Hypothetical protein SpyM3_1200 |
| AP53_1272 | yesM | Two-component sensor kinase yesM (EC 2.7.3.-), associated with MetSO reductase |
| AP53_1273 | yesN | Two-component response regulator yesN, associated with MetSO reductase |
| AP53_1274 | msrAB | Peptide methionine sulfoxide reductase MsrA (EC 1.8.4.11) / Peptide methionine sulfoxide reductase MsrB ([illegible] |
| AP53_1275 | tlpA | Thiol:disulfide oxidoreductase associated with MetSO reductase |
| AP53_1276 | ccdA | Cytochrome c-type biogenesis protein CcdA homolog, associated with MetSO reductase |
| AP53_1277 | - | Magnesium and cobalt transport protein CorA |
| AP53_1278 | - | Oxidoreductase, Gfo/Idh/MocA family |
| AP53_1279 | - | FIG01114815: hypothetical protein |
| AP53_1280 | valS | Valyl-tRNA synthetase (EC 6.1.1.9) |
| AP53_1281 | - | hypothetical protein |
| AP53_1282 | - | hypothetical protein |
| AP53_1283 | - | FIG01114159: hypothetical protein |
| AP53_1284 | - | Aldose 1-epimerase (EC 5.1.3.3) |
| AP53_rna61 | - | tRNA-Thr-GGT |
| AP53_1285 | aroA2 | 2-keto-3-deoxy-D-arabino-heptulosonate-7-phosphate synthase I beta (EC 2.5.1.54) |
| AP53_1286 | aroB | 3-dehydroquinate synthase (EC 4.2.3.4) |
| AP53_1287 | - | Uncharacterized conserved protein, contains double-stranded beta-helix domain |
| AP53_1288 | - | hypothetical protein |
| AP53_1289 | - | SAM-dependent methyltransferase HI0095 (UbiE paralog) |
| AP53_1290 | aroE | Shikimate/quinate 5-dehydrogenase I beta (EC 1.1.1.282) |
| AP53_1291 | lacZ | Beta-galactosidase (EC 3.2.1.23) |
| AP53_1292 | | DNA-binding response regulator, AraC family |
| AP53_1293 | | Two-component sensor kinase YesM (EC 2.7.3.-) |
| AP53_1294 | - | Integral membrane protein |
| AP53_1295 | - | Multiple sugar ABC transporter, substrate-binding protein |
| AP53_1296 | - | Multiple sugar ABC transporter, membrane-spanning permease protein MsmG |
| AP53_1297 | - | Multiple sugar ABC transporter, membrane-spanning permease protein MsmF |
| AP53_1298 | nanK | N-acetylmannosamine kinase (EC 2.7.1.60) |
| AP53_1299 | | FIG01114249: hypothetical protein |
| AP53_1300 | | Beta-glucosidase (EC 3.2.1.21) |
| AP53_1301 | hyl | Hyaluronoglucosaminidase (EC 3.2.1.35) |
| AP53_1302 | | Transcriptional repressor of arabinoside utilization operon, GntR family |
| AP53_1303 | | hypothetical protein |
| AP53_1304 | ams1 | Alpha-mannosidase (EC 3.2.1.24) |

| | | |
|---|---|---|
| AP53_1305 | rocA | Putative two-component responsible histidine kinase |
| AP53_1306 | rumA | RNA methyltransferase, TrmA family |
| AP53_1307 | prx | Paratox |
| AP53_1308 | spd3 | Streptococcal extracellular nuclease 3; Mitogenic factor 3 |
| AP53_1309 | | FIG01117556: hypothetical protein |
| AP53_1310 | - | Phage protein |
| AP53_1311 | - | Phage protein |
| AP53_1312 | - | Phage lysin |
| AP53_1313 | - | Phage holin |
| AP53_1314 | - | Phage protein |
| AP53_1315 | - | FIG01116522: hypothetical protein |
| AP53_1316 | - | Phage hyaluronidase |
| AP53_1317 | - | Hyaluronate lyase (phage associated) |
| AP53_1318 | - | Phage endopeptidase |
| AP53_1319 | - | Phage-related protein |
| AP53_1320 | - | Phage tail length tape-measure protein |
| AP53_1321 | - | Phage protein |
| AP53_1322 | - | Phage protein |
| AP53_1323 | - | Phage major tail shaft protein |
| AP53_1324 | - | Phage minor capsid protein |
| AP53_1325 | - | Phage minor capsid protein #Fam0017 |
| AP53_1326 | - | Phage minor capsid protein |
| AP53_1327 | - | Phage protein |
| AP53_1328 | - | Phage major capsid protein |
| AP53_1329 | - | Phage protein |
| AP53_1330 | - | Phage protein |
| AP53_1331 | - | hypothetical protein, phage associated |
| AP53_1332 | - | Phage minor capsid protein |
| AP53_1333 | - | Phage minor capsid protein #Fam0015 |
| AP53_1334 | - | Phage terminase large subunit |
| AP53_1335 | - | Phage protein |
| AP53_pseudo1337 | -/pseudogene | DNA modification methylase |
| AP53_1338 | - | FIG01114733: hypothetical protein |
| AP53_1339 | - | hypothetical protein |
| AP53_1340 | - | conserved hypothetical protein - phage associated |

| | | |
|---|---|---|
| AP53_1341 | - | Phage protein |
| AP53_1342 | - | Phage protein |
| AP53_1343 | - | Phage protein |
| AP53_1344 | - | FIG01115146: hypothetical protein |
| AP53_1345 | - | FIG01114622: hypothetical protein |
| AP53_1346 | - | Phage Holliday junction resolvase |
| AP53_1347 | | Single-stranded DNA-binding protein |
| AP53_1348 | | Single-stranded DNA-binding protein |
| AP53_1349 | - | hypothetical protein (phage associated) |
| AP53_1350 | - | FIG01116817: hypothetical protein |
| AP53_1351 | - | hypothetical phage protein |
| AP53_1352 | - | hypothetical phage protein |
| AP53_1353 | - | DNA replication protein DnaC |
| AP53_1354 | - | putative phage replication protein |
| AP53_1355 | - | hypothetical protein, phage associated |
| AP53_1356 | - | phage protein |
| AP53_1357 | - | conserved hypothetical protein - phage associated |
| AP53_1358 | - | hypothetical protein - phage associated |
| AP53_1359 | - | Phage antirepressor protein |
| AP53_1360 | - | FIG01117062: hypothetical protein |
| AP53_1361 | - | putative cI-like repressor - phage associated |
| AP53_1362 | - | Phage protein |
| AP53_1363 | - | Chromosome segregation ATPase |
| AP53_1364 | - | Phage integrase: site-specific recombinase |
| AP53_1365 | recX | Regulatory protein recX |
| AP53_1366 | - | Cysteinyl-tRNA synthetase related protein |
| AP53_rna62 | - | tRNA-Glu-TTC |
| AP53_rna63 | - | tRNA-Ile-GAT |
| AP53_rna64 | - | tRNA-Thr-TGT |
| AP53_rna65 | - | tRNA-Leu-TAG |
| AP53_rna66 | - | tRNA-Lys-TTT |
| AP53_rna67 | - | tRNA-Asp-GTC |
| AP53_rna68 | - | tRNA-Val-TAC |
| AP53_rna69 | - | 5S RNA |
| AP53_rna70 | - | Large Subunit Ribosomal RNA; lsuRNA; LSU rRNA |

| | | |
|---|---|---|
| AP53_rna71 | - | tRNA-Ala-TGC |
| AP53_rna72 | - | Small Subunit Ribosomal RNA; ssuRNA; SSU rRNA |
| AP53_1367 | raiA | Ribosomal subunit interface protein |
| AP53_1368 | comFC | Competence protein F homolog, phosphoribosyltransferase domain; protein YhgH required for utilization of |
| AP53_1369 | comFA | ComF operon protein A, DNA transporter ATPase |
| AP53_1370 | - | FIG000605: protein co-occurring with transport systems (COG1739) |
| AP53_1371 | cysM | Cysteine synthase (EC 2.5.1.47) |
| AP53_1372 | - | General stress protein 13 |
| AP53_1373 | - | Streptococcal lipoprotein rotamase A; Peptidyl-prolyl cis-trans isomerase (EC 5.2.1.8) |
| AP53_1374 | yvqC | Two component transcriptional regulator VraR |
| AP53_1375 | yvqE | Sensor histidine kinase VraS |
| AP53_1376 | yvqF | Transporter associated with VraSR |
| AP53_1377 | | Serine/threonine protein kinase PrkC, regulator of stationary phase |
| AP53_1378 | | Protein serine/threonine phosphatase PrpC, regulation of stationary phase |
| AP53_1379 | sunL | 16S rRNA (cytosine(967)-C(5))-methyltransferase (EC 2.1.1.176) |
| AP53_1380 | fmt | Methionyl-tRNA formyltransferase (EC 2.1.2.9) |
| AP53_1381 | priA | Helicase PriA essential for oriC/DnaA-independent DNA replication |
| AP53_1382 | rpoZ | DNA-directed RNA polymerase omega subunit (EC 2.7.7.6) |
| AP53_1383 | | Guanylate kinase (EC 2.7.4.8) |
| AP53_1384 | cvfA | Hydrolase (HAD superfamily) |
| AP53_1385 | - | Transcriptional regulators, LysR family |
| AP53_1386 | atoB | 3-ketoacyl-CoA thiolase (EC 2.3.1.16) @ Acetyl-CoA acetyltransferase (EC 2.3.1.9) |
| AP53_1387 | atoD1 | Butyrate-acetoacetate CoA-transferase subunit A (EC 2.8.3.9) |
| AP53_1388 | atoA | Butyrate-acetoacetate CoA-transferase subunit B (EC 2.8.3.9) |
| AP53_pseudo1389 | -/pseudogene | D-beta-hydroxybutyrate dehydrogenase (EC 1.1.1.30) |
| AP53_1391 | - | D-beta-hydroxybutyrate permease |
| AP53_1392 | luxS | S-ribosylhomocysteine lyase (EC 4.4.1.21) / Autoinducer-2 production protein LuxS |
| AP53_1393 | - | Hypothetical protein SPy1643 |
| AP53_1394 | - | FIG001721: Predicted N6-adenine-specific DNA methylase |
| AP53_1395 | gpsB | Cell division protein GpsB, coordinates the switch between cylindrical and septal cell wall synthesis by re-loc |
| AP53_1396 | - | FIG005686: hypothetical protein |
| AP53_1397 | recU | RecU Holliday junction resolvase |
| AP53_1398 | pbp1A | Multimodular transpeptidase-transglycosylase (EC 2.4.1.129) (EC 3.4.-.-) / Penicillin-binding protein 1A/1B ( |
| AP53_1399 | pepC | Aminopeptidase C (EC 3.4.22.40) |
| AP53_1400 | nadE | NAD synthetase (EC 6.3.1.5) |

| | | |
|---|---|---|
| AP53_1401 | pncB | Nicotinate phosphoribosyltransferase (EC 2.4.2.11) |
| AP53_1402 | - | Histidine transport protein (permease) |
| AP53_1403 | | Thioredoxin reductase (EC 1.8.1.9) |
| AP53_1404 | aapA | Hypothetical protein SPy1656 |
| AP53_1405 | tcyN | ABC-type multidrug transport system, ATPase component |
| AP53_1406 | yxeN | Cysteine ABC transporter, permease protein |
| AP53_1407 | mraW | FIG01116065: hypothetical protein |
| AP53_1408 | - | ATP-dependent RNA helicase YqfR |
| AP53_1409 | mraY | Phospho-N-acetylmuramoyl-pentapeptide-transferase (EC 2.7.8.13) |
| AP53_1410 | pbpX | Cell division protein FtsI [Peptidoglycan synthetase] (EC 2.4.1.129) |
| AP53_1411 | ftsL | Cell division protein FtsL |
| AP53_1412 | rmsH | rRNA small subunit methyltransferase H |
| AP53_1413 | - | FIG01115263: hypothetical protein |
| AP53_1414 | | Gamma-glutamyl phosphate reductase (EC 1.2.1.41) |
| AP53_1415 | SK | Glutamate 5-kinase (EC 2.7.2.11) |
| AP53_1416 | | ABC transporter permease protein |
| AP53_1417 | - | Methionine ABC transporter ATP-binding protein |
| AP53_1418 | - | FIG01113961: hypothetical protein |
| AP53_1419 | tkt | Transketolase (EC 2.2.1.1) |
| AP53_1420 | talC | Transaldolase (EC 2.2.1.2) |
| AP53_1421 | - | Trans-acting positive regulator |
| AP53_1422 | npx | NADH peroxidase (EC 1.11.1.1) |
| AP53_1423 | glpF | Glycerol uptake facilitator protein |
| AP53_1424 | glpO | Alpha-glycerophosphate oxidase (EC 1.1.3.21) |
| AP53_1425 | glpK | Glycerol kinase (EC 2.7.1.30) |
| AP53_1426 | - | FIG01116532: hypothetical protein |
| AP53_1427 | | FIG01116327: hypothetical protein |
| AP53_1428 | - | FIG01114118: hypothetical protein |
| AP53_1429 | | Glycyl-tRNA synthetase beta chain (EC 6.1.1.14) |
| AP53_1430 | | Glycyl-tRNA synthetase alpha chain (EC 6.1.1.14) |
| AP53_1431 | - | INTEGRAL MEMBRANE PROTEIN (Rhomboid family) |
| AP53_1432 | AKR | oxidoreductase of aldo/keto reductase family, subgroup 1 |
| AP53_1433 | nagA | N-acetylglucosamine-6-phosphate deacetylase (EC 3.5.1.25) |
| AP53_1434 | nptA | Sodium-dependent phosphate transporter |
| AP53_1435 | - | FIG01116130: hypothetical protein |

| | | |
|---|---|---|
| AP53_1436 | DegV | DegV family protein |
| AP53_1437 | | Transcriptional regulator, TetR family |
| AP53_1438 | - | Hydrolase (HAD superfamily) |
| AP53_1439 | - | INTEGRAL MEMBRANE PROTEIN (Rhomboid family) |
| AP53_1440 | - | hypothetical protein |
| AP53_1441 | lacD1 | Tagatose 1,6-diphosphate aldolase (EC 4.1.2.40) |
| AP53_1442 | lacC1 | Tagatose-6-phosphate kinase (EC 2.7.1.144) |
| AP53_1443 | lacB1 | Galactose-6-phosphate isomerase, LacB subunit (EC 5.3.1.26) |
| AP53_1444 | lacA1 | Galactose-6-phosphate isomerase, LacA subunit (EC 5.3.1.26) |
| AP53_1445 | gatC | PTS system, galactose-specific IIC component (EC 2.7.1.69) |
| AP53_1446 | gatB | PTS system, galactose-specific IIB component (EC 2.7.1.69) |
| AP53_1447 | gatA | PTS system, galactose-specific IIA component (EC 2.7.1.69) |
| AP53_1448 | lacR1 | Lactose phosphotransferase system repressor |
| AP53_1449 | copZ | Copper chaperone |
| AP53_1450 | copA | Copper-translocating P-type ATPase (EC 3.6.3.4) |
| AP53_1451 | copY | Negative transcriptional regulator-copper transport operon |
| AP53_1452 | sse | putative esterase |
| AP53_1453 | rbfA | Ribosome-binding factor A |
| AP53_1454 | infB | Translation initiation factor 2 |
| AP53_1455 | - | ribosomal protein L7Ae family protein |
| AP53_1456 | - | COG2740: Predicted nucleic-acid-binding protein implicated in transcription termination |
| AP53_1457 | nusA | Transcription termination protein NusA |
| AP53_1458 | rimP | FIG000325: clustered with transcription termination protein NusA |
| AP53_1459 | - | Mobile element protein |
| AP53_rna73 | - | tRNA-Ser-GGA |
| AP53_1460 | trmB | tRNA (guanine46-N7-)-methyltransferase (EC 2.1.1.33) |
| AP53_1461 | - | aminoglycoside phosphotransferase family protein |
| AP53_1462 | ecsB | ABC transporter, permease protein EscB |
| AP53_1463 | ecsA | ABC transporter, ATP-binding protein EcsA |
| AP53_1464 | hit | Histidine triad (HIT) nucleotide-binding protein, similarity with At5g48545 and yeast YDL125C (HNT1) |
| AP53_1465 | - | Hypothetical protein in cluster with Ecs transporter (in Streptococci) |
| AP53_1466 | lytR-F2 | Cell envelope-associated transcriptional attenuator LytR-CpsA-Psr, subfamily F2 (as in PMID19099556) |
| AP53_1467 | - | Acetyltransferase (EC 2.3.1.-) |
| AP53_1468 | - | ATPase YjeE, predicted to have essential role in cell wall biosynthesis |
| AP53_1469 | - | Guanine-hypoxanthine permease |

| | | |
|---|---|---|
| AP53_1470 | - | Hydrolase (HAD superfamily) |
| AP53_1471 | manL | PTS system, mannose-specific IIB component (EC 2.7.1.69) / PTS system, mannose-specific IIA component (E[illegible] |
| AP53_1472 | manM | PTS system, mannose-specific IIC component (EC 2.7.1.69) |
| AP53_1473 | manN | PTS system, mannose-specific IID component (EC 2.7.1.69) |
| AP53_1474 | - | Putative regulator of the mannose operon, ManO |
| AP53_1475 | - | Mobile element protein |
| AP53_1476 | serS | Seryl-tRNA synthetase (EC 6.1.1.11) |
| AP53_1477 | | Acetyl-coenzyme A carboxyl transferase alpha chain (EC 6.4.1.2) |
| AP53_1478 | | Acetyl-coenzyme A carboxyl transferase beta chain (EC 6.4.1.2) |
| AP53_1479 | | Biotin carboxylase of acetyl-CoA carboxylase (EC 6.3.4.14) |
| AP53_1480 | fabZ | (3R)-hydroxymyristoyl-[acyl carrier protein] dehydratase (EC 4.2.1.-) |
| AP53_1481 | accB | Biotin carboxyl carrier protein of acetyl-CoA carboxylase |
| AP53_1482 | fabF | 3-oxoacyl-[acyl-carrier-protein] synthase, KASII (EC 2.3.1.41) |
| AP53_1483 | fabG | 3-oxoacyl-[acyl-carrier protein] reductase (EC 1.1.1.100) |
| AP53_1484 | fabD | Malonyl CoA-acyl carrier protein transacylase (EC 2.3.1.39) |
| AP53_1485 | fabK | Enoyl-[acyl-carrier-protein] reductase [FMN] (EC 1.3.1.9) |
| AP53_1486 | acpP | Acyl carrier protein |
| AP53_1487 | fabH | 3-oxoacyl-[acyl-carrier-protein] synthase, KASIII (EC 2.3.1.41) |
| AP53_1488 | FabT | Transcriptional regulator of fatty acid biosynthesis FabT |
| AP53_1489 | | Trans-2,cis-3-Decenoyl-ACP isomerase |
| AP53_1490 | dnaJ | Chaperone protein DnaJ |
| AP53_1491 | dnaK | Chaperone protein DnaK |
| AP53_1492 | grpE | Heat shock protein GrpE |
| AP53_1493 | hrcA | Heat-inducible transcription repressor HrcA |
| AP53_1494 | - | putative N-acetyl-muramidase |
| AP53_1495 | - | D-alanyl-D-alanine carboxypeptidase (EC 3.4.16.4) |
| AP53_1496 | - | Phosphoglycerate mutase family |
| AP53_1497 | - | Integral membrane protein |
| AP53_1498 | - | hypothetical protein |
| AP53_1499 | gatB | Aspartyl-tRNA(Asn) amidotransferase subunit B (EC 6.3.5.6) @ Glutamyl-tRNA(Gln) amidotransferase subunit |
| AP53_1500 | gatA | Aspartyl-tRNA(Asn) amidotransferase subunit A (EC 6.3.5.6) @ Glutamyl-tRNA(Gln) amidotransferase subunit |
| AP53_1501 | gatC | Aspartyl-tRNA(Asn) amidotransferase subunit C (EC 6.3.5.6) @ Glutamyl-tRNA(Gln) amidotransferase subunit |
| AP53_1502 | - | Mobile element protein |
| AP53_1503 | - | Pyruvate,phosphate dikinase (EC 2.7.9.1) |
| AP53_1504 | - | Nicotinamidase (EC 3.5.1.19) |

| | | |
|---|---|---|
| AP53_1505 | codY | GTP-sensing transcriptional pleiotropic repressor codY |
| AP53_1506 | - | Aspartate aminotransferase (EC 2.6.1.1) |
| AP53_1507 | - | Universal stress protein family |
| AP53_1508 | - | Hydrolase (HAD superfamily) |
| AP53_1509 | asnB | L-asparaginase (EC 3.5.1.1) |
| AP53_1510 | - | ABC transporter permease protein |
| AP53_1511 | - | putative ABC transporter (ATP-binding protein) |
| AP53_1512 | recG | ATP-dependent DNA helicase RecG (EC 3.6.1.-) |
| AP53_1513 | cbiO | Duplicated ATPase component BL0693 of energizing module of predicted ECF transporter |
| AP53_1514 | cbiQ | Transmembrane component BL0694 of energizing module of predicted ECF transporter |
| AP53_1515 | cbiMN | Substrate-specific component BL0695 of predicted ECF transporter |
| AP53_1516 | cycC | Putative ABC transporter ATP-binding protein, spy1790 homolog |
| AP53_1517 | cycD | Putative ABC transporter (ATP-binding protein), spy1791 homolog |
| AP53_1518 | fhuC-htsC | Heme ABC transporter (Streptococcus), ATP-binding protein |
| AP53_1519 | fhuB-htsB | Heme ABC transporter (Streptococcus), permease protein |
| AP53_1520 | fhuD2-htsA | Heme transporter IsdDEF, lipoprotein IsdE |
| AP53_1521 | shp | Cell surface protein Shp, transfers heme from hemoglobin to apo-SiaA/HtsA |
| AP53_1522 | shr | Streptococcal cell surface hemoprotein receptor Shr |
| AP53_1523 | - | hypothetical protein |
| AP53_1524 | | Immunogenic secreted protein |
| AP53_1525 | alr | Alanine racemase (EC 5.1.1.1) |
| AP53_1526 | acpS | Holo-[acyl-carrier protein] synthase (EC 2.7.8.7) |
| AP53_1527 | secA | Protein export cytoplasm protein SecA ATPase RNA helicase (TC 3.A.5.1.1) |
| AP53_1528 | - | Mobile element protein |
| AP53_1529 | pmi | Mannose-6-phosphate isomerase (EC 5.3.1.8) |
| AP53_1530 | scrK | Fructokinase (EC 2.7.1.4) |
| AP53_1531 | endoS | Secreted Endo-beta-N-acetylglucosaminidase (EndoS) |
| AP53_1532 | scrA | PTS system, sucrose-specific IIB component (EC 2.7.1.69) / PTS system, sucrose-specific IIC component (EC 2. |
| AP53_1533 | scrB | Sucrose-6-phosphate hydrolase (EC 3.2.1.26) |
| AP53_1534 | scrR | Sucrose operon repressor ScrR, LacI family |
| AP53_1535 | nusB | Transcription termination protein NusB |
| AP53_1536 | - | Alkaline shock protein |
| AP53_1537 | efp | Translation elongation factor P |
| AP53_1538 | comEB | dCMP deaminase (EC 3.5.4.12); Late competence protein ComEB |
| AP53_1539 | pepP | Aminopeptidase YpdF (MP-, MA-, MS-, AP-, NP- specific) |

| | | |
|---|---|---|
| AP53_1540 | | Excinuclease ABC subunit A |
| AP53_1541 | corA | Magnesium and cobalt transport protein CorA |
| AP53_1542 | - | Integral membrane protein |
| AP53_1543 | rpsR | SSU ribosomal protein S18p @ SSU ribosomal protein S18p, zinc-independent |
| AP53_1544 | | Single-stranded DNA-binding protein |
| AP53_1545 | rpsF | SSU ribosomal protein S6p |
| AP53_1546 | - | hypothetical protein |
| AP53_1547 | mutY | A/G-specific adenine glycosylase (EC 3.2.2.-) |
| AP53_1548 | - | Transcriptional regulator, XRE family |
| AP53_1549 | trxA | Thioredoxin |
| AP53_1550 | - | Phosphatidylglycerophosphatase B (EC 3.1.3.27) |
| AP53_1551 | - | Recombination inhibitory protein MutS2 |
| AP53_1552 | - | Colicin V production protein |
| AP53_1553 | - | FIG01114225: hypothetical protein |
| AP53_1554 | | Ribonuclease HIII (EC 3.1.26.4) |
| AP53_1555 | spi | Putative signal peptidase I (EC 3.4.21.89) |
| AP53_1556 | recD | RecD-like DNA helicase YrrC |
| AP53_1557 | - | Putative membrane protein precursor SPs0273 |
| AP53_1558 | dinP | DNA polymerase IV (EC 2.7.7.7) |
| AP53_1559 | pflB | Pyruvate formate-lyase (EC 2.3.1.54) |
| AP53_1560 | - | hypothetical protein |
| AP53_1561 | - | Beta-lactamase, class C |
| AP53_1562 | cppA | C3-degrading proteinase |
| AP53_1563 | - | Integral membrane protein |
| AP53_1564 | gla | Aquaporin (Major Intrinsic Protein Family) |
| AP53_1565 | | transporter |
| AP53_1566 | | Transcriptional regulator, CrP/Fnr family |
| AP53_1567 | | Xaa-Pro dipeptidyl-peptidase (EC 3.4.14.11) |
| AP53_1568 | - | Transcriptional regulator, Cro/CI family |
| AP53_1569 | - | putative repressor - phage associated |
| AP53_1570 | - | Transcriptional regulatory protein |
| AP53_1571 | - | Transcriptional regulator, MerR family |
| AP53_1572 | dnaQ | DNA polymerase III epsilon subunit (EC 2.7.7.7) |
| AP53_1573 | - | Transcriptional regulator OrfX |
| AP53_1574 | - | NAD(FAD)-utilizing dehydrogenases |

| | | |
|---|---|---|
| AP53_1575 | | Deoxyribose-phosphate aldolase (EC 4.1.2.4) |
| AP53_1576 | nupC | Nucleoside permease NupC |
| AP53_1577 | udp | Uridine phosphorylase (EC 2.4.2.3) |
| AP53_1578 | crgR | Predicted regulator for deoxynucleoside utilization, GntR family |
| AP53_1579 | rpsN | SSU ribosomal protein S14p (S29e) @ SSU ribosomal protein S14p (S29e), zinc-independent |
| AP53_1580 | gcp | TsaD/Kae1/Qri7 protein, required for threonylcarbamoyladenosine t(6)A37 formation in tRNA |
| AP53_1581 | rimI | Ribosomal-protein-S18p-alanine acetyltransferase (EC 2.3.1.-) |
| AP53_1582 | - | Inactive homolog of metal-dependent proteases, putative molecular chaperone |
| AP53_1583 | - | Protein of unknown function DUF1447 |
| AP53_1584 | rnjA | Ribonuclease J1 (endonuclease and 5' exonuclease) |
| AP53_1585 | glnA | Glutamine synthetase type I (EC 6.3.1.2) |
| AP53_1586 | glnR | Transcriptional regulator, MerR family |
| AP53_1587 | - | FIG01114549: hypothetical protein |
| AP53_1588 | pgk | Phosphoglycerate kinase (EC 2.7.2.3) |
| AP53_1589 | lppC | Acid phosphatase (EC 3.1.3.2) |
| AP53_1590 | ybbK | Putative stomatin/prohibitin-family membrane protease subunit YbbK |
| AP53_1591 | - | Dihydroxyacetone kinase family protein |
| AP53_1592 | asp | FIG001802: Putative alkaline-shock protein |
| AP53_1593 | rpmB | LSU ribosomal protein L28p |
| AP53_1594 | fba | Fructose-bisphosphate aldolase class II (EC 4.1.2.13) |
| AP53_rna74 | - | tRNA-Leu-AAG |
| AP53_1595 | - | Hydrolases of the alpha/beta superfamily |
| AP53_1596 | - | hypothetical protein |
| AP53_1597 | pyrG | CTP synthase (EC 6.3.4.2) |
| AP53_1598 | rpoE | DNA-directed RNA polymerase delta subunit (EC 2.7.7.6) |
| AP53_1599 | tig | Cell division trigger factor (EC 5.2.1.8) |
| AP53_1600 | - | Potassium efflux system KefA protein / Small-conductance mechanosensitive channel |
| AP53_1601 | - | Hypothetical protein ywlG |
| AP53_1602 | - | hypothetical protein |
| AP53_1603 | - | Substrate-specific component PdxU2 of predicted pyridoxin-related ECF transporter |
| AP53_1604 | thiD | Novel pyridoxal kinase, thiD family (EC 2.7.1.35) |
| AP53_1605 | truA | tRNA pseudouridine synthase A (EC 4.2.1.70) |
| AP53_1606 | comX2 | Competence-specific sigma factor ComX |
| AP53_rna75 | - | tRNA-Arg-ACG |
| AP53_rna76 | - | tRNA-Asn-GTT |

| | | |
|---|---|---|
| AP53_rna77 | - | 5S RNA |
| AP53_rna78 | - | Large Subunit Ribosomal RNA; lsuRNA; LSU rRNA |
| AP53_rna79 | - | tRNA-Ala-TGC |
| AP53_rna80 | - | Small Subunit Ribosomal RNA; ssuRNA; SSU rRNA |
| AP53_1607 | - | Glycerate kinase (EC 2.7.1.31) |
| AP53_1608 | hsdR | Type I restriction-modification system, restriction subunit R (EC 3.1.21.3) |
| AP53_1609 | hsdS | Type I restriction-modification system, specificity subunit S (EC 3.1.21.3) |
| AP53_1610 | hsdM | Type I restriction-modification system, DNA-methyltransferase subunit M (EC 2.1.1.72) |
| AP53_1611 | | Transcriptional regulator DegU, LuxR family |
| AP53_1612 | | FIG00525014: hypothetical protein |
| AP53_1613 | - | Salivaricin A modification enzyme |
| AP53_pseudo1615 | salB/pseudogene | Lanthionine biosynthesis protein LanM/pseudogene causing stop codon at 177th aa |
| AP53_1616 | salA | Lantibiotic salivaricin A |
| AP53_1617 | lacG | 6-phospho-beta-galactosidase (EC 3.2.1.85) |
| AP53_1618 | lacE | PTS system, lactose-specific IIB component (EC 2.7.1.69) / PTS system, lactose-specific IIC component (EC 2.7 |
| AP53_1619 | lacF | PTS system, lactose-specific IIA component (EC 2.7.1.69) |
| AP53_1620 | lacD2 | Tagatose 1,6-diphosphate aldolase (EC 4.1.2.40) |
| AP53_1621 | lacC2 | Tagatose-6-phosphate kinase (EC 2.7.1.144) |
| AP53_1622 | lacB2 | Galactose-6-phosphate isomerase, LacB subunit (EC 5.3.1.26) |
| AP53_1623 | lacA2 | Galactose-6-phosphate isomerase, LacA subunit (EC 5.3.1.26) |
| AP53_1624 | lacR2 | Lactose phosphotransferase system repressor |
| AP53_1625 | - | DNA-damage-inducible protein J |
| AP53_1626 | - | Toxin to DNA-damage-inducible protein J |
| AP53_1627 | - | hypothetical protein |
| AP53_1628 | - | Integrase |
| AP53_1629 | - | Protein of unknown function DUF208 |
| AP53_1630 | rpsI | SSU ribosomal protein S9p (S16e) |
| AP53_1631 | rplM | LSU ribosomal protein L13p (L13Ae) |
| AP53_1632 | - | Transcriptional regulator, Cro/CI family |
| AP53_1633 | - | hypothetical protein |
| AP53_1634 | - | DegV family protein in cluster with TrmH family tRNA/rRNA methyltransferase YacO |
| AP53_1635 | - | Hypothetical protein DUF901, similar to C-terminal domain of ribosome protection-type Tc-resistance prote |
| AP53_1636 | - | 23S rRNA (guanosine-2'-O-) -methyltransferase rlmB (EC 2.1.1.-) ## LSU rRNA Gm2251 |
| AP53_1637 | - | Hypothetical protein, Spy1939 homolog |
| AP53_1638 | - | COG1939: Ribonuclease III family protein |

| | | |
|---|---|---|
| AP53_1639 | cysS | Cysteinyl-tRNA synthetase (EC 6.1.1.16) |
| AP53_1640 | | FIG01114632: hypothetical protein |
| AP53_1641 | cysE | Serine acetyltransferase (EC 2.3.1.30) |
| AP53_1642 | | FIG01114029: hypothetical protein |
| AP53_1643 | pnp | Polyribonucleotide nucleotidyltransferase (EC 2.7.7.8) |
| AP53_1644 | | Transaldolase (EC 2.2.1.2) |
| AP53_1645 | sgaT | Transport protein SgaT, putative |
| AP53_1646 | | PTS system IIB component |
| AP53_1647 | bglG | Transcription antiterminator, BglG family (EC 2.7.1.69) |
| AP53_1648 | rpsO | SSU ribosomal protein S15p (S13e) |
| AP53_1649 | hypo22 | FIG01114948: hypothetical protein |
| AP53_1650 | - | FIG01118158: hypothetical protein |
| AP53_1651 | def | Peptide deformylase (EC 3.5.1.88) |
| AP53_1652 | - | FIG01114677: hypothetical protein |
| AP53_1653 | marR | Transcriptional regulator, MarR family |
| AP53_1654 | polC | DNA polymerase III alpha subunit (EC 2.7.7.7) |
| AP53_1655 | proS | Prolyl-tRNA synthetase (EC 6.1.1.15), bacterial type |
| AP53_1656 | eep | Intramembrane protease RasP/YluC, implicated in cell division based on FtsL cleavage |
| AP53_1657 | | Phosphatidate cytidylyltransferase (EC 2.7.7.41) |
| AP53_1658 | | Undecaprenyl diphosphate synthase (EC 2.5.1.31) |
| AP53_1659 | yajC | Preprotein translocase subunit YajC (TC 3.A.5.1.1) |
| AP53_1660 | bta | putative transport accessory protein |
| AP53_1661 | pulA | Pullulanase (EC 3.2.1.41) |
| AP53_1662 | dexB | Glucan 1,6-alpha-glucosidase (EC 3.2.1.70) |
| AP53_1663 | | Multiple sugar ABC transporter, ATP-binding protein |
| AP53_1664 | lrp | Leucine rich protein |
| AP53_1665 | ska | Streptokinase |
| AP53_1666 | dtd | D-tyrosyl-tRNA(Tyr) deacylase |
| AP53_1667 | | GTP pyrophosphokinase (EC 2.7.6.5), (p)ppGpp synthetase I |
| AP53_1668 | sclA | Collagen-like surface protein |
| AP53_1669 | - | Ribonucleotide reduction protein NrdI |
| AP53_1670 | - | hypothetical protein |
| AP53_1671 | - | Exodeoxyribonuclease III (EC 3.1.11.2) |
| AP53_1672 | ptbA | PTS system, maltose and glucose-specific IIC component (EC 2.7.1.69) / PTS system, maltose and glucose-spe[illegible] |
| AP53_1673 | - | Ribosomal RNA small subunit methyltransferase E (EC 2.1.1.-) |

| | | |
|---|---|---|
| AP53_1674 | prmA | Ribosomal protein L11 methyltransferase (EC 2.1.1.-) |
| AP53_1675 | - | FIG01115320: hypothetical protein |
| AP53_1676 | - | 6-aminohexanoate-cyclic-dimer hydrolase (EC 3.5.2.12) |
| AP53_1677 | pabB | Para-aminobenzoate synthase, aminase component (EC 2.6.1.85) / Aminodeoxychorismate lyase (EC 4.1.3.3[illegible] |
| AP53_1678 | trpG | Anthranilate synthase, amidotransferase component (EC 4.1.3.27) @ Para-aminobenzoate synthase, amidot[illegible] |
| AP53_1679 | - | Chromosome segregation helicase |
| AP53_rna81 | - | tRNA-Lys-CTT |
| AP53_pseudo1680 | smeZ | Streptococcal mitogenic exotoxin Z (SmeZ) |
| AP53_1681 | dppA | Dipeptide-binding ABC transporter, periplasmic substrate-binding component (TC 3.A.1.5.2) |
| AP53_1682 | dppB | Dipeptide transport system permease protein DppB (TC 3.A.1.5.2) |
| AP53_1683 | dppC | Dipeptide transport system permease protein DppC (TC 3.A.1.5.2) |
| AP53_1684 | dppD | Dipeptide transport ATP-binding protein DppD (TC 3.A.1.5.2) |
| AP53_1685 | dppE | Dipeptide transport ATP-binding protein DppF (TC 3.A.1.5.2) |
| AP53_1686 | - | FIG01115696: hypothetical protein |
| AP53_1687 | htpA | Streptococcal histidine triad protein |
| AP53_1688 | lmb | Laminin-binding surface protein |
| AP53_1689 | - | hypothetical protein |
| AP53_1690 | fbaA | Fibronectin-binding protein |
| AP53_1691 | scpA | C5a peptidase (EC 3.4.21.-) |
| AP53_1692 | emm49 | Antiphagocytic M protein |
| AP53_1693 | pam | Antiphagocytic M protein |
| AP53_pseudo1695 | FcR/pseudogene | Antiphagocytic M protein/pseudogene causing stop codon at 85th aa |
| AP53_1696 | mga | M protein trans-acting positive regulator (Mga) |
| AP53_1697 | - | Hypothetical mga-associated protein |
| AP53_1698 | isp | Immunogenic secreted protein |
| AP53_1699 | ihk | Two-component system histidine kinase |
| AP53_1700 | irr | Two-component system response regulator |
| AP53_1701 | salY | ABC transporter permease protein |
| AP53_1702 | salX | ABC transporter ATP-binding protein |
| AP53_1703 | acrA | periplasmic component of efflux system |
| AP53_1704 | | FIG01115531: hypothetical protein |
| AP53_1705 | sHIP | FIG01116435: hypothetical protein |
| AP53_1706 | | FIG01114845: hypothetical protein |
| AP53_1707 | ropA;prsA | Peptidylproline cis-trans-isomerase (EC 5.2.1.8); Proteinase maturation protein (RopA) |
| AP53_1708 | spi | Streptopain inhibitor |

| | | |
|---|---|---|
| AP53_1709 | speB | Strepotococcal cysteine protease (Streptopain) (EC 3.4.22.10) / Streptococcal pyrogenic exotoxin B (SpeB) |
| AP53_1710 | | FIG01114305: hypothetical protein |
| AP53_1711 | | hypothetical protein |
| AP53_1712 | ropB | RopB; Rgg-like transcription regulator |
| AP53_1713 | spd | Streptodornase B; Mitogenic factor 1 |
| AP53_1714 | - | FIG01115319: hypothetical protein |
| AP53_1715 | ltrC | Low temperature requirement C protein |
| AP53_1716 | | Glycerol dehydrogenase (EC 1.1.1.6) |
| AP53_1717 | mipB | Transaldolase (EC 2.2.1.2) |
| AP53_1718 | pflD | Pyruvate formate-lyase (EC 2.3.1.54) |
| AP53_1719 | celB | PTS system, cellobiose-specific IIC component (EC 2.7.1.69) |
| AP53_1720 | celA | PTS system, cellobiose-specific IIB component (EC 2.7.1.69) |
| AP53_1721 | celC | PTS system, cellobiose-specific IIA component (EC 2.7.1.69) |
| AP53_1722 | DeoR | Deoxyribonucleoside regulator DeoR (transcriptional repressor) |
| AP53_1723 | | Transcriptional regulator, DeoR family |
| AP53_1724 | pflA | Pyruvate formate-lyase activating enzyme (EC 1.97.1.4) |
| AP53_1725 | - | FIG01114281: hypothetical protein |
| AP53_1726 | secE | Preprotein translocase subunit SecE (TC 3.A.5.1.1) |
| AP53_1727 | pbp2A | Multimodular transpeptidase-transglycosylase (EC 2.4.1.129) (EC 3.4.-.-) |
| AP53_1728 | - | Hypothetical protein SPy2060 |
| AP53_1729 | - | FIG01115477: hypothetical protein |
| AP53_1730 | - | Similar to ribosomal large subunit pseudouridine synthase D, Bacillus subtilis YhcT type |
| AP53_1731 | - | Hypothetical protein SPy2065 |
| AP53_1732 | - | Dipeptidase (EC 3.4.-.-) |
| AP53_1733 | - | transcriptional regulator, MutR family |
| AP53_1734 | - | Transcriptional regulator, MutR family |
| AP53_1735 | - | hypothetical protein |
| AP53_1736 | groEL | Heat shock protein 60 family chaperone GroEL |
| AP53_1737 | groES | Heat shock protein 60 family co-chaperone GroES |
| AP53_1738 | clpC | ATP-dependent Clp protease ATP-binding subunit ClpA |
| AP53_1739 | ctsR | Transcriptional regulator CtsR |
| AP53_1740 | - | hypothetical protein |
| AP53_1741 | cspA | Cold shock protein CspA |
| AP53_rna82 | - | tRNA-Cys-GCA |
| AP53_1742 | | Alkyl hydroperoxide reductase protein C (EC 1.6.4.-) |

| | | |
|---|---|---|
| AP53_1743 | | Alkyl hydroperoxide reductase protein F (EC 1.6.4.-) |
| AP53_1744 | | Imidazolonepropionase (EC 3.5.2.7) |
| AP53_1745 | | Urocanate hydratase (EC 4.2.1.49) |
| AP53_1746 | ftcD | Glutamate formiminotransferase (EC 2.1.2.5) @ Glutamate formyltransferase |
| AP53_1747 | fchA | Formiminotetrahydrofolate cyclodeaminase (EC 4.3.1.4) |
| AP53_1748 | fhs2 | Formate--tetrahydrofolate ligase (EC 6.3.4.3) |
| AP53_1749 | - | Conserved protein |
| AP53_1750 | - | putative cationic amino acid transporter protein |
| AP53_1751 | | Histidine ammonia-lyase (EC 4.3.1.3) |
| AP53_1752 | | Formiminoglutamase (EC 3.5.3.8) |
| AP53_1753 | - | Transcriptional regulator, LuxR family |
| AP53_1754 | rpsB | SSU ribosomal protein S2p (SAe) |
| AP53_1755 | tsf | Translation elongation factor Ts |
| AP53_1756 | pepO | Neutral endopeptidase O (EC 3.4.24.-) |
| AP53_1757 | treC | Trehalose-6-phosphate hydrolase (EC 3.2.1.93) |
| AP53_1758 | treP | PTS system, trehalose-specific IIB component (EC 2.7.1.69) / PTS system, trehalose-specific IIC component (E[illegible] |
| AP53_1759 | | Trehalose operon transcriptional repressor |
| AP53_1760 | - | FIG01115695: hypothetical protein |
| AP53_1761 | - | FIG01114717: hypothetical protein |
| AP53_1762 | - | Transcriptional regulator, HxlR family |
| AP53_1763 | - | Glyoxalase family protein |
| AP53_1764 | yaaA | UPF0246 protein YaaA |
| AP53_1765 | nrdG | Ribonucleotide reductase of class III (anaerobic), activating protein (EC 1.97.1.4) |
| AP53_1766 | - | Acetyltransferase (EC 2.3.1.-) |
| AP53_1767 | MviM | Putative oxidoreductase |
| AP53_1768 | hypo24 | FIG01115181: hypothetical protein |
| AP53_1769 | nrdD | Ribonucleotide reductase of class III (anaerobic), large subunit (EC 1.17.4.2) |
| AP53_1770 | - | Competence-induced protein Ccs4 |
| AP53_1771 | - | hypothetical protein |
| AP53_1772 | - | Ortholog of S. aureus MRSA252 (BX571856) SAR1694 |
| AP53_1773 | yggF | Putative Holliday junction resolvase YggF |
| AP53_1774 | - | Hypothetical protein possible functionally linked with Alanyl-tRNA synthetase |
| AP53_1775 | spxA2 | Transcriptional regulator SpxA2 |
| AP53_1776 | recA | RecA protein |
| AP53_1777 | cinA | Molybdopterin binding motif, CinA N-terminal domain / C-terminal domain of CinA type S |

| | | |
|---|---|---|
| AP53_1778 | tag | DNA-3-methyladenine glycosylase (EC 3.2.2.20) |
| AP53_1779 | ruvA | Holliday junction DNA helicase RuvA |
| AP53_1780 | lmrP | Multidrug resistance protein B |
| AP53_1781 | mutL | DNA mismatch repair protein MutL |
| AP53_1782 | - | Phage integrase: site-specific recombinase |
| AP53_1783 | - | hypothetical protein |
| AP53_1784 | - | Phage transcriptional repressor |
| AP53_1785 | - | conserved hypothetical protein - phage associated |
| AP53_1786 | - | kilA protein, putative phage-related DNA binding protein |
| AP53_1787 | - | Phage antirepressor protein |
| AP53_1788 | - | FIG01118151: hypothetical protein |
| AP53_1789 | - | Mobile element protein |
| AP53_1790 | - | FIG01118442: hypothetical protein |
| AP53_1791 | - | FIG01115767: hypothetical protein |
| AP53_1792 | - | phage protein |
| AP53_1793 | - | FIG01114419: hypothetical protein |
| AP53_1794 | - | FIG01114073: hypothetical protein |
| AP53_1795 | - | Phage protein |
| AP53_1796 | - | DNA primase, phage associated |
| AP53_1797 | - | FIG01114328: hypothetical protein |
| AP53_1798 | - | Phage protein |
| AP53_1799 | - | FIG01115461: hypothetical protein |
| AP53_1800 | - | FIG01115581: hypothetical protein |
| AP53_1801 | - | FIG01116340: hypothetical protein |
| AP53_1802 | - | Phage encoded transcriptional regulator, ArpU family |
| AP53_1803 | - | hypothetical protein |
| AP53_1804 | mutS | DNA mismatch repair protein MutS |
| AP53_1805 | ymcA | biofilm formation protein YmcA |
| AP53_1806 | argR | Arginine pathway regulatory protein ArgR, repressor of arg regulon |
| AP53_1807 | | Arginyl-tRNA synthetase (EC 6.1.1.19) |
| AP53_1808 | | bacteriocin immunity protein |
| AP53_1809 | - | Transporter |
| AP53_1810 | - | membrane protein, putative |
| AP53_1811 | - | Integral membrane protein |
| AP53_1812 | | Aspartyl-tRNA synthetase (EC 6.1.1.12) |
