## Supplementary figures and images for "Group A *Streptococcus* remains viable inside fibrin clots and gains access to human plasminogen for subsequent fibrinolysis and dissemination"

### Supplemental Videos

#### Slide 1
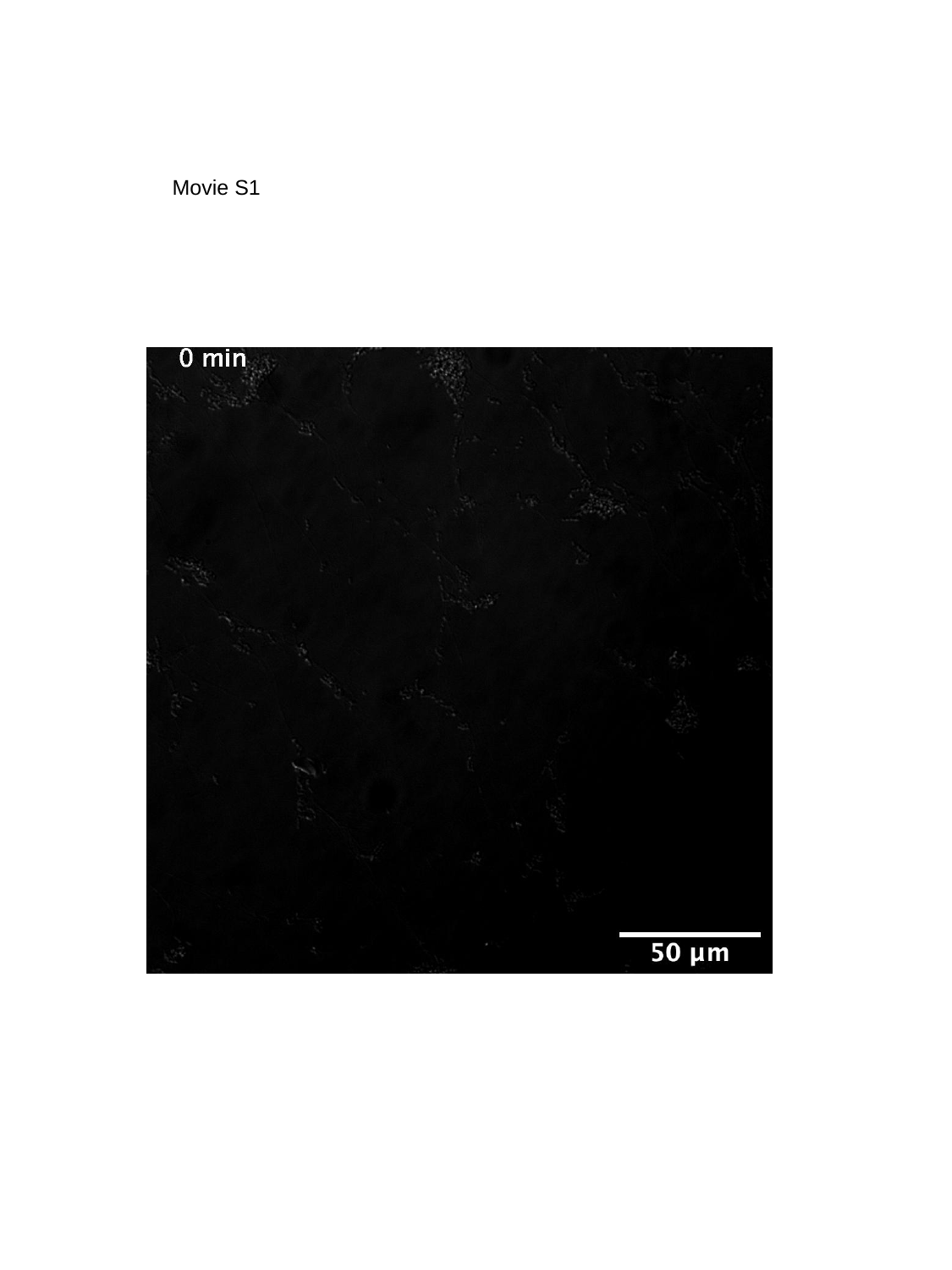

Movie S1

#### Slide 2
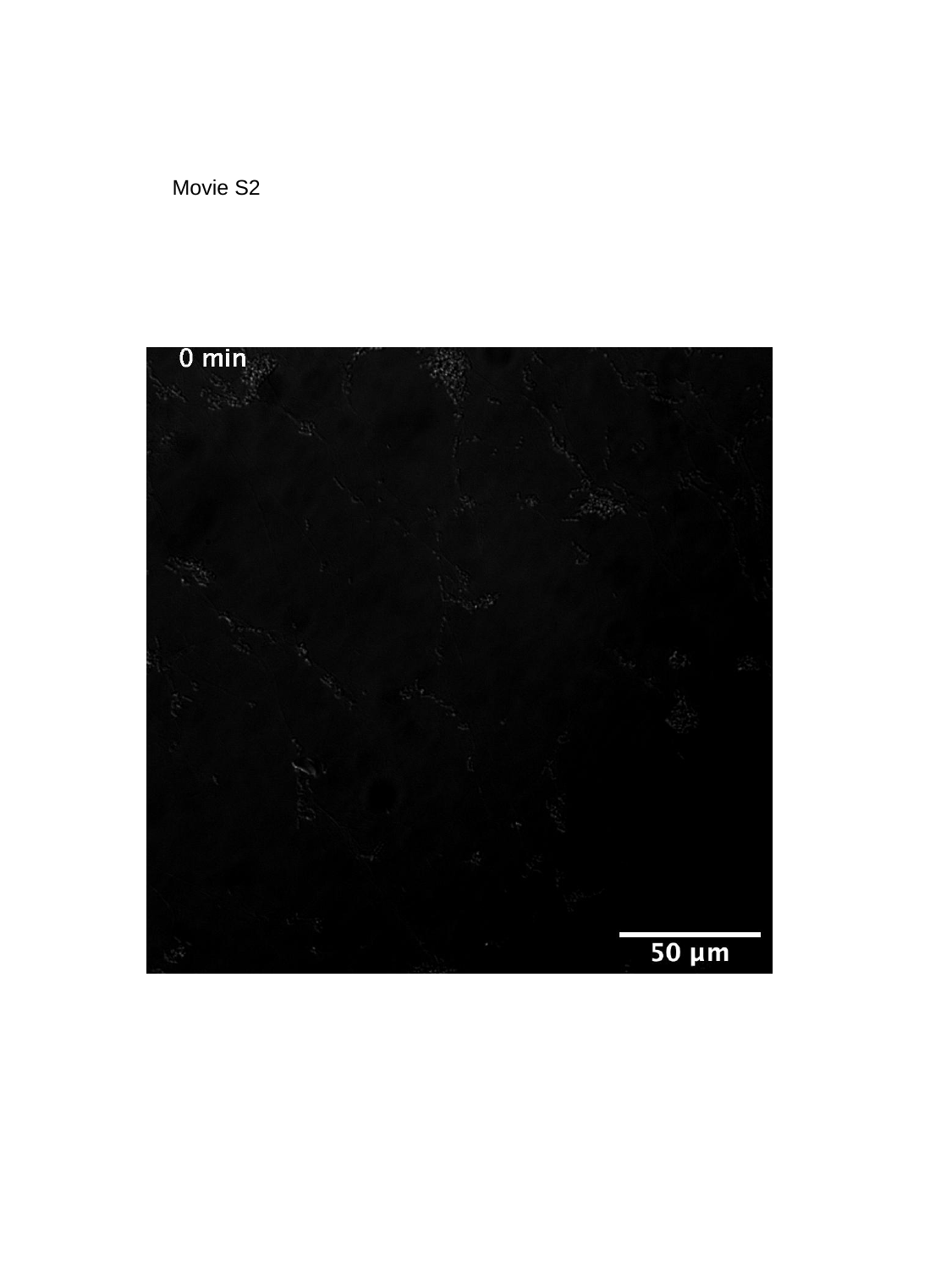

Movie S2

#### Slide 3
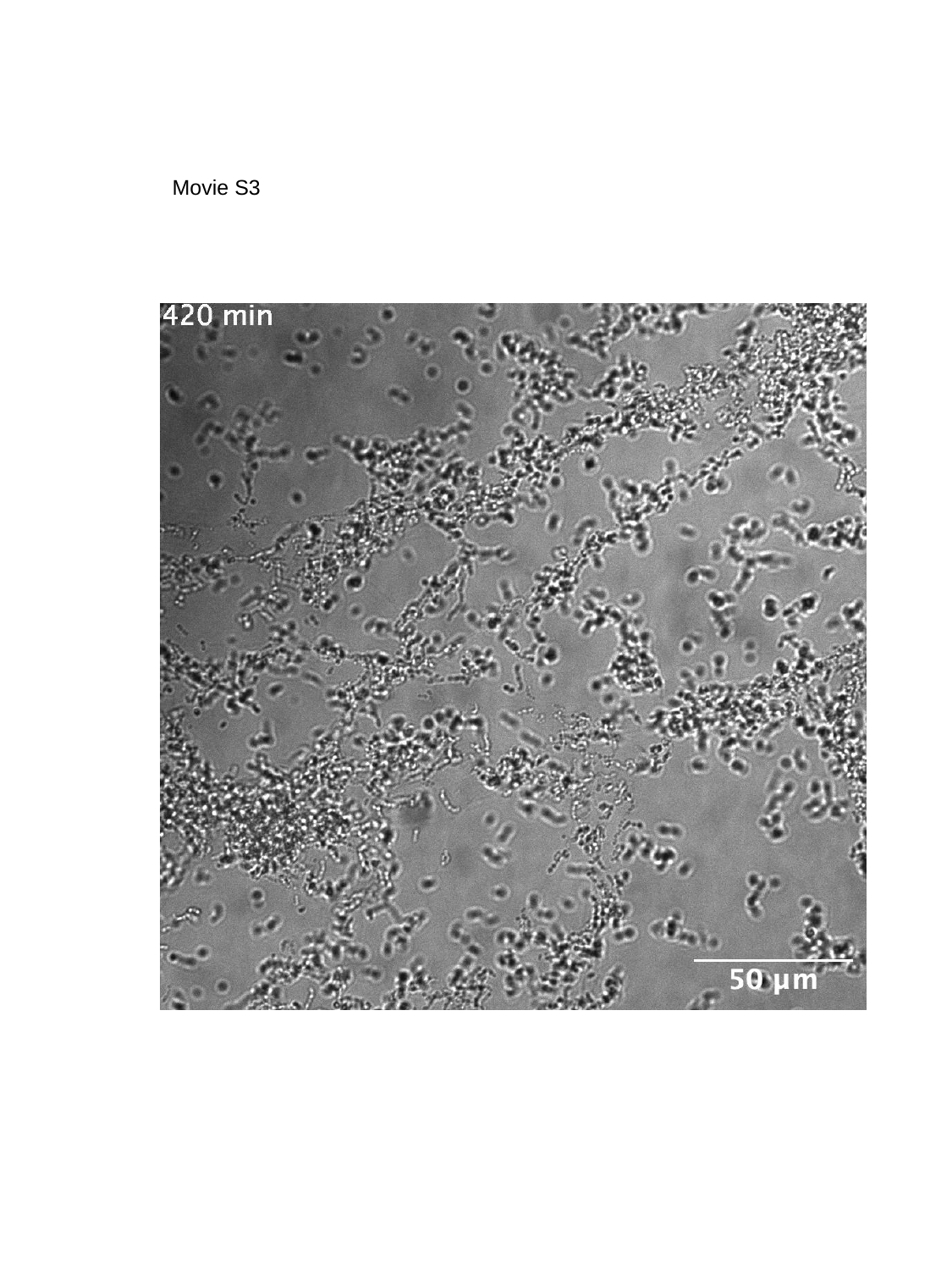

Movie S3

#### Slide 4
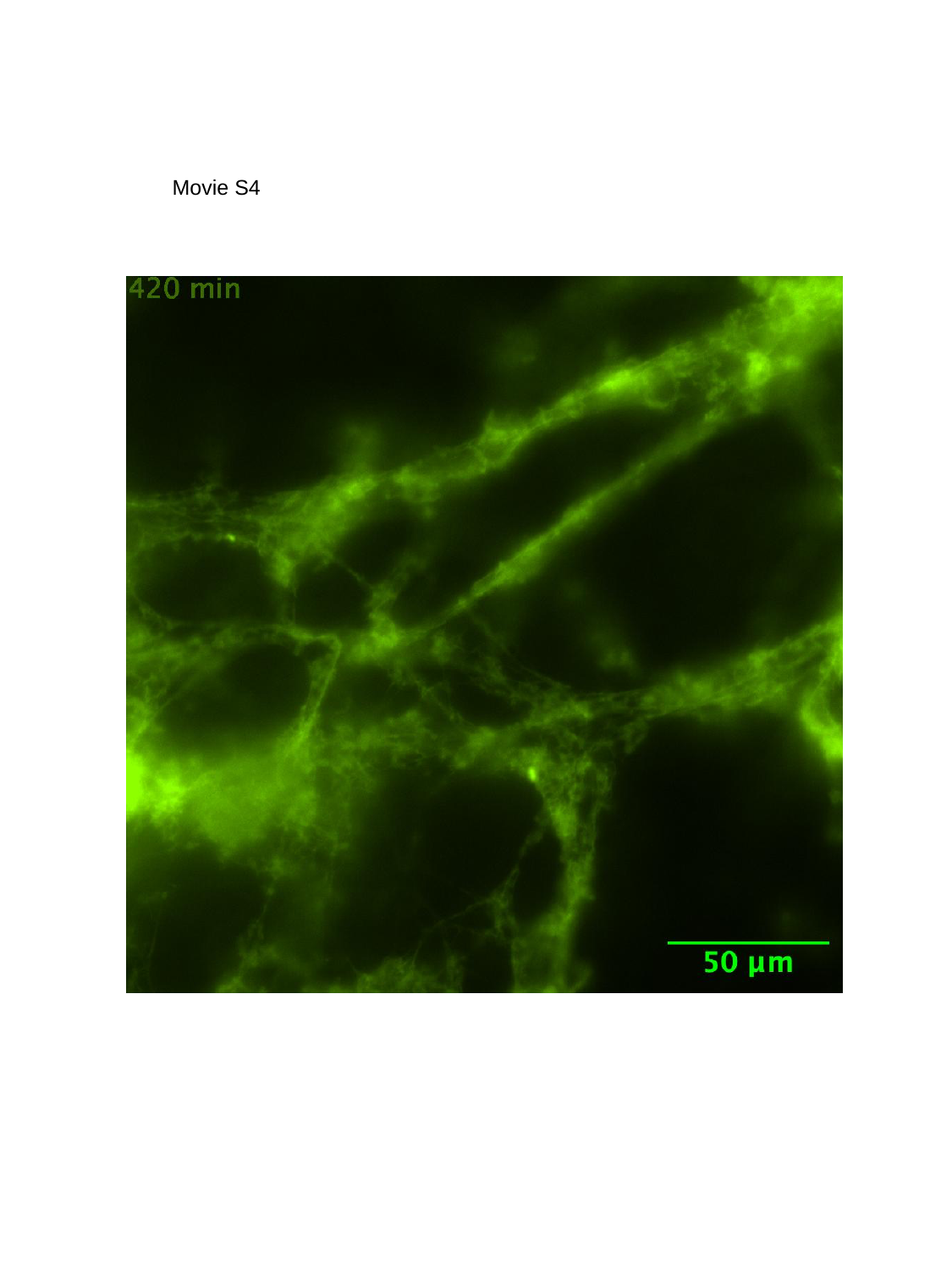

Movie S4

#### Slide 5
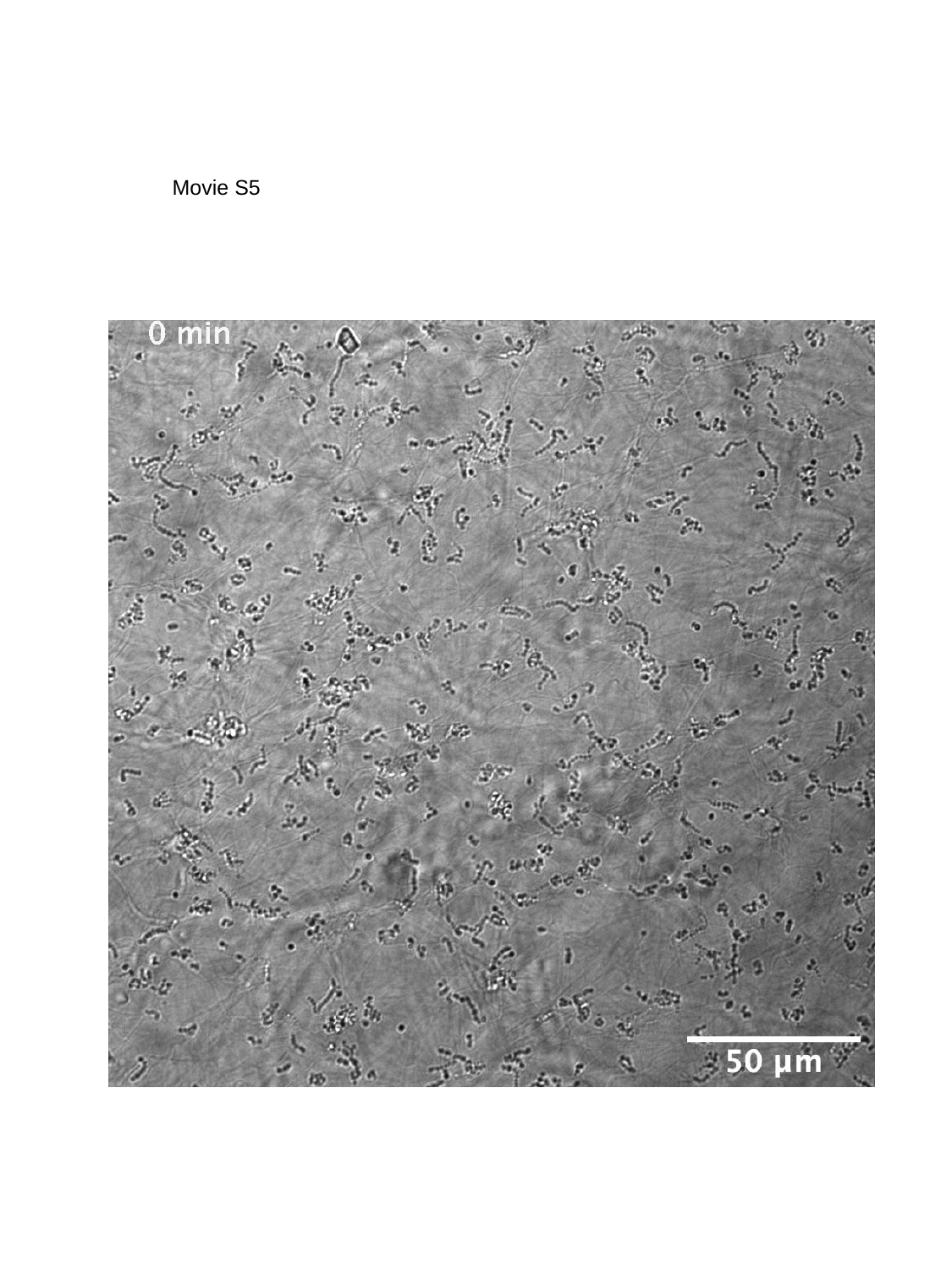

Movie S5
